# Principles of ion binding to RNA inferred from the analysis of a 1.55 Å resolution bacterial ribosome structure – Part I: Mg^2+^

**DOI:** 10.1101/2024.04.07.588377

**Authors:** Filip Leonarski, Anja Henning-Knechtel, Serdal Kirmizialtin, Eric Ennifar, Pascal Auffinger

## Abstract

The importance of Mg^2+^ ions for RNA structure and function can difficultly be overstated. Several attempts were made to establish a comprehensive Mg^2+^ binding site classification. However, such descriptions were hampered by poorly modelled ion binding sites. Recently, ribosome cryo-EM structures with resolutions < 2.0 Å allowed better descriptions of site-bound ions. However, in a recent cryo-EM 1.55 Å *E. coli* ribosome structure, incomplete ion assignments prevented a full understanding of their binding modes. We revisited this model to establish general binding principles applicable to any RNA of sufficient resolution. These principles rely on the 2.9 Å distance separating two Mg^2+^-bound *cis-*water molecules. By applying these rules, we could assign all Mg^2+^ ions bound with 2 to 4 non-water oxygens. We also uncovered unanticipated motifs where up to five adjacent nucleotides wrap around a single ion. The formation of these complex motifs involves a hierarchical dehydration of the Mg^2+^ ions, a process that plays a significant role in ribosome biogenesis and in the folding of large RNAs. These binding principles enhance our understanding of the roles of ions in RNA structure and will help refining the solvation shell of different ribosomes and of RNAs with complex topologies.

## INTRODUCTION

Since the first biochemical investigations on ribosomes, it has become clear that Mg^2+^ along with K^+^ and polyamines are essential for the structure and function of these systems (1–8). The first ribosome structure that allowed the assignment of mono- and divalent ions was the *Haloarcula marismortui* 50S large subunit (Hm-LSU) by Steitz and co-workers (9). A comprehensive study about the identification of Mg^2+^, Na^+^ and K^+^ ions was subsequently published by Klein and Steitz (10). Meanwhile, several hundreds of ribosomal cryo-EM/X-ray structures have been deposited to the PDB with resolutions in the 1.55 to > 4.0 Å range (11). The number of assigned Mg^2+^ ions is zero for the most cautious authors and reaches ≈1,260 Mg^2+^ per 70S ribosome (or one Mg^2+^ per three nucleotide; PDBid: 4Y4O) as in a 2.3 Å resolution *Thermus thermophilus* (12) structure or close to full neutrality as in another 3.1 Å resolution *Thermus thermophilus* (13) structure (PDBid: 4V6F; ≈2,970 Mg^2+^/70S). These numbers are in sharp contrast with the 138 Mg^2+^, 85 Na^+^, 2 K^+^, 5 Cd^2+^ cations and the 30 Cl^-^ and 4 acetate anions that were assigned in the latest 2013 refinement (14) of the Hm-LSU structure (PDBid: 4V9F; resolution: 2.4 Å).

According to current knowledge, 100-300 Mg^2+^ site-bound ions per ribosome seem a reasonable number depending on ribosome size and cryo-EM/X-ray experimental conditions (3,7). Larger Mg^2+^ numbers result usually from specific buffer conditions or from an excess of faith in the amount of information that can be extracted from experimental data. For RNA and other metal containing biomolecular systems, the latter bias has been documented (15–24). Some tools to pinpoint assignment issues have been developed (25–27) but none specific to nucleic acids besides the MgRNA attempt (28) that has been critically evaluated (15,16).

To get better insights into the biological functions of Mg^2+^ and K^+^ ions, it is essential to advance our understanding of their binding stereochemistry. For that, we re-examined the recent *Escherichia coli* 70S ribosomal cryo-EM structure (11) that was solved to a stunning 1.55 Å resolution (PDBid: 8b0x). This structure comprises 1 Zn^2+^, 361 Mg^2+^, 168 K^+^ and 11,461 water molecules. A rapid examination of 8b0x made us realize that ≈261 Mg^2+^ ions exhibit incomplete octahedral coordination shells and that ≈28 K^+^ displaying coordination distances close to 2.8 Å were wrongly assigned as Mg^2+^ ions (**Figure 1**). Such imprecise or incomplete assignments are a hurdle to database surveys (20,27,28) and to the development of artificial intelligence (AI) tools (29–33).

**Figure 1.**
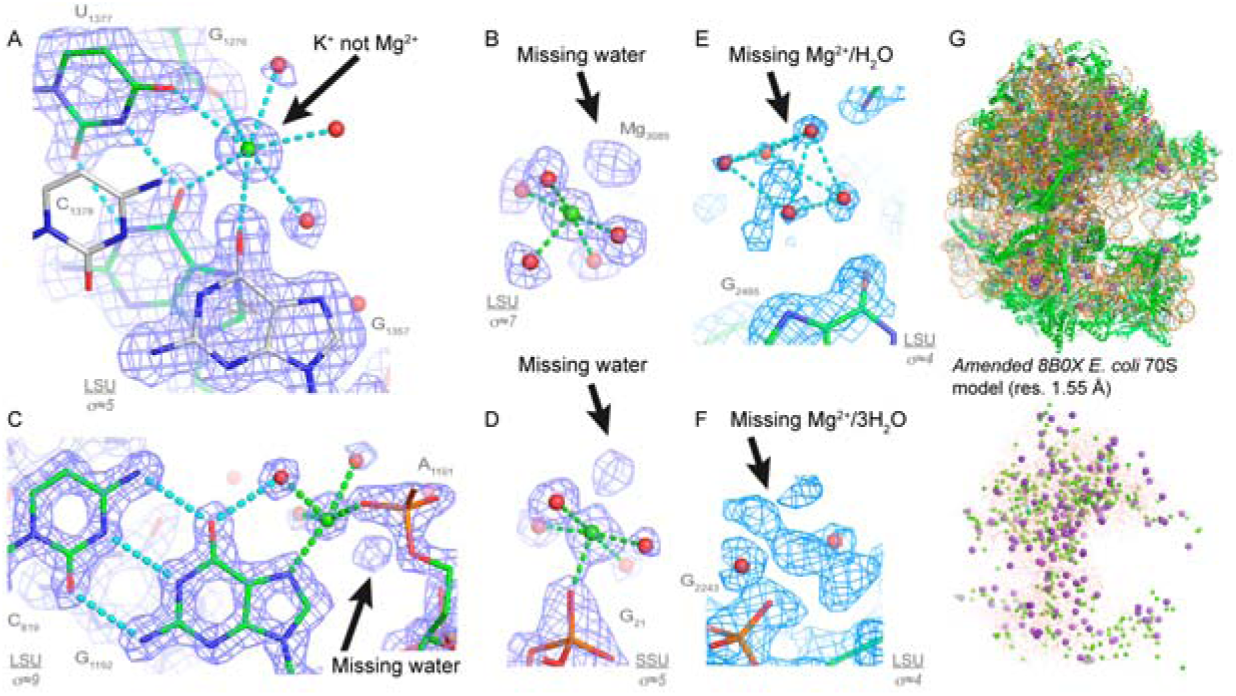
8b0x ion assignments and first shell completion issues. (**A**) This Mg^2+^ ion has been assigned to the deep grove of a G•U pair (a neighbouring C=G pair shows silver carbons). However, the heptahedral ion coordination with *d*(Mg^2+^…O) ≈2.8 Å implies that this ion is K^+^ and not Mg^2+^; see Figure 2 for distance criteria. (**B**) These 6O_w_, (**C**) O_ph_.N_b_.4O_w_ and (**D**) O_ph_.5O_w_ ions display missing first shell water molecules. (**E**) A Mg^2+^ and a water molecule belonging to a hexahydrated Mg^2+^ were not assigned. This 6O_w_ ion binds to a guanine Hoogsteen edge. (**F**) The densities of this O_ph_.5O_w_ ion and of three water molecules were left empty. See Figure 4 for Mg^2+^ binding site nomenclature. Green/cyan lines mark distances < 2.3 Å and in the 2.6–3.2 Å range. (**G**) View of the amended 8b0x model (top) and of its solvation shell (bottom) with 403 Mg^2+^ (green spheres), 231 K^+^ (purple spheres) and 21,397 water molecules (red dots). Sigma levels are indicated where appropriate.

The aim of the present study is to apply state-of-the-art stereochemical knowledge to define criteria to validate the ionic structure of small RNAs up to large ribosome structures. Indeed, correcting inaccurate assignments such as those described here and in earlier reports is an essential prerequisite for the understanding of ion binding features (15–17,19). This should help to build the foundation of a thorough “prior knowledge” Mg^2+^/K^+^ ion binding site classification and would help amending ion positions derived from earlier and sometimes less well resolved structures. For instance, the authors of the 8b0x structure based part of their Mg^2+^/K^+^ assignments on the 7k00 and 6qnr structures which led to some of the issues described herein (34,35).

Herein, we first present an updated classification of Mg^2+^ binding sites based on the MgRNA study (15,16,28) and the seminal investigations by Steitz and Klein (10). We illustrate each of the uncovered ion binding site category and discuss the formation of novel motifs such as those involving Mg^2+^…O2’ coordination. We propose also a classification of Mg^2+^…Mg^2+^ and Mg^2+^…K^+^ ion pairs that are recurrently observed in catalytic and non-catalytic systems. Based on the amended 8b0x structure, we could define a set of simple stereochemical rules to place Mg^2+^ ions at key locations. We stress the importance of Mg^2+^ bidentate clamps and demonstrate that by scanning the OP…OP distances < 3.4 Å, it is possible to assign close to 100% of these clamps. We infer that these Mg^2+^ based motifs are key to essential ribosome folding events occurring during biogenesis.

We present also rules to characterize the binding of “chelated” hexahydrated Mg(H_2_O)_6_^2+^ ions that are based on a correspondence between the local rRNA hydration structure and that of the Mg^2+^ hydration shell. Thus, we underline the importance of thoroughly refining the solvation shell of RNA systems even when the 1.5 Å resolution limit has been reached. However, even in this high-resolution ribosomal structure, the diffuse Mg(H_2_O) ^2+^ and K^+^ ions, which dominantly contribute to the charge neutralization process, still escape direct observation. Although, K^+^ ions are also key to the ribosome structure (15,35–38), we will discuss their binding features in a companion paper.

Of course, some drawbacks are associated with the use of cryo-EM techniques. Despite some attempts in that direction (39,40) and contrary to X-ray experiments, cryo-EM techniques do not produce differential maps. Thus, like for most structures deposited to the PDB, only indirect evidence were gathered to identify the ions. Though anomalous signals can be obtained for transition metals and K^+^ ions (35,41), no current technique produces signals allowing direct Mg^2+^ identification. Therefore, we mostly had to rely on stereochemistry. Apart these limitations, we felt that 8b0x is currently the best candidate to investigate the ribosomal ionic shell. Given conservation of the sequence and structure of the ribosome cores, we believe that a significant percentage of the chelated ions present in the *E. coli* ribosomes are conserved in other archaea, bacterial, and eukaryote cytosolic/mitochondrial ribosomes (10,42–44). Likewise, we infer that present stereochemical rules are not limited in their use to ribosomes and can be successfully applied to the exploration of the solvation shell of all nucleic acids as well as protein systems. Present study leads to clearer views of the ion binding rules that, in turn, may help better design structures used to start molecular dynamics (MD) simulations (29,45–51) and allow more accurate RNA 3D structure predictions.

## METHODS

### 8b0x cryo-EM buffers

The final cryo-EM buffer contained 25 mM Mg(OAc)_2_, 100 mM K(OAc), 50 mM HEPES, and 1 mM DTT for a pH of 7.4 (11). Thus, next to HEPES, DTT and possible contaminants (see below), only Mg^2+^ and K^+^ cations along with acetate anions and water molecules are part of the 8b0x solvation shell. For notes related to other *E. coli* ribosome structures with resolutions < 2.0 Å, see **SI**.

### Updated CSD histograms for Mg and K

Distance and angle histograms related to the Mg^2+^ and K^+^ ions were derived from the Cambridge Structural Database (CSD version 5.43, update November 2022; see **Figure 2/3**) (52,53). These histograms are updated versions of those presented earlier (15,16,54). Only good resolution structures were considered (R_factor ≤ 0.05 %). Disordered, error containing, polymeric and powder structures were discarded. All O/N atom types were considered with the exception of protonated nitrogens that do not contact cations. Charge constraints were not applied givent that the CSD attribution process is sometimes arbitrary (55). Only hexacoordinated Mg ions were considered since six is their dominant coordination number in biomolecules. For potassium (K), no restrictions on the coordination number of coordinated atoms were applied as the coordination number can range from six to nine (15,35,56). Hence, no restrictions on the number of coordinated atoms were applied for this ion. For additional potassium coordination features, see **SI**.

**Figure 2.**
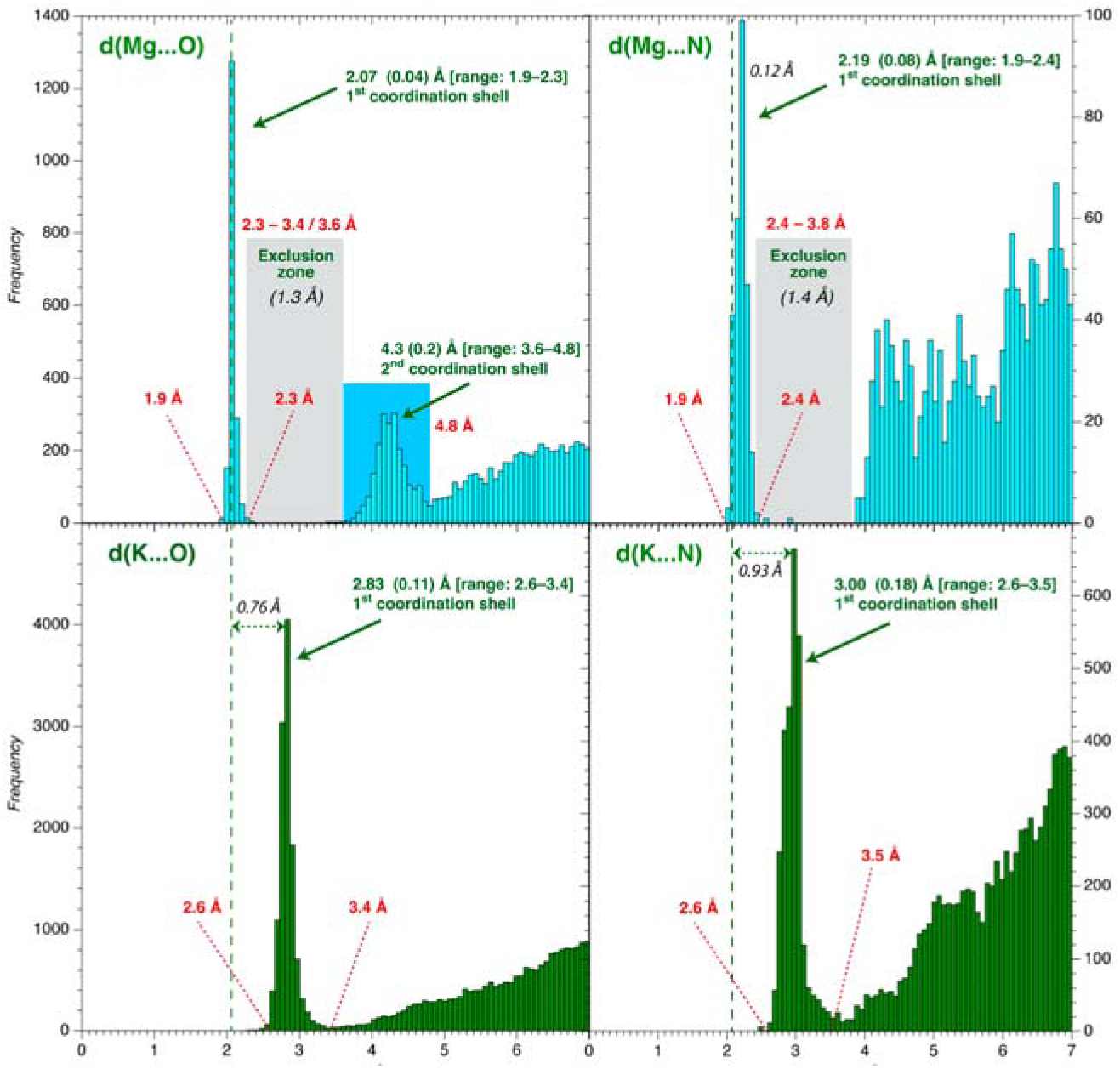
O/N coordination distance histograms for Mg/K derived from the CSD (version 5.43, update November 2022). (**Top**) See **Method** section for CSD search criteria. Only hexacoordinated Mg ions were considered; no coordination number constraints were placed on K. Standard deviations for the 1^st^ and 2^nd^ peaks are in parenthesis. All types of O and non-protonated N atoms were used to build these histograms (15,16,54). A vertical dashed line marks the average *d*(Mg…O) ≈2.07 Å coordination distance in all histograms. Mg exclusion zones are marked by grey rectangles. Note that we readjusted the 3.6 Å exclusion zone limit to 3.4 Å (see **SI)**.

### Mg coordination parameters

Average coordination distances for hexacoordinated Mg derived from the CSD histograms are *d*(Mg…O/N) ≈2.07±0.04 Å and *d*(Mg…N) ≈2.19±0.08 Å (**Figure 2**). A 2.3–3.6 Å exclusion zone was drawn for Mg specifying that no O/N atoms should be present in this distance range. However, given stereochemical constraints associated with highly chelated Mg^2+^ coordination motifs not represented in the CSD, we readjusted the upper limit of the exclusion zone from 3.6 to 3.4 Å (see **SI**). Slightly longer *d*(Mg…N) distances are observed in the CSD histograms. The origin of this offset remains unclear. It might in part be linked to the binding of bulky imidazole rings that are the main imino nitrogen containing biomolecular ligands (57). For Mg, the *cis-* and *trans-* coordination angles are θ(O…Mg…O) ≈90±3° and θ(O…Mg…O) ≈177±4°. The resulting average distances between oxygen atoms in *cis-* and *trans-* are *d*(O…O) ≈2.93±0.09 Å and ≈4.13±0.07 Å, respectively (**Figure 3**).

**Figure 3.**
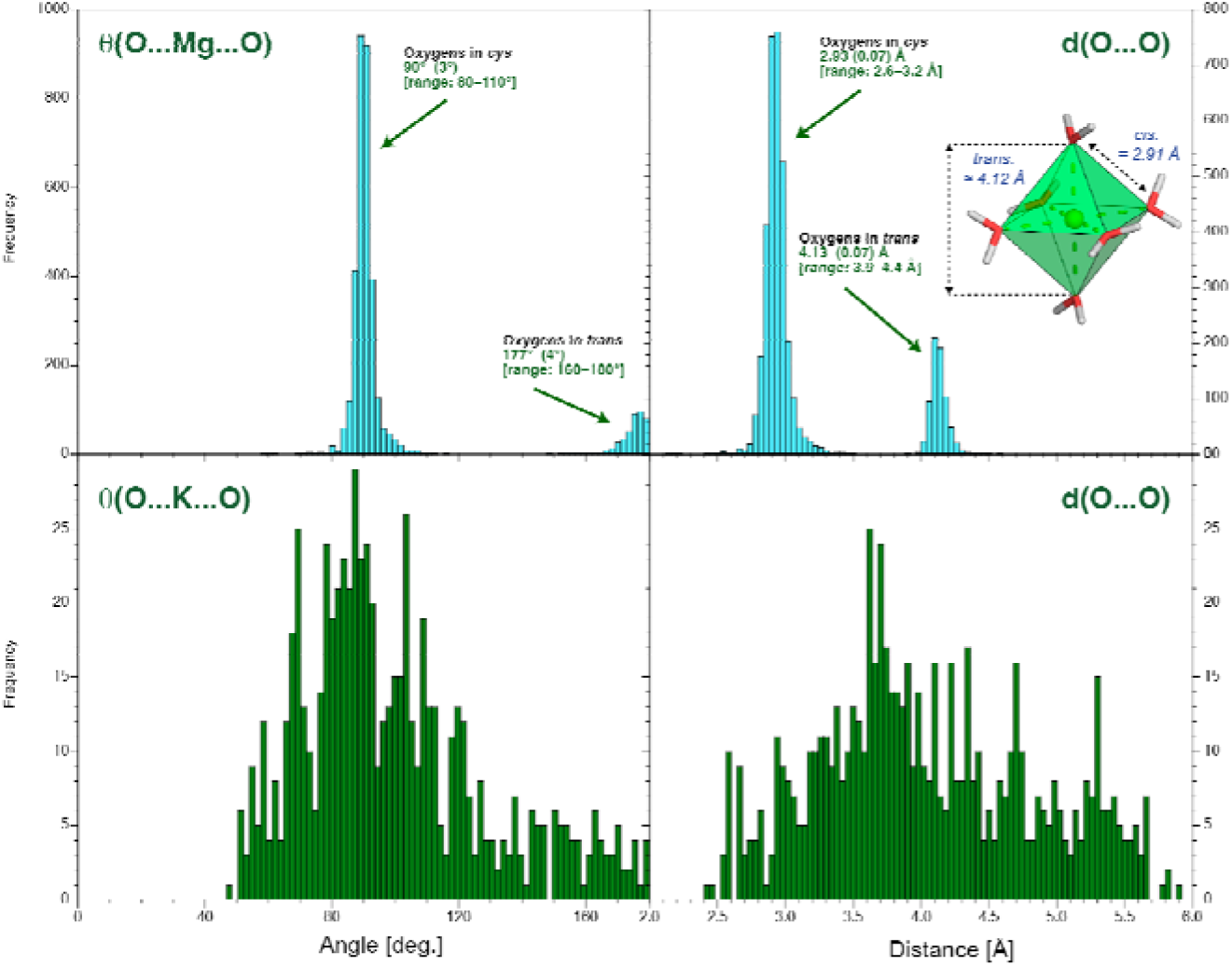
First shell oxygen coordination angle and *d*(O…O) distance histograms for Mg/K derived from the CSD. (**Left**) Coordination angle histograms for θ(O…Mg/K…O) and (**right**) related coordination distances for oxygen ligands in the ion first coordination shell derived from the CSD (52,53). The insert shows an ultra-high-accuracy Mg(H_2_O)_6_^2+^ X-ray structure showing the water octahedral arrangement around Mg^2+^ (121,122) and the *cis-*/*trans-* configurations. The angle histograms for K are marginaly useful given that the most frequent K ligands in the CSD are diethoxy (O-C-C-O) groups found in crown ethers and cryptands. These groups are rare in nucleic acids except when the cryo-EM/X-ray buffers contains ethylene glycol monomers or PEG fragments (123). Diethoxy groups were excluded from the CSD searches shown **Figures 2/3**.

### Ion binding site nomenclature

The ion binding site nomenclature, that has been extended to comprise amino acid atoms, is derived from the MgRNA study (15,16,28). O_ph_ corresponds to OP1/OP2 phosphate anionic oxygens, O_r_ to O2’/O4’/O3’/O5’ ribose and phosphate bridging oxygens, O_b_ to nucleobase O2/O4/O6 oxygens and N_b_ to non-protonated nucleobase N1/N3/N7 nitrogens. Ion binding sites are named through a combination of these categories. For instance, 2O_ph_.4O_w_ stands for a hexacoordinated ion bound to two O_ph_ atoms and four waters. The sites with two O_ph_ atoms in *cis-* or *trans-* and four water molecules are named *cis*-2O_ph_.4O_w_ and *trans-*2O_ph_.4O_w_. When three non-water atoms such as O_ph_ are attached to the ion, the corresponding isoforms become *fac-*3O_ph_.3O_w_ (*facial*) when the three O_ph_ pairs are in *cis-* and *mer-*3O_ph_.3O_w_ (*meridional*) when at least one O_ph_ pair is in *trans-* (28). When four non-water atoms are bound to the ion, the *cis-/trans-* terminology is used to name the respective orientation of the two bound water molecules in place of that used for coordinated O_ph_ atoms. To distinguish them from the cis-/trans-2O_ph_.4O_w_ types, the *<u>Cis</u>-/<u>Trans</u>-* prefix (underlined with a capital) are used. Finally, this *cis-/trans-/fac-/mer-/<u>Cis</u>-/<u>Trans</u>-* nomenclature can be extended to any O_ph_/O_b_/O_r_/N_b_ atom combinations. For proteins, we used O_bb_ (previously O_back_) for amino acid backbone oxygens, O_coo_ for Asp/Glu carboxyl oxygens, O_cno_ for Asn/Gln carbonyl oxygens, OH_prot_ for Ser/Thr/Tyr hydroxyl oxygens and N_His_ for non-protonated ND1/NE2 histidine nitrogens.

### Phenix refinements

We used the outputs of an in-house “*IonDiagnosis*” tool developed earlier (17) to identify stereochemical issues, missing coordinated waters, and ion misassignments (see **SI** for an 8b0x *“IonDiagnosis”* output). We “idealized” the Mg^2+^ coordination shell through the addition of *d*(Mg^2+^…O/N) ≈2.07/2.19 Å distance restraints that allowed an improved ion binding site assessment. Indeed, Mg^2+^ binding sites are definitely more difficult to identify when ions present one or more of the following flaws: *(i)* coordination distances that exceed the 2.3 Å limit, *(ii)* incomplete coordination shells or *(iii)* unassigned ions (**Figure 1**). After having defined appropriate distance restraints, the modified 8b0x structure was refined in Phenix (58). We refrained of using angle restraints that could have led, especially in a first investigation, to an inappropriate level of idealization (see next paragraph). This resulted in an amended 8b0x model. The coordinates of this 1.0 model are available from the authors upon request.

### Consistency criteria used to check the assigned Mg^2+^ ion binding sites

We used several criteria to estimate the consistency of the identified Mg^2+^ binding sites. First, we checked whether six coordination distances in the 1.9–2.3 Å range were present. Then, the octahedral character of the coordination shell was assessed by calculating the angular deviations of the bound atoms. For that purpose, the hexacoordinated Mg^2+^ ions were classified into one of three categories: “correct”, “slightly distorted” and “highly distorted”. These categories are determined based on the deviation of the cumulated (ligand…Mg^2+^…ligand) angular value associated with a regular coordination octahedron: less than 5° for “correct”, in the 5-10° range for “slightly distorted” and more than 10° for “highly distorted”, respectively. The ions in the “highly distorted” category display stereochemistry that are affected by local disorder, partial occupancy, or any other factors that might blur their coordination pattern. As such, we refrained to include these “distorted” ions in the “well-defined” binding site counts of our survey unless otherwise specified. However, we included them in our amended 8b0x model.

A second consistency criterion was defined as follows. If the density peak of a given ion is below an arbitrary 4.0 Å r.m.s.d. value as defined by the Coot visualization program (59), this ion was excluded from our “well-defined” Mg^2+^ binding site ensemble. Further, if some coordinating atoms are found in the 2.3–3.4 Å exclusion zone, we tried to correct the stereochemistry of the binding site by imposing local restraints. If we were unable to find any rational for these “exclusion zone” contacts, these ions were discarded. However, the ions associated with sites that were considered as “not-well-defined” might have been appropriately modelled but the experimental density maps were not sufficiently precise to support their assignment. The coherency criteria used to check the K^+^ assignments are described in the **SI** which also comprise an EXCEL file describing all “distorted” and “well-defined” ion binding sites in the amended 8b0x model.

## RESULTS

### Amendments made to the 8b0x solvent shell structure

Herein, we assume that the preferred Mg^2+^ coordination number is six in most nucleic acid environments with strict *d(Mg^2+^…O/N)* ≈*2.07/2.19 Å* coordination distances to oxygen atoms and to purine N7 and histidine ND1/NE2 nitrogens (**Figure 2**). Despite its exceptional 1.55 Å resolution, a rapid inspection of the 8b0x structure deposited to the PDB and of the corresponding *“IonDiagnosis”* file (see **SI**) reveals that only 96 of the 361 assigned Mg^2+^ are hexacoordinated with coordination distances in the 1.9–2.3 Å range.

To establish a comprehensive classification of Mg^2+^ ribosomal binding sites, we completed their solvation shell by adding 104 hexacoordinated Mg^2+^ ions to non-assigned density spots. We also added during subsequent Phenix refinements *d*(Mg^2+^…O/N) ≈2.07/2.19 Å coordination distance restraints. Through that process a much clearer coordination picture emerged, although additional “fixes” where necessary. For instance, 28 Mg^2+^ ions with coordination distances in better agreement with those of K^+^ were reassigned (**Figure 1A**). Likewise, two K^+^ ions were reassigned as Mg^2+^.

The coordination of a sub-category of Mg^2+^ ions appeared significantly distorted even when using distance restraints implying that the experimental densities are locally too blurred to allow a good modelling of these sites (see **Methods**). These ions were not included in the “well-defined” Mg^2+^ binding site categories described below. However, they were included in our amended 8b0x model. For some Mg^2+^, it was not possible to define a clear hexacoordinated pattern. This resulted in the deletion of ≈40 of the assigned 8b0x Mg^2+^ ions or their reassignment to water. It was also necessary to slightly modify the upper limit of the Mg^2+^ exclusion zone from 3.6 to 3.4 Å (see **SI**).

To summarize, all densities close to rRNA/r-protein atoms were inspected and allowed to recover ≈110 Mg^2+^ ions that were either misassigned as water molecules, K^+^ ions or not assigned at all (**Figure 1**). This resulted in a final model that contains a total of 403 Mg^2+^, 231 K^+^, and 21,397 water molecules that contrast with the 361 Mg^2+^, 168 K^+^, and 11,461 water molecules initially assigned. The two models share 299 Mg^2+^ common positions. Although it is documented that anions can bind to nucleic acids (60,61), we were unable to assign any of those (see **SI**).

### Mg^2+^ binding site categorization

Our ion identification process allowed a clear-cut classification of Mg^2+^ ion binding sites in terms of type and frequency. Since the resolution of cryo-EM structures weakens with the distance to the center of the ribosomal particle, the atomic models associated with some regions of the structure are less precise. This is the case for part of the SSU. However, we considered that the assigned 403 Mg^2+^ ions (290 “well-defined”) correspond to a significant part of the chelated divalent ions necessary to stabilize the bacterial ribosome. These ions differ from diffuse or weakly chelated ions that have too weak densities to be characterized in cryo-EM structures. Next, we describe the classification of all these uncovered sites along with their occurrences in the amended 8b0x structure (**Figure 4**).

- <u>6O_w_ coordination</u> (79 “well-defined” occurrences among 115): Although Mg[H_2_O] ^2+^ can form up to 12 hydrogen-bonds, the number of water-mediated contacts in 8b0x is in the 1–8 range with an average of ≈4 when we consider the 79 “well-defined” ions with density peaks > 4.0 and with an octahedral angle deviation < 10° (see **Methods**). The best hydrogen bond acceptors are the Hoogsteen (G)O6/(G)N7 atoms and the anionic OP1/OP2 phosphate oxygens (10,28). Importantly, depending on the structural context, any combination of OP1/OP2/O_b_/O_r_/N_b_ atoms can be part of the Mg[H_2_O]_6_^2+^ coordination shell (for more, see **SI**).

**Figure 4.**
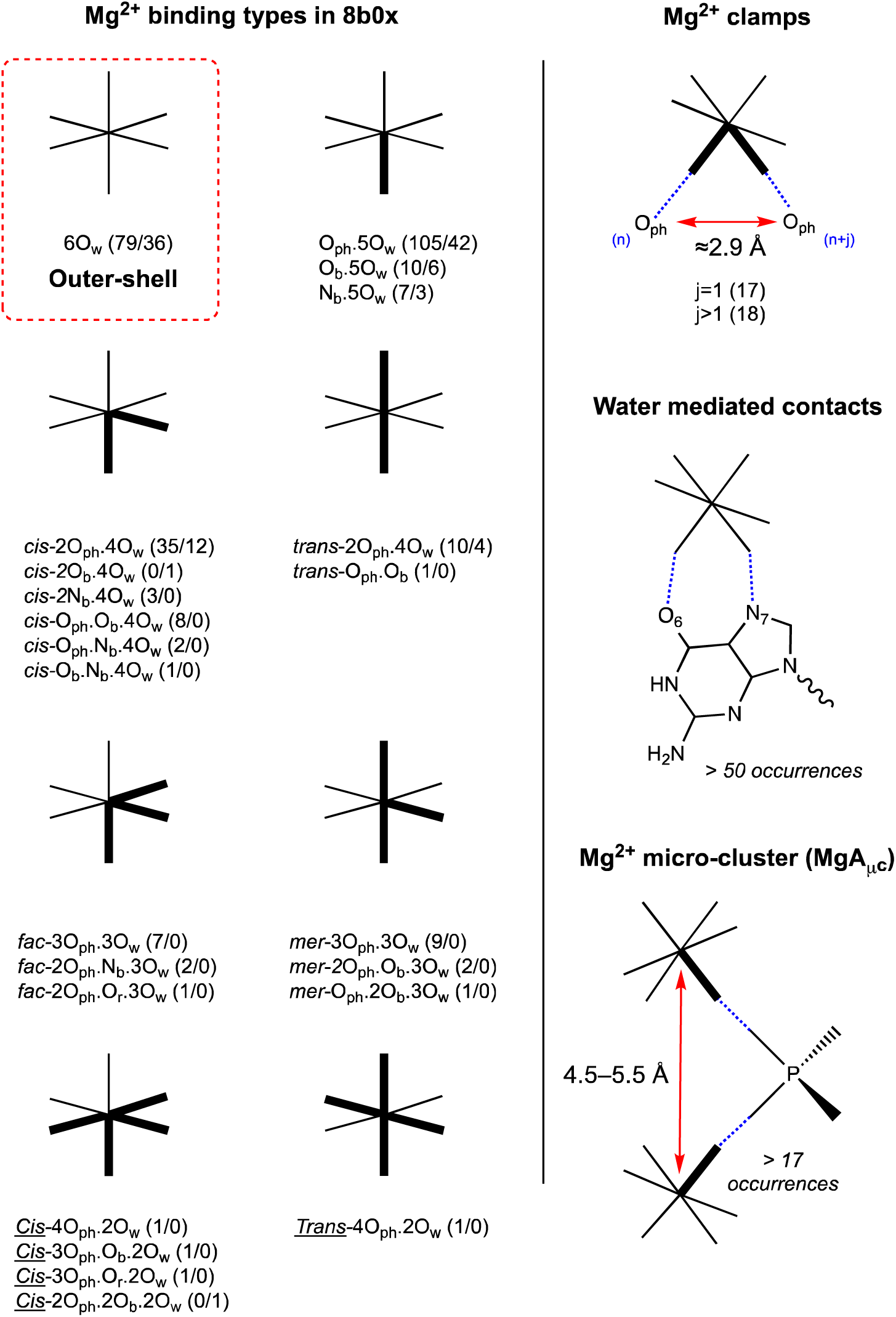
Schematics and occurrences of Mg^2+^ binding types observed in the amended 8b0x structure. (Left) In addition to the outer-shell 6O_w_ binding type, all the inner-shell Mg^2+^ ion binding types and occurrences uncovered in the 8b0x structure are shown (“well-defined” and “not-well-defined” binding site occurrences are given in parenthesis; we did not include direct binding to r-proteins that account for a total of 15 ions). (Right) The top panel shows a Mg^2+^ clamp where the ion coordinates in *cis-* to two adjacent (j=1) or distant (j>1) O_ph_ atoms. The middle panel shows a 6O_w_ outer-shell binding to a guanine Hoogsteen edge. The bottom panel shows a Mg^2+^ micro-cluster of the MgA_μc_ type (see Figure 14).

The most strongly bound 6O_w_ ion in 8b0x establishes eight water-mediated contacts to OP/O6/N7 atoms and appears to have been trapped during the folding of domain “III” (**Figure 5A**). This ion has to be considered as strongly chelated contrary to the common knowledge that suggest that 6O_w_ ions bind to helical grooves or the RNA surface and are exchangeable. In the present case, it is difficult to envisage an Mg[H_2_O]_6_^2+^ exchange with bulk ions unless domain “III” unfolds.

**Figure 5.**
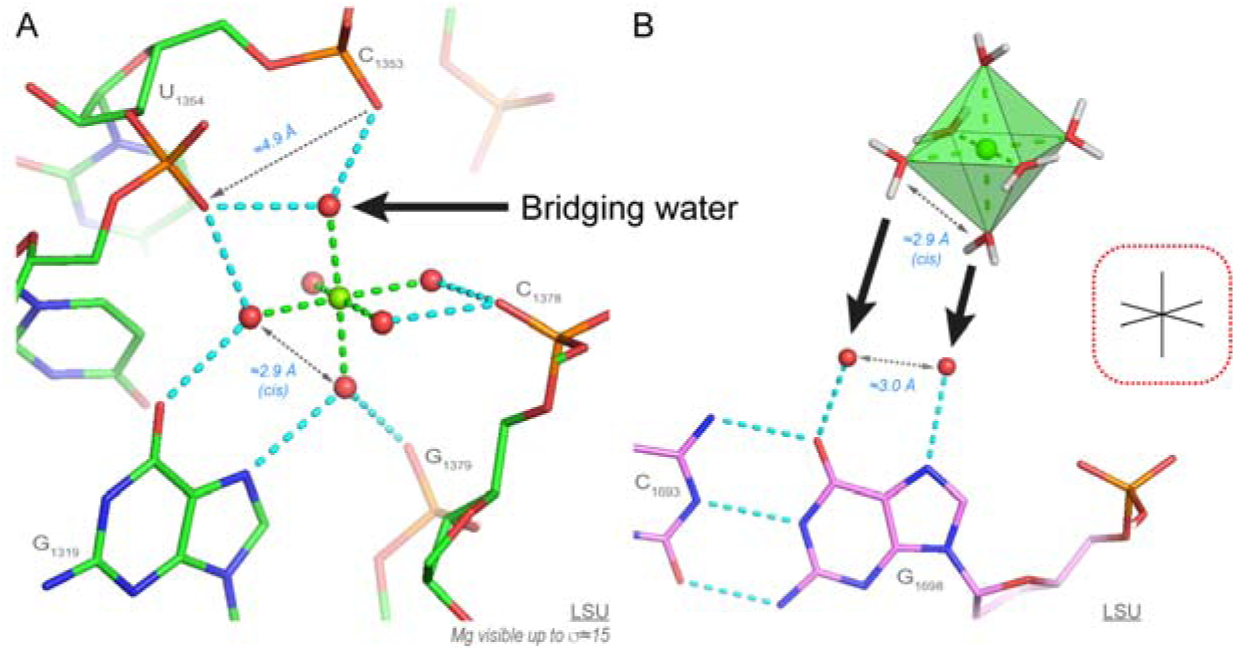
A 6O_w_ ion establishing eight water-mediated contacts to rRNA atoms and a Mg[H_2_O]_6_^2+^ binding to a guanine Hoogsteen edge. (**A**) This domain “III” 6O_w_ ion occupies a tight binding pocket where it forms eight water-mediated contacts. Note the binding of two first shell water molecules to a guanine Hoogsteen edge and the recurrent first shell water molecule bridging two consecutive phosphate groups. (**B**) Schematics illustrating how two water molecules bound to a guanine Hoogsteen edge and separated by ≈2.9 Å can be replaced by two first shell water molecules in *cis-* of a Mg[H_2_O]_6_^2+^ ion. Green/cyan lines mark distances < 2.3 Å and in the 2.6–3.2 Å range. For clarity, experimental densities and some water molecules were hidden.

Interestingly, when a Mg[H_2_O]_6_^2+^ ion establishes water-mediated contacts to a guanine Hoogsteen edge, the two O6/N7 bound water molecules can replace two Mg^2+^ water molecules coordinated in *cis-*. This process takes advantage of the similar ≈2.8–3.0 Å distance separating the coordinated guanine and Mg[H_2_O]_6_^2+^ water molecules in *cis-*. This relationship helps to understand why guanine Hoogsteen edges are favourable binding sites (**Figures 3/5B**).

- <u>O_ph_.5O_w_ coordination</u> (105 “well-defined” occurrences among 147): This coordination mode is the most frequently observed and involves OP1/OP2 atoms with a slight preference for OP2 atoms (46 versus 59). The coordinated water molecules establish a variable number of water-mediated contacts with rRNA/r-protein atoms. The most highly connected O_ph_.5O_w_ ion involves a direct O_ph_ bond completed by ten water-mediated contacts (**Figure 6A**). This stunning motif encompasses five consecutive nucleotides that wrap around the ion.

**Figure 6.**
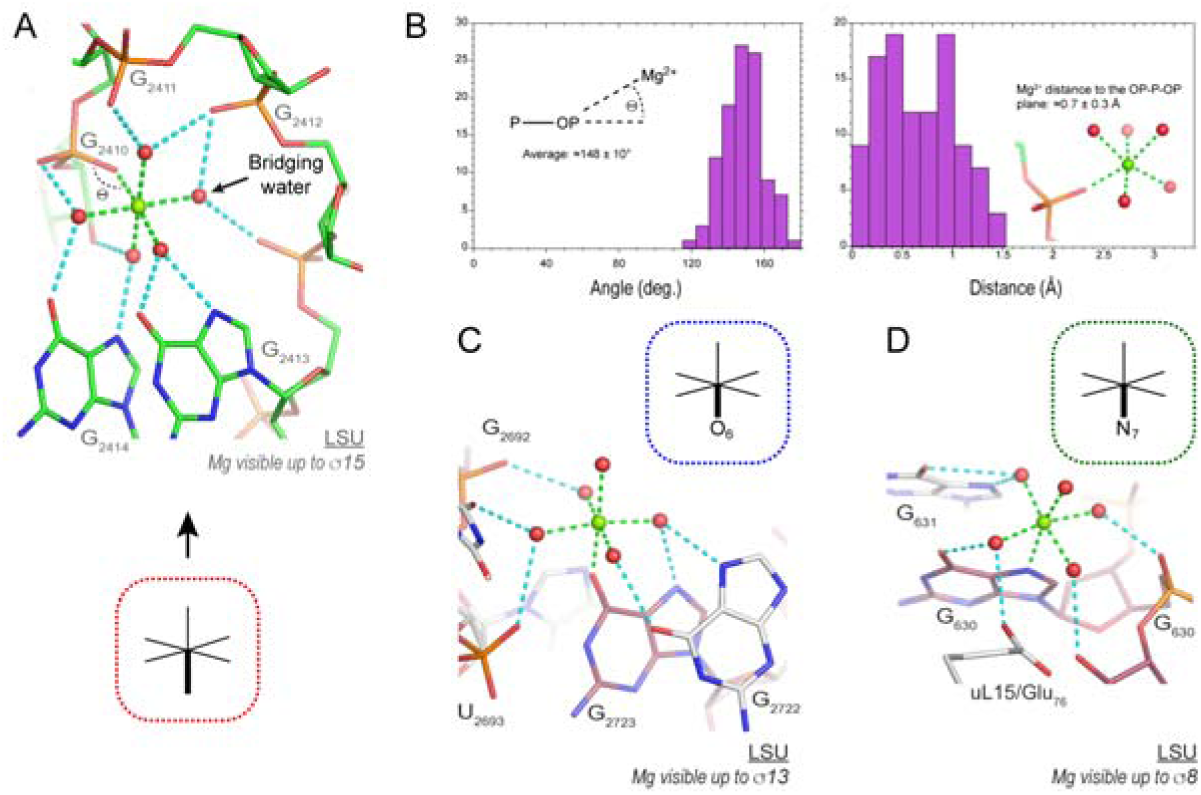
Examples of O_ph_/O_b_/N_b_.5O_w_ Mg^2+^ ions establishing water-mediated contacts to rRNA atoms. (**A**) This domain “V” O_ph_.5O_w_ ion is the most highly connected in 8b0x with one inner-shell and 10 outer-shell contacts. Strikingly, this ion establishes contacts with five consecutive nucleotides and displays at least one bridging water molecule similar to that shown Figure 5A. (**B**) The left histogram illustrates the P–OP…Mg^2+^ angular distribution with the average value calculated from the ensemble of 105 “well-defined” O_ph_.5O_w_ occurrences. The right histogram describes the Mg^2+^ distance to the OP–P–OP plane for the same 105 occurrences. (**C**) This domain “VI” O_b_.5O_w_ ion, although less connected, establishes inner- and outer-shell contacts to two consecutive guanine Hoogsteen edges. (**D**) This domain “I” N_b_.5O_w_ ion establishes inner- and outer-shell contacts to a guanine Hoogsteen edge and a r-protein. Green/cyan lines mark distances < 2.3 Å and in the 2.6–3.2 Å range. Experimental densities and some water molecules were hidden.

When bound to a single phosphate group, the (P–OP…Mg^2+^) angle is ≈148±10° (**Figure 6B**). The bound Mg^2+^ is mainly located in the OP1–P–OP2 plane with an average deviation of ≈0.7±0.3 Å (**Figure 6B**). We did not observe the rather elusive “bidentate” type of coordination (62,63) that implies that Mg^2+^ interacts with both anionic oxygens of carboxylate or phosphate groups (**Figure S1**). With 15 binding occurrences, the guanine Hoogsteen edge is, as for 6O_w_ ions, an excellent anchor-point for water-mediated contacts (**Figure 6A**). These rather strict constraints can be used to define rules for (semi)-automatic ion placement techniques (see below).

- <u>O_b_.5O_w_ coordination</u> (10 “well-defined” occurrences among 16): These rare patterns involve (G)O6, (U)O4, (PSU)O2 and (C)O2 atoms. Binding to O_b_ atoms is only possible in rare structural contexts the characteristics of which have not been established yet (**Figure 6C**). An example of a completely encapsulated O_b_.5O_w_ ion was reported in a P4-P6 group I intron structure (15,64). In most instances, the Mg^2+^ ion turns to the Hoogsteen edge but can also be oriented towards the nucleobase Watson-Crick edge depending on neighboring nucleotide conformations. The C=O…Mg^2+^ angle is ≈144° with Mg^2+^ strictly in the nucleobase plane in contrast to monovalent ions binding to the same atoms (15,65).
- <u>N_b_.5O_w_ coordination</u> (7 “well-defined” occurrences among 10): Only patterns involving (G/A)N7 atoms were observed excluding those that involve N1/N3 atoms (**Figure 6D**). Note that coordination to (A)N7 is only observed in higher-order motifs involving more than two-direct contacts (see below). The electron densities around the N7 atoms are often blurred and difficult to interpret as discussed previously (66). Occasionally, direct Mg^2+^ binding to N7 atoms was considered to be important for catalytic mechanisms (16,67,68).
- *<u>cis-</u>*<u>2O_ph_.4O_w_ coordination and variations</u> (35 “well-defined” occurrences among 47): *cis-*2O_ph_.4O_w_ coordination modes can be split into two main categories involving adjacent (17 occurrences) and sequence distant nucleotides (18 occurrences). The first category (**Figure 7A**) was comprised of bidentate Mg^2+^ clamp motifs (28,42,67,69). These motifs were also defined as 10-membered ring systems (Mg^2+^…OP–P–O5’–C5’–C4’–C3’–O3–P–OP) and involve any combination of OP1/OP2 atoms, the most common one involving the deep groove OP2/OP2 atoms. Other patterns comprise O_ph_ atoms from distant residues (**Figure 7B**). In a few instances, clamps involving a Mg^2+^ bridging an O_ph_(n) and an O_ph_(n+2) atom occur. These were considered as new motifs and called 16-membered ring systems but are simply a variation of the 10-membered ring system (30).

**Figure 7.**
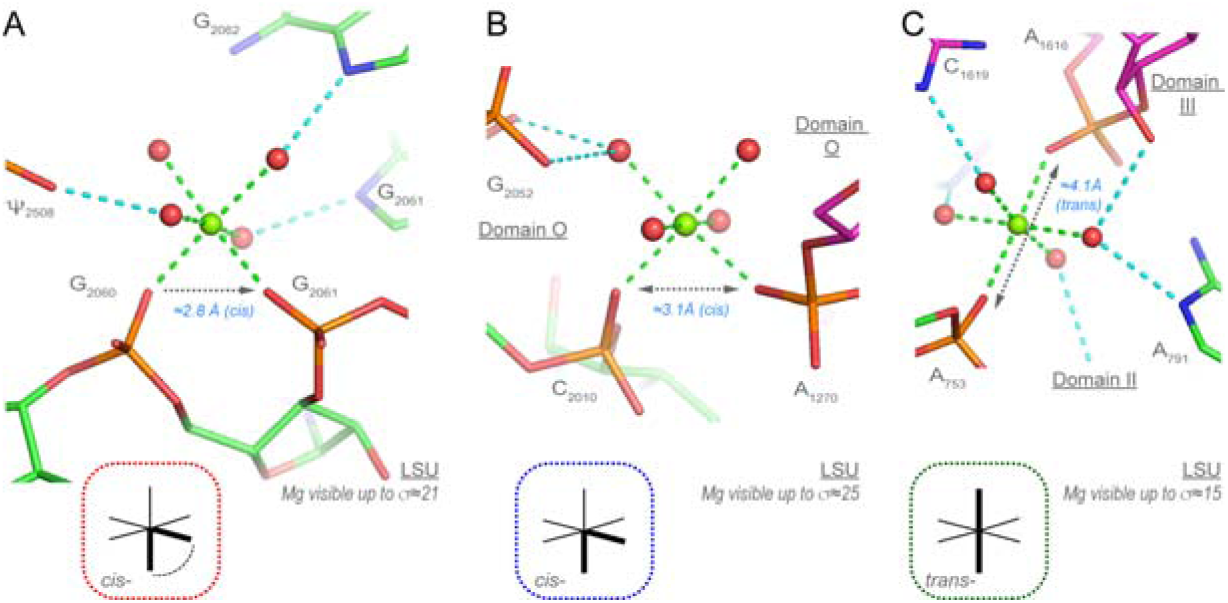
Two *cis-*2O_ph_.4O_w_ and one *trans-*2O_ph_.4O_w_ motifs. (**A**) These domain “V” phosphate groups form a bidentate Mg^2+^ clamp also defined as a Mg^2+^ 10-membered ring (42,67,69). (**B**) The two phosphate groups in *cis-* belong to domain “0”. (**C**) The two phosphate groups are in *trans-* and the associated Mg^2+^ joins LSU domains “II” and “III”. Green/cyan lines mark distances < 2.3 Å and in the 2.6–3.2 Å range. Experimental densities and some water molecules were hidden.

Importantly, there is a strict correspondence between the distance separating two Mg^2+^ first shell water molecules in *cis-* and the ≈2.9 Å distance separating two coordinated O_ph_ atoms (**Figures 3/7A**). Thus, Mg^2+^ ions are required to stabilize short O_ph_ contacts with *d*(O_ph_…O_ph_) < 3.4 Å. This distance range is associated with the Mg^2+^ first shell water molecules in a *cis-* configuration. As discussed below, all short *d*(O_ph_…O_ph_) ≈2.9 Å distances observed in 8b0x are stabilized directly or indirectly by a Mg^2+^ ion (for stabilization through both, direct and water-mediated interactions, see **Figure S2**). The *cis-*2N_b_.4O_w_ binding types are discussed below. Variations of these patterns with O_b_/N atoms replacing O_ph_ atoms are discussed in the **SI**.

- *<u>cis-</u>*<u>2N_b_.4O_w_ coordination</u> (3 “well-defined” occurrences): One SSU and two LSU *cis-*2N_b_.4O_w_ patterns involving N7 atoms were observed. They engage G/G or A/G nucleotide pairs (**Figures 8/S3**) and were named elsewhere as “purine N7 seats” (15,28). However, it might be more appropriate to name them “head-to-tail stacked purine” motifs (**Figures 8/S3**). In each instance, these ions that bind to two N7 atoms are also coordinated through water-mediated contacts to three or four OP2 atoms forming a total of 8 to 9 inner- and outer-shell contacts. This rare motif is involved in the folding and stabilization of the rRNA subunits and is recurrent in ribosomes from various kingdoms (16,28).

**Figure 8.**
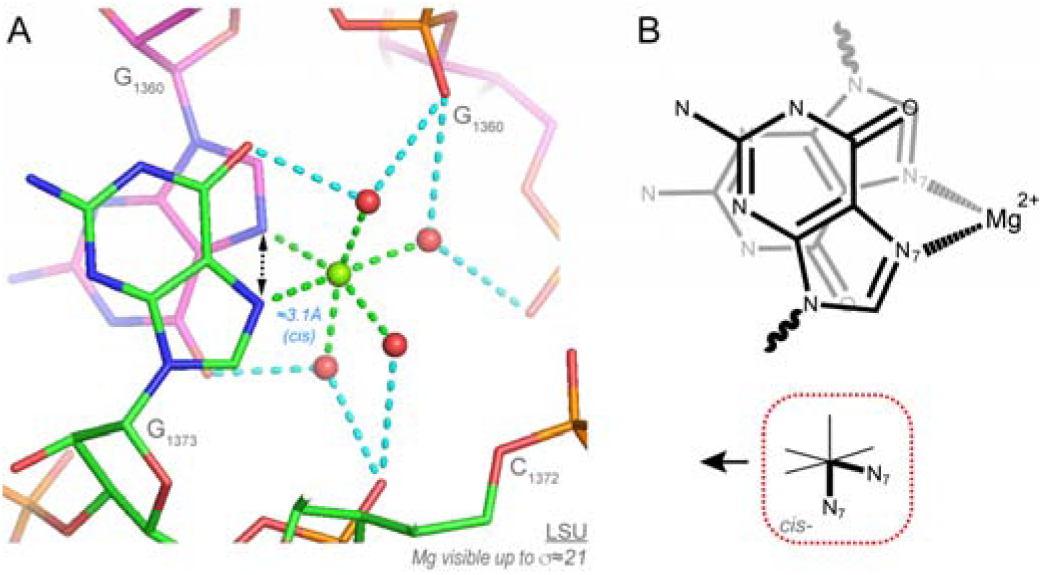
“Head-to-tail stacked purine” motif. (**A**) Two domain “III” guanines coordinate a Mg^2+^ through their N7 atoms in a *cis-*2N_b_.4O_w_ arrangement. As a result, a total of 9 inner- and outer-shell contacts are formed. The purines in these examples are separated by >10 nucleotides. Two similar *cis-*2N_b_.4O_w_ arrangements are described in **Figure S3B/C**. (**B**) Schematics showing the *cis-*2N_b_.4O_w_ “head-to-tail stacked purine” arrangement. Green/cyan lines mark distances < 2.3 Å and in the 2.6–3.2 Å range. Experimental densities and some water molecules were hidden.

Some of us questioned earlier the nature of the ion associated with the *cis-*2N_b_.4O_w_ motif and suggested that the assigned Mg^2+^ might be an hexahydrated Zn^2+^ given the very high-density peaks observed in some related X-ray structures and given that the affinity of Mg^2+^ for purine N7 atoms is much lower than that of Zn^2+^ (16,54). At this stage, this hypothesis cannot be excluded and Zn^2+^ might be present at similar locations in rRNA even as contaminants (see below).

- *<u>trans-</u>*<u>2O_ph_.4O_w_ coordination and variations</u> (10 “well-defined” occurrences among 14): These important patterns are less frequent than the *cis-* patterns. They appear more difficult to characterize given *d*(O_ph_…O_ph_) ≈4.2 Å (**Figure 3/7C**). Their architecture implies a greater flexibility and lower stability when compared to patterns in *cis-*. Variations of this motif involving the replacement of one or two O_ph_ atom(s) by O_b_/N_b_ atoms are rare (see **Figure S2A**).
- *<u>fac-</u>*<u>3O_ph_.3O_w_ and *mer-*3O_ph_.3O_w_ coordination and variations</u> (22 “well-defined” occurrences among 23): The 3O_ph_.3O_w_ arrangement comprises 7 *fac-* and 9 *mer-* arrangements. We believe these arrangements, most probably formed during ribosome biogenesis to and crucialfor the mature ribosome structure. Most of these occurrences comprise a Mg^2+^ bidentate motif (**Figure 9**) with only one exception in the SSU, where the bidentate motif involves a two-nucleotide insertion. Variations of these patterns are mentioned in the **SI**.
- *<u>Cis-/Trans-</u>*<u>4O_ph_.2O_w_ coordination and variations</u> (5 “well-defined” occurrences among 8): We observed five “well-defined” LSU binding sites in addition to three “poorly-defined” ones: two in the SSU and one in the uS2 r-protein region. For naming these binding sites, we used the *<u>Cis</u>-/<u>Trans</u>-* configuration of the two bound water molecules (for nomenclature details, see **Methods**). The first binding site is *<u>Trans</u>-*4O_ph_.2O_w_. Here, all four O_ph_ atoms are coplanar with Mg^2+^ (**Figure 10A**). The second is *<u>Cis</u>-*4O_ph_.2O_w_ (**Figure 10B**). This domain “V” site displays a unique pattern of three consecutive nucleotides that is completed by a fourth domain “II” phosphate group.

**Figure 9.**
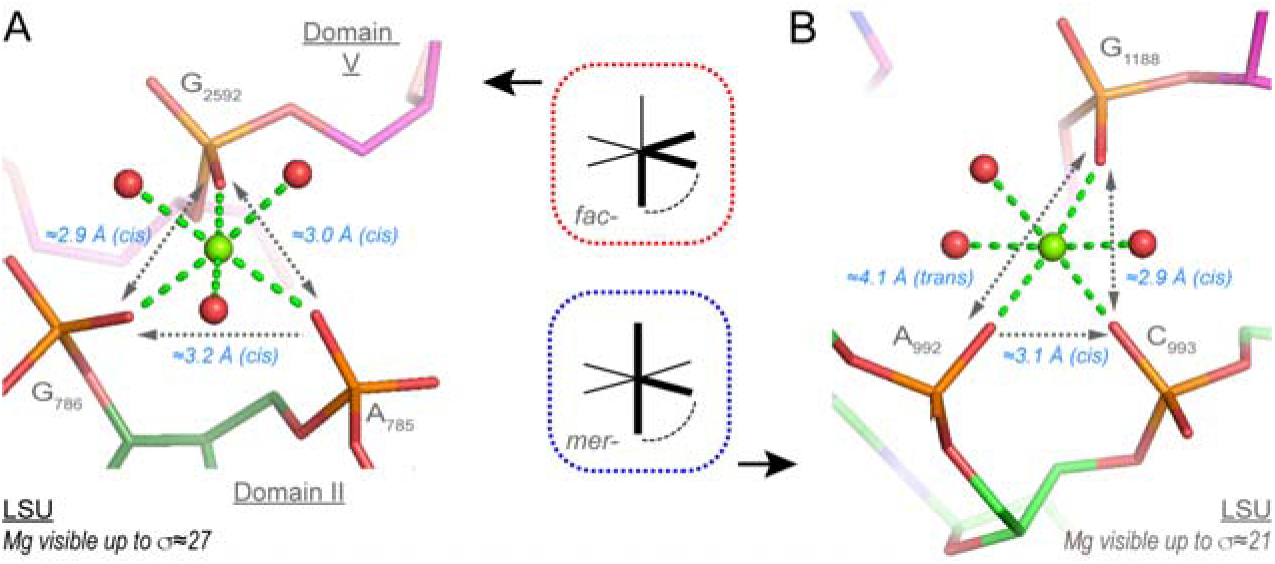
*fac-* and *mer-*3O_ph_.3O_w_ binding sites. (**A**) A *fac-*3O_ph_.3O_w_ ion joins domain “II” and “V”. (**B**) A view of a domain “II” *mer-*3O_ph_.3O_w_ ion. In both instances, a bidentate Mg^2+^ clamp is associated with a distant phosphate group. Green/cyan lines mark distances < 2.3 Å and in the 2.6–3.2 Å range. Experimental densities and some water molecules were hidden.

**Figure 10.**
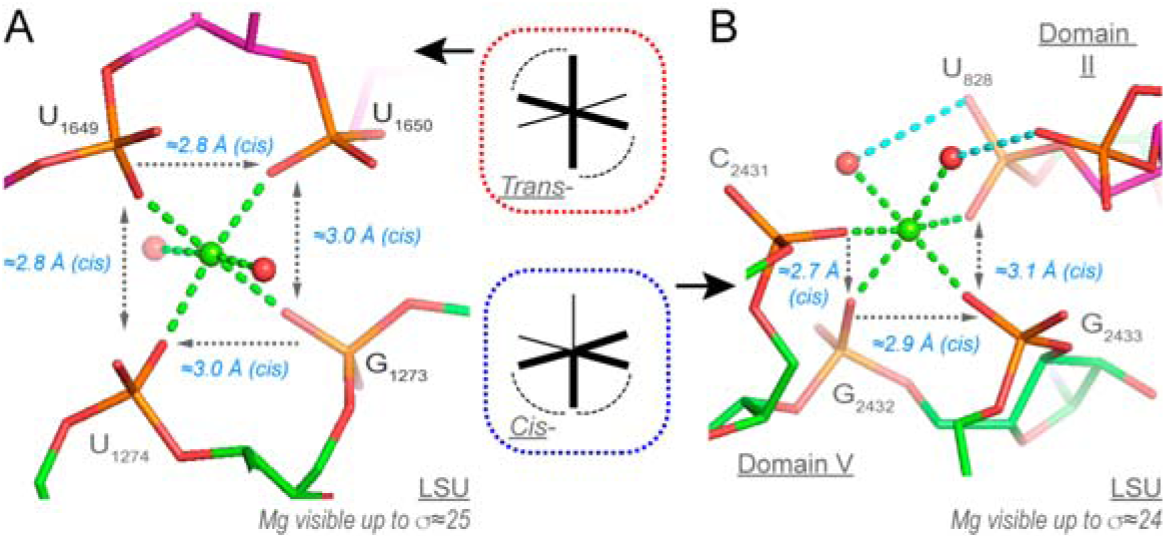
*<u>Trans</u>-* and *<u>Cis</u>-*2O_ph_.4O_w_ binding sites. (**A**) This “domain III” *<u>Trans</u>-*2O_ph_.4O_w_ ion involves two bidentate Mg^2+^ clamps. (**B**) A *<u>Cis</u>-*2O_ph_.4O_w_ ion joining domain “II” and “V” (the carbons are in green and purple, respectively). The two consecutive bidentate Mg^2+^ clamps form a rare pyramid motif described in (30) that involves three adjacent nucleotides. Green/cyan lines mark distances < 2.3 Å and in the 2.6–3.2 Å range. Experimental densities and some water molecules were hidden.

These intricate motifs mostly based on Mg^2+^ bidentate chelation are rare given the complexity of the folds and the difficulty in dehydrating multiple times a hexahydrated Mg[H_2_O] ^2+^ ion. Such motifs can only emerge in large RNAs where the nucleotide chain can fold back on itself as in for instance the ≈440 nucleotide long P4-P6 group I intron structure (PDBid: 8tjx; res. 2.44 Å) where a *<u>Trans</u>*- 4O_ph_.2O_w_ motif involving nucleotides (n, n+2, n+3, n+4) is observed (70). It is expected to observe additional variations of these motifs in other RNA structures. Variations of these patterns are discussed in the **SI**.

### Rare inner-shell Mg^2+^ binding to O2’ atoms

Three “well-defined” Mg^2+^…O2’ contacts were characterized (15). Two of those involve a bidentate contact to OP1/OP2 atoms where the O2’ atom is at 3.0/2.8 Å from the OP1/OP2 atoms of the next residue (**Figure 11**). Only two O2’(n)…OP1/OP2(n+1) contacts of the Mg^2+^…O2’-C2’-C3’-P- OP…Mg^2+^ (a six-membered ring) were localized in the entire 70S suggesting that these conformations cannot exist without the participation of Mg^2+^. The first binding type is *<u>Cis</u>-*3O_ph_.O_r_.2O_w_. It is the only binding type involving a chain of four consecutive nucleotides (**Figure 11A**). The second is *fac-*2O_ph_.O_r_.3O_w_ and comprises three consecutive nucleotides (**Figure 11B**). The third contact of the O_r_.5O_w_ type needs to be validated through the analysis of larger structural ensembles (**Figure S4C**). Overall, these occurrences imply that in a specific but rare context, O2’ atoms can coordinate to Mg^2+^. We note that two of these motifs induce turns associated with consecutive nucleotides that are probably impossible to achieve, like bidentate binding in general, without the involvement of a Mg^2+^ ion. None of the two involved O2’ atoms appear to be deprotonated; one of them is hydrogen bonded to a (C)OP2, the other to a (G)N7 atom.

**Figure 11.**
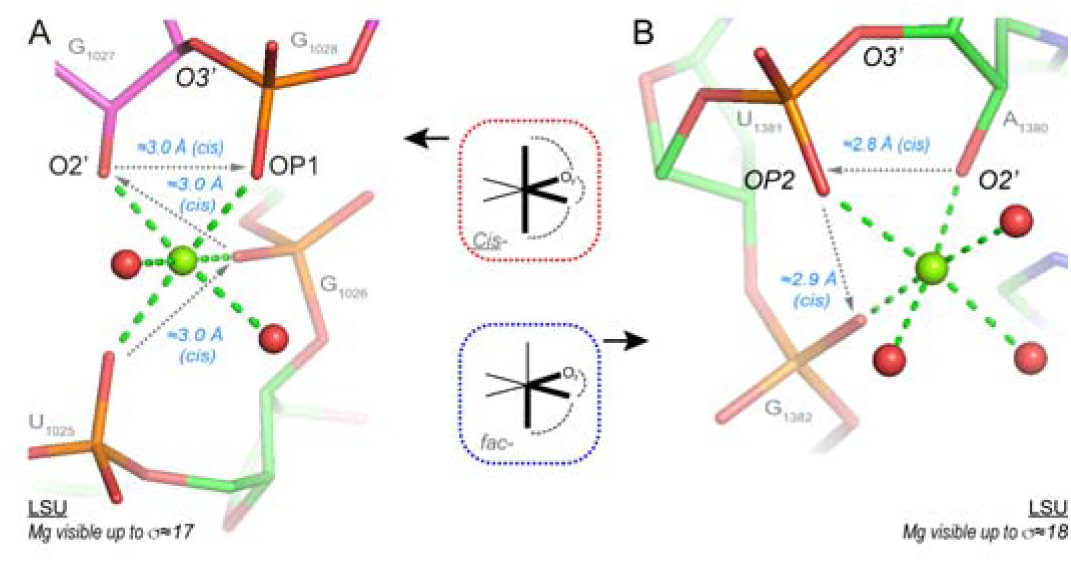
Two rare O2’…Mg^2+^ contacts. (**A**) This domain ”II” *<u>Cis</u>-*3O_ph_.O_r_.2O_w_ ion involves a turn formed by four consecutive nucleotides and a rare backbone conformation leading to a (n)O2’…(n+1)OP1 distance close to 3.0 Å. (**B**) This domain “III” *fac-*2O_ph_.O_r_.3O_w_ ion involves a turn formed by three consecutive nucleotides and a rare backbone conformation leading to a (n)O2’…(n+1)OP2 distance close to 2.8 Å. The two conformations involving OP1 and OP2 atoms are different. None of the O2’ atoms appear deprotonated. Green/cyan lines mark distances < 2.3 Å and in the 2.6–3.2 Å range. Experimental densities and some water molecules were hidden. See also **Figure S4C**.

### Anionic base pairs in 8b0x

According to a recent study (71), at least four anionic base pairs are present and conserved in bacterial ribosomes. In 8b0x, two and one cWW U(-)•G pairs are in the SSU and LSU, while a fourth cWW G•G(-) pair is in the LSU (**Figures 12/S5**). These pairs involve a deprotonated uridine/guanine nucleotide that carries a negative charge.

**Figure 12.**
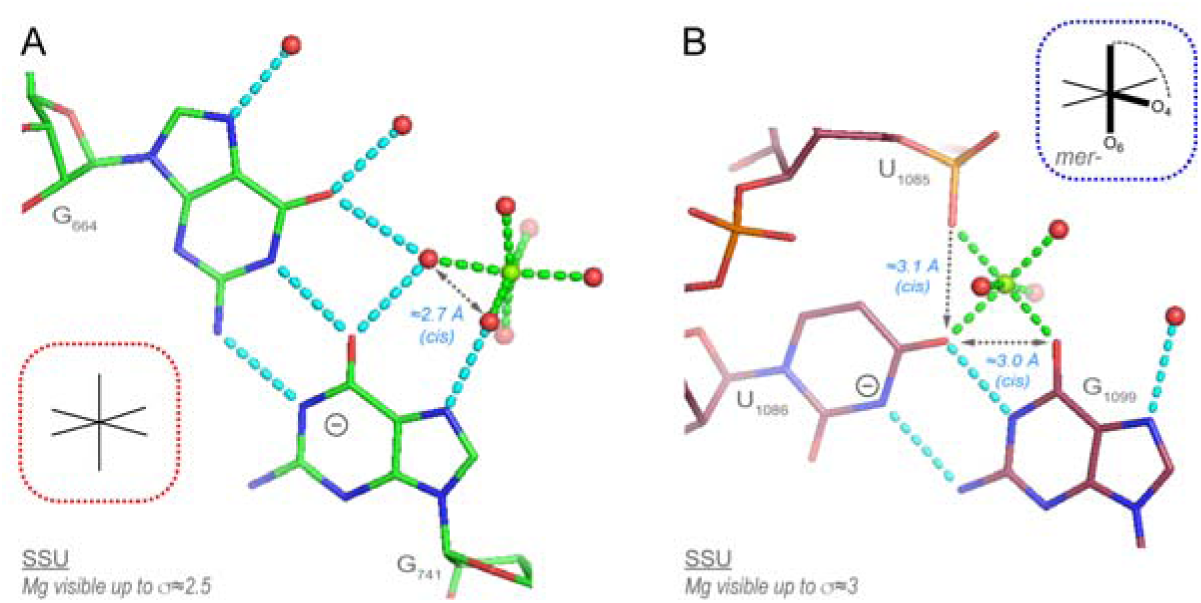
Two Mg^2+^ bound base pairs with a negatively charged guanine/uridine nucleobase. (**A**) The first G•G(-) pair involves a negatively charged guanine. The 6O_w_ ion binds to a guanine Hoogsteen edge Figure 5). (**B**) The U(-)•G pair involves a negatively charged uridine and is associated with a unique Mg^2+^ binding pattern of the *mer-*O_ph_.2O_b_.3O_w_ type. Green/cyan lines mark distances ≤ 2.3 Å and in the 2.6–3.2 Å range. Experimental densities and some water molecules were hidden. See also **Figure S5**.

One of these U(-)•G pairs displays a unique Mg^2+^ binding pattern of the *mer-*O_ph_.2O_b_.3O_w_ type (**Figures 12B**). These pairs related to negatively charged nucleobases are unique in the ribosomal ecosystem. We note the occurrence of an [A^+^•C]•ho5C base triple (**Figure S5**) associated with the ho5C base modification that benefits *E. coli* under oxidative stress conditions (72).

### Unexplained “metal” binding patterns

While most well-ordered 8b0x ion binding sites could be assigned to Mg^2+^/K^+^ ions, two of the unassigned density peaks could not be associated with these ions (**Figure 13**). In the SSU, an unidentified ion links the two N7 atoms of an “ion-mediated” A…G pair (**Figure 13A**). This position with a density peak distant by ≈2.1 Å from the two N7 atoms can only be occupied by a tetrahedral ion such as Zn^2+^. However, there is no clear-cut evidence for a Zn^2+^ assignment. Therefore, we placed a UNX ion with a 2N_b_.2O_w_ coordination in our model, UNX being the PDB code for an unknown atom/ion.

**Figure 13.**
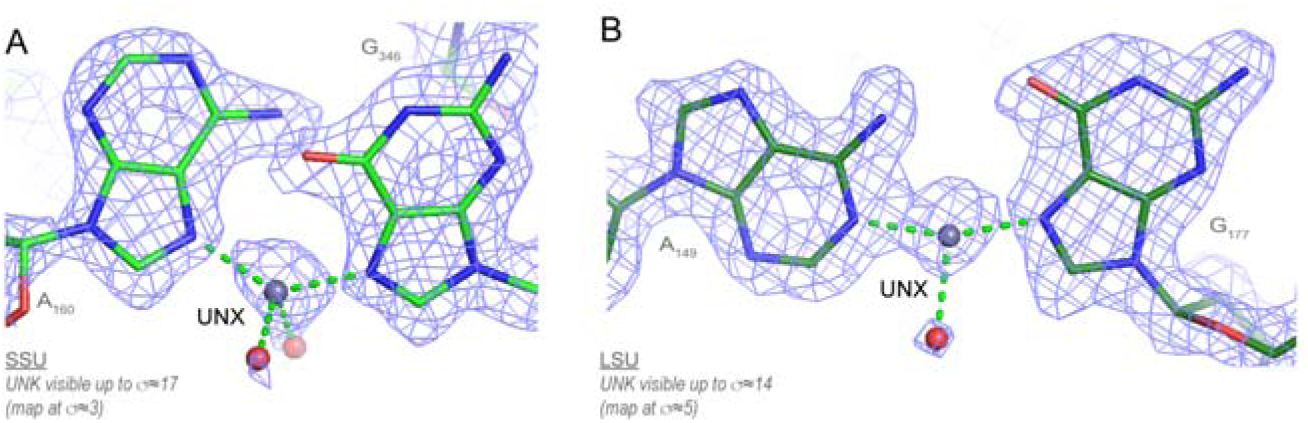
Unexplained ion binding patterns. (**A**) A UNX atom was placed in the density linking the two N7 atoms of an “A…G” pair that was left empty in 8b0x. UNX corresponds to the PDB code for unknown atom/ion. (**B**) Similarly, a UNX ion was placed in the density linking the (A)N1 and the (G)N7 atoms that was left empty in 8b0x. It is possible that these UNX ions are Zn^2+^ ions. Green lines mark distances < 2.3 Å.

In the LSU, another “ion-mediated” A…G pair is observed. A density peak occurs between the (A)N1 and the (G)N7 atoms with the A and G bases being strictly coplanar (**Figure 13B**). This pattern resembles to that of metal-mediated base pairs containing Ag, Hg or Au ions (41,73–75). Such “ion-mediated” base pairs have not been observed in the 7k00 *E. coli* ribosome used to build the 8b0x model. In the later structures, the adenine is shifted to establish a (A)N6…N7(G) hydrogen bond. These unusual base pairs suggest the presence of metal contaminants in 8b0x that are not present in other ribosomal structures or of excess Zn^2+^ ions that are difficult to identify through conventional techniques (76). We mentioned above and elsewhere the possible association of Zn^2+^ ions with the three characterized 2N_b_.4O_w_ motifs (16). This hypothesis has neither been confirmed nor invalidated. Noteworthy, the inclusion of molecules not added to the buffer are sometimes observed in cryo-EM structures as is the case for two polyamines mentioned in a recent 1.9 Å resolution *Human* 80S structure (77).

### Mg^2+^…Mg^2+^ ion pairs

An intriguing feature of the ribosomal ionic shell is related to the occurrence of numerous Mg^2+^…Mg^2+^, Mg^2+^…K^+^ and K^+^…K^+^ ion pairs. Although similar ion pairs have been observed in nucleic acid and protein systems (62,78–80), their characteristics were rarely discussed unless located in catalytic sites. In this study, an ion pair is defined by two metals sharing at least one and up to three phosphate groups, oxygen atoms, or water molecules. In 8b0x, none of these pairs are associated with r-proteins and we note that all the involved Mg^2+^ ions are hexacoordinated as expected.

Here, we propose a preliminary classification of Mg^2+^…Mg^2+^ pairs. Their limited occurrence in 8b0x suggests that more configurations remain to be uncovered through the exploration of additional RNA systems. Presently, we divided the observed Mg^2+^…Mg^2+^ pairs into four categories called micro- clusters or Mg^2+^-μc’s (2,6,42,81). The MgA_μc_ involves two Mg^2+^ ions that are separated by > 4.5 Å and are bridged by the same phosphate group. The MgB_μc_/MgC_μc_/MgD_μc_ clusters display inter-metal distances in the 2.4–3.8 Å range. In these groups, the metals can share from one to three oxygen atoms and/or phosphate groups. To differentiate them, we added a 1 to 3 suffix. For instance, the MgA_μc_ seen **Figure 14A** is named MgA_μc2_ since the Mg^2+^ ions share two PO_4_ groups. This nomenclature applies to Mg/Mg pairs as well as to Mg/K pairs.

- <u>Type I “Magnesium micro-clusters” (MgA_μc_):</u> Mg^2+^…OP-P-OP…Mg^2+^ with *d*(Mg^2+^…Mg^2+^) ≈4.5–5.3 Å (17 “well-defined” occurrences). In these MgA_μc_, the ions share one or two phosphate group(s) (**Figure 14A/S6A**). These arrangements were previously called Mg^2+^-μc (2,6,42,81) and resemble the bridges formed by carboxylate groups between two metal ions in enzymatic systems (62). In rRNA, such MgA_μc_ were considered to be part of the peptidyl transferase center of ribosomes from all kingdoms and were also found in the P4-P6 domain of the tetrahymena group I intron ribozyme (82) and the self-spliced group II intron from *Oceanobacillus iheyensis* (83) among other RNA structures. Most of these MgA_μc_ shares a common phosphate group while only two of them share two common phosphate groups (**Figure 14A**).
- <u>Type II (MgB_μc_):</u> Mg^2+^…O_w_…Mg^2+^ with *d*(Mg^2+^…Mg^2+^) ≈3.8 Å (2 “well-defined” occurrences). In the 1^st^ pair, the ions share a single water molecule and no RNA atoms (**Figure 14B**). The 2^nd^ pair involves one shared water molecule. In addition, two Mg^2+^ connected waters are separated by 2.07 Å suggesting the presence of an alternate conformation that must be interpreted with caution (**Figure S6B**).
- <u>Type III (MgC_μc_):</u> Mg^2+^…O_w_…Mg^2+^ with *d*(Mg^2+^…Mg^2+^) ≈2.8 Å (2 “well-defined” occurrences). Both ions of a 1^st^ pair share an OP1 atom and two water molecules (**Figure 14C**) while the two ions of a 2^nd^ pair share three water molecules (**Figure S6C**). These instances recall a documented example of an Mg^2+^…Mg^2+^ ion pair where the ions are separated by 2.7 Å (80). This pair appears in a structure of a 5S rRNA Loop E fragment at 1.5 Å resolution (**Figure S6D**). Earlier, we proposed that such an arrangement resulted from alternate conformations of two slightly offset Mg^2+^ binding patterns (84). However, considering the two identified MgC_μc_ pairs in 8b0x, we re-examined this hypothesis. It seems that in specific environments, *d*(Mg^2+^…Mg^2+^) ≈2.8 Å distances are stable. This is supported by a recent CSD structure containing such a Mg^2+^ pair (85), by the description of inorganic compounds involving stable Mg(I)…Mg(I) covalent bonds with *d*(Mg^+^…Mg^+^) ≈2.8 Å and by the implication of Mg pairs in enzymatic systems (86,87).
- <u>Type IV (MgD_μc_):</u> Mg^2+^…O_w_…Mg^2+^ with *d*(Mg^2+^…Mg^2+^) = 2.38 Å (1 “well-defined” occurrence). Here, the proximity of the two Mg^2+^ ions is astonishing (**Figure 14D**). This ion pair involves three bridging water molecules associated with surprisingly well-defined density patterns and resembles the ones described in **Figures S6E/F**. The first Mg^2+^ is of the *cis*-2O_ph_4O_w_ type; the second Mg^2+^ involves a deformed 6O_w_ coordination octahedron. Another Mg pair with *d*(Mg^2+^…Mg^2+^) ≈2.42 Å has been described based on a 1.52 Å resolution nuclease structure, although one of the ions is not hexacoordinated (88). Additional independent observations need to confirm the existence of ion pairs with *d*(Mg^2+^…Mg^2+^) <2.50 Å.

**Figure 14.**
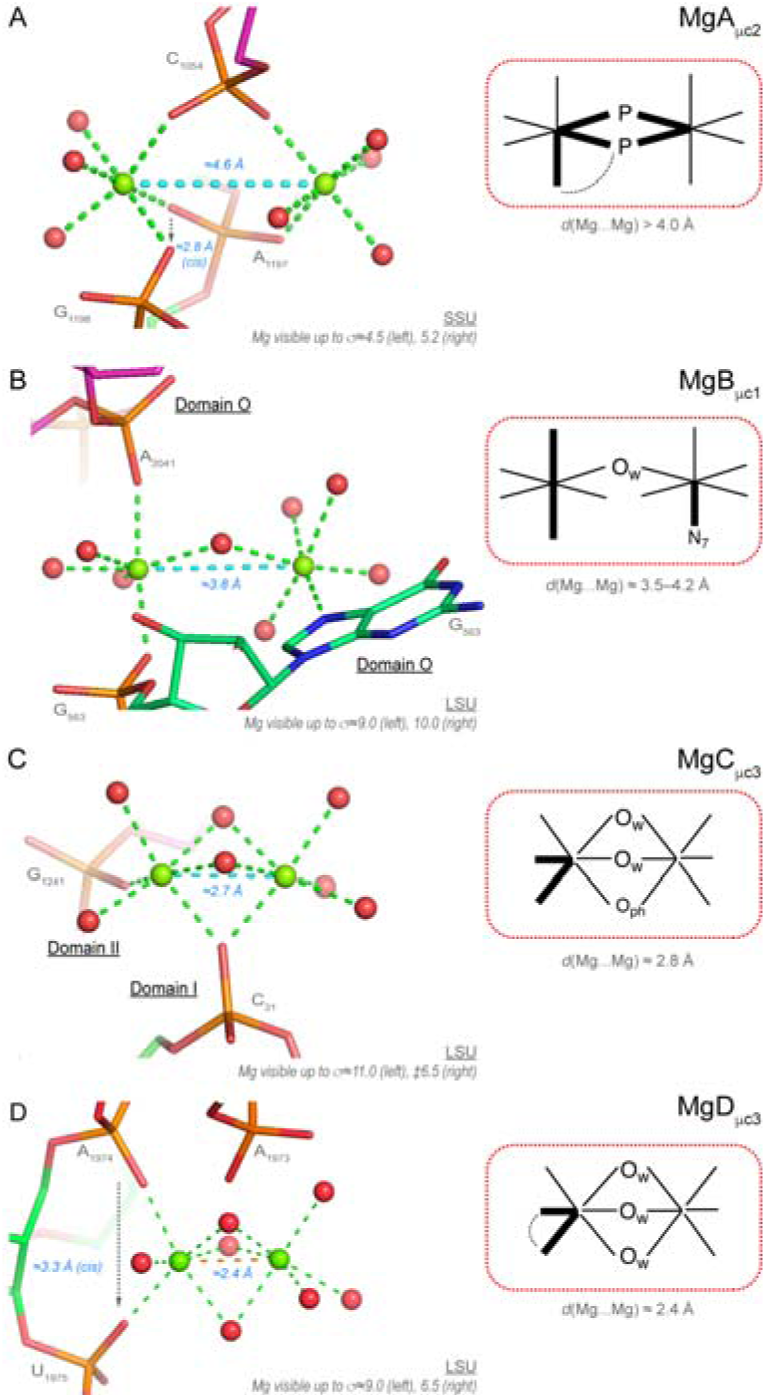
The four Mg^2+^…Mg^2+^ pair types. (**A**) The two bottom phosphate groups of this MgA_μc2_ pair belong to adjacent domain “3’C” nucleotides. The first ion is *fac-*3O_ph_.3O_w_, the second is *cis-*2O_ph_.4O_w_. (**B**) This MgB_μc1_ ion pair is located in the central domain “O”. This ion pair is connected to the OP2 and N7 atoms of a same guanine and involves a bridging water molecule. The first ion is *trans-*2O_ph_.4O_w_, the second is N_b_.5O_w_. (**C**) This MgC_μc3_ ion pair, that joins LSU domains “I/II”, involves two bridging waters and an O_ph_ atom in a regular arrangement associated with a short 2.7 Å inter-metal distance (see also **Figure S6**). The first ion is *cis-*2O_ph_.4O_w_, the second is O_ph_.5O_w_. (**D**) In the absence of corroborating examples, this domain “IV” MgD_μc3_ ion pair must be considered with caution. The first *cis-*2O_ph_.4O_w_ ion forms a bidentate phosphate clamp, the second is 6O_w_. Green/cyan lines mark distances < 2.3 Å and in the 2.6–3.2 Å range. Orange lines mark the inter-metal distance Experimental densities and some water molecules were hidden.

### Mg^2+^…K^+^ ion pairs (28 “well-defined” occurrences among 42)

Given the exceptional 8b0x resolution, over 200 binding occurrences of K^+^ ions to rRNA atoms were characterized. Among those, 28 “well-defined” Mg^2+^…K^+^ pairs with *d*(Mg^2+^…K^+^) ≈3.5–4.4 Å were identified. In many instances, the K^+^ ions share two Mg^2+^ water molecules (**Figure 15A**). A subcategory of these pairs involves 10 occurrences of a K^+^ ion bound to the deep groove of G•U pairs.

**Figure 15.**
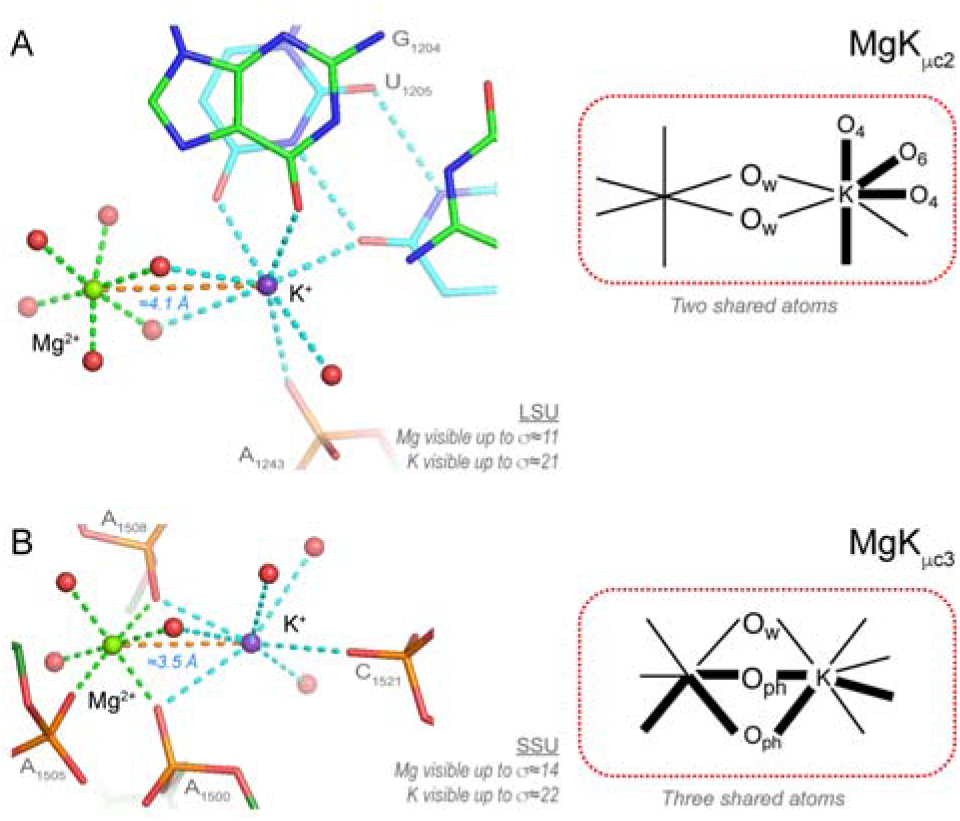
Examples of Mg^2+^…K^+^ pairs of type I and II. (**A**) This domain “II” MgK_μc2_ pair involves a Mg^2+^ of the 6O_w_ type and a heptacoordinated K^+^ that binds to the deep groove of a domain “II” G•U pair (carbon atoms in cyan). Two water molecules are bridging the ions. (**B**) This domain “III” MgK_μc3_ pair displays a shorter inter-metal distance. It involves a Mg^2+^ of the *mer-*3O_ph_.3O_w_ type and a heptacoordinated K^+^ (for density representations, see **Figure S7**). Three water molecules are bridging the ions. Green/cyan/orange lines mark distances ≤ 2.3 Å, in the 2.6–3.2 Å range and > 3.2 Å. Experimental densities and some water molecules were hidden.

Seven of these pairs involve a Mg^2+^ of the 6O_w_ type. A less frequent pattern where three oxygen atoms are shared by both hydration shells involve shorter inter-metal distances (**Figure 15B**). In all instances, the K^+^ density peak is significantly higher than that of the Mg^2+^ ion and of their surrounding water molecules (**Figure S7**). This suggests that examining the peak heights of solvent molecules in cryo-EM structures has some value, especially when the Mg^2+^/K^+^ ions are close in space.

### Mg^2+^ ions are rarely present at the SSU/LSU interface

Among the 408 Mg^2+^ ions in 8b0x, only a single O_ph_.5O_w_ Mg^2+^ ion contacts both subunits (**Figure S4A**). This suggests that Mg^2+^ ions are rarely present at the SSU/LSU interface and are therefore not of structural or functional importance for the subunit assembly. The presence of a single Mg^2+^ ion at the SSU/LSU interface might result from the elevated experimental Mg^2+^ concentration. Examination of other high-resolution structures is needed to assess this observation. Overall, this suggests that destabilization of the 70S particles observed at low Mg^2+^ concentrations is the result of the destabilization of each of the subunits leading to a fuzzy structural interface (7,89). Conversely, the association of ribosome particles forced by Mg^2+^ concentrations above 15 mM should result from subunit stabilization rather than from the presence of interfacial Mg^2+^ ions (3).

### Mg^2+^ bound to r-proteins are uncommon

Only one poorly defined *cis-*3O_coo_.2O_w_.N_His_ binding site buried in uS2 was observed (**Figure S4B**). This binding site is also present in the parent 8fto structure. No other r-protein binding site was characterized suggesting that r-proteins might have evolved to avoid competition for Mg^2+^ resources with rRNA. Overall, the need for Mg^2+^ to support proper r-protein functions is significantly smaller than that required by RNA. We note that this motif involves a “carboxyl clamp” similar to the phosphate clamps shown **Figure 7**. This suggests that that the binding of Mg^2+^ to proteins adopts rules similar to those described for RNA.

### Mg^2+^ bound at the rRNA/r-protein interfaces

More Mg^2+^ ions are observed at rRNA/r-protein interfaces (**Table S1** and **SI**). Five of these ions make direct contacts to rRNA and r-proteins while others form at least one direct contact to the rRNA with water-mediated contacts to the -proteins.

Eight ions are located at the LSU/uL2 interface of the peptidyl transferase center (PTC) that gathers the largest number of observed contacts (10,42,81). Petrov *et al.* noted the conservation of a Mg^2+^-μc at this interface that involves a N-terminal 18 amino acid long loop and discussed the importance of the Ala-Met-Asn sequence in the ribosomal assembly process (**Figure 16**). This interface is conserved in Archaea, Bacteria, and Eukarya (81). Note that the involved Mg^2+^ ions establish direct contacts through the binding of MgA_μc_’s to RNA phosphate groups but they form only water-mediated contacts to uL2. This suggests that Mg^2+^ must first shape the local RNA folds so that uL2 can bind later. A more detailed discussion of the rRNA/r-protein interfaces is beyond the scope of this paper.

**Figure 16.**
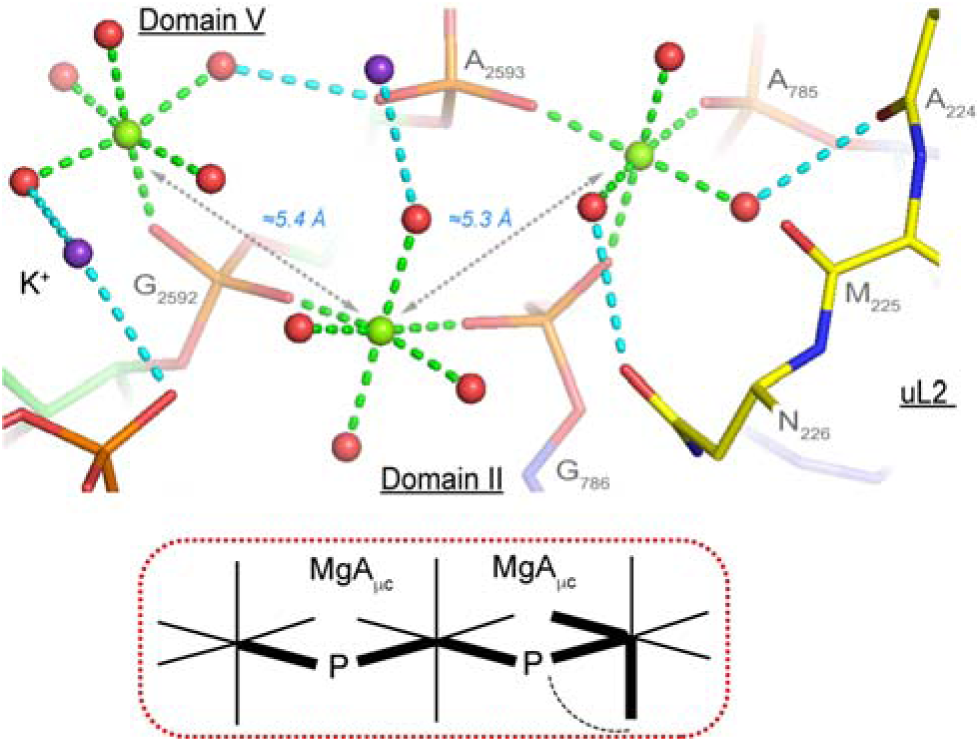
Partial view of the LSU/uL2 interface centred on two MgA_μc_’s. This view emphasizes the intricacy of the ion-binding patterns at the LSU/uL2 interface that involves the MgA_μc_ named D2 by Williams and co-workers (6,42,81). These ions link the LSU domains “II/V” with a N-terminal uL2 segment (yellow backbone). From this view, it can be hypothesized that the Mg^2+^ ions first establish a link between the two distant domains “II” and “V”. When this connection is established, uL2 binds and the LSU comes closer to its functional state. Note the presence of K^+^ ions (purple) at this interface. Green/cyan lines mark distances < 2.3 Å and in the 2.6–3.2 Å range. Experimental densities and some water molecules were hidden.

### Mg^2+^ linking intra-LSU domains

Twenty “well-defined” out of twenty-one Mg^2+^ ions coordinate with LSU domains through a combination of direct and water-mediated contacts (**Table S2** and **Figures 7/9/10/11/14/16**). A few of them involve Mg^2+^-μc’s as previously described (6,42,81). Interestingly, inter-domain contacts are not observed in the SSU suggesting a less complex folding process. Among the characterized intra-LSU links, seven of them involve domain “II/V”, none involve domain “VI” and only two are associated with domain “III”. This suggests that the concomitant binding of domain “II/V” as well as the interaction between domain “V” and r-protein uS2 are important Mg^2+^-dependent biogenesis steps (**Figure 16**) as discussed by Williams and colleagues (6,42,81).

### Description of a universal set of rules to identify Mg^2+^ binding sites

In this work, we inferred straightforward binding rules from the examination of the amended 8b0x structure. Next, we describe step by step these rules starting from those related to the detection of critical Mg^2+^ bidentate motifs.

Locating Mg^2+^ bidentate clamps (O_ph_…O_ph_): A unique feature of Mg^2+^ binding clamps is associated with a conserved O_ph_…O_ph_ distance close to ≈2.9±0.2 Å that strictly matches that separating Mg^2+^ water molecules in *cis-* (see **Figures 3/7/9/10/11/14**). Thus, we hypothesized that a simple method to locate Mg^2+^ bidentate clamps is to search for all O_ph_ pairs with *d*(O_ph_…O_ph_) < 3.4 Å. By discarding the poorly modelled pairs comprised of at least one O_ph_ atom with B-factors > 60 Å^2^, we located a total of 94 pairs (87 “well-defined” ones) comprising 36 OP1…OP1, 23 OP2….OP2 and 35 OP1…OP2 combinations. We observed that all 94 pairs were associated with Mg^2+^, 92 of them forming Mg^2+^ bidentate clamps. The two remaining motifs display a binding pattern involving a distinctive Mg^2+^ water-mediated contact stabilizing the short *d*(O_ph_…O_ph_) ≈2.9 Å distance (**Figure S2**). We propose that the latter configurations could correspond to intermediates in the formation of Mg^2+^ bidentate clamps.

We concluded that all *cis-*bidentate motifs involving a Mg^2+^ ion can be detected in rRNA fragments of appropriate resolution by collecting O_ph_…O_ph_ distances ≤ 3.4 Å. Obviously, all “higher-order” patterns of the *fac-/mer-/<u>Cis</u>-/<u>Trans</u>-* type can also be localized by checking these distances since most of them involve at least one Mg^2+^ *cis-* bidentate motif (the only exceptions being the less frequent motifs in *trans-*; see **Figure 7C**). As a result, the 94 uncovered *cis-*bindings involve a limited number of 68 “well-defined” Mg^2+^ ions that are all associated with essential rRNA folding patterns.

To emphasise even further the effectiveness of this stereochemical rule, we note that by just checking the O_ph_ pairs with *d*(O_ph_…O_ph_) ≤ 3.4 Å, we could identify at least 7 new Mg^2+^ bidentate motifs not assigned in the initial 8b0x structure. This illustrates the ability of this stereochemically based process to characterize Mg^2+^ binding sites that escaped from the experimentalist’s attention. It also suggests that this criterion is universal and can be applied to the search of major Mg^2+^ binding sites in all RNA systems as well as in proteins where the main Mg^2+^ ligands are carboxylate groups (see **Figure S4B**).

#### Locating *trans-*2O_ph_.4O_w_ motifs and their variations

This is probably the most difficult exercise given the ≈4.1 Å distances separating the O_ph_ atoms that overlap with inter-phosphate distances associated with other motifs (**Figure 3/7C**). However, this distance criterion might accelerate the identification of *trans-* patterns.

#### Locating 2O_b_.4O_w_ and O_ph_.O_b_/N_b_.4O_w_ motifs

The 2O_b_.4O_w_ binding type is rare. We found just one “poorly-defined” occurrence involving distant residues. In relation to this observation, it seems that Mg^2+^ is not an ideal ion for bridging O_b_ atoms from stacked base pairs or from consecutive nucleotides in opposition to the monovalent ions such as Na^+^/K^+^ (15). Similarly, only two “well-defined” O_ph_.4O_w_.N_His_ occurrences involving r-protein N_His_ atoms and no O_ph_.N_b_.4O_w_ binding types could be observed. Yet, the slightly more frequent O_ph_.O_b_.4O_w_ binding types can be located by monitoring *d*(O_ph_…O_b_) ≤ 3.4 Å distances that can also be of the “intra-nucleotide” type.

#### Locating 2N_b_.4O_w_ motifs

In most instances, although the stacking distance between two consecutive nucleotides is around ≈3.4 Å, the distance between two stacked purine N7 atoms is often > 3.8 Å, therefore excluding a bidentate binding to a Mg^2+^ ion. We found that in 8b0x the only nucleobase arrangement where N7 atoms are separated by less than 3.4 Å are analogous to the ones shown **Figures 8/S3**. Indeed, only three “head-to-tail purine stacks” were detected in 8b0x that bind Mg^2+^ ions or Zn^2+^ as suggested elsewhere (16,54). Therefore, searches for *cis-*2N_b_.4O_w_ binding patterns involving consecutive purines can be avoided in favour of searches for those involving non-consecutive stacked nucleotides.

Locating Mg^2+^ of the O_ph_/O_b_/N_b_.5O_w_ type: Locating these binding types is more difficult but the coordination geometry of the Mg^2+^ ions provides some hints as shown in **Figure 6B** and (15,16). For O_ph_.5O_w_, the Mg^2+^ ion must roughly be in the OP-P-OP plane and the P-OP…Mg^2+^ angle must be in the 120–160° range. For the O_b_/N_b_.5O_w_ variants, the Mg^2+^ ion must be in the nucleobase plane. Yet the angle constrains are slightly different with an average (C=O…Mg^2+^) angle of ≈144°. In this binding mode, the ion is approximately in the alignment with one of the O_b_ lone pairs as confirmed by recent simulation data (65). For N_b_.5O_w_ types on the other hand, Mg^2+^ ions are aligned along the N7 lone pair.

Locating Mg^2+^ of the 6O_w_ type: Defining binding rules for the less tightly bound 6O_w_ ions is challenging. Like for O_ph_/O_b_/N_b_.5O_w_, the Mg^2+^ ions of the 6O_w_ type that bind strongly to the rRNA establish a high number of water-mediated contacts. Some of these ions are associated with important structural folding “wedges” (**Figure 5A**). The fact that these Mg^2+^ ions use their first shell water molecules to bind to appropriate rRNA binding sites such as the guanine O6/N7 and the OP1/OP2 atoms suggests to scan the rRNA hydration shell to locate potential 6O_w_ binding sites. Indeed, a precise knowledge of the rRNA hydration shell will be helpful to locate the most important water-mediated Mg^2+^ binding spots. These can be derived from experimental structures and MD simulations for protein and DNA systems (46,90–93).

## DISCUSSION

### Stereochemistry is an efficient tool for assigning Mg^2+^ ions

Here, we revised the 8b0x structure and completed its solvation shell. The amended 8b0x structure, provides insights that could not be derived with the same level of accuracy from lower resolution structures. We confirmed the previously described Mg^2+^ binding patterns to O_b_/N_b_ atoms (10,15,16,28) and added a comprehensive description of Mg^2+^ binding to O_ph_ atoms. Our findings that establish a clear-cut classification of all the Mg^2+^ binding types and occurrences gathered from our amended model are summarized in **Figure 4**.

An important result of this study on the characterization of stereochemical Mg^2+^ binding principles appears to be surprisingly simple. We found that Mg^2+^ binding induces folding patterns where rRNA oxygens are separated by an average distance of 2.9 Å (**Figure 4**). By considering this principle, we observed that in every occurrence where *d*(O_ph_…O_ph_) ≤ 3.4 Å, the two O_ph_ atoms are linked by a Mg^2+^ through inner-shell contacts or for two of them through inner- and outer-shell contacts (**Figure S2**). Hence, we infer that in all instances, a Mg^2+^ ion is needed to stabilize configurations with short inter- O_ph_ distances. Indeed, given the proximity of two to four O_ph_ atoms, these configurations are associated with the foremost electronegative locations in RNA structures. The stabilization of such configurations by Na^+^/K^+^ ions with smaller charge density has not been observed in this ribosome nor elsewhere (69).

We estimate that by using this distance criterion, we reach ≈100% accuracy in the detection of *cis-* bidentate binding sites, a percentage that needs to be confirmed by scanning a larger variety of RNA structures. We also uncovered two novel *cis*-O_ph_.O_r_ motifs involving O2’ atoms, which to the best of our knowledge, have not been documented elsewhere. They involve a specific backbone conformation that allows the O2’ atom to be at ≈3.0 Å from an O_ph_ atom of a consecutive nucleotide (**Figure 11**). Interestingly, this binding principle is also applicable to proteins where *d*(COO^-^…^-^OOC) < 3.4 Å are pointing to *cis-*2O_coo_.4O_w_ Mg^2+^ ions, as observed in uS2 (**Figure S4B**), and to drugs as observed for tetracyclines bound to bacterial ribosomes where a rare 3O_ph_.O_w_.2O_tetracycline_ binding mode is observed. It involves a coordination with two oxygens separated by 2.6 Å (50). However, the efficiency of this criterion might diminish for structures with lower resolutions where the phosphate groups have not been accurately modelled, resulting in potential Mg^2+^ binding sites with *d*(O_ph_…O_ph_) > 3.4 Å. Stereochemical considerations are also determining for characterizing O_ph_.5O_w_ and 6O_w_ binding sites that involve a correspondence between the Mg^2+^ hydration shell and that of local rRNA folds associated for instance with guanine Hoogsteen edges (**Figures 4/5**). Information related to the density peak heights of solvent particles has also to be considered during the solvent identification process. In general K^+^ ions display higher densities than Mg^2+^ ions and bound water molecules. However, we observed that some immobilized water molecules involved in water-mediated contacts display higher densities than those associated with the more distant Mg^2+^ ion. In such instances, stereochemistry is of great help for interpreting the local structural arrangements. Indeed, the interpretation of experimental data has to comply with stereochemical rules. Disagreements should be solved through additional experiments or changes in theoretical paradigms.

### Structural ion pairs in ribosomes

The characterisation of intricate ion pairs might shed light on some features of these motifs in ribosomal and non-ribosomal RNA systems and their catalytic sites (78,94–97). Indeed, since the proposal by Steitz of a two-metal-ion catalytic mechanism for RNA splicing and RNase P hydrolysis (98), the interest for catalytic sites has grown and has been extended to numerous RNA systems such as Group I and II introns as well as spliceosomes (99,100). K^+^ ions were also shown to be present in catalytic sites and participate in catalytic mechanisms (101,102).

However, the structural roles of Mg^2+^…Mg^2+^ ion pairs have been less explored with the exception of some MgA_μc_ occurring in ribosomes (2,6,42,81). In 8b0x, we identified and classified several types of Mg^2+^…Mg^2+^/K^+^ ion pairs and showed how they “assemble” to form extensive ion chains (**Figure 16**). This suggests that ion pairs, including those comprising monovalent ions (15), recurrently appear in RNA systems though the relation between their formation and the ionic strength of the surrounding media is not understood. In that respect, it is important to note that the +2 net charge on the Mg^2+^ ions that are in close proximity might significantly decrease. It has been suggested that the formation of ion pairs results in a redistribution of the ion electronic charge towards the bound phosphate groups (62) that amplifies polarization and charge transfer effects. This effect has already been investigated for Mg^2+^ ions not involved in ion pairs (2,103).

### Hierarchical Mg^2+^ dehydration during ribosome biogenesis

The most intricate ion binding pockets described herein correspond to motifs embedded in the core of the ribosome structure during biogenesis. The formation of such motifs needs a level of structural complexity that is probably not accessible to the smaller RNAs. For instance, Mg^2+^ does not establish direct contacts with phosphate oxygens in Watson-Crick helical structures unless non-canonical base pairs are present like in the ribosomal 5S loop E motif (38,80,84). Similarly, very few direct Mg^2+^ ions contacts with the ≈76 nucleotide long tRNAs as observed in crystal structures with resolutions < 2.0 Å. an exception being a *cis*-2O_ph_.4O_w_ observed in the D-loop of a tRNA^Phe^ structure (PDBid: 1EHZ; res. 1.93 Å (104)). More complex motifs emerge in larger RNAs such as for instance the ≈ 400 nucleotide long Group I introns (105,106). Based on these observations, we infer that structural complexity is a prerequisite to the formation of the higher-order Mg^2+^ binding sites.

The apparently simple Mg^2+^ binding classification we propose underlines a great diversity of motifs if one considers the combinatorial possibilities associated with the nature and relative positions of the coordinating atoms. It also suggests a hierarchical dehydration path associated with the formation of the observed binding motifs (**Figure 17**). The principles described in this study will certainly benefit groups involved in RNA folding competitions such as RNA-puzzle (107,108) given that the techniques that are currently used rarely use information related to water or ion coordination.

**Figure 17.**
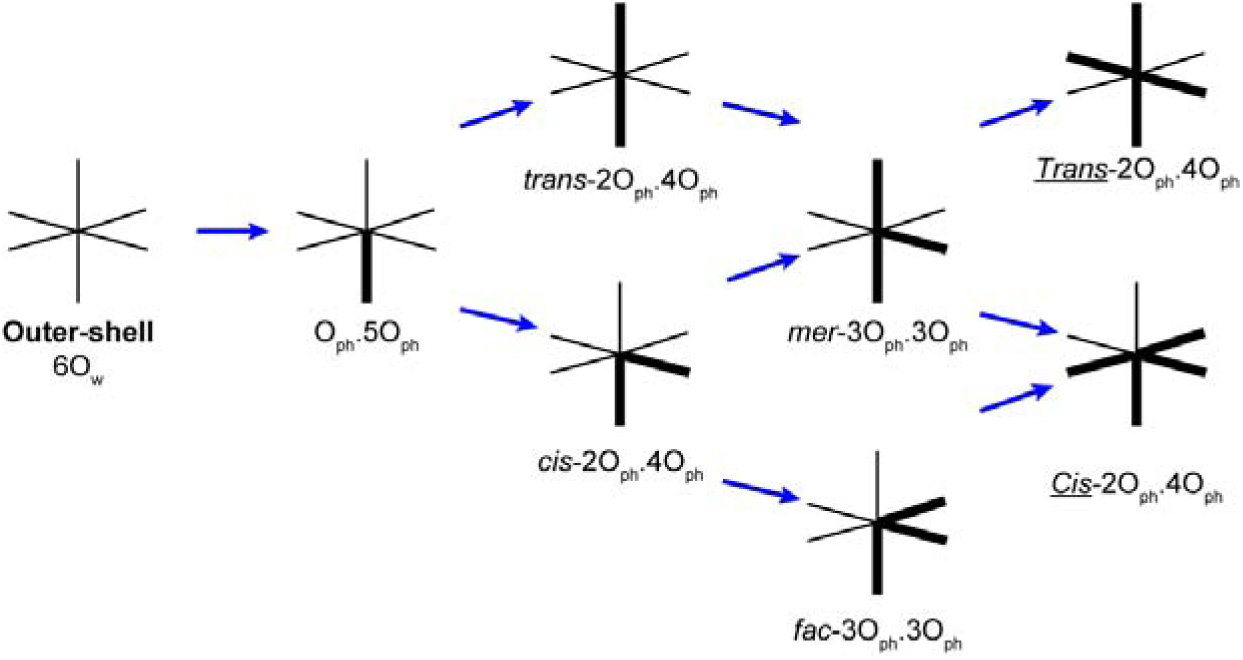
Sketch showing the hierarchy of dehydration of Mg^2+^ binding motifs. This sketch illustrates the path of going stepwise from an hexahydrated ion towards more complex motifs through various dehydration mechanisms.

### Considerations related to buffer and *in vivo* Mg^2+^ concentrations

Biochemical investigations suggested that optimal buffer conditions for ribosomes are 2–5 mM for Mg^2+^ and 60–150 mM for K^+^ when 2 mM of spermidine are present (3,7). Although the total Mg^2+^ concentration in the cell is 17–20 mM which is close to the 25 mM present in the 8b0x buffer, the concentration of non-chelated hexahydrated Mg^2+^ ions in a crowded intracellular environment is 1– 2 mM (109). The idea of an optimal Mg^2+^ concentration has been examined by several groups. The addition of weakly chelating agents to buffers was found to enhance RNA function and chemical stability as well as catalytic efficiency probably by sequestering an excess of divalent ions (109–111). Similarly, A low Mg^2+^ concentration affects RNA properties of some systems by preventing suitable folding whereas a high Mg^2+^ concentration reduces activity (112), in-line with results showing that excess Mg^2+^ reduces ribosomal translational activity and accuracy (7,113). Elsewhere, we speculated that excess Mg^2+^ might over-stabilize rRNA structures and grip the translation machinery by binding to the mRNA and tRNAs while a less organizing ion such as K^+^ would act as a lubricant favouring a smoother translation process thereby preserving an optimal turn-over rate (17).

Here, we infer that the optimal number of chelated Mg^2+^ comprise ≈100 *cis-* and *trans-* ions that are involved in the coordination of at least two non-water oxygen/nitrogen atoms and that most of them are incorporated during biogenesis. A proportion that remains to be determined of the ≈290 “O_ph_.5O_w_/6O_w_” ions that display limited or no access to the bulk solvent might similarly be encapsulated during biogenesis (**Figure 5A/6A**). Thus, it is not currently known if all the “O_ph_.5O_w_/6O_w_” ions are essential for ribosome activity or are the result of an excess of Mg^2+^ ions in the cryo-EM buffer. It is possible that the 1.55 Å resolution of this model is a byproduct of the high Mg^2+^ concentration that might have (over)saturated Mg^2+^ binding sites. Hence, one might wonder if the number of Mg^2+^ ions observed in 8b0x exceeds or not that necessary for a ribosome to perform its *in vivo* tasks (17,89,112).

To summarize, through a reinterpretation of the 8b0x structure, we were able to characterize the currently largest number of chelated Mg^2+^/K^+^ ions in a ribosome. Yet, the 634 Mg^2+^/K^+^ ions we could validate carry a total of ≈1,040 positive charges that is far from being sufficient to neutralize the ≈5,000 rRNA nucleotides. This remains true even if we consider the ≈525 excess positive charges carried by all the bound r-proteins or the ≈1,160 charges carried by all the positively charged Lys/Arg residues. For finding the “missing” ≈3,000 positive charges, theoretical methods such as MD simulations can be used (36–38,51,65,84,114–116).

### Amending existing structures

This study also raises the issue of amending existing PDB structures, a practice that should be better integrated in the structural community habits (19,117). It is clear that the 8b0x refinement process was not conducted to its end regarding the interpretation of its solvation shell structure. That is understandable given the amount of time needed to conduct such a task. Structures deposited to the PDB are a timely interpretation of the experimental data that can be corrected for flaws/omissions by using knowledge acquired after their deposition (23,118,119). A good illustration of this process is provided by the multiple corrections brought to the *Haloarcula marismortui* 50S structure first deposited to the PDB in 2000 and last refined in 2013 (9,14). In the present case, we provided significant improvements to the 8b0x bacterial ribosomal structure and suggest that a large subset of PDB structures could be amended by applying strategies like those described in this paper regarding their solvation shell and other important structural features.

## CONCLUSIONS

Through a careful analysis of the amended 8b0x bacterial ribosome structure, we analysed Mg^2+^ binding modes and derived Mg^2+^ binding principles based on simple stereochemical rules. The rules described herein should allow developing more accurate solvation shell models in small and large RNA. We provide also a complete description of all the major Mg^2+^ chelation sites that involve 2 to 4 non-water coordinating atoms. This description of intricate Mg^2+^ binding motifs comprising ion pairs will certainly enhance our views about RNA folding, structure and catalysis (120) while opening doors for new RNA design strategies involving Mg^2+^ ions. They will also facilitate RNA 3D structure prediction.

## SUPPLEMENTARY DATA

Supplementary Data are available at NAR online.

## Supporting information

SI_ion_diagnosis

SI_Excel_summary

## ACKNOWLEDGEMENT

P.A. wishes to thank Dr Yaser Hashem, Dr Quentin Vicens and Prof. Eric Westhof for comments on the manuscript as well as Dr Nawavi Naleem for computational assistance.

## FUNDING

This work was supported by the “Centre National de la Recherche Scientifique (CNRS)”. S.K. and A.H.K are supported by NYUAD AD181 faculty research grant and REF Grant RE317.

## CONFLICT OF INTEREST

None

## Notes

### Competing Interest Statement

The authors have declared no competing interest.

### Summary of Updates

Extensive corrections of abstract, text and figures in the manuscript and SI_1.

