## Supplementary material for "Principles of ion binding to RNA inferred from the analysis of a 1.55 Å resolution bacterial ribosome structure – Part I: Mg^2+^": SI_1

**Table of content:**

1) **I.** – Current high-resolution (< 2.0 Å) *E. coli* ribosomal structures.
2) **II.** – K coordination parameters derived from the CSD.
3) **III.** – More details on the *“IonDiagnosis*” tool.
4) **IV.** – Coherency criteria used to check the assigned K^+^ ion binding sites.
5) **V.** – Readjustment of the Mg^2+^ exclusion zone from 3.6 to 3.4 Å.
6) **VI.** – Considerations on acetate and HEPES binding.
7) **VII.** – Further details regarding Mg^2+^ binding types.
8) **VIII.** – Mg^2+^ bound at the rRNA/r-protein interfaces in detail.
9) **Table S1:** Mg^2+^ bound to r-proteins, mRNA and to the SSU/LSU interface.
10) **Table S2:** Twenty-one Mg^2+^ intra-domain links in LSU.
11) **Figure S1.** Occurrences of *syn-*/*anti-*O_ph_.5O_w_ binding types.
12) **Figure S2.** Two short O_ph_…O_ph_ pairs stabilized by a water-mediated Mg^2+^ contact.
13) **Figure S3.** Similarities between the three 2N_b_.4O_w_ motifs also called “head-to-tail stacked purine” found in 8b0x.
14) **Figure S4.** View of (i) an SSU/LSU linking Mg^2+^, (ii) an uS2 bidentate Mg^2+^ and (iii) a rare O2’ binding.
15) **Figure S5.** Two base pairs with a negatively charged uridine and one base triple with a positively charged adenine nucleobase.
16) **Figure S6.** Complementary view of Mg^2+^ pairs of the MgA_μc_, MgB_μc_, MgC_μc_ and MgD_μc_ types.
17) **Figure S7.** K^+^ displays higher density peaks than Mg^2+^ and bound water.
18) **References:**

**I. Current high-resolution (< 2.0 Å) *E. coli* ribosomal structures**

The **8b0x** cryo-EM structure is currently the ribosomal structure with the best resolution (1.55 Å). Yet, other *E. coli* ribosome cryo-EM structures were obtained with resolutions ≤ 2.0 Å such as **7k00** and **8fto** (1) that display resolutions of 1.98 and 1.85 Å, respectively, **8aye** with a 1.96 Å resolution (2), the **8cgk** antibiotic bound 50S structure with a 1.64 Å resolution and related structures (3), the 30S **8cgj** structure with a 1.79 Å resolution and the 30S **8cf1** antibiotic bound structure with a 1.82 Å resolution that contains a model of the 30S head that is absent in **8cgj** (3). Note that **8b0x** is derived from strain B, that **7k00/8fto/8yae** are derived from strain MRE 600 and **8cgk/8cf1** from strain BW25113.

The SSU/LSU chains display sequence gaps between these different strains leading to small shifts in residue numbering (4). Note that the following SSU G_79_…C_90_, A_205_…G_213_, C_841_…A_845_, C_1028_…G_1033_, and A_1534_…C_1554_ and LSU G_545_…C_550_, G_1055_…C_1108_, G_1175_…C_1178_, G_1873_…C_1875_, G_2103_…C_2194_, U_2908_…U_2925_ residues were not modelled in **8b0x**. The 2.1 Å resolution **4ybb** X-ray structure of the K-12 strain integrates all SSU/LSU residues (5). The more recent **7zp8** cryo-EM structure with a 2.20 Å resolution contains all 50S residues (6).

**II. K coordination parameters derived from the CSD**

The average coordination distances derived from the CSD histogrames are: *d*(K…O) ≈ 2.85±0.13 Å and *d*(K…N) ≈ 2.96±0.14 Å (**Figure 2**). Given its large radius, K^+^ can accommodate six to nine O/N atoms in its first coordination shell (7-11). Although K^+^ ions with a coordination number of six are rare, such ions can be found in specific rRNA binding pockets. They might also be associated with incomplete hydration shells. When only surrounded by water, their coordination number is seven (note that 21 potassium ions coordinating to at least three water molecules and only two fully hydrated K[H_2_O]_7_^+^ ions are present in the current version of the CSD). Contrary to Mg, the K angle histograms are not very useful given that the most frequent potassium ligands in the CSD are diethoxy (O-C-C-O) groups belonging to crown ethers and related molecules that we discarded from our searches (**Figure 3**). Given that the Mg^2+^ binding site description is already quite extensive, we refrained to discuss in this report K^+^ ion binding features that will be addressed in a companion paper.

**III. More details on the *“IonDiagnosis”* tool**

To analyse the **8b0x** structure, we improved the functionalities of a tool developed earlier to validate ion positions in ribosome structures (12). This PyMol (13) based program takes as input a PDBid code and retrieves automatically from the PDB the associated CIF files and, if they were deposited, the 2Fo‑Fc and Fo‑Fc maps for X‑ray structures or the electron microscopy density maps (emd maps) for cryo-EM structures. Note that the *“IonDiagnosis”* tool has been designed specifically to analyse nucleic acid structures in association with rRNA binding proteins (r-proteins).

In spirit, this program is alike “CheckMyMetal” (CMM) that is a valuable tool to assess the validity of ion assignments in biomolecular structures (14). Older versions of CMM did not accept large CIF files making it unsuitable for the analysis of most ribosomal structures. This has been corrected in its most recent incarnation that allows a rapid visualization of every ion binding site and their density patterns (15) and provides an analysis of the **8b0x** structure as a test example.

Currently, the “*IonDiagnosis*” tool checks several parameters related to the stereochemistry of each ion assigned in the analysed structure and marks them as correct (green), incorrect (red) or problematic (orange). For instance, a valid Mg^2+^ must have six octahedrally coordinated O/N ligands with coordination distances in the 1.9 Å < *d*(Mg^2+^…O/N) < 2.3 Å range. Deviations from these values marked in red or orange call for a reinspection of the binding site and eventually the addition of restraints during a subsequent refinement stage (see below). Second shell water-mediated contacts are also monitored with a hydrogen-bond distance limit of 3.0 Å. If water-mediated contacts in the vicinity of the ions are shorter than 2.4 Å, they are marked in red, calling for reinspection.

The “*IonDiagnosis*” tool also provides the density peak values for each non-hydrogen atom position. For X-ray and cryo-EM structures, it uses the *“phenix.map_value_at_point”* program and the mtz file provided by the PDB to analyse the 2Fo-Fc/Fo-Fc X-ray maps or cryo-EM the electron density maps. Additionally, an annotated Excel file that describes the different ion binding features is available as **SI**. Unfortunately, we have to mention that this program, despite our efforts, will not be publicly available given the complexity of the issues that are associated with the ion correct/incorrect ion assignments made in PDB current structures. A good first step in the evaluation of ion binding issues can currently be made by using the CMM web site (15).

**IV. Coherence criteria used to check the assigned K^+^ binding sites**

For K^+^ ions, we used a different strategy than for Mg^2+^ since the observed coordination patterns are sometimes incomplete with coordination numbers ≤ 5 although *d*(K^+^…O/N) are in a broad 2.6–3.2 Å range. Thus, we relied mostly on the fact that K^+^ had to coordinate to at least a combination of three phosphate/carbonyl oxygen atoms and/or non-protonated nitrogens. To distinguish these ions from water, we checked if they exhibit coordination numbers greater than four and demonstrate density peak values higher than those of surrounding water molecules or Mg^2+^ ions (**Figure 15**). Such criteria allow to identify a large part of the most strongly bound K^+^ ions but not the diffusely bound ions. A full analysis of K^+^ coordination features will be provided in a companion paper.

**V. Readjustment of the Mg^2+^ exclusion zone from 3.6 to 3.4 Å**

By examining the Mg^2+^ binding sites, it was necessary to adjust the 3.6 Å upper Mg^2+^ exclusion zone limit derived from the CSD histograms (**Figure 2**) since some rRNA atoms and solvent molecules were found to occupy the 3.4–3.6 Å region. In a few instances, we could modify the backbone angles of the residues contacting the ions to push oxygen atoms out of the 3.6 Å exclusion zone. In other instances, we had to assume that these distances result from a local environment with no equivalence in the CSD. A closer examination of these instances revealed that they occurred mainly for ions that are coordinated to at least three non-water ligands (**Figure 9/10**).

**VI. Considerations on acetate and HEPES binding**

We were unable to assign acetate (ACY/ACT) anions, although one acetate was modelled in the **8cgk** structure of an LSU at 1.64 Å resolution at the uL14 r-protein interface (3), four were modelled in the 2.4 Å resolution **4v9f** Hm-LSU structure (16) and another in the 1.35 Å resolution **4fen** riboswitch structure where it binds to the Watson-Crick site of a guanine (17). More high-resolution structures will be required to precisely determine acetate binding modes (18,19) and locate HEPES (EPE) molecules that are present at a 50 mM concentration in the final cryo-EM buffer although we assume that the latter rather bulky molecules might attach to the ribosome surface or remain in the bulk.

We also note that the main *in vivo* anion is Cl^-^ and not CH_3_COO^-^ although acetate is an essential metabolite for living organisms. Therefore, in the present system, the ≈150 mM acetate concentration is largely overestimated. The ≈50 mM HEPES and ≈1 mM DTT are non-natural additions to the system.

**VII. Further details regarding Mg^2+^ binding types**

Here we present additional details relative to the binding sites described **Figure 4**.

• 6O_w_: Given their great diversity, it is difficult to describe all 6O_w_ binding patterns despite some attempts made by the MgRNA study and the Klein and Steitz seminal investigation (20,21). However, a few of their characteristics can be mentioned. The best hydrogen bond acceptors are the Hoogsteen O6/N7 guanine atoms and the OP1/OP2 phosphate group oxygens (20,21). We note a slight preference for the OP2 atoms that are pointing towards the deep groove in helical fragments. Among these 79 “well-defined” ions, 45 of them contact both O6/N7 atoms and 9 contact only one of these atoms. Further, 16 ions contact OP1/OP2/O2’ backbone atoms while 63 contact a combination of nucleobase and backbone atoms (**Figure 5A**).

Besides the major groove oriented OP2 atoms, the binding to OP1 atoms is more characteristic of shallow groove contacts associated with diverse loops and interhelical motifs. **Figure 5A** shows a Mg^2+^ “bridging water” that connects two O_ph_ atoms from consecutive nucleotides. Such recurrent hydration patters seen in **Figure 5A** will be explored elsewhere.

Due to reduced accessibility, the O2’/O3’/O4’/O5’, the purine N3 atoms and the pyrimidine O2 atoms are rarely involved in water-mediated hydrogen-bonds. Thus, while a recurrent 6O_w_ binding pattern to the guanine Hoogsteen edge emerges, this pattern is not unique since numerous structural shifts associated with sequence variations can perturb it. For instance, when an ion binds to the deep groove of a Watson-Crick pair, it tends also to establish water-mediated contacts with the nearby OP2 atoms.

To summarize, there is a preference for the binding of 6O_w_ ions to guanine-rich major grooves (20,21). However, depending on the structural context, any combination of OP1/OP2/O_b_/O_r_/N_b_ atoms can be part of the 2^nd^ coordination shell of Mg[H_2_O]_6_^2+^ ions.

• O_ph_.5O_w_ coordination: Ninety-seven of the O_ph_.5O_w_ ions (**Figure S1**) are in *syn-* with respect of the phosphate groups and only 8 are in *anti-* if we consider a nomenclature developed for carboxylate groups (22); see also (23).

• *cis-*2O_ph_.4O_w_ coordination variations: Variations of this motif involve the replacement of one or two O_ph_ atom by O_b_/N_b_ atoms. Eight well-defined occurrences are of the *cis-*O_ph_.O_b_.4O_w_ type, two are of the *cis-*O_ph_.N_b_.4O_w_ type and two of the *cis-*O_ph_.4O_w_.N_His_ type while one poorly defined occurrence is of the *cis-*2O_b_.4O_w_ type and involves (C)O2 and (G)O6 atoms.

• *fac-*3O_ph_.3O_w_ and *mer-*3O_ph_.3O_w_ coordination variations: As variations of these motifs, two *mer-*2O_ph_.O_b_.3O_w_, two *fac-*2O_ph_.N_b_.3O_w_ and one of each of the *mer-*O_ph_.2O_b_.3O_w_, *fac-*2O_ph_.O_r_.3O_w_ (see **Figure 11B** for the latter) and *mer-*2O_ph_.O_cno_.3O_w_ binding sites were observed. Several of these motifs are associated with Mg^2+^…K^+^ pairs (see **Figure 15B**).

• *Cis-/Trans-*4O_ph_.2O_w_ coordination variations: Three “well-defined” variations involve O_b_/O_r_/N atoms that replace O_ph_ atoms. A *Cis-*3O_ph_.O_b_.2O_w_ site is associated with a (U)O4 atom, a *Cis-*3O_ph_.O_r_.2O_w_ site is associated with a rare O2’ coordination (see **Figure 11A**), a *Cis-2*O_ph_.O_bb_.2O_w_.N_His_ site joins the uL3/15 r-proteins to the LSU. Additionally, three “poorly-defined” site with Mg^2+^ having a density peak value < 4.0 were also characterized: a *Cis-*2O_ph_.2O_b_.2O_w_, a *Cis-*3O_coo_.2O_w_.N_His_ and a *Cis-*O_ph_.3O_bb_.2O_w_ binding site in uS2.

**VIII. Mg^2+^ bound at the rRNA/r-protein interfaces in detail**

Only five Mg^2+^ ions establish inner shell contacts to both rRNA and r-proteins (**Table S2**). These comprise: *(i)* a *Cis-*O_ph_.3O_bb_.2O_w_ ion that bridges SSU/uS13, *(ii)* a *cis-*O_ph_.4O_w_.N_His_ ion that bridges LSU/uL2, *(iii)* a second *cis-*O_ph_.4O_w_.N_His_ ion that bridges LSU/uL3, *(iv)* a *Cis-*2O_ph_.O_bb_.2O_w_.N_His_ ion that bridges LSU/uL3+bL17 and *(v)* a *cis-*2O_ph_.O_cno_.3O_w_ ion that bridges LSU/uL15. In addition, some Mg^2+^ are forming at least one contact to the rRNA and establish water-mediated contacts to rProteins. Seven such ions are located at the LSU/uL2 interface, three at the LSU/uL3 interface and three at the uL23 interface. Single instances of ions binding to L4/L13/L14/L17/L20/L27/L34/L35 were noted (**Table S2**). In addition, a Mg^2+^ of the O_coo_.5O_w_ type connects bS34 to the LSU through water-mediated contacts. For the SSU, similar single instances of ions binding to S2/S10/S14 are observed. It can be noted that nine hexahydrated Mg^2+^ ions are also located at rRNA/r-protein interfaces.

**Table S1: Mg^2+^ bound to r-proteins, mRNA and to the SSU/LSU interface.** The number of contacts is given in parenthesis.

| rRNA chains | r-protein/  rRNA; mRNA | Inner-shell contacts  (RNA) | Inner-shell contacts  (r-protein, mRNA, SSU/LSU) | Outer-shell contacts  (RNA) | Outer-shell contacts  (r-protein, mRNA, SSU/LSU) |
| --- | --- | --- | --- | --- | --- |
| SSU (chain A) | uS2 (chain B) | – | 9:3969 (4) | – | 9:3969 (2) |
| SSU (chain A) | uS10 (chain J) | A:1703 (1) | – | A:1703 (2) | A:1703 (1) |
| SSU (chain A/X) | uS11 (chain K) | – | – | 9:486 (5) | 9:486 (1) |
| SSU (chain A) | uS13 (chain M) | 9:1080 (1) | 9:1080 (3) | 9:1080 (1) | – |
| SSU (chain A) | uS14 (chain N) | –  – | –  – | A:1687 (3)  A:1692 (4) | A:1687 (2)  A:1692 (2) |
| SSU (chain A) | uS17 (chain Q) | – |  | A:1741 (3) | A:1741 (1) |
| SSU (chain A) | mRNA (chain X) | 9:2511 (1)  –  – | –  –  – | 9:2511 (2)  A:1611 (2)  X:101 (3) | –  A:1611 (1)  X:101 (3) |
| SSU (chain A)^a^ | LSU (chain a) | A:1638 (1) | – | – | A:1638 (2) |
| LSU (chain a) | uL2 (chain c) | –  c:304 (1)  9:376 (1)  9:2043 (1)  a:3153 (1)  a:3202 (1)  a:3224 (3)  a:3225 (2)  a:3305 (1) | –  c:304 (1)  –  –  –  –  –  –  – | –  c:304 (1)  9:376 (2)  9:2043 (3)  a:3153 (2)  a:3202 (3)  –  a:3225 (3)  a:3305 (4) | c:302 (2)  c:304 (2)  9:376 (1)  9:2043 (1)  a:3153 (3)  a:3202 (2)  a:3224 (2)  a:3225 (2)  a:3305 (2) |
| LSU (chain a) | uL3 (chain d) | d:301(1)  a:3147 (1)  a:3222 (1) | d:301 (1)  –  – | d:301 (3)  –  a:3222 (5) | d:301 (1)  a:3147 (2)  a:3222 (1) |
| LSU (chain a) | uL3 (chain d)/  bL17 (chain m) | a:3186 (2)  – | a:3186 (1)  a:3186 (1) | a:3186 (1)  – | –  – |
| LSU (chain a) | uL4 (chain e) | a:3082 (1) | – | a:3082 (4) | a:3082 (2) |
| LSU (chain a) | uL13 (chain i) | a:3093 (2) | – |  | a:30943(2) |
| LSU (chain a) | uL15 (chain k) | a:3094 (1)  a:3219 (2) | –  a:3219 (1) | a:3094 (1)  – | a:3094 (3)  – |
| LSU (chain a) | uL16 (chain l) | – | – | 9:486 (4) | 9:486 (1) |
| LSU (chain a) | bL17 (chain m) | (see also uL3)  – | –  – | a:3028 (3)  a:3095 (5) | a:3028 (1)  a:3095 (1) |
| LSU (chain a) | bL19 (chain o) | – | – | 9:7287 (1) | a:3364 (2) |
| LSU (chain a) | bL20 (chain p) | –  a:3166 (1) | –  – | a:3146 (1)  a:3166 (1) | a:3146 (2)  a:3166 (2) |
| LSU (chain a) | uL23 (chain s) | 9:2716 (1)  a:3332 (2)  a:3188 (1) | –  –  – | –  a:3332 (1)  a:3188 (3) | 9:2716 (1)  a:3332 (1)  a:3188 (1) |
| LSU (chain a) | bL27 (chain v) | a:3364 (1) | – | a:3364 (1) | a:3364 (2) |
| LSU (chain a) | bL34 (chain 1) | 9:1154 (1)  – | –  – | 9:1154 (2)  a:3171 (2) | 9:1154 (1)  A:3171 (1) |
| LSU (chain a) | bL35 (chain 2) | a:3321 (1) | – | a:3321 (1) | a:3321 (2) |

**Table S2: Twenty-one Mg^2+^ intra-domain links in LSU.** None were observed in the SSU.

| LSU Mg^2+^ ion | Domain I | Domain II | Domain III | Domain IV | Domain V | Domain VI | Total: | Figure: |
| --- | --- | --- | --- | --- | --- | --- | --- | --- |
| 9:6037^a^ | 1 | 1 | . | . | . | . | 2 | . |
| a:3152 | . | . | . | 1 | 1 | . | 2 | . |
| a:3201 | . | 2 | . | . | 1 | . | 3 | . |
| a:3223 | . | 1 | 1 | . | . | . | 2 | 7C |
| a:3224 | . | 2 | . | . | 1 | . | 3 | 8A/16 |
| a:3225 | . | 1 | . | . | 1 | . | 2 | 9A/16 |
| a:3230 | . | 1 | . | . | 2 | . | 3 | . |
| a:3236 | . | 1 | . | 1 | . | . | 2 | 6B |
| a:3237 | . | 1 | . | 1 | . | . | 2 | . |
| a:3238 | . | 1 | 1 | 2 | . | . | 4 | . |
| a:3308 | 1 | . | . | . | 1 | . | 2 | . |
| a:3309 | 1 | . | . | . | 1 | . | 2 | . |
| a:3322 | . | 1 | . | . | 1 | . | 2 | . |
| a:3324 | . | 2 | . | . | 1 | . | 3 | 16 |
| a:3325 | . | 1 | . | . | 2 | . | 3 | 16 |
| a:3328 | . | 1 | . | 2 | . | . | 3 | . |
| a:3329 | . | 1 | . | . | 3 | . | 4 | 10B |
| a:3330 | 1 | 1 | . | . | . | . | 2 | 14C |
| a:3333 | . | . | . | 3 | 1 | . | 4 | . |
| a:3336 | 1 | . | . | . | 1 | . | 2 | . |
| a:3363 | . | . | . | 1 | 1 | . | 2 | . |
| Total: | 5 | 16 | 2 | 11 | 17 | . | 51 | . |
| ^a^ This contact is less well-defined. | | | | | | | | |

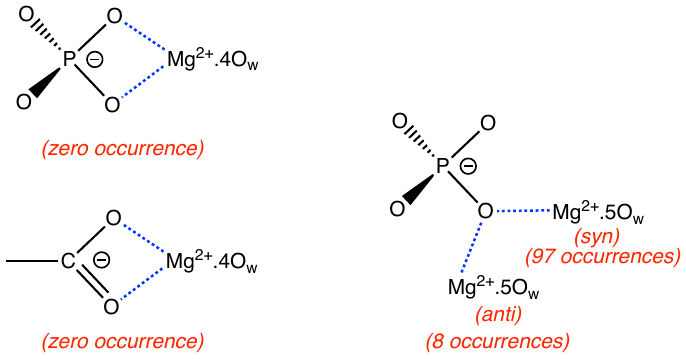

**Figure S1. Occurrences of *syn-*/*anti-*O_ph_.5O_w_ binding types.** (Left) These “bidentate” coordination types, mentionned in several instances (23,24), are not observed in **8b0x**. They seem inaccessile to Mg^2+^ ions and are not to be confused with the “bidentate” coordination type shown in **Figure 7A** that involves two different phosphate groups that are consecutive or not. (Right) The coordination in *syn-* to phosphate groups dominates over that in *anti-* (22). Present occurrence count refers to the 105 well-defined O_ph_.5O_w_ types located in the amended **8b0x** structure.

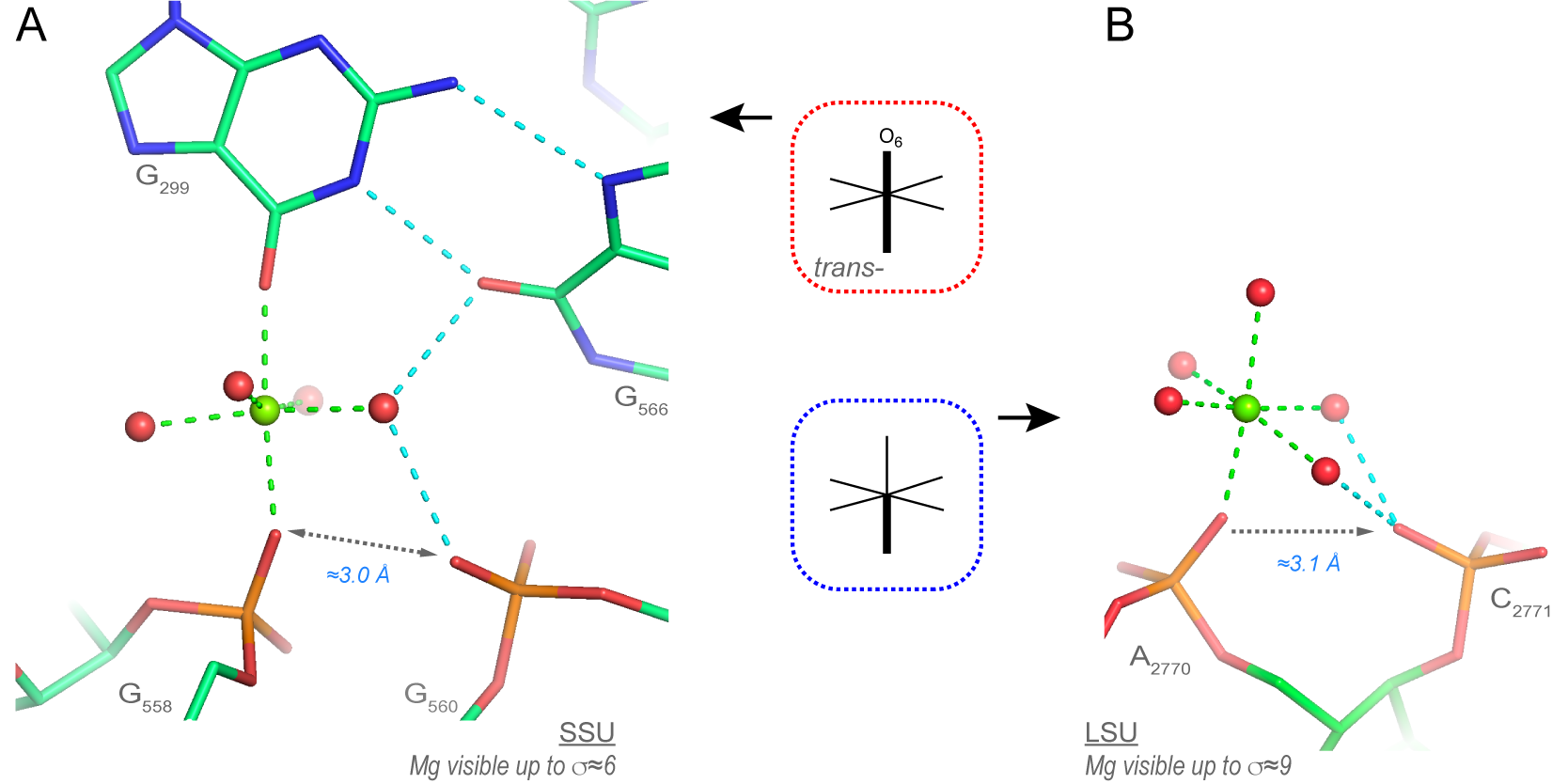

**Figure S2. Two short O_ph_…O_ph_ pairs stabilized by a water-mediated Mg^2+^ contact.** (**A/B**) These SSU domain “5’” and LSU domain “VI” Mg^2+^ ions are of the *trans-*O_ph_.O_b_.4O_w_ and O_ph_.5O_w_ types. These configurations might represent intermediate stages in the formation of Mg^2+^ bidentate clamps. Green/cyan lines mark distances ≤ 2.3 Å and in the 2.6–3.2 Å range. Experimental densities and some water molecules are not shown.

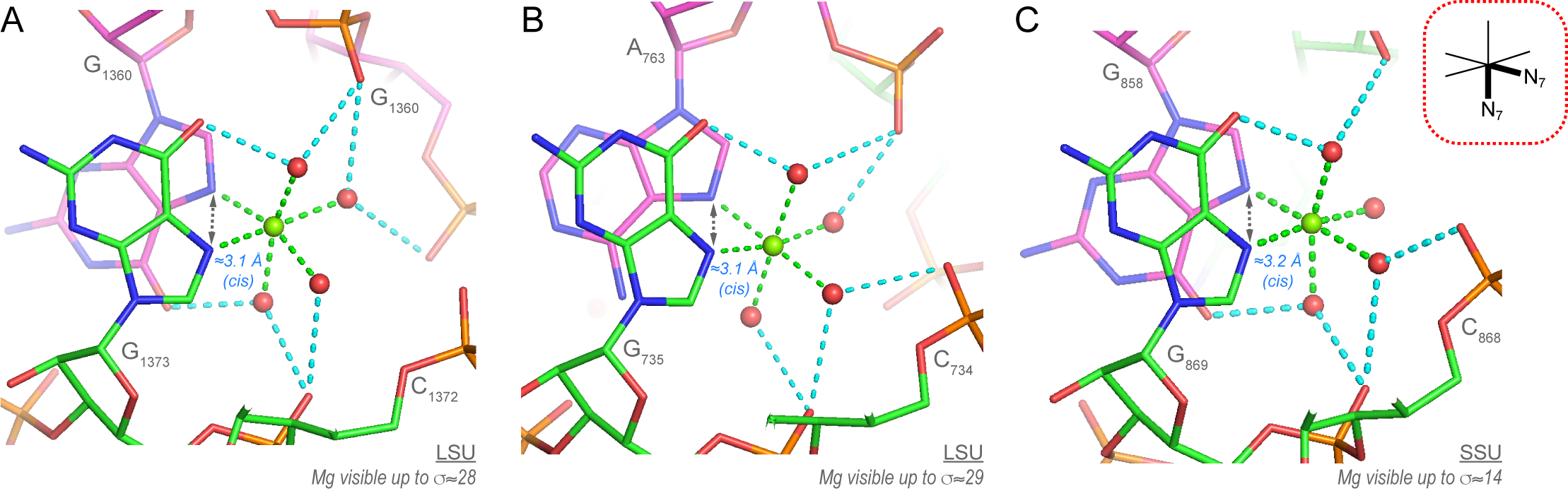

**Figure S3. Similarities between the three 2N_b_.4O_w_ motifs also called “head-to-tail stacked purine” found in 8b0x**. (**A/B/C**) These 2N_b_.4O_w_ ions are highly connected. They display two inner-shell contacts and up to 7 water-mediated contacts. The motif in **A** is the same as the one shown **Figure 8**. Green/cyan lines mark distances ≤ 2.3 Å and in the 2.6–3.2 Å range. Experimental densities and some water molecules were hidden.

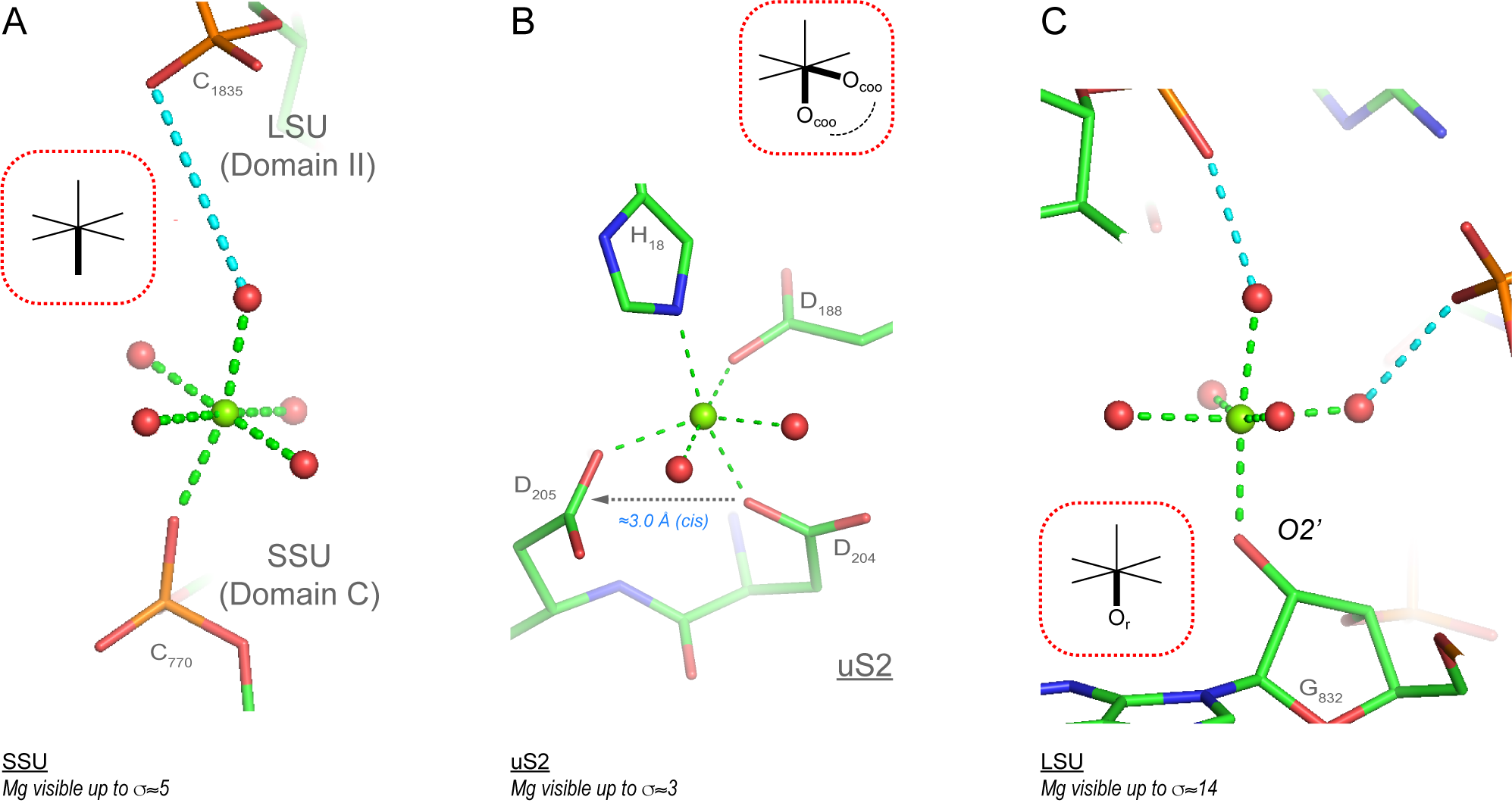

**Figure S4. View of (i) an SSU/LSU linking Mg^2+^, (ii) an uS2 bidentate Mg^2+^ and (iii) a rare O2’ binding.** (**A**) This is the only Mg^2+^ ion interacting with both the SSU and the LSU. This contact is probably marginal. (**B**) uS2 is the only **8b0x** r-protein with a detected Mg^2+^ binding site. Note that this Mg^2+^ links an N-terminal histidine to residues close to the C-terminal end and that two adjacent aspartates form a Mg^2+^ clamp similar to those found in rRNA. (**C**) Although “well defined”, this O_r_.5O_w_ coordination pattern is difficult to rationalize given that it occurs in an almost unconstrained environment and is unique among the >400 observed Mg^2+^ ions. Such a binding site, despite a clear density, remains hypothetical until similar occurrences in other high-resolution RNA structures are uncovered. Green/cyan lines mark distances ≤ 2.3 Å and in the 2.6–3.2 Å range. Experimental densities and some water molecules were hidden.

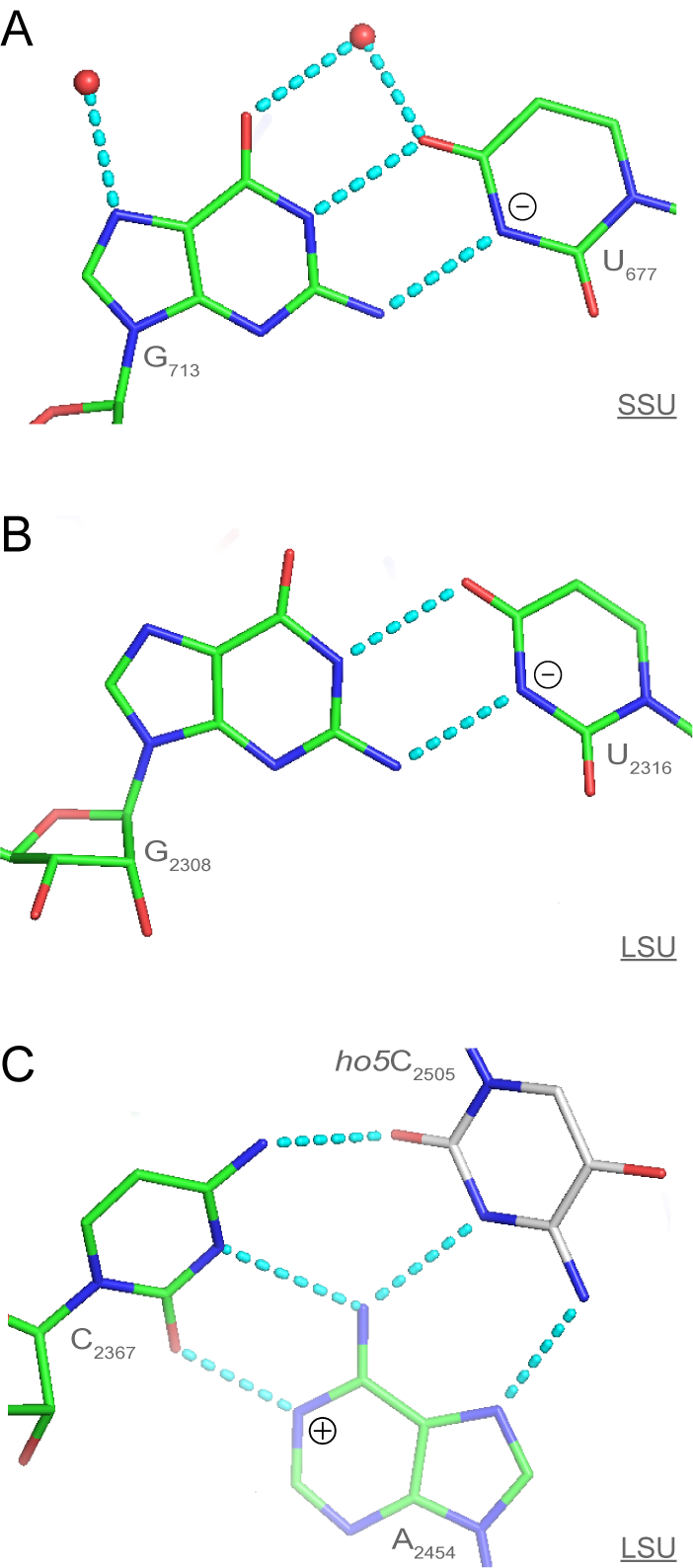

**Figure S5. Two base pairs with a negatively charged uridine and one base triple with a positively charged adenine nucleobase.** (**A/B**) Two G•U(-) pair with a negatively charged uridine. These base pairs are also described in reference (25). (**C**) In addition, we show the LSU C•A(+) pair with a protonated adenine that forms a [C•A^+^]•5hoC base triple. Note that the 5-hydroxycytidine (ho5C) residue was modelled as a C residue in **8b0x**. This was corrected in the amended model. Cyan lines mark distances in the 2.6–3.2 Å range (see also **Figure 12**). Experimental densities and some water molecules were hidden.

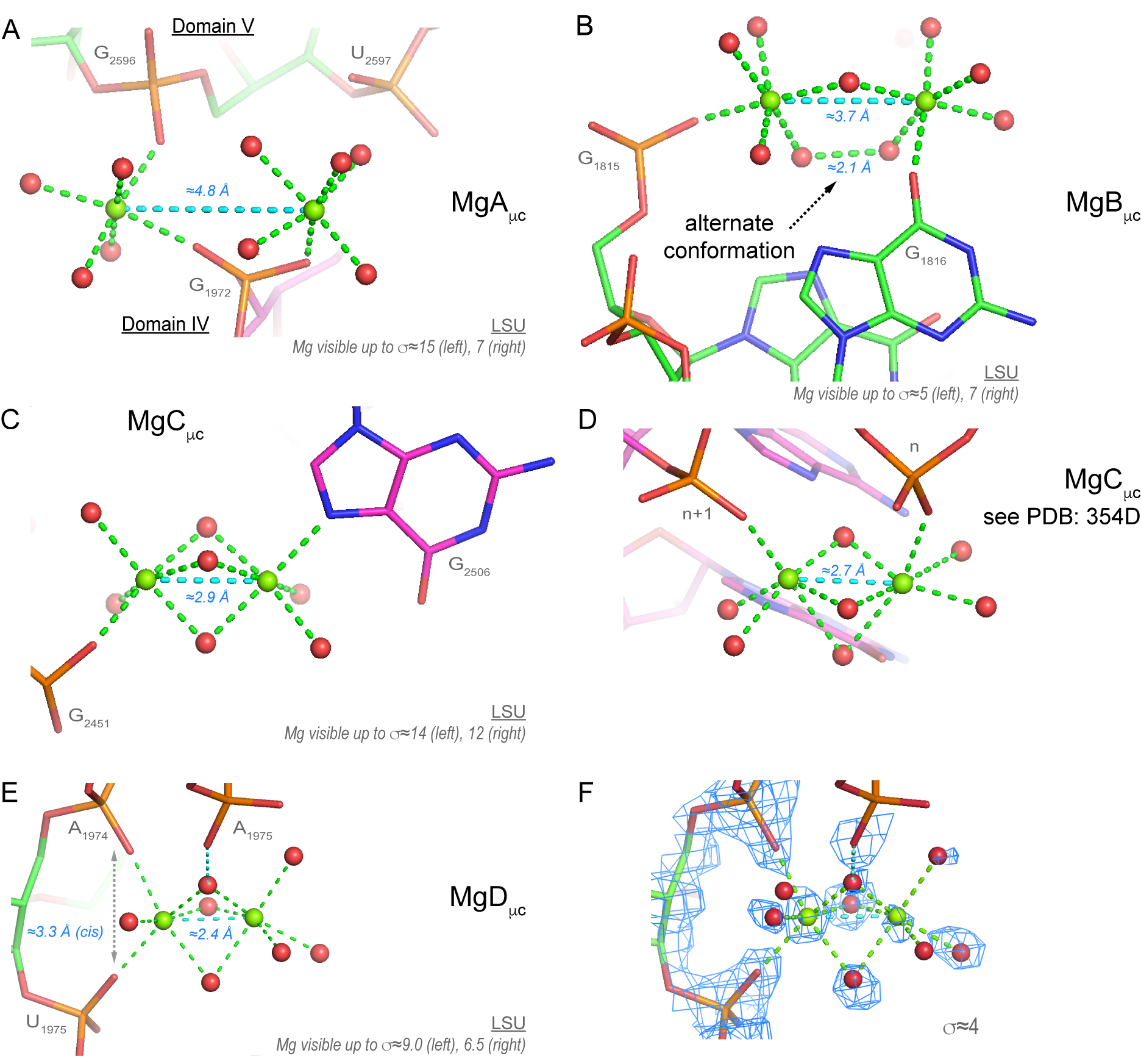

**Figure S6. Complementary view of Mg^2+^ pairs of the MgA_μc_, MgB_μc_, MgC_μc_ and MgD_μc_ types.** (**A**) This MgA_μc_ pair connects domain “IV” and “V”. The first ion is *cis-*2O_ph_.4O_w_ and the second is O_ph_.5O_w_ (see also **Figure 14**). (**B**) Note the 2.1 Å distance between two water molecules of a MgB_μc_ pair that suggests the presence of an alternate conformation. The first ion is O_ph_.5O_w_ and the second is O_b_.5O_w_. (**C**) This MgC_μc_ pair involves three bridging waters. (**D**) This MgC_μc_ pair is derived from a 5S rRNA loop E motif (PDBid: **354d**) solved at 1.5 Å resolution (26). Note the distorted coordination octahedrons of the second Mg^2+^. The X-ray density maps were not deposited to the PDB. (**E**) The arrangement of this MgD_μc_ pair remains speculative in the absence of corroborating examples showing ultrashort inter-metal distances close to ≈2.4 Å (see **Figure 14D**). (**F**) The density pattern associated with the MgD_μc_ pair. Green/cyan lines mark distances ≤ 2.3 Å and in the 2.6–3.2 Å range. Experimental densities and some water molecules were hidden in **A** to **E**.

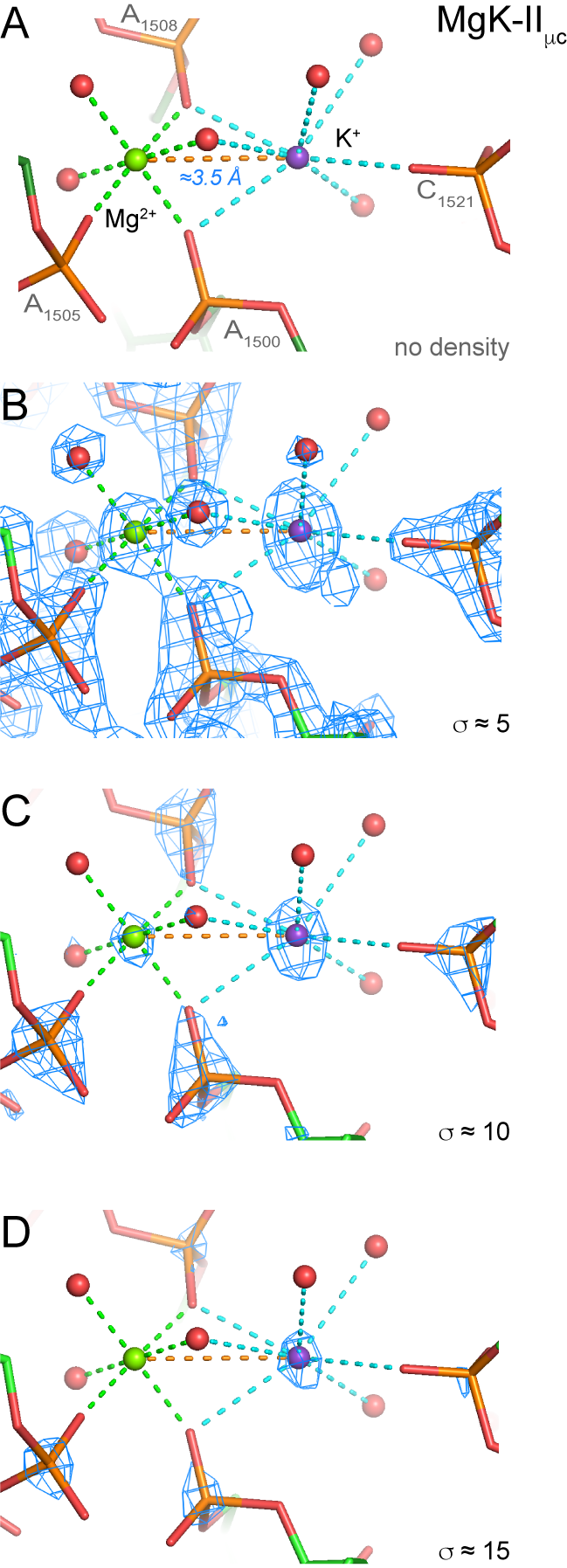

**Figure S7. K^+^ displays higher density peaks than Mg^2+^ and bound waters.** (**A**) This MgK_μc3_ pair involves a Mg^2+^ of the *mer-*3O_ph_.3O_w_ type and a heptacoordinated K^+^ (see **Figure 15B**). (**B/C/D**) Views at increasing σ levels emphasizing that the K^+^ density peak is higher than that of the Mg^2+^ ion and surrounding water molecules. Mg^2+^/K^+^ are green/purple. Green/cyan/orange lines mark distances ≤ 2.3 Å, in the 2.6–3.2 Å range and > 3.2 Å. Some water molecules were omitted from the views.
