## Supplementary material for "Principles of ion binding to RNA inferred from the analysis of a 1.55 Å resolution bacterial ribosome structure – Part I: Mg^2+^": SI_ion_diagnosis: 8B0X.ion_diagnosis_original.html

 Ion report: 8B0X  

### Ion report: 8B0X

#### date: 2023-11-10 17:28:24

 **→→→ This is a cryo-EM structure:**  **→→→ An EMDB map has been retrieved and is checked!** Authors: Filip Leonarski & Pascal Auffinger  
\*\*\* BETA version: unpublished \*\*\*  
  
→ reference 1: Leonarski F., D'Ascenzo L., Auffinger P. (2017) NAR, 45, 987-1004  
→ reference 2: Leonarski F., D'Ascenzo L., Auffinger P. (2019) RNA, 25, 173-192  
→ reference 3: Auffinger P., Ennifar E., D'Ascenzo L. (2021) RNA, 27, 243-252  
  
\_\_\_\_\_\_\_\_\_\_\_\_\_\_\_\_\_\_\_\_\_\_\_\_\_\_\_\_\_\_\_\_\_\_\_\_\_\_\_\_\_\_\_\_\_\_\_\_\_\_\_\_\_\_\_\_\_\_\_\_\_\_\_\_\_\_\_\_\_\_\_\_\_\_\_\_\_\_\_\_\_  

#### Table of content (ToC):

1) Number of ions of each type + coordination distance ranges used for validation;  
2) Ion validation table;  
3) Potential ion coordination issue list;  
4) Ion/ligand tables;  
5) Sorted coded contact list;  
6) B-factor and 2FoFc, FoFc density map summary;  
7) Potential and detected Mg^2+^ coodination clamps list;  
\_\_\_\_\_\_\_\_\_\_\_\_\_\_\_\_\_\_\_\_\_\_\_\_\_\_\_\_\_\_\_\_\_\_\_\_\_\_\_\_\_\_\_\_\_\_\_\_\_\_\_\_\_\_\_\_\_\_\_\_\_\_\_\_\_\_\_\_\_\_\_\_\_\_\_\_\_\_\_\_\_  
 **Notes:**  
**• 1^st^ coordination shell values correspond to an approximate coordination distance range;**  
**• Inner and outer exclusion zones (excl. zone) correspond to distance ranges where \*NO\* coordinating atoms should be present;**  
**→ These values are derived from Cambridge Structural database (CSD) searches (see reeferences above);**  
**→ *Coordination distances with sulfur (S) atoms are not yet checked;***  
**→ *Currently no CD/ZN (cadmium/zinc) validation;***  
\_\_\_\_\_\_\_\_\_\_\_\_\_\_\_\_\_\_\_\_\_\_\_\_\_\_\_\_\_\_\_\_\_\_\_\_\_\_\_\_\_\_\_\_\_\_\_\_\_\_\_\_\_\_\_\_\_\_\_\_\_\_\_\_\_\_\_\_\_\_\_\_\_\_\_\_\_\_\_\_\_  

#### 1) Number of ions of each type + coordination distance ranges used for validation

##### Go to ToC

| Count \| Ion list \| Inner excl. zone [Å] \| 1^st^ shell [Å] \| Outer excl. zone [Å] \| 2^nd^ shell [Å] \| |  |  |  |  |  |
| --- | --- | --- | --- | --- | --- |
| Magnesium (MG) | 361 | A:1606 A:1607 A:1608 A:1609 A:1610 A:1611 A:1612 A:1613 A:1614 A:1615 A:1616 A:1617 A:1618 A:1619 A:1620 A:1621 A:1622 A:1623 A:1624 A:1625 A:1626 A:1627 A:1628 A:1629 A:1630 A:1631 A:1632 A:1633 A:1634 A:1635 A:1636 A:1637 A:1638 A:1639 A:1640 A:1641 A:1642 A:1643 A:1668 A:1669 A:1670 A:1671 A:1672 A:1673 A:1674 A:1675 A:1676 A:1677 A:1678 A:1679 A:1680 A:1681 A:1682 A:1683 A:1684 A:1685 A:1686 A:1687 A:1688 A:1689 A:1690 A:1691 A:1692 A:1693 A:1694 A:1695 A:1696 A:1697 A:1698 A:1699 A:1700 A:1701 A:1702 A:1703 A:1704 A:1705 A:1706 A:1707 A:1708 A:1709 A:1710 A:1711 A:1712 A:1713 A:1714 A:1715 A:1716 A:1717 A:1718 A:1719 A:1720 A:1721 A:1741 A:1742 A:1743 A:1744 A:1745 X:101 a:3001 a:3002 a:3003 a:3004 a:3005 a:3006 a:3007 a:3008 a:3009 a:3010 a:3011 a:3012 a:3013 a:3014 a:3015 a:3016 a:3017 a:3018 a:3019 a:3020 a:3021 a:3022 a:3023 a:3024 a:3025 a:3026 a:3027 a:3028 a:3029 a:3030 a:3031 a:3032 a:3033 a:3034 a:3035 a:3036 a:3037 a:3038 a:3039 a:3040 a:3041 a:3042 a:3043 a:3044 a:3045 a:3046 a:3047 a:3048 a:3049 a:3050 a:3051 a:3052 a:3053 a:3054 a:3055 a:3065 a:3066 a:3067 a:3068 a:3069 a:3070 a:3071 a:3072 a:3073 a:3074 a:3075 a:3076 a:3077 a:3078 a:3079 a:3080 a:3081 a:3082 a:3083 a:3084 a:3085 a:3086 a:3087 a:3088 a:3089 a:3090 a:3091 a:3092 a:3093 a:3094 a:3095 a:3143 a:3144 a:3145 a:3146 a:3147 a:3148 a:3149 a:3150 a:3151 a:3152 a:3153 a:3154 a:3155 a:3156 a:3157 a:3158 a:3159 a:3160 a:3161 a:3162 a:3163 a:3164 a:3165 a:3166 a:3167 a:3168 a:3169 a:3170 a:3171 a:3172 a:3173 a:3174 a:3182 a:3183 a:3184 a:3185 a:3186 a:3187 a:3188 a:3189 a:3190 a:3191 a:3192 a:3193 a:3194 a:3195 a:3196 a:3197 a:3198 a:3199 a:3200 a:3201 a:3202 a:3203 a:3204 a:3205 a:3206 a:3207 a:3208 a:3209 a:3210 a:3211 a:3212 a:3213 a:3214 a:3215 a:3216 a:3217 a:3218 a:3219 a:3220 a:3221 a:3222 a:3223 a:3224 a:3225 a:3226 a:3227 a:3228 a:3229 a:3230 a:3231 a:3232 a:3233 a:3234 a:3235 a:3236 a:3237 a:3238 a:3239 a:3240 a:3241 a:3242 a:3243 a:3244 a:3245 a:3285 a:3286 a:3287 a:3288 a:3289 a:3290 a:3291 a:3292 a:3293 a:3294 a:3295 a:3296 a:3297 a:3298 a:3299 a:3300 a:3301 a:3302 a:3303 a:3304 a:3305 a:3306 a:3307 a:3308 a:3309 a:3310 a:3311 a:3312 a:3313 a:3314 a:3315 a:3316 a:3317 a:3318 a:3319 a:3320 a:3321 a:3322 a:3323 a:3324 a:3325 a:3326 a:3327 a:3328 a:3329 a:3330 a:3331 a:3332 a:3333 a:3334 a:3335 a:3336 a:3337 a:3348 a:3349 a:3350 a:3351 a:3352 a:3353 a:3354 a:3355 a:3356 a:3357 a:3358 a:3359 a:3360 a:3361 a:3362 a:3363 a:3364 a:3365 a:3366 b:201 b:202 b:203 b:204 b:205 c:302 c:304 d:301 z:101 | <1.9 | 1.9 - 2.3 | 2.3 - 3.6 >3.6 |
| Sodium (NA) | --- |  | <2.3 | 2.3 - 2.6 | 2.6 - 3.6 >3.6 |
| Potassium (K) | 168 | A:1601 A:1602 A:1603 A:1604 A:1605 A:1644 A:1645 A:1646 A:1647 A:1648 A:1649 A:1650 A:1651 A:1652 A:1653 A:1654 A:1655 A:1656 A:1657 A:1658 A:1659 A:1660 A:1661 A:1662 A:1663 A:1664 A:1665 A:1666 A:1667 A:1722 A:1723 A:1724 A:1725 A:1726 A:1727 A:1728 A:1729 A:1730 A:1731 A:1732 A:1733 A:1734 A:1735 A:1736 A:1737 A:1738 A:1739 A:1740 a:3056 a:3057 a:3058 a:3059 a:3060 a:3061 a:3062 a:3063 a:3064 a:3096 a:3097 a:3098 a:3099 a:3100 a:3101 a:3102 a:3103 a:3104 a:3105 a:3106 a:3107 a:3108 a:3109 a:3110 a:3111 a:3112 a:3113 a:3114 a:3115 a:3116 a:3117 a:3118 a:3119 a:3120 a:3121 a:3122 a:3123 a:3124 a:3125 a:3126 a:3127 a:3128 a:3129 a:3130 a:3131 a:3132 a:3133 a:3134 a:3135 a:3136 a:3137 a:3138 a:3139 a:3140 a:3141 a:3142 a:3175 a:3176 a:3177 a:3178 a:3179 a:3180 a:3181 a:3246 a:3247 a:3248 a:3249 a:3250 a:3251 a:3252 a:3253 a:3254 a:3255 a:3256 a:3257 a:3258 a:3259 a:3260 a:3261 a:3262 a:3263 a:3264 a:3265 a:3266 a:3267 a:3268 a:3269 a:3270 a:3271 a:3272 a:3273 a:3274 a:3275 a:3276 a:3277 a:3278 a:3279 a:3280 a:3281 a:3282 a:3283 a:3284 a:3338 a:3339 a:3340 a:3341 a:3342 a:3343 a:3344 a:3345 a:3346 a:3347 b:206 c:301 c:303 c:305 c:306 d:302 e:301 f:201 | <2.6 | 2.6 - 3.2 | 3.2 - 3.2 >3.2 |
| Caesium (CS) | --- |  | <2.9 | 2.9 - 3.6 | 3.6 - 3.6 >3.6 |
| Calcium (CA) | --- |  | <2.2 | 2.2 - 2.8 | 2.8 - 3.6 >3.6 |
| Strontium (SR) | --- |  | <2.4 | 2.4 - 2.8 | 2.8 - 4.0 >4.0 |
| Barium (BA) | --- |  | <2.6 | 2.6 - 3.2 | 3.2 - 3.2 >3.2 |
| Manganese (MN) | --- |  | <2.0 | 2.0 - 2.3 | 2.3 - 3.6 >3.6 |
| Cobalt (CO) | --- |  | <1.9 | 1.9 - 2.3 | 2.3 - 3.5 >3.5 |
| Zinc (ZN) | 1 | 3:101 | <1.9 | 1.9 - 2.4 | 2.4 - 3.4 >3.4 |
| Cadmium (CD) | --- |  | <2.4 | 2.4 - 2.8 | 2.8 - 2.8 >2.8 |
| Iridium (IRI) | --- |  | <2.0 | 2.0 - 2.3 | 2.3 - 3.6 >3.6 |
| Ammonium (NH4) | --- |  | <2.6 | 2.6 - 3.2 | 3.2 - 3.2 >3.2 |
| Chloride (CL) | --- |  | <2.8 | 2.8 - 3.4 | 3.4 - 3.4 >3.4 |
| Sulphate (SO4) | --- |  | <2.8 | 2.8 - 3.4 | 3.4 - 3.4 >3.4 |
| Phosphate (PO4) | --- |  | <2.8 | 2.8 - 3.4 | 3.4 - 3.4 >3.4 |
| **Total** | **530** |  |  |  |  |

---

\_\_\_\_\_\_\_\_\_\_\_\_\_\_\_\_\_\_\_\_\_\_\_\_\_\_\_\_\_\_\_\_\_\_\_\_\_\_\_\_\_\_\_\_\_\_\_\_\_\_\_\_\_\_\_\_\_\_\_\_\_\_\_\_  

#### 2) Ion validation table

##### Go to ToC

Atom type definitions:

• Nucleic acid:  
 O_ph_ → OP1/OP2 anionic phosphate oxygens;  
 O_r_ → O2'/O3'/O4'/O5' hydroxyl and ester oxygens;  
 O_b_ → O2/O4/O6 carbonyl oxygens;  
 N_b_ → N1_A_/N3_C_/N3_G_/N3_A_/N7 non-protonated nitrogens;  
• Protein:  
 O_back_ → protein backbone carbonyl group;  
 O_coo_ → ASP/GLU carboxyl group;  
 O_cno_ → ASN/GLN carbonyl group;  
 OH_prot_ → SER/THR/TYR hydroxyl group;  
 N_His_ → ND1/NE2 histidine atoms;  
• Water:  
 O_w_ → water oxygen;  
• NH groups:  
 -NH, -NH_2_, NH_3_(+), NH_4_(+) → groups from nucleic acids, proteins, ligands, or ammonium cations potentially binding to water or anions.;  
(suggests anion contact or poor binding site modeling);  
  

Notes:

The ions that are currently checked for octahedral coordination are Mg^2+^, Mn^2+^ and Na^+^;  
*→ Octahedral coordination is only checked for ions with more than three ligands in their 1^st^ coordination shell;*  
• "Distorded coordination octahedron" means a cumulated (ligand...ion...ligand) angle deviation > 10º;  
• "Slightly distorded coordination octahedron" means a cumulated (ligand...ion...ligand) angle deviation between 5 and 10º;  
• "Idealized" suggests that some water positions have been derived from a template (only applicable for water coordination) that are sometimes used for low-resolution structures;  
• "Idealized" might also indicate that strong restraints on the coordination distances were used;  
→ In the presence of an incomplete 1^st^ coordination shell:   
*1) For Mg^2+^ or Na^+^, consider adding a hydration shell and optimize the water to solute distances;*   
2) If preceding step is not conclusive, consider the presence of another ion/solvent molecule aat this position;   
→ If O3'/O5' atoms are mentioned in the 1^st^ coordination shell, this suggests a defficient local RNA backbone modelling;  
→ B-factors > 100 Å^2^ are marked as suspicious;  
→ Note that cryo-EM B-factors are not very reliable;  
→ Occupancies < 1.0 are marked as suspicious;  
→ 2Fo-Fc values < 1.0 are marked as suspicious;  
→ Fo-Fc values < -3.0 or > +3.0 are marked as suspicious with warnings at < -2.5 and > +2.5;  
  
  

| Ion | Occup. | B-factor | EM\_dens. | Ligands (too close) | Ligands (1^st^ shell) | Ligands (excl. zone) | List of contacts & potential issues |
| --- | --- | --- | --- | --- | --- | --- | --- |
| ZN:3:101 | 1.0 | 81.2 | 15.04 |  | 4 |  | N_His_3SG_Cys_ → ZN ion B\_factor > 80 ← |
| K:A:1601 | 1.0 | 30.0 | 3.36 |  | 3 |  | O_ph_2O_r_ → Coordination < 6 for K ← → 2Fo-Fc density between 1.0 and 4.0 ! |
| K:A:1602 | 1.0 | 30.0 | 4.24 | 1 | 3 |  | O_ph_2O_w_ - Exclusion zone contact(s): /O_w_/ → Coordination < 6 for K ← |
| K:A:1603 | 1.0 | 30.0 | 3.72 | 1 | 0 |  | ••• No first shell contacts ••• - Exclusion zone contact(s): /O6/ → First K coordination shell missing ← → 2Fo-Fc density between 1.0 and 4.0 ! |
| K:A:1604 | 1.0 | 30.0 | 10.02 |  | 4 |  | O_r_O_b_N_b_O_w_ → Coordination < 6 for K ← |
| K:A:1605 | 1.0 | 30.0 | 3.77 | 1 | 2 |  | 2O_b_ - Exclusion zone contact(s): /O6/ → Coordination < 6 for K ← → 2Fo-Fc density between 1.0 and 4.0 ! |
| **MG:A:1606** | 1.0 | 36.2 | 14.98 |  | 6 |  | O_ph_5O_w_ Octahedral coordination (4.4º) |
| **MG:A:1607** | 1.0 | 62.0 | 6.04 | 2 | 4 |  | O_ph_3O_w_ - Exclusion zone contact(s): /O_w_/O_w_/ → Missing octahedral coordination for MG ← → Slightly distorded coord. octahedron (7.2º) ← |
| **MG:A:1608** | 1.0 | 65.4 | 4.09 | 1 | 5 |  | O_ph_4O_w_ - Exclusion zone contact(s): /O_w_/ → Missing octahedral coordination for MG ← → Slightly distorded coord. octahedron (6.1º) ← |
| **MG:A:1609** | 1.0 | 38.6 | 10.91 |  | 0 | 9 | ••• No first shell contacts ••• - Exclusion zone contact(s): /O4/O6/O6/O_w_/O_w_/ → Consider K in place of MG (3 valid contact(s) for K vs. 0 for MG) |
| **MG:A:1610** | 1.0 | 63.7 | 4.16 | 3 | 3 |  | 3O_w_ - Exclusion zone contact(s): /O_w_/O_w_/O_w_/ → Missing octahedral coordination for MG ← → Slightly distorded coord. octahedron (7.3º) ← |
| **MG:A:1611** | 1.0 | 61.0 | 2.46 |  | 2 | 2 | 2O_w_ - Exclusion zone contact(s): /O_w_/O_w_/ → Missing octahedral coordination for MG ← → 2Fo-Fc density between 1.0 and 4.0 ! |
| **MG:A:1612** | 1.0 | 52.7 | 5.33 |  | 3 | 2 | N_b_2O_w_ - Exclusion zone contact(s): /O_w_/ → Missing octahedral coordination for MG ← |
| **MG:A:1613** | 1.0 | 45.0 | 11.34 | 2 | 3 | 1 | 3O_w_ - Exclusion zone contact(s): /O_w_/O_w_/O_w_/ → Missing octahedral coordination for MG ← → Slightly distorded coord. octahedron (6.3º) ← |
| **MG:A:1614** | 1.0 | 47.1 | 8.95 | 1 | 4 |  | 4O_w_ - Exclusion zone contact(s): /O_w_/ → Missing octahedral coordination for MG ← → Slightly distorded coord. octahedron (5.6º) ← |
| **MG:A:1615** | 1.0 | 56.7 | 4.83 |  | 4 | 1 | N_b_3O_w_ → Missing octahedral coordination for MG ← Octahedral coordination (4.1º) |
| **MG:A:1616** | 1.0 | 77.8 | 1.96 |  | 3 |  | 3O_w_ → Missing octahedral coordination for MG ← → 2Fo-Fc density between 1.0 and 4.0 ! |
| **MG:A:1617** | 1.0 | 70.9 | 2.21 |  | 1 | 1 | O_w_ - Exclusion zone contact(s): /O_w_/ → Missing octahedral coordination for MG ← → 2Fo-Fc density between 1.0 and 4.0 ! |
| **MG:A:1618** | 1.0 | 61.7 | 3.98 | 1 | 1 | 7 | O_w_ - Exclusion zone contact(s): /O2'/O3'/O4/O_w_/OP2/OP2/ → Consider K in place of MG (2 valid contact(s) for K vs. 0 for MG) → Missing octahedral coordination for MG ← → 2Fo-Fc density between 1.0 and 4.0 ! |
| **MG:A:1619** | 1.0 | 55.9 | 6.31 | 2 | 4 | 1 | 2O_ph_2O_w_ - Exclusion zone contact(s): /O_w_/O_w_/O_w_/ → Missing octahedral coordination for MG ← →→→ Distorted coord. octahedron (10.3º) ←←← |
| **MG:A:1620** | 1.0 | 43.8 | 11.33 |  | 5 |  | O_ph_4O_w_ → Missing octahedral coordination for MG ← Octahedral coordination (3.2º) |
| **MG:A:1621** | 1.0 | 72.6 | 1.86 |  | 0 | 5 | ••• No first shell contacts ••• - Exclusion zone contact(s): /O6/O_w_/O_w_/O_w_/O_w_/ → Consider K in place of MG (1 valid contact(s) for K vs. 0 for MG) → 2Fo-Fc density between 1.0 and 4.0 ! |
| **MG:A:1622** | 1.0 | 67.9 | 2.04 |  | 2 | 2 | 2O_w_ - Exclusion zone contact(s): /O_w_/O_w_/ → Missing octahedral coordination for MG ← → 2Fo-Fc density between 1.0 and 4.0 ! |
| **MG:A:1623** | 1.0 | 85.6 | 3.06 | 1 | 2 |  | O_ph_O_b_ - Exclusion zone contact(s): /O_w_/ → Missing octahedral coordination for MG ← → MG ion B\_factor > 80 ← → 2Fo-Fc density between 1.0 and 4.0 ! |
| **MG:A:1624** | 1.0 | 66.9 | 2.93 | 2 | 1 | 1 | O_ph_ - Exclusion zone contact(s): /O_w_/O_w_/OP1/ → Missing octahedral coordination for MG ← → 2Fo-Fc density between 1.0 and 4.0 ! |
| **MG:A:1625** | 1.0 | 79.6 | 3.11 | 2 | 0 |  | ••• No first shell contacts ••• - Exclusion zone contact(s): /O_w_/OP2/ → First MG coordination shell missing ← → 2Fo-Fc density between 1.0 and 4.0 ! |
| **MG:A:1626** | 1.0 | 66.9 | 3.39 |  | 2 |  | 2O_ph_ → Missing octahedral coordination for MG ← → 2Fo-Fc density between 1.0 and 4.0 ! |
| **MG:A:1627** | 1.0 | 54.3 | 6.10 |  | 6 |  | O_ph_O_b_4O_w_ → Slightly distorded coord. octahedron (7.6º) ← |
| **MG:A:1628** | 1.0 | 42.2 | 11.40 |  | 5 |  | O_ph_4O_w_ → Missing octahedral coordination for MG ← Octahedral coordination (4.4º) |
| **MG:A:1629** | 1.0 | 44.2 | 9.03 | 1 | 5 |  | 5O_w_ - Exclusion zone contact(s): /OP1/ → Missing octahedral coordination for MG ← → Slightly distorded coord. octahedron (8.4º) ← |
| **MG:A:1630** | 1.0 | 56.4 | 3.72 |  | 2 | 4 | 2O_w_ - Exclusion zone contact(s): /O_w_/O_w_/OP2/ → Missing octahedral coordination for MG ← → 2Fo-Fc density between 1.0 and 4.0 ! |
| **MG:A:1631** | 1.0 | 63.4 | 3.08 | 1 | 4 | 2 | 4O_w_ - Exclusion zone contact(s): /OP2/ → Missing octahedral coordination for MG ← → Slightly distorded coord. octahedron (9.6º) ← → 2Fo-Fc density between 1.0 and 4.0 ! |
| **MG:A:1632** | 1.0 | 39.9 | 13.03 |  | 5 | 2 | 5O_w_ - Exclusion zone contact(s): /OP2/ → Missing octahedral coordination for MG ← → Slightly distorded coord. octahedron (7.0º) ← |
| **MG:A:1633** | 1.0 | 53.9 | 7.12 | 1 | 4 | 1 | O_b_3O_w_ - Exclusion zone contact(s): /O_w_/O_w_/ → Missing octahedral coordination for MG ← → Slightly distorded coord. octahedron (6.2º) ← |
| **MG:A:1634** | 1.0 | 37.8 | 15.06 |  | 6 | 2 | 3O_ph_3O_w_ Octahedral coordination (2.5º) |
| **MG:A:1635** | 1.0 | 70.5 | 1.79 |  | 1 | 1 | O_w_ - Exclusion zone contact(s): /O_w_/ → Missing octahedral coordination for MG ← → 2Fo-Fc density between 1.0 and 4.0 ! |
| **MG:A:1636** | 1.0 | 51.6 | 7.15 |  | 0 | 6 | ••• No first shell contacts ••• - Exclusion zone contact(s): /O4/O4/O6/O_w_/O_w_/O_w_/ → Consider K in place of MG (2 valid contact(s) for K vs. 0 for MG) |
| **MG:A:1637** | 1.0 | 63.0 | 3.29 |  | 0 | 5 | ••• No first shell contacts ••• - Exclusion zone contact(s): /O_back_/O_w_/OP1/ → First MG coordination shell missing ← → 2Fo-Fc density between 1.0 and 4.0 ! |
| **MG:A:1638** | 1.0 | 53.7 | 5.46 |  | 6 | 1 | O_ph_5O_w_ - Exclusion zone contact(s): /O_w_/ → Slightly distorded coord. octahedron (5.2º) ← |
| **MG:A:1639** | 1.0 | 52.9 | 5.39 | 1 | 3 | 1 | 3O_w_ - Exclusion zone contact(s): /O_w_/O_w_/ → Missing octahedral coordination for MG ← →→→ Distorted coord. octahedron (12.1º) ←←← |
| **MG:A:1640** | 1.0 | 51.6 | 7.91 |  | 0 | 5 | ••• No first shell contacts ••• - Exclusion zone contact(s): /NH2_Arg_/NZ_Lys_/O_w_/OP1/OP2/ → Consider K in place of MG (3 valid contact(s) for K vs. 0 for MG) |
| **MG:A:1641** | 1.0 | 54.9 | 5.95 |  | 0 | 5 | ••• No first shell contacts ••• - Exclusion zone contact(s): /O2'/O_w_/O_w_/OP1/OP2/ → Consider K in place of MG (3 valid contact(s) for K vs. 0 for MG) |
| **MG:A:1642** | 1.0 | 61.0 | 3.73 |  | 6 |  | 2O_ph_4O_w_ Octahedral coordination (4.5º)  → 2Fo-Fc density between 1.0 and 4.0 ! |
| **MG:A:1643** | 1.0 | 30.0 | 2.92 |  | 0 | 4 | ••• No first shell contacts ••• - Exclusion zone contact(s): /O_w_/O_w_/OP2/ → First MG coordination shell missing ← → 2Fo-Fc density between 1.0 and 4.0 ! |
| K:A:1644 | 1.0 | 30.0 | 4.65 | 2 | 2 |  | 2O_w_ - Exclusion zone contact(s): /N7/OP1/ → Coordination < 6 for K ← |
| K:A:1645 | 1.0 | 30.0 | 4.57 |  | 4 |  | 4O_b_ → Coordination < 6 for K ← |
| K:A:1646 | 1.0 | 30.0 | 10.44 |  | 6 |  | O_ph_O_r_O_b_3O_w_ |
| K:A:1647 | 1.0 | 30.0 | 3.14 | 2 | 0 |  | ••• No first shell contacts ••• - Exclusion zone contact(s): /O2/O6/ → Consider MG in place of K (2 valid contact(s) for MG vs. 0 for K) → 2Fo-Fc density between 1.0 and 4.0 ! |
| K:A:1648 | 1.0 | 30.0 | 3.87 |  | 4 |  | O_r_O_b_N_b_O_w_ → Coordination < 6 for K ← → 2Fo-Fc density between 1.0 and 4.0 ! |
| K:A:1649 | 1.0 | 30.0 | 3.96 |  | 4 |  | O_ph_O_r_O_b_O_w_ → Coordination < 6 for K ← → 2Fo-Fc density between 1.0 and 4.0 ! |
| K:A:1650 | 1.0 | 30.0 | 12.02 |  | 5 |  | 4O_b_O_w_ → Coordination < 6 for K ← |
| K:A:1651 | 1.0 | 30.0 | 2.28 |  | 3 |  | 2O_b_O_w_ → Coordination < 6 for K ← → 2Fo-Fc density between 1.0 and 4.0 ! |
| K:A:1652 | 1.0 | 30.0 | 13.71 | 2 | 5 |  | 2O_r_3O_w_ - Exclusion zone contact(s): /O_w_/OP1/ → Coordination < 6 for K ← |
| K:A:1653 | 1.0 | 30.0 | 8.54 |  | 4 |  | 3O_b_O_w_ → Coordination < 6 for K ← |
| K:A:1654 | 1.0 | 30.0 | 8.14 |  | 4 |  | O_ph_N_b_2O_w_ → Coordination < 6 for K ← |
| K:A:1655 | 1.0 | 30.0 | 4.12 | 1 | 4 |  | 3O_b_O_w_ - Exclusion zone contact(s): /O_w_/ → Coordination < 6 for K ← |
| K:A:1656 | 1.0 | 30.0 | 3.10 |  | 4 |  | 4O_b_ → Coordination < 6 for K ← → 2Fo-Fc density between 1.0 and 4.0 ! |
| K:A:1657 | 1.0 | 30.0 | 2.41 |  | 5 |  | O_ph_2O_b_N_b_O_w_ → Coordination < 6 for K ← → 2Fo-Fc density between 1.0 and 4.0 ! |
| K:A:1658 | 1.0 | 30.0 | 3.45 |  | 5 |  | O_ph_O_r_N_b_2O_w_ → Coordination < 6 for K ← → 2Fo-Fc density between 1.0 and 4.0 ! |
| K:A:1659 | 1.0 | 30.0 | 7.90 | 1 | 4 |  | O_ph_O_b_2O_w_ - Exclusion zone contact(s): /O_w_/ → Coordination < 6 for K ← |
| K:A:1660 | 1.0 | 30.0 | 1.89 |  | 0 |  | ••• No first shell contacts ••• → First K coordination shell missing ← → 2Fo-Fc density between 1.0 and 4.0 ! |
| K:A:1661 | 1.0 | 30.0 | 2.46 |  | 3 |  | O_ph_2O_b_ → Coordination < 6 for K ← → 2Fo-Fc density between 1.0 and 4.0 ! |
| K:A:1662 | 1.0 | 30.0 | 10.49 | 1 | 4 |  | O_ph_O_r_O_b_O_w_ - Exclusion zone contact(s): /O_w_/ → Coordination < 6 for K ← |
| K:A:1663 | 1.0 | 30.0 | 5.03 |  | 6 |  | 3O_b_3O_w_ |
| K:A:1664 | 1.0 | 30.0 | 3.80 | 2 | 1 |  | O_b_ - Exclusion zone contact(s): /N7/N7/ → Coordination < 6 for K ← → 2Fo-Fc density between 1.0 and 4.0 ! |
| K:A:1665 | 1.0 | 30.0 | 6.81 |  | 6 |  | 2O_r_2O_b_2N_b_ |
| K:A:1666 | 1.0 | 30.0 | 8.06 | 1 | 4 |  | 2O_ph_O_b_N_b_ - Exclusion zone contact(s): /O_w_/ → Coordination < 6 for K ← |
| K:A:1667 | 1.0 | 30.0 | 12.79 |  | 4 |  | 3O_b_O_w_ → Coordination < 6 for K ← |
| **MG:A:1668** | 1.0 | 61.2 | 3.32 |  | 3 | 1 | 3O_w_ - Exclusion zone contact(s): /O_w_/ → Missing octahedral coordination for MG ← → 2Fo-Fc density between 1.0 and 4.0 ! |
| **MG:A:1669** | 1.0 | 46.0 | 9.74 | 1 | 5 |  | O_ph_4O_w_ - Exclusion zone contact(s): /O_w_/ → Missing octahedral coordination for MG ← → Slightly distorded coord. octahedron (7.1º) ← |
| **MG:A:1670** | 1.0 | 60.3 | 2.18 |  | 6 |  | 6O_w_ →→→ Distorted coord. octahedron (15.2º) ←←← → 2Fo-Fc density between 1.0 and 4.0 ! |
| **MG:A:1671** | 1.0 | 35.6 | 13.72 |  | 6 | 1 | 3O_ph_3O_w_ - Exclusion zone contact(s): /O3'/ → Slightly distorded coord. octahedron (5.4º) ← |
| **MG:A:1672** | 1.0 | 37.2 | 13.91 |  | 5 | 1 | 2O_ph_3O_w_ - Exclusion zone contact(s): /OP2/ → Missing octahedral coordination for MG ← → Slightly distorded coord. octahedron (5.9º) ← |
| **MG:A:1673** | 1.0 | 38.4 | 14.70 |  | 6 |  | O_ph_5O_w_ Octahedral coordination (4.7º) |
| **MG:A:1674** | 1.0 | 37.3 | 13.56 |  | 2 | 3 | 2N_b_ - Exclusion zone contact(s): /O_w_/ → Missing octahedral coordination for MG ← |
| **MG:A:1675** | 1.0 | 45.8 | 8.91 | 1 | 5 |  | O_ph_4O_w_ - Exclusion zone contact(s): /O_w_/ → Missing octahedral coordination for MG ← Octahedral coordination (3.9º) |
| **MG:A:1676** | 1.0 | 50.7 | 6.41 | 2 | 4 |  | 4O_w_ - Exclusion zone contact(s): /O_w_/O_w_/ → Missing octahedral coordination for MG ← → Slightly distorded coord. octahedron (5.2º) ← |
| **MG:A:1677** | 1.0 | 66.8 | 3.94 |  | 4 |  | O_ph_3O_w_ → Missing octahedral coordination for MG ← →→→ Distorted coord. octahedron (11.5º) ←←← → 2Fo-Fc density between 1.0 and 4.0 ! |
| **MG:A:1678** | 1.0 | 68.1 | 4.42 | 2 | 4 |  | 4O_w_ - Exclusion zone contact(s): /O_w_/O_w_/ → Missing octahedral coordination for MG ← → Slightly distorded coord. octahedron (6.3º) ← |
| **MG:A:1679** | 1.0 | 69.9 | 3.48 |  | 5 | 1 | 2O_ph_3O_w_ → Missing octahedral coordination for MG ← → Slightly distorded coord. octahedron (7.0º) ← → 2Fo-Fc density between 1.0 and 4.0 ! |
| **MG:A:1680** | 1.0 | 61.9 | 5.13 |  | 5 |  | O_ph_4O_w_ → Missing octahedral coordination for MG ← → Slightly distorded coord. octahedron (5.1º) ← |
| **MG:A:1681** | 1.0 | 47.7 | 9.46 |  | 6 |  | 3O_ph_3O_w_ Octahedral coordination (4.7º) |
| **MG:A:1682** | 1.0 | 51.9 | 6.96 |  | 0 | 10 | ••• No first shell contacts ••• - Exclusion zone contact(s): /O6/O6/O_w_/O_w_/O_w_/O_w_/O_w_/O_w_/ → Consider K in place of MG (2 valid contact(s) for K vs. 0 for MG) |
| **MG:A:1683** | 1.0 | 57.1 | 5.45 | 2 | 4 |  | 2O_ph_2O_w_ - Exclusion zone contact(s): /O_w_/O_w_/ → Missing octahedral coordination for MG ← →→→ Distorted coord. octahedron (15.7º) ←←← |
| **MG:A:1684** | 1.0 | 40.5 | 11.83 |  | 6 |  | 6O_w_ Octahedral coordination (4.8º) |
| **MG:A:1685** | 1.0 | 71.3 | 2.64 |  | 2 | 1 | 2O_w_ - Exclusion zone contact(s): /O_w_/ → Missing octahedral coordination for MG ← → 2Fo-Fc density between 1.0 and 4.0 ! |
| **MG:A:1686** | 1.0 | 65.2 | 4.38 | 2 | 4 |  | O_ph_3O_w_ - Exclusion zone contact(s): /O_w_/O_w_/ → Missing octahedral coordination for MG ← → Slightly distorded coord. octahedron (9.9º) ← |
| **MG:A:1687** | 1.0 | 80.1 | 2.52 |  | 4 |  | 4O_w_ → Missing octahedral coordination for MG ← → MG ion B\_factor > 80 ← → Slightly distorded coord. octahedron (7.2º) ← → 2Fo-Fc density between 1.0 and 4.0 ! |
| **MG:A:1688** | 1.0 | 76.1 | 1.75 |  | 3 |  | 3O_w_ → Missing octahedral coordination for MG ← → 2Fo-Fc density between 1.0 and 4.0 ! |
| **MG:A:1689** | 1.0 | 80.7 | 3.56 | 1 | 5 | 3 | O_ph_2O_b_2O_w_ - Exclusion zone contact(s): /O5'/OP1/ → Missing octahedral coordination for MG ← →→→ Distorted coord. octahedron (14.3º) ←←← → MG ion B\_factor > 80 ← → 2Fo-Fc density between 1.0 and 4.0 ! |
| **MG:A:1690** | 1.0 | 70.6 | 3.27 | 1 | 2 | 1 | 2O_w_ - Exclusion zone contact(s): /O6/O_w_/ → Missing octahedral coordination for MG ← → 2Fo-Fc density between 1.0 and 4.0 ! |
| **MG:A:1691** | 1.0 | 73.6 | 3.39 | 2 | 3 | 2 | O_ph_2O_w_ - Exclusion zone contact(s): /O_w_/O_w_/OP2/ → Missing octahedral coordination for MG ← → Slightly distorded coord. octahedron (7.5º) ← → 2Fo-Fc density between 1.0 and 4.0 ! |
| **MG:A:1692** | 1.0 | 73.4 | 3.38 | 2 | 2 | 1 | 2O_w_ - Exclusion zone contact(s): /O_w_/O_w_/O_w_/ → Missing octahedral coordination for MG ← →→→ Distorted coord. octahedron (11.6º) ←←← → 2Fo-Fc density between 1.0 and 4.0 ! |
| **MG:A:1693** | 1.0 | 61.4 | 4.45 | 2 | 3 | 1 | 3O_w_ - Exclusion zone contact(s): /O_w_/O_w_/OP2/ → Missing octahedral coordination for MG ← → Slightly distorded coord. octahedron (9.2º) ← |
| **MG:A:1694** | 1.0 | 77.0 | 2.23 |  | 2 |  | 2O_w_ → Missing octahedral coordination for MG ← → 2Fo-Fc density between 1.0 and 4.0 ! |
| **MG:A:1695** | 1.0 | 76.3 | 1.85 |  | 0 | 3 | ••• No first shell contacts ••• - Exclusion zone contact(s): /O2/OP2/ → First MG coordination shell missing ← → 2Fo-Fc density between 1.0 and 4.0 ! |
| **MG:A:1696** | 1.0 | 69.2 | 1.76 |  | 0 |  | ••• No first shell contacts ••• → First MG coordination shell missing ← → 2Fo-Fc density between 1.0 and 4.0 ! |
| **MG:A:1697** | 1.0 | 59.1 | 3.15 |  | 1 | 3 | O_w_ - Exclusion zone contact(s): /O_w_/O_w_/ → Missing octahedral coordination for MG ← → 2Fo-Fc density between 1.0 and 4.0 ! |
| **MG:A:1698** | 1.0 | 69.8 | 3.53 | 1 | 3 | 1 | 3O_w_ - Exclusion zone contact(s): /O_w_/OP2/ → Missing octahedral coordination for MG ← →→→ Distorted coord. octahedron (13.2º) ←←← → 2Fo-Fc density between 1.0 and 4.0 ! |
| **MG:A:1699** | 1.0 | 70.1 | 1.93 |  | 2 | 1 | O_ph_O_w_ - Exclusion zone contact(s): /O4/ → Missing octahedral coordination for MG ← → 2Fo-Fc density between 1.0 and 4.0 ! |
| **MG:A:1700** | 1.0 | 65.0 | 5.10 | 1 | 5 |  | 2O_ph_3O_w_ - Exclusion zone contact(s): /O_w_/ → Missing octahedral coordination for MG ← → Slightly distorded coord. octahedron (8.0º) ← |
| **MG:A:1701** | 1.0 | 59.2 | 4.56 |  | 4 | 4 | 3O_ph_O_w_ - Exclusion zone contact(s): /O_w_/O_w_/ → Missing octahedral coordination for MG ← → Slightly distorded coord. octahedron (8.4º) ← |
| **MG:A:1702** | 1.0 | 62.3 | 5.18 | 2 | 3 | 2 | 2O_ph_O_w_ - Exclusion zone contact(s): /O_w_/O_w_/O_w_/ → Missing octahedral coordination for MG ← → Slightly distorded coord. octahedron (6.8º) ← |
| **MG:A:1703** | 1.0 | 70.2 | 3.34 | 2 | 3 | 2 | 3O_w_ - Exclusion zone contact(s): /O_w_/O_w_/OP1/OP2/ → Consider K in place of MG (1 valid contact(s) for K vs. 0 for MG) → Missing octahedral coordination for MG ← → Slightly distorded coord. octahedron (9.7º) ← → 2Fo-Fc density between 1.0 and 4.0 ! |
| **MG:A:1704** | 1.0 | 73.6 | 2.33 | 1 | 1 |  | O_ph_ - Exclusion zone contact(s): /O_w_/ → Missing octahedral coordination for MG ← → 2Fo-Fc density between 1.0 and 4.0 ! |
| **MG:A:1705** | 1.0 | 72.4 | 1.82 |  | 1 | 3 | O_w_ - Exclusion zone contact(s): /N7/O_w_/O_w_/ → Missing octahedral coordination for MG ← → 2Fo-Fc density between 1.0 and 4.0 ! |
| **MG:A:1706** | 1.0 | 61.1 | 5.17 | 4 | 2 |  | O_ph_O_w_ - Exclusion zone contact(s): /O_w_/O_w_/O_w_/OP1/ → Missing octahedral coordination for MG ← → Slightly distorded coord. octahedron (9.7º) ← |
| **MG:A:1707** | 1.0 | 61.2 | 3.22 | 2 | 1 | 2 | O_w_ - Exclusion zone contact(s): /O_w_/O_w_/O_w_/OP2/ → Missing octahedral coordination for MG ← → 2Fo-Fc density between 1.0 and 4.0 ! |
| **MG:A:1708** | 1.0 | 69.6 | 3.16 | 2 | 3 | 2 | O_b_2O_w_ - Exclusion zone contact(s): /O6/O_w_/OP2/ → Missing octahedral coordination for MG ← →→→ Distorted coord. octahedron (13.2º) ←←← → 2Fo-Fc density between 1.0 and 4.0 ! |
| **MG:A:1709** | 1.0 | 73.9 | 1.80 |  | 1 | 3 | O_w_ - Exclusion zone contact(s): /N7/ → Consider K in place of MG (1 valid contact(s) for K vs. 0 for MG) → Missing octahedral coordination for MG ← → 2Fo-Fc density between 1.0 and 4.0 ! |
| **MG:A:1710** | 1.0 | 58.4 | 4.54 | 2 | 2 | 2 | O_b_O_w_ - Exclusion zone contact(s): /O_w_/O_w_/OP2/ → Missing octahedral coordination for MG ← Octahedral coordination (4.4º) |
| **MG:A:1711** | 1.0 | 75.3 | 1.05 |  | 0 |  | ••• No first shell contacts ••• → First MG coordination shell missing ← → 2Fo-Fc density between 1.0 and 4.0 ! |
| **MG:A:1712** | 1.0 | 80.3 | 1.74 |  | 1 |  | O_w_ → Missing octahedral coordination for MG ← → MG ion B\_factor > 80 ← → 2Fo-Fc density between 1.0 and 4.0 ! |
| **MG:A:1713** | 1.0 | 74.9 | 2.51 |  | 5 |  | O_ph_4O_w_ → Missing octahedral coordination for MG ← Octahedral coordination (4.8º)  → 2Fo-Fc density between 1.0 and 4.0 ! |
| **MG:A:1714** | 1.0 | 65.8 | 3.62 |  | 0 | 7 | ••• No first shell contacts ••• - Exclusion zone contact(s): /O2'/O3'/O_w_/O_w_/O_w_/O_w_/O_w_/ → Consider K in place of MG (1 valid contact(s) for K vs. 0 for MG) → 2Fo-Fc density between 1.0 and 4.0 ! |
| **MG:A:1715** | 1.0 | 82.4 | 1.31 |  | 0 |  | ••• No first shell contacts ••• → First MG coordination shell missing ← → MG ion B\_factor > 80 ← → 2Fo-Fc density between 1.0 and 4.0 ! |
| **MG:A:1716** | 1.0 | 72.1 | 3.97 | 2 | 4 |  | O_ph_3O_w_ - Exclusion zone contact(s): /O_w_/O_w_/ → Missing octahedral coordination for MG ← → Slightly distorded coord. octahedron (6.2º) ← → 2Fo-Fc density between 1.0 and 4.0 ! |
| **MG:A:1717** | 1.0 | 76.0 | 2.89 |  | 2 | 1 | 2O_ph_ - Exclusion zone contact(s): /O_w_/ → Missing octahedral coordination for MG ← → 2Fo-Fc density between 1.0 and 4.0 ! |
| **MG:A:1718** | 1.0 | 65.1 | 1.58 |  | 0 | 1 | ••• No first shell contacts ••• → First MG coordination shell missing ← → 2Fo-Fc density between 1.0 and 4.0 ! |
| **MG:A:1719** | 1.0 | 74.2 | 1.68 |  | 0 | 1 | ••• No first shell contacts ••• - Exclusion zone contact(s): /O_w_/ → First MG coordination shell missing ← → 2Fo-Fc density between 1.0 and 4.0 ! |
| **MG:A:1720** | 1.0 | 75.6 | 1.95 |  | 1 |  | O_ph_ → Missing octahedral coordination for MG ← → 2Fo-Fc density between 1.0 and 4.0 ! |
| **MG:A:1721** | 1.0 | 68.1 | 1.82 |  | 0 | 2 | ••• No first shell contacts ••• - Exclusion zone contact(s): /O_w_/OP2/ → Consider K in place of MG (1 valid contact(s) for K vs. 0 for MG) → 2Fo-Fc density between 1.0 and 4.0 ! |
| K:A:1722 | 1.0 | 30.0 | 4.05 |  | 5 |  | O_b_2N_b_O_back_O_cno_ → Coordination < 6 for K ← |
| K:A:1723 | 1.0 | 30.0 | 3.85 |  | 3 |  | O_ph_O_r_O_back_ → Coordination < 6 for K ← → 2Fo-Fc density between 1.0 and 4.0 ! |
| K:A:1724 | 1.0 | 30.0 | 21.79 |  | 7 |  | 3O_ph_O_r_3O_w_ |
| K:A:1725 | 1.0 | 30.0 | 6.14 |  | 4 |  | O_r_O_b_N_b_O_w_ → Coordination < 6 for K ← |
| K:A:1726 | 1.0 | 30.0 | 5.56 |  | 3 |  | O_r_O_b_Anion → Coordination < 6 for K ← |
| K:A:1727 | 1.0 | 30.0 | 6.36 | 1 | 4 |  | 2O_b_2O_w_ - Exclusion zone contact(s): /O6/ → Coordination < 6 for K ← |
| K:A:1728 | 1.0 | 30.0 | 7.60 | 1 | 4 |  | O_r_3O_b_ - Exclusion zone contact(s): /O2/ → Coordination < 6 for K ← |
| K:A:1729 | 1.0 | 30.0 | 4.50 |  | 5 |  | 2O_ph_2O_r_O_w_ → Coordination < 6 for K ← |
| K:A:1730 | 1.0 | 30.0 | 9.42 |  | 5 |  | 3O_ph_O_r_O_w_ → Coordination < 6 for K ← |
| K:A:1731 | 1.0 | 30.0 | 11.87 |  | 5 |  | 3O_b_2O_w_ → Coordination < 6 for K ← |
| K:A:1732 | 1.0 | 30.0 | 7.93 |  | 5 |  | 2O_b_3O_w_ → Coordination < 6 for K ← |
| K:A:1733 | 1.0 | 30.0 | 14.26 |  | 7 |  | 4O_r_O_b_N_b_O_w_ |
| K:A:1734 | 1.0 | 30.0 | 5.31 |  | 4 |  | 3O_b_O_w_ → Coordination < 6 for K ← |
| K:A:1735 | 1.0 | 30.0 | 7.16 | 2 | 6 |  | 2O_ph_2O_r_2O_w_ - Exclusion zone contact(s): /O_w_/ |
| K:A:1736 | 1.0 | 30.0 | 4.13 |  | 4 |  | 2O_ph_O_r_O_w_ → Coordination < 6 for K ← |
| K:A:1737 | 1.0 | 30.0 | 4.22 | 2 | 0 |  | ••• No first shell contacts ••• - Exclusion zone contact(s): /OP1/OP2/ → Consider MG in place of K (1 valid contact(s) for MG vs. 0 for K) |
| K:A:1738 | 1.0 | 30.0 | 8.28 | 1 | 6 |  | 3O_b_3O_w_ - Exclusion zone contact(s): /O_w_/ |
| K:A:1739 | 1.0 | 30.0 | 8.20 |  | 4 |  | 4O_b_ → Coordination < 6 for K ← |
| K:A:1740 | 1.0 | 30.0 | 9.09 |  | 5 |  | 3O_b_2O_w_ → Coordination < 6 for K ← |
| **MG:A:1741** | 1.0 | 66.4 | 2.50 |  | 3 | 1 | 3O_w_ - Exclusion zone contact(s): /O_w_/ → Missing octahedral coordination for MG ← → 2Fo-Fc density between 1.0 and 4.0 ! |
| **MG:A:1742** | 1.0 | 64.8 | 3.73 |  | 5 | 1 | O_ph_4O_w_ - Exclusion zone contact(s): /O_w_/ → Missing octahedral coordination for MG ← → Slightly distorded coord. octahedron (7.3º) ← → 2Fo-Fc density between 1.0 and 4.0 ! |
| **MG:A:1743** | 1.0 | 82.6 | 2.51 |  | 6 | 1 | O_ph_O_b_4O_w_ →→→ Distorted coord. octahedron (17.6º) ←←← → MG ion B\_factor > 80 ← → 2Fo-Fc density between 1.0 and 4.0 ! |
| **MG:A:1744** | 1.0 | 72.5 | 2.35 |  | 0 | 10 | ••• No first shell contacts ••• - Exclusion zone contact(s): /N4/O4/O6/O6/O_w_/O_w_/O_w_/ → Consider K in place of MG (2 valid contact(s) for K vs. 0 for MG) → 2Fo-Fc density between 1.0 and 4.0 ! |
| **MG:A:1745** | 1.0 | 61.7 | 2.86 |  | 6 |  | 6O_w_ →→→ Distorted coord. octahedron (16.9º) ←←← → 2Fo-Fc density between 1.0 and 4.0 ! |
| **MG:X:101** | 1.0 | 59.5 | 4.52 |  | 3 | 2 | 3O_w_ - Exclusion zone contact(s): /O_w_/O_w_/ → Missing octahedral coordination for MG ← |
| **MG:a:3001** | 1.0 | 41.9 | 12.42 |  | 5 |  | 5O_w_ → Missing octahedral coordination for MG ← → Slightly distorded coord. octahedron (6.6º) ← |
| **MG:a:3002** | 1.0 | 49.0 | 7.56 |  | 1 | 3 | O_w_ - Exclusion zone contact(s): /O_w_/O_w_/O_w_/ → Missing octahedral coordination for MG ← |
| **MG:a:3003** | 1.0 | 32.9 | 17.01 |  | 4 | 2 | O_ph_3O_w_ - Exclusion zone contact(s): /O_w_/O_w_/ → Missing octahedral coordination for MG ← → Slightly distorded coord. octahedron (7.5º) ← |
| **MG:a:3004** | 1.0 | 38.0 | 12.06 |  | 3 | 2 | O_ph_2O_w_ - Exclusion zone contact(s): /O_w_/OP2/ → Missing octahedral coordination for MG ← |
| **MG:a:3005** | 1.0 | 25.6 | 19.64 |  | 0 | 7 | ••• No first shell contacts ••• - Exclusion zone contact(s): /O4/O6/O6/O6/O_w_/O_w_/O_w_/ → Consider K in place of MG (4 valid contact(s) for K vs. 0 for MG) |
| **MG:a:3006** | 1.0 | 54.6 | 6.84 | 1 | 2 | 2 | 2O_w_ - Exclusion zone contact(s): /O_w_/O_w_/O_w_/ → Missing octahedral coordination for MG ← |
| **MG:a:3007** | 1.0 | 40.1 | 12.79 |  | 5 | 1 | O_ph_4O_w_ - Exclusion zone contact(s): /O_w_/ → Missing octahedral coordination for MG ← → Slightly distorded coord. octahedron (7.1º) ← |
| **MG:a:3008** | 1.0 | 42.3 | 13.35 |  | 6 |  | O_b_5O_w_ → Slightly distorded coord. octahedron (5.1º) ← |
| **MG:a:3009** | 1.0 | 39.8 | 13.00 |  | 4 | 2 | O_ph_3O_w_ - Exclusion zone contact(s): /O_w_/O_w_/ → Missing octahedral coordination for MG ← → Slightly distorded coord. octahedron (5.9º) ← |
| **MG:a:3010** | 1.0 | 53.1 | 7.11 | 1 | 3 | 2 | O_b_2O_w_ - Exclusion zone contact(s): /O_w_/O_w_/O_w_/ → Missing octahedral coordination for MG ← → Slightly distorded coord. octahedron (8.6º) ← |
| **MG:a:3011** | 1.0 | 52.4 | 6.55 | 1 | 1 | 3 | O_w_ - Exclusion zone contact(s): /O_w_/O_w_/O_w_/ → Missing octahedral coordination for MG ← |
| **MG:a:3012** | 1.0 | 51.3 | 6.47 |  | 1 | 1 | O_w_ - Exclusion zone contact(s): /OP2/ → Missing octahedral coordination for MG ← |
| **MG:a:3013** | 1.0 | 52.2 | 5.03 |  | 4 | 1 | 4O_w_ - Exclusion zone contact(s): /O_w_/ → Missing octahedral coordination for MG ← →→→ Distorted coord. octahedron (10.6º) ←←← |
| **MG:a:3014** | 1.0 | 45.7 | 9.65 |  | 3 | 3 | 3O_w_ - Exclusion zone contact(s): /N7/O_w_/O_w_/ → Missing octahedral coordination for MG ← |
| **MG:a:3015** | 1.0 | 44.0 | 12.18 |  | 1 | 1 | O_w_ - Exclusion zone contact(s): /OP2/ → Missing octahedral coordination for MG ← |
| **MG:a:3016** | 1.0 | 44.6 | 11.62 |  | 6 |  | 6O_w_ Octahedral coordination (4.5º) |
| **MG:a:3017** | 1.0 | 49.3 | 5.81 |  | 1 | 1 | O_w_ - Exclusion zone contact(s): /O_w_/ → Missing octahedral coordination for MG ← |
| **MG:a:3018** | 1.0 | 47.3 | 7.14 |  | 0 | 6 | ••• No first shell contacts ••• - Exclusion zone contact(s): /O4/O6/O6/O_w_/O_w_/O_w_/ → Consider K in place of MG (2 valid contact(s) for K vs. 0 for MG) |
| **MG:a:3019** | 1.0 | 52.5 | 9.76 |  | 6 |  | 6O_w_ → Slightly distorded coord. octahedron (5.8º) ← |
| **MG:a:3020** | 1.0 | 42.7 | 8.86 |  | 6 | 1 | O_ph_5O_w_ - Exclusion zone contact(s): /OP2/ → Slightly distorded coord. octahedron (7.8º) ← |
| **MG:a:3021** | 1.0 | 52.7 | 9.10 |  | 5 | 1 | 5O_w_ - Exclusion zone contact(s): /O_w_/ → Missing octahedral coordination for MG ← Octahedral coordination (4.9º) |
| **MG:a:3022** | 1.0 | 51.9 | 9.69 |  | 3 | 2 | 3O_w_ - Exclusion zone contact(s): /O4/O_w_/ → Missing octahedral coordination for MG ← |
| **MG:a:3023** | 1.0 | 48.2 | 6.30 |  | 2 | 2 | 2O_ph_ - Exclusion zone contact(s): /O_w_/ → Missing octahedral coordination for MG ← |
| **MG:a:3024** | 1.0 | 41.1 | 13.10 | 1 | 4 | 1 | 4O_w_ - Exclusion zone contact(s): /O_w_/OP2/ → Missing octahedral coordination for MG ← → Slightly distorded coord. octahedron (7.2º) ← |
| **MG:a:3025** | 1.0 | 34.7 | 18.36 |  | 5 | 1 | O_ph_4O_w_ - Exclusion zone contact(s): /O_w_/ → Missing octahedral coordination for MG ← → Slightly distorded coord. octahedron (7.4º) ← |
| **MG:a:3026** | 1.0 | 61.3 | 2.90 |  | 0 |  | ••• No first shell contacts ••• → First MG coordination shell missing ← → 2Fo-Fc density between 1.0 and 4.0 ! |
| **MG:a:3027** | 1.0 | 55.9 | 5.34 |  | 4 |  | O_ph_O_b_2O_w_ → Missing octahedral coordination for MG ← → Slightly distorded coord. octahedron (6.3º) ← |
| **MG:a:3028** | 1.0 | 47.5 | 11.37 |  | 3 |  | 3O_w_ → Missing octahedral coordination for MG ← |
| **MG:a:3029** | 1.0 | 40.0 | 9.04 |  | 0 | 6 | ••• No first shell contacts ••• - Exclusion zone contact(s): /O4/O6/O_w_/O_w_/ → Consider K in place of MG (1 valid contact(s) for K vs. 0 for MG) |
| **MG:a:3030** | 1.0 | 44.0 | 8.75 |  | 6 |  | 6O_w_ Octahedral coordination (4.0º) |
| **MG:a:3031** | 1.0 | 24.6 | 23.53 |  | 0 | 8 | ••• No first shell contacts ••• - Exclusion zone contact(s): /O4/O6/O6/O_w_/O_w_/O_w_/O_w_/ → Consider K in place of MG (2 valid contact(s) for K vs. 0 for MG) |
| **MG:a:3032** | 1.0 | 25.8 | 19.71 |  | 0 | 8 | ••• No first shell contacts ••• - Exclusion zone contact(s): /O4/O4/O6/O_w_/O_w_/O_w_/OP2/ → Consider K in place of MG (4 valid contact(s) for K vs. 0 for MG) |
| **MG:a:3033** | 1.0 | 51.1 | 9.12 | 2 | 2 | 2 | 2O_w_ - Exclusion zone contact(s): /O_w_/O_w_/O_w_/OP1/ → Missing octahedral coordination for MG ← →→→ Distorted coord. octahedron (10.8º) ←←← |
| **MG:a:3034** | 1.0 | 36.8 | 18.05 |  | 5 | 1 | 5O_w_ - Exclusion zone contact(s): /O_w_/ → Missing octahedral coordination for MG ← → Slightly distorded coord. octahedron (5.4º) ← |
| **MG:a:3035** | 1.0 | 46.5 | 10.90 |  | 5 | 1 | 5O_w_ - Exclusion zone contact(s): /O_w_/ → Missing octahedral coordination for MG ← → Slightly distorded coord. octahedron (5.5º) ← |
| **MG:a:3036** | 1.0 | 48.8 | 8.74 |  | 6 |  | O_ph_5O_w_ → Slightly distorded coord. octahedron (5.8º) ← |
| **MG:a:3037** | 1.0 | 53.5 | 9.07 |  | 3 |  | 3O_w_ → Missing octahedral coordination for MG ← |
| **MG:a:3038** | 1.0 | 48.1 | 14.16 |  | 4 | 2 | 4O_w_ - Exclusion zone contact(s): /O_w_/O_w_/ → Missing octahedral coordination for MG ← → Slightly distorded coord. octahedron (6.7º) ← |
| **MG:a:3039** | 1.0 | 57.3 | 5.96 | 2 | 2 | 2 | 2O_w_ - Exclusion zone contact(s): /O_w_/O_w_/O_w_/ → Missing octahedral coordination for MG ← → Slightly distorded coord. octahedron (9.9º) ← |
| **MG:a:3040** | 1.0 | 49.7 | 8.87 |  | 6 | 1 | 2O_ph_4O_w_ → Slightly distorded coord. octahedron (7.6º) ← |
| **MG:a:3041** | 1.0 | 56.6 | 5.53 | 2 | 2 | 1 | 2O_w_ - Exclusion zone contact(s): /O_w_/O_w_/O_w_/ → Missing octahedral coordination for MG ← →→→ Distorted coord. octahedron (19.5º) ←←← |
| **MG:a:3042** | 1.0 | 47.4 | 7.50 |  | 0 | 5 | ••• No first shell contacts ••• - Exclusion zone contact(s): /O4'/O_w_/O_w_/O_w_/ → First MG coordination shell missing ← |
| **MG:a:3043** | 1.0 | 59.2 | 3.73 |  | 1 | 4 | O_w_ - Exclusion zone contact(s): /O_w_/O_w_/O_w_/O_w_/ → Missing octahedral coordination for MG ← → 2Fo-Fc density between 1.0 and 4.0 ! |
| **MG:a:3044** | 1.0 | 24.2 | 25.59 |  | 0 | 6 | ••• No first shell contacts ••• - Exclusion zone contact(s): /N3_A_/O3'/O6/O_w_/O_w_/OP1/ → Consider K in place of MG (4 valid contact(s) for K vs. 0 for MG) |
| **MG:a:3045** | 1.0 | 37.3 | 13.36 | 1 | 0 | 7 | ••• No first shell contacts ••• - Exclusion zone contact(s): /O2'/O6/O_w_/O_w_/O_w_/O_w_/O_w_/ → Consider K in place of MG (1 valid contact(s) for K vs. 0 for MG) |
| **MG:a:3046** | 1.0 | 43.8 | 7.97 |  | 4 | 2 | 2O_ph_2O_w_ - Exclusion zone contact(s): /O_w_/ → Missing octahedral coordination for MG ← →→→ Distorted coord. octahedron (11.7º) ←←← |
| **MG:a:3047** | 1.0 | 50.3 | 6.01 |  | 6 |  | O_ph_5O_w_ → Slightly distorded coord. octahedron (9.5º) ← |
| **MG:a:3048** | 1.0 | 60.1 | 2.44 |  | 6 |  | O_ph_5O_w_ → Slightly distorded coord. octahedron (5.6º) ← → 2Fo-Fc density between 1.0 and 4.0 ! |
| **MG:a:3049** | 1.0 | 55.8 | 4.04 |  | 3 | 3 | O_ph_2O_w_ - Exclusion zone contact(s): /O_w_/O_w_/O_w_/ → Missing octahedral coordination for MG ← |
| **MG:a:3050** | 1.0 | 54.7 | 3.75 |  | 4 | 1 | O_b_3O_w_ - Exclusion zone contact(s): /O_w_/ → Missing octahedral coordination for MG ← → Slightly distorded coord. octahedron (7.0º) ← → 2Fo-Fc density between 1.0 and 4.0 ! |
| **MG:a:3051** | 1.0 | 33.2 | 12.44 |  | 0 | 6 | ••• No first shell contacts ••• - Exclusion zone contact(s): /O4/O6/O6/O_w_/O_w_/ → Consider K in place of MG (3 valid contact(s) for K vs. 0 for MG) |
| **MG:a:3052** | 1.0 | 51.6 | 5.95 |  | 0 | 3 | ••• No first shell contacts ••• - Exclusion zone contact(s): /OP1/OP2/ → Consider K in place of MG (1 valid contact(s) for K vs. 0 for MG) |
| **MG:a:3053** | 1.0 | 43.8 | 9.83 |  | 0 | 7 | ••• No first shell contacts ••• - Exclusion zone contact(s): /O2'/O6/O6/O_w_/O_w_/OP2/ → Consider K in place of MG (3 valid contact(s) for K vs. 0 for MG) |
| **MG:a:3054** | 1.0 | 30.4 | 16.41 |  | 0 | 7 | ••• No first shell contacts ••• - Exclusion zone contact(s): /O3'/O3'/O_w_/O_w_/O_w_/OP1/OP2/ → Consider K in place of MG (4 valid contact(s) for K vs. 0 for MG) |
| **MG:a:3055** | 1.0 | 41.0 | 13.28 |  | 0 | 6 | ••• No first shell contacts ••• - Exclusion zone contact(s): /O_w_/O_w_/OP1/OP2/ → Consider K in place of MG (1 valid contact(s) for K vs. 0 for MG) |
| K:a:3056 | 1.0 | 30.0 | 17.72 |  | 6 |  | O_r_3O_b_2O_w_ |
| K:a:3057 | 1.0 | 30.0 | 5.17 |  | 3 |  | 2O_b_O_w_ → Coordination < 6 for K ← |
| K:a:3058 | 1.0 | 30.0 | 16.63 |  | 6 |  | 2O_b_O_back_O_cno_OH_prot_O_w_ |
| K:a:3059 | 1.0 | 30.0 | 18.85 |  | 7 |  | 2O_ph_3O_r_2O_w_ |
| K:a:3060 | 1.0 | 30.0 | 11.63 |  | 4 |  | 3O_b_O_w_ → Coordination < 6 for K ← |
| K:a:3061 | 1.0 | 30.0 | 19.06 |  | 5 |  | 3O_b_2O_w_ → Coordination < 6 for K ← |
| K:a:3062 | 1.0 | 30.0 | 15.96 | 1 | 5 |  | O_ph_O_r_O_b_2O_w_ - Exclusion zone contact(s): /O6/ → Coordination < 6 for K ← |
| K:a:3063 | 1.0 | 30.0 | 15.89 | 1 | 5 |  | 3O_ph_O_r_O_w_ → Coordination < 6 for K ← |
| K:a:3064 | 1.0 | 30.0 | 13.40 | 1 | 5 |  | O_ph_O_b_3O_w_ - Exclusion zone contact(s): /O2'/ → Coordination < 6 for K ← |
| **MG:a:3065** | 1.0 | 49.9 | 8.47 |  | 4 | 2 | 4O_w_ - Exclusion zone contact(s): /O_w_/O_w_/ → Missing octahedral coordination for MG ← → Slightly distorded coord. octahedron (5.2º) ← |
| **MG:a:3066** | 1.0 | 48.5 | 7.62 |  | 1 | 2 | O_w_ - Exclusion zone contact(s): /O_w_/OP2/ → Missing octahedral coordination for MG ← |
| **MG:a:3067** | 1.0 | 51.9 | 8.14 |  | 6 |  | 6O_w_ → Slightly distorded coord. octahedron (7.9º) ← |
| **MG:a:3068** | 1.0 | 40.5 | 11.77 |  | 0 | 7 | ••• No first shell contacts ••• - Exclusion zone contact(s): /O2'/O3'/O_w_/O_w_/OP1/OP1/ → Consider K in place of MG (2 valid contact(s) for K vs. 0 for MG) |
| **MG:a:3069** | 1.0 | 44.7 | 11.61 |  | 0 | 6 | ••• No first shell contacts ••• - Exclusion zone contact(s): /O2'/O3'/O3'/O_w_/O_w_/OP1/ → Consider K in place of MG (4 valid contact(s) for K vs. 0 for MG) |
| **MG:a:3070** | 1.0 | 44.2 | 8.08 |  | 1 | 4 | O_w_ - Exclusion zone contact(s): /NE_Arg_/O_w_/OP2/ → Consider K in place of MG (1 valid contact(s) for K vs. 0 for MG) → Missing octahedral coordination for MG ← |
| **MG:a:3071** | 1.0 | 44.4 | 7.26 | 1 | 0 | 7 | ••• No first shell contacts ••• - Exclusion zone contact(s): /O2'/O3'/O_w_/O_w_/O_w_/O_w_/ → Consider K in place of MG (2 valid contact(s) for K vs. 0 for MG) |
| **MG:a:3072** | 1.0 | 46.5 | 10.50 |  | 3 |  | O_ph_2O_w_ → Missing octahedral coordination for MG ← |
| **MG:a:3073** | 1.0 | 52.9 | 6.78 |  | 1 | 4 | O_w_ - Exclusion zone contact(s): /O_w_/O_w_/OP1/OP2/ → Consider K in place of MG (1 valid contact(s) for K vs. 0 for MG) → Missing octahedral coordination for MG ← |
| **MG:a:3074** | 1.0 | 50.5 | 6.26 | 3 | 1 | 2 | O_w_ - Exclusion zone contact(s): /O_w_/O_w_/O_w_/O_w_/O_w_/ → Missing octahedral coordination for MG ← →→→ Distorted coord. octahedron (12.4º) ←←← |
| **MG:a:3075** | 1.0 | 47.2 | 6.54 |  | 3 | 1 | 3O_w_ - Exclusion zone contact(s): /O_w_/ → Missing octahedral coordination for MG ← |
| **MG:a:3076** | 1.0 | 53.4 | 6.14 |  | 4 | 1 | 4O_w_ - Exclusion zone contact(s): /O_w_/ → Missing octahedral coordination for MG ← → Slightly distorded coord. octahedron (7.9º) ← |
| **MG:a:3077** | 1.0 | 31.5 | 17.65 |  | 4 | 1 | 4O_w_ - Exclusion zone contact(s): /O_w_/ → Missing octahedral coordination for MG ← →→→ Distorted coord. octahedron (15.5º) ←←← |
| **MG:a:3078** | 1.0 | 32.7 | 13.17 |  | 6 | 1 | O_ph_5O_w_ → Slightly distorded coord. octahedron (7.0º) ← |
| **MG:a:3079** | 1.0 | 35.3 | 14.39 |  | 4 | 2 | O_ph_3O_w_ - Exclusion zone contact(s): /O_w_/O_w_/ → Missing octahedral coordination for MG ← → Slightly distorded coord. octahedron (8.1º) ← |
| **MG:a:3080** | 1.0 | 56.3 | 8.00 |  | 6 |  | O_ph_5O_w_ → Slightly distorded coord. octahedron (8.4º) ← |
| **MG:a:3081** | 1.0 | 47.2 | 8.66 |  | 3 | 1 | O_ph_2O_w_ - Exclusion zone contact(s): /O_w_/ → Missing octahedral coordination for MG ← |
| **MG:a:3082** | 1.0 | 27.0 | 23.77 |  | 6 |  | O_ph_5O_w_ Octahedral coordination (4.1º) |
| **MG:a:3083** | 1.0 | 24.4 | 24.35 |  | 5 | 1 | O_ph_O_b_3O_w_ - Exclusion zone contact(s): /O_w_/ → Missing octahedral coordination for MG ← Octahedral coordination (4.2º) |
| **MG:a:3084** | 1.0 | 33.6 | 17.54 |  | 6 |  | 2O_ph_4O_w_ Octahedral coordination (3.2º) |
| **MG:a:3085** | 1.0 | 41.0 | 15.64 |  | 5 |  | 5O_w_ → Missing octahedral coordination for MG ← → Slightly distorded coord. octahedron (6.6º) ← |
| **MG:a:3086** | 1.0 | 23.7 | 30.08 |  | 6 |  | O_ph_5O_w_ Octahedral coordination (5.0º) |
| **MG:a:3087** | 1.0 | 26.2 | 20.85 |  | 6 | 2 | 2O_ph_4O_w_ → Slightly distorded coord. octahedron (6.9º) ← |
| **MG:a:3088** | 1.0 | 24.4 | 27.54 |  | 6 |  | 2O_ph_N_b_3O_w_ → Slightly distorded coord. octahedron (5.1º) ← |
| **MG:a:3089** | 1.0 | 28.9 | 18.78 |  | 6 |  | 3O_ph_3O_w_ Octahedral coordination (3.2º) |
| **MG:a:3090** | 1.0 | 31.3 | 19.14 |  | 6 |  | O_ph_5O_w_ Octahedral coordination (4.7º) |
| **MG:a:3091** | 1.0 | 28.5 | 20.82 | 1 | 3 | 3 | O_ph_2O_w_ - Exclusion zone contact(s): /N7/O_w_/OP2/ → Missing octahedral coordination for MG ← Octahedral coordination (4.5º) |
| **MG:a:3092** | 1.0 | 25.2 | 27.20 |  | 4 | 2 | 4O_w_ - Exclusion zone contact(s): /O_w_/O_w_/ → Missing octahedral coordination for MG ← Octahedral coordination (4.6º) |
| **MG:a:3093** | 1.0 | 31.2 | 19.38 |  | 6 | 1 | 2O_ph_4O_w_ Octahedral coordination (4.9º) |
| **MG:a:3094** | 1.0 | 25.7 | 26.89 | 1 | 3 | 2 | 3O_w_ - Exclusion zone contact(s): /O_w_/O_w_/OP2/ → Missing octahedral coordination for MG ← → Slightly distorded coord. octahedron (7.2º) ← |
| **MG:a:3095** | 1.0 | 41.6 | 15.46 |  | 6 |  | 6O_w_ → Slightly distorded coord. octahedron (8.3º) ← |
| K:a:3096 | 1.0 | 30.0 | 10.55 |  | 4 |  | O_r_O_b_2O_w_ → Coordination < 6 for K ← |
| K:a:3097 | 1.0 | 30.0 | 7.61 |  | 2 |  | O_r_O_b_ → Coordination < 6 for K ← |
| K:a:3098 | 1.0 | 30.0 | 16.97 |  | 6 |  | 3O_b_2O_w_Anion |
| K:a:3099 | 1.0 | 30.0 | 22.15 |  | 5 |  | 3O_b_2O_w_ → Coordination < 6 for K ← |
| K:a:3100 | 1.0 | 30.0 | 30.41 |  | 7 |  | 2O_ph_O_r_4O_w_ |
| K:a:3101 | 1.0 | 30.0 | 19.70 |  | 3 |  | O_ph_O_r_N_b_ → Coordination < 6 for K ← |
| K:a:3102 | 1.0 | 30.0 | 21.64 | 1 | 5 |  | 3O_b_2O_w_ - Exclusion zone contact(s): /O_w_/ → Coordination < 6 for K ← |
| K:a:3103 | 1.0 | 30.0 | 7.32 | 2 | 1 |  | O_w_ - Exclusion zone contact(s): /O_w_/O_w_/ → Coordination < 6 for K ← |
| K:a:3104 | 1.0 | 30.0 | 14.32 |  | 7 |  | 2O_ph_O_r_N_b_3O_w_ |
| K:a:3105 | 1.0 | 30.0 | 28.96 |  | 5 |  | O_r_2O_b_N_b_ → Coordination < 6 for K ← |
| K:a:3106 | 1.0 | 30.0 | 14.63 |  | 6 |  | O_b_5O_w_ |
| K:a:3107 | 1.0 | 30.0 | 15.75 | 1 | 5 |  | 3O_b_O_w_Anion - Exclusion zone contact(s): /O_w_/ → Coordination < 6 for K ← |
| K:a:3108 | 1.0 | 30.0 | 6.48 |  | 2 |  | 2O_b_ → Coordination < 6 for K ← |
| K:a:3109 | 1.0 | 30.0 | 14.01 |  | 4 |  | 2O_ph_O_r_O_w_ → Coordination < 6 for K ← |
| K:a:3110 | 1.0 | 30.0 | 19.46 |  | 5 |  | O_ph_O_r_3O_w_ → Coordination < 6 for K ← |
| K:a:3111 | 1.0 | 30.0 | 31.46 |  | 7 |  | O_ph_O_r_5O_w_ |
| K:a:3112 | 1.0 | 30.0 | 25.24 |  | 6 |  | 5O_r_O_w_ |
| K:a:3113 | 1.0 | 30.0 | 22.51 |  | 7 |  | O_r_O_b_3N_b_2O_w_ |
| K:a:3114 | 1.0 | 30.0 | 39.62 |  | 5 |  | O_r_O_b_2N_b_O_w_ → Coordination < 6 for K ← |
| K:a:3115 | 1.0 | 30.0 | 16.92 | 1 | 3 |  | 3O_b_ - Exclusion zone contact(s): /O_w_/ → Coordination < 6 for K ← |
| K:a:3116 | 1.0 | 30.0 | 31.60 |  | 6 |  | O_r_3O_b_N_b_O_w_ |
| K:a:3117 | 1.0 | 30.0 | 13.58 |  | 5 |  | 2O_ph_2O_r_O_w_ → Coordination < 6 for K ← |
| K:a:3118 | 1.0 | 30.0 | 24.34 |  | 4 |  | 2O_ph_O_r_N_b_ → Coordination < 6 for K ← |
| K:a:3119 | 1.0 | 30.0 | 25.79 |  | 8 |  | 3O_ph_O_r_4O_w_ |
| K:a:3120 | 1.0 | 30.0 | 21.95 |  | 6 |  | 3O_ph_2O_r_O_w_ |
| K:a:3121 | 1.0 | 30.0 | 24.38 |  | 6 |  | O_ph_2O_r_N_b_2O_w_ |
| K:a:3122 | 1.0 | 30.0 | 25.71 | 1 | 3 |  | O_ph_N_b_O_w_ - Exclusion zone contact(s): /O_w_/ → Coordination < 6 for K ← |
| K:a:3123 | 1.0 | 30.0 | 31.82 |  | 7 |  | N_b_6O_w_ |
| K:a:3124 | 1.0 | 30.0 | 21.05 |  | 5 |  | 4O_b_O_w_ → Coordination < 6 for K ← |
| K:a:3125 | 1.0 | 30.0 | 23.85 |  | 5 |  | 4O_b_O_w_ → Coordination < 6 for K ← |
| K:a:3126 | 1.0 | 30.0 | 36.37 | 2 | 3 |  | O_ph_O_b_O_w_ - Exclusion zone contact(s): /O_back_/OP2/ → Coordination < 6 for K ← |
| K:a:3127 | 1.0 | 30.0 | 37.54 |  | 6 |  | O_r_2O_b_3O_w_ |
| K:a:3128 | 1.0 | 30.0 | 12.55 |  | 7 |  | O_r_2O_b_4O_w_ |
| K:a:3129 | 1.0 | 30.0 | 22.27 | 1 | 5 |  | 3O_b_2O_w_ - Exclusion zone contact(s): /OP2/ → Coordination < 6 for K ← |
| K:a:3130 | 1.0 | 30.0 | 21.64 |  | 6 |  | 3O_b_3O_w_ |
| K:a:3131 | 1.0 | 30.0 | 2.77 | 1 | 0 |  | ••• No first shell contacts ••• - Exclusion zone contact(s): /O4/ → First K coordination shell missing ← → 2Fo-Fc density between 1.0 and 4.0 ! |
| K:a:3132 | 1.0 | 30.0 | 13.33 |  | 5 |  | O_ph_O_r_2O_b_O_w_ → Coordination < 6 for K ← |
| K:a:3133 | 1.0 | 30.0 | 26.17 |  | 7 |  | O_ph_2O_r_O_b_2N_b_O_w_ |
| K:a:3134 | 1.0 | 30.0 | 29.50 |  | 6 |  | O_ph_2O_r_O_back_2O_w_ |
| K:a:3135 | 1.0 | 30.0 | 19.59 |  | 6 |  | 4O_b_2O_w_ |
| K:a:3136 | 1.0 | 30.0 | 8.59 |  | 5 |  | 3O_b_O_w_ → Coordination < 6 for K ← |
| K:a:3137 | 1.0 | 30.0 | 10.45 |  | 7 |  | 2O_b_N_b_3O_w_ |
| K:a:3138 | 1.0 | 30.0 | 9.78 | 1 | 3 |  | O_ph_2O_b_ - Exclusion zone contact(s): /O_w_/ → Coordination < 6 for K ← |
| K:a:3139 | 1.0 | 30.0 | 20.81 |  | 5 |  | 3O_b_2O_w_ → Coordination < 6 for K ← |
| K:a:3140 | 1.0 | 30.0 | 18.55 |  | 6 |  | O_ph_2O_r_3O_w_ |
| K:a:3141 | 1.0 | 30.0 | 25.68 |  | 8 |  | O_ph_5O_r_O_b_O_w_ |
| K:a:3142 | 1.0 | 30.0 | 18.36 | 1 | 5 |  | 3O_b_2O_w_ - Exclusion zone contact(s): /O2'/ → Coordination < 6 for K ← |
| **MG:a:3143** | 1.0 | 35.3 | 13.95 |  | 6 | 1 | 2O_ph_4O_w_ → Slightly distorded coord. octahedron (6.1º) ← |
| **MG:a:3144** | 1.0 | 34.8 | 16.23 | 1 | 4 | 2 | 2O_ph_O_r_O_w_ - Exclusion zone contact(s): /O_w_/OP2/ → Missing octahedral coordination for MG ← → Slightly distorded coord. octahedron (6.9º) ← |
| **MG:a:3145** | 1.0 | 31.0 | 16.70 |  | 5 | 1 | 2O_ph_O_r_2O_w_ - Exclusion zone contact(s): /O_w_/ → Missing octahedral coordination for MG ← Octahedral coordination (2.8º) |
| **MG:a:3146** | 1.0 | 48.7 | 8.52 |  | 4 | 2 | 4O_w_ - Exclusion zone contact(s): /O_w_/O_w_/ → Missing octahedral coordination for MG ← → Slightly distorded coord. octahedron (6.7º) ← |
| **MG:a:3147** | 1.0 | 40.3 | 13.11 |  | 6 |  | O_ph_5O_w_ →→→ Distorted coord. octahedron (10.7º) ←←← |
| **MG:a:3148** | 1.0 | 46.8 | 5.77 |  | 0 | 7 | ••• No first shell contacts ••• - Exclusion zone contact(s): /O_w_/O_w_/OP1/OP1/OP1/ → Consider K in place of MG (2 valid contact(s) for K vs. 0 for MG) |
| **MG:a:3149** | 1.0 | 44.0 | 13.81 |  | 4 | 1 | O_ph_3O_w_ - Exclusion zone contact(s): /O_w_/ → Missing octahedral coordination for MG ← → Slightly distorded coord. octahedron (6.9º) ← |
| **MG:a:3150** | 1.0 | 28.2 | 19.66 |  | 6 | 1 | 2O_ph_4O_w_ → Slightly distorded coord. octahedron (5.9º) ← |
| **MG:a:3151** | 1.0 | 51.1 | 3.98 |  | 2 | 2 | O_ph_O_w_ - Exclusion zone contact(s): /O_w_/O_w_/ → Missing octahedral coordination for MG ← → 2Fo-Fc density between 1.0 and 4.0 ! |
| **MG:a:3152** | 1.0 | 33.4 | 14.41 |  | 4 | 3 | 2O_ph_2O_w_ - Exclusion zone contact(s): /O_w_/O_w_/ → Missing octahedral coordination for MG ← → Slightly distorded coord. octahedron (7.2º) ← |
| **MG:a:3153** | 1.0 | 29.6 | 21.76 |  | 6 |  | O_b_5O_w_ Octahedral coordination (3.4º) |
| **MG:a:3154** | 1.0 | 41.7 | 11.81 |  | 6 |  | O_ph_5O_w_ Octahedral coordination (4.9º) |
| **MG:a:3155** | 1.0 | 52.6 | 6.99 |  | 4 | 2 | O_ph_3O_w_ - Exclusion zone contact(s): /O_w_/ → Missing octahedral coordination for MG ← Octahedral coordination (4.5º) |
| **MG:a:3156** | 1.0 | 41.7 | 10.51 |  | 6 | 1 | O_ph_5O_w_ Octahedral coordination (3.9º) |
| **MG:a:3157** | 1.0 | 41.5 | 16.19 |  | 5 | 1 | 5O_w_ - Exclusion zone contact(s): /O_w_/ → Missing octahedral coordination for MG ← Octahedral coordination (4.3º) |
| **MG:a:3158** | 1.0 | 44.2 | 7.82 |  | 2 | 5 | 2O_w_ - Exclusion zone contact(s): /O4/O6/O_w_/ → Consider K in place of MG (2 valid contact(s) for K vs. 0 for MG) → Missing octahedral coordination for MG ← |
| **MG:a:3159** | 1.0 | 44.9 | 11.28 | 1 | 4 | 2 | 2O_ph_2O_w_ - Exclusion zone contact(s): /O_w_/ → Missing octahedral coordination for MG ← → Slightly distorded coord. octahedron (6.7º) ← |
| **MG:a:3160** | 1.0 | 49.6 | 7.86 |  | 3 | 1 | 3O_w_ - Exclusion zone contact(s): /O_w_/ → Missing octahedral coordination for MG ← |
| **MG:a:3161** | 1.0 | 46.7 | 6.99 |  | 6 |  | O_ph_5O_w_ Octahedral coordination (4.5º) |
| **MG:a:3162** | 1.0 | 37.6 | 9.21 |  | 0 | 4 | ••• No first shell contacts ••• - Exclusion zone contact(s): /O4/O6/O6/O6/ → Consider K in place of MG (3 valid contact(s) for K vs. 0 for MG) |
| **MG:a:3163** | 1.0 | 40.4 | 12.83 | 1 | 3 | 2 | 3O_w_ - Exclusion zone contact(s): /O_w_/O_w_/O_w_/ → Missing octahedral coordination for MG ← → Slightly distorded coord. octahedron (6.1º) ← |
| **MG:a:3164** | 1.0 | 45.4 | 9.02 |  | 5 | 1 | O_ph_4O_w_ - Exclusion zone contact(s): /O_w_/ → Missing octahedral coordination for MG ← Octahedral coordination (3.0º) |
| **MG:a:3165** | 1.0 | 39.4 | 13.36 |  | 6 |  | O_ph_5O_w_ Octahedral coordination (3.7º) |
| **MG:a:3166** | 1.0 | 48.6 | 6.99 |  | 4 | 2 | O_ph_3O_w_ - Exclusion zone contact(s): /O_w_/ → Missing octahedral coordination for MG ← → Slightly distorded coord. octahedron (9.8º) ← |
| **MG:a:3167** | 1.0 | 29.0 | 21.82 |  | 4 | 3 | 2O_ph_2O_w_ - Exclusion zone contact(s): /O_w_/O_w_/ → Missing octahedral coordination for MG ← → Slightly distorded coord. octahedron (6.7º) ← |
| **MG:a:3168** | 1.0 | 54.0 | 4.11 | 1 | 1 | 2 | O_w_ - Exclusion zone contact(s): /O_w_/O_w_/O_w_/ → Missing octahedral coordination for MG ← |
| **MG:a:3169** | 1.0 | 41.3 | 10.93 |  | 0 | 8 | ••• No first shell contacts ••• - Exclusion zone contact(s): /N3_C_/ → Consider K in place of MG (2 valid contact(s) for K vs. 0 for MG) |
| **MG:a:3170** | 1.0 | 30.9 | 20.65 |  | 5 |  | O_ph_N_b_3O_w_ → Missing octahedral coordination for MG ← → Slightly distorded coord. octahedron (9.3º) ← |
| **MG:a:3171** | 1.0 | 38.6 | 14.33 |  | 4 | 1 | 4O_w_ - Exclusion zone contact(s): /O_w_/ → Missing octahedral coordination for MG ← → Slightly distorded coord. octahedron (7.6º) ← |
| **MG:a:3172** | 1.0 | 52.7 | 12.18 |  | 4 | 2 | N_b_3O_w_ - Exclusion zone contact(s): /O_w_/ → Missing octahedral coordination for MG ← → Slightly distorded coord. octahedron (5.4º) ← |
| **MG:a:3173** | 1.0 | 39.7 | 11.15 |  | 0 | 8 | ••• No first shell contacts ••• - Exclusion zone contact(s): /O2/O2'/O_w_/O_w_/O_w_/OP2/ → Consider K in place of MG (3 valid contact(s) for K vs. 0 for MG) |
| **MG:a:3174** | 1.0 | 40.5 | 10.71 |  | 6 | 1 | O_ph_5O_w_ - Exclusion zone contact(s): /O4/ →→→ Distorted coord. octahedron (11.0º) ←←← |
| K:a:3175 | 1.0 | 30.0 | 5.33 | 1 | 2 |  | 2O_b_ - Exclusion zone contact(s): /O_w_/ → Coordination < 6 for K ← |
| K:a:3176 | 1.0 | 30.0 | 15.71 | 1 | 5 |  | 2O_b_O_back_O_w_Anion - Exclusion zone contact(s): /O_w_/ → Coordination < 6 for K ← |
| K:a:3177 | 1.0 | 30.0 | 25.49 |  | 6 |  | 3O_b_N_b_2O_w_ |
| K:a:3178 | 1.0 | 30.0 | 21.05 |  | 5 |  | 2O_ph_OH_prot_2O_w_ → Coordination < 6 for K ← |
| K:a:3179 | 1.0 | 30.0 | 25.86 |  | 7 |  | 2O_r_2O_b_N_b_2O_w_ |
| K:a:3180 | 1.0 | 30.0 | 25.84 |  | 6 |  | 3O_b_3O_w_ |
| K:a:3181 | 1.0 | 30.0 | 25.41 | 2 | 4 |  | O_r_O_back_2O_w_ - Exclusion zone contact(s): /O2/O6/ → Coordination < 6 for K ← |
| **MG:a:3182** | 1.0 | 45.3 | 10.46 |  | 6 |  | 6O_w_ → Slightly distorded coord. octahedron (5.6º) ← |
| **MG:a:3183** | 1.0 | 31.9 | 14.73 | 1 | 5 | 1 | O_ph_4O_w_ - Exclusion zone contact(s): /O_w_/ → Missing octahedral coordination for MG ← Octahedral coordination (4.6º) |
| **MG:a:3184** | 1.0 | 31.8 | 20.47 |  | 5 | 1 | 5O_w_ - Exclusion zone contact(s): /O_w_/ → Missing octahedral coordination for MG ← → Slightly distorded coord. octahedron (5.1º) ← |
| **MG:a:3185** | 1.0 | 42.5 | 10.66 |  | 6 |  | O_ph_5O_w_ → Slightly distorded coord. octahedron (5.1º) ← |
| **MG:a:3186** | 1.0 | 26.6 | 22.58 |  | 6 | 1 | 2O_ph_O_back_2O_w_N_His_ Octahedral coordination (3.2º) |
| **MG:a:3187** | 1.0 | 45.5 | 10.38 | 1 | 4 | 2 | O_ph_3O_w_ - Exclusion zone contact(s): /O3'/O_w_/OP1/ → Missing octahedral coordination for MG ← → Slightly distorded coord. octahedron (8.8º) ← |
| **MG:a:3188** | 1.0 | 42.1 | 13.47 |  | 6 |  | O_ph_5O_w_ Octahedral coordination (4.2º) |
| **MG:a:3189** | 1.0 | 52.9 | 5.94 |  | 2 | 1 | 2O_w_ - Exclusion zone contact(s): /O_w_/ → Missing octahedral coordination for MG ← |
| **MG:a:3190** | 1.0 | 51.2 | 5.95 |  | 3 |  | 3O_w_ → Missing octahedral coordination for MG ← |
| **MG:a:3191** | 1.0 | 58.8 | 3.43 |  | 1 |  | O_w_ → Missing octahedral coordination for MG ← → 2Fo-Fc density between 1.0 and 4.0 ! |
| **MG:a:3192** | 1.0 | 41.5 | 13.91 | 1 | 3 | 2 | 2O_ph_O_w_ - Exclusion zone contact(s): /O_w_/O_w_/ → Missing octahedral coordination for MG ← Octahedral coordination (3.3º) |
| **MG:a:3193** | 1.0 | 44.8 | 11.00 |  | 6 |  | O_ph_5O_w_ → Slightly distorded coord. octahedron (5.8º) ← |
| **MG:a:3194** | 1.0 | 27.2 | 28.04 |  | 6 | 1 | 2O_ph_4O_w_ Octahedral coordination (4.4º) |
| **MG:a:3195** | 1.0 | 47.5 | 8.46 |  | 3 | 1 | O_ph_2O_w_ - Exclusion zone contact(s): /O_w_/ → Missing octahedral coordination for MG ← |
| **MG:a:3196** | 1.0 | 45.3 | 7.06 |  | 1 | 3 | N_b_ - Exclusion zone contact(s): /O6/O_w_/O_w_/ → Missing octahedral coordination for MG ← |
| **MG:a:3197** | 1.0 | 47.2 | 11.53 |  | 5 | 1 | 5O_w_ → Missing octahedral coordination for MG ← → Slightly distorded coord. octahedron (7.7º) ← |
| **MG:a:3198** | 1.0 | 40.9 | 14.13 |  | 6 |  | O_r_5O_w_ Octahedral coordination (5.0º) |
| **MG:a:3199** | 1.0 | 28.6 | 23.09 |  | 6 |  | O_ph_5O_w_ Octahedral coordination (4.9º) |
| **MG:a:3200** | 1.0 | 33.6 | 13.89 | 1 | 5 |  | O_ph_4O_w_ - Exclusion zone contact(s): /O_w_/ → Missing octahedral coordination for MG ← Octahedral coordination (4.2º) |
| **MG:a:3201** | 1.0 | 24.5 | 25.16 |  | 6 | 2 | 2O_ph_O_b_3O_w_ → Slightly distorded coord. octahedron (5.2º) ← |
| **MG:a:3202** | 1.0 | 29.6 | 19.97 | 1 | 3 | 1 | 3O_w_ - Exclusion zone contact(s): /O_w_/OP1/ → Missing octahedral coordination for MG ← → Slightly distorded coord. octahedron (6.8º) ← |
| **MG:a:3203** | 1.0 | 29.2 | 18.40 |  | 0 | 7 | ••• No first shell contacts ••• - Exclusion zone contact(s): /O2'/O3'/O4/O_w_/O_w_/O_w_/OP2/ → Consider K in place of MG (4 valid contact(s) for K vs. 0 for MG) |
| **MG:a:3204** | 1.0 | 43.4 | 17.39 |  | 4 |  | O_ph_3O_w_ → Missing octahedral coordination for MG ← Octahedral coordination (2.8º) |
| **MG:a:3205** | 1.0 | 33.1 | 19.97 |  | 6 |  | 6O_w_ Octahedral coordination (4.0º) |
| **MG:a:3206** | 1.0 | 51.3 | 9.73 |  | 5 | 1 | O_ph_4O_w_ - Exclusion zone contact(s): /O_w_/ → Missing octahedral coordination for MG ← → Slightly distorded coord. octahedron (6.6º) ← |
| **MG:a:3207** | 1.0 | 31.3 | 19.18 |  | 6 |  | 2O_ph_4O_w_ → Slightly distorded coord. octahedron (5.9º) ← |
| **MG:a:3208** | 1.0 | 56.1 | 3.90 | 1 | 2 | 1 | O_b_O_w_ - Exclusion zone contact(s): /O_w_/O_w_/ → Missing octahedral coordination for MG ← → 2Fo-Fc density between 1.0 and 4.0 ! |
| **MG:a:3209** | 1.0 | 40.1 | 13.99 |  | 4 |  | O_ph_3O_w_ → Missing octahedral coordination for MG ← → Slightly distorded coord. octahedron (6.3º) ← |
| **MG:a:3210** | 1.0 | 43.2 | 11.27 |  | 6 |  | 6O_w_ Octahedral coordination (3.7º) |
| **MG:a:3211** | 1.0 | 35.9 | 16.39 |  | 5 |  | O_b_4O_w_ → Missing octahedral coordination for MG ← → Slightly distorded coord. octahedron (8.7º) ← |
| **MG:a:3212** | 1.0 | 37.2 | 15.10 |  | 6 |  | 6O_w_ Octahedral coordination (3.1º) |
| **MG:a:3213** | 1.0 | 37.6 | 15.21 |  | 5 | 1 | 5O_w_ - Exclusion zone contact(s): /O_w_/ → Missing octahedral coordination for MG ← Octahedral coordination (4.7º) |
| **MG:a:3214** | 1.0 | 25.6 | 26.25 |  | 4 | 2 | 4O_w_ - Exclusion zone contact(s): /O_w_/O_w_/ → Missing octahedral coordination for MG ← → Slightly distorded coord. octahedron (6.0º) ← |
| **MG:a:3215** | 1.0 | 26.4 | 27.58 |  | 4 | 2 | 4O_w_ - Exclusion zone contact(s): /O_w_/OP2/ → Missing octahedral coordination for MG ← Octahedral coordination (3.4º) |
| **MG:a:3216** | 1.0 | 33.3 | 22.55 |  | 6 |  | 6O_w_ Octahedral coordination (3.9º) |
| **MG:a:3217** | 1.0 | 26.8 | 25.15 |  | 5 | 1 | 5O_w_ - Exclusion zone contact(s): /O_w_/ → Missing octahedral coordination for MG ← → Slightly distorded coord. octahedron (5.4º) ← |
| **MG:a:3218** | 1.0 | 38.8 | 12.18 |  | 5 |  | 5O_w_ → Missing octahedral coordination for MG ← Octahedral coordination (4.9º) |
| **MG:a:3219** | 1.0 | 39.0 | 12.40 |  | 6 | 1 | 2O_ph_O_cno_3O_w_ → Slightly distorded coord. octahedron (6.2º) ← |
| **MG:a:3220** | 1.0 | 22.8 | 28.90 |  | 5 | 1 | N_b_4O_w_ - Exclusion zone contact(s): /N7/ → Missing octahedral coordination for MG ← Octahedral coordination (2.2º) |
| **MG:a:3221** | 1.0 | 33.8 | 17.77 |  | 6 |  | 6O_w_ Octahedral coordination (3.6º) |
| **MG:a:3222** | 1.0 | 35.8 | 18.25 |  | 5 | 1 | O_b_4O_w_ - Exclusion zone contact(s): /O_w_/ → Missing octahedral coordination for MG ← Octahedral coordination (4.0º) |
| **MG:a:3223** | 1.0 | 23.4 | 25.39 |  | 5 | 1 | 2O_ph_3O_w_ - Exclusion zone contact(s): /O_w_/ → Missing octahedral coordination for MG ← Octahedral coordination (3.9º) |
| **MG:a:3224** | 1.0 | 24.7 | 27.42 |  | 5 | 1 | 3O_ph_2O_w_ - Exclusion zone contact(s): /O_w_/ → Missing octahedral coordination for MG ← → Slightly distorded coord. octahedron (8.7º) ← |
| **MG:a:3225** | 1.0 | 26.7 | 24.33 |  | 5 | 1 | 2O_ph_3O_w_ - Exclusion zone contact(s): /O_w_/ → Missing octahedral coordination for MG ← → Slightly distorded coord. octahedron (6.5º) ← |
| **MG:a:3226** | 1.0 | 29.3 | 21.27 |  | 5 | 1 | O_ph_4O_w_ - Exclusion zone contact(s): /O_w_/ → Missing octahedral coordination for MG ← → Slightly distorded coord. octahedron (7.3º) ← |
| **MG:a:3227** | 1.0 | 30.3 | 22.17 |  | 5 | 1 | O_ph_4O_w_ - Exclusion zone contact(s): /O_w_/ → Missing octahedral coordination for MG ← Octahedral coordination (3.5º) |
| **MG:a:3228** | 1.0 | 21.8 | 28.19 |  | 5 | 1 | O_ph_4O_w_ - Exclusion zone contact(s): /N7/ → Missing octahedral coordination for MG ← → Slightly distorded coord. octahedron (7.4º) ← |
| **MG:a:3229** | 1.0 | 37.4 | 19.61 |  | 6 |  | 6O_w_ → Slightly distorded coord. octahedron (5.6º) ← |
| **MG:a:3230** | 1.0 | 28.0 | 19.09 |  | 3 | 5 | 2O_ph_O_w_ - Exclusion zone contact(s): /O_w_/O_w_/OP1/OP2/ → Missing octahedral coordination for MG ← |
| **MG:a:3231** | 1.0 | 30.4 | 23.88 |  | 5 | 2 | 2O_ph_3O_w_ - Exclusion zone contact(s): /O_w_/ → Missing octahedral coordination for MG ← Octahedral coordination (3.6º) |
| **MG:a:3232** | 1.0 | 27.2 | 24.10 |  | 6 |  | O_ph_5O_w_ Octahedral coordination (4.0º) |
| **MG:a:3233** | 1.0 | 26.0 | 21.96 |  | 6 | 1 | 3O_ph_3O_w_ Octahedral coordination (3.0º) |
| **MG:a:3234** | 1.0 | 28.4 | 21.26 |  | 6 |  | O_ph_5O_w_ Octahedral coordination (3.2º) |
| **MG:a:3235** | 1.0 | 26.9 | 26.03 |  | 5 | 1 | 5O_w_ - Exclusion zone contact(s): /O_w_/ → Missing octahedral coordination for MG ← Octahedral coordination (3.5º) |
| **MG:a:3236** | 1.0 | 25.8 | 24.05 | 1 | 5 | 1 | 2O_ph_3O_w_ - Exclusion zone contact(s): /O_w_/ → Missing octahedral coordination for MG ← → Slightly distorded coord. octahedron (5.0º) ← |
| **MG:a:3237** | 1.0 | 26.6 | 25.58 |  | 6 |  | 2O_ph_4O_w_ Octahedral coordination (4.3º) |
| **MG:a:3238** | 1.0 | 27.5 | 23.74 |  | 6 |  | 4O_ph_2O_w_ Octahedral coordination (4.2º) |
| **MG:a:3239** | 1.0 | 32.2 | 16.71 |  | 5 | 1 | O_ph_4O_w_ - Exclusion zone contact(s): /O_w_/ → Missing octahedral coordination for MG ← Octahedral coordination (3.4º) |
| **MG:a:3240** | 1.0 | 28.9 | 18.70 |  | 6 |  | O_ph_5O_w_ → Slightly distorded coord. octahedron (5.0º) ← |
| **MG:a:3241** | 1.0 | 33.5 | 15.07 |  | 5 | 1 | 5O_w_ - Exclusion zone contact(s): /O_w_/ → Missing octahedral coordination for MG ← → Slightly distorded coord. octahedron (5.8º) ← |
| **MG:a:3242** | 1.0 | 20.9 | 27.91 |  | 5 | 2 | 2N_b_3O_w_ - Exclusion zone contact(s): /O_w_/ → Missing octahedral coordination for MG ← →→→ Distorted coord. octahedron (10.2º) ←←← |
| **MG:a:3243** | 1.0 | 33.9 | 15.23 |  | 5 | 1 | 2O_ph_3O_w_ - Exclusion zone contact(s): /O_w_/ → Missing octahedral coordination for MG ← Octahedral coordination (4.9º) |
| **MG:a:3244** | 1.0 | 30.5 | 15.30 |  | 6 |  | 3O_ph_3O_w_ → Slightly distorded coord. octahedron (5.7º) ← |
| **MG:a:3245** | 1.0 | 34.6 | 17.00 |  | 5 | 1 | O_ph_4O_w_ - Exclusion zone contact(s): /O_w_/ → Missing octahedral coordination for MG ← → Slightly distorded coord. octahedron (6.0º) ← |
| K:a:3246 | 1.0 | 30.0 | 12.67 | 2 | 3 |  | O_b_2O_w_ - Exclusion zone contact(s): /O6/ → Coordination < 6 for K ← |
| K:a:3247 | 1.0 | 30.0 | 14.30 |  | 6 |  | O_ph_2O_r_N_b_2O_w_ |
| K:a:3248 | 1.0 | 30.0 | 12.24 | 1 | 3 |  | 2O_ph_O_w_ - Exclusion zone contact(s): /O_w_/ → Coordination < 6 for K ← |
| K:a:3249 | 1.0 | 30.0 | 25.91 |  | 5 |  | 2O_ph_2O_r_O_w_ → Coordination < 6 for K ← |
| K:a:3250 | 1.0 | 30.0 | 13.51 |  | 3 |  | N_b_O_w_Anion → Coordination < 6 for K ← |
| K:a:3251 | 1.0 | 30.0 | 31.56 |  | 5 |  | 3O_b_2O_w_ → Coordination < 6 for K ← |
| K:a:3252 | 1.0 | 30.0 | 32.46 |  | 7 |  | 3O_b_4O_w_ |
| K:a:3253 | 1.0 | 30.0 | 37.84 | 1 | 5 |  | O_ph_N_b_3O_w_ - Exclusion zone contact(s): /OP2/ → Coordination < 6 for K ← |
| K:a:3254 | 1.0 | 30.0 | 30.71 |  | 6 |  | O_b_3N_b_O_w_Anion |
| K:a:3255 | 1.0 | 30.0 | 8.05 |  | 4 |  | O_ph_O_b_2O_w_ → Coordination < 6 for K ← |
| K:a:3256 | 1.0 | 30.0 | 16.15 | 1 | 5 |  | O_b_N_b_3O_w_ - Exclusion zone contact(s): /O_w_/ → Coordination < 6 for K ← |
| K:a:3257 | 1.0 | 30.0 | 21.39 |  | 5 |  | 3O_b_2O_w_ → Coordination < 6 for K ← |
| K:a:3258 | 1.0 | 30.0 | 16.80 |  | 4 |  | 3O_b_O_w_ → Coordination < 6 for K ← |
| K:a:3259 | 1.0 | 30.0 | 11.88 |  | 3 |  | 3O_b_ → Coordination < 6 for K ← |
| K:a:3260 | 1.0 | 30.0 | 10.91 |  | 4 |  | 3O_b_O_w_ → Coordination < 6 for K ← |
| K:a:3261 | 1.0 | 30.0 | 15.97 |  | 6 |  | 2O_b_N_b_3O_w_ |
| K:a:3262 | 1.0 | 30.0 | 15.73 |  | 3 |  | 3O_b_ → Coordination < 6 for K ← |
| K:a:3263 | 1.0 | 30.0 | 21.75 |  | 7 |  | 4O_b_3O_w_ |
| K:a:3264 | 1.0 | 30.0 | 22.79 |  | 8 |  | O_r_3O_b_3N_b_O_w_ |
| K:a:3265 | 1.0 | 30.0 | 31.55 |  | 6 |  | 3O_ph_O_r_2O_w_ |
| K:a:3266 | 1.0 | 30.0 | 9.94 |  | 3 |  | 2O_b_O_w_ → Coordination < 6 for K ← |
| K:a:3267 | 1.0 | 30.0 | 29.60 |  | 4 |  | O_ph_2O_b_O_w_ → Coordination < 6 for K ← |
| K:a:3268 | 1.0 | 30.0 | 3.22 |  | 1 |  | O_b_ → Coordination < 6 for K ← → 2Fo-Fc density between 1.0 and 4.0 ! |
| K:a:3269 | 1.0 | 30.0 | 17.24 |  | 5 |  | 2O_ph_2O_b_N_b_ → Coordination < 6 for K ← |
| K:a:3270 | 1.0 | 30.0 | 13.26 |  | 5 |  | O_ph_2O_r_O_b_O_w_ → Coordination < 6 for K ← |
| K:a:3271 | 1.0 | 30.0 | 9.36 |  | 3 |  | 3O_b_ → Coordination < 6 for K ← |
| K:a:3272 | 1.0 | 30.0 | 30.18 |  | 7 |  | O_ph_3O_r_N_b_2O_w_ |
| K:a:3273 | 1.0 | 30.0 | 17.21 |  | 5 |  | O_r_3O_b_O_w_ → Coordination < 6 for K ← |
| K:a:3274 | 1.0 | 30.0 | 13.43 |  | 6 |  | 2O_ph_O_r_3O_w_ |
| K:a:3275 | 1.0 | 30.0 | 31.63 |  | 7 |  | 2O_ph_O_r_O_back_3O_w_ |
| K:a:3276 | 1.0 | 30.0 | 11.91 |  | 4 |  | 2O_r_O_b_O_w_ → Coordination < 6 for K ← |
| K:a:3277 | 1.0 | 30.0 | 31.63 |  | 5 |  | O_ph_O_b_3O_w_ → Coordination < 6 for K ← |
| K:a:3278 | 1.0 | 30.0 | 22.48 |  | 7 |  | 2O_r_O_b_4O_w_ |
| K:a:3279 | 1.0 | 30.0 | 14.45 | 1 | 4 |  | O_ph_O_r_2O_w_ - Exclusion zone contact(s): /O_w_/ → Coordination < 6 for K ← |
| K:a:3280 | 1.0 | 30.0 | 5.71 |  | 3 |  | 2O_b_O_w_ → Coordination < 6 for K ← |
| K:a:3281 | 1.0 | 30.0 | 5.68 |  | 3 |  | O_b_2O_w_ → Coordination < 6 for K ← |
| K:a:3282 | 1.0 | 30.0 | 18.73 |  | 6 |  | 3O_b_3O_w_ |
| K:a:3283 | 1.0 | 30.0 | 10.74 |  | 4 |  | 2O_b_2O_w_ → Coordination < 6 for K ← |
| K:a:3284 | 1.0 | 30.0 | 31.09 |  | 6 |  | O_r_N_b_4O_w_ |
| **MG:a:3285** | 1.0 | 25.3 | 24.13 |  | 4 | 2 | O_ph_3O_w_ - Exclusion zone contact(s): /O_w_/O_w_/ → Missing octahedral coordination for MG ← Octahedral coordination (4.6º) |
| **MG:a:3286** | 1.0 | 39.7 | 13.95 |  | 6 |  | 6O_w_ Octahedral coordination (4.8º) |
| **MG:a:3287** | 1.0 | 28.4 | 21.66 |  | 3 | 3 | 2O_ph_O_w_ - Exclusion zone contact(s): /O_w_/O_w_/O_w_/ → Missing octahedral coordination for MG ← |
| **MG:a:3288** | 1.0 | 38.2 | 16.23 |  | 5 | 1 | O_ph_4O_w_ - Exclusion zone contact(s): /O_w_/ → Missing octahedral coordination for MG ← Octahedral coordination (4.4º) |
| **MG:a:3289** | 1.0 | 27.4 | 24.83 |  | 6 |  | O_ph_5O_w_ → Slightly distorded coord. octahedron (5.0º) ← |
| **MG:a:3290** | 1.0 | 28.1 | 23.76 |  | 4 | 1 | O_ph_3O_w_ - Exclusion zone contact(s): /O_w_/ → Missing octahedral coordination for MG ← → Slightly distorded coord. octahedron (6.2º) ← |
| **MG:a:3291** | 1.0 | 39.5 | 12.24 |  | 3 | 1 | 3O_w_ - Exclusion zone contact(s): /O_w_/ → Missing octahedral coordination for MG ← |
| **MG:a:3292** | 1.0 | 38.8 | 15.14 | 2 | 3 |  | O_ph_2O_w_ - Exclusion zone contact(s): /O_w_/O_w_/ → Missing octahedral coordination for MG ← →→→ Distorted coord. octahedron (10.3º) ←←← |
| **MG:a:3293** | 1.0 | 29.3 | 21.79 |  | 5 | 1 | O_ph_4O_w_ - Exclusion zone contact(s): /O_w_/ → Missing octahedral coordination for MG ← Octahedral coordination (2.3º) |
| **MG:a:3294** | 1.0 | 28.9 | 22.66 |  | 5 | 1 | 2O_ph_3O_w_ - Exclusion zone contact(s): /O_w_/ → Missing octahedral coordination for MG ← Octahedral coordination (3.4º) |
| **MG:a:3295** | 1.0 | 26.3 | 25.05 |  | 4 | 1 | 2O_ph_2O_w_ - Exclusion zone contact(s): /O_w_/ → Missing octahedral coordination for MG ← → Slightly distorded coord. octahedron (5.2º) ← |
| **MG:a:3296** | 1.0 | 27.3 | 24.43 |  | 5 | 2 | O_ph_4O_w_ - Exclusion zone contact(s): /O_w_/ → Missing octahedral coordination for MG ← Octahedral coordination (4.5º) |
| **MG:a:3297** | 1.0 | 26.1 | 24.95 |  | 6 |  | 6O_w_ → Slightly distorded coord. octahedron (6.8º) ← |
| **MG:a:3298** | 1.0 | 31.8 | 21.50 |  | 4 | 1 | O_ph_O_b_2O_w_ - Exclusion zone contact(s): /O_w_/ → Missing octahedral coordination for MG ← Octahedral coordination (4.3º) |
| **MG:a:3299** | 1.0 | 30.3 | 19.58 | 1 | 5 |  | O_ph_4O_w_ - Exclusion zone contact(s): /OP1/ → Missing octahedral coordination for MG ← Octahedral coordination (3.7º) |
| **MG:a:3300** | 1.0 | 29.6 | 18.16 |  | 5 | 1 | O_ph_4O_w_ - Exclusion zone contact(s): /O_w_/ → Missing octahedral coordination for MG ← → Slightly distorded coord. octahedron (6.9º) ← |
| **MG:a:3301** | 1.0 | 40.4 | 15.27 |  | 5 | 1 | 5O_w_ - Exclusion zone contact(s): /O_w_/ → Missing octahedral coordination for MG ← → Slightly distorded coord. octahedron (5.5º) ← |
| **MG:a:3302** | 1.0 | 28.7 | 23.55 |  | 3 | 3 | O_b_2O_w_ - Exclusion zone contact(s): /O_w_/O_w_/O_w_/ → Missing octahedral coordination for MG ← |
| **MG:a:3303** | 1.0 | 31.5 | 17.41 |  | 6 |  | O_ph_5O_w_ → Slightly distorded coord. octahedron (6.4º) ← |
| **MG:a:3304** | 1.0 | 39.7 | 15.25 |  | 4 | 1 | 2O_ph_2O_w_ - Exclusion zone contact(s): /O_w_/ → Missing octahedral coordination for MG ← → Slightly distorded coord. octahedron (7.1º) ← |
| **MG:a:3305** | 1.0 | 26.2 | 23.23 |  | 5 | 1 | O_ph_4O_w_ - Exclusion zone contact(s): /O_w_/ → Missing octahedral coordination for MG ← → Slightly distorded coord. octahedron (7.3º) ← |
| **MG:a:3306** | 1.0 | 31.2 | 17.19 | 1 | 4 |  | O_ph_3O_w_ - Exclusion zone contact(s): /OP2/ → Missing octahedral coordination for MG ← → Slightly distorded coord. octahedron (5.8º) ← |
| **MG:a:3307** | 1.0 | 42.6 | 7.75 | 1 | 3 |  | O_ph_2O_w_ - Exclusion zone contact(s): /O_w_/ → Missing octahedral coordination for MG ← Octahedral coordination (4.9º) |
| **MG:a:3308** | 1.0 | 48.5 | 9.39 |  | 5 |  | 2O_ph_3O_w_ → Missing octahedral coordination for MG ← → Slightly distorded coord. octahedron (6.6º) ← |
| **MG:a:3309** | 1.0 | 32.2 | 15.50 |  | 5 | 1 | 2O_ph_3O_w_ - Exclusion zone contact(s): /O_w_/ → Missing octahedral coordination for MG ← Octahedral coordination (4.7º) |
| **MG:a:3310** | 1.0 | 37.0 | 19.64 |  | 3 | 2 | O_ph_2O_w_ - Exclusion zone contact(s): /O_w_/O_w_/ → Missing octahedral coordination for MG ← |
| **MG:a:3311** | 1.0 | 25.8 | 23.39 |  | 6 |  | 6O_w_ → Slightly distorded coord. octahedron (8.1º) ← |
| **MG:a:3312** | 1.0 | 25.6 | 22.93 |  | 5 | 1 | O_ph_O_b_3O_w_ - Exclusion zone contact(s): /O_w_/ → Missing octahedral coordination for MG ← Octahedral coordination (4.0º) |
| **MG:a:3313** | 1.0 | 42.6 | 14.14 |  | 3 | 2 | O_ph_2O_w_ - Exclusion zone contact(s): /O_w_/O_w_/ → Missing octahedral coordination for MG ← |
| **MG:a:3314** | 1.0 | 25.8 | 21.68 |  | 6 |  | 3O_ph_3O_w_ Octahedral coordination (4.8º) |
| **MG:a:3315** | 1.0 | 33.4 | 17.26 |  | 4 | 2 | O_ph_3O_w_ - Exclusion zone contact(s): /O_w_/O_w_/ → Missing octahedral coordination for MG ← → Slightly distorded coord. octahedron (6.9º) ← |
| **MG:a:3316** | 1.0 | 38.2 | 11.86 |  | 4 | 1 | 2O_ph_2O_w_ - Exclusion zone contact(s): /O_w_/ → Missing octahedral coordination for MG ← → Slightly distorded coord. octahedron (8.1º) ← |
| **MG:a:3317** | 1.0 | 37.7 | 11.96 | 1 | 5 |  | O_b_4O_w_ - Exclusion zone contact(s): /OP2/ → Missing octahedral coordination for MG ← Octahedral coordination (3.8º) |
| **MG:a:3318** | 1.0 | 48.2 | 6.62 |  | 3 | 2 | 3O_w_ - Exclusion zone contact(s): /O_w_/O_w_/ → Missing octahedral coordination for MG ← |
| **MG:a:3319** | 1.0 | 33.2 | 19.85 |  | 6 |  | O_ph_5O_w_ Octahedral coordination (4.2º) |
| **MG:a:3320** | 1.0 | 29.6 | 20.07 |  | 5 | 1 | O_ph_4O_w_ - Exclusion zone contact(s): /O_w_/ → Missing octahedral coordination for MG ← Octahedral coordination (3.7º) |
| **MG:a:3321** | 1.0 | 28.4 | 17.91 |  | 5 | 1 | O_b_4O_w_ - Exclusion zone contact(s): /O_w_/ → Missing octahedral coordination for MG ← Octahedral coordination (4.0º) |
| **MG:a:3322** | 1.0 | 25.5 | 25.28 |  | 6 |  | 2O_ph_4O_w_ Octahedral coordination (4.1º) |
| **MG:a:3323** | 1.0 | 37.2 | 12.37 | 1 | 4 | 1 | O_ph_3O_w_ - Exclusion zone contact(s): /O_w_/O_w_/ → Missing octahedral coordination for MG ← → Slightly distorded coord. octahedron (5.1º) ← |
| **MG:a:3324** | 1.0 | 26.3 | 26.90 |  | 5 | 2 | 3O_ph_2O_w_ - Exclusion zone contact(s): /O3'/O_w_/ → Missing octahedral coordination for MG ← Octahedral coordination (3.8º) |
| **MG:a:3325** | 1.0 | 23.2 | 26.73 |  | 5 | 1 | 2O_ph_O_b_2O_w_ - Exclusion zone contact(s): /O_w_/ → Missing octahedral coordination for MG ← → Slightly distorded coord. octahedron (8.1º) ← |
| **MG:a:3326** | 1.0 | 23.7 | 25.77 |  | 6 |  | 3O_ph_3O_w_ Octahedral coordination (2.7º) |
| **MG:a:3327** | 1.0 | 25.9 | 23.15 |  | 5 | 1 | 3O_ph_2O_w_ - Exclusion zone contact(s): /O_w_/ → Missing octahedral coordination for MG ← → Slightly distorded coord. octahedron (8.9º) ← |
| **MG:a:3328** | 1.0 | 27.0 | 23.82 |  | 5 | 1 | 3O_ph_2O_w_ - Exclusion zone contact(s): /O_w_/ → Missing octahedral coordination for MG ← Octahedral coordination (4.4º) |
| **MG:a:3329** | 1.0 | 25.3 | 23.10 |  | 6 |  | 4O_ph_2O_w_ → Slightly distorded coord. octahedron (5.5º) ← |
| **MG:a:3330** | 1.0 | 42.7 | 7.69 |  | 0 | 5 | ••• No first shell contacts ••• - Exclusion zone contact(s): /O_w_/O_w_/O_w_/OP1/OP1/ → First MG coordination shell missing ← |
| **MG:a:3331** | 1.0 | 28.6 | 21.50 |  | 6 |  | 3O_ph_3O_w_ Octahedral coordination (4.9º) |
| **MG:a:3332** | 1.0 | 29.6 | 18.47 | 1 | 5 |  | O_b_4O_w_ - Exclusion zone contact(s): /OP2/ → Missing octahedral coordination for MG ← → Slightly distorded coord. octahedron (5.5º) ← |
| **MG:a:3333** | 1.0 | 24.2 | 23.66 |  | 5 | 1 | 3O_ph_O_b_O_w_ - Exclusion zone contact(s): /O_w_/ → Missing octahedral coordination for MG ← → Slightly distorded coord. octahedron (7.9º) ← |
| **MG:a:3334** | 1.0 | 35.3 | 15.57 |  | 4 | 1 | O_ph_3O_w_ - Exclusion zone contact(s): /O_w_/ → Missing octahedral coordination for MG ← → Slightly distorded coord. octahedron (8.2º) ← |
| **MG:a:3335** | 1.0 | 37.9 | 14.90 |  | 5 | 1 | O_ph_4O_w_ - Exclusion zone contact(s): /O_w_/ → Missing octahedral coordination for MG ← → Slightly distorded coord. octahedron (5.3º) ← |
| **MG:a:3336** | 1.0 | 26.0 | 25.15 |  | 5 | 1 | 2O_ph_3O_w_ - Exclusion zone contact(s): /O_w_/ → Missing octahedral coordination for MG ← → Slightly distorded coord. octahedron (8.1º) ← |
| **MG:a:3337** | 1.0 | 24.8 | 24.30 |  | 5 | 1 | O_ph_4O_w_ - Exclusion zone contact(s): /O_w_/ → Missing octahedral coordination for MG ← Octahedral coordination (4.4º) |
| K:a:3338 | 1.0 | 30.0 | 13.94 |  | 5 |  | 3O_b_2O_w_ → Coordination < 6 for K ← |
| K:a:3339 | 1.0 | 30.0 | 7.10 |  | 2 |  | 2O_b_ → Coordination < 6 for K ← |
| K:a:3340 | 1.0 | 30.0 | 12.70 |  | 4 |  | 3O_b_O_w_ → Coordination < 6 for K ← |
| K:a:3341 | 1.0 | 30.0 | 7.10 |  | 4 |  | O_ph_O_b_N_b_O_w_ → Coordination < 6 for K ← |
| K:a:3342 | 1.0 | 30.0 | 19.03 |  | 6 |  | O_ph_3O_r_N_b_O_w_ |
| K:a:3343 | 1.0 | 30.0 | 7.33 |  | 3 |  | 2O_b_O_w_ → Coordination < 6 for K ← |
| K:a:3344 | 1.0 | 30.0 | 12.81 |  | 4 |  | O_r_O_b_N_b_O_w_ → Coordination < 6 for K ← |
| K:a:3345 | 1.0 | 30.0 | 16.17 |  | 5 |  | O_r_4O_w_ → Coordination < 6 for K ← |
| K:a:3346 | 1.0 | 30.0 | 11.43 |  | 4 |  | 2O_ph_O_r_O_w_ → Coordination < 6 for K ← |
| K:a:3347 | 1.0 | 30.0 | 24.02 |  | 5 |  | O_ph_O_b_N_b_2O_w_ → Coordination < 6 for K ← |
| **MG:a:3348** | 1.0 | 23.2 | 28.23 |  | 6 |  | O_ph_5O_w_ Octahedral coordination (4.9º) |
| **MG:a:3349** | 1.0 | 28.1 | 21.46 |  | 4 | 1 | O_ph_3O_w_ - Exclusion zone contact(s): /O_w_/ → Missing octahedral coordination for MG ← Octahedral coordination (2.7º) |
| **MG:a:3350** | 1.0 | 51.7 | 9.18 |  | 5 |  | 5O_w_ → Missing octahedral coordination for MG ← Octahedral coordination (3.1º) |
| **MG:a:3351** | 1.0 | 34.5 | 16.29 |  | 5 |  | 2O_ph_3O_w_ → Missing octahedral coordination for MG ← Octahedral coordination (4.9º) |
| **MG:a:3352** | 1.0 | 55.2 | 5.06 |  | 3 | 1 | 3O_w_ - Exclusion zone contact(s): /O_w_/ → Missing octahedral coordination for MG ← |
| **MG:a:3353** | 1.0 | 43.4 | 14.26 |  | 5 | 1 | O_ph_4O_w_ → Missing octahedral coordination for MG ← Octahedral coordination (4.4º) |
| **MG:a:3354** | 1.0 | 44.6 | 7.19 |  | 0 | 4 | ••• No first shell contacts ••• - Exclusion zone contact(s): /O2'/O6/O_w_/ → Consider K in place of MG (2 valid contact(s) for K vs. 0 for MG) |
| **MG:a:3355** | 1.0 | 55.8 | 7.72 | 2 | 0 | 3 | ••• No first shell contacts ••• - Exclusion zone contact(s): /O_w_/O_w_/O_w_/O_w_/O_w_/ → First MG coordination shell missing ← |
| **MG:a:3356** | 1.0 | 45.1 | 6.77 |  | 3 | 2 | O_ph_2O_w_ - Exclusion zone contact(s): /O_w_/ → Missing octahedral coordination for MG ← |
| **MG:a:3357** | 1.0 | 42.8 | 8.99 |  | 6 |  | O_ph_5O_w_ Octahedral coordination (4.9º) |
| **MG:a:3358** | 1.0 | 30.2 | 15.49 |  | 0 | 6 | ••• No first shell contacts ••• - Exclusion zone contact(s): /O4/O6/O6/O_w_/O_w_/ → Consider K in place of MG (3 valid contact(s) for K vs. 0 for MG) |
| **MG:a:3359** | 1.0 | 48.5 | 8.30 |  | 4 | 2 | 4O_w_ - Exclusion zone contact(s): /O_w_/ → Missing octahedral coordination for MG ← → Slightly distorded coord. octahedron (8.2º) ← |
| **MG:a:3360** | 1.0 | 48.0 | 12.11 |  | 4 | 1 | 4O_w_ - Exclusion zone contact(s): /O_w_/ → Missing octahedral coordination for MG ← → Slightly distorded coord. octahedron (5.2º) ← |
| **MG:a:3361** | 1.0 | 50.4 | 8.55 |  | 4 | 1 | 4O_w_ - Exclusion zone contact(s): /O_w_/ → Missing octahedral coordination for MG ← Octahedral coordination (3.3º) |
| **MG:a:3362** | 1.0 | 52.9 | 5.74 |  | 2 | 3 | 2O_w_ - Exclusion zone contact(s): /O_w_/O_w_/ → Missing octahedral coordination for MG ← |
| **MG:a:3363** | 1.0 | 52.8 | 7.85 |  | 3 | 2 | 2O_ph_O_w_ - Exclusion zone contact(s): /O_w_/ → Missing octahedral coordination for MG ← |
| **MG:a:3364** | 1.0 | 55.2 | 6.85 |  | 3 | 1 | 3O_w_ - Exclusion zone contact(s): /OP2/ → Missing octahedral coordination for MG ← |
| **MG:a:3365** | 1.0 | 39.0 | 10.85 |  | 0 | 8 | ••• No first shell contacts ••• - Exclusion zone contact(s): /O4/O6/O6/O_w_/O_w_/ → Consider K in place of MG (2 valid contact(s) for K vs. 0 for MG) |
| **MG:a:3366** | 1.0 | 35.9 | 12.93 |  | 0 | 7 | ••• No first shell contacts ••• - Exclusion zone contact(s): /O4/O6/O_w_/O_w_/O_w_/ → Consider K in place of MG (3 valid contact(s) for K vs. 0 for MG) |
| **MG:b:201** | 1.0 | 67.3 | 2.19 |  | 0 |  | ••• No first shell contacts ••• → First MG coordination shell missing ← → 2Fo-Fc density between 1.0 and 4.0 ! |
| **MG:b:202** | 1.0 | 62.5 | 3.81 |  | 2 |  | 2O_ph_ → Missing octahedral coordination for MG ← → 2Fo-Fc density between 1.0 and 4.0 ! |
| **MG:b:203** | 1.0 | 55.1 | 2.90 |  | 0 | 3 | ••• No first shell contacts ••• - Exclusion zone contact(s): /O6/O6/ → Consider K in place of MG (1 valid contact(s) for K vs. 0 for MG) → 2Fo-Fc density between 1.0 and 4.0 ! |
| **MG:b:204** | 1.0 | 46.0 | 8.31 | 1 | 4 |  | O_ph_3O_w_ - Exclusion zone contact(s): /O_w_/ → Missing octahedral coordination for MG ← → Slightly distorded coord. octahedron (6.1º) ← |
| **MG:b:205** | 1.0 | 38.7 | 12.55 |  | 6 |  | O_b_5O_w_ → Slightly distorded coord. octahedron (5.0º) ← |
| K:b:206 | 1.0 | 30.0 | 6.39 | 1 | 2 |  | O_ph_N_b_ - Exclusion zone contact(s): /O6/ → Coordination < 6 for K ← |
| K:c:301 | 1.0 | 30.0 | 19.57 | 1 | 4 |  | O_back_3O_w_ - Exclusion zone contact(s): /O_back_/ → Coordination < 6 for K ← |
| **MG:c:302** | 1.0 | 52.1 | 6.08 |  | 4 | 1 | 4O_w_ → Missing octahedral coordination for MG ← → Slightly distorded coord. octahedron (5.9º) ← |
| K:c:303 | 1.0 | 30.0 | 27.12 |  | 7 |  | O_ph_O_back_O_coo_4O_w_ |
| **MG:c:304** | 1.0 | 22.8 | 25.47 |  | 6 |  | O_ph_4O_w_N_His_ Octahedral coordination (3.0º) |
| K:c:305 | 1.0 | 30.0 | 29.54 |  | 8 |  | 3O_back_5O_w_ |
| K:c:306 | 1.0 | 30.0 | 33.16 | 1 | 5 |  | 2O_back_OH_prot_2O_w_ - Exclusion zone contact(s): /O_back_/ → Coordination < 6 for K ← |
| **MG:d:301** | 1.0 | 26.6 | 23.54 | 1 | 2 | 3 | 2O_w_ - Exclusion zone contact(s): /NE2_His_/O_w_/O_w_/OP1/ → Missing octahedral coordination for MG ← |
| K:d:302 | 1.0 | 30.0 | 10.35 |  | 3 |  | O_back_O_cno_O_w_ → Coordination < 6 for K ← |
| K:e:301 | 1.0 | 30.0 | 10.62 | 2 | 1 |  | O_back_ - Exclusion zone contact(s): /O_back_/O_back_/ → Coordination < 6 for K ← |
| K:f:201 | 1.0 | 30.0 | 2.92 | 3 | 0 |  | ••• No first shell contacts ••• - Exclusion zone contact(s): /O_back_/O_back_/O_back_/ → Consider MG in place of K (1 valid contact(s) for MG vs. 0 for K) → 2Fo-Fc density between 1.0 and 4.0 ! |
| **MG:z:101** | 1.0 | 32.5 | 16.90 |  | 5 | 2 | O_coo_4O_w_ - Exclusion zone contact(s): /O_w_/OD2_Asp_/ → Missing octahedral coordination for MG ← → Slightly distorded coord. octahedron (5.1º) ← |

---

\_\_\_\_\_\_\_\_\_\_\_\_\_\_\_\_\_\_\_\_\_\_\_\_\_\_\_\_\_\_\_\_\_\_\_\_\_\_\_\_\_\_\_\_\_\_\_\_\_\_\_\_\_\_\_\_\_\_\_\_\_\_\_\_  

#### 3) Potential ion coordination issue list

##### Go to ToC

|  |  |  |
| --- | --- | --- |
| **Issue(s)/Comment(s):** | **Count** | **Affected ion(s)** |
| **1) Possible missatribution: K could be MG !!!** | 3 | A:1647  A:1737  f:201 |
| **1) Possible missatribution: MG could be K !!!** | 40 | A:1609  A:1618  A:1621  A:1636  A:1640  A:1641  A:1682  A:1703  A:1709  A:1714  A:1721  A:1744  a:3005  a:3018  a:3029  a:3031  a:3032  a:3044  a:3045  a:3051  a:3052  a:3053  a:3054  a:3055  a:3068  a:3069  a:3070  a:3071  a:3073  a:3148  a:3158  a:3162  a:3169  a:3173  a:3203  a:3354  a:3358  a:3365  a:3366  b:203 |
| **1) No misattribution(s) detected** | 487 | 3:101  A:1601  A:1602  A:1603  A:1604  A:1605  A:1606  A:1607  A:1608  A:1610  A:1611  A:1612  A:1613  A:1614  A:1615  A:1616  A:1617  A:1619  A:1620  A:1622  A:1623  A:1624  A:1625  A:1626  A:1627  A:1628  A:1629  A:1630  A:1631  A:1632  A:1633  A:1634  A:1635  A:1637  A:1638  A:1639  A:1642  A:1643  A:1644  A:1645  A:1646  A:1648  A:1649  A:1650  A:1651  A:1652  A:1653  A:1654  A:1655  A:1656  A:1657  A:1658  A:1659  A:1660  A:1661  A:1662  A:1663  A:1664  A:1665  A:1666  A:1667  A:1668  A:1669  A:1670  A:1671  A:1672  A:1673  A:1674  A:1675  A:1676  A:1677  A:1678  A:1679  A:1680  A:1681  A:1683  A:1684  A:1685  A:1686  A:1687  A:1688  A:1689  A:1690  A:1691  A:1692  A:1693  A:1694  A:1695  A:1696  A:1697  A:1698  A:1699  A:1700  A:1701  A:1702  A:1704  A:1705  A:1706  A:1707  A:1708  A:1710  A:1711  A:1712  A:1713  A:1715  A:1716  A:1717  A:1718  A:1719  A:1720  A:1722  A:1723  A:1724  A:1725  A:1726  A:1727  A:1728  A:1729  A:1730  A:1731  A:1732  A:1733  A:1734  A:1735  A:1736  A:1738  A:1739  A:1740  A:1741  A:1742  A:1743  A:1745  X:101  a:3001  a:3002  a:3003  a:3004  a:3006  a:3007  a:3008  a:3009  a:3010  a:3011  a:3012  a:3013  a:3014  a:3015  a:3016  a:3017  a:3019  a:3020  a:3021  a:3022  a:3023  a:3024  a:3025  a:3026  a:3027  a:3028  a:3030  a:3033  a:3034  a:3035  a:3036  a:3037  a:3038  a:3039  a:3040  a:3041  a:3042  a:3043  a:3046  a:3047  a:3048  a:3049  a:3050  a:3056  a:3057  a:3058  a:3059  a:3060  a:3061  a:3062  a:3063  a:3064  a:3065  a:3066  a:3067  a:3072  a:3074  a:3075  a:3076  a:3077  a:3078  a:3079  a:3080  a:3081  a:3082  a:3083  a:3084  a:3085  a:3086  a:3087  a:3088  a:3089  a:3090  a:3091  a:3092  a:3093  a:3094  a:3095  a:3096  a:3097  a:3098  a:3099  a:3100  a:3101  a:3102  a:3103  a:3104  a:3105  a:3106  a:3107  a:3108  a:3109  a:3110  a:3111  a:3112  a:3113  a:3114  a:3115  a:3116  a:3117  a:3118  a:3119  a:3120  a:3121  a:3122  a:3123  a:3124  a:3125  a:3126  a:3127  a:3128  a:3129  a:3130  a:3131  a:3132  a:3133  a:3134  a:3135  a:3136  a:3137  a:3138  a:3139  a:3140  a:3141  a:3142  a:3143  a:3144  a:3145  a:3146  a:3147  a:3149  a:3150  a:3151  a:3152  a:3153  a:3154  a:3155  a:3156  a:3157  a:3159  a:3160  a:3161  a:3163  a:3164  a:3165  a:3166  a:3167  a:3168  a:3170  a:3171  a:3172  a:3174  a:3175  a:3176  a:3177  a:3178  a:3179  a:3180  a:3181  a:3182  a:3183  a:3184  a:3185  a:3186  a:3187  a:3188  a:3189  a:3190  a:3191  a:3192  a:3193  a:3194  a:3195  a:3196  a:3197  a:3198  a:3199  a:3200  a:3201  a:3202  a:3204  a:3205  a:3206  a:3207  a:3208  a:3209  a:3210  a:3211  a:3212  a:3213  a:3214  a:3215  a:3216  a:3217  a:3218  a:3219  a:3220  a:3221  a:3222  a:3223  a:3224  a:3225  a:3226  a:3227  a:3228  a:3229  a:3230  a:3231  a:3232  a:3233  a:3234  a:3235  a:3236  a:3237  a:3238  a:3239  a:3240  a:3241  a:3242  a:3243  a:3244  a:3245  a:3246  a:3247  a:3248  a:3249  a:3250  a:3251  a:3252  a:3253  a:3254  a:3255  a:3256  a:3257  a:3258  a:3259  a:3260  a:3261  a:3262  a:3263  a:3264  a:3265  a:3266  a:3267  a:3268  a:3269  a:3270  a:3271  a:3272  a:3273  a:3274  a:3275  a:3276  a:3277  a:3278  a:3279  a:3280  a:3281  a:3282  a:3283  a:3284  a:3285  a:3286  a:3287  a:3288  a:3289  a:3290  a:3291  a:3292  a:3293  a:3294  a:3295  a:3296  a:3297  a:3298  a:3299  a:3300  a:3301  a:3302  a:3303  a:3304  a:3305  a:3306  a:3307  a:3308  a:3309  a:3310  a:3311  a:3312  a:3313  a:3314  a:3315  a:3316  a:3317  a:3318  a:3319  a:3320  a:3321  a:3322  a:3323  a:3324  a:3325  a:3326  a:3327  a:3328  a:3329  a:3330  a:3331  a:3332  a:3333  a:3334  a:3335  a:3336  a:3337  a:3338  a:3339  a:3340  a:3341  a:3342  a:3343  a:3344  a:3345  a:3346  a:3347  a:3348  a:3349  a:3350  a:3351  a:3352  a:3353  a:3355  a:3356  a:3357  a:3359  a:3360  a:3361  a:3362  a:3363  a:3364  b:201  b:202  b:204  b:205  b:206  c:301  c:302  c:303  c:304  c:305  c:306  d:301  d:302  e:301  z:101 |
| **2) K: First shell missing !!!** | 3 | A:1603  A:1660  a:3131 |
| **2) MG: First shell missing !!!** | 14 | A:1625  A:1637  A:1643  A:1695  A:1696  A:1711  A:1715  A:1718  A:1719  a:3026  a:3042  a:3330  a:3355  b:201 |
| **3) B\_factor > 80 for Mg!!!** | 6 | A:1623  A:1687  A:1689  A:1712  A:1715  A:1743 |
| **3) B\_factor > 80 for Zn !!!** | 1 | 3:101 |
| **4) Missing octahedral coordination for Mg !!!** | 232 | A:1607  A:1608  A:1610  A:1611  A:1612  A:1613  A:1614  A:1615  A:1616  A:1617  A:1618  A:1619  A:1620  A:1622  A:1623  A:1624  A:1626  A:1628  A:1629  A:1630  A:1631  A:1632  A:1633  A:1635  A:1639  A:1668  A:1669  A:1672  A:1674  A:1675  A:1676  A:1677  A:1678  A:1679  A:1680  A:1683  A:1685  A:1686  A:1687  A:1688  A:1689  A:1690  A:1691  A:1692  A:1693  A:1694  A:1697  A:1698  A:1699  A:1700  A:1701  A:1702  A:1703  A:1704  A:1705  A:1706  A:1707  A:1708  A:1709  A:1710  A:1712  A:1713  A:1716  A:1717  A:1720  A:1741  A:1742  X:101  a:3001  a:3002  a:3003  a:3004  a:3006  a:3007  a:3009  a:3010  a:3011  a:3012  a:3013  a:3014  a:3015  a:3017  a:3021  a:3022  a:3023  a:3024  a:3025  a:3027  a:3028  a:3033  a:3034  a:3035  a:3037  a:3038  a:3039  a:3041  a:3043  a:3046  a:3049  a:3050  a:3065  a:3066  a:3070  a:3072  a:3073  a:3074  a:3075  a:3076  a:3077  a:3079  a:3081  a:3083  a:3085  a:3091  a:3092  a:3094  a:3144  a:3145  a:3146  a:3149  a:3151  a:3152  a:3155  a:3157  a:3158  a:3159  a:3160  a:3163  a:3164  a:3166  a:3167  a:3168  a:3170  a:3171  a:3172  a:3183  a:3184  a:3187  a:3189  a:3190  a:3191  a:3192  a:3195  a:3196  a:3197  a:3200  a:3202  a:3204  a:3206  a:3208  a:3209  a:3211  a:3213  a:3214  a:3215  a:3217  a:3218  a:3220  a:3222  a:3223  a:3224  a:3225  a:3226  a:3227  a:3228  a:3230  a:3231  a:3235  a:3236  a:3239  a:3241  a:3242  a:3243  a:3245  a:3285  a:3287  a:3288  a:3290  a:3291  a:3292  a:3293  a:3294  a:3295  a:3296  a:3298  a:3299  a:3300  a:3301  a:3302  a:3304  a:3305  a:3306  a:3307  a:3308  a:3309  a:3310  a:3312  a:3313  a:3315  a:3316  a:3317  a:3318  a:3320  a:3321  a:3323  a:3324  a:3325  a:3327  a:3328  a:3332  a:3333  a:3334  a:3335  a:3336  a:3337  a:3349  a:3350  a:3351  a:3352  a:3353  a:3356  a:3359  a:3360  a:3361  a:3362  a:3363  a:3364  b:202  b:204  c:302  d:301  z:101 |
| **5) Mg octahedral coordination** | 93 | A:1606  A:1615  A:1620  A:1628  A:1634  A:1642  A:1673  A:1675  A:1681  A:1684  A:1710  A:1713  a:3016  a:3021  a:3030  a:3082  a:3083  a:3084  a:3086  a:3089  a:3090  a:3091  a:3092  a:3093  a:3145  a:3153  a:3154  a:3155  a:3156  a:3157  a:3161  a:3164  a:3165  a:3183  a:3186  a:3188  a:3192  a:3194  a:3198  a:3199  a:3200  a:3204  a:3205  a:3210  a:3212  a:3213  a:3215  a:3216  a:3218  a:3220  a:3221  a:3222  a:3223  a:3227  a:3231  a:3232  a:3233  a:3234  a:3235  a:3237  a:3238  a:3239  a:3243  a:3285  a:3286  a:3288  a:3293  a:3294  a:3296  a:3298  a:3299  a:3307  a:3309  a:3312  a:3314  a:3317  a:3319  a:3320  a:3321  a:3322  a:3324  a:3326  a:3328  a:3331  a:3337  a:3348  a:3349  a:3350  a:3351  a:3353  a:3357  a:3361  c:304 |
| **5) Slightly distorded coord. octahedron !** | 127 | A:1607  A:1608  A:1610  A:1613  A:1614  A:1627  A:1629  A:1631  A:1632  A:1633  A:1638  A:1669  A:1671  A:1672  A:1676  A:1678  A:1679  A:1680  A:1686  A:1687  A:1691  A:1693  A:1700  A:1701  A:1702  A:1703  A:1706  A:1716  A:1742  a:3001  a:3003  a:3007  a:3008  a:3009  a:3010  a:3019  a:3020  a:3024  a:3025  a:3027  a:3034  a:3035  a:3036  a:3038  a:3039  a:3040  a:3047  a:3048  a:3050  a:3065  a:3067  a:3076  a:3078  a:3079  a:3080  a:3085  a:3087  a:3088  a:3094  a:3095  a:3143  a:3144  a:3146  a:3149  a:3150  a:3152  a:3159  a:3163  a:3166  a:3167  a:3170  a:3171  a:3172  a:3182  a:3184  a:3185  a:3187  a:3193  a:3197  a:3201  a:3202  a:3206  a:3207  a:3209  a:3211  a:3214  a:3217  a:3219  a:3224  a:3225  a:3226  a:3228  a:3229  a:3236  a:3240  a:3241  a:3244  a:3245  a:3289  a:3290  a:3295  a:3297  a:3300  a:3301  a:3303  a:3304  a:3305  a:3306  a:3308  a:3311  a:3315  a:3316  a:3323  a:3325  a:3327  a:3329  a:3332  a:3333  a:3334  a:3335  a:3336  a:3359  a:3360  b:204  b:205  c:302  z:101 |
| **5) Distorted coordination octahedron !!!** | 21 | A:1619  A:1639  A:1670  A:1677  A:1683  A:1689  A:1692  A:1698  A:1708  A:1743  A:1745  a:3013  a:3033  a:3041  a:3046  a:3074  a:3077  a:3147  a:3174  a:3242  a:3292 |
| **6) Coordination < 6 for for K !** | 112 | A:1601  A:1602  A:1604  A:1605  A:1644  A:1645  A:1648  A:1649  A:1650  A:1651  A:1652  A:1653  A:1654  A:1655  A:1656  A:1657  A:1658  A:1659  A:1661  A:1662  A:1664  A:1666  A:1667  A:1722  A:1723  A:1725  A:1726  A:1727  A:1728  A:1729  A:1730  A:1731  A:1732  A:1734  A:1736  A:1739  A:1740  a:3057  a:3060  a:3061  a:3062  a:3063  a:3064  a:3096  a:3097  a:3099  a:3101  a:3102  a:3103  a:3105  a:3107  a:3108  a:3109  a:3110  a:3114  a:3115  a:3117  a:3118  a:3122  a:3124  a:3125  a:3126  a:3129  a:3132  a:3136  a:3138  a:3139  a:3142  a:3175  a:3176  a:3178  a:3181  a:3246  a:3248  a:3249  a:3250  a:3251  a:3253  a:3255  a:3256  a:3257  a:3258  a:3259  a:3260  a:3262  a:3266  a:3267  a:3268  a:3269  a:3270  a:3271  a:3273  a:3276  a:3277  a:3279  a:3280  a:3281  a:3283  a:3338  a:3339  a:3340  a:3341  a:3343  a:3344  a:3345  a:3346  a:3347  b:206  c:301  c:306  d:302  e:301 |
| **8) 2Fo-Fc density for K between 1.0 and 4.0 !** | 17 | A:1601  A:1603  A:1605  A:1647  A:1648  A:1649  A:1651  A:1656  A:1657  A:1658  A:1660  A:1661  A:1664  A:1723  a:3131  a:3268  f:201 |
| **8) 2Fo-Fc density for Mg between 1.0 and 4.0 !** | 65 | A:1611  A:1616  A:1617  A:1618  A:1621  A:1622  A:1623  A:1624  A:1625  A:1626  A:1630  A:1631  A:1635  A:1637  A:1642  A:1643  A:1668  A:1670  A:1677  A:1679  A:1685  A:1687  A:1688  A:1689  A:1690  A:1691  A:1692  A:1694  A:1695  A:1696  A:1697  A:1698  A:1699  A:1703  A:1704  A:1705  A:1707  A:1708  A:1709  A:1711  A:1712  A:1713  A:1714  A:1715  A:1716  A:1717  A:1718  A:1719  A:1720  A:1721  A:1741  A:1742  A:1743  A:1744  A:1745  a:3026  a:3043  a:3048  a:3050  a:3151  a:3191  a:3208  b:201  b:202  b:203 |
| **9) K...N7 excl. contact(s) !!!** | 3 | A:1644  A:1664  A:1664 |
| **9) K...O2 excl. contact(s) !!!** | 3 | A:1647  A:1728  a:3181 |
| **9) K...O2' excl. contact(s) !!!** | 2 | a:3064  a:3142 |
| **9) K...O4 excl. contact(s) !!!** | 1 | a:3131 |
| **9) K...O6 excl. contact(s) !!!** | 8 | A:1603  A:1605  A:1647  A:1727  a:3062  a:3181  a:3246  b:206 |
| **9) K...O_back_ excl. contact(s) !!!** | 8 | a:3126  c:301  c:306  e:301  e:301  f:201  f:201  f:201 |
| **9) K...O_w_ excl. contact(s) !!!** | 20 | A:1602  A:1652  A:1655  A:1659  A:1662  A:1666  A:1735  A:1738  a:3102  a:3103  a:3103  a:3107  a:3115  a:3122  a:3138  a:3175  a:3176  a:3248  a:3256  a:3279 |
| **9) K...OP1 excl. contact(s) !!!** | 3 | A:1644  A:1652  A:1737 |
| **9) K...OP2 excl. contact(s) !!!** | 4 | A:1737  a:3126  a:3129  a:3253 |
| **9) MG...N3_A_ excl. contact(s) !!!** | 1 | a:3044 |
| **9) MG...N3_C_ excl. contact(s) !!!** | 1 | a:3169 |
| **9) MG...N4 excl. contact(s) !!!** | 1 | A:1744 |
| **9) MG...N7 excl. contact(s) !!!** | 6 | A:1705  A:1709  a:3014  a:3091  a:3220  a:3228 |
| **9) MG...NE2_His_ excl. contact(s) !!!** | 1 | d:301 |
| **9) MG...NE_Arg_ excl. contact(s) !!!** | 1 | a:3070 |
| **9) MG...NH2_Arg_ excl. contact(s) !!!** | 1 | A:1640 |
| **9) MG...NZ_Lys_ excl. contact(s) !!!** | 1 | A:1640 |
| **9) MG...O2 excl. contact(s) !!!** | 2 | A:1695  a:3173 |
| **9) MG...O2' excl. contact(s) !!!** | 11 | A:1618  A:1641  A:1714  a:3045  a:3053  a:3068  a:3069  a:3071  a:3173  a:3203  a:3354 |
| **9) MG...O3' excl. contact(s) !!!** | 13 | A:1618  A:1671  A:1714  a:3044  a:3054  a:3054  a:3068  a:3069  a:3069  a:3071  a:3187  a:3203  a:3324 |
| **9) MG...O4 excl. contact(s) !!!** | 21 | A:1609  A:1618  A:1636  A:1636  A:1699  A:1744  a:3005  a:3018  a:3022  a:3029  a:3031  a:3032  a:3032  a:3051  a:3158  a:3162  a:3174  a:3203  a:3358  a:3365  a:3366 |
| **9) MG...O4' excl. contact(s) !!!** | 1 | a:3042 |
| **9) MG...O5' excl. contact(s) !!!** | 1 | A:1689 |
| **9) MG...O6 excl. contact(s) !!!** | 38 | A:1609  A:1609  A:1621  A:1636  A:1682  A:1682  A:1690  A:1708  A:1744  A:1744  a:3005  a:3005  a:3005  a:3018  a:3018  a:3029  a:3031  a:3031  a:3032  a:3044  a:3045  a:3051  a:3051  a:3053  a:3053  a:3158  a:3162  a:3162  a:3162  a:3196  a:3354  a:3358  a:3358  a:3365  a:3365  a:3366  b:203  b:203 |
| **9) MG...O_back_ excl. contact(s) !!!** | 1 | A:1637 |
| **9) MG...O_w_ excl. contact(s) !!!** | 373 | A:1607  A:1607  A:1608  A:1609  A:1609  A:1610  A:1610  A:1610  A:1611  A:1611  A:1612  A:1613  A:1613  A:1613  A:1614  A:1617  A:1618  A:1619  A:1619  A:1619  A:1621  A:1621  A:1621  A:1621  A:1622  A:1622  A:1623  A:1624  A:1624  A:1625  A:1630  A:1630  A:1633  A:1633  A:1635  A:1636  A:1636  A:1636  A:1637  A:1638  A:1639  A:1639  A:1640  A:1641  A:1641  A:1643  A:1643  A:1668  A:1669  A:1674  A:1675  A:1676  A:1676  A:1678  A:1678  A:1682  A:1682  A:1682  A:1682  A:1682  A:1682  A:1683  A:1683  A:1685  A:1686  A:1686  A:1690  A:1691  A:1691  A:1692  A:1692  A:1692  A:1693  A:1693  A:1697  A:1697  A:1698  A:1700  A:1701  A:1701  A:1702  A:1702  A:1702  A:1703  A:1703  A:1704  A:1705  A:1705  A:1706  A:1706  A:1706  A:1707  A:1707  A:1707  A:1708  A:1710  A:1710  A:1714  A:1714  A:1714  A:1714  A:1714  A:1716  A:1716  A:1717  A:1719  A:1721  A:1741  A:1742  A:1744  A:1744  A:1744  X:101  X:101  a:3002  a:3002  a:3002  a:3003  a:3003  a:3004  a:3005  a:3005  a:3005  a:3006  a:3006  a:3006  a:3007  a:3009  a:3009  a:3010  a:3010  a:3010  a:3011  a:3011  a:3011  a:3013  a:3014  a:3014  a:3017  a:3018  a:3018  a:3018  a:3021  a:3022  a:3023  a:3024  a:3025  a:3029  a:3029  a:3031  a:3031  a:3031  a:3031  a:3032  a:3032  a:3032  a:3033  a:3033  a:3033  a:3034  a:3035  a:3038  a:3038  a:3039  a:3039  a:3039  a:3041  a:3041  a:3041  a:3042  a:3042  a:3042  a:3043  a:3043  a:3043  a:3043  a:3044  a:3044  a:3045  a:3045  a:3045  a:3045  a:3045  a:3046  a:3049  a:3049  a:3049  a:3050  a:3051  a:3051  a:3053  a:3053  a:3054  a:3054  a:3054  a:3055  a:3055  a:3065  a:3065  a:3066  a:3068  a:3068  a:3069  a:3069  a:3070  a:3071  a:3071  a:3071  a:3071  a:3073  a:3073  a:3074  a:3074  a:3074  a:3074  a:3074  a:3075  a:3076  a:3077  a:3079  a:3079  a:3081  a:3083  a:3091  a:3092  a:3092  a:3094  a:3094  a:3144  a:3145  a:3146  a:3146  a:3148  a:3148  a:3149  a:3151  a:3151  a:3152  a:3152  a:3155  a:3157  a:3158  a:3159  a:3160  a:3163  a:3163  a:3163  a:3164  a:3166  a:3167  a:3167  a:3168  a:3168  a:3168  a:3171  a:3172  a:3173  a:3173  a:3173  a:3183  a:3184  a:3187  a:3189  a:3192  a:3192  a:3195  a:3196  a:3196  a:3200  a:3202  a:3203  a:3203  a:3203  a:3206  a:3208  a:3208  a:3213  a:3214  a:3214  a:3215  a:3217  a:3222  a:3223  a:3224  a:3225  a:3226  a:3227  a:3230  a:3230  a:3231  a:3235  a:3236  a:3239  a:3241  a:3242  a:3243  a:3245  a:3285  a:3285  a:3287  a:3287  a:3287  a:3288  a:3290  a:3291  a:3292  a:3292  a:3293  a:3294  a:3295  a:3296  a:3298  a:3300  a:3301  a:3302  a:3302  a:3302  a:3304  a:3305  a:3307  a:3309  a:3310  a:3310  a:3312  a:3313  a:3313  a:3315  a:3315  a:3316  a:3318  a:3318  a:3320  a:3321  a:3323  a:3323  a:3324  a:3325  a:3327  a:3328  a:3330  a:3330  a:3330  a:3333  a:3334  a:3335  a:3336  a:3337  a:3349  a:3352  a:3354  a:3355  a:3355  a:3355  a:3355  a:3355  a:3356  a:3358  a:3358  a:3359  a:3360  a:3361  a:3362  a:3362  a:3363  a:3365  a:3365  a:3366  a:3366  a:3366  b:204  d:301  d:301  z:101 |
| **9) MG...OD2_Asp_ excl. contact(s) !!!** | 1 | z:101 |
| **9) MG...OP1 excl. contact(s) !!!** | 27 | A:1624  A:1629  A:1637  A:1640  A:1641  A:1689  A:1703  A:1706  a:3033  a:3044  a:3052  a:3054  a:3055  a:3068  a:3068  a:3069  a:3073  a:3148  a:3148  a:3148  a:3187  a:3202  a:3230  a:3299  a:3330  a:3330  d:301 |
| **9) MG...OP2 excl. contact(s) !!!** | 43 | A:1618  A:1618  A:1625  A:1630  A:1631  A:1632  A:1640  A:1641  A:1643  A:1672  A:1691  A:1693  A:1695  A:1698  A:1703  A:1707  A:1708  A:1710  A:1721  a:3004  a:3012  a:3015  a:3020  a:3024  a:3032  a:3052  a:3053  a:3054  a:3055  a:3066  a:3070  a:3073  a:3091  a:3094  a:3144  a:3173  a:3203  a:3215  a:3230  a:3306  a:3317  a:3332  a:3364 |
| **10) K...N1_A_ contact(s)** | 5 | A:1604  A:1654  a:3114  a:3254  a:3264 |
| **10) K...N3 contact(s)** | 1 | a:3105 |
| **10) K...N3_A_ contact(s)** | 8 | A:1648  A:1665  a:3101  a:3105  a:3113  a:3122  a:3284  a:3344 |
| **10) K...N3_C_ contact(s)** | 1 | A:1722 |
| **10) K...N3_G_ contact(s)** | 4 | A:1665  a:3104  a:3133  a:3247 |
| **10) K...N3_U_ contact(s)** | 1 | a:3250 |
| **10) K...N4 contact(s)** | 2 | A:1726  a:3254 |
| **10) K...N6 contact(s)** | 2 | a:3098  a:3107 |
| **10) K...N7 contact(s)** | 31 | A:1657  A:1658  A:1666  A:1722  A:1725  A:1733  a:3113  a:3113  a:3114  a:3116  a:3118  a:3121  a:3123  a:3133  a:3137  a:3177  a:3179  a:3250  a:3253  a:3254  a:3254  a:3256  a:3261  a:3264  a:3264  a:3269  a:3272  a:3341  a:3342  a:3347  b:206 |
| **10) K...N_back_ contact(s)** | 1 | a:3176 |
| **10) K...O2 contact(s)** | 26 | A:1604  A:1645  A:1648  A:1651  A:1662  A:1665  A:1726  a:3056  a:3058  a:3058  a:3096  a:3097  a:3105  a:3116  a:3127  a:3127  a:3138  a:3138  a:3176  a:3176  a:3251  a:3263  a:3264  a:3267  a:3267  a:3344 |
| **10) K...O2' contact(s)** | 49 | A:1601  A:1601  A:1604  A:1648  A:1652  A:1662  A:1665  A:1665  A:1723  A:1725  A:1726  A:1728  A:1729  A:1730  A:1733  a:3056  a:3059  a:3062  a:3096  a:3097  a:3101  a:3104  a:3110  a:3112  a:3112  a:3112  a:3114  a:3117  a:3118  a:3121  a:3127  a:3128  a:3132  a:3133  a:3134  a:3140  a:3141  a:3141  a:3249  a:3272  a:3272  a:3273  a:3278  a:3279  a:3284  a:3342  a:3342  a:3344  a:3345 |
| **10) K...O3' contact(s)** | 26 | A:1646  A:1658  A:1724  A:1729  A:1733  A:1735  A:1735  A:1736  a:3063  a:3100  a:3109  a:3112  a:3117  a:3119  a:3120  a:3121  a:3133  a:3134  a:3140  a:3141  a:3141  a:3249  a:3265  a:3272  a:3278  a:3342 |
| **10) K...O4 contact(s)** | 72 | A:1645  A:1649  A:1650  A:1653  A:1653  A:1655  A:1656  A:1656  A:1667  A:1727  A:1728  A:1728  A:1731  A:1732  A:1734  A:1738  A:1739  A:1739  A:1740  a:3056  a:3060  a:3060  a:3062  a:3064  a:3098  a:3099  a:3102  a:3105  a:3107  a:3108  a:3115  a:3124  a:3125  a:3129  a:3129  a:3130  a:3132  a:3132  a:3135  a:3135  a:3136  a:3137  a:3139  a:3141  a:3142  a:3175  a:3177  a:3179  a:3180  a:3251  a:3252  a:3257  a:3258  a:3259  a:3260  a:3260  a:3262  a:3263  a:3263  a:3269  a:3271  a:3271  a:3273  a:3273  a:3277  a:3278  a:3280  a:3281  a:3282  a:3338  a:3338  a:3339 |
| **10) K...O4' contact(s)** | 14 | A:1649  A:1652  A:1733  a:3059  a:3113  a:3116  a:3120  a:3179  a:3181  a:3247  a:3264  a:3270  a:3275  a:3276 |
| **10) K...O5' contact(s)** | 12 | A:1733  a:3059  a:3105  a:3111  a:3112  a:3141  a:3179  a:3247  a:3270  a:3274  a:3276  a:3346 |
| **10) K...O6 contact(s)** | 137 | A:1605  A:1605  A:1645  A:1645  A:1646  A:1650  A:1650  A:1650  A:1651  A:1653  A:1655  A:1655  A:1656  A:1656  A:1657  A:1657  A:1659  A:1661  A:1661  A:1663  A:1663  A:1663  A:1664  A:1665  A:1666  A:1667  A:1667  A:1722  A:1725  A:1727  A:1728  A:1731  A:1731  A:1732  A:1733  A:1734  A:1734  A:1738  A:1738  A:1739  A:1739  A:1740  A:1740  a:3056  a:3057  a:3057  a:3060  a:3061  a:3061  a:3061  a:3098  a:3098  a:3099  a:3099  a:3102  a:3102  a:3106  a:3107  a:3107  a:3108  a:3113  a:3114  a:3115  a:3115  a:3116  a:3116  a:3124  a:3124  a:3124  a:3125  a:3125  a:3125  a:3126  a:3128  a:3128  a:3129  a:3130  a:3130  a:3133  a:3135  a:3135  a:3136  a:3136  a:3137  a:3139  a:3139  a:3142  a:3142  a:3175  a:3177  a:3177  a:3179  a:3180  a:3180  a:3246  a:3251  a:3252  a:3252  a:3254  a:3255  a:3256  a:3257  a:3257  a:3258  a:3258  a:3259  a:3259  a:3260  a:3261  a:3261  a:3262  a:3262  a:3263  a:3264  a:3264  a:3266  a:3266  a:3268  a:3269  a:3270  a:3271  a:3273  a:3276  a:3280  a:3282  a:3282  a:3283  a:3283  a:3338  a:3339  a:3340  a:3340  a:3340  a:3341  a:3343  a:3343  a:3347 |
| **10) K...O_back_ contact(s)** | 16 | A:1722  A:1723  a:3058  a:3134  a:3176  a:3181  a:3275  c:301  c:303  c:305  c:305  c:305  c:306  c:306  d:302  e:301 |
| **10) K...O_w_ contact(s)** | 261 | A:1602  A:1602  A:1604  A:1644  A:1644  A:1646  A:1646  A:1646  A:1648  A:1649  A:1650  A:1651  A:1652  A:1652  A:1652  A:1653  A:1654  A:1654  A:1655  A:1657  A:1658  A:1658  A:1659  A:1659  A:1662  A:1663  A:1663  A:1663  A:1667  A:1724  A:1724  A:1724  A:1725  A:1727  A:1727  A:1729  A:1730  A:1731  A:1731  A:1732  A:1732  A:1732  A:1733  A:1734  A:1735  A:1735  A:1736  A:1738  A:1738  A:1738  A:1740  A:1740  a:3056  a:3056  a:3057  a:3058  a:3059  a:3059  a:3060  a:3061  a:3061  a:3062  a:3062  a:3063  a:3064  a:3064  a:3064  a:3096  a:3096  a:3098  a:3098  a:3099  a:3099  a:3100  a:3100  a:3100  a:3100  a:3102  a:3102  a:3103  a:3104  a:3104  a:3104  a:3106  a:3106  a:3106  a:3106  a:3106  a:3107  a:3109  a:3110  a:3110  a:3110  a:3111  a:3111  a:3111  a:3111  a:3111  a:3112  a:3113  a:3113  a:3114  a:3116  a:3117  a:3119  a:3119  a:3119  a:3119  a:3120  a:3121  a:3121  a:3122  a:3123  a:3123  a:3123  a:3123  a:3123  a:3123  a:3124  a:3125  a:3126  a:3127  a:3127  a:3127  a:3128  a:3128  a:3128  a:3128  a:3129  a:3129  a:3130  a:3130  a:3130  a:3132  a:3133  a:3134  a:3134  a:3135  a:3135  a:3136  a:3137  a:3137  a:3137  a:3139  a:3139  a:3140  a:3140  a:3140  a:3141  a:3142  a:3142  a:3176  a:3177  a:3177  a:3178  a:3178  a:3179  a:3179  a:3180  a:3180  a:3180  a:3181  a:3181  a:3246  a:3246  a:3247  a:3247  a:3248  a:3249  a:3250  a:3251  a:3251  a:3252  a:3252  a:3252  a:3252  a:3253  a:3253  a:3253  a:3254  a:3255  a:3255  a:3256  a:3256  a:3256  a:3257  a:3257  a:3258  a:3260  a:3261  a:3261  a:3261  a:3263  a:3263  a:3263  a:3264  a:3265  a:3265  a:3266  a:3267  a:3270  a:3272  a:3272  a:3273  a:3274  a:3274  a:3274  a:3275  a:3275  a:3275  a:3276  a:3277  a:3277  a:3277  a:3278  a:3278  a:3278  a:3278  a:3279  a:3279  a:3280  a:3281  a:3281  a:3282  a:3282  a:3282  a:3283  a:3283  a:3284  a:3284  a:3284  a:3284  a:3338  a:3338  a:3340  a:3341  a:3342  a:3343  a:3344  a:3345  a:3345  a:3345  a:3345  a:3346  a:3347  a:3347  c:301  c:301  c:301  c:303  c:303  c:303  c:303  c:305  c:305  c:305  c:305  c:305  c:306  c:306  d:302 |
| **10) K...OD1_Asn_ contact(s)** | 3 | A:1722  a:3058  d:302 |
| **10) K...OE2_Glu_ contact(s)** | 1 | c:303 |
| **10) K...OG1_Thr_ contact(s)** | 1 | c:306 |
| **10) K...OG_Ser_ contact(s)** | 2 | a:3058  a:3178 |
| **10) K...OP1 contact(s)** | 42 | A:1602  A:1646  A:1649  A:1658  A:1662  A:1724  A:1724  A:1724  A:1729  A:1735  A:1736  a:3059  a:3062  a:3063  a:3100  a:3100  a:3104  a:3109  a:3110  a:3111  a:3119  a:3119  a:3120  a:3122  a:3126  a:3134  a:3138  a:3141  a:3178  a:3178  a:3249  a:3253  a:3265  a:3265  a:3265  a:3267  a:3270  a:3274  a:3274  a:3275  a:3279  b:206 |
| **10) K...OP2 contact(s)** | 48 | A:1601  A:1654  A:1657  A:1659  A:1661  A:1666  A:1666  A:1723  A:1729  A:1730  A:1730  A:1730  A:1735  A:1736  a:3059  a:3063  a:3063  a:3064  a:3101  a:3104  a:3109  a:3117  a:3117  a:3118  a:3118  a:3119  a:3120  a:3120  a:3121  a:3132  a:3133  a:3140  a:3247  a:3248  a:3248  a:3249  a:3255  a:3269  a:3269  a:3272  a:3275  a:3277  a:3341  a:3342  a:3346  a:3346  a:3347  c:303 |
| **10) MG...N7 contact(s)** | 11 | A:1612  A:1615  A:1674  A:1674  a:3088  a:3170  a:3172  a:3196  a:3220  a:3242  a:3242 |
| **10) MG...ND1_His_ contact(s)** | 1 | c:304 |
| **10) MG...NE2_His_ contact(s)** | 1 | a:3186 |
| **10) MG...O2 contact(s)** | 3 | A:1743  a:3222  a:3302 |
| **10) MG...O2' contact(s)** | 3 | a:3144  a:3145  a:3198 |
| **10) MG...O4 contact(s)** | 12 | A:1623  A:1689  A:1708  a:3027  a:3153  a:3211  a:3298  a:3317  a:3321  a:3332  a:3333  b:205 |
| **10) MG...O6 contact(s)** | 12 | A:1627  A:1633  A:1689  A:1710  a:3008  a:3010  a:3050  a:3083  a:3201  a:3208  a:3312  a:3325 |
| **10) MG...O_back_ contact(s)** | 1 | a:3186 |
| **10) MG...O_w_ contact(s)** | 1017 | A:1606  A:1606  A:1606  A:1606  A:1606  A:1607  A:1607  A:1607  A:1608  A:1608  A:1608  A:1608  A:1610  A:1610  A:1610  A:1611  A:1611  A:1612  A:1612  A:1613  A:1613  A:1613  A:1614  A:1614  A:1614  A:1614  A:1615  A:1615  A:1615  A:1616  A:1616  A:1616  A:1617  A:1618  A:1619  A:1619  A:1620  A:1620  A:1620  A:1620  A:1622  A:1622  A:1627  A:1627  A:1627  A:1627  A:1628  A:1628  A:1628  A:1628  A:1629  A:1629  A:1629  A:1629  A:1629  A:1630  A:1630  A:1631  A:1631  A:1631  A:1631  A:1632  A:1632  A:1632  A:1632  A:1632  A:1633  A:1633  A:1633  A:1634  A:1634  A:1634  A:1635  A:1638  A:1638  A:1638  A:1638  A:1638  A:1639  A:1639  A:1639  A:1642  A:1642  A:1642  A:1642  A:1668  A:1668  A:1668  A:1669  A:1669  A:1669  A:1669  A:1670  A:1670  A:1670  A:1670  A:1670  A:1670  A:1671  A:1671  A:1671  A:1672  A:1672  A:1672  A:1673  A:1673  A:1673  A:1673  A:1673  A:1675  A:1675  A:1675  A:1675  A:1676  A:1676  A:1676  A:1676  A:1677  A:1677  A:1677  A:1678  A:1678  A:1678  A:1678  A:1679  A:1679  A:1679  A:1680  A:1680  A:1680  A:1680  A:1681  A:1681  A:1681  A:1683  A:1683  A:1684  A:1684  A:1684  A:1684  A:1684  A:1684  A:1685  A:1685  A:1686  A:1686  A:1686  A:1687  A:1687  A:1687  A:1687  A:1688  A:1688  A:1688  A:1689  A:1689  A:1690  A:1690  A:1691  A:1691  A:1692  A:1692  A:1693  A:1693  A:1693  A:1694  A:1694  A:1697  A:1698  A:1698  A:1698  A:1699  A:1700  A:1700  A:1700  A:1701  A:1702  A:1703  A:1703  A:1703  A:1705  A:1706  A:1707  A:1708  A:1708  A:1709  A:1710  A:1712  A:1713  A:1713  A:1713  A:1713  A:1716  A:1716  A:1716  A:1741  A:1741  A:1741  A:1742  A:1742  A:1742  A:1742  A:1743  A:1743  A:1743  A:1743  A:1745  A:1745  A:1745  A:1745  A:1745  A:1745  X:101  X:101  X:101  a:3001  a:3001  a:3001  a:3001  a:3001  a:3002  a:3003  a:3003  a:3003  a:3004  a:3004  a:3006  a:3006  a:3007  a:3007  a:3007  a:3007  a:3008  a:3008  a:3008  a:3008  a:3008  a:3009  a:3009  a:3009  a:3010  a:3010  a:3011  a:3012  a:3013  a:3013  a:3013  a:3013  a:3014  a:3014  a:3014  a:3015  a:3016  a:3016  a:3016  a:3016  a:3016  a:3016  a:3017  a:3019  a:3019  a:3019  a:3019  a:3019  a:3019  a:3020  a:3020  a:3020  a:3020  a:3020  a:3021  a:3021  a:3021  a:3021  a:3021  a:3022  a:3022  a:3022  a:3024  a:3024  a:3024  a:3024  a:3025  a:3025  a:3025  a:3025  a:3027  a:3027  a:3028  a:3028  a:3028  a:3030  a:3030  a:3030  a:3030  a:3030  a:3030  a:3033  a:3033  a:3034  a:3034  a:3034  a:3034  a:3034  a:3035  a:3035  a:3035  a:3035  a:3035  a:3036  a:3036  a:3036  a:3036  a:3036  a:3037  a:3037  a:3037  a:3038  a:3038  a:3038  a:3038  a:3039  a:3039  a:3040  a:3040  a:3040  a:3040  a:3041  a:3041  a:3043  a:3046  a:3046  a:3047  a:3047  a:3047  a:3047  a:3047  a:3048  a:3048  a:3048  a:3048  a:3048  a:3049  a:3049  a:3050  a:3050  a:3050  a:3065  a:3065  a:3065  a:3065  a:3066  a:3067  a:3067  a:3067  a:3067  a:3067  a:3067  a:3070  a:3072  a:3072  a:3073  a:3074  a:3075  a:3075  a:3075  a:3076  a:3076  a:3076  a:3076  a:3077  a:3077  a:3077  a:3077  a:3078  a:3078  a:3078  a:3078  a:3078  a:3079  a:3079  a:3079  a:3080  a:3080  a:3080  a:3080  a:3080  a:3081  a:3081  a:3082  a:3082  a:3082  a:3082  a:3082  a:3083  a:3083  a:3083  a:3084  a:3084  a:3084  a:3084  a:3085  a:3085  a:3085  a:3085  a:3085  a:3086  a:3086  a:3086  a:3086  a:3086  a:3087  a:3087  a:3087  a:3087  a:3088  a:3088  a:3088  a:3089  a:3089  a:3089  a:3090  a:3090  a:3090  a:3090  a:3090  a:3091  a:3091  a:3092  a:3092  a:3092  a:3092  a:3093  a:3093  a:3093  a:3093  a:3094  a:3094  a:3094  a:3095  a:3095  a:3095  a:3095  a:3095  a:3095  a:3143  a:3143  a:3143  a:3143  a:3144  a:3145  a:3145  a:3146  a:3146  a:3146  a:3146  a:3147  a:3147  a:3147  a:3147  a:3147  a:3149  a:3149  a:3149  a:3150  a:3150  a:3150  a:3150  a:3151  a:3152  a:3152  a:3153  a:3153  a:3153  a:3153  a:3153  a:3154  a:3154  a:3154  a:3154  a:3154  a:3155  a:3155  a:3155  a:3156  a:3156  a:3156  a:3156  a:3156  a:3157  a:3157  a:3157  a:3157  a:3157  a:3158  a:3158  a:3159  a:3159  a:3160  a:3160  a:3160  a:3161  a:3161  a:3161  a:3161  a:3161  a:3163  a:3163  a:3163  a:3164  a:3164  a:3164  a:3164  a:3165  a:3165  a:3165  a:3165  a:3165  a:3166  a:3166  a:3166  a:3167  a:3167  a:3168  a:3170  a:3170  a:3170  a:3171  a:3171  a:3171  a:3171  a:3172  a:3172  a:3172  a:3174  a:3174  a:3174  a:3174  a:3174  a:3182  a:3182  a:3182  a:3182  a:3182  a:3182  a:3183  a:3183  a:3183  a:3183  a:3184  a:3184  a:3184  a:3184  a:3184  a:3185  a:3185  a:3185  a:3185  a:3185  a:3186  a:3186  a:3187  a:3187  a:3187  a:3188  a:3188  a:3188  a:3188  a:3188  a:3189  a:3189  a:3190  a:3190  a:3190  a:3191  a:3192  a:3193  a:3193  a:3193  a:3193  a:3193  a:3194  a:3194  a:3194  a:3194  a:3195  a:3195  a:3197  a:3197  a:3197  a:3197  a:3197  a:3198  a:3198  a:3198  a:3198  a:3198  a:3199  a:3199  a:3199  a:3199  a:3199  a:3200  a:3200  a:3200  a:3200  a:3201  a:3201  a:3201  a:3202  a:3202  a:3202  a:3204  a:3204  a:3204  a:3205  a:3205  a:3205  a:3205  a:3205  a:3205  a:3206  a:3206  a:3206  a:3206  a:3207  a:3207  a:3207  a:3207  a:3208  a:3209  a:3209  a:3209  a:3210  a:3210  a:3210  a:3210  a:3210  a:3210  a:3211  a:3211  a:3211  a:3211  a:3212  a:3212  a:3212  a:3212  a:3212  a:3212  a:3213  a:3213  a:3213  a:3213  a:3213  a:3214  a:3214  a:3214  a:3214  a:3215  a:3215  a:3215  a:3215  a:3216  a:3216  a:3216  a:3216  a:3216  a:3216  a:3217  a:3217  a:3217  a:3217  a:3217  a:3218  a:3218  a:3218  a:3218  a:3218  a:3219  a:3219  a:3219  a:3220  a:3220  a:3220  a:3220  a:3221  a:3221  a:3221  a:3221  a:3221  a:3221  a:3222  a:3222  a:3222  a:3222  a:3223  a:3223  a:3223  a:3224  a:3224  a:3225  a:3225  a:3225  a:3226  a:3226  a:3226  a:3226  a:3227  a:3227  a:3227  a:3227  a:3228  a:3228  a:3228  a:3228  a:3229  a:3229  a:3229  a:3229  a:3229  a:3229  a:3230  a:3231  a:3231  a:3231  a:3232  a:3232  a:3232  a:3232  a:3232  a:3233  a:3233  a:3233  a:3234  a:3234  a:3234  a:3234  a:3234  a:3235  a:3235  a:3235  a:3235  a:3235  a:3236  a:3236  a:3236  a:3237  a:3237  a:3237  a:3237  a:3238  a:3238  a:3239  a:3239  a:3239  a:3239  a:3240  a:3240  a:3240  a:3240  a:3240  a:3241  a:3241  a:3241  a:3241  a:3241  a:3242  a:3242  a:3242  a:3243  a:3243  a:3243  a:3244  a:3244  a:3244  a:3245  a:3245  a:3245  a:3245  a:3285  a:3285  a:3285  a:3286  a:3286  a:3286  a:3286  a:3286  a:3286  a:3287  a:3288  a:3288  a:3288  a:3288  a:3289  a:3289  a:3289  a:3289  a:3289  a:3290  a:3290  a:3290  a:3291  a:3291  a:3291  a:3292  a:3292  a:3293  a:3293  a:3293  a:3293  a:3294  a:3294  a:3294  a:3295  a:3295  a:3296  a:3296  a:3296  a:3296  a:3297  a:3297  a:3297  a:3297  a:3297  a:3297  a:3298  a:3298  a:3299  a:3299  a:3299  a:3299  a:3300  a:3300  a:3300  a:3300  a:3301  a:3301  a:3301  a:3301  a:3301  a:3302  a:3302  a:3303  a:3303  a:3303  a:3303  a:3303  a:3304  a:3304  a:3305  a:3305  a:3305  a:3305  a:3306  a:3306  a:3306  a:3307  a:3307  a:3308  a:3308  a:3308  a:3309  a:3309  a:3309  a:3310  a:3310  a:3311  a:3311  a:3311  a:3311  a:3311  a:3311  a:3312  a:3312  a:3312  a:3313  a:3313  a:3314  a:3314  a:3314  a:3315  a:3315  a:3315  a:3316  a:3316  a:3317  a:3317  a:3317  a:3317  a:3318  a:3318  a:3318  a:3319  a:3319  a:3319  a:3319  a:3319  a:3320  a:3320  a:3320  a:3320  a:3321  a:3321  a:3321  a:3321  a:3322  a:3322  a:3322  a:3322  a:3323  a:3323  a:3323  a:3324  a:3324  a:3325  a:3325  a:3326  a:3326  a:3326  a:3327  a:3327  a:3328  a:3328  a:3329  a:3329  a:3331  a:3331  a:3331  a:3332  a:3332  a:3332  a:3332  a:3333  a:3334  a:3334  a:3334  a:3335  a:3335  a:3335  a:3335  a:3336  a:3336  a:3336  a:3337  a:3337  a:3337  a:3337  a:3348  a:3348  a:3348  a:3348  a:3348  a:3349  a:3349  a:3349  a:3350  a:3350  a:3350  a:3350  a:3350  a:3351  a:3351  a:3351  a:3352  a:3352  a:3352  a:3353  a:3353  a:3353  a:3353  a:3356  a:3356  a:3357  a:3357  a:3357  a:3357  a:3357  a:3359  a:3359  a:3359  a:3359  a:3360  a:3360  a:3360  a:3360  a:3361  a:3361  a:3361  a:3361  a:3362  a:3362  a:3363  a:3364  a:3364  a:3364  b:204  b:204  b:204  b:205  b:205  b:205  b:205  b:205  c:302  c:302  c:302  c:302  c:304  c:304  c:304  c:304  d:301  d:301  z:101  z:101  z:101  z:101 |
| **10) MG...OD1_Asn_ contact(s)** | 1 | a:3219 |
| **10) MG...OD1_Asp_ contact(s)** | 1 | z:101 |
| **10) MG...OP1 contact(s)** | 128 | A:1606  A:1608  A:1619  A:1619  A:1620  A:1624  A:1626  A:1627  A:1628  A:1634  A:1634  A:1634  A:1638  A:1642  A:1683  A:1691  A:1700  A:1700  A:1701  A:1701  A:1704  A:1706  A:1713  A:1716  A:1717  A:1743  a:3003  a:3009  a:3020  a:3040  a:3040  a:3046  a:3048  a:3072  a:3079  a:3081  a:3083  a:3084  a:3084  a:3086  a:3087  a:3088  a:3089  a:3089  a:3091  a:3143  a:3143  a:3144  a:3147  a:3149  a:3150  a:3151  a:3152  a:3152  a:3154  a:3156  a:3159  a:3164  a:3167  a:3167  a:3174  a:3183  a:3187  a:3188  a:3192  a:3193  a:3194  a:3194  a:3195  a:3201  a:3201  a:3204  a:3206  a:3209  a:3223  a:3223  a:3224  a:3225  a:3225  a:3226  a:3227  a:3228  a:3230  a:3232  a:3233  a:3233  a:3236  a:3236  a:3237  a:3238  a:3238  a:3238  a:3239  a:3243  a:3243  a:3244  a:3244  a:3287  a:3289  a:3299  a:3304  a:3309  a:3309  a:3310  a:3313  a:3316  a:3320  a:3322  a:3322  a:3325  a:3325  a:3326  a:3326  a:3327  a:3328  a:3329  a:3329  a:3329  a:3329  a:3331  a:3331  a:3333  a:3336  a:3336  a:3349  a:3351  a:3356  a:3363 |
| **10) MG...OP2 contact(s)** | 131 | A:1607  A:1623  A:1626  A:1642  A:1669  A:1671  A:1671  A:1671  A:1672  A:1672  A:1673  A:1675  A:1677  A:1679  A:1679  A:1680  A:1681  A:1681  A:1681  A:1683  A:1686  A:1689  A:1699  A:1701  A:1702  A:1702  A:1717  A:1720  A:1742  a:3004  a:3007  a:3023  a:3023  a:3025  a:3027  a:3036  a:3046  a:3047  a:3049  a:3078  a:3080  a:3082  a:3087  a:3088  a:3089  a:3090  a:3093  a:3093  a:3144  a:3145  a:3145  a:3150  a:3155  a:3159  a:3161  a:3165  a:3166  a:3170  a:3185  a:3186  a:3186  a:3192  a:3199  a:3200  a:3207  a:3207  a:3219  a:3219  a:3224  a:3224  a:3230  a:3231  a:3231  a:3233  a:3234  a:3237  a:3238  a:3240  a:3244  a:3245  a:3285  a:3287  a:3288  a:3290  a:3292  a:3293  a:3294  a:3294  a:3295  a:3295  a:3296  a:3298  a:3300  a:3303  a:3304  a:3305  a:3306  a:3307  a:3308  a:3308  a:3312  a:3314  a:3314  a:3314  a:3315  a:3316  a:3319  a:3323  a:3324  a:3324  a:3324  a:3326  a:3327  a:3327  a:3328  a:3328  a:3331  a:3333  a:3333  a:3334  a:3335  a:3337  a:3348  a:3351  a:3353  a:3357  a:3363  b:202  b:202  b:204  c:304 |
| **10) ZN...ND1_His_ contact(s)** | 1 | 3:101 |
| **10) ZN...SG_Cys_ contact(s)** | 3 | 3:101  3:101  3:101 |

---

\_\_\_\_\_\_\_\_\_\_\_\_\_\_\_\_\_\_\_\_\_\_\_\_\_\_\_\_\_\_\_\_\_\_\_\_\_\_\_\_\_\_\_\_\_\_\_\_\_\_\_\_\_\_\_\_\_\_\_\_\_\_\_\_  

#### 4) The following *"ion/ligand"* tables list all *"ion coordination contacts"* up to 4.8 Å

*(4.8 Å corresponds to the second coordination shell limit)  
Go to ToC

### ZN:3:101

|  | Res. type | Atom | Residue | Distance | Occupancy | B-factor | Closest atom(s) to a 1^st^ shell water | EM\_dens. |
| --- | --- | --- | --- | --- | --- | --- | --- | --- |
| **Ion** | | | |  | 1.0 | 81.2 |  | 15.04 |
 1 | HIS | ND1 | 3:33 | 1.96 | 1.0 | 41.0 |  | 5.07 | 2 | CYS | SG | 3:14 | 2.30 | 1.0 | 49.6 |  | 12.34 | 3 | CYS | SG | 3:11 | 2.33 | 1.0 | 57.9 |  | 11.47 | 4 | CYS | SG | 3:27 | 2.40 | 1.0 | 52.5 |  | 10.15 | 5 | HIS | NE2 | 3:33 | 4.00 | 1.0 | 36.0 |  | 6.41 | 6 | CYS | O | 3:11 | 4.36 | 1.0 | 51.3 |  | 4.17 |

  
**1^st^ shell coordination angles and distances (*trans* in "blue"; *cys* in "green"; deviations in "orange"):**   
• 1-2(ND1/SG): 105; 1-3(ND1/SG): 117; 1-4(ND1/SG): 111; 2-3(SG/SG): 121; 2-4(SG/SG): 102; 3-4(SG/SG): 100;   
• 1-2(ND1/SG): 3.4; 1-3(ND1/SG): 3.7; 1-4(ND1/SG): 3.6; 2-3(SG/SG): 4.0; 2-4(SG/SG): 3.7; 3-4(SG/SG): 3.6;   
  
**1.50 ± 0.63: the average coodination distance of the 1^st^ six ligands is Mg^2+^ compatible**  
**0: Number of potential Water contacts**   
• Contact(s): /ND1_His_/SG_Cys_/SG_Cys_/SG_Cys_/  
• Contact code: N_His_3SG_Cys_  
→ ZN ion B\_factor > 80 ←  
Go to ToC

### K:A:1601

|  | Res. type | Atom | Residue | Distance | Occupancy | B-factor | Closest atom(s) to a 1^st^ shell water | EM\_dens. |
| --- | --- | --- | --- | --- | --- | --- | --- | --- |
| **Ion** | | | |  | 1.0 | 30.0 |  | 3.36 |
 1 | U | OP2 | A:268 | 2.81 | 1.0 | 66.5 |  | 2.53 | 2 | G | O2' | A:251 | 2.88 | 1.0 | 72.2 |  | 2.34 | 3 | G | O2' | A:266 | 2.95 | 1.0 | 69.0 |  | 2.14 | 4 | G | O3' | A:251 | 3.25 | 1.0 | 74.2 |  | 2.81 | 5 | G | O3' | A:266 | 3.68 | 1.0 | 65.7 |  | 3.79 | 6 | U | OP1 | A:268 | 3.77 | 1.0 | 69.1 |  | 1.71 | 7 | U | OP2 | A:252 | 3.82 | 1.0 | 70.6 |  | 2.88 | 8 | G | N9 | A:266 | 4.48 | 1.0 | 61.4 |  | 4.70 | 9 | C | O3' | A:267 | 4.79 | 1.0 | 68.6 |  | 3.17 | 10 | C | OP2 | A:267 | 4.79 | 1.0 | 66.2 |  | 3.20 |

  
**1^st^ shell coordination angles and distances (*trans* in "blue"; *cys* in "green"; deviations in "orange"):**   
• 1-2(OP2/O2'): 84; 1-3(OP2/O2'): 98; 2-3(O2'/O2'): 97;   
• 1-2(OP2/O2'): 3.8; 1-3(OP2/O2'): 4.3; 2-3(O2'/O2'): 4.4;   
  
**4.61 ± 1.47: the average coodination distance of the 1^st^ six ligands could correspond to an hexahydrated Mg^2+^.**  
**0: Number of potential Water contacts**   
• Contact(s): /O2'/O2'/OP2/  
• Contact code: O_ph_2O_r_  
→ Coordination < 6 for K ←  
→2Fo-Fc density between 1.0 and 4.0 !   
Go to ToC

### K:A:1602

|  | Res. type | Atom | Residue | Distance | Occupancy | B-factor | Closest atom(s) to a 1^st^ shell water | EM\_dens. |
| --- | --- | --- | --- | --- | --- | --- | --- | --- |
| **Ion** | | | |  | 1.0 | 30.0 |  | 4.24 |
 1 | Water 1 | | A:2192 | 2.59 | 1.0 | 72.2 | 2.55 [U:O4]A:268 // 2.93 [C:N4]A:269 | 1.78 | 2 | G | OP1 | A:266 | 2.85 | 1.0 | 68.7 |  | 2.61 | 3 | Water 2 | | A:4175 | 3.06 | 1.0 | 70.7 | 2.84 [MG:MG]A:1744 | 2.01 | 4 | Water 3 | | A:4284 | 3.17 | 1.0 | 70.4 |  | 1.68 | 5 | U | O4 | A:256 | 3.24 | 1.0 | 74.5 |  | 2.20 | 6 | **MG** | MG | A:1744 | 3.48 | 1.0 | 72.5 |  | 2.35 | 7 | G | O6 | A:257 | 3.61 | 1.0 | 72.0 |  | 2.75 | 8 | G | N2 | A:266 | 3.82 | 1.0 | 66.3 |  | 3.04 | 9 | G | OP2 | A:266 | 3.92 | 1.0 | 69.1 |  | 2.31 | 10 | Water 4 | | A:1911 | 4.16 | 1.0 | 69.3 |  | 2.37 | 11 | G | O5' | A:266 | 4.16 | 1.0 | 66.9 |  | 2.50 | 12 | **MG** | MG | A:1637 | 4.19 | 1.0 | 63.0 |  | 3.29 | 13 | U | O4 | A:268 | 4.27 | 1.0 | 67.3 |  | 2.56 | 14 | G | N3 | A:266 | 4.56 | 1.0 | 67.4 |  | 3.02 |

  
**1^st^ shell coordination angles and distances (*trans* in "blue"; *cys* in "green"; deviations in "orange"):**   
• 1-2(W/OP1): 89; 1-3(W/W): 125; 1-4(W/W): 137; 2-3(OP1/W): 114; 2-4(OP1/W): 73; 3-4(W/W): 97;   
• 1-2(W/OP1): 3.8; 1-3(W/W): 5.0; 1-4(W/W): 5.4; 2-3(OP1/W): 5.0; 2-4(OP1/W): 3.6; 3-4(W/W): 4.7;   
  
**4.30 ± 1.32: the average coodination distance of the 1^st^ six ligands could correspond to an hexahydrated Mg^2+^.**  
**6: Number of potential Water contacts**   
**3 : Number of second shell contacts involving water or ions**   
**50%: Percentage of satisfied second shell Water contacts**   
• Contact(s): /O_w_/O_w_/OP1/  
• Contact code: O_ph_2O_w_  
• Contact(s) in the exclusion zone: /O_w_/  
→ Coordination < 6 for K ←  
Go to ToC

### K:A:1603

|  | Res. type | Atom | Residue | Distance | Occupancy | B-factor | Closest atom(s) to a 1^st^ shell water | EM\_dens. |
| --- | --- | --- | --- | --- | --- | --- | --- | --- |
| **Ion** | | | |  | 1.0 | 30.0 |  | 3.72 |
 1 | G | O6 | A:111 | 2.58 | 1.0 | 53.2 |  | 3.74 | 2 | G | O6 | A:112 | 3.20 | 1.0 | 51.6 |  | 4.27 | 3 | G | N7 | A:111 | 3.35 | 1.0 | 50.9 |  | 4.50 | 4 | Water 1 | | A:2152 | 3.52 | 1.0 | 55.6 | 2.54 [A:OP2]A:109 | 1.96 | 5 | C | OP2 | A:110 | 3.54 | 1.0 | 52.0 |  | 5.17 | 6 | A | OP2 | A:109 | 3.67 | 1.0 | 48.1 |  | 4.81 | 7 | C | N4 | A:330 | 4.10 | 1.0 | 52.0 |  | 2.09 | 8 | G | N7 | A:112 | 4.17 | 1.0 | 45.7 |  | 4.84 | 9 | Water 2 | | A:3215 | 4.53 | 1.0 | 53.6 | 2.84 [A:OP2]A:109 | 2.67 | 10 | Water 3 | | A:1935 | 4.76 | 1.0 | 56.5 | 2.94 [G:N7]A:112 | 2.10 |

  
• • **4.69 ± 1.46: the average coodination distance of the 1^st^ six ligands could correspond to an hexahydrated Mg^2+^.**  
**2: Number of potential Water contacts**   
**1 : Number of second shell contacts involving water or ions**   
**50%: Percentage of satisfied second shell Water contacts**   
• Contact code: → No contacts ←   
• Contact(s) in the exclusion zone: /O6/  
→ First K coordination shell missing  
→2Fo-Fc density between 1.0 and 4.0 !   
Go to ToC

### K:A:1604

|  | Res. type | Atom | Residue | Distance | Occupancy | B-factor | Closest atom(s) to a 1^st^ shell water | EM\_dens. |
| --- | --- | --- | --- | --- | --- | --- | --- | --- |
| **Ion** | | | |  | 1.0 | 30.0 |  | 10.02 |
 1 | C | O2' | A:811 | 2.78 | 1.0 | 33.1 |  | 10.45 | 2 | Water 1 | | A:4647 | 2.80 | 1.0 | 30.0 |  | 2.99 | 3 | A | N1 | A:768 | 2.84 | 1.0 | 26.4 |  | 13.15 | 4 | C | O2 | A:811 | 3.05 | 1.0 | 32.4 |  | 10.56 | 5 | Water 2 | | A:4092 | 3.48 | 1.0 | 41.7 | 3.00 [G:O4']A:812 | 4.41 | 6 | A | N6 | A:768 | 4.03 | 1.0 | 25.2 |  | 12.65 | 7 | Water 3 | | A:2037 | 4.12 | 1.0 | 36.7 |  | 7.02 | 8 | Water 4 | | A:4395 | 4.34 | 1.0 | 46.0 |  | 3.49 | 9 | A | N1 | A:901 | 4.38 | 1.0 | 29.8 |  | 9.99 | 10 | Water 5 | | A:2143 | 4.57 | 1.0 | 40.7 |  | 5.35 | 11 | Water 6 | | A:4055 | 4.65 | 1.0 | 42.6 |  | 3.89 | 12 | C | N1 | A:811 | 4.66 | 1.0 | 29.4 |  | 12.16 | 13 | G | O4' | A:812 | 4.72 | 1.0 | 36.3 |  | 8.96 | 14 | A | N3 | A:768 | 4.78 | 1.0 | 23.8 |  | 12.43 |

  
**1^st^ shell coordination angles and distances (*trans* in "blue"; *cys* in "green"; deviations in "orange"):**   
• 1-2(O2'/W): 111; 1-3(O2'/N1): 140; 1-4(O2'/O2): 73; 2-3(W/N1): 90; 2-4(W/O2): 163; 3-4(N1/O2): 78;   
• 1-2(O2'/W): 4.6; 1-3(O2'/N1): 5.3; 1-4(O2'/O2): 3.5; 2-3(W/N1): 4.0; 2-4(W/O2): 5.8; 3-4(N1/O2): 3.7;   
  
**4.57 ± 1.50: the average coodination distance of the 1^st^ six ligands could correspond to an hexahydrated Mg^2+^.**  
**8: Number of potential Water contacts**   
**1 : Number of second shell contacts involving water or ions**   
**12%: Percentage of satisfied second shell Water contacts**   
• Contact(s): /N1_A_/O2/O2'/O_w_/  
• Contact code: O_r_O_b_N_b_O_w_  
→ Coordination < 6 for K ←  
Go to ToC

### K:A:1605

|  | Res. type | Atom | Residue | Distance | Occupancy | B-factor | Closest atom(s) to a 1^st^ shell water | EM\_dens. |
| --- | --- | --- | --- | --- | --- | --- | --- | --- |
| **Ion** | | | |  | 1.0 | 30.0 |  | 3.77 |
 1 | G | O6 | A:177 | 2.58 | 1.0 | 80.2 |  | 1.84 | 2 | G | O6 | A:145 | 2.85 | 1.0 | 84.6 |  | 2.10 | 3 | G | O6 | A:144 | 3.15 | 1.0 | 85.6 |  | 1.35 | 4 | A | OP2 | A:197 | 3.45 | 1.0 | 81.3 |  | 2.00 | 5 | C | N4 | A:178 | 3.57 | 1.0 | 80.4 |  | 2.60 | 6 | G | N1 | A:177 | 3.60 | 1.0 | 79.0 |  | 2.55 | 7 | G | N1 | A:145 | 3.70 | 1.0 | 83.8 |  | 3.09 | 8 | Water 1 | | A:3967 | 4.42 | 1.0 | 74.4 |  | 2.43 | 9 | A | OP2 | A:196 | 4.54 | 1.0 | 80.8 |  | 2.03 | 10 | G | O6 | A:146 | 4.54 | 1.0 | 82.8 |  | 1.88 |

  
**1^st^ shell coordination angles and distances (*trans* in "blue"; *cys* in "green"; deviations in "orange"):**   
• 1-2(O6/O6): 83; 1-3(O6/O6): 95; 2-3(O6/O6): 66;   
• 1-2(O6/O6): 3.6; 1-3(O6/O6): 4.3; 2-3(O6/O6): 3.3;   
  
**4.55 ± 1.45: the average coodination distance of the 1^st^ six ligands could correspond to an hexahydrated Mg^2+^.**  
**2: Number of potential Water contacts**   
**0 : Number of second shell contacts involving water or ions**   
**0%: Percentage of satisfied second shell Water contacts**   
• Contact(s): /O6/O6/  
• Contact code: 2O_b_  
• Contact(s) in the exclusion zone: /O6/  
→ Coordination < 6 for K ←  
→2Fo-Fc density between 1.0 and 4.0 !   
Go to ToC

### MG:A:1606

|  | Res. type | Atom | Residue | Distance | Occupancy | B-factor | Closest atom(s) to a 1^st^ shell water | EM\_dens. |
| --- | --- | --- | --- | --- | --- | --- | --- | --- |
| **Ion** | | | |  | 1.0 | 36.2 |  | 14.98 |
 1 | Water 1 | | A:2871 | 1.91 | 1.0 | 33.6 | 2.83 [C:OP2]A:817 | 8.25 | 2 | C | OP1 | A:578 | 2.07 | 1.0 | 35.5 |  | 8.54 | 3 | Water 2 | | A:2494 | 2.08 | 1.0 | 39.3 | 2.64 [U:OP1]A:820 | 8.19 | 4 | Water 3 | | A:3307 | 2.13 | 1.0 | 32.4 |  | 7.39 | 5 | Water 4 | | A:2649 | 2.17 | 1.0 | 34.7 | 2.68 [C:OP2]A:576 // 2.98 [G:O3']A:577 | 10.76 | 6 | Water 5 | | A:3599 | 2.25 | 1.0 | 33.3 |  | 9.49 | 7 | G | O3' | A:577 | 3.68 | 1.0 | 36.5 |  | 8.00 | 8 | C | OP2 | A:576 | 4.05 | 1.0 | 39.4 |  | 8.38 | 9 | Water 6 | | A:3959 | 4.16 | 1.0 | 45.5 | 2.96 [C:OP2]A:578 | 4.54 | 10 | U | OP1 | A:820 | 4.18 | 1.0 | 38.0 |  | 8.19 | 11 | C | OP2 | A:817 | 4.22 | 1.0 | 36.9 |  | 8.57 | 12 | Water 7 | | A:3806 | 4.27 | 1.0 | 43.1 |  | 5.68 | 13 | C | OP2 | A:578 | 4.31 | 1.0 | 31.7 |  | 10.24 | 14 | Water 8 | | A:4566 | 4.38 | 1.0 | 48.9 |  | 2.74 | 15 | Water 9 | | A:3908 | 4.38 | 1.0 | 42.4 |  | 5.38 | 16 | C | O5' | A:578 | 4.43 | 1.0 | 34.9 |  | 9.63 | 17 | G | O3' | A:575 | 4.80 | 1.0 | 35.7 |  | 9.27 |

  
**1^st^ shell coordination angles and distances (*trans* in "blue"; *cys* in "green"; deviations in "orange"):**   
• 1-2(W/OP1): 86º; 1-3(W/W): 94º; 1-4(W/W): 89º; 1-5(W/W): 88º; 1-6(W/W): *168º*; 2-3(OP1/W): 179º; 2-4(OP1/W): *84º*; 2-5(OP1/W): 91º; 2-6(OP1/W): *83º*; 3-4(W/W): *96º*; 3-5(W/W): 90º; 3-6(W/W): *98º*; 4-5(W/W): 174º; 4-6(W/W): *95º*; 5-6(W/W): 87º;   
• 1-2(W/OP1): 2.7Å; 1-3(W/W): 2.9Å; 1-4(W/W): 2.8Å; 1-5(W/W): 2.8Å; 1-6(W/W): 4.1Å; 2-3(OP1/W): 4.2Å; 2-4(OP1/W): 2.8Å; 2-5(OP1/W): 3.0Å; 2-6(OP1/W): 2.9Å; 3-4(W/W): 3.1Å; 3-5(W/W): 3.0Å; 3-6(W/W): *3.3Å*; 4-5(W/W): *4.3Å*; 4-6(W/W): *3.2Å*; 5-6(W/W): 3.0Å;   
  
**2.10 ± 0.10: the average coodination distance of the 1^st^ six ligands is Mg^2+^ compatible**  
**10: Number of potential Water contacts**   
**4 : Number of second shell contacts involving water or ions**   
**40%: Percentage of satisfied second shell Water contacts**   
• Contact(s): /O_w_/O_w_/O_w_/O_w_/O_w_/OP1/  
• Contact code: O_ph_5O_w_  
Ooctahedral coordination (4.4º)   
Go to ToC

### MG:A:1607

|  | Res. type | Atom | Residue | Distance | Occupancy | B-factor | Closest atom(s) to a 1^st^ shell water | EM\_dens. |
| --- | --- | --- | --- | --- | --- | --- | --- | --- |
| **Ion** | | | |  | 1.0 | 62.0 |  | 6.04 |
 1 | Water 1 | | A:3691 | 1.89 | 1.0 | 53.1 |  | 4.63 | 2 | Water 2 | | A:2865 | 1.90 | 1.0 | 49.3 | 2.93 [G:N7]A:354 | 4.75 | 3 | Water 3 | | A:3374 | 1.92 | 1.0 | 52.0 | 2.81 [A:N1]A:59 // 2.95 [C:O2]A:330 | 5.13 | 4 | C | OP2 | A:352 | 2.09 | 1.0 | 51.8 |  | 5.36 | 5 | Water 4 | | A:4374 | 2.13 | 1.0 | 51.4 |  | 4.28 | 6 | Water 5 | | A:3753 | 2.29 | 1.0 | 54.2 |  | 2.99 | 7 | Water 6 | | A:4085 | 3.66 | 1.0 | 52.2 |  | 2.52 | 8 | C | O2 | A:330 | 3.79 | 1.0 | 51.1 |  | 3.63 | 9 | C | O5' | A:352 | 3.84 | 1.0 | 54.3 |  | 3.63 | 10 | G | O4' | A:331 | 4.10 | 1.0 | 54.2 |  | 4.01 | 11 | Water 7 | | A:3871 | 4.14 | 1.0 | 57.6 | 2.93 [C:OP1]A:352 | 2.63 | 12 | G | N7 | A:354 | 4.39 | 1.0 | 47.3 |  | 6.47 | 13 | C | O4' | A:352 | 4.39 | 1.0 | 52.5 |  | 4.79 | 14 | C | OP1 | A:352 | 4.42 | 1.0 | 46.7 |  | 6.81 | 15 | G | O3' | A:351 | 4.46 | 1.0 | 55.5 |  | 4.00 | 16 | Water 8 | | A:4117 | 4.48 | 1.0 | 53.6 |  | 1.62 | 17 | A | N1 | A:59 | 4.51 | 1.0 | 48.4 |  | 5.35 | 18 | Water 9 | | A:4601 | 4.61 | 1.0 | 57.5 |  | 1.94 | 19 | G | OP1 | A:354 | 4.73 | 1.0 | 48.1 |  | 5.21 |

  
**1^st^ shell coordination angles and distances (*trans* in "blue"; *cys* in "green"; deviations in "orange"):**   
• 1-2(W/W): 177º; 1-3(W/W): 93º; 1-4(W/OP2): 93º; 1-5(W/W): *102º*; 1-6(W/W): *97º*; 2-3(W/W): 88º; 2-4(W/OP2): 86º; 2-5(W/W): *82º*; 2-6(W/W): *80º*; 3-4(W/OP2): *166º*; 3-5(W/W): 95º; 3-6(W/W): *84º*; 4-5(OP2/W): *97º*; 4-6(OP2/W): *83º*; 5-6(W/W): *162º*;   
• 1-2(W/W): *3.8Å*; 1-3(W/W): 2.8Å; 1-4(W/OP2): 2.9Å; 1-5(W/W): 3.1Å; 1-6(W/W): 3.1Å; 2-3(W/W): 2.7Å; 2-4(W/OP2): 2.7Å; 2-5(W/W): 2.6Å; 2-6(W/W): 2.7Å; 3-4(W/OP2): 4.0Å; 3-5(W/W): 3.0Å; 3-6(W/W): 2.8Å; 4-5(OP2/W): 3.2Å; 4-6(OP2/W): 2.9Å; 5-6(W/W): *4.4Å*;   
  
**2.04 ± 0.15: the average coodination distance of the 1^st^ six ligands is Mg^2+^ compatible**  
**10: Number of potential Water contacts**   
**3 : Number of second shell contacts involving water or ions**   
**30%: Percentage of satisfied second shell Water contacts**   
• Contact(s): /O_w_/O_w_/O_w_/OP2/  
• Contact code: O_ph_3O_w_  
• Contact(s) in the exclusion zone: /O_w_/O_w_/  
→ Missing octahedral coordination for MG ←  
→ Slightly distorded coord. octahedron (7.2º) ←  
Go to ToC

### MG:A:1608

|  | Res. type | Atom | Residue | Distance | Occupancy | B-factor | Closest atom(s) to a 1^st^ shell water | EM\_dens. |
| --- | --- | --- | --- | --- | --- | --- | --- | --- |
| **Ion** | | | |  | 1.0 | 65.4 |  | 4.09 |
 1 | Water 1 | | A:4086 | 1.89 | 1.0 | 60.9 | 2.99 [G:N7]A:402 | 3.12 | 2 | Water 2 | | A:3675 | 1.90 | 1.0 | 65.3 |  | 2.28 | 3 | A | OP1 | A:547 | 1.95 | 1.0 | 61.3 |  | 3.54 | 4 | Water 3 | | A:2759 | 1.99 | 1.0 | 63.1 | 2.79 [G:OP1]A:548 | 2.62 | 5 | Water 4 | | A:4375 | 2.16 | 1.0 | 62.6 |  | 2.60 | 6 | Water 5 | | A:4163 | 2.18 | 1.0 | 62.4 |  | 2.54 | 7 | G | OP1 | A:548 | 3.96 | 1.0 | 56.3 |  | 3.11 | 8 | A | OP2 | A:547 | 3.97 | 1.0 | 62.9 |  | 3.17 | 9 | Water 6 | | A:2531 | 4.12 | 1.0 | 64.1 |  | 1.89 | 10 | A | O5' | A:547 | 4.23 | 1.0 | 61.8 |  | 3.45 | 11 | G | O5' | A:548 | 4.29 | 1.0 | 58.2 |  | 2.48 | 12 | A | O3' | A:546 | 4.34 | 1.0 | 63.1 |  | 3.12 | 13 | Water 7 | | A:3210 | 4.40 | 1.0 | 62.4 |  | 1.91 | 14 | G | N7 | A:402 | 4.49 | 1.0 | 62.2 |  | 3.80 | 15 | Water 8 | | A:3512 | 4.51 | 1.0 | 59.2 |  | 1.72 | 16 | G | O6 | A:402 | 4.76 | 1.0 | 63.4 |  | 3.80 |

  
**1^st^ shell coordination angles and distances (*trans* in "blue"; *cys* in "green"; deviations in "orange"):**   
• 1-2(W/W): 91º; 1-3(W/OP1): *168º*; 1-4(W/W): 86º; 1-5(W/W): *73º*; 1-6(W/W): *96º*; 2-3(W/OP1): *97º*; 2-4(W/W): 176º; 2-5(W/W): *84º*; 2-6(W/W): 92º; 3-4(OP1/W): 86º; 3-5(OP1/W): *98º*; 3-6(OP1/W): 94º; 4-5(W/W): 94º; 4-6(W/W): 90º; 5-6(W/W): *168º*;   
• 1-2(W/W): 2.7Å; 1-3(W/OP1): 3.8Å; 1-4(W/W): 2.6Å; 1-5(W/W): 2.4Å; 1-6(W/W): 3.0Å; 2-3(W/OP1): 2.9Å; 2-4(W/W): 3.9Å; 2-5(W/W): 2.7Å; 2-6(W/W): 2.9Å; 3-4(OP1/W): 2.7Å; 3-5(OP1/W): 3.1Å; 3-6(OP1/W): 3.0Å; 4-5(W/W): 3.0Å; 4-6(W/W): 3.0Å; 5-6(W/W): *4.3Å*;   
  
**2.01 ± 0.12: the average coodination distance of the 1^st^ six ligands is Mg^2+^ compatible**  
**10: Number of potential Water contacts**   
**2 : Number of second shell contacts involving water or ions**   
**20%: Percentage of satisfied second shell Water contacts**   
• Contact(s): /O_w_/O_w_/O_w_/O_w_/OP1/  
• Contact code: O_ph_4O_w_  
• Contact(s) in the exclusion zone: /O_w_/  
→ Missing octahedral coordination for MG ←  
→ Slightly distorded coord. octahedron (6.1º) ←  
Go to ToC

### MG:A:1609

|  | Res. type | Atom | Residue | Distance | Occupancy | B-factor | Closest atom(s) to a 1^st^ shell water | EM\_dens. |
| --- | --- | --- | --- | --- | --- | --- | --- | --- |
| **Ion** | | | |  | 1.0 | 38.6 |  | 10.91 |
 1 | Water 1 | | A:2722 | 2.61 | 1.0 | 39.9 | 2.69 [G:N7]A:894 | 6.54 | 2 | Water 2 | | A:4050 | 2.62 | 1.0 | 41.9 | 2.98 [C:N4]A:893 | 6.70 | 3 | U | O4 | A:904 | 2.78 | 1.0 | 36.2 |  | 10.41 | 4 | G | O6 | A:895 | 2.92 | 1.0 | 36.6 |  | 10.10 | 5 | G | O6 | A:894 | 2.94 | 1.0 | 41.1 |  | 9.08 | 6 | Water 3 | | A:3066 | 3.43 | 1.0 | 42.0 | 2.76 [G:N7]A:895 | 6.19 | 7 | U | O4 | A:905 | 3.44 | 1.0 | 38.2 |  | 8.36 | 8 | Water 4 | | A:4640 | 3.47 | 1.0 | 42.5 |  | 3.32 | 9 | Water 5 | | A:2269 | 3.52 | 1.0 | 39.2 | 2.58 [G:O6]A:903 | 4.21 | 10 | Water 6 | | A:2516 | 3.98 | 1.0 | 37.0 | 2.64 [U:O4]A:904 | 7.54 | 11 | G | O6 | A:903 | 4.18 | 1.0 | 36.4 |  | 9.30 | 12 | G | N7 | A:894 | 4.44 | 1.0 | 38.7 |  | 7.09 | 13 | G | N7 | A:895 | 4.45 | 1.0 | 36.0 |  | 8.50 | 14 | Water 7 | | A:2410 | 4.60 | 1.0 | 44.5 |  | 4.24 | 15 | Water 8 | | A:4683 | 4.75 | 1.0 | 48.9 |  | 2.39 |

  
• • **2.89 ± 0.28: the average coodination distance of the 1^st^ six ligands is K^+^ compatible**  
**6: Number of potential Water contacts**   
**3 : Number of second shell contacts involving water or ions**   
**50%: Percentage of satisfied second shell Water contacts**   
• Contact code: → No contacts ←   
• Contact(s) in the exclusion zone: /O4/O6/O6/O_w_/O_w_/  
→ Consider K in place of MG (3 valid contact(s) for K vs. 0 for MG ←)  
Go to ToC

### MG:A:1610

|  | Res. type | Atom | Residue | Distance | Occupancy | B-factor | Closest atom(s) to a 1^st^ shell water | EM\_dens. |
| --- | --- | --- | --- | --- | --- | --- | --- | --- |
| **Ion** | | | |  | 1.0 | 63.7 |  | 4.16 |
 1 | Water 1 | | A:4686 | 1.81 | 1.0 | 49.8 |  | 2.29 | 2 | Water 2 | | A:2284 | 1.89 | 1.0 | 52.8 | 2.59 [C:OP1]A:797 | 4.04 | 3 | Water 3 | | A:4599 | 1.90 | 1.0 | 56.4 |  | 2.49 | 4 | Water 4 | | A:3905 | 1.98 | 1.0 | 51.0 | 2.94 [C:OP1]A:797 | 2.78 | 5 | Water 5 | | A:3111 | 2.07 | 1.0 | 47.6 | 2.77 [U:OP1]A:798 | 4.11 | 6 | Water 6 | | A:3385 | 2.08 | 1.0 | 53.0 | 2.82 [U:OP2]A:798 | 3.22 | 7 | C | OP1 | A:797 | 3.80 | 1.0 | 38.8 |  | 8.29 | 8 | U | OP1 | A:692 | 4.00 | 1.0 | 52.6 |  | 3.41 | 9 | U | OP1 | A:798 | 4.20 | 1.0 | 45.8 |  | 5.45 | 10 | U | OP2 | A:798 | 4.28 | 1.0 | 47.0 |  | 4.39 | 11 | Water 7 | | A:3381 | 4.37 | 1.0 | 55.2 |  | 1.44 | 12 | Water 8 | | K:305 | 4.43 | 1.0 | 43.2 |  | 4.36 | 13 | Water 9 | | U:107 | 4.54 | 1.0 | 51.6 |  | 2.72 | 14 | Water 10 | | A:2389 | 4.60 | 1.0 | 57.3 | 2.62 [U:OP1]A:692 | 1.62 | 15 | Water 11 | | A:2948 | 4.74 | 1.0 | 51.3 |  | 2.01 |

  
**1^st^ shell coordination angles and distances (*trans* in "blue"; *cys* in "green"; deviations in "orange"):**   
• 1-2(W/W): 171º; 1-3(W/W): *81º*; 1-4(W/W): 94º; 1-5(W/W): *84º*; 1-6(W/W): 91º; 2-3(W/W): 91º; 2-4(W/W): *85º*; 2-5(W/W): *100º*; 2-6(W/W): *96º*; 3-4(W/W): *100º*; 3-5(W/W): *95º*; 3-6(W/W): *170º*; 4-5(W/W): *164º*; 4-6(W/W): *75º*; 5-6(W/W): 90º;   
• 1-2(W/W): *3.7Å*; 1-3(W/W): 2.4Å; 1-4(W/W): 2.8Å; 1-5(W/W): 2.6Å; 1-6(W/W): 2.8Å; 2-3(W/W): 2.7Å; 2-4(W/W): 2.6Å; 2-5(W/W): 3.0Å; 2-6(W/W): 3.0Å; 3-4(W/W): 3.0Å; 3-5(W/W): 2.9Å; 3-6(W/W): 4.0Å; 4-5(W/W): 4.0Å; 4-6(W/W): 2.5Å; 5-6(W/W): 2.9Å;   
  
**1.95 ± 0.10: the average coodination distance of the 1^st^ six ligands is Mg^2+^ compatible**  
**12: Number of potential Water contacts**   
**4 : Number of second shell contacts involving water or ions**   
**33%: Percentage of satisfied second shell Water contacts**   
• Contact(s): /O_w_/O_w_/O_w_/  
• Contact code: 3O_w_  
• Contact(s) in the exclusion zone: /O_w_/O_w_/O_w_/  
→ Missing octahedral coordination for MG ←  
→ Slightly distorded coord. octahedron (7.3º) ←  
Go to ToC

### MG:A:1611

|  | Res. type | Atom | Residue | Distance | Occupancy | B-factor | Closest atom(s) to a 1^st^ shell water | EM\_dens. |
| --- | --- | --- | --- | --- | --- | --- | --- | --- |
| **Ion** | | | |  | 1.0 | 61.0 |  | 2.46 |
 1 | Water 1 | | X:206 | 1.91 | 1.0 | 53.5 | 2.73 [U:O2']X:17 | 2.17 | 2 | Water 2 | | A:3423 | 2.05 | 1.0 | 47.5 | 2.82 [G:N7]A:1497 | 3.39 | 3 | Water 3 | | A:4553 | 2.58 | 1.0 | 45.9 |  | 3.20 | 4 | Water 4 | | A:2826 | 2.74 | 1.0 | 45.3 | 2.71 [C:OP2]A:1496 | 2.47 | 5 | Water 5 | | A:3781 | 3.84 | 1.0 | 49.4 | 2.91 [C:OP2]A:1496 | 2.99 | 6 | UR3 | O4 | A:1498 | 4.16 | 1.0 | 42.5 |  | 8.02 | 7 | Water 6 | | A:2248 | 4.24 | 1.0 | 45.3 |  | 3.31 | 8 | Water 7 | | A:4513 | 4.25 | 1.0 | 50.6 |  | 3.60 | 9 | U | O2' | X:17 | 4.31 | 1.0 | 43.8 |  | 5.52 | 10 | G | N7 | A:1497 | 4.44 | 1.0 | 37.1 |  | 8.36 | 11 | Water 8 | | X:204 | 4.50 | 1.0 | 50.1 |  | 2.02 | 12 | Water 9 | | X:211 | 4.61 | 1.0 | 50.7 |  | 3.03 | 13 | C | OP2 | A:1496 | 4.67 | 1.0 | 45.8 |  | 4.14 | 14 | Water 10 | | X:218 | 4.80 | 1.0 | 48.9 |  | 1.82 |

  
**1^st^ shell coordination angles and distances (*trans* in "blue"; *cys* in "green"; deviations in "orange"):**   
• 1-2(W/W): 90º;   
• 1-2(W/W): 2.8Å;   
  
**2.88 ± 0.85: the average coodination distance of the 1^st^ six ligands is K^+^ compatible**  
**10: Number of potential Water contacts**   
**4 : Number of second shell contacts involving water or ions**   
**40%: Percentage of satisfied second shell Water contacts**   
• Contact(s): /O_w_/O_w_/  
• Contact code: 2O_w_  
• Contact(s) in the exclusion zone: /O_w_/O_w_/  
→ Missing octahedral coordination for MG ←  
→2Fo-Fc density between 1.0 and 4.0 !   
Go to ToC

### MG:A:1612

|  | Res. type | Atom | Residue | Distance | Occupancy | B-factor | Closest atom(s) to a 1^st^ shell water | EM\_dens. |
| --- | --- | --- | --- | --- | --- | --- | --- | --- |
| **Ion** | | | |  | 1.0 | 52.7 |  | 5.33 |
 1 | G | N7 | A:362 | 1.99 | 1.0 | 48.3 |  | 4.93 | 2 | Water 1 | | A:4277 | 2.00 | 1.0 | 57.6 |  | 2.45 | 3 | Water 2 | | A:4351 | 2.01 | 1.0 | 55.4 |  | 2.25 | 4 | Water 3 | | A:2011 | 2.47 | 1.0 | 55.1 | 2.47 [G:N7]A:361 | 2.26 | 5 | G | O6 | A:362 | 3.51 | 1.0 | 53.7 |  | 3.26 | 6 | Water 4 | | A:2393 | 3.86 | 1.0 | 53.3 |  | 1.81 | 7 | G | N7 | A:361 | 4.01 | 1.0 | 48.8 |  | 4.76 | 8 | Water 5 | | A:1986 | 4.28 | 1.0 | 46.2 |  | 3.58 | 9 | Water 6 | | A:3547 | 4.47 | 1.0 | 58.5 | 2.85 [G:O6]A:362 | 1.45 | 10 | G | OP2 | A:361 | 4.78 | 1.0 | 54.4 |  | 3.04 |

  
**1^st^ shell coordination angles and distances (*trans* in "blue"; *cys* in "green"; deviations in "orange"):**   
• 1-2(N7/W): *100º*; 1-3(N7/W): *101º*; 2-3(W/W): 93º;   
• 1-2(N7/W): 3.1Å; 1-3(N7/W): 3.1Å; 2-3(W/W): 2.9Å;   
  
**2.64 ± 0.76: the average coodination distance of the 1^st^ six ligands is K^+^ compatible**  
**8: Number of potential Water contacts**   
**1 : Number of second shell contacts involving water or ions**   
**12%: Percentage of satisfied second shell Water contacts**   
• Contact(s): /N7/O_w_/O_w_/  
• Contact code: N_b_2O_w_  
• Contact(s) in the exclusion zone: /O_w_/  
→ Missing octahedral coordination for MG ←  
Go to ToC

### MG:A:1613

|  | Res. type | Atom | Residue | Distance | Occupancy | B-factor | Closest atom(s) to a 1^st^ shell water | EM\_dens. |
| --- | --- | --- | --- | --- | --- | --- | --- | --- |
| **Ion** | | | |  | 1.0 | 45.0 |  | 11.34 |
 1 | Water 1 | | A:3824 | 1.89 | 1.0 | 37.1 | 2.92 [U:OP1]A:1527 | 6.29 | 2 | Water 2 | | A:2953 | 1.90 | 1.0 | 36.6 | 2.74 [G:OP1]A:730 // 2.86 [G:OP2]A:731 | 8.49 | 3 | Water 3 | | A:4591 | 1.99 | 1.0 | 38.8 |  | 5.84 | 4 | Water 4 | | A:3064 | 2.12 | 1.0 | 41.2 | 2.76 [G:O2']A:1526 | 8.70 | 5 | Water 5 | | A:2684 | 2.24 | 1.0 | 37.8 | 2.68 [G:OP1]A:731 | 8.85 | 6 | Water 6 | | A:3320 | 2.41 | 1.0 | 39.5 | 2.81 [A:N1]A:814 | 7.81 | 7 | Water 7 | | A:4664 | 4.02 | 1.0 | 47.4 |  | 3.59 | 8 | Water 8 | | A:3679 | 4.03 | 1.0 | 47.4 |  | 2.67 | 9 | G | OP1 | A:730 | 4.05 | 1.0 | 35.4 |  | 10.03 | 10 | Water 9 | | A:2704 | 4.09 | 1.0 | 42.1 | 2.69 [G:OP1]A:730 | 3.67 | 11 | G | OP2 | A:731 | 4.10 | 1.0 | 40.2 |  | 9.38 | 12 | Water 10 | | U:117 | 4.20 | 1.0 | 51.0 |  | 3.57 | 13 | G | O3' | A:1526 | 4.27 | 1.0 | 39.3 |  | 9.58 | 14 | Water 11 | | A:2390 | 4.40 | 1.0 | 50.0 | 2.62 [G:OP1]A:730 | 2.58 | 15 | U | OP1 | A:1527 | 4.40 | 1.0 | 36.8 |  | 8.53 | 16 | A | OP2 | A:816 | 4.42 | 1.0 | 28.2 |  | 9.96 | 17 | G | OP1 | A:731 | 4.45 | 1.0 | 38.4 |  | 9.93 | 18 | A | N1 | A:814 | 4.54 | 1.0 | 28.5 |  | 12.00 | 19 | G | O3' | A:730 | 4.60 | 1.0 | 36.6 |  | 10.09 | 20 | G | O2' | A:1526 | 4.69 | 1.0 | 31.6 |  | 12.21 | 21 | Water 12 | | A:4063 | 4.71 | 1.0 | 51.2 |  | 1.97 |

  
**1^st^ shell coordination angles and distances (*trans* in "blue"; *cys* in "green"; deviations in "orange"):**   
• 1-2(W/W): *84º*; 1-3(W/W): *98º*; 1-4(W/W): 93º; 1-5(W/W): 92º; 2-3(W/W): 93º; 2-4(W/W): 170º; 2-5(W/W): 92º; 3-4(W/W): *97º*; 3-5(W/W): *169º*; 4-5(W/W): *78º*;   
• 1-2(W/W): 2.5Å; 1-3(W/W): 2.9Å; 1-4(W/W): 2.9Å; 1-5(W/W): 3.0Å; 2-3(W/W): 2.8Å; 2-4(W/W): 4.0Å; 2-5(W/W): 3.0Å; 3-4(W/W): 3.1Å; 3-5(W/W): *4.2Å*; 4-5(W/W): 2.8Å;   
  
**2.09 ± 0.19: the average coodination distance of the 1^st^ six ligands is Mg^2+^ compatible**  
**12: Number of potential Water contacts**   
**6 : Number of second shell contacts involving water or ions**   
**50%: Percentage of satisfied second shell Water contacts**   
• Contact(s): /O_w_/O_w_/O_w_/  
• Contact code: 3O_w_  
• Contact(s) in the exclusion zone: /O_w_/O_w_/O_w_/  
→ Missing octahedral coordination for MG ←  
→ Slightly distorded coord. octahedron (6.3º) ←  
Go to ToC

### MG:A:1614

|  | Res. type | Atom | Residue | Distance | Occupancy | B-factor | Closest atom(s) to a 1^st^ shell water | EM\_dens. |
| --- | --- | --- | --- | --- | --- | --- | --- | --- |
| **Ion** | | | |  | 1.0 | 47.1 |  | 8.95 |
 1 | Water 1 | | A:3346 | 1.89 | 1.0 | 45.3 | 2.81 [U:O4]A:798 | 4.34 | 2 | Water 2 | | A:2955 | 1.95 | 1.0 | 40.5 | 2.74 [G:O6]A:800 | 7.03 | 3 | Water 3 | | A:3949 | 2.03 | 1.0 | 45.2 | 2.95 [G:N7]A:799 | 5.36 | 4 | Water 4 | | A:4668 | 2.09 | 1.0 | 44.0 |  | 5.15 | 5 | Water 5 | | K:321 | 2.10 | 1.0 | 41.9 |  | 5.36 | 6 | G | O6 | A:800 | 4.03 | 1.0 | 38.0 |  | 8.56 | 7 | U | O4 | A:798 | 4.09 | 1.0 | 41.2 |  | 6.44 | 8 | A | OP2 | A:781 | 4.14 | 1.0 | 40.1 |  | 6.58 | 9 | Water 6 | | A:2369 | 4.20 | 1.0 | 46.7 |  | 3.74 | 10 | G | O6 | A:799 | 4.28 | 1.0 | 40.5 |  | 8.24 | 11 | Water 7 | | A:3077 | 4.31 | 1.0 | 41.6 |  | 5.61 | 12 | Water 8 | | A:3577 | 4.39 | 1.0 | 44.0 |  | 4.41 | 13 | Water 9 | | A:2709 | 4.51 | 1.0 | 43.6 |  | 5.99 | 14 | Water 10 | | A:4053 | 4.54 | 1.0 | 40.7 |  | 6.13 | 15 | G | N7 | A:799 | 4.67 | 1.0 | 37.2 |  | 7.30 |

  
**1^st^ shell coordination angles and distances (*trans* in "blue"; *cys* in "green"; deviations in "orange"):**   
• 1-2(W/W): 173º; 1-3(W/W): 89º; 1-4(W/W): 94º; 1-5(W/W): *80º*; 2-3(W/W): 93º; 2-4(W/W): 93º; 2-5(W/W): *99º*; 3-4(W/W): 87º; 3-5(W/W): *167º*; 4-5(W/W): 88º;   
• 1-2(W/W): 3.8Å; 1-3(W/W): 2.7Å; 1-4(W/W): 2.9Å; 1-5(W/W): 2.6Å; 2-3(W/W): 2.9Å; 2-4(W/W): 2.9Å; 2-5(W/W): 3.1Å; 3-4(W/W): 2.8Å; 3-5(W/W): 4.1Å; 4-5(W/W): 2.9Å;   
  
**2.35 ± 0.76: the average coodination distance of the 1^st^ six ligands is Na^+^** compatible  
**10: Number of potential Water contacts**   
**3 : Number of second shell contacts involving water or ions**   
**30%: Percentage of satisfied second shell Water contacts**   
• Contact(s): /O_w_/O_w_/O_w_/O_w_/  
• Contact code: 4O_w_  
• Contact(s) in the exclusion zone: /O_w_/  
→ Missing octahedral coordination for MG ←  
→ Slightly distorded coord. octahedron (5.6º) ←  
Go to ToC

### MG:A:1615

|  | Res. type | Atom | Residue | Distance | Occupancy | B-factor | Closest atom(s) to a 1^st^ shell water | EM\_dens. |
| --- | --- | --- | --- | --- | --- | --- | --- | --- |
| **Ion** | | | |  | 1.0 | 56.7 |  | 4.83 |
 1 | Water 1 | | A:3981 | 1.91 | 1.0 | 56.0 |  | 1.98 | 2 | Water 2 | | A:3420 | 1.94 | 1.0 | 55.1 |  | 3.67 | 3 | Water 3 | | A:4669 | 2.16 | 1.0 | 59.9 |  | 2.21 | 4 | G | N7 | A:324 | 2.17 | 1.0 | 44.8 |  | 5.72 | 5 | G | O6 | A:324 | 3.50 | 1.0 | 50.4 |  | 3.96 | 6 | Water 4 | | A:3387 | 4.05 | 1.0 | 52.4 | 2.82 [U:OP2]A:323 | 2.41 | 7 | Water 5 | | A:2335 | 4.05 | 1.0 | 55.6 |  | 2.29 | 8 | U | OP2 | A:323 | 4.29 | 1.0 | 53.3 |  | 4.30 | 9 | Water 6 | | A:1856 | 4.59 | 1.0 | 58.6 | 2.34 [G:OP2]A:324 | 1.90 | 10 | G | OP2 | A:324 | 4.71 | 1.0 | 48.9 |  | 5.32 |

  
**1^st^ shell coordination angles and distances (*trans* in "blue"; *cys* in "green"; deviations in "orange"):**   
• 1-2(W/W): 88º; 1-3(W/W): *82º*; 1-4(W/N7): 93º; 2-3(W/W): 93º; 2-4(W/N7): 86º; 3-4(W/N7): 174º;   
• 1-2(W/W): 2.7Å; 1-3(W/W): 2.7Å; 1-4(W/N7): 3.0Å; 2-3(W/W): 3.0Å; 2-4(W/N7): 2.8Å; 3-4(W/N7): *4.3Å*;   
  
**2.62 ± 0.83: the average coodination distance of the 1^st^ six ligands is K^+^ compatible**  
**8: Number of potential Water contacts**   
**1 : Number of second shell contacts involving water or ions**   
**12%: Percentage of satisfied second shell Water contacts**   
• Contact(s): /N7/O_w_/O_w_/O_w_/  
• Contact code: N_b_3O_w_  
→ Missing octahedral coordination for MG ←  
Ooctahedral coordination (4.1º)   
Go to ToC

### MG:A:1616

|  | Res. type | Atom | Residue | Distance | Occupancy | B-factor | Closest atom(s) to a 1^st^ shell water | EM\_dens. |
| --- | --- | --- | --- | --- | --- | --- | --- | --- |
| **Ion** | | | |  | 1.0 | 77.8 |  | 1.96 |
 1 | Water 1 | | A:4643 | 2.07 | 1.0 | 67.9 |  | 2.31 | 2 | Water 2 | | A:2266 | 2.09 | 1.0 | 81.5 | 2.58 [G:OP2]A:450 | 1.91 | 3 | Water 3 | | A:3533 | 2.27 | 1.0 | 76.0 | 2.85 [G:N7]A:450 | 1.60 | 4 | G | OP2 | A:450 | 3.81 | 1.0 | 76.4 |  | 2.73 | 5 | G | N7 | A:481 | 3.92 | 1.0 | 71.9 |  | 3.32 | 6 | A | O2' | A:451 | 3.97 | 1.0 | 72.0 |  | 2.15 | 7 | G | O6 | A:481 | 3.99 | 1.0 | 75.5 |  | 2.11 | 8 | G | N7 | A:450 | 4.16 | 1.0 | 75.0 |  | 3.31 | 9 | G | OP2 | A:449 | 4.54 | 1.0 | 82.7 |  | 1.81 | 10 | Water 4 | | A:2247 | 4.62 | 1.0 | 78.5 |  | 2.01 |

  
**1^st^ shell coordination angles and distances (*trans* in "blue"; *cys* in "green"; deviations in "orange"):**   
• 1-2(W/W): 91º; 1-3(W/W): *77º*; 2-3(W/W): *96º*;   
• 1-2(W/W): 3.0Å; 1-3(W/W): 2.7Å; 2-3(W/W): *3.2Å*;   
  
**3.02 ± 0.88: the average coodination distance of the 1^st^ six ligands is K^+^ compatible**  
**6: Number of potential Water contacts**   
**2 : Number of second shell contacts involving water or ions**   
**33%: Percentage of satisfied second shell Water contacts**   
• Contact(s): /O_w_/O_w_/O_w_/  
• Contact code: 3O_w_  
→ Missing octahedral coordination for MG ←  
→2Fo-Fc density between 1.0 and 4.0 !   
Go to ToC

### MG:A:1617

|  | Res. type | Atom | Residue | Distance | Occupancy | B-factor | Closest atom(s) to a 1^st^ shell water | EM\_dens. |
| --- | --- | --- | --- | --- | --- | --- | --- | --- |
| **Ion** | | | |  | 1.0 | 70.9 |  | 2.21 |
 1 | Water 1 | | A:2808 | 1.90 | 1.0 | 70.3 | 2.71 [G:N7]A:617 | 2.25 | 2 | Water 2 | | A:1973 | 2.50 | 1.0 | 70.8 | 2.46 [A:OP2]A:621 | 1.58 | 3 | Water 3 | | A:1993 | 4.05 | 1.0 | 71.5 |  | 2.40 | 4 | G | N7 | A:617 | 4.22 | 1.0 | 68.3 |  | 2.84 | 5 | C | OP1 | A:620 | 4.30 | 1.0 | 68.4 |  | 3.03 | 6 | A | OP1 | A:621 | 4.30 | 1.0 | 60.5 |  | 3.46 | 7 | Water 4 | | A:1845 | 4.50 | 1.0 | 72.3 | 2.65 [C:OP1]A:620 | 1.48 | 8 | G | OP2 | A:617 | 4.55 | 1.0 | 68.6 |  | 2.02 | 9 | A | OP2 | A:621 | 4.57 | 1.0 | 67.8 |  | 1.52 |

  
• • **3.55 ± 0.97: the average coodination distance of the 1^st^ six ligands is K^+^ compatible**  
**6: Number of potential Water contacts**   
**2 : Number of second shell contacts involving water or ions**   
**33%: Percentage of satisfied second shell Water contacts**   
• Contact(s): /O_w_/  
• Contact code: O_w_  
• Contact(s) in the exclusion zone: /O_w_/  
→ Missing octahedral coordination for MG ←  
→2Fo-Fc density between 1.0 and 4.0 !   
Go to ToC

### MG:A:1618

|  | Res. type | Atom | Residue | Distance | Occupancy | B-factor | Closest atom(s) to a 1^st^ shell water | EM\_dens. |
| --- | --- | --- | --- | --- | --- | --- | --- | --- |
| **Ion** | | | |  | 1.0 | 61.7 |  | 3.98 |
 1 | Water 1 | | A:1981 | 1.81 | 1.0 | 58.3 | 2.46 [G:OP2]A:597 // 2.89 [U:O4]A:598 | 2.36 | 2 | Water 2 | | A:2730 | 1.98 | 1.0 | 60.8 | 2.69 [G:N7]A:597 // 2.85 [U:O4]A:598 | 2.75 | 3 | U | O4 | A:598 | 2.50 | 1.0 | 55.2 |  | 3.91 | 4 | G | OP2 | A:597 | 2.61 | 1.0 | 62.0 |  | 2.42 | 5 | A | OP2 | A:596 | 3.04 | 1.0 | 60.9 |  | 2.54 | 6 | A | O3' | A:595 | 3.35 | 1.0 | 65.1 |  | 3.30 | 7 | A | O2' | A:595 | 3.38 | 1.0 | 63.9 |  | 2.82 | 8 | Water 3 | | A:4346 | 3.50 | 1.0 | 65.5 |  | 1.73 | 9 | A | O5' | A:596 | 3.50 | 1.0 | 64.6 |  | 2.48 | 10 | G | N7 | A:597 | 4.29 | 1.0 | 54.7 |  | 4.64 | 11 | Water 4 | | A:3081 | 4.32 | 1.0 | 59.3 |  | 3.10 | 12 | A | O3' | A:596 | 4.71 | 1.0 | 63.0 |  | 2.39 |

  
**1^st^ shell coordination angles and distances (*trans* in "blue"; *cys* in "green"; deviations in "orange"):**   
• 1-2(W/W): *147º*;   
• 1-2(W/W): *3.6Å*;   
  
**2.55 ± 0.54: the average coodination distance of the 1^st^ six ligands is Na^+^** compatible  
**4: Number of potential Water contacts**   
**4 : Number of second shell contacts involving water or ions**   
**100%: Percentage of satisfied second shell Water contacts**   
• Contact(s): /O_w_/  
• Contact code: O_w_  
• Contact(s) in the exclusion zone: /O2'/O3'/O4/O_w_/OP2/OP2/  
→ Consider K in place of MG (2 valid contact(s) for K vs. 0 for MG ←)  
→ Missing octahedral coordination for MG ←  
→2Fo-Fc density between 1.0 and 4.0 !   
Go to ToC

### MG:A:1619

|  | Res. type | Atom | Residue | Distance | Occupancy | B-factor | Closest atom(s) to a 1^st^ shell water | EM\_dens. |
| --- | --- | --- | --- | --- | --- | --- | --- | --- |
| **Ion** | | | |  | 1.0 | 55.9 |  | 6.31 |
 1 | Water 1 | | A:2906 | 1.83 | 1.0 | 36.4 | 2.85 [C:O2']A:1521 | 7.46 | 2 | Water 2 | | A:2740 | 1.88 | 1.0 | 43.2 | 2.69 [C:OP2]A:783 | 6.00 | 3 | Water 3 | | A:4207 | 2.08 | 1.0 | 45.7 |  | 2.66 | 4 | A | OP1 | A:794 | 2.12 | 1.0 | 43.5 |  | 4.68 | 5 | A | OP1 | A:782 | 2.27 | 1.0 | 34.9 |  | 8.76 | 6 | Water 4 | | A:3603 | 2.27 | 1.0 | 42.4 | 2.86 [C:O2]A:1521 | 5.20 | 7 | Water 5 | | A:4562 | 2.60 | 1.0 | 49.0 |  | 2.43 | 8 | A | OP2 | A:794 | 3.81 | 1.0 | 48.7 |  | 3.47 | 9 | A | OP2 | A:782 | 3.82 | 1.0 | 38.2 |  | 7.57 | 10 | Water 6 | | A:2831 | 3.98 | 1.0 | 48.8 | 2.92 [U:O3']A:793 | 3.98 | 11 | Water 7 | | A:3817 | 4.04 | 1.0 | 53.3 | 2.92 [A:OP2]A:794 | 1.95 | 12 | C | OP2 | A:783 | 4.15 | 1.0 | 42.3 |  | 7.55 | 13 | A | O5' | A:782 | 4.18 | 1.0 | 37.6 |  | 8.68 | 14 | Water 8 | | A:3951 | 4.33 | 1.0 | 44.6 |  | 4.26 | 15 | C | O2 | A:1521 | 4.34 | 1.0 | 33.5 |  | 9.45 | 16 | U | O3' | A:793 | 4.40 | 1.0 | 51.5 |  | 4.20 | 17 | A | O5' | A:794 | 4.40 | 1.0 | 48.3 |  | 5.06 | 18 | U | O4' | A:1522 | 4.48 | 1.0 | 34.6 |  | 10.98 | 19 | C | O2' | A:1521 | 4.57 | 1.0 | 35.8 |  | 9.89 | 20 | A | O3' | A:781 | 4.69 | 1.0 | 33.9 |  | 12.42 |

  
**1^st^ shell coordination angles and distances (*trans* in "blue"; *cys* in "green"; deviations in "orange"):**   
• 1-2(W/W): *146º*; 1-3(W/W): *120º*; 1-4(W/OP1): 87º; 1-5(W/OP1): 93º; 1-6(W/W): *75º*; 2-3(W/W): 93º; 2-4(W/OP1): 91º; 2-5(W/OP1): *85º*; 2-6(W/W): *71º*; 3-4(W/OP1): *98º*; 3-5(W/OP1): 90º; 3-6(W/W): *165º*; 4-5(OP1/OP1): 171º; 4-6(OP1/W): *83º*; 5-6(OP1/W): 88º;   
• 1-2(W/W): *3.6Å*; 1-3(W/W): *3.4Å*; 1-4(W/OP1): 2.7Å; 1-5(W/OP1): 3.0Å; 1-6(W/W): 2.5Å; 2-3(W/W): 2.9Å; 2-4(W/OP1): 2.9Å; 2-5(W/OP1): 2.8Å; 2-6(W/W): 2.4Å; 3-4(W/OP1): 3.2Å; 3-5(W/OP1): 3.1Å; 3-6(W/W): *4.3Å*; 4-5(OP1/OP1): *4.4Å*; 4-6(OP1/W): 2.9Å; 5-6(OP1/W): 3.2Å;   
  
**2.07 ± 0.17: the average coodination distance of the 1^st^ six ligands is Mg^2+^ compatible**  
**8: Number of potential Water contacts**   
**3 : Number of second shell contacts involving water or ions**   
**38%: Percentage of satisfied second shell Water contacts**   
• Contact(s): /O_w_/O_w_/OP1/OP1/  
• Contact code: *cis-*2O_ph_2O_w_  
• Contact(s) in the exclusion zone: /O_w_/O_w_/O_w_/  
→ Missing octahedral coordination for MG ←  
→→→ Distorded coord. octahedron (10.3º) ←←←  
Go to ToC

### MG:A:1620

|  | Res. type | Atom | Residue | Distance | Occupancy | B-factor | Closest atom(s) to a 1^st^ shell water | EM\_dens. |
| --- | --- | --- | --- | --- | --- | --- | --- | --- |
| **Ion** | | | |  | 1.0 | 43.8 |  | 11.33 |
 1 | Water 1 | | A:3086 | 1.99 | 1.0 | 42.3 |  | 4.99 | 2 | Water 2 | | A:4017 | 2.03 | 1.0 | 42.2 |  | 6.17 | 3 | G | OP1 | A:21 | 2.10 | 1.0 | 36.2 |  | 7.63 | 4 | Water 3 | | A:3253 | 2.25 | 1.0 | 39.9 | 2.79 [A:N1]A:573 | 9.58 | 5 | Water 4 | | A:4282 | 2.25 | 1.0 | 43.6 |  | 6.03 | 6 | Water 5 | | A:3847 | 3.82 | 1.0 | 45.2 |  | 5.57 | 7 | G | OP2 | A:21 | 3.95 | 1.0 | 41.9 |  | 7.00 | 8 | U | O3' | A:20 | 4.15 | 1.0 | 39.6 |  | 8.39 | 9 | Water 6 | | A:3745 | 4.16 | 1.0 | 47.2 |  | 4.67 | 10 | A | N1 | A:573 | 4.21 | 1.0 | 32.7 |  | 10.42 | 11 | G | O2' | A:567 | 4.24 | 1.0 | 42.1 |  | 6.02 | 12 | Water 7 | | A:4618 | 4.38 | 1.0 | 47.9 |  | 2.97 | 13 | G | N3 | A:567 | 4.49 | 1.0 | 35.0 |  | 9.94 | 14 | Water 8 | | A:1812 | 4.49 | 1.0 | 42.7 |  | 3.60 | 15 | G | O5' | A:21 | 4.56 | 1.0 | 38.2 |  | 7.56 | 16 | G | O2' | A:568 | 4.64 | 1.0 | 36.3 |  | 9.52 | 17 | Water 9 | | A:3886 | 4.79 | 1.0 | 48.4 | 2.94 [G:O2']A:567 | 2.99 |

  
**1^st^ shell coordination angles and distances (*trans* in "blue"; *cys* in "green"; deviations in "orange"):**   
• 1-2(W/W): 172º; 1-3(W/OP1): *85º*; 1-4(W/W): *95º*; 1-5(W/W): *85º*; 2-3(W/OP1): 92º; 2-4(W/W): 92º; 2-5(W/W): 88º; 3-4(OP1/W): 89º; 3-5(OP1/W): 89º; 4-5(W/W): 178º;   
• 1-2(W/W): 4.0Å; 1-3(W/OP1): 2.8Å; 1-4(W/W): 3.1Å; 1-5(W/W): 2.9Å; 2-3(W/OP1): 3.0Å; 2-4(W/W): 3.1Å; 2-5(W/W): 3.0Å; 3-4(OP1/W): 3.0Å; 3-5(OP1/W): 3.1Å; 4-5(W/W): *4.5Å*;   
  
**2.41 ± 0.64: the average coodination distance of the 1^st^ six ligands is Na^+^** compatible  
**10: Number of potential Water contacts**   
**1 : Number of second shell contacts involving water or ions**   
**10%: Percentage of satisfied second shell Water contacts**   
• Contact(s): /O_w_/O_w_/O_w_/O_w_/OP1/  
• Contact code: O_ph_4O_w_  
→ Missing octahedral coordination for MG ←  
Ooctahedral coordination (3.2º)   
Go to ToC

### MG:A:1621

|  | Res. type | Atom | Residue | Distance | Occupancy | B-factor | Closest atom(s) to a 1^st^ shell water | EM\_dens. |
| --- | --- | --- | --- | --- | --- | --- | --- | --- |
| **Ion** | | | |  | 1.0 | 72.6 |  | 1.86 |
 1 | Water 1 | | A:2990 | 2.46 | 1.0 | 65.7 | 2.74 [U:OP2]A:610 | 2.40 | 2 | Water 2 | | A:4480 | 2.49 | 1.0 | 67.6 |  | 1.61 | 3 | Water 3 | | A:3141 | 2.94 | 1.0 | 69.8 | 2.77 [G:O6]A:626 | 1.67 | 4 | Water 4 | | A:4133 | 3.02 | 1.0 | 62.7 |  | 1.77 | 5 | G | O6 | A:627 | 3.03 | 1.0 | 70.6 |  | 2.91 | 6 | G | N7 | A:627 | 3.81 | 1.0 | 69.6 |  | 2.65 | 7 | Water 5 | | A:3778 | 3.94 | 1.0 | 61.9 |  | 2.21 | 8 | G | O6 | A:628 | 4.28 | 1.0 | 72.9 |  | 2.37 | 9 | G | O6 | A:626 | 4.38 | 1.0 | 67.3 |  | 2.62 | 10 | G | N7 | A:626 | 4.42 | 1.0 | 67.3 |  | 2.54 | 11 | U | OP2 | A:610 | 4.79 | 1.0 | 56.0 |  | 4.88 |

  
• • **2.96 ± 0.45: the average coodination distance of the 1^st^ six ligands is K^+^ compatible**  
**8: Number of potential Water contacts**   
**2 : Number of second shell contacts involving water or ions**   
**25%: Percentage of satisfied second shell Water contacts**   
• Contact code: → No contacts ←   
• Contact(s) in the exclusion zone: /O6/O_w_/O_w_/O_w_/O_w_/  
→ Consider K in place of MG (1 valid contact(s) for K vs. 0 for MG ←)  
→2Fo-Fc density between 1.0 and 4.0 !   
Go to ToC

### MG:A:1622

|  | Res. type | Atom | Residue | Distance | Occupancy | B-factor | Closest atom(s) to a 1^st^ shell water | EM\_dens. |
| --- | --- | --- | --- | --- | --- | --- | --- | --- |
| **Ion** | | | |  | 1.0 | 67.9 |  | 2.04 |
 1 | Water 1 | | O:128 | 2.00 | 1.0 | 62.2 |  | 2.22 | 2 | Water 2 | | A:4560 | 2.25 | 1.0 | 63.2 |  | 1.81 | 3 | Water 3 | | A:4062 | 2.36 | 1.0 | 61.5 | 2.99 [C:OP1]A:658 | 1.70 | 4 | Water 4 | | A:4152 | 2.53 | 1.0 | 58.0 |  | 1.67 | 5 | Water 5 | | A:4190 | 4.23 | 1.0 | 57.7 |  | 2.52 | 6 | Water 6 | | A:2131 | 4.24 | 1.0 | 50.3 |  | 2.07 | 7 | U | OP1 | A:657 | 4.33 | 1.0 | 62.8 |  | 3.05 | 8 | G | OP1 | A:742 | 4.44 | 1.0 | 43.5 |  | 4.58 | 9 | C | OP1 | A:658 | 4.45 | 1.0 | 62.1 |  | 2.83 | 10 | C | OP2 | A:658 | 4.50 | 1.0 | 63.2 |  | 3.09 | 11 | GLN | OE1 | O:35 | 4.58 | 1.0 | 59.1 |  | 1.34 |

  
**1^st^ shell coordination angles and distances (*trans* in "blue"; *cys* in "green"; deviations in "orange"):**   
• 1-2(W/W): *77º*;   
• 1-2(W/W): 2.6Å;   
  
**2.93 ± 0.93: the average coodination distance of the 1^st^ six ligands is K^+^ compatible**  
**12: Number of potential Water contacts**   
**1 : Number of second shell contacts involving water or ions**   
**8%: Percentage of satisfied second shell Water contacts**   
• Contact(s): /O_w_/O_w_/  
• Contact code: 2O_w_  
• Contact(s) in the exclusion zone: /O_w_/O_w_/  
→ Missing octahedral coordination for MG ←  
→2Fo-Fc density between 1.0 and 4.0 !   
Go to ToC

### MG:A:1623

|  | Res. type | Atom | Residue | Distance | Occupancy | B-factor | Closest atom(s) to a 1^st^ shell water | EM\_dens. |
| --- | --- | --- | --- | --- | --- | --- | --- | --- |
| **Ion** | | | |  | 1.0 | 85.6 |  | 3.06 |
 1 | Water 1 | | A:2491 | 1.87 | 1.0 | 87.6 | 2.94 [A:N7]A:179 | 2.20 | 2 | A | OP2 | A:195 | 1.95 | 1.0 | 85.3 |  | 1.29 | 3 | U | O4 | A:180 | 2.11 | 1.0 | 86.1 |  | 1.28 | 4 | Water 2 | | A:2765 | 3.84 | 1.0 | 80.1 | 2.70 [C:OP2]A:194 | 1.47 | 5 | Water 3 | | A:3615 | 3.88 | 1.0 | 85.7 |  | 1.66 | 6 | A | OP1 | A:195 | 3.93 | 1.0 | 82.4 |  | 2.23 | 7 | C | O5' | A:194 | 3.94 | 1.0 | 81.7 |  | 2.43 | 8 | C | OP1 | A:194 | 4.12 | 1.0 | 80.2 |  | 2.28 | 9 | C | O3' | A:194 | 4.13 | 1.0 | 79.7 |  | 3.56 | 10 | C | OP2 | A:194 | 4.24 | 1.0 | 82.6 |  | 1.90 | 11 | A | N7 | A:179 | 4.26 | 1.0 | 84.4 |  | 2.35 | 12 | A | N7 | A:181 | 4.31 | 1.0 | 85.3 |  | 1.51 | 13 | A | O5' | A:195 | 4.37 | 1.0 | 82.1 |  | 2.78 | 14 | A | OP2 | A:179 | 4.43 | 1.0 | 87.2 |  | 0.73 |

  
**1^st^ shell coordination angles and distances (*trans* in "blue"; *cys* in "green"; deviations in "orange"):**   
• 1-2(W/OP2): 91º; 1-3(W/O4): *83º*; 2-3(OP2/O4): 87º;   
• 1-2(W/OP2): 2.7Å; 1-3(W/O4): 2.6Å; 2-3(OP2/O4): 2.8Å;   
  
**2.93 ± 0.96: the average coodination distance of the 1^st^ six ligands is K^+^ compatible**  
**6: Number of potential Water contacts**   
**2 : Number of second shell contacts involving water or ions**   
**33%: Percentage of satisfied second shell Water contacts**   
• Contact(s): /O4/OP2/  
• Contact code: O_ph_O_b_  
• Contact(s) in the exclusion zone: /O_w_/  
→ Missing octahedral coordination for MG ←  
→ MG ion B\_factor > 80 ←→2Fo-Fc density between 1.0 and 4.0 !   
Go to ToC

### MG:A:1624

|  | Res. type | Atom | Residue | Distance | Occupancy | B-factor | Closest atom(s) to a 1^st^ shell water | EM\_dens. |
| --- | --- | --- | --- | --- | --- | --- | --- | --- |
| **Ion** | | | |  | 1.0 | 66.9 |  | 2.93 |
 1 | Water 1 | | A:4056 | 1.80 | 1.0 | 58.4 |  | 2.82 | 2 | Water 2 | | A:1853 | 1.85 | 1.0 | 55.8 | 2.34 [U:OP1]A:387 | 2.36 | 3 | A | OP1 | A:59 | 1.96 | 1.0 | 54.7 |  | 3.06 | 4 | U | OP1 | A:387 | 2.68 | 1.0 | 56.2 |  | 3.10 | 5 | C | O3' | A:386 | 3.60 | 1.0 | 55.2 |  | 4.22 | 6 | C | O3' | A:58 | 3.69 | 1.0 | 53.7 |  | 4.23 | 7 | A | OP2 | A:59 | 3.97 | 1.0 | 55.7 |  | 2.87 | 8 | Water 3 | | A:2937 | 4.33 | 1.0 | 59.5 | 2.73 [U:OP1]A:387 | 2.36 | 9 | G | OP1 | A:61 | 4.40 | 1.0 | 54.3 |  | 4.22 | 10 | A | O5' | A:59 | 4.41 | 1.0 | 57.3 |  | 3.24 | 11 | Water 4 | | A:2641 | 4.60 | 1.0 | 58.3 | 2.68 [A:OP2]A:59 | 1.52 | 12 | G | O5' | A:61 | 4.61 | 1.0 | 54.9 |  | 2.79 | 13 | U | O5' | A:387 | 4.77 | 1.0 | 59.1 |  | 2.81 |

  
**1^st^ shell coordination angles and distances (*trans* in "blue"; *cys* in "green"; deviations in "orange"):**   
• 1-2(W/W): *150º*; 1-3(W/OP1): *105º*; 2-3(W/OP1): 89º;   
• 1-2(W/W): *3.5Å*; 1-3(W/OP1): 3.0Å; 2-3(W/OP1): 2.7Å;   
  
**2.60 ± 0.80: the average coodination distance of the 1^st^ six ligands is Na^+^** compatible  
**4: Number of potential Water contacts**   
**1 : Number of second shell contacts involving water or ions**   
**25%: Percentage of satisfied second shell Water contacts**   
• Contact(s): /OP1/  
• Contact code: O_ph_  
• Contact(s) in the exclusion zone: /O_w_/O_w_/OP1/  
→ Missing octahedral coordination for MG ←  
• MG hydration shell issue: water too close (< 2.4 Å.)  
→2Fo-Fc density between 1.0 and 4.0 !   
Go to ToC

### MG:A:1625

|  | Res. type | Atom | Residue | Distance | Occupancy | B-factor | Closest atom(s) to a 1^st^ shell water | EM\_dens. |
| --- | --- | --- | --- | --- | --- | --- | --- | --- |
| **Ion** | | | |  | 1.0 | 79.6 |  | 3.11 |
 1 | Water 1 | | A:4518 | 1.82 | 1.0 | 71.4 |  | 2.58 | 2 | G | OP2 | A:100 | 1.87 | 1.0 | 85.1 |  | 3.15 | 3 | G | O5' | A:100 | 3.62 | 1.0 | 86.8 |  | 2.40 | 4 | G | OP1 | A:100 | 4.20 | 1.0 | 88.1 |  | 1.42 | 5 | Water 2 | | A:3698 | 4.24 | 1.0 | 74.3 |  | 1.57 | 6 | C | O3' | A:99 | 4.28 | 1.0 | 93.2 |  | 1.43 | 7 | A | N7 | A:101 | 4.34 | 1.0 | 78.4 |  | 2.52 | 8 | A | OP2 | A:101 | 4.34 | 1.0 | 79.9 |  | 2.16 | 9 | G | N7 | A:100 | 4.35 | 1.0 | 86.4 |  | 2.67 | 10 | G | O6 | A:64 | 4.42 | 1.0 | 75.4 |  | 2.65 | 11 | G | N7 | A:64 | 4.53 | 1.0 | 73.4 |  | 3.68 |

  
**1^st^ shell coordination angles and distances (*trans* in "blue"; *cys* in "green"; deviations in "orange"):**   
• 1-2(W/OP2): *167º*;   
• 1-2(W/OP2): *3.7Å*;   
  
**3.34 ± 1.08: the average coodination distance of the 1^st^ six ligands is K^+^ compatible**  
**4: Number of potential Water contacts**   
**0 : Number of second shell contacts involving water or ions**   
**0%: Percentage of satisfied second shell Water contacts**   
• Contact code: → No contacts ←   
• Contact(s) in the exclusion zone: /O_w_/OP2/  
→ Firts MG coordination shell missing ←  
→2Fo-Fc density between 1.0 and 4.0 !   
Go to ToC

### MG:A:1626

|  | Res. type | Atom | Residue | Distance | Occupancy | B-factor | Closest atom(s) to a 1^st^ shell water | EM\_dens. |
| --- | --- | --- | --- | --- | --- | --- | --- | --- |
| **Ion** | | | |  | 1.0 | 66.9 |  | 3.39 |
 1 | C | OP2 | A:175 | 1.93 | 1.0 | 76.9 |  | 1.69 | 2 | A | OP1 | A:174 | 2.12 | 1.0 | 71.9 |  | 3.58 | 3 | C | OP1 | A:175 | 3.90 | 1.0 | 73.5 |  | 2.22 | 4 | A | OP2 | A:174 | 4.08 | 1.0 | 74.3 |  | 2.09 | 5 | A | O3' | A:174 | 4.15 | 1.0 | 71.8 |  | 3.99 | 6 | C | O5' | A:175 | 4.34 | 1.0 | 75.8 |  | 2.48 | 7 | A | O5' | A:174 | 4.34 | 1.0 | 73.7 |  | 2.71 | 8 | A | OP1 | A:197 | 4.37 | 1.0 | 80.5 |  | 1.70 | 9 | U | O3' | A:173 | 4.56 | 1.0 | 75.2 |  | 2.94 |

  
**1^st^ shell coordination angles and distances (*trans* in "blue"; *cys* in "green"; deviations in "orange"):**   
• 1-2(OP2/OP1): 88º;   
• 1-2(OP2/OP1): 2.8Å;   
  
**3.42 ± 1.00: the average coodination distance of the 1^st^ six ligands is K^+^ compatible**  
**0: Number of potential Water contacts**   
• Contact(s): /OP1/OP2/  
• Contact code: *cis-*2O_ph_  
→ Missing octahedral coordination for MG ←  
→2Fo-Fc density between 1.0 and 4.0 !   
Go to ToC

### MG:A:1627

|  | Res. type | Atom | Residue | Distance | Occupancy | B-factor | Closest atom(s) to a 1^st^ shell water | EM\_dens. |
| --- | --- | --- | --- | --- | --- | --- | --- | --- |
| **Ion** | | | |  | 1.0 | 54.3 |  | 6.10 |
 1 | G | O6 | A:299 | 1.93 | 1.0 | 48.1 |  | 5.51 | 2 | Water 1 | | A:2728 | 1.93 | 1.0 | 43.6 | 2.78 [A:OP1]A:560 // 2.85 [G:O6]A:566 | 5.64 | 3 | Water 2 | | A:3178 | 1.93 | 1.0 | 47.6 |  | 4.44 | 4 | Water 3 | | A:2928 | 1.99 | 1.0 | 47.9 | 2.73 [G:O2']A:297 | 4.68 | 5 | Water 4 | | A:2598 | 2.06 | 1.0 | 46.5 | 2.79 [A:N1]A:300 | 4.02 | 6 | G | OP1 | A:558 | 2.16 | 1.0 | 39.5 |  | 8.10 | 7 | G | O3' | A:557 | 3.87 | 1.0 | 41.1 |  | 6.73 | 8 | G | O2' | A:297 | 4.01 | 1.0 | 48.3 |  | 4.23 | 9 | G | O6 | A:566 | 4.09 | 1.0 | 44.0 |  | 5.75 | 10 | A | N1 | A:300 | 4.10 | 1.0 | 44.1 |  | 6.06 | 11 | A | OP1 | A:560 | 4.12 | 1.0 | 42.4 |  | 6.48 | 12 | G | OP2 | A:558 | 4.26 | 1.0 | 48.6 |  | 7.21 | 13 | Water 5 | | A:3904 | 4.40 | 1.0 | 50.0 |  | 3.01 | 14 | A | N7 | A:298 | 4.51 | 1.0 | 46.1 |  | 5.28 | 15 | Water 6 | | A:2786 | 4.56 | 1.0 | 41.7 | 2.70 [G:OP2]A:558 | 5.80 | 16 | G | O5' | A:558 | 4.62 | 1.0 | 45.8 |  | 6.05 | 17 | G | N7 | A:299 | 4.65 | 1.0 | 42.2 |  | 5.96 | 18 | Water 7 | | E:237 | 4.76 | 1.0 | 58.7 |  | 1.42 |

  
**1^st^ shell coordination angles and distances (*trans* in "blue"; *cys* in "green"; deviations in "orange"):**   
• 1-2(O6/W): *96º*; 1-3(O6/W): 92º; 1-4(O6/W): *100º*; 1-5(O6/W): *84º*; 1-6(O6/OP1): 178º; 2-3(W/W): 92º; 2-4(W/W): *159º*; 2-5(W/W): 86º; 2-6(W/OP1): *82º*; 3-4(W/W): *74º*; 3-5(W/W): 175º; 3-6(W/OP1): 89º; 4-5(W/W): *109º*; 4-6(W/OP1): *82º*; 5-6(W/OP1): *95º*;   
• 1-2(O6/W): 2.9Å; 1-3(O6/W): 2.8Å; 1-4(O6/W): 3.0Å; 1-5(O6/W): 2.7Å; 1-6(O6/OP1): 4.1Å; 2-3(W/W): 2.8Å; 2-4(W/W): 3.9Å; 2-5(W/W): 2.7Å; 2-6(W/OP1): 2.7Å; 3-4(W/W): 2.4Å; 3-5(W/W): 4.0Å; 3-6(W/OP1): 2.9Å; 4-5(W/W): *3.3Å*; 4-6(W/OP1): 2.7Å; 5-6(W/OP1): 3.1Å;   
  
**2.00 ± 0.08: the average coodination distance of the 1^st^ six ligands is Mg^2+^ compatible**  
**8: Number of potential Water contacts**   
**4 : Number of second shell contacts involving water or ions**   
**50%: Percentage of satisfied second shell Water contacts**   
• Contact(s): /O6/O_w_/O_w_/O_w_/O_w_/OP1/  
• Contact code: O_ph_O_b_4O_w_  
→ Slightly distorded coord. octahedron (7.6º) ←  
Go to ToC

### MG:A:1628

|  | Res. type | Atom | Residue | Distance | Occupancy | B-factor | Closest atom(s) to a 1^st^ shell water | EM\_dens. |
| --- | --- | --- | --- | --- | --- | --- | --- | --- |
| **Ion** | | | |  | 1.0 | 42.2 |  | 11.40 |
 1 | Water 1 | | A:2811 | 1.98 | 1.0 | 38.2 | 2.71 [A:OP1]A:573 | 8.03 | 2 | Water 2 | | A:4244 | 2.02 | 1.0 | 39.2 |  | 7.10 | 3 | Water 3 | | A:3872 | 2.06 | 1.0 | 39.3 |  | 6.75 | 4 | A | OP1 | A:572 | 2.19 | 1.0 | 37.9 |  | 7.95 | 5 | Water 4 | | A:2452 | 2.27 | 1.0 | 42.7 | 2.63 [A:N1]A:819 | 5.64 | 6 | Water 5 | | A:4260 | 4.01 | 1.0 | 34.6 |  | 8.96 | 7 | Water 6 | | A:3467 | 4.05 | 1.0 | 39.1 | 2.84 [A:OP1]A:573 | 6.13 | 8 | Water 7 | | A:3502 | 4.18 | 1.0 | 43.4 |  | 6.22 | 9 | A | O5' | A:572 | 4.22 | 1.0 | 35.4 |  | 8.03 | 10 | A | N1 | A:819 | 4.29 | 1.0 | 35.3 |  | 8.70 | 11 | U | O3' | A:571 | 4.30 | 1.0 | 33.9 |  | 8.79 | 12 | A | OP1 | A:573 | 4.36 | 1.0 | 38.9 |  | 7.89 | 13 | Water 8 | | A:4653 | 4.67 | 1.0 | 49.6 |  | 2.77 | 14 | A | OP2 | A:572 | 4.69 | 1.0 | 40.8 |  | 8.92 | 15 | Water 9 | | A:2497 | 4.79 | 1.0 | 44.9 |  | 3.23 |

  
**1^st^ shell coordination angles and distances (*trans* in "blue"; *cys* in "green"; deviations in "orange"):**   
• 1-2(W/W): 89º; 1-3(W/W): 170º; 1-4(W/OP1): 87º; 1-5(W/W): *97º*; 2-3(W/W): *84º*; 2-4(W/OP1): 93º; 2-5(W/W): 174º; 3-4(W/OP1): 87º; 3-5(W/W): 90º; 4-5(OP1/W): 85º;   
• 1-2(W/W): 2.8Å; 1-3(W/W): 4.0Å; 1-4(W/OP1): 2.9Å; 1-5(W/W): 3.2Å; 2-3(W/W): 2.7Å; 2-4(W/OP1): 3.0Å; 2-5(W/W): *4.3Å*; 3-4(W/OP1): 2.9Å; 3-5(W/W): 3.1Å; 4-5(OP1/W): 3.0Å;   
  
**2.42 ± 0.72: the average coodination distance of the 1^st^ six ligands is Na^+^** compatible  
**10: Number of potential Water contacts**   
**2 : Number of second shell contacts involving water or ions**   
**20%: Percentage of satisfied second shell Water contacts**   
• Contact(s): /O_w_/O_w_/O_w_/O_w_/OP1/  
• Contact code: O_ph_4O_w_  
→ Missing octahedral coordination for MG ←  
Ooctahedral coordination (4.4º)   
Go to ToC

### MG:A:1629

|  | Res. type | Atom | Residue | Distance | Occupancy | B-factor | Closest atom(s) to a 1^st^ shell water | EM\_dens. |
| --- | --- | --- | --- | --- | --- | --- | --- | --- |
| **Ion** | | | |  | 1.0 | 44.2 |  | 9.03 |
 1 | C | OP1 | A:576 | 1.86 | 1.0 | 38.9 |  | 9.06 | 2 | Water 1 | | A:2755 | 1.95 | 1.0 | 39.8 |  | 5.72 | 3 | Water 2 | | A:4302 | 1.96 | 1.0 | 43.7 |  | 4.53 | 4 | Water 3 | | A:4645 | 2.02 | 1.0 | 42.1 |  | 5.99 | 5 | Water 4 | | A:3417 | 2.20 | 1.0 | 34.4 | 2.84 [G:OP2]A:577 | 7.59 | 6 | Water 5 | | A:3041 | 2.27 | 1.0 | 35.5 | 2.75 [U:OP1]A:820 | 8.07 | 7 | C | OP2 | A:576 | 3.83 | 1.0 | 39.4 |  | 8.38 | 8 | Water 6 | | A:2212 | 3.93 | 1.0 | 37.6 | 2.74 [C:OP1]A:817 | 7.54 | 9 | Water 7 | | A:2891 | 3.98 | 1.0 | 38.2 | 2.84 [C:O5']A:576 | 7.55 | 10 | Water 8 | | A:3496 | 4.07 | 1.0 | 41.9 |  | 5.44 | 11 | C | O5' | A:576 | 4.14 | 1.0 | 43.6 |  | 8.54 | 12 | U | OP1 | A:820 | 4.21 | 1.0 | 38.0 |  | 8.19 | 13 | G | O3' | A:575 | 4.24 | 1.0 | 35.7 |  | 9.27 | 14 | Water 9 | | A:2649 | 4.57 | 1.0 | 34.7 | 2.68 [C:OP2]A:576 | 10.76 | 15 | C | OP1 | A:817 | 4.59 | 1.0 | 34.9 |  | 7.36 | 16 | G | OP2 | A:577 | 4.76 | 1.0 | 38.1 |  | 8.96 | 17 | Water 10 | | A:3631 | 4.77 | 1.0 | 45.3 |  | 3.04 | 18 | G | O5' | A:575 | 4.79 | 1.0 | 43.6 |  | 7.09 |

  
**1^st^ shell coordination angles and distances (*trans* in "blue"; *cys* in "green"; deviations in "orange"):**   
• 1-2(OP1/W): 90º; 1-3(OP1/W): *107º*; 1-4(OP1/W): *167º*; 1-5(OP1/W): 88º; 1-6(OP1/W): *103º*; 2-3(W/W): *84º*; 2-4(W/W): *84º*; 2-5(W/W): 86º; 2-6(W/W): *166º*; 3-4(W/W): *84º*; 3-5(W/W): *162º*; 3-6(W/W): 87º; 4-5(W/W): *80º*; 4-6(W/W): *84º*; 5-6(W/W): *99º*;   
• 1-2(OP1/W): 2.7Å; 1-3(OP1/W): 3.1Å; 1-4(OP1/W): 3.9Å; 1-5(OP1/W): 2.8Å; 1-6(OP1/W): *3.2Å*; 2-3(W/W): 2.6Å; 2-4(W/W): 2.7Å; 2-5(W/W): 2.8Å; 2-6(W/W): 4.2Å; 3-4(W/W): 2.7Å; 3-5(W/W): 4.1Å; 3-6(W/W): 2.9Å; 4-5(W/W): 2.7Å; 4-6(W/W): 2.9Å; 5-6(W/W): *3.4Å*;   
  
**2.04 ± 0.14: the average coodination distance of the 1^st^ six ligands is Mg^2+^ compatible**  
**10: Number of potential Water contacts**   
**2 : Number of second shell contacts involving water or ions**   
**20%: Percentage of satisfied second shell Water contacts**   
• Contact(s): /O_w_/O_w_/O_w_/O_w_/O_w_/  
• Contact code: 5O_w_  
• Contact(s) in the exclusion zone: /OP1/  
→ Missing octahedral coordination for MG ←  
→ Slightly distorded coord. octahedron (8.4º) ←  
Go to ToC

### MG:A:1630

|  | Res. type | Atom | Residue | Distance | Occupancy | B-factor | Closest atom(s) to a 1^st^ shell water | EM\_dens. |
| --- | --- | --- | --- | --- | --- | --- | --- | --- |
| **Ion** | | | |  | 1.0 | 56.4 |  | 3.72 |
 1 | Water 1 | | A:3964 | 1.95 | 1.0 | 55.8 | 2.96 [C:OP2]A:580 | 2.56 | 2 | Water 2 | | A:3095 | 2.25 | 1.0 | 44.5 | 2.76 [C:OP2]A:758 | 3.51 | 3 | Water 3 | | A:2289 | 2.33 | 1.0 | 51.5 | 2.59 [U:OP2]A:757 | 4.22 | 4 | Water 4 | | A:1996 | 2.43 | 1.0 | 46.4 | 2.47 [C:OP2]A:758 | 3.27 | 5 | C | OP2 | A:580 | 2.43 | 1.0 | 47.3 |  | 4.50 | 6 | Water 5 | | A:2398 | 3.42 | 1.0 | 48.0 | 2.62 [G:OP2]A:581 | 1.94 | 7 | C | O5' | A:580 | 3.72 | 1.0 | 48.2 |  | 4.34 | 8 | Water 6 | | A:2707 | 3.78 | 1.0 | 47.4 | 2.69 [G:OP2]A:581 | 2.49 | 9 | C | OP2 | A:758 | 3.78 | 1.0 | 49.9 |  | 4.68 | 10 | G | OP2 | A:581 | 4.05 | 1.0 | 46.3 |  | 4.26 | 11 | G | N7 | A:581 | 4.12 | 1.0 | 36.5 |  | 8.67 | 12 | Water 7 | | A:4625 | 4.19 | 1.0 | 50.3 |  | 2.48 | 13 | C | OP1 | A:580 | 4.21 | 1.0 | 47.2 |  | 2.81 | 14 | Water 8 | | A:3914 | 4.40 | 1.0 | 53.3 |  | 2.79 | 15 | U | OP2 | A:757 | 4.56 | 1.0 | 39.0 |  | 6.49 |

  
**1^st^ shell coordination angles and distances (*trans* in "blue"; *cys* in "green"; deviations in "orange"):**   
• 1-2(W/W): *160º*;   
• 1-2(W/W): 4.1Å;   
  
**2.47 ± 0.46: the average coodination distance of the 1^st^ six ligands is Na^+^** compatible  
**10: Number of potential Water contacts**   
**5 : Number of second shell contacts involving water or ions**   
**50%: Percentage of satisfied second shell Water contacts**   
• Contact(s): /O_w_/O_w_/  
• Contact code: 2O_w_  
• Contact(s) in the exclusion zone: /O_w_/O_w_/OP2/  
→ Missing octahedral coordination for MG ←  
→2Fo-Fc density between 1.0 and 4.0 !   
Go to ToC

### MG:A:1631

|  | Res. type | Atom | Residue | Distance | Occupancy | B-factor | Closest atom(s) to a 1^st^ shell water | EM\_dens. |
| --- | --- | --- | --- | --- | --- | --- | --- | --- |
| **Ion** | | | |  | 1.0 | 63.4 |  | 3.08 |
 1 | G | OP2 | A:818 | 1.88 | 1.0 | 46.4 |  | 4.95 | 2 | Water 1 | | A:4208 | 2.09 | 1.0 | 52.5 |  | 2.19 | 3 | Water 2 | | A:3193 | 2.12 | 1.0 | 48.4 | 2.78 [U:OP1]A:1527 | 3.76 | 4 | Water 3 | | A:3679 | 2.13 | 1.0 | 47.4 | 2.45 [K:K]A:1735 | 2.67 | 5 | Water 4 | | A:4063 | 2.22 | 1.0 | 51.2 |  | 1.97 | 6 | **K** | K | A:1735 | 2.32 | 1.0 | 30.0 |  | 7.16 | 7 | C | O3' | A:817 | 3.56 | 1.0 | 42.6 |  | 7.12 | 8 | Water 5 | | A:3632 | 3.66 | 1.0 | 52.4 |  | 1.58 | 9 | Water 6 | | A:3532 | 4.04 | 1.0 | 51.0 |  | 1.93 | 10 | G | O5' | A:818 | 4.18 | 1.0 | 45.7 |  | 5.88 | 11 | G | OP1 | A:818 | 4.22 | 1.0 | 42.5 |  | 5.75 | 12 | U | OP1 | A:1528 | 4.34 | 1.0 | 40.2 |  | 5.99 | 13 | Water 7 | | A:2390 | 4.38 | 1.0 | 50.0 |  | 2.58 | 14 | Water 8 | | A:3824 | 4.45 | 1.0 | 37.1 | 2.92 [U:OP1]A:1527 | 6.29 | 15 | U | OP1 | A:1527 | 4.54 | 1.0 | 36.8 |  | 8.53 | 16 | Water 9 | | A:4664 | 4.70 | 1.0 | 47.4 |  | 3.59 | 17 | C | O2' | A:817 | 4.71 | 1.0 | 40.2 |  | 7.85 |

  
**1^st^ shell coordination angles and distances (*trans* in "blue"; *cys* in "green"; deviations in "orange"):**   
• 1-2(OP2/W): *99º*; 1-3(OP2/W): *170º*; 1-4(OP2/W): 91º; 1-5(OP2/W): 93º; 2-3(W/W): 91º; 2-4(W/W): *158º*; 2-5(W/W): *115º*; 3-4(W/W): *79º*; 3-5(W/W): *84º*; 4-5(W/W): *82º*;   
• 1-2(OP2/W): 3.0Å; 1-3(OP2/W): 4.0Å; 1-4(OP2/W): 2.9Å; 1-5(OP2/W): 3.0Å; 2-3(W/W): 3.0Å; 2-4(W/W): 4.2Å; 2-5(W/W): *3.6Å*; 3-4(W/W): 2.7Å; 3-5(W/W): 2.9Å; 4-5(W/W): 2.9Å;   
  
**2.13 ± 0.13: the average coodination distance of the 1^st^ six ligands is Mg^2+^ compatible**  
**8: Number of potential Water contacts**   
**2 : Number of second shell contacts involving water or ions**   
**25%: Percentage of satisfied second shell Water contacts**   
• Contact(s): /O_w_/O_w_/O_w_/O_w_/  
• Contact code: 4O_w_  
• Contact(s) in the exclusion zone: /OP2/  
→ Missing octahedral coordination for MG ←  
→ Slightly distorded coord. octahedron (9.6º) ←→2Fo-Fc density between 1.0 and 4.0 !   
Go to ToC

### MG:A:1632

|  | Res. type | Atom | Residue | Distance | Occupancy | B-factor | Closest atom(s) to a 1^st^ shell water | EM\_dens. |
| --- | --- | --- | --- | --- | --- | --- | --- | --- |
| **Ion** | | | |  | 1.0 | 39.9 |  | 13.03 |
 1 | Water 1 | | A:2516 | 2.03 | 1.0 | 37.0 | 2.64 [U:O4]A:904 | 7.54 | 2 | Water 2 | | A:4068 | 2.06 | 1.0 | 43.1 | 2.99 [A:OP2]A:892 | 6.77 | 3 | Water 3 | | A:2757 | 2.12 | 1.0 | 37.8 | 2.70 [A:N1]A:906 | 8.17 | 4 | Water 4 | | A:3161 | 2.27 | 1.0 | 38.3 | 2.77 [G:OP2]A:903 | 7.81 | 5 | Water 5 | | A:2435 | 2.30 | 1.0 | 31.8 | 2.63 [U:O4]A:905 | 9.86 | 6 | U | OP2 | A:891 | 2.31 | 1.0 | 41.5 |  | 8.31 | 7 | U | O5' | A:891 | 3.48 | 1.0 | 44.0 |  | 5.46 | 8 | A | OP2 | A:892 | 3.72 | 1.0 | 46.2 |  | 5.87 | 9 | A | N1 | A:906 | 4.09 | 1.0 | 36.9 |  | 8.42 | 10 | U | OP1 | A:891 | 4.09 | 1.0 | 44.1 |  | 5.19 | 11 | U | O4 | A:904 | 4.15 | 1.0 | 36.2 |  | 10.41 | 12 | Water 6 | | A:3401 | 4.24 | 1.0 | 44.7 |  | 4.55 | 13 | G | OP2 | A:903 | 4.31 | 1.0 | 38.5 |  | 8.95 | 14 | Water 7 | | A:4050 | 4.38 | 1.0 | 41.9 |  | 6.70 | 15 | U | O4 | A:905 | 4.40 | 1.0 | 38.2 |  | 8.36 | 16 | Water 8 | | A:2114 | 4.46 | 1.0 | 48.8 | 2.53 [G:OP2]A:903 | 2.91 | 17 | Water 9 | | A:4640 | 4.52 | 1.0 | 42.5 |  | 3.32 | 18 | Water 10 | | A:3985 | 4.59 | 1.0 | 37.0 |  | 5.86 | 19 | Water 11 | | A:2651 | 4.64 | 1.0 | 44.5 | 2.68 [U:OP1]A:891 | 3.28 | 20 | G | O3' | A:890 | 4.66 | 1.0 | 44.0 |  | 4.65 | 21 | Water 12 | | A:2793 | 4.74 | 1.0 | 42.5 |  | 6.87 |

  
**1^st^ shell coordination angles and distances (*trans* in "blue"; *cys* in "green"; deviations in "orange"):**   
• 1-2(W/W): 94º; 1-3(W/W): *98º*; 1-4(W/W): 90º; 1-5(W/W): *83º*; 2-3(W/W): *98º*; 2-4(W/W): *97º*; 2-5(W/W): 177º; 3-4(W/W): *162º*; 3-5(W/W): *81º*; 4-5(W/W): *84º*;   
• 1-2(W/W): 3.0Å; 1-3(W/W): 3.1Å; 1-4(W/W): 3.1Å; 1-5(W/W): 2.9Å; 2-3(W/W): 3.1Å; 2-4(W/W): *3.3Å*; 2-5(W/W): *4.4Å*; 3-4(W/W): *4.3Å*; 3-5(W/W): 2.9Å; 4-5(W/W): 3.1Å;   
  
**2.18 ± 0.11: the average coodination distance of the 1^st^ six ligands is Mg^2+^ compatible**  
**10: Number of potential Water contacts**   
**5 : Number of second shell contacts involving water or ions**   
**50%: Percentage of satisfied second shell Water contacts**   
• Contact(s): /O_w_/O_w_/O_w_/O_w_/O_w_/  
• Contact code: 5O_w_  
• Contact(s) in the exclusion zone: /OP2/  
→ Missing octahedral coordination for MG ←  
→ Slightly distorded coord. octahedron (7.0º) ←  
Go to ToC

### MG:A:1633

|  | Res. type | Atom | Residue | Distance | Occupancy | B-factor | Closest atom(s) to a 1^st^ shell water | EM\_dens. |
| --- | --- | --- | --- | --- | --- | --- | --- | --- |
| **Ion** | | | |  | 1.0 | 53.9 |  | 7.12 |
 1 | Water 1 | | A:3508 | 1.88 | 1.0 | 45.2 | 2.84 [G:N7]A:1416 | 4.75 | 2 | Water 2 | | A:2742 | 1.94 | 1.0 | 39.0 | 2.73 [G:O6]A:1482 | 7.04 | 3 | Water 3 | | A:3214 | 2.03 | 1.0 | 43.2 |  | 4.47 | 4 | G | O6 | A:1417 | 2.19 | 1.0 | 43.0 |  | 7.19 | 5 | Water 4 | | A:3500 | 2.26 | 1.0 | 51.7 | 2.84 [G:OP2]A:1415 | 3.61 | 6 | Water 5 | | A:2691 | 2.41 | 1.0 | 47.5 | 2.69 [G:N7]A:1417 | 5.19 | 7 | Water 6 | | A:3717 | 3.90 | 1.0 | 48.2 | 2.89 [G:O6]A:1416 | 3.32 | 8 | G | N7 | A:1416 | 3.93 | 1.0 | 39.8 |  | 7.01 | 9 | G | N7 | A:1417 | 3.94 | 1.0 | 39.1 |  | 8.16 | 10 | G | O6 | A:1482 | 3.95 | 1.0 | 42.2 |  | 8.34 | 11 | Water 7 | | A:3270 | 4.01 | 1.0 | 46.3 |  | 3.96 | 12 | Water 8 | | A:4673 | 4.12 | 1.0 | 54.2 |  | 1.90 | 13 | Water 9 | | A:1992 | 4.20 | 1.0 | 48.6 | 2.47 [G:OP2]A:1416 | 2.00 | 14 | G | OP2 | A:1416 | 4.28 | 1.0 | 42.2 |  | 5.05 | 15 | Water 10 | | A:3921 | 4.42 | 1.0 | 52.8 |  | 3.32 | 16 | G | O6 | A:1416 | 4.57 | 1.0 | 37.5 |  | 9.02 | 17 | Water 11 | | A:3661 | 4.58 | 1.0 | 52.1 |  | 2.96 | 18 | G | OP2 | A:1415 | 4.60 | 1.0 | 46.0 |  | 4.46 |

  
**1^st^ shell coordination angles and distances (*trans* in "blue"; *cys* in "green"; deviations in "orange"):**   
• 1-2(W/W): 173º; 1-3(W/W): 89º; 1-4(W/O6): *102º*; 1-5(W/W): 89º; 2-3(W/W): 86º; 2-4(W/O6): *81º*; 2-5(W/W): 88º; 3-4(W/O6): *82º*; 3-5(W/W): *99º*; 4-5(O6/W): *170º*;   
• 1-2(W/W): 3.8Å; 1-3(W/W): 2.7Å; 1-4(W/O6): 3.2Å; 1-5(W/W): 2.9Å; 2-3(W/W): 2.7Å; 2-4(W/O6): 2.7Å; 2-5(W/W): 2.9Å; 3-4(W/O6): 2.8Å; 3-5(W/W): *3.3Å*; 4-5(O6/W): *4.4Å*;   
  
**2.12 ± 0.18: the average coodination distance of the 1^st^ six ligands is Mg^2+^ compatible**  
**10: Number of potential Water contacts**   
**4 : Number of second shell contacts involving water or ions**   
**40%: Percentage of satisfied second shell Water contacts**   
• Contact(s): /O6/O_w_/O_w_/O_w_/  
• Contact code: O_b_3O_w_  
• Contact(s) in the exclusion zone: /O_w_/O_w_/  
→ Missing octahedral coordination for MG ←  
→ Slightly distorded coord. octahedron (6.2º) ←  
Go to ToC

### MG:A:1634

|  | Res. type | Atom | Residue | Distance | Occupancy | B-factor | Closest atom(s) to a 1^st^ shell water | EM\_dens. |
| --- | --- | --- | --- | --- | --- | --- | --- | --- |
| **Ion** | | | |  | 1.0 | 37.8 |  | 15.06 |
 1 | A | OP1 | A:1500 | 1.99 | 1.0 | 32.4 |  | 10.66 | 2 | A | OP1 | A:1508 | 2.06 | 1.0 | 31.1 |  | 11.65 | 3 | Water 1 | | A:4005 | 2.12 | 1.0 | 36.2 |  | 10.57 | 4 | G | OP1 | A:1505 | 2.12 | 1.0 | 36.2 |  | 8.72 | 5 | Water 2 | | A:2940 | 2.15 | 1.0 | 28.4 | 2.73 [C:OP2]A:1501 | 11.11 | 6 | Water 3 | | A:2991 | 2.15 | 1.0 | 34.3 | 2.74 [G:OP1]A:1504 // 2.99 [A:O3']A:1507 | 8.62 | 7 | **K** | K | A:1724 | 3.46 | 1.0 | 30.0 |  | 21.79 | 8 | A | O3' | A:1507 | 3.49 | 1.0 | 37.1 |  | 7.42 | 9 | A | OP2 | A:1500 | 4.02 | 1.0 | 34.8 |  | 9.74 | 10 | Water 4 | | A:4185 | 4.14 | 1.0 | 32.5 |  | 11.09 | 11 | U | O2' | A:1506 | 4.17 | 1.0 | 39.5 |  | 10.04 | 12 | A | O5' | A:1508 | 4.28 | 1.0 | 36.0 |  | 7.83 | 13 | G | OP1 | A:1504 | 4.29 | 1.0 | 39.7 |  | 8.38 | 14 | A | O5' | A:1500 | 4.31 | 1.0 | 31.7 |  | 10.03 | 15 | A | O3' | A:1499 | 4.34 | 1.0 | 37.5 |  | 9.19 | 16 | A | OP2 | A:1508 | 4.35 | 1.0 | 35.5 |  | 9.31 | 17 | G | OP2 | A:1505 | 4.41 | 1.0 | 33.4 |  | 10.88 | 18 | G | O5' | A:1505 | 4.42 | 1.0 | 38.1 |  | 9.48 | 19 | G | O3' | A:1504 | 4.44 | 1.0 | 37.7 |  | 9.05 | 20 | Water 5 | | A:4333 | 4.57 | 1.0 | 38.6 |  | 6.38 | 21 | C | OP2 | A:1501 | 4.60 | 1.0 | 36.7 |  | 9.87 | 22 | G | O2' | A:1504 | 4.72 | 1.0 | 33.8 |  | 10.32 |

  
**1^st^ shell coordination angles and distances (*trans* in "blue"; *cys* in "green"; deviations in "orange"):**   
• 1-2(OP1/OP1): 91º; 1-3(OP1/W): 94º; 1-4(OP1/OP1): 89º; 1-5(OP1/W): 88º; 1-6(OP1/W): 174º; 2-3(OP1/W): 91º; 2-4(OP1/OP1): 177º; 2-5(OP1/W): 91º; 2-6(OP1/W): 94º; 3-4(W/OP1): 93º; 3-5(W/W): 177º; 3-6(W/W): 90º; 4-5(OP1/W): 86º; 4-6(OP1/W): 86º; 5-6(W/W): 88º;   
• 1-2(OP1/OP1): 2.9Å; 1-3(OP1/W): 3.0Å; 1-4(OP1/OP1): 2.9Å; 1-5(OP1/W): 2.9Å; 1-6(OP1/W): 4.1Å; 2-3(OP1/W): 3.0Å; 2-4(OP1/OP1): 4.2Å; 2-5(OP1/W): 3.0Å; 2-6(OP1/W): 3.1Å; 3-4(W/OP1): 3.1Å; 3-5(W/W): *4.3Å*; 3-6(W/W): 3.0Å; 4-5(OP1/W): 2.9Å; 4-6(OP1/W): 2.9Å; 5-6(W/W): 3.0Å;   
  
**2.10 ± 0.06: the average coodination distance of the 1^st^ six ligands is Mg^2+^ compatible**  
**6: Number of potential Water contacts**   
**3 : Number of second shell contacts involving water or ions**   
**50%: Percentage of satisfied second shell Water contacts**   
• Contact(s): /O_w_/O_w_/O_w_/OP1/OP1/OP1/  
• Contact code: *mer-*3O_ph_3O_w_  
Ooctahedral coordination (2.5º)   
Go to ToC

### MG:A:1635

|  | Res. type | Atom | Residue | Distance | Occupancy | B-factor | Closest atom(s) to a 1^st^ shell water | EM\_dens. |
| --- | --- | --- | --- | --- | --- | --- | --- | --- |
| **Ion** | | | |  | 1.0 | 70.5 |  | 1.79 |
 1 | Water 1 | | A:3787 | 1.96 | 1.0 | 73.8 | 2.91 [G:N7]A:259 | 1.67 | 2 | Water 2 | | A:4440 | 2.55 | 1.0 | 68.3 |  | 1.73 | 3 | G | OP2 | A:259 | 3.67 | 1.0 | 71.1 |  | 1.79 | 4 | ASN | OD1 | T:78 | 4.16 | 1.0 | 72.5 |  | 2.06 | 5 | G | N7 | A:259 | 4.48 | 1.0 | 67.8 |  | 3.47 | 6 | G | OP2 | A:258 | 4.60 | 1.0 | 72.2 |  | 1.52 | 7 | G | O6 | A:260 | 4.64 | 1.0 | 66.4 |  | 2.67 |

  
• • **3.57 ± 0.99: the average coodination distance of the 1^st^ six ligands is K^+^ compatible**  
**4: Number of potential Water contacts**   
**1 : Number of second shell contacts involving water or ions**   
**25%: Percentage of satisfied second shell Water contacts**   
• Contact(s): /O_w_/  
• Contact code: O_w_  
• Contact(s) in the exclusion zone: /O_w_/  
→ Missing octahedral coordination for MG ←  
→2Fo-Fc density between 1.0 and 4.0 !   
Go to ToC

### MG:A:1636

|  | Res. type | Atom | Residue | Distance | Occupancy | B-factor | Closest atom(s) to a 1^st^ shell water | EM\_dens. |
| --- | --- | --- | --- | --- | --- | --- | --- | --- |
| **Ion** | | | |  | 1.0 | 51.6 |  | 7.15 |
 1 | G | O6 | A:645 | 2.57 | 1.0 | 59.3 |  | 4.29 | 2 | Water 1 | | A:2370 | 2.65 | 1.0 | 60.8 | 2.30 [MG:MG]A:1678 | 3.60 | 3 | Water 2 | | A:2720 | 2.82 | 1.0 | 59.8 | 2.16 [MG:MG]A:1678 | 3.77 | 4 | Water 3 | | A:2908 | 3.05 | 1.0 | 59.5 | 2.73 [A:N1]A:595 // 2.94 [C:N4]A:643 | 4.01 | 5 | U | O4 | A:594 | 3.08 | 1.0 | 63.5 |  | 3.70 | 6 | U | O4 | A:644 | 3.17 | 1.0 | 57.2 |  | 3.25 | 7 | Water 4 | | A:4678 | 3.65 | 1.0 | 63.3 | 1.86 [MG:MG]A:1678 | 2.27 | 8 | **MG** | MG | A:1678 | 3.75 | 1.0 | 68.1 |  | 4.42 | 9 | G | N7 | A:645 | 3.97 | 1.0 | 57.2 |  | 4.03 | 10 | A | N1 | A:595 | 4.08 | 1.0 | 62.2 |  | 3.15 | 11 | U | O4 | A:593 | 4.17 | 1.0 | 70.6 |  | 2.85 | 12 | Water 5 | | A:3834 | 4.45 | 1.0 | 60.2 | 1.88 [MG:MG]A:1678 | 3.55 | 13 | Water 6 | | A:3406 | 4.49 | 1.0 | 57.0 |  | 2.30 | 14 | G | O6 | A:646 | 4.54 | 1.0 | 64.9 |  | 3.97 | 15 | Water 7 | | A:2245 | 4.70 | 1.0 | 57.3 |  | 2.75 | 16 | Water 8 | | A:2036 | 4.76 | 1.0 | 63.8 | 2.48 [U:O4]A:593 | 2.20 |

  
• • **2.89 ± 0.23: the average coodination distance of the 1^st^ six ligands is K^+^ compatible**  
**6: Number of potential Water contacts**   
**4 : Number of second shell contacts involving water or ions**   
**67%: Percentage of satisfied second shell Water contacts**   
• Contact code: → No contacts ←   
• Contact(s) in the exclusion zone: /O4/O4/O6/O_w_/O_w_/O_w_/  
→ Consider K in place of MG (2 valid contact(s) for K vs. 0 for MG ←)  
Go to ToC

### MG:A:1637

|  | Res. type | Atom | Residue | Distance | Occupancy | B-factor | Closest atom(s) to a 1^st^ shell water | EM\_dens. |
| --- | --- | --- | --- | --- | --- | --- | --- | --- |
| **Ion** | | | |  | 1.0 | 63.0 |  | 3.29 |
 1 | G | OP1 | A:266 | 2.49 | 1.0 | 68.7 |  | 2.61 | 2 | LEU | O | Q:67 | 2.55 | 1.0 | 65.8 |  | 2.40 | 3 | Water 1 | | A:4284 | 3.10 | 1.0 | 70.4 |  | 1.68 | 4 | G | O3' | A:265 | 3.51 | 1.0 | 63.0 |  | 4.25 | 5 | Water 2 | | A:1911 | 3.54 | 1.0 | 69.3 | 2.40 [G:OP2]A:255 // 2.66 [G:OP1]A:266 | 2.37 | 6 | G | OP2 | A:255 | 3.71 | 1.0 | 75.5 |  | 2.21 | 7 | G | OP2 | A:266 | 3.95 | 1.0 | 69.1 |  | 2.31 | 8 | **K** | K | A:1602 | 4.19 | 1.0 | 30.0 |  | 4.24 | 9 | G | OP1 | A:255 | 4.23 | 1.0 | 71.3 |  | 2.60 | 10 | U | OP2 | A:256 | 4.54 | 1.0 | 74.8 |  | 1.79 | 11 | G | O5' | A:266 | 4.80 | 1.0 | 66.9 |  | 2.50 |

  
• • **3.15 ± 0.48: the average coodination distance of the 1^st^ six ligands is K^+^ compatible**  
**6: Number of potential Water contacts**   
**2 : Number of second shell contacts involving water or ions**   
**33%: Percentage of satisfied second shell Water contacts**   
• Contact code: → No contacts ←   
• Contact(s) in the exclusion zone: /O_back_/O_w_/OP1/  
→ Firts MG coordination shell missing ←  
→2Fo-Fc density between 1.0 and 4.0 !   
Go to ToC

### MG:A:1638

|  | Res. type | Atom | Residue | Distance | Occupancy | B-factor | Closest atom(s) to a 1^st^ shell water | EM\_dens. |
| --- | --- | --- | --- | --- | --- | --- | --- | --- |
| **Ion** | | | |  | 1.0 | 53.7 |  | 5.46 |
 1 | Water 1 | | a:8650 | 1.99 | 1.0 | 46.0 | 2.95 [C:OP1]a:1835 | 5.14 | 2 | Water 2 | | A:3664 | 2.00 | 1.0 | 42.1 |  | 5.71 | 3 | Water 3 | | A:4272 | 2.08 | 1.0 | 46.8 |  | 3.56 | 4 | Water 4 | | a:10144 | 2.11 | 1.0 | 48.8 |  | 3.75 | 5 | C | OP1 | A:770 | 2.12 | 1.0 | 37.8 |  | 8.68 | 6 | Water 5 | | A:3157 | 2.27 | 1.0 | 44.3 |  | 4.06 | 7 | Water 6 | | A:3173 | 2.96 | 1.0 | 50.8 |  | 2.34 | 8 | G | O3' | A:769 | 4.07 | 1.0 | 37.7 |  | 8.45 | 9 | Water 7 | | A:3448 | 4.08 | 1.0 | 40.8 | 2.99 [G:O3']A:769 | 6.47 | 10 | C | O5' | A:770 | 4.27 | 1.0 | 34.1 |  | 10.55 | 11 | Water 8 | | a:6564 | 4.31 | 1.0 | 44.2 |  | 3.00 | 12 | Water 9 | | A:2257 | 4.31 | 1.0 | 43.3 |  | 6.07 | 13 | Water 10 | | A:3837 | 4.35 | 1.0 | 45.4 |  | 3.85 | 14 | Water 11 | | A:2520 | 4.39 | 1.0 | 40.2 | 2.92 [C:OP2]A:770 | 6.51 | 15 | Water 12 | | a:3818 | 4.49 | 1.0 | 48.0 | 2.54 [C:OP1]a:1835 | 2.95 | 16 | Water 13 | | A:3821 | 4.49 | 1.0 | 48.6 |  | 2.30 | 17 | C | OP2 | A:770 | 4.63 | 1.0 | 34.8 |  | 10.67 | 18 | C | OP1 | a:1835 | 4.65 | 1.0 | 43.5 |  | 6.11 | 19 | Water 14 | | a:9462 | 4.69 | 1.0 | 46.8 |  | 1.31 |

  
**1^st^ shell coordination angles and distances (*trans* in "blue"; *cys* in "green"; deviations in "orange"):**   
• 1-2(W/W): 93º; 1-3(W/W): 90º; 1-4(W/W): 93º; 1-5(W/OP1): *99º*; 1-6(W/W): 177º; 2-3(W/W): 176º; 2-4(W/W): *100º*; 2-5(W/OP1): 89º; 2-6(W/W): 87º; 3-4(W/W): *77º*; 3-5(W/OP1): 93º; 3-6(W/W): 90º; 4-5(W/OP1): *165º*; 4-6(W/W): 90º; 5-6(OP1/W): *78º*;   
• 1-2(W/W): 2.9Å; 1-3(W/W): 2.9Å; 1-4(W/W): 3.0Å; 1-5(W/OP1): 3.1Å; 1-6(W/W): *4.3Å*; 2-3(W/W): 4.1Å; 2-4(W/W): 3.1Å; 2-5(W/OP1): 2.9Å; 2-6(W/W): 2.9Å; 3-4(W/W): 2.6Å; 3-5(W/OP1): 3.1Å; 3-6(W/W): 3.1Å; 4-5(W/OP1): 4.2Å; 4-6(W/W): 3.1Å; 5-6(OP1/W): 2.8Å;   
  
**2.09 ± 0.09: the average coodination distance of the 1^st^ six ligands is Mg^2+^ compatible**  
**10: Number of potential Water contacts**   
**1 : Number of second shell contacts involving water or ions**   
**10%: Percentage of satisfied second shell Water contacts**   
• Contact(s): /O_w_/O_w_/O_w_/O_w_/O_w_/OP1/  
• Contact code: O_ph_5O_w_  
• Contact(s) in the exclusion zone: /O_w_/  
→ Slightly distorded coord. octahedron (5.2º) ←  
Go to ToC

### MG:A:1639

|  | Res. type | Atom | Residue | Distance | Occupancy | B-factor | Closest atom(s) to a 1^st^ shell water | EM\_dens. |
| --- | --- | --- | --- | --- | --- | --- | --- | --- |
| **Ion** | | | |  | 1.0 | 52.9 |  | 5.39 |
 1 | Water 1 | | A:4503 | 1.86 | 1.0 | 52.7 |  | 4.11 | 2 | Water 2 | | A:2984 | 2.00 | 1.0 | 43.8 | 2.74 [G:OP1]A:570 | 4.43 | 3 | Water 3 | | A:3386 | 2.10 | 1.0 | 41.9 | 2.82 [G:O6]A:570 | 4.94 | 4 | Water 4 | | A:3067 | 2.20 | 1.0 | 40.2 | 2.76 [G:N7]A:570 | 6.75 | 5 | Water 5 | | A:1962 | 2.48 | 1.0 | 42.1 | 2.45 [G:OP1]A:570 // 2.81 [U:O4]A:571 | 6.61 | 6 | C | OP1 | A:569 | 3.71 | 1.0 | 44.8 |  | 4.49 | 7 | G | OP1 | A:570 | 3.84 | 1.0 | 40.3 |  | 4.69 | 8 | G | N7 | A:570 | 3.97 | 1.0 | 33.8 |  | 10.43 | 9 | Water 6 | | A:4679 | 3.98 | 1.0 | 56.0 |  | 1.96 | 10 | Water 7 | | A:4507 | 4.02 | 1.0 | 44.0 |  | 4.12 | 11 | Water 8 | | A:3352 | 4.04 | 1.0 | 41.6 |  | 5.10 | 12 | Water 9 | | A:4575 | 4.21 | 1.0 | 51.7 |  | 1.54 | 13 | G | O6 | A:570 | 4.27 | 1.0 | 35.3 |  | 10.59 | 14 | Water 10 | | A:4179 | 4.39 | 1.0 | 42.1 |  | 5.78 | 15 | U | O4 | A:571 | 4.53 | 1.0 | 35.9 |  | 9.50 | 16 | Water 11 | | A:2391 | 4.78 | 1.0 | 47.9 |  | 4.00 | 17 | Water 12 | | A:2434 | 4.78 | 1.0 | 53.0 |  | 2.01 |

  
**1^st^ shell coordination angles and distances (*trans* in "blue"; *cys* in "green"; deviations in "orange"):**   
• 1-2(W/W): *112º*; 1-3(W/W): *74º*; 1-4(W/W): *161º*; 2-3(W/W): 170º; 2-4(W/W): 87º; 3-4(W/W): 88º;   
• 1-2(W/W): *3.2Å*; 1-3(W/W): 2.4Å; 1-4(W/W): 4.0Å; 2-3(W/W): 4.1Å; 2-4(W/W): 2.9Å; 3-4(W/W): 3.0Å;   
  
**2.39 ± 0.62: the average coodination distance of the 1^st^ six ligands is Na^+^** compatible  
**10: Number of potential Water contacts**   
**5 : Number of second shell contacts involving water or ions**   
**50%: Percentage of satisfied second shell Water contacts**   
• Contact(s): /O_w_/O_w_/O_w_/  
• Contact code: 3O_w_  
• Contact(s) in the exclusion zone: /O_w_/O_w_/  
→ Missing octahedral coordination for MG ←  
→→→ Distorded coord. octahedron (12.1º) ←←←  
Go to ToC

### MG:A:1640

|  | Res. type | Atom | Residue | Distance | Occupancy | B-factor | Closest atom(s) to a 1^st^ shell water | EM\_dens. |
| --- | --- | --- | --- | --- | --- | --- | --- | --- |
| **Ion** | | | |  | 1.0 | 51.6 |  | 7.91 |
 1 | C | OP1 | A:795 | 2.65 | 1.0 | 41.3 |  | 7.40 | 2 | C | OP2 | A:796 | 2.74 | 1.0 | 44.5 |  | 7.81 | 3 | LYS | NZ | K:126 | 2.81 | 1.0 | 38.6 |  | 6.59 | 4 | Water 1 | | A:3577 | 3.14 | 1.0 | 44.0 | 2.86 [C:OP1]A:795 | 4.41 | 5 | ARG | NH2 | K:129 | 3.24 | 1.0 | 40.4 |  | 7.79 | 6 | Water 2 | | K:321 | 3.65 | 1.0 | 41.9 |  | 5.36 | 7 | U | O3' | A:1522 | 3.70 | 1.0 | 35.5 |  | 9.88 | 8 | C | O5' | A:795 | 3.80 | 1.0 | 37.7 |  | 8.21 | 9 | Water 3 | | A:2709 | 3.99 | 1.0 | 43.6 | 2.69 [C:OP2]A:796 | 5.99 | 10 | G | OP1 | A:1523 | 4.09 | 1.0 | 34.2 |  | 8.98 | 11 | Water 4 | | A:2182 | 4.19 | 1.0 | 46.2 | 2.55 [C:OP1]A:796 | 4.62 | 12 | C | OP1 | A:796 | 4.45 | 1.0 | 37.0 |  | 9.28 | 13 | Water 5 | | A:4053 | 4.50 | 1.0 | 40.7 |  | 6.13 | 14 | C | O3' | A:795 | 4.55 | 1.0 | 37.7 |  | 10.46 |

  
• • **3.04 ± 0.35: the average coodination distance of the 1^st^ six ligands is K^+^ compatible**  
**4: Number of potential Water contacts**   
**1 : Number of second shell contacts involving water or ions**   
**25%: Percentage of satisfied second shell Water contacts**   
• Contact code: → No contacts ←   
• Contact(s) in the exclusion zone: /NH2_Arg_/NZ_Lys_/O_w_/OP1/OP2/  
→ Consider K in place of MG (3 valid contact(s) for K vs. 0 for MG ←)  
Go to ToC

### MG:A:1641

|  | Res. type | Atom | Residue | Distance | Occupancy | B-factor | Closest atom(s) to a 1^st^ shell water | EM\_dens. |
| --- | --- | --- | --- | --- | --- | --- | --- | --- |
| **Ion** | | | |  | 1.0 | 54.9 |  | 5.95 |
 1 | C | OP1 | A:811 | 2.60 | 1.0 | 38.7 |  | 7.06 | 2 | C | O2' | A:897 | 2.63 | 1.0 | 40.5 |  | 8.93 | 3 | G | OP2 | A:812 | 2.65 | 1.0 | 35.1 |  | 9.17 | 4 | Water 1 | | A:2820 | 3.14 | 1.0 | 45.2 | 2.71 [G:OP2]A:812 | 4.11 | 5 | Water 2 | | A:2577 | 3.30 | 1.0 | 40.7 | 2.66 [G:OP2]A:812 // 2.84 [C:O2]A:897 | 6.10 | 6 | C | O5' | A:811 | 3.69 | 1.0 | 36.0 |  | 9.39 | 7 | G | O4' | A:898 | 3.70 | 1.0 | 37.1 |  | 9.64 | 8 | Water 3 | | A:3297 | 3.77 | 1.0 | 49.4 | 2.80 [C:O2']A:897 | 2.95 | 9 | C | OP2 | A:811 | 4.47 | 1.0 | 35.7 |  | 7.29 | 10 | C | O3' | A:811 | 4.54 | 1.0 | 38.1 |  | 8.31 | 11 | Water 4 | | A:3839 | 4.63 | 1.0 | 41.8 | 2.92 [C:OP2]A:811 | 3.87 | 12 | C | O2 | A:897 | 4.74 | 1.0 | 34.7 |  | 8.63 | 13 | G | O5' | A:898 | 4.75 | 1.0 | 45.4 |  | 6.16 | 14 | C | O3' | A:897 | 4.77 | 1.0 | 40.1 |  | 6.84 |

  
• • **3.00 ± 0.41: the average coodination distance of the 1^st^ six ligands is K^+^ compatible**  
**4: Number of potential Water contacts**   
**3 : Number of second shell contacts involving water or ions**   
**75%: Percentage of satisfied second shell Water contacts**   
• Contact code: → No contacts ←   
• Contact(s) in the exclusion zone: /O2'/O_w_/O_w_/OP1/OP2/  
→ Consider K in place of MG (3 valid contact(s) for K vs. 0 for MG ←)  
Go to ToC

### MG:A:1642

|  | Res. type | Atom | Residue | Distance | Occupancy | B-factor | Closest atom(s) to a 1^st^ shell water | EM\_dens. |
| --- | --- | --- | --- | --- | --- | --- | --- | --- |
| **Ion** | | | |  | 1.0 | 61.0 |  | 3.73 |
 1 | Water 1 | | A:3892 | 1.94 | 1.0 | 51.4 |  | 2.78 | 2 | Water 2 | | A:3445 | 1.95 | 1.0 | 53.7 |  | 2.55 | 3 | Water 3 | | A:1846 | 1.98 | 1.0 | 48.5 | 2.33 [U:OP2]A:49 | 2.23 | 4 | C | OP2 | A:48 | 2.16 | 1.0 | 55.2 |  | 4.51 | 5 | Water 4 | | A:4145 | 2.23 | 1.0 | 52.6 |  | 2.37 | 6 | G | OP1 | A:115 | 2.24 | 1.0 | 48.7 |  | 4.62 | 7 | U | O3' | A:114 | 3.64 | 1.0 | 48.0 |  | 4.68 | 8 | C | O3' | A:47 | 3.88 | 1.0 | 55.4 |  | 4.29 | 9 | Water 5 | | A:4554 | 4.01 | 1.0 | 58.5 |  | 1.83 | 10 | U | OP2 | A:49 | 4.03 | 1.0 | 56.9 |  | 2.18 | 11 | Water 6 | | A:3968 | 4.24 | 1.0 | 59.6 | 2.96 [C:OP1]A:48 | 1.34 | 12 | G | OP2 | A:115 | 4.37 | 1.0 | 48.7 |  | 4.18 | 13 | C | OP1 | A:48 | 4.44 | 1.0 | 56.6 |  | 3.04 | 14 | C | O5' | A:48 | 4.51 | 1.0 | 56.0 |  | 3.64 | 15 | Water 7 | | A:1805 | 4.53 | 1.0 | 57.6 | 2.21 [C:OP1]A:48 | 1.83 | 16 | A | OP1 | A:51 | 4.56 | 1.0 | 53.8 |  | 1.84 | 17 | Water 8 | | A:4361 | 4.58 | 1.0 | 49.4 |  | 2.44 | 18 | G | O5' | A:115 | 4.62 | 1.0 | 45.2 |  | 6.14 |

  
**1^st^ shell coordination angles and distances (*trans* in "blue"; *cys* in "green"; deviations in "orange"):**   
• 1-2(W/W): 90º; 1-3(W/W): 90º; 1-4(W/OP2): 91º; 1-5(W/W): 86º; 1-6(W/OP1): *167º*; 2-3(W/W): 177º; 2-4(W/OP2): 87º; 2-5(W/W): 90º; 2-6(W/OP1): *98º*; 3-4(W/OP2): *96º*; 3-5(W/W): 87º; 3-6(W/OP1): *81º*; 4-5(OP2/W): 176º; 4-6(OP2/OP1): *98º*; 5-6(W/OP1): *85º*;   
• 1-2(W/W): 2.8Å; 1-3(W/W): 2.8Å; 1-4(W/OP2): 2.9Å; 1-5(W/W): 2.9Å; 1-6(W/OP1): 4.2Å; 2-3(W/W): 3.9Å; 2-4(W/OP2): 2.8Å; 2-5(W/W): 3.0Å; 2-6(W/OP1): 3.2Å; 3-4(W/OP2): 3.1Å; 3-5(W/W): 2.9Å; 3-6(W/OP1): 2.8Å; 4-5(OP2/W): *4.4Å*; 4-6(OP2/OP1): *3.3Å*; 5-6(W/OP1): 3.0Å;   
  
**2.09 ± 0.13: the average coodination distance of the 1^st^ six ligands is Mg^2+^ compatible**  
**8: Number of potential Water contacts**   
**1 : Number of second shell contacts involving water or ions**   
**12%: Percentage of satisfied second shell Water contacts**   
• Contact(s): /O_w_/O_w_/O_w_/O_w_/OP1/OP2/  
• Contact code: *cis-*2O_ph_4O_w_  
• MG hydration shell issue: water too close (< 2.4 Å.)  
Ooctahedral coordination (4.5º) →2Fo-Fc density between 1.0 and 4.0 !   
Go to ToC

### MG:A:1643

|  | Res. type | Atom | Residue | Distance | Occupancy | B-factor | Closest atom(s) to a 1^st^ shell water | EM\_dens. |
| --- | --- | --- | --- | --- | --- | --- | --- | --- |
| **Ion** | | | |  | 1.0 | 30.0 |  | 2.92 |
 1 | C | OP2 | A:536 | 2.37 | 1.0 | 63.8 |  | 3.81 | 2 | Water 1 | | A:4401 | 2.59 | 1.0 | 63.8 |  | 1.94 | 3 | Water 2 | | A:3593 | 3.33 | 1.0 | 72.5 |  | 1.68 | 4 | Water 3 | | A:4237 | 3.50 | 1.0 | 71.1 |  | 1.93 | 5 | C | O5' | A:536 | 3.89 | 1.0 | 65.3 |  | 3.65 | 6 | G | OP2 | A:537 | 3.95 | 1.0 | 69.0 |  | 2.91 | 7 | G | N7 | A:537 | 4.07 | 1.0 | 68.3 |  | 3.08 | 8 | Water 4 | | A:1835 | 4.21 | 1.0 | 65.2 | 2.31 [A:OP1]A:535 | 1.55 | 9 | C | OP1 | A:536 | 4.42 | 1.0 | 66.6 |  | 2.85 | 10 | A | OP1 | A:535 | 4.78 | 1.0 | 66.3 |  | 2.73 |

  
• • **3.27 ± 0.60: the average coodination distance of the 1^st^ six ligands is K^+^ compatible**  
**6: Number of potential Water contacts**   
**0 : Number of second shell contacts involving water or ions**   
**0%: Percentage of satisfied second shell Water contacts**   
• Contact code: → No contacts ←   
• Contact(s) in the exclusion zone: /O_w_/O_w_/OP2/  
→ Firts MG coordination shell missing ←  
→2Fo-Fc density between 1.0 and 4.0 !   
Go to ToC

### K:A:1644

|  | Res. type | Atom | Residue | Distance | Occupancy | B-factor | Closest atom(s) to a 1^st^ shell water | EM\_dens. |
| --- | --- | --- | --- | --- | --- | --- | --- | --- |
| **Ion** | | | |  | 1.0 | 30.0 |  | 4.65 |
 1 | G | N7 | A:1222 | 2.57 | 1.0 | 75.1 |  | 2.91 | 2 | C | OP1 | A:979 | 2.60 | 1.0 | 73.9 |  | 4.18 | 3 | Water 1 | | A:3746 | 2.92 | 1.0 | 75.2 |  | 2.52 | 4 | Water 2 | | A:2055 | 3.09 | 1.0 | 76.4 | 1.94 [MG:MG]A:1689 | 2.27 | 5 | C | N4 | A:1322 | 3.38 | 1.0 | 71.6 |  | 3.65 | 6 | **MG** | MG | A:1689 | 3.56 | 1.0 | 80.7 |  | 3.56 | 7 | Water 3 | | A:4181 | 3.66 | 1.0 | 75.0 |  | 1.70 | 8 | G | O6 | A:1222 | 3.66 | 1.0 | 75.2 |  | 2.35 | 9 | A | O3' | A:978 | 3.69 | 1.0 | 74.9 |  | 3.24 | 10 | Water 4 | | A:3305 | 3.92 | 1.0 | 78.0 | 2.80 [G:N7]A:1221 | 1.68 | 11 | G | N7 | A:1221 | 4.10 | 1.0 | 78.2 |  | 2.72 | 12 | C | O5' | A:979 | 4.28 | 1.0 | 75.1 |  | 2.30 | 13 | G | OP2 | A:1222 | 4.29 | 1.0 | 75.8 |  | 2.23 | 14 | A | O2' | A:978 | 4.79 | 1.0 | 76.4 |  | 2.42 |

  
**1^st^ shell coordination angles and distances (*trans* in "blue"; *cys* in "green"; deviations in "orange"):**   
• 1-2(N7/OP1): 109; 1-3(N7/W): 92; 1-4(N7/W): 79; 2-3(OP1/W): 79; 2-4(OP1/W): 62; 3-4(W/W): 133;   
• 1-2(N7/OP1): 4.2; 1-3(N7/W): 4.0; 1-4(N7/W): 3.6; 2-3(OP1/W): 3.5; 2-4(OP1/W): 2.9; 3-4(W/W): 5.5;   
  
**4.24 ± 1.32: the average coodination distance of the 1^st^ six ligands could correspond to an hexahydrated Mg^2+^.**  
**6: Number of potential Water contacts**   
**1 : Number of second shell contacts involving water or ions**   
**17%: Percentage of satisfied second shell Water contacts**   
• Contact(s): /O_w_/O_w_/  
• Contact code: 2O_w_  
• Contact(s) in the exclusion zone: /N7/OP1/  
→ Coordination < 6 for K ←  
Go to ToC

### K:A:1645

|  | Res. type | Atom | Residue | Distance | Occupancy | B-factor | Closest atom(s) to a 1^st^ shell water | EM\_dens. |
| --- | --- | --- | --- | --- | --- | --- | --- | --- |
| **Ion** | | | |  | 1.0 | 30.0 |  | 4.57 |
 1 | G | O6 | A:1206 | 2.61 | 1.0 | 69.8 |  | 2.26 | 2 | C | O2 | A:1200 | 2.71 | 1.0 | 65.7 |  | 2.69 | 3 | U | O4 | A:1052 | 2.71 | 1.0 | 70.3 |  | 2.39 | 4 | 2MG | O6 | A:1207 | 2.91 | 1.0 | 72.4 |  | 2.55 | 5 | U | N3 | A:1205 | 3.64 | 1.0 | 63.5 |  | 3.39 | 6 | C | N4 | A:1051 | 3.75 | 1.0 | 70.9 |  | 2.72 | 7 | G | N7 | A:1206 | 3.80 | 1.0 | 66.7 |  | 3.02 | 8 | U | N3 | A:1052 | 3.83 | 1.0 | 67.8 |  | 3.08 | 9 | U | O4 | A:1205 | 3.90 | 1.0 | 68.9 |  | 2.26 | 10 | C | N3 | A:1200 | 3.98 | 1.0 | 64.3 |  | 3.82 | 11 | 2MG | N1 | A:1207 | 4.37 | 1.0 | 71.3 |  | 3.89 | 12 | C | O2' | A:1200 | 4.39 | 1.0 | 63.8 |  | 2.27 | 13 | C | N1 | A:1200 | 4.41 | 1.0 | 63.4 |  | 3.73 | 14 | U | O2 | A:1205 | 4.59 | 1.0 | 67.0 |  | 2.66 | 15 | 2MG | N7 | A:1207 | 4.78 | 1.0 | 71.3 |  | 2.75 |

  
**1^st^ shell coordination angles and distances (*trans* in "blue"; *cys* in "green"; deviations in "orange"):**   
• 1-2(O6/O2): 94; 1-3(O6/O4): 80; 1-4(O6/O6): 71; 2-3(O2/O4): 72; 2-4(O2/O6): 160; 3-4(O4/O6): 91;   
• 1-2(O6/O2): 3.9; 1-3(O6/O4): 3.4; 1-4(O6/O6): 3.2; 2-3(O2/O4): 3.2; 2-4(O2/O6): 5.5; 3-4(O4/O6): 4.0;   
  
**4.33 ± 1.39: the average coodination distance of the 1^st^ six ligands could correspond to an hexahydrated Mg^2+^.**  
**0: Number of potential Water contacts**   
• Contact(s): /O2/O4/O6/O6/  
• Contact code: 4O_b_  
→ Coordination < 6 for K ←  
Go to ToC

### K:A:1646

|  | Res. type | Atom | Residue | Distance | Occupancy | B-factor | Closest atom(s) to a 1^st^ shell water | EM\_dens. |
| --- | --- | --- | --- | --- | --- | --- | --- | --- |
| **Ion** | | | |  | 1.0 | 30.0 |  | 10.44 |
 1 | G | O6 | A:1079 | 2.76 | 1.0 | 40.0 |  | 7.51 | 2 | Water 1 | | A:3289 | 2.76 | 1.0 | 47.5 |  | 3.98 | 3 | Water 2 | | A:2385 | 2.78 | 1.0 | 52.7 | 2.62 [G:O2']A:1077 | 3.25 | 4 | Water 3 | | A:3586 | 3.00 | 1.0 | 43.6 | 2.86 [G:O2']A:1077 | 5.09 | 5 | A | O3' | A:865 | 3.10 | 1.0 | 38.4 |  | 7.62 | 6 | C | OP1 | A:866 | 3.12 | 1.0 | 45.4 |  | 4.82 | 7 | G | O2' | A:1077 | 4.07 | 1.0 | 46.5 |  | 4.66 | 8 | Water 4 | | A:4602 | 4.31 | 1.0 | 49.0 |  | 1.58 | 9 | Water 5 | | A:3037 | 4.57 | 1.0 | 53.3 | 2.75 [C:OP2]A:866 | 1.72 | 10 | Water 6 | | A:2963 | 4.62 | 1.0 | 51.4 |  | 1.90 | 11 | C | OP2 | A:866 | 4.64 | 1.0 | 45.7 |  | 5.48 | 12 | A | O2' | A:865 | 4.80 | 1.0 | 36.5 |  | 8.83 |

  
**1^st^ shell coordination angles and distances (*trans* in "blue"; *cys* in "green"; deviations in "orange"):**   
• 1-2(O6/W): 82; 1-3(O6/W): 78; 1-4(O6/W): 74; 1-5(O6/O3'): 103; 1-6(O6/OP1): 150; 2-3(W/W): 84; 2-4(W/W): 153; 2-5(W/O3'): 106; 2-6(W/OP1): 106; 3-4(W/W): 79; 3-5(W/O3'): 170; 3-6(W/OP1): 131; 4-5(W/O3'): 92; 4-6(W/OP1): 101; 5-6(O3'/OP1): 47;   
• 1-2(O6/W): 3.6; 1-3(O6/W): 3.5; 1-4(O6/W): 3.5; 1-5(O6/O3'): 4.6; 1-6(O6/OP1): 5.7; 2-3(W/W): 3.7; 2-4(W/W): 5.6; 2-5(W/O3'): 4.7; 2-6(W/OP1): 4.7; 3-4(W/W): 3.7; 3-5(W/O3'): 5.9; 3-6(W/OP1): 5.4; 4-5(W/O3'): 4.4; 4-6(W/OP1): 4.7; 5-6(O3'/OP1): 2.5;   
  
**4.32 ± 1.41: the average coodination distance of the 1^st^ six ligands could correspond to an hexahydrated Mg^2+^.**  
**8: Number of potential Water contacts**   
**2 : Number of second shell contacts involving water or ions**   
**25%: Percentage of satisfied second shell Water contacts**   
• Contact(s): /O3'/O6/O_w_/O_w_/O_w_/OP1/  
• Contact code: O_ph_O_r_O_b_3O_w_  
Go to ToC

### K:A:1647

|  | Res. type | Atom | Residue | Distance | Occupancy | B-factor | Closest atom(s) to a 1^st^ shell water | EM\_dens. |
| --- | --- | --- | --- | --- | --- | --- | --- | --- |
| **Ion** | | | |  | 1.0 | 30.0 |  | 3.14 |
 1 | G | O6 | A:976 | 2.21 | 1.0 | 67.4 |  | 2.26 | 2 | C | O2 | A:1359 | 2.29 | 1.0 | 70.1 |  | 2.21 | 3 | A | OP2 | A:1362 | 3.53 | 1.0 | 72.6 |  | 1.63 | 4 | G | OP2 | A:1361 | 3.91 | 1.0 | 76.2 |  | 1.16 | 5 | A | N7 | A:1362 | 3.98 | 1.0 | 68.0 |  | 3.24 | 6 | Water 1 | | A:1959 | 4.17 | 1.0 | 66.7 | 2.51 [G:N1]A:976 | 1.78 | 7 | G | N1 | A:976 | 4.21 | 1.0 | 65.3 |  | 3.69 | 8 | C | N3 | A:1359 | 4.27 | 1.0 | 66.2 |  | 3.77 | 9 | Water 2 | | A:2338 | 4.36 | 1.0 | 68.1 | 2.61 [C:N3]A:1359 | 2.56 | 10 | C | N1 | A:1359 | 4.44 | 1.0 | 64.8 |  | 4.54 | 11 | C | O2' | A:1359 | 4.56 | 1.0 | 70.1 |  | 2.30 | 12 | U | O4 | A:1358 | 4.75 | 1.0 | 69.1 |  | 3.04 |

  
**1^st^ shell coordination angles and distances (*trans* in "blue"; *cys* in "green"; deviations in "orange"):**   
• 1-2(O6/O2): 88;   
• 1-2(O6/O2): 3.1;   
  
**4.76 ± 1.65: the average coodination distance of the 1^st^ six ligands could correspond to an hexahydrated Mg^2+^.**  
**2: Number of potential Water contacts**   
**1 : Number of second shell contacts involving water or ions**   
**50%: Percentage of satisfied second shell Water contacts**   
• Contact code: → No contacts ←   
• Contact(s) in the exclusion zone: /O2/O6/  
→ Consider MG in place of K (2 valid contact(s) for MG vs. 0 for K ←)  
→2Fo-Fc density between 1.0 and 4.0 !   
Go to ToC

### K:A:1648

|  | Res. type | Atom | Residue | Distance | Occupancy | B-factor | Closest atom(s) to a 1^st^ shell water | EM\_dens. |
| --- | --- | --- | --- | --- | --- | --- | --- | --- |
| **Ion** | | | |  | 1.0 | 30.0 |  | 3.87 |
 1 | C | O2' | A:1322 | 2.66 | 1.0 | 75.0 |  | 1.76 | 2 | C | O2 | A:1322 | 2.90 | 1.0 | 73.0 |  | 2.96 | 3 | A | N3 | A:1319 | 3.05 | 1.0 | 79.4 |  | 2.55 | 4 | Water 1 | | A:2068 | 3.09 | 1.0 | 77.7 | 2.51 [A:O4']A:978 | 1.79 | 5 | C | O3' | A:1322 | 3.45 | 1.0 | 71.9 |  | 3.84 | 6 | Water 2 | | A:1937 | 3.81 | 1.0 | 77.9 | 2.42 [U:OP2]A:1321 // 2.91 [G:OP2]A:1323 | 1.62 | 7 | G | O4' | A:1323 | 3.85 | 1.0 | 76.8 |  | 2.59 | 8 | A | O2' | A:1319 | 3.90 | 1.0 | 81.5 |  | 1.24 | 9 | C | N1 | A:1322 | 4.32 | 1.0 | 70.3 |  | 4.49 | 10 | G | O5' | A:1323 | 4.35 | 1.0 | 77.3 |  | 2.57 | 11 | A | O4' | A:978 | 4.39 | 1.0 | 77.0 |  | 2.52 | 12 | A | N3 | A:978 | 4.54 | 1.0 | 76.0 |  | 2.63 | 13 | G | OP2 | A:1323 | 4.56 | 1.0 | 78.8 |  | 1.29 | 14 | U | OP2 | A:1321 | 4.67 | 1.0 | 75.7 |  | 2.13 |

  
**1^st^ shell coordination angles and distances (*trans* in "blue"; *cys* in "green"; deviations in "orange"):**   
• 1-2(O2'/O2): 71; 1-3(O2'/N3): 171; 1-4(O2'/W): 108; 2-3(O2/N3): 112; 2-4(O2/W): 57; 3-4(N3/W): 80;   
• 1-2(O2'/O2): 3.2; 1-3(O2'/N3): 5.7; 1-4(O2'/W): 4.7; 2-3(O2/N3): 4.9; 2-4(O2/W): 2.8; 3-4(N3/W): 3.9;   
  
**4.45 ± 1.39: the average coodination distance of the 1^st^ six ligands could correspond to an hexahydrated Mg^2+^.**  
**4: Number of potential Water contacts**   
**3 : Number of second shell contacts involving water or ions**   
**75%: Percentage of satisfied second shell Water contacts**   
• Contact(s): /N3_A_/O2/O2'/O_w_/  
• Contact code: O_r_O_b_N_b_O_w_  
→ Coordination < 6 for K ←  
→2Fo-Fc density between 1.0 and 4.0 !   
Go to ToC

### K:A:1649

|  | Res. type | Atom | Residue | Distance | Occupancy | B-factor | Closest atom(s) to a 1^st^ shell water | EM\_dens. |
| --- | --- | --- | --- | --- | --- | --- | --- | --- |
| **Ion** | | | |  | 1.0 | 30.0 |  | 3.96 |
 1 | Water 1 | | A:4498 | 2.68 | 1.0 | 61.3 |  | 1.76 | 2 | U | O4 | A:793 | 2.69 | 1.0 | 54.6 |  | 2.40 | 3 | G | O4' | A:1517 | 2.84 | 1.0 | 49.6 |  | 4.99 | 4 | C | OP1 | a:1924 | 3.08 | 1.0 | 53.9 |  | 3.92 | 5 | Water 2 | | A:2091 | 3.64 | 1.0 | 51.9 | 2.52 [G:OP1]A:1517 | 1.63 | 6 | C | OP2 | a:1924 | 3.98 | 1.0 | 53.4 |  | 3.58 | 7 | G | OP1 | A:1517 | 4.03 | 1.0 | 47.5 |  | 3.42 | 8 | G | N9 | A:1517 | 4.22 | 1.0 | 38.2 |  | 7.44 | 9 | G | O5' | A:1517 | 4.42 | 1.0 | 50.7 |  | 2.44 | 10 | A | O3' | a:1923 | 4.45 | 1.0 | 52.2 |  | 4.99 | 11 | Water 3 | | A:2372 | 4.65 | 1.0 | 50.5 | 2.62 [G:OP1]A:1517 | 1.86 |

  
**1^st^ shell coordination angles and distances (*trans* in "blue"; *cys* in "green"; deviations in "orange"):**   
• 1-2(W/O4): 154; 1-3(W/O4'): 71; 1-4(W/OP1): 103; 2-3(O4/O4'): 94; 2-4(O4/OP1): 93; 3-4(O4'/OP1): 71;   
• 1-2(W/O4): 5.2; 1-3(W/O4'): 3.2; 1-4(W/OP1): 4.5; 2-3(O4/O4'): 4.1; 2-4(O4/OP1): 4.2; 3-4(O4'/OP1): 3.4;   
  
**4.52 ± 1.48: the average coodination distance of the 1^st^ six ligands could correspond to an hexahydrated Mg^2+^.**  
**4: Number of potential Water contacts**   
**1 : Number of second shell contacts involving water or ions**   
**25%: Percentage of satisfied second shell Water contacts**   
• Contact(s): /O4/O4'/O_w_/OP1/  
• Contact code: O_ph_O_r_O_b_O_w_  
→ Coordination < 6 for K ←  
→2Fo-Fc density between 1.0 and 4.0 !   
Go to ToC

### K:A:1650

|  | Res. type | Atom | Residue | Distance | Occupancy | B-factor | Closest atom(s) to a 1^st^ shell water | EM\_dens. |
| --- | --- | --- | --- | --- | --- | --- | --- | --- |
| **Ion** | | | |  | 1.0 | 30.0 |  | 12.02 |
 1 | Water 1 | | A:2022 | 2.82 | 1.0 | 44.0 | 2.48 [G:N7]A:22 | 4.68 | 2 | U | O4 | A:12 | 2.86 | 1.0 | 41.2 |  | 6.89 | 3 | G | O6 | A:22 | 2.94 | 1.0 | 41.0 |  | 7.74 | 4 | G | O6 | A:11 | 2.97 | 1.0 | 44.6 |  | 6.89 | 5 | G | O6 | A:21 | 3.11 | 1.0 | 41.3 |  | 6.23 | 6 | Water 2 | | A:4533 | 3.22 | 1.0 | 51.0 |  | 3.71 | 7 | Water 3 | | A:3998 | 3.37 | 1.0 | 46.4 |  | 2.87 | 8 | C | N4 | A:23 | 3.64 | 1.0 | 30.6 |  | 8.32 | 9 | Water 4 | | A:4545 | 3.96 | 1.0 | 53.8 |  | 1.66 | 10 | G | N7 | A:22 | 4.17 | 1.0 | 33.0 |  | 8.50 | 11 | Water 5 | | A:2216 | 4.18 | 1.0 | 52.0 |  | 2.71 | 12 | Water 6 | | A:4676 | 4.25 | 1.0 | 56.8 |  | 2.34 | 13 | Water 7 | | A:2838 | 4.51 | 1.0 | 47.5 | 2.72 [G:N1]A:21 | 3.83 | 14 | Water 8 | | A:4542 | 4.59 | 1.0 | 51.9 |  | 2.12 |

  
**1^st^ shell coordination angles and distances (*trans* in "blue"; *cys* in "green"; deviations in "orange"):**   
• 1-2(W/O4): 158; 1-3(W/O6): 83; 1-4(W/O6): 115; 1-5(W/O6): 81; 2-3(O4/O6): 77; 2-4(O4/O6): 69; 2-5(O4/O6): 89; 3-4(O6/O6): 77; 3-5(O6/O6): 87; 4-5(O6/O6): 155;   
• 1-2(W/O4): 5.6; 1-3(W/O6): 3.8; 1-4(W/O6): 4.9; 1-5(W/O6): 3.9; 2-3(O4/O6): 3.6; 2-4(O4/O6): 3.3; 2-5(O4/O6): 4.2; 3-4(O6/O6): 3.7; 3-5(O6/O6): 4.2; 4-5(O6/O6): 5.9;   
  
**4.16 ± 1.24: the average coodination distance of the 1^st^ six ligands could correspond to an hexahydrated Mg^2+^.**  
**6: Number of potential Water contacts**   
**1 : Number of second shell contacts involving water or ions**   
**17%: Percentage of satisfied second shell Water contacts**   
• Contact(s): /O4/O6/O6/O6/O_w_/  
• Contact code: 4O_b_O_w_  
→ Coordination < 6 for K ←  
Go to ToC

### K:A:1651

|  | Res. type | Atom | Residue | Distance | Occupancy | B-factor | Closest atom(s) to a 1^st^ shell water | EM\_dens. |
| --- | --- | --- | --- | --- | --- | --- | --- | --- |
| **Ion** | | | |  | 1.0 | 30.0 |  | 2.28 |
 1 | Water 1 | | A:4006 | 2.90 | 1.0 | 75.6 |  | 1.58 | 2 | U | O2 | A:531 | 2.92 | 1.0 | 78.1 |  | 2.28 | 3 | G | O6 | A:517 | 2.94 | 1.0 | 77.0 |  | 2.31 | 4 | Water 2 | | A:1811 | 4.12 | 1.0 | 77.0 |  | 1.91 | 5 | PSU | OP2 | A:516 | 4.14 | 1.0 | 76.4 |  | 3.32 | 6 | U | N3 | A:531 | 4.16 | 1.0 | 80.0 |  | 2.20 | 7 | G | OP2 | A:515 | 4.42 | 1.0 | 74.1 |  | 2.08 | 8 | G | N7 | A:517 | 4.51 | 1.0 | 76.8 |  | 3.01 |

  
**1^st^ shell coordination angles and distances (*trans* in "blue"; *cys* in "green"; deviations in "orange"):**   
• 1-2(W/O2): 136; 1-3(W/O6): 67; 2-3(O2/O6): 73;   
• 1-2(W/O2): 5.4; 1-3(W/O6): 3.2; 2-3(O2/O6): 3.5;   
  
**5.02 ± 1.64: the average coodination distance of the 1^st^ six ligands could correspond to an hexahydrated Mg^2+^.**  
**4: Number of potential Water contacts**   
**0 : Number of second shell contacts involving water or ions**   
**0%: Percentage of satisfied second shell Water contacts**   
• Contact(s): /O2/O6/O_w_/  
• Contact code: 2O_b_O_w_  
→ Coordination < 6 for K ←  
→2Fo-Fc density between 1.0 and 4.0 !   
Go to ToC

### K:A:1652

|  | Res. type | Atom | Residue | Distance | Occupancy | B-factor | Closest atom(s) to a 1^st^ shell water | EM\_dens. |
| --- | --- | --- | --- | --- | --- | --- | --- | --- |
| **Ion** | | | |  | 1.0 | 30.0 |  | 13.71 |
 1 | C | OP1 | A:1501 | 2.52 | 1.0 | 38.4 |  | 9.86 | 2 | Water 1 | | A:2297 | 2.55 | 1.0 | 47.4 | 2.59 [G:OP2]A:1504 | 4.14 | 3 | Water 2 | | A:2417 | 2.77 | 1.0 | 47.5 |  | 5.10 | 4 | A | O2' | A:1507 | 2.79 | 1.0 | 40.3 |  | 8.76 | 5 | Water 3 | | A:2441 | 3.02 | 1.0 | 48.5 | 2.63 [A:OP1]A:1502 | 3.35 | 6 | Water 4 | | A:4215 | 3.04 | 1.0 | 48.7 |  | 2.30 | 7 | A | O4' | A:1508 | 3.11 | 1.0 | 37.7 |  | 7.52 | 8 | Water 5 | | A:1803 | 3.84 | 1.0 | 54.1 |  | 2.28 | 9 | G | OP2 | A:1504 | 3.91 | 1.0 | 43.8 |  | 7.64 | 10 | A | OP2 | A:1502 | 4.01 | 1.0 | 44.2 |  | 5.43 | 11 | Water 6 | | A:3543 | 4.37 | 1.0 | 49.3 |  | 2.79 | 12 | C | O5' | A:1501 | 4.56 | 1.0 | 34.0 |  | 8.18 | 13 | A | OP1 | A:1502 | 4.61 | 1.0 | 41.1 |  | 5.94 | 14 | C | OP2 | A:1501 | 4.66 | 1.0 | 36.7 |  | 9.87 | 15 | A | O2' | A:1508 | 4.79 | 1.0 | 40.6 |  | 6.68 |

  
**1^st^ shell coordination angles and distances (*trans* in "blue"; *cys* in "green"; deviations in "orange"):**   
• 1-2(OP1/W): 80; 1-3(OP1/W): 149; 1-4(OP1/O2'): 102; 1-5(OP1/W): 118; 1-6(OP1/W): 89; 1-7(OP1/O4'): 78; 2-3(W/W): 130; 2-4(W/O2'): 83; 2-5(W/W): 68; 2-6(W/W): 110; 2-7(W/O4'): 140; 3-4(W/O2'): 80; 3-5(W/W): 84; 3-6(W/W): 85; 3-7(W/O4'): 73; 4-5(O2'/W): 124; 4-6(O2'/W): 165; 4-7(O2'/O4'): 69; 5-6(W/W): 56; 5-7(W/O4'): 152; 6-7(W/O4'): 104;   
• 1-2(OP1/W): 3.3; 1-3(OP1/W): 5.1; 1-4(OP1/O2'): 4.1; 1-5(OP1/W): 4.8; 1-6(OP1/W): 3.9; 1-7(OP1/O4'): 3.6; 2-3(W/W): 4.8; 2-4(W/O2'): 3.5; 2-5(W/W): 3.1; 2-6(W/W): 4.6; 2-7(W/O4'): 5.3; 3-4(W/O2'): 3.6; 3-5(W/W): 3.9; 3-6(W/W): 3.9; 3-7(W/O4'): 3.5; 4-5(O2'/W): 5.1; 4-6(O2'/W): 5.8; 4-7(O2'/O4'): 3.3; 5-6(W/W): 2.9; 5-7(W/O4'): 5.9; 6-7(W/O4'): 4.8;   
  
**3.94 ± 1.22: the average coodination distance of the 1^st^ six ligands could correspond to an hexahydrated Mg^2+^.**  
**10: Number of potential Water contacts**   
**2 : Number of second shell contacts involving water or ions**   
**20%: Percentage of satisfied second shell Water contacts**   
• Contact(s): /O2'/O4'/O_w_/O_w_/O_w_/  
• Contact code: 2O_r_3O_w_  
• Contact(s) in the exclusion zone: /O_w_/OP1/  
→ Coordination < 6 for K ←  
Go to ToC

### K:A:1653

|  | Res. type | Atom | Residue | Distance | Occupancy | B-factor | Closest atom(s) to a 1^st^ shell water | EM\_dens. |
| --- | --- | --- | --- | --- | --- | --- | --- | --- |
| **Ion** | | | |  | 1.0 | 30.0 |  | 8.54 |
 1 | G | O6 | A:1074 | 2.79 | 1.0 | 51.9 |  | 4.73 | 2 | U | O4 | A:1073 | 2.89 | 1.0 | 53.8 |  | 3.68 | 3 | U | O4 | A:1083 | 2.92 | 1.0 | 45.7 |  | 5.41 | 4 | Water 1 | | A:4219 | 3.04 | 1.0 | 55.1 |  | 2.81 | 5 | Water 2 | | A:4014 | 3.27 | 1.0 | 54.3 |  | 4.13 | 6 | A | N6 | A:1082 | 3.65 | 1.0 | 36.1 |  | 6.89 | 7 | G | N2 | A:1084 | 3.81 | 1.0 | 44.7 |  | 5.39 | 8 | U | N3 | A:1083 | 3.84 | 1.0 | 37.1 |  | 7.78 | 9 | G | N7 | A:1074 | 4.23 | 1.0 | 46.4 |  | 5.75 | 10 | Water 3 | | A:3155 | 4.35 | 1.0 | 53.0 | 2.77 [G:N1]A:1084 | 4.08 | 11 | Water 4 | | A:3212 | 4.42 | 1.0 | 45.0 |  | 3.88 | 12 | U | O4 | A:1075 | 4.61 | 1.0 | 46.4 |  | 5.10 |

  
**1^st^ shell coordination angles and distances (*trans* in "blue"; *cys* in "green"; deviations in "orange"):**   
• 1-2(O6/O4): 75; 1-3(O6/O4): 83; 1-4(O6/W): 83; 2-3(O4/O4): 119; 2-4(O4/W): 84; 3-4(O4/W): 148;   
• 1-2(O6/O4): 3.4; 1-3(O6/O4): 3.8; 1-4(O6/W): 3.9; 2-3(O4/O4): 5.0; 2-4(O4/W): 4.0; 3-4(O4/W): 5.7;   
  
**4.37 ± 1.34: the average coodination distance of the 1^st^ six ligands could correspond to an hexahydrated Mg^2+^.**  
**4: Number of potential Water contacts**   
**0 : Number of second shell contacts involving water or ions**   
**0%: Percentage of satisfied second shell Water contacts**   
• Contact(s): /O4/O4/O6/O_w_/  
• Contact code: 3O_b_O_w_  
→ Coordination < 6 for K ←  
Go to ToC

### K:A:1654

|  | Res. type | Atom | Residue | Distance | Occupancy | B-factor | Closest atom(s) to a 1^st^ shell water | EM\_dens. |
| --- | --- | --- | --- | --- | --- | --- | --- | --- |
| **Ion** | | | |  | 1.0 | 30.0 |  | 8.14 |
 1 | A | N1 | A:780 | 2.84 | 1.0 | 32.8 |  | 11.77 | 2 | Water 1 | | A:4433 | 2.85 | 1.0 | 46.2 |  | 3.88 | 3 | Water 2 | | A:4597 | 2.90 | 1.0 | 44.2 |  | 5.85 | 4 | A | OP2 | A:802 | 3.19 | 1.0 | 46.9 |  | 6.03 | 5 | G | O6 | A:803 | 3.28 | 1.0 | 35.4 |  | 9.51 | 6 | U | OP2 | A:801 | 3.44 | 1.0 | 45.4 |  | 5.05 | 7 | G | N2 | A:776 | 3.52 | 1.0 | 38.2 |  | 7.13 | 8 | U | O5' | A:801 | 3.60 | 1.0 | 45.7 |  | 4.31 | 9 | Water 3 | | A:2431 | 3.66 | 1.0 | 42.7 | 2.63 [U:OP2]A:801 | 3.95 | 10 | U | O3' | A:801 | 4.44 | 1.0 | 42.9 |  | 4.85 | 11 | Water 4 | | A:4536 | 4.51 | 1.0 | 45.4 |  | 5.10 | 12 | Water 5 | | A:3626 | 4.65 | 1.0 | 45.1 |  | 3.02 | 13 | U | OP1 | A:801 | 4.77 | 1.0 | 44.2 |  | 3.79 | 14 | Water 6 | | A:2931 | 4.79 | 1.0 | 36.7 |  | 6.81 | 15 | A | N3 | A:780 | 4.79 | 1.0 | 33.8 |  | 9.75 |

  
**1^st^ shell coordination angles and distances (*trans* in "blue"; *cys* in "green"; deviations in "orange"):**   
• 1-2(N1/W): 140; 1-3(N1/W): 117; 1-4(N1/OP2): 118; 2-3(W/W): 70; 2-4(W/OP2): 89; 3-4(W/OP2): 116;   
• 1-2(N1/W): 5.3; 1-3(N1/W): 4.9; 1-4(N1/OP2): 5.2; 2-3(W/W): 3.3; 2-4(W/OP2): 4.2; 3-4(W/OP2): 5.2;   
  
**4.27 ± 1.28: the average coodination distance of the 1^st^ six ligands could correspond to an hexahydrated Mg^2+^.**  
**4: Number of potential Water contacts**   
**0 : Number of second shell contacts involving water or ions**   
**0%: Percentage of satisfied second shell Water contacts**   
• Contact(s): /N1_A_/O_w_/O_w_/OP2/  
• Contact code: O_ph_N_b_2O_w_  
→ Coordination < 6 for K ←  
Go to ToC

### K:A:1655

|  | Res. type | Atom | Residue | Distance | Occupancy | B-factor | Closest atom(s) to a 1^st^ shell water | EM\_dens. |
| --- | --- | --- | --- | --- | --- | --- | --- | --- |
| **Ion** | | | |  | 1.0 | 30.0 |  | 4.12 |
 1 | Water 1 | | A:3689 | 2.33 | 1.0 | 60.8 |  | 2.36 | 2 | G | O6 | A:852 | 2.71 | 1.0 | 54.0 |  | 5.08 | 3 | Water 2 | | A:2440 | 2.88 | 1.0 | 54.7 | 2.63 [G:N7]A:852 | 2.52 | 4 | U | O4 | A:834 | 3.06 | 1.0 | 50.0 |  | 4.36 | 5 | G | O6 | A:851 | 3.19 | 1.0 | 57.0 |  | 4.50 | 6 | Water 3 | | A:2064 | 3.30 | 1.0 | 66.7 | 2.51 [G:N7]A:851 | 2.17 | 7 | U | O4 | A:835 | 3.38 | 1.0 | 52.9 |  | 4.55 | 8 | U | N3 | A:834 | 3.91 | 1.0 | 44.8 |  | 5.40 | 9 | Water 4 | | A:2897 | 4.02 | 1.0 | 55.9 | 2.73 [G:O6]A:833 | 2.01 | 10 | G | O6 | A:833 | 4.10 | 1.0 | 51.2 |  | 4.48 | 11 | G | N7 | A:852 | 4.24 | 1.0 | 51.0 |  | 5.79 | 12 | Water 5 | | A:2582 | 4.43 | 1.0 | 51.8 |  | 1.80 | 13 | G | N7 | A:851 | 4.61 | 1.0 | 56.2 |  | 4.60 |

  
**1^st^ shell coordination angles and distances (*trans* in "blue"; *cys* in "green"; deviations in "orange"):**   
• 1-2(W/O6): 130; 1-3(W/W): 104; 1-4(W/O4): 105; 1-5(W/O6): 61; 2-3(O6/W): 81; 2-4(O6/O4): 82; 2-5(O6/O6): 72; 3-4(W/O4): 151; 3-5(W/O6): 79; 4-5(O4/O6): 117;   
• 1-2(W/O6): 4.6; 1-3(W/W): 4.1; 1-4(W/O4): 4.3; 1-5(W/O6): 2.9; 2-3(O6/W): 3.6; 2-4(O6/O4): 3.8; 2-5(O6/O6): 3.5; 3-4(W/O4): 5.8; 3-5(W/O6): 3.9; 4-5(O4/O6): 5.3;   
  
**4.13 ± 1.30: the average coodination distance of the 1^st^ six ligands could correspond to an hexahydrated Mg^2+^.**  
**6: Number of potential Water contacts**   
**2 : Number of second shell contacts involving water or ions**   
**33%: Percentage of satisfied second shell Water contacts**   
• Contact(s): /O4/O6/O6/O_w_/  
• Contact code: 3O_b_O_w_  
• Contact(s) in the exclusion zone: /O_w_/  
→ Coordination < 6 for K ←  
Go to ToC

### K:A:1656

|  | Res. type | Atom | Residue | Distance | Occupancy | B-factor | Closest atom(s) to a 1^st^ shell water | EM\_dens. |
| --- | --- | --- | --- | --- | --- | --- | --- | --- |
| **Ion** | | | |  | 1.0 | 30.0 |  | 3.10 |
 1 | G | O6 | A:849 | 2.86 | 1.0 | 80.5 |  | 2.14 | 2 | G | O6 | A:836 | 3.04 | 1.0 | 62.7 |  | 3.09 | 3 | U | O4 | A:837 | 3.04 | 1.0 | 66.1 |  | 2.24 | 4 | U | O4 | A:850 | 3.10 | 1.0 | 70.6 |  | 2.82 | 5 | Water 1 | | A:3355 | 3.21 | 1.0 | 63.4 | 2.81 [G:N7]A:836 | 2.08 | 6 | Water 2 | | A:3689 | 3.72 | 1.0 | 60.8 |  | 2.36 | 7 | U | N3 | A:837 | 3.76 | 1.0 | 62.7 |  | 4.43 | 8 | U | N3 | A:850 | 3.97 | 1.0 | 67.2 |  | 4.49 | 9 | G | N7 | A:836 | 4.48 | 1.0 | 56.2 |  | 4.10 | 10 | G | N7 | A:849 | 4.53 | 1.0 | 79.4 |  | 2.61 | 11 | Water 3 | | A:2064 | 4.59 | 1.0 | 66.7 |  | 2.17 |

  
**1^st^ shell coordination angles and distances (*trans* in "blue"; *cys* in "green"; deviations in "orange"):**   
• 1-2(O6/O6): 73; 1-3(O6/O4): 77; 1-4(O6/O4): 77; 2-3(O6/O4): 81; 2-4(O6/O4): 70; 3-4(O4/O4): 145;   
• 1-2(O6/O6): 3.5; 1-3(O6/O4): 3.7; 1-4(O6/O4): 3.7; 2-3(O6/O4): 3.9; 2-4(O6/O4): 3.5; 3-4(O4/O4): 5.9;   
  
**4.45 ± 1.36: the average coodination distance of the 1^st^ six ligands could correspond to an hexahydrated Mg^2+^.**  
**4: Number of potential Water contacts**   
**1 : Number of second shell contacts involving water or ions**   
**25%: Percentage of satisfied second shell Water contacts**   
• Contact(s): /O4/O4/O6/O6/  
• Contact code: 4O_b_  
→ Coordination < 6 for K ←  
→2Fo-Fc density between 1.0 and 4.0 !   
Go to ToC

### K:A:1657

|  | Res. type | Atom | Residue | Distance | Occupancy | B-factor | Closest atom(s) to a 1^st^ shell water | EM\_dens. |
| --- | --- | --- | --- | --- | --- | --- | --- | --- |
| **Ion** | | | |  | 1.0 | 30.0 |  | 2.41 |
 1 | G | O6 | A:1304 | 2.72 | 1.0 | 80.3 |  | 1.53 | 2 | G | O6 | A:1305 | 2.78 | 1.0 | 77.5 |  | 2.18 | 3 | C | OP2 | A:1303 | 2.85 | 1.0 | 77.3 |  | 3.57 | 4 | Water 1 | | A:2133 | 3.12 | 1.0 | 78.9 |  | 2.10 | 5 | G | N7 | A:1304 | 3.16 | 1.0 | 77.2 |  | 3.65 | 6 | G | N1 | A:1305 | 4.39 | 1.0 | 74.7 |  | 3.46 | 7 | C | O5' | A:1303 | 4.59 | 1.0 | 81.1 |  | 2.90 | 8 | C | N4 | A:1303 | 4.62 | 1.0 | 76.8 |  | 2.69 |

  
**1^st^ shell coordination angles and distances (*trans* in "blue"; *cys* in "green"; deviations in "orange"):**   
• 1-2(O6/O6): 69; 1-3(O6/OP2): 152; 1-4(O6/W): 104; 1-5(O6/N7): 62; 2-3(O6/OP2): 102; 2-4(O6/W): 50; 2-5(O6/N7): 64; 3-4(OP2/W): 55; 3-5(OP2/N7): 90; 4-5(W/N7): 53;   
• 1-2(O6/O6): 3.1; 1-3(O6/OP2): 5.4; 1-4(O6/W): 4.6; 1-5(O6/N7): 3.1; 2-3(O6/OP2): 4.4; 2-4(O6/W): 2.5; 2-5(O6/N7): 3.2; 3-4(OP2/W): 2.8; 3-5(OP2/N7): 4.2; 4-5(W/N7): 2.8;   
  
**4.71 ± 1.64: the average coodination distance of the 1^st^ six ligands could correspond to an hexahydrated Mg^2+^.**  
**2: Number of potential Water contacts**   
**0 : Number of second shell contacts involving water or ions**   
**0%: Percentage of satisfied second shell Water contacts**   
• Contact(s): /N7/O6/O6/O_w_/OP2/  
• Contact code: O_ph_2O_b_N_b_O_w_  
→ Coordination < 6 for K ←  
→2Fo-Fc density between 1.0 and 4.0 !   
Go to ToC

### K:A:1658

|  | Res. type | Atom | Residue | Distance | Occupancy | B-factor | Closest atom(s) to a 1^st^ shell water | EM\_dens. |
| --- | --- | --- | --- | --- | --- | --- | --- | --- |
| **Ion** | | | |  | 1.0 | 30.0 |  | 3.45 |
 1 | Water 1 | | A:2233 | 2.66 | 1.0 | 63.8 | 2.57 [U:O2']A:943 | 1.87 | 2 | G | OP1 | A:944 | 2.89 | 1.0 | 62.4 |  | 4.22 | 3 | G | N7 | A:1233 | 2.89 | 1.0 | 63.2 |  | 3.42 | 4 | Water 2 | | A:4238 | 3.12 | 1.0 | 65.6 |  | 2.44 | 5 | U | O3' | A:943 | 3.15 | 1.0 | 60.3 |  | 4.70 | 6 | U | O4 | A:1232 | 3.88 | 1.0 | 61.6 |  | 3.18 | 7 | G | O6 | A:1233 | 3.93 | 1.0 | 63.2 |  | 3.13 | 8 | Water 3 | | A:3759 | 3.98 | 1.0 | 68.7 | 2.05 [MG:MG]A:1691 | 2.24 | 9 | Water 4 | | A:2995 | 4.25 | 1.0 | 69.5 | 1.96 [MG:MG]A:1691 | 1.89 | 10 | **MG** | MG | A:1691 | 4.32 | 1.0 | 73.6 |  | 3.39 | 11 | U | O2' | A:943 | 4.59 | 1.0 | 59.3 |  | 3.72 | 12 | G | O5' | A:944 | 4.67 | 1.0 | 64.9 |  | 3.65 | 13 | G | OP2 | A:944 | 4.68 | 1.0 | 62.7 |  | 3.56 |

  
**1^st^ shell coordination angles and distances (*trans* in "blue"; *cys* in "green"; deviations in "orange"):**   
• 1-2(W/OP1): 96; 1-3(W/N7): 120; 1-4(W/W): 114; 1-5(W/O3'): 64; 2-3(OP1/N7): 144; 2-4(OP1/W): 70; 2-5(OP1/O3'): 49; 3-4(N7/W): 89; 3-5(N7/O3'): 140; 4-5(W/O3'): 58;   
• 1-2(W/OP1): 4.1; 1-3(W/N7): 4.8; 1-4(W/W): 4.8; 1-5(W/O3'): 3.1; 2-3(OP1/N7): 5.5; 2-4(OP1/W): 3.5; 2-5(OP1/O3'): 2.5; 3-4(N7/W): 4.2; 3-5(N7/O3'): 5.7; 4-5(W/O3'): 3.0;   
  
**4.41 ± 1.40: the average coodination distance of the 1^st^ six ligands could correspond to an hexahydrated Mg^2+^.**  
**6: Number of potential Water contacts**   
**2 : Number of second shell contacts involving water or ions**   
**33%: Percentage of satisfied second shell Water contacts**   
• Contact(s): /N7/O3'/O_w_/O_w_/OP1/  
• Contact code: O_ph_O_r_N_b_2O_w_  
→ Coordination < 6 for K ←  
→2Fo-Fc density between 1.0 and 4.0 !   
Go to ToC

### K:A:1659

|  | Res. type | Atom | Residue | Distance | Occupancy | B-factor | Closest atom(s) to a 1^st^ shell water | EM\_dens. |
| --- | --- | --- | --- | --- | --- | --- | --- | --- |
| **Ion** | | | |  | 1.0 | 30.0 |  | 7.90 |
 1 | Water 1 | | A:3147 | 2.57 | 1.0 | 48.5 | 2.77 [G:N7]A:800 | 2.42 | 2 | A | OP2 | A:780 | 2.74 | 1.0 | 37.2 |  | 7.99 | 3 | G | O6 | A:800 | 2.84 | 1.0 | 38.0 |  | 8.56 | 4 | Water 2 | | A:2983 | 2.89 | 1.0 | 41.7 | 2.74 [A:N7]A:780 | 4.48 | 5 | Water 3 | | A:2929 | 3.07 | 1.0 | 44.0 |  | 4.09 | 6 | G | N7 | A:800 | 3.50 | 1.0 | 40.0 |  | 7.04 | 7 | Water 4 | | A:3949 | 3.94 | 1.0 | 45.2 | 2.95 [G:N7]A:799 | 5.36 | 8 | Water 5 | | A:3226 | 4.08 | 1.0 | 52.9 |  | 2.15 | 9 | A | N7 | A:780 | 4.19 | 1.0 | 31.7 |  | 10.36 | 10 | Water 6 | | A:2955 | 4.29 | 1.0 | 40.5 |  | 7.03 | 11 | A | O5' | A:780 | 4.45 | 1.0 | 36.7 |  | 8.81 | 12 | Water 7 | | A:2369 | 4.70 | 1.0 | 46.7 |  | 3.74 | 13 | G | N7 | A:799 | 4.77 | 1.0 | 37.2 |  | 7.30 |

  
**1^st^ shell coordination angles and distances (*trans* in "blue"; *cys* in "green"; deviations in "orange"):**   
• 1-2(W/OP2): 158; 1-3(W/O6): 106; 1-4(W/W): 77; 1-5(W/W): 111; 2-3(OP2/O6): 88; 2-4(OP2/W): 84; 2-5(OP2/W): 56; 3-4(O6/W): 96; 3-5(O6/W): 143; 4-5(W/W): 88;   
• 1-2(W/OP2): 5.2; 1-3(W/O6): 4.3; 1-4(W/W): 3.4; 1-5(W/W): 4.6; 2-3(OP2/O6): 3.9; 2-4(OP2/W): 3.8; 2-5(OP2/W): 2.7; 3-4(O6/W): 4.2; 3-5(O6/W): 5.6; 4-5(W/W): 4.1;   
  
**4.27 ± 1.38: the average coodination distance of the 1^st^ six ligands could correspond to an hexahydrated Mg^2+^.**  
**10: Number of potential Water contacts**   
**3 : Number of second shell contacts involving water or ions**   
**30%: Percentage of satisfied second shell Water contacts**   
• Contact(s): /O6/O_w_/O_w_/OP2/  
• Contact code: O_ph_O_b_2O_w_  
• Contact(s) in the exclusion zone: /O_w_/  
→ Coordination < 6 for K ←  
Go to ToC

### K:A:1660

|  | Res. type | Atom | Residue | Distance | Occupancy | B-factor | Closest atom(s) to a 1^st^ shell water | EM\_dens. |
| --- | --- | --- | --- | --- | --- | --- | --- | --- |
| **Ion** | | | |  | 1.0 | 30.0 |  | 1.89 |
 1 | **K** | K | A:1661 | 3.93 | 1.0 | 30.0 |  | 2.46 | 2 | A | OP2 | A:415 | 4.13 | 1.0 | 101.5 |  | 1.09 | 3 | U | O4 | A:426 | 4.40 | 1.0 | 90.8 |  | 1.59 | 4 | G | O6 | A:425 | 4.46 | 1.0 | 95.0 |  | 2.10 | 5 | G | N7 | A:416 | 4.49 | 1.0 | 97.8 |  | 2.06 | 6 | G | O6 | A:417 | 4.61 | 1.0 | 92.5 |  | 2.16 | 7 | G | O6 | A:416 | 4.74 | 1.0 | 96.3 |  | 2.24 | 8 | C | N4 | A:418 | 4.77 | 1.0 | 91.3 |  | 1.84 |

  
• • **5.92 ± 1.74: the average coodination distance of the 1^st^ six ligands could correspond to an hexahydrated Mg^2+^.**  
**0: Number of potential Water contacts**   
• Contact code: → No contacts ←   
→ First K coordination shell missing  
→2Fo-Fc density between 1.0 and 4.0 !   
Go to ToC

### K:A:1661

|  | Res. type | Atom | Residue | Distance | Occupancy | B-factor | Closest atom(s) to a 1^st^ shell water | EM\_dens. |
| --- | --- | --- | --- | --- | --- | --- | --- | --- |
| **Ion** | | | |  | 1.0 | 30.0 |  | 2.46 |
 1 | G | O6 | A:416 | 2.61 | 1.0 | 96.3 |  | 2.24 | 2 | A | OP2 | A:414 | 3.06 | 1.0 | 101.0 |  | 1.13 | 3 | G | O6 | A:428 | 3.10 | 1.0 | 93.4 |  | 1.71 | 4 | U | O4 | A:427 | 3.32 | 1.0 | 91.2 |  | 1.21 | 5 | A | N7 | A:415 | 3.36 | 1.0 | 99.1 |  | 1.89 | 6 | G | N1 | A:428 | 3.62 | 1.0 | 92.4 |  | 3.14 | 7 | A | OP2 | A:415 | 3.83 | 1.0 | 101.5 |  | 1.09 | 8 | G | N7 | A:416 | 3.89 | 1.0 | 97.8 |  | 2.06 | 9 | **K** | K | A:1660 | 3.93 | 1.0 | 30.0 |  | 1.89 | 10 | U | O4 | A:426 | 4.14 | 1.0 | 90.8 |  | 1.59 | 11 | A | OP1 | A:414 | 4.55 | 1.0 | 100.9 |  | 1.19 | 12 | A | O5' | A:414 | 4.58 | 1.0 | 101.6 |  | 2.09 | 13 | G | O6 | A:417 | 4.76 | 1.0 | 92.5 |  | 2.16 |

  
**1^st^ shell coordination angles and distances (*trans* in "blue"; *cys* in "green"; deviations in "orange"):**   
• 1-2(O6/OP2): 150; 1-3(O6/O6): 64; 2-3(OP2/O6): 87;   
• 1-2(O6/OP2): 5.5; 1-3(O6/O6): 3.1; 2-3(OP2/O6): 4.3;   
  
**4.46 ± 1.37: the average coodination distance of the 1^st^ six ligands could correspond to an hexahydrated Mg^2+^.**  
**0: Number of potential Water contacts**   
• Contact(s): /O6/O6/OP2/  
• Contact code: O_ph_2O_b_  
→ Coordination < 6 for K ←  
→2Fo-Fc density between 1.0 and 4.0 !   
Go to ToC

### K:A:1662

|  | Res. type | Atom | Residue | Distance | Occupancy | B-factor | Closest atom(s) to a 1^st^ shell water | EM\_dens. |
| --- | --- | --- | --- | --- | --- | --- | --- | --- |
| **Ion** | | | |  | 1.0 | 30.0 |  | 10.49 |
 1 | Water 1 | | a:9568 | 2.40 | 1.0 | 44.9 |  | 4.11 | 2 | C | O2 | A:899 | 2.62 | 1.0 | 44.0 |  | 8.20 | 3 | C | OP1 | a:1834 | 2.72 | 1.0 | 40.1 |  | 7.60 | 4 | C | O2' | A:770 | 2.93 | 1.0 | 34.0 |  | 9.79 | 5 | Water 2 | | A:3087 | 3.20 | 1.0 | 47.7 | 2.76 [G:OP1]A:771 | 2.67 | 6 | C | O3' | A:770 | 3.40 | 1.0 | 37.9 |  | 9.99 | 7 | Water 3 | | A:2610 | 3.66 | 1.0 | 46.5 |  | 4.24 | 8 | C | N3 | A:899 | 4.18 | 1.0 | 32.7 |  | 9.94 | 9 | C | O5' | a:1834 | 4.18 | 1.0 | 35.9 |  | 9.87 | 10 | C | OP2 | a:1834 | 4.21 | 1.0 | 37.1 |  | 9.69 | 11 | G | OP1 | A:771 | 4.37 | 1.0 | 40.5 |  | 6.50 | 12 | A | N3 | A:900 | 4.45 | 1.0 | 31.9 |  | 9.75 | 13 | A | O4' | A:900 | 4.46 | 1.0 | 40.4 |  | 10.40 | 14 | G | O5' | A:771 | 4.52 | 1.0 | 36.7 |  | 8.70 | 15 | C | OP2 | a:1835 | 4.58 | 1.0 | 42.7 |  | 6.26 | 16 | Water 4 | | A:3105 | 4.63 | 1.0 | 40.6 | 2.76 [A:N3]A:900 | 4.35 | 17 | C | N1 | A:899 | 4.69 | 1.0 | 34.0 |  | 10.89 | 18 | C | O2' | A:899 | 4.74 | 1.0 | 32.5 |  | 9.15 |

  
**1^st^ shell coordination angles and distances (*trans* in "blue"; *cys* in "green"; deviations in "orange"):**   
• 1-2(W/O2): 149; 1-3(W/OP1): 111; 1-4(W/O2'): 103; 1-5(W/W): 92; 2-3(O2/OP1): 77; 2-4(O2/O2'): 65; 2-5(O2/W): 119; 3-4(OP1/O2'): 142; 3-5(OP1/W): 82; 4-5(O2'/W): 115;   
• 1-2(W/O2): 4.8; 1-3(W/OP1): 4.2; 1-4(W/O2'): 4.2; 1-5(W/W): 4.1; 2-3(O2/OP1): 3.3; 2-4(O2/O2'): 3.0; 2-5(O2/W): 5.0; 3-4(OP1/O2'): 5.3; 3-5(OP1/W): 3.9; 4-5(O2'/W): 5.2;   
  
**4.19 ± 1.37: the average coodination distance of the 1^st^ six ligands could correspond to an hexahydrated Mg^2+^.**  
**6: Number of potential Water contacts**   
**1 : Number of second shell contacts involving water or ions**   
**17%: Percentage of satisfied second shell Water contacts**   
• Contact(s): /O2/O2'/O_w_/OP1/  
• Contact code: O_ph_O_r_O_b_O_w_  
• Contact(s) in the exclusion zone: /O_w_/  
→ Coordination < 6 for K ←  
Go to ToC

### K:A:1663

|  | Res. type | Atom | Residue | Distance | Occupancy | B-factor | Closest atom(s) to a 1^st^ shell water | EM\_dens. |
| --- | --- | --- | --- | --- | --- | --- | --- | --- |
| **Ion** | | | |  | 1.0 | 30.0 |  | 5.03 |
 1 | G | O6 | A:666 | 2.85 | 1.0 | 42.2 |  | 4.98 | 2 | G | O6 | A:741 | 2.89 | 1.0 | 40.2 |  | 6.13 | 3 | Water 1 | | A:3552 | 2.91 | 1.0 | 51.9 | 2.85 [G:N7]A:666 | 2.25 | 4 | Water 2 | | A:3324 | 3.01 | 1.0 | 43.9 |  | 4.47 | 5 | Water 3 | | A:1866 | 3.19 | 1.0 | 44.2 | 2.36 [G:N7]A:741 | 4.63 | 6 | G | O6 | A:664 | 3.20 | 1.0 | 46.0 |  | 4.51 | 7 | U | O4 | A:740 | 3.80 | 1.0 | 47.0 |  | 4.63 | 8 | Water 4 | | A:2833 | 3.92 | 1.0 | 53.2 |  | 2.24 | 9 | Water 5 | | A:2286 | 4.05 | 1.0 | 54.8 |  | 1.72 | 10 | G | N7 | A:666 | 4.23 | 1.0 | 33.9 |  | 7.90 | 11 | G | N7 | A:741 | 4.27 | 1.0 | 38.2 |  | 7.53 |

  
**1^st^ shell coordination angles and distances (*trans* in "blue"; *cys* in "green"; deviations in "orange"):**   
• 1-2(O6/O6): 75; 1-3(O6/W): 77; 1-4(O6/W): 123; 1-5(O6/W): 83; 1-6(O6/O6): 135; 2-3(O6/W): 91; 2-4(O6/W): 59; 2-5(O6/W): 72; 2-6(O6/O6): 71; 3-4(W/W): 131; 3-5(W/W): 156; 3-6(W/O6): 74; 4-5(W/W): 53; 4-6(W/O6): 60; 5-6(W/O6): 113;   
• 1-2(O6/O6): 3.5; 1-3(O6/W): 3.6; 1-4(O6/W): 5.2; 1-5(O6/W): 4.0; 1-6(O6/O6): 5.6; 2-3(O6/W): 4.1; 2-4(O6/W): 2.9; 2-5(O6/W): 3.6; 2-6(O6/O6): 3.6; 3-4(W/W): 5.4; 3-5(W/W): 6.0; 3-6(W/O6): 3.7; 4-5(W/W): 2.8; 4-6(W/O6): 3.1; 5-6(W/O6): 5.3;   
  
**4.30 ± 1.32: the average coodination distance of the 1^st^ six ligands could correspond to an hexahydrated Mg^2+^.**  
**8: Number of potential Water contacts**   
**2 : Number of second shell contacts involving water or ions**   
**25%: Percentage of satisfied second shell Water contacts**   
• Contact(s): /O6/O6/O6/O_w_/O_w_/O_w_/  
• Contact code: 3O_b_3O_w_  
• K hydration shell issue: water too close (< 2.4 Å.)  
Go to ToC

### K:A:1664

|  | Res. type | Atom | Residue | Distance | Occupancy | B-factor | Closest atom(s) to a 1^st^ shell water | EM\_dens. |
| --- | --- | --- | --- | --- | --- | --- | --- | --- |
| **Ion** | | | |  | 1.0 | 30.0 |  | 3.80 |
 1 | G | N7 | A:945 | 2.55 | 1.0 | 68.6 |  | 2.95 | 2 | G | N7 | A:944 | 2.59 | 1.0 | 61.6 |  | 3.87 | 3 | G | O6 | A:945 | 3.12 | 1.0 | 69.5 |  | 2.47 | 4 | G | OP2 | A:944 | 3.23 | 1.0 | 62.7 |  | 3.56 | 5 | U | OP2 | A:943 | 3.51 | 1.0 | 62.2 |  | 2.72 | 6 | G | N7 | A:1337 | 3.57 | 1.0 | 71.5 |  | 3.21 | 7 | U | O5' | A:943 | 4.00 | 1.0 | 61.8 |  | 3.06 | 8 | G | O6 | A:944 | 4.10 | 1.0 | 62.1 |  | 3.35 | 9 | G | O5' | A:944 | 4.54 | 1.0 | 64.9 |  | 3.65 | 10 | U | OP1 | A:943 | 4.63 | 1.0 | 60.5 |  | 3.29 |

  
**1^st^ shell coordination angles and distances (*trans* in "blue"; *cys* in "green"; deviations in "orange"):**   
• 1-2(N7/N7): 89; 1-3(N7/O6): 64; 2-3(N7/O6): 152;   
• 1-2(N7/N7): 3.6; 1-3(N7/O6): 3.1; 2-3(N7/O6): 5.6;   
  
**4.45 ± 1.43: the average coodination distance of the 1^st^ six ligands could correspond to an hexahydrated Mg^2+^.**  
**0: Number of potential Water contacts**   
• Contact(s): /O6/  
• Contact code: O_b_  
• Contact(s) in the exclusion zone: /N7/N7/  
→ Coordination < 6 for K ←  
→2Fo-Fc density between 1.0 and 4.0 !   
Go to ToC

### K:A:1665

|  | Res. type | Atom | Residue | Distance | Occupancy | B-factor | Closest atom(s) to a 1^st^ shell water | EM\_dens. |
| --- | --- | --- | --- | --- | --- | --- | --- | --- |
| **Ion** | | | |  | 1.0 | 30.0 |  | 6.81 |
 1 | G | O6 | A:971 | 2.77 | 1.0 | 62.9 |  | 3.08 | 2 | U | O2' | A:1364 | 2.88 | 1.0 | 70.5 |  | 2.24 | 3 | U | O2 | A:1364 | 2.91 | 1.0 | 71.4 |  | 2.68 | 4 | G | O2' | A:1233 | 2.96 | 1.0 | 61.3 |  | 3.66 | 5 | A | N3 | A:949 | 3.17 | 1.0 | 62.4 |  | 4.28 | 6 | G | N3 | A:1233 | 3.18 | 1.0 | 63.6 |  | 3.67 | 7 | G | N7 | A:971 | 3.36 | 1.0 | 61.4 |  | 3.86 | 8 | G | N2 | A:1233 | 3.84 | 1.0 | 59.3 |  | 4.28 | 9 | C | O4' | A:1234 | 4.00 | 1.0 | 66.7 |  | 2.94 | 10 | U | O3' | A:1364 | 4.08 | 1.0 | 69.7 |  | 2.22 | 11 | U | N1 | A:1364 | 4.32 | 1.0 | 66.9 |  | 3.34 | 12 | A | O2' | A:949 | 4.66 | 1.0 | 61.8 |  | 2.98 |

  
**1^st^ shell coordination angles and distances (*trans* in "blue"; *cys* in "green"; deviations in "orange"):**   
• 1-2(O6/O2'): 60; 1-3(O6/O2): 94; 1-4(O6/O2'): 137; 1-5(O6/N3): 73; 1-6(O6/N3): 135; 2-3(O2'/O2): 76; 2-4(O2'/O2'): 99; 2-5(O2'/N3): 131; 2-6(O2'/N3): 155; 3-4(O2/O2'): 119; 3-5(O2/N3): 92; 3-6(O2/N3): 83; 4-5(O2'/N3): 128; 4-6(O2'/N3): 80; 5-6(N3/N3): 61;   
• 1-2(O6/O2'): 2.8; 1-3(O6/O2): 4.1; 1-4(O6/O2'): 5.3; 1-5(O6/N3): 3.6; 1-6(O6/N3): 5.5; 2-3(O2'/O2): 3.6; 2-4(O2'/O2'): 4.4; 2-5(O2'/N3): 5.5; 2-6(O2'/N3): 5.9; 3-4(O2/O2'): 5.0; 3-5(O2/N3): 4.4; 3-6(O2/N3): 4.0; 4-5(O2'/N3): 5.5; 4-6(O2'/N3): 4.0; 5-6(N3/N3): 3.2;   
  
**4.18 ± 1.26: the average coodination distance of the 1^st^ six ligands could correspond to an hexahydrated Mg^2+^.**  
**0: Number of potential Water contacts**   
• Contact(s): /N3_A_/N3_G_/O2/O2'/O2'/O6/  
• Contact code: 2O_r_2O_b_2N_b_  
Go to ToC

### K:A:1666

|  | Res. type | Atom | Residue | Distance | Occupancy | B-factor | Closest atom(s) to a 1^st^ shell water | EM\_dens. |
| --- | --- | --- | --- | --- | --- | --- | --- | --- |
| **Ion** | | | |  | 1.0 | 30.0 |  | 8.06 |
 1 | Water 1 | | A:2121 | 2.58 | 1.0 | 49.3 | 2.53 [G:O6]A:108 // 2.77 [A:N1]A:109 | 3.22 | 2 | G | O6 | A:326 | 2.94 | 1.0 | 49.2 |  | 4.78 | 3 | G | N7 | A:108 | 2.99 | 1.0 | 47.2 |  | 6.05 | 4 | G | OP2 | A:108 | 3.04 | 1.0 | 45.8 |  | 5.62 | 5 | G | OP2 | A:107 | 3.11 | 1.0 | 50.2 |  | 4.27 | 6 | A | N1 | A:109 | 3.37 | 1.0 | 38.6 |  | 6.97 | 7 | Water 2 | | A:3902 | 3.43 | 1.0 | 56.3 |  | 2.77 | 8 | G | O5' | A:107 | 3.52 | 1.0 | 48.2 |  | 5.53 | 9 | G | N7 | A:107 | 4.06 | 1.0 | 44.9 |  | 5.40 | 10 | Water 3 | | A:3695 | 4.24 | 1.0 | 51.1 |  | 2.50 | 11 | Water 4 | | A:3482 | 4.28 | 1.0 | 44.2 |  | 4.93 | 12 | G | OP1 | A:107 | 4.43 | 1.0 | 52.8 |  | 3.69 | 13 | G | O6 | A:108 | 4.59 | 1.0 | 44.7 |  | 5.39 | 14 | G | O5' | A:108 | 4.61 | 1.0 | 45.5 |  | 6.26 | 15 | Water 5 | | A:4446 | 4.78 | 1.0 | 54.2 |  | 2.95 |

  
**1^st^ shell coordination angles and distances (*trans* in "blue"; *cys* in "green"; deviations in "orange"):**   
• 1-2(W/O6): 83; 1-3(W/N7): 66; 1-4(W/OP2): 128; 1-5(W/OP2): 122; 2-3(O6/N7): 133; 2-4(O6/OP2): 83; 2-5(O6/OP2): 81; 3-4(N7/OP2): 89; 3-5(N7/OP2): 145; 4-5(OP2/OP2): 105;   
• 1-2(W/O6): 3.7; 1-3(W/N7): 3.0; 1-4(W/OP2): 5.1; 1-5(W/OP2): 5.0; 2-3(O6/N7): 5.4; 2-4(O6/OP2): 4.0; 2-5(O6/OP2): 3.9; 3-4(N7/OP2): 4.2; 3-5(N7/OP2): 5.8; 4-5(OP2/OP2): 4.9;   
  
**4.16 ± 1.25: the average coodination distance of the 1^st^ six ligands could correspond to an hexahydrated Mg^2+^.**  
**4: Number of potential Water contacts**   
**2 : Number of second shell contacts involving water or ions**   
**50%: Percentage of satisfied second shell Water contacts**   
• Contact(s): /N7/O6/OP2/OP2/  
• Contact code: 2O_ph_O_b_N_b_  
• Contact(s) in the exclusion zone: /O_w_/  
→ Coordination < 6 for K ←  
Go to ToC

### K:A:1667

|  | Res. type | Atom | Residue | Distance | Occupancy | B-factor | Closest atom(s) to a 1^st^ shell water | EM\_dens. |
| --- | --- | --- | --- | --- | --- | --- | --- | --- |
| **Ion** | | | |  | 1.0 | 30.0 |  | 12.79 |
 1 | G | O6 | A:1523 | 2.73 | 1.0 | 32.8 |  | 12.09 | 2 | Water 1 | | A:3364 | 2.88 | 1.0 | 34.6 |  | 8.91 | 3 | U | O4 | A:1512 | 3.08 | 1.0 | 32.8 |  | 11.11 | 4 | G | O6 | A:1511 | 3.14 | 1.0 | 31.2 |  | 10.57 | 5 | U | O4 | A:1522 | 3.28 | 1.0 | 32.4 |  | 10.35 | 6 | Water 2 | | A:2697 | 3.50 | 1.0 | 44.6 | 2.69 [U:O4]A:1522 | 2.91 | 7 | Water 3 | | A:4077 | 3.63 | 1.0 | 42.2 | 2.99 [C:N4]A:1510 | 3.18 | 8 | C | N4 | A:1524 | 3.69 | 1.0 | 23.2 |  | 11.11 | 9 | G | N7 | A:1523 | 4.29 | 1.0 | 24.3 |  | 12.62 | 10 | Water 4 | | A:3597 | 4.58 | 1.0 | 43.8 |  | 3.68 | 11 | A | N1 | A:1513 | 4.74 | 1.0 | 28.4 |  | 10.03 |

  
**1^st^ shell coordination angles and distances (*trans* in "blue"; *cys* in "green"; deviations in "orange"):**   
• 1-2(O6/W): 122; 1-3(O6/O4): 80; 1-4(O6/O6): 71; 2-3(W/O4): 56; 2-4(W/O6): 60; 3-4(O4/O6): 70;   
• 1-2(O6/W): 4.9; 1-3(O6/O4): 3.7; 1-4(O6/O6): 3.4; 2-3(W/O4): 2.8; 2-4(W/O6): 3.0; 3-4(O4/O6): 3.6;   
  
**4.32 ± 1.30: the average coodination distance of the 1^st^ six ligands could correspond to an hexahydrated Mg^2+^.**  
**6: Number of potential Water contacts**   
**2 : Number of second shell contacts involving water or ions**   
**33%: Percentage of satisfied second shell Water contacts**   
• Contact(s): /O4/O6/O6/O_w_/  
• Contact code: 3O_b_O_w_  
→ Coordination < 6 for K ←  
Go to ToC

### MG:A:1668

|  | Res. type | Atom | Residue | Distance | Occupancy | B-factor | Closest atom(s) to a 1^st^ shell water | EM\_dens. |
| --- | --- | --- | --- | --- | --- | --- | --- | --- |
| **Ion** | | | |  | 1.0 | 61.2 |  | 3.32 |
 1 | Water 1 | | A:4620 | 2.06 | 1.0 | 55.5 |  | 1.95 | 2 | Water 2 | | A:2560 | 2.07 | 1.0 | 52.5 | 2.65 [G:O6]A:305 | 3.47 | 3 | Water 3 | | A:3032 | 2.08 | 1.0 | 49.9 | 2.75 [G:N7]A:305 | 3.28 | 4 | Water 4 | | A:3390 | 2.36 | 1.0 | 52.2 | 2.82 [C:OP1]A:290 // 2.91 [U:OP2]A:291 | 4.48 | 5 | Water 5 | | A:4420 | 4.06 | 1.0 | 53.6 |  | 1.74 | 6 | Water 6 | | A:4102 | 4.08 | 1.0 | 57.5 |  | 2.14 | 7 | U | OP1 | A:291 | 4.22 | 1.0 | 52.1 |  | 3.56 | 8 | G | N7 | A:305 | 4.36 | 1.0 | 43.4 |  | 5.64 | 9 | U | OP2 | A:291 | 4.40 | 1.0 | 43.2 |  | 6.91 | 10 | Water 7 | | A:3751 | 4.43 | 1.0 | 51.9 | 2.90 [C:OP1]A:290 | 1.63 | 11 | G | O6 | A:305 | 4.52 | 1.0 | 51.3 |  | 5.41 | 12 | C | OP1 | A:290 | 4.60 | 1.0 | 46.7 |  | 3.82 |

  
**1^st^ shell coordination angles and distances (*trans* in "blue"; *cys* in "green"; deviations in "orange"):**   
• 1-2(W/W): *97º*; 1-3(W/W): *165º*; 2-3(W/W): *73º*;   
• 1-2(W/W): 3.1Å; 1-3(W/W): 4.1Å; 2-3(W/W): 2.5Å;   
  
**2.79 ± 0.91: the average coodination distance of the 1^st^ six ligands is K^+^ compatible**  
**12: Number of potential Water contacts**   
**4 : Number of second shell contacts involving water or ions**   
**33%: Percentage of satisfied second shell Water contacts**   
• Contact(s): /O_w_/O_w_/O_w_/  
• Contact code: 3O_w_  
• Contact(s) in the exclusion zone: /O_w_/  
→ Missing octahedral coordination for MG ←  
→2Fo-Fc density between 1.0 and 4.0 !   
Go to ToC

### MG:A:1669

|  | Res. type | Atom | Residue | Distance | Occupancy | B-factor | Closest atom(s) to a 1^st^ shell water | EM\_dens. |
| --- | --- | --- | --- | --- | --- | --- | --- | --- |
| **Ion** | | | |  | 1.0 | 46.0 |  | 9.74 |
 1 | Water 1 | | A:3795 | 1.89 | 1.0 | 33.0 | 2.91 [G:N7]A:769 | 7.56 | 2 | Water 2 | | A:4659 | 1.97 | 1.0 | 36.5 |  | 5.60 | 3 | Water 3 | | A:3668 | 2.15 | 1.0 | 41.9 |  | 5.85 | 4 | A | OP2 | A:768 | 2.16 | 1.0 | 33.9 |  | 10.46 | 5 | Water 4 | | A:3139 | 2.21 | 1.0 | 39.7 | 2.77 [U:OP1]A:804 | 6.07 | 6 | Water 5 | | A:3245 | 2.21 | 1.0 | 46.0 | 2.79 [A:N7]A:768 | 6.65 | 7 | Water 6 | | A:2736 | 4.06 | 1.0 | 37.2 | 2.90 [U:OP1]A:804 | 8.38 | 8 | A | N7 | A:768 | 4.08 | 1.0 | 25.1 |  | 12.88 | 9 | A | O5' | A:768 | 4.12 | 1.0 | 32.7 |  | 9.99 | 10 | Water 7 | | A:3489 | 4.15 | 1.0 | 53.2 |  | 2.83 | 11 | Water 8 | | A:1829 | 4.28 | 1.0 | 42.3 | 2.30 [A:OP2]A:767 | 4.23 | 12 | U | OP1 | A:804 | 4.37 | 1.0 | 36.9 |  | 8.57 | 13 | Water 9 | | A:4091 | 4.40 | 1.0 | 46.2 |  | 2.95 | 14 | A | O3' | A:767 | 4.52 | 1.0 | 35.2 |  | 9.69 | 15 | A | OP1 | A:768 | 4.55 | 1.0 | 39.7 |  | 7.97 | 16 | G | N7 | A:769 | 4.59 | 1.0 | 29.1 |  | 10.34 | 17 | A | OP2 | A:767 | 4.60 | 1.0 | 34.8 |  | 7.02 | 18 | A | O5' | A:767 | 4.64 | 1.0 | 37.2 |  | 9.37 | 19 | Water 10 | | A:4065 | 4.78 | 1.0 | 46.1 | 2.99 [U:OP1]A:804 | 4.66 | 20 | C | OP2 | A:805 | 4.79 | 1.0 | 41.0 |  | 6.37 |

  
**1^st^ shell coordination angles and distances (*trans* in "blue"; *cys* in "green"; deviations in "orange"):**   
• 1-2(W/W): 92º; 1-3(W/W): 176º; 1-4(W/OP2): 93º; 1-5(W/W): *97º*; 1-6(W/W): 89º; 2-3(W/W): 91º; 2-4(W/OP2): 174º; 2-5(W/W): *79º*; 2-6(W/W): *81º*; 3-4(W/OP2): *84º*; 3-5(W/W): *80º*; 3-6(W/W): 95º; 4-5(OP2/W): *102º*; 4-6(OP2/W): *98º*; 5-6(W/W): *159º*;   
• 1-2(W/W): 2.8Å; 1-3(W/W): 4.0Å; 1-4(W/OP2): 2.9Å; 1-5(W/W): 3.1Å; 1-6(W/W): 2.9Å; 2-3(W/W): 2.9Å; 2-4(W/OP2): 4.1Å; 2-5(W/W): 2.7Å; 2-6(W/W): 2.7Å; 3-4(W/OP2): 2.9Å; 3-5(W/W): 2.8Å; 3-6(W/W): *3.2Å*; 4-5(OP2/W): *3.4Å*; 4-6(OP2/W): *3.3Å*; 5-6(W/W): *4.3Å*;   
  
**2.10 ± 0.12: the average coodination distance of the 1^st^ six ligands is Mg^2+^ compatible**  
**10: Number of potential Water contacts**   
**3 : Number of second shell contacts involving water or ions**   
**30%: Percentage of satisfied second shell Water contacts**   
• Contact(s): /O_w_/O_w_/O_w_/O_w_/OP2/  
• Contact code: O_ph_4O_w_  
• Contact(s) in the exclusion zone: /O_w_/  
→ Missing octahedral coordination for MG ←  
→ Slightly distorded coord. octahedron (7.1º) ←  
Go to ToC

### MG:A:1670

|  | Res. type | Atom | Residue | Distance | Occupancy | B-factor | Closest atom(s) to a 1^st^ shell water | EM\_dens. |
| --- | --- | --- | --- | --- | --- | --- | --- | --- |
| **Ion** | | | |  | 1.0 | 60.3 |  | 2.18 |
 1 | Water 1 | | A:3347 | 1.92 | 1.0 | 46.7 | 2.81 [A:OP1]A:814 | 3.20 | 2 | Water 2 | | A:3815 | 2.08 | 1.0 | 45.9 | 2.92 [U:OP2]A:1512 | 2.30 | 3 | Water 3 | | A:3026 | 2.09 | 1.0 | 49.8 | 2.75 [G:OP1]A:1511 | 2.52 | 4 | Water 4 | | A:2331 | 2.21 | 1.0 | 48.7 | 2.61 [G:OP1]A:903 // 2.70 [U:OP1]A:1512 | 3.98 | 5 | Water 5 | | A:4033 | 2.23 | 1.0 | 45.3 | 2.98 [G:OP1]A:903 | 3.21 | 6 | Water 6 | | A:4652 | 2.29 | 1.0 | 48.1 |  | 2.55 | 7 | Water 7 | | A:2660 | 3.92 | 1.0 | 55.5 | 2.68 [G:OP1]A:1511 | 1.54 | 8 | G | OP1 | A:1511 | 3.98 | 1.0 | 41.5 |  | 6.80 | 9 | G | OP1 | A:903 | 4.15 | 1.0 | 37.8 |  | 6.58 | 10 | A | OP1 | A:814 | 4.21 | 1.0 | 38.1 |  | 7.24 | 11 | Water 8 | | A:2083 | 4.21 | 1.0 | 46.4 | 2.51 [A:OP1]A:814 | 4.33 | 12 | U | OP1 | A:1512 | 4.27 | 1.0 | 38.4 |  | 7.49 | 13 | U | OP2 | A:1512 | 4.36 | 1.0 | 33.7 |  | 9.22 | 14 | Water 9 | | A:2293 | 4.40 | 1.0 | 42.2 | 2.59 [U:OP1]A:1512 | 5.24 | 15 | Water 10 | | A:2651 | 4.42 | 1.0 | 44.5 | 2.84 [G:OP1]A:903 | 3.28 | 16 | Water 11 | | A:4265 | 4.59 | 1.0 | 48.4 |  | 1.61 | 17 | G | O3' | A:1511 | 4.65 | 1.0 | 35.2 |  | 10.48 | 18 | Water 12 | | A:2424 | 4.67 | 1.0 | 40.9 | 2.92 [A:OP1]A:814 | 7.30 | 19 | Water 13 | | A:3401 | 4.80 | 1.0 | 44.7 |  | 4.55 | 20 | Water 14 | | A:4588 | 4.80 | 1.0 | 54.8 |  | 2.08 |

  
**1^st^ shell coordination angles and distances (*trans* in "blue"; *cys* in "green"; deviations in "orange"):**   
• 1-2(W/W): *116º*; 1-3(W/W): *98º*; 1-4(W/W): *85º*; 1-5(W/W): *79º*; 1-6(W/W): *154º*; 2-3(W/W): 91º; 2-4(W/W): *77º*; 2-5(W/W): *148º*; 2-6(W/W): *74º*; 3-4(W/W): *168º*; 3-5(W/W): *115º*; 3-6(W/W): *106º*; 4-5(W/W): *77º*; 4-6(W/W): *74º*; 5-6(W/W): *81º*;   
• 1-2(W/W): *3.4Å*; 1-3(W/W): 3.0Å; 1-4(W/W): 2.8Å; 1-5(W/W): 2.7Å; 1-6(W/W): 4.1Å; 2-3(W/W): 3.0Å; 2-4(W/W): 2.7Å; 2-5(W/W): 4.1Å; 2-6(W/W): 2.6Å; 3-4(W/W): *4.3Å*; 3-5(W/W): *3.6Å*; 3-6(W/W): *3.5Å*; 4-5(W/W): 2.7Å; 4-6(W/W): 2.7Å; 5-6(W/W): 2.9Å;   
  
**2.14 ± 0.12: the average coodination distance of the 1^st^ six ligands is Mg^2+^ compatible**  
**12: Number of potential Water contacts**   
**6 : Number of second shell contacts involving water or ions**   
**50%: Percentage of satisfied second shell Water contacts**   
• Contact(s): /O_w_/O_w_/O_w_/O_w_/O_w_/O_w_/  
• Contact code: 6O_w_  
→→→ Distorded coord. octahedron (15.2º) ←←←  
→2Fo-Fc density between 1.0 and 4.0 !   
Go to ToC

### MG:A:1671

|  | Res. type | Atom | Residue | Distance | Occupancy | B-factor | Closest atom(s) to a 1^st^ shell water | EM\_dens. |
| --- | --- | --- | --- | --- | --- | --- | --- | --- |
| **Ion** | | | |  | 1.0 | 35.6 |  | 13.72 |
 1 | A | OP2 | A:1499 | 2.03 | 1.0 | 35.4 |  | 9.22 | 2 | Water 1 | | A:3250 | 2.03 | 1.0 | 32.3 | 2.87 [G:O2']A:1504 | 9.60 | 3 | A | OP2 | A:1500 | 2.07 | 1.0 | 34.8 |  | 9.74 | 4 | Water 2 | | A:3830 | 2.08 | 1.0 | 31.1 |  | 11.91 | 5 | Water 3 | | A:3875 | 2.09 | 1.0 | 36.9 | 2.93 [UR3:O2']A:1498 | 9.33 | 6 | G | OP2 | A:1505 | 2.17 | 1.0 | 33.4 |  | 10.88 | 7 | UR3 | O3' | A:1498 | 3.31 | 1.0 | 35.9 |  | 8.35 | 8 | G | O3' | A:1504 | 3.87 | 1.0 | 37.7 |  | 9.05 | 9 | G | OP1 | A:1505 | 4.08 | 1.0 | 36.2 |  | 8.72 | 10 | A | O5' | A:1499 | 4.09 | 1.0 | 36.8 |  | 10.48 | 11 | A | O3' | A:1499 | 4.18 | 1.0 | 37.5 |  | 9.19 | 12 | Water 4 | | A:4185 | 4.21 | 1.0 | 32.5 |  | 11.09 | 13 | G | O2' | A:1504 | 4.30 | 1.0 | 33.8 |  | 10.32 | 14 | A | OP1 | A:1499 | 4.32 | 1.0 | 39.2 |  | 7.86 | 15 | UR3 | O2' | A:1498 | 4.33 | 1.0 | 34.4 |  | 8.80 | 16 | Water 5 | | A:4054 | 4.34 | 1.0 | 38.5 |  | 6.55 | 17 | A | O5' | A:1500 | 4.37 | 1.0 | 31.7 |  | 10.03 | 18 | A | OP1 | A:1500 | 4.43 | 1.0 | 32.4 |  | 10.66 | 19 | Water 6 | | A:2445 | 4.45 | 1.0 | 43.6 |  | 4.81 | 20 | Water 7 | | X:212 | 4.57 | 1.0 | 49.2 |  | 3.03 | 21 | A | N7 | A:1500 | 4.61 | 1.0 | 31.9 |  | 9.67 | 22 | G | O5' | A:1505 | 4.63 | 1.0 | 38.1 |  | 9.48 | 23 | Water 8 | | A:3742 | 4.68 | 1.0 | 41.1 |  | 6.60 |

  
**1^st^ shell coordination angles and distances (*trans* in "blue"; *cys* in "green"; deviations in "orange"):**   
• 1-2(OP2/W): *169º*; 1-3(OP2/OP2): 86º; 1-4(OP2/W): *96º*; 1-5(OP2/W): *102º*; 1-6(OP2/OP2): 92º; 2-3(W/OP2): 86º; 2-4(W/W): *77º*; 2-5(W/W): 87º; 2-6(W/OP2): 95º; 3-4(OP2/W): 89º; 3-5(OP2/W): 172º; 3-6(OP2/OP2): 88º; 4-5(W/W): 89º; 4-6(W/OP2): 172º; 5-6(W/OP2): 92º;   
• 1-2(OP2/W): 4.0Å; 1-3(OP2/OP2): 2.8Å; 1-4(OP2/W): 3.1Å; 1-5(OP2/W): 3.2Å; 1-6(OP2/OP2): 3.0Å; 2-3(W/OP2): 2.8Å; 2-4(W/W): 2.6Å; 2-5(W/W): 2.8Å; 2-6(W/OP2): 3.1Å; 3-4(OP2/W): 2.9Å; 3-5(OP2/W): 4.2Å; 3-6(OP2/OP2): 3.0Å; 4-5(W/W): 2.9Å; 4-6(W/OP2): *4.2Å*; 5-6(W/OP2): 3.1Å;   
  
**2.08 ± 0.05: the average coodination distance of the 1^st^ six ligands is Mg^2+^ compatible**  
**6: Number of potential Water contacts**   
**2 : Number of second shell contacts involving water or ions**   
**33%: Percentage of satisfied second shell Water contacts**   
• Contact(s): /O_w_/O_w_/O_w_/OP2/OP2/OP2/  
• Contact code: *fac-*3O_ph_3O_w_  
• Contact(s) in the exclusion zone: /O3'/  
→ Slightly distorded coord. octahedron (5.4º) ←  
Go to ToC

### MG:A:1672

|  | Res. type | Atom | Residue | Distance | Occupancy | B-factor | Closest atom(s) to a 1^st^ shell water | EM\_dens. |
| --- | --- | --- | --- | --- | --- | --- | --- | --- |
| **Ion** | | | |  | 1.0 | 37.2 |  | 13.91 |
 1 | A | OP2 | A:573 | 2.01 | 1.0 | 34.0 |  | 9.91 | 2 | Water 1 | | A:3319 | 2.02 | 1.0 | 35.3 | 2.80 [U:OP1]A:571 // 2.96 [A:OP2]A:572 | 8.44 | 3 | Water 2 | | A:4260 | 2.02 | 1.0 | 34.6 |  | 8.96 | 4 | A | OP2 | A:574 | 2.05 | 1.0 | 39.7 |  | 8.28 | 5 | Water 3 | | A:3929 | 2.27 | 1.0 | 37.6 |  | 8.76 | 6 | A | OP2 | A:572 | 2.32 | 1.0 | 40.8 |  | 8.92 | 7 | A | OP1 | A:572 | 3.74 | 1.0 | 37.9 |  | 7.95 | 8 | A | OP1 | A:573 | 3.83 | 1.0 | 38.9 |  | 7.89 | 9 | A | O5' | A:572 | 3.95 | 1.0 | 35.4 |  | 8.03 | 10 | A | OP1 | A:574 | 4.02 | 1.0 | 40.0 |  | 6.42 | 11 | A | O5' | A:573 | 4.15 | 1.0 | 38.8 |  | 7.89 | 12 | Water 4 | | A:2803 | 4.22 | 1.0 | 48.2 | 2.71 [A:OP1]A:574 | 2.88 | 13 | Water 5 | | A:4309 | 4.26 | 1.0 | 50.6 |  | 2.11 | 14 | Water 6 | | A:4242 | 4.35 | 1.0 | 43.5 |  | 5.67 | 15 | A | O3' | A:573 | 4.37 | 1.0 | 36.7 |  | 7.91 | 16 | U | OP1 | A:571 | 4.37 | 1.0 | 37.2 |  | 7.54 | 17 | A | O5' | A:574 | 4.42 | 1.0 | 38.6 |  | 7.38 | 18 | A | O3' | A:572 | 4.46 | 1.0 | 43.0 |  | 7.16 | 19 | Water 7 | | A:2195 | 4.62 | 1.0 | 41.5 |  | 4.48 | 20 | Water 8 | | A:2811 | 4.68 | 1.0 | 38.2 | 2.71 [A:OP1]A:573 | 8.03 | 21 | U | O3' | A:571 | 4.74 | 1.0 | 33.9 |  | 8.79 | 22 | U | OP2 | A:571 | 4.75 | 1.0 | 42.4 |  | 6.95 |

  
**1^st^ shell coordination angles and distances (*trans* in "blue"; *cys* in "green"; deviations in "orange"):**   
• 1-2(OP2/W): 173º; 1-3(OP2/W): *103º*; 1-4(OP2/OP2): 93º; 1-5(OP2/W): 87º; 2-3(W/W): *83º*; 2-4(W/OP2): 89º; 2-5(W/W): 87º; 3-4(W/OP2): *97º*; 3-5(W/W): *169º*; 4-5(OP2/W): 87º;   
• 1-2(OP2/W): 4.0Å; 1-3(OP2/W): 3.2Å; 1-4(OP2/OP2): 3.0Å; 1-5(OP2/W): 3.0Å; 2-3(W/W): 2.7Å; 2-4(W/OP2): 2.9Å; 2-5(W/W): 2.9Å; 3-4(W/OP2): 3.1Å; 3-5(W/W): *4.3Å*; 4-5(OP2/W): 3.0Å;   
  
**2.12 ± 0.13: the average coodination distance of the 1^st^ six ligands is Mg^2+^ compatible**  
**6: Number of potential Water contacts**   
**2 : Number of second shell contacts involving water or ions**   
**33%: Percentage of satisfied second shell Water contacts**   
• Contact(s): /O_w_/O_w_/O_w_/OP2/OP2/  
• Contact code: *cis-*2O_ph_3O_w_  
• Contact(s) in the exclusion zone: /OP2/  
→ Missing octahedral coordination for MG ←  
→ Slightly distorded coord. octahedron (5.9º) ←  
Go to ToC

### MG:A:1673

|  | Res. type | Atom | Residue | Distance | Occupancy | B-factor | Closest atom(s) to a 1^st^ shell water | EM\_dens. |
| --- | --- | --- | --- | --- | --- | --- | --- | --- |
| **Ion** | | | |  | 1.0 | 38.4 |  | 14.70 |
 1 | Water 1 | | A:3859 | 1.98 | 1.0 | 32.9 |  | 8.57 | 2 | A | OP2 | A:814 | 2.02 | 1.0 | 32.2 |  | 10.18 | 3 | Water 2 | | A:3198 | 2.02 | 1.0 | 30.9 | 2.78 [A:N7]A:816 | 13.07 | 4 | Water 3 | | A:4194 | 2.03 | 1.0 | 32.5 |  | 9.81 | 5 | Water 4 | | A:2733 | 2.05 | 1.0 | 32.3 | 2.81 [G:OP1]A:577 | 10.34 | 6 | Water 5 | | A:3251 | 2.23 | 1.0 | 31.1 | 2.79 [A:OP1]A:815 | 10.44 | 7 | A | OP1 | A:814 | 3.79 | 1.0 | 38.1 |  | 7.24 | 8 | Water 6 | | A:2254 | 4.13 | 1.0 | 40.9 |  | 6.65 | 9 | A | O5' | A:814 | 4.17 | 1.0 | 37.2 |  | 7.45 | 10 | G | OP1 | A:577 | 4.18 | 1.0 | 35.3 |  | 9.28 | 11 | Water 7 | | A:4685 | 4.26 | 1.0 | 53.4 |  | 2.18 | 12 | Water 8 | | A:2424 | 4.30 | 1.0 | 40.9 | 2.92 [A:OP1]A:814 | 7.30 | 13 | A | N7 | A:816 | 4.39 | 1.0 | 27.0 |  | 12.40 | 14 | G | OP2 | A:577 | 4.43 | 1.0 | 38.1 |  | 8.96 | 15 | U | O3' | A:813 | 4.47 | 1.0 | 35.2 |  | 8.15 | 16 | A | OP2 | A:815 | 4.50 | 1.0 | 37.8 |  | 9.10 | 17 | A | OP1 | A:815 | 4.60 | 1.0 | 35.5 |  | 9.54 |

  
**1^st^ shell coordination angles and distances (*trans* in "blue"; *cys* in "green"; deviations in "orange"):**   
• 1-2(W/OP2): 94º; 1-3(W/W): 176º; 1-4(W/W): *85º*; 1-5(W/W): 94º; 1-6(W/W): 94º; 2-3(OP2/W): 88º; 2-4(OP2/W): 179º; 2-5(OP2/W): *83º*; 2-6(OP2/W): *97º*; 3-4(W/W): 93º; 3-5(W/W): 90º; 3-6(W/W): *82º*; 4-5(W/W): *96º*; 4-6(W/W): *84º*; 5-6(W/W): 172º;   
• 1-2(W/OP2): 2.9Å; 1-3(W/W): 4.0Å; 1-4(W/W): 2.7Å; 1-5(W/W): 2.9Å; 1-6(W/W): 3.1Å; 2-3(OP2/W): 2.8Å; 2-4(OP2/W): 4.0Å; 2-5(OP2/W): 2.7Å; 2-6(OP2/W): 3.2Å; 3-4(W/W): 2.9Å; 3-5(W/W): 2.9Å; 3-6(W/W): 2.8Å; 4-5(W/W): 3.0Å; 4-6(W/W): 2.9Å; 5-6(W/W): *4.3Å*;   
  
**2.05 ± 0.08: the average coodination distance of the 1^st^ six ligands is Mg^2+^ compatible**  
**10: Number of potential Water contacts**   
**3 : Number of second shell contacts involving water or ions**   
**30%: Percentage of satisfied second shell Water contacts**   
• Contact(s): /O_w_/O_w_/O_w_/O_w_/O_w_/OP2/  
• Contact code: O_ph_5O_w_  
Ooctahedral coordination (4.7º)   
Go to ToC

### MG:A:1674

|  | Res. type | Atom | Residue | Distance | Occupancy | B-factor | Closest atom(s) to a 1^st^ shell water | EM\_dens. |
| --- | --- | --- | --- | --- | --- | --- | --- | --- |
| **Ion** | | | |  | 1.0 | 37.3 |  | 13.56 |
 1 | G | N7 | A:869 | 2.10 | 1.0 | 40.5 |  | 6.37 | 2 | G | N7 | A:858 | 2.22 | 1.0 | 45.6 |  | 6.06 | 3 | Water 1 | | A:1897 | 2.34 | 1.0 | 46.5 | 2.39 [G:O6]A:858 | 4.20 | 4 | G | O6 | A:858 | 3.44 | 1.0 | 50.2 |  | 5.97 | 5 | G | O6 | A:869 | 3.58 | 1.0 | 49.0 |  | 4.57 | 6 | G | OP2 | A:869 | 3.98 | 1.0 | 43.1 |  | 6.15 | 7 | C | OP2 | A:868 | 4.15 | 1.0 | 42.4 |  | 6.11 | 8 | G | OP2 | A:858 | 4.34 | 1.0 | 54.9 |  | 3.15 | 9 | Water 2 | | A:3882 | 4.58 | 1.0 | 50.6 | 2.94 [C:OP2]A:868 | 2.64 |

  
**1^st^ shell coordination angles and distances (*trans* in "blue"; *cys* in "green"; deviations in "orange"):**   
• 1-2(N7/N7): 91º;   
• 1-2(N7/N7): 3.1Å;   
  
**2.94 ± 0.74: the average coodination distance of the 1^st^ six ligands is K^+^ compatible**  
**2: Number of potential Water contacts**   
**1 : Number of second shell contacts involving water or ions**   
**50%: Percentage of satisfied second shell Water contacts**   
• Contact(s): /N7/N7/  
• Contact code: *cis-*2N_b_  
• Contact(s) in the exclusion zone: /O_w_/  
→ Missing octahedral coordination for MG ←  
• MG hydration shell issue: water too close (< 2.4 Å.)  
Go to ToC

### MG:A:1675

|  | Res. type | Atom | Residue | Distance | Occupancy | B-factor | Closest atom(s) to a 1^st^ shell water | EM\_dens. |
| --- | --- | --- | --- | --- | --- | --- | --- | --- |
| **Ion** | | | |  | 1.0 | 45.8 |  | 8.91 |
 1 | Water 1 | | A:3151 | 1.89 | 1.0 | 45.0 |  | 5.48 | 2 | Water 2 | | A:3933 | 1.95 | 1.0 | 45.0 |  | 5.16 | 3 | Water 3 | | A:2786 | 1.96 | 1.0 | 41.7 | 2.70 [G:OP2]A:558 | 5.80 | 4 | Water 4 | | A:4488 | 2.01 | 1.0 | 44.6 |  | 3.35 | 5 | A | OP2 | A:560 | 2.14 | 1.0 | 42.2 |  | 8.22 | 6 | Water 5 | | A:2942 | 2.28 | 1.0 | 41.4 | 2.73 [A:OP1]A:563 | 6.40 | 7 | A | O3' | A:559 | 3.70 | 1.0 | 39.8 |  | 6.90 | 8 | U | OP1 | A:562 | 3.99 | 1.0 | 48.3 |  | 4.22 | 9 | G | OP1 | A:557 | 4.12 | 1.0 | 46.0 |  | 4.87 | 10 | Water 6 | | A:4000 | 4.17 | 1.0 | 50.9 |  | 2.27 | 11 | Water 7 | | A:2598 | 4.19 | 1.0 | 46.5 |  | 4.02 | 12 | Water 8 | | A:2677 | 4.25 | 1.0 | 45.8 |  | 3.99 | 13 | A | OP1 | A:560 | 4.33 | 1.0 | 42.4 |  | 6.48 | 14 | G | OP2 | A:558 | 4.36 | 1.0 | 48.6 |  | 7.21 | 15 | A | O5' | A:560 | 4.49 | 1.0 | 46.6 |  | 4.62 | 16 | A | OP1 | A:563 | 4.56 | 1.0 | 43.6 |  | 5.43 | 17 | A | O2' | A:559 | 4.57 | 1.0 | 45.0 |  | 5.72 |

  
**1^st^ shell coordination angles and distances (*trans* in "blue"; *cys* in "green"; deviations in "orange"):**   
• 1-2(W/W): 175º; 1-3(W/W): 91º; 1-4(W/W): *99º*; 1-5(W/OP2): 87º; 1-6(W/W): 88º; 2-3(W/W): 93º; 2-4(W/W): *81º*; 2-5(W/OP2): 92º; 2-6(W/W): 87º; 3-4(W/W): 89º; 3-5(W/OP2): 95º; 3-6(W/W): 176º; 4-5(W/OP2): 173º; 4-6(W/W): 87º; 5-6(OP2/W): 89º;   
• 1-2(W/W): 3.8Å; 1-3(W/W): 2.7Å; 1-4(W/W): 3.0Å; 1-5(W/OP2): 2.8Å; 1-6(W/W): 2.9Å; 2-3(W/W): 2.8Å; 2-4(W/W): 2.6Å; 2-5(W/OP2): 3.0Å; 2-6(W/W): 2.9Å; 3-4(W/W): 2.8Å; 3-5(W/OP2): 3.0Å; 3-6(W/W): *4.2Å*; 4-5(W/OP2): 4.1Å; 4-6(W/W): 3.0Å; 5-6(OP2/W): 3.1Å;   
  
**2.04 ± 0.13: the average coodination distance of the 1^st^ six ligands is Mg^2+^ compatible**  
**10: Number of potential Water contacts**   
**2 : Number of second shell contacts involving water or ions**   
**20%: Percentage of satisfied second shell Water contacts**   
• Contact(s): /O_w_/O_w_/O_w_/O_w_/OP2/  
• Contact code: O_ph_4O_w_  
• Contact(s) in the exclusion zone: /O_w_/  
→ Missing octahedral coordination for MG ←  
Ooctahedral coordination (3.9º)   
Go to ToC

### MG:A:1676

|  | Res. type | Atom | Residue | Distance | Occupancy | B-factor | Closest atom(s) to a 1^st^ shell water | EM\_dens. |
| --- | --- | --- | --- | --- | --- | --- | --- | --- |
| **Ion** | | | |  | 1.0 | 50.7 |  | 6.41 |
 1 | Water 1 | | A:4047 | 1.82 | 1.0 | 49.2 | 2.98 [G:N7]A:357 | 4.50 | 2 | Water 2 | | A:2450 | 1.88 | 1.0 | 48.6 | 2.63 [C:OP2]A:355 | 4.70 | 3 | Water 3 | | A:2328 | 1.94 | 1.0 | 49.1 | 2.60 [C:OP2]A:355 // 2.87 [A:OP2]A:356 | 3.80 | 4 | Water 4 | | A:4621 | 2.00 | 1.0 | 53.1 |  | 3.90 | 5 | Water 5 | | A:2922 | 2.08 | 1.0 | 49.9 | 2.73 [G:O6]A:357 | 6.04 | 6 | Water 6 | | A:2918 | 2.14 | 1.0 | 51.2 | 2.73 [A:N7]A:356 | 4.74 | 7 | C | OP2 | A:355 | 3.73 | 1.0 | 53.4 |  | 4.41 | 8 | G | O6 | A:357 | 3.99 | 1.0 | 55.0 |  | 4.11 | 9 | Water 7 | | A:3298 | 4.15 | 1.0 | 50.8 | 2.80 [A:OP2]A:356 | 3.15 | 10 | Water 8 | | A:3254 | 4.25 | 1.0 | 53.4 | 2.79 [A:OP2]A:356 | 2.08 | 11 | G | N7 | A:357 | 4.29 | 1.0 | 52.3 |  | 4.15 | 12 | A | N7 | A:356 | 4.31 | 1.0 | 50.8 |  | 4.61 | 13 | Water 9 | | A:1869 | 4.36 | 1.0 | 58.4 |  | 2.50 | 14 | A | OP2 | A:356 | 4.38 | 1.0 | 57.8 |  | 3.37 | 15 | C | O2' | A:352 | 4.43 | 1.0 | 52.0 |  | 4.96 | 16 | C | O2 | A:352 | 4.59 | 1.0 | 53.4 |  | 4.22 | 17 | Water 10 | | A:3514 | 4.77 | 1.0 | 48.1 | 2.84 [C:OP2]A:355 | 3.95 | 18 | Water 11 | | A:3573 | 4.77 | 1.0 | 59.6 | 2.99 [G:O6]A:357 | 1.66 |

  
**1^st^ shell coordination angles and distances (*trans* in "blue"; *cys* in "green"; deviations in "orange"):**   
• 1-2(W/W): 176º; 1-3(W/W): *99º*; 1-4(W/W): 92º; 1-5(W/W): *96º*; 1-6(W/W): 89º; 2-3(W/W): *77º*; 2-4(W/W): 89º; 2-5(W/W): 89º; 2-6(W/W): 90º; 3-4(W/W): 88º; 3-5(W/W): *165º*; 3-6(W/W): *99º*; 4-5(W/W): 87º; 4-6(W/W): 172º; 5-6(W/W): 85º;   
• 1-2(W/W): *3.7Å*; 1-3(W/W): 2.9Å; 1-4(W/W): 2.7Å; 1-5(W/W): 2.9Å; 1-6(W/W): 2.8Å; 2-3(W/W): 2.4Å; 2-4(W/W): 2.7Å; 2-5(W/W): 2.8Å; 2-6(W/W): 2.9Å; 3-4(W/W): 2.7Å; 3-5(W/W): 4.0Å; 3-6(W/W): 3.1Å; 4-5(W/W): 2.8Å; 4-6(W/W): 4.1Å; 5-6(W/W): 2.9Å;   
  
**1.98 ± 0.11: the average coodination distance of the 1^st^ six ligands is Mg^2+^ compatible**  
**12: Number of potential Water contacts**   
**6 : Number of second shell contacts involving water or ions**   
**50%: Percentage of satisfied second shell Water contacts**   
• Contact(s): /O_w_/O_w_/O_w_/O_w_/  
• Contact code: 4O_w_  
• Contact(s) in the exclusion zone: /O_w_/O_w_/  
→ Missing octahedral coordination for MG ←  
→ Slightly distorded coord. octahedron (5.2º) ←  
Go to ToC

### MG:A:1677

|  | Res. type | Atom | Residue | Distance | Occupancy | B-factor | Closest atom(s) to a 1^st^ shell water | EM\_dens. |
| --- | --- | --- | --- | --- | --- | --- | --- | --- |
| **Ion** | | | |  | 1.0 | 66.8 |  | 3.94 |
 1 | Water 1 | | A:4250 | 1.95 | 1.0 | 60.0 |  | 2.49 | 2 | Water 2 | | A:3277 | 2.03 | 1.0 | 57.5 | 2.92 [G:O2']A:587 | 2.71 | 3 | Water 3 | | A:3657 | 2.04 | 1.0 | 62.8 | 2.88 [G:OP2]A:645 | 2.55 | 4 | G | OP2 | A:588 | 2.08 | 1.0 | 51.9 |  | 4.67 | 5 | G | O2' | A:587 | 4.00 | 1.0 | 49.0 |  | 2.88 | 6 | Water 4 | | A:3135 | 4.01 | 1.0 | 60.4 |  | 2.28 | 7 | Water 5 | | A:3222 | 4.03 | 1.0 | 51.7 |  | 2.97 | 8 | G | OP1 | A:588 | 4.08 | 1.0 | 55.3 |  | 3.42 | 9 | U | OP1 | A:644 | 4.18 | 1.0 | 55.9 |  | 4.03 | 10 | G | OP2 | A:645 | 4.23 | 1.0 | 59.4 |  | 3.62 | 11 | G | OP1 | A:645 | 4.30 | 1.0 | 60.0 |  | 3.16 | 12 | G | O3' | A:587 | 4.34 | 1.0 | 51.6 |  | 3.29 | 13 | Water 6 | | A:4366 | 4.38 | 1.0 | 59.7 |  | 2.04 | 14 | G | O5' | A:588 | 4.46 | 1.0 | 52.0 |  | 4.25 | 15 | Water 7 | | A:4275 | 4.75 | 1.0 | 58.1 |  | 1.64 |

  
**1^st^ shell coordination angles and distances (*trans* in "blue"; *cys* in "green"; deviations in "orange"):**   
• 1-2(W/W): 93º; 1-3(W/W): *71º*; 1-4(W/OP2): *98º*; 2-3(W/W): *77º*; 2-4(W/OP2): 86º; 3-4(W/OP2): *159º*;   
• 1-2(W/W): 2.9Å; 1-3(W/W): 2.3Å; 1-4(W/OP2): 3.0Å; 2-3(W/W): 2.5Å; 2-4(W/OP2): 2.8Å; 3-4(W/OP2): 4.1Å;   
  
**2.69 ± 0.93: the average coodination distance of the 1^st^ six ligands is K^+^ compatible**  
**8: Number of potential Water contacts**   
**2 : Number of second shell contacts involving water or ions**   
**25%: Percentage of satisfied second shell Water contacts**   
• Contact(s): /O_w_/O_w_/O_w_/OP2/  
• Contact code: O_ph_3O_w_  
→ Missing octahedral coordination for MG ←  
→→→ Distorded coord. octahedron (11.5º) ←←←  
→2Fo-Fc density between 1.0 and 4.0 !   
Go to ToC

### MG:A:1678

|  | Res. type | Atom | Residue | Distance | Occupancy | B-factor | Closest atom(s) to a 1^st^ shell water | EM\_dens. |
| --- | --- | --- | --- | --- | --- | --- | --- | --- |
| **Ion** | | | |  | 1.0 | 68.1 |  | 4.42 |
 1 | Water 1 | | A:4678 | 1.86 | 1.0 | 63.3 |  | 2.27 | 2 | Water 2 | | A:3834 | 1.88 | 1.0 | 60.2 | 2.92 [G:N7]A:646 | 3.55 | 3 | Water 3 | | A:3641 | 1.95 | 1.0 | 66.3 | 2.87 [G:O6]A:592 | 2.29 | 4 | Water 4 | | A:4657 | 2.01 | 1.0 | 59.3 |  | 2.59 | 5 | Water 5 | | A:2720 | 2.16 | 1.0 | 59.8 | 2.69 [G:N7]A:645 // 2.82 [MG:MG]A:1636 | 3.77 | 6 | Water 6 | | A:2370 | 2.30 | 1.0 | 60.8 | 2.62 [G:O6]A:646 // 2.65 [MG:MG]A:1636 | 3.60 | 7 | **MG** | MG | A:1636 | 3.75 | 1.0 | 51.6 |  | 7.15 | 8 | Water 7 | | A:2672 | 3.90 | 1.0 | 55.5 |  | 3.34 | 9 | Water 8 | | A:2570 | 4.04 | 1.0 | 57.7 |  | 2.92 | 10 | G | N7 | A:645 | 4.21 | 1.0 | 57.2 |  | 4.03 | 11 | G | O6 | A:646 | 4.27 | 1.0 | 64.9 |  | 3.97 | 12 | Water 9 | | A:3406 | 4.32 | 1.0 | 57.0 |  | 2.30 | 13 | Water 10 | | A:4654 | 4.43 | 1.0 | 60.5 |  | 2.01 | 14 | G | O6 | A:592 | 4.43 | 1.0 | 69.7 |  | 2.67 | 15 | G | O6 | A:645 | 4.50 | 1.0 | 59.3 |  | 4.29 | 16 | Water 11 | | A:4366 | 4.60 | 1.0 | 59.7 |  | 2.04 | 17 | U | O4 | A:593 | 4.64 | 1.0 | 70.6 |  | 2.85 | 18 | G | N7 | A:646 | 4.64 | 1.0 | 62.7 |  | 3.94 |

  
**1^st^ shell coordination angles and distances (*trans* in "blue"; *cys* in "green"; deviations in "orange"):**   
• 1-2(W/W): 171º; 1-3(W/W): 85º; 1-4(W/W): 86º; 1-5(W/W): *84º*; 1-6(W/W): 89º; 2-3(W/W): *98º*; 2-4(W/W): *102º*; 2-5(W/W): 92º; 2-6(W/W): *84º*; 3-4(W/W): 88º; 3-5(W/W): *166º*; 3-6(W/W): *83º*; 4-5(W/W): *99º*; 4-6(W/W): 170º; 5-6(W/W): 89º;   
• 1-2(W/W): *3.7Å*; 1-3(W/W): 2.6Å; 1-4(W/W): 2.6Å; 1-5(W/W): 2.7Å; 1-6(W/W): 2.9Å; 2-3(W/W): 2.9Å; 2-4(W/W): 3.0Å; 2-5(W/W): 2.9Å; 2-6(W/W): 2.8Å; 3-4(W/W): 2.8Å; 3-5(W/W): 4.1Å; 3-6(W/W): 2.8Å; 4-5(W/W): 3.2Å; 4-6(W/W): *4.3Å*; 5-6(W/W): 3.1Å;   
  
**2.03 ± 0.16: the average coodination distance of the 1^st^ six ligands is Mg^2+^ compatible**  
**12: Number of potential Water contacts**   
**6 : Number of second shell contacts involving water or ions**   
**50%: Percentage of satisfied second shell Water contacts**   
• Contact(s): /O_w_/O_w_/O_w_/O_w_/  
• Contact code: 4O_w_  
• Contact(s) in the exclusion zone: /O_w_/O_w_/  
→ Missing octahedral coordination for MG ←  
→ Slightly distorded coord. octahedron (6.3º) ←  
Go to ToC

### MG:A:1679

|  | Res. type | Atom | Residue | Distance | Occupancy | B-factor | Closest atom(s) to a 1^st^ shell water | EM\_dens. |
| --- | --- | --- | --- | --- | --- | --- | --- | --- |
| **Ion** | | | |  | 1.0 | 69.9 |  | 3.48 |
 1 | A | OP2 | A:509 | 2.01 | 1.0 | 64.9 |  | 3.19 | 2 | Water 1 | | A:2609 | 2.03 | 1.0 | 63.8 |  | 1.94 | 3 | Water 2 | | A:3068 | 2.08 | 1.0 | 59.9 |  | 2.98 | 4 | Water 3 | | A:2829 | 2.15 | 1.0 | 67.1 | 2.71 [U:O3']A:508 // 2.80 [A:N7]A:510 | 2.85 | 5 | A | OP2 | A:510 | 2.16 | 1.0 | 63.8 |  | 3.07 | 6 | U | O3' | A:508 | 3.45 | 1.0 | 65.2 |  | 3.40 | 7 | G | OP1 | A:506 | 3.92 | 1.0 | 62.1 |  | 2.26 | 8 | A | O5' | A:509 | 3.96 | 1.0 | 62.1 |  | 3.26 | 9 | C | OP2 | A:507 | 4.05 | 1.0 | 62.0 |  | 2.60 | 10 | A | N7 | A:510 | 4.29 | 1.0 | 61.4 |  | 3.34 | 11 | A | O5' | A:510 | 4.33 | 1.0 | 63.3 |  | 3.92 | 12 | A | OP1 | A:509 | 4.46 | 1.0 | 64.9 |  | 2.22 | 13 | A | OP1 | A:510 | 4.47 | 1.0 | 64.3 |  | 3.15 | 14 | U | O2' | A:508 | 4.55 | 1.0 | 62.3 |  | 2.99 | 15 | A | O3' | A:509 | 4.56 | 1.0 | 68.7 |  | 2.73 | 16 | Water 4 | | A:2451 | 4.70 | 1.0 | 64.3 |  | 1.64 |

  
**1^st^ shell coordination angles and distances (*trans* in "blue"; *cys* in "green"; deviations in "orange"):**   
• 1-2(OP2/W): 90º; 1-3(OP2/W): *164º*; 1-4(OP2/W): 87º; 1-5(OP2/OP2): *83º*; 2-3(W/W): 88º; 2-4(W/W): *164º*; 2-5(W/OP2): *79º*; 3-4(W/W): 91º; 3-5(W/OP2): *81º*; 4-5(W/OP2): 85º;   
• 1-2(OP2/W): 2.9Å; 1-3(OP2/W): 4.1Å; 1-4(OP2/W): 2.9Å; 1-5(OP2/OP2): 2.8Å; 2-3(W/W): 2.9Å; 2-4(W/W): 4.1Å; 2-5(W/OP2): 2.7Å; 3-4(W/W): 3.0Å; 3-5(W/OP2): 2.8Å; 4-5(W/OP2): 2.9Å;   
  
**2.31 ± 0.51: the average coodination distance of the 1^st^ six ligands is Na^+^** compatible  
**6: Number of potential Water contacts**   
**2 : Number of second shell contacts involving water or ions**   
**33%: Percentage of satisfied second shell Water contacts**   
• Contact(s): /O_w_/O_w_/O_w_/OP2/OP2/  
• Contact code: *cis-*2O_ph_3O_w_  
→ Missing octahedral coordination for MG ←  
→ Slightly distorded coord. octahedron (7.0º) ←→2Fo-Fc density between 1.0 and 4.0 !   
Go to ToC

### MG:A:1680

|  | Res. type | Atom | Residue | Distance | Occupancy | B-factor | Closest atom(s) to a 1^st^ shell water | EM\_dens. |
| --- | --- | --- | --- | --- | --- | --- | --- | --- |
| **Ion** | | | |  | 1.0 | 61.9 |  | 5.13 |
 1 | A | OP2 | A:608 | 1.90 | 1.0 | 60.9 |  | 2.86 | 2 | Water 1 | | A:3356 | 1.90 | 1.0 | 61.0 | 2.82 [A:N1]A:630 | 3.55 | 3 | Water 2 | | A:2621 | 2.04 | 1.0 | 53.7 | 2.67 [A:N7]A:609 | 4.30 | 4 | Water 3 | | A:3642 | 2.10 | 1.0 | 57.5 | 3.00 [A:N7]A:608 | 2.86 | 5 | Water 4 | | A:2280 | 2.16 | 1.0 | 61.7 | 2.59 [U:O4]A:610 | 3.37 | 6 | A | O5' | A:608 | 3.66 | 1.0 | 54.4 |  | 4.16 | 7 | Water 5 | | A:2194 | 3.91 | 1.0 | 54.8 |  | 2.42 | 8 | A | OP1 | A:608 | 4.01 | 1.0 | 56.6 |  | 2.51 | 9 | A | OP1 | A:607 | 4.08 | 1.0 | 60.6 |  | 2.46 | 10 | A | N7 | A:609 | 4.13 | 1.0 | 50.5 |  | 5.23 | 11 | Water 6 | | A:2932 | 4.18 | 1.0 | 67.4 | 2.73 [U:O4]A:610 | 2.16 | 12 | Water 7 | | A:3284 | 4.25 | 1.0 | 63.9 | 2.80 [A:OP1]A:608 | 2.51 | 13 | U | O4 | A:610 | 4.26 | 1.0 | 57.8 |  | 3.15 | 14 | A | N1 | A:630 | 4.35 | 1.0 | 69.0 |  | 3.46 | 15 | A | O3' | A:607 | 4.39 | 1.0 | 55.8 |  | 4.19 | 16 | A | N7 | A:608 | 4.46 | 1.0 | 51.6 |  | 4.37 | 17 | Water 8 | | A:3768 | 4.64 | 1.0 | 62.8 | 2.90 [C:O2]A:631 | 1.96 | 18 | C | O2 | A:631 | 4.73 | 1.0 | 70.7 |  | 2.58 | 19 | A | OP2 | A:609 | 4.79 | 1.0 | 55.6 |  | 3.63 |

  
**1^st^ shell coordination angles and distances (*trans* in "blue"; *cys* in "green"; deviations in "orange"):**   
• 1-2(OP2/W): *95º*; 1-3(OP2/W): *101º*; 1-4(OP2/W): 91º; 1-5(OP2/W): 172º; 2-3(W/W): 88º; 2-4(W/W): 172º; 2-5(W/W): *84º*; 3-4(W/W): 95º; 3-5(W/W): 87º; 4-5(W/W): 89º;   
• 1-2(OP2/W): 2.8Å; 1-3(OP2/W): 3.1Å; 1-4(OP2/W): 2.9Å; 1-5(OP2/W): 4.1Å; 2-3(W/W): 2.7Å; 2-4(W/W): 4.0Å; 2-5(W/W): 2.7Å; 3-4(W/W): 3.0Å; 3-5(W/W): 2.9Å; 4-5(W/W): 3.0Å;   
  
**2.30 ± 0.62: the average coodination distance of the 1^st^ six ligands is Mg^2+^ compatible**  
**8: Number of potential Water contacts**   
**4 : Number of second shell contacts involving water or ions**   
**50%: Percentage of satisfied second shell Water contacts**   
• Contact(s): /O_w_/O_w_/O_w_/O_w_/OP2/  
• Contact code: O_ph_4O_w_  
→ Missing octahedral coordination for MG ←  
→ Slightly distorded coord. octahedron (5.1º) ←  
Go to ToC

### MG:A:1681

|  | Res. type | Atom | Residue | Distance | Occupancy | B-factor | Closest atom(s) to a 1^st^ shell water | EM\_dens. |
| --- | --- | --- | --- | --- | --- | --- | --- | --- |
| **Ion** | | | |  | 1.0 | 47.7 |  | 9.46 |
 1 | Water 1 | | A:2976 | 1.94 | 1.0 | 51.2 | 2.68 [K:K]A:1733 | 5.09 | 2 | G | OP2 | A:289 | 2.00 | 1.0 | 42.5 |  | 8.25 | 3 | A | OP2 | A:116 | 2.00 | 1.0 | 45.6 |  | 5.98 | 4 | Water 2 | | A:3382 | 2.04 | 1.0 | 46.3 |  | 6.29 | 5 | Water 3 | | A:2965 | 2.14 | 1.0 | 48.5 |  | 5.07 | 6 | G | OP2 | A:117 | 2.14 | 1.0 | 45.6 |  | 4.85 | 7 | G | O5' | A:289 | 3.83 | 1.0 | 42.6 |  | 6.94 | 8 | **K** | K | A:1733 | 3.94 | 1.0 | 30.0 |  | 14.26 | 9 | A | OP1 | A:116 | 4.15 | 1.0 | 43.9 |  | 6.91 | 10 | Water 4 | | A:2890 | 4.15 | 1.0 | 50.0 |  | 2.97 | 11 | G | O3' | A:115 | 4.27 | 1.0 | 48.5 |  | 6.77 | 12 | U | O4 | A:118 | 4.27 | 1.0 | 48.1 |  | 5.96 | 13 | G | OP1 | A:289 | 4.32 | 1.0 | 42.4 |  | 5.79 | 14 | A | O5' | A:116 | 4.36 | 1.0 | 52.5 |  | 4.98 | 15 | A | O3' | A:116 | 4.36 | 1.0 | 50.4 |  | 4.43 | 16 | Water 5 | | A:3485 | 4.39 | 1.0 | 54.5 | 2.84 [G:OP1]A:117 | 1.62 | 17 | A | O3' | A:288 | 4.42 | 1.0 | 46.9 |  | 5.27 | 18 | G | OP1 | A:117 | 4.42 | 1.0 | 49.5 |  | 3.47 | 19 | G | O5' | A:117 | 4.45 | 1.0 | 48.3 |  | 5.22 | 20 | G | N7 | A:117 | 4.49 | 1.0 | 42.0 |  | 7.75 | 21 | Water 6 | | A:2265 | 4.57 | 1.0 | 49.6 | 2.58 [G:OP1]A:289 | 2.64 | 22 | A | O5' | A:288 | 4.68 | 1.0 | 47.9 |  | 4.13 | 23 | Water 7 | | A:4465 | 4.75 | 1.0 | 54.8 |  | 1.72 | 24 | A | OP2 | A:288 | 4.76 | 1.0 | 47.8 |  | 3.68 |

  
**1^st^ shell coordination angles and distances (*trans* in "blue"; *cys* in "green"; deviations in "orange"):**   
• 1-2(W/OP2): 88º; 1-3(W/OP2): 91º; 1-4(W/W): 178º; 1-5(W/W): 95º; 1-6(W/OP2): 94º; 2-3(OP2/OP2): *99º*; 2-4(OP2/W): 93º; 2-5(OP2/W): 91º; 2-6(OP2/OP2): 170º; 3-4(OP2/W): 90º; 3-5(OP2/W): *169º*; 3-6(OP2/OP2): 90º; 4-5(W/W): *83º*; 4-6(W/OP2): *85º*; 5-6(W/OP2): *80º*;   
• 1-2(W/OP2): 2.7Å; 1-3(W/OP2): 2.8Å; 1-4(W/W): 4.0Å; 1-5(W/W): 3.0Å; 1-6(W/OP2): 3.0Å; 2-3(OP2/OP2): 3.0Å; 2-4(OP2/W): 2.9Å; 2-5(OP2/W): 2.9Å; 2-6(OP2/OP2): 4.1Å; 3-4(OP2/W): 2.9Å; 3-5(OP2/W): 4.1Å; 3-6(OP2/OP2): 2.9Å; 4-5(W/W): 2.8Å; 4-6(W/OP2): 2.8Å; 5-6(W/OP2): 2.7Å;   
  
**2.04 ± 0.07: the average coodination distance of the 1^st^ six ligands is Mg^2+^ compatible**  
**6: Number of potential Water contacts**   
**1 : Number of second shell contacts involving water or ions**   
**17%: Percentage of satisfied second shell Water contacts**   
• Contact(s): /O_w_/O_w_/O_w_/OP2/OP2/OP2/  
• Contact code: *mer-*3O_ph_3O_w_  
Ooctahedral coordination (4.7º)   
Go to ToC

### MG:A:1682

|  | Res. type | Atom | Residue | Distance | Occupancy | B-factor | Closest atom(s) to a 1^st^ shell water | EM\_dens. |
| --- | --- | --- | --- | --- | --- | --- | --- | --- |
| **Ion** | | | |  | 1.0 | 51.9 |  | 6.96 |
 1 | Water 1 | | A:2665 | 2.67 | 1.0 | 43.5 | 2.68 [G:N7]A:773 | 6.07 | 2 | G | O6 | A:773 | 2.69 | 1.0 | 40.8 |  | 9.35 | 3 | Water 2 | | A:3426 | 2.71 | 1.0 | 50.8 | 2.83 [G:OP1]A:803 | 4.30 | 4 | Water 3 | | A:3780 | 2.71 | 1.0 | 47.8 | 2.91 [G:N7]A:774 | 4.31 | 5 | Water 4 | | A:4097 | 2.83 | 1.0 | 48.4 | 3.00 [G:OP1]A:803 | 2.96 | 6 | Water 5 | | A:2775 | 2.90 | 1.0 | 49.5 |  | 2.12 | 7 | G | O6 | A:774 | 3.00 | 1.0 | 39.2 |  | 8.69 | 8 | Water 6 | | A:2337 | 3.21 | 1.0 | 37.7 | 2.11 [MG:MG]A:1684 | 7.95 | 9 | G | N7 | A:774 | 3.47 | 1.0 | 37.9 |  | 6.48 | 10 | Water 7 | | A:4534 | 3.55 | 1.0 | 38.5 | 2.13 [MG:MG]A:1684 | 7.69 | 11 | G | N7 | A:773 | 3.97 | 1.0 | 35.9 |  | 8.53 | 12 | Water 8 | | A:4672 | 3.98 | 1.0 | 52.8 |  | 1.50 | 13 | U | O4 | A:772 | 4.47 | 1.0 | 32.5 |  | 11.38 | 14 | G | OP1 | A:803 | 4.62 | 1.0 | 40.9 |  | 7.46 | 15 | **MG** | MG | A:1684 | 4.72 | 1.0 | 40.5 |  | 11.83 |

  
• • **2.75 ± 0.08: the average coodination distance of the 1^st^ six ligands is K^+^ compatible**  
**10: Number of potential Water contacts**   
**4 : Number of second shell contacts involving water or ions**   
**40%: Percentage of satisfied second shell Water contacts**   
• Contact code: → No contacts ←   
• Contact(s) in the exclusion zone: /O6/O6/O_w_/O_w_/O_w_/O_w_/O_w_/O_w_/  
→ Consider K in place of MG (2 valid contact(s) for K vs. 0 for MG ←)  
Go to ToC

### MG:A:1683

|  | Res. type | Atom | Residue | Distance | Occupancy | B-factor | Closest atom(s) to a 1^st^ shell water | EM\_dens. |
| --- | --- | --- | --- | --- | --- | --- | --- | --- |
| **Ion** | | | |  | 1.0 | 57.1 |  | 5.45 |
 1 | Water 1 | | A:1917 | 1.83 | 1.0 | 50.4 |  | 3.27 | 2 | Water 2 | | A:2945 | 1.90 | 1.0 | 54.8 | 2.73 [A:O2']A:329 // 2.94 [A:N3]A:329 | 2.58 | 3 | G | OP2 | A:331 | 2.03 | 1.0 | 50.8 |  | 5.14 | 4 | Water 3 | | A:3215 | 2.07 | 1.0 | 53.6 | 2.84 [A:OP2]A:109 | 2.67 | 5 | Water 4 | | A:3252 | 2.08 | 1.0 | 47.5 |  | 3.09 | 6 | A | OP1 | A:109 | 2.23 | 1.0 | 52.2 |  | 4.15 | 7 | A | OP2 | A:109 | 3.80 | 1.0 | 48.1 |  | 4.81 | 8 | Water 5 | | A:3487 | 3.85 | 1.0 | 52.2 |  | 1.98 | 9 | Water 6 | | A:2152 | 4.03 | 1.0 | 55.6 | 2.54 [A:OP2]A:109 | 1.96 | 10 | A | O5' | A:109 | 4.22 | 1.0 | 46.3 |  | 5.85 | 11 | A | O2' | A:329 | 4.22 | 1.0 | 50.1 |  | 5.90 | 12 | G | OP1 | A:331 | 4.26 | 1.0 | 53.0 |  | 3.17 | 13 | Water 7 | | A:2238 | 4.29 | 1.0 | 44.7 |  | 4.32 | 14 | C | O3' | A:330 | 4.34 | 1.0 | 54.5 |  | 3.85 | 15 | G | O5' | A:331 | 4.34 | 1.0 | 47.9 |  | 6.13 | 16 | Water 8 | | A:3578 | 4.40 | 1.0 | 52.6 |  | 3.12 | 17 | A | N3 | A:329 | 4.46 | 1.0 | 40.7 |  | 6.63 | 18 | Water 9 | | A:3570 | 4.60 | 1.0 | 53.7 | 2.86 [G:O3']A:108 | 2.32 | 19 | G | O3' | A:108 | 4.65 | 1.0 | 51.8 |  | 4.04 |

  
**1^st^ shell coordination angles and distances (*trans* in "blue"; *cys* in "green"; deviations in "orange"):**   
• 1-2(W/W): *155º*; 1-3(W/OP2): *81º*; 1-4(W/W): *77º*; 1-5(W/W): 86º; 1-6(W/OP1): *72º*; 2-3(W/OP2): *99º*; 2-4(W/W): *128º*; 2-5(W/W): *69º*; 2-6(W/OP1): *108º*; 3-4(OP2/W): 86º; 3-5(OP2/W): *104º*; 3-6(OP2/OP1): *152º*; 4-5(W/W): *160º*; 4-6(W/OP1): *83º*; 5-6(W/OP1): *81º*;   
• 1-2(W/W): *3.6Å*; 1-3(W/OP2): 2.5Å; 1-4(W/W): 2.4Å; 1-5(W/W): 2.7Å; 1-6(W/OP1): 2.4Å; 2-3(W/OP2): 3.0Å; 2-4(W/W): *3.6Å*; 2-5(W/W): 2.3Å; 2-6(W/OP1): *3.3Å*; 3-4(OP2/W): 2.8Å; 3-5(OP2/W): *3.2Å*; 3-6(OP2/OP1): 4.1Å; 4-5(W/W): 4.1Å; 4-6(W/OP1): 2.9Å; 5-6(W/OP1): 2.8Å;   
  
**2.02 ± 0.13: the average coodination distance of the 1^st^ six ligands is Mg^2+^ compatible**  
**8: Number of potential Water contacts**   
**3 : Number of second shell contacts involving water or ions**   
**38%: Percentage of satisfied second shell Water contacts**   
• Contact(s): /O_w_/O_w_/OP1/OP2/  
• Contact code: *trans-*2O_ph_2O_w_  
• Contact(s) in the exclusion zone: /O_w_/O_w_/  
→ Missing octahedral coordination for MG ←  
→→→ Distorded coord. octahedron (15.7º) ←←←  
Go to ToC

### MG:A:1684

|  | Res. type | Atom | Residue | Distance | Occupancy | B-factor | Closest atom(s) to a 1^st^ shell water | EM\_dens. |
| --- | --- | --- | --- | --- | --- | --- | --- | --- |
| **Ion** | | | |  | 1.0 | 40.5 |  | 11.83 |
 1 | Water 1 | | A:2483 | 2.01 | 1.0 | 35.2 | 2.64 [G:OP1]A:803 | 7.47 | 2 | Water 2 | | A:4576 | 2.03 | 1.0 | 35.8 |  | 8.16 | 3 | Water 3 | | A:3189 | 2.10 | 1.0 | 39.4 | 2.78 [G:O6]A:771 | 8.57 | 4 | Water 4 | | A:2337 | 2.11 | 1.0 | 37.7 | 2.61 [U:O4]A:772 | 7.95 | 5 | Water 5 | | A:4534 | 2.13 | 1.0 | 38.5 |  | 7.69 | 6 | Water 6 | | A:2654 | 2.14 | 1.0 | 36.7 | 2.68 [G:N7]A:771 | 8.46 | 7 | Water 7 | | A:2889 | 3.86 | 1.0 | 34.6 |  | 8.02 | 8 | Water 8 | | A:4065 | 4.07 | 1.0 | 46.1 |  | 4.66 | 9 | U | O4 | A:772 | 4.15 | 1.0 | 32.5 |  | 11.38 | 10 | Water 9 | | A:4598 | 4.30 | 1.0 | 44.7 |  | 5.42 | 11 | G | OP1 | A:803 | 4.31 | 1.0 | 40.9 |  | 7.46 | 12 | G | N7 | A:771 | 4.32 | 1.0 | 30.7 |  | 10.79 | 13 | Water 10 | | A:4097 | 4.36 | 1.0 | 48.4 | 2.83 [MG:MG]A:1682 | 2.96 | 14 | Water 11 | | A:3703 | 4.37 | 1.0 | 39.5 |  | 8.22 | 15 | G | O6 | A:771 | 4.49 | 1.0 | 35.0 |  | 10.47 | 16 | Water 12 | | A:3426 | 4.56 | 1.0 | 50.8 | 2.71 [MG:MG]A:1682 | 4.30 | 17 | U | OP2 | A:804 | 4.68 | 1.0 | 37.7 |  | 8.12 | 18 | **MG** | MG | A:1682 | 4.72 | 1.0 | 51.9 |  | 6.96 |

  
**1^st^ shell coordination angles and distances (*trans* in "blue"; *cys* in "green"; deviations in "orange"):**   
• 1-2(W/W): 93º; 1-3(W/W): 173º; 1-4(W/W): 92º; 1-5(W/W): 88º; 1-6(W/W): 90º; 2-3(W/W): 94º; 2-4(W/W): *168º*; 2-5(W/W): 91º; 2-6(W/W): 94º; 3-4(W/W): *82º*; 3-5(W/W): 94º; 3-6(W/W): 88º; 4-5(W/W): *78º*; 4-6(W/W): *98º*; 5-6(W/W): 175º;   
• 1-2(W/W): 2.9Å; 1-3(W/W): 4.1Å; 1-4(W/W): 3.0Å; 1-5(W/W): 2.9Å; 1-6(W/W): 2.9Å; 2-3(W/W): 3.0Å; 2-4(W/W): 4.1Å; 2-5(W/W): 3.0Å; 2-6(W/W): 3.0Å; 3-4(W/W): 2.8Å; 3-5(W/W): 3.1Å; 3-6(W/W): 2.9Å; 4-5(W/W): 2.7Å; 4-6(W/W): 3.2Å; 5-6(W/W): *4.3Å*;   
  
**2.09 ± 0.05: the average coodination distance of the 1^st^ six ligands is Mg^2+^ compatible**  
**12: Number of potential Water contacts**   
**4 : Number of second shell contacts involving water or ions**   
**33%: Percentage of satisfied second shell Water contacts**   
• Contact(s): /O_w_/O_w_/O_w_/O_w_/O_w_/O_w_/  
• Contact code: 6O_w_  
Ooctahedral coordination (4.8º)   
Go to ToC

### MG:A:1685

|  | Res. type | Atom | Residue | Distance | Occupancy | B-factor | Closest atom(s) to a 1^st^ shell water | EM\_dens. |
| --- | --- | --- | --- | --- | --- | --- | --- | --- |
| **Ion** | | | |  | 1.0 | 71.3 |  | 2.64 |
 1 | Water 1 | | A:4049 | 1.90 | 1.0 | 64.0 | 2.98 [U:OP1]A:952 | 2.07 | 2 | Water 2 | | A:3339 | 1.97 | 1.0 | 64.1 | 2.81 [G:N7]A:963 | 1.95 | 3 | Water 3 | | A:1830 | 2.48 | 1.0 | 68.6 | 2.30 [G:OP2]A:953 | 1.83 | 4 | Water 4 | | A:2625 | 3.71 | 1.0 | 68.8 |  | 2.21 | 5 | G | OP1 | A:953 | 4.23 | 1.0 | 68.5 |  | 2.69 | 6 | G | OP2 | A:953 | 4.26 | 1.0 | 68.9 |  | 2.55 | 7 | U | OP1 | A:952 | 4.54 | 1.0 | 62.8 |  | 3.76 | 8 | G | N7 | A:963 | 4.64 | 1.0 | 56.2 |  | 4.08 | 9 | G | O6 | A:963 | 4.73 | 1.0 | 60.4 |  | 3.25 |

  
**1^st^ shell coordination angles and distances (*trans* in "blue"; *cys* in "green"; deviations in "orange"):**   
• 1-2(W/W): *77º*;   
• 1-2(W/W): 2.4Å;   
  
**3.09 ± 1.01: the average coodination distance of the 1^st^ six ligands is K^+^ compatible**  
**8: Number of potential Water contacts**   
**3 : Number of second shell contacts involving water or ions**   
**38%: Percentage of satisfied second shell Water contacts**   
• Contact(s): /O_w_/O_w_/  
• Contact code: 2O_w_  
• Contact(s) in the exclusion zone: /O_w_/  
→ Missing octahedral coordination for MG ←  
• MG hydration shell issue: water too close (< 2.4 Å.)  
→2Fo-Fc density between 1.0 and 4.0 !   
Go to ToC

### MG:A:1686

|  | Res. type | Atom | Residue | Distance | Occupancy | B-factor | Closest atom(s) to a 1^st^ shell water | EM\_dens. |
| --- | --- | --- | --- | --- | --- | --- | --- | --- |
| **Ion** | | | |  | 1.0 | 65.2 |  | 4.38 |
 1 | Water 1 | | A:3106 | 1.80 | 1.0 | 58.2 | 2.85 [C:OP1]A:1109 // 2.87 [U:O2]A:1189 | 3.27 | 2 | Water 2 | | A:3316 | 1.88 | 1.0 | 61.2 | 2.92 [A:N7]A:1111 | 3.34 | 3 | A | OP2 | A:1110 | 1.98 | 1.0 | 55.3 |  | 5.24 | 4 | Water 3 | | A:4577 | 1.98 | 1.0 | 60.8 |  | 2.86 | 5 | Water 4 | | A:1948 | 2.10 | 1.0 | 62.8 | 2.43 [A:N7]A:1110 | 3.63 | 6 | Water 5 | | A:2416 | 2.26 | 1.0 | 61.6 |  | 2.17 | 7 | A | O5' | A:1110 | 3.83 | 1.0 | 57.9 |  | 4.49 | 8 | C | OP2 | A:1109 | 3.85 | 1.0 | 55.9 |  | 3.39 | 9 | U | O2 | A:1189 | 3.95 | 1.0 | 62.7 |  | 2.74 | 10 | C | OP1 | A:1109 | 4.00 | 1.0 | 58.8 |  | 2.96 | 11 | A | N7 | A:1110 | 4.22 | 1.0 | 58.3 |  | 3.90 | 12 | Water 6 | | A:3743 | 4.25 | 1.0 | 61.3 |  | 2.33 | 13 | A | N7 | A:1111 | 4.26 | 1.0 | 58.2 |  | 3.84 | 14 | A | OP1 | A:1110 | 4.27 | 1.0 | 61.7 |  | 3.10 | 15 | C | O3' | A:1109 | 4.42 | 1.0 | 59.5 |  | 2.69 | 16 | Water 7 | | A:3341 | 4.53 | 1.0 | 64.4 |  | 2.07 | 17 | Water 8 | | A:2847 | 4.66 | 1.0 | 60.9 |  | 2.76 | 18 | C | O5' | A:1109 | 4.71 | 1.0 | 53.8 |  | 5.52 |

  
**1^st^ shell coordination angles and distances (*trans* in "blue"; *cys* in "green"; deviations in "orange"):**   
• 1-2(W/W): *160º*; 1-3(W/OP2): 94º; 1-4(W/W): 87º; 1-5(W/W): *112º*; 1-6(W/W): *80º*; 2-3(W/OP2): 93º; 2-4(W/W): *80º*; 2-5(W/W): *84º*; 2-6(W/W): *83º*; 3-4(OP2/W): *162º*; 3-5(OP2/W): *100º*; 3-6(OP2/W): *76º*; 4-5(W/W): *96º*; 4-6(W/W): 86º; 5-6(W/W): *167º*;   
• 1-2(W/W): *3.6Å*; 1-3(W/OP2): 2.8Å; 1-4(W/W): 2.6Å; 1-5(W/W): *3.2Å*; 1-6(W/W): 2.6Å; 2-3(W/OP2): 2.8Å; 2-4(W/W): 2.5Å; 2-5(W/W): 2.7Å; 2-6(W/W): 2.8Å; 3-4(OP2/W): 3.9Å; 3-5(OP2/W): 3.1Å; 3-6(OP2/W): 2.6Å; 4-5(W/W): 3.0Å; 4-6(W/W): 2.9Å; 5-6(W/W): *4.3Å*;   
  
**2.00 ± 0.15: the average coodination distance of the 1^st^ six ligands is Mg^2+^ compatible**  
**10: Number of potential Water contacts**   
**4 : Number of second shell contacts involving water or ions**   
**40%: Percentage of satisfied second shell Water contacts**   
• Contact(s): /O_w_/O_w_/O_w_/OP2/  
• Contact code: O_ph_3O_w_  
• Contact(s) in the exclusion zone: /O_w_/O_w_/  
→ Missing octahedral coordination for MG ←  
→ Slightly distorded coord. octahedron (9.9º) ←  
Go to ToC

### MG:A:1687

|  | Res. type | Atom | Residue | Distance | Occupancy | B-factor | Closest atom(s) to a 1^st^ shell water | EM\_dens. |
| --- | --- | --- | --- | --- | --- | --- | --- | --- |
| **Ion** | | | |  | 1.0 | 80.1 |  | 2.52 |
 1 | Water 1 | | A:4379 | 1.92 | 1.0 | 70.6 |  | 2.04 | 2 | Water 2 | | A:4205 | 2.00 | 1.0 | 78.8 |  | 1.69 | 3 | Water 3 | | A:2952 | 2.00 | 1.0 | 66.5 | 2.74 [A:OP1]A:1204 | 2.05 | 4 | Water 4 | | N:203 | 2.04 | 1.0 | 80.6 | 2.66 [ALA:O]N:2 // 2.84 [GLN:OE1]N:4 | 1.49 | 5 | ALA | O | N:2 | 3.83 | 1.0 | 75.1 |  | 2.18 | 6 | G | OP1 | A:1048 | 3.85 | 1.0 | 82.1 |  | 2.48 | 7 | Water 5 | | A:2098 | 3.87 | 1.0 | 77.1 | 2.92 [U:OP1]A:1049 | 1.63 | 8 | G | OP2 | A:1048 | 4.11 | 1.0 | 81.7 |  | 2.95 | 9 | U | OP1 | A:1049 | 4.29 | 1.0 | 73.6 |  | 2.06 | 10 | A | OP2 | A:1204 | 4.43 | 1.0 | 67.2 |  | 2.10 | 11 | A | OP1 | A:1204 | 4.45 | 1.0 | 68.4 |  | 1.30 | 12 | GLN | OE1 | N:4 | 4.59 | 1.0 | 80.6 |  | 1.78 |

  
**1^st^ shell coordination angles and distances (*trans* in "blue"; *cys* in "green"; deviations in "orange"):**   
• 1-2(W/W): *164º*; 1-3(W/W): 92º; 1-4(W/W): 90º; 2-3(W/W): *76º*; 2-4(W/W): *82º*; 3-4(W/W): 93º;   
• 1-2(W/W): 3.9Å; 1-3(W/W): 2.8Å; 1-4(W/W): 2.8Å; 2-3(W/W): 2.5Å; 2-4(W/W): 2.6Å; 3-4(W/W): 2.9Å;   
  
**2.61 ± 0.87: the average coodination distance of the 1^st^ six ligands is K^+^ compatible**  
**10: Number of potential Water contacts**   
**3 : Number of second shell contacts involving water or ions**   
**30%: Percentage of satisfied second shell Water contacts**   
• Contact(s): /O_w_/O_w_/O_w_/O_w_/  
• Contact code: 4O_w_  
→ Missing octahedral coordination for MG ←  
→ MG ion B\_factor > 80 ←→ Slightly distorded coord. octahedron (7.2º) ←→2Fo-Fc density between 1.0 and 4.0 !   
Go to ToC

### MG:A:1688

|  | Res. type | Atom | Residue | Distance | Occupancy | B-factor | Closest atom(s) to a 1^st^ shell water | EM\_dens. |
| --- | --- | --- | --- | --- | --- | --- | --- | --- |
| **Ion** | | | |  | 1.0 | 76.1 |  | 1.75 |
 1 | Water 1 | | A:3916 | 1.99 | 1.0 | 72.1 | 2.95 [G:N7]A:1353 | 1.60 | 2 | Water 2 | | A:4129 | 2.00 | 1.0 | 71.9 |  | 1.74 | 3 | Water 3 | | A:4430 | 2.06 | 1.0 | 69.0 |  | 1.61 | 4 | G | N7 | A:1353 | 4.02 | 1.0 | 71.5 |  | 2.64 | 5 | G | O6 | A:1353 | 4.40 | 1.0 | 74.1 |  | 2.10 | 6 | U | O4 | A:1354 | 4.54 | 1.0 | 70.8 |  | 2.40 |

  
**1^st^ shell coordination angles and distances (*trans* in "blue"; *cys* in "green"; deviations in "orange"):**   
• 1-2(W/W): *118º*; 1-3(W/W): 89º; 2-3(W/W): *96º*;   
• 1-2(W/W): *3.4Å*; 1-3(W/W): 2.8Å; 2-3(W/W): 3.0Å;   
  
**3.17 ± 1.16: the average coodination distance of the 1^st^ six ligands is K^+^ compatible**  
**6: Number of potential Water contacts**   
**1 : Number of second shell contacts involving water or ions**   
**17%: Percentage of satisfied second shell Water contacts**   
• Contact(s): /O_w_/O_w_/O_w_/  
• Contact code: 3O_w_  
→ Missing octahedral coordination for MG ←  
→2Fo-Fc density between 1.0 and 4.0 !   
Go to ToC

### MG:A:1689

|  | Res. type | Atom | Residue | Distance | Occupancy | B-factor | Closest atom(s) to a 1^st^ shell water | EM\_dens. |
| --- | --- | --- | --- | --- | --- | --- | --- | --- |
| **Ion** | | | |  | 1.0 | 80.7 |  | 3.56 |
 1 | C | OP1 | A:979 | 1.87 | 1.0 | 73.9 |  | 4.18 | 2 | Water 1 | | A:1813 | 1.92 | 1.0 | 76.7 |  | 2.33 | 3 | Water 2 | | A:2055 | 1.94 | 1.0 | 76.4 | 2.91 [G:O6]A:1221 | 2.27 | 4 | U | O4 | A:981 | 2.01 | 1.0 | 72.7 |  | 3.04 | 5 | C | OP2 | A:980 | 2.05 | 1.0 | 76.7 |  | 2.22 | 6 | G | O6 | A:1222 | 2.06 | 1.0 | 75.2 |  | 2.35 | 7 | C | O5' | A:979 | 3.30 | 1.0 | 75.1 |  | 2.30 | 8 | C | OP2 | A:979 | 3.55 | 1.0 | 74.6 |  | 3.00 | 9 | **K** | K | A:1644 | 3.56 | 1.0 | 30.0 |  | 4.65 | 10 | Water 3 | | A:3305 | 4.26 | 1.0 | 78.0 |  | 1.68 | 11 | C | OP1 | A:980 | 4.27 | 1.0 | 72.5 |  | 3.33 | 12 | A | O3' | A:978 | 4.28 | 1.0 | 74.9 |  | 3.24 | 13 | G | N7 | A:1222 | 4.30 | 1.0 | 75.1 |  | 2.91 | 14 | C | O3' | A:979 | 4.33 | 1.0 | 75.2 |  | 3.23 | 15 | G | O6 | A:1221 | 4.39 | 1.0 | 79.0 |  | 2.59 | 16 | C | O5' | A:980 | 4.40 | 1.0 | 75.3 |  | 3.03 | 17 | U | O2 | A:982 | 4.49 | 1.0 | 69.1 |  | 3.20 | 18 | A | O2' | A:977 | 4.61 | 1.0 | 70.9 |  | 1.84 |

  
**1^st^ shell coordination angles and distances (*trans* in "blue"; *cys* in "green"; deviations in "orange"):**   
• 1-2(OP1/W): 172º; 1-3(OP1/W): *101º*; 1-4(OP1/O4): *104º*; 1-5(OP1/OP2): *112º*; 1-6(OP1/O6): *100º*; 2-3(W/W): *72º*; 2-4(W/O4): *83º*; 2-5(W/OP2): *70º*; 2-6(W/O6): *77º*; 3-4(W/O4): *154º*; 3-5(W/OP2): *78º*; 3-6(W/O6): *82º*; 4-5(O4/OP2): *97º*; 4-6(O4/O6): 89º; 5-6(OP2/O6): *145º*;   
• 1-2(OP1/W): *3.8Å*; 1-3(OP1/W): 2.9Å; 1-4(OP1/O4): 3.1Å; 1-5(OP1/OP2): *3.3Å*; 1-6(OP1/O6): 3.0Å; 2-3(W/W): 2.3Å; 2-4(W/O4): 2.6Å; 2-5(W/OP2): 2.3Å; 2-6(W/O6): 2.5Å; 3-4(W/O4): 3.9Å; 3-5(W/OP2): 2.5Å; 3-6(W/O6): 2.6Å; 4-5(O4/OP2): 3.0Å; 4-6(O4/O6): 2.9Å; 5-6(OP2/O6): 3.9Å;   
  
**1.98 ± 0.07: the average coodination distance of the 1^st^ six ligands is Mg^2+^ compatible**  
**4: Number of potential Water contacts**   
**1 : Number of second shell contacts involving water or ions**   
**25%: Percentage of satisfied second shell Water contacts**   
• Contact(s): /O4/O6/O_w_/O_w_/OP2/  
• Contact code: O_ph_2O_b_2O_w_  
• Contact(s) in the exclusion zone: /O5'/OP1/  
→ Missing octahedral coordination for MG ←  
→→→ Distorded coord. octahedron (14.3º) ←←←  
→ MG ion B\_factor > 80 ←→2Fo-Fc density between 1.0 and 4.0 !   
Go to ToC

### MG:A:1690

|  | Res. type | Atom | Residue | Distance | Occupancy | B-factor | Closest atom(s) to a 1^st^ shell water | EM\_dens. |
| --- | --- | --- | --- | --- | --- | --- | --- | --- |
| **Ion** | | | |  | 1.0 | 70.6 |  | 3.27 |
 1 | Water 1 | | A:2919 | 1.87 | 1.0 | 71.2 | 2.73 [G:O6]A:1370 // 2.75 [G:N7]A:1370 | 2.87 | 2 | Water 2 | | A:3609 | 1.96 | 1.0 | 70.3 | 2.86 [G:O6]A:1370 | 2.90 | 3 | Water 3 | | A:4186 | 1.97 | 1.0 | 68.0 |  | 2.50 | 4 | G | O6 | A:1370 | 2.34 | 1.0 | 70.4 |  | 3.13 | 5 | Water 4 | | A:2340 | 3.76 | 1.0 | 67.6 |  | 2.15 | 6 | G | N7 | A:1370 | 3.82 | 1.0 | 70.4 |  | 2.98 | 7 | G | O6 | A:1371 | 4.11 | 1.0 | 71.4 |  | 2.34 | 8 | Water 5 | | A:1818 | 4.14 | 1.0 | 66.4 |  | 2.48 | 9 | U | O4 | A:1351 | 4.38 | 1.0 | 68.9 |  | 2.82 | 10 | Water 6 | | A:1916 | 4.48 | 1.0 | 63.1 |  | 2.96 | 11 | Water 7 | | A:4522 | 4.65 | 1.0 | 67.3 |  | 1.89 | 12 | Water 8 | | I:204 | 4.68 | 1.0 | 65.3 |  | 1.96 | 13 | G | N7 | A:1371 | 4.72 | 1.0 | 69.0 |  | 3.24 |

  
**1^st^ shell coordination angles and distances (*trans* in "blue"; *cys* in "green"; deviations in "orange"):**   
• 1-2(W/W): 95º; 1-3(W/W): *82º*; 2-3(W/W): 172º;   
• 1-2(W/W): 2.8Å; 1-3(W/W): 2.5Å; 2-3(W/W): 3.9Å;   
  
**2.62 ± 0.84: the average coodination distance of the 1^st^ six ligands is K^+^ compatible**  
**8: Number of potential Water contacts**   
**3 : Number of second shell contacts involving water or ions**   
**38%: Percentage of satisfied second shell Water contacts**   
• Contact(s): /O_w_/O_w_/  
• Contact code: 2O_w_  
• Contact(s) in the exclusion zone: /O6/O_w_/  
→ Missing octahedral coordination for MG ←  
→2Fo-Fc density between 1.0 and 4.0 !   
Go to ToC

### MG:A:1691

|  | Res. type | Atom | Residue | Distance | Occupancy | B-factor | Closest atom(s) to a 1^st^ shell water | EM\_dens. |
| --- | --- | --- | --- | --- | --- | --- | --- | --- |
| **Ion** | | | |  | 1.0 | 73.6 |  | 3.39 |
 1 | Water 1 | | A:3495 | 1.88 | 1.0 | 70.2 | 2.84 [A:N7]A:946 | 2.99 | 2 | Water 2 | | A:2396 | 1.89 | 1.0 | 69.2 | 2.62 [A:OP2]A:946 | 2.51 | 3 | Water 3 | | A:2995 | 1.96 | 1.0 | 69.5 | 2.74 [G:OP2]A:945 | 1.89 | 4 | G | OP1 | A:944 | 1.97 | 1.0 | 62.4 |  | 4.22 | 5 | Water 4 | | A:3759 | 2.05 | 1.0 | 68.7 |  | 2.24 | 6 | G | OP2 | A:945 | 2.31 | 1.0 | 70.2 |  | 2.95 | 7 | G | OP2 | A:944 | 3.47 | 1.0 | 62.7 |  | 3.56 | 8 | G | O5' | A:944 | 3.68 | 1.0 | 64.9 |  | 3.65 | 9 | Water 5 | | A:4322 | 3.97 | 1.0 | 68.8 |  | 1.50 | 10 | G | O5' | A:945 | 4.25 | 1.0 | 68.9 |  | 3.53 | 11 | **K** | K | A:1658 | 4.32 | 1.0 | 30.0 |  | 3.45 | 12 | A | OP2 | A:946 | 4.37 | 1.0 | 71.9 |  | 2.37 | 13 | U | O3' | A:943 | 4.41 | 1.0 | 60.3 |  | 4.70 | 14 | A | N7 | A:946 | 4.46 | 1.0 | 69.6 |  | 2.87 | 15 | G | OP1 | A:945 | 4.56 | 1.0 | 68.6 |  | 2.97 | 16 | Water 6 | | A:4238 | 4.61 | 1.0 | 65.6 |  | 2.44 | 17 | G | O3' | A:944 | 4.74 | 1.0 | 64.6 |  | 3.24 |

  
**1^st^ shell coordination angles and distances (*trans* in "blue"; *cys* in "green"; deviations in "orange"):**   
• 1-2(W/W): *80º*; 1-3(W/W): *161º*; 1-4(W/OP1): *103º*; 1-5(W/W): 86º; 2-3(W/W): *82º*; 2-4(W/OP1): 172º; 2-5(W/W): *95º*; 3-4(W/OP1): *95º*; 3-5(W/W): 89º; 4-5(OP1/W): 92º;   
• 1-2(W/W): 2.4Å; 1-3(W/W): *3.8Å*; 1-4(W/OP1): 3.0Å; 1-5(W/W): 2.7Å; 2-3(W/W): 2.5Å; 2-4(W/OP1): 3.9Å; 2-5(W/W): 2.9Å; 3-4(W/OP1): 2.9Å; 3-5(W/W): 2.8Å; 4-5(OP1/W): 2.9Å;   
  
**2.01 ± 0.14: the average coodination distance of the 1^st^ six ligands is Mg^2+^ compatible**  
**8: Number of potential Water contacts**   
**3 : Number of second shell contacts involving water or ions**   
**38%: Percentage of satisfied second shell Water contacts**   
• Contact(s): /O_w_/O_w_/OP1/  
• Contact code: O_ph_2O_w_  
• Contact(s) in the exclusion zone: /O_w_/O_w_/OP2/  
→ Missing octahedral coordination for MG ←  
→ Slightly distorded coord. octahedron (7.5º) ←→2Fo-Fc density between 1.0 and 4.0 !   
Go to ToC

### MG:A:1692

|  | Res. type | Atom | Residue | Distance | Occupancy | B-factor | Closest atom(s) to a 1^st^ shell water | EM\_dens. |
| --- | --- | --- | --- | --- | --- | --- | --- | --- |
| **Ion** | | | |  | 1.0 | 73.4 |  | 3.38 |
 1 | Water 1 | | N:206 | 1.85 | 1.0 | 73.4 |  | 2.00 | 2 | Water 2 | | N:205 | 1.89 | 1.0 | 78.7 | 2.92 [SER:O]N:58 | 2.10 | 3 | Water 3 | | A:4130 | 1.93 | 1.0 | 72.8 |  | 2.92 | 4 | Water 4 | | A:2790 | 1.97 | 1.0 | 68.6 | 2.70 [C:N3]A:980 | 2.12 | 5 | Water 5 | | N:208 | 2.44 | 1.0 | 75.5 |  | 1.76 | 6 | C | O2 | A:980 | 3.86 | 1.0 | 73.8 |  | 2.50 | 7 | GLN | O | N:60 | 3.95 | 1.0 | 78.9 |  | 1.82 | 8 | C | N3 | A:980 | 4.02 | 1.0 | 72.2 |  | 2.85 | 9 | G | O6 | A:1361 | 4.09 | 1.0 | 75.0 |  | 1.72 | 10 | Water 6 | | A:3667 | 4.11 | 1.0 | 70.7 | 2.96 [C:N4]A:979 | 1.97 | 11 | Water 7 | | A:3269 | 4.30 | 1.0 | 74.4 |  | 2.03 | 12 | SER | O | N:58 | 4.35 | 1.0 | 77.5 |  | 1.96 | 13 | A | N7 | A:1360 | 4.56 | 1.0 | 73.1 |  | 3.02 | 14 | A | OP1 | A:1360 | 4.64 | 1.0 | 70.3 |  | 3.32 | 15 | C | O3' | A:1359 | 4.66 | 1.0 | 71.0 |  | 3.10 |

  
**1^st^ shell coordination angles and distances (*trans* in "blue"; *cys* in "green"; deviations in "orange"):**   
• 1-2(W/W): *79º*; 1-3(W/W): *75º*; 1-4(W/W): 175º; 2-3(W/W): *82º*; 2-4(W/W): *105º*; 3-4(W/W): *105º*;   
• 1-2(W/W): 2.4Å; 1-3(W/W): 2.3Å; 1-4(W/W): 3.8Å; 2-3(W/W): 2.5Å; 2-4(W/W): 3.1Å; 3-4(W/W): 3.1Å;   
  
**2.32 ± 0.71: the average coodination distance of the 1^st^ six ligands is Na^+^** compatible  
**10: Number of potential Water contacts**   
**2 : Number of second shell contacts involving water or ions**   
**20%: Percentage of satisfied second shell Water contacts**   
• Contact(s): /O_w_/O_w_/  
• Contact code: 2O_w_  
• Contact(s) in the exclusion zone: /O_w_/O_w_/O_w_/  
→ Missing octahedral coordination for MG ←  
→→→ Distorded coord. octahedron (11.6º) ←←←  
→2Fo-Fc density between 1.0 and 4.0 !   
Go to ToC

### MG:A:1693

|  | Res. type | Atom | Residue | Distance | Occupancy | B-factor | Closest atom(s) to a 1^st^ shell water | EM\_dens. |
| --- | --- | --- | --- | --- | --- | --- | --- | --- |
| **Ion** | | | |  | 1.0 | 61.4 |  | 4.45 |
 1 | C | OP2 | A:1069 | 1.84 | 1.0 | 52.1 |  | 5.02 | 2 | Water 1 | | A:4225 | 1.85 | 1.0 | 56.0 |  | 2.79 | 3 | Water 2 | | A:3424 | 1.96 | 1.0 | 54.1 | 2.82 [G:O6]A:1094 | 3.54 | 4 | Water 3 | | A:3760 | 1.99 | 1.0 | 55.0 | 2.90 [G:N7]A:1094 | 2.73 | 5 | Water 4 | | A:3563 | 2.05 | 1.0 | 58.7 |  | 1.92 | 6 | Water 5 | | A:3943 | 2.38 | 1.0 | 55.2 |  | 2.61 | 7 | C | O5' | A:1069 | 3.95 | 1.0 | 49.8 |  | 5.22 | 8 | C | OP1 | A:1069 | 4.02 | 1.0 | 55.8 |  | 4.06 | 9 | G | N7 | A:1094 | 4.21 | 1.0 | 56.7 |  | 3.65 | 10 | Water 6 | | A:4607 | 4.21 | 1.0 | 57.5 |  | 1.47 | 11 | G | O6 | A:1094 | 4.29 | 1.0 | 57.3 |  | 4.69 | 12 | G | O3' | A:1068 | 4.31 | 1.0 | 53.8 |  | 3.68 | 13 | U | O2 | A:1085 | 4.36 | 1.0 | 54.5 |  | 4.78 | 14 | Water 7 | | A:3187 | 4.62 | 1.0 | 55.0 | 2.78 [C:OP1]A:1069 | 1.98 | 15 | Water 8 | | A:4299 | 4.75 | 1.0 | 55.5 |  | 2.35 |

  
**1^st^ shell coordination angles and distances (*trans* in "blue"; *cys* in "green"; deviations in "orange"):**   
• 1-2(OP2/W): *167º*; 1-3(OP2/W): *98º*; 1-4(OP2/W): *99º*; 1-5(OP2/W): 95º; 2-3(W/W): 87º; 2-4(W/W): 92º; 2-5(W/W): *73º*; 3-4(W/W): *103º*; 3-5(W/W): *85º*; 4-5(W/W): *163º*;   
• 1-2(OP2/W): *3.7Å*; 1-3(OP2/W): 2.9Å; 1-4(OP2/W): 2.9Å; 1-5(OP2/W): 2.9Å; 2-3(W/W): 2.6Å; 2-4(W/W): 2.8Å; 2-5(W/W): 2.3Å; 3-4(W/W): 3.1Å; 3-5(W/W): 2.7Å; 4-5(W/W): 4.0Å;   
  
**2.01 ± 0.18: the average coodination distance of the 1^st^ six ligands is Mg^2+^ compatible**  
**10: Number of potential Water contacts**   
**2 : Number of second shell contacts involving water or ions**   
**20%: Percentage of satisfied second shell Water contacts**   
• Contact(s): /O_w_/O_w_/O_w_/  
• Contact code: 3O_w_  
• Contact(s) in the exclusion zone: /O_w_/O_w_/OP2/  
→ Missing octahedral coordination for MG ←  
→ Slightly distorded coord. octahedron (9.2º) ←  
Go to ToC

### MG:A:1694

|  | Res. type | Atom | Residue | Distance | Occupancy | B-factor | Closest atom(s) to a 1^st^ shell water | EM\_dens. |
| --- | --- | --- | --- | --- | --- | --- | --- | --- |
| **Ion** | | | |  | 1.0 | 77.0 |  | 2.23 |
 1 | Water 1 | | A:4080 | 1.93 | 1.0 | 80.8 | 2.99 [A:N7]A:1254 | 1.49 | 2 | Water 2 | | A:4113 | 2.07 | 1.0 | 81.8 |  | 1.75 | 3 | G | N7 | A:1279 | 4.13 | 1.0 | 85.9 |  | 2.60 | 4 | A | OP2 | A:1254 | 4.14 | 1.0 | 78.5 |  | 2.18 | 5 | A | N7 | A:1254 | 4.19 | 1.0 | 79.2 |  | 2.84 |

  
**1^st^ shell coordination angles and distances (*trans* in "blue"; *cys* in "green"; deviations in "orange"):**   
• 1-2(W/W): 94º;   
• 1-2(W/W): 2.9Å;   
  
**3.29 ± 1.06: the average coodination distance of the 1^st^ six ligands is K^+^ compatible**  
**4: Number of potential Water contacts**   
**1 : Number of second shell contacts involving water or ions**   
**25%: Percentage of satisfied second shell Water contacts**   
• Contact(s): /O_w_/O_w_/  
• Contact code: 2O_w_  
→ Missing octahedral coordination for MG ←  
→2Fo-Fc density between 1.0 and 4.0 !   
Go to ToC

### MG:A:1695

|  | Res. type | Atom | Residue | Distance | Occupancy | B-factor | Closest atom(s) to a 1^st^ shell water | EM\_dens. |
| --- | --- | --- | --- | --- | --- | --- | --- | --- |
| **Ion** | | | |  | 1.0 | 76.3 |  | 1.85 |
 1 | A | OP2 | A:1238 | 2.59 | 1.0 | 81.2 |  | 1.56 | 2 | U | O2 | A:1335 | 3.24 | 1.0 | 80.2 |  | 1.43 | 3 | U | O4 | A:1301 | 3.41 | 1.0 | 85.2 |  | 1.76 | 4 | A | N7 | A:1299 | 3.66 | 1.0 | 85.4 |  | 2.29 | 5 | G | N2 | A:1300 | 3.86 | 1.0 | 77.7 |  | 3.04 | 6 | A | OP1 | A:1238 | 4.28 | 1.0 | 79.2 |  | 1.82 | 7 | C | O3' | A:1237 | 4.32 | 1.0 | 74.8 |  | 3.82 | 8 | G | N3 | A:1300 | 4.70 | 1.0 | 79.2 |  | 3.75 |

  
• • **3.51 ± 0.53: the average coodination distance of the 1^st^ six ligands is K^+^ compatible**  
**0: Number of potential Water contacts**   
• Contact code: → No contacts ←   
• Contact(s) in the exclusion zone: /O2/OP2/  
→ Firts MG coordination shell missing ←  
→2Fo-Fc density between 1.0 and 4.0 !   
Go to ToC

### MG:A:1696

|  | Res. type | Atom | Residue | Distance | Occupancy | B-factor | Closest atom(s) to a 1^st^ shell water | EM\_dens. |
| --- | --- | --- | --- | --- | --- | --- | --- | --- |
| **Ion** | | | |  | 1.0 | 69.2 |  | 1.76 |
 1 | C | OP1 | A:1200 | 3.86 | 1.0 | 65.9 |  | 2.16 | 2 | C | OP2 | A:1051 | 3.99 | 1.0 | 76.1 |  | 2.15 | 3 | C | O2' | A:962 | 4.21 | 1.0 | 66.3 |  | 2.49 | 4 | C | OP1 | A:1051 | 4.34 | 1.0 | 74.7 |  | 1.88 | 5 | U | OP2 | A:1052 | 4.44 | 1.0 | 71.6 |  | 2.00 | 6 | Water 1 | | A:2228 | 4.45 | 1.0 | 61.1 |  | 2.03 | 7 | Water 2 | | A:2077 | 4.58 | 1.0 | 66.0 | 2.51 [C:O2']A:962 | 2.29 |

  
• • **4.21 ± 0.22: the average coodination distance of the 1^st^ six ligands could correspond to an hexahydrated Mg^2+^.**  
**2: Number of potential Water contacts**   
**0 : Number of second shell contacts involving water or ions**   
**0%: Percentage of satisfied second shell Water contacts**   
• Contact code: → No contacts ←   
→ Firts MG coordination shell missing ←  
→2Fo-Fc density between 1.0 and 4.0 !   
Go to ToC

### MG:A:1697

|  | Res. type | Atom | Residue | Distance | Occupancy | B-factor | Closest atom(s) to a 1^st^ shell water | EM\_dens. |
| --- | --- | --- | --- | --- | --- | --- | --- | --- |
| **Ion** | | | |  | 1.0 | 59.1 |  | 3.15 |
 1 | Water 1 | | A:3408 | 2.00 | 1.0 | 47.4 | 2.82 [G:N7]A:1077 | 3.75 | 2 | Water 2 | | A:3229 | 2.76 | 1.0 | 47.3 | 2.79 [G:OP2]A:1079 | 4.38 | 3 | Water 3 | | E:234 | 3.28 | 1.0 | 51.2 |  | 2.27 | 4 | G | OP1 | A:1079 | 3.42 | 1.0 | 48.0 |  | 4.70 | 5 | G | N7 | A:1077 | 3.77 | 1.0 | 37.7 |  | 7.85 | 6 | G | O6 | A:1077 | 4.07 | 1.0 | 48.4 |  | 3.90 | 7 | G | OP2 | A:1079 | 4.16 | 1.0 | 39.7 |  | 6.59 | 8 | Water 4 | | A:2240 | 4.38 | 1.0 | 49.8 |  | 2.09 | 9 | Water 5 | | A:2468 | 4.61 | 1.0 | 52.2 | 2.64 [G:OP1]A:1079 // 2.68 [ARG:NH2]E:138 | 2.02 |

  
• • **3.22 ± 0.68: the average coodination distance of the 1^st^ six ligands is K^+^ compatible**  
**6: Number of potential Water contacts**   
**2 : Number of second shell contacts involving water or ions**   
**33%: Percentage of satisfied second shell Water contacts**   
• Contact(s): /O_w_/  
• Contact code: O_w_  
• Contact(s) in the exclusion zone: /O_w_/O_w_/  
→ Missing octahedral coordination for MG ←  
→2Fo-Fc density between 1.0 and 4.0 !   
Go to ToC

### MG:A:1698

|  | Res. type | Atom | Residue | Distance | Occupancy | B-factor | Closest atom(s) to a 1^st^ shell water | EM\_dens. |
| --- | --- | --- | --- | --- | --- | --- | --- | --- |
| **Ion** | | | |  | 1.0 | 69.8 |  | 3.53 |
 1 | Water 1 | | A:1834 | 1.83 | 1.0 | 68.6 | 2.31 [A:OP2]A:937 | 2.10 | 2 | Water 2 | | A:3670 | 1.92 | 1.0 | 69.2 | 2.88 [G:N7]A:939 | 2.56 | 3 | Water 3 | | A:2430 | 1.95 | 1.0 | 62.4 | 2.63 [A:N7]A:937 | 2.31 | 4 | Water 4 | | A:2299 | 2.02 | 1.0 | 61.2 | 2.60 [A:OP2]A:937 | 2.70 | 5 | A | OP2 | A:937 | 2.32 | 1.0 | 67.1 |  | 2.57 | 6 | A | O5' | A:937 | 4.09 | 1.0 | 62.7 |  | 4.37 | 7 | A | N7 | A:938 | 4.18 | 1.0 | 62.0 |  | 4.16 | 8 | G | O6 | A:939 | 4.21 | 1.0 | 64.8 |  | 2.92 | 9 | A | N7 | A:937 | 4.26 | 1.0 | 63.2 |  | 4.09 | 10 | Water 5 | | A:2285 | 4.44 | 1.0 | 60.4 |  | 2.79 | 11 | A | OP1 | A:937 | 4.65 | 1.0 | 67.3 |  | 1.84 | 12 | G | N7 | A:939 | 4.67 | 1.0 | 62.2 |  | 3.94 | 13 | C | O3' | A:936 | 4.69 | 1.0 | 69.2 |  | 1.17 |

  
**1^st^ shell coordination angles and distances (*trans* in "blue"; *cys* in "green"; deviations in "orange"):**   
• 1-2(W/W): *80º*; 1-3(W/W): *159º*; 1-4(W/W): *72º*; 2-3(W/W): *110º*; 2-4(W/W): *81º*; 3-4(W/W): 91º;   
• 1-2(W/W): 2.4Å; 1-3(W/W): *3.7Å*; 1-4(W/W): 2.3Å; 2-3(W/W): 3.2Å; 2-4(W/W): 2.6Å; 3-4(W/W): 2.8Å;   
  
**2.36 ± 0.79: the average coodination distance of the 1^st^ six ligands is Na^+^** compatible  
**8: Number of potential Water contacts**   
**4 : Number of second shell contacts involving water or ions**   
**50%: Percentage of satisfied second shell Water contacts**   
• Contact(s): /O_w_/O_w_/O_w_/  
• Contact code: 3O_w_  
• Contact(s) in the exclusion zone: /O_w_/OP2/  
→ Missing octahedral coordination for MG ←  
→→→ Distorded coord. octahedron (13.2º) ←←←  
• MG hydration shell issue: water too close (< 2.4 Å.)  
→2Fo-Fc density between 1.0 and 4.0 !   
Go to ToC

### MG:A:1699

|  | Res. type | Atom | Residue | Distance | Occupancy | B-factor | Closest atom(s) to a 1^st^ shell water | EM\_dens. |
| --- | --- | --- | --- | --- | --- | --- | --- | --- |
| **Ion** | | | |  | 1.0 | 70.1 |  | 1.93 |
 1 | A | OP2 | A:964 | 1.90 | 1.0 | 61.2 |  | 2.56 | 2 | Water 1 | | A:4318 | 2.03 | 1.0 | 62.1 |  | 1.63 | 3 | U | O4 | A:965 | 2.48 | 1.0 | 63.8 |  | 1.42 | 4 | A | OP1 | A:964 | 3.94 | 1.0 | 61.8 |  | 2.64 | 5 | G | O3' | A:963 | 4.04 | 1.0 | 64.3 |  | 2.87 | 6 | G | O5' | A:963 | 4.12 | 1.0 | 61.7 |  | 2.91 | 7 | G | OP1 | A:963 | 4.28 | 1.0 | 62.8 |  | 1.74 | 8 | A | O5' | A:964 | 4.33 | 1.0 | 56.8 |  | 4.88 | 9 | G | OP2 | A:963 | 4.56 | 1.0 | 61.7 |  | 2.73 |

  
**1^st^ shell coordination angles and distances (*trans* in "blue"; *cys* in "green"; deviations in "orange"):**   
• 1-2(OP2/W): *108º*;   
• 1-2(OP2/W): 3.2Å;   
  
**3.09 ± 0.96: the average coodination distance of the 1^st^ six ligands is K^+^ compatible**  
**2: Number of potential Water contacts**   
**0 : Number of second shell contacts involving water or ions**   
**0%: Percentage of satisfied second shell Water contacts**   
• Contact(s): /O_w_/OP2/  
• Contact code: O_ph_O_w_  
• Contact(s) in the exclusion zone: /O4/  
→ Missing octahedral coordination for MG ←  
→2Fo-Fc density between 1.0 and 4.0 !   
Go to ToC

### MG:A:1700

|  | Res. type | Atom | Residue | Distance | Occupancy | B-factor | Closest atom(s) to a 1^st^ shell water | EM\_dens. |
| --- | --- | --- | --- | --- | --- | --- | --- | --- |
| **Ion** | | | |  | 1.0 | 65.0 |  | 5.10 |
 1 | Water 1 | | A:2153 | 1.82 | 1.0 | 58.4 |  | 2.79 | 2 | Water 2 | | A:2485 | 1.92 | 1.0 | 57.7 | 2.64 [U:OP2]A:965 // 2.79 [G:OP1]A:1198 | 3.04 | 3 | A | OP1 | A:964 | 1.93 | 1.0 | 61.8 |  | 2.64 | 4 | Water 3 | | A:3359 | 1.98 | 1.0 | 62.7 |  | 2.22 | 5 | Water 4 | | A:2926 | 2.06 | 1.0 | 56.0 | 2.73 [U:OP2]A:1199 | 3.80 | 6 | U | OP1 | A:1199 | 2.23 | 1.0 | 56.5 |  | 3.81 | 7 | U | OP2 | A:1199 | 3.69 | 1.0 | 60.0 |  | 3.53 | 8 | G | O3' | A:963 | 3.98 | 1.0 | 64.3 |  | 2.87 | 9 | U | OP2 | A:965 | 4.01 | 1.0 | 61.1 |  | 3.64 | 10 | G | O3' | A:1198 | 4.03 | 1.0 | 57.5 |  | 4.82 | 11 | A | O5' | A:964 | 4.11 | 1.0 | 56.8 |  | 4.88 | 12 | G | OP1 | A:1198 | 4.21 | 1.0 | 58.1 |  | 4.08 | 13 | A | OP2 | A:964 | 4.44 | 1.0 | 61.2 |  | 2.56 | 14 | Water 5 | | A:2228 | 4.50 | 1.0 | 61.1 |  | 2.03 | 15 | U | O5' | A:1199 | 4.65 | 1.0 | 60.8 |  | 3.03 |

  
**1^st^ shell coordination angles and distances (*trans* in "blue"; *cys* in "green"; deviations in "orange"):**   
• 1-2(W/W): 173º; 1-3(W/OP1): 87º; 1-4(W/W): *80º*; 1-5(W/W): 88º; 1-6(W/OP1): *77º*; 2-3(W/OP1): *100º*; 2-4(W/W): *98º*; 2-5(W/W): 85º; 2-6(W/OP1): *106º*; 3-4(OP1/W): 92º; 3-5(OP1/W): 171º; 3-6(OP1/OP1): 85º; 4-5(W/W): 94º; 4-6(W/OP1): *157º*; 5-6(W/OP1): 87º;   
• 1-2(W/W): *3.7Å*; 1-3(W/OP1): 2.6Å; 1-4(W/W): 2.4Å; 1-5(W/W): 2.7Å; 1-6(W/OP1): 2.5Å; 2-3(W/OP1): 2.9Å; 2-4(W/W): 2.9Å; 2-5(W/W): 2.7Å; 2-6(W/OP1): *3.3Å*; 3-4(OP1/W): 2.8Å; 3-5(OP1/W): 4.0Å; 3-6(OP1/OP1): 2.8Å; 4-5(W/W): 3.0Å; 4-6(W/OP1): 4.1Å; 5-6(W/OP1): 2.9Å;   
  
**1.99 ± 0.13: the average coodination distance of the 1^st^ six ligands is Mg^2+^ compatible**  
**8: Number of potential Water contacts**   
**3 : Number of second shell contacts involving water or ions**   
**38%: Percentage of satisfied second shell Water contacts**   
• Contact(s): /O_w_/O_w_/O_w_/OP1/OP1/  
• Contact code: *cis-*2O_ph_3O_w_  
• Contact(s) in the exclusion zone: /O_w_/  
→ Missing octahedral coordination for MG ←  
→ Slightly distorded coord. octahedron (8.0º) ←  
Go to ToC

### MG:A:1701

|  | Res. type | Atom | Residue | Distance | Occupancy | B-factor | Closest atom(s) to a 1^st^ shell water | EM\_dens. |
| --- | --- | --- | --- | --- | --- | --- | --- | --- |
| **Ion** | | | |  | 1.0 | 59.2 |  | 4.56 |
 1 | A | OP1 | A:1197 | 1.92 | 1.0 | 56.4 |  | 4.97 | 2 | C | OP1 | A:1054 | 1.95 | 1.0 | 64.6 |  | 3.29 | 3 | G | OP2 | A:1198 | 2.07 | 1.0 | 57.0 |  | 3.58 | 4 | Water 1 | | A:3950 | 2.26 | 1.0 | 59.0 |  | 3.17 | 5 | Water 2 | | A:3191 | 2.30 | 1.0 | 59.7 | 2.78 [A:O2']A:1196 | 2.39 | 6 | Water 3 | | A:3242 | 2.33 | 1.0 | 56.2 |  | 3.08 | 7 | C | OP2 | A:1054 | 3.59 | 1.0 | 60.1 |  | 4.49 | 8 | G | OP1 | A:1198 | 3.60 | 1.0 | 58.1 |  | 4.08 | 9 | A | O2' | A:1196 | 3.64 | 1.0 | 64.4 |  | 2.77 | 10 | Water 4 | | A:2118 | 3.69 | 1.0 | 60.5 | 1.95 [MG:MG]A:1702 | 3.61 | 11 | A | OP2 | A:1197 | 3.85 | 1.0 | 59.9 |  | 2.85 | 12 | A | O3' | A:1197 | 3.94 | 1.0 | 57.7 |  | 4.17 | 13 | C | O5' | A:1054 | 4.05 | 1.0 | 63.6 |  | 3.34 | 14 | A | O5' | A:1197 | 4.20 | 1.0 | 58.6 |  | 4.28 | 15 | **MG** | MG | A:1702 | 4.24 | 1.0 | 62.3 |  | 5.18 | 16 | A | O3' | A:1196 | 4.31 | 1.0 | 63.9 |  | 2.95 | 17 | G | O3' | A:1053 | 4.35 | 1.0 | 61.6 |  | 4.24 | 18 | G | O5' | A:1198 | 4.48 | 1.0 | 58.6 |  | 3.43 |

  
**1^st^ shell coordination angles and distances (*trans* in "blue"; *cys* in "green"; deviations in "orange"):**   
• 1-2(OP1/OP1): *108º*; 1-3(OP1/OP2): 89º; 1-4(OP1/W): 90º; 2-3(OP1/OP2): *99º*; 2-4(OP1/W): *160º*; 3-4(OP2/W): 92º;   
• 1-2(OP1/OP1): 3.1Å; 1-3(OP1/OP2): 2.8Å; 1-4(OP1/W): 3.0Å; 2-3(OP1/OP2): 3.1Å; 2-4(OP1/W): 4.1Å; 3-4(OP2/W): 3.1Å;   
  
**2.14 ± 0.17: the average coodination distance of the 1^st^ six ligands is Mg^2+^ compatible**  
**6: Number of potential Water contacts**   
**1 : Number of second shell contacts involving water or ions**   
**17%: Percentage of satisfied second shell Water contacts**   
• Contact(s): /O_w_/OP1/OP1/OP2/  
• Contact code: *fac-*3O_ph_O_w_  
• Contact(s) in the exclusion zone: /O_w_/O_w_/  
→ Missing octahedral coordination for MG ←  
→ Slightly distorded coord. octahedron (8.4º) ←  
Go to ToC

### MG:A:1702

|  | Res. type | Atom | Residue | Distance | Occupancy | B-factor | Closest atom(s) to a 1^st^ shell water | EM\_dens. |
| --- | --- | --- | --- | --- | --- | --- | --- | --- |
| **Ion** | | | |  | 1.0 | 62.3 |  | 5.18 |
 1 | Water 1 | | A:3474 | 1.88 | 1.0 | 60.9 |  | 2.62 | 2 | Water 2 | | A:1985 | 1.89 | 1.0 | 60.4 |  | 2.99 | 3 | A | OP2 | A:1197 | 1.91 | 1.0 | 59.9 |  | 2.85 | 4 | C | OP2 | A:1054 | 1.94 | 1.0 | 60.1 |  | 4.49 | 5 | Water 3 | | A:2118 | 1.95 | 1.0 | 60.5 | 2.53 [G:O2']A:1053 // 2.59 [G:OP2]A:1198 | 3.61 | 6 | Water 4 | | A:2460 | 2.44 | 1.0 | 55.5 | 2.64 [G:N7]A:1198 | 4.31 | 7 | G | O3' | A:1053 | 3.45 | 1.0 | 61.6 |  | 4.24 | 8 | A | OP1 | A:1197 | 3.65 | 1.0 | 56.4 |  | 4.97 | 9 | C | OP1 | A:1054 | 3.70 | 1.0 | 64.6 |  | 3.29 | 10 | A | O3' | A:1196 | 3.81 | 1.0 | 63.9 |  | 2.95 | 11 | G | O2' | A:1053 | 3.85 | 1.0 | 59.1 |  | 3.30 | 12 | G | OP2 | A:1198 | 3.85 | 1.0 | 57.0 |  | 3.58 | 13 | Water 5 | | A:4255 | 4.03 | 1.0 | 59.0 |  | 1.68 | 14 | **MG** | MG | A:1701 | 4.24 | 1.0 | 59.2 |  | 4.56 | 15 | G | N7 | A:1198 | 4.33 | 1.0 | 56.8 |  | 4.96 | 16 | A | O5' | A:1197 | 4.36 | 1.0 | 58.6 |  | 4.28 | 17 | C | O5' | A:1054 | 4.37 | 1.0 | 63.6 |  | 3.34 | 18 | Water 6 | | A:3600 | 4.39 | 1.0 | 68.8 |  | 1.40 | 19 | U | O4 | A:1199 | 4.48 | 1.0 | 59.1 |  | 3.68 | 20 | A | O2' | A:1196 | 4.75 | 1.0 | 64.4 |  | 2.77 |

  
**1^st^ shell coordination angles and distances (*trans* in "blue"; *cys* in "green"; deviations in "orange"):**   
• 1-2(W/W): 91º; 1-3(W/OP2): *165º*; 1-4(W/OP2): *96º*; 1-5(W/W): 90º; 2-3(W/OP2): *81º*; 2-4(W/OP2): 90º; 2-5(W/W): *168º*; 3-4(OP2/OP2): *97º*; 3-5(OP2/W): *96º*; 4-5(OP2/W): *102º*;   
• 1-2(W/W): 2.7Å; 1-3(W/OP2): *3.8Å*; 1-4(W/OP2): 2.8Å; 1-5(W/W): 2.7Å; 2-3(W/OP2): 2.5Å; 2-4(W/OP2): 2.7Å; 2-5(W/W): 3.8Å; 3-4(OP2/OP2): 2.9Å; 3-5(OP2/W): 2.9Å; 4-5(OP2/W): 3.0Å;   
  
**2.00 ± 0.20: the average coodination distance of the 1^st^ six ligands is Mg^2+^ compatible**  
**8: Number of potential Water contacts**   
**3 : Number of second shell contacts involving water or ions**   
**38%: Percentage of satisfied second shell Water contacts**   
• Contact(s): /O_w_/OP2/OP2/  
• Contact code: *cis-*2O_ph_O_w_  
• Contact(s) in the exclusion zone: /O_w_/O_w_/O_w_/  
→ Missing octahedral coordination for MG ←  
→ Slightly distorded coord. octahedron (6.8º) ←  
Go to ToC

### MG:A:1703

|  | Res. type | Atom | Residue | Distance | Occupancy | B-factor | Closest atom(s) to a 1^st^ shell water | EM\_dens. |
| --- | --- | --- | --- | --- | --- | --- | --- | --- |
| **Ion** | | | |  | 1.0 | 70.2 |  | 3.34 |
 1 | Water 1 | | A:4612 | 1.79 | 1.0 | 58.6 |  | 2.69 | 2 | C | OP1 | A:972 | 1.80 | 1.0 | 61.3 |  | 3.73 | 3 | Water 2 | | A:2014 | 2.01 | 1.0 | 60.8 | 2.48 [C:OP2]A:972 | 2.36 | 4 | Water 3 | | J:209 | 2.10 | 1.0 | 59.0 | 2.82 [ASP:OD1]J:60 | 2.39 | 5 | Water 4 | | A:2967 | 2.14 | 1.0 | 63.7 |  | 2.31 | 6 | Water 5 | | A:3233 | 2.37 | 1.0 | 60.8 | 2.79 [G:O2']A:971 // 2.89 [C:OP1]A:1366 | 3.24 | 7 | C | OP2 | A:972 | 3.12 | 1.0 | 58.6 |  | 3.32 | 8 | C | O5' | A:972 | 3.87 | 1.0 | 58.3 |  | 3.34 | 9 | G | O3' | A:971 | 4.05 | 1.0 | 64.0 |  | 3.53 | 10 | Water 6 | | I:206 | 4.08 | 1.0 | 64.8 |  | 1.64 | 11 | Water 7 | | A:1854 | 4.19 | 1.0 | 63.2 |  | 1.99 | 12 | G | O2' | A:971 | 4.34 | 1.0 | 57.8 |  | 4.86 | 13 | ASP | OD1 | J:60 | 4.37 | 1.0 | 64.5 |  | 2.44 | 14 | Water 8 | | A:2741 | 4.37 | 1.0 | 62.0 |  | 2.37 | 15 | Water 9 | | J:208 | 4.51 | 1.0 | 62.8 | 2.88 [ASP:OD1]J:60 | 3.00 | 16 | Water 10 | | J:203 | 4.62 | 1.0 | 63.2 |  | 1.57 | 17 | C | OP1 | A:1366 | 4.77 | 1.0 | 63.4 |  | 2.99 |

  
**1^st^ shell coordination angles and distances (*trans* in "blue"; *cys* in "green"; deviations in "orange"):**   
• 1-2(W/OP1): *163º*; 1-3(W/W): 89º; 1-4(W/W): *79º*; 1-5(W/W): *78º*; 2-3(OP1/W): *105º*; 2-4(OP1/W): *110º*; 2-5(OP1/W): 87º; 3-4(W/W): 88º; 3-5(W/W): *167º*; 4-5(W/W): 86º;   
• 1-2(W/OP1): *3.6Å*; 1-3(W/W): 2.7Å; 1-4(W/W): 2.5Å; 1-5(W/W): 2.5Å; 2-3(OP1/W): 3.0Å; 2-4(OP1/W): *3.2Å*; 2-5(OP1/W): 2.7Å; 3-4(W/W): 2.9Å; 3-5(W/W): 4.1Å; 4-5(W/W): 2.9Å;   
  
**2.04 ± 0.20: the average coodination distance of the 1^st^ six ligands is Mg^2+^ compatible**  
**10: Number of potential Water contacts**   
**4 : Number of second shell contacts involving water or ions**   
**40%: Percentage of satisfied second shell Water contacts**   
• Contact(s): /O_w_/O_w_/O_w_/  
• Contact code: 3O_w_  
• Contact(s) in the exclusion zone: /O_w_/O_w_/OP1/OP2/  
→ Consider K in place of MG (1 valid contact(s) for K vs. 0 for MG ←)  
→ Missing octahedral coordination for MG ←  
→ Slightly distorded coord. octahedron (9.7º) ←→2Fo-Fc density between 1.0 and 4.0 !   
Go to ToC

### MG:A:1704

|  | Res. type | Atom | Residue | Distance | Occupancy | B-factor | Closest atom(s) to a 1^st^ shell water | EM\_dens. |
| --- | --- | --- | --- | --- | --- | --- | --- | --- |
| **Ion** | | | |  | 1.0 | 73.6 |  | 2.33 |
 1 | Water 1 | | A:1806 | 1.88 | 1.0 | 73.8 |  | 1.78 | 2 | C | OP1 | A:980 | 2.02 | 1.0 | 72.5 |  | 3.33 | 3 | C | O3' | A:979 | 3.84 | 1.0 | 75.2 |  | 3.23 | 4 | A | OP2 | A:1219 | 4.08 | 1.0 | 90.9 |  | 1.14 | 5 | C | O5' | A:980 | 4.19 | 1.0 | 75.3 |  | 3.03 | 6 | Water 2 | | A:3305 | 4.48 | 1.0 | 78.0 |  | 1.68 | 7 | A | N7 | A:1219 | 4.49 | 1.0 | 88.8 |  | 2.35 | 8 | C | OP2 | A:980 | 4.53 | 1.0 | 76.7 |  | 2.22 | 9 | G | O6 | A:1220 | 4.68 | 1.0 | 84.4 |  | 1.84 | 10 | G | N7 | A:1220 | 4.68 | 1.0 | 82.6 |  | 2.66 |

  
**1^st^ shell coordination angles and distances (*trans* in "blue"; *cys* in "green"; deviations in "orange"):**   
• 1-2(W/OP1): *70º*;   
• 1-2(W/OP1): 2.2Å;   
  
**3.41 ± 1.05: the average coodination distance of the 1^st^ six ligands is K^+^ compatible**  
**4: Number of potential Water contacts**   
**0 : Number of second shell contacts involving water or ions**   
**0%: Percentage of satisfied second shell Water contacts**   
• Contact(s): /OP1/  
• Contact code: O_ph_  
• Contact(s) in the exclusion zone: /O_w_/  
→ Missing octahedral coordination for MG ←  
→2Fo-Fc density between 1.0 and 4.0 !   
Go to ToC

### MG:A:1705

|  | Res. type | Atom | Residue | Distance | Occupancy | B-factor | Closest atom(s) to a 1^st^ shell water | EM\_dens. |
| --- | --- | --- | --- | --- | --- | --- | --- | --- |
| **Ion** | | | |  | 1.0 | 72.4 |  | 1.82 |
 1 | Water 1 | | A:4644 | 1.96 | 1.0 | 67.0 |  | 1.47 | 2 | Water 2 | | A:4039 | 2.46 | 1.0 | 61.9 |  | 1.51 | 3 | Water 3 | | A:3595 | 2.66 | 1.0 | 65.2 | 2.86 [G:OP2]A:1190 | 1.44 | 4 | G | N7 | A:1190 | 3.39 | 1.0 | 60.1 |  | 2.93 | 5 | G | OP1 | A:1061 | 4.23 | 1.0 | 55.9 |  | 4.03 | 6 | G | OP2 | A:1190 | 4.39 | 1.0 | 61.5 |  | 3.12 | 7 | U | OP2 | A:1189 | 4.44 | 1.0 | 61.0 |  | 3.77 | 8 | Water 4 | | A:3677 | 4.49 | 1.0 | 66.7 | 2.88 [U:OP2]A:1189 | 1.51 | 9 | U | OP1 | A:1189 | 4.69 | 1.0 | 59.6 |  | 3.64 |

  
• • **3.18 ± 0.90: the average coodination distance of the 1^st^ six ligands is K^+^ compatible**  
**6: Number of potential Water contacts**   
**1 : Number of second shell contacts involving water or ions**   
**17%: Percentage of satisfied second shell Water contacts**   
• Contact(s): /O_w_/  
• Contact code: O_w_  
• Contact(s) in the exclusion zone: /N7/O_w_/O_w_/  
→ Missing octahedral coordination for MG ←  
→2Fo-Fc density between 1.0 and 4.0 !   
Go to ToC

### MG:A:1706

|  | Res. type | Atom | Residue | Distance | Occupancy | B-factor | Closest atom(s) to a 1^st^ shell water | EM\_dens. |
| --- | --- | --- | --- | --- | --- | --- | --- | --- |
| **Ion** | | | |  | 1.0 | 61.1 |  | 5.17 |
 1 | Water 1 | | A:1836 | 1.83 | 1.0 | 57.2 |  | 2.71 | 2 | G | OP1 | A:1068 | 1.84 | 1.0 | 53.6 |  | 5.15 | 3 | Water 2 | | A:2448 | 1.85 | 1.0 | 55.6 |  | 3.90 | 4 | Water 3 | | A:3493 | 1.89 | 1.0 | 52.0 |  | 3.18 | 5 | G | OP1 | A:1094 | 1.98 | 1.0 | 57.2 |  | 3.04 | 6 | Water 4 | | A:3772 | 2.03 | 1.0 | 57.7 | 2.90 [C:O4']A:1388 | 2.50 | 7 | G | OP2 | A:1068 | 3.61 | 1.0 | 54.5 |  | 4.67 | 8 | G | O2' | A:1387 | 3.79 | 1.0 | 56.0 |  | 3.36 | 9 | A | O3' | A:1067 | 3.86 | 1.0 | 56.8 |  | 3.18 | 10 | Water 5 | | A:3556 | 4.16 | 1.0 | 58.9 |  | 2.11 | 11 | G | O5' | A:1094 | 4.18 | 1.0 | 55.1 |  | 3.31 | 12 | G | OP2 | A:1094 | 4.20 | 1.0 | 60.6 |  | 2.66 | 13 | G | O5' | A:1068 | 4.30 | 1.0 | 52.4 |  | 3.88 | 14 | Water 6 | | A:4191 | 4.37 | 1.0 | 60.0 |  | 1.84 | 15 | A | O3' | A:1093 | 4.38 | 1.0 | 61.6 |  | 3.50 | 16 | C | O4' | A:1388 | 4.44 | 1.0 | 54.6 |  | 3.73 | 17 | Water 7 | | A:4444 | 4.47 | 1.0 | 57.3 |  | 3.40 |

  
**1^st^ shell coordination angles and distances (*trans* in "blue"; *cys* in "green"; deviations in "orange"):**   
• 1-2(W/OP1): 174º; 1-3(W/W): *83º*; 1-4(W/W): *80º*; 1-5(W/OP1): *75º*; 1-6(W/W): *82º*; 2-3(OP1/W): 91º; 2-4(OP1/W): *106º*; 2-5(OP1/OP1): *105º*; 2-6(OP1/W): *98º*; 3-4(W/W): *161º*; 3-5(W/OP1): 88º; 3-6(W/W): *82º*; 4-5(W/OP1): 94º; 4-6(W/W): 88º; 5-6(OP1/W): *156º*;   
• 1-2(W/OP1): *3.7Å*; 1-3(W/W): 2.4Å; 1-4(W/W): 2.4Å; 1-5(W/OP1): 2.3Å; 1-6(W/W): 2.5Å; 2-3(OP1/W): 2.6Å; 2-4(OP1/W): 3.0Å; 2-5(OP1/OP1): 3.0Å; 2-6(OP1/W): 2.9Å; 3-4(W/W): *3.7Å*; 3-5(W/OP1): 2.7Å; 3-6(W/W): 2.5Å; 4-5(W/OP1): 2.8Å; 4-6(W/W): 2.7Å; 5-6(OP1/W): 3.9Å;   
  
**1.90 ± 0.07: the average coodination distance of the 1^st^ six ligands is Mg^2+^ compatible**  
**8: Number of potential Water contacts**   
**1 : Number of second shell contacts involving water or ions**   
**12%: Percentage of satisfied second shell Water contacts**   
• Contact(s): /O_w_/OP1/  
• Contact code: O_ph_O_w_  
• Contact(s) in the exclusion zone: /O_w_/O_w_/O_w_/OP1/  
→ Missing octahedral coordination for MG ←  
→ Slightly distorded coord. octahedron (9.7º) ←  
Go to ToC

### MG:A:1707

|  | Res. type | Atom | Residue | Distance | Occupancy | B-factor | Closest atom(s) to a 1^st^ shell water | EM\_dens. |
| --- | --- | --- | --- | --- | --- | --- | --- | --- |
| **Ion** | | | |  | 1.0 | 61.2 |  | 3.22 |
 1 | U | OP2 | A:1083 | 1.87 | 1.0 | 44.4 |  | 6.39 | 2 | Water 1 | | A:2799 | 1.90 | 1.0 | 55.3 | 2.95 [G:N7]A:1084 | 2.68 | 3 | Water 2 | | A:2629 | 1.95 | 1.0 | 52.5 | 2.96 [G:O6]A:1084 | 2.49 | 4 | Water 3 | | A:3733 | 2.45 | 1.0 | 55.9 | 2.89 [U:OP1]A:1083 | 2.26 | 5 | Water 4 | | A:4299 | 2.45 | 1.0 | 55.5 |  | 2.35 | 6 | U | OP1 | A:1083 | 3.65 | 1.0 | 50.2 |  | 4.16 | 7 | Water 5 | | A:4051 | 3.82 | 1.0 | 56.3 |  | 1.66 | 8 | Water 6 | | A:3622 | 3.86 | 1.0 | 54.5 |  | 1.52 | 9 | U | O5' | A:1083 | 4.08 | 1.0 | 46.5 |  | 6.36 | 10 | A | O3' | A:1082 | 4.23 | 1.0 | 49.3 |  | 4.89 | 11 | U | O2 | A:1085 | 4.35 | 1.0 | 54.5 |  | 4.78 | 12 | A | OP1 | A:1082 | 4.36 | 1.0 | 47.9 |  | 4.61 | 13 | Water 7 | | A:4083 | 4.38 | 1.0 | 59.2 |  | 1.64 | 14 | Water 8 | | A:1843 | 4.44 | 1.0 | 56.0 |  | 1.56 | 15 | Water 9 | | A:4141 | 4.45 | 1.0 | 55.8 |  | 3.69 | 16 | G | N7 | A:1084 | 4.51 | 1.0 | 48.7 |  | 5.28 | 17 | A | O5' | A:1082 | 4.57 | 1.0 | 43.9 |  | 5.70 | 18 | G | O6 | A:1084 | 4.61 | 1.0 | 50.0 |  | 3.98 |

  
**1^st^ shell coordination angles and distances (*trans* in "blue"; *cys* in "green"; deviations in "orange"):**   
• 1-2(OP2/W): 92º; 1-3(OP2/W): 89º; 2-3(W/W): 85º;   
• 1-2(OP2/W): 2.7Å; 1-3(OP2/W): 2.7Å; 2-3(W/W): 2.6Å;   
  
**2.38 ± 0.62: the average coodination distance of the 1^st^ six ligands is Na^+^** compatible  
**8: Number of potential Water contacts**   
**3 : Number of second shell contacts involving water or ions**   
**38%: Percentage of satisfied second shell Water contacts**   
• Contact(s): /O_w_/  
• Contact code: O_w_  
• Contact(s) in the exclusion zone: /O_w_/O_w_/O_w_/OP2/  
→ Missing octahedral coordination for MG ←  
→2Fo-Fc density between 1.0 and 4.0 !   
Go to ToC

### MG:A:1708

|  | Res. type | Atom | Residue | Distance | Occupancy | B-factor | Closest atom(s) to a 1^st^ shell water | EM\_dens. |
| --- | --- | --- | --- | --- | --- | --- | --- | --- |
| **Ion** | | | |  | 1.0 | 69.6 |  | 3.16 |
 1 | Water 1 | | A:2508 | 1.87 | 1.0 | 63.2 | 2.64 [G:O6]A:1099 | 1.97 | 2 | U | OP2 | A:1085 | 1.88 | 1.0 | 59.8 |  | 2.22 | 3 | Water 2 | | A:2551 | 1.96 | 1.0 | 58.0 | 2.65 [C:O2]A:1100 | 3.30 | 4 | Water 3 | | A:3009 | 1.96 | 1.0 | 62.5 | 2.75 [G:O6]A:1099 | 2.30 | 5 | U | O4 | A:1086 | 2.07 | 1.0 | 61.8 |  | 2.46 | 6 | G | O6 | A:1099 | 2.44 | 1.0 | 64.0 |  | 4.19 | 7 | U | OP1 | A:1085 | 3.59 | 1.0 | 57.1 |  | 2.64 | 8 | G | O3' | A:1084 | 3.88 | 1.0 | 51.5 |  | 3.10 | 9 | Water 4 | | A:2234 | 3.91 | 1.0 | 60.5 |  | 2.48 | 10 | C | O2 | A:1100 | 4.22 | 1.0 | 63.0 |  | 2.80 | 11 | U | O5' | A:1085 | 4.31 | 1.0 | 55.4 |  | 3.82 | 12 | Water 5 | | A:2136 | 4.36 | 1.0 | 56.5 | 2.98 [G:O3']A:1084 | 2.59 | 13 | G | O6 | A:1087 | 4.60 | 1.0 | 62.4 |  | 3.66 | 14 | Water 6 | | A:2351 | 4.80 | 1.0 | 63.8 | 2.98 [G:O6]A:1099 | 1.62 |

  
**1^st^ shell coordination angles and distances (*trans* in "blue"; *cys* in "green"; deviations in "orange"):**   
• 1-2(W/OP2): *99º*; 1-3(W/W): *97º*; 1-4(W/W): *73º*; 1-5(W/O4): *153º*; 2-3(OP2/W): *105º*; 2-4(OP2/W): *101º*; 2-5(OP2/O4): *102º*; 3-4(W/W): *154º*; 3-5(W/O4): 94º; 4-5(W/O4): 86º;   
• 1-2(W/OP2): 2.8Å; 1-3(W/W): 2.9Å; 1-4(W/W): 2.3Å; 1-5(W/O4): 3.8Å; 2-3(OP2/W): 3.0Å; 2-4(OP2/W): 3.0Å; 2-5(OP2/O4): 3.1Å; 3-4(W/W): 3.8Å; 3-5(W/O4): 2.9Å; 4-5(W/O4): 2.8Å;   
  
**2.03 ± 0.19: the average coodination distance of the 1^st^ six ligands is Mg^2+^ compatible**  
**6: Number of potential Water contacts**   
**3 : Number of second shell contacts involving water or ions**   
**50%: Percentage of satisfied second shell Water contacts**   
• Contact(s): /O4/O_w_/O_w_/  
• Contact code: O_b_2O_w_  
• Contact(s) in the exclusion zone: /O6/O_w_/OP2/  
→ Missing octahedral coordination for MG ←  
→→→ Distorded coord. octahedron (13.2º) ←←←  
→2Fo-Fc density between 1.0 and 4.0 !   
Go to ToC

### MG:A:1709

|  | Res. type | Atom | Residue | Distance | Occupancy | B-factor | Closest atom(s) to a 1^st^ shell water | EM\_dens. |
| --- | --- | --- | --- | --- | --- | --- | --- | --- |
| **Ion** | | | |  | 1.0 | 73.9 |  | 1.80 |
 1 | Water 1 | | A:4537 | 2.12 | 1.0 | 67.3 |  | 1.76 | 2 | G | N7 | A:1088 | 2.75 | 1.0 | 65.1 |  | 3.44 | 3 | Water 2 | | A:4415 | 3.46 | 1.0 | 67.6 |  | 1.65 | 4 | G | O6 | A:1088 | 3.54 | 1.0 | 66.7 |  | 2.98 | 5 | G | N7 | A:1087 | 3.83 | 1.0 | 61.1 |  | 3.94 | 6 | G | N7 | A:1089 | 4.07 | 1.0 | 65.1 |  | 3.06 | 7 | G | OP2 | A:1087 | 4.25 | 1.0 | 66.1 |  | 2.29 | 8 | G | OP2 | A:1088 | 4.42 | 1.0 | 68.0 |  | 2.04 | 9 | G | O6 | A:1089 | 4.50 | 1.0 | 66.7 |  | 3.12 | 10 | G | O6 | A:1087 | 4.69 | 1.0 | 62.4 |  | 3.66 | 11 | G | O5' | A:1087 | 4.77 | 1.0 | 63.1 |  | 3.42 |

  
• • **3.30 ± 0.66: the average coodination distance of the 1^st^ six ligands is K^+^ compatible**  
**4: Number of potential Water contacts**   
**0 : Number of second shell contacts involving water or ions**   
**0%: Percentage of satisfied second shell Water contacts**   
• Contact(s): /O_w_/  
• Contact code: O_w_  
• Contact(s) in the exclusion zone: /N7/  
→ Consider K in place of MG (1 valid contact(s) for K vs. 0 for MG ←)  
→ Missing octahedral coordination for MG ←  
→2Fo-Fc density between 1.0 and 4.0 !   
Go to ToC

### MG:A:1710

|  | Res. type | Atom | Residue | Distance | Occupancy | B-factor | Closest atom(s) to a 1^st^ shell water | EM\_dens. |
| --- | --- | --- | --- | --- | --- | --- | --- | --- |
| **Ion** | | | |  | 1.0 | 58.4 |  | 4.54 |
 1 | U | OP2 | A:1095 | 1.87 | 1.0 | 60.0 |  | 3.66 | 2 | Water 1 | | A:2305 | 1.88 | 1.0 | 57.8 |  | 2.52 | 3 | G | O6 | A:1108 | 2.12 | 1.0 | 53.3 |  | 5.14 | 4 | Water 2 | | A:4140 | 2.29 | 1.0 | 61.0 |  | 2.26 | 5 | Water 3 | | A:2614 | 2.33 | 1.0 | 56.7 | 2.67 [G:O2']A:1094 | 2.91 | 6 | G | N1 | A:1108 | 3.42 | 1.0 | 47.7 |  | 6.00 | 7 | G | O3' | A:1094 | 4.00 | 1.0 | 60.0 |  | 2.35 | 8 | U | O5' | A:1095 | 4.11 | 1.0 | 56.8 |  | 4.20 | 9 | G | O2' | A:1094 | 4.22 | 1.0 | 53.7 |  | 4.21 | 10 | C | OP2 | A:1096 | 4.23 | 1.0 | 67.5 |  | 1.84 | 11 | U | OP1 | A:1095 | 4.31 | 1.0 | 60.8 |  | 3.29 | 12 | C | N3 | A:1109 | 4.62 | 1.0 | 53.5 |  | 4.89 | 13 | Water 4 | | A:2758 | 4.79 | 1.0 | 59.0 |  | 1.72 |

  
**1^st^ shell coordination angles and distances (*trans* in "blue"; *cys* in "green"; deviations in "orange"):**   
• 1-2(OP2/W): 175º; 1-3(OP2/O6): *97º*; 1-4(OP2/W): 92º; 2-3(W/O6): *81º*; 2-4(W/W): 92º; 3-4(O6/W): 92º;   
• 1-2(OP2/W): *3.7Å*; 1-3(OP2/O6): 3.0Å; 1-4(OP2/W): 3.0Å; 2-3(W/O6): 2.6Å; 2-4(W/W): 3.0Å; 3-4(O6/W): 3.2Å;   
  
**2.32 ± 0.52: the average coodination distance of the 1^st^ six ligands is Na^+^** compatible  
**6: Number of potential Water contacts**   
**1 : Number of second shell contacts involving water or ions**   
**17%: Percentage of satisfied second shell Water contacts**   
• Contact(s): /O6/O_w_/  
• Contact code: O_b_O_w_  
• Contact(s) in the exclusion zone: /O_w_/O_w_/OP2/  
→ Missing octahedral coordination for MG ←  
Ooctahedral coordination (4.4º)   
Go to ToC

### MG:A:1711

|  | Res. type | Atom | Residue | Distance | Occupancy | B-factor | Closest atom(s) to a 1^st^ shell water | EM\_dens. |
| --- | --- | --- | --- | --- | --- | --- | --- | --- |
| **Ion** | | | |  | 1.0 | 75.3 |  | 1.05 |
 1 | G | N7 | A:1133 | 4.46 | 1.0 | 133.4 |  | 1.24 | 2 | G | OP2 | A:1133 | 4.71 | 1.0 | 133.9 |  | 0.93 |

  
• • **4.59 ± 0.12: the average coodination distance of the 1^st^ six ligands could correspond to an hexahydrated Mg^2+^.**  
**0: Number of potential Water contacts**   
• Contact code: → No contacts ←   
→ Firts MG coordination shell missing ←  
→2Fo-Fc density between 1.0 and 4.0 !   
Go to ToC

### MG:A:1712

|  | Res. type | Atom | Residue | Distance | Occupancy | B-factor | Closest atom(s) to a 1^st^ shell water | EM\_dens. |
| --- | --- | --- | --- | --- | --- | --- | --- | --- |
| **Ion** | | | |  | 1.0 | 80.3 |  | 1.74 |
 1 | Water 1 | | A:2480 | 2.02 | 1.0 | 90.8 | 2.64 [G:O6]A:1177 | 1.49 | 2 | A | N7 | A:1176 | 4.04 | 1.0 | 96.0 |  | 1.88 | 3 | G | O6 | A:1177 | 4.13 | 1.0 | 93.7 |  | 1.85 | 4 | G | N7 | A:1175 | 4.18 | 1.0 | 96.2 |  | 2.29 | 5 | G | OP2 | A:1175 | 4.31 | 1.0 | 98.6 |  | 1.12 | 6 | A | OP2 | A:1176 | 4.34 | 1.0 | 95.4 |  | 1.45 | 7 | G | O2' | A:1181 | 4.58 | 1.0 | 86.7 |  | 1.66 |

  
• • **3.84 ± 0.82: the average coodination distance of the 1^st^ six ligands could correspond to an hexahydrated Mg^2+^.**  
**2: Number of potential Water contacts**   
**1 : Number of second shell contacts involving water or ions**   
**50%: Percentage of satisfied second shell Water contacts**   
• Contact(s): /O_w_/  
• Contact code: O_w_  
→ Missing octahedral coordination for MG ←  
→ MG ion B\_factor > 80 ←→2Fo-Fc density between 1.0 and 4.0 !   
Go to ToC

### MG:A:1713

|  | Res. type | Atom | Residue | Distance | Occupancy | B-factor | Closest atom(s) to a 1^st^ shell water | EM\_dens. |
| --- | --- | --- | --- | --- | --- | --- | --- | --- |
| **Ion** | | | |  | 1.0 | 74.9 |  | 2.51 |
 1 | Water 1 | | A:3470 | 1.96 | 1.0 | 62.3 |  | 1.81 | 2 | Water 2 | | A:4178 | 1.99 | 1.0 | 64.4 |  | 2.11 | 3 | Water 3 | | A:3194 | 2.01 | 1.0 | 66.4 | 2.78 [U:OP2]A:1345 | 2.07 | 4 | C | OP1 | A:934 | 2.08 | 1.0 | 65.6 |  | 2.48 | 5 | Water 4 | | A:4462 | 2.23 | 1.0 | 64.9 |  | 1.72 | 6 | Water 5 | | A:4176 | 3.89 | 1.0 | 62.8 |  | 2.41 | 7 | C | O5' | A:934 | 4.10 | 1.0 | 63.2 |  | 3.60 | 8 | U | OP2 | A:1345 | 4.23 | 1.0 | 68.9 |  | 1.85 | 9 | U | OP1 | A:1345 | 4.27 | 1.0 | 61.9 |  | 3.64 | 10 | C | OP2 | A:934 | 4.37 | 1.0 | 63.7 |  | 3.56 | 11 | G | O3' | A:933 | 4.47 | 1.0 | 68.9 |  | 1.71 | 12 | Water 6 | | A:4596 | 4.59 | 1.0 | 65.4 |  | 1.52 | 13 | U | O2' | A:1345 | 4.71 | 1.0 | 65.4 |  | 2.43 |

  
**1^st^ shell coordination angles and distances (*trans* in "blue"; *cys* in "green"; deviations in "orange"):**   
• 1-2(W/W): *99º*; 1-3(W/W): 176º; 1-4(W/OP1): 89º; 1-5(W/W): *84º*; 2-3(W/W): 85º; 2-4(W/OP1): *96º*; 2-5(W/W): *169º*; 3-4(W/OP1): 90º; 3-5(W/W): 92º; 4-5(OP1/W): 95º;   
• 1-2(W/W): 3.0Å; 1-3(W/W): 4.0Å; 1-4(W/OP1): 2.8Å; 1-5(W/W): 2.8Å; 2-3(W/W): 2.7Å; 2-4(W/OP1): 3.0Å; 2-5(W/W): *4.2Å*; 3-4(W/OP1): 2.9Å; 3-5(W/W): 3.0Å; 4-5(OP1/W): 3.2Å;   
  
**2.36 ± 0.69: the average coodination distance of the 1^st^ six ligands is Na^+^** compatible  
**10: Number of potential Water contacts**   
**1 : Number of second shell contacts involving water or ions**   
**10%: Percentage of satisfied second shell Water contacts**   
• Contact(s): /O_w_/O_w_/O_w_/O_w_/OP1/  
• Contact code: O_ph_4O_w_  
→ Missing octahedral coordination for MG ←  
Ooctahedral coordination (4.8º) →2Fo-Fc density between 1.0 and 4.0 !   
Go to ToC

### MG:A:1714

|  | Res. type | Atom | Residue | Distance | Occupancy | B-factor | Closest atom(s) to a 1^st^ shell water | EM\_dens. |
| --- | --- | --- | --- | --- | --- | --- | --- | --- |
| **Ion** | | | |  | 1.0 | 65.8 |  | 3.62 |
 1 | Water 1 | | A:4466 | 2.37 | 1.0 | 61.0 |  | 2.21 | 2 | Water 2 | | A:3572 | 2.82 | 1.0 | 63.0 | 2.86 [5MC:OP2]A:967 | 1.52 | 3 | Water 3 | | A:2690 | 2.85 | 1.0 | 55.0 |  | 3.04 | 4 | Water 4 | | A:2129 | 2.89 | 1.0 | 61.4 | 2.53 [A:N3]A:968 | 1.87 | 5 | A | O2' | A:968 | 3.09 | 1.0 | 56.2 |  | 2.82 | 6 | Water 5 | | A:2016 | 3.20 | 1.0 | 59.5 | 2.48 [A:OP1]A:969 | 1.73 | 7 | A | O3' | A:968 | 3.30 | 1.0 | 61.0 |  | 3.04 | 8 | A | OP1 | A:969 | 3.81 | 1.0 | 58.9 |  | 3.98 | 9 | Water 6 | | A:2367 | 3.87 | 1.0 | 60.1 |  | 1.96 | 10 | A | O2' | A:1197 | 4.32 | 1.0 | 57.7 |  | 3.86 | 11 | 5MC | OP2 | A:967 | 4.60 | 1.0 | 60.2 |  | 3.05 | 12 | A | N3 | A:968 | 4.70 | 1.0 | 57.5 |  | 3.49 |

  
• • **2.87 ± 0.26: the average coodination distance of the 1^st^ six ligands is K^+^ compatible**  
**10: Number of potential Water contacts**   
**3 : Number of second shell contacts involving water or ions**   
**30%: Percentage of satisfied second shell Water contacts**   
• Contact code: → No contacts ←   
• Contact(s) in the exclusion zone: /O2'/O3'/O_w_/O_w_/O_w_/O_w_/O_w_/  
→ Consider K in place of MG (1 valid contact(s) for K vs. 0 for MG ←)  
→2Fo-Fc density between 1.0 and 4.0 !   
Go to ToC

### MG:A:1715

|  | Res. type | Atom | Residue | Distance | Occupancy | B-factor | Closest atom(s) to a 1^st^ shell water | EM\_dens. |
| --- | --- | --- | --- | --- | --- | --- | --- | --- |
| **Ion** | | | |  | 1.0 | 82.4 |  | 1.31 |
 1 | U | O4 | A:989 | 3.77 | 1.0 | 99.0 |  | 1.19 | 2 | G | O6 | A:988 | 3.85 | 1.0 | 96.8 |  | 1.41 | 3 | G | O6 | A:987 | 4.12 | 1.0 | 91.9 |  | 1.56 | 4 | G | N7 | A:987 | 4.40 | 1.0 | 92.1 |  | 2.15 | 5 | G | N7 | A:988 | 4.64 | 1.0 | 95.3 |  | 2.10 |

  
• • **4.16 ± 0.33: the average coodination distance of the 1^st^ six ligands could correspond to an hexahydrated Mg^2+^.**  
**0: Number of potential Water contacts**   
• Contact code: → No contacts ←   
→ Firts MG coordination shell missing ←  
→ MG ion B\_factor > 80 ←→2Fo-Fc density between 1.0 and 4.0 !   
Go to ToC

### MG:A:1716

|  | Res. type | Atom | Residue | Distance | Occupancy | B-factor | Closest atom(s) to a 1^st^ shell water | EM\_dens. |
| --- | --- | --- | --- | --- | --- | --- | --- | --- |
| **Ion** | | | |  | 1.0 | 72.1 |  | 3.97 |
 1 | Water 1 | | A:2102 | 1.81 | 1.0 | 68.5 |  | 2.90 | 2 | Water 2 | | A:2099 | 1.86 | 1.0 | 69.1 | 2.52 [G:OP1]A:951 | 2.80 | 3 | U | OP1 | A:1224 | 1.95 | 1.0 | 71.7 |  | 3.22 | 4 | Water 3 | | A:2954 | 2.00 | 1.0 | 68.7 | 2.89 [A:N7]A:977 | 2.36 | 5 | Water 4 | | A:4646 | 2.01 | 1.0 | 67.9 |  | 2.73 | 6 | Water 5 | | A:2950 | 2.29 | 1.0 | 67.2 | 2.74 [U:OP2]A:952 // 2.92 [U:OP2]A:1224 | 2.76 | 7 | U | OP2 | A:1224 | 3.89 | 1.0 | 70.7 |  | 2.67 | 8 | Water 6 | | A:3315 | 3.95 | 1.0 | 65.5 | 2.80 [G:OP1]A:951 | 1.90 | 9 | G | OP1 | A:951 | 4.09 | 1.0 | 63.4 |  | 3.27 | 10 | C | O3' | A:1223 | 4.18 | 1.0 | 70.4 |  | 3.98 | 11 | Water 7 | | A:3213 | 4.35 | 1.0 | 69.2 |  | 1.64 | 12 | U | O5' | A:1224 | 4.36 | 1.0 | 74.8 |  | 2.41 | 13 | A | N7 | A:977 | 4.38 | 1.0 | 71.0 |  | 3.33 | 14 | C | O2' | A:1223 | 4.45 | 1.0 | 72.3 |  | 2.41 | 15 | U | OP2 | A:952 | 4.65 | 1.0 | 62.7 |  | 2.55 | 16 | Water 8 | | A:2053 | 4.72 | 1.0 | 71.8 | 2.50 [U:OP2]A:1224 | 1.59 |

  
**1^st^ shell coordination angles and distances (*trans* in "blue"; *cys* in "green"; deviations in "orange"):**   
• 1-2(W/W): 171º; 1-3(W/OP1): *84º*; 1-4(W/W): 92º; 1-5(W/W): 91º; 1-6(W/W): *75º*; 2-3(W/OP1): *97º*; 2-4(W/W): *98º*; 2-5(W/W): 87º; 2-6(W/W): *96º*; 3-4(OP1/W): 88º; 3-5(OP1/W): 174º; 3-6(OP1/W): *83º*; 4-5(W/W): *95º*; 4-6(W/W): *165º*; 5-6(W/W): 93º;   
• 1-2(W/W): *3.7Å*; 1-3(W/OP1): 2.5Å; 1-4(W/W): 2.7Å; 1-5(W/W): 2.7Å; 1-6(W/W): 2.5Å; 2-3(W/OP1): 2.8Å; 2-4(W/W): 2.9Å; 2-5(W/W): 2.7Å; 2-6(W/W): 3.1Å; 3-4(OP1/W): 2.7Å; 3-5(OP1/W): 4.0Å; 3-6(OP1/W): 2.8Å; 4-5(W/W): 3.0Å; 4-6(W/W): *4.3Å*; 5-6(W/W): 3.1Å;   
  
**1.99 ± 0.16: the average coodination distance of the 1^st^ six ligands is Mg^2+^ compatible**  
**10: Number of potential Water contacts**   
**4 : Number of second shell contacts involving water or ions**   
**40%: Percentage of satisfied second shell Water contacts**   
• Contact(s): /O_w_/O_w_/O_w_/OP1/  
• Contact code: O_ph_3O_w_  
• Contact(s) in the exclusion zone: /O_w_/O_w_/  
→ Missing octahedral coordination for MG ←  
→ Slightly distorded coord. octahedron (6.2º) ←→2Fo-Fc density between 1.0 and 4.0 !   
Go to ToC

### MG:A:1717

|  | Res. type | Atom | Residue | Distance | Occupancy | B-factor | Closest atom(s) to a 1^st^ shell water | EM\_dens. |
| --- | --- | --- | --- | --- | --- | --- | --- | --- |
| **Ion** | | | |  | 1.0 | 76.0 |  | 2.89 |
 1 | G | OP2 | A:1304 | 1.94 | 1.0 | 81.1 |  | 1.66 | 2 | C | OP1 | A:1303 | 1.96 | 1.0 | 80.6 |  | 2.58 | 3 | Water 1 | | A:2133 | 3.12 | 1.0 | 78.9 | 2.79 [C:OP2]A:1303 | 2.10 | 4 | C | OP2 | A:1303 | 3.81 | 1.0 | 77.3 |  | 3.57 | 5 | G | O2' | A:1242 | 3.97 | 1.0 | 80.4 |  | 2.94 | 6 | G | OP1 | A:1304 | 3.99 | 1.0 | 79.2 |  | 2.11 | 7 | C | O5' | A:1303 | 4.09 | 1.0 | 81.1 |  | 2.90 | 8 | G | O5' | A:1304 | 4.21 | 1.0 | 79.5 |  | 2.28 | 9 | C | O3' | A:1303 | 4.33 | 1.0 | 78.5 |  | 3.00 | 10 | C | O3' | A:1302 | 4.39 | 1.0 | 83.7 |  | 2.67 | 11 | G | N7 | A:1305 | 4.76 | 1.0 | 76.1 |  | 2.80 |

  
**1^st^ shell coordination angles and distances (*trans* in "blue"; *cys* in "green"; deviations in "orange"):**   
• 1-2(OP2/OP1): 86º;   
• 1-2(OP2/OP1): 2.7Å;   
  
**3.13 ± 0.88: the average coodination distance of the 1^st^ six ligands is K^+^ compatible**  
**2: Number of potential Water contacts**   
**1 : Number of second shell contacts involving water or ions**   
**50%: Percentage of satisfied second shell Water contacts**   
• Contact(s): /OP1/OP2/  
• Contact code: *cis-*2O_ph_  
• Contact(s) in the exclusion zone: /O_w_/  
→ Missing octahedral coordination for MG ←  
→2Fo-Fc density between 1.0 and 4.0 !   
Go to ToC

### MG:A:1718

|  | Res. type | Atom | Residue | Distance | Occupancy | B-factor | Closest atom(s) to a 1^st^ shell water | EM\_dens. |
| --- | --- | --- | --- | --- | --- | --- | --- | --- |
| **Ion** | | | |  | 1.0 | 65.1 |  | 1.58 |
 1 | U | O2 | A:1224 | 3.54 | 1.0 | 70.2 |  | 2.90 | 2 | GLY | O | M:99 | 3.99 | 1.0 | 77.4 |  | 1.09 | 3 | G | OP1 | A:1323 | 4.49 | 1.0 | 78.7 |  | 1.83 |

  
• • **4.01 ± 0.39: the average coodination distance of the 1^st^ six ligands could correspond to an hexahydrated Mg^2+^.**  
**2: Number of potential Water contacts**   
**0 : Number of second shell contacts involving water or ions**   
**0%: Percentage of satisfied second shell Water contacts**   
• Contact code: → No contacts ←   
→ Firts MG coordination shell missing ←  
→2Fo-Fc density between 1.0 and 4.0 !   
Go to ToC

### MG:A:1719

|  | Res. type | Atom | Residue | Distance | Occupancy | B-factor | Closest atom(s) to a 1^st^ shell water | EM\_dens. |
| --- | --- | --- | --- | --- | --- | --- | --- | --- |
| **Ion** | | | |  | 1.0 | 74.2 |  | 1.68 |
 1 | Water 1 | | A:3686 | 2.35 | 1.0 | 80.8 | 2.88 [A:N7]A:1329 | 1.62 | 2 | G | O6 | A:1331 | 3.65 | 1.0 | 79.5 |  | 1.41 | 3 | A | OP2 | A:1329 | 3.93 | 1.0 | 82.9 |  | 1.60 | 4 | C | OP2 | A:1328 | 4.50 | 1.0 | 84.2 |  | 1.55 | 5 | U | O4 | A:1330 | 4.67 | 1.0 | 79.1 |  | 2.37 | 6 | A | N7 | A:1329 | 4.74 | 1.0 | 80.4 |  | 2.53 |

  
• • **3.97 ± 0.83: the average coodination distance of the 1^st^ six ligands could correspond to an hexahydrated Mg^2+^.**  
**2: Number of potential Water contacts**   
**1 : Number of second shell contacts involving water or ions**   
**50%: Percentage of satisfied second shell Water contacts**   
• Contact code: → No contacts ←   
• Contact(s) in the exclusion zone: /O_w_/  
→ Firts MG coordination shell missing ←  
→2Fo-Fc density between 1.0 and 4.0 !   
Go to ToC

### MG:A:1720

|  | Res. type | Atom | Residue | Distance | Occupancy | B-factor | Closest atom(s) to a 1^st^ shell water | EM\_dens. |
| --- | --- | --- | --- | --- | --- | --- | --- | --- |
| **Ion** | | | |  | 1.0 | 75.6 |  | 1.95 |
 1 | G | OP2 | A:1334 | 2.17 | 1.0 | 78.7 |  | 1.86 | 2 | G | N7 | A:1300 | 3.82 | 1.0 | 79.1 |  | 2.70 | 3 | G | O5' | A:1334 | 4.18 | 1.0 | 77.5 |  | 2.36 | 4 | G | N7 | A:1334 | 4.50 | 1.0 | 75.6 |  | 3.23 | 5 | G | OP1 | A:1334 | 4.50 | 1.0 | 76.3 |  | 1.39 | 6 | A | O3' | A:1333 | 4.58 | 1.0 | 75.9 |  | 2.84 | 7 | A | O5' | A:1333 | 4.65 | 1.0 | 80.5 |  | 1.75 |

  
• • **3.96 ± 0.84: the average coodination distance of the 1^st^ six ligands could correspond to an hexahydrated Mg^2+^.**  
**0: Number of potential Water contacts**   
• Contact(s): /OP2/  
• Contact code: O_ph_  
→ Missing octahedral coordination for MG ←  
→2Fo-Fc density between 1.0 and 4.0 !   
Go to ToC

### MG:A:1721

|  | Res. type | Atom | Residue | Distance | Occupancy | B-factor | Closest atom(s) to a 1^st^ shell water | EM\_dens. |
| --- | --- | --- | --- | --- | --- | --- | --- | --- |
| **Ion** | | | |  | 1.0 | 68.1 |  | 1.82 |
 1 | A | OP2 | A:1357 | 2.85 | 1.0 | 70.3 |  | 2.55 | 2 | Water 1 | | A:2338 | 2.95 | 1.0 | 68.1 | 2.61 [C:N3]A:1359 | 2.56 | 3 | A | OP1 | A:1357 | 4.15 | 1.0 | 69.0 |  | 2.05 | 4 | C | N4 | A:1359 | 4.18 | 1.0 | 66.0 |  | 3.37 | 5 | G | OP1 | A:1356 | 4.55 | 1.0 | 69.4 |  | 1.74 | 6 | A | O2' | A:1254 | 4.58 | 1.0 | 78.0 |  | 2.78 | 7 | C | N3 | A:1359 | 4.64 | 1.0 | 66.2 |  | 3.77 | 8 | G | O3' | A:1356 | 4.76 | 1.0 | 67.1 |  | 4.71 | 9 | Water 2 | | A:1840 | 4.78 | 1.0 | 70.5 |  | 1.83 |

  
• • **3.88 ± 0.71: the average coodination distance of the 1^st^ six ligands could correspond to an hexahydrated Mg^2+^.**  
**2: Number of potential Water contacts**   
**1 : Number of second shell contacts involving water or ions**   
**50%: Percentage of satisfied second shell Water contacts**   
• Contact code: → No contacts ←   
• Contact(s) in the exclusion zone: /O_w_/OP2/  
→ Consider K in place of MG (1 valid contact(s) for K vs. 0 for MG ←)  
→2Fo-Fc density between 1.0 and 4.0 !   
Go to ToC

### K:A:1722

|  | Res. type | Atom | Residue | Distance | Occupancy | B-factor | Closest atom(s) to a 1^st^ shell water | EM\_dens. |
| --- | --- | --- | --- | --- | --- | --- | --- | --- |
| **Ion** | | | |  | 1.0 | 30.0 |  | 4.05 |
 1 | ASN | OD1 | L:46 | 2.74 | 1.0 | 64.6 |  | 2.85 | 2 | PRO | O | L:45 | 2.80 | 1.0 | 65.2 |  | 2.50 | 3 | G | O6 | A:529 | 2.86 | 1.0 | 70.2 |  | 2.46 | 4 | G | N7 | A:529 | 3.02 | 1.0 | 67.0 |  | 3.58 | 5 | C | N3 | A:518 | 3.17 | 1.0 | 71.3 |  | 3.08 | 6 | C | O2 | A:518 | 3.47 | 1.0 | 69.9 |  | 2.51 | 7 | C | N4 | A:528 | 4.03 | 1.0 | 59.7 |  | 4.09 | 8 | ASN | N | L:46 | 4.09 | 1.0 | 62.0 |  | 3.49 | 9 | C | N1 | A:518 | 4.70 | 1.0 | 69.9 |  | 3.68 |

  
**1^st^ shell coordination angles and distances (*trans* in "blue"; *cys* in "green"; deviations in "orange"):**   
• 1-2(OD1/W): 81; 1-3(OD1/O6): 73; 1-4(OD1/N7): 102; 1-5(OD1/N3): 161; 2-3(W/O6): 97; 2-4(W/N7): 157; 2-5(W/N3): 102; 3-4(O6/N7): 63; 3-5(O6/N3): 88; 4-5(N7/N3): 68;   
• 1-2(OD1/W): 3.6; 1-3(OD1/O6): 3.3; 1-4(OD1/N7): 4.5; 1-5(OD1/N3): 5.8; 2-3(W/O6): 4.2; 2-4(W/N7): 5.7; 2-5(W/N3): 4.6; 3-4(O6/N7): 3.1; 3-5(O6/N3): 4.2; 4-5(N7/N3): 3.5;   
  
**4.36 ± 1.39: the average coodination distance of the 1^st^ six ligands could correspond to an hexahydrated Mg^2+^.**  
**2: Number of potential Water contacts**   
**0 : Number of second shell contacts involving water or ions**   
**0%: Percentage of satisfied second shell Water contacts**   
• Contact(s): /N3_C_/N7/O6/O_back_/OD1_Asn_/  
• Contact code: O_b_2N_b_O_back_O_cno_  
→ Coordination < 6 for K ←  
Go to ToC

### K:A:1723

|  | Res. type | Atom | Residue | Distance | Occupancy | B-factor | Closest atom(s) to a 1^st^ shell water | EM\_dens. |
| --- | --- | --- | --- | --- | --- | --- | --- | --- |
| **Ion** | | | |  | 1.0 | 30.0 |  | 3.85 |
 1 | U | O2' | A:1224 | 2.88 | 1.0 | 73.4 |  | 2.27 | 2 | C | OP2 | A:1322 | 3.09 | 1.0 | 74.1 |  | 2.39 | 3 | GLN | O | M:100 | 3.09 | 1.0 | 75.3 |  | 2.62 | 4 | U | O3' | A:1224 | 3.20 | 1.0 | 74.6 |  | 2.60 | 5 | A | OP1 | A:1225 | 3.31 | 1.0 | 75.2 |  | 2.94 | 6 | GLY | O | M:99 | 4.06 | 1.0 | 77.4 |  | 1.09 | 7 | ARG | N | M:101 | 4.34 | 1.0 | 72.2 |  | 2.83 | 8 | A | O5' | A:1225 | 4.46 | 1.0 | 74.8 |  | 3.48 | 9 | C | OP1 | A:1322 | 4.54 | 1.0 | 76.2 |  | 2.63 | 10 | C | O5' | A:1322 | 4.73 | 1.0 | 74.3 |  | 3.19 |

  
**1^st^ shell coordination angles and distances (*trans* in "blue"; *cys* in "green"; deviations in "orange"):**   
• 1-2(O2'/OP2): 67; 1-3(O2'/W): 170; 2-3(OP2/W): 122;   
• 1-2(O2'/OP2): 3.3; 1-3(O2'/W): 5.9; 2-3(OP2/W): 5.4;   
  
**4.74 ± 1.53: the average coodination distance of the 1^st^ six ligands could correspond to an hexahydrated Mg^2+^.**  
**4: Number of potential Water contacts**   
**0 : Number of second shell contacts involving water or ions**   
**0%: Percentage of satisfied second shell Water contacts**   
• Contact(s): /O2'/O_back_/OP2/  
• Contact code: O_ph_O_r_O_back_  
→ Coordination < 6 for K ←  
→2Fo-Fc density between 1.0 and 4.0 !   
Go to ToC

### K:A:1724

|  | Res. type | Atom | Residue | Distance | Occupancy | B-factor | Closest atom(s) to a 1^st^ shell water | EM\_dens. |
| --- | --- | --- | --- | --- | --- | --- | --- | --- |
| **Ion** | | | |  | 1.0 | 30.0 |  | 21.79 |
 1 | Water 1 | | A:4369 | 2.63 | 1.0 | 43.0 |  | 5.37 | 2 | A | OP1 | A:1508 | 2.68 | 1.0 | 31.1 |  | 11.65 | 3 | C | OP1 | A:1521 | 2.84 | 1.0 | 41.1 |  | 8.17 | 4 | A | OP1 | A:1500 | 2.87 | 1.0 | 32.4 |  | 10.66 | 5 | Water 2 | | A:4005 | 3.07 | 1.0 | 36.2 | 2.12 [MG:MG]A:1634 | 10.57 | 6 | Water 3 | | A:4595 | 3.09 | 1.0 | 50.4 |  | 3.61 | 7 | A | O3' | A:1499 | 3.19 | 1.0 | 37.5 |  | 9.19 | 8 | Water 4 | | A:4333 | 3.26 | 1.0 | 38.6 |  | 6.38 | 9 | **MG** | MG | A:1634 | 3.46 | 1.0 | 37.8 |  | 15.06 | 10 | C | OP2 | A:1521 | 4.08 | 1.0 | 35.8 |  | 10.04 | 11 | A | OP2 | A:1508 | 4.17 | 1.0 | 35.5 |  | 9.31 | 12 | A | O5' | A:1508 | 4.20 | 1.0 | 36.0 |  | 7.83 | 13 | Water 5 | | A:3230 | 4.20 | 1.0 | 41.1 |  | 7.60 | 14 | Water 6 | | A:2964 | 4.21 | 1.0 | 48.1 |  | 3.80 | 15 | A | O5' | A:1500 | 4.41 | 1.0 | 31.7 |  | 10.03 | 16 | Water 7 | | A:4185 | 4.56 | 1.0 | 32.5 |  | 11.09 | 17 | A | O2' | A:1499 | 4.57 | 1.0 | 37.8 |  | 7.44 | 18 | Water 8 | | A:2868 | 4.58 | 1.0 | 52.8 |  | 1.91 | 19 | C | O3' | A:1520 | 4.75 | 1.0 | 34.8 |  | 9.04 |

  
**1^st^ shell coordination angles and distances (*trans* in "blue"; *cys* in "green"; deviations in "orange"):**   
• 1-2(W/OP1): 120; 1-3(W/OP1): 85; 1-4(W/OP1): 106; 1-5(W/W): 165; 1-6(W/W): 79; 1-7(W/O3'): 78; 2-3(OP1/OP1): 151; 2-4(OP1/OP1): 62; 2-5(OP1/W): 62; 2-6(OP1/W): 90; 2-7(OP1/O3'): 111; 3-4(OP1/OP1): 130; 3-5(OP1/W): 99; 3-6(OP1/W): 79; 3-7(OP1/O3'): 89; 4-5(OP1/W): 61; 4-6(OP1/W): 151; 4-7(OP1/O3'): 49; 5-6(W/W): 116; 5-7(W/O3'): 88; 6-7(W/O3'): 154;   
• 1-2(W/OP1): 4.6; 1-3(W/OP1): 3.7; 1-4(W/OP1): 4.4; 1-5(W/W): 5.6; 1-6(W/W): 3.7; 1-7(W/O3'): 3.7; 2-3(OP1/OP1): 5.3; 2-4(OP1/OP1): 2.9; 2-5(OP1/W): 3.0; 2-6(OP1/W): 4.1; 2-7(OP1/O3'): 4.8; 3-4(OP1/OP1): 5.2; 3-5(OP1/W): 4.5; 3-6(OP1/W): 3.8; 3-7(OP1/O3'): 4.2; 4-5(OP1/W): 3.0; 4-6(OP1/W): 5.8; 4-7(OP1/O3'): 2.5; 5-6(W/W): 5.2; 5-7(W/O3'): 4.3; 6-7(W/O3'): 6.1;   
  
**3.94 ± 1.16: the average coodination distance of the 1^st^ six ligands could correspond to an hexahydrated Mg^2+^.**  
**8: Number of potential Water contacts**   
**1 : Number of second shell contacts involving water or ions**   
**12%: Percentage of satisfied second shell Water contacts**   
• Contact(s): /O3'/O_w_/O_w_/O_w_/OP1/OP1/OP1/  
• Contact code: 3O_ph_O_r_3O_w_  
Go to ToC

### K:A:1725

|  | Res. type | Atom | Residue | Distance | Occupancy | B-factor | Closest atom(s) to a 1^st^ shell water | EM\_dens. |
| --- | --- | --- | --- | --- | --- | --- | --- | --- |
| **Ion** | | | |  | 1.0 | 30.0 |  | 6.14 |
 1 | U | O2' | A:789 | 2.63 | 1.0 | 47.1 |  | 6.40 | 2 | G | O6 | A:791 | 2.93 | 1.0 | 42.8 |  | 8.39 | 3 | A | N7 | A:790 | 3.00 | 1.0 | 44.8 |  | 5.80 | 4 | Water 1 | | X:213 | 3.11 | 1.0 | 54.0 |  | 2.66 | 5 | Water 2 | | A:3911 | 3.64 | 1.0 | 44.4 |  | 3.53 | 6 | U | O3' | A:789 | 3.84 | 1.0 | 49.2 |  | 4.25 | 7 | G | N7 | A:791 | 3.99 | 1.0 | 40.8 |  | 6.37 | 8 | A | N6 | A:790 | 4.15 | 1.0 | 41.7 |  | 6.39 | 9 | A | OP1 | X:15 | 4.59 | 1.0 | 63.4 |  | 1.27 |

  
**1^st^ shell coordination angles and distances (*trans* in "blue"; *cys* in "green"; deviations in "orange"):**   
• 1-2(O2'/O6): 86; 1-3(O2'/N7): 83; 1-4(O2'/W): 163; 2-3(O6/N7): 77; 2-4(O6/W): 84; 3-4(N7/W): 82;   
• 1-2(O2'/O6): 3.8; 1-3(O2'/N7): 3.7; 1-4(O2'/W): 5.7; 2-3(O6/N7): 3.7; 2-4(O6/W): 4.0; 3-4(N7/W): 4.0;   
  
**4.55 ± 1.45: the average coodination distance of the 1^st^ six ligands could correspond to an hexahydrated Mg^2+^.**  
**4: Number of potential Water contacts**   
**0 : Number of second shell contacts involving water or ions**   
**0%: Percentage of satisfied second shell Water contacts**   
• Contact(s): /N7/O2'/O6/O_w_/  
• Contact code: O_r_O_b_N_b_O_w_  
→ Coordination < 6 for K ←  
Go to ToC

### K:A:1726

|  | Res. type | Atom | Residue | Distance | Occupancy | B-factor | Closest atom(s) to a 1^st^ shell water | EM\_dens. |
| --- | --- | --- | --- | --- | --- | --- | --- | --- |
| **Ion** | | | |  | 1.0 | 30.0 |  | 5.56 |
 1 | C | O2 | A:880 | 2.79 | 1.0 | 36.9 |  | 7.99 | 2 | C | O2' | A:880 | 2.87 | 1.0 | 39.3 |  | 7.01 | 3 | C | N4 | A:576 | 3.16 | 1.0 | 36.1 |  | 5.85 | 4 | C | OP1 | A:758 | 3.29 | 1.0 | 39.4 |  | 5.73 | 5 | Water 1 | | A:4617 | 3.53 | 1.0 | 52.0 |  | 2.01 | 6 | Water 2 | | A:4590 | 3.65 | 1.0 | 47.1 |  | 3.54 | 7 | G | N2 | A:575 | 3.70 | 1.0 | 30.1 |  | 8.58 | 8 | G | O4' | A:881 | 3.98 | 1.0 | 38.1 |  | 9.89 | 9 | C | O5' | A:758 | 4.23 | 1.0 | 41.1 |  | 5.61 | 10 | C | N1 | A:880 | 4.29 | 1.0 | 31.7 |  | 9.36 | 11 | C | O3' | A:758 | 4.48 | 1.0 | 41.5 |  | 5.65 | 12 | G | N3 | A:821 | 4.60 | 1.0 | 35.1 |  | 9.28 | 13 | Water 3 | | A:2980 | 4.72 | 1.0 | 49.7 |  | 2.53 | 14 | C | O4' | A:880 | 4.72 | 1.0 | 40.3 |  | 7.73 |

  
**1^st^ shell coordination angles and distances (*trans* in "blue"; *cys* in "green"; deviations in "orange"):**   
• 1-2(O2/O2'): 81; 1-3(O2/N4): 133; 2-3(O2'/N4): 75;   
• 1-2(O2/O2'): 3.7; 1-3(O2/N4): 5.5; 2-3(O2'/N4): 3.7;   
  
**4.50 ± 1.37: the average coodination distance of the 1^st^ six ligands could correspond to an hexahydrated Mg^2+^.**  
**4: Number of potential Water contacts**   
**0 : Number of second shell contacts involving water or ions**   
**0%: Percentage of satisfied second shell Water contacts**   
• Contact(s): /N4/O2/O2'/  
• Contact code: O_r_O_b_Anion  
→ Coordination < 6 for K ←  
Go to ToC

### K:A:1727

|  | Res. type | Atom | Residue | Distance | Occupancy | B-factor | Closest atom(s) to a 1^st^ shell water | EM\_dens. |
| --- | --- | --- | --- | --- | --- | --- | --- | --- |
| **Ion** | | | |  | 1.0 | 30.0 |  | 6.36 |
 1 | G | O6 | A:105 | 2.59 | 1.0 | 52.0 |  | 5.62 | 2 | Water 1 | | A:3040 | 2.80 | 1.0 | 56.5 | 2.75 [G:N7]A:105 | 2.96 | 3 | Water 2 | | A:3566 | 2.81 | 1.0 | 59.1 | 2.86 [G:O6]A:61 | 2.47 | 4 | U | O4 | A:62 | 2.90 | 1.0 | 53.8 |  | 3.94 | 5 | G | O6 | A:104 | 2.99 | 1.0 | 57.4 |  | 4.73 | 6 | G | O6 | A:61 | 3.33 | 1.0 | 51.5 |  | 4.38 | 7 | Water 3 | | A:4608 | 3.40 | 1.0 | 63.7 |  | 2.17 | 8 | C | N4 | A:106 | 3.78 | 1.0 | 45.2 |  | 5.26 | 9 | G | N7 | A:105 | 4.08 | 1.0 | 49.5 |  | 5.16 | 10 | Water 4 | | A:1988 | 4.45 | 1.0 | 55.4 |  | 2.59 |

  
**1^st^ shell coordination angles and distances (*trans* in "blue"; *cys* in "green"; deviations in "orange"):**   
• 1-2(O6/W): 90; 1-3(O6/W): 122; 1-4(O6/O4): 83; 1-5(O6/O6): 80; 2-3(W/W): 111; 2-4(W/O4): 171; 2-5(W/O6): 78; 3-4(W/O4): 69; 3-5(W/O6): 155; 4-5(O4/O6): 106;   
• 1-2(O6/W): 3.8; 1-3(O6/W): 4.7; 1-4(O6/O4): 3.6; 1-5(O6/O6): 3.6; 2-3(W/W): 4.6; 2-4(W/O4): 5.7; 2-5(W/O6): 3.6; 3-4(W/O4): 3.2; 3-5(W/O6): 5.7; 4-5(O4/O6): 4.7;   
  
**4.10 ± 1.26: the average coodination distance of the 1^st^ six ligands could correspond to an hexahydrated Mg^2+^.**  
**6: Number of potential Water contacts**   
**2 : Number of second shell contacts involving water or ions**   
**33%: Percentage of satisfied second shell Water contacts**   
• Contact(s): /O4/O6/O_w_/O_w_/  
• Contact code: 2O_b_2O_w_  
• Contact(s) in the exclusion zone: /O6/  
→ Coordination < 6 for K ←  
Go to ToC

### K:A:1728

|  | Res. type | Atom | Residue | Distance | Occupancy | B-factor | Closest atom(s) to a 1^st^ shell water | EM\_dens. |
| --- | --- | --- | --- | --- | --- | --- | --- | --- |
| **Ion** | | | |  | 1.0 | 30.0 |  | 7.60 |
 1 | C | O2 | A:372 | 2.49 | 1.0 | 61.9 |  | 3.23 | 2 | C | O2' | A:372 | 2.66 | 1.0 | 61.7 |  | 3.30 | 3 | U | O4 | A:375 | 2.76 | 1.0 | 60.3 |  | 3.86 | 4 | G | O6 | A:376 | 2.93 | 1.0 | 60.4 |  | 3.71 | 5 | U | O4 | A:387 | 3.00 | 1.0 | 60.2 |  | 2.73 | 6 | Water 1 | | A:3541 | 3.28 | 1.0 | 60.5 | 2.85 [G:N7]A:376 | 2.54 | 7 | Water 2 | | A:3846 | 3.29 | 1.0 | 62.0 |  | 2.31 | 8 | U | N3 | A:387 | 4.02 | 1.0 | 54.7 |  | 4.06 | 9 | C | N3 | A:372 | 4.27 | 1.0 | 57.8 |  | 4.03 | 10 | C | N1 | A:372 | 4.31 | 1.0 | 60.6 |  | 4.15 | 11 | Water 3 | | A:2453 | 4.36 | 1.0 | 60.2 |  | 2.42 | 12 | G | N7 | A:376 | 4.48 | 1.0 | 56.5 |  | 3.53 |

  
**1^st^ shell coordination angles and distances (*trans* in "blue"; *cys* in "green"; deviations in "orange"):**   
• 1-2(O2/O2'): 68; 1-3(O2/O4): 83; 1-4(O2/O6): 105; 1-5(O2/O4): 77; 2-3(O2'/O4): 142; 2-4(O2'/O6): 137; 2-5(O2'/O4): 60; 3-4(O4/O6): 73; 3-5(O4/O4): 137; 4-5(O6/O4): 77;   
• 1-2(O2/O2'): 2.9; 1-3(O2/O4): 3.5; 1-4(O2/O6): 4.3; 1-5(O2/O4): 3.4; 2-3(O2'/O4): 5.1; 2-4(O2'/O6): 5.2; 2-5(O2'/O4): 2.9; 3-4(O4/O6): 3.4; 3-5(O4/O4): 5.4; 4-5(O6/O4): 3.7;   
  
**4.07 ± 1.28: the average coodination distance of the 1^st^ six ligands could correspond to an hexahydrated Mg^2+^.**  
**4: Number of potential Water contacts**   
**1 : Number of second shell contacts involving water or ions**   
**25%: Percentage of satisfied second shell Water contacts**   
• Contact(s): /O2'/O4/O4/O6/  
• Contact code: O_r_3O_b_  
• Contact(s) in the exclusion zone: /O2/  
→ Coordination < 6 for K ←  
Go to ToC

### K:A:1729

|  | Res. type | Atom | Residue | Distance | Occupancy | B-factor | Closest atom(s) to a 1^st^ shell water | EM\_dens. |
| --- | --- | --- | --- | --- | --- | --- | --- | --- |
| **Ion** | | | |  | 1.0 | 30.0 |  | 4.50 |
 1 | A | OP2 | A:914 | 2.69 | 1.0 | 50.4 |  | 2.80 | 2 | Water 1 | | A:2402 | 2.95 | 1.0 | 56.1 |  | 1.58 | 3 | G7M | OP1 | A:527 | 3.00 | 1.0 | 64.9 |  | 1.54 | 4 | U | O2' | A:12 | 3.05 | 1.0 | 44.8 |  | 5.34 | 5 | U | O3' | A:12 | 3.12 | 1.0 | 45.6 |  | 5.24 | 6 | Water 2 | | A:3232 | 3.29 | 1.0 | 53.0 |  | 1.99 | 7 | C | O3' | A:526 | 3.50 | 1.0 | 58.9 |  | 2.67 | 8 | U | OP1 | A:13 | 4.04 | 1.0 | 48.7 |  | 4.04 | 9 | G7M | OP2 | A:527 | 4.33 | 1.0 | 63.0 |  | 1.94 | 10 | U | O5' | A:13 | 4.47 | 1.0 | 45.2 |  | 5.09 | 11 | A | OP1 | A:914 | 4.60 | 1.0 | 50.0 |  | 2.98 | 12 | A | O5' | A:914 | 4.65 | 1.0 | 50.1 |  | 1.96 |

  
**1^st^ shell coordination angles and distances (*trans* in "blue"; *cys* in "green"; deviations in "orange"):**   
• 1-2(OP2/W): 168; 1-3(OP2/OP1): 132; 1-4(OP2/O2'): 63; 1-5(OP2/O3'): 107; 2-3(W/OP1): 52; 2-4(W/O2'): 120; 2-5(W/O3'): 71; 3-4(OP1/O2'): 148; 3-5(OP1/O3'): 121; 4-5(O2'/O3'): 54;   
• 1-2(OP2/W): 5.6; 1-3(OP2/OP1): 5.2; 1-4(OP2/O2'): 3.0; 1-5(OP2/O3'): 4.7; 2-3(W/OP1): 2.6; 2-4(W/O2'): 5.2; 2-5(W/O3'): 3.5; 3-4(OP1/O2'): 5.8; 3-5(OP1/O3'): 5.3; 4-5(O2'/O3'): 2.8;   
  
**4.27 ± 1.31: the average coodination distance of the 1^st^ six ligands could correspond to an hexahydrated Mg^2+^.**  
**4: Number of potential Water contacts**   
**0 : Number of second shell contacts involving water or ions**   
**0%: Percentage of satisfied second shell Water contacts**   
• Contact(s): /O2'/O3'/O_w_/OP1/OP2/  
• Contact code: 2O_ph_2O_r_O_w_  
→ Coordination < 6 for K ←  
Go to ToC

### K:A:1730

|  | Res. type | Atom | Residue | Distance | Occupancy | B-factor | Closest atom(s) to a 1^st^ shell water | EM\_dens. |
| --- | --- | --- | --- | --- | --- | --- | --- | --- |
| **Ion** | | | |  | 1.0 | 30.0 |  | 9.42 |
 1 | G | OP2 | A:567 | 2.64 | 1.0 | 43.3 |  | 6.00 | 2 | C | OP2 | A:564 | 2.77 | 1.0 | 47.7 |  | 5.23 | 3 | A | O2' | A:563 | 2.89 | 1.0 | 42.8 |  | 5.88 | 4 | U | OP2 | A:565 | 2.92 | 1.0 | 52.7 |  | 3.86 | 5 | Water 1 | | A:2495 | 3.17 | 1.0 | 51.1 | 2.64 [U:OP1]A:565 | 2.20 | 6 | Water 2 | | A:2882 | 3.23 | 1.0 | 48.2 | 2.72 [G:N7]A:567 | 3.28 | 7 | Water 3 | | L:203 | 3.38 | 1.0 | 52.6 |  | 2.46 | 8 | A | O3' | A:563 | 3.61 | 1.0 | 44.5 |  | 5.66 | 9 | G | O3' | A:566 | 3.77 | 1.0 | 46.5 |  | 4.88 | 10 | G | O2' | A:566 | 4.10 | 1.0 | 45.5 |  | 6.02 | 11 | Water 4 | | A:3318 | 4.10 | 1.0 | 51.7 |  | 2.18 | 12 | G | O5' | A:567 | 4.29 | 1.0 | 47.7 |  | 3.91 | 13 | U | OP1 | A:565 | 4.36 | 1.0 | 49.5 |  | 3.58 | 14 | C | O5' | A:564 | 4.42 | 1.0 | 46.4 |  | 4.85 | 15 | G | N7 | A:567 | 4.42 | 1.0 | 37.8 |  | 7.63 | 16 | C | O3' | A:564 | 4.80 | 1.0 | 46.5 |  | 5.36 |

  
**1^st^ shell coordination angles and distances (*trans* in "blue"; *cys* in "green"; deviations in "orange"):**   
• 1-2(OP2/OP2): 169; 1-3(OP2/O2'): 84; 1-4(OP2/OP2): 88; 1-5(OP2/W): 55; 2-3(OP2/O2'): 84; 2-4(OP2/OP2): 89; 2-5(OP2/W): 133; 3-4(O2'/OP2): 81; 3-5(O2'/W): 129; 4-5(OP2/W): 70;   
• 1-2(OP2/OP2): 5.4; 1-3(OP2/O2'): 3.7; 1-4(OP2/OP2): 3.9; 1-5(OP2/W): 2.7; 2-3(OP2/O2'): 3.8; 2-4(OP2/OP2): 4.0; 2-5(OP2/W): 5.5; 3-4(O2'/OP2): 3.8; 3-5(O2'/W): 5.5; 4-5(OP2/W): 3.5;   
  
**4.10 ± 1.24: the average coodination distance of the 1^st^ six ligands could correspond to an hexahydrated Mg^2+^.**  
**6: Number of potential Water contacts**   
**2 : Number of second shell contacts involving water or ions**   
**33%: Percentage of satisfied second shell Water contacts**   
• Contact(s): /O2'/O_w_/OP2/OP2/OP2/  
• Contact code: 3O_ph_O_r_O_w_  
→ Coordination < 6 for K ←  
Go to ToC

### K:A:1731

|  | Res. type | Atom | Residue | Distance | Occupancy | B-factor | Closest atom(s) to a 1^st^ shell water | EM\_dens. |
| --- | --- | --- | --- | --- | --- | --- | --- | --- |
| **Ion** | | | |  | 1.0 | 30.0 |  | 11.87 |
 1 | G | O6 | A:925 | 2.87 | 1.0 | 41.0 |  | 6.60 | 2 | Water 1 | | A:3079 | 2.88 | 1.0 | 43.2 | 2.76 [G:N7]A:925 | 5.09 | 3 | U | O4 | A:1390 | 2.99 | 1.0 | 43.4 |  | 7.73 | 4 | G | O6 | A:927 | 3.13 | 1.0 | 42.3 |  | 7.54 | 5 | Water 2 | | A:3616 | 3.15 | 1.0 | 51.7 | 2.87 [C:N4]A:924 | 4.17 | 6 | Water 3 | | A:3455 | 3.21 | 1.0 | 52.9 |  | 2.72 | 7 | Water 4 | | A:4629 | 3.30 | 1.0 | 58.7 |  | 2.04 | 8 | U | O4 | A:1391 | 3.35 | 1.0 | 42.4 |  | 5.95 | 9 | Water 5 | | A:1980 | 3.63 | 1.0 | 44.5 | 2.46 [G:N7]A:927 | 3.68 | 10 | G | N7 | A:925 | 4.17 | 1.0 | 34.6 |  | 9.08 | 11 | Water 6 | | A:2277 | 4.25 | 1.0 | 55.8 |  | 2.67 | 12 | G | N7 | A:927 | 4.57 | 1.0 | 38.0 |  | 7.70 |

  
**1^st^ shell coordination angles and distances (*trans* in "blue"; *cys* in "green"; deviations in "orange"):**   
• 1-2(O6/W): 88; 1-3(O6/O4): 122; 1-4(O6/O6): 69; 1-5(O6/W): 115; 2-3(W/O4): 149; 2-4(W/O6): 122; 2-5(W/W): 92; 3-4(O4/O6): 76; 3-5(O4/W): 74; 4-5(O6/W): 146;   
• 1-2(O6/W): 4.0; 1-3(O6/O4): 5.1; 1-4(O6/O6): 3.4; 1-5(O6/W): 5.1; 2-3(W/O4): 5.7; 2-4(W/O6): 5.3; 2-5(W/W): 4.3; 3-4(O4/O6): 3.8; 3-5(O4/W): 3.7; 4-5(O6/W): 6.0;   
  
**4.15 ± 1.21: the average coodination distance of the 1^st^ six ligands could correspond to an hexahydrated Mg^2+^.**  
**8: Number of potential Water contacts**   
**2 : Number of second shell contacts involving water or ions**   
**25%: Percentage of satisfied second shell Water contacts**   
• Contact(s): /O4/O6/O6/O_w_/O_w_/  
• Contact code: 3O_b_2O_w_  
→ Coordination < 6 for K ←  
Go to ToC

### K:A:1732

|  | Res. type | Atom | Residue | Distance | Occupancy | B-factor | Closest atom(s) to a 1^st^ shell water | EM\_dens. |
| --- | --- | --- | --- | --- | --- | --- | --- | --- |
| **Ion** | | | |  | 1.0 | 30.0 |  | 7.93 |
 1 | Water 1 | | A:3473 | 2.79 | 1.0 | 56.7 | 2.84 [G:N7]A:1058 | 3.84 | 2 | U | O4 | A:1199 | 2.85 | 1.0 | 59.1 |  | 3.68 | 3 | G | O6 | A:1058 | 2.89 | 1.0 | 56.9 |  | 3.87 | 4 | Water 2 | | A:1955 | 2.92 | 1.0 | 64.7 | 2.44 [G:OP2]A:1057 | 2.73 | 5 | Water 3 | | A:3813 | 3.09 | 1.0 | 57.9 | 2.92 [G:O6]A:1198 | 2.37 | 6 | G | O6 | A:1198 | 3.26 | 1.0 | 55.8 |  | 4.13 | 7 | C | N4 | A:1059 | 3.46 | 1.0 | 56.0 |  | 4.07 | 8 | G | N2 | A:1053 | 3.49 | 1.0 | 56.8 |  | 4.17 | 9 | G | OP2 | A:1057 | 4.14 | 1.0 | 60.2 |  | 3.52 | 10 | Water 4 | | A:4423 | 4.25 | 1.0 | 56.0 |  | 2.06 | 11 | G | N3 | A:1053 | 4.30 | 1.0 | 57.9 |  | 4.98 | 12 | Water 5 | | A:3474 | 4.31 | 1.0 | 60.9 |  | 2.62 | 13 | Water 6 | | A:4060 | 4.36 | 1.0 | 60.0 | 2.99 [C:N4]A:1059 | 2.39 | 14 | G | N7 | A:1058 | 4.38 | 1.0 | 57.5 |  | 4.18 | 15 | Water 7 | | A:3525 | 4.52 | 1.0 | 60.2 | 2.85 [G:N3]A:1053 | 2.79 | 16 | Water 8 | | A:2460 | 4.79 | 1.0 | 55.5 |  | 4.31 |

  
**1^st^ shell coordination angles and distances (*trans* in "blue"; *cys* in "green"; deviations in "orange"):**   
• 1-2(W/O4): 164; 1-3(W/O6): 83; 1-4(W/W): 83; 1-5(W/W): 97; 2-3(O4/O6): 81; 2-4(O4/W): 107; 2-5(O4/W): 91; 3-4(O6/W): 116; 3-5(O6/W): 133; 4-5(W/W): 110;   
• 1-2(W/O4): 5.6; 1-3(W/O6): 3.8; 1-4(W/W): 3.8; 1-5(W/W): 4.4; 2-3(O4/O6): 3.7; 2-4(O4/W): 4.6; 2-5(O4/W): 4.2; 3-4(O6/W): 4.9; 3-5(O6/W): 5.5; 4-5(W/W): 4.9;   
  
**4.13 ± 1.23: the average coodination distance of the 1^st^ six ligands could correspond to an hexahydrated Mg^2+^.**  
**6: Number of potential Water contacts**   
**3 : Number of second shell contacts involving water or ions**   
**50%: Percentage of satisfied second shell Water contacts**   
• Contact(s): /O4/O6/O_w_/O_w_/O_w_/  
• Contact code: 2O_b_3O_w_  
→ Coordination < 6 for K ←  
Go to ToC

### K:A:1733

|  | Res. type | Atom | Residue | Distance | Occupancy | B-factor | Closest atom(s) to a 1^st^ shell water | EM\_dens. |
| --- | --- | --- | --- | --- | --- | --- | --- | --- |
| **Ion** | | | |  | 1.0 | 30.0 |  | 14.26 |
 1 | Water 1 | | A:2976 | 2.68 | 1.0 | 51.2 | 1.94 [MG:MG]A:1681 | 5.09 | 2 | G | O2' | A:115 | 2.82 | 1.0 | 47.5 |  | 4.96 | 3 | G | O3' | A:115 | 2.88 | 1.0 | 48.5 |  | 6.77 | 4 | G | N7 | A:117 | 2.98 | 1.0 | 42.0 |  | 7.75 | 5 | G | O6 | A:117 | 3.06 | 1.0 | 46.1 |  | 6.99 | 6 | G | O4' | A:289 | 3.15 | 1.0 | 47.4 |  | 5.87 | 7 | G | O5' | A:289 | 3.16 | 1.0 | 42.6 |  | 6.94 | 8 | A | OP2 | A:116 | 3.31 | 1.0 | 45.6 |  | 5.98 | 9 | A | N7 | A:116 | 3.58 | 1.0 | 44.6 |  | 5.95 | 10 | **MG** | MG | A:1681 | 3.94 | 1.0 | 47.7 |  | 9.46 | 11 | G | N3 | A:115 | 4.11 | 1.0 | 36.9 |  | 8.05 | 12 | G | OP2 | A:289 | 4.13 | 1.0 | 42.5 |  | 8.25 | 13 | U | O4 | A:118 | 4.18 | 1.0 | 48.1 |  | 5.96 | 14 | A | O5' | A:116 | 4.68 | 1.0 | 52.5 |  | 4.98 | 15 | G | O4' | A:115 | 4.72 | 1.0 | 45.2 |  | 6.97 | 16 | Water 2 | | A:2596 | 4.79 | 1.0 | 46.9 |  | 3.52 |

  
**1^st^ shell coordination angles and distances (*trans* in "blue"; *cys* in "green"; deviations in "orange"):**   
• 1-2(W/O2'): 135; 1-3(W/O3'): 101; 1-4(W/N7): 64; 1-5(W/O6): 101; 1-6(W/O4'): 95; 1-7(W/O5'): 66; 2-3(O2'/O3'): 57; 2-4(O2'/N7): 158; 2-5(O2'/O6): 116; 2-6(O2'/O4'): 84; 2-7(O2'/O5'): 77; 3-4(O3'/N7): 114; 3-5(O3'/O6): 150; 3-6(O3'/O4'): 137; 3-7(O3'/O5'): 96; 4-5(N7/O6): 60; 4-6(N7/O4'): 109; 4-7(N7/O5'): 125; 5-6(O6/O4'): 60; 5-7(O6/O5'): 111; 6-7(O4'/O5'): 55;   
• 1-2(W/O2'): 5.1; 1-3(W/O3'): 4.3; 1-4(W/N7): 3.0; 1-5(W/O6): 4.4; 1-6(W/O4'): 4.3; 1-7(W/O5'): 3.2; 2-3(O2'/O3'): 2.7; 2-4(O2'/N7): 5.7; 2-5(O2'/O6): 5.0; 2-6(O2'/O4'): 4.0; 2-7(O2'/O5'): 3.7; 3-4(O3'/N7): 4.9; 3-5(O3'/O6): 5.7; 3-6(O3'/O4'): 5.6; 3-7(O3'/O5'): 4.5; 4-5(N7/O6): 3.0; 4-6(N7/O4'): 5.0; 4-7(N7/O5'): 5.5; 5-6(O6/O4'): 3.1; 5-7(O6/O5'): 5.1; 6-7(O4'/O5'): 2.9;   
  
**4.01 ± 1.18: the average coodination distance of the 1^st^ six ligands could correspond to an hexahydrated Mg^2+^.**  
**2: Number of potential Water contacts**   
**1 : Number of second shell contacts involving water or ions**   
**50%: Percentage of satisfied second shell Water contacts**   
• Contact(s): /N7/O2'/O3'/O4'/O5'/O6/O_w_/  
• Contact code: 4O_r_O_b_N_b_O_w_  
Go to ToC

### K:A:1734

|  | Res. type | Atom | Residue | Distance | Occupancy | B-factor | Closest atom(s) to a 1^st^ shell water | EM\_dens. |
| --- | --- | --- | --- | --- | --- | --- | --- | --- |
| **Ion** | | | |  | 1.0 | 30.0 |  | 5.31 |
 1 | Water 1 | | A:3128 | 2.83 | 1.0 | 55.6 | 2.77 [G:N7]A:293 | 2.33 | 2 | G | O6 | A:293 | 2.83 | 1.0 | 46.4 |  | 5.44 | 3 | G | O6 | A:305 | 2.94 | 1.0 | 51.3 |  | 5.41 | 4 | U | O4 | A:304 | 3.00 | 1.0 | 48.3 |  | 4.89 | 5 | Water 2 | | A:3557 | 3.84 | 1.0 | 52.5 |  | 2.15 | 6 | U | N3 | A:304 | 3.88 | 1.0 | 40.3 |  | 7.73 | 7 | A | N6 | A:303 | 3.97 | 1.0 | 41.4 |  | 5.95 | 8 | G | N1 | A:305 | 4.17 | 1.0 | 44.6 |  | 6.63 | 9 | G | N7 | A:293 | 4.43 | 1.0 | 47.3 |  | 5.42 | 10 | Water 3 | | A:2560 | 4.47 | 1.0 | 52.5 |  | 3.47 | 11 | U | O4 | A:294 | 4.70 | 1.0 | 49.6 |  | 4.21 | 12 | G | OP2 | A:292 | 4.77 | 1.0 | 47.7 |  | 5.43 |

  
**1^st^ shell coordination angles and distances (*trans* in "blue"; *cys* in "green"; deviations in "orange"):**   
• 1-2(W/O6): 80; 1-3(W/O6): 105; 1-4(W/O4): 165; 2-3(O6/O6): 69; 2-4(O6/O4): 85; 3-4(O6/O4): 69;   
• 1-2(W/O6): 3.7; 1-3(W/O6): 4.6; 1-4(W/O4): 5.8; 2-3(O6/O6): 3.3; 2-4(O6/O4): 3.9; 3-4(O6/O4): 3.4;   
  
**4.58 ± 1.46: the average coodination distance of the 1^st^ six ligands could correspond to an hexahydrated Mg^2+^.**  
**4: Number of potential Water contacts**   
**1 : Number of second shell contacts involving water or ions**   
**25%: Percentage of satisfied second shell Water contacts**   
• Contact(s): /O4/O6/O6/O_w_/  
• Contact code: 3O_b_O_w_  
→ Coordination < 6 for K ←  
Go to ToC

### K:A:1735

|  | Res. type | Atom | Residue | Distance | Occupancy | B-factor | Closest atom(s) to a 1^st^ shell water | EM\_dens. |
| --- | --- | --- | --- | --- | --- | --- | --- | --- |
| **Ion** | | | |  | 1.0 | 30.0 |  | 7.16 |
 1 | **MG** | MG | A:1631 | 2.32 | 1.0 | 63.4 |  | 3.08 | 2 | Water 1 | | A:3679 | 2.45 | 1.0 | 47.4 |  | 2.67 | 3 | Water 2 | | A:2817 | 2.76 | 1.0 | 45.2 |  | 5.61 | 4 | C | O3' | A:817 | 2.90 | 1.0 | 42.6 |  | 7.12 | 5 | U | OP1 | A:1528 | 2.93 | 1.0 | 40.2 |  | 5.99 | 6 | G | OP2 | A:818 | 2.93 | 1.0 | 46.4 |  | 4.95 | 7 | Water 3 | | A:3193 | 3.12 | 1.0 | 48.4 |  | 3.76 | 8 | U | O3' | A:1527 | 3.16 | 1.0 | 40.9 |  | 8.50 | 9 | Water 4 | | A:4208 | 3.25 | 1.0 | 52.5 |  | 2.19 | 10 | C | O2' | A:817 | 3.27 | 1.0 | 40.2 |  | 7.85 | 11 | G | O5' | A:818 | 4.11 | 1.0 | 45.7 |  | 5.88 | 12 | Water 5 | | A:3532 | 4.37 | 1.0 | 51.0 | 2.85 [U:OP2]A:1528 | 1.93 | 13 | Water 6 | | A:4063 | 4.37 | 1.0 | 51.2 |  | 1.97 | 14 | U | OP2 | A:1528 | 4.47 | 1.0 | 39.0 |  | 7.68 |

  
**1^st^ shell coordination angles and distances (*trans* in "blue"; *cys* in "green"; deviations in "orange"):**   
• 1-2(MG/W): 53; 1-3(MG/W): 156; 1-4(MG/O3'): 85; 1-5(MG/OP1): 111; 1-6(MG/OP2): 40; 1-7(MG/W): 43; 1-8(MG/O3'): 129; 2-3(W/W): 137; 2-4(W/O3'): 69; 2-5(W/OP1): 139; 2-6(W/OP2): 64; 2-7(W/W): 56; 2-8(W/O3'): 108; 3-4(W/O3'): 82; 3-5(W/OP1): 75; 3-6(W/OP2): 118; 3-7(W/W): 158; 3-8(W/O3'): 73; 4-5(O3'/OP1): 153; 4-6(O3'/OP2): 50; 4-7(O3'/W): 119; 4-8(O3'/O3'): 136; 5-6(OP1/OP2): 133; 5-7(OP1/W): 86; 5-8(OP1/O3'): 48; 6-7(OP2/W): 82; 6-8(OP2/O3'): 169; 7-8(W/O3'): 86;   
• 1-2(MG/W): 2.1; 1-3(MG/W): 5.0; 1-4(MG/O3'): 3.6; 1-5(MG/OP1): 4.3; 1-6(MG/OP2): 1.9; 1-7(MG/W): 2.1; 1-8(MG/O3'): 5.0; 2-3(W/W): 4.9; 2-4(W/O3'): 3.0; 2-5(W/OP1): 5.0; 2-6(W/OP2): 2.9; 2-7(W/W): 2.7; 2-8(W/O3'): 4.6; 3-4(W/O3'): 3.7; 3-5(W/OP1): 3.5; 3-6(W/OP2): 4.9; 3-7(W/W): 5.8; 3-8(W/O3'): 3.5; 4-5(O3'/OP1): 5.7; 4-6(O3'/OP2): 2.5; 4-7(O3'/W): 5.2; 4-8(O3'/O3'): 5.6; 5-6(OP1/OP2): 5.4; 5-7(OP1/W): 4.1; 5-8(OP1/O3'): 2.5; 6-7(OP2/W): 4.0; 6-8(OP2/O3'): 6.1; 7-8(W/O3'): 4.3;   
  
**3.76 ± 1.13: the average coodination distance of the 1^st^ six ligands could correspond to an hexahydrated Mg^2+^.**  
**6: Number of potential Water contacts**   
**0 : Number of second shell contacts involving water or ions**   
**0%: Percentage of satisfied second shell Water contacts**   
• Contact(s): /O3'/O3'/O_w_/O_w_/OP1/OP2/  
• Contact code: 2O_ph_2O_r_2O_w_  
• Contact(s) in the exclusion zone: /O_w_/  
Go to ToC

### K:A:1736

|  | Res. type | Atom | Residue | Distance | Occupancy | B-factor | Closest atom(s) to a 1^st^ shell water | EM\_dens. |
| --- | --- | --- | --- | --- | --- | --- | --- | --- |
| **Ion** | | | |  | 1.0 | 30.0 |  | 4.13 |
 1 | A | OP1 | A:315 | 2.66 | 1.0 | 50.8 |  | 3.06 | 2 | Water 1 | | A:3889 | 2.88 | 1.0 | 53.4 |  | 2.52 | 3 | G | OP2 | A:318 | 2.96 | 1.0 | 50.6 |  | 2.79 | 4 | C | O3' | A:314 | 3.08 | 1.0 | 49.4 |  | 4.10 | 5 | Water 2 | | A:4169 | 3.43 | 1.0 | 55.2 |  | 2.13 | 6 | G | OP1 | A:318 | 4.25 | 1.0 | 54.9 |  | 2.56 | 7 | Water 3 | | A:4660 | 4.29 | 1.0 | 54.8 |  | 1.74 | 8 | A | O5' | A:315 | 4.45 | 1.0 | 43.3 |  | 6.11 | 9 | Water 4 | | A:2542 | 4.58 | 1.0 | 55.6 |  | 2.29 | 10 | Water 5 | | A:3729 | 4.60 | 1.0 | 57.9 |  | 1.65 | 11 | A | OP2 | A:315 | 4.62 | 1.0 | 49.6 |  | 5.19 |

  
**1^st^ shell coordination angles and distances (*trans* in "blue"; *cys* in "green"; deviations in "orange"):**   
• 1-2(OP1/W): 98; 1-3(OP1/OP2): 114; 1-4(OP1/O3'): 51; 2-3(W/OP2): 93; 2-4(W/O3'): 64; 3-4(OP2/O3'): 146;   
• 1-2(OP1/W): 4.2; 1-3(OP1/OP2): 4.7; 1-4(OP1/O3'): 2.5; 2-3(W/OP2): 4.2; 2-4(W/O3'): 3.2; 3-4(OP2/O3'): 5.8;   
  
**4.67 ± 1.56: the average coodination distance of the 1^st^ six ligands could correspond to an hexahydrated Mg^2+^.**  
**6: Number of potential Water contacts**   
**0 : Number of second shell contacts involving water or ions**   
**0%: Percentage of satisfied second shell Water contacts**   
• Contact(s): /O3'/O_w_/OP1/OP2/  
• Contact code: 2O_ph_O_r_O_w_  
→ Coordination < 6 for K ←  
Go to ToC

### K:A:1737

|  | Res. type | Atom | Residue | Distance | Occupancy | B-factor | Closest atom(s) to a 1^st^ shell water | EM\_dens. |
| --- | --- | --- | --- | --- | --- | --- | --- | --- |
| **Ion** | | | |  | 1.0 | 30.0 |  | 4.22 |
 1 | G | OP2 | A:760 | 2.29 | 1.0 | 44.4 |  | 3.55 | 2 | A | OP1 | A:759 | 2.43 | 1.0 | 45.1 |  | 4.05 | 3 | A | OP2 | A:759 | 3.56 | 1.0 | 44.2 |  | 5.51 | 4 | G | O6 | A:761 | 3.86 | 1.0 | 40.0 |  | 5.95 | 5 | A | O5' | A:759 | 3.92 | 1.0 | 41.3 |  | 8.27 | 6 | Water 1 | | A:4323 | 3.93 | 1.0 | 47.5 |  | 2.43 | 7 | G | N7 | A:761 | 4.19 | 1.0 | 39.0 |  | 6.55 | 8 | Water 2 | | A:4414 | 4.29 | 1.0 | 49.8 |  | 2.23 | 9 | C | N3 | A:576 | 4.46 | 1.0 | 36.4 |  | 7.79 | 10 | G | O5' | A:760 | 4.48 | 1.0 | 49.0 |  | 4.77 | 11 | G | OP1 | A:760 | 4.61 | 1.0 | 49.2 |  | 3.73 | 12 | A | O3' | A:759 | 4.62 | 1.0 | 39.9 |  | 6.14 | 13 | Water 3 | | A:3005 | 4.66 | 1.0 | 56.3 |  | 2.10 | 14 | C | O3' | A:758 | 4.76 | 1.0 | 41.5 |  | 5.65 |

  
**1^st^ shell coordination angles and distances (*trans* in "blue"; *cys* in "green"; deviations in "orange"):**   
• 1-2(OP2/OP1): 94;   
• 1-2(OP2/OP1): 3.5;   
  
**4.74 ± 1.60: the average coodination distance of the 1^st^ six ligands could correspond to an hexahydrated Mg^2+^.**  
**4: Number of potential Water contacts**   
**0 : Number of second shell contacts involving water or ions**   
**0%: Percentage of satisfied second shell Water contacts**   
• Contact code: → No contacts ←   
• Contact(s) in the exclusion zone: /OP1/OP2/  
→ Consider MG in place of K (1 valid contact(s) for MG vs. 0 for K ←)  
Go to ToC

### K:A:1738

|  | Res. type | Atom | Residue | Distance | Occupancy | B-factor | Closest atom(s) to a 1^st^ shell water | EM\_dens. |
| --- | --- | --- | --- | --- | --- | --- | --- | --- |
| **Ion** | | | |  | 1.0 | 30.0 |  | 8.28 |
 1 | Water 1 | | A:4639 | 2.55 | 1.0 | 61.8 |  | 2.35 | 2 | Water 2 | | A:2439 | 2.62 | 1.0 | 49.9 | 2.63 [G:O6]A:887 | 3.40 | 3 | G | O6 | A:886 | 2.76 | 1.0 | 42.9 |  | 7.28 | 4 | U | O4 | A:911 | 2.90 | 1.0 | 48.6 |  | 5.84 | 5 | G | O6 | A:885 | 2.97 | 1.0 | 40.5 |  | 7.88 | 6 | Water 3 | | A:4571 | 2.97 | 1.0 | 56.6 |  | 1.58 | 7 | Water 4 | | A:2724 | 3.18 | 1.0 | 45.4 | 2.69 [G:N7]A:886 | 4.27 | 8 | G | O6 | A:887 | 3.72 | 1.0 | 42.6 |  | 6.13 | 9 | Water 5 | | A:2933 | 4.04 | 1.0 | 49.7 | 2.73 [G:N7]A:885 | 1.93 | 10 | G | N7 | A:886 | 4.26 | 1.0 | 33.5 |  | 8.14 | 11 | Water 6 | | A:2501 | 4.35 | 1.0 | 58.8 |  | 1.95 | 12 | G | N7 | A:885 | 4.74 | 1.0 | 35.2 |  | 7.88 |

  
**1^st^ shell coordination angles and distances (*trans* in "blue"; *cys* in "green"; deviations in "orange"):**   
• 1-2(W/W): 98; 1-3(W/O6): 162; 1-4(W/O4): 100; 1-5(W/O6): 86; 1-6(W/W): 63; 1-7(W/W): 100; 2-3(W/O6): 98; 2-4(W/O4): 114; 2-5(W/O6): 141; 2-6(W/W): 48; 2-7(W/W): 60; 3-4(O6/O4): 79; 3-5(O6/O6): 77; 3-6(O6/W): 134; 3-7(O6/W): 82; 4-5(O4/O6): 103; 4-6(O4/W): 87; 4-7(O4/W): 160; 5-6(O6/W): 149; 5-7(O6/W): 81; 6-7(W/W): 99;   
• 1-2(W/W): 3.9; 1-3(W/O6): 5.2; 1-4(W/O4): 4.2; 1-5(W/O6): 3.8; 1-6(W/W): 2.9; 1-7(W/W): 4.4; 2-3(W/O6): 4.1; 2-4(W/O4): 4.6; 2-5(W/O6): 5.3; 2-6(W/W): 2.3; 2-7(W/W): 3.0; 3-4(O6/O4): 3.6; 3-5(O6/O6): 3.6; 3-6(O6/W): 5.3; 3-7(O6/W): 3.9; 4-5(O4/O6): 4.6; 4-6(O4/W): 4.1; 4-7(O4/W): 6.0; 5-6(O6/W): 5.7; 5-7(O6/W): 4.0; 6-7(W/W): 4.7;   
  
**3.95 ± 1.21: the average coodination distance of the 1^st^ six ligands could correspond to an hexahydrated Mg^2+^.**  
**8: Number of potential Water contacts**   
**2 : Number of second shell contacts involving water or ions**   
**25%: Percentage of satisfied second shell Water contacts**   
• Contact(s): /O4/O6/O6/O_w_/O_w_/O_w_/  
• Contact code: 3O_b_3O_w_  
• Contact(s) in the exclusion zone: /O_w_/  
Go to ToC

### K:A:1739

|  | Res. type | Atom | Residue | Distance | Occupancy | B-factor | Closest atom(s) to a 1^st^ shell water | EM\_dens. |
| --- | --- | --- | --- | --- | --- | --- | --- | --- |
| **Ion** | | | |  | 1.0 | 30.0 |  | 8.20 |
 1 | U | O4 | A:1485 | 2.73 | 1.0 | 41.4 |  | 8.28 | 2 | G | O6 | A:1415 | 2.78 | 1.0 | 44.6 |  | 7.37 | 3 | U | O4 | A:1414 | 2.89 | 1.0 | 51.8 |  | 3.40 | 4 | G | O6 | A:1486 | 3.13 | 1.0 | 45.4 |  | 4.99 | 5 | Water 1 | | A:3661 | 3.33 | 1.0 | 52.1 | 2.88 [G:N7]A:1415 | 2.96 | 6 | Water 2 | | A:3717 | 3.75 | 1.0 | 48.2 | 2.89 [G:O6]A:1416 | 3.32 | 7 | U | N3 | A:1485 | 3.79 | 1.0 | 33.2 |  | 8.69 | 8 | U | N3 | A:1414 | 3.92 | 1.0 | 45.5 |  | 6.55 | 9 | G | N7 | A:1415 | 4.43 | 1.0 | 40.5 |  | 6.31 | 10 | G | N7 | A:1486 | 4.60 | 1.0 | 43.3 |  | 6.16 | 11 | Water 3 | | A:3887 | 4.61 | 1.0 | 55.6 |  | 2.12 | 12 | G | O6 | A:1416 | 4.76 | 1.0 | 37.5 |  | 9.02 |

  
**1^st^ shell coordination angles and distances (*trans* in "blue"; *cys* in "green"; deviations in "orange"):**   
• 1-2(O4/O6): 83; 1-3(O4/O4): 152; 1-4(O4/O6): 118; 2-3(O6/O4): 125; 2-4(O6/O6): 70; 3-4(O4/O6): 73;   
• 1-2(O4/O6): 3.6; 1-3(O4/O4): 5.5; 1-4(O4/O6): 5.0; 2-3(O6/O4): 5.0; 2-4(O6/O6): 3.4; 3-4(O4/O6): 3.6;   
  
**4.39 ± 1.37: the average coodination distance of the 1^st^ six ligands could correspond to an hexahydrated Mg^2+^.**  
**4: Number of potential Water contacts**   
**2 : Number of second shell contacts involving water or ions**   
**50%: Percentage of satisfied second shell Water contacts**   
• Contact(s): /O4/O4/O6/O6/  
• Contact code: 4O_b_  
→ Coordination < 6 for K ←  
Go to ToC

### K:A:1740

|  | Res. type | Atom | Residue | Distance | Occupancy | B-factor | Closest atom(s) to a 1^st^ shell water | EM\_dens. |
| --- | --- | --- | --- | --- | --- | --- | --- | --- |
| **Ion** | | | |  | 1.0 | 30.0 |  | 9.09 |
 1 | Water 1 | | A:2717 | 2.84 | 1.0 | 46.0 | 2.69 [G:N7]A:1419 | 2.50 | 2 | U | O4 | A:1481 | 2.92 | 1.0 | 44.3 |  | 7.89 | 3 | G | O6 | A:1419 | 2.97 | 1.0 | 42.2 |  | 6.37 | 4 | G | O6 | A:1482 | 2.98 | 1.0 | 42.2 |  | 8.34 | 5 | Water 2 | | A:4197 | 3.06 | 1.0 | 49.2 |  | 2.73 | 6 | A | N6 | A:1480 | 3.70 | 1.0 | 37.3 |  | 7.56 | 7 | Water 3 | | A:3978 | 3.93 | 1.0 | 46.9 | 2.96 [G:N1]A:1482 | 3.39 | 8 | U | N3 | A:1481 | 3.97 | 1.0 | 39.2 |  | 7.77 | 9 | Water 4 | | A:1821 | 4.05 | 1.0 | 50.1 |  | 2.21 | 10 | Water 5 | | A:3921 | 4.10 | 1.0 | 52.8 |  | 3.32 | 11 | Water 6 | | A:2742 | 4.38 | 1.0 | 39.0 |  | 7.04 | 12 | U | O4 | A:1420 | 4.50 | 1.0 | 47.5 |  | 4.10 | 13 | G | N7 | A:1419 | 4.54 | 1.0 | 43.1 |  | 5.96 | 14 | Water 7 | | A:2353 | 4.77 | 1.0 | 54.1 |  | 3.61 |

  
**1^st^ shell coordination angles and distances (*trans* in "blue"; *cys* in "green"; deviations in "orange"):**   
• 1-2(W/O4): 147; 1-3(W/O6): 71; 1-4(W/O6): 117; 1-5(W/W): 108; 2-3(O4/O6): 81; 2-4(O4/O6): 67; 2-5(O4/W): 70; 3-4(O6/O6): 73; 3-5(O6/W): 117; 4-5(O6/W): 134;   
• 1-2(W/O4): 5.5; 1-3(W/O6): 3.4; 1-4(W/O6): 5.0; 1-5(W/W): 4.8; 2-3(O4/O6): 3.8; 2-4(O4/O6): 3.3; 2-5(O4/W): 3.4; 3-4(O6/O6): 3.5; 3-5(O6/W): 5.1; 4-5(O6/W): 5.5;   
  
**4.40 ± 1.37: the average coodination distance of the 1^st^ six ligands could correspond to an hexahydrated Mg^2+^.**  
**6: Number of potential Water contacts**   
**2 : Number of second shell contacts involving water or ions**   
**33%: Percentage of satisfied second shell Water contacts**   
• Contact(s): /O4/O6/O6/O_w_/O_w_/  
• Contact code: 3O_b_2O_w_  
→ Coordination < 6 for K ←  
Go to ToC

### MG:A:1741

|  | Res. type | Atom | Residue | Distance | Occupancy | B-factor | Closest atom(s) to a 1^st^ shell water | EM\_dens. |
| --- | --- | --- | --- | --- | --- | --- | --- | --- |
| **Ion** | | | |  | 1.0 | 66.4 |  | 2.50 |
 1 | Water 1 | | Q:108 | 2.00 | 1.0 | 69.2 |  | 2.09 | 2 | Water 2 | | A:3131 | 2.04 | 1.0 | 58.3 | 2.77 [C:OP1]A:234 | 2.94 | 3 | Water 3 | | A:4359 | 2.22 | 1.0 | 63.0 |  | 2.85 | 4 | Water 4 | | Q:109 | 2.89 | 1.0 | 64.7 |  | 2.00 | 5 | C | OP1 | A:267 | 3.80 | 1.0 | 66.7 |  | 3.46 | 6 | G | N3 | A:265 | 3.80 | 1.0 | 63.9 |  | 3.18 | 7 | G | O2' | A:265 | 3.85 | 1.0 | 66.8 |  | 2.82 | 8 | Water 5 | | A:2849 | 4.37 | 1.0 | 59.9 |  | 3.10 | 9 | LYS | O | Q:69 | 4.43 | 1.0 | 63.5 |  | 3.15 | 10 | SER | O | Q:68 | 4.54 | 1.0 | 64.4 |  | 3.97 | 11 | C | OP2 | A:267 | 4.64 | 1.0 | 66.2 |  | 3.20 | 12 | C | OP1 | A:234 | 4.69 | 1.0 | 58.0 |  | 3.56 |

  
**1^st^ shell coordination angles and distances (*trans* in "blue"; *cys* in "green"; deviations in "orange"):**   
• 1-2(W/W): *71º*; 1-3(W/W): *136º*; 2-3(W/W): *83º*;   
• 1-2(W/W): 2.3Å; 1-3(W/W): 3.9Å; 2-3(W/W): 2.8Å;   
  
**2.79 ± 0.77: the average coodination distance of the 1^st^ six ligands is K^+^ compatible**  
**8: Number of potential Water contacts**   
**1 : Number of second shell contacts involving water or ions**   
**12%: Percentage of satisfied second shell Water contacts**   
• Contact(s): /O_w_/O_w_/O_w_/  
• Contact code: 3O_w_  
• Contact(s) in the exclusion zone: /O_w_/  
→ Missing octahedral coordination for MG ←  
→2Fo-Fc density between 1.0 and 4.0 !   
Go to ToC

### MG:A:1742

|  | Res. type | Atom | Residue | Distance | Occupancy | B-factor | Closest atom(s) to a 1^st^ shell water | EM\_dens. |
| --- | --- | --- | --- | --- | --- | --- | --- | --- |
| **Ion** | | | |  | 1.0 | 64.8 |  | 3.73 |
 1 | Water 1 | | A:2107 | 1.95 | 1.0 | 55.0 | 2.52 [A:OP2]A:1434 | 3.48 | 2 | A | OP2 | A:1433 | 2.00 | 1.0 | 54.4 |  | 2.91 | 3 | Water 2 | | A:2655 | 2.08 | 1.0 | 56.4 | 2.89 [A:N7]A:1434 | 2.23 | 4 | Water 3 | | o:261 | 2.08 | 1.0 | 58.8 |  | 2.83 | 5 | Water 4 | | A:2252 | 2.16 | 1.0 | 57.9 | 2.99 [G:O2']A:1432 | 2.61 | 6 | Water 5 | | A:2282 | 2.43 | 1.0 | 58.4 | 2.59 [G:O6]A:1435 | 2.87 | 7 | A | O5' | A:1433 | 3.91 | 1.0 | 52.1 |  | 3.12 | 8 | Water 6 | | A:4519 | 4.02 | 1.0 | 61.5 |  | 1.86 | 9 | Water 7 | | A:2503 | 4.12 | 1.0 | 56.6 |  | 2.72 | 10 | A | OP2 | A:1434 | 4.15 | 1.0 | 54.9 |  | 3.30 | 11 | Water 8 | | A:4389 | 4.24 | 1.0 | 60.4 |  | 1.60 | 12 | A | OP1 | A:1433 | 4.30 | 1.0 | 51.4 |  | 2.69 | 13 | G | O2' | A:1432 | 4.33 | 1.0 | 48.4 |  | 3.96 | 14 | A | N7 | A:1434 | 4.35 | 1.0 | 53.5 |  | 3.71 | 15 | G | O6 | A:1435 | 4.42 | 1.0 | 52.1 |  | 5.36 | 16 | G | O3' | A:1432 | 4.42 | 1.0 | 49.3 |  | 4.74 | 17 | Water 9 | | A:2586 | 4.67 | 1.0 | 49.5 | 2.98 [G:O2']A:1432 | 3.86 |

  
**1^st^ shell coordination angles and distances (*trans* in "blue"; *cys* in "green"; deviations in "orange"):**   
• 1-2(W/OP2): *101º*; 1-3(W/W): 88º; 1-4(W/W): 95º; 1-5(W/W): 171º; 2-3(OP2/W): *82º*; 2-4(OP2/W): *100º*; 2-5(OP2/W): *76º*; 3-4(W/W): 176º; 3-5(W/W): *84º*; 4-5(W/W): 94º;   
• 1-2(W/OP2): 3.0Å; 1-3(W/W): 2.8Å; 1-4(W/W): 3.0Å; 1-5(W/W): 4.1Å; 2-3(OP2/W): 2.7Å; 2-4(OP2/W): 3.1Å; 2-5(OP2/W): 2.6Å; 3-4(W/W): 4.2Å; 3-5(W/W): 2.8Å; 4-5(W/W): 3.1Å;   
  
**2.12 ± 0.16: the average coodination distance of the 1^st^ six ligands is Mg^2+^ compatible**  
**10: Number of potential Water contacts**   
**4 : Number of second shell contacts involving water or ions**   
**40%: Percentage of satisfied second shell Water contacts**   
• Contact(s): /O_w_/O_w_/O_w_/O_w_/OP2/  
• Contact code: O_ph_4O_w_  
• Contact(s) in the exclusion zone: /O_w_/  
→ Missing octahedral coordination for MG ←  
→ Slightly distorded coord. octahedron (7.3º) ←→2Fo-Fc density between 1.0 and 4.0 !   
Go to ToC

### MG:A:1743

|  | Res. type | Atom | Residue | Distance | Occupancy | B-factor | Closest atom(s) to a 1^st^ shell water | EM\_dens. |
| --- | --- | --- | --- | --- | --- | --- | --- | --- |
| **Ion** | | | |  | 1.0 | 82.6 |  | 2.51 |
 1 | Water 1 | | A:4119 | 1.91 | 1.0 | 78.9 |  | 2.05 | 2 | PSU | O2 | A:516 | 1.92 | 1.0 | 76.8 |  | 2.04 | 3 | Water 2 | | A:1801 | 1.94 | 1.0 | 74.4 |  | 2.05 | 4 | Water 3 | | A:1811 | 1.99 | 1.0 | 77.0 | 2.35 [G:O6]A:517 | 1.91 | 5 | A | OP1 | A:533 | 2.03 | 1.0 | 72.8 |  | 2.80 | 6 | Water 4 | | A:3185 | 2.05 | 1.0 | 78.2 | 2.78 [G:O6]A:515 | 1.59 | 7 | A | O5' | A:533 | 3.59 | 1.0 | 74.5 |  | 1.96 | 8 | G | O6 | A:515 | 3.76 | 1.0 | 72.2 |  | 2.59 | 9 | G | O6 | A:517 | 3.91 | 1.0 | 77.0 |  | 2.31 | 10 | G | N7 | A:515 | 4.10 | 1.0 | 74.4 |  | 2.86 | 11 | A | O3' | A:532 | 4.32 | 1.0 | 83.3 |  | 1.72 | 12 | A | OP2 | A:533 | 4.37 | 1.0 | 74.7 |  | 2.52 | 13 | A | N7 | A:533 | 4.52 | 1.0 | 70.7 |  | 3.22 |

  
**1^st^ shell coordination angles and distances (*trans* in "blue"; *cys* in "green"; deviations in "orange"):**   
• 1-2(W/O2): *104º*; 1-3(W/W): *72º*; 1-4(W/W): *75º*; 1-5(W/OP1): *121º*; 1-6(W/W): *83º*; 2-3(O2/W): 171º; 2-4(O2/W): *77º*; 2-5(O2/OP1): *111º*; 2-6(O2/W): *110º*; 3-4(W/W): 95º; 3-5(W/OP1): *67º*; 3-6(W/W): *77º*; 4-5(W/OP1): *69º*; 4-6(W/W): *158º*; 5-6(OP1/W): *124º*;   
• 1-2(W/O2): 3.0Å; 1-3(W/W): 2.3Å; 1-4(W/W): 2.4Å; 1-5(W/OP1): *3.4Å*; 1-6(W/W): 2.6Å; 2-3(O2/W): 3.9Å; 2-4(O2/W): 2.4Å; 2-5(O2/OP1): *3.3Å*; 2-6(O2/W): *3.3Å*; 3-4(W/W): 2.9Å; 3-5(W/OP1): 2.2Å; 3-6(W/W): 2.5Å; 4-5(W/OP1): 2.3Å; 4-6(W/W): 4.0Å; 5-6(OP1/W): *3.6Å*;   
  
**1.97 ± 0.05: the average coodination distance of the 1^st^ six ligands is Mg^2+^ compatible**  
**8: Number of potential Water contacts**   
**2 : Number of second shell contacts involving water or ions**   
**25%: Percentage of satisfied second shell Water contacts**   
• Contact(s): /O2/O_w_/O_w_/O_w_/O_w_/OP1/  
• Contact code: O_ph_O_b_4O_w_  
→→→ Distorded coord. octahedron (17.6º) ←←←  
• MG hydration shell issue: water too close (< 2.4 Å.)  
→ MG ion B\_factor > 80 ←→2Fo-Fc density between 1.0 and 4.0 !   
Go to ToC

### MG:A:1744

|  | Res. type | Atom | Residue | Distance | Occupancy | B-factor | Closest atom(s) to a 1^st^ shell water | EM\_dens. |
| --- | --- | --- | --- | --- | --- | --- | --- | --- |
| **Ion** | | | |  | 1.0 | 72.5 |  | 2.35 |
 1 | Water 1 | | A:2806 | 2.37 | 1.0 | 65.5 | 2.71 [G:O6]A:259 | 1.89 | 2 | Water 2 | | A:4175 | 2.84 | 1.0 | 70.7 |  | 2.01 | 3 | Water 3 | | A:3202 | 2.98 | 1.0 | 71.3 | 2.78 [G:N7]A:258 | 1.76 | 4 | U | O4 | A:268 | 3.08 | 1.0 | 67.3 |  | 2.56 | 5 | G | O6 | A:258 | 3.10 | 1.0 | 70.7 |  | 2.58 | 6 | G | O6 | A:257 | 3.27 | 1.0 | 72.0 |  | 2.75 | 7 | **K** | K | A:1602 | 3.48 | 1.0 | 30.0 |  | 4.24 | 8 | G | O6 | A:259 | 3.50 | 1.0 | 67.6 |  | 3.28 | 9 | G | OP2 | A:266 | 3.53 | 1.0 | 69.1 |  | 2.31 | 10 | Water 4 | | A:2192 | 3.90 | 1.0 | 72.2 | 2.55 [U:O4]A:268 // 2.93 [C:N4]A:269 | 1.78 | 11 | G | N7 | A:258 | 4.42 | 1.0 | 69.3 |  | 2.97 | 12 | G | OP1 | A:266 | 4.76 | 1.0 | 68.7 |  | 2.61 |

  
• • **2.94 ± 0.28: the average coodination distance of the 1^st^ six ligands is K^+^ compatible**  
**6: Number of potential Water contacts**   
**2 : Number of second shell contacts involving water or ions**   
**33%: Percentage of satisfied second shell Water contacts**   
• Contact code: → No contacts ←   
• Contact(s) in the exclusion zone: /N4/O4/O6/O6/O_w_/O_w_/O_w_/  
→ Consider K in place of MG (2 valid contact(s) for K vs. 0 for MG ←)  
→2Fo-Fc density between 1.0 and 4.0 !   
Go to ToC

### MG:A:1745

|  | Res. type | Atom | Residue | Distance | Occupancy | B-factor | Closest atom(s) to a 1^st^ shell water | EM\_dens. |
| --- | --- | --- | --- | --- | --- | --- | --- | --- |
| **Ion** | | | |  | 1.0 | 61.7 |  | 2.86 |
 1 | Water 1 | | A:4558 | 1.91 | 1.0 | 54.6 |  | 2.14 | 2 | Water 2 | | E:231 | 2.05 | 1.0 | 55.0 |  | 1.70 | 3 | Water 3 | | A:4308 | 2.05 | 1.0 | 49.9 |  | 2.81 | 4 | Water 4 | | A:4676 | 2.11 | 1.0 | 56.8 |  | 2.34 | 5 | Water 5 | | A:4542 | 2.12 | 1.0 | 51.9 |  | 2.12 | 6 | Water 6 | | A:3459 | 2.25 | 1.0 | 50.4 | 2.84 [G:N7]A:21 | 1.52 | 7 | Water 7 | | A:4546 | 3.81 | 1.0 | 52.7 |  | 2.37 | 8 | Water 8 | | A:3256 | 4.10 | 1.0 | 50.8 | 2.79 [G:OP2]A:21 | 2.56 | 9 | Water 9 | | A:2029 | 4.16 | 1.0 | 51.0 |  | 3.89 | 10 | GLY | O | E:129 | 4.25 | 1.0 | 49.1 |  | 4.55 | 11 | G | N7 | A:21 | 4.29 | 1.0 | 34.9 |  | 7.63 | 12 | Water 10 | | A:2094 | 4.35 | 1.0 | 48.0 | 2.52 [U:OP2]A:20 | 2.40 | 13 | U | OP2 | A:20 | 4.54 | 1.0 | 35.5 |  | 8.36 | 14 | G | OP2 | A:21 | 4.56 | 1.0 | 41.9 |  | 7.00 | 15 | U | OP2 | A:561 | 4.66 | 1.0 | 53.2 |  | 3.06 |

  
**1^st^ shell coordination angles and distances (*trans* in "blue"; *cys* in "green"; deviations in "orange"):**   
• 1-2(W/W): *99º*; 1-3(W/W): *71º*; 1-4(W/W): *106º*; 1-5(W/W): *168º*; 1-6(W/W): *123º*; 2-3(W/W): 90º; 2-4(W/W): *79º*; 2-5(W/W): *73º*; 2-6(W/W): *126º*; 3-4(W/W): *169º*; 3-5(W/W): *101º*; 3-6(W/W): *75º*; 4-5(W/W): *81º*; 4-6(W/W): *115º*; 5-6(W/W): *60º*;   
• 1-2(W/W): 3.0Å; 1-3(W/W): 2.3Å; 1-4(W/W): *3.2Å*; 1-5(W/W): 4.0Å; 1-6(W/W): *3.7Å*; 2-3(W/W): 2.9Å; 2-4(W/W): 2.7Å; 2-5(W/W): 2.5Å; 2-6(W/W): 3.8Å; 3-4(W/W): 4.1Å; 3-5(W/W): *3.2Å*; 3-6(W/W): 2.6Å; 4-5(W/W): 2.7Å; 4-6(W/W): *3.7Å*; 5-6(W/W): 2.2Å;   
  
**2.08 ± 0.10: the average coodination distance of the 1^st^ six ligands is Mg^2+^ compatible**  
**12: Number of potential Water contacts**   
**1 : Number of second shell contacts involving water or ions**   
**8%: Percentage of satisfied second shell Water contacts**   
• Contact(s): /O_w_/O_w_/O_w_/O_w_/O_w_/O_w_/  
• Contact code: 6O_w_  
→→→ Distorded coord. octahedron (16.9º) ←←←  
→2Fo-Fc density between 1.0 and 4.0 !   
Go to ToC

### MG:X:101

|  | Res. type | Atom | Residue | Distance | Occupancy | B-factor | Closest atom(s) to a 1^st^ shell water | EM\_dens. |
| --- | --- | --- | --- | --- | --- | --- | --- | --- |
| **Ion** | | | |  | 1.0 | 59.5 |  | 4.52 |
 1 | Water 1 | | A:2444 | 2.16 | 1.0 | 49.3 | 2.63 [G:N7]A:1401 // 2.68 [G:OP2]A:1401 | 3.90 | 2 | Water 2 | | X:209 | 2.19 | 1.0 | 49.9 | 2.79 [U:OP2]X:17 | 3.30 | 3 | Water 3 | | X:201 | 2.26 | 1.0 | 53.6 | 2.46 [A:N7]X:18 | 3.11 | 4 | Water 4 | | A:3030 | 2.33 | 1.0 | 46.1 | 2.75 [G:O6]A:1401 | 5.21 | 5 | Water 5 | | A:3867 | 3.04 | 1.0 | 45.4 |  | 3.03 | 6 | Water 6 | | X:216 | 3.82 | 1.0 | 53.3 |  | 2.42 | 7 | A | OP2 | X:18 | 3.84 | 1.0 | 43.8 |  | 7.17 | 8 | G | N7 | A:1401 | 4.12 | 1.0 | 38.0 |  | 7.30 | 9 | U | OP2 | X:17 | 4.23 | 1.0 | 44.3 |  | 5.77 | 10 | Water 7 | | X:207 | 4.29 | 1.0 | 44.5 |  | 6.56 | 11 | A | N7 | X:18 | 4.29 | 1.0 | 40.8 |  | 5.55 | 12 | Water 8 | | A:2747 | 4.33 | 1.0 | 49.6 |  | 2.87 | 13 | Water 9 | | A:2594 | 4.42 | 1.0 | 46.6 | 2.66 [G:OP2]A:1401 // 2.80 [A:OP2]X:18 | 3.75 | 14 | U | O5' | X:17 | 4.44 | 1.0 | 46.5 |  | 8.06 | 15 | G | OP2 | A:1401 | 4.49 | 1.0 | 46.6 |  | 5.05 | 16 | Water 10 | | A:4610 | 4.57 | 1.0 | 52.9 |  | 2.30 | 17 | G | O6 | A:1401 | 4.59 | 1.0 | 40.3 |  | 6.78 |

  
**1^st^ shell coordination angles and distances (*trans* in "blue"; *cys* in "green"; deviations in "orange"):**   
• 1-2(W/W): *144º*; 1-3(W/W): *77º*; 2-3(W/W): *81º*;   
• 1-2(W/W): 4.1Å; 1-3(W/W): 2.8Å; 2-3(W/W): 2.9Å;   
  
**2.63 ± 0.61: the average coodination distance of the 1^st^ six ligands is K^+^ compatible**  
**12: Number of potential Water contacts**   
**5 : Number of second shell contacts involving water or ions**   
**42%: Percentage of satisfied second shell Water contacts**   
• Contact(s): /O_w_/O_w_/O_w_/  
• Contact code: 3O_w_  
• Contact(s) in the exclusion zone: /O_w_/O_w_/  
→ Missing octahedral coordination for MG ←  
Go to ToC

### MG:a:3001

|  | Res. type | Atom | Residue | Distance | Occupancy | B-factor | Closest atom(s) to a 1^st^ shell water | EM\_dens. |
| --- | --- | --- | --- | --- | --- | --- | --- | --- |
| **Ion** | | | |  | 1.0 | 41.9 |  | 12.42 |
 1 | Water 1 | | a:7742 | 2.08 | 1.0 | 32.9 |  | 9.20 | 2 | Water 2 | | a:5632 | 2.08 | 1.0 | 38.4 | 2.72 [G:N7]a:1281 | 6.82 | 3 | Water 3 | | a:3920 | 2.12 | 1.0 | 38.2 | 2.56 [G:O6]a:1281 | 6.26 | 4 | Water 4 | | a:10175 | 2.15 | 1.0 | 35.2 |  | 6.86 | 5 | Water 5 | | a:10031 | 2.26 | 1.0 | 42.2 |  | 6.98 | 6 | Water 6 | | a:3748 | 3.79 | 1.0 | 39.8 |  | 5.82 | 7 | Water 7 | | a:10119 | 3.96 | 1.0 | 39.3 |  | 5.57 | 8 | Water 8 | | a:4796 | 4.12 | 1.0 | 41.8 |  | 5.17 | 9 | Water 9 | | a:8173 | 4.13 | 1.0 | 43.5 |  | 3.53 | 10 | G | OP1 | a:1290 | 4.16 | 1.0 | 33.0 |  | 9.82 | 11 | G | O6 | a:1282 | 4.20 | 1.0 | 36.0 |  | 8.69 | 12 | G | N7 | a:1281 | 4.21 | 1.0 | 25.8 |  | 11.30 | 13 | Water 10 | | a:8353 | 4.25 | 1.0 | 34.9 |  | 7.26 | 14 | G | O6 | a:1281 | 4.26 | 1.0 | 30.8 |  | 11.21 | 15 | Water 11 | | a:3721 | 4.57 | 1.0 | 40.4 | 2.52 [G:OP1]a:1290 | 5.85 | 16 | Water 12 | | a:8968 | 4.61 | 1.0 | 37.4 |  | 7.99 |

  
**1^st^ shell coordination angles and distances (*trans* in "blue"; *cys* in "green"; deviations in "orange"):**   
• 1-2(W/W): *80º*; 1-3(W/W): 87º; 1-4(W/W): 89º; 1-5(W/W): *162º*; 2-3(W/W): 87º; 2-4(W/W): 86º; 2-5(W/W): *82º*; 3-4(W/W): 173º; 3-5(W/W): *97º*; 4-5(W/W): *85º*;   
• 1-2(W/W): 2.7Å; 1-3(W/W): 2.9Å; 1-4(W/W): 3.0Å; 1-5(W/W): *4.3Å*; 2-3(W/W): 2.9Å; 2-4(W/W): 2.9Å; 2-5(W/W): 2.9Å; 3-4(W/W): *4.3Å*; 3-5(W/W): *3.3Å*; 4-5(W/W): 3.0Å;   
  
**2.41 ± 0.62: the average coodination distance of the 1^st^ six ligands is Na^+^** compatible  
**12: Number of potential Water contacts**   
**2 : Number of second shell contacts involving water or ions**   
**17%: Percentage of satisfied second shell Water contacts**   
• Contact(s): /O_w_/O_w_/O_w_/O_w_/O_w_/  
• Contact code: 5O_w_  
→ Missing octahedral coordination for MG ←  
→ Slightly distorded coord. octahedron (6.6º) ←  
Go to ToC

### MG:a:3002

|  | Res. type | Atom | Residue | Distance | Occupancy | B-factor | Closest atom(s) to a 1^st^ shell water | EM\_dens. |
| --- | --- | --- | --- | --- | --- | --- | --- | --- |
| **Ion** | | | |  | 1.0 | 49.0 |  | 7.56 |
 1 | Water 1 | | a:10264 | 2.03 | 1.0 | 48.0 |  | 5.00 | 2 | Water 2 | | a:5210 | 2.39 | 1.0 | 35.9 | 2.69 [G:O6]a:1294 | 8.86 | 3 | Water 3 | | a:9541 | 2.46 | 1.0 | 39.3 |  | 8.55 | 4 | Water 4 | | a:8591 | 2.46 | 1.0 | 34.2 |  | 8.57 | 5 | Water 5 | | a:5891 | 3.64 | 1.0 | 38.7 |  | 5.52 | 6 | Water 6 | | a:8857 | 4.06 | 1.0 | 38.4 |  | 8.11 | 7 | Water 7 | | a:9094 | 4.20 | 1.0 | 41.0 |  | 5.75 | 8 | Water 8 | | a:5115 | 4.29 | 1.0 | 39.4 |  | 8.11 | 9 | Water 9 | | a:3768 | 4.30 | 1.0 | 35.4 |  | 8.99 | 10 | Water 10 | | a:3574 | 4.42 | 1.0 | 44.4 |  | 4.80 | 11 | G | O6 | a:1294 | 4.75 | 1.0 | 29.2 |  | 11.32 |

  
• • **2.84 ± 0.74: the average coodination distance of the 1^st^ six ligands is K^+^ compatible**  
**12: Number of potential Water contacts**   
**1 : Number of second shell contacts involving water or ions**   
**8%: Percentage of satisfied second shell Water contacts**   
• Contact(s): /O_w_/  
• Contact code: O_w_  
• Contact(s) in the exclusion zone: /O_w_/O_w_/O_w_/  
→ Missing octahedral coordination for MG ←  
Go to ToC

### MG:a:3003

|  | Res. type | Atom | Residue | Distance | Occupancy | B-factor | Closest atom(s) to a 1^st^ shell water | EM\_dens. |
| --- | --- | --- | --- | --- | --- | --- | --- | --- |
| **Ion** | | | |  | 1.0 | 32.9 |  | 17.01 |
 1 | Water 1 | | a:8412 | 1.97 | 1.0 | 26.7 | 2.92 [G:N3]a:2713 | 11.38 | 2 | C | OP1 | a:1641 | 2.00 | 1.0 | 27.3 |  | 11.52 | 3 | Water 2 | | a:4058 | 2.09 | 1.0 | 31.4 | 2.58 [C:OP2]a:1299 // 2.94 [G:O2']a:2713 | 13.25 | 4 | Water 3 | | a:9093 | 2.18 | 1.0 | 31.1 |  | 10.61 | 5 | Water 4 | | a:8890 | 2.30 | 1.0 | 34.2 |  | 10.10 | 6 | Water 5 | | a:6388 | 2.33 | 1.0 | 34.0 | 2.77 [G:OP2]a:1298 | 10.66 | 7 | Water 6 | | a:9518 | 3.78 | 1.0 | 42.3 |  | 4.15 | 8 | C | O5' | a:1641 | 4.00 | 1.0 | 27.0 |  | 14.44 | 9 | C | OP2 | a:1641 | 4.06 | 1.0 | 28.3 |  | 11.83 | 10 | Water 7 | | a:3709 | 4.08 | 1.0 | 39.8 |  | 7.41 | 11 | G | O2' | a:2713 | 4.14 | 1.0 | 28.0 |  | 13.45 | 12 | C | OP2 | a:1299 | 4.27 | 1.0 | 30.0 |  | 12.41 | 13 | G | N3 | a:2713 | 4.37 | 1.0 | 21.4 |  | 15.28 | 14 | C | O3' | a:1640 | 4.46 | 1.0 | 29.0 |  | 12.80 | 15 | Water 8 | | a:10188 | 4.50 | 1.0 | 41.5 |  | 4.84 | 16 | Water 9 | | a:9709 | 4.52 | 1.0 | 42.4 |  | 5.22 | 17 | Water 10 | | a:9651 | 4.52 | 1.0 | 37.7 |  | 6.92 | 18 | Water 11 | | a:8815 | 4.69 | 1.0 | 31.5 | 2.97 [C:OP2]a:1299 | 8.43 | 19 | G | OP2 | a:1298 | 4.74 | 1.0 | 29.2 |  | 11.07 |

  
**1^st^ shell coordination angles and distances (*trans* in "blue"; *cys* in "green"; deviations in "orange"):**   
• 1-2(W/OP1): *97º*; 1-3(W/W): *102º*; 1-4(W/W): 85º; 2-3(OP1/W): *97º*; 2-4(OP1/W): 92º; 3-4(W/W): *168º*;   
• 1-2(W/OP1): 3.0Å; 1-3(W/W): 3.2Å; 1-4(W/W): 2.8Å; 2-3(OP1/W): 3.1Å; 2-4(OP1/W): 3.0Å; 3-4(W/W): *4.2Å*;   
  
**2.15 ± 0.14: the average coodination distance of the 1^st^ six ligands is Mg^2+^ compatible**  
**10: Number of potential Water contacts**   
**4 : Number of second shell contacts involving water or ions**   
**40%: Percentage of satisfied second shell Water contacts**   
• Contact(s): /O_w_/O_w_/O_w_/OP1/  
• Contact code: O_ph_3O_w_  
• Contact(s) in the exclusion zone: /O_w_/O_w_/  
→ Missing octahedral coordination for MG ←  
→ Slightly distorded coord. octahedron (7.5º) ←  
Go to ToC

### MG:a:3004

|  | Res. type | Atom | Residue | Distance | Occupancy | B-factor | Closest atom(s) to a 1^st^ shell water | EM\_dens. |
| --- | --- | --- | --- | --- | --- | --- | --- | --- |
| **Ion** | | | |  | 1.0 | 38.0 |  | 12.06 |
 1 | Water 1 | | a:5026 | 2.10 | 1.0 | 28.6 | 2.67 [G:O6]a:1301 | 13.12 | 2 | Water 2 | | a:9490 | 2.18 | 1.0 | 37.3 |  | 6.57 | 3 | C | OP2 | a:1641 | 2.27 | 1.0 | 28.3 |  | 11.83 | 4 | Water 3 | | a:8815 | 2.45 | 1.0 | 31.5 | 2.97 [C:OP2]a:1299 | 8.43 | 5 | C | OP2 | a:1300 | 2.47 | 1.0 | 34.1 |  | 13.25 | 6 | Water 4 | | a:6295 | 3.67 | 1.0 | 47.5 | 2.90 [C:O5']a:1300 | 5.78 | 7 | Water 5 | | a:4160 | 4.06 | 1.0 | 44.7 | 2.92 [C:O2']a:2714 | 5.79 | 8 | C | O5' | a:1640 | 4.23 | 1.0 | 23.6 |  | 12.71 | 9 | C | O3' | a:1640 | 4.25 | 1.0 | 29.0 |  | 12.80 | 10 | C | OP1 | a:1641 | 4.46 | 1.0 | 27.3 |  | 11.52 | 11 | Water 6 | | a:8890 | 4.47 | 1.0 | 34.2 | 2.98 [C:OP1]a:1641 | 10.10 | 12 | C | OP2 | a:1299 | 4.48 | 1.0 | 30.0 |  | 12.41 | 13 | C | OP1 | a:1300 | 4.54 | 1.0 | 37.9 |  | 10.20 | 14 | Water 7 | | a:5971 | 4.55 | 1.0 | 41.4 | 2.74 [C:OP1]a:1640 | 4.61 | 15 | C | OP2 | a:1640 | 4.59 | 1.0 | 30.2 |  | 12.28 | 16 | G | O6 | a:1301 | 4.59 | 1.0 | 27.0 |  | 14.91 | 17 | C | OP1 | a:1640 | 4.62 | 1.0 | 28.6 |  | 14.13 | 18 | C | O2' | a:2714 | 4.64 | 1.0 | 26.0 |  | 13.21 | 19 | C | O5' | a:1300 | 4.69 | 1.0 | 26.8 |  | 11.43 | 20 | C | O5' | a:1641 | 4.70 | 1.0 | 27.0 |  | 14.44 | 21 | Water 8 | | a:7860 | 4.72 | 1.0 | 34.5 | 2.87 [G:O6]a:1301 | 7.07 | 22 | C | O3' | a:1299 | 4.80 | 1.0 | 30.2 |  | 10.65 |

  
**1^st^ shell coordination angles and distances (*trans* in "blue"; *cys* in "green"; deviations in "orange"):**   
• 1-2(W/W): *76º*; 1-3(W/OP2): *81º*; 2-3(W/OP2): *95º*;   
• 1-2(W/W): 2.6Å; 1-3(W/OP2): 2.8Å; 2-3(W/OP2): *3.3Å*;   
  
**2.52 ± 0.53: the average coodination distance of the 1^st^ six ligands is Na^+^** compatible  
**8: Number of potential Water contacts**   
**3 : Number of second shell contacts involving water or ions**   
**38%: Percentage of satisfied second shell Water contacts**   
• Contact(s): /O_w_/O_w_/OP2/  
• Contact code: O_ph_2O_w_  
• Contact(s) in the exclusion zone: /O_w_/OP2/  
→ Missing octahedral coordination for MG ←  
Go to ToC

### MG:a:3005

|  | Res. type | Atom | Residue | Distance | Occupancy | B-factor | Closest atom(s) to a 1^st^ shell water | EM\_dens. |
| --- | --- | --- | --- | --- | --- | --- | --- | --- |
| **Ion** | | | |  | 1.0 | 25.6 |  | 19.64 |
 1 | G | O6 | a:1358 | 2.77 | 1.0 | 25.9 |  | 13.49 | 2 | Water 1 | | a:4863 | 2.82 | 1.0 | 45.2 | 2.66 [G:O6]a:1376 | 5.82 | 3 | Water 2 | | a:10237 | 2.85 | 1.0 | 53.0 |  | 3.46 | 4 | Water 3 | | a:6799 | 2.95 | 1.0 | 35.3 | 2.79 [G:N7]a:1358 | 8.83 | 5 | U | O4 | a:1377 | 3.00 | 1.0 | 31.8 |  | 12.60 | 6 | G | O6 | a:1357 | 3.13 | 1.0 | 27.2 |  | 14.03 | 7 | G | O6 | a:1376 | 3.17 | 1.0 | 30.7 |  | 10.91 | 8 | Water 4 | | a:4129 | 4.09 | 1.0 | 35.9 |  | 7.38 | 9 | G | N7 | a:1358 | 4.21 | 1.0 | 20.6 |  | 14.45 | 10 | Water 5 | | a:4880 | 4.41 | 1.0 | 42.4 | 2.66 [U:O4]a:1377 | 4.28 | 11 | C | N3 | a:1378 | 4.74 | 1.0 | 19.9 |  | 15.23 |

  
• • **2.92 ± 0.12: the average coodination distance of the 1^st^ six ligands is K^+^ compatible**  
**6: Number of potential Water contacts**   
**2 : Number of second shell contacts involving water or ions**   
**33%: Percentage of satisfied second shell Water contacts**   
• Contact code: → No contacts ←   
• Contact(s) in the exclusion zone: /O4/O6/O6/O6/O_w_/O_w_/O_w_/  
→ Consider K in place of MG (4 valid contact(s) for K vs. 0 for MG ←)  
Go to ToC

### MG:a:3006

|  | Res. type | Atom | Residue | Distance | Occupancy | B-factor | Closest atom(s) to a 1^st^ shell water | EM\_dens. |
| --- | --- | --- | --- | --- | --- | --- | --- | --- |
| **Ion** | | | |  | 1.0 | 54.6 |  | 6.84 |
 1 | Water 1 | | a:5716 | 1.88 | 1.0 | 35.9 | 2.72 [G:N7]a:1661 | 7.63 | 2 | Water 2 | | a:10061 | 2.22 | 1.0 | 41.7 |  | 4.37 | 3 | Water 3 | | a:4228 | 2.25 | 1.0 | 35.9 | 2.60 [G:N7]a:1662 | 7.89 | 4 | Water 4 | | a:4251 | 2.33 | 1.0 | 24.6 | 2.61 [G:O6]a:1661 // 2.91 [G:O6]a:1662 | 13.91 | 5 | Water 5 | | a:9755 | 2.89 | 1.0 | 36.3 |  | 5.37 | 6 | Water 6 | | a:7675 | 3.63 | 1.0 | 41.2 |  | 4.13 | 7 | Water 7 | | a:6317 | 4.27 | 1.0 | 36.6 |  | 6.58 | 8 | G | O6 | a:1662 | 4.30 | 1.0 | 26.5 |  | 14.53 | 9 | G | O6 | a:1661 | 4.31 | 1.0 | 27.4 |  | 15.01 | 10 | G | N7 | a:1661 | 4.34 | 1.0 | 17.9 |  | 14.79 | 11 | Water 8 | | a:6894 | 4.34 | 1.0 | 38.6 |  | 7.15 | 12 | Water 9 | | d:506 | 4.35 | 1.0 | 44.8 |  | 3.72 | 13 | G | N7 | a:1662 | 4.39 | 1.0 | 20.6 |  | 12.32 |

  
**1^st^ shell coordination angles and distances (*trans* in "blue"; *cys* in "green"; deviations in "orange"):**   
• 1-2(W/W): *165º*; 1-3(W/W): 87º; 2-3(W/W): 93º;   
• 1-2(W/W): 4.1Å; 1-3(W/W): 2.9Å; 2-3(W/W): *3.2Å*;   
  
**2.53 ± 0.57: the average coodination distance of the 1^st^ six ligands is Na^+^** compatible  
**12: Number of potential Water contacts**   
**4 : Number of second shell contacts involving water or ions**   
**33%: Percentage of satisfied second shell Water contacts**   
• Contact(s): /O_w_/O_w_/  
• Contact code: 2O_w_  
• Contact(s) in the exclusion zone: /O_w_/O_w_/O_w_/  
→ Missing octahedral coordination for MG ←  
Go to ToC

### MG:a:3007

|  | Res. type | Atom | Residue | Distance | Occupancy | B-factor | Closest atom(s) to a 1^st^ shell water | EM\_dens. |
| --- | --- | --- | --- | --- | --- | --- | --- | --- |
| **Ion** | | | |  | 1.0 | 40.1 |  | 12.79 |
 1 | Water 1 | | a:9140 | 2.03 | 1.0 | 32.6 |  | 9.23 | 2 | Water 2 | | a:9457 | 2.17 | 1.0 | 31.6 |  | 7.97 | 3 | U | OP2 | a:1664 | 2.19 | 1.0 | 31.0 |  | 13.00 | 4 | Water 3 | | a:10066 | 2.20 | 1.0 | 39.3 |  | 7.67 | 5 | Water 4 | | a:9428 | 2.29 | 1.0 | 33.7 |  | 10.21 | 6 | Water 5 | | a:5480 | 2.32 | 1.0 | 37.4 | 2.71 [A:OP1]a:1680 | 9.58 | 7 | U | O5' | a:1664 | 3.93 | 1.0 | 27.4 |  | 12.04 | 8 | Water 6 | | a:9759 | 4.24 | 1.0 | 44.0 |  | 4.30 | 9 | Water 7 | | a:8934 | 4.25 | 1.0 | 37.4 |  | 5.56 | 10 | Water 8 | | a:9936 | 4.26 | 1.0 | 42.4 |  | 4.72 | 11 | Water 9 | | a:7624 | 4.32 | 1.0 | 32.0 |  | 12.38 | 12 | U | OP1 | a:1664 | 4.33 | 1.0 | 31.7 |  | 9.86 | 13 | Water 10 | | a:9955 | 4.35 | 1.0 | 41.8 |  | 5.08 | 14 | Water 11 | | a:9295 | 4.38 | 1.0 | 44.4 |  | 3.25 | 15 | G | O3' | a:1663 | 4.66 | 1.0 | 26.4 |  | 13.23 | 16 | Water 12 | | a:6401 | 4.67 | 1.0 | 37.4 |  | 5.84 | 17 | G | OP2 | a:1665 | 4.68 | 1.0 | 35.6 |  | 10.47 | 18 | G | OP2 | a:1663 | 4.73 | 1.0 | 29.7 |  | 12.51 | 19 | A | OP1 | a:1680 | 4.74 | 1.0 | 35.9 |  | 10.82 |

  
**1^st^ shell coordination angles and distances (*trans* in "blue"; *cys* in "green"; deviations in "orange"):**   
• 1-2(W/W): 90º; 1-3(W/OP2): 91º; 1-4(W/W): *96º*; 1-5(W/W): *169º*; 2-3(W/OP2): *104º*; 2-4(W/W): 92º; 2-5(W/W): *100º*; 3-4(OP2/W): *163º*; 3-5(OP2/W): 87º; 4-5(W/W): *83º*;   
• 1-2(W/W): 3.0Å; 1-3(W/OP2): 3.0Å; 1-4(W/W): 3.2Å; 1-5(W/W): *4.3Å*; 2-3(W/OP2): *3.4Å*; 2-4(W/W): 3.1Å; 2-5(W/W): *3.4Å*; 3-4(OP2/W): *4.3Å*; 3-5(OP2/W): 3.1Å; 4-5(W/W): 3.0Å;   
  
**2.20 ± 0.09: the average coodination distance of the 1^st^ six ligands is Mg^2+^ compatible**  
**10: Number of potential Water contacts**   
**1 : Number of second shell contacts involving water or ions**   
**10%: Percentage of satisfied second shell Water contacts**   
• Contact(s): /O_w_/O_w_/O_w_/O_w_/OP2/  
• Contact code: O_ph_4O_w_  
• Contact(s) in the exclusion zone: /O_w_/  
→ Missing octahedral coordination for MG ←  
→ Slightly distorded coord. octahedron (7.1º) ←  
Go to ToC

### MG:a:3008

|  | Res. type | Atom | Residue | Distance | Occupancy | B-factor | Closest atom(s) to a 1^st^ shell water | EM\_dens. |
| --- | --- | --- | --- | --- | --- | --- | --- | --- |
| **Ion** | | | |  | 1.0 | 42.3 |  | 13.35 |
 1 | Water 1 | | a:10160 | 1.94 | 1.0 | 34.9 |  | 10.91 | 2 | Water 2 | | a:6474 | 2.01 | 1.0 | 26.5 | 2.77 [G:O6]a:2722 | 12.79 | 3 | Water 3 | | a:4583 | 2.08 | 1.0 | 28.4 | 2.64 [G:OP2]a:2692 | 11.02 | 4 | Water 4 | | a:6604 | 2.13 | 1.0 | 28.2 | 2.78 [G:N7]a:2722 | 9.16 | 5 | G | O6 | a:2723 | 2.22 | 1.0 | 25.7 |  | 14.87 | 6 | Water 5 | | a:5615 | 2.27 | 1.0 | 23.3 | 2.72 [U:O4]a:2717 | 14.40 | 7 | Water 6 | | a:10113 | 3.85 | 1.0 | 40.0 |  | 7.33 | 8 | Water 7 | | a:5497 | 3.89 | 1.0 | 34.2 |  | 8.56 | 9 | Water 8 | | a:9521 | 4.03 | 1.0 | 32.3 |  | 9.47 | 10 | Water 9 | | a:9301 | 4.04 | 1.0 | 34.0 |  | 8.58 | 11 | G | OP2 | a:2692 | 4.05 | 1.0 | 29.2 |  | 12.47 | 12 | G | O6 | a:2722 | 4.14 | 1.0 | 27.1 |  | 13.87 | 13 | G | N7 | a:2722 | 4.19 | 1.0 | 22.5 |  | 14.96 | 14 | G | N7 | a:2723 | 4.23 | 1.0 | 20.1 |  | 15.35 | 15 | Water 10 | | a:5315 | 4.28 | 1.0 | 31.3 |  | 12.77 | 16 | Water 11 | | a:6424 | 4.42 | 1.0 | 29.9 | 2.77 [U:OP1]a:2693 | 10.63 | 17 | U | OP1 | a:2693 | 4.46 | 1.0 | 26.1 |  | 12.92 | 18 | U | O4 | a:2717 | 4.69 | 1.0 | 32.5 |  | 12.34 | 19 | Water 12 | | a:7875 | 4.70 | 1.0 | 39.0 |  | 6.79 | 20 | Water 13 | | a:4711 | 4.76 | 1.0 | 37.6 |  | 6.78 |

  
**1^st^ shell coordination angles and distances (*trans* in "blue"; *cys* in "green"; deviations in "orange"):**   
• 1-2(W/W): *99º*; 1-3(W/W): 90º; 1-4(W/W): 94º; 1-5(W/O6): 172º; 1-6(W/W): 94º; 2-3(W/W): *170º*; 2-4(W/W): *95º*; 2-5(W/O6): 88º; 2-6(W/W): 85º; 3-4(W/W): 89º; 3-5(W/O6): *83º*; 3-6(W/W): 90º; 4-5(W/O6): 89º; 4-6(W/W): 171º; 5-6(O6/W): *82º*;   
• 1-2(W/W): 3.0Å; 1-3(W/W): 2.8Å; 1-4(W/W): 3.0Å; 1-5(W/O6): 4.1Å; 1-6(W/W): 3.1Å; 2-3(W/W): 4.1Å; 2-4(W/W): 3.1Å; 2-5(W/O6): 2.9Å; 2-6(W/W): 2.9Å; 3-4(W/W): 2.9Å; 3-5(W/O6): 2.8Å; 3-6(W/W): 3.1Å; 4-5(W/O6): 3.1Å; 4-6(W/W): *4.4Å*; 5-6(O6/W): 2.9Å;   
  
**2.11 ± 0.11: the average coodination distance of the 1^st^ six ligands is Mg^2+^ compatible**  
**10: Number of potential Water contacts**   
**4 : Number of second shell contacts involving water or ions**   
**40%: Percentage of satisfied second shell Water contacts**   
• Contact(s): /O6/O_w_/O_w_/O_w_/O_w_/O_w_/  
• Contact code: O_b_5O_w_  
→ Slightly distorded coord. octahedron (5.1º) ←  
Go to ToC

### MG:a:3009

|  | Res. type | Atom | Residue | Distance | Occupancy | B-factor | Closest atom(s) to a 1^st^ shell water | EM\_dens. |
| --- | --- | --- | --- | --- | --- | --- | --- | --- |
| **Ion** | | | |  | 1.0 | 39.8 |  | 13.00 |
 1 | G | OP1 | a:1830 | 1.98 | 1.0 | 23.5 |  | 14.72 | 2 | Water 1 | | a:6430 | 2.01 | 1.0 | 39.1 | 2.77 [A:OP2]a:1775 | 7.11 | 3 | Water 2 | | a:5533 | 2.02 | 1.0 | 36.2 | 2.71 [G:OP2]a:1977 | 8.27 | 4 | Water 3 | | a:9637 | 2.29 | 1.0 | 39.0 |  | 7.97 | 5 | Water 4 | | a:4149 | 2.33 | 1.0 | 39.2 | 2.59 [A:OP1]a:1774 | 8.77 | 6 | Water 5 | | a:3780 | 2.39 | 1.0 | 37.9 | 2.53 [G:OP2]a:1976 | 7.98 | 7 | G | O5' | a:1830 | 3.87 | 1.0 | 22.4 |  | 13.71 | 8 | G | OP2 | a:1830 | 4.04 | 1.0 | 28.8 |  | 15.01 | 9 | A | OP2 | a:1775 | 4.19 | 1.0 | 29.0 |  | 10.52 | 10 | G | OP2 | a:1977 | 4.21 | 1.0 | 27.6 |  | 11.15 | 11 | Water 6 | | a:9027 | 4.36 | 1.0 | 42.0 |  | 6.59 | 12 | Water 7 | | a:9266 | 4.39 | 1.0 | 24.8 |  | 14.50 | 13 | U | O3' | a:1829 | 4.45 | 1.0 | 23.5 |  | 14.64 | 14 | G | OP2 | a:1976 | 4.56 | 1.0 | 26.9 |  | 12.40 | 15 | A | OP1 | a:1774 | 4.60 | 1.0 | 33.1 |  | 8.39 | 16 | Water 8 | | a:8709 | 4.79 | 1.0 | 38.2 |  | 5.59 | 17 | Water 9 | | c:582 | 4.80 | 1.0 | 43.6 |  | 2.99 |

  
**1^st^ shell coordination angles and distances (*trans* in "blue"; *cys* in "green"; deviations in "orange"):**   
• 1-2(OP1/W): *102º*; 1-3(OP1/W): 92º; 1-4(OP1/W): *96º*; 2-3(W/W): 93º; 2-4(W/W): *84º*; 3-4(W/W): 172º;   
• 1-2(OP1/W): 3.1Å; 1-3(OP1/W): 2.9Å; 1-4(OP1/W): 3.2Å; 2-3(W/W): 2.9Å; 2-4(W/W): 2.9Å; 3-4(W/W): *4.3Å*;   
  
**2.17 ± 0.17: the average coodination distance of the 1^st^ six ligands is Mg^2+^ compatible**  
**10: Number of potential Water contacts**   
**4 : Number of second shell contacts involving water or ions**   
**40%: Percentage of satisfied second shell Water contacts**   
• Contact(s): /O_w_/O_w_/O_w_/OP1/  
• Contact code: O_ph_3O_w_  
• Contact(s) in the exclusion zone: /O_w_/O_w_/  
→ Missing octahedral coordination for MG ←  
→ Slightly distorded coord. octahedron (5.9º) ←  
Go to ToC

### MG:a:3010

|  | Res. type | Atom | Residue | Distance | Occupancy | B-factor | Closest atom(s) to a 1^st^ shell water | EM\_dens. |
| --- | --- | --- | --- | --- | --- | --- | --- | --- |
| **Ion** | | | |  | 1.0 | 53.1 |  | 7.11 |
 1 | Water 1 | | a:4748 | 1.79 | 1.0 | 36.4 | 2.65 [A:OP1]a:1804 | 7.97 | 2 | Water 2 | | a:8842 | 2.08 | 1.0 | 43.7 | 2.97 [U:O2]a:2207 | 4.43 | 3 | G | O6 | a:1816 | 2.13 | 1.0 | 24.5 |  | 15.21 | 4 | Water 3 | | a:6475 | 2.28 | 1.0 | 32.8 |  | 10.92 | 5 | Water 4 | | a:7904 | 2.44 | 1.0 | 29.3 | 2.87 [A:N1]a:1817 | 8.65 | 6 | Water 5 | | a:7009 | 2.93 | 1.0 | 40.2 |  | 3.99 | 7 | Water 6 | | a:4970 | 3.75 | 1.0 | 34.5 |  | 9.71 | 8 | A | OP1 | a:1804 | 3.89 | 1.0 | 35.1 |  | 8.53 | 9 | A | N1 | a:1817 | 4.12 | 1.0 | 20.2 |  | 14.99 | 10 | U | O2 | a:2207 | 4.51 | 1.0 | 42.5 |  | 4.53 | 11 | Water 7 | | a:7092 | 4.55 | 1.0 | 31.8 |  | 10.65 | 12 | Water 8 | | a:5423 | 4.58 | 1.0 | 45.7 |  | 3.50 | 13 | Water 9 | | a:5359 | 4.59 | 1.0 | 42.1 |  | 5.49 | 14 | A | OP2 | a:1804 | 4.73 | 1.0 | 30.1 |  | 9.66 |

  
**1^st^ shell coordination angles and distances (*trans* in "blue"; *cys* in "green"; deviations in "orange"):**   
• 1-2(W/W): 92º; 1-3(W/O6): *103º*; 1-4(W/W): 173º; 2-3(W/O6): *165º*; 2-4(W/W): 87º; 3-4(O6/W): *78º*;   
• 1-2(W/W): 2.8Å; 1-3(W/O6): 3.1Å; 1-4(W/W): 4.1Å; 2-3(W/O6): 4.2Å; 2-4(W/W): 3.0Å; 3-4(O6/W): 2.8Å;   
  
**2.28 ± 0.35: the average coodination distance of the 1^st^ six ligands is Mg^2+^ compatible**  
**10: Number of potential Water contacts**   
**3 : Number of second shell contacts involving water or ions**   
**30%: Percentage of satisfied second shell Water contacts**   
• Contact(s): /O6/O_w_/O_w_/  
• Contact code: O_b_2O_w_  
• Contact(s) in the exclusion zone: /O_w_/O_w_/O_w_/  
→ Missing octahedral coordination for MG ←  
→ Slightly distorded coord. octahedron (8.6º) ←  
Go to ToC

### MG:a:3011

|  | Res. type | Atom | Residue | Distance | Occupancy | B-factor | Closest atom(s) to a 1^st^ shell water | EM\_dens. |
| --- | --- | --- | --- | --- | --- | --- | --- | --- |
| **Ion** | | | |  | 1.0 | 52.4 |  | 6.55 |
 1 | Water 1 | | a:7558 | 1.84 | 1.0 | 41.6 | 2.85 [G:OP1]a:2601 | 5.43 | 2 | Water 2 | | a:5165 | 1.99 | 1.0 | 39.2 | 2.69 [G:N7]a:2599 | 7.47 | 3 | Water 3 | | a:9597 | 2.44 | 1.0 | 43.1 |  | 4.08 | 4 | Water 4 | | a:5456 | 2.65 | 1.0 | 38.4 |  | 4.96 | 5 | Water 5 | | a:4481 | 3.52 | 1.0 | 48.6 | 2.63 [G:O6]a:2599 | 4.40 | 6 | Water 6 | | a:9127 | 3.95 | 1.0 | 41.2 |  | 5.72 | 7 | Water 7 | | a:9777 | 4.08 | 1.0 | 35.9 |  | 8.43 | 8 | G | N7 | a:2599 | 4.12 | 1.0 | 22.0 |  | 12.80 | 9 | Water 8 | | a:7884 | 4.25 | 1.0 | 39.8 | 2.87 [G:OP1]a:2601 | 7.10 | 10 | G | OP1 | a:1908 | 4.32 | 1.0 | 37.7 |  | 6.23 | 11 | Water 9 | | a:5349 | 4.34 | 1.0 | 41.3 |  | 4.06 | 12 | Water 10 | | a:6857 | 4.37 | 1.0 | 41.9 |  | 6.02 | 13 | G | OP1 | a:2601 | 4.38 | 1.0 | 31.0 |  | 12.35 | 14 | U | O4 | a:1975 | 4.47 | 1.0 | 42.7 |  | 5.89 | 15 | G | O6 | a:2599 | 4.68 | 1.0 | 34.0 |  | 13.11 |

  
**1^st^ shell coordination angles and distances (*trans* in "blue"; *cys* in "green"; deviations in "orange"):**   
• 1-2(W/W): 86º;   
• 1-2(W/W): 2.6Å;   
  
**2.73 ± 0.77: the average coodination distance of the 1^st^ six ligands is K^+^ compatible**  
**12: Number of potential Water contacts**   
**3 : Number of second shell contacts involving water or ions**   
**25%: Percentage of satisfied second shell Water contacts**   
• Contact(s): /O_w_/  
• Contact code: O_w_  
• Contact(s) in the exclusion zone: /O_w_/O_w_/O_w_/  
→ Missing octahedral coordination for MG ←  
Go to ToC

### MG:a:3012

|  | Res. type | Atom | Residue | Distance | Occupancy | B-factor | Closest atom(s) to a 1^st^ shell water | EM\_dens. |
| --- | --- | --- | --- | --- | --- | --- | --- | --- |
| **Ion** | | | |  | 1.0 | 51.3 |  | 6.47 |
 1 | Water 1 | | a:5538 | 2.03 | 1.0 | 41.3 | 2.71 [C:OP2]a:1946 | 5.72 | 2 | C | OP2 | a:1945 | 2.33 | 1.0 | 39.1 |  | 7.10 | 3 | C | OP1 | a:1945 | 3.86 | 1.0 | 39.0 |  | 8.36 | 4 | Water 2 | | a:5677 | 4.01 | 1.0 | 35.6 | 2.91 [C:OP2]a:1945 | 7.92 | 5 | U | O2' | a:1947 | 4.04 | 1.0 | 38.7 |  | 8.80 | 6 | U | OP2 | a:1948 | 4.16 | 1.0 | 44.5 |  | 5.27 | 7 | C | OP2 | a:1946 | 4.29 | 1.0 | 37.6 |  | 7.11 | 8 | Water 3 | | a:3803 | 4.33 | 1.0 | 45.3 |  | 7.94 | 9 | U | O3' | a:1944 | 4.45 | 1.0 | 37.5 |  | 8.85 | 10 | U | O3' | a:1947 | 4.49 | 1.0 | 39.9 |  | 6.71 | 11 | C | O5' | a:1945 | 4.67 | 1.0 | 38.2 |  | 8.79 |

  
• • **3.40 ± 0.88: the average coodination distance of the 1^st^ six ligands is K^+^ compatible**  
**4: Number of potential Water contacts**   
**2 : Number of second shell contacts involving water or ions**   
**50%: Percentage of satisfied second shell Water contacts**   
• Contact(s): /O_w_/  
• Contact code: O_w_  
• Contact(s) in the exclusion zone: /OP2/  
→ Missing octahedral coordination for MG ←  
Go to ToC

### MG:a:3013

|  | Res. type | Atom | Residue | Distance | Occupancy | B-factor | Closest atom(s) to a 1^st^ shell water | EM\_dens. |
| --- | --- | --- | --- | --- | --- | --- | --- | --- |
| **Ion** | | | |  | 1.0 | 52.2 |  | 5.03 |
 1 | Water 1 | | a:7073 | 1.97 | 1.0 | 41.4 | 2.81 [U:OP2]a:1959 | 4.74 | 2 | Water 2 | | a:9678 | 2.08 | 1.0 | 41.2 |  | 5.83 | 3 | Water 3 | | a:10124 | 2.10 | 1.0 | 44.7 | 2.55 [MG:MG]a:3053 | 4.51 | 4 | Water 4 | | a:5801 | 2.27 | 1.0 | 36.3 | 2.73 [G:N7]a:1952 | 7.48 | 5 | Water 5 | | a:5527 | 2.32 | 1.0 | 36.9 | 2.71 [G:O6]a:1952 // 2.95 [MG:MG]a:3053 | 7.75 | 6 | **MG** | MG | a:3053 | 3.86 | 1.0 | 43.8 |  | 9.83 | 7 | Water 6 | | a:5143 | 3.88 | 1.0 | 46.6 |  | 4.31 | 8 | Water 7 | | a:10045 | 4.10 | 1.0 | 41.8 |  | 6.71 | 9 | Water 8 | | a:3791 | 4.19 | 1.0 | 36.7 |  | 7.85 | 10 | Water 9 | | a:6403 | 4.22 | 1.0 | 35.4 |  | 6.42 | 11 | Water 10 | | a:4627 | 4.34 | 1.0 | 36.5 |  | 7.58 | 12 | U | OP2 | a:1959 | 4.48 | 1.0 | 32.5 |  | 9.12 | 13 | G | O6 | a:1952 | 4.53 | 1.0 | 32.9 |  | 10.97 | 14 | G | N7 | a:1952 | 4.73 | 1.0 | 23.0 |  | 11.70 |

  
**1^st^ shell coordination angles and distances (*trans* in "blue"; *cys* in "green"; deviations in "orange"):**   
• 1-2(W/W): 86º; 1-3(W/W): 94º; 1-4(W/W): *167º*; 2-3(W/W): *167º*; 2-4(W/W): *104º*; 3-4(W/W): *75º*;   
• 1-2(W/W): 2.8Å; 1-3(W/W): 3.0Å; 1-4(W/W): *4.2Å*; 2-3(W/W): 4.1Å; 2-4(W/W): *3.4Å*; 3-4(W/W): 2.6Å;   
  
**2.43 ± 0.65: the average coodination distance of the 1^st^ six ligands is Na^+^** compatible  
**10: Number of potential Water contacts**   
**5 : Number of second shell contacts involving water or ions**   
**50%: Percentage of satisfied second shell Water contacts**   
• Contact(s): /O_w_/O_w_/O_w_/O_w_/  
• Contact code: 4O_w_  
• Contact(s) in the exclusion zone: /O_w_/  
→ Missing octahedral coordination for MG ←  
→→→ Distorded coord. octahedron (10.6º) ←←←  
Go to ToC

### MG:a:3014

|  | Res. type | Atom | Residue | Distance | Occupancy | B-factor | Closest atom(s) to a 1^st^ shell water | EM\_dens. |
| --- | --- | --- | --- | --- | --- | --- | --- | --- |
| **Ion** | | | |  | 1.0 | 45.7 |  | 9.65 |
 1 | Water 1 | | a:9019 | 1.94 | 1.0 | 36.2 | 3.00 [G:O6]a:563 | 6.53 | 2 | Water 2 | | a:9391 | 1.96 | 1.0 | 31.9 |  | 8.72 | 3 | Water 3 | | a:3777 | 2.23 | 1.0 | 36.9 | 2.53 [A:OP1]a:2041 | 6.80 | 4 | G | N7 | a:563 | 2.39 | 1.0 | 20.6 |  | 12.33 | 5 | Water 4 | | a:9848 | 2.48 | 1.0 | 44.1 |  | 6.29 | 6 | Water 5 | | a:10062 | 2.52 | 1.0 | 40.0 |  | 7.12 | 7 | G | O6 | a:563 | 3.70 | 1.0 | 32.4 |  | 10.51 | 8 | Water 6 | | a:7011 | 4.12 | 1.0 | 32.0 |  | 9.76 | 9 | Water 7 | | a:6269 | 4.20 | 1.0 | 40.8 |  | 6.65 | 10 | G | OP2 | a:563 | 4.22 | 1.0 | 30.7 |  | 10.83 | 11 | A | OP1 | a:2041 | 4.32 | 1.0 | 28.9 |  | 13.20 | 12 | Water 8 | | a:4866 | 4.43 | 1.0 | 38.4 |  | 6.77 | 13 | Water 9 | | a:3901 | 4.52 | 1.0 | 36.4 | 2.56 [G:OP2]a:563 // 2.62 [A:OP1]a:2041 | 6.67 | 14 | C | OP2 | a:562 | 4.72 | 1.0 | 31.2 |  | 9.37 | 15 | Water 10 | | a:5525 | 4.76 | 1.0 | 37.0 |  | 6.74 |

  
**1^st^ shell coordination angles and distances (*trans* in "blue"; *cys* in "green"; deviations in "orange"):**   
• 1-2(W/W): *96º*; 1-3(W/W): *166º*; 2-3(W/W): 92º;   
• 1-2(W/W): 2.9Å; 1-3(W/W): 4.1Å; 2-3(W/W): 3.0Å;   
  
**2.25 ± 0.23: the average coodination distance of the 1^st^ six ligands is Mg^2+^ compatible**  
**10: Number of potential Water contacts**   
**2 : Number of second shell contacts involving water or ions**   
**20%: Percentage of satisfied second shell Water contacts**   
• Contact(s): /O_w_/O_w_/O_w_/  
• Contact code: 3O_w_  
• Contact(s) in the exclusion zone: /N7/O_w_/O_w_/  
→ Missing octahedral coordination for MG ←  
Go to ToC

### MG:a:3015

|  | Res. type | Atom | Residue | Distance | Occupancy | B-factor | Closest atom(s) to a 1^st^ shell water | EM\_dens. |
| --- | --- | --- | --- | --- | --- | --- | --- | --- |
| **Ion** | | | |  | 1.0 | 44.0 |  | 12.18 |
 1 | Water 1 | | a:7674 | 2.28 | 1.0 | 39.5 | 2.86 [C:OP1]a:2054 | 5.19 | 2 | A | OP2 | a:2055 | 2.46 | 1.0 | 22.0 |  | 14.67 | 3 | Water 2 | | a:6316 | 3.72 | 1.0 | 37.9 | 2.76 [A:OP1]a:2055 | 8.12 | 4 | Water 3 | | a:4931 | 3.82 | 1.0 | 38.7 | 2.67 [G:OP2]a:2582 | 5.34 | 5 | A | N7 | a:2055 | 4.05 | 1.0 | 16.3 |  | 18.41 | 6 | G | OP2 | a:2582 | 4.07 | 1.0 | 25.1 |  | 11.94 | 7 | G | OP1 | a:2582 | 4.28 | 1.0 | 27.1 |  | 15.89 | 8 | A | OP1 | a:2055 | 4.29 | 1.0 | 27.3 |  | 12.56 | 9 | C | OP2 | a:2054 | 4.41 | 1.0 | 23.2 |  | 15.23 | 10 | C | O3' | a:2054 | 4.42 | 1.0 | 25.2 |  | 16.10 | 11 | Water 4 | | a:3676 | 4.52 | 1.0 | 47.7 |  | 5.68 | 12 | C | O5' | a:2054 | 4.57 | 1.0 | 26.6 |  | 13.98 | 13 | C | OP1 | a:2054 | 4.62 | 1.0 | 27.9 |  | 12.78 | 14 | Water 5 | | a:6092 | 4.69 | 1.0 | 44.2 | 2.75 [C:OP1]a:2054 | 3.71 |

  
• • **3.40 ± 0.74: the average coodination distance of the 1^st^ six ligands is K^+^ compatible**  
**6: Number of potential Water contacts**   
**3 : Number of second shell contacts involving water or ions**   
**50%: Percentage of satisfied second shell Water contacts**   
• Contact(s): /O_w_/  
• Contact code: O_w_  
• Contact(s) in the exclusion zone: /OP2/  
→ Missing octahedral coordination for MG ←  
Go to ToC

### MG:a:3016

|  | Res. type | Atom | Residue | Distance | Occupancy | B-factor | Closest atom(s) to a 1^st^ shell water | EM\_dens. |
| --- | --- | --- | --- | --- | --- | --- | --- | --- |
| **Ion** | | | |  | 1.0 | 44.6 |  | 11.62 |
 1 | Water 1 | | a:9745 | 1.92 | 1.0 | 37.7 |  | 6.94 | 2 | Water 2 | | a:4487 | 1.98 | 1.0 | 36.7 | 2.63 [A:OP2]a:2390 | 7.81 | 3 | Water 3 | | a:9757 | 2.04 | 1.0 | 42.4 |  | 6.84 | 4 | Water 4 | | a:9224 | 2.06 | 1.0 | 43.8 |  | 7.49 | 5 | Water 5 | | a:5906 | 2.19 | 1.0 | 30.8 | 2.74 [A:N7]a:2390 | 9.84 | 6 | Water 6 | | a:4267 | 2.26 | 1.0 | 37.3 | 2.61 [U:O4]a:2391 | 9.38 | 7 | A | OP2 | a:2390 | 3.88 | 1.0 | 34.1 |  | 8.93 | 8 | Water 7 | | a:3559 | 3.97 | 1.0 | 42.7 |  | 4.74 | 9 | Water 8 | | a:4613 | 4.16 | 1.0 | 38.5 |  | 6.58 | 10 | Water 9 | | a:9454 | 4.20 | 1.0 | 32.8 |  | 7.60 | 11 | Water 10 | | a:10082 | 4.23 | 1.0 | 43.2 |  | 5.89 | 12 | A | N7 | a:2390 | 4.34 | 1.0 | 22.8 |  | 13.38 | 13 | Water 11 | | a:7788 | 4.52 | 1.0 | 40.3 |  | 5.60 | 14 | U | O4 | a:2391 | 4.53 | 1.0 | 29.6 |  | 12.57 | 15 | Water 12 | | a:6959 | 4.58 | 1.0 | 41.5 |  | 4.89 |

  
**1^st^ shell coordination angles and distances (*trans* in "blue"; *cys* in "green"; deviations in "orange"):**   
• 1-2(W/W): 174º; 1-3(W/W): 91º; 1-4(W/W): 90º; 1-5(W/W): 90º; 1-6(W/W): 88º; 2-3(W/W): *95º*; 2-4(W/W): 89º; 2-5(W/W): 90º; 2-6(W/W): 86º; 3-4(W/W): *99º*; 3-5(W/W): 94º; 3-6(W/W): 171º; 4-5(W/W): *167º*; 4-6(W/W): 90º; 5-6(W/W): *78º*;   
• 1-2(W/W): 3.9Å; 1-3(W/W): 2.8Å; 1-4(W/W): 2.8Å; 1-5(W/W): 2.9Å; 1-6(W/W): 2.9Å; 2-3(W/W): 3.0Å; 2-4(W/W): 2.8Å; 2-5(W/W): 2.9Å; 2-6(W/W): 2.9Å; 3-4(W/W): 3.1Å; 3-5(W/W): 3.1Å; 3-6(W/W): *4.3Å*; 4-5(W/W): *4.2Å*; 4-6(W/W): 3.1Å; 5-6(W/W): 2.8Å;   
  
**2.08 ± 0.12: the average coodination distance of the 1^st^ six ligands is Mg^2+^ compatible**  
**12: Number of potential Water contacts**   
**3 : Number of second shell contacts involving water or ions**   
**25%: Percentage of satisfied second shell Water contacts**   
• Contact(s): /O_w_/O_w_/O_w_/O_w_/O_w_/O_w_/  
• Contact code: 6O_w_  
Ooctahedral coordination (4.5º)   
Go to ToC

### MG:a:3017

|  | Res. type | Atom | Residue | Distance | Occupancy | B-factor | Closest atom(s) to a 1^st^ shell water | EM\_dens. |
| --- | --- | --- | --- | --- | --- | --- | --- | --- |
| **Ion** | | | |  | 1.0 | 49.3 |  | 5.81 |
 1 | Water 1 | | a:3913 | 2.27 | 1.0 | 37.8 | 2.56 [G:OP2]a:2294 | 8.31 | 2 | Water 2 | | a:9728 | 2.48 | 1.0 | 47.1 |  | 3.41 | 3 | Water 3 | | a:6237 | 3.63 | 1.0 | 43.2 |  | 3.72 | 4 | Water 4 | | a:5099 | 3.83 | 1.0 | 44.8 | 2.71 [G:OP2]a:2294 | 3.70 | 5 | G | OP2 | a:2294 | 4.01 | 1.0 | 41.3 |  | 6.42 | 6 | G | OP2 | a:2341 | 4.14 | 1.0 | 42.7 |  | 8.14 | 7 | Water 5 | | a:8233 | 4.20 | 1.0 | 36.5 | 2.90 [C:OP1]a:2389 | 8.02 | 8 | C | OP1 | a:2389 | 4.25 | 1.0 | 41.1 |  | 7.32 | 9 | Water 6 | | a:6314 | 4.26 | 1.0 | 42.5 | 2.76 [G:OP2]a:2293 // 2.80 [G:OP2]a:2294 | 6.52 | 10 | C | OP2 | a:2389 | 4.47 | 1.0 | 34.4 |  | 9.30 | 11 | G | OP1 | a:2293 | 4.48 | 1.0 | 38.1 |  | 7.77 | 12 | Water 7 | | a:9532 | 4.74 | 1.0 | 39.3 |  | 4.21 | 13 | G | OP2 | a:2293 | 4.79 | 1.0 | 39.0 |  | 6.60 |

  
• • **3.39 ± 0.74: the average coodination distance of the 1^st^ six ligands is K^+^ compatible**  
**8: Number of potential Water contacts**   
**2 : Number of second shell contacts involving water or ions**   
**25%: Percentage of satisfied second shell Water contacts**   
• Contact(s): /O_w_/  
• Contact code: O_w_  
• Contact(s) in the exclusion zone: /O_w_/  
→ Missing octahedral coordination for MG ←  
Go to ToC

### MG:a:3018

|  | Res. type | Atom | Residue | Distance | Occupancy | B-factor | Closest atom(s) to a 1^st^ shell water | EM\_dens. |
| --- | --- | --- | --- | --- | --- | --- | --- | --- |
| **Ion** | | | |  | 1.0 | 47.3 |  | 7.14 |
 1 | Water 1 | | a:5978 | 2.35 | 1.0 | 41.9 | 2.74 [G:O6]a:2404 | 5.31 | 2 | Water 2 | | a:4743 | 2.63 | 1.0 | 40.0 | 2.65 [G:N7]a:2419 | 6.35 | 3 | G | O6 | a:2419 | 2.73 | 1.0 | 27.5 |  | 11.75 | 4 | Water 3 | | a:5297 | 2.74 | 1.0 | 41.4 | 2.70 [G:N7]a:2418 | 6.43 | 5 | U | O4 | a:2405 | 2.93 | 1.0 | 40.8 |  | 8.35 | 6 | G | O6 | a:2418 | 3.40 | 1.0 | 30.8 |  | 10.24 | 7 | G | O6 | a:2404 | 3.63 | 1.0 | 36.3 |  | 9.58 | 8 | Water 4 | | a:6175 | 3.98 | 1.0 | 27.6 | 2.75 [G:O6]a:2418 | 13.65 | 9 | G | N7 | a:2419 | 4.07 | 1.0 | 22.8 |  | 11.76 | 10 | G | N7 | a:2418 | 4.35 | 1.0 | 26.1 |  | 10.94 | 11 | Water 5 | | a:3771 | 4.42 | 1.0 | 50.3 | 2.53 [U:O4]a:2405 | 3.67 | 12 | Water 6 | | a:5012 | 4.64 | 1.0 | 42.8 |  | 4.56 |

  
• • **2.80 ± 0.32: the average coodination distance of the 1^st^ six ligands is K^+^ compatible**  
**6: Number of potential Water contacts**   
**3 : Number of second shell contacts involving water or ions**   
**50%: Percentage of satisfied second shell Water contacts**   
• Contact code: → No contacts ←   
• Contact(s) in the exclusion zone: /O4/O6/O6/O_w_/O_w_/O_w_/  
→ Consider K in place of MG (2 valid contact(s) for K vs. 0 for MG ←)  
Go to ToC

### MG:a:3019

|  | Res. type | Atom | Residue | Distance | Occupancy | B-factor | Closest atom(s) to a 1^st^ shell water | EM\_dens. |
| --- | --- | --- | --- | --- | --- | --- | --- | --- |
| **Ion** | | | |  | 1.0 | 52.5 |  | 9.76 |
 1 | Water 1 | | a:6264 | 2.01 | 1.0 | 35.9 | 2.76 [G:O6]a:2499 | 7.22 | 2 | Water 2 | | a:10279 | 2.03 | 1.0 | 39.9 |  | 5.70 | 3 | Water 3 | | a:10218 | 2.15 | 1.0 | 44.6 |  | 5.24 | 4 | Water 4 | | a:5603 | 2.23 | 1.0 | 40.2 | 2.72 [G:O6]a:2459 | 6.79 | 5 | Water 5 | | a:5661 | 2.26 | 1.0 | 37.6 | 2.72 [G:O6]a:2498 | 7.99 | 6 | Water 6 | | a:8635 | 2.27 | 1.0 | 40.7 | 2.95 [G:N7]a:2498 | 6.70 | 7 | Water 7 | | a:9949 | 3.82 | 1.0 | 34.2 |  | 9.29 | 8 | Water 8 | | a:9286 | 4.06 | 1.0 | 33.7 |  | 7.19 | 9 | Water 9 | | a:9000 | 4.15 | 1.0 | 35.2 | 2.99 [C:N4]a:2460 | 6.99 | 10 | G | O6 | a:2499 | 4.18 | 1.0 | 27.8 |  | 12.68 | 11 | G | O6 | a:2498 | 4.37 | 1.0 | 34.4 |  | 12.46 | 12 | G | N7 | a:2498 | 4.43 | 1.0 | 28.9 |  | 9.83 | 13 | G | O6 | a:2459 | 4.62 | 1.0 | 24.0 |  | 15.27 | 14 | Water 10 | | a:5277 | 4.69 | 1.0 | 37.6 |  | 10.33 | 15 | Water 11 | | a:5118 | 4.75 | 1.0 | 45.5 |  | 3.28 |

  
**1^st^ shell coordination angles and distances (*trans* in "blue"; *cys* in "green"; deviations in "orange"):**   
• 1-2(W/W): *100º*; 1-3(W/W): 170º; 1-4(W/W): *85º*; 1-5(W/W): 88º; 1-6(W/W): 90º; 2-3(W/W): 88º; 2-4(W/W): 89º; 2-5(W/W): *168º*; 2-6(W/W): 95º; 3-4(W/W): 90º; 3-5(W/W): *85º*; 3-6(W/W): *95º*; 4-5(W/W): *100º*; 4-6(W/W): 174º; 5-6(W/W): *77º*;   
• 1-2(W/W): 3.1Å; 1-3(W/W): 4.1Å; 1-4(W/W): 2.9Å; 1-5(W/W): 3.0Å; 1-6(W/W): 3.0Å; 2-3(W/W): 2.9Å; 2-4(W/W): 3.0Å; 2-5(W/W): *4.3Å*; 2-6(W/W): 3.2Å; 3-4(W/W): 3.1Å; 3-5(W/W): 3.0Å; 3-6(W/W): *3.3Å*; 4-5(W/W): *3.4Å*; 4-6(W/W): *4.5Å*; 5-6(W/W): 2.8Å;   
  
**2.16 ± 0.10: the average coodination distance of the 1^st^ six ligands is Mg^2+^ compatible**  
**12: Number of potential Water contacts**   
**4 : Number of second shell contacts involving water or ions**   
**33%: Percentage of satisfied second shell Water contacts**   
• Contact(s): /O_w_/O_w_/O_w_/O_w_/O_w_/O_w_/  
• Contact code: 6O_w_  
→ Slightly distorded coord. octahedron (5.8º) ←  
Go to ToC

### MG:a:3020

|  | Res. type | Atom | Residue | Distance | Occupancy | B-factor | Closest atom(s) to a 1^st^ shell water | EM\_dens. |
| --- | --- | --- | --- | --- | --- | --- | --- | --- |
| **Ion** | | | |  | 1.0 | 42.7 |  | 8.86 |
 1 | Water 1 | | a:4210 | 1.96 | 1.0 | 35.8 | 2.60 [C:OP2]a:2771 | 6.28 | 2 | Water 2 | | a:5782 | 1.97 | 1.0 | 41.7 | 2.73 [U:O4]a:2773 | 5.37 | 3 | Water 3 | | a:9298 | 1.98 | 1.0 | 42.5 |  | 3.87 | 4 | A | OP1 | a:2770 | 2.05 | 1.0 | 34.7 |  | 8.17 | 5 | Water 4 | | a:3424 | 2.06 | 1.0 | 32.8 | 2.35 [C:OP2]a:2771 | 8.09 | 6 | Water 5 | | a:5487 | 2.17 | 1.0 | 37.2 | 2.71 [U:O4]a:2772 // 2.95 [U:O4]a:2773 | 7.27 | 7 | C | OP2 | a:2771 | 3.36 | 1.0 | 34.6 |  | 9.20 | 8 | Water 6 | | a:8154 | 3.78 | 1.0 | 36.6 |  | 7.38 | 9 | U | O4 | a:2773 | 4.03 | 1.0 | 36.3 |  | 9.28 | 10 | A | O3' | a:2769 | 4.12 | 1.0 | 31.4 |  | 10.37 | 11 | A | OP2 | a:2770 | 4.13 | 1.0 | 39.5 |  | 6.02 | 12 | A | O5' | a:2770 | 4.53 | 1.0 | 31.9 |  | 9.10 | 13 | U | O4 | a:2772 | 4.59 | 1.0 | 31.2 |  | 10.10 | 14 | Water 7 | | a:6087 | 4.68 | 1.0 | 42.2 |  | 5.46 | 15 | G | N7 | a:2649 | 4.80 | 1.0 | 24.8 |  | 11.21 |

  
**1^st^ shell coordination angles and distances (*trans* in "blue"; *cys* in "green"; deviations in "orange"):**   
• 1-2(W/W): *105º*; 1-3(W/W): *97º*; 1-4(W/OP1): *84º*; 1-5(W/W): *83º*; 1-6(W/W): *168º*; 2-3(W/W): *75º*; 2-4(W/OP1): *169º*; 2-5(W/W): *96º*; 2-6(W/W): *81º*; 3-4(W/OP1): *98º*; 3-5(W/W): 170º; 3-6(W/W): 95º; 4-5(OP1/W): 92º; 4-6(OP1/W): 91º; 5-6(W/W): 86º;   
• 1-2(W/W): 3.1Å; 1-3(W/W): 2.9Å; 1-4(W/OP1): 2.7Å; 1-5(W/W): 2.7Å; 1-6(W/W): 4.1Å; 2-3(W/W): 2.4Å; 2-4(W/OP1): 4.0Å; 2-5(W/W): 3.0Å; 2-6(W/W): 2.7Å; 3-4(W/OP1): 3.0Å; 3-5(W/W): 4.0Å; 3-6(W/W): 3.1Å; 4-5(OP1/W): 2.9Å; 4-6(OP1/W): 3.0Å; 5-6(W/W): 2.9Å;   
  
**2.03 ± 0.07: the average coodination distance of the 1^st^ six ligands is Mg^2+^ compatible**  
**10: Number of potential Water contacts**   
**5 : Number of second shell contacts involving water or ions**   
**50%: Percentage of satisfied second shell Water contacts**   
• Contact(s): /O_w_/O_w_/O_w_/O_w_/O_w_/OP1/  
• Contact code: O_ph_5O_w_  
• Contact(s) in the exclusion zone: /OP2/  
• MG hydration shell issue: water too close (< 2.4 Å.)  
→ Slightly distorded coord. octahedron (7.8º) ←  
Go to ToC

### MG:a:3021

|  | Res. type | Atom | Residue | Distance | Occupancy | B-factor | Closest atom(s) to a 1^st^ shell water | EM\_dens. |
| --- | --- | --- | --- | --- | --- | --- | --- | --- |
| **Ion** | | | |  | 1.0 | 52.7 |  | 9.10 |
 1 | Water 1 | | a:10267 | 1.92 | 1.0 | 48.0 |  | 4.12 | 2 | Water 2 | | a:7816 | 1.96 | 1.0 | 35.4 | 2.87 [G:O6]a:2554 | 7.48 | 3 | Water 3 | | a:6708 | 2.06 | 1.0 | 43.3 | 2.79 [G:O6]a:2553 | 5.79 | 4 | Water 4 | | a:10278 | 2.08 | 1.0 | 45.9 |  | 4.89 | 5 | Water 5 | | a:4375 | 2.26 | 1.0 | 36.0 | 2.62 [G:O6]a:2561 | 7.74 | 6 | Water 6 | | a:6354 | 2.49 | 1.0 | 31.1 | 2.76 [G:N7]a:2553 | 7.93 | 7 | G | O6 | a:2553 | 4.03 | 1.0 | 32.7 |  | 11.93 | 8 | Water 7 | | a:8617 | 4.18 | 1.0 | 38.1 |  | 5.86 | 9 | G | O6 | a:2561 | 4.24 | 1.0 | 39.1 |  | 11.13 | 10 | Water 8 | | a:8623 | 4.30 | 1.0 | 38.0 |  | 5.71 | 11 | Water 9 | | a:7737 | 4.31 | 1.0 | 40.0 |  | 6.29 | 12 | Water 10 | | a:10041 | 4.37 | 1.0 | 36.8 |  | 7.47 | 13 | Water 11 | | a:3612 | 4.40 | 1.0 | 42.4 |  | 4.97 | 14 | Water 12 | | a:3986 | 4.42 | 1.0 | 43.2 |  | 4.67 | 15 | G | O6 | a:2554 | 4.63 | 1.0 | 26.7 |  | 14.57 | 16 | G | N7 | a:2553 | 4.65 | 1.0 | 22.8 |  | 12.53 | 17 | Water 13 | | a:4934 | 4.79 | 1.0 | 34.2 |  | 4.96 |

  
**1^st^ shell coordination angles and distances (*trans* in "blue"; *cys* in "green"; deviations in "orange"):**   
• 1-2(W/W): 92º; 1-3(W/W): *169º*; 1-4(W/W): 94º; 1-5(W/W): *101º*; 2-3(W/W): *82º*; 2-4(W/W): 174º; 2-5(W/W): 92º; 3-4(W/W): 92º; 3-5(W/W): 89º; 4-5(W/W): 87º;   
• 1-2(W/W): 2.8Å; 1-3(W/W): 4.0Å; 1-4(W/W): 2.9Å; 1-5(W/W): *3.2Å*; 2-3(W/W): 2.6Å; 2-4(W/W): 4.0Å; 2-5(W/W): 3.0Å; 3-4(W/W): 3.0Å; 3-5(W/W): 3.0Å; 4-5(W/W): 3.0Å;   
  
**2.13 ± 0.20: the average coodination distance of the 1^st^ six ligands is Mg^2+^ compatible**  
**12: Number of potential Water contacts**   
**4 : Number of second shell contacts involving water or ions**   
**33%: Percentage of satisfied second shell Water contacts**   
• Contact(s): /O_w_/O_w_/O_w_/O_w_/O_w_/  
• Contact code: 5O_w_  
• Contact(s) in the exclusion zone: /O_w_/  
→ Missing octahedral coordination for MG ←  
Ooctahedral coordination (4.9º)   
Go to ToC

### MG:a:3022

|  | Res. type | Atom | Residue | Distance | Occupancy | B-factor | Closest atom(s) to a 1^st^ shell water | EM\_dens. |
| --- | --- | --- | --- | --- | --- | --- | --- | --- |
| **Ion** | | | |  | 1.0 | 51.9 |  | 9.69 |
 1 | Water 1 | | a:8311 | 1.93 | 1.0 | 37.8 | 2.91 [G:OP2]a:2578 | 7.76 | 2 | Water 2 | | a:10249 | 2.06 | 1.0 | 44.7 |  | 4.68 | 3 | Water 3 | | a:9496 | 2.09 | 1.0 | 47.7 | 2.73 [K:K]a:3107 | 5.75 | 4 | Water 4 | | a:6947 | 2.34 | 1.0 | 36.5 | 2.80 [G:N7]a:2578 | 6.95 | 5 | U | O4 | a:2575 | 2.40 | 1.0 | 22.9 |  | 15.39 | 6 | Water 5 | | a:4919 | 3.80 | 1.0 | 44.1 |  | 5.70 | 7 | Water 6 | | a:8478 | 4.18 | 1.0 | 40.5 |  | 9.27 | 8 | **K** | K | a:3107 | 4.18 | 1.0 | 30.0 |  | 15.75 | 9 | G | N7 | a:2578 | 4.26 | 1.0 | 16.6 |  | 15.42 | 10 | G | OP2 | a:2578 | 4.41 | 1.0 | 31.5 |  | 11.28 | 11 | G | N7 | a:2574 | 4.70 | 1.0 | 16.3 |  | 14.01 |

  
**1^st^ shell coordination angles and distances (*trans* in "blue"; *cys* in "green"; deviations in "orange"):**   
• 1-2(W/W): *82º*; 1-3(W/W): 170º; 2-3(W/W): *99º*;   
• 1-2(W/W): 2.6Å; 1-3(W/W): 4.0Å; 2-3(W/W): 3.2Å;   
  
**2.44 ± 0.63: the average coodination distance of the 1^st^ six ligands is Na^+^** compatible  
**10: Number of potential Water contacts**   
**3 : Number of second shell contacts involving water or ions**   
**30%: Percentage of satisfied second shell Water contacts**   
• Contact(s): /O_w_/O_w_/O_w_/  
• Contact code: 3O_w_  
• Contact(s) in the exclusion zone: /O4/O_w_/  
→ Missing octahedral coordination for MG ←  
Go to ToC

### MG:a:3023

|  | Res. type | Atom | Residue | Distance | Occupancy | B-factor | Closest atom(s) to a 1^st^ shell water | EM\_dens. |
| --- | --- | --- | --- | --- | --- | --- | --- | --- |
| **Ion** | | | |  | 1.0 | 48.2 |  | 6.30 |
 1 | G | OP2 | a:2586 | 2.11 | 1.0 | 33.6 |  | 8.32 | 2 | G | OP2 | a:2587 | 2.12 | 1.0 | 37.7 |  | 9.11 | 3 | Water 1 | | a:5990 | 2.47 | 1.0 | 43.2 | 2.74 [G:N7]a:2587 | 4.47 | 4 | Water 2 | | a:7411 | 3.50 | 1.0 | 43.5 |  | 4.06 | 5 | G | O3' | a:2585 | 3.66 | 1.0 | 30.1 |  | 11.16 | 6 | Water 3 | | a:6451 | 4.06 | 1.0 | 39.7 |  | 4.33 | 7 | G | O5' | a:2586 | 4.30 | 1.0 | 33.5 |  | 8.85 | 8 | G | O5' | a:2587 | 4.32 | 1.0 | 35.1 |  | 11.69 | 9 | G | O3' | a:2586 | 4.42 | 1.0 | 31.5 |  | 9.41 | 10 | G | OP1 | a:2587 | 4.47 | 1.0 | 38.6 |  | 6.48 | 11 | G | N7 | a:2587 | 4.48 | 1.0 | 28.8 |  | 8.44 | 12 | G | OP1 | a:2586 | 4.51 | 1.0 | 34.4 |  | 8.34 | 13 | G | O2' | a:2585 | 4.51 | 1.0 | 33.5 |  | 11.03 | 14 | U | O4 | a:2588 | 4.56 | 1.0 | 48.0 |  | 1.54 |

  
**1^st^ shell coordination angles and distances (*trans* in "blue"; *cys* in "green"; deviations in "orange"):**   
• 1-2(OP2/OP2): *83º*;   
• 1-2(OP2/OP2): 2.8Å;   
  
**2.99 ± 0.78: the average coodination distance of the 1^st^ six ligands is K^+^ compatible**  
**6: Number of potential Water contacts**   
**1 : Number of second shell contacts involving water or ions**   
**17%: Percentage of satisfied second shell Water contacts**   
• Contact(s): /OP2/OP2/  
• Contact code: *cis-*2O_ph_  
• Contact(s) in the exclusion zone: /O_w_/  
→ Missing octahedral coordination for MG ←  
Go to ToC

### MG:a:3024

|  | Res. type | Atom | Residue | Distance | Occupancy | B-factor | Closest atom(s) to a 1^st^ shell water | EM\_dens. |
| --- | --- | --- | --- | --- | --- | --- | --- | --- |
| **Ion** | | | |  | 1.0 | 41.1 |  | 13.10 |
 1 | A | OP2 | a:2686 | 1.78 | 1.0 | 32.7 |  | 8.86 | 2 | Water 1 | | a:6132 | 2.00 | 1.0 | 33.4 | 2.75 [C:OP2]a:2687 | 7.67 | 3 | Water 2 | | a:8522 | 2.10 | 1.0 | 33.2 |  | 9.58 | 4 | Water 3 | | a:6891 | 2.12 | 1.0 | 36.7 | 2.80 [G:OP2]a:2726 | 9.15 | 5 | Water 4 | | a:5672 | 2.16 | 1.0 | 29.8 | 2.72 [G:OP2]a:2726 | 10.57 | 6 | Water 5 | | a:8832 | 2.38 | 1.0 | 35.7 |  | 7.58 | 7 | C | O3' | a:2685 | 3.63 | 1.0 | 34.6 |  | 8.71 | 8 | Water 6 | | a:9813 | 3.85 | 1.0 | 44.0 |  | 3.63 | 9 | A | OP1 | a:2686 | 3.89 | 1.0 | 34.8 |  | 9.82 | 10 | G | OP2 | a:2726 | 3.95 | 1.0 | 28.6 |  | 12.29 | 11 | Water 7 | | o:239 | 4.22 | 1.0 | 42.5 |  | 7.15 | 12 | C | OP2 | a:2687 | 4.24 | 1.0 | 31.8 |  | 11.30 | 13 | A | O5' | a:2686 | 4.24 | 1.0 | 27.0 |  | 10.65 | 14 | A | OP1 | a:2725 | 4.31 | 1.0 | 32.3 |  | 8.88 | 15 | Water 8 | | a:4716 | 4.33 | 1.0 | 41.7 |  | 7.33 | 16 | Water 9 | | a:4470 | 4.41 | 1.0 | 43.2 | 2.63 [A:OP1]a:2686 | 4.22 | 17 | Water 10 | | a:4304 | 4.48 | 1.0 | 30.4 | 2.61 [C:OP2]a:2687 | 13.18 | 18 | C | O2' | a:2685 | 4.61 | 1.0 | 30.3 |  | 9.01 | 19 | Water 11 | | a:7202 | 4.77 | 1.0 | 32.6 | 2.82 [G:OP2]a:2726 | 7.60 |

  
**1^st^ shell coordination angles and distances (*trans* in "blue"; *cys* in "green"; deviations in "orange"):**   
• 1-2(OP2/W): 94º; 1-3(OP2/W): *98º*; 1-4(OP2/W): 92º; 1-5(OP2/W): 171º; 2-3(W/W): *83º*; 2-4(W/W): *105º*; 2-5(W/W): 91º; 3-4(W/W): *167º*; 3-5(W/W): 91º; 4-5(W/W): *79º*;   
• 1-2(OP2/W): 2.8Å; 1-3(OP2/W): 2.9Å; 1-4(OP2/W): 2.8Å; 1-5(OP2/W): 3.9Å; 2-3(W/W): 2.7Å; 2-4(W/W): *3.3Å*; 2-5(W/W): 3.0Å; 3-4(W/W): 4.2Å; 3-5(W/W): 3.0Å; 4-5(W/W): 2.7Å;   
  
**2.09 ± 0.18: the average coodination distance of the 1^st^ six ligands is Mg^2+^ compatible**  
**10: Number of potential Water contacts**   
**3 : Number of second shell contacts involving water or ions**   
**30%: Percentage of satisfied second shell Water contacts**   
• Contact(s): /O_w_/O_w_/O_w_/O_w_/  
• Contact code: 4O_w_  
• Contact(s) in the exclusion zone: /O_w_/OP2/  
→ Missing octahedral coordination for MG ←  
→ Slightly distorded coord. octahedron (7.2º) ←  
Go to ToC

### MG:a:3025

|  | Res. type | Atom | Residue | Distance | Occupancy | B-factor | Closest atom(s) to a 1^st^ shell water | EM\_dens. |
| --- | --- | --- | --- | --- | --- | --- | --- | --- |
| **Ion** | | | |  | 1.0 | 34.7 |  | 18.36 |
 1 | G | OP2 | a:2690 | 1.94 | 1.0 | 26.1 |  | 12.93 | 2 | Water 1 | | a:7850 | 1.96 | 1.0 | 32.0 |  | 8.78 | 3 | Water 2 | | a:7248 | 2.03 | 1.0 | 31.1 | 2.82 [G:O6]a:2692 // 2.89 [G:OP2]a:2723 | 10.80 | 4 | Water 3 | | a:7399 | 2.10 | 1.0 | 30.8 | 2.83 [U:O4]a:2691 | 10.33 | 5 | Water 4 | | a:8285 | 2.15 | 1.0 | 30.7 |  | 9.88 | 6 | Water 5 | | a:9304 | 2.33 | 1.0 | 32.3 |  | 10.43 | 7 | Water 6 | | a:7333 | 3.94 | 1.0 | 44.4 | 2.83 [G:OP1]a:2723 | 5.07 | 8 | G | OP1 | a:2690 | 4.01 | 1.0 | 30.4 |  | 10.91 | 9 | G | O6 | a:2692 | 4.01 | 1.0 | 29.0 |  | 15.69 | 10 | G | O3' | a:2689 | 4.21 | 1.0 | 26.5 |  | 13.30 | 11 | Water 7 | | a:5087 | 4.23 | 1.0 | 32.7 |  | 10.44 | 12 | G | O5' | a:2690 | 4.27 | 1.0 | 29.3 |  | 12.83 | 13 | G | OP2 | a:2723 | 4.29 | 1.0 | 32.7 |  | 11.36 | 14 | Water 8 | | a:10072 | 4.35 | 1.0 | 37.4 |  | 7.65 | 15 | G | OP1 | a:2723 | 4.36 | 1.0 | 29.1 |  | 11.94 | 16 | Water 9 | | a:6174 | 4.36 | 1.0 | 42.3 | 2.75 [G:OP1]a:2690 | 5.91 | 17 | Water 10 | | a:6631 | 4.37 | 1.0 | 31.7 |  | 9.97 | 18 | G | O5' | a:2723 | 4.69 | 1.0 | 30.1 |  | 11.99 | 19 | U | O4 | a:2691 | 4.79 | 1.0 | 21.4 |  | 17.66 |

  
**1^st^ shell coordination angles and distances (*trans* in "blue"; *cys* in "green"; deviations in "orange"):**   
• 1-2(OP2/W): 95º; 1-3(OP2/W): 95º; 1-4(OP2/W): *102º*; 1-5(OP2/W): 90º; 2-3(W/W): *96º*; 2-4(W/W): *159º*; 2-5(W/W): 88º; 3-4(W/W): 94º; 3-5(W/W): 173º; 4-5(W/W): *79º*;   
• 1-2(OP2/W): 2.9Å; 1-3(OP2/W): 2.9Å; 1-4(OP2/W): 3.1Å; 1-5(OP2/W): 2.9Å; 2-3(W/W): 3.0Å; 2-4(W/W): 4.0Å; 2-5(W/W): 2.9Å; 3-4(W/W): 3.0Å; 3-5(W/W): 4.2Å; 4-5(W/W): 2.7Å;   
  
**2.08 ± 0.13: the average coodination distance of the 1^st^ six ligands is Mg^2+^ compatible**  
**10: Number of potential Water contacts**   
**3 : Number of second shell contacts involving water or ions**   
**30%: Percentage of satisfied second shell Water contacts**   
• Contact(s): /O_w_/O_w_/O_w_/O_w_/OP2/  
• Contact code: O_ph_4O_w_  
• Contact(s) in the exclusion zone: /O_w_/  
→ Missing octahedral coordination for MG ←  
→ Slightly distorded coord. octahedron (7.4º) ←  
Go to ToC

### MG:a:3026

|  | Res. type | Atom | Residue | Distance | Occupancy | B-factor | Closest atom(s) to a 1^st^ shell water | EM\_dens. |
| --- | --- | --- | --- | --- | --- | --- | --- | --- |
| **Ion** | | | |  | 1.0 | 61.3 |  | 2.90 |
 1 | Water 1 | | a:7861 | 3.87 | 1.0 | 44.9 |  | 4.37 | 2 | Water 2 | | a:3557 | 4.13 | 1.0 | 44.2 |  | 4.09 | 3 | G | N7 | a:2739 | 4.23 | 1.0 | 35.9 |  | 6.46 | 4 | A | OP2 | a:2738 | 4.31 | 1.0 | 45.5 |  | 4.14 | 5 | A | OP2 | a:2769 | 4.38 | 1.0 | 37.8 |  | 6.63 | 6 | A | N6 | a:2740 | 4.55 | 1.0 | 31.8 |  | 10.03 | 7 | A | N7 | a:2740 | 4.62 | 1.0 | 35.6 |  | 6.87 | 8 | A | OP1 | a:2769 | 4.64 | 1.0 | 36.9 |  | 4.44 | 9 | G | OP2 | a:2739 | 4.67 | 1.0 | 48.7 |  | 2.58 |

  
• • **4.25 ± 0.21: the average coodination distance of the 1^st^ six ligands could correspond to an hexahydrated Mg^2+^.**  
**4: Number of potential Water contacts**   
**0 : Number of second shell contacts involving water or ions**   
**0%: Percentage of satisfied second shell Water contacts**   
• Contact code: → No contacts ←   
→ Firts MG coordination shell missing ←  
→2Fo-Fc density between 1.0 and 4.0 !   
Go to ToC

### MG:a:3027

|  | Res. type | Atom | Residue | Distance | Occupancy | B-factor | Closest atom(s) to a 1^st^ shell water | EM\_dens. |
| --- | --- | --- | --- | --- | --- | --- | --- | --- |
| **Ion** | | | |  | 1.0 | 55.9 |  | 5.34 |
 1 | Water 1 | | a:8732 | 1.91 | 1.0 | 46.8 |  | 3.52 | 2 | A | OP2 | a:2752 | 1.92 | 1.0 | 49.5 |  | 6.29 | 3 | Water 2 | | a:6979 | 2.06 | 1.0 | 48.3 | 2.87 [A:N7]a:2757 | 4.11 | 4 | U | O4 | a:2758 | 2.13 | 1.0 | 45.5 |  | 5.26 | 5 | A | OP1 | a:2752 | 3.63 | 1.0 | 52.7 |  | 2.92 | 6 | G | O3' | a:2751 | 4.19 | 1.0 | 52.1 |  | 4.02 | 7 | A | O5' | a:2752 | 4.27 | 1.0 | 50.9 |  | 4.34 | 8 | A | N7 | a:2757 | 4.37 | 1.0 | 45.2 |  | 5.30 | 9 | A | OP2 | a:2757 | 4.59 | 1.0 | 49.9 |  | 3.84 | 10 | G | N7 | a:2751 | 4.61 | 1.0 | 43.6 |  | 6.55 |

  
**1^st^ shell coordination angles and distances (*trans* in "blue"; *cys* in "green"; deviations in "orange"):**   
• 1-2(W/OP2): *169º*; 1-3(W/W): *79º*; 1-4(W/O4): 94º; 2-3(OP2/W): 93º; 2-4(OP2/O4): 93º; 3-4(W/O4): *84º*;   
• 1-2(W/OP2): 3.8Å; 1-3(W/W): 2.5Å; 1-4(W/O4): 3.0Å; 2-3(OP2/W): 2.9Å; 2-4(OP2/O4): 2.9Å; 3-4(W/O4): 2.8Å;   
  
**2.64 ± 0.92: the average coodination distance of the 1^st^ six ligands is K^+^ compatible**  
**4: Number of potential Water contacts**   
**1 : Number of second shell contacts involving water or ions**   
**25%: Percentage of satisfied second shell Water contacts**   
• Contact(s): /O4/O_w_/O_w_/OP2/  
• Contact code: O_ph_O_b_2O_w_  
→ Missing octahedral coordination for MG ←  
→ Slightly distorded coord. octahedron (6.3º) ←  
Go to ToC

### MG:a:3028

|  | Res. type | Atom | Residue | Distance | Occupancy | B-factor | Closest atom(s) to a 1^st^ shell water | EM\_dens. |
| --- | --- | --- | --- | --- | --- | --- | --- | --- |
| **Ion** | | | |  | 1.0 | 47.5 |  | 11.37 |
 1 | Water 1 | | m:216 | 1.92 | 1.0 | 42.6 | 2.74 [SER:OG]m:6 | 3.97 | 2 | Water 2 | | a:8375 | 2.18 | 1.0 | 39.5 | 2.92 [G:OP1]a:2842 | 6.30 | 3 | Water 3 | | a:3476 | 2.28 | 1.0 | 34.4 | 2.43 [C:OP1]a:2878 | 9.88 | 4 | Water 4 | | a:7888 | 3.90 | 1.0 | 38.4 | 2.87 [G:OP1]a:2823 | 6.47 | 5 | C | OP1 | a:2878 | 4.04 | 1.0 | 40.4 |  | 7.54 | 6 | G | OP1 | a:2842 | 4.06 | 1.0 | 38.7 |  | 5.53 | 7 | G | OP1 | a:2823 | 4.16 | 1.0 | 30.8 |  | 8.46 | 8 | SER | OG | m:6 | 4.43 | 1.0 | 41.0 |  | 4.57 | 9 | A | OP1 | a:2824 | 4.47 | 1.0 | 35.0 |  | 12.14 | 10 | ARG | O | m:4 | 4.50 | 1.0 | 28.2 |  | 12.63 | 11 | A | O3' | a:2877 | 4.54 | 1.0 | 36.6 |  | 10.20 | 12 | C | OP2 | a:2878 | 4.60 | 1.0 | 31.5 |  | 8.73 |

  
**1^st^ shell coordination angles and distances (*trans* in "blue"; *cys* in "green"; deviations in "orange"):**   
• 1-2(W/W): 86º; 1-3(W/W): 88º; 2-3(W/W): 92º;   
• 1-2(W/W): 2.8Å; 1-3(W/W): 2.9Å; 2-3(W/W): *3.2Å*;   
  
**3.06 ± 0.94: the average coodination distance of the 1^st^ six ligands is K^+^ compatible**  
**8: Number of potential Water contacts**   
**4 : Number of second shell contacts involving water or ions**   
**50%: Percentage of satisfied second shell Water contacts**   
• Contact(s): /O_w_/O_w_/O_w_/  
• Contact code: 3O_w_  
→ Missing octahedral coordination for MG ←  
Go to ToC

### MG:a:3029

|  | Res. type | Atom | Residue | Distance | Occupancy | B-factor | Closest atom(s) to a 1^st^ shell water | EM\_dens. |
| --- | --- | --- | --- | --- | --- | --- | --- | --- |
| **Ion** | | | |  | 1.0 | 40.0 |  | 9.04 |
 1 | U | O4 | a:2851 | 2.79 | 1.0 | 23.9 |  | 15.48 | 2 | Water 1 | | a:8265 | 2.83 | 1.0 | 32.8 | 2.91 [G:OP2]a:2850 | 9.08 | 3 | Water 2 | | a:4602 | 3.07 | 1.0 | 37.7 | 2.64 [G:OP2]a:2850 | 7.73 | 4 | G | O6 | a:2852 | 3.38 | 1.0 | 36.5 |  | 7.91 | 5 | Water 3 | | a:5730 | 3.53 | 1.0 | 39.6 | 2.73 [G:O6]a:2850 // 2.91 [K:K]a:3257 | 5.18 | 6 | G | N7 | a:2850 | 3.58 | 1.0 | 19.3 |  | 14.97 | 7 | Water 4 | | a:8069 | 3.65 | 1.0 | 40.0 | 2.89 [G:O6]a:2852 | 5.80 | 8 | Water 5 | | a:5015 | 3.75 | 1.0 | 43.8 |  | 4.83 | 9 | **K** | K | a:3257 | 4.01 | 1.0 | 30.0 |  | 21.39 | 10 | G | OP2 | a:2850 | 4.48 | 1.0 | 30.1 |  | 11.21 | 11 | G | O6 | a:2850 | 4.63 | 1.0 | 25.6 |  | 14.84 |

  
• • **3.20 ± 0.32: the average coodination distance of the 1^st^ six ligands is K^+^ compatible**  
**6: Number of potential Water contacts**   
**4 : Number of second shell contacts involving water or ions**   
**67%: Percentage of satisfied second shell Water contacts**   
• Contact code: → No contacts ←   
• Contact(s) in the exclusion zone: /O4/O6/O_w_/O_w_/  
→ Consider K in place of MG (1 valid contact(s) for K vs. 0 for MG ←)  
Go to ToC

### MG:a:3030

|  | Res. type | Atom | Residue | Distance | Occupancy | B-factor | Closest atom(s) to a 1^st^ shell water | EM\_dens. |
| --- | --- | --- | --- | --- | --- | --- | --- | --- |
| **Ion** | | | |  | 1.0 | 44.0 |  | 8.75 |
 1 | Water 1 | | a:9912 | 1.95 | 1.0 | 39.3 |  | 7.14 | 2 | Water 2 | | a:7488 | 1.98 | 1.0 | 37.1 | 2.84 [G:OP1]a:2697 | 7.38 | 3 | Water 3 | | a:6420 | 2.10 | 1.0 | 33.3 | 2.77 [A:O2']a:2854 | 8.90 | 4 | Water 4 | | a:9510 | 2.13 | 1.0 | 39.4 |  | 6.51 | 5 | Water 5 | | a:3781 | 2.20 | 1.0 | 35.0 | 2.53 [G:OP2]a:2698 | 6.97 | 6 | Water 6 | | a:4625 | 2.21 | 1.0 | 36.6 | 2.64 [U:OP2]a:2711 | 5.07 | 7 | U | OP1 | a:2711 | 4.08 | 1.0 | 40.5 |  | 6.65 | 8 | G | OP1 | a:2697 | 4.10 | 1.0 | 30.2 |  | 8.79 | 9 | G | OP2 | a:2698 | 4.12 | 1.0 | 30.8 |  | 10.24 | 10 | Water 7 | | a:6583 | 4.22 | 1.0 | 40.0 | 2.78 [U:OP1]a:2711 | 4.57 | 11 | Water 8 | | a:3648 | 4.26 | 1.0 | 36.9 | 2.50 [G:OP1]a:2697 | 6.82 | 12 | U | OP2 | a:2711 | 4.30 | 1.0 | 32.5 |  | 9.26 | 13 | A | O2' | a:2854 | 4.32 | 1.0 | 36.1 |  | 9.67 | 14 | G | OP1 | a:2698 | 4.55 | 1.0 | 37.2 |  | 7.45 | 15 | Water 9 | | a:9297 | 4.57 | 1.0 | 47.7 |  | 3.22 |

  
**1^st^ shell coordination angles and distances (*trans* in "blue"; *cys* in "green"; deviations in "orange"):**   
• 1-2(W/W): 88º; 1-3(W/W): 87º; 1-4(W/W): *96º*; 1-5(W/W): *95º*; 1-6(W/W): 174º; 2-3(W/W): *99º*; 2-4(W/W): 173º; 2-5(W/W): *84º*; 2-6(W/W): 91º; 3-4(W/W): 87º; 3-5(W/W): 176º; 3-6(W/W): 87º; 4-5(W/W): 89º; 4-6(W/W): 86º; 5-6(W/W): 90º;   
• 1-2(W/W): 2.7Å; 1-3(W/W): 2.8Å; 1-4(W/W): 3.0Å; 1-5(W/W): 3.1Å; 1-6(W/W): 4.2Å; 2-3(W/W): 3.1Å; 2-4(W/W): 4.1Å; 2-5(W/W): 2.8Å; 2-6(W/W): 3.0Å; 3-4(W/W): 2.9Å; 3-5(W/W): *4.3Å*; 3-6(W/W): 3.0Å; 4-5(W/W): 3.0Å; 4-6(W/W): 3.0Å; 5-6(W/W): 3.1Å;   
  
**2.09 ± 0.10: the average coodination distance of the 1^st^ six ligands is Mg^2+^ compatible**  
**12: Number of potential Water contacts**   
**4 : Number of second shell contacts involving water or ions**   
**33%: Percentage of satisfied second shell Water contacts**   
• Contact(s): /O_w_/O_w_/O_w_/O_w_/O_w_/O_w_/  
• Contact code: 6O_w_  
Ooctahedral coordination (4.0º)   
Go to ToC

### MG:a:3031

|  | Res. type | Atom | Residue | Distance | Occupancy | B-factor | Closest atom(s) to a 1^st^ shell water | EM\_dens. |
| --- | --- | --- | --- | --- | --- | --- | --- | --- |
| **Ion** | | | |  | 1.0 | 24.6 |  | 23.53 |
 1 | G | O6 | a:376 | 2.71 | 1.0 | 32.0 |  | 12.24 | 2 | Water 1 | | a:7375 | 2.84 | 1.0 | 29.8 |  | 10.38 | 3 | G | O6 | a:375 | 2.88 | 1.0 | 32.9 |  | 10.18 | 4 | Water 2 | | a:8125 | 2.92 | 1.0 | 34.2 | 1.88 [MG:MG]a:3163 | 8.22 | 5 | Water 3 | | a:4816 | 2.93 | 1.0 | 37.2 | 2.66 [A:OP2]a:374 | 6.96 | 6 | Water 4 | | a:5670 | 3.06 | 1.0 | 30.7 | 2.37 [MG:MG]a:3163 | 11.01 | 7 | U | O4 | a:399 | 3.21 | 1.0 | 34.0 |  | 10.06 | 8 | Water 5 | | a:10254 | 3.49 | 1.0 | 38.5 | 2.12 [MG:MG]a:3163 | 7.27 | 9 | **MG** | MG | a:3163 | 3.63 | 1.0 | 40.4 |  | 12.83 | 10 | G | N7 | a:376 | 4.11 | 1.0 | 27.3 |  | 12.16 | 11 | Water 6 | | a:7950 | 4.24 | 1.0 | 30.1 |  | 9.34 | 12 | G | N7 | a:375 | 4.27 | 1.0 | 28.4 |  | 10.79 | 13 | A | OP2 | a:374 | 4.56 | 1.0 | 36.9 |  | 8.38 |

  
• • **2.89 ± 0.11: the average coodination distance of the 1^st^ six ligands is K^+^ compatible**  
**8: Number of potential Water contacts**   
**3 : Number of second shell contacts involving water or ions**   
**38%: Percentage of satisfied second shell Water contacts**   
• Contact code: → No contacts ←   
• Contact(s) in the exclusion zone: /O4/O6/O6/O_w_/O_w_/O_w_/O_w_/  
→ Consider K in place of MG (2 valid contact(s) for K vs. 0 for MG ←)  
Go to ToC

### MG:a:3032

|  | Res. type | Atom | Residue | Distance | Occupancy | B-factor | Closest atom(s) to a 1^st^ shell water | EM\_dens. |
| --- | --- | --- | --- | --- | --- | --- | --- | --- |
| **Ion** | | | |  | 1.0 | 25.8 |  | 19.71 |
 1 | G | O6 | a:1204 | 2.67 | 1.0 | 31.0 |  | 12.78 | 2 | U | O4 | a:1244 | 2.69 | 1.0 | 31.9 |  | 10.65 | 3 | U | O4 | a:1205 | 2.72 | 1.0 | 33.2 |  | 8.25 | 4 | Water 1 | | a:8318 | 2.94 | 1.0 | 31.4 | 2.04 [MG:MG]a:3210 | 10.36 | 5 | A | OP2 | a:1243 | 3.01 | 1.0 | 45.0 |  | 6.57 | 6 | Water 2 | | a:8198 | 3.09 | 1.0 | 38.6 | 2.05 [MG:MG]a:3210 | 7.89 | 7 | Water 3 | | a:6595 | 3.13 | 1.0 | 42.4 | 2.78 [A:OP1]a:1243 | 5.06 | 8 | Water 4 | | a:5820 | 3.70 | 1.0 | 44.3 |  | 4.94 | 9 | **MG** | MG | a:3210 | 4.02 | 1.0 | 43.2 |  | 11.27 | 10 | Water 5 | | a:4352 | 4.24 | 1.0 | 32.1 | 1.99 [MG:MG]a:3210 | 9.21 | 11 | A | N1 | a:1206 | 4.25 | 1.0 | 29.9 |  | 8.96 | 12 | G | N7 | a:1204 | 4.28 | 1.0 | 26.4 |  | 12.76 | 13 | A | OP1 | a:1243 | 4.35 | 1.0 | 40.5 |  | 6.67 | 14 | Water 6 | | a:10060 | 4.75 | 1.0 | 41.2 | 2.25 [MG:MG]a:3210 | 5.37 |

  
• • **2.85 ± 0.17: the average coodination distance of the 1^st^ six ligands is K^+^ compatible**  
**4: Number of potential Water contacts**   
**2 : Number of second shell contacts involving water or ions**   
**50%: Percentage of satisfied second shell Water contacts**   
• Contact code: → No contacts ←   
• Contact(s) in the exclusion zone: /O4/O4/O6/O_w_/O_w_/O_w_/OP2/  
→ Consider K in place of MG (4 valid contact(s) for K vs. 0 for MG ←)  
Go to ToC

### MG:a:3033

|  | Res. type | Atom | Residue | Distance | Occupancy | B-factor | Closest atom(s) to a 1^st^ shell water | EM\_dens. |
| --- | --- | --- | --- | --- | --- | --- | --- | --- |
| **Ion** | | | |  | 1.0 | 51.1 |  | 9.12 |
 1 | G | OP1 | a:1949 | 1.82 | 1.0 | 42.2 |  | 7.12 | 2 | Water 1 | | a:4693 | 1.85 | 1.0 | 40.1 | 2.65 [U:O4]a:1950 | 6.34 | 3 | Water 2 | | a:7637 | 2.03 | 1.0 | 36.9 | 2.85 [G:N7]a:1949 | 5.98 | 4 | Water 3 | | a:6419 | 2.27 | 1.0 | 36.5 | 2.77 [U:OP1]a:1960 | 7.67 | 5 | Water 4 | | a:10027 | 2.36 | 1.0 | 45.5 |  | 4.54 | 6 | Water 5 | | a:3736 | 2.37 | 1.0 | 41.7 | 2.52 [U:OP2]a:1960 | 5.19 | 7 | Water 6 | | a:10045 | 3.66 | 1.0 | 41.8 |  | 6.71 | 8 | G | OP2 | a:1949 | 3.72 | 1.0 | 34.9 |  | 7.54 | 9 | Water 7 | | a:4627 | 3.95 | 1.0 | 36.5 |  | 7.58 | 10 | U | O3' | a:1948 | 3.99 | 1.0 | 36.0 |  | 7.89 | 11 | U | OP2 | a:1960 | 4.18 | 1.0 | 36.7 |  | 8.63 | 12 | G | O5' | a:1949 | 4.21 | 1.0 | 37.2 |  | 7.17 | 13 | G | N7 | a:1949 | 4.29 | 1.0 | 30.3 |  | 10.61 | 14 | Water 8 | | a:3803 | 4.37 | 1.0 | 45.3 | 2.54 [G:OP2]a:1949 | 7.94 | 15 | U | O4 | a:1950 | 4.41 | 1.0 | 37.3 |  | 8.88 | 16 | U | OP1 | a:1960 | 4.42 | 1.0 | 34.2 |  | 10.45 | 17 | Water 9 | | a:6864 | 4.46 | 1.0 | 43.8 |  | 4.60 | 18 | Water 10 | | a:9593 | 4.63 | 1.0 | 42.8 |  | 4.37 | 19 | Water 11 | | a:9678 | 4.73 | 1.0 | 41.2 |  | 5.83 | 20 | Water 12 | | a:4055 | 4.78 | 1.0 | 43.1 |  | 5.40 |

  
**1^st^ shell coordination angles and distances (*trans* in "blue"; *cys* in "green"; deviations in "orange"):**   
• 1-2(OP1/W): *103º*; 1-3(OP1/W): *97º*; 1-4(OP1/W): 93º; 2-3(W/W): *82º*; 2-4(W/W): *157º*; 3-4(W/W): *80º*;   
• 1-2(OP1/W): 2.9Å; 1-3(OP1/W): 2.9Å; 1-4(OP1/W): 3.0Å; 2-3(W/W): 2.5Å; 2-4(W/W): 4.0Å; 3-4(W/W): 2.8Å;   
  
**2.12 ± 0.23: the average coodination distance of the 1^st^ six ligands is Mg^2+^ compatible**  
**10: Number of potential Water contacts**   
**4 : Number of second shell contacts involving water or ions**   
**40%: Percentage of satisfied second shell Water contacts**   
• Contact(s): /O_w_/O_w_/  
• Contact code: 2O_w_  
• Contact(s) in the exclusion zone: /O_w_/O_w_/O_w_/OP1/  
→ Missing octahedral coordination for MG ←  
→→→ Distorded coord. octahedron (10.8º) ←←←  
Go to ToC

### MG:a:3034

|  | Res. type | Atom | Residue | Distance | Occupancy | B-factor | Closest atom(s) to a 1^st^ shell water | EM\_dens. |
| --- | --- | --- | --- | --- | --- | --- | --- | --- |
| **Ion** | | | |  | 1.0 | 36.8 |  | 18.05 |
 1 | Water 1 | | a:6988 | 1.90 | 1.0 | 30.7 | 2.80 [G:O6]a:973 | 11.01 | 2 | Water 2 | | a:9907 | 1.95 | 1.0 | 28.8 |  | 10.78 | 3 | Water 3 | | a:10111 | 2.14 | 1.0 | 35.8 |  | 9.96 | 4 | Water 4 | | a:10033 | 2.18 | 1.0 | 29.4 |  | 10.70 | 5 | Water 5 | | a:10077 | 2.27 | 1.0 | 33.2 |  | 10.96 | 6 | Water 6 | | a:7250 | 2.34 | 1.0 | 26.4 | 2.82 [A:N1]a:974 | 12.66 | 7 | Water 7 | | a:5759 | 4.18 | 1.0 | 32.1 |  | 9.13 | 8 | Water 8 | | a:10231 | 4.22 | 1.0 | 43.6 |  | 7.42 | 9 | U | O4 | a:972 | 4.26 | 1.0 | 21.9 |  | 16.20 | 10 | A | N1 | a:974 | 4.27 | 1.0 | 16.1 |  | 17.63 | 11 | Water 9 | | a:5182 | 4.28 | 1.0 | 46.6 |  | 4.81 | 12 | Water 10 | | a:7419 | 4.38 | 1.0 | 34.3 |  | 9.35 | 13 | Water 11 | | a:7929 | 4.42 | 1.0 | 33.5 |  | 8.14 | 14 | G | O6 | a:973 | 4.43 | 1.0 | 22.5 |  | 17.67 | 15 | Water 12 | | a:6304 | 4.46 | 1.0 | 30.1 |  | 12.55 | 16 | Water 13 | | a:10289 | 4.48 | 1.0 | 42.0 |  | 4.60 | 17 | Water 14 | | a:6973 | 4.49 | 1.0 | 29.5 |  | 9.53 | 18 | Water 15 | | a:7178 | 4.50 | 1.0 | 31.7 |  | 12.30 | 19 | Water 16 | | a:4091 | 4.70 | 1.0 | 37.5 |  | 5.68 |

  
**1^st^ shell coordination angles and distances (*trans* in "blue"; *cys* in "green"; deviations in "orange"):**   
• 1-2(W/W): 92º; 1-3(W/W): *96º*; 1-4(W/W): 92º; 1-5(W/W): 172º; 2-3(W/W): *170º*; 2-4(W/W): 91º; 2-5(W/W): 93º; 3-4(W/W): *82º*; 3-5(W/W): *79º*; 4-5(W/W): 93º;   
• 1-2(W/W): 2.8Å; 1-3(W/W): 3.0Å; 1-4(W/W): 2.9Å; 1-5(W/W): 4.2Å; 2-3(W/W): 4.1Å; 2-4(W/W): 3.0Å; 2-5(W/W): 3.1Å; 3-4(W/W): 2.8Å; 3-5(W/W): 2.8Å; 4-5(W/W): *3.2Å*;   
  
**2.13 ± 0.16: the average coodination distance of the 1^st^ six ligands is Mg^2+^ compatible**  
**12: Number of potential Water contacts**   
**2 : Number of second shell contacts involving water or ions**   
**17%: Percentage of satisfied second shell Water contacts**   
• Contact(s): /O_w_/O_w_/O_w_/O_w_/O_w_/  
• Contact code: 5O_w_  
• Contact(s) in the exclusion zone: /O_w_/  
→ Missing octahedral coordination for MG ←  
→ Slightly distorded coord. octahedron (5.4º) ←  
Go to ToC

### MG:a:3035

|  | Res. type | Atom | Residue | Distance | Occupancy | B-factor | Closest atom(s) to a 1^st^ shell water | EM\_dens. |
| --- | --- | --- | --- | --- | --- | --- | --- | --- |
| **Ion** | | | |  | 1.0 | 46.5 |  | 10.90 |
 1 | Water 1 | | a:10299 | 1.94 | 1.0 | 41.3 |  | 6.39 | 2 | Water 2 | | a:5101 | 2.06 | 1.0 | 34.8 | 2.68 [G:O6]a:379 // 2.68 [U:O4]a:395 | 7.66 | 3 | Water 3 | | a:6062 | 2.07 | 1.0 | 43.8 | 2.75 [G:O6]a:380 | 7.44 | 4 | Water 4 | | a:7947 | 2.19 | 1.0 | 29.9 | 2.88 [G:N7]a:379 | 9.87 | 5 | Water 5 | | a:9956 | 2.23 | 1.0 | 33.0 |  | 8.20 | 6 | Water 6 | | a:10039 | 2.48 | 1.0 | 39.8 |  | 6.96 | 7 | Water 7 | | a:6671 | 3.80 | 1.0 | 31.5 |  | 9.37 | 8 | Water 8 | | a:5983 | 4.01 | 1.0 | 40.0 |  | 5.26 | 9 | Water 9 | | a:10253 | 4.18 | 1.0 | 46.8 |  | 3.90 | 10 | G | O6 | a:379 | 4.21 | 1.0 | 23.4 |  | 13.70 | 11 | G | O6 | a:380 | 4.28 | 1.0 | 29.6 |  | 13.86 | 12 | Water 10 | | a:5295 | 4.29 | 1.0 | 34.2 |  | 10.30 | 13 | U | O4 | a:395 | 4.45 | 1.0 | 37.9 |  | 7.94 | 14 | G | N7 | a:379 | 4.60 | 1.0 | 22.9 |  | 13.42 | 15 | Water 11 | | a:8247 | 4.60 | 1.0 | 37.4 |  | 8.56 | 16 | Water 12 | | a:9267 | 4.67 | 1.0 | 33.9 |  | 7.12 |

  
**1^st^ shell coordination angles and distances (*trans* in "blue"; *cys* in "green"; deviations in "orange"):**   
• 1-2(W/W): *169º*; 1-3(W/W): 93º; 1-4(W/W): 91º; 1-5(W/W): 90º; 2-3(W/W): *81º*; 2-4(W/W): *98º*; 2-5(W/W): *95º*; 3-4(W/W): *97º*; 3-5(W/W): 171º; 4-5(W/W): 91º;   
• 1-2(W/W): 4.0Å; 1-3(W/W): 2.9Å; 1-4(W/W): 3.0Å; 1-5(W/W): 2.9Å; 2-3(W/W): 2.7Å; 2-4(W/W): *3.2Å*; 2-5(W/W): 3.2Å; 3-4(W/W): 3.2Å; 3-5(W/W): *4.3Å*; 4-5(W/W): 3.2Å;   
  
**2.16 ± 0.17: the average coodination distance of the 1^st^ six ligands is Mg^2+^ compatible**  
**12: Number of potential Water contacts**   
**4 : Number of second shell contacts involving water or ions**   
**33%: Percentage of satisfied second shell Water contacts**   
• Contact(s): /O_w_/O_w_/O_w_/O_w_/O_w_/  
• Contact code: 5O_w_  
• Contact(s) in the exclusion zone: /O_w_/  
→ Missing octahedral coordination for MG ←  
→ Slightly distorded coord. octahedron (5.5º) ←  
Go to ToC

### MG:a:3036

|  | Res. type | Atom | Residue | Distance | Occupancy | B-factor | Closest atom(s) to a 1^st^ shell water | EM\_dens. |
| --- | --- | --- | --- | --- | --- | --- | --- | --- |
| **Ion** | | | |  | 1.0 | 48.8 |  | 8.74 |
 1 | Water 1 | | a:9355 | 1.95 | 1.0 | 38.7 |  | 5.98 | 2 | Water 2 | | a:8580 | 1.96 | 1.0 | 36.4 | 2.94 [G:N7]a:776 | 7.09 | 3 | Water 3 | | a:9605 | 2.08 | 1.0 | 38.9 |  | 6.83 | 4 | Water 4 | | a:4511 | 2.13 | 1.0 | 34.8 | 2.63 [G:O6]a:776 | 7.37 | 5 | Water 5 | | a:10132 | 2.20 | 1.0 | 41.3 |  | 8.35 | 6 | A | OP2 | a:687 | 2.25 | 1.0 | 25.5 |  | 12.45 | 7 | G | O6 | a:776 | 4.02 | 1.0 | 21.6 |  | 15.70 | 8 | Water 6 | | a:4505 | 4.02 | 1.0 | 33.7 |  | 10.51 | 9 | G | N7 | a:776 | 4.20 | 1.0 | 17.1 |  | 15.80 | 10 | Water 7 | | a:9861 | 4.26 | 1.0 | 40.3 |  | 4.50 | 11 | Water 8 | | 1:113 | 4.35 | 1.0 | 41.9 |  | 3.81 | 12 | A | O5' | a:687 | 4.43 | 1.0 | 31.9 |  | 7.69 | 13 | Water 9 | | 1:149 | 4.54 | 1.0 | 43.6 |  | 3.44 | 14 | G | O3' | a:686 | 4.58 | 1.0 | 24.2 |  | 14.40 | 15 | A | OP1 | a:687 | 4.61 | 1.0 | 36.2 |  | 8.36 |

  
**1^st^ shell coordination angles and distances (*trans* in "blue"; *cys* in "green"; deviations in "orange"):**   
• 1-2(W/W): *99º*; 1-3(W/W): 89º; 1-4(W/W): 173º; 1-5(W/W): 91º; 1-6(W/OP2): 93º; 2-3(W/W): *166º*; 2-4(W/W): 85º; 2-5(W/W): 90º; 2-6(W/OP2): *97º*; 3-4(W/W): 88º; 3-5(W/W): *79º*; 3-6(W/OP2): 93º; 4-5(W/W): *95º*; 4-6(W/OP2): *81º*; 5-6(W/OP2): 172º;   
• 1-2(W/W): 3.0Å; 1-3(W/W): 2.8Å; 1-4(W/W): 4.1Å; 1-5(W/W): 3.0Å; 1-6(W/OP2): 3.1Å; 2-3(W/W): 4.0Å; 2-4(W/W): 2.8Å; 2-5(W/W): 2.9Å; 2-6(W/OP2): 3.2Å; 3-4(W/W): 2.9Å; 3-5(W/W): 2.7Å; 3-6(W/OP2): 3.2Å; 4-5(W/W): 3.2Å; 4-6(W/OP2): 2.8Å; 5-6(W/OP2): *4.4Å*;   
  
**2.10 ± 0.11: the average coodination distance of the 1^st^ six ligands is Mg^2+^ compatible**  
**10: Number of potential Water contacts**   
**2 : Number of second shell contacts involving water or ions**   
**20%: Percentage of satisfied second shell Water contacts**   
• Contact(s): /O_w_/O_w_/O_w_/O_w_/O_w_/OP2/  
• Contact code: O_ph_5O_w_  
→ Slightly distorded coord. octahedron (5.8º) ←  
Go to ToC

### MG:a:3037

|  | Res. type | Atom | Residue | Distance | Occupancy | B-factor | Closest atom(s) to a 1^st^ shell water | EM\_dens. |
| --- | --- | --- | --- | --- | --- | --- | --- | --- |
| **Ion** | | | |  | 1.0 | 53.5 |  | 9.07 |
 1 | Water 1 | | a:5508 | 1.99 | 1.0 | 33.0 | 2.71 [G:O6]a:979 | 8.42 | 2 | Water 2 | | a:4415 | 2.20 | 1.0 | 30.1 | 2.62 [G:N7]a:980 | 11.06 | 3 | Water 3 | | a:9671 | 2.21 | 1.0 | 37.6 |  | 4.68 | 4 | Water 4 | | a:6133 | 4.00 | 1.0 | 32.0 |  | 9.70 | 5 | Water 5 | | a:8849 | 4.03 | 1.0 | 29.4 | 2.98 [G:OP2]a:978 | 11.91 | 6 | G | O6 | a:980 | 4.10 | 1.0 | 30.0 |  | 15.91 | 7 | Water 6 | | a:6384 | 4.11 | 1.0 | 30.7 |  | 10.37 | 8 | G | OP2 | a:978 | 4.17 | 1.0 | 31.4 |  | 11.57 | 9 | G | O6 | a:979 | 4.21 | 1.0 | 26.3 |  | 15.52 | 10 | G | N7 | a:979 | 4.22 | 1.0 | 22.3 |  | 13.09 | 11 | Water 7 | | a:8561 | 4.32 | 1.0 | 39.1 |  | 6.94 | 12 | G | N7 | a:980 | 4.35 | 1.0 | 21.1 |  | 14.35 | 13 | Water 8 | | a:5494 | 4.62 | 1.0 | 35.1 |  | 5.35 | 14 | Water 9 | | a:6450 | 4.75 | 1.0 | 31.8 |  | 8.88 | 15 | Water 10 | | a:6265 | 4.80 | 1.0 | 41.8 |  | 7.08 |

  
**1^st^ shell coordination angles and distances (*trans* in "blue"; *cys* in "green"; deviations in "orange"):**   
• 1-2(W/W): *95º*; 1-3(W/W): 90º; 2-3(W/W): 171º;   
• 1-2(W/W): 3.1Å; 1-3(W/W): 3.0Å; 2-3(W/W): *4.4Å*;   
  
**3.09 ± 0.96: the average coodination distance of the 1^st^ six ligands is K^+^ compatible**  
**10: Number of potential Water contacts**   
**3 : Number of second shell contacts involving water or ions**   
**30%: Percentage of satisfied second shell Water contacts**   
• Contact(s): /O_w_/O_w_/O_w_/  
• Contact code: 3O_w_  
→ Missing octahedral coordination for MG ←  
Go to ToC

### MG:a:3038

|  | Res. type | Atom | Residue | Distance | Occupancy | B-factor | Closest atom(s) to a 1^st^ shell water | EM\_dens. |
| --- | --- | --- | --- | --- | --- | --- | --- | --- |
| **Ion** | | | |  | 1.0 | 48.1 |  | 14.16 |
 1 | Water 1 | | a:9903 | 2.10 | 1.0 | 38.4 |  | 8.56 | 2 | Water 2 | | a:3694 | 2.16 | 1.0 | 34.4 | 2.51 [G:OP1]a:580 // 2.86 [G:OP2]a:581 | 10.79 | 3 | Water 3 | | a:10071 | 2.18 | 1.0 | 36.0 |  | 8.45 | 4 | Water 4 | | a:9613 | 2.22 | 1.0 | 41.1 |  | 7.17 | 5 | Water 5 | | a:6669 | 2.38 | 1.0 | 35.9 | 2.78 [U:OP2]a:2021 | 8.04 | 6 | Water 6 | | a:7245 | 2.56 | 1.0 | 33.6 | 2.82 [U:O2']a:1257 | 5.46 | 7 | Water 7 | | a:4565 | 3.82 | 1.0 | 27.4 | 2.64 [G:OP2]a:581 // 2.71 [U:OP2]a:2021 | 10.77 | 8 | U | O2' | a:1257 | 4.02 | 1.0 | 27.4 |  | 14.73 | 9 | Water 8 | | a:10261 | 4.10 | 1.0 | 43.8 |  | 3.94 | 10 | Water 9 | | a:8727 | 4.24 | 1.0 | 21.2 |  | 17.77 | 11 | G | OP1 | a:580 | 4.31 | 1.0 | 25.2 |  | 15.33 | 12 | Water 10 | | a:7340 | 4.34 | 1.0 | 32.8 |  | 8.98 | 13 | G | OP2 | a:581 | 4.42 | 1.0 | 27.0 |  | 14.88 | 14 | U | OP2 | a:2021 | 4.42 | 1.0 | 22.6 |  | 15.04 | 15 | U | OP1 | a:2020 | 4.44 | 1.0 | 31.6 |  | 9.82 | 16 | Water 11 | | a:9314 | 4.62 | 1.0 | 37.2 |  | 7.17 | 17 | Water 12 | | a:5744 | 4.65 | 1.0 | 33.1 |  | 9.55 | 18 | Water 13 | | a:10209 | 4.75 | 1.0 | 33.0 |  | 9.21 |

  
**1^st^ shell coordination angles and distances (*trans* in "blue"; *cys* in "green"; deviations in "orange"):**   
• 1-2(W/W): *81º*; 1-3(W/W): 86º; 1-4(W/W): *83º*; 2-3(W/W): 92º; 2-4(W/W): *163º*; 3-4(W/W): 89º;   
• 1-2(W/W): 2.8Å; 1-3(W/W): 2.9Å; 1-4(W/W): 2.9Å; 2-3(W/W): 3.1Å; 2-4(W/W): *4.3Å*; 3-4(W/W): 3.1Å;   
  
**2.27 ± 0.16: the average coodination distance of the 1^st^ six ligands is Mg^2+^ compatible**  
**12: Number of potential Water contacts**   
**4 : Number of second shell contacts involving water or ions**   
**33%: Percentage of satisfied second shell Water contacts**   
• Contact(s): /O_w_/O_w_/O_w_/O_w_/  
• Contact code: 4O_w_  
• Contact(s) in the exclusion zone: /O_w_/O_w_/  
→ Missing octahedral coordination for MG ←  
→ Slightly distorded coord. octahedron (6.7º) ←  
Go to ToC

### MG:a:3039

|  | Res. type | Atom | Residue | Distance | Occupancy | B-factor | Closest atom(s) to a 1^st^ shell water | EM\_dens. |
| --- | --- | --- | --- | --- | --- | --- | --- | --- |
| **Ion** | | | |  | 1.0 | 57.3 |  | 5.96 |
 1 | Water 1 | | a:5300 | 1.80 | 1.0 | 41.9 | 1.93 [MG:MG]a:3040 | 4.60 | 2 | Water 2 | | a:9908 | 1.86 | 1.0 | 39.2 |  | 8.44 | 3 | Water 3 | | a:5035 | 2.03 | 1.0 | 37.6 | 2.07 [MG:MG]a:3040 | 7.56 | 4 | Water 4 | | a:5719 | 2.22 | 1.0 | 33.3 | 2.01 [MG:MG]a:3040 | 8.01 | 5 | **MG** | MG | a:3040 | 2.34 | 1.0 | 49.7 |  | 8.87 | 6 | Water 5 | | a:10083 | 2.63 | 1.0 | 43.0 |  | 3.84 | 7 | Water 6 | | a:5683 | 3.75 | 1.0 | 38.7 |  | 8.28 | 8 | A | OP1 | a:1974 | 3.76 | 1.0 | 34.3 |  | 7.00 | 9 | Water 7 | | a:9070 | 3.82 | 1.0 | 39.9 | 2.15 [MG:MG]a:3040 | 5.63 | 10 | Water 8 | | a:8112 | 3.97 | 1.0 | 39.5 |  | 7.45 | 11 | Water 9 | | a:8274 | 3.97 | 1.0 | 43.1 |  | 6.14 | 12 | U | OP1 | a:1975 | 4.01 | 1.0 | 36.4 |  | 8.58 | 13 | Water 10 | | a:3519 | 4.08 | 1.0 | 37.0 |  | 8.18 | 14 | A | OP1 | a:1973 | 4.26 | 1.0 | 34.4 |  | 9.58 | 15 | Water 11 | | a:8458 | 4.35 | 1.0 | 42.0 |  | 6.15 | 16 | Water 12 | | a:9270 | 4.49 | 1.0 | 41.8 |  | 7.67 | 17 | Water 13 | | a:5451 | 4.54 | 1.0 | 29.8 |  | 11.18 |

  
**1^st^ shell coordination angles and distances (*trans* in "blue"; *cys* in "green"; deviations in "orange"):**   
• 1-2(W/W): *159º*; 1-3(W/W): 91º; 1-4(W/W): 92º; 2-3(W/W): *103º*; 2-4(W/W): *105º*; 3-4(W/W): *83º*;   
• 1-2(W/W): *3.6Å*; 1-3(W/W): 2.7Å; 1-4(W/W): 2.9Å; 2-3(W/W): 3.0Å; 2-4(W/W): *3.2Å*; 3-4(W/W): 2.8Å;   
  
**2.15 ± 0.29: the average coodination distance of the 1^st^ six ligands is Mg^2+^ compatible**  
**10: Number of potential Water contacts**   
**3 : Number of second shell contacts involving water or ions**   
**30%: Percentage of satisfied second shell Water contacts**   
• Contact(s): /O_w_/O_w_/  
• Contact code: 2O_w_  
• Contact(s) in the exclusion zone: /O_w_/O_w_/O_w_/  
→ Missing octahedral coordination for MG ←  
→ Slightly distorded coord. octahedron (9.9º) ←  
Go to ToC

### MG:a:3040

|  | Res. type | Atom | Residue | Distance | Occupancy | B-factor | Closest atom(s) to a 1^st^ shell water | EM\_dens. |
| --- | --- | --- | --- | --- | --- | --- | --- | --- |
| **Ion** | | | |  | 1.0 | 49.7 |  | 8.87 |
 1 | Water 1 | | a:5300 | 1.93 | 1.0 | 41.9 | 1.80 [MG:MG]a:3039 | 4.60 | 2 | Water 2 | | a:5719 | 2.01 | 1.0 | 33.3 | 2.22 [MG:MG]a:3039 | 8.01 | 3 | A | OP1 | a:1974 | 2.07 | 1.0 | 34.3 |  | 7.00 | 4 | Water 3 | | a:5035 | 2.07 | 1.0 | 37.6 | 2.03 [MG:MG]a:3039 | 7.56 | 5 | U | OP1 | a:1975 | 2.12 | 1.0 | 36.4 |  | 8.58 | 6 | Water 4 | | a:9070 | 2.15 | 1.0 | 39.9 |  | 5.63 | 7 | **MG** | MG | a:3039 | 2.34 | 1.0 | 57.3 |  | 5.96 | 8 | Water 5 | | a:9908 | 4.03 | 1.0 | 39.2 | 1.86 [MG:MG]a:3039 | 8.44 | 9 | Water 6 | | a:10083 | 4.08 | 1.0 | 43.0 | 2.63 [MG:MG]a:3039 | 3.84 | 10 | A | O3' | a:1973 | 4.08 | 1.0 | 33.9 |  | 10.02 | 11 | Water 7 | | a:5451 | 4.15 | 1.0 | 29.8 | 2.86 [U:OP2]a:1975 | 11.18 | 12 | Water 8 | | a:8458 | 4.24 | 1.0 | 42.0 | 2.93 [A:OP2]a:1974 | 6.15 | 13 | A | O5' | a:1974 | 4.27 | 1.0 | 35.1 |  | 8.00 | 14 | A | OP1 | a:1973 | 4.27 | 1.0 | 34.4 |  | 9.58 | 15 | U | OP2 | a:1975 | 4.36 | 1.0 | 32.4 |  | 11.06 | 16 | A | O3' | a:1974 | 4.41 | 1.0 | 29.8 |  | 12.94 | 17 | U | O5' | a:1975 | 4.46 | 1.0 | 32.5 |  | 11.56 | 18 | A | OP2 | a:1974 | 4.50 | 1.0 | 33.9 |  | 7.99 |

  
**1^st^ shell coordination angles and distances (*trans* in "blue"; *cys* in "green"; deviations in "orange"):**   
• 1-2(W/W): *95º*; 1-3(W/OP1): 90º; 1-4(W/W): 87º; 1-5(W/OP1): *160º*; 1-6(W/W): *78º*; 2-3(W/OP1): *84º*; 2-4(W/W): 87º; 2-5(W/OP1): *99º*; 2-6(W/W): 171º; 3-4(OP1/W): *170º*; 3-5(OP1/OP1): *106º*; 3-6(OP1/W): 91º; 4-5(W/OP1): *79º*; 4-6(W/W): *98º*; 5-6(OP1/W): 89º;   
• 1-2(W/W): 2.9Å; 1-3(W/OP1): 2.8Å; 1-4(W/W): 2.7Å; 1-5(W/OP1): 4.0Å; 1-6(W/W): 2.6Å; 2-3(W/OP1): 2.7Å; 2-4(W/W): 2.8Å; 2-5(W/OP1): 3.1Å; 2-6(W/W): 4.2Å; 3-4(OP1/W): 4.1Å; 3-5(OP1/OP1): *3.3Å*; 3-6(OP1/W): 3.0Å; 4-5(W/OP1): 2.7Å; 4-6(W/W): 3.2Å; 5-6(OP1/W): 3.0Å;   
  
**2.06 ± 0.07: the average coodination distance of the 1^st^ six ligands is Mg^2+^ compatible**  
**8: Number of potential Water contacts**   
**3 : Number of second shell contacts involving water or ions**   
**38%: Percentage of satisfied second shell Water contacts**   
• Contact(s): /O_w_/O_w_/O_w_/O_w_/OP1/OP1/  
• Contact code: *cis-*2O_ph_4O_w_  
→ Slightly distorded coord. octahedron (7.6º) ←  
Go to ToC

### MG:a:3041

|  | Res. type | Atom | Residue | Distance | Occupancy | B-factor | Closest atom(s) to a 1^st^ shell water | EM\_dens. |
| --- | --- | --- | --- | --- | --- | --- | --- | --- |
| **Ion** | | | |  | 1.0 | 56.6 |  | 5.53 |
 1 | Water 1 | | a:7271 | 1.80 | 1.0 | 30.3 | 2.83 [G:N7]a:799 | 11.30 | 2 | Water 2 | | a:4580 | 1.84 | 1.0 | 38.5 | 2.64 [G:O6]a:800 | 5.18 | 3 | Water 3 | | a:10287 | 2.22 | 1.0 | 41.2 |  | 4.57 | 4 | Water 4 | | a:10192 | 2.23 | 1.0 | 40.1 |  | 4.43 | 5 | Water 5 | | a:7980 | 2.87 | 1.0 | 30.3 |  | 10.32 | 6 | Water 6 | | a:4894 | 3.60 | 1.0 | 29.7 |  | 11.18 | 7 | G | N7 | a:799 | 4.05 | 1.0 | 17.2 |  | 16.64 | 8 | Water 7 | | a:8148 | 4.26 | 1.0 | 28.3 |  | 11.90 | 9 | G | O6 | a:800 | 4.42 | 1.0 | 22.4 |  | 16.66 | 10 | Water 8 | | a:5257 | 4.43 | 1.0 | 34.3 |  | 8.84 | 11 | G | O6 | a:799 | 4.44 | 1.0 | 22.9 |  | 16.49 | 12 | Water 9 | | e:468 | 4.52 | 1.0 | 39.0 |  | 6.64 | 13 | Water 10 | | a:9249 | 4.59 | 1.0 | 43.0 |  | 5.40 | 14 | Water 11 | | a:9084 | 4.73 | 1.0 | 32.9 |  | 8.95 |

  
**1^st^ shell coordination angles and distances (*trans* in "blue"; *cys* in "green"; deviations in "orange"):**   
• 1-2(W/W): *116º*; 1-3(W/W): *85º*; 1-4(W/W): *99º*; 2-3(W/W): *109º*; 2-4(W/W): *132º*; 3-4(W/W): *106º*;   
• 1-2(W/W): 3.1Å; 1-3(W/W): 2.7Å; 1-4(W/W): 3.1Å; 2-3(W/W): *3.3Å*; 2-4(W/W): *3.7Å*; 3-4(W/W): *3.6Å*;   
  
**2.43 ± 0.63: the average coodination distance of the 1^st^ six ligands is Na^+^** compatible  
**12: Number of potential Water contacts**   
**2 : Number of second shell contacts involving water or ions**   
**17%: Percentage of satisfied second shell Water contacts**   
• Contact(s): /O_w_/O_w_/  
• Contact code: 2O_w_  
• Contact(s) in the exclusion zone: /O_w_/O_w_/O_w_/  
→ Missing octahedral coordination for MG ←  
→→→ Distorded coord. octahedron (19.5º) ←←←  
Go to ToC

### MG:a:3042

|  | Res. type | Atom | Residue | Distance | Occupancy | B-factor | Closest atom(s) to a 1^st^ shell water | EM\_dens. |
| --- | --- | --- | --- | --- | --- | --- | --- | --- |
| **Ion** | | | |  | 1.0 | 47.4 |  | 7.50 |
 1 | Water 1 | | a:7670 | 2.95 | 1.0 | 30.2 | 2.85 [A:N7]a:1656 // 2.87 [K:K]a:3344 | 11.12 | 2 | Water 2 | | a:7698 | 2.96 | 1.0 | 35.5 |  | 7.10 | 3 | Water 3 | | a:9734 | 3.13 | 1.0 | 37.9 |  | 5.24 | 4 | **K** | K | a:3344 | 3.21 | 1.0 | 30.0 |  | 12.81 | 5 | G | O4' | a:1655 | 3.29 | 1.0 | 27.2 |  | 14.25 | 6 | A | O5' | a:1656 | 3.64 | 1.0 | 29.7 |  | 12.13 | 7 | A | N3 | a:1654 | 3.80 | 1.0 | 18.9 |  | 14.91 | 8 | A | O2' | a:1654 | 4.14 | 1.0 | 32.3 |  | 11.46 | 9 | A | O4' | a:1656 | 4.26 | 1.0 | 25.3 |  | 17.19 | 10 | A | N7 | a:1656 | 4.33 | 1.0 | 17.6 |  | 16.84 | 11 | Water 4 | | a:9896 | 4.45 | 1.0 | 33.1 |  | 7.72 | 12 | A | N1 | a:2009 | 4.59 | 1.0 | 18.0 |  | 18.82 | 13 | G | O3' | a:1655 | 4.62 | 1.0 | 29.9 |  | 12.77 | 14 | A | O2' | a:2827 | 4.72 | 1.0 | 26.8 |  | 14.09 | 15 | Water 5 | | a:8172 | 4.73 | 1.0 | 26.0 | 2.97 [A:N1]a:2009 | 13.67 |

  
• • **3.20 ± 0.23: the average coodination distance of the 1^st^ six ligands is K^+^ compatible**  
**6: Number of potential Water contacts**   
**2 : Number of second shell contacts involving water or ions**   
**33%: Percentage of satisfied second shell Water contacts**   
• Contact code: → No contacts ←   
• Contact(s) in the exclusion zone: /O4'/O_w_/O_w_/O_w_/  
→ Firts MG coordination shell missing ←  
Go to ToC

### MG:a:3043

|  | Res. type | Atom | Residue | Distance | Occupancy | B-factor | Closest atom(s) to a 1^st^ shell water | EM\_dens. |
| --- | --- | --- | --- | --- | --- | --- | --- | --- |
| **Ion** | | | |  | 1.0 | 59.2 |  | 3.73 |
 1 | Water 1 | | a:4917 | 2.24 | 1.0 | 40.9 | 2.67 [G:OP1]a:248 | 6.92 | 2 | Water 2 | | a:8369 | 2.33 | 1.0 | 42.9 | 2.92 [C:O2']a:2399 | 4.62 | 3 | Water 3 | | a:10294 | 2.38 | 1.0 | 42.7 |  | 4.47 | 4 | Water 4 | | a:4552 | 2.53 | 1.0 | 36.4 | 2.64 [G:OP2]a:248 // 2.82 [C:OP2]a:249 | 7.27 | 5 | Water 5 | | a:9029 | 2.78 | 1.0 | 39.5 |  | 5.74 | 6 | Water 6 | | a:4654 | 4.00 | 1.0 | 38.4 | 2.65 [G:OP2]a:248 | 6.24 | 7 | C | O2' | a:2399 | 4.15 | 1.0 | 34.0 |  | 7.12 | 8 | G | OP1 | a:248 | 4.30 | 1.0 | 27.9 |  | 11.40 | 9 | G | OP2 | a:248 | 4.36 | 1.0 | 33.5 |  | 12.13 | 10 | C | OP2 | a:249 | 4.47 | 1.0 | 29.9 |  | 11.45 | 11 | Water 7 | | a:7196 | 4.53 | 1.0 | 33.5 | 2.82 [C:OP2]a:249 | 8.72 | 12 | C | O4' | a:2399 | 4.79 | 1.0 | 35.9 |  | 11.34 |

  
• • **2.71 ± 0.60: the average coodination distance of the 1^st^ six ligands is K^+^ compatible**  
**12: Number of potential Water contacts**   
**5 : Number of second shell contacts involving water or ions**   
**42%: Percentage of satisfied second shell Water contacts**   
• Contact(s): /O_w_/  
• Contact code: O_w_  
• Contact(s) in the exclusion zone: /O_w_/O_w_/O_w_/O_w_/  
→ Missing octahedral coordination for MG ←  
→2Fo-Fc density between 1.0 and 4.0 !   
Go to ToC

### MG:a:3044

|  | Res. type | Atom | Residue | Distance | Occupancy | B-factor | Closest atom(s) to a 1^st^ shell water | EM\_dens. |
| --- | --- | --- | --- | --- | --- | --- | --- | --- |
| **Ion** | | | |  | 1.0 | 24.2 |  | 25.59 |
 1 | G | O6 | a:2829 | 2.62 | 1.0 | 26.6 |  | 13.94 | 2 | Water 1 | | a:10036 | 2.64 | 1.0 | 48.6 |  | 4.42 | 3 | U | O3' | a:2626 | 2.88 | 1.0 | 34.7 |  | 12.37 | 4 | Water 2 | | d:461 | 2.93 | 1.0 | 31.8 |  | 10.81 | 5 | A | N3 | a:2830 | 2.97 | 1.0 | 23.0 |  | 13.15 | 6 | G | OP1 | a:2627 | 3.19 | 1.0 | 29.6 |  | 11.74 | 7 | Water 3 | | a:4380 | 3.86 | 1.0 | 39.2 | 2.62 [G:OP1]a:2627 | 3.20 | 8 | G | N7 | a:2829 | 4.25 | 1.0 | 24.4 |  | 12.72 | 9 | Water 4 | | a:7229 | 4.27 | 1.0 | 29.5 |  | 11.53 | 10 | Water 5 | | a:3680 | 4.33 | 1.0 | 42.0 |  | 5.10 | 11 | A | N1 | a:2830 | 4.33 | 1.0 | 19.0 |  | 17.20 | 12 | G | O5' | a:2627 | 4.37 | 1.0 | 32.3 |  | 11.10 | 13 | U | O2' | a:2626 | 4.56 | 1.0 | 28.4 |  | 12.71 |

  
• • **2.87 ± 0.20: the average coodination distance of the 1^st^ six ligands is K^+^ compatible**  
**4: Number of potential Water contacts**   
**0 : Number of second shell contacts involving water or ions**   
**0%: Percentage of satisfied second shell Water contacts**   
• Contact code: → No contacts ←   
• Contact(s) in the exclusion zone: /N3_A_/O3'/O6/O_w_/O_w_/OP1/  
→ Consider K in place of MG (4 valid contact(s) for K vs. 0 for MG ←)  
Go to ToC

### MG:a:3045

|  | Res. type | Atom | Residue | Distance | Occupancy | B-factor | Closest atom(s) to a 1^st^ shell water | EM\_dens. |
| --- | --- | --- | --- | --- | --- | --- | --- | --- |
| **Ion** | | | |  | 1.0 | 37.3 |  | 13.36 |
 1 | Water 1 | | a:7702 | 1.87 | 1.0 | 37.6 | 2.86 [G:O2']a:1268 | 8.63 | 2 | Water 2 | | a:6163 | 2.71 | 1.0 | 31.9 | 2.89 [G:OP2]a:2014 | 6.70 | 3 | Water 3 | | a:8293 | 2.87 | 1.0 | 20.8 | 2.00 [MG:MG]a:3235 | 15.59 | 4 | Water 4 | | a:3462 | 3.10 | 1.0 | 23.5 | 2.19 [MG:MG]a:3235 | 14.34 | 5 | G | O6 | a:2016 | 3.13 | 1.0 | 24.8 |  | 15.16 | 6 | G | O2' | a:1268 | 3.22 | 1.0 | 33.5 |  | 8.79 | 7 | Water 5 | | a:7699 | 3.23 | 1.0 | 32.2 | 2.86 [G:OP2]a:2014 | 11.61 | 8 | Water 6 | | a:9967 | 3.51 | 1.0 | 26.0 | 1.92 [MG:MG]a:3235 | 14.35 | 9 | **MG** | MG | a:3235 | 3.80 | 1.0 | 26.9 |  | 26.03 | 10 | U | O4 | a:2015 | 4.09 | 1.0 | 24.2 |  | 14.38 | 11 | Water 7 | | r:217 | 4.48 | 1.0 | 38.2 |  | 8.09 | 12 | U | OP2 | a:1269 | 4.52 | 1.0 | 27.5 |  | 12.88 | 13 | G | OP2 | a:2014 | 4.52 | 1.0 | 25.4 |  | 14.20 |

  
• • **2.82 ± 0.46: the average coodination distance of the 1^st^ six ligands is K^+^ compatible**  
**8: Number of potential Water contacts**   
**4 : Number of second shell contacts involving water or ions**   
**50%: Percentage of satisfied second shell Water contacts**   
• Contact code: → No contacts ←   
• Contact(s) in the exclusion zone: /O2'/O6/O_w_/O_w_/O_w_/O_w_/O_w_/  
→ Consider K in place of MG (1 valid contact(s) for K vs. 0 for MG ←)  
Go to ToC

### MG:a:3046

|  | Res. type | Atom | Residue | Distance | Occupancy | B-factor | Closest atom(s) to a 1^st^ shell water | EM\_dens. |
| --- | --- | --- | --- | --- | --- | --- | --- | --- |
| **Ion** | | | |  | 1.0 | 43.8 |  | 7.97 |
 1 | Water 1 | | a:6372 | 1.94 | 1.0 | 43.5 |  | 4.17 | 2 | Water 2 | | a:3445 | 2.05 | 1.0 | 38.3 |  | 5.63 | 3 | U | OP2 | a:2651 | 2.23 | 1.0 | 33.5 |  | 9.83 | 4 | C | OP1 | a:2650 | 2.29 | 1.0 | 30.6 |  | 10.29 | 5 | Water 3 | | a:6335 | 2.34 | 1.0 | 42.8 |  | 5.14 | 6 | Water 4 | | a:9543 | 3.44 | 1.0 | 41.3 |  | 3.79 | 7 | Water 5 | | a:4508 | 3.61 | 1.0 | 37.9 | 2.63 [U:OP1]a:2651 | 6.26 | 8 | U | OP1 | a:2651 | 4.02 | 1.0 | 35.0 |  | 8.05 | 9 | C | O5' | a:2650 | 4.15 | 1.0 | 30.7 |  | 11.46 | 10 | C | OP2 | a:2650 | 4.28 | 1.0 | 31.6 |  | 9.78 | 11 | Water 6 | | a:4133 | 4.36 | 1.0 | 31.9 |  | 10.89 | 12 | Water 7 | | a:3416 | 4.38 | 1.0 | 47.4 |  | 3.72 | 13 | U | O5' | a:2651 | 4.51 | 1.0 | 32.7 |  | 10.96 | 14 | C | O3' | a:2650 | 4.55 | 1.0 | 29.7 |  | 11.86 | 15 | G | O3' | a:2649 | 4.77 | 1.0 | 30.3 |  | 11.17 |

  
**1^st^ shell coordination angles and distances (*trans* in "blue"; *cys* in "green"; deviations in "orange"):**   
• 1-2(W/W): *79º*; 1-3(W/OP2): 92º; 1-4(W/OP1): *81º*; 2-3(W/OP2): *160º*; 2-4(W/OP1): *67º*; 3-4(OP2/OP1): 94º;   
• 1-2(W/W): 2.5Å; 1-3(W/OP2): 3.0Å; 1-4(W/OP1): 2.8Å; 2-3(W/OP2): *4.2Å*; 2-4(W/OP1): 2.4Å; 3-4(OP2/OP1): *3.3Å*;   
  
**2.38 ± 0.50: the average coodination distance of the 1^st^ six ligands is Na^+^** compatible  
**8: Number of potential Water contacts**   
**0 : Number of second shell contacts involving water or ions**   
**0%: Percentage of satisfied second shell Water contacts**   
• Contact(s): /O_w_/O_w_/OP1/OP2/  
• Contact code: *cis-*2O_ph_2O_w_  
• Contact(s) in the exclusion zone: /O_w_/  
→ Missing octahedral coordination for MG ←  
→→→ Distorded coord. octahedron (11.7º) ←←←  
Go to ToC

### MG:a:3047

|  | Res. type | Atom | Residue | Distance | Occupancy | B-factor | Closest atom(s) to a 1^st^ shell water | EM\_dens. |
| --- | --- | --- | --- | --- | --- | --- | --- | --- |
| **Ion** | | | |  | 1.0 | 50.3 |  | 6.01 |
 1 | Water 1 | | a:4446 | 1.91 | 1.0 | 38.1 | 2.63 [A:N7]a:1903 | 6.90 | 2 | Water 2 | | a:8870 | 1.92 | 1.0 | 39.6 | 2.99 [U:OP2]a:1902 | 5.77 | 3 | Water 3 | | a:10189 | 1.96 | 1.0 | 43.5 |  | 4.91 | 4 | Water 4 | | a:9365 | 2.13 | 1.0 | 44.8 |  | 3.43 | 5 | Water 5 | | a:7802 | 2.25 | 1.0 | 40.1 | 2.89 [U:OP1]a:1902 | 6.01 | 6 | A | OP2 | a:1903 | 2.29 | 1.0 | 33.8 |  | 8.02 | 7 | Water 6 | | a:9043 | 3.95 | 1.0 | 39.2 |  | 6.38 | 8 | Water 7 | | a:10026 | 4.00 | 1.0 | 45.2 |  | 2.80 | 9 | A | OP1 | a:1903 | 4.03 | 1.0 | 36.7 |  | 9.28 | 10 | Water 8 | | a:9551 | 4.10 | 1.0 | 52.7 |  | 2.25 | 11 | Water 9 | | a:9558 | 4.11 | 1.0 | 43.3 |  | 4.39 | 12 | A | N7 | a:1903 | 4.33 | 1.0 | 25.6 |  | 12.46 | 13 | Water 10 | | a:8490 | 4.35 | 1.0 | 48.3 | 2.93 [A:OP1]a:1903 | 2.11 | 14 | U | OP1 | a:1902 | 4.50 | 1.0 | 38.9 |  | 7.14 | 15 | U | O3' | a:1902 | 4.58 | 1.0 | 34.8 |  | 8.06 | 16 | A | O5' | a:1903 | 4.62 | 1.0 | 37.0 |  | 10.38 | 17 | U | O5' | a:1902 | 4.62 | 1.0 | 44.6 |  | 7.27 | 18 | U | OP2 | a:1902 | 4.76 | 1.0 | 37.0 |  | 7.78 |

  
**1^st^ shell coordination angles and distances (*trans* in "blue"; *cys* in "green"; deviations in "orange"):**   
• 1-2(W/W): *103º*; 1-3(W/W): *105º*; 1-4(W/W): *98º*; 1-5(W/W): *163º*; 1-6(W/OP2): 86º; 2-3(W/W): *82º*; 2-4(W/W): *159º*; 2-5(W/W): *82º*; 2-6(W/OP2): 89º; 3-4(W/W): *97º*; 3-5(W/W): 92º; 3-6(W/OP2): *168º*; 4-5(W/W): *78º*; 4-6(W/OP2): 88º; 5-6(W/OP2): *78º*;   
• 1-2(W/W): 3.0Å; 1-3(W/W): 3.1Å; 1-4(W/W): 3.0Å; 1-5(W/W): 4.1Å; 1-6(W/OP2): 2.9Å; 2-3(W/W): 2.6Å; 2-4(W/W): 4.0Å; 2-5(W/W): 2.7Å; 2-6(W/OP2): 3.0Å; 3-4(W/W): 3.1Å; 3-5(W/W): 3.0Å; 3-6(W/OP2): *4.2Å*; 4-5(W/W): 2.7Å; 4-6(W/OP2): 3.1Å; 5-6(W/OP2): 2.9Å;   
  
**2.08 ± 0.15: the average coodination distance of the 1^st^ six ligands is Mg^2+^ compatible**  
**10: Number of potential Water contacts**   
**3 : Number of second shell contacts involving water or ions**   
**30%: Percentage of satisfied second shell Water contacts**   
• Contact(s): /O_w_/O_w_/O_w_/O_w_/O_w_/OP2/  
• Contact code: O_ph_5O_w_  
→ Slightly distorded coord. octahedron (9.5º) ←  
Go to ToC

### MG:a:3048

|  | Res. type | Atom | Residue | Distance | Occupancy | B-factor | Closest atom(s) to a 1^st^ shell water | EM\_dens. |
| --- | --- | --- | --- | --- | --- | --- | --- | --- |
| **Ion** | | | |  | 1.0 | 60.1 |  | 2.44 |
 1 | Water 1 | | a:9341 | 1.94 | 1.0 | 46.3 |  | 2.92 | 2 | Water 2 | | a:6852 | 1.98 | 1.0 | 48.3 |  | 2.30 | 3 | Water 3 | | a:4479 | 2.11 | 1.0 | 53.8 | 2.63 [U:OP2]a:1442 | 2.59 | 4 | A | OP1 | a:1441 | 2.12 | 1.0 | 37.6 |  | 5.32 | 5 | Water 4 | | a:6299 | 2.15 | 1.0 | 49.9 |  | 2.11 | 6 | Water 5 | | a:7473 | 2.26 | 1.0 | 48.4 |  | 2.58 | 7 | Water 6 | | a:6100 | 4.22 | 1.0 | 53.8 |  | 2.68 | 8 | A | OP2 | a:1441 | 4.30 | 1.0 | 39.1 |  | 6.88 | 9 | Water 7 | | a:10292 | 4.32 | 1.0 | 49.2 |  | 2.60 | 10 | U | OP2 | a:1442 | 4.32 | 1.0 | 40.5 |  | 6.11 | 11 | Water 8 | | a:4915 | 4.38 | 1.0 | 39.9 | 2.67 [A:OP2]a:1441 | 6.43 | 12 | A | O5' | a:1441 | 4.39 | 1.0 | 42.0 |  | 6.55 | 13 | Water 9 | | a:4771 | 4.43 | 1.0 | 49.0 |  | 3.07 | 14 | U | O3' | a:1440 | 4.46 | 1.0 | 43.2 |  | 6.08 | 15 | U | OP1 | a:1470 | 4.47 | 1.0 | 48.6 |  | 2.93 | 16 | Water 10 | | a:7604 | 4.47 | 1.0 | 45.9 |  | 2.49 |

  
**1^st^ shell coordination angles and distances (*trans* in "blue"; *cys* in "green"; deviations in "orange"):**   
• 1-2(W/W): 86º; 1-3(W/W): 89º; 1-4(W/OP1): *97º*; 1-5(W/W): 174º; 1-6(W/W): 87º; 2-3(W/W): 173º; 2-4(W/OP1): 86º; 2-5(W/W): *100º*; 2-6(W/W): *100º*; 3-4(W/OP1): 90º; 3-5(W/W): *85º*; 3-6(W/W): *85º*; 4-5(OP1/W): *80º*; 4-6(OP1/W): 173º; 5-6(W/W): 95º;   
• 1-2(W/W): 2.7Å; 1-3(W/W): 2.8Å; 1-4(W/OP1): 3.1Å; 1-5(W/W): 4.1Å; 1-6(W/W): 2.9Å; 2-3(W/W): 4.1Å; 2-4(W/OP1): 2.8Å; 2-5(W/W): 3.2Å; 2-6(W/W): *3.2Å*; 3-4(W/OP1): 3.0Å; 3-5(W/W): 2.9Å; 3-6(W/W): 2.9Å; 4-5(OP1/W): 2.8Å; 4-6(OP1/W): *4.4Å*; 5-6(W/W): *3.2Å*;   
  
**2.09 ± 0.11: the average coodination distance of the 1^st^ six ligands is Mg^2+^ compatible**  
**10: Number of potential Water contacts**   
**1 : Number of second shell contacts involving water or ions**   
**10%: Percentage of satisfied second shell Water contacts**   
• Contact(s): /O_w_/O_w_/O_w_/O_w_/O_w_/OP1/  
• Contact code: O_ph_5O_w_  
→ Slightly distorded coord. octahedron (5.6º) ←→2Fo-Fc density between 1.0 and 4.0 !   
Go to ToC

### MG:a:3049

|  | Res. type | Atom | Residue | Distance | Occupancy | B-factor | Closest atom(s) to a 1^st^ shell water | EM\_dens. |
| --- | --- | --- | --- | --- | --- | --- | --- | --- |
| **Ion** | | | |  | 1.0 | 55.8 |  | 4.04 |
 1 | A | OP2 | a:1553 | 1.96 | 1.0 | 41.3 |  | 6.58 | 2 | Water 1 | | a:9185 | 2.20 | 1.0 | 44.0 |  | 3.97 | 3 | Water 2 | | a:7786 | 2.23 | 1.0 | 46.4 | 2.92 [U:O2']a:1722 | 3.44 | 4 | Water 3 | | a:9606 | 2.40 | 1.0 | 53.2 |  | 2.40 | 5 | Water 4 | | a:9458 | 2.53 | 1.0 | 44.1 |  | 3.82 | 6 | Water 5 | | a:7226 | 2.73 | 1.0 | 47.5 | 2.82 [A:N7]a:1553 | 3.72 | 7 | A | O5' | a:1553 | 3.70 | 1.0 | 43.4 |  | 5.29 | 8 | Water 6 | | a:4575 | 4.12 | 1.0 | 44.3 |  | 3.36 | 9 | A | OP1 | a:1553 | 4.18 | 1.0 | 43.8 |  | 4.02 | 10 | A | N7 | a:1553 | 4.32 | 1.0 | 30.6 |  | 9.45 | 11 | C | O3' | a:1552 | 4.37 | 1.0 | 43.0 |  | 6.08 | 12 | A | N7 | a:1554 | 4.59 | 1.0 | 32.1 |  | 7.54 | 13 | U | O2' | a:1722 | 4.70 | 1.0 | 49.4 |  | 4.87 |

  
**1^st^ shell coordination angles and distances (*trans* in "blue"; *cys* in "green"; deviations in "orange"):**   
• 1-2(OP2/W): *100º*; 1-3(OP2/W): 86º; 2-3(W/W): *169º*;   
• 1-2(OP2/W): 3.2Å; 1-3(OP2/W): 2.9Å; 2-3(W/W): *4.4Å*;   
  
**2.34 ± 0.25: the average coodination distance of the 1^st^ six ligands is Na^+^** compatible  
**10: Number of potential Water contacts**   
**2 : Number of second shell contacts involving water or ions**   
**20%: Percentage of satisfied second shell Water contacts**   
• Contact(s): /O_w_/O_w_/OP2/  
• Contact code: O_ph_2O_w_  
• Contact(s) in the exclusion zone: /O_w_/O_w_/O_w_/  
→ Missing octahedral coordination for MG ←  
Go to ToC

### MG:a:3050

|  | Res. type | Atom | Residue | Distance | Occupancy | B-factor | Closest atom(s) to a 1^st^ shell water | EM\_dens. |
| --- | --- | --- | --- | --- | --- | --- | --- | --- |
| **Ion** | | | |  | 1.0 | 54.7 |  | 3.75 |
 1 | Water 1 | | a:6652 | 1.98 | 1.0 | 44.9 |  | 3.70 | 2 | Water 2 | | a:4690 | 2.00 | 1.0 | 43.9 | 2.65 [U:O4]a:1522 | 4.46 | 3 | Water 3 | | a:4223 | 2.01 | 1.0 | 47.5 | 2.60 [A:OP2]a:1471 | 3.63 | 4 | G | O6 | a:1523 | 2.09 | 1.0 | 42.5 |  | 5.88 | 5 | Water 4 | | a:3684 | 2.31 | 1.0 | 48.0 | 2.51 [A:N1]a:1524 | 4.02 | 6 | A | OP2 | a:1471 | 3.82 | 1.0 | 41.8 |  | 5.52 | 7 | U | O4 | a:1522 | 3.86 | 1.0 | 41.8 |  | 7.05 | 8 | A | N1 | a:1524 | 4.07 | 1.0 | 39.5 |  | 6.23 | 9 | Water 5 | | a:9389 | 4.75 | 1.0 | 48.3 |  | 3.35 |

  
**1^st^ shell coordination angles and distances (*trans* in "blue"; *cys* in "green"; deviations in "orange"):**   
• 1-2(W/W): *101º*; 1-3(W/W): *170º*; 1-4(W/O6): 86º; 2-3(W/W): 87º; 2-4(W/O6): 94º; 3-4(W/O6): *100º*;   
• 1-2(W/W): 3.1Å; 1-3(W/W): 4.0Å; 1-4(W/O6): 2.8Å; 2-3(W/W): 2.8Å; 2-4(W/O6): 3.0Å; 3-4(W/O6): 3.2Å;   
  
**2.37 ± 0.66: the average coodination distance of the 1^st^ six ligands is Na^+^** compatible  
**8: Number of potential Water contacts**   
**3 : Number of second shell contacts involving water or ions**   
**38%: Percentage of satisfied second shell Water contacts**   
• Contact(s): /O6/O_w_/O_w_/O_w_/  
• Contact code: O_b_3O_w_  
• Contact(s) in the exclusion zone: /O_w_/  
→ Missing octahedral coordination for MG ←  
→ Slightly distorded coord. octahedron (7.0º) ←→2Fo-Fc density between 1.0 and 4.0 !   
Go to ToC

### MG:a:3051

|  | Res. type | Atom | Residue | Distance | Occupancy | B-factor | Closest atom(s) to a 1^st^ shell water | EM\_dens. |
| --- | --- | --- | --- | --- | --- | --- | --- | --- |
| **Ion** | | | |  | 1.0 | 33.2 |  | 12.44 |
 1 | G | O6 | a:1521 | 2.82 | 1.0 | 45.9 |  | 6.07 | 2 | Water 1 | | a:6069 | 2.85 | 1.0 | 37.0 | 2.75 [G:N7]a:1473 | 6.14 | 3 | Water 2 | | a:6704 | 2.87 | 1.0 | 42.2 | 2.79 [A:OP2]a:1472 | 5.77 | 4 | G | O6 | a:1473 | 2.98 | 1.0 | 38.2 |  | 6.72 | 5 | U | O4 | a:1522 | 2.99 | 1.0 | 41.8 |  | 7.05 | 6 | Water 3 | | a:9389 | 3.53 | 1.0 | 48.3 |  | 3.35 | 7 | A | OP2 | a:1472 | 4.24 | 1.0 | 40.3 |  | 6.15 | 8 | Water 4 | | a:4690 | 4.48 | 1.0 | 43.9 | 2.65 [U:O4]a:1522 | 4.46 | 9 | G | N7 | a:1473 | 4.57 | 1.0 | 33.3 |  | 7.55 |

  
• • **3.01 ± 0.24: the average coodination distance of the 1^st^ six ligands is K^+^ compatible**  
**6: Number of potential Water contacts**   
**2 : Number of second shell contacts involving water or ions**   
**33%: Percentage of satisfied second shell Water contacts**   
• Contact code: → No contacts ←   
• Contact(s) in the exclusion zone: /O4/O6/O6/O_w_/O_w_/  
→ Consider K in place of MG (3 valid contact(s) for K vs. 0 for MG ←)  
Go to ToC

### MG:a:3052

|  | Res. type | Atom | Residue | Distance | Occupancy | B-factor | Closest atom(s) to a 1^st^ shell water | EM\_dens. |
| --- | --- | --- | --- | --- | --- | --- | --- | --- |
| **Ion** | | | |  | 1.0 | 51.6 |  | 5.95 |
 1 | G | OP1 | a:1684 | 2.52 | 1.0 | 35.9 |  | 8.23 | 2 | U | OP2 | a:1685 | 2.80 | 1.0 | 37.8 |  | 7.09 | 3 | G | N1 | a:1701 | 3.48 | 1.0 | 33.5 |  | 10.46 | 4 | G | O5' | a:1684 | 3.74 | 1.0 | 30.9 |  | 10.21 | 5 | Water 1 | | a:4699 | 3.89 | 1.0 | 44.2 | 2.65 [G:OP2]a:1684 | 6.36 | 6 | U | OP1 | a:1685 | 3.95 | 1.0 | 41.3 |  | 5.76 | 7 | G | O6 | a:1701 | 3.96 | 1.0 | 41.1 |  | 7.90 | 8 | G | OP2 | a:1684 | 4.07 | 1.0 | 32.9 |  | 8.81 | 9 | **K** | K | a:3270 | 4.17 | 1.0 | 30.0 |  | 13.26 | 10 | G | O3' | a:1684 | 4.24 | 1.0 | 36.4 |  | 9.11 | 11 | Water 2 | | a:9598 | 4.48 | 1.0 | 49.9 | 2.76 [K:K]a:3270 | 4.34 | 12 | G | N3 | a:1701 | 4.48 | 1.0 | 33.1 |  | 8.87 |

  
• • **3.40 ± 0.55: the average coodination distance of the 1^st^ six ligands is K^+^ compatible**  
**2: Number of potential Water contacts**   
**1 : Number of second shell contacts involving water or ions**   
**50%: Percentage of satisfied second shell Water contacts**   
• Contact code: → No contacts ←   
• Contact(s) in the exclusion zone: /OP1/OP2/  
→ Consider K in place of MG (1 valid contact(s) for K vs. 0 for MG ←)  
Go to ToC

### MG:a:3053

|  | Res. type | Atom | Residue | Distance | Occupancy | B-factor | Closest atom(s) to a 1^st^ shell water | EM\_dens. |
| --- | --- | --- | --- | --- | --- | --- | --- | --- |
| **Ion** | | | |  | 1.0 | 43.8 |  | 9.83 |
 1 | Water 1 | | a:10124 | 2.55 | 1.0 | 44.7 | 2.10 [MG:MG]a:3013 | 4.51 | 2 | G | O6 | a:1954 | 2.68 | 1.0 | 31.5 |  | 12.19 | 3 | G | O2' | a:1958 | 2.83 | 1.0 | 28.9 |  | 11.30 | 4 | U | OP2 | a:1959 | 2.85 | 1.0 | 32.5 |  | 9.12 | 5 | Water 2 | | a:5527 | 2.95 | 1.0 | 36.9 | 2.32 [MG:MG]a:3013 | 7.75 | 6 | G | O6 | a:1953 | 3.20 | 1.0 | 30.0 |  | 11.41 | 7 | Water 3 | | a:7073 | 3.44 | 1.0 | 41.4 | 1.97 [MG:MG]a:3013 | 4.74 | 8 | Water 4 | | a:3791 | 3.85 | 1.0 | 36.7 |  | 7.85 | 9 | **MG** | MG | a:3013 | 3.86 | 1.0 | 52.2 |  | 5.03 | 10 | Water 5 | | a:8246 | 4.03 | 1.0 | 43.3 | 2.90 [U:OP1]a:1959 | 6.17 | 11 | Water 6 | | a:4051 | 4.09 | 1.0 | 31.8 | 2.58 [G:O6]a:1954 | 8.73 | 12 | G | O3' | a:1958 | 4.21 | 1.0 | 36.3 |  | 10.14 | 13 | Water 7 | | a:5018 | 4.48 | 1.0 | 32.3 | 2.77 [U:OP2]a:1959 // 2.93 [C:N4]a:1961 | 9.12 | 14 | U | OP1 | a:1959 | 4.61 | 1.0 | 34.8 |  | 10.83 |

  
• • **2.84 ± 0.21: the average coodination distance of the 1^st^ six ligands is K^+^ compatible**  
**4: Number of potential Water contacts**   
**2 : Number of second shell contacts involving water or ions**   
**50%: Percentage of satisfied second shell Water contacts**   
• Contact code: → No contacts ←   
• Contact(s) in the exclusion zone: /O2'/O6/O6/O_w_/O_w_/OP2/  
→ Consider K in place of MG (3 valid contact(s) for K vs. 0 for MG ←)  
Go to ToC

### MG:a:3054

|  | Res. type | Atom | Residue | Distance | Occupancy | B-factor | Closest atom(s) to a 1^st^ shell water | EM\_dens. |
| --- | --- | --- | --- | --- | --- | --- | --- | --- |
| **Ion** | | | |  | 1.0 | 30.4 |  | 16.41 |
 1 | Water 1 | | a:9540 | 2.80 | 1.0 | 38.2 |  | 6.06 | 2 | C | OP1 | a:2583 | 2.81 | 1.0 | 27.0 |  | 12.53 | 3 | G | OP2 | a:2582 | 2.85 | 1.0 | 25.1 |  | 11.94 | 4 | Water 2 | | a:10139 | 2.90 | 1.0 | 41.3 |  | 5.39 | 5 | G | O3' | a:2582 | 3.02 | 1.0 | 31.8 |  | 13.56 | 6 | A | O3' | a:2581 | 3.17 | 1.0 | 27.2 |  | 13.43 | 7 | Water 3 | | a:6943 | 3.36 | 1.0 | 32.9 | 2.15 [MG:MG]a:3304 | 8.40 | 8 | Water 4 | | a:9271 | 3.66 | 1.0 | 34.6 | 2.36 [MG:MG]a:3304 | 9.35 | 9 | **MG** | MG | a:3304 | 4.18 | 1.0 | 39.7 |  | 15.25 | 10 | Water 5 | | a:4931 | 4.27 | 1.0 | 38.7 | 2.67 [G:OP2]a:2582 | 5.34 | 11 | Water 6 | | d:417 | 4.34 | 1.0 | 44.1 |  | 6.83 | 12 | Water 7 | | d:497 | 4.43 | 1.0 | 35.7 |  | 7.95 | 13 | G | O5' | a:2582 | 4.48 | 1.0 | 29.0 |  | 14.31 | 14 | C | O5' | a:2583 | 4.58 | 1.0 | 29.9 |  | 12.99 | 15 | C | OP2 | a:2616 | 4.64 | 1.0 | 35.9 |  | 7.26 | 16 | C | OP2 | a:2583 | 4.65 | 1.0 | 29.6 |  | 12.65 |

  
• • **2.93 ± 0.13: the average coodination distance of the 1^st^ six ligands is K^+^ compatible**  
**4: Number of potential Water contacts**   
**0 : Number of second shell contacts involving water or ions**   
**0%: Percentage of satisfied second shell Water contacts**   
• Contact code: → No contacts ←   
• Contact(s) in the exclusion zone: /O3'/O3'/O_w_/O_w_/O_w_/OP1/OP2/  
→ Consider K in place of MG (4 valid contact(s) for K vs. 0 for MG ←)  
Go to ToC

### MG:a:3055

|  | Res. type | Atom | Residue | Distance | Occupancy | B-factor | Closest atom(s) to a 1^st^ shell water | EM\_dens. |
| --- | --- | --- | --- | --- | --- | --- | --- | --- |
| **Ion** | | | |  | 1.0 | 41.0 |  | 13.28 |
 1 | C | OP2 | a:516 | 2.59 | 1.0 | 24.6 |  | 16.46 | 2 | Water 1 | | p:256 | 2.64 | 1.0 | 41.3 | 2.52 [MG:MG]a:3146 | 8.58 | 3 | A | OP1 | a:515 | 2.76 | 1.0 | 27.7 |  | 12.97 | 4 | Water 2 | | a:9372 | 3.16 | 1.0 | 37.2 |  | 7.30 | 5 | Water 3 | | a:7310 | 3.45 | 1.0 | 30.5 | 2.92 [C:OP2]a:516 | 11.97 | 6 | A | O5' | a:515 | 3.55 | 1.0 | 29.5 |  | 12.19 | 7 | Water 4 | | a:4047 | 3.87 | 1.0 | 34.7 | 2.11 [MG:MG]a:3146 | 8.50 | 8 | Water 5 | | p:235 | 4.07 | 1.0 | 41.2 |  | 7.09 | 9 | A | OP2 | a:515 | 4.08 | 1.0 | 26.2 |  | 11.38 | 10 | Water 6 | | a:6727 | 4.28 | 1.0 | 43.4 | 2.79 [A:OP2]a:515 | 5.13 | 11 | **MG** | MG | a:3146 | 4.40 | 1.0 | 48.7 |  | 8.52 | 12 | Water 7 | | a:4325 | 4.57 | 1.0 | 31.2 | 2.61 [A:OP1]a:515 | 12.23 | 13 | Water 8 | | p:223 | 4.58 | 1.0 | 40.5 | 2.23 [MG:MG]a:3146 | 7.21 | 14 | Water 9 | | a:10146 | 4.70 | 1.0 | 44.6 |  | 3.73 | 15 | C | O5' | a:516 | 4.76 | 1.0 | 24.7 |  | 13.77 | 16 | A | O3' | a:515 | 4.78 | 1.0 | 28.9 |  | 13.28 |

  
• • **3.02 ± 0.38: the average coodination distance of the 1^st^ six ligands is K^+^ compatible**  
**6: Number of potential Water contacts**   
**2 : Number of second shell contacts involving water or ions**   
**33%: Percentage of satisfied second shell Water contacts**   
• Contact code: → No contacts ←   
• Contact(s) in the exclusion zone: /O_w_/O_w_/OP1/OP2/  
→ Consider K in place of MG (1 valid contact(s) for K vs. 0 for MG ←)  
Go to ToC

### K:a:3056

|  | Res. type | Atom | Residue | Distance | Occupancy | B-factor | Closest atom(s) to a 1^st^ shell water | EM\_dens. |
| --- | --- | --- | --- | --- | --- | --- | --- | --- |
| **Ion** | | | |  | 1.0 | 30.0 |  | 17.72 |
 1 | G | O6 | a:871 | 2.75 | 1.0 | 36.7 |  | 10.35 | 2 | C | O2 | a:867 | 2.77 | 1.0 | 37.1 |  | 8.18 | 3 | U | O4 | a:870 | 2.78 | 1.0 | 42.9 |  | 9.21 | 4 | Water 1 | | a:5392 | 2.90 | 1.0 | 42.7 |  | 5.59 | 5 | Water 2 | | a:7035 | 3.09 | 1.0 | 39.4 | 2.81 [G:N7]a:871 | 6.55 | 6 | C | O2' | a:867 | 3.10 | 1.0 | 40.3 |  | 7.79 | 7 | A | N6 | a:911 | 4.11 | 1.0 | 24.0 |  | 11.99 | 8 | G | N7 | a:871 | 4.13 | 1.0 | 29.6 |  | 8.90 | 9 | C | N1 | a:867 | 4.25 | 1.0 | 30.4 |  | 9.36 | 10 | Water 3 | | a:6145 | 4.39 | 1.0 | 43.4 |  | 3.99 | 11 | C | N3 | a:867 | 4.40 | 1.0 | 34.2 |  | 9.51 | 12 | U | O4 | a:872 | 4.41 | 1.0 | 36.7 |  | 6.64 | 13 | C | O3' | a:867 | 4.74 | 1.0 | 49.0 |  | 4.45 |

  
**1^st^ shell coordination angles and distances (*trans* in "blue"; *cys* in "green"; deviations in "orange"):**   
• 1-2(O6/O2): 95; 1-3(O6/O4): 70; 1-4(O6/W): 170; 1-5(O6/W): 85; 1-6(O6/O2'): 116; 2-3(O2/O4): 79; 2-4(O2/W): 93; 2-5(O2/W): 151; 2-6(O2/O2'): 75; 3-4(O4/W): 106; 3-5(O4/W): 74; 3-6(O4/O2'): 153; 4-5(W/W): 85; 4-6(W/O2'): 71; 5-6(W/O2'): 131;   
• 1-2(O6/O2): 4.1; 1-3(O6/O4): 3.2; 1-4(O6/W): 5.6; 1-5(O6/W): 4.0; 1-6(O6/O2'): 5.0; 2-3(O2/O4): 3.5; 2-4(O2/W): 4.1; 2-5(O2/W): 5.7; 2-6(O2/O2'): 3.6; 3-4(O4/W): 4.5; 3-5(O4/W): 3.5; 3-6(O4/O2'): 5.7; 4-5(W/W): 4.0; 4-6(W/O2'): 3.5; 5-6(W/O2'): 5.6;   
  
**4.27 ± 1.38: the average coodination distance of the 1^st^ six ligands could correspond to an hexahydrated Mg^2+^.**  
**4: Number of potential Water contacts**   
**1 : Number of second shell contacts involving water or ions**   
**25%: Percentage of satisfied second shell Water contacts**   
• Contact(s): /O2/O2'/O4/O6/O_w_/O_w_/  
• Contact code: O_r_3O_b_2O_w_  
Go to ToC

### K:a:3057

|  | Res. type | Atom | Residue | Distance | Occupancy | B-factor | Closest atom(s) to a 1^st^ shell water | EM\_dens. |
| --- | --- | --- | --- | --- | --- | --- | --- | --- |
| **Ion** | | | |  | 1.0 | 30.0 |  | 5.17 |
 1 | G | O6 | a:857 | 2.64 | 1.0 | 37.2 |  | 8.59 | 2 | G | O6 | a:858 | 2.90 | 1.0 | 32.0 |  | 11.10 | 3 | Water 1 | | a:7575 | 2.92 | 1.0 | 42.1 | 2.85 [G:N7]a:857 | 4.39 | 4 | Water 2 | | a:4760 | 3.61 | 1.0 | 37.0 | 2.65 [G:N7]a:858 | 5.80 | 5 | Water 3 | | a:9247 | 3.82 | 1.0 | 48.5 |  | 3.49 | 6 | C | N4 | a:856 | 3.93 | 1.0 | 33.3 |  | 7.09 | 7 | C | N4 | a:923 | 3.97 | 1.0 | 23.4 |  | 11.71 | 8 | C | N4 | a:924 | 3.99 | 1.0 | 23.0 |  | 10.69 | 9 | G | N7 | a:858 | 4.04 | 1.0 | 32.1 |  | 9.64 | 10 | G | N7 | a:857 | 4.21 | 1.0 | 33.7 |  | 8.65 | 11 | Water 4 | | a:5953 | 4.48 | 1.0 | 40.9 | 2.74 [C:N4]a:923 | 4.77 | 12 | Water 5 | | a:9657 | 4.74 | 1.0 | 44.7 |  | 5.04 |

  
**1^st^ shell coordination angles and distances (*trans* in "blue"; *cys* in "green"; deviations in "orange"):**   
• 1-2(O6/O6): 73; 1-3(O6/W): 89; 2-3(O6/W): 126;   
• 1-2(O6/O6): 3.3; 1-3(O6/W): 3.9; 2-3(O6/W): 5.2;   
  
**4.63 ± 1.47: the average coodination distance of the 1^st^ six ligands could correspond to an hexahydrated Mg^2+^.**  
**6: Number of potential Water contacts**   
**2 : Number of second shell contacts involving water or ions**   
**33%: Percentage of satisfied second shell Water contacts**   
• Contact(s): /O6/O6/O_w_/  
• Contact code: 2O_b_O_w_  
→ Coordination < 6 for K ←  
Go to ToC

### K:a:3058

|  | Res. type | Atom | Residue | Distance | Occupancy | B-factor | Closest atom(s) to a 1^st^ shell water | EM\_dens. |
| --- | --- | --- | --- | --- | --- | --- | --- | --- |
| **Ion** | | | |  | 1.0 | 30.0 |  | 16.63 |
 1 | Water 1 | | a:9173 | 2.67 | 1.0 | 45.5 |  | 4.29 | 2 | U | O2 | a:328 | 2.75 | 1.0 | 40.8 |  | 8.85 | 3 | ASN | OD1 | t:69 | 2.82 | 1.0 | 45.3 |  | 7.86 | 4 | C | O2 | a:335 | 2.88 | 1.0 | 38.9 |  | 8.89 | 5 | SER | OG | t:68 | 3.02 | 1.0 | 44.2 |  | 7.45 | 6 | SER | O | t:68 | 3.16 | 1.0 | 37.9 |  | 9.26 | 7 | U | O2' | a:328 | 3.26 | 1.0 | 41.1 |  | 7.38 | 8 | Water 2 | | a:8901 | 3.31 | 1.0 | 41.5 |  | 4.93 | 9 | U | N1 | a:328 | 4.45 | 1.0 | 31.9 |  | 9.72 | 10 | C | N1 | a:335 | 4.59 | 1.0 | 32.1 |  | 8.58 | 11 | C | N3 | a:335 | 4.60 | 1.0 | 32.5 |  | 8.27 |

  
**1^st^ shell coordination angles and distances (*trans* in "blue"; *cys* in "green"; deviations in "orange"):**   
• 1-2(W/O2): 117; 1-3(W/OD1): 91; 1-4(W/O2): 87; 1-5(W/OG): 134; 1-6(W/W): 70; 2-3(O2/OD1): 120; 2-4(O2/O2): 81; 2-5(O2/OG): 104; 2-6(O2/W): 159; 3-4(OD1/O2): 156; 3-5(OD1/OG): 84; 3-6(OD1/W): 78; 4-5(O2/OG): 80; 4-6(O2/W): 79; 5-6(OG/W): 65;   
• 1-2(W/O2): 4.6; 1-3(W/OD1): 3.9; 1-4(W/O2): 3.8; 1-5(W/OG): 5.2; 1-6(W/W): 3.3; 2-3(O2/OD1): 4.8; 2-4(O2/O2): 3.7; 2-5(O2/OG): 4.6; 2-6(O2/W): 5.8; 3-4(OD1/O2): 5.6; 3-5(OD1/OG): 3.9; 3-6(OD1/W): 3.8; 4-5(O2/OG): 3.8; 4-6(O2/W): 3.9; 5-6(OG/W): 3.3;   
  
**3.98 ± 1.18: the average coodination distance of the 1^st^ six ligands could correspond to an hexahydrated Mg^2+^.**  
**6: Number of potential Water contacts**   
**0 : Number of second shell contacts involving water or ions**   
**0%: Percentage of satisfied second shell Water contacts**   
• Contact(s): /O2/O2/O_back_/O_w_/OD1_Asn_/OG_Ser_/  
• Contact code: 2O_b_O_back_O_cno_OH_prot_O_w_  
Go to ToC

### K:a:3059

|  | Res. type | Atom | Residue | Distance | Occupancy | B-factor | Closest atom(s) to a 1^st^ shell water | EM\_dens. |
| --- | --- | --- | --- | --- | --- | --- | --- | --- |
| **Ion** | | | |  | 1.0 | 30.0 |  | 18.85 |
 1 | U | O5' | a:1975 | 2.82 | 1.0 | 32.5 |  | 11.56 | 2 | U | OP2 | a:1975 | 2.84 | 1.0 | 32.4 |  | 11.06 | 3 | Water 1 | | a:9326 | 2.95 | 1.0 | 41.4 |  | 4.18 | 4 | G | O2' | a:1828 | 2.96 | 1.0 | 25.9 |  | 15.26 | 5 | Water 2 | | a:4622 | 3.03 | 1.0 | 43.3 |  | 4.94 | 6 | C | OP1 | a:1906 | 3.04 | 1.0 | 36.9 |  | 9.03 | 7 | U | O4' | a:1975 | 3.05 | 1.0 | 33.9 |  | 8.96 | 8 | Water 3 | | a:9639 | 3.34 | 1.0 | 43.8 |  | 3.68 | 9 | G | O3' | a:1828 | 3.93 | 1.0 | 23.6 |  | 16.29 | 10 | Water 4 | | a:6096 | 4.26 | 1.0 | 45.7 |  | 3.95 | 11 | U | OP1 | a:1975 | 4.33 | 1.0 | 36.4 |  | 8.58 | 12 | Water 5 | | a:7996 | 4.43 | 1.0 | 40.6 |  | 5.39 | 13 | Water 6 | | a:5451 | 4.58 | 1.0 | 29.8 |  | 11.18 | 14 | A | O3' | a:1974 | 4.60 | 1.0 | 29.8 |  | 12.94 | 15 | Water 7 | | a:7500 | 4.64 | 1.0 | 36.0 |  | 6.64 | 16 | G | O4' | a:1828 | 4.74 | 1.0 | 25.2 |  | 15.03 | 17 | U | N1 | a:1975 | 4.79 | 1.0 | 37.3 |  | 10.03 |

  
**1^st^ shell coordination angles and distances (*trans* in "blue"; *cys* in "green"; deviations in "orange"):**   
• 1-2(O5'/OP2): 52; 1-3(O5'/W): 163; 1-4(O5'/O2'): 71; 1-5(O5'/W): 116; 1-6(O5'/OP1): 135; 1-7(O5'/O4'): 58; 2-3(OP2/W): 142; 2-4(OP2/O2'): 71; 2-5(OP2/W): 122; 2-6(OP2/OP1): 90; 2-7(OP2/O4'): 105; 3-4(W/O2'): 103; 3-5(W/W): 67; 3-6(W/OP1): 61; 3-7(W/O4'): 112; 4-5(O2'/W): 168; 4-6(O2'/OP1): 122; 4-7(O2'/O4'): 109; 5-6(W/OP1): 62; 5-7(W/O4'): 70; 6-7(OP1/O4'): 130;   
• 1-2(O5'/OP2): 2.5; 1-3(O5'/W): 5.7; 1-4(O5'/O2'): 3.4; 1-5(O5'/W): 5.0; 1-6(O5'/OP1): 5.4; 1-7(O5'/O4'): 2.8; 2-3(OP2/W): 5.5; 2-4(OP2/O2'): 3.4; 2-5(OP2/W): 5.1; 2-6(OP2/OP1): 4.2; 2-7(OP2/O4'): 4.7; 3-4(W/O2'): 4.6; 3-5(W/W): 3.3; 3-6(W/OP1): 3.1; 3-7(W/O4'): 5.0; 4-5(O2'/W): 6.0; 4-6(O2'/OP1): 5.2; 4-7(O2'/O4'): 4.9; 5-6(W/OP1): 3.1; 5-7(W/O4'): 3.5; 6-7(OP1/O4'): 5.5;   
  
**4.01 ± 1.17: the average coodination distance of the 1^st^ six ligands could correspond to an hexahydrated Mg^2+^.**  
**6: Number of potential Water contacts**   
**0 : Number of second shell contacts involving water or ions**   
**0%: Percentage of satisfied second shell Water contacts**   
• Contact(s): /O2'/O4'/O5'/O_w_/O_w_/OP1/OP2/  
• Contact code: 2O_ph_3O_r_2O_w_  
Go to ToC

### K:a:3060

|  | Res. type | Atom | Residue | Distance | Occupancy | B-factor | Closest atom(s) to a 1^st^ shell water | EM\_dens. |
| --- | --- | --- | --- | --- | --- | --- | --- | --- |
| **Ion** | | | |  | 1.0 | 30.0 |  | 11.63 |
 1 | G | O6 | a:1220 | 2.62 | 1.0 | 38.4 |  | 9.66 | 2 | Water 1 | | a:5371 | 2.84 | 1.0 | 41.0 | 2.70 [G:N7]a:1220 | 5.59 | 3 | U | O4 | a:1219 | 2.91 | 1.0 | 36.3 |  | 9.84 | 4 | U | O4 | a:1233 | 2.94 | 1.0 | 42.3 |  | 7.19 | 5 | Water 2 | | a:5020 | 3.53 | 1.0 | 43.7 |  | 4.59 | 6 | Water 3 | | a:10180 | 3.58 | 1.0 | 45.2 |  | 3.70 | 7 | U | N3 | a:1233 | 3.83 | 1.0 | 29.6 |  | 8.85 | 8 | A | N6 | a:1232 | 3.94 | 1.0 | 30.8 |  | 7.69 | 9 | G | N7 | a:1220 | 3.98 | 1.0 | 27.9 |  | 7.85 | 10 | G | O6 | a:1234 | 4.10 | 1.0 | 32.8 |  | 9.95 | 11 | Water 4 | | a:9392 | 4.51 | 1.0 | 40.9 |  | 4.68 | 12 | U | O4 | a:1221 | 4.58 | 1.0 | 39.4 |  | 6.86 | 13 | Water 5 | | a:10028 | 4.62 | 1.0 | 50.8 |  | 4.87 |

  
**1^st^ shell coordination angles and distances (*trans* in "blue"; *cys* in "green"; deviations in "orange"):**   
• 1-2(O6/W): 83; 1-3(O6/O4): 110; 1-4(O6/O4): 81; 2-3(W/O4): 71; 2-4(W/O4): 148; 3-4(O4/O4): 141;   
• 1-2(O6/W): 3.6; 1-3(O6/O4): 4.5; 1-4(O6/O4): 3.6; 2-3(W/O4): 3.3; 2-4(W/O4): 5.6; 3-4(O4/O4): 5.5;   
  
**4.36 ± 1.37: the average coodination distance of the 1^st^ six ligands could correspond to an hexahydrated Mg^2+^.**  
**6: Number of potential Water contacts**   
**1 : Number of second shell contacts involving water or ions**   
**17%: Percentage of satisfied second shell Water contacts**   
• Contact(s): /O4/O4/O6/O_w_/  
• Contact code: 3O_b_O_w_  
→ Coordination < 6 for K ←  
Go to ToC

### K:a:3061

|  | Res. type | Atom | Residue | Distance | Occupancy | B-factor | Closest atom(s) to a 1^st^ shell water | EM\_dens. |
| --- | --- | --- | --- | --- | --- | --- | --- | --- |
| **Ion** | | | |  | 1.0 | 30.0 |  | 19.06 |
 1 | G | O6 | a:1977 | 2.73 | 1.0 | 29.9 |  | 14.28 | 2 | G | O6 | a:1833 | 2.74 | 1.0 | 33.4 |  | 13.31 | 3 | Water 1 | | a:5077 | 2.84 | 1.0 | 37.6 |  | 8.46 | 4 | Water 2 | | a:8222 | 3.10 | 1.0 | 33.1 | 2.90 [G:N7]a:1833 | 9.68 | 5 | G | O6 | a:1976 | 3.16 | 1.0 | 27.9 |  | 13.27 | 6 | C | N4 | a:1834 | 3.73 | 1.0 | 20.2 |  | 12.51 | 7 | C | N4 | a:1978 | 3.75 | 1.0 | 18.7 |  | 13.24 | 8 | C | N4 | a:1832 | 3.81 | 1.0 | 24.5 |  | 12.82 | 9 | Water 3 | | a:6771 | 3.82 | 1.0 | 37.5 | 2.79 [G:N7]a:1977 | 7.17 | 10 | Water 4 | | a:8223 | 4.00 | 1.0 | 31.0 |  | 9.85 | 11 | Water 5 | | a:9027 | 4.46 | 1.0 | 42.0 | 3.00 [A:OP1]a:1831 | 6.59 | 12 | A | OP1 | a:1831 | 4.50 | 1.0 | 32.3 |  | 9.28 | 13 | G | N7 | a:1977 | 4.57 | 1.0 | 25.1 |  | 12.87 | 14 | G | N7 | a:1833 | 4.79 | 1.0 | 24.7 |  | 11.61 |

  
**1^st^ shell coordination angles and distances (*trans* in "blue"; *cys* in "green"; deviations in "orange"):**   
• 1-2(O6/O6): 68; 1-3(O6/W): 125; 1-4(O6/W): 106; 1-5(O6/O6): 80; 2-3(O6/W): 129; 2-4(O6/W): 71; 2-5(O6/O6): 139; 3-4(W/W): 59; 3-5(W/O6): 53; 4-5(W/O6): 95;   
• 1-2(O6/O6): 3.1; 1-3(O6/W): 4.9; 1-4(O6/W): 4.7; 1-5(O6/O6): 3.8; 2-3(O6/W): 5.0; 2-4(O6/W): 3.4; 2-5(O6/O6): 5.5; 3-4(W/W): 2.9; 3-5(W/O6): 2.7; 4-5(W/O6): 4.6;   
  
**4.31 ± 1.34: the average coodination distance of the 1^st^ six ligands could correspond to an hexahydrated Mg^2+^.**  
**4: Number of potential Water contacts**   
**1 : Number of second shell contacts involving water or ions**   
**25%: Percentage of satisfied second shell Water contacts**   
• Contact(s): /O6/O6/O6/O_w_/O_w_/  
• Contact code: 3O_b_2O_w_  
→ Coordination < 6 for K ←  
Go to ToC

### K:a:3062

|  | Res. type | Atom | Residue | Distance | Occupancy | B-factor | Closest atom(s) to a 1^st^ shell water | EM\_dens. |
| --- | --- | --- | --- | --- | --- | --- | --- | --- |
| **Ion** | | | |  | 1.0 | 30.0 |  | 15.96 |
 1 | G | O6 | a:1458 | 2.58 | 1.0 | 37.2 |  | 9.07 | 2 | Water 1 | | a:8062 | 2.90 | 1.0 | 41.0 | 2.19 [MG:MG]a:3357 | 7.21 | 3 | A | OP1 | a:1455 | 2.91 | 1.0 | 41.4 |  | 6.53 | 4 | G | O2' | a:1454 | 2.92 | 1.0 | 41.7 |  | 5.94 | 5 | U | O4 | a:1459 | 3.00 | 1.0 | 42.7 |  | 8.02 | 6 | Water 2 | | a:6250 | 3.01 | 1.0 | 43.5 | 2.76 [G:O6]a:1457 | 5.38 | 7 | A | OP2 | a:1455 | 3.71 | 1.0 | 47.4 |  | 3.29 | 8 | Water 3 | | a:5261 | 3.72 | 1.0 | 35.5 | 2.05 [MG:MG]a:3357 | 8.07 | 9 | Water 4 | | a:3764 | 4.00 | 1.0 | 38.4 | 2.24 [MG:MG]a:3357 | 8.06 | 10 | G | N7 | a:1458 | 4.16 | 1.0 | 33.0 |  | 9.47 | 11 | G | O6 | a:2706 | 4.17 | 1.0 | 37.4 |  | 9.40 | 12 | G | O3' | a:1454 | 4.24 | 1.0 | 48.1 |  | 5.81 | 13 | **MG** | MG | a:3357 | 4.41 | 1.0 | 42.8 |  | 8.99 | 14 | C | O2' | a:1456 | 4.50 | 1.0 | 41.2 |  | 8.09 | 15 | G | O6 | a:1457 | 4.66 | 1.0 | 37.6 |  | 7.62 |

  
**1^st^ shell coordination angles and distances (*trans* in "blue"; *cys* in "green"; deviations in "orange"):**   
• 1-2(O6/W): 90; 1-3(O6/OP1): 174; 1-4(O6/O2'): 106; 1-5(O6/O4): 74; 1-6(O6/W): 79; 2-3(W/OP1): 86; 2-4(W/O2'): 147; 2-5(W/O4): 81; 2-6(W/W): 74; 3-4(OP1/O2'): 75; 3-5(OP1/O4): 101; 3-6(OP1/W): 104; 4-5(O2'/O4): 76; 4-6(O2'/W): 136; 5-6(O4/W): 143;   
• 1-2(O6/W): 3.9; 1-3(O6/OP1): 5.5; 1-4(O6/O2'): 4.4; 1-5(O6/O4): 3.4; 1-6(O6/W): 3.6; 2-3(W/OP1): 4.0; 2-4(W/O2'): 5.6; 2-5(W/O4): 3.8; 2-6(W/W): 3.6; 3-4(OP1/O2'): 3.6; 3-5(OP1/O4): 4.6; 3-6(OP1/W): 4.7; 4-5(O2'/O4): 3.7; 4-6(O2'/W): 5.5; 5-6(O4/W): 5.7;   
  
**4.13 ± 1.27: the average coodination distance of the 1^st^ six ligands could correspond to an hexahydrated Mg^2+^.**  
**6: Number of potential Water contacts**   
**3 : Number of second shell contacts involving water or ions**   
**50%: Percentage of satisfied second shell Water contacts**   
• Contact(s): /O2'/O4/O_w_/O_w_/OP1/  
• Contact code: O_ph_O_r_O_b_2O_w_  
• Contact(s) in the exclusion zone: /O6/  
→ Coordination < 6 for K ←  
Go to ToC

### K:a:3063

|  | Res. type | Atom | Residue | Distance | Occupancy | B-factor | Closest atom(s) to a 1^st^ shell water | EM\_dens. |
| --- | --- | --- | --- | --- | --- | --- | --- | --- |
| **Ion** | | | |  | 1.0 | 30.0 |  | 15.89 |
 1 | **MG** | MG | a:3363 | 2.50 | 1.0 | 52.8 |  | 7.85 | 2 | U | OP2 | a:2558 | 2.65 | 1.0 | 36.8 |  | 9.74 | 3 | Water 1 | | a:3741 | 2.94 | 1.0 | 39.8 | 2.52 [G:OP2]a:2557 | 6.96 | 4 | A | O3' | a:1674 | 3.01 | 1.0 | 32.1 |  | 11.66 | 5 | OMU | OP2 | a:2556 | 3.02 | 1.0 | 36.0 |  | 6.41 | 6 | G | OP1 | a:1675 | 3.02 | 1.0 | 36.1 |  | 10.99 | 7 | Water 2 | | a:8576 | 3.37 | 1.0 | 41.2 |  | 5.98 | 8 | OMU | O5' | a:2556 | 3.70 | 1.0 | 35.0 |  | 12.39 | 9 | Water 3 | | a:3956 | 3.84 | 1.0 | 38.5 |  | 5.04 | 10 | U | OP1 | a:2558 | 4.09 | 1.0 | 26.5 |  | 11.78 | 11 | G | OP2 | a:2557 | 4.41 | 1.0 | 35.6 |  | 10.27 | 12 | A | O2' | a:1674 | 4.45 | 1.0 | 32.3 |  | 10.74 | 13 | G | OP2 | a:1675 | 4.51 | 1.0 | 24.8 |  | 12.44 | 14 | OMU | N1 | a:2556 | 4.75 | 1.0 | 21.1 |  | 21.80 |

  
**1^st^ shell coordination angles and distances (*trans* in "blue"; *cys* in "green"; deviations in "orange"):**   
• 1-2(MG/OP2): 170; 1-3(MG/W): 82; 1-4(MG/O3'): 89; 1-5(MG/OP2): 44; 1-6(MG/OP1): 47; 2-3(OP2/W): 94; 2-4(OP2/O3'): 99; 2-5(OP2/OP2): 127; 2-6(OP2/OP1): 143; 3-4(W/O3'): 70; 3-5(W/OP2): 90; 3-6(W/OP1): 91; 4-5(O3'/OP2): 131; 4-6(O3'/OP1): 49; 5-6(OP2/OP1): 89;   
• 1-2(MG/OP2): 5.1; 1-3(MG/W): 3.6; 1-4(MG/O3'): 3.9; 1-5(MG/OP2): 2.1; 1-6(MG/OP1): 2.3; 2-3(OP2/W): 4.1; 2-4(OP2/O3'): 4.3; 2-5(OP2/OP2): 5.1; 2-6(OP2/OP1): 5.4; 3-4(W/O3'): 3.4; 3-5(W/OP2): 4.2; 3-6(W/OP1): 4.2; 4-5(O3'/OP2): 5.5; 4-6(O3'/OP1): 2.5; 5-6(OP2/OP1): 4.3;   
  
**4.04 ± 1.23: the average coodination distance of the 1^st^ six ligands could correspond to an hexahydrated Mg^2+^.**  
**4: Number of potential Water contacts**   
**1 : Number of second shell contacts involving water or ions**   
**25%: Percentage of satisfied second shell Water contacts**   
• Contact(s): /O3'/O_w_/OP1/OP2/OP2/  
• Contact code: 3O_ph_O_r_O_w_  
→ Coordination < 6 for K ←  
Go to ToC

### K:a:3064

|  | Res. type | Atom | Residue | Distance | Occupancy | B-factor | Closest atom(s) to a 1^st^ shell water | EM\_dens. |
| --- | --- | --- | --- | --- | --- | --- | --- | --- |
| **Ion** | | | |  | 1.0 | 30.0 |  | 13.40 |
 1 | U | O2' | a:2853 | 2.59 | 1.0 | 39.5 |  | 6.24 | 2 | Water 1 | | a:8382 | 2.68 | 1.0 | 49.7 | 2.92 [G:N7]a:2868 | 3.88 | 3 | Water 2 | | a:9545 | 2.78 | 1.0 | 41.0 |  | 4.51 | 4 | U | O4 | a:2869 | 2.82 | 1.0 | 40.5 |  | 9.09 | 5 | A | OP2 | a:2854 | 2.94 | 1.0 | 32.2 |  | 9.67 | 6 | Water 3 | | a:6318 | 3.11 | 1.0 | 47.1 | 2.76 [A:OP1]a:2854 | 4.47 | 7 | A | OP1 | a:2854 | 3.44 | 1.0 | 38.5 |  | 8.01 | 8 | U | O2 | a:2870 | 3.72 | 1.0 | 40.0 |  | 8.14 | 9 | Water 4 | | a:8987 | 3.75 | 1.0 | 43.2 |  | 4.41 | 10 | U | O3' | a:2853 | 3.79 | 1.0 | 34.0 |  | 9.20 | 11 | **K** | K | a:3129 | 4.30 | 1.0 | 30.0 |  | 22.27 | 12 | G | N7 | a:2868 | 4.43 | 1.0 | 34.4 |  | 8.38 |

  
**1^st^ shell coordination angles and distances (*trans* in "blue"; *cys* in "green"; deviations in "orange"):**   
• 1-2(O2'/W): 133; 1-3(O2'/W): 82; 1-4(O2'/O4): 138; 1-5(O2'/OP2): 84; 1-6(O2'/W): 74; 2-3(W/W): 87; 2-4(W/O4): 87; 2-5(W/OP2): 134; 2-6(W/W): 76; 3-4(W/O4): 90; 3-5(W/OP2): 131; 3-6(W/W): 125; 4-5(O4/OP2): 71; 4-6(O4/W): 139; 5-6(OP2/W): 96;   
• 1-2(O2'/W): 4.8; 1-3(O2'/W): 3.5; 1-4(O2'/O4): 5.1; 1-5(O2'/OP2): 3.7; 1-6(O2'/W): 3.4; 2-3(W/W): 3.8; 2-4(W/O4): 3.8; 2-5(W/OP2): 5.2; 2-6(W/W): 3.6; 3-4(W/O4): 3.9; 3-5(W/OP2): 5.2; 3-6(W/W): 5.2; 4-5(O4/OP2): 3.3; 4-6(O4/W): 5.6; 5-6(OP2/W): 4.5;   
  
**4.01 ± 1.23: the average coodination distance of the 1^st^ six ligands could correspond to an hexahydrated Mg^2+^.**  
**6: Number of potential Water contacts**   
**2 : Number of second shell contacts involving water or ions**   
**33%: Percentage of satisfied second shell Water contacts**   
• Contact(s): /O4/O_w_/O_w_/O_w_/OP2/  
• Contact code: O_ph_O_b_3O_w_  
• Contact(s) in the exclusion zone: /O2'/  
→ Coordination < 6 for K ←  
Go to ToC

### MG:a:3065

|  | Res. type | Atom | Residue | Distance | Occupancy | B-factor | Closest atom(s) to a 1^st^ shell water | EM\_dens. |
| --- | --- | --- | --- | --- | --- | --- | --- | --- |
| **Ion** | | | |  | 1.0 | 49.9 |  | 8.47 |
 1 | Water 1 | | a:10283 | 1.99 | 1.0 | 44.6 |  | 4.48 | 2 | Water 2 | | a:5872 | 2.00 | 1.0 | 30.8 | 2.73 [A:OP2]a:962 // 2.86 [G:N7]a:964 | 9.46 | 3 | Water 3 | | a:9098 | 2.09 | 1.0 | 40.2 |  | 4.57 | 4 | Water 4 | | a:9892 | 2.12 | 1.0 | 38.9 |  | 5.15 | 5 | Water 5 | | a:8036 | 2.30 | 1.0 | 37.4 |  | 6.13 | 6 | Water 6 | | a:4328 | 2.40 | 1.0 | 36.5 | 2.61 [C:OP2]a:963 // 2.75 [G:OP2]a:964 | 7.91 | 7 | Water 7 | | a:10204 | 3.76 | 1.0 | 48.8 |  | 4.16 | 8 | C | OP2 | a:963 | 3.86 | 1.0 | 28.6 |  | 9.80 | 9 | Water 8 | | a:5457 | 4.01 | 1.0 | 45.3 |  | 3.48 | 10 | A | OP2 | a:962 | 4.21 | 1.0 | 31.4 |  | 10.71 | 11 | Water 9 | | a:4158 | 4.25 | 1.0 | 42.5 | 2.60 [G:OP2]a:964 | 5.87 | 12 | G | N7 | a:964 | 4.26 | 1.0 | 19.9 |  | 14.96 | 13 | Water 10 | | a:4719 | 4.36 | 1.0 | 29.6 |  | 7.58 | 14 | Water 11 | | a:9478 | 4.45 | 1.0 | 48.8 |  | 4.62 | 15 | G | OP2 | a:964 | 4.57 | 1.0 | 26.0 |  | 12.63 | 16 | A | OP1 | a:962 | 4.71 | 1.0 | 30.7 |  | 10.33 | 17 | A | O5' | a:962 | 4.79 | 1.0 | 33.4 |  | 11.37 |

  
**1^st^ shell coordination angles and distances (*trans* in "blue"; *cys* in "green"; deviations in "orange"):**   
• 1-2(W/W): 92º; 1-3(W/W): 91º; 1-4(W/W): *95º*; 2-3(W/W): 91º; 2-4(W/W): *167º*; 3-4(W/W): *99º*;   
• 1-2(W/W): 2.9Å; 1-3(W/W): 2.9Å; 1-4(W/W): 3.0Å; 2-3(W/W): 2.9Å; 2-4(W/W): 4.1Å; 3-4(W/W): 3.2Å;   
  
**2.15 ± 0.15: the average coodination distance of the 1^st^ six ligands is Mg^2+^ compatible**  
**12: Number of potential Water contacts**   
**4 : Number of second shell contacts involving water or ions**   
**33%: Percentage of satisfied second shell Water contacts**   
• Contact(s): /O_w_/O_w_/O_w_/O_w_/  
• Contact code: 4O_w_  
• Contact(s) in the exclusion zone: /O_w_/O_w_/  
→ Missing octahedral coordination for MG ←  
→ Slightly distorded coord. octahedron (5.2º) ←  
Go to ToC

### MG:a:3066

|  | Res. type | Atom | Residue | Distance | Occupancy | B-factor | Closest atom(s) to a 1^st^ shell water | EM\_dens. |
| --- | --- | --- | --- | --- | --- | --- | --- | --- |
| **Ion** | | | |  | 1.0 | 48.5 |  | 7.62 |
 1 | Water 1 | | a:9710 | 2.18 | 1.0 | 47.1 |  | 4.22 | 2 | Water 2 | | a:3796 | 2.33 | 1.0 | 36.2 | 2.54 [G:O6]a:2260 | 7.37 | 3 | G | OP2 | a:2259 | 2.40 | 1.0 | 45.5 |  | 4.33 | 4 | Water 3 | | a:3842 | 3.92 | 1.0 | 35.9 |  | 6.67 | 5 | G | O5' | a:2259 | 4.00 | 1.0 | 42.2 |  | 6.09 | 6 | Water 4 | | a:6840 | 4.09 | 1.0 | 43.7 | 2.80 [C:OP2]a:2258 | 6.01 | 7 | G | N7 | a:2260 | 4.11 | 1.0 | 28.8 |  | 10.54 | 8 | G | N7 | a:2259 | 4.14 | 1.0 | 28.8 |  | 10.56 | 9 | C | OP2 | a:2258 | 4.15 | 1.0 | 42.2 |  | 6.10 | 10 | Water 5 | | a:9953 | 4.38 | 1.0 | 44.4 |  | 3.84 | 11 | G | O6 | a:2260 | 4.40 | 1.0 | 29.0 |  | 13.36 | 12 | Water 6 | | a:8908 | 4.53 | 1.0 | 43.4 |  | 3.16 | 13 | C | O5' | a:2258 | 4.58 | 1.0 | 41.0 |  | 6.24 | 14 | G | OP1 | a:2259 | 4.66 | 1.0 | 45.0 |  | 2.73 | 15 | C | O3' | a:2258 | 4.77 | 1.0 | 42.4 |  | 5.00 |

  
• • **3.15 ± 0.85: the average coodination distance of the 1^st^ six ligands is K^+^ compatible**  
**8: Number of potential Water contacts**   
**2 : Number of second shell contacts involving water or ions**   
**25%: Percentage of satisfied second shell Water contacts**   
• Contact(s): /O_w_/  
• Contact code: O_w_  
• Contact(s) in the exclusion zone: /O_w_/OP2/  
→ Missing octahedral coordination for MG ←  
Go to ToC

### MG:a:3067

|  | Res. type | Atom | Residue | Distance | Occupancy | B-factor | Closest atom(s) to a 1^st^ shell water | EM\_dens. |
| --- | --- | --- | --- | --- | --- | --- | --- | --- |
| **Ion** | | | |  | 1.0 | 51.9 |  | 8.14 |
 1 | Water 1 | | a:5391 | 1.90 | 1.0 | 37.1 | 2.70 [A:N7]a:663 | 9.27 | 2 | Water 2 | | a:7656 | 1.95 | 1.0 | 38.2 | 2.85 [G:O6]a:664 | 7.32 | 3 | Water 3 | | a:10225 | 2.09 | 1.0 | 45.6 |  | 5.19 | 4 | Water 4 | | a:7719 | 2.14 | 1.0 | 34.0 | 2.86 [G:N7]a:664 | 8.96 | 5 | Water 5 | | a:10068 | 2.23 | 1.0 | 36.7 | 3.00 [K:K]a:3130 | 6.31 | 6 | Water 6 | | a:10291 | 2.25 | 1.0 | 43.7 |  | 4.12 | 7 | Water 7 | | a:4734 | 3.89 | 1.0 | 35.5 |  | 6.31 | 8 | Water 8 | | a:6404 | 3.91 | 1.0 | 36.8 |  | 6.85 | 9 | Water 9 | | a:4346 | 4.23 | 1.0 | 45.2 |  | 4.27 | 10 | A | N7 | a:663 | 4.26 | 1.0 | 18.8 |  | 12.88 | 11 | Water 10 | | a:5350 | 4.31 | 1.0 | 40.0 |  | 5.67 | 12 | G | O6 | a:664 | 4.35 | 1.0 | 26.5 |  | 13.76 | 13 | Water 11 | | a:9993 | 4.35 | 1.0 | 42.8 |  | 5.67 | 14 | Water 12 | | a:4338 | 4.38 | 1.0 | 40.7 |  | 5.15 | 15 | G | N7 | a:664 | 4.44 | 1.0 | 20.6 |  | 12.76 | 16 | **K** | K | a:3130 | 4.44 | 1.0 | 30.0 |  | 21.64 | 17 | Water 13 | | a:9823 | 4.72 | 1.0 | 39.2 |  | 5.60 |

  
**1^st^ shell coordination angles and distances (*trans* in "blue"; *cys* in "green"; deviations in "orange"):**   
• 1-2(W/W): 91º; 1-3(W/W): 95º; 1-4(W/W): 87º; 1-5(W/W): *163º*; 1-6(W/W): *100º*; 2-3(W/W): *164º*; 2-4(W/W): *98º*; 2-5(W/W): 93º; 2-6(W/W): 90º; 3-4(W/W): *96º*; 3-5(W/W): 86º; 3-6(W/W): *75º*; 4-5(W/W): *77º*; 4-6(W/W): *169º*; 5-6(W/W): *96º*;   
• 1-2(W/W): 2.8Å; 1-3(W/W): 2.9Å; 1-4(W/W): 2.8Å; 1-5(W/W): 4.1Å; 1-6(W/W): 3.2Å; 2-3(W/W): 4.0Å; 2-4(W/W): 3.1Å; 2-5(W/W): 3.0Å; 2-6(W/W): 3.0Å; 3-4(W/W): 3.2Å; 3-5(W/W): 2.9Å; 3-6(W/W): 2.6Å; 4-5(W/W): 2.7Å; 4-6(W/W): *4.4Å*; 5-6(W/W): *3.3Å*;   
  
**2.09 ± 0.13: the average coodination distance of the 1^st^ six ligands is Mg^2+^ compatible**  
**12: Number of potential Water contacts**   
**4 : Number of second shell contacts involving water or ions**   
**33%: Percentage of satisfied second shell Water contacts**   
• Contact(s): /O_w_/O_w_/O_w_/O_w_/O_w_/O_w_/  
• Contact code: 6O_w_  
→ Slightly distorded coord. octahedron (7.9º) ←  
Go to ToC

### MG:a:3068

|  | Res. type | Atom | Residue | Distance | Occupancy | B-factor | Closest atom(s) to a 1^st^ shell water | EM\_dens. |
| --- | --- | --- | --- | --- | --- | --- | --- | --- |
| **Ion** | | | |  | 1.0 | 40.5 |  | 11.77 |
 1 | C | OP1 | a:192 | 2.74 | 1.0 | 26.0 |  | 15.75 | 2 | Water 1 | | a:5462 | 2.81 | 1.0 | 43.6 | 2.86 [C:O3']a:680 | 3.99 | 3 | A | O3' | a:191 | 2.99 | 1.0 | 21.8 |  | 15.81 | 4 | Water 2 | | a:10138 | 3.11 | 1.0 | 39.6 |  | 4.64 | 5 | C | O2' | a:680 | 3.21 | 1.0 | 26.6 |  | 14.00 | 6 | U | OP1 | a:2247 | 3.24 | 1.0 | 23.5 |  | 14.54 | 7 | G | O3' | a:2246 | 3.54 | 1.0 | 25.0 |  | 16.15 | 8 | Water 3 | | a:7189 | 3.81 | 1.0 | 28.4 | 1.84 [MG:MG]a:3159 | 10.78 | 9 | Water 4 | | a:9973 | 3.95 | 1.0 | 39.8 |  | 5.43 | 10 | **MG** | MG | a:3336 | 3.95 | 1.0 | 26.0 |  | 25.15 | 11 | Water 5 | | a:7142 | 4.15 | 1.0 | 39.0 | 2.27 [MG:MG]a:3159 | 5.12 | 12 | U | OP2 | a:2247 | 4.24 | 1.0 | 27.0 |  | 11.36 | 13 | C | O5' | a:192 | 4.31 | 1.0 | 28.4 |  | 13.72 | 14 | C | O3' | a:680 | 4.60 | 1.0 | 26.8 |  | 13.76 | 15 | Water 6 | | a:4961 | 4.61 | 1.0 | 22.5 | 2.29 [MG:MG]a:3336 | 16.55 | 16 | **MG** | MG | a:3159 | 4.61 | 1.0 | 44.9 |  | 11.28 | 17 | C | OP2 | a:192 | 4.70 | 1.0 | 22.8 |  | 17.54 | 18 | A | O2' | a:191 | 4.71 | 1.0 | 25.6 |  | 15.44 | 19 | Water 7 | | a:4401 | 4.76 | 1.0 | 19.9 | 2.12 [MG:MG]a:3336 | 16.39 |

  
• • **3.02 ± 0.19: the average coodination distance of the 1^st^ six ligands is K^+^ compatible**  
**4: Number of potential Water contacts**   
**1 : Number of second shell contacts involving water or ions**   
**25%: Percentage of satisfied second shell Water contacts**   
• Contact code: → No contacts ←   
• Contact(s) in the exclusion zone: /O2'/O3'/O_w_/O_w_/OP1/OP1/  
→ Consider K in place of MG (2 valid contact(s) for K vs. 0 for MG ←)  
Go to ToC

### MG:a:3069

|  | Res. type | Atom | Residue | Distance | Occupancy | B-factor | Closest atom(s) to a 1^st^ shell water | EM\_dens. |
| --- | --- | --- | --- | --- | --- | --- | --- | --- |
| **Ion** | | | |  | 1.0 | 44.7 |  | 11.61 |
 1 | Water 1 | | a:6307 | 2.50 | 1.0 | 32.3 | 2.76 [U:OP1]a:743 | 9.43 | 2 | A | OP1 | a:1786 | 2.73 | 1.0 | 26.2 |  | 13.65 | 3 | G | O2' | a:2611 | 2.95 | 1.0 | 27.2 |  | 13.23 | 4 | A | O3' | a:1785 | 3.09 | 1.0 | 24.6 |  | 13.09 | 5 | G | O3' | a:2611 | 3.15 | 1.0 | 32.2 |  | 10.28 | 6 | Water 2 | | a:5282 | 3.18 | 1.0 | 35.3 |  | 8.00 | 7 | Water 3 | | a:9045 | 3.62 | 1.0 | 37.7 | 3.00 [U:OP1]a:743 | 7.04 | 8 | U | OP1 | a:743 | 3.63 | 1.0 | 30.3 |  | 10.46 | 9 | G | OP1 | a:2612 | 4.15 | 1.0 | 36.4 |  | 9.30 | 10 | Water 4 | | a:5246 | 4.18 | 1.0 | 25.8 | 2.85 [A:OP1]a:1786 | 13.26 | 11 | Water 5 | | a:7478 | 4.31 | 1.0 | 42.8 |  | 5.26 | 12 | A | O5' | a:1786 | 4.33 | 1.0 | 30.5 |  | 13.35 | 13 | C | O3' | a:742 | 4.38 | 1.0 | 33.9 |  | 13.05 | 14 | A | O2' | a:1785 | 4.53 | 1.0 | 25.5 |  | 14.44 | 15 | Water 6 | | a:9063 | 4.54 | 1.0 | 45.2 |  | 3.09 | 16 | G | O5' | a:2612 | 4.64 | 1.0 | 35.7 |  | 8.86 | 17 | A | OP2 | a:1786 | 4.74 | 1.0 | 24.1 |  | 14.17 |

  
• • **2.94 ± 0.24: the average coodination distance of the 1^st^ six ligands is K^+^ compatible**  
**4: Number of potential Water contacts**   
**1 : Number of second shell contacts involving water or ions**   
**25%: Percentage of satisfied second shell Water contacts**   
• Contact code: → No contacts ←   
• Contact(s) in the exclusion zone: /O2'/O3'/O3'/O_w_/O_w_/OP1/  
→ Consider K in place of MG (4 valid contact(s) for K vs. 0 for MG ←)  
Go to ToC

### MG:a:3070

|  | Res. type | Atom | Residue | Distance | Occupancy | B-factor | Closest atom(s) to a 1^st^ shell water | EM\_dens. |
| --- | --- | --- | --- | --- | --- | --- | --- | --- |
| **Ion** | | | |  | 1.0 | 44.2 |  | 8.08 |
 1 | Water 1 | | a:4758 | 2.12 | 1.0 | 31.8 | 2.65 [C:OP1]a:1253 | 10.02 | 2 | Water 2 | | a:4066 | 2.68 | 1.0 | 38.2 | 2.58 [C:OP1]a:814 | 5.90 | 3 | G | OP2 | a:1254 | 3.05 | 1.0 | 24.5 |  | 12.12 | 4 | ARG | NE | p:13 | 3.30 | 1.0 | 34.9 |  | 6.71 | 5 | Water 3 | | p:214 | 3.41 | 1.0 | 47.4 | 2.75 [TYR:OH]p:32 | 3.47 | 6 | HIS | NE2 | p:14 | 3.78 | 1.0 | 26.9 |  | 12.03 | 7 | TYR | OH | p:32 | 3.84 | 1.0 | 31.2 |  | 13.07 | 8 | C | OP1 | a:1253 | 3.91 | 1.0 | 26.2 |  | 11.83 | 9 | G | OP1 | a:1254 | 4.11 | 1.0 | 27.8 |  | 12.52 | 10 | C | OP1 | a:814 | 4.40 | 1.0 | 24.7 |  | 13.34 | 11 | Water 4 | | a:5354 | 4.50 | 1.0 | 26.0 | 2.90 [C:OP1]a:814 | 13.16 | 12 | Water 5 | | p:246 | 4.73 | 1.0 | 29.2 |  | 12.80 | 13 | C | O5' | a:1253 | 4.76 | 1.0 | 26.5 |  | 15.30 | 14 | C | O3' | a:1253 | 4.80 | 1.0 | 28.3 |  | 15.86 |

  
• • **3.06 ± 0.53: the average coodination distance of the 1^st^ six ligands is K^+^ compatible**  
**6: Number of potential Water contacts**   
**3 : Number of second shell contacts involving water or ions**   
**50%: Percentage of satisfied second shell Water contacts**   
• Contact(s): /O_w_/  
• Contact code: O_w_  
• Contact(s) in the exclusion zone: /NE_Arg_/O_w_/OP2/  
→ Consider K in place of MG (1 valid contact(s) for K vs. 0 for MG ←)  
→ Missing octahedral coordination for MG ←  
Go to ToC

### MG:a:3071

|  | Res. type | Atom | Residue | Distance | Occupancy | B-factor | Closest atom(s) to a 1^st^ shell water | EM\_dens. |
| --- | --- | --- | --- | --- | --- | --- | --- | --- |
| **Ion** | | | |  | 1.0 | 44.4 |  | 7.26 |
 1 | Water 1 | | a:5834 | 1.78 | 1.0 | 40.0 | 2.73 [C:O2']a:2059 // 2.77 [PSU:O2']a:2508 | 7.81 | 2 | Water 2 | | a:9602 | 2.70 | 1.0 | 41.1 |  | 6.78 | 3 | Water 3 | | a:6426 | 2.90 | 1.0 | 29.3 |  | 10.02 | 4 | Water 4 | | a:8013 | 2.90 | 1.0 | 43.3 | 2.88 [PSU:O2']a:2508 | 5.40 | 5 | C | O3' | a:2059 | 2.91 | 1.0 | 29.1 |  | 14.03 | 6 | C | O2' | a:2059 | 3.06 | 1.0 | 22.4 |  | 18.56 | 7 | PSU | O3' | a:2508 | 3.44 | 1.0 | 37.7 |  | 12.59 | 8 | G | OP2 | a:2060 | 3.45 | 1.0 | 26.2 |  | 15.08 | 9 | PSU | O2' | a:2508 | 3.74 | 1.0 | 35.6 |  | 7.58 | 10 | Water 5 | | a:10270 | 3.99 | 1.0 | 47.5 |  | 4.97 | 11 | G | OP1 | a:2509 | 4.56 | 1.0 | 47.7 |  | 2.62 | 12 | G | O5' | a:2060 | 4.61 | 1.0 | 24.6 |  | 14.20 | 13 | Water 6 | | a:8044 | 4.62 | 1.0 | 27.8 | 2.89 [G:OP2]a:2060 | 14.68 | 14 | PSU | O4' | a:2508 | 4.65 | 1.0 | 34.2 |  | 10.21 | 15 | A | OP1 | a:2058 | 4.69 | 1.0 | 25.7 |  | 15.34 |

  
• • **2.71 ± 0.43: the average coodination distance of the 1^st^ six ligands is K^+^ compatible**  
**8: Number of potential Water contacts**   
**3 : Number of second shell contacts involving water or ions**   
**38%: Percentage of satisfied second shell Water contacts**   
• Contact code: → No contacts ←   
• Contact(s) in the exclusion zone: /O2'/O3'/O_w_/O_w_/O_w_/O_w_/  
→ Consider K in place of MG (2 valid contact(s) for K vs. 0 for MG ←)  
Go to ToC

### MG:a:3072

|  | Res. type | Atom | Residue | Distance | Occupancy | B-factor | Closest atom(s) to a 1^st^ shell water | EM\_dens. |
| --- | --- | --- | --- | --- | --- | --- | --- | --- |
| **Ion** | | | |  | 1.0 | 46.5 |  | 10.50 |
 1 | Water 1 | | a:9647 | 2.12 | 1.0 | 32.0 |  | 7.54 | 2 | G | OP1 | a:955 | 2.18 | 1.0 | 31.3 |  | 7.89 | 3 | Water 2 | | a:9161 | 2.29 | 1.0 | 34.2 |  | 6.60 | 4 | G | O5' | a:955 | 3.62 | 1.0 | 28.7 |  | 9.96 | 5 | Water 3 | | a:9650 | 4.13 | 1.0 | 38.5 |  | 5.82 | 6 | G | O3' | a:954 | 4.26 | 1.0 | 30.8 |  | 11.21 | 7 | Water 4 | | l:342 | 4.29 | 1.0 | 43.6 |  | 4.84 | 8 | Water 5 | | a:10003 | 4.32 | 1.0 | 39.0 |  | 5.92 | 9 | Water 6 | | a:8097 | 4.36 | 1.0 | 40.4 |  | 7.19 | 10 | Water 7 | | a:9048 | 4.46 | 1.0 | 46.4 |  | 4.65 | 11 | G | OP2 | a:955 | 4.61 | 1.0 | 34.5 |  | 9.04 | 12 | Water 8 | | a:4464 | 4.62 | 1.0 | 33.6 |  | 8.99 | 13 | Water 9 | | a:8045 | 4.63 | 1.0 | 33.9 |  | 8.82 | 14 | A | OP1 | a:863 | 4.64 | 1.0 | 29.9 |  | 10.01 | 15 | Water 10 | | a:9390 | 4.78 | 1.0 | 41.0 |  | 5.10 |

  
**1^st^ shell coordination angles and distances (*trans* in "blue"; *cys* in "green"; deviations in "orange"):**   
• 1-2(W/OP1): *95º*; 1-3(W/W): 86º; 2-3(OP1/W): *96º*;   
• 1-2(W/OP1): 3.2Å; 1-3(W/W): 3.0Å; 2-3(OP1/W): *3.3Å*;   
  
**3.10 ± 0.92: the average coodination distance of the 1^st^ six ligands is K^+^ compatible**  
**6: Number of potential Water contacts**   
**0 : Number of second shell contacts involving water or ions**   
**0%: Percentage of satisfied second shell Water contacts**   
• Contact(s): /O_w_/O_w_/OP1/  
• Contact code: O_ph_2O_w_  
→ Missing octahedral coordination for MG ←  
Go to ToC

### MG:a:3073

|  | Res. type | Atom | Residue | Distance | Occupancy | B-factor | Closest atom(s) to a 1^st^ shell water | EM\_dens. |
| --- | --- | --- | --- | --- | --- | --- | --- | --- |
| **Ion** | | | |  | 1.0 | 52.9 |  | 6.78 |
 1 | Water 1 | | a:7440 | 2.28 | 1.0 | 36.5 | 2.84 [G:O6]a:1934 | 5.91 | 2 | Water 2 | | a:6941 | 2.34 | 1.0 | 37.5 | 2.80 [G:OP1]a:1972 | 7.51 | 3 | Water 3 | | a:10131 | 2.36 | 1.0 | 44.7 |  | 4.55 | 4 | G | OP2 | a:1972 | 2.38 | 1.0 | 32.3 |  | 10.06 | 5 | G | OP1 | a:1972 | 3.19 | 1.0 | 27.0 |  | 10.42 | 6 | G | O5' | a:1972 | 4.14 | 1.0 | 27.8 |  | 14.37 | 7 | U | OP2 | a:2597 | 4.20 | 1.0 | 30.8 |  | 9.95 | 8 | Water 4 | | a:9416 | 4.41 | 1.0 | 46.0 |  | 4.30 | 9 | C | O3' | a:1971 | 4.48 | 1.0 | 33.0 |  | 11.78 | 10 | Water 5 | | a:7782 | 4.64 | 1.0 | 41.7 |  | 6.14 | 11 | Water 6 | | a:7664 | 4.65 | 1.0 | 36.2 | 2.89 [U:OP2]a:2597 | 6.19 | 12 | Water 7 | | a:8801 | 4.66 | 1.0 | 33.8 | 2.97 [G:OP1]a:2596 | 8.49 | 13 | G | OP1 | a:2596 | 4.72 | 1.0 | 26.6 |  | 13.05 | 14 | G | O6 | a:1934 | 4.79 | 1.0 | 34.3 |  | 10.05 | 15 | A | OP2 | a:1973 | 4.79 | 1.0 | 26.8 |  | 11.77 |

  
• • **2.78 ± 0.68: the average coodination distance of the 1^st^ six ligands is K^+^ compatible**  
**6: Number of potential Water contacts**   
**2 : Number of second shell contacts involving water or ions**   
**33%: Percentage of satisfied second shell Water contacts**   
• Contact(s): /O_w_/  
• Contact code: O_w_  
• Contact(s) in the exclusion zone: /O_w_/O_w_/OP1/OP2/  
→ Consider K in place of MG (1 valid contact(s) for K vs. 0 for MG ←)  
→ Missing octahedral coordination for MG ←  
Go to ToC

### MG:a:3074

|  | Res. type | Atom | Residue | Distance | Occupancy | B-factor | Closest atom(s) to a 1^st^ shell water | EM\_dens. |
| --- | --- | --- | --- | --- | --- | --- | --- | --- |
| **Ion** | | | |  | 1.0 | 50.5 |  | 6.26 |
 1 | Water 1 | | a:8939 | 1.79 | 1.0 | 37.6 | 2.98 [A:OP2]a:1636 | 5.51 | 2 | Water 2 | | a:10075 | 1.81 | 1.0 | 39.5 |  | 5.09 | 3 | Water 3 | | a:8453 | 1.88 | 1.0 | 36.7 | 2.92 [G:N7]a:1630 | 7.85 | 4 | Water 4 | | a:3910 | 2.16 | 1.0 | 31.0 | 2.56 [G:O6]a:1630 | 7.69 | 5 | Water 5 | | a:3611 | 2.66 | 1.0 | 36.5 | 2.49 [U:O4]a:1631 | 7.63 | 6 | Water 6 | | a:5228 | 3.39 | 1.0 | 32.4 |  | 8.16 | 7 | Water 7 | | a:3858 | 3.78 | 1.0 | 41.7 |  | 3.91 | 8 | U | O4 | a:1631 | 4.24 | 1.0 | 31.2 |  | 12.41 | 9 | G | O6 | a:1630 | 4.26 | 1.0 | 29.5 |  | 13.72 | 10 | Water 8 | | a:3817 | 4.27 | 1.0 | 40.2 |  | 5.87 | 11 | Water 9 | | a:9804 | 4.29 | 1.0 | 36.0 |  | 6.21 | 12 | G | N7 | a:1630 | 4.33 | 1.0 | 25.6 |  | 12.13 | 13 | Water 10 | | a:8538 | 4.44 | 1.0 | 42.1 |  | 4.37 | 14 | A | OP2 | a:1636 | 4.64 | 1.0 | 34.1 |  | 8.45 | 15 | Water 11 | | a:4416 | 4.74 | 1.0 | 38.0 |  | 8.23 |

  
**1^st^ shell coordination angles and distances (*trans* in "blue"; *cys* in "green"; deviations in "orange"):**   
• 1-2(W/W): *84º*; 1-3(W/W): *157º*; 1-4(W/W): *105º*; 2-3(W/W): *103º*; 2-4(W/W): *103º*; 3-4(W/W): 94º;   
• 1-2(W/W): 2.4Å; 1-3(W/W): *3.6Å*; 1-4(W/W): 3.2Å; 2-3(W/W): 2.9Å; 2-4(W/W): 3.1Å; 3-4(W/W): 3.0Å;   
  
**2.28 ± 0.58: the average coodination distance of the 1^st^ six ligands is Mg^2+^ compatible**  
**12: Number of potential Water contacts**   
**4 : Number of second shell contacts involving water or ions**   
**33%: Percentage of satisfied second shell Water contacts**   
• Contact(s): /O_w_/  
• Contact code: O_w_  
• Contact(s) in the exclusion zone: /O_w_/O_w_/O_w_/O_w_/O_w_/  
→ Missing octahedral coordination for MG ←  
→→→ Distorded coord. octahedron (12.4º) ←←←  
Go to ToC

### MG:a:3075

|  | Res. type | Atom | Residue | Distance | Occupancy | B-factor | Closest atom(s) to a 1^st^ shell water | EM\_dens. |
| --- | --- | --- | --- | --- | --- | --- | --- | --- |
| **Ion** | | | |  | 1.0 | 47.2 |  | 6.54 |
 1 | Water 1 | | a:5332 | 1.95 | 1.0 | 41.3 | 2.70 [G:O6]a:2629 | 7.05 | 2 | Water 2 | | a:4633 | 2.20 | 1.0 | 43.9 | 2.64 [G:O6]a:2781 | 5.37 | 3 | Water 3 | | a:5858 | 2.24 | 1.0 | 33.3 | 2.73 [G:N7]a:2629 | 9.11 | 4 | Water 4 | | a:4450 | 2.30 | 1.0 | 33.7 | 2.63 [G:N7]a:2628 | 7.59 | 5 | G | O6 | a:2629 | 4.16 | 1.0 | 24.2 |  | 13.94 | 6 | G | N7 | a:2628 | 4.18 | 1.0 | 24.8 |  | 13.64 | 7 | Water 5 | | a:6557 | 4.27 | 1.0 | 39.5 |  | 6.25 | 8 | Water 6 | | a:4977 | 4.38 | 1.0 | 42.8 |  | 4.18 | 9 | Water 7 | | a:8084 | 4.38 | 1.0 | 37.5 |  | 6.96 | 10 | G | N7 | a:2629 | 4.48 | 1.0 | 22.1 |  | 13.94 | 11 | G | O6 | a:2628 | 4.51 | 1.0 | 26.3 |  | 13.26 | 12 | G | O6 | a:2781 | 4.71 | 1.0 | 26.7 |  | 14.54 | 13 | Water 8 | | a:9442 | 4.72 | 1.0 | 41.7 |  | 3.84 |

  
**1^st^ shell coordination angles and distances (*trans* in "blue"; *cys* in "green"; deviations in "orange"):**   
• 1-2(W/W): *79º*; 1-3(W/W): *84º*; 2-3(W/W): 92º;   
• 1-2(W/W): 2.6Å; 1-3(W/W): 2.8Å; 2-3(W/W): 3.2Å;   
  
**2.84 ± 0.95: the average coodination distance of the 1^st^ six ligands is K^+^ compatible**  
**8: Number of potential Water contacts**   
**4 : Number of second shell contacts involving water or ions**   
**50%: Percentage of satisfied second shell Water contacts**   
• Contact(s): /O_w_/O_w_/O_w_/  
• Contact code: 3O_w_  
• Contact(s) in the exclusion zone: /O_w_/  
→ Missing octahedral coordination for MG ←  
Go to ToC

### MG:a:3076

|  | Res. type | Atom | Residue | Distance | Occupancy | B-factor | Closest atom(s) to a 1^st^ shell water | EM\_dens. |
| --- | --- | --- | --- | --- | --- | --- | --- | --- |
| **Ion** | | | |  | 1.0 | 53.4 |  | 6.14 |
 1 | Water 1 | | a:6497 | 1.98 | 1.0 | 40.0 | 2.77 [G:O6]a:925 // 2.85 [G:O6]a:926 | 5.08 | 2 | Water 2 | | a:10272 | 2.01 | 1.0 | 43.4 |  | 4.49 | 3 | Water 3 | | a:10269 | 2.04 | 1.0 | 48.2 |  | 4.31 | 4 | Water 4 | | a:5191 | 2.27 | 1.0 | 39.3 | 2.69 [G:N7]a:925 | 5.51 | 5 | Water 5 | | a:6805 | 2.33 | 1.0 | 43.3 | 2.79 [G:N7]a:926 | 4.63 | 6 | Water 6 | | a:9101 | 4.20 | 1.0 | 45.5 |  | 4.22 | 7 | G | N7 | a:925 | 4.23 | 1.0 | 28.3 |  | 10.02 | 8 | G | O6 | a:926 | 4.29 | 1.0 | 37.4 |  | 6.70 | 9 | Water 7 | | a:5601 | 4.32 | 1.0 | 38.7 |  | 6.85 | 10 | G | O6 | a:925 | 4.32 | 1.0 | 39.7 |  | 7.65 | 11 | Water 8 | | a:9732 | 4.52 | 1.0 | 35.5 |  | 6.81 | 12 | G | N7 | a:926 | 4.58 | 1.0 | 36.0 |  | 6.57 |

  
**1^st^ shell coordination angles and distances (*trans* in "blue"; *cys* in "green"; deviations in "orange"):**   
• 1-2(W/W): 92º; 1-3(W/W): *164º*; 1-4(W/W): *80º*; 2-3(W/W): *78º*; 2-4(W/W): 86º; 3-4(W/W): 87º;   
• 1-2(W/W): 2.9Å; 1-3(W/W): 4.0Å; 1-4(W/W): 2.7Å; 2-3(W/W): 2.5Å; 2-4(W/W): 2.9Å; 3-4(W/W): 3.0Å;   
  
**2.47 ± 0.78: the average coodination distance of the 1^st^ six ligands is Na^+^** compatible  
**12: Number of potential Water contacts**   
**4 : Number of second shell contacts involving water or ions**   
**33%: Percentage of satisfied second shell Water contacts**   
• Contact(s): /O_w_/O_w_/O_w_/O_w_/  
• Contact code: 4O_w_  
• Contact(s) in the exclusion zone: /O_w_/  
→ Missing octahedral coordination for MG ←  
→ Slightly distorded coord. octahedron (7.9º) ←  
Go to ToC

### MG:a:3077

|  | Res. type | Atom | Residue | Distance | Occupancy | B-factor | Closest atom(s) to a 1^st^ shell water | EM\_dens. |
| --- | --- | --- | --- | --- | --- | --- | --- | --- |
| **Ion** | | | |  | 1.0 | 31.5 |  | 17.65 |
 1 | Water 1 | | a:8513 | 2.11 | 1.0 | 41.5 | 2.93 [U:O4]a:29 | 5.27 | 2 | Water 2 | | a:3891 | 2.12 | 1.0 | 37.1 | 2.55 [G:O6]a:473 | 7.45 | 3 | Water 3 | | a:9836 | 2.17 | 1.0 | 37.9 |  | 8.47 | 4 | Water 4 | | a:5873 | 2.25 | 1.0 | 37.7 | 2.73 [U:OP2]a:29 | 7.62 | 5 | Water 5 | | a:7120 | 2.33 | 1.0 | 42.1 | 2.81 [A:OP2]a:447 | 6.85 | 6 | G | O6 | a:474 | 4.14 | 1.0 | 29.0 |  | 11.89 | 7 | A | OP2 | a:447 | 4.27 | 1.0 | 25.5 |  | 12.71 | 8 | A | OP2 | a:28 | 4.30 | 1.0 | 28.1 |  | 11.61 | 9 | Water 6 | | a:4009 | 4.31 | 1.0 | 38.3 |  | 6.39 | 10 | U | O4 | a:29 | 4.41 | 1.0 | 25.3 |  | 13.84 | 11 | G | O6 | a:30 | 4.44 | 1.0 | 28.5 |  | 12.86 | 12 | G | O6 | a:473 | 4.45 | 1.0 | 30.8 |  | 12.34 | 13 | U | OP2 | a:29 | 4.53 | 1.0 | 25.5 |  | 12.51 | 14 | Water 7 | | a:3673 | 4.59 | 1.0 | 36.8 | 2.51 [A:OP2]a:447 | 6.49 | 15 | Water 8 | | a:6823 | 4.72 | 1.0 | 41.4 | 2.79 [A:OP2]a:28 | 4.00 |

  
**1^st^ shell coordination angles and distances (*trans* in "blue"; *cys* in "green"; deviations in "orange"):**   
• 1-2(W/W): *142º*; 1-3(W/W): 88º; 1-4(W/W): *75º*; 2-3(W/W): 90º; 2-4(W/W): *143º*; 3-4(W/W): 89º;   
• 1-2(W/W): 4.0Å; 1-3(W/W): 3.0Å; 1-4(W/W): 2.7Å; 2-3(W/W): 3.0Å; 2-4(W/W): 4.1Å; 3-4(W/W): 3.1Å;   
  
**2.52 ± 0.73: the average coodination distance of the 1^st^ six ligands is Na^+^** compatible  
**10: Number of potential Water contacts**   
**4 : Number of second shell contacts involving water or ions**   
**40%: Percentage of satisfied second shell Water contacts**   
• Contact(s): /O_w_/O_w_/O_w_/O_w_/  
• Contact code: 4O_w_  
• Contact(s) in the exclusion zone: /O_w_/  
→ Missing octahedral coordination for MG ←  
→→→ Distorded coord. octahedron (15.5º) ←←←  
Go to ToC

### MG:a:3078

|  | Res. type | Atom | Residue | Distance | Occupancy | B-factor | Closest atom(s) to a 1^st^ shell water | EM\_dens. |
| --- | --- | --- | --- | --- | --- | --- | --- | --- |
| **Ion** | | | |  | 1.0 | 32.7 |  | 13.17 |
 1 | U | OP2 | a:120 | 1.94 | 1.0 | 31.9 |  | 10.24 | 2 | Water 1 | | a:9349 | 1.94 | 1.0 | 32.9 | 2.88 [K:K]a:3140 | 9.50 | 3 | Water 2 | | a:8583 | 2.12 | 1.0 | 31.6 | 2.94 [G:O6]a:123 | 8.60 | 4 | Water 3 | | a:4495 | 2.16 | 1.0 | 34.8 | 2.63 [G:OP1]a:121 | 9.58 | 5 | Water 4 | | a:5542 | 2.20 | 1.0 | 34.7 | 2.71 [G:O6]a:122 | 8.89 | 6 | Water 5 | | a:6715 | 2.21 | 1.0 | 32.3 | 2.79 [G:N7]a:122 // 2.85 [G:OP2]a:122 | 12.11 | 7 | **K** | K | a:3140 | 3.57 | 1.0 | 30.0 |  | 18.55 | 8 | U | OP1 | a:120 | 3.90 | 1.0 | 34.6 |  | 8.27 | 9 | A | O3' | a:119 | 4.10 | 1.0 | 36.1 |  | 12.96 | 10 | Water 6 | | a:3856 | 4.11 | 1.0 | 46.9 |  | 4.10 | 11 | U | O5' | a:120 | 4.34 | 1.0 | 36.9 |  | 9.80 | 12 | G | N7 | a:122 | 4.38 | 1.0 | 23.9 |  | 11.10 | 13 | G | OP1 | a:121 | 4.40 | 1.0 | 35.1 |  | 9.45 | 14 | G | O6 | a:122 | 4.49 | 1.0 | 31.0 |  | 13.29 | 15 | Water 7 | | a:6098 | 4.49 | 1.0 | 31.0 | 2.75 [G:OP2]a:122 | 9.65 | 16 | G | O6 | a:123 | 4.57 | 1.0 | 29.0 |  | 11.80 | 17 | G | OP2 | a:122 | 4.63 | 1.0 | 36.1 |  | 8.70 | 18 | Water 8 | | a:5113 | 4.68 | 1.0 | 30.0 | 2.86 [U:OP1]a:120 | 11.55 | 19 | G | N7 | a:121 | 4.75 | 1.0 | 33.8 |  | 9.63 |

  
**1^st^ shell coordination angles and distances (*trans* in "blue"; *cys* in "green"; deviations in "orange"):**   
• 1-2(OP2/W): *104º*; 1-3(OP2/W): 93º; 1-4(OP2/W): *97º*; 1-5(OP2/W): 173º; 1-6(OP2/W): 93º; 2-3(W/W): 88º; 2-4(W/W): 93º; 2-5(W/W): *81º*; 2-6(W/W): *161º*; 3-4(W/W): *169º*; 3-5(W/W): *83º*; 3-6(W/W): *83º*; 4-5(W/W): 86º; 4-6(W/W): 93º; 5-6(W/W): *81º*;   
• 1-2(OP2/W): 3.1Å; 1-3(OP2/W): 2.9Å; 1-4(OP2/W): 3.1Å; 1-5(OP2/W): 4.1Å; 1-6(OP2/W): 3.0Å; 2-3(W/W): 2.8Å; 2-4(W/W): 3.0Å; 2-5(W/W): 2.7Å; 2-6(W/W): 4.1Å; 3-4(W/W): *4.3Å*; 3-5(W/W): 2.9Å; 3-6(W/W): 2.9Å; 4-5(W/W): 3.0Å; 4-6(W/W): 3.2Å; 5-6(W/W): 2.9Å;   
  
**2.10 ± 0.12: the average coodination distance of the 1^st^ six ligands is Mg^2+^ compatible**  
**10: Number of potential Water contacts**   
**6 : Number of second shell contacts involving water or ions**   
**60%: Percentage of satisfied second shell Water contacts**   
• Contact(s): /O_w_/O_w_/O_w_/O_w_/O_w_/OP2/  
• Contact code: O_ph_5O_w_  
→ Slightly distorded coord. octahedron (7.0º) ←  
Go to ToC

### MG:a:3079

|  | Res. type | Atom | Residue | Distance | Occupancy | B-factor | Closest atom(s) to a 1^st^ shell water | EM\_dens. |
| --- | --- | --- | --- | --- | --- | --- | --- | --- |
| **Ion** | | | |  | 1.0 | 35.3 |  | 14.39 |
 1 | U | OP1 | a:120 | 1.95 | 1.0 | 34.6 |  | 8.27 | 2 | Water 1 | | a:7441 | 2.04 | 1.0 | 38.7 |  | 7.51 | 3 | Water 2 | | a:5113 | 2.09 | 1.0 | 30.0 | 2.68 [G:O6]a:124 | 11.55 | 4 | Water 3 | | a:7046 | 2.12 | 1.0 | 33.9 |  | 8.12 | 5 | Water 4 | | a:7995 | 2.38 | 1.0 | 35.5 | 2.88 [G:OP2]a:123 | 8.88 | 6 | Water 5 | | a:6098 | 2.56 | 1.0 | 31.0 | 2.99 [G:OP2]a:123 | 9.65 | 7 | U | O5' | a:120 | 3.86 | 1.0 | 36.9 |  | 9.80 | 8 | G | O6 | a:124 | 4.02 | 1.0 | 26.1 |  | 11.87 | 9 | Water 6 | | a:8583 | 4.02 | 1.0 | 31.6 | 2.95 [U:OP2]a:120 | 8.60 | 10 | Water 7 | | a:4559 | 4.04 | 1.0 | 34.0 |  | 9.41 | 11 | Water 8 | | a:3633 | 4.19 | 1.0 | 45.4 |  | 4.45 | 12 | U | OP2 | a:120 | 4.20 | 1.0 | 31.9 |  | 10.24 | 13 | G | OP2 | a:123 | 4.30 | 1.0 | 33.8 |  | 8.88 | 14 | Water 9 | | a:5736 | 4.30 | 1.0 | 39.9 | 2.98 [G:OP2]a:123 | 4.01 | 15 | A | O3' | a:119 | 4.40 | 1.0 | 36.1 |  | 12.96 | 16 | Water 10 | | a:5600 | 4.52 | 1.0 | 30.8 | 2.89 [G:O6]a:124 | 10.23 | 17 | G | N7 | a:123 | 4.68 | 1.0 | 24.5 |  | 12.77 |

  
**1^st^ shell coordination angles and distances (*trans* in "blue"; *cys* in "green"; deviations in "orange"):**   
• 1-2(OP1/W): 91º; 1-3(OP1/W): 90º; 1-4(OP1/W): 87º; 2-3(W/W): *157º*; 2-4(W/W): *111º*; 3-4(W/W): 91º;   
• 1-2(OP1/W): 2.8Å; 1-3(OP1/W): 2.9Å; 1-4(OP1/W): 2.8Å; 2-3(W/W): 4.0Å; 2-4(W/W): *3.4Å*; 3-4(W/W): 3.0Å;   
  
**2.19 ± 0.21: the average coodination distance of the 1^st^ six ligands is Mg^2+^ compatible**  
**10: Number of potential Water contacts**   
**3 : Number of second shell contacts involving water or ions**   
**30%: Percentage of satisfied second shell Water contacts**   
• Contact(s): /O_w_/O_w_/O_w_/OP1/  
• Contact code: O_ph_3O_w_  
• Contact(s) in the exclusion zone: /O_w_/O_w_/  
→ Missing octahedral coordination for MG ←  
→ Slightly distorded coord. octahedron (8.1º) ←  
Go to ToC

### MG:a:3080

|  | Res. type | Atom | Residue | Distance | Occupancy | B-factor | Closest atom(s) to a 1^st^ shell water | EM\_dens. |
| --- | --- | --- | --- | --- | --- | --- | --- | --- |
| **Ion** | | | |  | 1.0 | 56.3 |  | 8.00 |
 1 | Water 1 | | a:4166 | 1.96 | 1.0 | 39.0 | 2.60 [A:N7]a:219 | 7.53 | 2 | Water 2 | | a:6831 | 2.06 | 1.0 | 46.4 |  | 5.07 | 3 | Water 3 | | a:7638 | 2.10 | 1.0 | 44.2 | 2.85 [A:OP1]a:428 | 5.21 | 4 | Water 4 | | a:9092 | 2.18 | 1.0 | 42.8 |  | 3.86 | 5 | A | OP2 | a:218 | 2.19 | 1.0 | 33.5 |  | 6.72 | 6 | Water 5 | | a:7726 | 2.25 | 1.0 | 46.5 | 2.86 [A:N7]a:218 | 4.78 | 7 | A | O5' | a:218 | 4.02 | 1.0 | 39.2 |  | 6.98 | 8 | A | N7 | a:218 | 4.06 | 1.0 | 31.2 |  | 8.83 | 9 | A | N7 | a:219 | 4.14 | 1.0 | 32.3 |  | 8.82 | 10 | A | OP1 | a:218 | 4.38 | 1.0 | 42.3 |  | 4.42 | 11 | Water 6 | | a:7149 | 4.42 | 1.0 | 43.3 | 2.82 [G:O6]a:220 | 5.30 | 12 | G | O6 | a:220 | 4.44 | 1.0 | 41.3 |  | 7.29 | 13 | A | OP1 | a:428 | 4.52 | 1.0 | 43.6 |  | 4.11 | 14 | A | O3' | a:217 | 4.62 | 1.0 | 40.1 |  | 6.34 | 15 | A | O5' | a:217 | 4.63 | 1.0 | 40.3 |  | 7.02 |

  
**1^st^ shell coordination angles and distances (*trans* in "blue"; *cys* in "green"; deviations in "orange"):**   
• 1-2(W/W): *168º*; 1-3(W/W): *98º*; 1-4(W/W): 93º; 1-5(W/OP2): 92º; 1-6(W/W): *105º*; 2-3(W/W): 86º; 2-4(W/W): *77º*; 2-5(W/OP2): *82º*; 2-6(W/W): 85º; 3-4(W/W): *81º*; 3-5(W/OP2): *165º*; 3-6(W/W): 89º; 4-5(W/OP2): 87º; 4-6(W/W): *160º*; 5-6(OP2/W): *99º*;   
• 1-2(W/W): 4.0Å; 1-3(W/W): 3.1Å; 1-4(W/W): 3.0Å; 1-5(W/OP2): 3.0Å; 1-6(W/W): *3.4Å*; 2-3(W/W): 2.8Å; 2-4(W/W): 2.6Å; 2-5(W/OP2): 2.8Å; 2-6(W/W): 2.9Å; 3-4(W/W): 2.8Å; 3-5(W/OP2): *4.2Å*; 3-6(W/W): 3.1Å; 4-5(W/OP2): 3.0Å; 4-6(W/W): *4.4Å*; 5-6(OP2/W): *3.4Å*;   
  
**2.12 ± 0.10: the average coodination distance of the 1^st^ six ligands is Mg^2+^ compatible**  
**10: Number of potential Water contacts**   
**3 : Number of second shell contacts involving water or ions**   
**30%: Percentage of satisfied second shell Water contacts**   
• Contact(s): /O_w_/O_w_/O_w_/O_w_/O_w_/OP2/  
• Contact code: O_ph_5O_w_  
→ Slightly distorded coord. octahedron (8.4º) ←  
Go to ToC

### MG:a:3081

|  | Res. type | Atom | Residue | Distance | Occupancy | B-factor | Closest atom(s) to a 1^st^ shell water | EM\_dens. |
| --- | --- | --- | --- | --- | --- | --- | --- | --- |
| **Ion** | | | |  | 1.0 | 47.2 |  | 8.66 |
 1 | U | OP1 | a:419 | 1.95 | 1.0 | 37.6 |  | 7.77 | 2 | Water 1 | | a:8048 | 2.03 | 1.0 | 46.0 |  | 4.34 | 3 | Water 2 | | a:4489 | 2.06 | 1.0 | 40.4 | 2.63 [G:N7]a:230 | 5.89 | 4 | Water 3 | | a:7463 | 2.33 | 1.0 | 42.0 |  | 5.08 | 5 | C | O3' | a:418 | 3.77 | 1.0 | 35.7 |  | 8.81 | 6 | U | O5' | a:419 | 4.10 | 1.0 | 39.3 |  | 7.60 | 7 | Water 4 | | a:3506 | 4.12 | 1.0 | 46.0 |  | 3.15 | 8 | G | N7 | a:230 | 4.14 | 1.0 | 38.5 |  | 7.19 | 9 | Water 5 | | a:6097 | 4.31 | 1.0 | 38.8 |  | 6.38 | 10 | U | OP2 | a:419 | 4.45 | 1.0 | 39.7 |  | 6.01 | 11 | Water 6 | | a:7164 | 4.57 | 1.0 | 41.3 |  | 5.39 |

  
**1^st^ shell coordination angles and distances (*trans* in "blue"; *cys* in "green"; deviations in "orange"):**   
• 1-2(OP1/W): 93º; 1-3(OP1/W): *101º*; 2-3(W/W): 92º;   
• 1-2(OP1/W): 2.9Å; 1-3(OP1/W): 3.1Å; 2-3(W/W): 2.9Å;   
  
**2.71 ± 0.88: the average coodination distance of the 1^st^ six ligands is K^+^ compatible**  
**6: Number of potential Water contacts**   
**1 : Number of second shell contacts involving water or ions**   
**17%: Percentage of satisfied second shell Water contacts**   
• Contact(s): /O_w_/O_w_/OP1/  
• Contact code: O_ph_2O_w_  
• Contact(s) in the exclusion zone: /O_w_/  
→ Missing octahedral coordination for MG ←  
Go to ToC

### MG:a:3082

|  | Res. type | Atom | Residue | Distance | Occupancy | B-factor | Closest atom(s) to a 1^st^ shell water | EM\_dens. |
| --- | --- | --- | --- | --- | --- | --- | --- | --- |
| **Ion** | | | |  | 1.0 | 27.0 |  | 23.77 |
 1 | Water 1 | | a:5227 | 1.99 | 1.0 | 24.7 | 2.69 [U:OP2]a:451 // 2.71 [G:OP2]a:452 | 15.28 | 2 | G | OP2 | a:450 | 2.00 | 1.0 | 25.5 |  | 12.36 | 3 | Water 2 | | a:6160 | 2.07 | 1.0 | 25.4 |  | 13.76 | 4 | Water 3 | | e:406 | 2.14 | 1.0 | 22.9 | 2.58 [GLY:O]e:81 | 16.16 | 5 | Water 4 | | e:433 | 2.16 | 1.0 | 29.8 | 2.77 [GLY:O]e:81 | 15.05 | 6 | Water 5 | | a:7599 | 2.27 | 1.0 | 23.6 | 2.85 [G:OP2]a:452 // 2.92 [SER:O]e:80 | 15.23 | 7 | GLY | O | e:81 | 3.72 | 1.0 | 28.5 |  | 15.31 | 8 | Water 6 | | a:3988 | 3.82 | 1.0 | 27.1 | 2.57 [G:OP1]a:450 // 2.73 [U:OP2]a:451 | 13.57 | 9 | Water 7 | | a:4032 | 3.85 | 1.0 | 29.8 | 2.84 [G:OP2]a:452 | 9.70 | 10 | Water 8 | | a:9257 | 3.94 | 1.0 | 36.8 |  | 6.83 | 11 | G | OP2 | a:452 | 4.01 | 1.0 | 30.5 |  | 12.04 | 12 | G | O5' | a:450 | 4.08 | 1.0 | 23.2 |  | 16.61 | 13 | G | OP1 | a:450 | 4.15 | 1.0 | 24.2 |  | 14.28 | 14 | U | OP2 | a:451 | 4.29 | 1.0 | 30.6 |  | 12.90 | 15 | A | OP1 | a:449 | 4.39 | 1.0 | 27.6 |  | 12.71 | 16 | Water 9 | | e:475 | 4.42 | 1.0 | 35.4 |  | 6.73 | 17 | A | O3' | a:449 | 4.44 | 1.0 | 24.8 |  | 12.06 | 18 | SER | O | e:80 | 4.48 | 1.0 | 26.5 |  | 15.83 | 19 | A | OP2 | a:449 | 4.71 | 1.0 | 20.8 |  | 14.29 |

  
**1^st^ shell coordination angles and distances (*trans* in "blue"; *cys* in "green"; deviations in "orange"):**   
• 1-2(W/OP2): *96º*; 1-3(W/W): 91º; 1-4(W/W): *169º*; 1-5(W/W): 88º; 1-6(W/W): *80º*; 2-3(OP2/W): 85º; 2-4(OP2/W): 93º; 2-5(OP2/W): 95º; 2-6(OP2/W): 175º; 3-4(W/W): 94º; 3-5(W/W): 179º; 3-6(W/W): 92º; 4-5(W/W): 86º; 4-6(W/W): 90º; 5-6(W/W): 88º;   
• 1-2(W/OP2): 3.0Å; 1-3(W/W): 2.9Å; 1-4(W/W): 4.1Å; 1-5(W/W): 2.9Å; 1-6(W/W): 2.8Å; 2-3(OP2/W): 2.8Å; 2-4(OP2/W): 3.0Å; 2-5(OP2/W): 3.1Å; 2-6(OP2/W): *4.3Å*; 3-4(W/W): 3.1Å; 3-5(W/W): *4.2Å*; 3-6(W/W): 3.1Å; 4-5(W/W): 2.9Å; 4-6(W/W): 3.1Å; 5-6(W/W): 3.1Å;   
  
**2.10 ± 0.10: the average coodination distance of the 1^st^ six ligands is Mg^2+^ compatible**  
**10: Number of potential Water contacts**   
**6 : Number of second shell contacts involving water or ions**   
**60%: Percentage of satisfied second shell Water contacts**   
• Contact(s): /O_w_/O_w_/O_w_/O_w_/O_w_/OP2/  
• Contact code: O_ph_5O_w_  
Ooctahedral coordination (4.1º)   
Go to ToC

### MG:a:3083

|  | Res. type | Atom | Residue | Distance | Occupancy | B-factor | Closest atom(s) to a 1^st^ shell water | EM\_dens. |
| --- | --- | --- | --- | --- | --- | --- | --- | --- |
| **Ion** | | | |  | 1.0 | 24.4 |  | 24.35 |
 1 | A | OP1 | a:453 | 1.99 | 1.0 | 23.9 |  | 15.02 | 2 | G | O6 | a:450 | 2.06 | 1.0 | 23.6 |  | 17.24 | 3 | Water 1 | | a:6294 | 2.19 | 1.0 | 22.2 | 2.76 [A:N7]a:449 | 19.00 | 4 | Water 2 | | a:8049 | 2.23 | 1.0 | 21.6 | 2.89 [A:OP1]a:454 | 17.37 | 5 | Water 3 | | a:8800 | 2.24 | 1.0 | 22.6 | 2.96 [K:K]a:3100 | 16.66 | 6 | Water 4 | | a:5422 | 2.32 | 1.0 | 21.3 | 2.70 [G:N7]a:450 | 15.55 | 7 | G | N7 | a:450 | 3.87 | 1.0 | 17.1 |  | 15.48 | 8 | A | OP2 | a:453 | 4.00 | 1.0 | 24.9 |  | 15.45 | 9 | A | O5' | a:453 | 4.05 | 1.0 | 22.9 |  | 15.46 | 10 | **K** | K | a:3100 | 4.10 | 1.0 | 30.0 |  | 30.41 | 11 | Water 5 | | a:9924 | 4.18 | 1.0 | 33.2 |  | 10.54 | 12 | Water 6 | | a:3409 | 4.18 | 1.0 | 35.2 |  | 8.58 | 13 | Water 7 | | e:418 | 4.27 | 1.0 | 26.9 | 2.74 [A:OP2]a:449 | 13.62 | 14 | A | N7 | a:449 | 4.32 | 1.0 | 18.9 |  | 16.39 | 15 | G | O3' | a:452 | 4.42 | 1.0 | 25.9 |  | 13.14 | 16 | Water 8 | | e:475 | 4.44 | 1.0 | 35.4 | 2.98 [K:K]a:3100 | 6.73 | 17 | A | OP1 | a:454 | 4.45 | 1.0 | 30.4 |  | 8.59 | 18 | Water 9 | | a:4032 | 4.52 | 1.0 | 29.8 | 2.58 [A:OP2]a:453 | 9.70 | 19 | Water 10 | | a:4822 | 4.55 | 1.0 | 25.5 |  | 13.46 | 20 | Water 11 | | a:3555 | 4.55 | 1.0 | 43.6 | 2.47 [A:OP1]a:454 | 5.05 | 21 | A | OP2 | a:449 | 4.74 | 1.0 | 20.8 |  | 14.29 |

  
**1^st^ shell coordination angles and distances (*trans* in "blue"; *cys* in "green"; deviations in "orange"):**   
• 1-2(OP1/O6): 87º; 1-3(OP1/W): 174º; 1-4(OP1/W): 95º; 1-5(OP1/W): 89º; 2-3(O6/W): 88º; 2-4(O6/W): 91º; 2-5(O6/W): 173º; 3-4(W/W): 86º; 3-5(W/W): *97º*; 4-5(W/W): *85º*;   
• 1-2(OP1/O6): 2.8Å; 1-3(OP1/W): 4.2Å; 1-4(OP1/W): 3.1Å; 1-5(OP1/W): 3.0Å; 2-3(O6/W): 2.9Å; 2-4(O6/W): 3.1Å; 2-5(O6/W): *4.3Å*; 3-4(W/W): 3.0Å; 3-5(W/W): *3.3Å*; 4-5(W/W): 3.0Å;   
  
**2.17 ± 0.12: the average coodination distance of the 1^st^ six ligands is Mg^2+^ compatible**  
**8: Number of potential Water contacts**   
**4 : Number of second shell contacts involving water or ions**   
**50%: Percentage of satisfied second shell Water contacts**   
• Contact(s): /O6/O_w_/O_w_/O_w_/OP1/  
• Contact code: O_ph_O_b_3O_w_  
• Contact(s) in the exclusion zone: /O_w_/  
→ Missing octahedral coordination for MG ←  
Ooctahedral coordination (4.2º)   
Go to ToC

### MG:a:3084

|  | Res. type | Atom | Residue | Distance | Occupancy | B-factor | Closest atom(s) to a 1^st^ shell water | EM\_dens. |
| --- | --- | --- | --- | --- | --- | --- | --- | --- |
| **Ion** | | | |  | 1.0 | 33.6 |  | 17.54 |
 1 | Water 1 | | a:6291 | 1.94 | 1.0 | 31.5 | 2.89 [A:OP1]a:528 | 8.58 | 2 | C | OP1 | a:527 | 2.02 | 1.0 | 34.5 |  | 11.32 | 3 | A | OP1 | a:526 | 2.03 | 1.0 | 32.3 |  | 10.72 | 4 | Water 2 | | a:8272 | 2.12 | 1.0 | 30.4 |  | 9.02 | 5 | Water 3 | | a:9550 | 2.21 | 1.0 | 34.8 |  | 9.24 | 6 | Water 4 | | a:8836 | 2.24 | 1.0 | 36.5 |  | 7.90 | 7 | Water 5 | | a:8176 | 4.05 | 1.0 | 40.3 | 2.90 [C:OP2]a:527 | 5.10 | 8 | A | O5' | a:526 | 4.07 | 1.0 | 32.3 |  | 10.71 | 9 | C | O5' | a:527 | 4.12 | 1.0 | 28.9 |  | 14.20 | 10 | C | OP2 | a:527 | 4.15 | 1.0 | 30.9 |  | 12.31 | 11 | A | OP2 | a:526 | 4.19 | 1.0 | 30.6 |  | 11.66 | 12 | Water 6 | | a:9860 | 4.25 | 1.0 | 47.3 |  | 4.53 | 13 | A | OP1 | a:528 | 4.36 | 1.0 | 30.0 |  | 11.03 | 14 | Water 7 | | a:8321 | 4.44 | 1.0 | 40.2 |  | 5.01 | 15 | Water 8 | | a:8278 | 4.45 | 1.0 | 43.6 | 2.91 [A:OP1]a:528 | 3.88 | 16 | A | O3' | a:526 | 4.47 | 1.0 | 34.3 |  | 12.52 | 17 | U | O3' | a:525 | 4.47 | 1.0 | 33.0 |  | 11.03 | 18 | Water 9 | | a:3490 | 4.71 | 1.0 | 43.9 | 2.71 [A:OP2]a:526 | 5.40 | 19 | Water 10 | | a:6559 | 4.74 | 1.0 | 43.5 |  | 5.44 |

  
**1^st^ shell coordination angles and distances (*trans* in "blue"; *cys* in "green"; deviations in "orange"):**   
• 1-2(W/OP1): *96º*; 1-3(W/OP1): 88º; 1-4(W/W): 175º; 1-5(W/W): 91º; 1-6(W/W): 87º; 2-3(OP1/OP1): 89º; 2-4(OP1/W): 89º; 2-5(OP1/W): *96º*; 2-6(OP1/W): 176º; 3-4(OP1/W): 94º; 3-5(OP1/W): 175º; 3-6(OP1/W): 88º; 4-5(W/W): 86º; 4-6(W/W): 88º; 5-6(W/W): 88º;   
• 1-2(W/OP1): 2.9Å; 1-3(W/OP1): 2.8Å; 1-4(W/W): 4.1Å; 1-5(W/W): 3.0Å; 1-6(W/W): 2.9Å; 2-3(OP1/OP1): 2.8Å; 2-4(OP1/W): 2.9Å; 2-5(OP1/W): 3.1Å; 2-6(OP1/W): *4.3Å*; 3-4(OP1/W): 3.0Å; 3-5(OP1/W): *4.2Å*; 3-6(OP1/W): 3.0Å; 4-5(W/W): 3.0Å; 4-6(W/W): 3.0Å; 5-6(W/W): 3.1Å;   
  
**2.09 ± 0.11: the average coodination distance of the 1^st^ six ligands is Mg^2+^ compatible**  
**8: Number of potential Water contacts**   
**1 : Number of second shell contacts involving water or ions**   
**12%: Percentage of satisfied second shell Water contacts**   
• Contact(s): /O_w_/O_w_/O_w_/O_w_/OP1/OP1/  
• Contact code: *cis-*2O_ph_4O_w_  
Ooctahedral coordination (3.2º)   
Go to ToC

### MG:a:3085

|  | Res. type | Atom | Residue | Distance | Occupancy | B-factor | Closest atom(s) to a 1^st^ shell water | EM\_dens. |
| --- | --- | --- | --- | --- | --- | --- | --- | --- |
| **Ion** | | | |  | 1.0 | 41.0 |  | 15.64 |
 1 | Water 1 | | a:8813 | 1.91 | 1.0 | 38.8 | 2.97 [G:OP1]a:561 | 9.11 | 2 | Water 2 | | a:6348 | 2.06 | 1.0 | 33.1 | 2.76 [G:OP1]a:2042 | 12.30 | 3 | Water 3 | | a:9225 | 2.08 | 1.0 | 39.3 |  | 8.10 | 4 | Water 4 | | a:10170 | 2.09 | 1.0 | 34.8 |  | 9.69 | 5 | Water 5 | | a:7730 | 2.16 | 1.0 | 36.7 | 2.86 [C:OP1]a:562 | 9.15 | 6 | Water 6 | | a:6310 | 3.75 | 1.0 | 39.5 |  | 7.37 | 7 | Water 7 | | a:7526 | 3.85 | 1.0 | 35.3 |  | 7.73 | 8 | Water 8 | | a:8038 | 3.96 | 1.0 | 42.9 | 2.88 [A:OP1]a:1010 | 6.93 | 9 | A | OP1 | a:1010 | 4.16 | 1.0 | 35.8 |  | 8.39 | 10 | C | OP2 | a:562 | 4.26 | 1.0 | 31.2 |  | 9.37 | 11 | Water 9 | | a:9182 | 4.28 | 1.0 | 37.2 |  | 8.30 | 12 | Water 10 | | a:10223 | 4.34 | 1.0 | 46.6 |  | 4.73 | 13 | Water 11 | | a:9776 | 4.42 | 1.0 | 38.3 |  | 6.51 | 14 | C | OP1 | a:562 | 4.43 | 1.0 | 35.4 |  | 10.59 | 15 | G | OP1 | a:561 | 4.46 | 1.0 | 34.1 |  | 9.68 | 16 | Water 12 | | i:204 | 4.53 | 1.0 | 42.2 | 2.99 [G:OP1]a:561 | 6.07 | 17 | Water 13 | | a:4866 | 4.56 | 1.0 | 38.4 |  | 6.77 | 18 | G | OP1 | a:2042 | 4.66 | 1.0 | 25.9 |  | 11.48 |

  
**1^st^ shell coordination angles and distances (*trans* in "blue"; *cys* in "green"; deviations in "orange"):**   
• 1-2(W/W): *169º*; 1-3(W/W): 92º; 1-4(W/W): *98º*; 1-5(W/W): 90º; 2-3(W/W): *84º*; 2-4(W/W): 92º; 2-5(W/W): *80º*; 3-4(W/W): 92º; 3-5(W/W): *77º*; 4-5(W/W): *167º*;   
• 1-2(W/W): 4.0Å; 1-3(W/W): 2.9Å; 1-4(W/W): 3.0Å; 1-5(W/W): 2.9Å; 2-3(W/W): 2.8Å; 2-4(W/W): 3.0Å; 2-5(W/W): 2.7Å; 3-4(W/W): 3.0Å; 3-5(W/W): 2.6Å; 4-5(W/W): *4.2Å*;   
  
**2.34 ± 0.63: the average coodination distance of the 1^st^ six ligands is Na^+^** compatible  
**12: Number of potential Water contacts**   
**3 : Number of second shell contacts involving water or ions**   
**25%: Percentage of satisfied second shell Water contacts**   
• Contact(s): /O_w_/O_w_/O_w_/O_w_/O_w_/  
• Contact code: 5O_w_  
→ Missing octahedral coordination for MG ←  
→ Slightly distorded coord. octahedron (6.6º) ←  
Go to ToC

### MG:a:3086

|  | Res. type | Atom | Residue | Distance | Occupancy | B-factor | Closest atom(s) to a 1^st^ shell water | EM\_dens. |
| --- | --- | --- | --- | --- | --- | --- | --- | --- |
| **Ion** | | | |  | 1.0 | 23.7 |  | 30.08 |
 1 | Water 1 | | a:7373 | 1.97 | 1.0 | 24.4 | 2.83 [G:OP2]a:810 | 19.02 | 2 | U | OP1 | a:569 | 2.10 | 1.0 | 19.7 |  | 15.76 | 3 | Water 2 | | a:5429 | 2.21 | 1.0 | 23.1 | 2.70 [G:OP2]a:811 | 18.62 | 4 | Water 3 | | a:8158 | 2.23 | 1.0 | 18.5 | 2.89 [G:N7]a:811 | 17.64 | 5 | Water 4 | | a:7780 | 2.26 | 1.0 | 20.6 | 2.86 [U:O4]a:812 | 17.13 | 6 | Water 5 | | k:243 | 2.29 | 1.0 | 23.7 | 2.93 [HIS:ND1]k:35 | 18.06 | 7 | U | O3' | a:568 | 3.79 | 1.0 | 22.0 |  | 15.11 | 8 | Water 6 | | a:8850 | 4.07 | 1.0 | 27.8 |  | 15.29 | 9 | G | OP2 | a:810 | 4.12 | 1.0 | 22.1 |  | 15.63 | 10 | U | O5' | a:569 | 4.21 | 1.0 | 23.3 |  | 18.75 | 11 | U | O4 | a:812 | 4.23 | 1.0 | 24.6 |  | 19.18 | 12 | Water 7 | | k:224 | 4.35 | 1.0 | 27.5 |  | 16.30 | 13 | G | N7 | a:811 | 4.40 | 1.0 | 13.6 |  | 20.26 | 14 | Water 8 | | a:5966 | 4.43 | 1.0 | 24.0 |  | 17.24 | 15 | G | OP2 | a:811 | 4.53 | 1.0 | 22.4 |  | 16.02 | 16 | HIS | ND1 | k:35 | 4.56 | 1.0 | 27.4 |  | 9.82 | 17 | U | OP2 | a:569 | 4.57 | 1.0 | 24.7 |  | 15.21 | 18 | Water 9 | | a:6576 | 4.57 | 1.0 | 29.5 |  | 10.81 |

  
**1^st^ shell coordination angles and distances (*trans* in "blue"; *cys* in "green"; deviations in "orange"):**   
• 1-2(W/OP1): 95º; 1-3(W/W): 90º; 1-4(W/W): *98º*; 1-5(W/W): 177º; 1-6(W/W): *85º*; 2-3(OP1/W): *98º*; 2-4(OP1/W): *165º*; 2-5(OP1/W): 85º; 2-6(OP1/W): *84º*; 3-4(W/W): 90º; 3-5(W/W): 93º; 3-6(W/W): 174º; 4-5(W/W): *82º*; 4-6(W/W): 88º; 5-6(W/W): 92º;   
• 1-2(W/OP1): 3.0Å; 1-3(W/W): 3.0Å; 1-4(W/W): 3.2Å; 1-5(W/W): *4.2Å*; 1-6(W/W): 2.9Å; 2-3(OP1/W): *3.3Å*; 2-4(OP1/W): *4.3Å*; 2-5(OP1/W): 3.0Å; 2-6(OP1/W): 3.0Å; 3-4(W/W): 3.1Å; 3-5(W/W): *3.2Å*; 3-6(W/W): *4.5Å*; 4-5(W/W): 3.0Å; 4-6(W/W): 3.1Å; 5-6(W/W): *3.3Å*;   
  
**2.18 ± 0.11: the average coodination distance of the 1^st^ six ligands is Mg^2+^ compatible**  
**10: Number of potential Water contacts**   
**5 : Number of second shell contacts involving water or ions**   
**50%: Percentage of satisfied second shell Water contacts**   
• Contact(s): /O_w_/O_w_/O_w_/O_w_/O_w_/OP1/  
• Contact code: O_ph_5O_w_  
Ooctahedral coordination (5.0º)   
Go to ToC

### MG:a:3087

|  | Res. type | Atom | Residue | Distance | Occupancy | B-factor | Closest atom(s) to a 1^st^ shell water | EM\_dens. |
| --- | --- | --- | --- | --- | --- | --- | --- | --- |
| **Ion** | | | |  | 1.0 | 26.2 |  | 20.85 |
 1 | A | OP2 | a:1267 | 1.93 | 1.0 | 22.3 |  | 14.96 | 2 | U | OP1 | a:2619 | 2.01 | 1.0 | 29.3 |  | 13.34 | 3 | Water 1 | | a:8162 | 2.01 | 1.0 | 25.6 | 2.94 [U:OP2]a:2619 | 11.00 | 4 | Water 2 | | a:5993 | 2.15 | 1.0 | 24.5 | 2.74 [U:O2']a:1269 | 15.69 | 5 | Water 3 | | a:5214 | 2.21 | 1.0 | 29.7 | 2.69 [A:OP1]a:2618 | 12.36 | 6 | Water 4 | | a:6209 | 2.22 | 1.0 | 27.6 | 2.76 [A:OP1]a:1267 | 12.19 | 7 | A | OP1 | a:1267 | 3.52 | 1.0 | 24.8 |  | 14.32 | 8 | U | OP2 | a:2619 | 3.59 | 1.0 | 29.1 |  | 11.28 | 9 | Water 5 | | a:5929 | 3.82 | 1.0 | 37.9 |  | 8.79 | 10 | Water 6 | | a:5290 | 3.91 | 1.0 | 33.2 |  | 12.22 | 11 | A | O3' | a:1266 | 4.10 | 1.0 | 25.9 |  | 14.31 | 12 | U | O5' | a:2619 | 4.15 | 1.0 | 33.2 |  | 12.35 | 13 | Water 7 | | a:9948 | 4.21 | 1.0 | 38.5 |  | 7.37 | 14 | A | O5' | a:1267 | 4.25 | 1.0 | 21.8 |  | 14.86 | 15 | U | O2' | a:1269 | 4.26 | 1.0 | 27.5 |  | 15.44 | 16 | Water 8 | | a:9766 | 4.35 | 1.0 | 40.7 |  | 6.97 | 17 | A | O3' | a:2618 | 4.36 | 1.0 | 27.6 |  | 12.95 | 18 | Water 9 | | a:9672 | 4.50 | 1.0 | 40.6 |  | 4.97 | 19 | A | OP1 | a:2618 | 4.51 | 1.0 | 27.6 |  | 11.65 | 20 | Water 10 | | a:5016 | 4.79 | 1.0 | 35.8 |  | 8.96 |

  
**1^st^ shell coordination angles and distances (*trans* in "blue"; *cys* in "green"; deviations in "orange"):**   
• 1-2(OP2/OP1): *96º*; 1-3(OP2/W): 94º; 1-4(OP2/W): *97º*; 1-5(OP2/W): 175º; 1-6(OP2/W): 94º; 2-3(OP1/W): *98º*; 2-4(OP1/W): *167º*; 2-5(OP1/W): 89º; 2-6(OP1/W): 94º; 3-4(W/W): *83º*; 3-5(W/W): 86º; 3-6(W/W): *164º*; 4-5(W/W): *79º*; 4-6(W/W): *83º*; 5-6(W/W): *85º*;   
• 1-2(OP2/OP1): 2.9Å; 1-3(OP2/W): 2.9Å; 1-4(OP2/W): 3.1Å; 1-5(OP2/W): 4.1Å; 1-6(OP2/W): 3.1Å; 2-3(OP1/W): 3.0Å; 2-4(OP1/W): 4.1Å; 2-5(OP1/W): 2.9Å; 2-6(OP1/W): 3.1Å; 3-4(W/W): 2.8Å; 3-5(W/W): 2.9Å; 3-6(W/W): 4.2Å; 4-5(W/W): 2.8Å; 4-6(W/W): 2.9Å; 5-6(W/W): 3.0Å;   
  
**2.09 ± 0.11: the average coodination distance of the 1^st^ six ligands is Mg^2+^ compatible**  
**8: Number of potential Water contacts**   
**4 : Number of second shell contacts involving water or ions**   
**50%: Percentage of satisfied second shell Water contacts**   
• Contact(s): /O_w_/O_w_/O_w_/O_w_/OP1/OP2/  
• Contact code: *cis-*2O_ph_4O_w_  
→ Slightly distorded coord. octahedron (6.9º) ←  
Go to ToC

### MG:a:3088

|  | Res. type | Atom | Residue | Distance | Occupancy | B-factor | Closest atom(s) to a 1^st^ shell water | EM\_dens. |
| --- | --- | --- | --- | --- | --- | --- | --- | --- |
| **Ion** | | | |  | 1.0 | 24.4 |  | 27.54 |
 1 | Water 1 | | a:6019 | 2.11 | 1.0 | 20.2 | 2.74 [A:OP2]a:195 | 17.41 | 2 | A | OP1 | a:197 | 2.12 | 1.0 | 22.5 |  | 17.42 | 3 | Water 2 | | a:7688 | 2.12 | 1.0 | 21.0 | 2.86 [G:N7]a:194 | 18.41 | 4 | C | OP2 | a:198 | 2.14 | 1.0 | 21.1 |  | 16.35 | 5 | Water 3 | | a:7619 | 2.16 | 1.0 | 22.6 |  | 15.37 | 6 | A | N7 | a:195 | 2.25 | 1.0 | 18.1 |  | 17.74 | 7 | G | OP2 | a:194 | 4.08 | 1.0 | 30.5 |  | 11.62 | 8 | G | N7 | a:194 | 4.14 | 1.0 | 18.1 |  | 16.02 | 9 | C | OP1 | a:198 | 4.16 | 1.0 | 30.1 |  | 11.83 | 10 | Water 4 | | a:5025 | 4.20 | 1.0 | 29.5 | 2.92 [C:OP1]a:198 | 14.25 | 11 | A | OP2 | a:197 | 4.23 | 1.0 | 27.3 |  | 14.63 | 12 | A | O5' | a:197 | 4.23 | 1.0 | 20.0 |  | 16.61 | 13 | **K** | K | a:3249 | 4.29 | 1.0 | 30.0 |  | 25.91 | 14 | A | O3' | a:197 | 4.34 | 1.0 | 26.5 |  | 14.38 | 15 | A | OP2 | a:195 | 4.36 | 1.0 | 27.8 |  | 12.05 | 16 | C | O5' | a:198 | 4.50 | 1.0 | 22.5 |  | 15.92 | 17 | A | O3' | a:196 | 4.51 | 1.0 | 25.2 |  | 13.34 | 18 | Water 5 | | a:6738 | 4.73 | 1.0 | 39.5 |  | 6.81 |

  
**1^st^ shell coordination angles and distances (*trans* in "blue"; *cys* in "green"; deviations in "orange"):**   
• 1-2(W/OP1): 86º; 1-3(W/W): 94º; 1-4(W/OP2): *168º*; 1-5(W/W): *85º*; 1-6(W/N7): 92º; 2-3(OP1/W): 173º; 2-4(OP1/OP2): 95º; 2-5(OP1/W): 93º; 2-6(OP1/N7): *84º*; 3-4(W/OP2): 88º; 3-5(W/W): 94º; 3-6(W/N7): 89º; 4-5(OP2/W): *83º*; 4-6(OP2/N7): *100º*; 5-6(W/N7): 175º;   
• 1-2(W/OP1): 2.9Å; 1-3(W/W): 3.1Å; 1-4(W/OP2): *4.2Å*; 1-5(W/W): 2.9Å; 1-6(W/N7): 3.1Å; 2-3(OP1/W): *4.2Å*; 2-4(OP1/OP2): 3.1Å; 2-5(OP1/W): 3.1Å; 2-6(OP1/N7): 2.9Å; 3-4(W/OP2): 2.9Å; 3-5(W/W): 3.1Å; 3-6(W/N7): 3.1Å; 4-5(OP2/W): 2.9Å; 4-6(OP2/N7): *3.4Å*; 5-6(W/N7): *4.4Å*;   
  
**2.15 ± 0.05: the average coodination distance of the 1^st^ six ligands is Mg^2+^ compatible**  
**6: Number of potential Water contacts**   
**2 : Number of second shell contacts involving water or ions**   
**33%: Percentage of satisfied second shell Water contacts**   
• Contact(s): /N7/O_w_/O_w_/O_w_/OP1/OP2/  
• Contact code: *cis-*2O_ph_N_b_3O_w_  
→ Slightly distorded coord. octahedron (5.1º) ←  
Go to ToC

### MG:a:3089

|  | Res. type | Atom | Residue | Distance | Occupancy | B-factor | Closest atom(s) to a 1^st^ shell water | EM\_dens. |
| --- | --- | --- | --- | --- | --- | --- | --- | --- |
| **Ion** | | | |  | 1.0 | 28.9 |  | 18.78 |
 1 | C | OP2 | a:1317 | 1.98 | 1.0 | 41.0 |  | 7.26 | 2 | C | OP1 | a:1316 | 2.01 | 1.0 | 26.7 |  | 11.61 | 3 | G | OP1 | a:1334 | 2.04 | 1.0 | 33.5 |  | 8.25 | 4 | Water 1 | | a:5104 | 2.04 | 1.0 | 30.7 |  | 8.52 | 5 | Water 2 | | a:6936 | 2.20 | 1.0 | 30.8 | 2.93 [C:OP2]a:1316 | 10.73 | 6 | Water 3 | | a:9261 | 2.25 | 1.0 | 35.3 |  | 11.08 | 7 | Water 4 | | a:7566 | 3.75 | 1.0 | 38.2 |  | 7.62 | 8 | C | OP2 | a:1316 | 3.76 | 1.0 | 29.5 |  | 11.43 | 9 | C | OP1 | a:1317 | 3.89 | 1.0 | 30.7 |  | 9.14 | 10 | G | O5' | a:1334 | 4.01 | 1.0 | 27.6 |  | 9.89 | 11 | C | O5' | a:1316 | 4.06 | 1.0 | 32.6 |  | 11.30 | 12 | Water 5 | | a:9083 | 4.09 | 1.0 | 42.9 |  | 5.84 | 13 | Water 6 | | a:5539 | 4.16 | 1.0 | 46.2 | 2.71 [G:OP2]a:1334 | 5.09 | 14 | G | OP2 | a:1334 | 4.19 | 1.0 | 26.0 |  | 11.33 | 15 | C | O3' | a:1316 | 4.30 | 1.0 | 35.7 |  | 10.03 | 16 | Water 7 | | a:9132 | 4.32 | 1.0 | 37.7 |  | 6.36 | 17 | C | O5' | a:1317 | 4.33 | 1.0 | 34.9 |  | 8.60 | 18 | U | O3' | a:1315 | 4.45 | 1.0 | 29.4 |  | 10.65 | 19 | G | O3' | a:1333 | 4.47 | 1.0 | 30.9 |  | 10.26 | 20 | Water 8 | | a:4083 | 4.47 | 1.0 | 43.1 |  | 3.59 | 21 | Water 9 | | a:7571 | 4.50 | 1.0 | 42.5 |  | 4.44 | 22 | Water 10 | | a:7086 | 4.69 | 1.0 | 24.6 | 2.85 [C:OP2]a:1316 // 2.86 [G:O5']a:1334 | 13.01 |

  
**1^st^ shell coordination angles and distances (*trans* in "blue"; *cys* in "green"; deviations in "orange"):**   
• 1-2(OP2/OP1): 92º; 1-3(OP2/OP1): *168º*; 1-4(OP2/W): 88º; 1-5(OP2/W): *84º*; 1-6(OP2/W): *95º*; 2-3(OP1/OP1): *98º*; 2-4(OP1/W): 179º; 2-5(OP1/W): 90º; 2-6(OP1/W): 91º; 3-4(OP1/W): *82º*; 3-5(OP1/W): 89º; 3-6(OP1/W): 91º; 4-5(W/W): 89º; 4-6(W/W): 90º; 5-6(W/W): 179º;   
• 1-2(OP2/OP1): 2.9Å; 1-3(OP2/OP1): 4.0Å; 1-4(OP2/W): 2.8Å; 1-5(OP2/W): 2.8Å; 1-6(OP2/W): 3.1Å; 2-3(OP1/OP1): 3.1Å; 2-4(OP1/W): 4.1Å; 2-5(OP1/W): 3.0Å; 2-6(OP1/W): 3.0Å; 3-4(OP1/W): 2.7Å; 3-5(OP1/W): 3.0Å; 3-6(OP1/W): 3.1Å; 4-5(W/W): 3.0Å; 4-6(W/W): 3.0Å; 5-6(W/W): *4.4Å*;   
  
**2.09 ± 0.10: the average coodination distance of the 1^st^ six ligands is Mg^2+^ compatible**  
**6: Number of potential Water contacts**   
**1 : Number of second shell contacts involving water or ions**   
**17%: Percentage of satisfied second shell Water contacts**   
• Contact(s): /O_w_/O_w_/O_w_/OP1/OP1/OP2/  
• Contact code: *mer-*3O_ph_3O_w_  
Ooctahedral coordination (3.2º)   
Go to ToC

### MG:a:3090

|  | Res. type | Atom | Residue | Distance | Occupancy | B-factor | Closest atom(s) to a 1^st^ shell water | EM\_dens. |
| --- | --- | --- | --- | --- | --- | --- | --- | --- |
| **Ion** | | | |  | 1.0 | 31.3 |  | 19.14 |
 1 | A | OP2 | a:1678 | 2.05 | 1.0 | 31.0 |  | 11.82 | 2 | Water 1 | | a:7815 | 2.09 | 1.0 | 28.1 | 2.87 [A:N7]a:1679 | 12.77 | 3 | Water 2 | | a:5221 | 2.14 | 1.0 | 24.4 | 2.69 [C:OP2]a:1677 // 2.78 [A:N7]a:1678 | 12.85 | 4 | Water 3 | | a:8055 | 2.16 | 1.0 | 30.6 |  | 11.06 | 5 | Water 4 | | a:9441 | 2.25 | 1.0 | 30.8 |  | 10.99 | 6 | Water 5 | | a:9863 | 2.27 | 1.0 | 31.3 |  | 12.00 | 7 | A | O5' | a:1678 | 3.75 | 1.0 | 32.6 |  | 11.64 | 8 | A | N7 | a:1678 | 4.05 | 1.0 | 25.3 |  | 15.36 | 9 | Water 6 | | a:6929 | 4.10 | 1.0 | 38.1 | 2.80 [A:OP1]a:1678 | 7.42 | 10 | Water 7 | | a:10288 | 4.12 | 1.0 | 43.8 |  | 3.52 | 11 | C | OP2 | a:1677 | 4.13 | 1.0 | 32.5 |  | 10.05 | 12 | A | OP1 | a:1678 | 4.14 | 1.0 | 28.7 |  | 12.25 | 13 | Water 8 | | a:4363 | 4.20 | 1.0 | 44.0 |  | 4.01 | 14 | A | N7 | a:1679 | 4.32 | 1.0 | 22.8 |  | 15.95 | 15 | Water 9 | | a:7349 | 4.38 | 1.0 | 31.3 |  | 12.02 | 16 | C | O3' | a:1677 | 4.51 | 1.0 | 31.1 |  | 12.53 | 17 | C | O5' | a:1677 | 4.51 | 1.0 | 36.5 |  | 10.51 |

  
**1^st^ shell coordination angles and distances (*trans* in "blue"; *cys* in "green"; deviations in "orange"):**   
• 1-2(OP2/W): *96º*; 1-3(OP2/W): 94º; 1-4(OP2/W): 86º; 1-5(OP2/W): 92º; 1-6(OP2/W): 175º; 2-3(W/W): *101º*; 2-4(W/W): 172º; 2-5(W/W): 86º; 2-6(W/W): 88º; 3-4(W/W): 86º; 3-5(W/W): 170º; 3-6(W/W): 87º; 4-5(W/W): 86º; 4-6(W/W): 89º; 5-6(W/W): 86º;   
• 1-2(OP2/W): 3.1Å; 1-3(OP2/W): 3.1Å; 1-4(OP2/W): 2.9Å; 1-5(OP2/W): 3.1Å; 1-6(OP2/W): *4.3Å*; 2-3(W/W): *3.3Å*; 2-4(W/W): *4.2Å*; 2-5(W/W): 3.0Å; 2-6(W/W): 3.0Å; 3-4(W/W): 2.9Å; 3-5(W/W): *4.4Å*; 3-6(W/W): 3.0Å; 4-5(W/W): 3.0Å; 4-6(W/W): 3.1Å; 5-6(W/W): 3.1Å;   
  
**2.16 ± 0.08: the average coodination distance of the 1^st^ six ligands is Mg^2+^ compatible**  
**10: Number of potential Water contacts**   
**3 : Number of second shell contacts involving water or ions**   
**30%: Percentage of satisfied second shell Water contacts**   
• Contact(s): /O_w_/O_w_/O_w_/O_w_/O_w_/OP2/  
• Contact code: O_ph_5O_w_  
Ooctahedral coordination (4.7º)   
Go to ToC

### MG:a:3091

|  | Res. type | Atom | Residue | Distance | Occupancy | B-factor | Closest atom(s) to a 1^st^ shell water | EM\_dens. |
| --- | --- | --- | --- | --- | --- | --- | --- | --- |
| **Ion** | | | |  | 1.0 | 28.5 |  | 20.82 |
 1 | U | OP2 | a:511 | 1.81 | 1.0 | 30.4 |  | 10.86 | 2 | Water 1 | | a:8835 | 2.13 | 1.0 | 24.6 |  | 11.84 | 3 | C | OP1 | a:510 | 2.22 | 1.0 | 29.8 |  | 10.76 | 4 | Water 2 | | a:5760 | 2.26 | 1.0 | 27.5 | 2.73 [C:O3']a:509 | 13.67 | 5 | Water 3 | | a:6363 | 2.32 | 1.0 | 31.5 | 2.77 [G:O6]a:512 | 11.76 | 6 | G | N7 | a:512 | 2.35 | 1.0 | 20.1 |  | 17.04 | 7 | C | O3' | a:509 | 3.48 | 1.0 | 30.6 |  | 10.54 | 8 | G | O6 | a:512 | 3.86 | 1.0 | 36.9 |  | 4.70 | 9 | Water 4 | | a:3788 | 3.88 | 1.0 | 34.1 | 2.53 [U:OP1]a:511 | 7.60 | 10 | C | O3' | a:510 | 4.14 | 1.0 | 23.2 |  | 13.26 | 11 | U | OP1 | a:511 | 4.15 | 1.0 | 28.0 |  | 12.34 | 12 | U | O5' | a:511 | 4.20 | 1.0 | 32.8 |  | 12.15 | 13 | Water 5 | | a:8710 | 4.25 | 1.0 | 42.2 |  | 5.60 | 14 | C | O5' | a:510 | 4.35 | 1.0 | 33.6 |  | 11.18 | 15 | C | OP2 | a:510 | 4.52 | 1.0 | 34.2 |  | 8.98 | 16 | Water 6 | | a:5492 | 4.70 | 1.0 | 32.9 |  | 8.74 | 17 | Water 7 | | a:9810 | 4.73 | 1.0 | 45.2 |  | 4.31 |

  
**1^st^ shell coordination angles and distances (*trans* in "blue"; *cys* in "green"; deviations in "orange"):**   
• 1-2(OP2/W): *98º*; 1-3(OP2/OP1): 91º; 1-4(OP2/W): 95º; 2-3(W/OP1): 171º; 2-4(W/W): 89º; 3-4(OP1/W): 92º;   
• 1-2(OP2/W): 3.0Å; 1-3(OP2/OP1): 2.9Å; 1-4(OP2/W): 3.0Å; 2-3(W/OP1): *4.3Å*; 2-4(W/W): 3.1Å; 3-4(OP1/W): *3.2Å*;   
  
**2.18 ± 0.18: the average coodination distance of the 1^st^ six ligands is Mg^2+^ compatible**  
**6: Number of potential Water contacts**   
**2 : Number of second shell contacts involving water or ions**   
**33%: Percentage of satisfied second shell Water contacts**   
• Contact(s): /O_w_/O_w_/OP1/  
• Contact code: O_ph_2O_w_  
• Contact(s) in the exclusion zone: /N7/O_w_/OP2/  
→ Missing octahedral coordination for MG ←  
Ooctahedral coordination (4.5º)   
Go to ToC

### MG:a:3092

|  | Res. type | Atom | Residue | Distance | Occupancy | B-factor | Closest atom(s) to a 1^st^ shell water | EM\_dens. |
| --- | --- | --- | --- | --- | --- | --- | --- | --- |
| **Ion** | | | |  | 1.0 | 25.2 |  | 27.20 |
 1 | Water 1 | | a:6018 | 2.11 | 1.0 | 22.5 | 2.74 [A:OP1]a:975 | 17.37 | 2 | Water 2 | | a:4150 | 2.16 | 1.0 | 20.4 | 2.59 [A:N1]a:823 // 2.81 [A:O2']a:974 | 18.46 | 3 | Water 3 | | k:272 | 2.16 | 1.0 | 19.7 |  | 17.22 | 4 | Water 4 | | a:8814 | 2.19 | 1.0 | 23.0 | 2.97 [A:N1]a:822 | 17.05 | 5 | Water 5 | | a:4947 | 2.30 | 1.0 | 23.1 | 2.67 [U:OP2]a:571 // 2.89 [G:OP2]a:572 | 17.59 | 6 | Water 6 | | a:4851 | 2.34 | 1.0 | 19.9 | 2.66 [G:OP1]a:572 // 2.95 [A:OP2]a:975 | 18.42 | 7 | Water 7 | | a:5144 | 4.25 | 1.0 | 26.9 | 2.68 [U:OP2]a:571 | 14.02 | 8 | A | OP1 | a:975 | 4.25 | 1.0 | 24.2 |  | 14.77 | 9 | Water 8 | | k:260 | 4.26 | 1.0 | 29.4 |  | 10.86 | 10 | Water 9 | | a:9228 | 4.28 | 1.0 | 30.5 |  | 11.71 | 11 | Water 10 | | a:10137 | 4.33 | 1.0 | 31.0 |  | 13.09 | 12 | A | N1 | a:823 | 4.35 | 1.0 | 19.8 |  | 17.65 | 13 | A | O2' | a:974 | 4.40 | 1.0 | 28.9 |  | 16.63 | 14 | A | N1 | a:822 | 4.45 | 1.0 | 17.9 |  | 18.69 | 15 | G | OP1 | a:572 | 4.46 | 1.0 | 19.3 |  | 19.52 | 16 | G | OP2 | a:572 | 4.52 | 1.0 | 24.5 |  | 15.68 | 17 | Water 11 | | a:5441 | 4.54 | 1.0 | 32.1 |  | 14.15 | 18 | U | OP2 | a:571 | 4.62 | 1.0 | 18.3 |  | 18.08 | 19 | A | OP2 | a:975 | 4.62 | 1.0 | 26.1 |  | 14.75 |

  
**1^st^ shell coordination angles and distances (*trans* in "blue"; *cys* in "green"; deviations in "orange"):**   
• 1-2(W/W): 88º; 1-3(W/W): 95º; 1-4(W/W): *99º*; 2-3(W/W): 175º; 2-4(W/W): *84º*; 3-4(W/W): 92º;   
• 1-2(W/W): 3.0Å; 1-3(W/W): 3.1Å; 1-4(W/W): *3.3Å*; 2-3(W/W): *4.3Å*; 2-4(W/W): 2.9Å; 3-4(W/W): 3.1Å;   
  
**2.21 ± 0.08: the average coodination distance of the 1^st^ six ligands is Mg^2+^ compatible**  
**12: Number of potential Water contacts**   
**8 : Number of second shell contacts involving water or ions**   
**67%: Percentage of satisfied second shell Water contacts**   
• Contact(s): /O_w_/O_w_/O_w_/O_w_/  
• Contact code: 4O_w_  
• Contact(s) in the exclusion zone: /O_w_/O_w_/  
→ Missing octahedral coordination for MG ←  
Ooctahedral coordination (4.6º)   
Go to ToC

### MG:a:3093

|  | Res. type | Atom | Residue | Distance | Occupancy | B-factor | Closest atom(s) to a 1^st^ shell water | EM\_dens. |
| --- | --- | --- | --- | --- | --- | --- | --- | --- |
| **Ion** | | | |  | 1.0 | 31.2 |  | 19.38 |
 1 | A | OP2 | a:1011 | 1.98 | 1.0 | 28.4 |  | 12.79 | 2 | Water 1 | | a:7038 | 2.02 | 1.0 | 32.0 |  | 10.82 | 3 | Water 2 | | a:3662 | 2.04 | 1.0 | 31.8 | 2.50 [C:OP2]a:1009 | 9.30 | 4 | A | OP2 | a:1012 | 2.06 | 1.0 | 26.8 |  | 12.60 | 5 | Water 3 | | a:5514 | 2.14 | 1.0 | 27.7 | 2.71 [A:N7]a:1012 | 11.65 | 6 | Water 4 | | a:7840 | 2.30 | 1.0 | 27.4 | 2.87 [C:OP2]a:1008 | 11.46 | 7 | A | O3' | a:1010 | 3.53 | 1.0 | 27.3 |  | 11.87 | 8 | A | O5' | a:1011 | 3.86 | 1.0 | 32.7 |  | 11.64 | 9 | C | OP2 | a:1008 | 4.14 | 1.0 | 34.2 |  | 8.82 | 10 | A | N7 | a:1012 | 4.16 | 1.0 | 22.7 |  | 12.53 | 11 | C | OP2 | a:1009 | 4.18 | 1.0 | 29.2 |  | 11.89 | 12 | Water 5 | | a:5045 | 4.24 | 1.0 | 34.8 | 2.68 [A:OP1]a:1012 | 7.96 | 13 | A | OP1 | a:1012 | 4.31 | 1.0 | 27.0 |  | 12.41 | 14 | A | O3' | a:1011 | 4.32 | 1.0 | 27.1 |  | 11.55 | 15 | A | O5' | a:1012 | 4.39 | 1.0 | 31.4 |  | 10.94 | 16 | C | OP1 | a:1008 | 4.41 | 1.0 | 33.9 |  | 8.30 | 17 | A | OP1 | a:1011 | 4.44 | 1.0 | 29.1 |  | 11.33 | 18 | Water 6 | | a:4428 | 4.45 | 1.0 | 33.4 |  | 8.31 | 19 | A | O2' | a:1010 | 4.60 | 1.0 | 32.8 |  | 11.75 | 20 | Water 7 | | i:202 | 4.65 | 1.0 | 40.7 | 2.37 [LYS:NZ]i:39 | 3.54 | 21 | Water 8 | | a:9015 | 4.67 | 1.0 | 39.5 | 3.00 [A:O2']a:1010 | 6.93 |

  
**1^st^ shell coordination angles and distances (*trans* in "blue"; *cys* in "green"; deviations in "orange"):**   
• 1-2(OP2/W): 89º; 1-3(OP2/W): *96º*; 1-4(OP2/OP2): 93º; 1-5(OP2/W): *101º*; 1-6(OP2/W): 176º; 2-3(W/W): 87º; 2-4(W/OP2): 90º; 2-5(W/W): *169º*; 2-6(W/W): 88º; 3-4(W/OP2): *170º*; 3-5(W/W): 88º; 3-6(W/W): 86º; 4-5(OP2/W): 94º; 4-6(OP2/W): *85º*; 5-6(W/W): *83º*;   
• 1-2(OP2/W): 2.8Å; 1-3(OP2/W): 3.0Å; 1-4(OP2/OP2): 2.9Å; 1-5(OP2/W): 3.2Å; 1-6(OP2/W): *4.3Å*; 2-3(W/W): 2.8Å; 2-4(W/OP2): 2.9Å; 2-5(W/W): 4.1Å; 2-6(W/W): 3.0Å; 3-4(W/OP2): 4.1Å; 3-5(W/W): 2.9Å; 3-6(W/W): 3.0Å; 4-5(OP2/W): 3.1Å; 4-6(OP2/W): 2.9Å; 5-6(W/W): 2.9Å;   
  
**2.09 ± 0.10: the average coodination distance of the 1^st^ six ligands is Mg^2+^ compatible**  
**8: Number of potential Water contacts**   
**3 : Number of second shell contacts involving water or ions**   
**38%: Percentage of satisfied second shell Water contacts**   
• Contact(s): /O_w_/O_w_/O_w_/O_w_/OP2/OP2/  
• Contact code: *cis-*2O_ph_4O_w_  
Ooctahedral coordination (4.9º)   
Go to ToC

### MG:a:3094

|  | Res. type | Atom | Residue | Distance | Occupancy | B-factor | Closest atom(s) to a 1^st^ shell water | EM\_dens. |
| --- | --- | --- | --- | --- | --- | --- | --- | --- |
| **Ion** | | | |  | 1.0 | 25.7 |  | 26.89 |
 1 | A | OP2 | a:945 | 1.90 | 1.0 | 21.5 |  | 15.29 | 2 | Water 1 | | a:8972 | 2.06 | 1.0 | 25.5 |  | 17.48 | 3 | Water 2 | | a:8638 | 2.12 | 1.0 | 20.2 |  | 15.74 | 4 | Water 3 | | k:225 | 2.24 | 1.0 | 22.6 | 2.82 [HIS:O]k:35 | 18.09 | 5 | Water 4 | | k:214 | 2.37 | 1.0 | 21.8 | 2.70 [ARG:O]k:33 // 2.90 [LYS:O]k:36 | 16.63 | 6 | Water 5 | | a:4126 | 2.45 | 1.0 | 23.2 | 2.59 [U:OP1]a:834 // 3.00 [LYS:O]k:36 | 17.78 | 7 | A | OP1 | a:945 | 3.83 | 1.0 | 24.7 |  | 13.97 | 8 | LYS | O | k:36 | 3.98 | 1.0 | 21.9 |  | 16.74 | 9 | A | O5' | a:945 | 4.06 | 1.0 | 19.6 |  | 18.03 | 10 | Water 6 | | a:8092 | 4.13 | 1.0 | 26.9 |  | 15.61 | 11 | Water 7 | | a:4864 | 4.17 | 1.0 | 31.2 | 2.70 [ARG:O]k:33 | 14.72 | 12 | Water 8 | | a:4932 | 4.17 | 1.0 | 25.6 |  | 14.68 | 13 | Water 9 | | a:4152 | 4.25 | 1.0 | 29.1 |  | 12.00 | 14 | HIS | O | k:35 | 4.26 | 1.0 | 30.9 |  | 14.67 | 15 | G | O3' | a:944 | 4.29 | 1.0 | 24.9 |  | 17.18 | 16 | ARG | O | k:33 | 4.41 | 1.0 | 26.0 |  | 16.28 | 17 | U | OP1 | a:834 | 4.61 | 1.0 | 24.0 |  | 13.97 |

  
**1^st^ shell coordination angles and distances (*trans* in "blue"; *cys* in "green"; deviations in "orange"):**   
• 1-2(OP2/W): *98º*; 1-3(OP2/W): 94º; 1-4(OP2/W): *100º*; 2-3(W/W): 92º; 2-4(W/W): 86º; 3-4(W/W): *166º*;   
• 1-2(OP2/W): 3.0Å; 1-3(OP2/W): 2.9Å; 1-4(OP2/W): 3.2Å; 2-3(W/W): 3.0Å; 2-4(W/W): 2.9Å; 3-4(W/W): *4.3Å*;   
  
**2.19 ± 0.19: the average coodination distance of the 1^st^ six ligands is Mg^2+^ compatible**  
**10: Number of potential Water contacts**   
**5 : Number of second shell contacts involving water or ions**   
**50%: Percentage of satisfied second shell Water contacts**   
• Contact(s): /O_w_/O_w_/O_w_/  
• Contact code: 3O_w_  
• Contact(s) in the exclusion zone: /O_w_/O_w_/OP2/  
→ Missing octahedral coordination for MG ←  
→ Slightly distorded coord. octahedron (7.2º) ←  
Go to ToC

### MG:a:3095

|  | Res. type | Atom | Residue | Distance | Occupancy | B-factor | Closest atom(s) to a 1^st^ shell water | EM\_dens. |
| --- | --- | --- | --- | --- | --- | --- | --- | --- |
| **Ion** | | | |  | 1.0 | 41.6 |  | 15.46 |
 1 | Water 1 | | a:3491 | 1.99 | 1.0 | 36.6 | 2.43 [G:O6]a:1653 | 7.26 | 2 | Water 2 | | a:8231 | 2.04 | 1.0 | 28.7 | 2.90 [G:N7]a:1653 | 9.83 | 3 | Water 3 | | a:10196 | 2.06 | 1.0 | 33.1 |  | 7.76 | 4 | Water 4 | | a:4264 | 2.22 | 1.0 | 33.6 | 2.61 [G:OP2]a:2008 // 2.93 [ASN:OD1]m:11 | 9.60 | 5 | Water 5 | | a:5560 | 2.25 | 1.0 | 35.7 | 2.71 [G:OP1]a:2008 | 7.35 | 6 | Water 6 | | a:4462 | 2.26 | 1.0 | 36.0 | 2.63 [A:N7]a:1652 | 10.97 | 7 | Water 7 | | a:5073 | 3.99 | 1.0 | 32.3 |  | 10.09 | 8 | Water 8 | | a:6199 | 4.07 | 1.0 | 41.1 |  | 6.75 | 9 | Water 9 | | a:8521 | 4.17 | 1.0 | 41.0 |  | 4.31 | 10 | G | OP2 | a:2008 | 4.26 | 1.0 | 30.7 |  | 11.72 | 11 | Water 10 | | a:5081 | 4.26 | 1.0 | 40.4 |  | 6.01 | 12 | G | O6 | a:1653 | 4.27 | 1.0 | 24.7 |  | 16.93 | 13 | A | N7 | a:1652 | 4.28 | 1.0 | 21.9 |  | 15.12 | 14 | ASN | OD1 | m:11 | 4.29 | 1.0 | 31.9 |  | 12.56 | 15 | G | OP1 | a:2008 | 4.52 | 1.0 | 28.8 |  | 10.84 | 16 | G | N7 | a:1653 | 4.54 | 1.0 | 20.8 |  | 13.62 | 17 | Water 11 | | a:4744 | 4.67 | 1.0 | 35.1 |  | 8.01 |

  
**1^st^ shell coordination angles and distances (*trans* in "blue"; *cys* in "green"; deviations in "orange"):**   
• 1-2(W/W): *82º*; 1-3(W/W): *156º*; 1-4(W/W): *83º*; 1-5(W/W): *106º*; 1-6(W/W): *103º*; 2-3(W/W): *84º*; 2-4(W/W): 89º; 2-5(W/W): 171º; 2-6(W/W): 87º; 3-4(W/W): *77º*; 3-5(W/W): 87º; 3-6(W/W): *95º*; 4-5(W/W): 87º; 4-6(W/W): 172º; 5-6(W/W): *96º*;   
• 1-2(W/W): 2.6Å; 1-3(W/W): 4.0Å; 1-4(W/W): 2.8Å; 1-5(W/W): *3.4Å*; 1-6(W/W): *3.3Å*; 2-3(W/W): 2.7Å; 2-4(W/W): 3.0Å; 2-5(W/W): *4.3Å*; 2-6(W/W): 3.0Å; 3-4(W/W): 2.7Å; 3-5(W/W): 3.0Å; 3-6(W/W): 3.2Å; 4-5(W/W): 3.1Å; 4-6(W/W): *4.5Å*; 5-6(W/W): *3.4Å*;   
  
**2.14 ± 0.11: the average coodination distance of the 1^st^ six ligands is Mg^2+^ compatible**  
**12: Number of potential Water contacts**   
**6 : Number of second shell contacts involving water or ions**   
**50%: Percentage of satisfied second shell Water contacts**   
• Contact(s): /O_w_/O_w_/O_w_/O_w_/O_w_/O_w_/  
• Contact code: 6O_w_  
→ Slightly distorded coord. octahedron (8.3º) ←  
Go to ToC

### K:a:3096

|  | Res. type | Atom | Residue | Distance | Occupancy | B-factor | Closest atom(s) to a 1^st^ shell water | EM\_dens. |
| --- | --- | --- | --- | --- | --- | --- | --- | --- |
| **Ion** | | | |  | 1.0 | 30.0 |  | 10.55 |
 1 | Water 1 | | c:559 | 2.62 | 1.0 | 53.3 |  | 1.34 | 2 | U | O2 | a:1695 | 2.82 | 1.0 | 28.3 |  | 13.15 | 3 | Water 2 | | a:6336 | 2.84 | 1.0 | 49.1 | 2.76 [U:O4']a:1695 | 2.33 | 4 | C | O2' | a:1832 | 2.84 | 1.0 | 29.5 |  | 12.66 | 5 | Water 3 | | a:8884 | 3.42 | 1.0 | 39.9 | 2.97 [C:O3']a:1832 | 3.71 | 6 | C | O4' | a:1832 | 3.67 | 1.0 | 33.1 |  | 10.17 | 7 | Water 4 | | c:546 | 3.99 | 1.0 | 41.6 |  | 4.26 | 8 | **K** | K | a:3097 | 4.15 | 1.0 | 30.0 |  | 7.61 | 9 | A | N1 | a:1831 | 4.28 | 1.0 | 19.5 |  | 17.55 | 10 | HIS | ND1 | c:15 | 4.32 | 1.0 | 22.4 |  | 13.11 | 11 | HIS | NE2 | c:15 | 4.33 | 1.0 | 27.1 |  | 12.15 | 12 | A | N3 | a:1831 | 4.43 | 1.0 | 18.4 |  | 16.33 | 13 | U | N1 | a:1695 | 4.47 | 1.0 | 27.8 |  | 12.23 | 14 | Water 5 | | a:9964 | 4.50 | 1.0 | 47.8 |  | 1.64 | 15 | U | O4' | a:1695 | 4.59 | 1.0 | 39.2 |  | 10.62 | 16 | C | O3' | a:1832 | 4.67 | 1.0 | 34.4 |  | 9.86 | 17 | Water 6 | | c:406 | 4.69 | 1.0 | 42.9 | 2.45 [HIS:ND1]c:15 | 4.18 |

  
**1^st^ shell coordination angles and distances (*trans* in "blue"; *cys* in "green"; deviations in "orange"):**   
• 1-2(W/O2): 137; 1-3(W/W): 83; 1-4(W/O2'): 106; 2-3(O2/W): 82; 2-4(O2/O2'): 114; 3-4(W/O2'): 95;   
• 1-2(W/O2): 5.1; 1-3(W/W): 3.6; 1-4(W/O2'): 4.4; 2-3(O2/W): 3.7; 2-4(O2/O2'): 4.8; 3-4(W/O2'): 4.2;   
  
**4.39 ± 1.42: the average coodination distance of the 1^st^ six ligands could correspond to an hexahydrated Mg^2+^.**  
**8: Number of potential Water contacts**   
**2 : Number of second shell contacts involving water or ions**   
**25%: Percentage of satisfied second shell Water contacts**   
• Contact(s): /O2/O2'/O_w_/O_w_/  
• Contact code: O_r_O_b_2O_w_  
→ Coordination < 6 for K ←  
Go to ToC

### K:a:3097

|  | Res. type | Atom | Residue | Distance | Occupancy | B-factor | Closest atom(s) to a 1^st^ shell water | EM\_dens. |
| --- | --- | --- | --- | --- | --- | --- | --- | --- |
| **Ion** | | | |  | 1.0 | 30.0 |  | 7.61 |
 1 | C | O2' | a:1832 | 2.85 | 1.0 | 29.5 |  | 12.66 | 2 | C | O2 | a:1832 | 2.89 | 1.0 | 27.1 |  | 15.02 | 3 | U | N3 | a:1695 | 3.27 | 1.0 | 19.5 |  | 16.21 | 4 | U | O4 | a:1695 | 3.72 | 1.0 | 27.4 |  | 13.32 | 5 | Water 1 | | a:5717 | 3.73 | 1.0 | 48.7 |  | 2.57 | 6 | G | O4' | a:1833 | 3.81 | 1.0 | 35.5 |  | 11.01 | 7 | Water 2 | | a:8252 | 3.84 | 1.0 | 45.8 | 2.90 [G:N2]a:1979 | 4.71 | 8 | U | N1 | a:1695 | 4.07 | 1.0 | 27.8 |  | 12.23 | 9 | **K** | K | a:3096 | 4.15 | 1.0 | 30.0 |  | 10.55 | 10 | U | O2 | a:1695 | 4.19 | 1.0 | 28.3 |  | 13.15 | 11 | A | N1 | a:1831 | 4.31 | 1.0 | 19.5 |  | 17.55 | 12 | C | N1 | a:1832 | 4.34 | 1.0 | 23.8 |  | 14.75 | 13 | Water 3 | | a:9269 | 4.67 | 1.0 | 44.2 |  | 3.03 | 14 | U | O4' | a:1695 | 4.76 | 1.0 | 39.2 |  | 10.62 |

  
**1^st^ shell coordination angles and distances (*trans* in "blue"; *cys* in "green"; deviations in "orange"):**   
• 1-2(O2'/O2): 82;   
• 1-2(O2'/O2): 3.8;   
  
**4.70 ± 1.44: the average coodination distance of the 1^st^ six ligands could correspond to an hexahydrated Mg^2+^.**  
**4: Number of potential Water contacts**   
**1 : Number of second shell contacts involving water or ions**   
**25%: Percentage of satisfied second shell Water contacts**   
• Contact(s): /O2/O2'/  
• Contact code: O_r_O_b_  
→ Coordination < 6 for K ←  
Go to ToC

### K:a:3098

|  | Res. type | Atom | Residue | Distance | Occupancy | B-factor | Closest atom(s) to a 1^st^ shell water | EM\_dens. |
| --- | --- | --- | --- | --- | --- | --- | --- | --- |
| **Ion** | | | |  | 1.0 | 30.0 |  | 16.97 |
 1 | G | O6 | a:602 | 2.80 | 1.0 | 30.0 |  | 11.39 | 2 | Water 1 | | a:6581 | 2.85 | 1.0 | 39.7 | 2.78 [G:N7]a:602 | 7.78 | 3 | G | O6 | a:658 | 2.88 | 1.0 | 32.5 |  | 9.46 | 4 | U | O4 | a:659 | 2.95 | 1.0 | 32.9 |  | 10.62 | 5 | Water 2 | | a:9844 | 3.09 | 1.0 | 47.4 |  | 3.64 | 6 | A | N6 | a:601 | 3.13 | 1.0 | 22.0 |  | 11.78 | 7 | C | N4 | a:603 | 3.85 | 1.0 | 24.7 |  | 9.98 | 8 | U | N3 | a:659 | 3.89 | 1.0 | 19.1 |  | 11.72 | 9 | Water 3 | | a:6835 | 3.98 | 1.0 | 41.3 |  | 6.29 | 10 | Water 4 | | a:7892 | 4.18 | 1.0 | 46.0 |  | 3.40 | 11 | G | N7 | a:602 | 4.34 | 1.0 | 21.4 |  | 13.35 | 12 | U | O4 | a:660 | 4.56 | 1.0 | 31.6 |  | 11.60 | 13 | G | N7 | a:658 | 4.79 | 1.0 | 30.2 |  | 8.18 |

  
**1^st^ shell coordination angles and distances (*trans* in "blue"; *cys* in "green"; deviations in "orange"):**   
• 1-2(O6/W): 84; 1-3(O6/O6): 68; 1-4(O6/O4): 81; 1-5(O6/W): 122; 1-6(O6/N6): 70; 2-3(W/O6): 109; 2-4(W/O4): 162; 2-5(W/W): 73; 2-6(W/N6): 75; 3-4(O6/O4): 75; 3-5(O6/W): 71; 3-6(O6/N6): 137; 4-5(O4/W): 124; 4-6(O4/N6): 90; 5-6(W/N6): 144;   
• 1-2(O6/W): 3.8; 1-3(O6/O6): 3.2; 1-4(O6/O4): 3.7; 1-5(O6/W): 5.2; 1-6(O6/N6): 3.4; 2-3(W/O6): 4.7; 2-4(W/O4): 5.7; 2-5(W/W): 3.5; 2-6(W/N6): 3.6; 3-4(O6/O4): 3.6; 3-5(O6/W): 3.5; 3-6(O6/N6): 5.6; 4-5(O4/W): 5.3; 4-6(O4/N6): 4.3; 5-6(W/N6): 5.9;   
  
**4.24 ± 1.31: the average coodination distance of the 1^st^ six ligands could correspond to an hexahydrated Mg^2+^.**  
**4: Number of potential Water contacts**   
**1 : Number of second shell contacts involving water or ions**   
**25%: Percentage of satisfied second shell Water contacts**   
• Contact(s): /N6/O4/O6/O6/O_w_/O_w_/  
• Contact code: 3O_b_2O_w_Anion  
Go to ToC

### K:a:3099

|  | Res. type | Atom | Residue | Distance | Occupancy | B-factor | Closest atom(s) to a 1^st^ shell water | EM\_dens. |
| --- | --- | --- | --- | --- | --- | --- | --- | --- |
| **Ion** | | | |  | 1.0 | 30.0 |  | 22.15 |
 1 | Water 1 | | a:4561 | 2.60 | 1.0 | 43.7 | 2.64 [U:O4]a:660 | 6.63 | 2 | Water 2 | | a:9198 | 2.72 | 1.0 | 33.8 |  | 9.84 | 3 | G | O6 | a:599 | 2.76 | 1.0 | 28.6 |  | 13.75 | 4 | G | O6 | a:661 | 2.87 | 1.0 | 31.5 |  | 13.07 | 5 | U | O4 | a:600 | 2.97 | 1.0 | 33.3 |  | 11.83 | 6 | Water 3 | | a:4895 | 3.23 | 1.0 | 43.3 |  | 5.02 | 7 | Water 4 | | a:6646 | 3.27 | 1.0 | 40.8 | 2.78 [G:N7]a:599 | 4.68 | 8 | U | O4 | a:660 | 3.46 | 1.0 | 31.6 |  | 11.60 | 9 | G | N7 | a:661 | 4.39 | 1.0 | 22.3 |  | 12.27 | 10 | Water 5 | | a:8105 | 4.48 | 1.0 | 42.5 |  | 4.00 | 11 | G | N7 | a:599 | 4.66 | 1.0 | 21.8 |  | 14.32 |

  
**1^st^ shell coordination angles and distances (*trans* in "blue"; *cys* in "green"; deviations in "orange"):**   
• 1-2(W/W): 82; 1-3(W/O6): 162; 1-4(W/O6): 130; 1-5(W/O4): 104; 2-3(W/O6): 105; 2-4(W/O6): 84; 2-5(W/O4): 162; 3-4(O6/O6): 67; 3-5(O6/O4): 74; 4-5(O6/O4): 79;   
• 1-2(W/W): 3.5; 1-3(W/O6): 5.3; 1-4(W/O6): 5.0; 1-5(W/O4): 4.4; 2-3(W/O6): 4.4; 2-4(W/O6): 3.8; 2-5(W/O4): 5.6; 3-4(O6/O6): 3.1; 3-5(O6/O4): 3.5; 4-5(O6/O4): 3.7;   
  
**3.98 ± 1.19: the average coodination distance of the 1^st^ six ligands could correspond to an hexahydrated Mg^2+^.**  
**8: Number of potential Water contacts**   
**2 : Number of second shell contacts involving water or ions**   
**25%: Percentage of satisfied second shell Water contacts**   
• Contact(s): /O4/O6/O6/O_w_/O_w_/  
• Contact code: 3O_b_2O_w_  
→ Coordination < 6 for K ←  
Go to ToC

### K:a:3100

|  | Res. type | Atom | Residue | Distance | Occupancy | B-factor | Closest atom(s) to a 1^st^ shell water | EM\_dens. |
| --- | --- | --- | --- | --- | --- | --- | --- | --- |
| **Ion** | | | |  | 1.0 | 30.0 |  | 30.41 |
 1 | Water 1 | | a:3642 | 2.68 | 1.0 | 30.6 | 2.50 [A:OP1]a:470 | 14.28 | 2 | A | OP1 | a:471 | 2.82 | 1.0 | 34.0 |  | 11.01 | 3 | G | O3' | a:452 | 2.87 | 1.0 | 25.9 |  | 13.14 | 4 | Water 2 | | a:8800 | 2.96 | 1.0 | 22.6 | 2.24 [MG:MG]a:3083 | 16.66 | 5 | Water 3 | | e:475 | 2.98 | 1.0 | 35.4 |  | 6.73 | 6 | A | OP1 | a:453 | 3.04 | 1.0 | 23.9 |  | 15.02 | 7 | Water 4 | | e:418 | 3.17 | 1.0 | 26.9 | 2.65 [SER:OG]e:80 | 13.62 | 8 | A | OP2 | a:471 | 3.25 | 1.0 | 23.7 |  | 15.41 | 9 | SER | OG | e:80 | 3.90 | 1.0 | 29.9 |  | 12.93 | 10 | G | O2' | a:452 | 3.92 | 1.0 | 22.4 |  | 16.16 | 11 | **MG** | MG | a:3083 | 4.10 | 1.0 | 24.4 |  | 24.35 | 12 | A | O3' | a:470 | 4.17 | 1.0 | 31.0 |  | 11.56 | 13 | Water 5 | | e:423 | 4.36 | 1.0 | 33.5 | 2.70 [SER:OG]e:80 | 10.43 | 14 | A | O5' | a:453 | 4.46 | 1.0 | 22.9 |  | 15.46 | 15 | Water 6 | | a:5422 | 4.61 | 1.0 | 21.3 | 2.32 [MG:MG]a:3083 | 15.55 | 16 | A | OP1 | a:470 | 4.70 | 1.0 | 29.1 |  | 11.26 | 17 | Water 7 | | a:4822 | 4.71 | 1.0 | 25.5 | 2.66 [A:OP2]a:471 | 13.46 | 18 | A | OP2 | a:453 | 4.76 | 1.0 | 24.9 |  | 15.45 |

  
**1^st^ shell coordination angles and distances (*trans* in "blue"; *cys* in "green"; deviations in "orange"):**   
• 1-2(W/OP1): 93; 1-3(W/O3'): 96; 1-4(W/W): 154; 1-5(W/W): 88; 1-6(W/OP1): 144; 1-7(W/W): 104; 2-3(OP1/O3'): 156; 2-4(OP1/W): 70; 2-5(OP1/W): 129; 2-6(OP1/OP1): 123; 2-7(OP1/W): 75; 3-4(O3'/W): 109; 3-5(O3'/W): 74; 3-6(O3'/OP1): 49; 3-7(O3'/W): 125; 4-5(W/W): 88; 4-6(W/OP1): 59; 4-7(W/W): 54; 5-6(W/OP1): 75; 5-7(W/W): 56; 6-7(OP1/W): 93;   
• 1-2(W/OP1): 4.0; 1-3(W/O3'): 4.1; 1-4(W/W): 5.5; 1-5(W/W): 3.9; 1-6(W/OP1): 5.4; 1-7(W/W): 4.6; 2-3(OP1/O3'): 5.6; 2-4(OP1/W): 3.3; 2-5(OP1/W): 5.2; 2-6(OP1/OP1): 5.2; 2-7(OP1/W): 3.7; 3-4(O3'/W): 4.7; 3-5(O3'/W): 3.5; 3-6(O3'/OP1): 2.5; 3-7(O3'/W): 5.3; 4-5(W/W): 4.1; 4-6(W/OP1): 3.0; 4-7(W/W): 2.8; 5-6(W/OP1): 3.6; 5-7(W/W): 2.9; 6-7(OP1/W): 4.5;   
  
**3.96 ± 1.16: the average coodination distance of the 1^st^ six ligands could correspond to an hexahydrated Mg^2+^.**  
**8: Number of potential Water contacts**   
**3 : Number of second shell contacts involving water or ions**   
**38%: Percentage of satisfied second shell Water contacts**   
• Contact(s): /O3'/O_w_/O_w_/O_w_/O_w_/OP1/OP1/  
• Contact code: 2O_ph_O_r_4O_w_  
Go to ToC

### K:a:3101

|  | Res. type | Atom | Residue | Distance | Occupancy | B-factor | Closest atom(s) to a 1^st^ shell water | EM\_dens. |
| --- | --- | --- | --- | --- | --- | --- | --- | --- |
| **Ion** | | | |  | 1.0 | 30.0 |  | 19.70 |
 1 | A | N3 | a:2576 | 2.93 | 1.0 | 26.9 |  | 11.55 | 2 | A | O2' | a:2576 | 3.04 | 1.0 | 24.4 |  | 14.39 | 3 | C | OP2 | a:2577 | 3.11 | 1.0 | 31.5 |  | 10.61 | 4 | U | OP1 | a:2510 | 3.21 | 1.0 | 57.2 |  | 1.63 | 5 | Water 1 | | a:5178 | 3.21 | 1.0 | 37.7 |  | 6.87 | 6 | A | O4' | a:2457 | 3.39 | 1.0 | 28.4 |  | 11.40 | 7 | A | O3' | a:2576 | 3.67 | 1.0 | 26.5 |  | 13.86 | 8 | Water 2 | | a:9436 | 4.05 | 1.0 | 44.5 |  | 3.03 | 9 | A | O2' | a:2457 | 4.10 | 1.0 | 30.1 |  | 14.30 | 10 | Water 3 | | d:493 | 4.23 | 1.0 | 41.4 |  | 5.31 | 11 | Water 4 | | a:8211 | 4.49 | 1.0 | 44.4 |  | 4.49 | 12 | Water 5 | | a:3472 | 4.55 | 1.0 | 44.1 |  | 3.13 | 13 | C | O2' | a:2456 | 4.59 | 1.0 | 37.8 |  | 8.29 |

  
**1^st^ shell coordination angles and distances (*trans* in "blue"; *cys* in "green"; deviations in "orange"):**   
• 1-2(N3/O2'): 79; 1-3(N3/OP2): 140; 2-3(O2'/OP2): 71;   
• 1-2(N3/O2'): 3.8; 1-3(N3/OP2): 5.7; 2-3(O2'/OP2): 3.6;   
  
**4.43 ± 1.34: the average coodination distance of the 1^st^ six ligands could correspond to an hexahydrated Mg^2+^.**  
**4: Number of potential Water contacts**   
**0 : Number of second shell contacts involving water or ions**   
**0%: Percentage of satisfied second shell Water contacts**   
• Contact(s): /N3_A_/O2'/OP2/  
• Contact code: O_ph_O_r_N_b_  
→ Coordination < 6 for K ←  
Go to ToC

### K:a:3102

|  | Res. type | Atom | Residue | Distance | Occupancy | B-factor | Closest atom(s) to a 1^st^ shell water | EM\_dens. |
| --- | --- | --- | --- | --- | --- | --- | --- | --- |
| **Ion** | | | |  | 1.0 | 30.0 |  | 21.64 |
 1 | Water 1 | | a:4161 | 2.49 | 1.0 | 33.5 | 2.60 [G:O6]a:2701 | 9.21 | 2 | U | O4 | a:2702 | 2.74 | 1.0 | 28.7 |  | 14.77 | 3 | G | O6 | a:2713 | 2.81 | 1.0 | 23.9 |  | 15.97 | 4 | Water 2 | | a:6862 | 2.95 | 1.0 | 36.5 | 2.80 [G:N7]a:2713 | 8.21 | 5 | Water 3 | | a:8480 | 3.06 | 1.0 | 40.6 |  | 4.55 | 6 | G | O6 | a:2712 | 3.07 | 1.0 | 29.6 |  | 12.43 | 7 | G | O6 | a:2701 | 3.47 | 1.0 | 20.5 |  | 15.66 | 8 | U | N3 | a:2702 | 3.81 | 1.0 | 19.8 |  | 14.86 | 9 | Water 4 | | a:9792 | 3.91 | 1.0 | 46.5 |  | 3.62 | 10 | Water 5 | | a:3708 | 4.07 | 1.0 | 40.3 |  | 3.99 | 11 | Water 6 | | a:4782 | 4.08 | 1.0 | 38.1 |  | 4.38 | 12 | Water 7 | | a:6732 | 4.36 | 1.0 | 31.6 |  | 10.34 | 13 | G | N7 | a:2713 | 4.37 | 1.0 | 20.7 |  | 13.96 | 14 | Water 8 | | a:5409 | 4.65 | 1.0 | 46.1 |  | 4.30 |

  
**1^st^ shell coordination angles and distances (*trans* in "blue"; *cys* in "green"; deviations in "orange"):**   
• 1-2(W/O4): 79; 1-3(W/O6): 115; 1-4(W/W): 105; 1-5(W/W): 110; 1-6(W/O6): 170; 2-3(O4/O6): 82; 2-4(O4/W): 164; 2-5(O4/W): 82; 2-6(O4/O6): 101; 3-4(O6/W): 83; 3-5(O6/W): 127; 3-6(O6/O6): 75; 4-5(W/W): 110; 4-6(W/O6): 77; 5-6(W/O6): 60;   
• 1-2(W/O4): 3.3; 1-3(W/O6): 4.5; 1-4(W/W): 4.3; 1-5(W/W): 4.6; 1-6(W/O6): 5.5; 2-3(O4/O6): 3.6; 2-4(O4/W): 5.6; 2-5(O4/W): 3.8; 2-6(O4/O6): 4.5; 3-4(O6/W): 3.8; 3-5(O6/W): 5.3; 3-6(O6/O6): 3.6; 4-5(W/W): 4.9; 4-6(W/O6): 3.8; 5-6(W/O6): 3.1;   
  
**4.07 ± 1.26: the average coodination distance of the 1^st^ six ligands could correspond to an hexahydrated Mg^2+^.**  
**6: Number of potential Water contacts**   
**2 : Number of second shell contacts involving water or ions**   
**33%: Percentage of satisfied second shell Water contacts**   
• Contact(s): /O4/O6/O6/O_w_/O_w_/  
• Contact code: 3O_b_2O_w_  
• Contact(s) in the exclusion zone: /O_w_/  
→ Coordination < 6 for K ←  
Go to ToC

### K:a:3103

|  | Res. type | Atom | Residue | Distance | Occupancy | B-factor | Closest atom(s) to a 1^st^ shell water | EM\_dens. |
| --- | --- | --- | --- | --- | --- | --- | --- | --- |
| **Ion** | | | |  | 1.0 | 30.0 |  | 7.32 |
 1 | Water 1 | | a:6527 | 2.25 | 1.0 | 30.0 | 2.78 [PSU:O2]a:2584 | 9.34 | 2 | Water 2 | | a:6946 | 2.26 | 1.0 | 40.7 |  | 7.29 | 3 | Water 3 | | a:3464 | 2.69 | 1.0 | 38.9 |  | 7.55 | 4 | Water 4 | | a:6519 | 3.73 | 1.0 | 30.3 | 2.78 [G:OP2]a:2580 | 10.86 | 5 | C | OP2 | a:2511 | 4.01 | 1.0 | 41.0 |  | 4.39 | 6 | Water 5 | | a:8552 | 4.11 | 1.0 | 39.7 |  | 5.37 | 7 | G | N7 | a:2580 | 4.36 | 1.0 | 36.8 |  | 5.73 | 8 | G | O6 | a:2512 | 4.62 | 1.0 | 27.7 |  | 13.80 | 9 | PSU | O2 | a:2584 | 4.71 | 1.0 | 27.3 |  | 15.05 | 10 | G | OP2 | a:2580 | 4.75 | 1.0 | 26.2 |  | 14.54 | 11 | G | O6 | a:2580 | 4.75 | 1.0 | 42.7 |  | 5.56 |

  
**1^st^ shell coordination angles and distances (*trans* in "blue"; *cys* in "green"; deviations in "orange"):**   
• 1-2(W/W): 84; 1-3(W/W): 64; 2-3(W/W): 147;   
• 1-2(W/W): 3.0; 1-3(W/W): 2.7; 2-3(W/W): 4.8;   
  
**4.67 ± 1.70: the average coodination distance of the 1^st^ six ligands could correspond to an hexahydrated Mg^2+^.**  
**10: Number of potential Water contacts**   
**2 : Number of second shell contacts involving water or ions**   
**20%: Percentage of satisfied second shell Water contacts**   
• Contact(s): /O_w_/  
• Contact code: O_w_  
• Contact(s) in the exclusion zone: /O_w_/O_w_/  
→ Coordination < 6 for K ←  
Go to ToC

### K:a:3104

|  | Res. type | Atom | Residue | Distance | Occupancy | B-factor | Closest atom(s) to a 1^st^ shell water | EM\_dens. |
| --- | --- | --- | --- | --- | --- | --- | --- | --- |
| **Ion** | | | |  | 1.0 | 30.0 |  | 14.32 |
 1 | Water 1 | | a:5927 | 2.67 | 1.0 | 39.3 | 2.74 [A:N1]a:1030 | 5.63 | 2 | G | O2' | a:2468 | 2.95 | 1.0 | 35.9 |  | 8.27 | 3 | Water 2 | | a:5618 | 2.99 | 1.0 | 34.4 | 2.96 [G:N3]a:2491 | 8.13 | 4 | Water 3 | | a:9528 | 3.01 | 1.0 | 42.6 |  | 7.40 | 5 | G | N3 | a:2468 | 3.04 | 1.0 | 28.2 |  | 10.85 | 6 | A | OP2 | a:1128 | 3.08 | 1.0 | 36.3 |  | 7.50 | 7 | A | OP1 | a:1128 | 3.11 | 1.0 | 33.8 |  | 8.81 | 8 | C | O4' | a:2469 | 3.72 | 1.0 | 36.4 |  | 8.83 | 9 | Water 4 | | a:7262 | 3.78 | 1.0 | 46.5 |  | 4.66 | 10 | Water 5 | | a:6386 | 3.89 | 1.0 | 38.0 |  | 6.95 | 11 | G | O3' | a:1127 | 4.43 | 1.0 | 35.8 |  | 9.63 | 12 | A | N1 | a:1030 | 4.73 | 1.0 | 26.2 |  | 13.77 | 13 | G | N3 | a:2491 | 4.73 | 1.0 | 24.0 |  | 13.14 |

  
**1^st^ shell coordination angles and distances (*trans* in "blue"; *cys* in "green"; deviations in "orange"):**   
• 1-2(W/O2'): 129; 1-3(W/W): 83; 1-4(W/W): 141; 1-5(W/N3): 92; 1-6(W/OP2): 101; 1-7(W/OP1): 72; 2-3(O2'/W): 147; 2-4(O2'/W): 90; 2-5(O2'/N3): 91; 2-6(O2'/OP2): 66; 2-7(O2'/OP1): 114; 3-4(W/W): 59; 3-5(W/N3): 80; 3-6(W/OP2): 121; 3-7(W/OP1): 79; 4-5(W/N3): 92; 4-6(W/OP2): 90; 4-7(W/OP1): 89; 5-6(N3/OP2): 156; 5-7(N3/OP1): 155; 6-7(OP2/OP1): 48;   
• 1-2(W/O2'): 5.1; 1-3(W/W): 3.8; 1-4(W/W): 5.4; 1-5(W/N3): 4.1; 1-6(W/OP2): 4.4; 1-7(W/OP1): 3.4; 2-3(O2'/W): 5.7; 2-4(O2'/W): 4.2; 2-5(O2'/N3): 4.3; 2-6(O2'/OP2): 3.3; 2-7(O2'/OP1): 5.1; 3-4(W/W): 3.0; 3-5(W/N3): 3.9; 3-6(W/OP2): 5.3; 3-7(W/OP1): 3.9; 4-5(W/N3): 4.4; 4-6(W/OP2): 4.3; 4-7(W/OP1): 4.3; 5-6(N3/OP2): 6.0; 5-7(N3/OP1): 6.0; 6-7(OP2/OP1): 2.5;   
  
**4.10 ± 1.22: the average coodination distance of the 1^st^ six ligands could correspond to an hexahydrated Mg^2+^.**  
**6: Number of potential Water contacts**   
**2 : Number of second shell contacts involving water or ions**   
**33%: Percentage of satisfied second shell Water contacts**   
• Contact(s): /N3_G_/O2'/O_w_/O_w_/O_w_/OP1/OP2/  
• Contact code: 2O_ph_O_r_N_b_3O_w_  
Go to ToC

### K:a:3105

|  | Res. type | Atom | Residue | Distance | Occupancy | B-factor | Closest atom(s) to a 1^st^ shell water | EM\_dens. |
| --- | --- | --- | --- | --- | --- | --- | --- | --- |
| **Ion** | | | |  | 1.0 | 30.0 |  | 28.96 |
 1 | A | O5' | a:1940 | 2.60 | 1.0 | 33.1 |  | 11.81 | 2 | 5MC | N3 | a:1966 | 2.84 | 1.0 | 26.7 |  | 13.39 | 3 | A | N3 | a:1940 | 2.89 | 1.0 | 32.4 |  | 11.39 | 4 | U | O4 | a:1944 | 2.92 | 1.0 | 37.8 |  | 9.43 | 5 | 5MC | O2 | a:1966 | 2.94 | 1.0 | 37.0 |  | 11.88 | 6 | A | OP2 | a:1940 | 3.46 | 1.0 | 35.4 |  | 7.86 | 7 | A | O4' | a:1940 | 3.70 | 1.0 | 31.9 |  | 10.01 | 8 | C | N4 | a:1965 | 3.77 | 1.0 | 28.5 |  | 11.85 | 9 | G | O2' | a:1939 | 3.89 | 1.0 | 32.3 |  | 11.94 | 10 | G | N3 | a:1939 | 4.34 | 1.0 | 28.3 |  | 12.03 | 11 | G | O3' | a:1939 | 4.41 | 1.0 | 35.5 |  | 10.88 | 12 | Water 1 | | a:4074 | 4.47 | 1.0 | 31.1 | 2.58 [G:O2']a:1939 | 10.17 | 13 | Water 2 | | a:6580 | 4.58 | 1.0 | 35.6 | 2.78 [U:N3]a:1944 | 7.41 | 14 | A | N1 | a:1940 | 4.63 | 1.0 | 23.8 |  | 13.70 | 15 | 5MC | N1 | a:1966 | 4.66 | 1.0 | 27.8 |  | 10.49 | 16 | Water 3 | | a:4043 | 4.77 | 1.0 | 39.2 |  | 7.66 |

  
**1^st^ shell coordination angles and distances (*trans* in "blue"; *cys* in "green"; deviations in "orange"):**   
• 1-2(O5'/N3): 116; 1-3(O5'/N3): 97; 1-4(O5'/O4): 105; 1-5(O5'/O2): 160; 2-3(N3/N3): 118; 2-4(N3/O4): 121; 2-5(N3/O2): 46; 3-4(N3/O4): 96; 3-5(N3/O2): 89; 4-5(O4/O2): 94;   
• 1-2(O5'/N3): 4.6; 1-3(O5'/N3): 4.1; 1-4(O5'/O4): 4.4; 1-5(O5'/O2): 5.5; 2-3(N3/N3): 4.9; 2-4(N3/O4): 5.0; 2-5(N3/O2): 2.2; 3-4(N3/O4): 4.3; 3-5(N3/O2): 4.1; 4-5(O4/O2): 4.3;   
  
**4.19 ± 1.30: the average coodination distance of the 1^st^ six ligands could correspond to an hexahydrated Mg^2+^.**  
**0: Number of potential Water contacts**   
• Contact(s): /N3/N3_A_/O2/O4/O5'/  
• Contact code: O_r_2O_b_N_b_  
→ Coordination < 6 for K ←  
Go to ToC

### K:a:3106

|  | Res. type | Atom | Residue | Distance | Occupancy | B-factor | Closest atom(s) to a 1^st^ shell water | EM\_dens. |
| --- | --- | --- | --- | --- | --- | --- | --- | --- |
| **Ion** | | | |  | 1.0 | 30.0 |  | 14.63 |
 1 | Water 1 | | a:4604 | 2.73 | 1.0 | 39.3 | 2.64 [A:OP2]a:1775 | 7.87 | 2 | G | O6 | a:1979 | 2.86 | 1.0 | 29.3 |  | 13.11 | 3 | Water 2 | | a:9027 | 2.91 | 1.0 | 42.0 |  | 6.59 | 4 | Water 3 | | a:8384 | 2.99 | 1.0 | 37.5 | 2.92 [G:N7]a:1979 | 8.55 | 5 | Water 4 | | a:8709 | 2.99 | 1.0 | 38.2 | 2.96 [C:N4]a:1978 | 5.59 | 6 | Water 5 | | a:7955 | 3.11 | 1.0 | 33.3 |  | 9.72 | 7 | U | O4 | a:1980 | 3.36 | 1.0 | 32.1 |  | 11.50 | 8 | Water 6 | | a:8626 | 3.64 | 1.0 | 41.3 | 2.95 [U:O4]a:1980 | 5.01 | 9 | Water 7 | | a:7224 | 4.12 | 1.0 | 42.9 | 2.82 [U:O4]a:1980 | 7.03 | 10 | Water 8 | | a:6836 | 4.22 | 1.0 | 27.4 | 2.95 [U:N3]a:1980 | 11.85 | 11 | A | OP2 | a:1775 | 4.24 | 1.0 | 29.0 |  | 10.52 | 12 | G | N7 | a:1979 | 4.35 | 1.0 | 25.5 |  | 10.29 | 13 | Water 9 | | a:6430 | 4.48 | 1.0 | 39.1 | 2.77 [A:OP2]a:1775 | 7.11 | 14 | Water 10 | | a:7835 | 4.76 | 1.0 | 45.6 |  | 3.22 |

  
**1^st^ shell coordination angles and distances (*trans* in "blue"; *cys* in "green"; deviations in "orange"):**   
• 1-2(W/O6): 147; 1-3(W/W): 97; 1-4(W/W): 97; 1-5(W/W): 107; 1-6(W/W): 80; 2-3(O6/W): 105; 2-4(O6/W): 87; 2-5(O6/W): 104; 2-6(O6/W): 91; 3-4(W/W): 131; 3-5(W/W): 69; 3-6(W/W): 60; 4-5(W/W): 62; 4-6(W/W): 169; 5-6(W/W): 128;   
• 1-2(W/O6): 5.4; 1-3(W/W): 4.2; 1-4(W/W): 4.3; 1-5(W/W): 4.6; 1-6(W/W): 3.8; 2-3(O6/W): 4.6; 2-4(O6/W): 4.0; 2-5(O6/W): 4.6; 2-6(O6/W): 4.3; 3-4(W/W): 5.4; 3-5(W/W): 3.3; 3-6(W/W): 3.0; 4-5(W/W): 3.1; 4-6(W/W): 6.1; 5-6(W/W): 5.5;   
  
**4.10 ± 1.23: the average coodination distance of the 1^st^ six ligands could correspond to an hexahydrated Mg^2+^.**  
**12: Number of potential Water contacts**   
**4 : Number of second shell contacts involving water or ions**   
**33%: Percentage of satisfied second shell Water contacts**   
• Contact(s): /O6/O_w_/O_w_/O_w_/O_w_/O_w_/  
• Contact code: O_b_5O_w_  
Go to ToC

### K:a:3107

|  | Res. type | Atom | Residue | Distance | Occupancy | B-factor | Closest atom(s) to a 1^st^ shell water | EM\_dens. |
| --- | --- | --- | --- | --- | --- | --- | --- | --- |
| **Ion** | | | |  | 1.0 | 30.0 |  | 15.75 |
 1 | Water 1 | | a:8848 | 2.46 | 1.0 | 47.6 |  | 3.65 | 2 | Water 2 | | a:9496 | 2.73 | 1.0 | 47.7 | 2.09 [MG:MG]a:3022 | 5.75 | 3 | G | O6 | a:2574 | 2.89 | 1.0 | 22.3 |  | 15.96 | 4 | G | O6 | a:2573 | 2.98 | 1.0 | 24.2 |  | 14.50 | 5 | U | O4 | a:2518 | 3.05 | 1.0 | 23.6 |  | 15.83 | 6 | A | N6 | a:2517 | 3.19 | 1.0 | 17.3 |  | 13.59 | 7 | Water 3 | | a:6303 | 3.62 | 1.0 | 36.0 |  | 8.88 | 8 | U | O4 | a:2575 | 3.81 | 1.0 | 22.9 |  | 15.39 | 9 | **MG** | MG | a:3022 | 4.18 | 1.0 | 51.9 |  | 9.69 | 10 | Water 4 | | a:4985 | 4.19 | 1.0 | 38.7 |  | 4.43 | 11 | G | N7 | a:2574 | 4.23 | 1.0 | 16.3 |  | 14.01 | 12 | A | N1 | a:2517 | 4.57 | 1.0 | 21.1 |  | 17.39 |

  
**1^st^ shell coordination angles and distances (*trans* in "blue"; *cys* in "green"; deviations in "orange"):**   
• 1-2(W/W): 99; 1-3(W/O6): 138; 1-4(W/O6): 93; 1-5(W/O4): 64; 1-6(W/N6): 93; 2-3(W/O6): 117; 2-4(W/O6): 126; 2-5(W/O4): 125; 2-6(W/N6): 65; 3-4(O6/O6): 83; 3-5(O6/O4): 77; 3-6(O6/N6): 85; 4-5(O6/O4): 108; 4-6(O6/N6): 167; 5-6(O4/N6): 64;   
• 1-2(W/W): 4.0; 1-3(W/O6): 5.0; 1-4(W/O6): 4.0; 1-5(W/O4): 3.0; 1-6(W/N6): 4.1; 2-3(W/O6): 4.8; 2-4(W/O6): 5.1; 2-5(W/O4): 5.1; 2-6(W/N6): 3.2; 3-4(O6/O6): 3.9; 3-5(O6/O4): 3.7; 3-6(O6/N6): 4.1; 4-5(O6/O4): 4.9; 4-6(O6/N6): 6.1; 5-6(O4/N6): 3.3;   
  
**4.12 ± 1.28: the average coodination distance of the 1^st^ six ligands could correspond to an hexahydrated Mg^2+^.**  
**6: Number of potential Water contacts**   
**1 : Number of second shell contacts involving water or ions**   
**17%: Percentage of satisfied second shell Water contacts**   
• Contact(s): /N6/O4/O6/O6/O_w_/  
• Contact code: 3O_b_O_w_Anion  
• Contact(s) in the exclusion zone: /O_w_/  
→ Coordination < 6 for K ←  
Go to ToC

### K:a:3108

|  | Res. type | Atom | Residue | Distance | Occupancy | B-factor | Closest atom(s) to a 1^st^ shell water | EM\_dens. |
| --- | --- | --- | --- | --- | --- | --- | --- | --- |
| **Ion** | | | |  | 1.0 | 30.0 |  | 6.48 |
 1 | G | O6 | a:709 | 2.83 | 1.0 | 38.2 |  | 7.43 | 2 | U | O4 | a:726 | 3.03 | 1.0 | 40.3 |  | 6.35 | 3 | Water 1 | | a:3805 | 3.20 | 1.0 | 43.2 | 2.54 [G:N7]a:709 | 5.58 | 4 | Water 2 | | a:3651 | 3.29 | 1.0 | 44.8 | 2.50 [A:OP2]a:708 | 3.92 | 5 | C | N4 | a:725 | 3.66 | 1.0 | 46.5 |  | 4.67 | 6 | G | O6 | a:710 | 3.77 | 1.0 | 42.4 |  | 4.92 | 7 | G | N2 | a:727 | 3.88 | 1.0 | 31.9 |  | 7.98 | 8 | U | N3 | a:726 | 3.96 | 1.0 | 30.5 |  | 8.82 | 9 | G | N7 | a:709 | 4.35 | 1.0 | 29.5 |  | 8.67 | 10 | A | OP2 | a:708 | 4.47 | 1.0 | 38.3 |  | 6.44 | 11 | G | N3 | a:727 | 4.75 | 1.0 | 31.3 |  | 9.77 | 12 | A | OP2 | a:707 | 4.78 | 1.0 | 33.5 |  | 8.69 |

  
**1^st^ shell coordination angles and distances (*trans* in "blue"; *cys* in "green"; deviations in "orange"):**   
• 1-2(O6/O4): 77;   
• 1-2(O6/O4): 3.6;   
  
**4.61 ± 1.41: the average coodination distance of the 1^st^ six ligands could correspond to an hexahydrated Mg^2+^.**  
**4: Number of potential Water contacts**   
**2 : Number of second shell contacts involving water or ions**   
**50%: Percentage of satisfied second shell Water contacts**   
• Contact(s): /O4/O6/  
• Contact code: 2O_b_  
→ Coordination < 6 for K ←  
Go to ToC

### K:a:3109

|  | Res. type | Atom | Residue | Distance | Occupancy | B-factor | Closest atom(s) to a 1^st^ shell water | EM\_dens. |
| --- | --- | --- | --- | --- | --- | --- | --- | --- |
| **Ion** | | | |  | 1.0 | 30.0 |  | 14.01 |
 1 | G7M | OP2 | a:2073 | 2.73 | 1.0 | 42.4 |  | 6.59 | 2 | U | OP1 | a:2248 | 2.77 | 1.0 | 32.7 |  | 13.63 | 3 | Water 1 | | a:9752 | 2.96 | 1.0 | 44.4 |  | 3.88 | 4 | A | O3' | a:197 | 3.12 | 1.0 | 26.5 |  | 14.38 | 5 | A | O2' | a:197 | 3.34 | 1.0 | 26.2 |  | 14.35 | 6 | Water 2 | | a:6738 | 3.35 | 1.0 | 39.5 |  | 6.81 | 7 | A | OP1 | a:2074 | 3.37 | 1.0 | 37.4 |  | 5.61 | 8 | C | OP1 | a:198 | 3.46 | 1.0 | 30.1 |  | 11.83 | 9 | G7M | O5' | a:2073 | 3.54 | 1.0 | 38.4 |  | 7.19 | 10 | Water 3 | | a:8219 | 3.60 | 1.0 | 39.8 | 2.90 [A:OP1]a:2074 | 7.52 | 11 | Water 4 | | a:3717 | 3.65 | 1.0 | 28.9 | 2.52 [C:OP1]a:198 | 12.68 | 12 | A | OP2 | a:2074 | 3.79 | 1.0 | 33.5 |  | 8.09 | 13 | U | OP2 | a:2248 | 4.33 | 1.0 | 29.9 |  | 11.64 | 14 | G7M | O3' | a:2073 | 4.36 | 1.0 | 32.2 |  | 9.51 | 15 | Water 5 | | a:6042 | 4.58 | 1.0 | 38.4 | 2.74 [A:O2']a:197 | 8.45 | 16 | C | O5' | a:198 | 4.70 | 1.0 | 22.5 |  | 15.92 | 17 | U | O3' | a:2072 | 4.70 | 1.0 | 37.6 |  | 9.12 |

  
**1^st^ shell coordination angles and distances (*trans* in "blue"; *cys* in "green"; deviations in "orange"):**   
• 1-2(OP2/OP1): 126; 1-3(OP2/W): 88; 1-4(OP2/O3'): 70; 2-3(OP1/W): 77; 2-4(OP1/O3'): 79; 3-4(W/O3'): 129;   
• 1-2(OP2/OP1): 4.9; 1-3(OP2/W): 4.0; 1-4(OP2/O3'): 3.4; 2-3(OP1/W): 3.6; 2-4(OP1/O3'): 3.8; 3-4(W/O3'): 5.5;   
  
**4.18 ± 1.25: the average coodination distance of the 1^st^ six ligands could correspond to an hexahydrated Mg^2+^.**  
**4: Number of potential Water contacts**   
**0 : Number of second shell contacts involving water or ions**   
**0%: Percentage of satisfied second shell Water contacts**   
• Contact(s): /O3'/O_w_/OP1/OP2/  
• Contact code: 2O_ph_O_r_O_w_  
→ Coordination < 6 for K ←  
Go to ToC

### K:a:3110

|  | Res. type | Atom | Residue | Distance | Occupancy | B-factor | Closest atom(s) to a 1^st^ shell water | EM\_dens. |
| --- | --- | --- | --- | --- | --- | --- | --- | --- |
| **Ion** | | | |  | 1.0 | 30.0 |  | 19.46 |
 1 | G | OP1 | a:2451 | 2.81 | 1.0 | 24.2 |  | 13.46 | 2 | Water 1 | | a:7190 | 2.85 | 1.0 | 22.0 | 1.98 [MG:MG]a:3353 | 14.61 | 3 | U | O2' | a:569 | 2.98 | 1.0 | 21.9 |  | 16.63 | 4 | Water 2 | | a:8040 | 3.06 | 1.0 | 30.6 |  | 11.92 | 5 | Water 3 | | a:4008 | 3.13 | 1.0 | 29.3 |  | 13.55 | 6 | Water 4 | | a:3457 | 3.22 | 1.0 | 32.2 | 2.40 [G:OP1]a:810 | 10.16 | 7 | Water 5 | | a:8014 | 3.29 | 1.0 | 35.9 | 2.15 [MG:MG]a:3353 | 7.46 | 8 | G | OP2 | a:2451 | 3.63 | 1.0 | 27.2 |  | 14.53 | 9 | **MG** | MG | a:3353 | 3.89 | 1.0 | 43.4 |  | 14.26 | 10 | Water 6 | | a:4615 | 3.90 | 1.0 | 33.5 | 2.80 [K:K]a:3111 | 8.50 | 11 | Water 7 | | a:8656 | 4.22 | 1.0 | 34.7 |  | 8.96 | 12 | G | OP1 | a:2450 | 4.26 | 1.0 | 32.2 |  | 13.70 | 13 | Water 8 | | a:7561 | 4.27 | 1.0 | 35.3 |  | 7.72 | 14 | Water 9 | | a:4567 | 4.49 | 1.0 | 38.8 |  | 5.99 | 15 | Water 10 | | a:9330 | 4.52 | 1.0 | 32.1 | 2.22 [MG:MG]a:3172 | 9.80 | 16 | **K** | K | a:3111 | 4.55 | 1.0 | 30.0 |  | 31.46 | 17 | U | O4' | a:569 | 4.59 | 1.0 | 24.4 |  | 19.24 | 18 | G | O5' | a:2451 | 4.67 | 1.0 | 24.4 |  | 13.31 | 19 | G | OP1 | a:810 | 4.70 | 1.0 | 24.5 |  | 14.25 | 20 | **MG** | MG | a:3172 | 4.78 | 1.0 | 52.7 |  | 12.18 |

  
**1^st^ shell coordination angles and distances (*trans* in "blue"; *cys* in "green"; deviations in "orange"):**   
• 1-2(OP1/W): 94; 1-3(OP1/O2'): 97; 1-4(OP1/W): 106; 1-5(OP1/W): 51; 2-3(W/O2'): 167; 2-4(W/W): 83; 2-5(W/W): 126; 3-4(O2'/W): 88; 3-5(O2'/W): 58; 4-5(W/W): 73;   
• 1-2(OP1/W): 4.1; 1-3(OP1/O2'): 4.3; 1-4(OP1/W): 4.7; 1-5(OP1/W): 2.6; 2-3(W/O2'): 5.8; 2-4(W/W): 3.9; 2-5(W/W): 5.3; 3-4(O2'/W): 4.2; 3-5(O2'/W): 3.0; 4-5(W/W): 3.7;   
  
**4.16 ± 1.23: the average coodination distance of the 1^st^ six ligands could correspond to an hexahydrated Mg^2+^.**  
**10: Number of potential Water contacts**   
**3 : Number of second shell contacts involving water or ions**   
**30%: Percentage of satisfied second shell Water contacts**   
• Contact(s): /O2'/O_w_/O_w_/O_w_/OP1/  
• Contact code: O_ph_O_r_3O_w_  
→ Coordination < 6 for K ←  
Go to ToC

### K:a:3111

|  | Res. type | Atom | Residue | Distance | Occupancy | B-factor | Closest atom(s) to a 1^st^ shell water | EM\_dens. |
| --- | --- | --- | --- | --- | --- | --- | --- | --- |
| **Ion** | | | |  | 1.0 | 30.0 |  | 31.46 |
 1 | G | OP1 | a:2451 | 2.74 | 1.0 | 24.2 |  | 13.46 | 2 | Water 1 | | a:4615 | 2.80 | 1.0 | 33.5 |  | 8.50 | 3 | G | O5' | a:2451 | 2.87 | 1.0 | 24.4 |  | 13.31 | 4 | Water 2 | | a:6561 | 2.92 | 1.0 | 22.0 | 2.84 [A:O5']a:2452 | 16.66 | 5 | Water 3 | | a:7008 | 2.96 | 1.0 | 20.7 | 2.14 [MG:MG]a:3325 | 18.60 | 6 | Water 4 | | a:6699 | 3.11 | 1.0 | 19.6 | 2.23 [MG:MG]a:3325 | 18.66 | 7 | Water 5 | | a:7482 | 3.18 | 1.0 | 36.0 |  | 7.79 | 8 | A | OP1 | a:2452 | 3.21 | 1.0 | 19.4 |  | 18.69 | 9 | **MG** | MG | a:3325 | 3.78 | 1.0 | 23.2 |  | 26.73 | 10 | Water 6 | | a:8014 | 4.16 | 1.0 | 35.9 | 2.88 [G:OP2]a:2451 | 7.46 | 11 | Water 7 | | a:4008 | 4.19 | 1.0 | 29.3 |  | 13.55 | 12 | Water 8 | | a:7561 | 4.24 | 1.0 | 35.3 |  | 7.72 | 13 | A | O5' | a:2452 | 4.27 | 1.0 | 22.1 |  | 16.61 | 14 | G | O3' | a:2450 | 4.30 | 1.0 | 25.4 |  | 13.58 | 15 | **K** | K | a:3110 | 4.55 | 1.0 | 30.0 |  | 19.46 | 16 | G | OP2 | a:2451 | 4.59 | 1.0 | 27.2 |  | 14.53 | 17 | G | O3' | a:2451 | 4.62 | 1.0 | 22.5 |  | 17.27 |

  
**1^st^ shell coordination angles and distances (*trans* in "blue"; *cys* in "green"; deviations in "orange"):**   
• 1-2(OP1/W): 84; 1-3(OP1/O5'): 52; 1-4(OP1/W): 83; 1-5(OP1/W): 164; 1-6(OP1/W): 129; 1-7(OP1/W): 105; 2-3(W/O5'): 132; 2-4(W/W): 91; 2-5(W/W): 107; 2-6(W/W): 80; 2-7(W/W): 92; 3-4(O5'/W): 98; 3-5(O5'/W): 114; 3-6(O5'/W): 139; 3-7(O5'/W): 86; 4-5(W/W): 109; 4-6(W/W): 50; 4-7(W/W): 171; 5-6(W/W): 66; 5-7(W/W): 63; 6-7(W/W): 123;   
• 1-2(OP1/W): 3.7; 1-3(OP1/O5'): 2.4; 1-4(OP1/W): 3.8; 1-5(OP1/W): 5.7; 1-6(OP1/W): 5.3; 1-7(OP1/W): 4.7; 2-3(W/O5'): 5.2; 2-4(W/W): 4.1; 2-5(W/W): 4.6; 2-6(W/W): 3.8; 2-7(W/W): 4.3; 3-4(O5'/W): 4.4; 3-5(O5'/W): 4.9; 3-6(O5'/W): 5.6; 3-7(O5'/W): 4.1; 4-5(W/W): 4.8; 4-6(W/W): 2.6; 4-7(W/W): 6.1; 5-6(W/W): 3.3; 5-7(W/W): 3.2; 6-7(W/W): 5.5;   
  
**3.96 ± 1.16: the average coodination distance of the 1^st^ six ligands could correspond to an hexahydrated Mg^2+^.**  
**10: Number of potential Water contacts**   
**3 : Number of second shell contacts involving water or ions**   
**30%: Percentage of satisfied second shell Water contacts**   
• Contact(s): /O5'/O_w_/O_w_/O_w_/O_w_/O_w_/OP1/  
• Contact code: O_ph_O_r_5O_w_  
Go to ToC

### K:a:3112

|  | Res. type | Atom | Residue | Distance | Occupancy | B-factor | Closest atom(s) to a 1^st^ shell water | EM\_dens. |
| --- | --- | --- | --- | --- | --- | --- | --- | --- |
| **Ion** | | | |  | 1.0 | 30.0 |  | 25.24 |
 1 | Water 1 | | a:4826 | 2.61 | 1.0 | 26.5 | 2.66 [C:OP1]a:1969 // 2.67 [C:OP2]a:1971 | 10.99 | 2 | A | O2' | a:1970 | 2.94 | 1.0 | 30.6 |  | 11.91 | 3 | G | O2' | a:2596 | 2.97 | 1.0 | 26.0 |  | 14.02 | 4 | A | O3' | a:1970 | 3.04 | 1.0 | 30.4 |  | 10.24 | 5 | 5MU | O2' | a:1943 | 3.06 | 1.0 | 27.7 |  | 12.74 | 6 | C | O5' | a:1971 | 3.12 | 1.0 | 32.2 |  | 10.28 | 7 | 5MU | O2 | a:1943 | 3.28 | 1.0 | 29.9 |  | 13.18 | 8 | 5MU | O3' | a:1943 | 3.64 | 1.0 | 28.8 |  | 10.72 | 9 | C | OP2 | a:1971 | 3.80 | 1.0 | 30.2 |  | 9.16 | 10 | G | O4' | a:2596 | 4.12 | 1.0 | 29.0 |  | 13.34 | 11 | C | OP1 | a:1969 | 4.20 | 1.0 | 36.0 |  | 10.01 | 12 | G | O3' | a:2596 | 4.21 | 1.0 | 27.8 |  | 13.42 | 13 | U | OP1 | a:1944 | 4.28 | 1.0 | 31.4 |  | 9.64 | 14 | 5MU | N1 | a:1943 | 4.56 | 1.0 | 22.8 |  | 18.62 |

  
**1^st^ shell coordination angles and distances (*trans* in "blue"; *cys* in "green"; deviations in "orange"):**   
• 1-2(W/O2'): 72; 1-3(W/O2'): 153; 1-4(W/O3'): 77; 1-5(W/O2'): 74; 1-6(W/O5'): 81; 2-3(O2'/O2'): 81; 2-4(O2'/O3'): 54; 2-5(O2'/O2'): 119; 2-6(O2'/O5'): 100; 3-4(O2'/O3'): 88; 3-5(O2'/O2'): 120; 3-6(O2'/O5'): 105; 4-5(O3'/O2'): 151; 4-6(O3'/O5'): 47; 5-6(O2'/O5'): 122;   
• 1-2(W/O2'): 3.3; 1-3(W/O2'): 5.4; 1-4(W/O3'): 3.5; 1-5(W/O2'): 3.4; 1-6(W/O5'): 3.8; 2-3(O2'/O2'): 3.8; 2-4(O2'/O3'): 2.7; 2-5(O2'/O2'): 5.2; 2-6(O2'/O5'): 4.7; 3-4(O2'/O3'): 4.2; 3-5(O2'/O2'): 5.2; 3-6(O2'/O5'): 4.8; 4-5(O3'/O2'): 5.9; 4-6(O3'/O5'): 2.5; 5-6(O2'/O5'): 5.4;   
  
**4.11 ± 1.23: the average coodination distance of the 1^st^ six ligands could correspond to an hexahydrated Mg^2+^.**  
**2: Number of potential Water contacts**   
**2 : Number of second shell contacts involving water or ions**   
**100%: Percentage of satisfied second shell Water contacts**   
• Contact(s): /O2'/O2'/O2'/O3'/O5'/O_w_/  
• Contact code: 5O_r_O_w_  
Go to ToC

### K:a:3113

|  | Res. type | Atom | Residue | Distance | Occupancy | B-factor | Closest atom(s) to a 1^st^ shell water | EM\_dens. |
| --- | --- | --- | --- | --- | --- | --- | --- | --- |
| **Ion** | | | |  | 1.0 | 30.0 |  | 22.51 |
 1 | Water 1 | | a:9476 | 2.81 | 1.0 | 38.9 |  | 7.49 | 2 | Water 2 | | a:8808 | 2.88 | 1.0 | 42.5 | 2.97 [A:O3']a:2546 | 3.49 | 3 | A | N3 | a:2546 | 2.90 | 1.0 | 20.9 |  | 13.85 | 4 | G | N7 | a:2547 | 2.93 | 1.0 | 25.4 |  | 13.24 | 5 | G | O6 | a:2548 | 2.93 | 1.0 | 32.6 |  | 13.82 | 6 | A | O4' | a:2546 | 2.99 | 1.0 | 31.6 |  | 12.61 | 7 | G | N7 | a:2548 | 3.09 | 1.0 | 21.8 |  | 14.04 | 8 | G | O6 | a:2547 | 3.87 | 1.0 | 28.4 |  | 13.92 | 9 | Water 3 | | a:10040 | 4.08 | 1.0 | 44.3 |  | 3.92 | 10 | A | O3' | a:2546 | 4.49 | 1.0 | 29.5 |  | 12.15 |

  
**1^st^ shell coordination angles and distances (*trans* in "blue"; *cys* in "green"; deviations in "orange"):**   
• 1-2(W/W): 70; 1-3(W/N3): 78; 1-4(W/N7): 137; 1-5(W/O6): 70; 1-6(W/O4'): 132; 1-7(W/N7): 63; 2-3(W/N3): 104; 2-4(W/N7): 90; 2-5(W/O6): 126; 2-6(W/O4'): 83; 2-7(W/N7): 69; 3-4(N3/N7): 145; 3-5(N3/O6): 102; 3-6(N3/O4'): 71; 3-7(N3/N7): 141; 4-5(N7/O6): 96; 4-6(N7/O4'): 79; 4-7(N7/N7): 74; 5-6(O6/O4'): 151; 5-7(O6/N7): 61; 6-7(O4'/N7): 141;   
• 1-2(W/W): 3.2; 1-3(W/N3): 3.6; 1-4(W/N7): 5.3; 1-5(W/O6): 3.3; 1-6(W/O4'): 5.3; 1-7(W/N7): 3.1; 2-3(W/N3): 4.5; 2-4(W/N7): 4.1; 2-5(W/O6): 5.2; 2-6(W/O4'): 3.9; 2-7(W/N7): 3.4; 3-4(N3/N7): 5.6; 3-5(N3/O6): 4.5; 3-6(N3/O4'): 3.4; 3-7(N3/N7): 5.6; 4-5(N7/O6): 4.3; 4-6(N7/O4'): 3.8; 4-7(N7/N7): 3.6; 5-6(O6/O4'): 5.7; 5-7(O6/N7): 3.1; 6-7(O4'/N7): 5.7;   
  
**4.07 ± 1.23: the average coodination distance of the 1^st^ six ligands could correspond to an hexahydrated Mg^2+^.**  
**4: Number of potential Water contacts**   
**1 : Number of second shell contacts involving water or ions**   
**25%: Percentage of satisfied second shell Water contacts**   
• Contact(s): /N3_A_/N7/N7/O4'/O6/O_w_/O_w_/  
• Contact code: O_r_O_b_3N_b_2O_w_  
Go to ToC

### K:a:3114

|  | Res. type | Atom | Residue | Distance | Occupancy | B-factor | Closest atom(s) to a 1^st^ shell water | EM\_dens. |
| --- | --- | --- | --- | --- | --- | --- | --- | --- |
| **Ion** | | | |  | 1.0 | 30.0 |  | 39.62 |
 1 | Water 1 | | a:5209 | 2.78 | 1.0 | 29.8 |  | 14.62 | 2 | G | N7 | a:2057 | 2.87 | 1.0 | 16.3 |  | 19.59 | 3 | A | O2' | a:2055 | 2.97 | 1.0 | 29.7 |  | 16.01 | 4 | A | N1 | a:2618 | 2.99 | 1.0 | 19.9 |  | 17.09 | 5 | G | O6 | a:2057 | 3.12 | 1.0 | 22.5 |  | 17.72 | 6 | A | N3 | a:2055 | 3.36 | 1.0 | 20.7 |  | 18.30 | 7 | A | N6 | a:2056 | 3.39 | 1.0 | 14.7 |  | 16.34 | 8 | A | N7 | a:2056 | 3.43 | 1.0 | 17.6 |  | 16.85 | 9 | A | N1 | a:2581 | 4.51 | 1.0 | 20.2 |  | 17.51 | 10 | A | N7 | a:2581 | 4.54 | 1.0 | 20.8 |  | 15.40 | 11 | A | N1 | a:2056 | 4.59 | 1.0 | 17.0 |  | 18.19 |

  
**1^st^ shell coordination angles and distances (*trans* in "blue"; *cys* in "green"; deviations in "orange"):**   
• 1-2(W/N7): 83; 1-3(W/O2'): 58; 1-4(W/N1): 117; 1-5(W/O6): 139; 2-3(N7/O2'): 139; 2-4(N7/N1): 133; 2-5(N7/O6): 61; 3-4(O2'/N1): 82; 3-5(O2'/O6): 160; 4-5(N1/O6): 79;   
• 1-2(W/N7): 3.8; 1-3(W/O2'): 2.8; 1-4(W/N1): 4.9; 1-5(W/O6): 5.5; 2-3(N7/O2'): 5.5; 2-4(N7/N1): 5.4; 2-5(N7/O6): 3.1; 3-4(O2'/N1): 3.9; 3-5(O2'/O6): 6.0; 4-5(N1/O6): 3.9;   
  
**4.15 ± 1.23: the average coodination distance of the 1^st^ six ligands could correspond to an hexahydrated Mg^2+^.**  
**2: Number of potential Water contacts**   
**0 : Number of second shell contacts involving water or ions**   
**0%: Percentage of satisfied second shell Water contacts**   
• Contact(s): /N1_A_/N7/O2'/O6/O_w_/  
• Contact code: O_r_O_b_2N_b_O_w_  
→ Coordination < 6 for K ←  
Go to ToC

### K:a:3115

|  | Res. type | Atom | Residue | Distance | Occupancy | B-factor | Closest atom(s) to a 1^st^ shell water | EM\_dens. |
| --- | --- | --- | --- | --- | --- | --- | --- | --- |
| **Ion** | | | |  | 1.0 | 30.0 |  | 16.92 |
 1 | Water 1 | | a:8282 | 2.49 | 1.0 | 37.3 | 2.91 [G:N7]a:2441 | 8.13 | 2 | U | O4 | a:2249 | 2.68 | 1.0 | 27.8 |  | 13.52 | 3 | G | O6 | a:2440 | 2.76 | 1.0 | 23.9 |  | 16.41 | 4 | G | O6 | a:2441 | 2.93 | 1.0 | 26.7 |  | 14.71 | 5 | Water 2 | | a:9622 | 3.38 | 1.0 | 45.3 | 2.05 [MG:MG]a:3160 | 5.83 | 6 | G | N7 | a:2441 | 3.92 | 1.0 | 16.7 |  | 15.19 | 7 | G | N7 | a:2440 | 3.96 | 1.0 | 17.8 |  | 15.14 | 8 | Water 3 | | a:7609 | 4.04 | 1.0 | 39.3 | 2.22 [MG:MG]a:3160 | 4.31 | 9 | Water 4 | | a:6341 | 4.14 | 1.0 | 45.6 | 2.76 [U:O4]a:2442 | 4.09 | 10 | **MG** | MG | a:3160 | 4.31 | 1.0 | 49.6 |  | 7.86 | 11 | U | O4 | a:2248 | 4.67 | 1.0 | 26.0 |  | 15.43 | 12 | U | O4 | a:2442 | 4.70 | 1.0 | 25.7 |  | 14.92 |

  
**1^st^ shell coordination angles and distances (*trans* in "blue"; *cys* in "green"; deviations in "orange"):**   
• 1-2(W/O4): 90; 1-3(W/O6): 97; 1-4(W/O6): 92; 2-3(O4/O6): 99; 2-4(O4/O6): 174; 3-4(O6/O6): 76;   
• 1-2(W/O4): 3.7; 1-3(W/O6): 3.9; 1-4(W/O6): 3.9; 2-3(O4/O6): 4.1; 2-4(O4/O6): 5.6; 3-4(O6/O6): 3.5;   
  
**4.36 ± 1.43: the average coodination distance of the 1^st^ six ligands could correspond to an hexahydrated Mg^2+^.**  
**6: Number of potential Water contacts**   
**3 : Number of second shell contacts involving water or ions**   
**50%: Percentage of satisfied second shell Water contacts**   
• Contact(s): /O4/O6/O6/  
• Contact code: 3O_b_  
• Contact(s) in the exclusion zone: /O_w_/  
→ Coordination < 6 for K ←  
Go to ToC

### K:a:3116

|  | Res. type | Atom | Residue | Distance | Occupancy | B-factor | Closest atom(s) to a 1^st^ shell water | EM\_dens. |
| --- | --- | --- | --- | --- | --- | --- | --- | --- |
| **Ion** | | | |  | 1.0 | 30.0 |  | 31.60 |
 1 | G | O6 | a:2451 | 2.70 | 1.0 | 25.7 |  | 15.19 | 2 | C | O4' | a:2505 | 2.79 | 1.0 | 25.5 |  | 12.59 | 3 | G | O6 | a:2065 | 2.86 | 1.0 | 33.7 |  | 13.05 | 4 | G | N7 | a:2451 | 2.89 | 1.0 | 19.7 |  | 14.42 | 5 | Water 1 | | a:8023 | 3.03 | 1.0 | 34.7 | 2.88 [PSU:N1]a:2508 | 11.70 | 6 | C | O2 | a:2505 | 3.16 | 1.0 | 25.9 |  | 15.16 | 7 | PSU | O2 | a:2508 | 3.23 | 1.0 | 26.2 |  | 10.05 | 8 | G | N7 | a:2065 | 3.28 | 1.0 | 27.5 |  | 13.22 | 9 | C | N1 | a:2505 | 3.86 | 1.0 | 19.9 |  | 16.26 | 10 | C | N3 | a:2505 | 4.55 | 1.0 | 17.0 |  | 18.07 | 11 | Water 2 | | a:5100 | 4.65 | 1.0 | 38.3 |  | 6.39 | 12 | C | O5' | a:2505 | 4.68 | 1.0 | 23.9 |  | 14.19 |

  
**1^st^ shell coordination angles and distances (*trans* in "blue"; *cys* in "green"; deviations in "orange"):**   
• 1-2(O6/O4'): 122; 1-3(O6/O6): 93; 1-4(O6/N7): 66; 1-5(O6/W): 136; 1-6(O6/O2): 81; 2-3(O4'/O6): 126; 2-4(O4'/N7): 67; 2-5(O4'/W): 91; 2-6(O4'/O2): 67; 3-4(O6/N7): 159; 3-5(O6/W): 88; 3-6(O6/O2): 82; 4-5(N7/W): 110; 4-6(N7/O2): 91; 5-6(W/O2): 142;   
• 1-2(O6/O4'): 4.8; 1-3(O6/O6): 4.0; 1-4(O6/N7): 3.1; 1-5(O6/W): 5.3; 1-6(O6/O2): 3.8; 2-3(O4'/O6): 5.0; 2-4(O4'/N7): 3.1; 2-5(O4'/W): 4.1; 2-6(O4'/O2): 3.3; 3-4(O6/N7): 5.7; 3-5(O6/W): 4.1; 3-6(O6/O2): 3.9; 4-5(N7/W): 4.8; 4-6(N7/O2): 4.3; 5-6(W/O2): 5.8;   
  
**3.99 ± 1.18: the average coodination distance of the 1^st^ six ligands could correspond to an hexahydrated Mg^2+^.**  
**2: Number of potential Water contacts**   
**1 : Number of second shell contacts involving water or ions**   
**50%: Percentage of satisfied second shell Water contacts**   
• Contact(s): /N7/O2/O4'/O6/O6/O_w_/  
• Contact code: O_r_3O_b_N_b_O_w_  
Go to ToC

### K:a:3117

|  | Res. type | Atom | Residue | Distance | Occupancy | B-factor | Closest atom(s) to a 1^st^ shell water | EM\_dens. |
| --- | --- | --- | --- | --- | --- | --- | --- | --- |
| **Ion** | | | |  | 1.0 | 30.0 |  | 13.58 |
 1 | C | OP2 | a:2444 | 2.80 | 1.0 | 32.2 |  | 9.86 | 2 | A | OP2 | a:2604 | 2.80 | 1.0 | 28.4 |  | 12.65 | 3 | A | O3' | a:2443 | 2.92 | 1.0 | 31.0 |  | 11.77 | 4 | Water 1 | | a:3786 | 2.99 | 1.0 | 30.0 | 1.90 [MG:MG]a:3305 | 10.91 | 5 | A | O2' | a:2443 | 3.05 | 1.0 | 32.6 |  | 11.21 | 6 | Water 2 | | c:478 | 3.29 | 1.0 | 26.2 | 2.20 [MG:MG]a:3305 | 13.14 | 7 | GLU | OE2 | c:236 | 3.49 | 1.0 | 43.8 |  | 6.80 | 8 | GLU | OE1 | c:236 | 3.66 | 1.0 | 46.1 |  | 4.35 | 9 | A | OP1 | a:2604 | 3.85 | 1.0 | 30.9 |  | 10.40 | 10 | **MG** | MG | a:3305 | 3.96 | 1.0 | 26.2 |  | 23.23 | 11 | Water 3 | | a:4168 | 3.99 | 1.0 | 26.9 |  | 10.84 | 12 | Water 4 | | a:4231 | 3.99 | 1.0 | 46.2 | 2.86 [GLU:OE1]c:236 | 4.02 | 13 | Water 5 | | a:3538 | 4.01 | 1.0 | 25.5 |  | 14.16 | 14 | C | OP1 | a:2444 | 4.10 | 1.0 | 30.8 |  | 12.41 | 15 | G | O3' | a:2603 | 4.34 | 1.0 | 28.5 |  | 13.38 | 16 | Water 6 | | c:407 | 4.34 | 1.0 | 31.6 | 2.38 [MG:MG]a:3305 | 10.43 | 17 | Water 7 | | c:426 | 4.47 | 1.0 | 30.9 | 2.58 [GLU:OE2]c:236 | 11.97 | 18 | C | O5' | a:2444 | 4.69 | 1.0 | 27.7 |  | 13.02 | 19 | G | OP2 | a:2592 | 4.77 | 1.0 | 22.7 |  | 13.56 |

  
**1^st^ shell coordination angles and distances (*trans* in "blue"; *cys* in "green"; deviations in "orange"):**   
• 1-2(OP2/OP2): 126; 1-3(OP2/O3'): 52; 1-4(OP2/W): 113; 1-5(OP2/O2'): 105; 2-3(OP2/O3'): 116; 2-4(OP2/W): 121; 2-5(OP2/O2'): 76; 3-4(O3'/W): 94; 3-5(O3'/O2'): 54; 4-5(W/O2'): 85;   
• 1-2(OP2/OP2): 5.0; 1-3(OP2/O3'): 2.5; 1-4(OP2/W): 4.8; 1-5(OP2/O2'): 4.6; 2-3(OP2/O3'): 4.9; 2-4(OP2/W): 5.0; 2-5(OP2/O2'): 3.6; 3-4(O3'/W): 4.3; 3-5(O3'/O2'): 2.7; 4-5(W/O2'): 4.1;   
  
**4.17 ± 1.25: the average coodination distance of the 1^st^ six ligands could correspond to an hexahydrated Mg^2+^.**  
**4: Number of potential Water contacts**   
**2 : Number of second shell contacts involving water or ions**   
**50%: Percentage of satisfied second shell Water contacts**   
• Contact(s): /O2'/O3'/O_w_/OP2/OP2/  
• Contact code: 2O_ph_2O_r_O_w_  
→ Coordination < 6 for K ←  
Go to ToC

### K:a:3118

|  | Res. type | Atom | Residue | Distance | Occupancy | B-factor | Closest atom(s) to a 1^st^ shell water | EM\_dens. |
| --- | --- | --- | --- | --- | --- | --- | --- | --- |
| **Ion** | | | |  | 1.0 | 30.0 |  | 24.34 |
 1 | G | O2' | a:2254 | 2.69 | 1.0 | 31.6 |  | 12.18 | 2 | A | OP2 | a:2501 | 2.74 | 1.0 | 24.5 |  | 14.20 | 3 | OMG | OP2 | a:2255 | 2.88 | 1.0 | 37.5 |  | 7.22 | 4 | A | N7 | a:2501 | 2.89 | 1.0 | 24.0 |  | 13.19 | 5 | G | O3' | a:2254 | 3.41 | 1.0 | 29.9 |  | 11.30 | 6 | Water 1 | | a:6867 | 3.50 | 1.0 | 33.0 |  | 12.46 | 7 | C | OP2 | a:2500 | 3.52 | 1.0 | 28.3 |  | 13.97 | 8 | Water 2 | | a:5402 | 3.53 | 1.0 | 42.6 |  | 6.38 | 9 | C | O5' | a:2500 | 3.53 | 1.0 | 26.6 |  | 13.64 | 10 | C | OP1 | a:2500 | 3.99 | 1.0 | 31.1 |  | 12.72 | 11 | Water 3 | | a:7064 | 4.04 | 1.0 | 32.0 |  | 12.65 | 12 | OMG | OP1 | a:2255 | 4.51 | 1.0 | 29.8 |  | 11.39 | 13 | C | O3' | a:2500 | 4.56 | 1.0 | 24.9 |  | 13.48 | 14 | A | O5' | a:2501 | 4.64 | 1.0 | 29.7 |  | 12.67 |

  
**1^st^ shell coordination angles and distances (*trans* in "blue"; *cys* in "green"; deviations in "orange"):**   
• 1-2(O2'/OP2): 84; 1-3(O2'/OP2): 93; 1-4(O2'/N7): 165; 2-3(OP2/OP2): 119; 2-4(OP2/N7): 89; 3-4(OP2/N7): 102;   
• 1-2(O2'/OP2): 3.6; 1-3(O2'/OP2): 4.0; 1-4(O2'/N7): 5.5; 2-3(OP2/OP2): 4.9; 2-4(OP2/N7): 3.9; 3-4(OP2/N7): 4.5;   
  
**4.19 ± 1.28: the average coodination distance of the 1^st^ six ligands could correspond to an hexahydrated Mg^2+^.**  
**4: Number of potential Water contacts**   
**0 : Number of second shell contacts involving water or ions**   
**0%: Percentage of satisfied second shell Water contacts**   
• Contact(s): /N7/O2'/OP2/OP2/  
• Contact code: 2O_ph_O_r_N_b_  
→ Coordination < 6 for K ←  
Go to ToC

### K:a:3119

|  | Res. type | Atom | Residue | Distance | Occupancy | B-factor | Closest atom(s) to a 1^st^ shell water | EM\_dens. |
| --- | --- | --- | --- | --- | --- | --- | --- | --- |
| **Ion** | | | |  | 1.0 | 30.0 |  | 25.79 |
 1 | Water 1 | | a:8476 | 2.76 | 1.0 | 30.6 |  | 12.02 | 2 | C | OP1 | a:2431 | 2.83 | 1.0 | 29.3 |  | 12.16 | 3 | Water 2 | | a:3814 | 2.93 | 1.0 | 24.3 | 2.22 [MG:MG]a:3329 | 18.96 | 4 | A | O3' | a:2430 | 3.08 | 1.0 | 32.9 |  | 12.62 | 5 | U | OP2 | a:829 | 3.08 | 1.0 | 25.2 |  | 14.42 | 6 | G | OP1 | a:2433 | 3.09 | 1.0 | 25.2 |  | 12.03 | 7 | Water 3 | | a:9068 | 3.11 | 1.0 | 33.3 |  | 9.58 | 8 | Water 4 | | a:7520 | 3.17 | 1.0 | 32.3 | 2.36 [MG:MG]a:3230 | 10.85 | 9 | G | OP2 | a:2433 | 3.38 | 1.0 | 24.6 |  | 11.90 | 10 | A | O2' | a:2430 | 3.63 | 1.0 | 28.9 |  | 13.68 | 11 | U | OP1 | a:829 | 3.64 | 1.0 | 26.1 |  | 13.52 | 12 | **MG** | MG | a:3329 | 3.70 | 1.0 | 25.3 |  | 23.10 | 13 | **MG** | MG | a:3230 | 4.11 | 1.0 | 28.0 |  | 19.09 | 14 | G | O5' | a:2433 | 4.51 | 1.0 | 23.0 |  | 13.75 | 15 | C | O5' | a:2431 | 4.56 | 1.0 | 22.2 |  | 15.84 | 16 | Water 5 | | a:9485 | 4.59 | 1.0 | 29.4 |  | 9.65 | 17 | Water 6 | | a:7941 | 4.64 | 1.0 | 37.1 | 2.88 [C:OP2]a:2431 | 6.40 | 18 | C | OP2 | a:2431 | 4.66 | 1.0 | 29.5 |  | 12.57 |

  
**1^st^ shell coordination angles and distances (*trans* in "blue"; *cys* in "green"; deviations in "orange"):**   
• 1-2(W/OP1): 89; 1-3(W/W): 83; 1-4(W/O3'): 73; 1-5(W/OP2): 80; 1-6(W/OP1): 139; 1-7(W/W): 70; 1-8(W/W): 118; 2-3(OP1/W): 60; 2-4(OP1/O3'): 49; 2-5(OP1/OP2): 110; 2-6(OP1/OP1): 59; 2-7(OP1/W): 153; 2-8(OP1/W): 149; 3-4(W/O3'): 104; 3-5(W/OP2): 50; 3-6(W/OP1): 60; 3-7(W/W): 131; 3-8(W/W): 133; 4-5(O3'/OP2): 145; 4-6(O3'/OP1): 99; 4-7(O3'/W): 106; 4-8(O3'/W): 121; 5-6(OP2/OP1): 88; 5-7(OP2/W): 84; 5-8(OP2/W): 91; 6-7(OP1/W): 147; 6-8(OP1/W): 100; 7-8(W/W): 48;   
• 1-2(W/OP1): 3.9; 1-3(W/W): 3.8; 1-4(W/O3'): 3.5; 1-5(W/OP2): 3.7; 1-6(W/OP1): 5.5; 1-7(W/W): 3.4; 1-8(W/W): 5.1; 2-3(OP1/W): 2.9; 2-4(OP1/O3'): 2.5; 2-5(OP1/OP2): 4.8; 2-6(OP1/OP1): 2.9; 2-7(OP1/W): 5.8; 2-8(OP1/W): 5.8; 3-4(W/O3'): 4.8; 3-5(W/OP2): 2.5; 3-6(W/OP1): 3.0; 3-7(W/W): 5.5; 3-8(W/W): 5.6; 4-5(O3'/OP2): 5.9; 4-6(O3'/OP1): 4.7; 4-7(O3'/W): 4.9; 4-8(O3'/W): 5.4; 5-6(OP2/OP1): 4.3; 5-7(OP2/W): 4.1; 5-8(OP2/W): 4.5; 6-7(OP1/W): 5.9; 6-8(OP1/W): 4.8; 7-8(W/W): 2.6;   
  
**4.01 ± 1.17: the average coodination distance of the 1^st^ six ligands could correspond to an hexahydrated Mg^2+^.**  
**8: Number of potential Water contacts**   
**2 : Number of second shell contacts involving water or ions**   
**25%: Percentage of satisfied second shell Water contacts**   
• Contact(s): /O3'/O_w_/O_w_/O_w_/O_w_/OP1/OP1/OP2/  
• Contact code: 3O_ph_O_r_4O_w_  
Go to ToC

### K:a:3120

|  | Res. type | Atom | Residue | Distance | Occupancy | B-factor | Closest atom(s) to a 1^st^ shell water | EM\_dens. |
| --- | --- | --- | --- | --- | --- | --- | --- | --- |
| **Ion** | | | |  | 1.0 | 30.0 |  | 21.95 |
 1 | A | OP2 | a:1787 | 2.72 | 1.0 | 23.8 |  | 13.28 | 2 | U | OP2 | a:1986 | 2.90 | 1.0 | 44.1 |  | 5.80 | 3 | Water 1 | | a:4388 | 2.91 | 1.0 | 32.2 |  | 11.03 | 4 | U | O4' | a:1986 | 3.12 | 1.0 | 34.8 |  | 9.36 | 5 | A | OP1 | a:1985 | 3.15 | 1.0 | 36.5 |  | 8.94 | 6 | G | O3' | a:1984 | 3.16 | 1.0 | 35.2 |  | 9.38 | 7 | Water 2 | | a:9981 | 3.23 | 1.0 | 28.6 |  | 10.03 | 8 | A | O5' | a:1985 | 3.62 | 1.0 | 32.2 |  | 7.80 | 9 | G | O2' | a:1984 | 3.93 | 1.0 | 33.5 |  | 8.07 | 10 | Water 3 | | a:9920 | 3.98 | 1.0 | 27.7 |  | 12.70 | 11 | A | O5' | a:1787 | 4.20 | 1.0 | 25.0 |  | 14.32 | 12 | U | O5' | a:1986 | 4.26 | 1.0 | 35.0 |  | 9.94 | 13 | Water 4 | | a:6552 | 4.38 | 1.0 | 24.1 |  | 12.63 | 14 | **K** | K | a:3121 | 4.56 | 1.0 | 30.0 |  | 24.38 | 15 | U | N1 | a:1986 | 4.79 | 1.0 | 25.4 |  | 13.27 |

  
**1^st^ shell coordination angles and distances (*trans* in "blue"; *cys* in "green"; deviations in "orange"):**   
• 1-2(OP2/OP2): 101; 1-3(OP2/W): 135; 1-4(OP2/O4'): 87; 1-5(OP2/OP1): 92; 1-6(OP2/O3'): 132; 2-3(OP2/W): 108; 2-4(OP2/O4'): 71; 2-5(OP2/OP1): 96; 2-6(OP2/O3'): 67; 3-4(W/O4'): 71; 3-5(W/OP1): 118; 3-6(W/O3'): 92; 4-5(O4'/OP1): 166; 4-6(O4'/O3'): 127; 5-6(OP1/O3'): 46;   
• 1-2(OP2/OP2): 4.3; 1-3(OP2/W): 5.2; 1-4(OP2/O4'): 4.0; 1-5(OP2/OP1): 4.2; 1-6(OP2/O3'): 5.4; 2-3(OP2/W): 4.7; 2-4(OP2/O4'): 3.5; 2-5(OP2/OP1): 4.5; 2-6(OP2/O3'): 3.4; 3-4(W/O4'): 3.5; 3-5(W/OP1): 5.2; 3-6(W/O3'): 4.4; 4-5(O4'/OP1): 6.2; 4-6(O4'/O3'): 5.6; 5-6(OP1/O3'): 2.5;   
  
**4.14 ± 1.23: the average coodination distance of the 1^st^ six ligands could correspond to an hexahydrated Mg^2+^.**  
**4: Number of potential Water contacts**   
**0 : Number of second shell contacts involving water or ions**   
**0%: Percentage of satisfied second shell Water contacts**   
• Contact(s): /O3'/O4'/O_w_/OP1/OP2/OP2/  
• Contact code: 3O_ph_2O_r_O_w_  
Go to ToC

### K:a:3121

|  | Res. type | Atom | Residue | Distance | Occupancy | B-factor | Closest atom(s) to a 1^st^ shell water | EM\_dens. |
| --- | --- | --- | --- | --- | --- | --- | --- | --- |
| **Ion** | | | |  | 1.0 | 30.0 |  | 24.38 |
 1 | A | O3' | a:1786 | 2.73 | 1.0 | 23.7 |  | 15.62 | 2 | A | O2' | a:1786 | 2.78 | 1.0 | 27.9 |  | 16.67 | 3 | A | N7 | a:1985 | 2.86 | 1.0 | 24.5 |  | 12.48 | 4 | Water 1 | | a:5704 | 2.91 | 1.0 | 40.8 |  | 9.95 | 5 | A | OP2 | a:1787 | 3.05 | 1.0 | 23.8 |  | 13.28 | 6 | Water 2 | | a:4284 | 3.15 | 1.0 | 36.0 |  | 9.16 | 7 | Water 3 | | a:9537 | 3.24 | 1.0 | 35.5 |  | 6.68 | 8 | A | OP1 | a:1985 | 4.01 | 1.0 | 36.5 |  | 8.94 | 9 | Water 4 | | a:9981 | 4.16 | 1.0 | 28.6 |  | 10.03 | 10 | **K** | K | a:3120 | 4.56 | 1.0 | 30.0 |  | 21.95 | 11 | A | O5' | a:1787 | 4.58 | 1.0 | 25.0 |  | 14.32 | 12 | A | OP1 | a:1787 | 4.58 | 1.0 | 25.7 |  | 15.14 | 13 | A | O4' | a:1786 | 4.68 | 1.0 | 23.6 |  | 16.93 |

  
**1^st^ shell coordination angles and distances (*trans* in "blue"; *cys* in "green"; deviations in "orange"):**   
• 1-2(O3'/O2'): 59; 1-3(O3'/N7): 99; 1-4(O3'/W): 114; 1-5(O3'/OP2): 50; 1-6(O3'/W): 78; 2-3(O2'/N7): 88; 2-4(O2'/W): 57; 2-5(O2'/OP2): 104; 2-6(O2'/W): 95; 3-4(N7/W): 94; 3-5(N7/OP2): 81; 3-6(N7/W): 173; 4-5(W/OP2): 161; 4-6(W/W): 93; 5-6(OP2/W): 92;   
• 1-2(O3'/O2'): 2.7; 1-3(O3'/N7): 4.2; 1-4(O3'/W): 4.7; 1-5(O3'/OP2): 2.5; 1-6(O3'/W): 3.7; 2-3(O2'/N7): 3.9; 2-4(O2'/W): 2.7; 2-5(O2'/OP2): 4.6; 2-6(O2'/W): 4.4; 3-4(N7/W): 4.2; 3-5(N7/OP2): 3.8; 3-6(N7/W): 6.0; 4-5(W/OP2): 5.9; 4-6(W/W): 4.4; 5-6(OP2/W): 4.5;   
  
**4.12 ± 1.27: the average coodination distance of the 1^st^ six ligands could correspond to an hexahydrated Mg^2+^.**  
**6: Number of potential Water contacts**   
**0 : Number of second shell contacts involving water or ions**   
**0%: Percentage of satisfied second shell Water contacts**   
• Contact(s): /N7/O2'/O3'/O_w_/O_w_/OP2/  
• Contact code: O_ph_2O_r_N_b_2O_w_  
Go to ToC

### K:a:3122

|  | Res. type | Atom | Residue | Distance | Occupancy | B-factor | Closest atom(s) to a 1^st^ shell water | EM\_dens. |
| --- | --- | --- | --- | --- | --- | --- | --- | --- |
| **Ion** | | | |  | 1.0 | 30.0 |  | 25.71 |
 1 | Water 1 | | a:5702 | 2.53 | 1.0 | 30.9 | 2.83 [U:O2]a:1780 | 10.97 | 2 | A | OP1 | a:785 | 2.80 | 1.0 | 25.1 |  | 15.58 | 3 | A | N3 | a:1789 | 2.83 | 1.0 | 15.4 |  | 18.55 | 4 | Water 2 | | a:4288 | 2.93 | 1.0 | 27.0 |  | 16.01 | 5 | U | O2 | a:1780 | 3.49 | 1.0 | 30.9 |  | 14.90 | 6 | C | O4' | a:1790 | 3.49 | 1.0 | 26.8 |  | 17.81 | 7 | A | N6 | a:1787 | 3.59 | 1.0 | 16.6 |  | 18.36 | 8 | A | O5' | a:785 | 3.80 | 1.0 | 23.1 |  | 16.93 | 9 | A | O2' | a:1789 | 4.07 | 1.0 | 25.0 |  | 17.59 | 10 | C | N1 | a:1790 | 4.34 | 1.0 | 15.7 |  | 18.99 | 11 | A | O3' | a:784 | 4.41 | 1.0 | 18.9 |  | 18.98 | 12 | A | O5' | a:784 | 4.53 | 1.0 | 28.5 |  | 13.02 | 13 | A | N1 | a:1789 | 4.60 | 1.0 | 19.1 |  | 15.15 | 14 | U | N1 | a:1780 | 4.80 | 1.0 | 16.9 |  | 14.77 |

  
**1^st^ shell coordination angles and distances (*trans* in "blue"; *cys* in "green"; deviations in "orange"):**   
• 1-2(W/OP1): 98; 1-3(W/N3): 127; 1-4(W/W): 96; 2-3(OP1/N3): 112; 2-4(OP1/W): 133; 3-4(N3/W): 94;   
• 1-2(W/OP1): 4.0; 1-3(W/N3): 4.8; 1-4(W/W): 4.1; 2-3(OP1/N3): 4.7; 2-4(OP1/W): 5.3; 3-4(N3/W): 4.2;   
  
**4.24 ± 1.32: the average coodination distance of the 1^st^ six ligands could correspond to an hexahydrated Mg^2+^.**  
**4: Number of potential Water contacts**   
**1 : Number of second shell contacts involving water or ions**   
**25%: Percentage of satisfied second shell Water contacts**   
• Contact(s): /N3_A_/O_w_/OP1/  
• Contact code: O_ph_N_b_O_w_  
• Contact(s) in the exclusion zone: /O_w_/  
→ Coordination < 6 for K ←  
Go to ToC

### K:a:3123

|  | Res. type | Atom | Residue | Distance | Occupancy | B-factor | Closest atom(s) to a 1^st^ shell water | EM\_dens. |
| --- | --- | --- | --- | --- | --- | --- | --- | --- |
| **Ion** | | | |  | 1.0 | 30.0 |  | 31.82 |
 1 | Water 1 | | a:6008 | 2.78 | 1.0 | 33.0 | 2.74 [G:O6]a:2060 | 12.79 | 2 | Water 2 | | a:6426 | 2.78 | 1.0 | 29.3 | 2.77 [A:OP2]a:2058 | 10.02 | 3 | Water 3 | | a:10270 | 2.84 | 1.0 | 47.5 |  | 4.97 | 4 | Water 4 | | a:6753 | 2.96 | 1.0 | 21.9 | 2.79 [A:OP2]a:2058 | 17.07 | 5 | Water 5 | | a:4523 | 3.01 | 1.0 | 22.2 | 2.03 [MG:MG]a:3194 | 17.29 | 6 | A | N7 | a:2581 | 3.11 | 1.0 | 20.8 |  | 15.40 | 7 | Water 6 | | a:7727 | 3.14 | 1.0 | 23.9 | 2.23 [MG:MG]a:3194 | 14.24 | 8 | G | N7 | a:2060 | 3.21 | 1.0 | 20.1 |  | 16.11 | 9 | G | OP2 | a:2057 | 4.07 | 1.0 | 24.9 |  | 15.70 | 10 | **MG** | MG | a:3194 | 4.14 | 1.0 | 27.2 |  | 28.04 | 11 | Water 7 | | a:9602 | 4.42 | 1.0 | 41.1 |  | 6.78 | 12 | A | OP1 | a:2581 | 4.47 | 1.0 | 24.1 |  | 13.82 | 13 | A | OP2 | a:2058 | 4.48 | 1.0 | 23.6 |  | 15.69 | 14 | Water 8 | | a:8194 | 4.51 | 1.0 | 40.5 | 2.90 [G:OP1]a:2057 | 6.34 | 15 | Water 9 | | a:4611 | 4.52 | 1.0 | 24.8 | 2.05 [MG:MG]a:3194 | 15.56 | 16 | G | OP1 | a:2057 | 4.62 | 1.0 | 20.6 |  | 15.05 | 17 | G | O6 | a:2060 | 4.66 | 1.0 | 27.8 |  | 14.10 | 18 | Water 10 | | a:5209 | 4.80 | 1.0 | 29.8 | 2.69 [G:OP2]a:2057 | 14.62 |

  
**1^st^ shell coordination angles and distances (*trans* in "blue"; *cys* in "green"; deviations in "orange"):**   
• 1-2(W/W): 120; 1-3(W/W): 81; 1-4(W/W): 154; 1-5(W/W): 97; 1-6(W/N7): 83; 1-7(W/W): 64; 2-3(W/W): 68; 2-4(W/W): 73; 2-5(W/W): 136; 2-6(W/N7): 127; 2-7(W/W): 117; 3-4(W/W): 125; 3-5(W/W): 100; 3-6(W/N7): 162; 3-7(W/W): 50; 4-5(W/W): 83; 4-6(W/N7): 72; 4-7(W/W): 134; 5-6(W/N7): 74; 5-7(W/W): 59; 6-7(N7/W): 116;   
• 1-2(W/W): 4.8; 1-3(W/W): 3.7; 1-4(W/W): 5.6; 1-5(W/W): 4.4; 1-6(W/N7): 3.9; 1-7(W/W): 3.2; 2-3(W/W): 3.1; 2-4(W/W): 3.4; 2-5(W/W): 5.4; 2-6(W/N7): 5.3; 2-7(W/W): 5.0; 3-4(W/W): 5.1; 3-5(W/W): 4.5; 3-6(W/N7): 5.9; 3-7(W/W): 2.5; 4-5(W/W): 3.9; 4-6(W/N7): 3.6; 4-7(W/W): 5.6; 5-6(W/N7): 3.7; 5-7(W/W): 3.0; 6-7(N7/W): 5.3;   
  
**3.97 ± 1.16: the average coodination distance of the 1^st^ six ligands could correspond to an hexahydrated Mg^2+^.**  
**12: Number of potential Water contacts**   
**5 : Number of second shell contacts involving water or ions**   
**42%: Percentage of satisfied second shell Water contacts**   
• Contact(s): /N7/O_w_/O_w_/O_w_/O_w_/O_w_/O_w_/  
• Contact code: N_b_6O_w_  
Go to ToC

### K:a:3124

|  | Res. type | Atom | Residue | Distance | Occupancy | B-factor | Closest atom(s) to a 1^st^ shell water | EM\_dens. |
| --- | --- | --- | --- | --- | --- | --- | --- | --- |
| **Ion** | | | |  | 1.0 | 30.0 |  | 21.05 |
 1 | G | O6 | a:651 | 2.73 | 1.0 | 32.0 |  | 11.97 | 2 | Water 1 | | a:7167 | 2.86 | 1.0 | 36.8 | 2.82 [G:N7]a:651 | 7.82 | 3 | G | O6 | a:640 | 2.90 | 1.0 | 32.3 |  | 9.20 | 4 | G | O6 | a:650 | 3.00 | 1.0 | 35.9 |  | 9.64 | 5 | U | O4 | a:641 | 3.10 | 1.0 | 34.3 |  | 10.24 | 6 | C | N4 | a:652 | 3.71 | 1.0 | 23.9 |  | 11.87 | 7 | U | N3 | a:641 | 3.95 | 1.0 | 27.7 |  | 10.53 | 8 | G | N7 | a:651 | 4.09 | 1.0 | 26.6 |  | 9.87 | 9 | G | N7 | a:650 | 4.49 | 1.0 | 30.6 |  | 9.47 |

  
**1^st^ shell coordination angles and distances (*trans* in "blue"; *cys* in "green"; deviations in "orange"):**   
• 1-2(O6/W): 92; 1-3(O6/O6): 71; 1-4(O6/O6): 75; 1-5(O6/O4): 80; 2-3(W/O6): 113; 2-4(W/O6): 85; 2-5(W/O4): 168; 3-4(O6/O6): 142; 3-5(O6/O4): 72; 4-5(O6/O4): 85;   
• 1-2(O6/W): 4.0; 1-3(O6/O6): 3.3; 1-4(O6/O6): 3.5; 1-5(O6/O4): 3.7; 2-3(W/O6): 4.8; 2-4(W/O6): 3.9; 2-5(W/O4): 5.9; 3-4(O6/O6): 5.6; 3-5(O6/O4): 3.5; 4-5(O6/O4): 4.1;   
  
**4.39 ± 1.39: the average coodination distance of the 1^st^ six ligands could correspond to an hexahydrated Mg^2+^.**  
**2: Number of potential Water contacts**   
**1 : Number of second shell contacts involving water or ions**   
**50%: Percentage of satisfied second shell Water contacts**   
• Contact(s): /O4/O6/O6/O6/O_w_/  
• Contact code: 4O_b_O_w_  
→ Coordination < 6 for K ←  
Go to ToC

### K:a:3125

|  | Res. type | Atom | Residue | Distance | Occupancy | B-factor | Closest atom(s) to a 1^st^ shell water | EM\_dens. |
| --- | --- | --- | --- | --- | --- | --- | --- | --- |
| **Ion** | | | |  | 1.0 | 30.0 |  | 23.85 |
 1 | G | O6 | a:952 | 2.72 | 1.0 | 31.6 |  | 11.59 | 2 | U | O4 | a:969 | 2.85 | 1.0 | 31.4 |  | 12.93 | 3 | G | O6 | a:968 | 2.85 | 1.0 | 30.9 |  | 13.63 | 4 | G | O6 | a:951 | 2.88 | 1.0 | 29.6 |  | 12.68 | 5 | Water 1 | | a:7383 | 3.10 | 1.0 | 35.0 | 2.83 [G:N7]a:952 | 7.76 | 6 | Water 2 | | a:9950 | 3.66 | 1.0 | 45.4 |  | 4.57 | 7 | U | N3 | a:969 | 3.82 | 1.0 | 17.6 |  | 13.44 | 8 | C | N4 | a:970 | 3.83 | 1.0 | 20.4 |  | 12.65 | 9 | Water 3 | | a:7122 | 3.93 | 1.0 | 35.5 |  | 8.86 | 10 | Water 4 | | a:7661 | 4.03 | 1.0 | 28.2 |  | 12.55 | 11 | G | N7 | a:952 | 4.38 | 1.0 | 23.9 |  | 11.55 |

  
**1^st^ shell coordination angles and distances (*trans* in "blue"; *cys* in "green"; deviations in "orange"):**   
• 1-2(O6/O4): 84; 1-3(O6/O6): 75; 1-4(O6/O6): 73; 1-5(O6/W): 82; 2-3(O4/O6): 68; 2-4(O4/O6): 104; 2-5(O4/W): 166; 3-4(O6/O6): 147; 3-5(O6/W): 106; 4-5(O6/W): 73;   
• 1-2(O6/O4): 3.7; 1-3(O6/O6): 3.4; 1-4(O6/O6): 3.3; 1-5(O6/W): 3.8; 2-3(O4/O6): 3.2; 2-4(O4/O6): 4.5; 2-5(O4/W): 5.9; 3-4(O6/O6): 5.5; 3-5(O6/W): 4.8; 4-5(O6/W): 3.6;   
  
**4.28 ± 1.34: the average coodination distance of the 1^st^ six ligands could correspond to an hexahydrated Mg^2+^.**  
**4: Number of potential Water contacts**   
**1 : Number of second shell contacts involving water or ions**   
**25%: Percentage of satisfied second shell Water contacts**   
• Contact(s): /O4/O6/O6/O6/O_w_/  
• Contact code: 4O_b_O_w_  
→ Coordination < 6 for K ←  
Go to ToC

### K:a:3126

|  | Res. type | Atom | Residue | Distance | Occupancy | B-factor | Closest atom(s) to a 1^st^ shell water | EM\_dens. |
| --- | --- | --- | --- | --- | --- | --- | --- | --- |
| **Ion** | | | |  | 1.0 | 30.0 |  | 36.37 |
 1 | THR | O | c:219 | 2.34 | 1.0 | 26.2 |  | 20.61 | 2 | C | OP2 | a:1792 | 2.49 | 1.0 | 24.4 |  | 18.15 | 3 | A | OP1 | a:1791 | 2.71 | 1.0 | 24.7 |  | 15.72 | 4 | Water 1 | | a:4366 | 2.75 | 1.0 | 24.0 | 2.62 [U:O4]a:1829 | 16.86 | 5 | G | O6 | a:1830 | 2.76 | 1.0 | 24.5 |  | 18.57 | 6 | A | O5' | a:1791 | 3.33 | 1.0 | 19.0 |  | 16.19 | 7 | Water 2 | | a:9209 | 3.44 | 1.0 | 28.7 |  | 11.96 | 8 | G | N1 | a:1830 | 3.85 | 1.0 | 15.6 |  | 18.92 | 9 | A | O3' | a:1791 | 3.89 | 1.0 | 26.3 |  | 14.96 | 10 | A | OP2 | a:1791 | 4.03 | 1.0 | 22.2 |  | 15.19 | 11 | C | OP1 | a:1792 | 4.18 | 1.0 | 23.4 |  | 15.66 | 12 | U | O4 | a:1829 | 4.64 | 1.0 | 24.0 |  | 15.87 |

  
**1^st^ shell coordination angles and distances (*trans* in "blue"; *cys* in "green"; deviations in "orange"):**   
• 1-2(W/OP2): 94; 1-3(W/OP1): 87; 1-4(W/W): 111; 1-5(W/O6): 178; 2-3(OP2/OP1): 135; 2-4(OP2/W): 99; 2-5(OP2/O6): 84; 3-4(OP1/W): 122; 3-5(OP1/O6): 95; 4-5(W/O6): 68;   
• 1-2(W/OP2): 3.5; 1-3(W/OP1): 3.5; 1-4(W/W): 4.2; 1-5(W/O6): 5.1; 2-3(OP2/OP1): 4.8; 2-4(OP2/W): 4.0; 2-5(OP2/O6): 3.5; 3-4(OP1/W): 4.8; 3-5(OP1/O6): 4.0; 4-5(W/O6): 3.1;   
  
**3.95 ± 1.27: the average coodination distance of the 1^st^ six ligands could correspond to an hexahydrated Mg^2+^.**  
**6: Number of potential Water contacts**   
**1 : Number of second shell contacts involving water or ions**   
**17%: Percentage of satisfied second shell Water contacts**   
• Contact(s): /O6/O_w_/OP1/  
• Contact code: O_ph_O_b_O_w_  
• Contact(s) in the exclusion zone: /O_back_/OP2/  
→ Coordination < 6 for K ←  
Go to ToC

### K:a:3127

|  | Res. type | Atom | Residue | Distance | Occupancy | B-factor | Closest atom(s) to a 1^st^ shell water | EM\_dens. |
| --- | --- | --- | --- | --- | --- | --- | --- | --- |
| **Ion** | | | |  | 1.0 | 30.0 |  | 37.54 |
 1 | U | O2 | a:2026 | 2.68 | 1.0 | 26.3 |  | 16.87 | 2 | Water 1 | | a:7646 | 2.69 | 1.0 | 29.4 | 2.85 [A:N3]a:2058 | 14.27 | 3 | Water 2 | | a:9666 | 2.85 | 1.0 | 24.1 |  | 15.96 | 4 | Water 3 | | a:9157 | 2.86 | 1.0 | 27.1 |  | 15.76 | 5 | C | O2' | a:2620 | 2.98 | 1.0 | 23.1 |  | 17.26 | 6 | C | O2 | a:2620 | 3.09 | 1.0 | 24.0 |  | 18.73 | 7 | U | O4' | a:2621 | 3.73 | 1.0 | 31.0 |  | 16.13 | 8 | G | N2 | a:2057 | 4.05 | 1.0 | 15.0 |  | 19.51 | 9 | Water 4 | | a:6854 | 4.21 | 1.0 | 22.5 |  | 15.09 | 10 | U | O4 | a:2038 | 4.24 | 1.0 | 27.2 |  | 16.79 | 11 | Water 5 | | z:208 | 4.55 | 1.0 | 26.5 | 2.97 [U:N3]a:2026 | 10.95 | 12 | C | N1 | a:2620 | 4.61 | 1.0 | 20.9 |  | 18.56 | 13 | A | N3 | a:2058 | 4.63 | 1.0 | 18.0 |  | 16.92 | 14 | U | N1 | a:2026 | 4.70 | 1.0 | 17.5 |  | 17.67 | 15 | U | O2' | a:2026 | 4.78 | 1.0 | 27.1 |  | 13.77 |

  
**1^st^ shell coordination angles and distances (*trans* in "blue"; *cys* in "green"; deviations in "orange"):**   
• 1-2(O2/W): 91; 1-3(O2/W): 84; 1-4(O2/W): 145; 1-5(O2/O2'): 65; 1-6(O2/O2): 122; 2-3(W/W): 144; 2-4(W/W): 87; 2-5(W/O2'): 125; 2-6(W/O2): 79; 3-4(W/W): 78; 3-5(W/O2'): 85; 3-6(W/O2): 133; 4-5(W/O2'): 140; 4-6(W/O2): 91; 5-6(O2'/O2): 75;   
• 1-2(O2/W): 3.8; 1-3(O2/W): 3.7; 1-4(O2/W): 5.3; 1-5(O2/O2'): 3.1; 1-6(O2/O2): 5.1; 2-3(W/W): 5.3; 2-4(W/W): 3.8; 2-5(W/O2'): 5.0; 2-6(W/O2): 3.7; 3-4(W/W): 3.6; 3-5(W/O2'): 3.9; 3-6(W/O2): 5.5; 4-5(W/O2'): 5.5; 4-6(W/O2): 4.3; 5-6(O2'/O2): 3.7;   
  
**4.15 ± 1.32: the average coodination distance of the 1^st^ six ligands could correspond to an hexahydrated Mg^2+^.**  
**6: Number of potential Water contacts**   
**1 : Number of second shell contacts involving water or ions**   
**17%: Percentage of satisfied second shell Water contacts**   
• Contact(s): /O2/O2/O2'/O_w_/O_w_/O_w_/  
• Contact code: O_r_2O_b_3O_w_  
Go to ToC

### K:a:3128

|  | Res. type | Atom | Residue | Distance | Occupancy | B-factor | Closest atom(s) to a 1^st^ shell water | EM\_dens. |
| --- | --- | --- | --- | --- | --- | --- | --- | --- |
| **Ion** | | | |  | 1.0 | 30.0 |  | 12.55 |
 1 | Water 1 | | a:9017 | 2.74 | 1.0 | 40.5 | 3.00 [C:OP1]a:334 | 3.76 | 2 | A | O2' | a:332 | 2.80 | 1.0 | 35.7 |  | 7.06 | 3 | G | O6 | a:326 | 2.85 | 1.0 | 39.3 |  | 7.81 | 4 | Water 2 | | a:6602 | 2.88 | 1.0 | 45.1 |  | 4.68 | 5 | Water 3 | | a:6780 | 2.88 | 1.0 | 45.9 | 2.79 [G:N7]a:327 | 3.81 | 6 | G | O6 | a:327 | 3.08 | 1.0 | 38.3 |  | 8.47 | 7 | Water 4 | | a:3432 | 3.09 | 1.0 | 46.6 | 2.36 [G:N7]a:326 | 3.53 | 8 | C | N4 | a:335 | 4.21 | 1.0 | 31.2 |  | 8.91 | 9 | G | N7 | a:327 | 4.29 | 1.0 | 38.2 |  | 7.27 | 10 | G | N7 | a:326 | 4.37 | 1.0 | 35.0 |  | 6.48 | 11 | C | OP1 | a:334 | 4.52 | 1.0 | 37.9 |  | 6.63 | 12 | Water 5 | | a:4802 | 4.75 | 1.0 | 43.8 |  | 3.46 |

  
**1^st^ shell coordination angles and distances (*trans* in "blue"; *cys* in "green"; deviations in "orange"):**   
• 1-2(W/O2'): 83; 1-3(W/O6): 73; 1-4(W/W): 169; 1-5(W/W): 142; 1-6(W/O6): 116; 1-7(W/W): 85; 2-3(O2'/O6): 138; 2-4(O2'/W): 86; 2-5(O2'/W): 134; 2-6(O2'/O6): 86; 2-7(O2'/W): 144; 3-4(O6/W): 116; 3-5(O6/W): 80; 3-6(O6/O6): 74; 3-7(O6/W): 68; 4-5(W/W): 49; 4-6(W/O6): 64; 4-7(W/W): 104; 5-6(W/O6): 81; 5-7(W/W): 60; 6-7(O6/W): 129;   
• 1-2(W/O2'): 3.7; 1-3(W/O6): 3.3; 1-4(W/W): 5.6; 1-5(W/W): 5.3; 1-6(W/O6): 4.9; 1-7(W/W): 3.9; 2-3(O2'/O6): 5.3; 2-4(O2'/W): 3.9; 2-5(O2'/W): 5.2; 2-6(O2'/O6): 4.0; 2-7(O2'/W): 5.6; 3-4(O6/W): 4.9; 3-5(O6/W): 3.7; 3-6(O6/O6): 3.6; 3-7(O6/W): 3.3; 4-5(W/W): 2.4; 4-6(W/O6): 3.2; 4-7(W/W): 4.7; 5-6(W/O6): 3.9; 5-7(W/W): 3.0; 6-7(O6/W): 5.6;   
  
**4.09 ± 1.29: the average coodination distance of the 1^st^ six ligands could correspond to an hexahydrated Mg^2+^.**  
**8: Number of potential Water contacts**   
**3 : Number of second shell contacts involving water or ions**   
**38%: Percentage of satisfied second shell Water contacts**   
• Contact(s): /O2'/O6/O6/O_w_/O_w_/O_w_/O_w_/  
• Contact code: O_r_2O_b_4O_w_  
• K hydration shell issue: water too close (< 2.4 Å.)  
Go to ToC

### K:a:3129

|  | Res. type | Atom | Residue | Distance | Occupancy | B-factor | Closest atom(s) to a 1^st^ shell water | EM\_dens. |
| --- | --- | --- | --- | --- | --- | --- | --- | --- |
| **Ion** | | | |  | 1.0 | 30.0 |  | 22.27 |
 1 | A | OP2 | a:2854 | 2.56 | 1.0 | 32.2 |  | 9.67 | 2 | U | O4 | a:2870 | 2.77 | 1.0 | 34.4 |  | 9.57 | 3 | U | O4 | a:2869 | 2.93 | 1.0 | 40.5 |  | 9.09 | 4 | Water 1 | | a:5199 | 2.96 | 1.0 | 38.1 | 2.69 [A:OP2]a:2855 // 2.86 [G:N7]a:2856 | 7.92 | 5 | G | O6 | a:2856 | 3.01 | 1.0 | 37.6 |  | 9.52 | 6 | Water 2 | | a:8987 | 3.16 | 1.0 | 43.2 | 2.99 [G:O6]a:2868 | 4.41 | 7 | A | N7 | a:2855 | 3.27 | 1.0 | 26.4 |  | 12.18 | 8 | A | OP2 | a:2855 | 3.60 | 1.0 | 31.5 |  | 10.06 | 9 | A | O5' | a:2854 | 3.93 | 1.0 | 33.7 |  | 10.26 | 10 | A | OP1 | a:2854 | 4.21 | 1.0 | 38.5 |  | 8.01 | 11 | Water 3 | | a:5308 | 4.23 | 1.0 | 41.4 | 2.70 [A:OP2]a:2855 | 6.23 | 12 | G | N7 | a:2856 | 4.27 | 1.0 | 29.9 |  | 10.94 | 13 | **K** | K | a:3064 | 4.30 | 1.0 | 30.0 |  | 13.40 | 14 | G | O6 | a:2868 | 4.40 | 1.0 | 37.2 |  | 7.65 |

  
**1^st^ shell coordination angles and distances (*trans* in "blue"; *cys* in "green"; deviations in "orange"):**   
• 1-2(OP2/O4): 77; 1-3(OP2/O4): 74; 1-4(OP2/W): 136; 1-5(OP2/O6): 145; 1-6(OP2/W): 92; 2-3(O4/O4): 70; 2-4(O4/W): 144; 2-5(O4/O6): 78; 2-6(O4/W): 136; 3-4(O4/W): 124; 3-5(O4/O6): 75; 3-6(O4/W): 66; 4-5(W/O6): 76; 4-6(W/W): 68; 5-6(O6/W): 90;   
• 1-2(OP2/O4): 3.3; 1-3(OP2/O4): 3.3; 1-4(OP2/W): 5.1; 1-5(OP2/O6): 5.3; 1-6(OP2/W): 4.1; 2-3(O4/O4): 3.3; 2-4(O4/W): 5.5; 2-5(O4/O6): 3.6; 2-6(O4/W): 5.5; 3-4(O4/W): 5.2; 3-5(O4/O6): 3.6; 3-6(O4/W): 3.3; 4-5(W/O6): 3.7; 4-6(W/W): 3.4; 5-6(O6/W): 4.4;   
  
**4.05 ± 1.22: the average coodination distance of the 1^st^ six ligands could correspond to an hexahydrated Mg^2+^.**  
**4: Number of potential Water contacts**   
**3 : Number of second shell contacts involving water or ions**   
**75%: Percentage of satisfied second shell Water contacts**   
• Contact(s): /O4/O4/O6/O_w_/O_w_/  
• Contact code: 3O_b_2O_w_  
• Contact(s) in the exclusion zone: /OP2/  
→ Coordination < 6 for K ←  
Go to ToC

### K:a:3130

|  | Res. type | Atom | Residue | Distance | Occupancy | B-factor | Closest atom(s) to a 1^st^ shell water | EM\_dens. |
| --- | --- | --- | --- | --- | --- | --- | --- | --- |
| **Ion** | | | |  | 1.0 | 30.0 |  | 21.64 |
 1 | Water 1 | | a:8758 | 2.75 | 1.0 | 41.8 | 2.96 [G:O6]a:666 | 6.49 | 2 | Water 2 | | a:5049 | 2.79 | 1.0 | 31.4 | 2.68 [G:N7]a:665 | 10.35 | 3 | U | O4 | a:596 | 2.89 | 1.0 | 29.0 |  | 13.18 | 4 | G | O6 | a:665 | 2.89 | 1.0 | 25.6 |  | 13.84 | 5 | Water 3 | | a:10068 | 3.00 | 1.0 | 36.7 | 2.23 [MG:MG]a:3067 | 6.31 | 6 | G | O6 | a:664 | 3.05 | 1.0 | 26.5 |  | 13.76 | 7 | G | O6 | a:666 | 3.47 | 1.0 | 28.7 |  | 13.03 | 8 | U | O4 | a:595 | 3.57 | 1.0 | 33.1 |  | 12.04 | 9 | Water 4 | | a:7656 | 3.75 | 1.0 | 38.2 | 1.95 [MG:MG]a:3067 | 7.32 | 10 | Water 5 | | a:4981 | 4.05 | 1.0 | 32.0 | 2.67 [G:N7]a:666 | 8.78 | 11 | G | N7 | a:665 | 4.08 | 1.0 | 20.2 |  | 13.00 | 12 | Water 6 | | a:4338 | 4.16 | 1.0 | 40.7 |  | 5.15 | 13 | Water 7 | | a:7719 | 4.38 | 1.0 | 34.0 | 2.14 [MG:MG]a:3067 | 8.96 | 14 | **MG** | MG | a:3067 | 4.44 | 1.0 | 51.9 |  | 8.14 | 15 | Water 8 | | a:4346 | 4.63 | 1.0 | 45.2 | 2.62 [U:O4]a:595 | 4.27 | 16 | G | N7 | a:666 | 4.71 | 1.0 | 16.8 |  | 14.75 | 17 | G | N7 | a:664 | 4.77 | 1.0 | 20.6 |  | 12.76 |

  
**1^st^ shell coordination angles and distances (*trans* in "blue"; *cys* in "green"; deviations in "orange"):**   
• 1-2(W/W): 80; 1-3(W/O4): 110; 1-4(W/O6): 112; 1-5(W/W): 77; 1-6(W/O6): 159; 2-3(W/O4): 165; 2-4(W/O6): 88; 2-5(W/W): 90; 2-6(W/O6): 81; 3-4(O4/O6): 77; 3-5(O4/W): 104; 3-6(O4/O6): 91; 4-5(O6/W): 171; 4-6(O6/O6): 77; 5-6(W/O6): 94;   
• 1-2(W/W): 3.5; 1-3(W/O4): 4.6; 1-4(W/O6): 4.7; 1-5(W/W): 3.6; 1-6(W/O6): 5.7; 2-3(W/O4): 5.6; 2-4(W/O6): 4.0; 2-5(W/W): 4.1; 2-6(W/O6): 3.8; 3-4(O4/O6): 3.6; 3-5(O4/W): 4.6; 3-6(O4/O6): 4.2; 4-5(O6/W): 5.9; 4-6(O6/O6): 3.7; 5-6(W/O6): 4.4;   
  
**4.06 ± 1.22: the average coodination distance of the 1^st^ six ligands could correspond to an hexahydrated Mg^2+^.**  
**6: Number of potential Water contacts**   
**3 : Number of second shell contacts involving water or ions**   
**50%: Percentage of satisfied second shell Water contacts**   
• Contact(s): /O4/O6/O6/O_w_/O_w_/O_w_/  
• Contact code: 3O_b_3O_w_  
Go to ToC

### K:a:3131

|  | Res. type | Atom | Residue | Distance | Occupancy | B-factor | Closest atom(s) to a 1^st^ shell water | EM\_dens. |
| --- | --- | --- | --- | --- | --- | --- | --- | --- |
| **Ion** | | | |  | 1.0 | 30.0 |  | 2.77 |
 1 | U | O4 | a:931 | 2.44 | 1.0 | 37.0 |  | 7.69 | 2 | U | O4 | a:848 | 3.40 | 1.0 | 73.2 |  | 1.03 | 3 | A | N7 | a:930 | 4.29 | 1.0 | 34.2 |  | 9.16 | 4 | G | O6 | a:932 | 4.31 | 1.0 | 43.3 |  | 5.35 | 5 | U | N3 | a:931 | 4.57 | 1.0 | 31.1 |  | 7.85 | 6 | A | N6 | a:930 | 4.61 | 1.0 | 33.1 |  | 8.89 |

  
• • **3.93 ± 0.78: the average coodination distance of the 1^st^ six ligands could correspond to an hexahydrated Mg^2+^.**  
**0: Number of potential Water contacts**   
• Contact code: → No contacts ←   
• Contact(s) in the exclusion zone: /O4/  
→ First K coordination shell missing  
→2Fo-Fc density between 1.0 and 4.0 !   
Go to ToC

### K:a:3132

|  | Res. type | Atom | Residue | Distance | Occupancy | B-factor | Closest atom(s) to a 1^st^ shell water | EM\_dens. |
| --- | --- | --- | --- | --- | --- | --- | --- | --- |
| **Ion** | | | |  | 1.0 | 30.0 |  | 13.33 |
 1 | U | O2' | a:390 | 2.63 | 1.0 | 44.2 |  | 7.06 | 2 | Water 1 | | a:9323 | 2.68 | 1.0 | 40.5 |  | 6.60 | 3 | U | O4 | a:392 | 2.80 | 1.0 | 36.2 |  | 9.17 | 4 | A | OP2 | a:391 | 2.93 | 1.0 | 37.5 |  | 7.04 | 5 | U | O4 | a:387 | 3.11 | 1.0 | 52.7 |  | 4.04 | 6 | G | O6 | a:388 | 3.28 | 1.0 | 41.8 |  | 5.82 | 7 | A | N7 | a:391 | 3.63 | 1.0 | 28.9 |  | 11.15 | 8 | U | O3' | a:390 | 3.75 | 1.0 | 38.3 |  | 7.57 | 9 | Water 2 | | a:7988 | 4.39 | 1.0 | 47.6 |  | 3.86 | 10 | U | O2 | a:390 | 4.53 | 1.0 | 48.3 |  | 4.05 |

  
**1^st^ shell coordination angles and distances (*trans* in "blue"; *cys* in "green"; deviations in "orange"):**   
• 1-2(O2'/W): 88; 1-3(O2'/O4): 162; 1-4(O2'/OP2): 81; 1-5(O2'/O4): 92; 2-3(W/O4): 90; 2-4(W/OP2): 139; 2-5(W/O4): 115; 3-4(O4/OP2): 88; 3-5(O4/O4): 105; 4-5(OP2/O4): 105;   
• 1-2(O2'/W): 3.7; 1-3(O2'/O4): 5.4; 1-4(O2'/OP2): 3.6; 1-5(O2'/O4): 4.1; 2-3(W/O4): 3.9; 2-4(W/OP2): 5.3; 2-5(W/O4): 4.9; 3-4(O4/OP2): 4.0; 3-5(O4/O4): 4.7; 4-5(OP2/O4): 4.8;   
  
**4.14 ± 1.28: the average coodination distance of the 1^st^ six ligands could correspond to an hexahydrated Mg^2+^.**  
**2: Number of potential Water contacts**   
**0 : Number of second shell contacts involving water or ions**   
**0%: Percentage of satisfied second shell Water contacts**   
• Contact(s): /O2'/O4/O4/O_w_/OP2/  
• Contact code: O_ph_O_r_2O_b_O_w_  
→ Coordination < 6 for K ←  
Go to ToC

### K:a:3133

|  | Res. type | Atom | Residue | Distance | Occupancy | B-factor | Closest atom(s) to a 1^st^ shell water | EM\_dens. |
| --- | --- | --- | --- | --- | --- | --- | --- | --- |
| **Ion** | | | |  | 1.0 | 30.0 |  | 26.17 |
 1 | G | O3' | a:1024 | 2.68 | 1.0 | 34.9 |  | 10.71 | 2 | U | OP2 | a:1025 | 2.72 | 1.0 | 30.2 |  | 9.92 | 3 | G | N7 | a:1026 | 2.76 | 1.0 | 26.9 |  | 12.63 | 4 | G | O2' | a:1024 | 3.00 | 1.0 | 36.8 |  | 10.02 | 5 | G | O6 | a:1026 | 3.03 | 1.0 | 28.2 |  | 10.56 | 6 | G | N3 | a:1027 | 3.07 | 1.0 | 26.6 |  | 12.42 | 7 | Water 1 | | a:8955 | 3.10 | 1.0 | 39.2 | 2.99 [C:OP2]a:1137 | 6.87 | 8 | G | O2' | a:1027 | 3.27 | 1.0 | 33.3 |  | 11.69 | 9 | **MG** | MG | a:3144 | 3.90 | 1.0 | 34.8 |  | 16.23 | 10 | Water 2 | | a:6103 | 4.13 | 1.0 | 33.9 | 2.39 [MG:MG]a:3144 | 9.70 | 11 | U | O5' | a:1025 | 4.34 | 1.0 | 33.0 |  | 10.56 | 12 | U | OP1 | a:1025 | 4.37 | 1.0 | 35.0 |  | 10.05 | 13 | C | OP2 | a:1137 | 4.57 | 1.0 | 34.9 |  | 8.34 |

  
**1^st^ shell coordination angles and distances (*trans* in "blue"; *cys* in "green"; deviations in "orange"):**   
• 1-2(O3'/OP2): 55; 1-3(O3'/N7): 87; 1-4(O3'/O2'): 57; 1-5(O3'/O6): 93; 1-6(O3'/N3): 171; 1-7(O3'/W): 109; 2-3(OP2/N7): 85; 2-4(OP2/O2'): 107; 2-5(OP2/O6): 136; 2-6(OP2/N3): 134; 2-7(OP2/W): 93; 3-4(N7/O2'): 115; 3-5(N7/O6): 64; 3-6(N7/N3): 93; 3-7(N7/W): 159; 4-5(O2'/O6): 66; 4-6(O2'/N3): 115; 4-7(O2'/W): 86; 5-6(O6/N3): 79; 5-7(O6/W): 127; 6-7(N3/W): 73;   
• 1-2(O3'/OP2): 2.5; 1-3(O3'/N7): 3.8; 1-4(O3'/O2'): 2.7; 1-5(O3'/O6): 4.1; 1-6(O3'/N3): 5.7; 1-7(O3'/W): 4.7; 2-3(OP2/N7): 3.7; 2-4(OP2/O2'): 4.6; 2-5(OP2/O6): 5.3; 2-6(OP2/N3): 5.3; 2-7(OP2/W): 4.2; 3-4(N7/O2'): 4.9; 3-5(N7/O6): 3.1; 3-6(N7/N3): 4.2; 3-7(N7/W): 5.8; 4-5(O2'/O6): 3.3; 4-6(O2'/N3): 5.1; 4-7(O2'/W): 4.1; 5-6(O6/N3): 3.9; 5-7(O6/W): 5.5; 6-7(N3/W): 3.7;   
  
**3.94 ± 1.16: the average coodination distance of the 1^st^ six ligands could correspond to an hexahydrated Mg^2+^.**  
**2: Number of potential Water contacts**   
**1 : Number of second shell contacts involving water or ions**   
**50%: Percentage of satisfied second shell Water contacts**   
• Contact(s): /N3_G_/N7/O2'/O3'/O6/O_w_/OP2/  
• Contact code: O_ph_2O_r_O_b_2N_b_O_w_  
Go to ToC

### K:a:3134

|  | Res. type | Atom | Residue | Distance | Occupancy | B-factor | Closest atom(s) to a 1^st^ shell water | EM\_dens. |
| --- | --- | --- | --- | --- | --- | --- | --- | --- |
| **Ion** | | | |  | 1.0 | 30.0 |  | 29.50 |
 1 | GLY | O | k:32 | 2.60 | 1.0 | 27.1 |  | 17.41 | 2 | Water 1 | | a:9733 | 2.69 | 1.0 | 45.6 |  | 5.30 | 3 | A | O2' | a:943 | 2.79 | 1.0 | 28.3 |  | 16.61 | 4 | G | OP1 | a:1193 | 2.94 | 1.0 | 31.9 |  | 12.29 | 5 | A | O3' | a:943 | 3.02 | 1.0 | 32.1 |  | 12.32 | 6 | Water 2 | | a:5054 | 3.05 | 1.0 | 37.6 |  | 7.94 | 7 | G | O3' | a:1192 | 3.22 | 1.0 | 28.2 |  | 13.16 | 8 | Water 3 | | k:210 | 3.93 | 1.0 | 39.1 |  | 4.56 | 9 | G | OP1 | a:944 | 4.01 | 1.0 | 27.5 |  | 11.92 | 10 | G | O5' | a:944 | 4.72 | 1.0 | 28.4 |  | 14.30 | 11 | G | OP2 | a:1193 | 4.76 | 1.0 | 28.5 |  | 12.34 | 12 | G | O5' | a:1193 | 4.78 | 1.0 | 28.8 |  | 15.19 |

  
**1^st^ shell coordination angles and distances (*trans* in "blue"; *cys* in "green"; deviations in "orange"):**   
• 1-2(W/W): 79; 1-3(W/O2'): 115; 1-4(W/OP1): 110; 1-5(W/O3'): 99; 1-6(W/W): 154; 2-3(W/O2'): 136; 2-4(W/OP1): 131; 2-5(W/O3'): 82; 2-6(W/W): 88; 3-4(O2'/OP1): 84; 3-5(O2'/O3'): 56; 3-6(O2'/W): 90; 4-5(OP1/O3'): 138; 4-6(OP1/W): 63; 5-6(O3'/W): 101;   
• 1-2(W/W): 3.4; 1-3(W/O2'): 4.6; 1-4(W/OP1): 4.6; 1-5(W/O3'): 4.3; 1-6(W/W): 5.5; 2-3(W/O2'): 5.1; 2-4(W/OP1): 5.1; 2-5(W/O3'): 3.7; 2-6(W/W): 4.0; 3-4(O2'/OP1): 3.8; 3-5(O2'/O3'): 2.7; 3-6(O2'/W): 4.1; 4-5(OP1/O3'): 5.6; 4-6(OP1/W): 3.1; 5-6(O3'/W): 4.7;   
  
**4.04 ± 1.25: the average coodination distance of the 1^st^ six ligands could correspond to an hexahydrated Mg^2+^.**  
**8: Number of potential Water contacts**   
**0 : Number of second shell contacts involving water or ions**   
**0%: Percentage of satisfied second shell Water contacts**   
• Contact(s): /O2'/O3'/O_back_/O_w_/O_w_/OP1/  
• Contact code: O_ph_2O_r_O_back_2O_w_  
Go to ToC

### K:a:3135

|  | Res. type | Atom | Residue | Distance | Occupancy | B-factor | Closest atom(s) to a 1^st^ shell water | EM\_dens. |
| --- | --- | --- | --- | --- | --- | --- | --- | --- |
| **Ion** | | | |  | 1.0 | 30.0 |  | 19.59 |
 1 | G | O6 | a:533 | 2.89 | 1.0 | 26.8 |  | 14.64 | 2 | U | O4 | a:534 | 2.92 | 1.0 | 28.5 |  | 13.90 | 3 | G | O6 | a:561 | 2.95 | 1.0 | 25.4 |  | 12.94 | 4 | U | O4 | a:560 | 3.06 | 1.0 | 32.1 |  | 10.89 | 5 | Water 1 | | a:3905 | 3.19 | 1.0 | 36.4 | 2.56 [G:N7]a:561 | 6.31 | 6 | Water 2 | | a:5280 | 3.19 | 1.0 | 23.6 | 2.77 [G:O6]a:535 | 11.02 | 7 | G | O6 | a:535 | 3.57 | 1.0 | 27.0 |  | 12.55 | 8 | U | N3 | a:560 | 3.79 | 1.0 | 17.9 |  | 13.68 | 9 | Water 3 | | a:6750 | 4.09 | 1.0 | 36.9 |  | 6.74 | 10 | G | N7 | a:561 | 4.30 | 1.0 | 24.6 |  | 13.47 | 11 | Water 4 | | a:5901 | 4.50 | 1.0 | 38.4 |  | 4.02 | 12 | Water 5 | | a:9254 | 4.54 | 1.0 | 40.2 |  | 4.12 |

  
**1^st^ shell coordination angles and distances (*trans* in "blue"; *cys* in "green"; deviations in "orange"):**   
• 1-2(O6/O4): 77; 1-3(O6/O6): 76; 1-4(O6/O4): 160; 1-5(O6/W): 114; 1-6(O6/W): 128; 2-3(O4/O6): 78; 2-4(O4/O4): 102; 2-5(O4/W): 153; 2-6(O4/W): 52; 3-4(O6/O4): 124; 3-5(O6/W): 81; 3-6(O6/W): 100; 4-5(O4/W): 76; 4-6(O4/W): 51; 5-6(W/W): 116;   
• 1-2(O6/O4): 3.6; 1-3(O6/O6): 3.6; 1-4(O6/O4): 5.9; 1-5(O6/W): 5.1; 1-6(O6/W): 5.5; 2-3(O4/O6): 3.7; 2-4(O4/O4): 4.7; 2-5(O4/W): 5.9; 2-6(O4/W): 2.7; 3-4(O6/O4): 5.3; 3-5(O6/W): 4.0; 3-6(O6/W): 4.7; 4-5(O4/W): 3.8; 4-6(O4/W): 2.7; 5-6(W/W): 5.4;   
  
**4.26 ± 1.28: the average coodination distance of the 1^st^ six ligands could correspond to an hexahydrated Mg^2+^.**  
**4: Number of potential Water contacts**   
**2 : Number of second shell contacts involving water or ions**   
**50%: Percentage of satisfied second shell Water contacts**   
• Contact(s): /O4/O4/O6/O6/O_w_/O_w_/  
• Contact code: 4O_b_2O_w_  
Go to ToC

### K:a:3136

|  | Res. type | Atom | Residue | Distance | Occupancy | B-factor | Closest atom(s) to a 1^st^ shell water | EM\_dens. |
| --- | --- | --- | --- | --- | --- | --- | --- | --- |
| **Ion** | | | |  | 1.0 | 30.0 |  | 8.59 |
 1 | G | O6 | a:2345 | 2.78 | 1.0 | 42.9 |  | 6.39 | 2 | U | O4 | a:2295 | 2.91 | 1.0 | 41.0 |  | 6.88 | 3 | **K** | K | a:3137 | 3.01 | 1.0 | 30.0 |  | 10.45 | 4 | G | O6 | a:2294 | 3.05 | 1.0 | 40.0 |  | 8.54 | 5 | Water 1 | | a:8216 | 3.11 | 1.0 | 45.4 | 2.90 [G:N7]a:2345 | 3.57 | 6 | A | N6 | a:2344 | 3.23 | 1.0 | 38.0 |  | 5.23 | 7 | Water 2 | | a:5107 | 3.71 | 1.0 | 44.7 |  | 4.02 | 8 | C | N4 | a:2346 | 3.84 | 1.0 | 37.6 |  | 5.97 | 9 | G | N7 | a:2345 | 4.18 | 1.0 | 41.6 |  | 5.88 | 10 | Water 3 | | a:6237 | 4.23 | 1.0 | 43.2 |  | 3.72 | 11 | A | N7 | a:2344 | 4.48 | 1.0 | 43.0 |  | 4.82 | 12 | A | N1 | a:2344 | 4.66 | 1.0 | 40.5 |  | 6.85 | 13 | Water 4 | | a:3533 | 4.69 | 1.0 | 43.7 |  | 4.09 |

  
**1^st^ shell coordination angles and distances (*trans* in "blue"; *cys* in "green"; deviations in "orange"):**   
• 1-2(O6/O4): 77; 1-3(O6/K): 119; 1-4(O6/O6): 69; 1-5(O6/W): 91; 2-3(O4/K): 62; 2-4(O4/O6): 74; 2-5(O4/W): 162; 3-4(K/O6): 58; 3-5(K/W): 135; 4-5(O6/W): 114;   
• 1-2(O6/O4): 3.6; 1-3(O6/K): 5.0; 1-4(O6/O6): 3.3; 1-5(O6/W): 4.2; 2-3(O4/K): 3.1; 2-4(O4/O6): 3.6; 2-5(O4/W): 5.9; 3-4(K/O6): 2.9; 3-5(K/W): 5.7; 4-5(O6/W): 5.2;   
  
**4.27 ± 1.30: the average coodination distance of the 1^st^ six ligands could correspond to an hexahydrated Mg^2+^.**  
**4: Number of potential Water contacts**   
**1 : Number of second shell contacts involving water or ions**   
**25%: Percentage of satisfied second shell Water contacts**   
• Contact(s): /O4/O6/O6/O_w_/  
• Contact code: 3O_b_O_w_  
→ Coordination < 6 for K ←  
Go to ToC

### K:a:3137

|  | Res. type | Atom | Residue | Distance | Occupancy | B-factor | Closest atom(s) to a 1^st^ shell water | EM\_dens. |
| --- | --- | --- | --- | --- | --- | --- | --- | --- |
| **Ion** | | | |  | 1.0 | 30.0 |  | 10.45 |
 1 | G | N7 | a:2294 | 2.77 | 1.0 | 35.3 |  | 7.45 | 2 | Water 1 | | a:6314 | 2.82 | 1.0 | 42.5 | 2.76 [G:OP2]a:2293 | 6.52 | 3 | Water 2 | | a:5107 | 2.88 | 1.0 | 44.7 | 2.68 [G:O6]a:2293 | 4.02 | 4 | G | O6 | a:2294 | 2.95 | 1.0 | 40.0 |  | 8.54 | 5 | Water 3 | | a:3533 | 2.98 | 1.0 | 43.7 | 2.46 [G:OP2]a:2293 | 4.09 | 6 | **K** | K | a:3136 | 3.01 | 1.0 | 30.0 |  | 8.59 | 7 | U | O4 | a:2295 | 3.07 | 1.0 | 41.0 |  | 6.88 | 8 | Water 4 | | a:6237 | 3.50 | 1.0 | 43.2 |  | 3.72 | 9 | G | N7 | a:2293 | 3.69 | 1.0 | 32.3 |  | 9.18 | 10 | G | O6 | a:2293 | 3.89 | 1.0 | 36.5 |  | 9.29 | 11 | A | O2' | a:2291 | 4.48 | 1.0 | 43.0 |  | 5.94 | 12 | G | OP2 | a:2293 | 4.51 | 1.0 | 39.0 |  | 6.60 |

  
**1^st^ shell coordination angles and distances (*trans* in "blue"; *cys* in "green"; deviations in "orange"):**   
• 1-2(N7/W): 84; 1-3(N7/W): 107; 1-4(N7/O6): 64; 1-5(N7/W): 139; 1-6(N7/K): 118; 1-7(N7/O4): 80; 2-3(W/W): 137; 2-4(W/O6): 146; 2-5(W/W): 65; 2-6(W/K): 134; 2-7(W/O4): 91; 3-4(W/O6): 68; 3-5(W/W): 82; 3-6(W/K): 78; 3-7(W/O4): 131; 4-5(O6/W): 148; 4-6(O6/K): 62; 4-7(O6/O4): 73; 5-6(W/K): 103; 5-7(W/O4): 124; 6-7(K/O4): 57;   
• 1-2(N7/W): 3.7; 1-3(N7/W): 4.6; 1-4(N7/O6): 3.0; 1-5(N7/W): 5.4; 1-6(N7/K): 5.0; 1-7(N7/O4): 3.8; 2-3(W/W): 5.3; 2-4(W/O6): 5.5; 2-5(W/W): 3.1; 2-6(W/K): 5.4; 2-7(W/O4): 4.2; 3-4(W/O6): 3.2; 3-5(W/W): 3.8; 3-6(W/K): 3.7; 3-7(W/O4): 5.4; 4-5(O6/W): 5.7; 4-6(O6/K): 3.1; 4-7(O6/O4): 3.6; 5-6(W/K): 4.7; 5-7(W/O4): 5.3; 6-7(K/O4): 2.9;   
  
**4.00 ± 1.18: the average coodination distance of the 1^st^ six ligands could correspond to an hexahydrated Mg^2+^.**  
**8: Number of potential Water contacts**   
**3 : Number of second shell contacts involving water or ions**   
**38%: Percentage of satisfied second shell Water contacts**   
• Contact(s): /N7/O4/O6/O_w_/O_w_/O_w_/  
• Contact code: 2O_b_N_b_3O_w_  
Go to ToC

### K:a:3138

|  | Res. type | Atom | Residue | Distance | Occupancy | B-factor | Closest atom(s) to a 1^st^ shell water | EM\_dens. |
| --- | --- | --- | --- | --- | --- | --- | --- | --- |
| **Ion** | | | |  | 1.0 | 30.0 |  | 9.78 |
 1 | Water 1 | | a:4581 | 2.52 | 1.0 | 33.3 | 2.64 [U:O2']a:829 // 2.87 [G:O3']a:2250 | 7.82 | 2 | U | OP1 | a:2249 | 2.83 | 1.0 | 34.7 |  | 10.31 | 3 | U | O2 | a:2072 | 3.02 | 1.0 | 36.7 |  | 10.76 | 4 | U | O2 | a:829 | 3.15 | 1.0 | 34.0 |  | 10.81 | 5 | U | O2' | a:2072 | 3.47 | 1.0 | 35.8 |  | 8.53 | 6 | Water 2 | | a:6042 | 3.55 | 1.0 | 38.4 | 2.85 [U:OP2]a:2249 | 8.45 | 7 | A | N6 | a:2434 | 3.64 | 1.0 | 27.2 |  | 12.64 | 8 | U | O2' | a:829 | 3.83 | 1.0 | 38.2 |  | 7.32 | 9 | U | OP2 | a:2249 | 3.90 | 1.0 | 32.4 |  | 11.54 | 10 | Water 3 | | a:7546 | 4.39 | 1.0 | 45.0 | 2.85 [U:O2']a:2072 | 4.91 | 11 | Water 4 | | a:5133 | 4.41 | 1.0 | 35.5 |  | 7.05 | 12 | U | N1 | a:2072 | 4.63 | 1.0 | 26.0 |  | 12.19 | 13 | G | O3' | a:2250 | 4.66 | 1.0 | 30.6 |  | 11.27 | 14 | Water 5 | | a:4779 | 4.73 | 1.0 | 43.6 |  | 3.14 | 15 | U | O5' | a:2249 | 4.77 | 1.0 | 30.4 |  | 12.71 |

  
**1^st^ shell coordination angles and distances (*trans* in "blue"; *cys* in "green"; deviations in "orange"):**   
• 1-2(W/OP1): 134; 1-3(W/O2): 99; 1-4(W/O2): 61; 2-3(OP1/O2): 105; 2-4(OP1/O2): 84; 3-4(O2/O2): 81;   
• 1-2(W/OP1): 4.9; 1-3(W/O2): 4.2; 1-4(W/O2): 2.9; 2-3(OP1/O2): 4.6; 2-4(OP1/O2): 4.0; 3-4(O2/O2): 4.0;   
  
**4.34 ± 1.34: the average coodination distance of the 1^st^ six ligands could correspond to an hexahydrated Mg^2+^.**  
**4: Number of potential Water contacts**   
**3 : Number of second shell contacts involving water or ions**   
**75%: Percentage of satisfied second shell Water contacts**   
• Contact(s): /O2/O2/OP1/  
• Contact code: O_ph_2O_b_  
• Contact(s) in the exclusion zone: /O_w_/  
→ Coordination < 6 for K ←  
Go to ToC

### K:a:3139

|  | Res. type | Atom | Residue | Distance | Occupancy | B-factor | Closest atom(s) to a 1^st^ shell water | EM\_dens. |
| --- | --- | --- | --- | --- | --- | --- | --- | --- |
| **Ion** | | | |  | 1.0 | 30.0 |  | 20.81 |
 1 | Water 1 | | a:10051 | 2.67 | 1.0 | 47.6 |  | 3.46 | 2 | G | O6 | a:941 | 2.75 | 1.0 | 25.6 |  | 15.40 | 3 | Water 2 | | a:4887 | 2.89 | 1.0 | 44.3 | 2.66 [G:O6]a:940 | 5.61 | 4 | U | O4 | a:841 | 3.01 | 1.0 | 23.4 |  | 15.14 | 5 | G | O6 | a:942 | 3.03 | 1.0 | 24.8 |  | 15.15 | 6 | G | O6 | a:940 | 3.25 | 1.0 | 32.1 |  | 11.46 | 7 | Water 3 | | a:4141 | 3.68 | 1.0 | 38.2 | 2.59 [G:N7]a:942 | 5.80 | 8 | U | N3 | a:841 | 3.98 | 1.0 | 16.0 |  | 17.80 | 9 | G | N7 | a:941 | 4.23 | 1.0 | 20.7 |  | 12.38 | 10 | Water 4 | | a:5648 | 4.27 | 1.0 | 40.1 |  | 4.83 | 11 | Water 5 | | a:9031 | 4.37 | 1.0 | 39.5 |  | 5.11 | 12 | Water 6 | | a:10296 | 4.46 | 1.0 | 45.3 |  | 3.34 | 13 | Water 7 | | a:8474 | 4.52 | 1.0 | 39.8 |  | 7.57 | 14 | G | N7 | a:942 | 4.53 | 1.0 | 17.6 |  | 13.94 |

  
**1^st^ shell coordination angles and distances (*trans* in "blue"; *cys* in "green"; deviations in "orange"):**   
• 1-2(W/O6): 160; 1-3(W/W): 81; 1-4(W/O4): 103; 1-5(W/O6): 90; 2-3(O6/W): 117; 2-4(O6/O4): 76; 2-5(O6/O6): 71; 3-4(W/O4): 77; 3-5(W/O6): 150; 4-5(O4/O6): 78;   
• 1-2(W/O6): 5.3; 1-3(W/W): 3.6; 1-4(W/O4): 4.4; 1-5(W/O6): 4.0; 2-3(O6/W): 4.8; 2-4(O6/O4): 3.6; 2-5(O6/O6): 3.3; 3-4(W/O4): 3.7; 3-5(W/O6): 5.7; 4-5(O4/O6): 3.8;   
  
**4.21 ± 1.31: the average coodination distance of the 1^st^ six ligands could correspond to an hexahydrated Mg^2+^.**  
**6: Number of potential Water contacts**   
**2 : Number of second shell contacts involving water or ions**   
**33%: Percentage of satisfied second shell Water contacts**   
• Contact(s): /O4/O6/O6/O_w_/O_w_/  
• Contact code: 3O_b_2O_w_  
→ Coordination < 6 for K ←  
Go to ToC

### K:a:3140

|  | Res. type | Atom | Residue | Distance | Occupancy | B-factor | Closest atom(s) to a 1^st^ shell water | EM\_dens. |
| --- | --- | --- | --- | --- | --- | --- | --- | --- |
| **Ion** | | | |  | 1.0 | 30.0 |  | 18.55 |
 1 | Water 1 | | a:8679 | 2.71 | 1.0 | 39.4 | 2.95 [G:OP2]a:117 | 6.14 | 2 | Water 2 | | a:6025 | 2.81 | 1.0 | 36.2 | 2.86 [A:N3]a:119 | 8.72 | 3 | A | O2' | a:119 | 2.84 | 1.0 | 33.2 |  | 11.39 | 4 | A | O3' | a:119 | 2.85 | 1.0 | 36.1 |  | 12.96 | 5 | Water 3 | | a:9349 | 2.88 | 1.0 | 32.9 | 1.94 [MG:MG]a:3078 | 9.50 | 6 | U | OP2 | a:120 | 3.13 | 1.0 | 31.9 |  | 10.24 | 7 | Water 4 | | a:4495 | 3.48 | 1.0 | 34.8 | 2.16 [MG:MG]a:3078 | 9.58 | 8 | **MG** | MG | a:3078 | 3.57 | 1.0 | 32.7 |  | 13.17 | 9 | Water 5 | | a:8024 | 4.20 | 1.0 | 34.8 |  | 8.62 | 10 | A | N3 | a:119 | 4.25 | 1.0 | 25.9 |  | 11.15 | 11 | G | OP2 | a:117 | 4.26 | 1.0 | 31.6 |  | 11.43 | 12 | Water 6 | | a:3856 | 4.44 | 1.0 | 46.9 |  | 4.10 | 13 | U | O5' | a:120 | 4.58 | 1.0 | 36.9 |  | 9.80 | 14 | G | OP1 | a:121 | 4.72 | 1.0 | 35.1 |  | 9.45 | 15 | U | OP1 | a:120 | 4.74 | 1.0 | 34.6 |  | 8.27 |

  
**1^st^ shell coordination angles and distances (*trans* in "blue"; *cys* in "green"; deviations in "orange"):**   
• 1-2(W/W): 123; 1-3(W/O2'): 69; 1-4(W/O3'): 120; 1-5(W/W): 79; 1-6(W/OP2): 135; 2-3(W/O2'): 98; 2-4(W/O3'): 89; 2-5(W/W): 151; 2-6(W/OP2): 102; 3-4(O2'/O3'): 57; 3-5(O2'/W): 108; 3-6(O2'/OP2): 102; 4-5(O3'/W): 94; 4-6(O3'/OP2): 49; 5-6(W/OP2): 61;   
• 1-2(W/W): 4.9; 1-3(W/O2'): 3.2; 1-4(W/O3'): 4.8; 1-5(W/W): 3.6; 1-6(W/OP2): 5.4; 2-3(W/O2'): 4.3; 2-4(W/O3'): 4.0; 2-5(W/W): 5.5; 2-6(W/OP2): 4.6; 3-4(O2'/O3'): 2.7; 3-5(O2'/W): 4.6; 3-6(O2'/OP2): 4.6; 4-5(O3'/W): 4.2; 4-6(O3'/OP2): 2.5; 5-6(W/OP2): 3.1;   
  
**4.04 ± 1.22: the average coodination distance of the 1^st^ six ligands could correspond to an hexahydrated Mg^2+^.**  
**8: Number of potential Water contacts**   
**4 : Number of second shell contacts involving water or ions**   
**50%: Percentage of satisfied second shell Water contacts**   
• Contact(s): /O2'/O3'/O_w_/O_w_/O_w_/OP2/  
• Contact code: O_ph_2O_r_3O_w_  
Go to ToC

### K:a:3141

|  | Res. type | Atom | Residue | Distance | Occupancy | B-factor | Closest atom(s) to a 1^st^ shell water | EM\_dens. |
| --- | --- | --- | --- | --- | --- | --- | --- | --- |
| **Ion** | | | |  | 1.0 | 30.0 |  | 25.68 |
 1 | Water 1 | | a:9660 | 2.73 | 1.0 | 39.2 |  | 6.30 | 2 | U | O4 | a:1399 | 2.86 | 1.0 | 32.7 |  | 11.47 | 3 | U | O2' | a:1604 | 2.88 | 1.0 | 31.3 |  | 14.19 | 4 | A | O5' | a:1397 | 2.95 | 1.0 | 37.0 |  | 9.28 | 5 | U | O2' | a:1396 | 2.96 | 1.0 | 33.5 |  | 9.92 | 6 | U | O3' | a:1396 | 3.08 | 1.0 | 26.8 |  | 11.86 | 7 | U | O3' | a:1604 | 3.09 | 1.0 | 30.7 |  | 11.41 | 8 | A | OP1 | a:1605 | 3.15 | 1.0 | 31.3 |  | 9.79 | 9 | A | OP1 | a:1397 | 3.29 | 1.0 | 33.2 |  | 8.65 | 10 | A | O5' | a:1605 | 4.16 | 1.0 | 29.2 |  | 13.51 | 11 | **MG** | MG | a:3244 | 4.17 | 1.0 | 30.5 |  | 15.30 | 12 | Water 2 | | a:5172 | 4.22 | 1.0 | 42.4 |  | 3.73 | 13 | Water 3 | | a:6698 | 4.34 | 1.0 | 32.8 | 2.20 [MG:MG]a:3244 | 9.63 | 14 | Water 4 | | a:9229 | 4.54 | 1.0 | 40.7 |  | 5.80 | 15 | G | OP1 | a:1343 | 4.57 | 1.0 | 30.5 |  | 9.70 | 16 | G | OP2 | a:1343 | 4.73 | 1.0 | 26.8 |  | 11.54 | 17 | U | O2 | a:1604 | 4.75 | 1.0 | 30.5 |  | 12.86 | 18 | A | OP2 | a:1397 | 4.79 | 1.0 | 36.5 |  | 9.58 |

  
**1^st^ shell coordination angles and distances (*trans* in "blue"; *cys* in "green"; deviations in "orange"):**   
• 1-2(W/O4): 76; 1-3(W/O2'): 111; 1-4(W/O5'): 94; 1-5(W/O2'): 161; 1-6(W/O3'): 125; 1-7(W/O3'): 74; 1-8(W/OP1): 65; 2-3(O4/O2'): 89; 2-4(O4/O5'): 89; 2-5(O4/O2'): 91; 2-6(O4/O3'): 129; 2-7(O4/O3'): 115; 2-8(O4/OP1): 140; 3-4(O2'/O5'): 153; 3-5(O2'/O2'): 81; 3-6(O2'/O3'): 116; 3-7(O2'/O3'): 54; 3-8(O2'/OP1): 98; 4-5(O5'/O2'): 72; 4-6(O5'/O3'): 49; 4-7(O5'/O3'): 149; 4-8(O5'/OP1): 102; 5-6(O2'/O3'): 54; 5-7(O2'/O3'): 124; 5-8(O2'/OP1): 129; 6-7(O3'/O3'): 115; 6-8(O3'/OP1): 83; 7-8(O3'/OP1): 47;   
• 1-2(W/O4): 3.4; 1-3(W/O2'): 4.6; 1-4(W/O5'): 4.2; 1-5(W/O2'): 5.6; 1-6(W/O3'): 5.2; 1-7(W/O3'): 3.5; 1-8(W/OP1): 3.2; 2-3(O4/O2'): 4.0; 2-4(O4/O5'): 4.1; 2-5(O4/O2'): 4.2; 2-6(O4/O3'): 5.4; 2-7(O4/O3'): 5.0; 2-8(O4/OP1): 5.7; 3-4(O2'/O5'): 5.7; 3-5(O2'/O2'): 3.8; 3-6(O2'/O3'): 5.1; 3-7(O2'/O3'): 2.7; 3-8(O2'/OP1): 4.5; 4-5(O5'/O2'): 3.5; 4-6(O5'/O3'): 2.5; 4-7(O5'/O3'): 5.8; 4-8(O5'/OP1): 4.7; 5-6(O2'/O3'): 2.8; 5-7(O2'/O3'): 5.3; 5-8(O2'/OP1): 5.5; 6-7(O3'/O3'): 5.2; 6-8(O3'/OP1): 4.1; 7-8(O3'/OP1): 2.5;   
  
**3.95 ± 1.15: the average coodination distance of the 1^st^ six ligands could correspond to an hexahydrated Mg^2+^.**  
**2: Number of potential Water contacts**   
**0 : Number of second shell contacts involving water or ions**   
**0%: Percentage of satisfied second shell Water contacts**   
• Contact(s): /O2'/O2'/O3'/O3'/O4/O5'/O_w_/OP1/  
• Contact code: O_ph_5O_r_O_b_O_w_  
Go to ToC

### K:a:3142

|  | Res. type | Atom | Residue | Distance | Occupancy | B-factor | Closest atom(s) to a 1^st^ shell water | EM\_dens. |
| --- | --- | --- | --- | --- | --- | --- | --- | --- |
| **Ion** | | | |  | 1.0 | 30.0 |  | 18.36 |
 1 | A | O2' | a:13 | 2.58 | 1.0 | 36.8 |  | 9.67 | 2 | Water 1 | | a:5075 | 2.62 | 1.0 | 43.1 |  | 4.71 | 3 | G | O6 | a:15 | 2.76 | 1.0 | 24.0 |  | 13.00 | 4 | U | O4 | a:525 | 2.87 | 1.0 | 30.5 |  | 13.50 | 5 | Water 2 | | a:9932 | 3.08 | 1.0 | 46.2 |  | 3.25 | 6 | G | O6 | a:524 | 3.13 | 1.0 | 28.9 |  | 14.17 | 7 | C | N4 | a:16 | 3.30 | 1.0 | 19.1 |  | 14.46 | 8 | U | N3 | a:525 | 3.79 | 1.0 | 18.4 |  | 14.18 | 9 | Water 3 | | a:3884 | 4.30 | 1.0 | 31.9 |  | 8.29 | 10 | G | N7 | a:15 | 4.35 | 1.0 | 19.6 |  | 14.03 | 11 | Water 4 | | a:8918 | 4.37 | 1.0 | 40.0 | 2.98 [C:N4]a:16 | 6.24 | 12 | Water 5 | | a:8596 | 4.57 | 1.0 | 37.3 |  | 6.82 | 13 | C | N3 | a:16 | 4.77 | 1.0 | 17.6 |  | 15.64 |

  
**1^st^ shell coordination angles and distances (*trans* in "blue"; *cys* in "green"; deviations in "orange"):**   
• 1-2(O2'/W): 150; 1-3(O2'/O6): 72; 1-4(O2'/O4): 121; 1-5(O2'/W): 83; 1-6(O2'/O6): 153; 2-3(W/O6): 139; 2-4(W/O4): 77; 2-5(W/W): 74; 2-6(W/O6): 55; 3-4(O6/O4): 79; 3-5(O6/W): 134; 3-6(O6/O6): 85; 4-5(O4/W): 83; 4-6(O4/O6): 66; 5-6(W/O6): 124;   
• 1-2(O2'/W): 5.0; 1-3(O2'/O6): 3.1; 1-4(O2'/O4): 4.7; 1-5(O2'/W): 3.8; 1-6(O2'/O6): 5.5; 2-3(W/O6): 5.0; 2-4(W/O4): 3.4; 2-5(W/W): 3.5; 2-6(W/O6): 2.7; 3-4(O6/O4): 3.6; 3-5(O6/W): 5.4; 3-6(O6/O6): 4.0; 4-5(O4/W): 3.9; 4-6(O4/O6): 3.3; 5-6(W/O6): 5.5;   
  
**4.02 ± 1.24: the average coodination distance of the 1^st^ six ligands could correspond to an hexahydrated Mg^2+^.**  
**4: Number of potential Water contacts**   
**0 : Number of second shell contacts involving water or ions**   
**0%: Percentage of satisfied second shell Water contacts**   
• Contact(s): /O4/O6/O6/O_w_/O_w_/  
• Contact code: 3O_b_2O_w_  
• Contact(s) in the exclusion zone: /O2'/  
→ Coordination < 6 for K ←  
Go to ToC

### MG:a:3143

|  | Res. type | Atom | Residue | Distance | Occupancy | B-factor | Closest atom(s) to a 1^st^ shell water | EM\_dens. |
| --- | --- | --- | --- | --- | --- | --- | --- | --- |
| **Ion** | | | |  | 1.0 | 35.3 |  | 13.95 |
 1 | Water 1 | | a:6948 | 1.94 | 1.0 | 31.8 |  | 10.94 | 2 | Water 2 | | a:4631 | 1.94 | 1.0 | 33.7 |  | 9.78 | 3 | C | OP1 | a:393 | 1.99 | 1.0 | 36.2 |  | 7.34 | 4 | G | OP1 | a:375 | 2.09 | 1.0 | 32.0 |  | 9.86 | 5 | Water 3 | | a:9359 | 2.20 | 1.0 | 33.4 |  | 9.55 | 6 | Water 4 | | a:4636 | 2.26 | 1.0 | 31.5 | 2.64 [G:OP1]a:409 | 10.57 | 7 | A | O3' | a:374 | 3.40 | 1.0 | 33.7 |  | 9.98 | 8 | C | OP2 | a:393 | 3.85 | 1.0 | 33.5 |  | 9.14 | 9 | Water 5 | | a:6906 | 4.02 | 1.0 | 35.2 |  | 9.01 | 10 | Water 6 | | a:5747 | 4.04 | 1.0 | 37.2 | 2.82 [G:OP2]a:375 | 8.85 | 11 | U | O3' | a:392 | 4.14 | 1.0 | 38.3 |  | 9.97 | 12 | Water 7 | | a:5734 | 4.28 | 1.0 | 37.1 | 2.73 [U:O3']a:392 | 5.89 | 13 | G | O5' | a:375 | 4.31 | 1.0 | 37.9 |  | 9.34 | 14 | G | OP2 | a:375 | 4.36 | 1.0 | 35.2 |  | 7.68 | 15 | G | O3' | a:408 | 4.39 | 1.0 | 36.0 |  | 7.72 | 16 | C | O5' | a:393 | 4.43 | 1.0 | 34.5 |  | 8.77 | 17 | G | OP1 | a:409 | 4.50 | 1.0 | 37.8 |  | 8.35 | 18 | Water 8 | | a:9382 | 4.58 | 1.0 | 41.4 |  | 5.60 | 19 | G | O2' | a:408 | 4.77 | 1.0 | 36.6 |  | 8.70 | 20 | A | O2' | a:374 | 4.78 | 1.0 | 36.9 |  | 9.95 |

  
**1^st^ shell coordination angles and distances (*trans* in "blue"; *cys* in "green"; deviations in "orange"):**   
• 1-2(W/W): *95º*; 1-3(W/OP1): 91º; 1-4(W/OP1): *97º*; 1-5(W/W): 93º; 1-6(W/W): 175º; 2-3(W/OP1): *84º*; 2-4(W/OP1): *168º*; 2-5(W/W): *84º*; 2-6(W/W): *82º*; 3-4(OP1/OP1): *99º*; 3-5(OP1/W): *168º*; 3-6(OP1/W): 93º; 4-5(OP1/W): 92º; 4-6(OP1/W): 86º; 5-6(W/W): *82º*;   
• 1-2(W/W): 2.9Å; 1-3(W/OP1): 2.8Å; 1-4(W/OP1): 3.0Å; 1-5(W/W): 3.0Å; 1-6(W/W): 4.2Å; 2-3(W/OP1): 2.6Å; 2-4(W/OP1): 4.0Å; 2-5(W/W): 2.8Å; 2-6(W/W): 2.8Å; 3-4(OP1/OP1): 3.1Å; 3-5(OP1/W): 4.2Å; 3-6(OP1/W): 3.1Å; 4-5(OP1/W): 3.1Å; 4-6(OP1/W): 3.0Å; 5-6(W/W): 2.9Å;   
  
**2.07 ± 0.12: the average coodination distance of the 1^st^ six ligands is Mg^2+^ compatible**  
**8: Number of potential Water contacts**   
**1 : Number of second shell contacts involving water or ions**   
**12%: Percentage of satisfied second shell Water contacts**   
• Contact(s): /O_w_/O_w_/O_w_/O_w_/OP1/OP1/  
• Contact code: *cis-*2O_ph_4O_w_  
→ Slightly distorded coord. octahedron (6.1º) ←  
Go to ToC

### MG:a:3144

|  | Res. type | Atom | Residue | Distance | Occupancy | B-factor | Closest atom(s) to a 1^st^ shell water | EM\_dens. |
| --- | --- | --- | --- | --- | --- | --- | --- | --- |
| **Ion** | | | |  | 1.0 | 34.8 |  | 16.23 |
 1 | G | OP2 | a:1026 | 1.81 | 1.0 | 33.2 |  | 9.72 | 2 | G | OP1 | a:1028 | 1.95 | 1.0 | 38.7 |  | 7.32 | 3 | U | OP2 | a:1025 | 2.05 | 1.0 | 30.2 |  | 9.92 | 4 | G | O2' | a:1027 | 2.06 | 1.0 | 33.3 |  | 11.69 | 5 | Water 1 | | a:5265 | 2.17 | 1.0 | 30.7 |  | 11.04 | 6 | Water 2 | | a:6103 | 2.39 | 1.0 | 33.9 | 2.86 [C:OP2]a:1137 | 9.70 | 7 | G | O3' | a:1027 | 3.59 | 1.0 | 34.2 |  | 10.88 | 8 | Water 3 | | a:9758 | 3.86 | 1.0 | 46.2 |  | 4.20 | 9 | **K** | K | a:3133 | 3.90 | 1.0 | 30.0 |  | 26.17 | 10 | U | O5' | a:1025 | 4.00 | 1.0 | 33.0 |  | 10.56 | 11 | U | O3' | a:1025 | 4.05 | 1.0 | 35.9 |  | 9.78 | 12 | C | OP2 | a:1137 | 4.08 | 1.0 | 34.9 |  | 8.34 | 13 | G | OP1 | a:1026 | 4.10 | 1.0 | 32.5 |  | 11.18 | 14 | G | O5' | a:1026 | 4.12 | 1.0 | 34.6 |  | 11.17 | 15 | G | O5' | a:1028 | 4.13 | 1.0 | 39.2 |  | 6.15 | 16 | U | OP1 | a:1025 | 4.13 | 1.0 | 35.0 |  | 10.05 | 17 | Water 4 | | a:9222 | 4.16 | 1.0 | 39.1 |  | 5.10 | 18 | G | OP2 | a:1028 | 4.36 | 1.0 | 38.5 |  | 7.38 | 19 | G | O3' | a:1024 | 4.52 | 1.0 | 34.9 |  | 10.71 | 20 | G | N7 | a:1026 | 4.53 | 1.0 | 26.9 |  | 12.63 | 21 | Water 5 | | a:8955 | 4.71 | 1.0 | 39.2 | 2.99 [C:OP2]a:1137 | 6.87 |

  
**1^st^ shell coordination angles and distances (*trans* in "blue"; *cys* in "green"; deviations in "orange"):**   
• 1-2(OP2/OP1): *97º*; 1-3(OP2/OP2): *100º*; 1-4(OP2/O2'): *95º*; 1-5(OP2/W): 91º; 2-3(OP1/OP2): *161º*; 2-4(OP1/O2'): 89º; 2-5(OP1/W): *81º*; 3-4(OP2/O2'): *97º*; 3-5(OP2/W): 90º; 4-5(O2'/W): *169º*;   
• 1-2(OP2/OP1): 2.8Å; 1-3(OP2/OP2): 3.0Å; 1-4(OP2/O2'): 2.9Å; 1-5(OP2/W): 2.9Å; 2-3(OP1/OP2): 3.9Å; 2-4(OP1/O2'): 2.8Å; 2-5(OP1/W): 2.7Å; 3-4(OP2/O2'): 3.1Å; 3-5(OP2/W): 3.0Å; 4-5(O2'/W): *4.2Å*;   
  
**2.07 ± 0.18: the average coodination distance of the 1^st^ six ligands is Mg^2+^ compatible**  
**4: Number of potential Water contacts**   
**1 : Number of second shell contacts involving water or ions**   
**25%: Percentage of satisfied second shell Water contacts**   
• Contact(s): /O2'/O_w_/OP1/OP2/  
• Contact code: *trans-*2O_ph_O_r_O_w_  
• Contact(s) in the exclusion zone: /O_w_/OP2/  
→ Missing octahedral coordination for MG ←  
→ Slightly distorded coord. octahedron (6.9º) ←  
Go to ToC

### MG:a:3145

|  | Res. type | Atom | Residue | Distance | Occupancy | B-factor | Closest atom(s) to a 1^st^ shell water | EM\_dens. |
| --- | --- | --- | --- | --- | --- | --- | --- | --- |
| **Ion** | | | |  | 1.0 | 31.0 |  | 16.70 |
 1 | U | OP2 | a:1381 | 2.02 | 1.0 | 26.9 |  | 11.07 | 2 | Water 1 | | a:7078 | 2.02 | 1.0 | 28.9 |  | 10.64 | 3 | Water 2 | | a:7497 | 2.03 | 1.0 | 29.0 | 2.99 [G:N7]a:1382 | 11.55 | 4 | A | O2' | a:1380 | 2.05 | 1.0 | 26.4 |  | 13.19 | 5 | G | OP2 | a:1382 | 2.12 | 1.0 | 30.9 |  | 12.05 | 6 | Water 3 | | a:9682 | 2.37 | 1.0 | 31.4 |  | 10.23 | 7 | A | O3' | a:1380 | 3.66 | 1.0 | 32.0 |  | 8.50 | 8 | G | N7 | a:1382 | 3.72 | 1.0 | 23.2 |  | 14.07 | 9 | Water 4 | | a:4059 | 4.08 | 1.0 | 39.9 | 2.58 [G:OP1]a:1382 | 4.00 | 10 | Water 5 | | a:9339 | 4.16 | 1.0 | 33.9 |  | 8.01 | 11 | U | O5' | a:1381 | 4.25 | 1.0 | 33.1 |  | 9.96 | 12 | Water 6 | | a:7469 | 4.34 | 1.0 | 35.5 |  | 7.93 | 13 | Water 7 | | a:10222 | 4.37 | 1.0 | 46.0 |  | 3.81 | 14 | G | OP1 | a:1382 | 4.37 | 1.0 | 35.5 |  | 6.20 | 15 | U | OP1 | a:1381 | 4.40 | 1.0 | 31.3 |  | 9.06 | 16 | G | O5' | a:1382 | 4.43 | 1.0 | 33.7 |  | 9.07 | 17 | U | O3' | a:1381 | 4.47 | 1.0 | 36.2 |  | 6.99 | 18 | Water 8 | | a:7464 | 4.54 | 1.0 | 38.3 |  | 10.03 |

  
**1^st^ shell coordination angles and distances (*trans* in "blue"; *cys* in "green"; deviations in "orange"):**   
• 1-2(OP2/W): 88º; 1-3(OP2/W): 178º; 1-4(OP2/O2'): 88º; 1-5(OP2/OP2): 93º; 2-3(W/W): 91º; 2-4(W/O2'): 175º; 2-5(W/OP2): 93º; 3-4(W/O2'): 93º; 3-5(W/OP2): 86º; 4-5(O2'/OP2): *84º*;   
• 1-2(OP2/W): 2.8Å; 1-3(OP2/W): 4.0Å; 1-4(OP2/O2'): 2.8Å; 1-5(OP2/OP2): 3.0Å; 2-3(W/W): 2.9Å; 2-4(W/O2'): 4.1Å; 2-5(W/OP2): 3.0Å; 3-4(W/O2'): 3.0Å; 3-5(W/OP2): 2.8Å; 4-5(O2'/OP2): 2.8Å;   
  
**2.10 ± 0.12: the average coodination distance of the 1^st^ six ligands is Mg^2+^ compatible**  
**6: Number of potential Water contacts**   
**1 : Number of second shell contacts involving water or ions**   
**17%: Percentage of satisfied second shell Water contacts**   
• Contact(s): /O2'/O_w_/O_w_/OP2/OP2/  
• Contact code: *cis-*2O_ph_O_r_2O_w_  
• Contact(s) in the exclusion zone: /O_w_/  
→ Missing octahedral coordination for MG ←  
Ooctahedral coordination (2.8º)   
Go to ToC

### MG:a:3146

|  | Res. type | Atom | Residue | Distance | Occupancy | B-factor | Closest atom(s) to a 1^st^ shell water | EM\_dens. |
| --- | --- | --- | --- | --- | --- | --- | --- | --- |
| **Ion** | | | |  | 1.0 | 48.7 |  | 8.52 |
 1 | Water 1 | | p:264 | 1.91 | 1.0 | 37.1 |  | 7.15 | 2 | Water 2 | | a:4047 | 2.11 | 1.0 | 34.7 | 2.58 [A:OP1]a:515 | 8.50 | 3 | Water 3 | | a:10301 | 2.16 | 1.0 | 42.1 |  | 4.86 | 4 | Water 4 | | p:223 | 2.23 | 1.0 | 40.5 | 2.88 [SER:O]p:29 | 7.21 | 5 | Water 5 | | a:10046 | 2.32 | 1.0 | 37.2 |  | 8.83 | 6 | Water 6 | | p:256 | 2.52 | 1.0 | 41.3 | 2.64 [MG:MG]a:3055 | 8.58 | 7 | Water 7 | | a:7808 | 4.01 | 1.0 | 41.9 |  | 7.08 | 8 | A | OP1 | a:515 | 4.07 | 1.0 | 27.7 |  | 12.97 | 9 | Water 8 | | p:250 | 4.21 | 1.0 | 39.7 |  | 5.33 | 10 | Water 9 | | a:4325 | 4.27 | 1.0 | 31.2 | 2.61 [A:OP1]a:515 | 12.23 | 11 | Water 10 | | a:3822 | 4.36 | 1.0 | 40.6 |  | 7.93 | 12 | Water 11 | | p:248 | 4.38 | 1.0 | 43.2 |  | 3.87 | 13 | **MG** | MG | a:3055 | 4.40 | 1.0 | 41.0 |  | 13.28 | 14 | Water 12 | | a:5461 | 4.61 | 1.0 | 26.8 |  | 12.18 | 15 | Water 13 | | a:6727 | 4.71 | 1.0 | 43.4 |  | 5.13 | 16 | SER | O | p:29 | 4.77 | 1.0 | 31.4 |  | 13.90 |

  
**1^st^ shell coordination angles and distances (*trans* in "blue"; *cys* in "green"; deviations in "orange"):**   
• 1-2(W/W): *98º*; 1-3(W/W): 95º; 1-4(W/W): 87º; 2-3(W/W): *99º*; 2-4(W/W): 93º; 3-4(W/W): *168º*;   
• 1-2(W/W): 3.0Å; 1-3(W/W): 3.0Å; 1-4(W/W): 2.8Å; 2-3(W/W): *3.2Å*; 2-4(W/W): 3.2Å; 3-4(W/W): *4.4Å*;   
  
**2.21 ± 0.19: the average coodination distance of the 1^st^ six ligands is Mg^2+^ compatible**  
**12: Number of potential Water contacts**   
**3 : Number of second shell contacts involving water or ions**   
**25%: Percentage of satisfied second shell Water contacts**   
• Contact(s): /O_w_/O_w_/O_w_/O_w_/  
• Contact code: 4O_w_  
• Contact(s) in the exclusion zone: /O_w_/O_w_/  
→ Missing octahedral coordination for MG ←  
→ Slightly distorded coord. octahedron (6.7º) ←  
Go to ToC

### MG:a:3147

|  | Res. type | Atom | Residue | Distance | Occupancy | B-factor | Closest atom(s) to a 1^st^ shell water | EM\_dens. |
| --- | --- | --- | --- | --- | --- | --- | --- | --- |
| **Ion** | | | |  | 1.0 | 40.3 |  | 13.11 |
 1 | U | OP1 | a:1999 | 1.94 | 1.0 | 31.4 |  | 11.75 | 2 | Water 1 | | a:5138 | 2.01 | 1.0 | 37.2 |  | 4.81 | 3 | Water 2 | | a:8251 | 2.04 | 1.0 | 33.7 |  | 7.20 | 4 | Water 3 | | a:5558 | 2.06 | 1.0 | 38.1 |  | 7.50 | 5 | Water 4 | | a:8482 | 2.08 | 1.0 | 34.6 | 2.97 [THR:O]d:129 | 8.42 | 6 | Water 5 | | d:459 | 2.26 | 1.0 | 32.9 | 2.85 [THR:O]d:129 | 8.14 | 7 | C | O3' | a:1998 | 3.73 | 1.0 | 31.4 |  | 12.41 | 8 | U | OP2 | a:1999 | 4.00 | 1.0 | 27.6 |  | 13.30 | 9 | THR | O | d:129 | 4.22 | 1.0 | 30.6 |  | 12.62 | 10 | Water 6 | | a:8183 | 4.23 | 1.0 | 38.7 |  | 6.45 | 11 | Water 7 | | a:9715 | 4.26 | 1.0 | 43.8 |  | 5.05 | 12 | U | OP1 | a:2515 | 4.30 | 1.0 | 28.4 |  | 10.44 | 13 | Water 8 | | a:9237 | 4.38 | 1.0 | 34.7 |  | 9.25 | 14 | U | O5' | a:1999 | 4.39 | 1.0 | 29.1 |  | 12.86 | 15 | C | OP1 | a:2001 | 4.45 | 1.0 | 33.9 |  | 8.17 | 16 | Water 9 | | a:8879 | 4.46 | 1.0 | 30.5 |  | 10.69 | 17 | Water 10 | | a:6858 | 4.50 | 1.0 | 23.8 |  | 13.61 |

  
**1^st^ shell coordination angles and distances (*trans* in "blue"; *cys* in "green"; deviations in "orange"):**   
• 1-2(OP1/W): 86º; 1-3(OP1/W): 94º; 1-4(OP1/W): 85º; 1-5(OP1/W): 93º; 1-6(OP1/W): 172º; 2-3(W/W): *67º*; 2-4(W/W): *123º*; 2-5(W/W): *160º*; 2-6(W/W): *100º*; 3-4(W/W): *170º*; 3-5(W/W): 94º; 3-6(W/W): *84º*; 4-5(W/W): *76º*; 4-6(W/W): *96º*; 5-6(W/W): *79º*;   
• 1-2(OP1/W): 2.7Å; 1-3(OP1/W): 2.9Å; 1-4(OP1/W): 2.7Å; 1-5(OP1/W): 2.9Å; 1-6(OP1/W): 4.2Å; 2-3(W/W): 2.2Å; 2-4(W/W): *3.6Å*; 2-5(W/W): 4.0Å; 2-6(W/W): *3.3Å*; 3-4(W/W): 4.1Å; 3-5(W/W): 3.0Å; 3-6(W/W): 2.9Å; 4-5(W/W): 2.5Å; 4-6(W/W): *3.2Å*; 5-6(W/W): 2.8Å;   
  
**2.06 ± 0.10: the average coodination distance of the 1^st^ six ligands is Mg^2+^ compatible**  
**10: Number of potential Water contacts**   
**2 : Number of second shell contacts involving water or ions**   
**20%: Percentage of satisfied second shell Water contacts**   
• Contact(s): /O_w_/O_w_/O_w_/O_w_/O_w_/OP1/  
• Contact code: O_ph_5O_w_  
→→→ Distorded coord. octahedron (10.7º) ←←←  
Go to ToC

### MG:a:3148

|  | Res. type | Atom | Residue | Distance | Occupancy | B-factor | Closest atom(s) to a 1^st^ shell water | EM\_dens. |
| --- | --- | --- | --- | --- | --- | --- | --- | --- |
| **Ion** | | | |  | 1.0 | 46.8 |  | 5.77 |
 1 | U | OP1 | a:1623 | 2.44 | 1.0 | 41.1 |  | 6.83 | 2 | C | OP1 | a:739 | 2.81 | 1.0 | 37.8 |  | 8.16 | 3 | Water 1 | | a:9724 | 2.92 | 1.0 | 43.0 |  | 4.21 | 4 | Water 2 | | a:8377 | 3.09 | 1.0 | 43.4 | 2.92 [C:OP1]a:739 | 5.21 | 5 | G | OP1 | a:1624 | 3.19 | 1.0 | 41.2 |  | 9.18 | 6 | G | OP2 | a:1624 | 3.44 | 1.0 | 39.1 |  | 5.91 | 7 | Water 3 | | a:3578 | 3.45 | 1.0 | 41.4 | 2.66 [U:OP2]a:1623 // 2.97 [G:OP2]a:1624 | 6.35 | 8 | U | OP2 | a:1623 | 3.75 | 1.0 | 31.8 |  | 10.41 | 9 | Water 4 | | a:3444 | 3.76 | 1.0 | 41.0 | 2.39 [G:OP2]a:1624 | 4.26 | 10 | C | O3' | a:738 | 3.91 | 1.0 | 33.2 |  | 10.98 | 11 | C | OP2 | a:739 | 3.94 | 1.0 | 30.8 |  | 9.40 | 12 | U | O5' | a:1623 | 3.99 | 1.0 | 32.4 |  | 11.25 |

  
• • **2.98 ± 0.32: the average coodination distance of the 1^st^ six ligands is K^+^ compatible**  
**4: Number of potential Water contacts**   
**1 : Number of second shell contacts involving water or ions**   
**25%: Percentage of satisfied second shell Water contacts**   
• Contact code: → No contacts ←   
• Contact(s) in the exclusion zone: /O_w_/O_w_/OP1/OP1/OP1/  
→ Consider K in place of MG (2 valid contact(s) for K vs. 0 for MG ←)  
Go to ToC

### MG:a:3149

|  | Res. type | Atom | Residue | Distance | Occupancy | B-factor | Closest atom(s) to a 1^st^ shell water | EM\_dens. |
| --- | --- | --- | --- | --- | --- | --- | --- | --- |
| **Ion** | | | |  | 1.0 | 44.0 |  | 13.81 |
 1 | Water 1 | | a:4395 | 2.07 | 1.0 | 33.6 | 2.62 [C:OP1]a:734 | 9.71 | 2 | G | OP1 | a:762 | 2.09 | 1.0 | 27.7 |  | 11.49 | 3 | Water 2 | | a:4177 | 2.10 | 1.0 | 28.6 | 2.60 [A:OP2]a:763 | 10.30 | 4 | Water 3 | | a:8500 | 2.20 | 1.0 | 32.8 |  | 9.75 | 5 | Water 4 | | a:8481 | 2.34 | 1.0 | 35.1 |  | 8.30 | 6 | C | OP2 | a:734 | 3.80 | 1.0 | 31.4 |  | 10.47 | 7 | C | OP1 | a:733 | 3.90 | 1.0 | 27.8 |  | 12.09 | 8 | Water 5 | | a:9768 | 3.92 | 1.0 | 35.5 |  | 7.35 | 9 | C | OP1 | a:734 | 4.08 | 1.0 | 34.7 |  | 11.05 | 10 | Water 6 | | a:7814 | 4.17 | 1.0 | 33.9 |  | 7.51 | 11 | G | O5' | a:762 | 4.18 | 1.0 | 28.6 |  | 13.91 | 12 | G | OP2 | a:762 | 4.22 | 1.0 | 25.2 |  | 13.35 | 13 | Water 7 | | a:5732 | 4.24 | 1.0 | 37.8 | 2.73 [C:OP1]a:733 | 8.19 | 14 | Water 8 | | a:3425 | 4.36 | 1.0 | 36.5 | 2.77 [G:O3']a:761 | 8.10 | 15 | A | OP2 | a:763 | 4.43 | 1.0 | 23.5 |  | 14.39 | 16 | Water 9 | | a:9904 | 4.48 | 1.0 | 47.2 |  | 5.30 | 17 | G | O3' | a:761 | 4.53 | 1.0 | 27.1 |  | 12.19 | 18 | Water 10 | | a:6942 | 4.72 | 1.0 | 31.4 |  | 9.18 | 19 | Water 11 | | a:9807 | 4.74 | 1.0 | 40.1 |  | 4.41 | 20 | Water 12 | | a:4849 | 4.75 | 1.0 | 24.2 | 2.66 [A:OP2]a:763 | 15.40 |

  
**1^st^ shell coordination angles and distances (*trans* in "blue"; *cys* in "green"; deviations in "orange"):**   
• 1-2(W/OP1): 87º; 1-3(W/W): *105º*; 1-4(W/W): *167º*; 2-3(OP1/W): 92º; 2-4(OP1/W): 86º; 3-4(W/W): 86º;   
• 1-2(W/OP1): 2.9Å; 1-3(W/W): *3.3Å*; 1-4(W/W): *4.2Å*; 2-3(OP1/W): 3.0Å; 2-4(OP1/W): 2.9Å; 3-4(W/W): 2.9Å;   
  
**2.43 ± 0.62: the average coodination distance of the 1^st^ six ligands is Na^+^** compatible  
**8: Number of potential Water contacts**   
**2 : Number of second shell contacts involving water or ions**   
**25%: Percentage of satisfied second shell Water contacts**   
• Contact(s): /O_w_/O_w_/O_w_/OP1/  
• Contact code: O_ph_3O_w_  
• Contact(s) in the exclusion zone: /O_w_/  
→ Missing octahedral coordination for MG ←  
→ Slightly distorded coord. octahedron (6.9º) ←  
Go to ToC

### MG:a:3150

|  | Res. type | Atom | Residue | Distance | Occupancy | B-factor | Closest atom(s) to a 1^st^ shell water | EM\_dens. |
| --- | --- | --- | --- | --- | --- | --- | --- | --- |
| **Ion** | | | |  | 1.0 | 28.2 |  | 19.66 |
 1 | C | OP1 | a:789 | 2.00 | 1.0 | 22.9 |  | 14.54 | 2 | Water 1 | | a:7405 | 2.00 | 1.0 | 23.7 | 2.84 [A:OP1]a:791 | 16.01 | 3 | Water 2 | | a:5036 | 2.07 | 1.0 | 29.9 |  | 12.46 | 4 | U | OP2 | a:792 | 2.08 | 1.0 | 37.4 |  | 8.76 | 5 | Water 3 | | a:5152 | 2.12 | 1.0 | 24.0 | 2.68 [C:OP2]a:793 | 15.49 | 6 | Water 4 | | a:7759 | 2.30 | 1.0 | 28.9 |  | 12.95 | 7 | C | OP2 | a:789 | 3.57 | 1.0 | 23.3 |  | 16.06 | 8 | U | O5' | a:792 | 3.63 | 1.0 | 35.5 |  | 9.73 | 9 | C | O3' | a:788 | 4.13 | 1.0 | 24.1 |  | 15.30 | 10 | U | OP1 | a:792 | 4.33 | 1.0 | 34.1 |  | 8.83 | 11 | C | O5' | a:789 | 4.35 | 1.0 | 22.8 |  | 19.68 | 12 | Water 5 | | a:5825 | 4.41 | 1.0 | 40.5 | 2.73 [C:OP2]a:793 | 8.43 | 13 | A | O3' | a:791 | 4.45 | 1.0 | 35.6 |  | 8.58 | 14 | A | OP1 | a:791 | 4.48 | 1.0 | 30.4 |  | 13.03 | 15 | C | OP2 | a:793 | 4.49 | 1.0 | 35.6 |  | 9.40 | 16 | Water 6 | | a:10024 | 4.50 | 1.0 | 38.1 |  | 5.95 |

  
**1^st^ shell coordination angles and distances (*trans* in "blue"; *cys* in "green"; deviations in "orange"):**   
• 1-2(OP1/W): *96º*; 1-3(OP1/W): *82º*; 1-4(OP1/OP2): *163º*; 1-5(OP1/W): 92º; 1-6(OP1/W): *83º*; 2-3(W/W): 93º; 2-4(W/OP2): *98º*; 2-5(W/W): 93º; 2-6(W/W): 179º; 3-4(W/OP2): 87º; 3-5(W/W): 172º; 3-6(W/W): 87º; 4-5(OP2/W): *97º*; 4-6(OP2/W): *82º*; 5-6(W/W): 86º;   
• 1-2(OP1/W): 3.0Å; 1-3(OP1/W): 2.7Å; 1-4(OP1/OP2): 4.0Å; 1-5(OP1/W): 3.0Å; 1-6(OP1/W): 2.9Å; 2-3(W/W): 3.0Å; 2-4(W/OP2): 3.1Å; 2-5(W/W): 3.0Å; 2-6(W/W): *4.3Å*; 3-4(W/OP2): 2.9Å; 3-5(W/W): 4.2Å; 3-6(W/W): 3.0Å; 4-5(OP2/W): 3.1Å; 4-6(OP2/W): 2.9Å; 5-6(W/W): 3.0Å;   
  
**2.09 ± 0.10: the average coodination distance of the 1^st^ six ligands is Mg^2+^ compatible**  
**8: Number of potential Water contacts**   
**2 : Number of second shell contacts involving water or ions**   
**25%: Percentage of satisfied second shell Water contacts**   
• Contact(s): /O_w_/O_w_/O_w_/O_w_/OP1/OP2/  
• Contact code: *trans-*2O_ph_4O_w_  
→ Slightly distorded coord. octahedron (5.9º) ←  
Go to ToC

### MG:a:3151

|  | Res. type | Atom | Residue | Distance | Occupancy | B-factor | Closest atom(s) to a 1^st^ shell water | EM\_dens. |
| --- | --- | --- | --- | --- | --- | --- | --- | --- |
| **Ion** | | | |  | 1.0 | 51.1 |  | 3.98 |
 1 | Water 1 | | a:9623 | 1.97 | 1.0 | 44.5 |  | 3.53 | 2 | A | OP1 | a:701 | 2.11 | 1.0 | 34.3 |  | 8.46 | 3 | Water 2 | | a:9635 | 2.41 | 1.0 | 35.5 |  | 5.01 | 4 | Water 3 | | a:6130 | 3.27 | 1.0 | 45.9 | 2.75 [G:OP2]a:702 | 4.12 | 5 | A | O5' | a:701 | 4.02 | 1.0 | 30.8 |  | 11.56 | 6 | Water 4 | | a:4067 | 4.16 | 1.0 | 37.2 |  | 7.04 | 7 | C | O5' | a:700 | 4.40 | 1.0 | 29.6 |  | 11.61 | 8 | A | OP2 | a:701 | 4.47 | 1.0 | 32.6 |  | 8.76 | 9 | C | O3' | a:700 | 4.47 | 1.0 | 30.2 |  | 11.57 | 10 | U | O4' | a:1556 | 4.52 | 1.0 | 45.2 |  | 4.72 | 11 | G | OP2 | a:702 | 4.54 | 1.0 | 36.2 |  | 9.19 | 12 | C | OP2 | a:700 | 4.63 | 1.0 | 33.2 |  | 8.80 | 13 | C | OP1 | a:700 | 4.70 | 1.0 | 38.0 |  | 8.89 |

  
**1^st^ shell coordination angles and distances (*trans* in "blue"; *cys* in "green"; deviations in "orange"):**   
• 1-2(W/OP1): *106º*;   
• 1-2(W/OP1): *3.3Å*;   
  
**2.99 ± 0.88: the average coodination distance of the 1^st^ six ligands is K^+^ compatible**  
**8: Number of potential Water contacts**   
**1 : Number of second shell contacts involving water or ions**   
**12%: Percentage of satisfied second shell Water contacts**   
• Contact(s): /O_w_/OP1/  
• Contact code: O_ph_O_w_  
• Contact(s) in the exclusion zone: /O_w_/O_w_/  
→ Missing octahedral coordination for MG ←  
→2Fo-Fc density between 1.0 and 4.0 !   
Go to ToC

### MG:a:3152

|  | Res. type | Atom | Residue | Distance | Occupancy | B-factor | Closest atom(s) to a 1^st^ shell water | EM\_dens. |
| --- | --- | --- | --- | --- | --- | --- | --- | --- |
| **Ion** | | | |  | 1.0 | 33.4 |  | 14.41 |
 1 | G | OP1 | a:2596 | 2.01 | 1.0 | 26.6 |  | 13.05 | 2 | Water 1 | | a:8801 | 2.08 | 1.0 | 33.8 |  | 8.49 | 3 | G | OP1 | a:1972 | 2.16 | 1.0 | 27.0 |  | 10.42 | 4 | Water 2 | | a:9750 | 2.25 | 1.0 | 31.2 |  | 10.18 | 5 | Water 3 | | a:8752 | 2.32 | 1.0 | 29.8 | 2.99 [C:O3']a:1971 | 11.41 | 6 | Water 4 | | a:7254 | 2.37 | 1.0 | 33.2 | 2.82 [G:OP1]a:1977 | 11.00 | 7 | Water 5 | | a:6941 | 3.41 | 1.0 | 37.5 |  | 7.51 | 8 | C | O3' | a:2595 | 3.74 | 1.0 | 30.1 |  | 11.92 | 9 | C | O3' | a:1971 | 3.86 | 1.0 | 33.0 |  | 11.78 | 10 | G | OP2 | a:2596 | 4.12 | 1.0 | 29.8 |  | 13.83 | 11 | Water 6 | | a:3716 | 4.12 | 1.0 | 40.6 |  | 5.92 | 12 | Water 7 | | a:6682 | 4.13 | 1.0 | 36.1 | 2.78 [G:OP2]a:2596 | 9.54 | 13 | G | O5' | a:1972 | 4.24 | 1.0 | 27.8 |  | 14.37 | 14 | Water 8 | | a:4886 | 4.38 | 1.0 | 42.1 | 2.66 [G:OP1]a:1977 | 4.55 | 15 | G | O5' | a:2596 | 4.47 | 1.0 | 28.5 |  | 14.04 | 16 | Water 9 | | a:9371 | 4.48 | 1.0 | 32.2 |  | 8.63 | 17 | Water 10 | | a:10157 | 4.51 | 1.0 | 45.2 |  | 4.46 | 18 | Water 11 | | a:6964 | 4.53 | 1.0 | 36.2 |  | 9.22 | 19 | G | OP2 | a:1972 | 4.66 | 1.0 | 32.3 |  | 10.06 | 20 | G | OP1 | a:1977 | 4.73 | 1.0 | 33.7 |  | 11.45 | 21 | Water 12 | | a:8475 | 4.75 | 1.0 | 32.5 |  | 7.18 |

  
**1^st^ shell coordination angles and distances (*trans* in "blue"; *cys* in "green"; deviations in "orange"):**   
• 1-2(OP1/W): 93º; 1-3(OP1/OP1): 85º; 1-4(OP1/W): *100º*; 2-3(W/OP1): *97º*; 2-4(W/W): 95º; 3-4(OP1/W): *167º*;   
• 1-2(OP1/W): 3.0Å; 1-3(OP1/OP1): 2.8Å; 1-4(OP1/W): *3.3Å*; 2-3(W/OP1): 3.2Å; 2-4(W/W): 3.2Å; 3-4(OP1/W): *4.4Å*;   
  
**2.20 ± 0.13: the average coodination distance of the 1^st^ six ligands is Mg^2+^ compatible**  
**8: Number of potential Water contacts**   
**2 : Number of second shell contacts involving water or ions**   
**25%: Percentage of satisfied second shell Water contacts**   
• Contact(s): /O_w_/O_w_/OP1/OP1/  
• Contact code: *cis-*2O_ph_2O_w_  
• Contact(s) in the exclusion zone: /O_w_/O_w_/  
→ Missing octahedral coordination for MG ←  
→ Slightly distorded coord. octahedron (7.2º) ←  
Go to ToC

### MG:a:3153

|  | Res. type | Atom | Residue | Distance | Occupancy | B-factor | Closest atom(s) to a 1^st^ shell water | EM\_dens. |
| --- | --- | --- | --- | --- | --- | --- | --- | --- |
| **Ion** | | | |  | 1.0 | 29.6 |  | 21.76 |
 1 | Water 1 | | c:428 | 1.97 | 1.0 | 24.7 | 2.61 [ASN:OD1]c:239 | 14.18 | 2 | Water 2 | | a:8589 | 1.98 | 1.0 | 23.8 |  | 12.82 | 3 | U | O4 | a:2597 | 2.14 | 1.0 | 28.9 |  | 13.78 | 4 | Water 3 | | a:5649 | 2.19 | 1.0 | 25.5 | 2.72 [G:OP2]a:2596 | 14.68 | 5 | Water 4 | | a:9067 | 2.19 | 1.0 | 22.2 |  | 15.67 | 6 | Water 5 | | a:5032 | 2.28 | 1.0 | 28.0 | 2.68 [C:OP2]a:2595 // 2.96 [ARG:O]c:238 | 13.13 | 7 | GLU | O | c:236 | 4.05 | 1.0 | 41.6 |  | 9.75 | 8 | ASN | OD1 | c:239 | 4.09 | 1.0 | 35.2 |  | 10.90 | 9 | G | N7 | a:2596 | 4.15 | 1.0 | 22.4 |  | 15.53 | 10 | Water 6 | | a:9777 | 4.26 | 1.0 | 35.9 |  | 8.43 | 11 | G | O6 | a:2596 | 4.29 | 1.0 | 30.4 |  | 15.11 | 12 | Water 7 | | a:4481 | 4.50 | 1.0 | 48.6 |  | 4.40 | 13 | C | OP2 | a:2595 | 4.51 | 1.0 | 26.6 |  | 13.65 | 14 | G | OP2 | a:2596 | 4.59 | 1.0 | 29.8 |  | 13.83 | 15 | Water 8 | | a:7664 | 4.60 | 1.0 | 36.2 | 2.85 [G:OP2]a:2596 | 6.19 | 16 | ARG | O | c:238 | 4.64 | 1.0 | 35.4 |  | 12.55 |

  
**1^st^ shell coordination angles and distances (*trans* in "blue"; *cys* in "green"; deviations in "orange"):**   
• 1-2(W/W): 94º; 1-3(W/O4): 93º; 1-4(W/W): 172º; 1-5(W/W): 92º; 1-6(W/W): *85º*; 2-3(W/O4): 91º; 2-4(W/W): 88º; 2-5(W/W): 174º; 2-6(W/W): 90º; 3-4(O4/W): *95º*; 3-5(O4/W): 88º; 3-6(O4/W): 177º; 4-5(W/W): 85º; 4-6(W/W): 87º; 5-6(W/W): 91º;   
• 1-2(W/W): 2.9Å; 1-3(W/O4): 3.0Å; 1-4(W/W): 4.1Å; 1-5(W/W): 3.0Å; 1-6(W/W): 2.9Å; 2-3(W/O4): 2.9Å; 2-4(W/W): 2.9Å; 2-5(W/W): 4.2Å; 2-6(W/W): 3.0Å; 3-4(O4/W): 3.2Å; 3-5(O4/W): 3.0Å; 3-6(O4/W): *4.4Å*; 4-5(W/W): 3.0Å; 4-6(W/W): 3.1Å; 5-6(W/W): 3.2Å;   
  
**2.12 ± 0.11: the average coodination distance of the 1^st^ six ligands is Mg^2+^ compatible**  
**10: Number of potential Water contacts**   
**4 : Number of second shell contacts involving water or ions**   
**40%: Percentage of satisfied second shell Water contacts**   
• Contact(s): /O4/O_w_/O_w_/O_w_/O_w_/O_w_/  
• Contact code: O_b_5O_w_  
Ooctahedral coordination (3.4º)   
Go to ToC

### MG:a:3154

|  | Res. type | Atom | Residue | Distance | Occupancy | B-factor | Closest atom(s) to a 1^st^ shell water | EM\_dens. |
| --- | --- | --- | --- | --- | --- | --- | --- | --- |
| **Ion** | | | |  | 1.0 | 41.7 |  | 11.81 |
 1 | G | OP1 | a:2547 | 1.90 | 1.0 | 29.6 |  | 10.69 | 2 | Water 1 | | a:3915 | 1.93 | 1.0 | 36.0 | 2.85 [A:O2']a:2770 | 8.68 | 3 | Water 2 | | a:8501 | 2.05 | 1.0 | 36.2 |  | 7.38 | 4 | Water 3 | | a:7367 | 2.09 | 1.0 | 39.0 |  | 7.03 | 5 | Water 4 | | a:10151 | 2.26 | 1.0 | 45.2 |  | 5.54 | 6 | Water 5 | | a:7937 | 2.27 | 1.0 | 35.6 | 2.88 [C:OP1]a:2771 | 7.07 | 7 | G | OP2 | a:2547 | 3.94 | 1.0 | 31.9 |  | 8.54 | 8 | A | O3' | a:2546 | 4.19 | 1.0 | 29.5 |  | 12.15 | 9 | Water 6 | | a:4747 | 4.20 | 1.0 | 43.0 |  | 6.93 | 10 | G | O5' | a:2547 | 4.30 | 1.0 | 30.3 |  | 13.11 | 11 | Water 7 | | a:10191 | 4.39 | 1.0 | 38.6 |  | 3.53 | 12 | Water 8 | | i:232 | 4.54 | 1.0 | 41.1 |  | 5.45 | 13 | A | O2' | a:2770 | 4.57 | 1.0 | 33.8 |  | 9.32 | 14 | C | OP1 | a:2771 | 4.58 | 1.0 | 38.2 |  | 8.94 | 15 | Water 9 | | i:235 | 4.70 | 1.0 | 43.1 |  | 4.31 | 16 | A | O3' | a:2770 | 4.72 | 1.0 | 38.3 |  | 9.39 |

  
**1^st^ shell coordination angles and distances (*trans* in "blue"; *cys* in "green"; deviations in "orange"):**   
• 1-2(OP1/W): *84º*; 1-3(OP1/W): *96º*; 1-4(OP1/W): 90º; 1-5(OP1/W): *168º*; 1-6(OP1/W): 91º; 2-3(W/W): 94º; 2-4(W/W): 173º; 2-5(W/W): *96º*; 2-6(W/W): 89º; 3-4(W/W): 90º; 3-5(W/W): *96º*; 3-6(W/W): 173º; 4-5(W/W): 89º; 4-6(W/W): 87º; 5-6(W/W): *77º*;   
• 1-2(OP1/W): 2.6Å; 1-3(OP1/W): 2.9Å; 1-4(OP1/W): 2.8Å; 1-5(OP1/W): 4.1Å; 1-6(OP1/W): 3.0Å; 2-3(W/W): 2.9Å; 2-4(W/W): 4.0Å; 2-5(W/W): 3.1Å; 2-6(W/W): 3.0Å; 3-4(W/W): 2.9Å; 3-5(W/W): *3.2Å*; 3-6(W/W): *4.3Å*; 4-5(W/W): 3.0Å; 4-6(W/W): 3.0Å; 5-6(W/W): 2.8Å;   
  
**2.08 ± 0.14: the average coodination distance of the 1^st^ six ligands is Mg^2+^ compatible**  
**10: Number of potential Water contacts**   
**2 : Number of second shell contacts involving water or ions**   
**20%: Percentage of satisfied second shell Water contacts**   
• Contact(s): /O_w_/O_w_/O_w_/O_w_/O_w_/OP1/  
• Contact code: O_ph_5O_w_  
Ooctahedral coordination (4.9º)   
Go to ToC

### MG:a:3155

|  | Res. type | Atom | Residue | Distance | Occupancy | B-factor | Closest atom(s) to a 1^st^ shell water | EM\_dens. |
| --- | --- | --- | --- | --- | --- | --- | --- | --- |
| **Ion** | | | |  | 1.0 | 52.6 |  | 6.99 |
 1 | Water 1 | | a:9145 | 2.00 | 1.0 | 40.8 |  | 5.39 | 2 | Water 2 | | a:5598 | 2.01 | 1.0 | 34.8 | 2.72 [G:N7]a:2648 // 2.81 [K:K]a:3246 | 8.75 | 3 | G | OP2 | a:2647 | 2.14 | 1.0 | 31.9 |  | 12.47 | 4 | Water 3 | | a:9663 | 2.25 | 1.0 | 43.2 |  | 3.93 | 5 | **K** | K | a:3246 | 2.42 | 1.0 | 30.0 |  | 12.67 | 6 | Water 4 | | a:5167 | 2.59 | 1.0 | 36.4 | 2.69 [G:N7]a:2647 | 6.45 | 7 | Water 5 | | a:4443 | 4.04 | 1.0 | 43.1 |  | 5.24 | 8 | G | O5' | a:2647 | 4.09 | 1.0 | 34.0 |  | 10.45 | 9 | G | O6 | a:2649 | 4.23 | 1.0 | 37.2 |  | 6.33 | 10 | Water 6 | | a:7201 | 4.27 | 1.0 | 38.8 | 2.90 [G:O6]a:2649 | 5.50 | 11 | G | N7 | a:2647 | 4.35 | 1.0 | 26.7 |  | 12.55 | 12 | G | OP1 | a:2647 | 4.38 | 1.0 | 37.2 |  | 7.76 | 13 | Water 7 | | a:8881 | 4.38 | 1.0 | 38.9 | 2.97 [G:OP1]a:2647 | 6.13 | 14 | Water 8 | | a:5537 | 4.48 | 1.0 | 42.9 |  | 6.35 | 15 | G | N7 | a:2648 | 4.48 | 1.0 | 22.5 |  | 12.49 | 16 | G | O6 | a:2648 | 4.56 | 1.0 | 33.5 |  | 12.33 | 17 | G | O3' | a:2646 | 4.60 | 1.0 | 31.8 |  | 10.68 |

  
**1^st^ shell coordination angles and distances (*trans* in "blue"; *cys* in "green"; deviations in "orange"):**   
• 1-2(W/W): 174º; 1-3(W/OP2): 94º; 1-4(W/W): 85º; 2-3(W/OP2): 91º; 2-4(W/W): *98º*; 3-4(OP2/W): 93º;   
• 1-2(W/W): 4.0Å; 1-3(W/OP2): 3.0Å; 1-4(W/W): 2.9Å; 2-3(W/OP2): 3.0Å; 2-4(W/W): *3.2Å*; 3-4(OP2/W): 3.2Å;   
  
**2.24 ± 0.21: the average coodination distance of the 1^st^ six ligands is Mg^2+^ compatible**  
**8: Number of potential Water contacts**   
**3 : Number of second shell contacts involving water or ions**   
**38%: Percentage of satisfied second shell Water contacts**   
• Contact(s): /O_w_/O_w_/O_w_/OP2/  
• Contact code: O_ph_3O_w_  
• Contact(s) in the exclusion zone: /O_w_/  
→ Missing octahedral coordination for MG ←  
Ooctahedral coordination (4.5º)   
Go to ToC

### MG:a:3156

|  | Res. type | Atom | Residue | Distance | Occupancy | B-factor | Closest atom(s) to a 1^st^ shell water | EM\_dens. |
| --- | --- | --- | --- | --- | --- | --- | --- | --- |
| **Ion** | | | |  | 1.0 | 41.7 |  | 10.51 |
 1 | A | OP1 | a:1239 | 1.94 | 1.0 | 36.5 |  | 8.24 | 2 | Water 1 | | a:6783 | 1.98 | 1.0 | 38.7 | 2.79 [A:OP2]a:1239 | 6.66 | 3 | Water 2 | | a:8212 | 2.01 | 1.0 | 36.5 |  | 6.17 | 4 | Water 3 | | a:8726 | 2.07 | 1.0 | 41.9 |  | 6.99 | 5 | Water 4 | | a:7912 | 2.07 | 1.0 | 35.5 |  | 7.65 | 6 | Water 5 | | a:6598 | 2.28 | 1.0 | 40.3 | 2.78 [A:OP2]a:477 | 5.66 | 7 | A | OP2 | a:1239 | 3.57 | 1.0 | 35.2 |  | 6.95 | 8 | Water 6 | | a:6045 | 4.16 | 1.0 | 41.3 | 2.74 [A:OP1]a:477 | 6.29 | 9 | Water 7 | | a:3415 | 4.20 | 1.0 | 40.5 | 2.31 [G:OP1]a:476 | 5.24 | 10 | G | O3' | a:1238 | 4.22 | 1.0 | 35.8 |  | 9.34 | 11 | A | O5' | a:1239 | 4.22 | 1.0 | 34.5 |  | 8.53 | 12 | A | OP2 | a:477 | 4.25 | 1.0 | 42.1 |  | 7.05 | 13 | Water 8 | | a:9959 | 4.31 | 1.0 | 41.4 |  | 4.19 | 14 | G | OP1 | a:476 | 4.34 | 1.0 | 37.6 |  | 6.65 | 15 | Water 9 | | a:7034 | 4.47 | 1.0 | 37.7 |  | 6.56 | 16 | A | OP1 | a:477 | 4.77 | 1.0 | 33.3 |  | 8.42 |

  
**1^st^ shell coordination angles and distances (*trans* in "blue"; *cys* in "green"; deviations in "orange"):**   
• 1-2(OP1/W): 93º; 1-3(OP1/W): 94º; 1-4(OP1/W): 95º; 1-5(OP1/W): 91º; 1-6(OP1/W): 179º; 2-3(W/W): 172º; 2-4(W/W): 93º; 2-5(W/W): 87º; 2-6(W/W): 88º; 3-4(W/W): *85º*; 3-5(W/W): 95º; 3-6(W/W): *84º*; 4-5(W/W): 174º; 4-6(W/W): *85º*; 5-6(W/W): 89º;   
• 1-2(OP1/W): 2.8Å; 1-3(OP1/W): 2.9Å; 1-4(OP1/W): 3.0Å; 1-5(OP1/W): 2.9Å; 1-6(OP1/W): *4.2Å*; 2-3(W/W): 4.0Å; 2-4(W/W): 2.9Å; 2-5(W/W): 2.8Å; 2-6(W/W): 3.0Å; 3-4(W/W): 2.8Å; 3-5(W/W): 3.0Å; 3-6(W/W): 2.9Å; 4-5(W/W): 4.1Å; 4-6(W/W): 2.9Å; 5-6(W/W): 3.1Å;   
  
**2.06 ± 0.11: the average coodination distance of the 1^st^ six ligands is Mg^2+^ compatible**  
**10: Number of potential Water contacts**   
**2 : Number of second shell contacts involving water or ions**   
**20%: Percentage of satisfied second shell Water contacts**   
• Contact(s): /O_w_/O_w_/O_w_/O_w_/O_w_/OP1/  
• Contact code: O_ph_5O_w_  
Ooctahedral coordination (3.9º)   
Go to ToC

### MG:a:3157

|  | Res. type | Atom | Residue | Distance | Occupancy | B-factor | Closest atom(s) to a 1^st^ shell water | EM\_dens. |
| --- | --- | --- | --- | --- | --- | --- | --- | --- |
| **Ion** | | | |  | 1.0 | 41.5 |  | 16.19 |
 1 | Water 1 | | a:4457 | 1.99 | 1.0 | 29.6 | 2.63 [U:O4]a:18 | 10.74 | 2 | Water 2 | | a:6123 | 2.01 | 1.0 | 29.0 | 2.75 [G:O6]a:17 | 11.09 | 3 | Water 3 | | a:6848 | 2.05 | 1.0 | 31.6 | 2.80 [G:N7]a:17 | 10.90 | 4 | Water 4 | | a:4891 | 2.18 | 1.0 | 31.4 | 2.67 [U:O4]a:521 | 9.38 | 5 | Water 5 | | a:10284 | 2.29 | 1.0 | 34.7 |  | 8.93 | 6 | Water 6 | | a:10246 | 2.38 | 1.0 | 33.2 |  | 8.72 | 7 | Water 7 | | a:6094 | 4.06 | 1.0 | 35.1 |  | 9.23 | 8 | Water 8 | | a:5909 | 4.08 | 1.0 | 33.9 |  | 8.29 | 9 | U | O4 | a:18 | 4.11 | 1.0 | 25.3 |  | 14.27 | 10 | Water 9 | | a:7338 | 4.24 | 1.0 | 39.3 |  | 7.80 | 11 | G | N7 | a:17 | 4.25 | 1.0 | 21.3 |  | 15.22 | 12 | G | O6 | a:17 | 4.33 | 1.0 | 24.3 |  | 15.20 | 13 | U | O4 | a:521 | 4.36 | 1.0 | 30.5 |  | 10.56 | 14 | Water 10 | | a:10245 | 4.37 | 1.0 | 43.2 |  | 5.85 | 15 | Water 11 | | a:8918 | 4.43 | 1.0 | 40.0 |  | 6.24 |

  
**1^st^ shell coordination angles and distances (*trans* in "blue"; *cys* in "green"; deviations in "orange"):**   
• 1-2(W/W): *95º*; 1-3(W/W): 89º; 1-4(W/W): 94º; 1-5(W/W): *97º*; 2-3(W/W): 87º; 2-4(W/W): 89º; 2-5(W/W): *167º*; 3-4(W/W): 175º; 3-5(W/W): 91º; 4-5(W/W): 92º;   
• 1-2(W/W): 3.0Å; 1-3(W/W): 2.8Å; 1-4(W/W): 3.0Å; 1-5(W/W): *3.2Å*; 2-3(W/W): 2.8Å; 2-4(W/W): 2.9Å; 2-5(W/W): *4.3Å*; 3-4(W/W): *4.2Å*; 3-5(W/W): 3.1Å; 4-5(W/W): *3.2Å*;   
  
**2.15 ± 0.15: the average coodination distance of the 1^st^ six ligands is Mg^2+^ compatible**  
**12: Number of potential Water contacts**   
**4 : Number of second shell contacts involving water or ions**   
**33%: Percentage of satisfied second shell Water contacts**   
• Contact(s): /O_w_/O_w_/O_w_/O_w_/O_w_/  
• Contact code: 5O_w_  
• Contact(s) in the exclusion zone: /O_w_/  
→ Missing octahedral coordination for MG ←  
Ooctahedral coordination (4.3º)   
Go to ToC

### MG:a:3158

|  | Res. type | Atom | Residue | Distance | Occupancy | B-factor | Closest atom(s) to a 1^st^ shell water | EM\_dens. |
| --- | --- | --- | --- | --- | --- | --- | --- | --- |
| **Ion** | | | |  | 1.0 | 44.2 |  | 7.82 |
 1 | Water 1 | | a:5224 | 1.99 | 1.0 | 37.8 | 2.69 [G:O6]a:24 | 6.24 | 2 | Water 2 | | a:9105 | 2.10 | 1.0 | 43.8 |  | 3.51 | 3 | Water 3 | | a:8984 | 2.44 | 1.0 | 37.3 | 2.92 [K:K]a:3250 | 6.59 | 4 | G | O6 | a:24 | 2.79 | 1.0 | 26.3 |  | 13.58 | 5 | U | O4 | a:25 | 2.93 | 1.0 | 26.3 |  | 13.10 | 6 | **K** | K | a:3250 | 3.31 | 1.0 | 30.0 |  | 13.51 | 7 | A | OP2 | a:514 | 3.55 | 1.0 | 34.9 |  | 11.65 | 8 | Water 4 | | a:7165 | 4.37 | 1.0 | 35.4 | 2.82 [U:O4]a:25 | 6.32 | 9 | A | OP2 | a:513 | 4.54 | 1.0 | 31.7 |  | 10.67 | 10 | Water 5 | | a:6727 | 4.66 | 1.0 | 43.4 |  | 5.13 | 11 | G | N7 | a:24 | 4.68 | 1.0 | 20.9 |  | 14.66 |

  
**1^st^ shell coordination angles and distances (*trans* in "blue"; *cys* in "green"; deviations in "orange"):**   
• 1-2(W/W): 91º;   
• 1-2(W/W): 2.9Å;   
  
**2.59 ± 0.47: the average coodination distance of the 1^st^ six ligands is Na^+^** compatible  
**6: Number of potential Water contacts**   
**2 : Number of second shell contacts involving water or ions**   
**33%: Percentage of satisfied second shell Water contacts**   
• Contact(s): /O_w_/O_w_/  
• Contact code: 2O_w_  
• Contact(s) in the exclusion zone: /O4/O6/O_w_/  
→ Consider K in place of MG (2 valid contact(s) for K vs. 0 for MG ←)  
→ Missing octahedral coordination for MG ←  
Go to ToC

### MG:a:3159

|  | Res. type | Atom | Residue | Distance | Occupancy | B-factor | Closest atom(s) to a 1^st^ shell water | EM\_dens. |
| --- | --- | --- | --- | --- | --- | --- | --- | --- |
| **Ion** | | | |  | 1.0 | 44.9 |  | 11.28 |
 1 | Water 1 | | a:7189 | 1.84 | 1.0 | 28.4 |  | 10.78 | 2 | Water 2 | | a:4772 | 2.00 | 1.0 | 33.5 | 2.66 [A:OP2]a:2075 | 9.69 | 3 | U | OP2 | a:2247 | 2.22 | 1.0 | 27.0 |  | 11.36 | 4 | A | OP1 | a:2075 | 2.26 | 1.0 | 28.3 |  | 11.63 | 5 | Water 3 | | a:7142 | 2.27 | 1.0 | 39.0 | 2.81 [U:OP1]a:2247 | 5.12 | 6 | A | OP2 | a:2075 | 3.42 | 1.0 | 28.0 |  | 13.48 | 7 | U | OP1 | a:2247 | 3.56 | 1.0 | 23.5 |  | 14.54 | 8 | Water 4 | | a:4474 | 3.71 | 1.0 | 36.1 |  | 8.65 | 9 | Water 5 | | a:5238 | 3.73 | 1.0 | 34.8 |  | 6.44 | 10 | Water 6 | | a:5120 | 4.16 | 1.0 | 40.5 |  | 5.51 | 11 | A | O5' | a:2075 | 4.26 | 1.0 | 28.9 |  | 13.86 | 12 | Water 7 | | a:5714 | 4.33 | 1.0 | 45.8 | 2.72 [A:OP2]a:2075 | 4.93 | 13 | U | O5' | a:2247 | 4.37 | 1.0 | 23.9 |  | 14.58 | 14 | Water 8 | | a:7932 | 4.44 | 1.0 | 34.5 |  | 8.29 | 15 | G | O3' | a:2246 | 4.47 | 1.0 | 25.0 |  | 16.15 | 16 | A | O3' | a:2074 | 4.47 | 1.0 | 29.6 |  | 12.56 | 17 | **MG** | MG | a:3068 | 4.61 | 1.0 | 40.5 |  | 11.77 | 18 | Water 9 | | a:8219 | 4.64 | 1.0 | 39.8 |  | 7.52 |

  
**1^st^ shell coordination angles and distances (*trans* in "blue"; *cys* in "green"; deviations in "orange"):**   
• 1-2(W/W): *159º*; 1-3(W/OP2): 87º; 1-4(W/OP1): 87º; 1-5(W/W): *109º*; 2-3(W/OP2): 90º; 2-4(W/OP1): 95º; 2-5(W/W): 92º; 3-4(OP2/OP1): 174º; 3-5(OP2/W): *96º*; 4-5(OP1/W): 87º;   
• 1-2(W/W): *3.8Å*; 1-3(W/OP2): 2.8Å; 1-4(W/OP1): 2.8Å; 1-5(W/W): *3.3Å*; 2-3(W/OP2): 3.0Å; 2-4(W/OP1): 3.1Å; 2-5(W/W): 3.1Å; 3-4(OP2/OP1): *4.5Å*; 3-5(OP2/W): *3.3Å*; 4-5(OP1/W): 3.1Å;   
  
**2.33 ± 0.51: the average coodination distance of the 1^st^ six ligands is Na^+^** compatible  
**6: Number of potential Water contacts**   
**2 : Number of second shell contacts involving water or ions**   
**33%: Percentage of satisfied second shell Water contacts**   
• Contact(s): /O_w_/O_w_/OP1/OP2/  
• Contact code: *cis-*2O_ph_2O_w_  
• Contact(s) in the exclusion zone: /O_w_/  
→ Missing octahedral coordination for MG ←  
→ Slightly distorded coord. octahedron (6.7º) ←  
Go to ToC

### MG:a:3160

|  | Res. type | Atom | Residue | Distance | Occupancy | B-factor | Closest atom(s) to a 1^st^ shell water | EM\_dens. |
| --- | --- | --- | --- | --- | --- | --- | --- | --- |
| **Ion** | | | |  | 1.0 | 49.6 |  | 7.86 |
 1 | Water 1 | | a:8978 | 1.97 | 1.0 | 41.3 | 2.99 [A:OP2]a:2074 | 5.56 | 2 | Water 2 | | a:9622 | 2.05 | 1.0 | 45.3 |  | 5.83 | 3 | Water 3 | | a:7609 | 2.22 | 1.0 | 39.3 | 2.85 [U:O4]a:2249 | 4.31 | 4 | Water 4 | | a:4569 | 2.39 | 1.0 | 40.8 | 2.64 [A:N7]a:2075 // 2.77 [A:OP2]a:2075 | 7.72 | 5 | Water 5 | | a:5714 | 3.77 | 1.0 | 45.8 | 2.72 [A:OP2]a:2075 | 4.93 | 6 | **K** | K | a:3115 | 4.31 | 1.0 | 30.0 |  | 16.92 | 7 | U | OP2 | a:2248 | 4.31 | 1.0 | 29.9 |  | 11.64 | 8 | A | N7 | a:2075 | 4.39 | 1.0 | 20.3 |  | 19.08 | 9 | U | O4 | a:2249 | 4.42 | 1.0 | 27.8 |  | 13.52 | 10 | Water 6 | | a:3719 | 4.50 | 1.0 | 37.2 |  | 6.99 | 11 | Water 7 | | a:8111 | 4.55 | 1.0 | 43.5 | 2.89 [U:OP2]a:2248 | 4.81 | 12 | A | OP2 | a:2075 | 4.71 | 1.0 | 28.0 |  | 13.48 | 13 | A | OP2 | a:2074 | 4.72 | 1.0 | 33.5 |  | 8.09 | 14 | Water 8 | | a:8219 | 4.79 | 1.0 | 39.8 |  | 7.52 |

  
**1^st^ shell coordination angles and distances (*trans* in "blue"; *cys* in "green"; deviations in "orange"):**   
• 1-2(W/W): 94º; 1-3(W/W): *84º*; 2-3(W/W): 92º;   
• 1-2(W/W): 2.9Å; 1-3(W/W): 2.8Å; 2-3(W/W): 3.1Å;   
  
**2.78 ± 0.91: the average coodination distance of the 1^st^ six ligands is K^+^ compatible**  
**10: Number of potential Water contacts**   
**5 : Number of second shell contacts involving water or ions**   
**50%: Percentage of satisfied second shell Water contacts**   
• Contact(s): /O_w_/O_w_/O_w_/  
• Contact code: 3O_w_  
• Contact(s) in the exclusion zone: /O_w_/  
→ Missing octahedral coordination for MG ←  
Go to ToC

### MG:a:3161

|  | Res. type | Atom | Residue | Distance | Occupancy | B-factor | Closest atom(s) to a 1^st^ shell water | EM\_dens. |
| --- | --- | --- | --- | --- | --- | --- | --- | --- |
| **Ion** | | | |  | 1.0 | 46.7 |  | 6.99 |
 1 | Water 1 | | a:8200 | 1.93 | 1.0 | 40.0 |  | 5.49 | 2 | Water 2 | | a:7378 | 1.97 | 1.0 | 40.5 | 2.83 [G:OP1]a:338 | 5.61 | 3 | Water 3 | | a:7228 | 1.98 | 1.0 | 43.7 | 2.82 [G:N7]a:298 | 5.28 | 4 | Water 4 | | a:6200 | 1.98 | 1.0 | 42.2 | 2.75 [U:OP2]a:296 | 4.84 | 5 | G | OP2 | a:297 | 2.04 | 1.0 | 42.6 |  | 5.75 | 6 | Water 5 | | a:8444 | 2.11 | 1.0 | 46.6 | 2.92 [G:N7]a:297 | 4.83 | 7 | G | O5' | a:297 | 3.86 | 1.0 | 47.2 |  | 4.74 | 8 | Water 6 | | a:4718 | 4.11 | 1.0 | 44.6 | 2.65 [G:OP1]a:297 | 3.56 | 9 | G | OP1 | a:338 | 4.22 | 1.0 | 47.0 |  | 3.50 | 10 | G | OP1 | a:297 | 4.25 | 1.0 | 48.7 |  | 4.82 | 11 | G | N7 | a:298 | 4.30 | 1.0 | 40.2 |  | 6.00 | 12 | U | O3' | a:296 | 4.49 | 1.0 | 44.7 |  | 5.50 | 13 | U | OP2 | a:296 | 4.54 | 1.0 | 44.5 |  | 4.89 | 14 | G | N7 | a:297 | 4.60 | 1.0 | 40.3 |  | 6.67 | 15 | G | O6 | a:298 | 4.63 | 1.0 | 44.1 |  | 5.11 |

  
**1^st^ shell coordination angles and distances (*trans* in "blue"; *cys* in "green"; deviations in "orange"):**   
• 1-2(W/W): 89º; 1-3(W/W): 89º; 1-4(W/W): *97º*; 1-5(W/OP2): 94º; 1-6(W/W): *169º*; 2-3(W/W): *85º*; 2-4(W/W): 90º; 2-5(W/OP2): 175º; 2-6(W/W): *82º*; 3-4(W/W): 172º; 3-5(W/OP2): 92º; 3-6(W/W): *84º*; 4-5(W/OP2): 93º; 4-6(W/W): 89º; 5-6(OP2/W): *95º*;   
• 1-2(W/W): 2.7Å; 1-3(W/W): 2.7Å; 1-4(W/W): 2.9Å; 1-5(W/OP2): 2.9Å; 1-6(W/W): 4.0Å; 2-3(W/W): 2.7Å; 2-4(W/W): 2.8Å; 2-5(W/OP2): 4.0Å; 2-6(W/W): 2.7Å; 3-4(W/W): 3.9Å; 3-5(W/OP2): 2.9Å; 3-6(W/W): 2.7Å; 4-5(W/OP2): 2.9Å; 4-6(W/W): 2.9Å; 5-6(OP2/W): 3.1Å;   
  
**2.00 ± 0.06: the average coodination distance of the 1^st^ six ligands is Mg^2+^ compatible**  
**10: Number of potential Water contacts**   
**4 : Number of second shell contacts involving water or ions**   
**40%: Percentage of satisfied second shell Water contacts**   
• Contact(s): /O_w_/O_w_/O_w_/O_w_/O_w_/OP2/  
• Contact code: O_ph_5O_w_  
Ooctahedral coordination (4.5º)   
Go to ToC

### MG:a:3162

|  | Res. type | Atom | Residue | Distance | Occupancy | B-factor | Closest atom(s) to a 1^st^ shell water | EM\_dens. |
| --- | --- | --- | --- | --- | --- | --- | --- | --- |
| **Ion** | | | |  | 1.0 | 37.6 |  | 9.21 |
 1 | G | O6 | a:313 | 2.68 | 1.0 | 46.3 |  | 5.06 | 2 | U | O4 | a:304 | 2.84 | 1.0 | 49.4 |  | 4.22 | 3 | G | O6 | a:312 | 3.09 | 1.0 | 45.0 |  | 6.13 | 4 | G | O6 | a:303 | 3.24 | 1.0 | 48.8 |  | 4.11 | 5 | U | N3 | a:304 | 3.76 | 1.0 | 41.8 |  | 5.23 | 6 | C | N4 | a:305 | 4.00 | 1.0 | 39.8 |  | 6.18 | 7 | A | O2' | a:310 | 4.07 | 1.0 | 44.5 |  | 3.82 | 8 | G | N7 | a:313 | 4.15 | 1.0 | 45.6 |  | 5.31 | 9 | G | N7 | a:312 | 4.30 | 1.0 | 34.0 |  | 7.80 |

  
• • **3.27 ± 0.47: the average coodination distance of the 1^st^ six ligands is K^+^ compatible**  
**0: Number of potential Water contacts**   
• Contact code: → No contacts ←   
• Contact(s) in the exclusion zone: /O4/O6/O6/O6/  
→ Consider K in place of MG (3 valid contact(s) for K vs. 0 for MG ←)  
Go to ToC

### MG:a:3163

|  | Res. type | Atom | Residue | Distance | Occupancy | B-factor | Closest atom(s) to a 1^st^ shell water | EM\_dens. |
| --- | --- | --- | --- | --- | --- | --- | --- | --- |
| **Ion** | | | |  | 1.0 | 40.4 |  | 12.83 |
 1 | Water 1 | | a:8125 | 1.88 | 1.0 | 34.2 | 2.89 [G:N7]a:376 // 2.92 [MG:MG]a:3031 | 8.22 | 2 | Water 2 | | a:5341 | 2.07 | 1.0 | 34.0 | 2.70 [G:OP2]a:375 | 9.02 | 3 | Water 3 | | a:10254 | 2.12 | 1.0 | 38.5 |  | 7.27 | 4 | Water 4 | | a:7848 | 2.16 | 1.0 | 32.8 | 2.87 [G:OP2]a:375 | 8.62 | 5 | Water 5 | | a:5670 | 2.37 | 1.0 | 30.7 | 2.72 [A:OP2]a:374 // 2.78 [G:N7]a:375 | 11.01 | 6 | Water 6 | | a:9760 | 2.39 | 1.0 | 39.8 |  | 6.64 | 7 | **MG** | MG | a:3031 | 3.63 | 1.0 | 24.6 |  | 23.53 | 8 | G | OP2 | a:375 | 3.98 | 1.0 | 35.2 |  | 7.68 | 9 | Water 7 | | a:7950 | 4.21 | 1.0 | 30.1 |  | 9.34 | 10 | Water 8 | | a:7449 | 4.26 | 1.0 | 43.6 |  | 3.84 | 11 | Water 9 | | a:7375 | 4.28 | 1.0 | 29.8 | 2.84 [MG:MG]a:3031 | 10.38 | 12 | G | N7 | a:375 | 4.39 | 1.0 | 28.4 |  | 10.79 | 13 | Water 10 | | a:6306 | 4.41 | 1.0 | 31.8 |  | 9.62 | 14 | G | N7 | a:376 | 4.43 | 1.0 | 27.3 |  | 12.16 | 15 | Water 11 | | a:10118 | 4.44 | 1.0 | 46.7 |  | 4.11 | 16 | A | OP2 | a:374 | 4.55 | 1.0 | 36.9 |  | 8.38 | 17 | Water 12 | | a:6489 | 4.74 | 1.0 | 34.9 |  | 6.79 |

  
**1^st^ shell coordination angles and distances (*trans* in "blue"; *cys* in "green"; deviations in "orange"):**   
• 1-2(W/W): 170º; 1-3(W/W): *98º*; 1-4(W/W): 89º; 2-3(W/W): 91º; 2-4(W/W): *81º*; 3-4(W/W): 173º;   
• 1-2(W/W): 3.9Å; 1-3(W/W): 3.0Å; 1-4(W/W): 2.8Å; 2-3(W/W): 3.0Å; 2-4(W/W): 2.8Å; 3-4(W/W): *4.3Å*;   
  
**2.17 ± 0.17: the average coodination distance of the 1^st^ six ligands is Mg^2+^ compatible**  
**12: Number of potential Water contacts**   
**6 : Number of second shell contacts involving water or ions**   
**50%: Percentage of satisfied second shell Water contacts**   
• Contact(s): /O_w_/O_w_/O_w_/  
• Contact code: 3O_w_  
• Contact(s) in the exclusion zone: /O_w_/O_w_/O_w_/  
→ Missing octahedral coordination for MG ←  
→ Slightly distorded coord. octahedron (6.1º) ←  
Go to ToC

### MG:a:3164

|  | Res. type | Atom | Residue | Distance | Occupancy | B-factor | Closest atom(s) to a 1^st^ shell water | EM\_dens. |
| --- | --- | --- | --- | --- | --- | --- | --- | --- |
| **Ion** | | | |  | 1.0 | 45.4 |  | 9.02 |
 1 | Water 1 | | a:6509 | 1.95 | 1.0 | 38.8 |  | 5.45 | 2 | G | OP1 | a:376 | 1.97 | 1.0 | 39.1 |  | 6.52 | 3 | Water 2 | | a:8761 | 2.03 | 1.0 | 40.2 |  | 4.90 | 4 | Water 3 | | a:3808 | 2.21 | 1.0 | 37.4 | 2.54 [U:OP2]a:392 | 6.67 | 5 | Water 4 | | a:10023 | 2.23 | 1.0 | 41.3 |  | 4.32 | 6 | Water 5 | | a:3482 | 2.36 | 1.0 | 40.7 | 2.43 [G:OP2]a:377 | 6.45 | 7 | G | OP2 | a:376 | 3.89 | 1.0 | 37.1 |  | 8.33 | 8 | G | O5' | a:376 | 3.90 | 1.0 | 37.3 |  | 8.62 | 9 | Water 6 | | a:7813 | 3.94 | 1.0 | 32.4 |  | 8.23 | 10 | Water 7 | | a:4459 | 3.99 | 1.0 | 46.1 | 2.95 [U:OP2]a:392 | 4.38 | 11 | U | OP2 | a:392 | 4.22 | 1.0 | 34.5 |  | 7.22 | 12 | G | OP2 | a:377 | 4.27 | 1.0 | 36.8 |  | 7.61 | 13 | Water 8 | | a:6489 | 4.28 | 1.0 | 34.9 | 2.77 [G:OP2]a:376 | 6.79 | 14 | Water 9 | | a:3958 | 4.37 | 1.0 | 38.2 |  | 7.52 | 15 | G | O3' | a:375 | 4.48 | 1.0 | 36.0 |  | 9.02 | 16 | Water 10 | | a:5803 | 4.52 | 1.0 | 41.9 |  | 4.47 | 17 | Water 11 | | a:9382 | 4.75 | 1.0 | 41.4 |  | 5.60 |

  
**1^st^ shell coordination angles and distances (*trans* in "blue"; *cys* in "green"; deviations in "orange"):**   
• 1-2(W/OP1): 90º; 1-3(W/W): 173º; 1-4(W/W): 93º; 1-5(W/W): 91º; 2-3(OP1/W): *95º*; 2-4(OP1/W): 92º; 2-5(OP1/W): 178º; 3-4(W/W): 91º; 3-5(W/W): *84º*; 4-5(W/W): 87º;   
• 1-2(W/OP1): 2.8Å; 1-3(W/W): 4.0Å; 1-4(W/W): 3.0Å; 1-5(W/W): 3.0Å; 2-3(OP1/W): 3.0Å; 2-4(OP1/W): 3.0Å; 2-5(OP1/W): *4.2Å*; 3-4(W/W): 3.0Å; 3-5(W/W): 2.8Å; 4-5(W/W): 3.0Å;   
  
**2.12 ± 0.15: the average coodination distance of the 1^st^ six ligands is Mg^2+^ compatible**  
**10: Number of potential Water contacts**   
**2 : Number of second shell contacts involving water or ions**   
**20%: Percentage of satisfied second shell Water contacts**   
• Contact(s): /O_w_/O_w_/O_w_/O_w_/OP1/  
• Contact code: O_ph_4O_w_  
• Contact(s) in the exclusion zone: /O_w_/  
→ Missing octahedral coordination for MG ←  
Ooctahedral coordination (3.0º)   
Go to ToC

### MG:a:3165

|  | Res. type | Atom | Residue | Distance | Occupancy | B-factor | Closest atom(s) to a 1^st^ shell water | EM\_dens. |
| --- | --- | --- | --- | --- | --- | --- | --- | --- |
| **Ion** | | | |  | 1.0 | 39.4 |  | 13.36 |
 1 | Water 1 | | a:5551 | 1.99 | 1.0 | 30.8 | 2.71 [C:OP2]a:413 | 9.54 | 2 | G | OP2 | a:410 | 2.07 | 1.0 | 33.4 |  | 9.47 | 3 | Water 2 | | a:10174 | 2.11 | 1.0 | 36.8 |  | 8.50 | 4 | Water 3 | | a:8052 | 2.21 | 1.0 | 31.7 | 2.89 [G:OP2]a:409 | 10.18 | 5 | Water 4 | | a:9316 | 2.24 | 1.0 | 34.7 |  | 9.00 | 6 | Water 5 | | a:4016 | 2.26 | 1.0 | 36.2 | 2.58 [G:OP1]a:409 | 9.12 | 7 | Water 6 | | a:9118 | 3.75 | 1.0 | 36.0 |  | 8.23 | 8 | G | OP1 | a:410 | 3.99 | 1.0 | 34.9 |  | 8.72 | 9 | Water 7 | | a:5023 | 4.06 | 1.0 | 38.5 | 2.98 [U:OP1]a:392 | 4.95 | 10 | Water 8 | | a:5668 | 4.20 | 1.0 | 37.8 |  | 7.30 | 11 | G | O3' | a:409 | 4.23 | 1.0 | 33.9 |  | 10.29 | 12 | A | O2' | a:391 | 4.31 | 1.0 | 34.3 |  | 9.47 | 13 | G | O5' | a:409 | 4.38 | 1.0 | 33.1 |  | 9.51 | 14 | C | OP2 | a:413 | 4.39 | 1.0 | 37.6 |  | 8.11 | 15 | G | OP1 | a:409 | 4.39 | 1.0 | 37.8 |  | 8.35 | 16 | Water 9 | | a:8562 | 4.47 | 1.0 | 40.7 | 2.99 [C:OP2]a:413 | 6.05 | 17 | G | O5' | a:410 | 4.49 | 1.0 | 28.7 |  | 11.00 | 18 | G | OP2 | a:409 | 4.51 | 1.0 | 34.4 |  | 8.78 | 19 | U | OP1 | a:392 | 4.52 | 1.0 | 37.6 |  | 7.94 | 20 | A | O3' | a:391 | 4.56 | 1.0 | 36.8 |  | 8.84 | 21 | Water 10 | | a:5620 | 4.67 | 1.0 | 39.2 | 2.72 [C:OP2]a:413 | 6.59 |

  
**1^st^ shell coordination angles and distances (*trans* in "blue"; *cys* in "green"; deviations in "orange"):**   
• 1-2(W/OP2): *99º*; 1-3(W/W): 87º; 1-4(W/W): 175º; 1-5(W/W): 93º; 1-6(W/W): 92º; 2-3(OP2/W): 173º; 2-4(OP2/W): 86º; 2-5(OP2/W): 90º; 2-6(OP2/W): 93º; 3-4(W/W): 87º; 3-5(W/W): 88º; 3-6(W/W): 89º; 4-5(W/W): 87º; 4-6(W/W): 87º; 5-6(W/W): 173º;   
• 1-2(W/OP2): 3.1Å; 1-3(W/W): 2.8Å; 1-4(W/W): 4.2Å; 1-5(W/W): 3.1Å; 1-6(W/W): 3.1Å; 2-3(OP2/W): 4.2Å; 2-4(OP2/W): 2.9Å; 2-5(OP2/W): 3.1Å; 2-6(OP2/W): 3.2Å; 3-4(W/W): 3.0Å; 3-5(W/W): 3.0Å; 3-6(W/W): 3.1Å; 4-5(W/W): 3.1Å; 4-6(W/W): 3.1Å; 5-6(W/W): *4.5Å*;   
  
**2.15 ± 0.10: the average coodination distance of the 1^st^ six ligands is Mg^2+^ compatible**  
**10: Number of potential Water contacts**   
**3 : Number of second shell contacts involving water or ions**   
**30%: Percentage of satisfied second shell Water contacts**   
• Contact(s): /O_w_/O_w_/O_w_/O_w_/O_w_/OP2/  
• Contact code: O_ph_5O_w_  
Ooctahedral coordination (3.7º)   
Go to ToC

### MG:a:3166

|  | Res. type | Atom | Residue | Distance | Occupancy | B-factor | Closest atom(s) to a 1^st^ shell water | EM\_dens. |
| --- | --- | --- | --- | --- | --- | --- | --- | --- |
| **Ion** | | | |  | 1.0 | 48.6 |  | 6.99 |
 1 | Water 1 | | a:7489 | 1.94 | 1.0 | 39.3 | 2.97 [C:OP1]a:445 | 6.74 | 2 | Water 2 | | a:9779 | 2.03 | 1.0 | 41.4 |  | 6.30 | 3 | G | OP2 | a:446 | 2.08 | 1.0 | 30.9 |  | 11.22 | 4 | Water 3 | | a:7682 | 2.22 | 1.0 | 38.7 |  | 4.87 | 5 | Water 4 | | a:5607 | 2.80 | 1.0 | 37.7 | 2.72 [G:OP1]a:446 // 2.97 [ARG:O]p:3 | 6.63 | 6 | Water 5 | | a:4812 | 3.43 | 1.0 | 40.1 | 2.66 [C:OP2]a:445 | 5.02 | 7 | G | OP1 | a:446 | 3.90 | 1.0 | 28.8 |  | 11.72 | 8 | G | O5' | a:446 | 4.15 | 1.0 | 24.4 |  | 13.36 | 9 | U | O2' | a:1201 | 4.26 | 1.0 | 31.7 |  | 8.78 | 10 | ARG | O | p:3 | 4.38 | 1.0 | 30.9 |  | 10.46 | 11 | C | OP1 | a:445 | 4.43 | 1.0 | 32.7 |  | 10.65 | 12 | C | OP2 | a:445 | 4.53 | 1.0 | 27.1 |  | 11.88 | 13 | C | O3' | a:445 | 4.56 | 1.0 | 28.7 |  | 12.26 | 14 | Water 6 | | a:7265 | 4.67 | 1.0 | 43.8 |  | 4.73 |

  
**1^st^ shell coordination angles and distances (*trans* in "blue"; *cys* in "green"; deviations in "orange"):**   
• 1-2(W/W): 94º; 1-3(W/OP2): 90º; 1-4(W/W): *166º*; 2-3(W/OP2): *151º*; 2-4(W/W): 86º; 3-4(OP2/W): *83º*;   
• 1-2(W/W): 2.9Å; 1-3(W/OP2): 2.8Å; 1-4(W/W): 4.1Å; 2-3(W/OP2): 4.0Å; 2-4(W/W): 2.9Å; 3-4(OP2/W): 2.9Å;   
  
**2.42 ± 0.53: the average coodination distance of the 1^st^ six ligands is Na^+^** compatible  
**10: Number of potential Water contacts**   
**4 : Number of second shell contacts involving water or ions**   
**40%: Percentage of satisfied second shell Water contacts**   
• Contact(s): /O_w_/O_w_/O_w_/OP2/  
• Contact code: O_ph_3O_w_  
• Contact(s) in the exclusion zone: /O_w_/  
→ Missing octahedral coordination for MG ←  
→ Slightly distorded coord. octahedron (9.8º) ←  
Go to ToC

### MG:a:3167

|  | Res. type | Atom | Residue | Distance | Occupancy | B-factor | Closest atom(s) to a 1^st^ shell water | EM\_dens. |
| --- | --- | --- | --- | --- | --- | --- | --- | --- |
| **Ion** | | | |  | 1.0 | 29.0 |  | 21.82 |
 1 | Water 1 | | a:8787 | 1.94 | 1.0 | 26.8 | 2.97 [G:OP2]a:587 | 13.59 | 2 | G | OP1 | a:1252 | 2.02 | 1.0 | 25.3 |  | 13.03 | 3 | U | OP1 | a:1251 | 2.07 | 1.0 | 27.0 |  | 11.58 | 4 | Water 2 | | a:6674 | 2.15 | 1.0 | 17.4 | 2.78 [U:O4]a:813 | 18.64 | 5 | Water 3 | | a:3695 | 2.39 | 1.0 | 19.9 | 2.51 [A:OP2]a:588 | 16.85 | 6 | Water 4 | | a:6169 | 2.39 | 1.0 | 25.8 | 2.75 [G:OP1]a:587 // 2.94 [A:OP2]a:588 | 13.42 | 7 | U | O3' | a:1251 | 3.51 | 1.0 | 26.2 |  | 13.65 | 8 | A | OP2 | a:588 | 3.63 | 1.0 | 26.0 |  | 13.12 | 9 | G | OP2 | a:587 | 3.94 | 1.0 | 26.1 |  | 15.85 | 10 | Water 5 | | a:4418 | 4.11 | 1.0 | 35.2 |  | 9.05 | 11 | G | O3' | a:1250 | 4.16 | 1.0 | 25.5 |  | 12.92 | 12 | G | OP2 | a:1252 | 4.16 | 1.0 | 30.4 |  | 14.87 | 13 | Water 6 | | p:215 | 4.23 | 1.0 | 29.4 |  | 9.52 | 14 | U | OP2 | a:1251 | 4.32 | 1.0 | 26.7 |  | 11.92 | 15 | U | O4 | a:813 | 4.36 | 1.0 | 24.8 |  | 16.79 | 16 | G | O5' | a:1252 | 4.36 | 1.0 | 25.4 |  | 13.54 | 17 | U | O5' | a:1251 | 4.45 | 1.0 | 26.1 |  | 13.51 | 18 | Water 7 | | a:3937 | 4.49 | 1.0 | 34.8 |  | 9.38 | 19 | G | OP1 | a:587 | 4.54 | 1.0 | 22.9 |  | 14.86 |

  
**1^st^ shell coordination angles and distances (*trans* in "blue"; *cys* in "green"; deviations in "orange"):**   
• 1-2(W/OP1): *100º*; 1-3(W/OP1): *95º*; 1-4(W/W): *165º*; 2-3(OP1/OP1): *95º*; 2-4(OP1/W): 91º; 3-4(OP1/W): 94º;   
• 1-2(W/OP1): 3.0Å; 1-3(W/OP1): 3.0Å; 1-4(W/W): 4.1Å; 2-3(OP1/OP1): 3.0Å; 2-4(OP1/W): 3.0Å; 3-4(OP1/W): 3.1Å;   
  
**2.16 ± 0.17: the average coodination distance of the 1^st^ six ligands is Mg^2+^ compatible**  
**8: Number of potential Water contacts**   
**5 : Number of second shell contacts involving water or ions**   
**62%: Percentage of satisfied second shell Water contacts**   
• Contact(s): /O_w_/O_w_/OP1/OP1/  
• Contact code: *cis-*2O_ph_2O_w_  
• Contact(s) in the exclusion zone: /O_w_/O_w_/  
→ Missing octahedral coordination for MG ←  
→ Slightly distorded coord. octahedron (6.7º) ←  
Go to ToC

### MG:a:3168

|  | Res. type | Atom | Residue | Distance | Occupancy | B-factor | Closest atom(s) to a 1^st^ shell water | EM\_dens. |
| --- | --- | --- | --- | --- | --- | --- | --- | --- |
| **Ion** | | | |  | 1.0 | 54.0 |  | 4.11 |
 1 | Water 1 | | a:10280 | 1.88 | 1.0 | 43.1 |  | 3.88 | 2 | Water 2 | | a:7522 | 2.14 | 1.0 | 36.6 | 2.84 [G:N7]a:463 | 8.64 | 3 | Water 3 | | a:5435 | 2.38 | 1.0 | 35.2 | 2.70 [G:OP2]a:465 | 7.59 | 4 | Water 4 | | a:4645 | 3.36 | 1.0 | 44.7 | 2.64 [G:O6]a:463 | 6.01 | 5 | G | N7 | a:463 | 3.71 | 1.0 | 18.8 |  | 14.78 | 6 | Water 5 | | 1:144 | 3.78 | 1.0 | 38.6 |  | 5.09 | 7 | Water 6 | | a:4997 | 3.98 | 1.0 | 39.0 |  | 6.08 | 8 | G | OP1 | a:465 | 4.03 | 1.0 | 25.2 |  | 12.53 | 9 | Water 7 | | a:5384 | 4.15 | 1.0 | 36.0 |  | 8.10 | 10 | G | OP2 | a:465 | 4.40 | 1.0 | 24.4 |  | 14.71 | 11 | G | O6 | a:463 | 4.51 | 1.0 | 24.3 |  | 13.83 |

  
**1^st^ shell coordination angles and distances (*trans* in "blue"; *cys* in "green"; deviations in "orange"):**   
• 1-2(W/W): *73º*;   
• 1-2(W/W): 2.4Å;   
  
**2.87 ± 0.76: the average coodination distance of the 1^st^ six ligands is K^+^ compatible**  
**10: Number of potential Water contacts**   
**3 : Number of second shell contacts involving water or ions**   
**30%: Percentage of satisfied second shell Water contacts**   
• Contact(s): /O_w_/  
• Contact code: O_w_  
• Contact(s) in the exclusion zone: /O_w_/O_w_/O_w_/  
→ Missing octahedral coordination for MG ←  
Go to ToC

### MG:a:3169

|  | Res. type | Atom | Residue | Distance | Occupancy | B-factor | Closest atom(s) to a 1^st^ shell water | EM\_dens. |
| --- | --- | --- | --- | --- | --- | --- | --- | --- |
| **Ion** | | | |  | 1.0 | 41.3 |  | 10.93 |
 1 | Water 1 | | a:6152 | 2.68 | 1.0 | 43.7 | 2.93 [U:OP1]a:2783 | 4.50 | 2 | Water 2 | | a:6868 | 2.87 | 1.0 | 36.5 | 2.80 [A:OP2]a:2046 | 11.06 | 3 | Water 3 | | a:10257 | 2.92 | 1.0 | 47.6 |  | 4.08 | 4 | C | N3 | a:527 | 2.97 | 1.0 | 21.5 |  | 13.88 | 5 | Water 4 | | a:7266 | 3.13 | 1.0 | 32.2 |  | 6.99 | 6 | C | O2 | a:527 | 3.17 | 1.0 | 28.9 |  | 13.87 | 7 | Water 5 | | a:4626 | 3.26 | 1.0 | 43.2 | 2.64 [U:OP2]a:2783 | 5.16 | 8 | U | OP2 | a:2783 | 3.55 | 1.0 | 32.3 |  | 9.69 | 9 | U | OP1 | a:2783 | 4.11 | 1.0 | 36.5 |  | 7.67 | 10 | A | OP2 | a:2046 | 4.60 | 1.0 | 32.4 |  | 10.51 | 11 | Water 6 | | a:6962 | 4.63 | 1.0 | 39.9 |  | 3.85 |

  
• • **2.96 ± 0.16: the average coodination distance of the 1^st^ six ligands is K^+^ compatible**  
**8: Number of potential Water contacts**   
**2 : Number of second shell contacts involving water or ions**   
**25%: Percentage of satisfied second shell Water contacts**   
• Contact code: → No contacts ←   
→ Consider K in place of MG (2 valid contact(s) for K vs. 0 for MG ←)  
Go to ToC

### MG:a:3170

|  | Res. type | Atom | Residue | Distance | Occupancy | B-factor | Closest atom(s) to a 1^st^ shell water | EM\_dens. |
| --- | --- | --- | --- | --- | --- | --- | --- | --- |
| **Ion** | | | |  | 1.0 | 30.9 |  | 20.65 |
 1 | A | OP2 | a:1191 | 2.05 | 1.0 | 22.7 |  | 15.35 | 2 | Water 1 | | k:255 | 2.20 | 1.0 | 28.9 |  | 9.66 | 3 | Water 2 | | a:6273 | 2.25 | 1.0 | 23.8 | 2.76 [G:O6]a:1192 | 17.22 | 4 | G | N7 | a:1192 | 2.26 | 1.0 | 18.1 |  | 16.70 | 5 | Water 3 | | a:6928 | 2.29 | 1.0 | 31.8 |  | 9.13 | 6 | A | OP1 | a:1191 | 3.76 | 1.0 | 25.6 |  | 11.91 | 7 | A | O5' | a:1191 | 3.81 | 1.0 | 24.9 |  | 15.99 | 8 | G | O6 | a:1192 | 3.92 | 1.0 | 25.2 |  | 18.04 | 9 | Water 4 | | a:9557 | 4.13 | 1.0 | 40.5 |  | 6.25 | 10 | Water 5 | | a:6177 | 4.35 | 1.0 | 36.2 | 2.83 [GLY:O]k:26 | 8.36 | 11 | Water 6 | | a:7260 | 4.36 | 1.0 | 33.7 |  | 6.85 | 12 | Water 7 | | a:3834 | 4.44 | 1.0 | 33.4 |  | 10.55 | 13 | G | OP2 | a:1192 | 4.46 | 1.0 | 25.1 |  | 14.02 | 14 | GLY | O | k:26 | 4.47 | 1.0 | 36.8 |  | 11.40 | 15 | Water 8 | | a:8518 | 4.47 | 1.0 | 41.1 |  | 4.86 | 16 | U | O3' | a:1190 | 4.54 | 1.0 | 26.1 |  | 14.97 | 17 | Water 9 | | a:10276 | 4.68 | 1.0 | 41.3 |  | 3.77 | 18 | U | OP2 | a:1190 | 4.69 | 1.0 | 28.9 |  | 12.42 |

  
**1^st^ shell coordination angles and distances (*trans* in "blue"; *cys* in "green"; deviations in "orange"):**   
• 1-2(OP2/W): *157º*; 1-3(OP2/W): 86º; 1-4(OP2/N7): *112º*; 1-5(OP2/W): *80º*; 2-3(W/W): 90º; 2-4(W/N7): 91º; 2-5(W/W): *77º*; 3-4(W/N7): 92º; 3-5(W/W): *84º*; 4-5(N7/W): *167º*;   
• 1-2(OP2/W): 4.2Å; 1-3(OP2/W): 2.9Å; 1-4(OP2/N7): *3.6Å*; 1-5(OP2/W): 2.8Å; 2-3(W/W): 3.1Å; 2-4(W/N7): 3.2Å; 2-5(W/W): 2.8Å; 3-4(W/N7): *3.2Å*; 3-5(W/W): 3.0Å; 4-5(N7/W): *4.5Å*;   
  
**2.47 ± 0.58: the average coodination distance of the 1^st^ six ligands is Na^+^** compatible  
**6: Number of potential Water contacts**   
**1 : Number of second shell contacts involving water or ions**   
**17%: Percentage of satisfied second shell Water contacts**   
• Contact(s): /N7/O_w_/O_w_/O_w_/OP2/  
• Contact code: O_ph_N_b_3O_w_  
→ Missing octahedral coordination for MG ←  
→ Slightly distorded coord. octahedron (9.3º) ←  
Go to ToC

### MG:a:3171

|  | Res. type | Atom | Residue | Distance | Occupancy | B-factor | Closest atom(s) to a 1^st^ shell water | EM\_dens. |
| --- | --- | --- | --- | --- | --- | --- | --- | --- |
| **Ion** | | | |  | 1.0 | 38.6 |  | 14.33 |
 1 | Water 1 | | z:251 | 2.03 | 1.0 | 38.0 |  | 12.01 | 2 | Water 2 | | a:5092 | 2.05 | 1.0 | 29.0 | 2.68 [C:OP1]a:2059 | 11.49 | 3 | Water 3 | | z:209 | 2.28 | 1.0 | 26.7 | 2.65 [GLN:OE1]z:5 | 11.99 | 4 | Water 4 | | a:10207 | 2.30 | 1.0 | 35.1 |  | 8.58 | 5 | Water 5 | | a:5095 | 2.40 | 1.0 | 31.4 | 2.68 [C:OP2]a:2059 | 10.46 | 6 | Water 6 | | a:6051 | 3.73 | 1.0 | 30.5 |  | 12.94 | 7 | C | OP2 | a:2059 | 4.10 | 1.0 | 22.4 |  | 14.43 | 8 | Water 7 | | a:4361 | 4.15 | 1.0 | 33.5 | 2.90 [G:OP1]a:2061 | 11.02 | 9 | C | OP1 | a:2059 | 4.15 | 1.0 | 30.4 |  | 13.63 | 10 | Water 8 | | a:3852 | 4.18 | 1.0 | 29.0 |  | 13.09 | 11 | Water 9 | | z:218 | 4.28 | 1.0 | 30.8 |  | 13.77 | 12 | Water 10 | | a:5250 | 4.29 | 1.0 | 32.4 |  | 13.54 | 13 | GLN | OE1 | z:5 | 4.32 | 1.0 | 31.5 |  | 15.56 | 14 | Water 11 | | a:8227 | 4.45 | 1.0 | 31.0 |  | 12.28 | 15 | G | OP1 | a:2061 | 4.46 | 1.0 | 26.5 |  | 12.14 | 16 | Water 12 | | a:9352 | 4.49 | 1.0 | 23.2 |  | 17.34 | 17 | Water 13 | | a:5065 | 4.58 | 1.0 | 24.8 | 2.68 [C:OP2]a:2059 | 14.58 | 18 | Water 14 | | a:10057 | 4.61 | 1.0 | 36.0 |  | 9.45 | 19 | Water 15 | | a:7206 | 4.76 | 1.0 | 35.7 |  | 9.41 |

  
**1^st^ shell coordination angles and distances (*trans* in "blue"; *cys* in "green"; deviations in "orange"):**   
• 1-2(W/W): *100º*; 1-3(W/W): 91º; 1-4(W/W): *79º*; 2-3(W/W): 87º; 2-4(W/W): *168º*; 3-4(W/W): *81º*;   
• 1-2(W/W): 3.1Å; 1-3(W/W): 3.1Å; 1-4(W/W): 2.8Å; 2-3(W/W): 3.0Å; 2-4(W/W): *4.3Å*; 3-4(W/W): 3.0Å;   
  
**2.47 ± 0.58: the average coodination distance of the 1^st^ six ligands is Na^+^** compatible  
**12: Number of potential Water contacts**   
**3 : Number of second shell contacts involving water or ions**   
**25%: Percentage of satisfied second shell Water contacts**   
• Contact(s): /O_w_/O_w_/O_w_/O_w_/  
• Contact code: 4O_w_  
• Contact(s) in the exclusion zone: /O_w_/  
→ Missing octahedral coordination for MG ←  
→ Slightly distorded coord. octahedron (7.6º) ←  
Go to ToC

### MG:a:3172

|  | Res. type | Atom | Residue | Distance | Occupancy | B-factor | Closest atom(s) to a 1^st^ shell water | EM\_dens. |
| --- | --- | --- | --- | --- | --- | --- | --- | --- |
| **Ion** | | | |  | 1.0 | 52.7 |  | 12.18 |
 1 | Water 1 | | a:9603 | 1.96 | 1.0 | 30.0 | 2.18 [MG:MG]a:3353 | 13.07 | 2 | G | N7 | a:2506 | 2.04 | 1.0 | 20.6 |  | 15.91 | 3 | Water 2 | | a:9330 | 2.22 | 1.0 | 32.1 |  | 9.80 | 4 | Water 3 | | a:7190 | 2.23 | 1.0 | 22.0 | 1.98 [MG:MG]a:3353 | 14.61 | 5 | Water 4 | | a:6080 | 2.55 | 1.0 | 26.1 | 2.84 [G:O6]a:2506 | 14.48 | 6 | **MG** | MG | a:3353 | 2.86 | 1.0 | 43.4 |  | 14.26 | 7 | G | O6 | a:2506 | 3.81 | 1.0 | 26.1 |  | 13.64 | 8 | Water 5 | | a:4500 | 3.99 | 1.0 | 27.7 | 2.73 [G:OP1]a:2506 | 14.91 | 9 | G | OP1 | a:2450 | 4.12 | 1.0 | 32.2 |  | 13.70 | 10 | G | OP2 | a:2450 | 4.25 | 1.0 | 24.5 |  | 15.39 | 11 | Water 6 | | a:8014 | 4.28 | 1.0 | 35.9 | 2.15 [MG:MG]a:3353 | 7.46 | 12 | Water 7 | | a:8656 | 4.38 | 1.0 | 34.7 |  | 8.96 | 13 | Water 8 | | a:3457 | 4.40 | 1.0 | 32.2 |  | 10.16 | 14 | G | OP1 | a:2506 | 4.48 | 1.0 | 30.3 |  | 13.98 | 15 | Water 9 | | a:5938 | 4.53 | 1.0 | 29.9 | 2.24 [MG:MG]a:3353 | 9.28 | 16 | U | O4' | a:578 | 4.56 | 1.0 | 28.2 |  | 15.96 | 17 | G | OP2 | a:2451 | 4.64 | 1.0 | 27.2 |  | 14.53 | 18 | U | O2' | a:578 | 4.70 | 1.0 | 24.0 |  | 12.23 | 19 | **K** | K | a:3110 | 4.78 | 1.0 | 30.0 |  | 19.46 |

  
**1^st^ shell coordination angles and distances (*trans* in "blue"; *cys* in "green"; deviations in "orange"):**   
• 1-2(W/N7): *104º*; 1-3(W/W): 90º; 1-4(W/W): 88º; 2-3(N7/W): 92º; 2-4(N7/W): *167º*; 3-4(W/W): 92º;   
• 1-2(W/N7): 3.2Å; 1-3(W/W): 3.0Å; 1-4(W/W): 2.9Å; 2-3(N7/W): 3.1Å; 2-4(N7/W): *4.2Å*; 3-4(W/W): 3.2Å;   
  
**2.31 ± 0.31: the average coodination distance of the 1^st^ six ligands is Na^+^** compatible  
**8: Number of potential Water contacts**   
**3 : Number of second shell contacts involving water or ions**   
**38%: Percentage of satisfied second shell Water contacts**   
• Contact(s): /N7/O_w_/O_w_/O_w_/  
• Contact code: N_b_3O_w_  
• Contact(s) in the exclusion zone: /O_w_/  
→ Missing octahedral coordination for MG ←  
→ Slightly distorded coord. octahedron (5.4º) ←  
Go to ToC

### MG:a:3173

|  | Res. type | Atom | Residue | Distance | Occupancy | B-factor | Closest atom(s) to a 1^st^ shell water | EM\_dens. |
| --- | --- | --- | --- | --- | --- | --- | --- | --- |
| **Ion** | | | |  | 1.0 | 39.7 |  | 11.15 |
 1 | Water 1 | | a:9781 | 2.76 | 1.0 | 23.5 |  | 13.12 | 2 | U | O2' | a:575 | 2.81 | 1.0 | 22.6 |  | 14.26 | 3 | Water 2 | | a:7353 | 2.87 | 1.0 | 25.7 | 2.83 [A:O2']a:576 | 14.99 | 4 | U | O2 | a:575 | 2.88 | 1.0 | 23.6 |  | 18.61 | 5 | A | OP2 | a:2035 | 3.05 | 1.0 | 27.3 |  | 11.57 | 6 | Water 3 | | a:4354 | 3.18 | 1.0 | 41.3 | 2.62 [A:OP2]a:2035 | 3.53 | 7 | A | O2' | a:576 | 3.58 | 1.0 | 34.9 |  | 11.80 | 8 | 6MZ | O2' | a:2034 | 3.58 | 1.0 | 28.7 |  | 16.06 | 9 | 6MZ | O3' | a:2034 | 3.67 | 1.0 | 24.0 |  | 17.63 | 10 | Water 4 | | a:6953 | 3.73 | 1.0 | 42.1 | 2.80 [A:OP2]a:2035 | 6.60 | 11 | A | OP2 | a:577 | 3.97 | 1.0 | 26.5 |  | 13.15 | 12 | A | O3' | a:576 | 4.00 | 1.0 | 25.9 |  | 14.50 | 13 | A | OP1 | a:577 | 4.03 | 1.0 | 27.2 |  | 14.60 | 14 | A | OP1 | a:2035 | 4.20 | 1.0 | 22.6 |  | 14.68 | 15 | U | N1 | a:575 | 4.29 | 1.0 | 18.4 |  | 17.46 | 16 | C | O2' | a:2503 | 4.68 | 1.0 | 29.1 |  | 14.03 | 17 | U | O2 | a:573 | 4.74 | 1.0 | 19.8 |  | 17.28 | 18 | Water 5 | | a:7936 | 4.76 | 1.0 | 22.0 | 2.88 [A:OP1]a:577 | 15.41 |

  
• • **2.92 ± 0.14: the average coodination distance of the 1^st^ six ligands is K^+^ compatible**  
**6: Number of potential Water contacts**   
**2 : Number of second shell contacts involving water or ions**   
**33%: Percentage of satisfied second shell Water contacts**   
• Contact code: → No contacts ←   
• Contact(s) in the exclusion zone: /O2/O2'/O_w_/O_w_/O_w_/OP2/  
→ Consider K in place of MG (3 valid contact(s) for K vs. 0 for MG ←)  
Go to ToC

### MG:a:3174

|  | Res. type | Atom | Residue | Distance | Occupancy | B-factor | Closest atom(s) to a 1^st^ shell water | EM\_dens. |
| --- | --- | --- | --- | --- | --- | --- | --- | --- |
| **Ion** | | | |  | 1.0 | 40.5 |  | 10.71 |
 1 | Water 1 | | a:5096 | 1.95 | 1.0 | 31.6 | 2.68 [U:O4]a:2032 | 8.33 | 2 | A | OP1 | a:2037 | 2.08 | 1.0 | 30.8 |  | 7.59 | 3 | Water 2 | | a:8065 | 2.10 | 1.0 | 30.5 | 2.89 [G:O6]a:2031 | 11.34 | 4 | Water 3 | | a:7777 | 2.21 | 1.0 | 33.6 | 2.86 [G:N7]a:2031 | 8.34 | 5 | Water 4 | | a:9489 | 2.22 | 1.0 | 38.7 |  | 6.89 | 6 | Water 5 | | a:9885 | 2.30 | 1.0 | 41.5 |  | 5.11 | 7 | U | O4 | a:2032 | 3.30 | 1.0 | 24.8 |  | 13.45 | 8 | Water 6 | | a:8725 | 3.92 | 1.0 | 38.1 |  | 4.00 | 9 | Water 7 | | a:8579 | 4.08 | 1.0 | 34.1 | 2.94 [A:O5']a:2037 | 9.48 | 10 | Water 8 | | a:9673 | 4.12 | 1.0 | 42.7 |  | 5.19 | 11 | A | O5' | a:2037 | 4.21 | 1.0 | 29.0 |  | 12.83 | 12 | Water 9 | | a:3939 | 4.39 | 1.0 | 36.4 |  | 5.49 | 13 | G | O3' | a:2036 | 4.39 | 1.0 | 34.3 |  | 10.37 | 14 | A | OP2 | a:2037 | 4.41 | 1.0 | 27.0 |  | 11.55 | 15 | G | N7 | a:2031 | 4.42 | 1.0 | 17.8 |  | 15.35 | 16 | Water 10 | | a:4963 | 4.49 | 1.0 | 32.4 |  | 10.98 | 17 | G | O6 | a:2033 | 4.50 | 1.0 | 24.7 |  | 15.93 | 18 | G | O6 | a:2031 | 4.64 | 1.0 | 20.0 |  | 16.64 |

  
**1^st^ shell coordination angles and distances (*trans* in "blue"; *cys* in "green"; deviations in "orange"):**   
• 1-2(W/OP1): 90º; 1-3(W/W): *120º*; 1-4(W/W): 94º; 1-5(W/W): *160º*; 1-6(W/W): *80º*; 2-3(OP1/W): *96º*; 2-4(OP1/W): 171º; 2-5(OP1/W): 93º; 2-6(OP1/W): *103º*; 3-4(W/W): *75º*; 3-5(W/W): *79º*; 3-6(W/W): *153º*; 4-5(W/W): 86º; 4-6(W/W): 85º; 5-6(W/W): *81º*;   
• 1-2(W/OP1): 2.9Å; 1-3(W/W): *3.5Å*; 1-4(W/W): 3.0Å; 1-5(W/W): 4.1Å; 1-6(W/W): 2.7Å; 2-3(OP1/W): 3.1Å; 2-4(OP1/W): *4.3Å*; 2-5(OP1/W): 3.1Å; 2-6(OP1/W): *3.4Å*; 3-4(W/W): 2.6Å; 3-5(W/W): 2.8Å; 3-6(W/W): *4.3Å*; 4-5(W/W): 3.0Å; 4-6(W/W): 3.1Å; 5-6(W/W): 2.9Å;   
  
**2.14 ± 0.12: the average coodination distance of the 1^st^ six ligands is Mg^2+^ compatible**  
**10: Number of potential Water contacts**   
**3 : Number of second shell contacts involving water or ions**   
**30%: Percentage of satisfied second shell Water contacts**   
• Contact(s): /O_w_/O_w_/O_w_/O_w_/O_w_/OP1/  
• Contact code: O_ph_5O_w_  
• Contact(s) in the exclusion zone: /O4/  
→→→ Distorded coord. octahedron (11.0º) ←←←  
Go to ToC

### K:a:3175

|  | Res. type | Atom | Residue | Distance | Occupancy | B-factor | Closest atom(s) to a 1^st^ shell water | EM\_dens. |
| --- | --- | --- | --- | --- | --- | --- | --- | --- |
| **Ion** | | | |  | 1.0 | 30.0 |  | 5.33 |
 1 | Water 1 | | a:9413 | 2.49 | 1.0 | 40.0 |  | 4.69 | 2 | G | O6 | a:730 | 2.70 | 1.0 | 28.9 |  | 10.69 | 3 | U | O4 | a:705 | 2.78 | 1.0 | 34.0 |  | 9.57 | 4 | Water 2 | | a:4420 | 3.30 | 1.0 | 43.1 | 2.62 [A:OP2]a:729 // 2.65 [G:N7]a:730 | 5.92 | 5 | G | N2 | a:706 | 3.36 | 1.0 | 27.5 |  | 10.34 | 6 | A | N6 | a:732 | 3.49 | 1.0 | 21.2 |  | 13.48 | 7 | U | N3 | a:705 | 3.63 | 1.0 | 23.2 |  | 12.84 | 8 | U | O4 | a:704 | 3.69 | 1.0 | 34.8 |  | 8.61 | 9 | Water 3 | | a:6777 | 3.69 | 1.0 | 41.5 | 2.79 [G:O2']a:728 | 5.74 | 10 | Water 4 | | a:4788 | 4.17 | 1.0 | 43.3 |  | 4.18 | 11 | A | OP2 | a:729 | 4.18 | 1.0 | 34.1 |  | 9.41 | 12 | G | N7 | a:730 | 4.22 | 1.0 | 23.5 |  | 13.11 | 13 | G | O2' | a:728 | 4.27 | 1.0 | 38.2 |  | 8.34 | 14 | A | N7 | a:729 | 4.75 | 1.0 | 24.4 |  | 13.55 | 15 | A | N1 | a:732 | 4.77 | 1.0 | 20.9 |  | 13.89 |

  
**1^st^ shell coordination angles and distances (*trans* in "blue"; *cys* in "green"; deviations in "orange"):**   
• 1-2(W/O6): 158; 1-3(W/O4): 72; 2-3(O6/O4): 88;   
• 1-2(W/O6): 5.1; 1-3(W/O4): 3.1; 2-3(O6/O4): 3.8;   
  
**4.24 ± 1.32: the average coodination distance of the 1^st^ six ligands could correspond to an hexahydrated Mg^2+^.**  
**4: Number of potential Water contacts**   
**2 : Number of second shell contacts involving water or ions**   
**50%: Percentage of satisfied second shell Water contacts**   
• Contact(s): /O4/O6/  
• Contact code: 2O_b_  
• Contact(s) in the exclusion zone: /O_w_/  
→ Coordination < 6 for K ←  
Go to ToC

### K:a:3176

|  | Res. type | Atom | Residue | Distance | Occupancy | B-factor | Closest atom(s) to a 1^st^ shell water | EM\_dens. |
| --- | --- | --- | --- | --- | --- | --- | --- | --- |
| **Ion** | | | |  | 1.0 | 30.0 |  | 15.71 |
 1 | Water 1 | | a:6925 | 2.52 | 1.0 | 29.2 | 2.80 [C:O2']a:586 | 12.12 | 2 | Water 2 | | a:9858 | 2.78 | 1.0 | 34.0 |  | 8.98 | 3 | C | O2 | a:1259 | 2.83 | 1.0 | 21.9 |  | 16.66 | 4 | C | O2 | a:586 | 2.88 | 1.0 | 24.9 |  | 18.21 | 5 | ILE | O | e:77 | 3.18 | 1.0 | 22.0 |  | 17.91 | 6 | ARG | N | e:79 | 3.19 | 1.0 | 15.9 |  | 18.01 | 7 | G | N2 | a:1258 | 3.77 | 1.0 | 15.7 |  | 19.36 | 8 | C | O2' | a:586 | 3.99 | 1.0 | 23.6 |  | 16.94 | 9 | TRP | O | e:78 | 4.12 | 1.0 | 27.0 |  | 16.50 | 10 | C | O2' | a:1259 | 4.18 | 1.0 | 25.7 |  | 17.36 | 11 | C | N1 | a:1259 | 4.29 | 1.0 | 19.4 |  | 17.11 | 12 | Water 3 | | a:4516 | 4.43 | 1.0 | 29.6 |  | 9.97 | 13 | C | N1 | a:586 | 4.56 | 1.0 | 13.9 |  | 17.78 | 14 | Water 4 | | e:413 | 4.56 | 1.0 | 33.0 | 2.62 [TRP:O]e:78 | 10.62 | 15 | C | N3 | a:1259 | 4.76 | 1.0 | 15.4 |  | 17.16 |

  
**1^st^ shell coordination angles and distances (*trans* in "blue"; *cys* in "green"; deviations in "orange"):**   
• 1-2(W/W): 95; 1-3(W/O2): 151; 1-4(W/O2): 81; 1-5(W/W): 61; 1-6(W/N): 92; 2-3(W/O2): 109; 2-4(W/O2): 99; 2-5(W/W): 150; 2-6(W/N): 88; 3-4(O2/O2): 79; 3-5(O2/W): 100; 3-6(O2/N): 104; 4-5(O2/W): 95; 4-6(O2/N): 171; 5-6(W/N): 76;   
• 1-2(W/W): 3.9; 1-3(W/O2): 5.2; 1-4(W/O2): 3.5; 1-5(W/W): 3.0; 1-6(W/N): 4.1; 2-3(W/O2): 4.6; 2-4(W/O2): 4.3; 2-5(W/W): 5.8; 2-6(W/N): 4.2; 3-4(O2/O2): 3.6; 3-5(O2/W): 4.6; 3-6(O2/N): 4.8; 4-5(O2/W): 4.5; 4-6(O2/N): 6.0; 5-6(W/N): 3.9;   
  
**4.19 ± 1.33: the average coodination distance of the 1^st^ six ligands could correspond to an hexahydrated Mg^2+^.**  
**6: Number of potential Water contacts**   
**1 : Number of second shell contacts involving water or ions**   
**17%: Percentage of satisfied second shell Water contacts**   
• Contact(s): /N_back_/O2/O2/O_back_/O_w_/  
• Contact code: 2O_b_O_back_O_w_Anion  
• Contact(s) in the exclusion zone: /O_w_/  
→ Coordination < 6 for K ←  
Go to ToC

### K:a:3177

|  | Res. type | Atom | Residue | Distance | Occupancy | B-factor | Closest atom(s) to a 1^st^ shell water | EM\_dens. |
| --- | --- | --- | --- | --- | --- | --- | --- | --- |
| **Ion** | | | |  | 1.0 | 30.0 |  | 25.49 |
 1 | Water 1 | | a:6532 | 2.86 | 1.0 | 32.0 |  | 8.29 | 2 | G | O6 | a:2042 | 2.95 | 1.0 | 20.6 |  | 16.77 | 3 | G | O6 | a:530 | 2.96 | 1.0 | 29.2 |  | 11.94 | 4 | U | O4 | a:2043 | 2.97 | 1.0 | 21.4 |  | 14.74 | 5 | Water 2 | | a:8196 | 3.03 | 1.0 | 31.8 |  | 11.09 | 6 | G | N7 | a:2042 | 3.06 | 1.0 | 16.8 |  | 17.31 | 7 | Water 3 | | a:5839 | 3.43 | 1.0 | 24.8 |  | 14.11 | 8 | A | N7 | a:2041 | 3.61 | 1.0 | 19.0 |  | 17.99 | 9 | Water 4 | | a:6398 | 3.66 | 1.0 | 28.7 | 2.90 [G:N1]a:530 | 11.85 | 10 | Water 5 | | a:8118 | 3.82 | 1.0 | 36.9 |  | 6.01 | 11 | Water 6 | | a:4618 | 4.46 | 1.0 | 31.6 |  | 8.18 |

  
**1^st^ shell coordination angles and distances (*trans* in "blue"; *cys* in "green"; deviations in "orange"):**   
• 1-2(W/O6): 117; 1-3(W/O6): 83; 1-4(W/O4): 156; 1-5(W/W): 83; 1-6(W/N7): 126; 2-3(O6/O6): 123; 2-4(O6/O4): 71; 2-5(O6/W): 138; 2-6(O6/N7): 61; 3-4(O6/O4): 75; 3-5(O6/W): 94; 3-6(O6/N7): 148; 4-5(O4/W): 106; 4-6(O4/N7): 78; 5-6(W/N7): 77;   
• 1-2(W/O6): 5.0; 1-3(W/O6): 3.8; 1-4(W/O4): 5.7; 1-5(W/W): 3.9; 1-6(W/N7): 5.3; 2-3(O6/O6): 5.2; 2-4(O6/O4): 3.4; 2-5(O6/W): 5.6; 2-6(O6/N7): 3.0; 3-4(O6/O4): 3.6; 3-5(O6/W): 4.4; 3-6(O6/N7): 5.8; 4-5(O4/W): 4.8; 4-6(O4/N7): 3.8; 5-6(W/N7): 3.8;   
  
**4.15 ± 1.23: the average coodination distance of the 1^st^ six ligands could correspond to an hexahydrated Mg^2+^.**  
**6: Number of potential Water contacts**   
**0 : Number of second shell contacts involving water or ions**   
**0%: Percentage of satisfied second shell Water contacts**   
• Contact(s): /N7/O4/O6/O6/O_w_/O_w_/  
• Contact code: 3O_b_N_b_2O_w_  
Go to ToC

### K:a:3178

|  | Res. type | Atom | Residue | Distance | Occupancy | B-factor | Closest atom(s) to a 1^st^ shell water | EM\_dens. |
| --- | --- | --- | --- | --- | --- | --- | --- | --- |
| **Ion** | | | |  | 1.0 | 30.0 |  | 21.05 |
 1 | A | OP1 | a:687 | 2.71 | 1.0 | 36.2 |  | 8.36 | 2 | U | OP1 | a:688 | 2.81 | 1.0 | 27.8 |  | 13.29 | 3 | Water 1 | | a:9374 | 2.91 | 1.0 | 31.6 |  | 9.89 | 4 | Water 2 | | 1:120 | 2.92 | 1.0 | 37.8 |  | 7.13 | 5 | SER | OG | 1:15 | 3.03 | 1.0 | 35.6 |  | 7.94 | 6 | A | O5' | a:687 | 3.39 | 1.0 | 31.9 |  | 7.69 | 7 | U | O5' | a:688 | 3.58 | 1.0 | 29.0 |  | 13.44 | 8 | Water 3 | | 1:140 | 4.03 | 1.0 | 42.1 |  | 7.73 | 9 | Water 4 | | a:3572 | 4.08 | 1.0 | 35.9 |  | 10.13 | 10 | Water 5 | | a:7385 | 4.12 | 1.0 | 31.2 |  | 12.26 | 11 | A | OP2 | a:687 | 4.30 | 1.0 | 25.5 |  | 12.45 | 12 | Water 6 | | a:6402 | 4.68 | 1.0 | 42.2 |  | 6.73 | 13 | A | O3' | a:687 | 4.70 | 1.0 | 28.6 |  | 15.12 |

  
**1^st^ shell coordination angles and distances (*trans* in "blue"; *cys* in "green"; deviations in "orange"):**   
• 1-2(OP1/OP1): 122; 1-3(OP1/W): 87; 1-4(OP1/W): 132; 1-5(OP1/OG): 77; 2-3(OP1/W): 98; 2-4(OP1/W): 105; 2-5(OP1/OG): 159; 3-4(W/W): 94; 3-5(W/OG): 75; 4-5(W/OG): 57;   
• 1-2(OP1/OP1): 4.8; 1-3(OP1/W): 3.9; 1-4(OP1/W): 5.1; 1-5(OP1/OG): 3.6; 2-3(OP1/W): 4.3; 2-4(OP1/W): 4.6; 2-5(OP1/OG): 5.7; 3-4(W/W): 4.3; 3-5(W/OG): 3.6; 4-5(W/OG): 2.8;   
  
**4.23 ± 1.32: the average coodination distance of the 1^st^ six ligands could correspond to an hexahydrated Mg^2+^.**  
**6: Number of potential Water contacts**   
**0 : Number of second shell contacts involving water or ions**   
**0%: Percentage of satisfied second shell Water contacts**   
• Contact(s): /O_w_/O_w_/OG_Ser_/OP1/OP1/  
• Contact code: 2O_ph_OH_prot_2O_w_  
→ Coordination < 6 for K ←  
Go to ToC

### K:a:3179

|  | Res. type | Atom | Residue | Distance | Occupancy | B-factor | Closest atom(s) to a 1^st^ shell water | EM\_dens. |
| --- | --- | --- | --- | --- | --- | --- | --- | --- |
| **Ion** | | | |  | 1.0 | 30.0 |  | 25.86 |
 1 | U | O4 | a:2244 | 2.72 | 1.0 | 27.5 |  | 14.55 | 2 | Water 1 | | a:6822 | 2.92 | 1.0 | 33.9 |  | 10.57 | 3 | G | O6 | a:2243 | 2.93 | 1.0 | 30.0 |  | 11.20 | 4 | G | O4' | a:2242 | 2.94 | 1.0 | 27.1 |  | 14.51 | 5 | Water 2 | | a:9978 | 2.96 | 1.0 | 37.6 |  | 7.90 | 6 | G | O5' | a:2242 | 3.03 | 1.0 | 33.3 |  | 7.60 | 7 | G | N7 | a:2243 | 3.08 | 1.0 | 23.3 |  | 12.74 | 8 | G | O3' | a:2241 | 3.33 | 1.0 | 33.2 |  | 8.43 | 9 | G | O2' | a:2241 | 3.99 | 1.0 | 35.8 |  | 10.19 | 10 | G | OP2 | a:2242 | 4.25 | 1.0 | 34.9 |  | 8.13 |

  
**1^st^ shell coordination angles and distances (*trans* in "blue"; *cys* in "green"; deviations in "orange"):**   
• 1-2(O4/W): 114; 1-3(O4/O6): 75; 1-4(O4/O4'): 81; 1-5(O4/W): 86; 1-6(O4/O5'): 133; 1-7(O4/N7): 95; 2-3(W/O6): 75; 2-4(W/O4'): 148; 2-5(W/W): 71; 2-6(W/O5'): 100; 2-7(W/N7): 118; 3-4(O6/O4'): 137; 3-5(O6/W): 129; 3-6(O6/O5'): 149; 3-7(O6/N7): 61; 4-5(O4'/W): 83; 4-6(O4'/O5'): 55; 4-7(O4'/N7): 87; 5-6(W/O5'): 76; 5-7(W/N7): 170; 6-7(O5'/N7): 97;   
• 1-2(O4/W): 4.7; 1-3(O4/O6): 3.4; 1-4(O4/O4'): 3.7; 1-5(O4/W): 3.9; 1-6(O4/O5'): 5.3; 1-7(O4/N7): 4.3; 2-3(W/O6): 3.6; 2-4(W/O4'): 5.6; 2-5(W/W): 3.4; 2-6(W/O5'): 4.6; 2-7(W/N7): 5.1; 3-4(O6/O4'): 5.4; 3-5(O6/W): 5.3; 3-6(O6/O5'): 5.7; 3-7(O6/N7): 3.1; 4-5(O4'/W): 3.9; 4-6(O4'/O5'): 2.7; 4-7(O4'/N7): 4.1; 5-6(W/O5'): 3.7; 5-7(W/N7): 6.0; 6-7(O5'/N7): 4.6;   
  
**3.99 ± 1.17: the average coodination distance of the 1^st^ six ligands could correspond to an hexahydrated Mg^2+^.**  
**4: Number of potential Water contacts**   
**0 : Number of second shell contacts involving water or ions**   
**0%: Percentage of satisfied second shell Water contacts**   
• Contact(s): /N7/O4/O4'/O5'/O6/O_w_/O_w_/  
• Contact code: 2O_r_2O_b_N_b_2O_w_  
Go to ToC

### K:a:3180

|  | Res. type | Atom | Residue | Distance | Occupancy | B-factor | Closest atom(s) to a 1^st^ shell water | EM\_dens. |
| --- | --- | --- | --- | --- | --- | --- | --- | --- |
| **Ion** | | | |  | 1.0 | 30.0 |  | 25.84 |
 1 | G | O6 | a:787 | 2.66 | 1.0 | 23.0 |  | 16.68 | 2 | U | O4 | a:781 | 2.81 | 1.0 | 20.7 |  | 17.70 | 3 | G | O6 | a:780 | 2.87 | 1.0 | 20.1 |  | 18.04 | 4 | Water 1 | | a:8021 | 2.95 | 1.0 | 43.6 |  | 5.87 | 5 | Water 2 | | c:402 | 2.98 | 1.0 | 34.5 | 2.33 [ASP:OD1]c:229 | 7.34 | 6 | Water 3 | | a:6352 | 3.03 | 1.0 | 30.9 | 2.76 [G:N7]a:787 | 11.41 | 7 | G | N1 | a:782 | 3.62 | 1.0 | 14.6 |  | 17.62 | 8 | C | N4 | a:788 | 3.69 | 1.0 | 15.2 |  | 16.89 | 9 | Water 4 | | a:7300 | 3.88 | 1.0 | 36.5 | 2.83 [G:N7]a:780 | 6.20 | 10 | Water 5 | | a:6761 | 4.07 | 1.0 | 36.7 | 2.79 [G:O6]a:782 | 7.73 | 11 | G | O6 | a:782 | 4.28 | 1.0 | 23.2 |  | 17.01 | 12 | ASP | OD1 | c:229 | 4.43 | 1.0 | 34.2 |  | 9.97 | 13 | G | N7 | a:787 | 4.43 | 1.0 | 17.5 |  | 15.55 | 14 | G | N3 | a:782 | 4.65 | 1.0 | 15.9 |  | 17.52 | 15 | G | N7 | a:780 | 4.75 | 1.0 | 17.9 |  | 17.58 |

  
**1^st^ shell coordination angles and distances (*trans* in "blue"; *cys* in "green"; deviations in "orange"):**   
• 1-2(O6/O4): 84; 1-3(O6/O6): 76; 1-4(O6/W): 127; 1-5(O6/W): 131; 1-6(O6/W): 80; 2-3(O4/O6): 71; 2-4(O4/W): 106; 2-5(O4/W): 89; 2-6(O4/W): 154; 3-4(O6/W): 59; 3-5(O6/W): 145; 3-6(O6/W): 124; 4-5(W/W): 102; 4-6(W/W): 100; 5-6(W/W): 86;   
• 1-2(O6/O4): 3.7; 1-3(O6/O6): 3.4; 1-4(O6/W): 5.0; 1-5(O6/W): 5.1; 1-6(O6/W): 3.7; 2-3(O4/O6): 3.3; 2-4(O4/W): 4.6; 2-5(O4/W): 4.1; 2-6(O4/W): 5.7; 3-4(O6/W): 2.9; 3-5(O6/W): 5.6; 3-6(O6/W): 5.2; 4-5(W/W): 4.6; 4-6(W/W): 4.6; 5-6(W/W): 4.1;   
  
**4.10 ± 1.25: the average coodination distance of the 1^st^ six ligands could correspond to an hexahydrated Mg^2+^.**  
**6: Number of potential Water contacts**   
**2 : Number of second shell contacts involving water or ions**   
**33%: Percentage of satisfied second shell Water contacts**   
• Contact(s): /O4/O6/O6/O_w_/O_w_/O_w_/  
• Contact code: 3O_b_3O_w_  
• K hydration shell issue: water too close (< 2.4 Å.)  
Go to ToC

### K:a:3181

|  | Res. type | Atom | Residue | Distance | Occupancy | B-factor | Closest atom(s) to a 1^st^ shell water | EM\_dens. |
| --- | --- | --- | --- | --- | --- | --- | --- | --- |
| **Ion** | | | |  | 1.0 | 30.0 |  | 25.41 |
 1 | G | O6 | a:1290 | 2.54 | 1.0 | 31.3 |  | 12.25 | 2 | U | O2 | a:1328 | 2.57 | 1.0 | 27.4 |  | 14.56 | 3 | GLY | O | m:105 | 2.86 | 1.0 | 28.7 |  | 11.99 | 4 | Water 1 | | a:8443 | 2.90 | 1.0 | 30.2 |  | 12.06 | 5 | Water 2 | | a:9749 | 2.99 | 1.0 | 40.2 |  | 5.10 | 6 | A | O4' | a:1329 | 3.09 | 1.0 | 34.6 |  | 12.35 | 7 | Water 3 | | m:253 | 3.23 | 1.0 | 41.2 |  | 4.46 | 8 | A | N6 | a:1289 | 3.31 | 1.0 | 19.4 |  | 13.97 | 9 | U | O2' | a:1328 | 3.68 | 1.0 | 27.7 |  | 11.88 | 10 | Water 4 | | a:4830 | 4.18 | 1.0 | 33.7 |  | 6.49 | 11 | U | N1 | a:1328 | 4.59 | 1.0 | 20.5 |  | 15.12 | 12 | Water 5 | | a:5773 | 4.74 | 1.0 | 44.1 |  | 3.95 |

  
**1^st^ shell coordination angles and distances (*trans* in "blue"; *cys* in "green"; deviations in "orange"):**   
• 1-2(O6/O2): 74; 1-3(O6/W): 92; 1-4(O6/W): 128; 1-5(O6/W): 153; 1-6(O6/O4'): 91; 2-3(O2/W): 129; 2-4(O2/W): 71; 2-5(O2/W): 125; 2-6(O2/O4'): 88; 3-4(W/W): 82; 3-5(W/W): 90; 3-6(W/O4'): 142; 4-5(W/W): 79; 4-6(W/O4'): 125; 5-6(W/O4'): 72;   
• 1-2(O6/O2): 3.1; 1-3(O6/W): 3.9; 1-4(O6/W): 4.9; 1-5(O6/W): 5.4; 1-6(O6/O4'): 4.0; 2-3(O2/W): 4.9; 2-4(O2/W): 3.2; 2-5(O2/W): 4.9; 2-6(O2/O4'): 4.0; 3-4(W/W): 3.8; 3-5(W/W): 4.1; 3-6(W/O4'): 5.6; 4-5(W/W): 3.8; 4-6(W/O4'): 5.3; 5-6(W/O4'): 3.6;   
  
**3.91 ± 1.17: the average coodination distance of the 1^st^ six ligands could correspond to an hexahydrated Mg^2+^.**  
**8: Number of potential Water contacts**   
**0 : Number of second shell contacts involving water or ions**   
**0%: Percentage of satisfied second shell Water contacts**   
• Contact(s): /O4'/O_back_/O_w_/O_w_/  
• Contact code: O_r_O_back_2O_w_  
• Contact(s) in the exclusion zone: /O2/O6/  
→ Coordination < 6 for K ←  
Go to ToC

### MG:a:3182

|  | Res. type | Atom | Residue | Distance | Occupancy | B-factor | Closest atom(s) to a 1^st^ shell water | EM\_dens. |
| --- | --- | --- | --- | --- | --- | --- | --- | --- |
| **Ion** | | | |  | 1.0 | 45.3 |  | 10.46 |
 1 | Water 1 | | a:5963 | 1.97 | 1.0 | 40.8 | 2.74 [C:OP2]a:1994 | 6.07 | 2 | Water 2 | | a:7432 | 2.11 | 1.0 | 41.9 |  | 5.13 | 3 | Water 3 | | a:7886 | 2.12 | 1.0 | 39.8 | 2.87 [C:OP2]a:1994 | 8.72 | 4 | Water 4 | | a:7627 | 2.13 | 1.0 | 33.2 | 2.85 [G:O2']a:1669 | 8.73 | 5 | Water 5 | | a:9812 | 2.19 | 1.0 | 33.3 |  | 7.69 | 6 | Water 6 | | a:9983 | 2.20 | 1.0 | 41.4 |  | 3.53 | 7 | C | OP2 | a:1994 | 3.67 | 1.0 | 37.6 |  | 8.45 | 8 | G | O2' | a:1669 | 4.12 | 1.0 | 23.6 |  | 14.43 | 9 | Water 7 | | a:4135 | 4.32 | 1.0 | 30.7 |  | 10.18 | 10 | Water 8 | | a:3549 | 4.33 | 1.0 | 41.8 |  | 4.70 | 11 | Water 9 | | j:255 | 4.49 | 1.0 | 46.1 |  | 4.40 | 12 | G | OP2 | a:1993 | 4.56 | 1.0 | 38.6 |  | 6.72 | 13 | Water 10 | | a:6760 | 4.61 | 1.0 | 35.4 |  | 9.61 |

  
**1^st^ shell coordination angles and distances (*trans* in "blue"; *cys* in "green"; deviations in "orange"):**   
• 1-2(W/W): 93º; 1-3(W/W): *96º*; 1-4(W/W): 93º; 1-5(W/W): 171º; 1-6(W/W): 92º; 2-3(W/W): *167º*; 2-4(W/W): *97º*; 2-5(W/W): *79º*; 2-6(W/W): *82º*; 3-4(W/W): 92º; 3-5(W/W): 90º; 3-6(W/W): 89º; 4-5(W/W): 94º; 4-6(W/W): 174º; 5-6(W/W): *81º*;   
• 1-2(W/W): 3.0Å; 1-3(W/W): 3.0Å; 1-4(W/W): 3.0Å; 1-5(W/W): 4.1Å; 1-6(W/W): 3.0Å; 2-3(W/W): *4.2Å*; 2-4(W/W): 3.2Å; 2-5(W/W): 2.8Å; 2-6(W/W): 2.8Å; 3-4(W/W): 3.0Å; 3-5(W/W): 3.0Å; 3-6(W/W): 3.0Å; 4-5(W/W): 3.2Å; 4-6(W/W): *4.3Å*; 5-6(W/W): 2.8Å;   
  
**2.12 ± 0.08: the average coodination distance of the 1^st^ six ligands is Mg^2+^ compatible**  
**12: Number of potential Water contacts**   
**3 : Number of second shell contacts involving water or ions**   
**25%: Percentage of satisfied second shell Water contacts**   
• Contact(s): /O_w_/O_w_/O_w_/O_w_/O_w_/O_w_/  
• Contact code: 6O_w_  
→ Slightly distorded coord. octahedron (5.6º) ←  
Go to ToC

### MG:a:3183

|  | Res. type | Atom | Residue | Distance | Occupancy | B-factor | Closest atom(s) to a 1^st^ shell water | EM\_dens. |
| --- | --- | --- | --- | --- | --- | --- | --- | --- |
| **Ion** | | | |  | 1.0 | 31.9 |  | 14.73 |
 1 | Water 1 | | a:5255 | 1.90 | 1.0 | 28.2 | 2.69 [G:N7]a:2413 | 10.15 | 2 | Water 2 | | a:5266 | 1.97 | 1.0 | 31.4 | 2.69 [G:O2']a:2409 // 2.90 [G:N7]a:2414 | 10.69 | 3 | Water 3 | | a:7041 | 2.08 | 1.0 | 29.3 | 2.81 [U:OP2]a:2412 | 9.41 | 4 | Water 4 | | a:7768 | 2.18 | 1.0 | 31.5 | 2.94 [U:OP2]a:2412 | 11.63 | 5 | Water 5 | | a:4112 | 2.19 | 1.0 | 31.1 | 2.59 [A:OP2]a:2410 // 2.94 [G:O6]a:2414 | 9.00 | 6 | A | OP1 | a:2410 | 2.26 | 1.0 | 37.9 |  | 8.53 | 7 | A | OP2 | a:2410 | 3.59 | 1.0 | 33.8 |  | 9.52 | 8 | Water 6 | | a:8866 | 3.87 | 1.0 | 27.6 | 2.97 [U:O4]a:2412 | 10.50 | 9 | U | OP2 | a:2412 | 3.98 | 1.0 | 39.6 |  | 7.81 | 10 | G | O2' | a:2409 | 4.09 | 1.0 | 39.5 |  | 7.96 | 11 | G | N7 | a:2413 | 4.17 | 1.0 | 29.7 |  | 9.75 | 12 | G | N7 | a:2414 | 4.19 | 1.0 | 29.9 |  | 10.00 | 13 | G | O6 | a:2414 | 4.33 | 1.0 | 36.5 |  | 10.35 | 14 | G | O3' | a:2409 | 4.43 | 1.0 | 35.9 |  | 8.62 | 15 | A | O5' | a:2410 | 4.46 | 1.0 | 38.8 |  | 9.66 | 16 | G | O6 | a:2413 | 4.52 | 1.0 | 36.1 |  | 9.69 | 17 | U | O4 | a:2412 | 4.63 | 1.0 | 34.6 |  | 11.24 | 18 | G | OP2 | a:2413 | 4.71 | 1.0 | 39.9 |  | 6.54 |

  
**1^st^ shell coordination angles and distances (*trans* in "blue"; *cys* in "green"; deviations in "orange"):**   
• 1-2(W/W): 93º; 1-3(W/W): 94º; 1-4(W/W): *99º*; 1-5(W/W): 90º; 1-6(W/OP1): 177º; 2-3(W/W): 92º; 2-4(W/W): *168º*; 2-5(W/W): 94º; 2-6(W/OP1): *84º*; 3-4(W/W): 86º; 3-5(W/W): 172º; 3-6(W/OP1): 86º; 4-5(W/W): 87º; 4-6(W/OP1): *84º*; 5-6(W/OP1): 90º;   
• 1-2(W/W): 2.8Å; 1-3(W/W): 2.9Å; 1-4(W/W): 3.1Å; 1-5(W/W): 2.9Å; 1-6(W/OP1): 4.2Å; 2-3(W/W): 2.9Å; 2-4(W/W): 4.1Å; 2-5(W/W): 3.1Å; 2-6(W/OP1): 2.8Å; 3-4(W/W): 2.9Å; 3-5(W/W): *4.3Å*; 3-6(W/OP1): 3.0Å; 4-5(W/W): 3.0Å; 4-6(W/OP1): 3.0Å; 5-6(W/OP1): 3.1Å;   
  
**2.09 ± 0.13: the average coodination distance of the 1^st^ six ligands is Mg^2+^ compatible**  
**10: Number of potential Water contacts**   
**7 : Number of second shell contacts involving water or ions**   
**70%: Percentage of satisfied second shell Water contacts**   
• Contact(s): /O_w_/O_w_/O_w_/O_w_/OP1/  
• Contact code: O_ph_4O_w_  
• Contact(s) in the exclusion zone: /O_w_/  
→ Missing octahedral coordination for MG ←  
Ooctahedral coordination (4.6º)   
Go to ToC

### MG:a:3184

|  | Res. type | Atom | Residue | Distance | Occupancy | B-factor | Closest atom(s) to a 1^st^ shell water | EM\_dens. |
| --- | --- | --- | --- | --- | --- | --- | --- | --- |
| **Ion** | | | |  | 1.0 | 31.8 |  | 20.47 |
 1 | Water 1 | | a:8994 | 2.13 | 1.0 | 26.6 |  | 11.06 | 2 | Water 2 | | a:3447 | 2.13 | 1.0 | 28.3 | 2.40 [G:O6]a:2006 | 13.24 | 3 | Water 3 | | a:9854 | 2.19 | 1.0 | 29.0 |  | 11.80 | 4 | Water 4 | | a:5890 | 2.21 | 1.0 | 29.1 | 2.73 [G:N7]a:2006 | 12.26 | 5 | Water 5 | | a:10271 | 2.23 | 1.0 | 29.0 |  | 11.56 | 6 | Water 6 | | a:4621 | 2.32 | 1.0 | 26.5 | 2.64 [G:O6]a:2008 | 14.40 | 7 | Water 7 | | a:8907 | 3.90 | 1.0 | 30.2 |  | 8.06 | 8 | G | O6 | a:2006 | 3.92 | 1.0 | 23.3 |  | 17.46 | 9 | Water 8 | | a:5625 | 4.01 | 1.0 | 30.0 |  | 13.15 | 10 | Water 9 | | m:208 | 4.07 | 1.0 | 36.6 |  | 6.88 | 11 | G | N7 | a:2006 | 4.15 | 1.0 | 18.0 |  | 16.28 | 12 | Water 10 | | a:7312 | 4.16 | 1.0 | 31.4 |  | 10.05 | 13 | Water 11 | | a:9896 | 4.21 | 1.0 | 33.1 |  | 7.72 | 14 | Water 12 | | a:5660 | 4.28 | 1.0 | 37.4 |  | 8.44 | 15 | Water 13 | | a:8076 | 4.45 | 1.0 | 31.1 |  | 8.59 | 16 | Water 14 | | a:9855 | 4.49 | 1.0 | 43.4 |  | 5.25 | 17 | Water 15 | | a:8172 | 4.54 | 1.0 | 26.0 |  | 13.67 | 18 | G | O6 | a:2008 | 4.72 | 1.0 | 23.2 |  | 15.84 |

  
**1^st^ shell coordination angles and distances (*trans* in "blue"; *cys* in "green"; deviations in "orange"):**   
• 1-2(W/W): *96º*; 1-3(W/W): *81º*; 1-4(W/W): 94º; 1-5(W/W): *168º*; 2-3(W/W): 176º; 2-4(W/W): 86º; 2-5(W/W): 94º; 3-4(W/W): 93º; 3-5(W/W): 89º; 4-5(W/W): 94º;   
• 1-2(W/W): 3.2Å; 1-3(W/W): 2.8Å; 1-4(W/W): 3.2Å; 1-5(W/W): *4.3Å*; 2-3(W/W): *4.3Å*; 2-4(W/W): 3.0Å; 2-5(W/W): *3.2Å*; 3-4(W/W): 3.2Å; 3-5(W/W): 3.1Å; 4-5(W/W): *3.2Å*;   
  
**2.20 ± 0.06: the average coodination distance of the 1^st^ six ligands is Mg^2+^ compatible**  
**12: Number of potential Water contacts**   
**3 : Number of second shell contacts involving water or ions**   
**25%: Percentage of satisfied second shell Water contacts**   
• Contact(s): /O_w_/O_w_/O_w_/O_w_/O_w_/  
• Contact code: 5O_w_  
• Contact(s) in the exclusion zone: /O_w_/  
→ Missing octahedral coordination for MG ←  
• MG hydration shell issue: water too close (< 2.4 Å.)  
→ Slightly distorded coord. octahedron (5.1º) ←  
Go to ToC

### MG:a:3185

|  | Res. type | Atom | Residue | Distance | Occupancy | B-factor | Closest atom(s) to a 1^st^ shell water | EM\_dens. |
| --- | --- | --- | --- | --- | --- | --- | --- | --- |
| **Ion** | | | |  | 1.0 | 42.5 |  | 10.66 |
 1 | Water 1 | | a:5272 | 1.93 | 1.0 | 32.1 | 2.87 [G:O2']a:1238 | 7.09 | 2 | A | OP2 | a:1239 | 1.93 | 1.0 | 35.2 |  | 6.95 | 3 | Water 2 | | a:7902 | 2.16 | 1.0 | 31.5 |  | 7.06 | 4 | Water 3 | | a:5608 | 2.17 | 1.0 | 40.0 | 2.72 [G:O6]a:1212 | 7.61 | 5 | Water 4 | | a:8147 | 2.23 | 1.0 | 32.0 | 2.95 [A:O4']a:330 | 9.25 | 6 | Water 5 | | a:4079 | 2.29 | 1.0 | 34.7 | 2.59 [A:OP1]a:330 | 8.73 | 7 | A | O5' | a:1239 | 3.74 | 1.0 | 34.5 |  | 8.53 | 8 | G | O3' | a:1238 | 3.94 | 1.0 | 35.8 |  | 9.34 | 9 | Water 6 | | a:6783 | 3.96 | 1.0 | 38.7 | 2.85 [A:OP1]a:1239 | 6.66 | 10 | Water 7 | | a:7098 | 4.10 | 1.0 | 42.1 |  | 4.97 | 11 | G | O2' | a:1238 | 4.16 | 1.0 | 33.3 |  | 9.72 | 12 | G | N7 | a:1212 | 4.24 | 1.0 | 28.9 |  | 9.13 | 13 | Water 8 | | a:9947 | 4.25 | 1.0 | 41.6 |  | 5.07 | 14 | A | OP1 | a:330 | 4.36 | 1.0 | 43.5 |  | 5.62 | 15 | A | O5' | a:330 | 4.40 | 1.0 | 39.0 |  | 5.97 | 16 | G | O6 | a:1212 | 4.43 | 1.0 | 36.2 |  | 8.07 | 17 | A | OP1 | a:1239 | 4.44 | 1.0 | 36.5 |  | 8.24 | 18 | A | O4' | a:330 | 4.45 | 1.0 | 41.7 |  | 8.05 | 19 | Water 9 | | a:5148 | 4.62 | 1.0 | 36.2 | 2.75 [A:OP1]a:330 | 7.79 | 20 | Water 10 | | a:6045 | 4.65 | 1.0 | 41.3 |  | 6.29 |

  
**1^st^ shell coordination angles and distances (*trans* in "blue"; *cys* in "green"; deviations in "orange"):**   
• 1-2(W/OP2): 89º; 1-3(W/W): 90º; 1-4(W/W): 90º; 1-5(W/W): 89º; 1-6(W/W): 177º; 2-3(OP2/W): 89º; 2-4(OP2/W): *108º*; 2-5(OP2/W): 171º; 2-6(OP2/W): 91º; 3-4(W/W): *163º*; 3-5(W/W): *82º*; 3-6(W/W): 93º; 4-5(W/W): *81º*; 4-6(W/W): 87º; 5-6(W/W): 92º;   
• 1-2(W/OP2): 2.7Å; 1-3(W/W): 2.9Å; 1-4(W/W): 2.9Å; 1-5(W/W): 2.9Å; 1-6(W/W): *4.2Å*; 2-3(OP2/W): 2.9Å; 2-4(OP2/W): *3.3Å*; 2-5(OP2/W): 4.1Å; 2-6(OP2/W): 3.0Å; 3-4(W/W): *4.3Å*; 3-5(W/W): 2.9Å; 3-6(W/W): *3.2Å*; 4-5(W/W): 2.8Å; 4-6(W/W): 3.1Å; 5-6(W/W): *3.2Å*;   
  
**2.12 ± 0.14: the average coodination distance of the 1^st^ six ligands is Mg^2+^ compatible**  
**10: Number of potential Water contacts**   
**4 : Number of second shell contacts involving water or ions**   
**40%: Percentage of satisfied second shell Water contacts**   
• Contact(s): /O_w_/O_w_/O_w_/O_w_/O_w_/OP2/  
• Contact code: O_ph_5O_w_  
→ Slightly distorded coord. octahedron (5.1º) ←  
Go to ToC

### MG:a:3186

|  | Res. type | Atom | Residue | Distance | Occupancy | B-factor | Closest atom(s) to a 1^st^ shell water | EM\_dens. |
| --- | --- | --- | --- | --- | --- | --- | --- | --- |
| **Ion** | | | |  | 1.0 | 26.6 |  | 22.58 |
 1 | A | OP2 | a:2825 | 1.97 | 1.0 | 28.3 |  | 13.89 | 2 | G | OP2 | a:2826 | 1.97 | 1.0 | 23.5 |  | 14.90 | 3 | GLY | O | d:115 | 2.20 | 1.0 | 21.9 |  | 17.15 | 4 | HIS | NE2 | m:3 | 2.26 | 1.0 | 25.1 |  | 12.95 | 5 | Water 1 | | a:6128 | 2.26 | 1.0 | 22.3 | 2.75 [G:N7]a:2826 | 16.02 | 6 | Water 2 | | a:8723 | 2.29 | 1.0 | 24.0 |  | 16.18 | 7 | A | O3' | a:2824 | 3.54 | 1.0 | 33.2 |  | 10.24 | 8 | A | O3' | a:2825 | 4.06 | 1.0 | 20.4 |  | 17.22 | 9 | A | O5' | a:2825 | 4.14 | 1.0 | 24.4 |  | 14.44 | 10 | G | OP1 | a:2826 | 4.24 | 1.0 | 26.2 |  | 16.30 | 11 | HIS | ND1 | m:3 | 4.28 | 1.0 | 18.0 |  | 15.65 | 12 | G | O5' | a:2826 | 4.31 | 1.0 | 23.4 |  | 13.23 | 13 | A | OP1 | a:2825 | 4.39 | 1.0 | 28.1 |  | 9.66 | 14 | G | N7 | a:2826 | 4.39 | 1.0 | 20.9 |  | 15.38 | 15 | Water 3 | | m:243 | 4.50 | 1.0 | 25.9 |  | 12.29 | 16 | Water 4 | | a:6640 | 4.77 | 1.0 | 29.1 |  | 12.82 |

  
**1^st^ shell coordination angles and distances (*trans* in "blue"; *cys* in "green"; deviations in "orange"):**   
• 1-2(OP2/OP2): 90º; 1-3(OP2/W): 87º; 1-4(OP2/NE2): *96º*; 1-5(OP2/W): 93º; 1-6(OP2/W): 177º; 2-3(OP2/W): 89º; 2-4(OP2/NE2): 174º; 2-5(OP2/W): 94º; 2-6(OP2/W): 87º; 3-4(W/NE2): 89º; 3-5(W/W): 177º; 3-6(W/W): 95º; 4-5(NE2/W): 88º; 4-6(NE2/W): 87º; 5-6(W/W): 85º;   
• 1-2(OP2/OP2): 2.8Å; 1-3(OP2/W): 2.9Å; 1-4(OP2/NE2): 3.1Å; 1-5(OP2/W): 3.1Å; 1-6(OP2/W): *4.3Å*; 2-3(OP2/W): 2.9Å; 2-4(OP2/NE2): *4.2Å*; 2-5(OP2/W): 3.1Å; 2-6(OP2/W): 3.0Å; 3-4(W/NE2): 3.1Å; 3-5(W/W): *4.5Å*; 3-6(W/W): *3.3Å*; 4-5(NE2/W): 3.1Å; 4-6(NE2/W): 3.1Å; 5-6(W/W): 3.1Å;   
  
**2.16 ± 0.14: the average coodination distance of the 1^st^ six ligands is Mg^2+^ compatible**  
**6: Number of potential Water contacts**   
**1 : Number of second shell contacts involving water or ions**   
**17%: Percentage of satisfied second shell Water contacts**   
• Contact(s): /NE2_His_/O_back_/O_w_/O_w_/OP2/OP2/  
• Contact code: *cis-*2O_ph_O_back_2O_w_N_His_  
Ooctahedral coordination (3.2º)   
Go to ToC

### MG:a:3187

|  | Res. type | Atom | Residue | Distance | Occupancy | B-factor | Closest atom(s) to a 1^st^ shell water | EM\_dens. |
| --- | --- | --- | --- | --- | --- | --- | --- | --- |
| **Ion** | | | |  | 1.0 | 45.5 |  | 10.38 |
 1 | C | OP1 | a:2389 | 1.89 | 1.0 | 41.1 |  | 7.32 | 2 | Water 1 | | a:9299 | 1.95 | 1.0 | 39.6 |  | 6.08 | 3 | Water 2 | | a:8233 | 2.01 | 1.0 | 36.5 |  | 8.02 | 4 | Water 3 | | a:3864 | 2.13 | 1.0 | 35.8 | 2.78 [G:OP1]a:2341 | 8.18 | 5 | G | OP1 | a:2293 | 2.16 | 1.0 | 38.1 |  | 7.77 | 6 | Water 4 | | a:4131 | 2.42 | 1.0 | 43.3 | 2.59 [U:O2']a:2388 | 7.03 | 7 | U | O3' | a:2388 | 3.28 | 1.0 | 35.2 |  | 7.26 | 8 | Water 5 | | a:3913 | 3.81 | 1.0 | 37.8 |  | 8.31 | 9 | C | O5' | a:2389 | 3.91 | 1.0 | 40.2 |  | 6.22 | 10 | A | O3' | a:2292 | 3.97 | 1.0 | 34.7 |  | 8.19 | 11 | Water 6 | | a:5395 | 4.18 | 1.0 | 38.6 |  | 4.34 | 12 | G | OP1 | a:2341 | 4.30 | 1.0 | 41.9 |  | 6.34 | 13 | C | OP2 | a:2389 | 4.31 | 1.0 | 34.4 |  | 9.30 | 14 | Water 7 | | a:3559 | 4.36 | 1.0 | 42.7 | 2.47 [G:OP1]a:2329 | 4.74 | 15 | Water 8 | | a:8658 | 4.40 | 1.0 | 43.6 |  | 6.56 | 16 | Water 9 | | a:9728 | 4.42 | 1.0 | 47.1 |  | 3.41 | 17 | G | OP2 | a:2293 | 4.43 | 1.0 | 39.0 |  | 6.60 | 18 | G | OP1 | a:2329 | 4.52 | 1.0 | 47.9 |  | 3.31 | 19 | G | O5' | a:2293 | 4.55 | 1.0 | 37.9 |  | 8.58 | 20 | U | O2' | a:2388 | 4.76 | 1.0 | 34.0 |  | 9.53 |

  
**1^st^ shell coordination angles and distances (*trans* in "blue"; *cys* in "green"; deviations in "orange"):**   
• 1-2(OP1/W): *160º*; 1-3(OP1/W): *96º*; 1-4(OP1/W): *78º*; 1-5(OP1/OP1): *104º*; 2-3(W/W): 87º; 2-4(W/W): *83º*; 2-5(W/OP1): *96º*; 3-4(W/W): *80º*; 3-5(W/OP1): 89º; 4-5(W/OP1): *170º*;   
• 1-2(OP1/W): *3.8Å*; 1-3(OP1/W): 2.9Å; 1-4(OP1/W): 2.5Å; 1-5(OP1/OP1): 3.2Å; 2-3(W/W): 2.7Å; 2-4(W/W): 2.7Å; 2-5(W/OP1): 3.0Å; 3-4(W/W): 2.7Å; 3-5(W/OP1): 2.9Å; 4-5(W/OP1): *4.3Å*;   
  
**2.09 ± 0.17: the average coodination distance of the 1^st^ six ligands is Mg^2+^ compatible**  
**8: Number of potential Water contacts**   
**2 : Number of second shell contacts involving water or ions**   
**25%: Percentage of satisfied second shell Water contacts**   
• Contact(s): /O_w_/O_w_/O_w_/OP1/  
• Contact code: O_ph_3O_w_  
• Contact(s) in the exclusion zone: /O3'/O_w_/OP1/  
→ Missing octahedral coordination for MG ←  
→ Slightly distorded coord. octahedron (8.8º) ←  
Go to ToC

### MG:a:3188

|  | Res. type | Atom | Residue | Distance | Occupancy | B-factor | Closest atom(s) to a 1^st^ shell water | EM\_dens. |
| --- | --- | --- | --- | --- | --- | --- | --- | --- |
| **Ion** | | | |  | 1.0 | 42.1 |  | 13.47 |
 1 | Water 1 | | s:212 | 1.95 | 1.0 | 36.9 | 2.84 [ASN:OD1]s:59 | 7.62 | 2 | Water 2 | | a:6797 | 2.02 | 1.0 | 36.3 | 2.79 [C:OP2]a:1401 | 7.39 | 3 | Water 3 | | a:8749 | 2.02 | 1.0 | 36.2 | 2.96 [G:OP2]a:1345 | 7.50 | 4 | C | OP1 | a:1400 | 2.05 | 1.0 | 35.5 |  | 10.08 | 5 | Water 4 | | a:8740 | 2.20 | 1.0 | 34.6 | 2.96 [G:OP1]a:1345 | 8.25 | 6 | Water 5 | | a:10213 | 2.27 | 1.0 | 41.3 |  | 5.55 | 7 | C | OP2 | a:1400 | 3.67 | 1.0 | 35.0 |  | 10.19 | 8 | ASN | OD1 | s:59 | 4.01 | 1.0 | 42.3 |  | 6.31 | 9 | C | O5' | a:1400 | 4.24 | 1.0 | 30.7 |  | 12.08 | 10 | U | O3' | a:1399 | 4.39 | 1.0 | 30.5 |  | 9.57 | 11 | Water 6 | | a:3518 | 4.49 | 1.0 | 43.3 |  | 3.51 | 12 | G | OP1 | a:1345 | 4.60 | 1.0 | 35.3 |  | 7.99 | 13 | C | OP2 | a:1401 | 4.61 | 1.0 | 40.9 |  | 7.73 | 14 | G | OP2 | a:1345 | 4.63 | 1.0 | 34.0 |  | 7.98 | 15 | Water 7 | | s:211 | 4.76 | 1.0 | 37.4 |  | 7.33 |

  
**1^st^ shell coordination angles and distances (*trans* in "blue"; *cys* in "green"; deviations in "orange"):**   
• 1-2(W/W): 90º; 1-3(W/W): 91º; 1-4(W/OP1): 92º; 1-5(W/W): 178º; 1-6(W/W): *96º*; 2-3(W/W): 171º; 2-4(W/OP1): 87º; 2-5(W/W): 89º; 2-6(W/W): 87º; 3-4(W/OP1): *102º*; 3-5(W/W): 89º; 3-6(W/W): *84º*; 4-5(OP1/W): 90º; 4-6(OP1/W): 170º; 5-6(W/W): *82º*;   
• 1-2(W/W): 2.8Å; 1-3(W/W): 2.8Å; 1-4(W/OP1): 2.9Å; 1-5(W/W): 4.2Å; 1-6(W/W): 3.2Å; 2-3(W/W): 4.0Å; 2-4(W/OP1): 2.8Å; 2-5(W/W): 3.0Å; 2-6(W/W): 3.0Å; 3-4(W/OP1): 3.2Å; 3-5(W/W): 3.0Å; 3-6(W/W): 2.9Å; 4-5(OP1/W): 3.0Å; 4-6(OP1/W): *4.3Å*; 5-6(W/W): 2.9Å;   
  
**2.09 ± 0.11: the average coodination distance of the 1^st^ six ligands is Mg^2+^ compatible**  
**10: Number of potential Water contacts**   
**4 : Number of second shell contacts involving water or ions**   
**40%: Percentage of satisfied second shell Water contacts**   
• Contact(s): /O_w_/O_w_/O_w_/O_w_/O_w_/OP1/  
• Contact code: O_ph_5O_w_  
Ooctahedral coordination (4.2º)   
Go to ToC

### MG:a:3189

|  | Res. type | Atom | Residue | Distance | Occupancy | B-factor | Closest atom(s) to a 1^st^ shell water | EM\_dens. |
| --- | --- | --- | --- | --- | --- | --- | --- | --- |
| **Ion** | | | |  | 1.0 | 52.9 |  | 5.94 |
 1 | Water 1 | | a:6114 | 2.04 | 1.0 | 41.6 | 2.75 [G:O6]a:2298 | 7.89 | 2 | Water 2 | | a:6161 | 2.16 | 1.0 | 44.4 | 2.75 [G:N7]a:2298 | 5.88 | 3 | Water 3 | | n:203 | 2.50 | 1.0 | 52.6 |  | 3.85 | 4 | G | N7 | a:2297 | 3.77 | 1.0 | 43.1 |  | 6.82 | 5 | Water 4 | | a:6498 | 3.84 | 1.0 | 43.1 |  | 5.07 | 6 | Water 5 | | a:7724 | 4.25 | 1.0 | 48.2 |  | 4.75 | 7 | G | N7 | a:2298 | 4.26 | 1.0 | 38.0 |  | 7.00 | 8 | G | O6 | a:2298 | 4.29 | 1.0 | 46.9 |  | 5.83 | 9 | G | O6 | a:2297 | 4.34 | 1.0 | 43.9 |  | 6.16 | 10 | Water 6 | | a:4195 | 4.41 | 1.0 | 46.4 |  | 4.06 | 11 | A | OP1 | a:2339 | 4.42 | 1.0 | 43.8 |  | 3.85 | 12 | Water 7 | | a:8149 | 4.68 | 1.0 | 46.9 |  | 3.57 |

  
**1^st^ shell coordination angles and distances (*trans* in "blue"; *cys* in "green"; deviations in "orange"):**   
• 1-2(W/W): *95º*;   
• 1-2(W/W): 3.1Å;   
  
**3.09 ± 0.88: the average coodination distance of the 1^st^ six ligands is K^+^ compatible**  
**10: Number of potential Water contacts**   
**2 : Number of second shell contacts involving water or ions**   
**20%: Percentage of satisfied second shell Water contacts**   
• Contact(s): /O_w_/O_w_/  
• Contact code: 2O_w_  
• Contact(s) in the exclusion zone: /O_w_/  
→ Missing octahedral coordination for MG ←  
Go to ToC

### MG:a:3190

|  | Res. type | Atom | Residue | Distance | Occupancy | B-factor | Closest atom(s) to a 1^st^ shell water | EM\_dens. |
| --- | --- | --- | --- | --- | --- | --- | --- | --- |
| **Ion** | | | |  | 1.0 | 51.2 |  | 5.95 |
 1 | Water 1 | | a:6716 | 2.00 | 1.0 | 41.7 | 2.79 [G:N7]a:178 | 6.04 | 2 | Water 2 | | a:9466 | 2.10 | 1.0 | 40.9 |  | 5.04 | 3 | Water 3 | | a:3579 | 2.30 | 1.0 | 42.6 | 2.48 [G:OP1]a:177 | 4.45 | 4 | G | OP1 | a:177 | 4.07 | 1.0 | 40.8 |  | 5.93 | 5 | C | N4 | a:179 | 4.08 | 1.0 | 28.9 |  | 8.08 | 6 | A | OP2 | a:213 | 4.17 | 1.0 | 48.2 |  | 2.93 | 7 | Water 4 | | a:5378 | 4.24 | 1.0 | 41.4 | 2.90 [G:OP2]a:178 | 6.44 | 8 | G | N7 | a:178 | 4.27 | 1.0 | 28.0 |  | 9.36 | 9 | G | OP2 | a:178 | 4.53 | 1.0 | 38.2 |  | 7.02 | 10 | G | O6 | a:178 | 4.61 | 1.0 | 36.7 |  | 9.73 |

  
**1^st^ shell coordination angles and distances (*trans* in "blue"; *cys* in "green"; deviations in "orange"):**   
• 1-2(W/W): *84º*; 1-3(W/W): 86º; 2-3(W/W): *74º*;   
• 1-2(W/W): 2.8Å; 1-3(W/W): 2.9Å; 2-3(W/W): 2.6Å;   
  
**3.12 ± 0.99: the average coodination distance of the 1^st^ six ligands is K^+^ compatible**  
**6: Number of potential Water contacts**   
**2 : Number of second shell contacts involving water or ions**   
**33%: Percentage of satisfied second shell Water contacts**   
• Contact(s): /O_w_/O_w_/O_w_/  
• Contact code: 3O_w_  
→ Missing octahedral coordination for MG ←  
Go to ToC

### MG:a:3191

|  | Res. type | Atom | Residue | Distance | Occupancy | B-factor | Closest atom(s) to a 1^st^ shell water | EM\_dens. |
| --- | --- | --- | --- | --- | --- | --- | --- | --- |
| **Ion** | | | |  | 1.0 | 58.8 |  | 3.43 |
 1 | Water 1 | | a:4776 | 2.24 | 1.0 | 45.9 | 2.66 [U:O4]a:1711 | 4.28 | 2 | Water 2 | | a:4353 | 3.91 | 1.0 | 46.1 |  | 3.73 | 3 | U | O4 | a:1711 | 4.03 | 1.0 | 41.7 |  | 6.04 | 4 | Water 3 | | a:7392 | 4.26 | 1.0 | 43.8 |  | 5.14 | 5 | U | O4 | a:1749 | 4.33 | 1.0 | 48.7 |  | 4.82 | 6 | G | O6 | a:1712 | 4.45 | 1.0 | 46.3 |  | 5.74 |

  
• • **3.87 ± 0.75: the average coodination distance of the 1^st^ six ligands could correspond to an hexahydrated Mg^2+^.**  
**6: Number of potential Water contacts**   
**1 : Number of second shell contacts involving water or ions**   
**17%: Percentage of satisfied second shell Water contacts**   
• Contact(s): /O_w_/  
• Contact code: O_w_  
→ Missing octahedral coordination for MG ←  
→2Fo-Fc density between 1.0 and 4.0 !   
Go to ToC

### MG:a:3192

|  | Res. type | Atom | Residue | Distance | Occupancy | B-factor | Closest atom(s) to a 1^st^ shell water | EM\_dens. |
| --- | --- | --- | --- | --- | --- | --- | --- | --- |
| **Ion** | | | |  | 1.0 | 41.5 |  | 13.91 |
 1 | Water 1 | | a:4610 | 1.90 | 1.0 | 35.0 | 2.64 [A:OP1]a:992 | 7.99 | 2 | G | OP1 | a:1159 | 2.04 | 1.0 | 31.2 |  | 9.48 | 3 | Water 2 | | a:7836 | 2.05 | 1.0 | 38.8 |  | 5.92 | 4 | C | OP2 | a:1160 | 2.18 | 1.0 | 32.7 |  | 9.71 | 5 | Water 3 | | a:4374 | 2.48 | 1.0 | 33.8 | 2.62 [G:OP2]a:1159 | 7.33 | 6 | G | OP2 | a:1159 | 3.57 | 1.0 | 25.8 |  | 11.72 | 7 | Water 4 | | a:8904 | 3.81 | 1.0 | 37.1 |  | 7.24 | 8 | A | OP1 | a:992 | 4.07 | 1.0 | 32.6 |  | 10.40 | 9 | G | O5' | a:1159 | 4.12 | 1.0 | 29.1 |  | 12.44 | 10 | C | OP1 | a:1160 | 4.14 | 1.0 | 35.1 |  | 7.35 | 11 | G | O3' | a:1159 | 4.30 | 1.0 | 31.3 |  | 11.29 | 12 | A | O3' | a:1158 | 4.41 | 1.0 | 32.3 |  | 10.10 | 13 | Water 5 | | a:10110 | 4.50 | 1.0 | 48.2 |  | 3.47 | 14 | C | O5' | a:1160 | 4.60 | 1.0 | 30.5 |  | 11.63 | 15 | C | OP1 | a:993 | 4.73 | 1.0 | 28.5 |  | 13.42 |

  
**1^st^ shell coordination angles and distances (*trans* in "blue"; *cys* in "green"; deviations in "orange"):**   
• 1-2(W/OP1): 91º; 1-3(W/W): 91º; 1-4(W/OP2): 87º; 2-3(OP1/W): 177º; 2-4(OP1/OP2): *98º*; 3-4(W/OP2): 85º;   
• 1-2(W/OP1): 2.8Å; 1-3(W/W): 2.8Å; 1-4(W/OP2): 2.8Å; 2-3(OP1/W): 4.1Å; 2-4(OP1/OP2): 3.2Å; 3-4(W/OP2): 2.9Å;   
  
**2.37 ± 0.57: the average coodination distance of the 1^st^ six ligands is Na^+^** compatible  
**6: Number of potential Water contacts**   
**2 : Number of second shell contacts involving water or ions**   
**33%: Percentage of satisfied second shell Water contacts**   
• Contact(s): /O_w_/OP1/OP2/  
• Contact code: *cis-*2O_ph_O_w_  
• Contact(s) in the exclusion zone: /O_w_/O_w_/  
→ Missing octahedral coordination for MG ←  
Ooctahedral coordination (3.3º)   
Go to ToC

### MG:a:3193

|  | Res. type | Atom | Residue | Distance | Occupancy | B-factor | Closest atom(s) to a 1^st^ shell water | EM\_dens. |
| --- | --- | --- | --- | --- | --- | --- | --- | --- |
| **Ion** | | | |  | 1.0 | 44.8 |  | 11.00 |
 1 | C | OP1 | a:1378 | 1.95 | 1.0 | 36.1 |  | 7.83 | 2 | Water 1 | | a:6249 | 2.01 | 1.0 | 35.8 | 2.88 [G:O2']a:122 // 2.99 [G:N3]a:122 | 6.89 | 3 | Water 2 | | a:9407 | 2.04 | 1.0 | 37.2 |  | 7.27 | 4 | Water 3 | | a:9376 | 2.15 | 1.0 | 38.3 |  | 7.85 | 5 | Water 4 | | a:7839 | 2.22 | 1.0 | 33.9 | 2.87 [G:O2']a:123 | 9.54 | 6 | Water 5 | | a:8308 | 2.25 | 1.0 | 40.4 |  | 6.20 | 7 | C | OP2 | a:1378 | 4.04 | 1.0 | 32.9 |  | 10.82 | 8 | G | O4' | a:123 | 4.13 | 1.0 | 29.9 |  | 12.96 | 9 | C | O5' | a:1378 | 4.15 | 1.0 | 33.1 |  | 10.58 | 10 | Water 6 | | a:6414 | 4.32 | 1.0 | 42.0 |  | 6.65 | 11 | U | O3' | a:1377 | 4.36 | 1.0 | 30.8 |  | 10.88 | 12 | Water 7 | | a:6737 | 4.41 | 1.0 | 40.9 |  | 6.74 | 13 | Water 8 | | a:3649 | 4.47 | 1.0 | 40.2 |  | 5.69 | 14 | Water 9 | | a:4130 | 4.54 | 1.0 | 39.1 | 2.59 [C:OP2]a:1378 | 6.20 | 15 | G | N3 | a:122 | 4.54 | 1.0 | 26.3 |  | 11.92 | 16 | G | O2' | a:122 | 4.55 | 1.0 | 38.3 |  | 7.75 | 17 | Water 10 | | a:5848 | 4.57 | 1.0 | 38.5 |  | 7.12 | 18 | Water 11 | | a:8677 | 4.65 | 1.0 | 30.2 | 2.95 [C:OP2]a:1378 | 12.94 | 19 | G | O2' | a:123 | 4.68 | 1.0 | 30.2 |  | 11.31 |

  
**1^st^ shell coordination angles and distances (*trans* in "blue"; *cys* in "green"; deviations in "orange"):**   
• 1-2(OP1/W): 88º; 1-3(OP1/W): *101º*; 1-4(OP1/W): *170º*; 1-5(OP1/W): 91º; 1-6(OP1/W): 88º; 2-3(W/W): *165º*; 2-4(W/W): *83º*; 2-5(W/W): *101º*; 2-6(W/W): *83º*; 3-4(W/W): 89º; 3-5(W/W): 90º; 3-6(W/W): 87º; 4-5(W/W): 85º; 4-6(W/W): *97º*; 5-6(W/W): 176º;   
• 1-2(OP1/W): 2.8Å; 1-3(OP1/W): 3.1Å; 1-4(OP1/W): 4.1Å; 1-5(OP1/W): 3.0Å; 1-6(OP1/W): 2.9Å; 2-3(W/W): 4.0Å; 2-4(W/W): 2.7Å; 2-5(W/W): *3.3Å*; 2-6(W/W): 2.8Å; 3-4(W/W): 2.9Å; 3-5(W/W): 3.0Å; 3-6(W/W): 2.9Å; 4-5(W/W): 3.0Å; 4-6(W/W): *3.3Å*; 5-6(W/W): *4.5Å*;   
  
**2.10 ± 0.11: the average coodination distance of the 1^st^ six ligands is Mg^2+^ compatible**  
**10: Number of potential Water contacts**   
**3 : Number of second shell contacts involving water or ions**   
**30%: Percentage of satisfied second shell Water contacts**   
• Contact(s): /O_w_/O_w_/O_w_/O_w_/O_w_/OP1/  
• Contact code: O_ph_5O_w_  
→ Slightly distorded coord. octahedron (5.8º) ←  
Go to ToC

### MG:a:3194

|  | Res. type | Atom | Residue | Distance | Occupancy | B-factor | Closest atom(s) to a 1^st^ shell water | EM\_dens. |
| --- | --- | --- | --- | --- | --- | --- | --- | --- |
| **Ion** | | | |  | 1.0 | 27.2 |  | 28.04 |
 1 | Water 1 | | a:4523 | 2.03 | 1.0 | 22.2 | 2.63 [G:OP2]a:2057 | 17.29 | 2 | Water 2 | | a:4611 | 2.05 | 1.0 | 24.8 | 2.64 [G:OP1]a:2057 | 15.56 | 3 | A | OP1 | a:2581 | 2.13 | 1.0 | 24.1 |  | 13.82 | 4 | G | OP1 | a:2580 | 2.14 | 1.0 | 23.4 |  | 16.75 | 5 | Water 3 | | a:4408 | 2.21 | 1.0 | 21.0 | 2.62 [A:OP2]a:2056 // 2.73 [C:O3']a:2579 | 16.12 | 6 | Water 4 | | a:7727 | 2.23 | 1.0 | 23.9 |  | 14.24 | 7 | C | O3' | a:2579 | 3.57 | 1.0 | 25.9 |  | 16.50 | 8 | Water 5 | | a:6176 | 3.91 | 1.0 | 25.2 | 2.75 [A:OP2]a:2056 | 14.03 | 9 | A | OP2 | a:2581 | 4.00 | 1.0 | 24.9 |  | 13.56 | 10 | **K** | K | a:3123 | 4.14 | 1.0 | 30.0 |  | 31.82 | 11 | G | OP2 | a:2057 | 4.15 | 1.0 | 24.9 |  | 15.70 | 12 | G | OP1 | a:2057 | 4.17 | 1.0 | 20.6 |  | 15.05 | 13 | G | O3' | a:2580 | 4.24 | 1.0 | 25.1 |  | 12.94 | 14 | A | OP2 | a:2056 | 4.31 | 1.0 | 24.2 |  | 16.22 | 15 | Water 6 | | a:8466 | 4.32 | 1.0 | 33.6 | 2.93 [G:OP2]a:2580 | 5.46 | 16 | Water 7 | | a:10270 | 4.33 | 1.0 | 47.5 | 2.84 [K:K]a:3123 | 4.97 | 17 | G | O5' | a:2580 | 4.38 | 1.0 | 25.0 |  | 14.76 | 18 | G | OP2 | a:2580 | 4.46 | 1.0 | 26.2 |  | 14.54 | 19 | A | O5' | a:2581 | 4.54 | 1.0 | 22.3 |  | 16.87 | 20 | Water 8 | | a:6008 | 4.77 | 1.0 | 33.0 | 2.78 [K:K]a:3123 | 12.79 |

  
**1^st^ shell coordination angles and distances (*trans* in "blue"; *cys* in "green"; deviations in "orange"):**   
• 1-2(W/W): 94º; 1-3(W/OP1): *80º*; 1-4(W/OP1): 172º; 1-5(W/W): 88º; 1-6(W/W): 91º; 2-3(W/OP1): 172º; 2-4(W/OP1): 93º; 2-5(W/W): 93º; 2-6(W/W): 92º; 3-4(OP1/OP1): 93º; 3-5(OP1/W): 92º; 3-6(OP1/W): *82º*; 4-5(OP1/W): 89º; 4-6(OP1/W): 92º; 5-6(W/W): 174º;   
• 1-2(W/W): 3.0Å; 1-3(W/OP1): 2.7Å; 1-4(W/OP1): 4.2Å; 1-5(W/W): 2.9Å; 1-6(W/W): 3.0Å; 2-3(W/OP1): 4.2Å; 2-4(W/OP1): 3.0Å; 2-5(W/W): 3.1Å; 2-6(W/W): 3.1Å; 3-4(OP1/OP1): 3.1Å; 3-5(OP1/W): 3.1Å; 3-6(OP1/W): 2.9Å; 4-5(OP1/W): 3.0Å; 4-6(OP1/W): 3.1Å; 5-6(W/W): *4.4Å*;   
  
**2.13 ± 0.07: the average coodination distance of the 1^st^ six ligands is Mg^2+^ compatible**  
**8: Number of potential Water contacts**   
**4 : Number of second shell contacts involving water or ions**   
**50%: Percentage of satisfied second shell Water contacts**   
• Contact(s): /O_w_/O_w_/O_w_/O_w_/OP1/OP1/  
• Contact code: *cis-*2O_ph_4O_w_  
Ooctahedral coordination (4.4º)   
Go to ToC

### MG:a:3195

|  | Res. type | Atom | Residue | Distance | Occupancy | B-factor | Closest atom(s) to a 1^st^ shell water | EM\_dens. |
| --- | --- | --- | --- | --- | --- | --- | --- | --- |
| **Ion** | | | |  | 1.0 | 47.5 |  | 8.46 |
 1 | Water 1 | | a:7793 | 1.94 | 1.0 | 40.1 | 3.00 [A:N1]a:1286 | 5.48 | 2 | A | OP1 | a:1330 | 2.01 | 1.0 | 31.7 |  | 8.48 | 3 | Water 2 | | a:6767 | 2.06 | 1.0 | 48.1 |  | 3.54 | 4 | Water 3 | | a:8893 | 2.44 | 1.0 | 44.3 |  | 4.14 | 5 | Water 4 | | a:7550 | 4.07 | 1.0 | 38.3 |  | 6.95 | 6 | A | OP2 | a:1330 | 4.13 | 1.0 | 36.2 |  | 6.27 | 7 | A | O3' | a:1329 | 4.17 | 1.0 | 30.6 |  | 9.79 | 8 | G | O6 | a:489 | 4.22 | 1.0 | 40.9 |  | 5.17 | 9 | A | O5' | a:1330 | 4.44 | 1.0 | 32.3 |  | 8.40 | 10 | A | OP2 | a:1323 | 4.47 | 1.0 | 40.4 |  | 5.33 | 11 | Water 5 | | a:7704 | 4.50 | 1.0 | 41.3 | 2.94 [A:O3']a:1329 | 5.17 | 12 | Water 6 | | a:7263 | 4.54 | 1.0 | 37.6 | 2.97 [A:OP2]a:1330 | 6.20 | 13 | A | N1 | a:1286 | 4.61 | 1.0 | 32.5 |  | 7.53 | 14 | Water 7 | | a:10013 | 4.71 | 1.0 | 38.2 |  | 5.35 |

  
**1^st^ shell coordination angles and distances (*trans* in "blue"; *cys* in "green"; deviations in "orange"):**   
• 1-2(W/OP1): 93º; 1-3(W/W): 176º; 2-3(OP1/W): 86º;   
• 1-2(W/OP1): 2.9Å; 1-3(W/W): 4.0Å; 2-3(OP1/W): 2.8Å;   
  
**2.77 ± 0.95: the average coodination distance of the 1^st^ six ligands is K^+^ compatible**  
**8: Number of potential Water contacts**   
**1 : Number of second shell contacts involving water or ions**   
**12%: Percentage of satisfied second shell Water contacts**   
• Contact(s): /O_w_/O_w_/OP1/  
• Contact code: O_ph_2O_w_  
• Contact(s) in the exclusion zone: /O_w_/  
→ Missing octahedral coordination for MG ←  
Go to ToC

### MG:a:3196

|  | Res. type | Atom | Residue | Distance | Occupancy | B-factor | Closest atom(s) to a 1^st^ shell water | EM\_dens. |
| --- | --- | --- | --- | --- | --- | --- | --- | --- |
| **Ion** | | | |  | 1.0 | 45.3 |  | 7.06 |
 1 | G | N7 | a:630 | 2.15 | 1.0 | 29.4 |  | 10.55 | 2 | Water 1 | | a:4229 | 2.86 | 1.0 | 42.6 | 2.60 [G:OP2]a:630 | 5.05 | 3 | Water 2 | | a:5340 | 2.86 | 1.0 | 36.2 | 2.70 [G:N7]a:631 | 5.87 | 4 | G | O6 | a:630 | 3.38 | 1.0 | 27.4 |  | 10.22 | 5 | GLU | OE1 | k:76 | 3.74 | 1.0 | 42.9 |  | 5.62 | 6 | A | O2' | a:629 | 4.23 | 1.0 | 38.0 |  | 7.42 | 7 | G | OP2 | a:630 | 4.24 | 1.0 | 35.7 |  | 7.45 | 8 | Water 3 | | a:4099 | 4.57 | 1.0 | 38.0 |  | 4.49 | 9 | G | N7 | a:631 | 4.64 | 1.0 | 27.1 |  | 12.01 | 10 | A | O3' | a:629 | 4.79 | 1.0 | 41.8 |  | 7.97 | 11 | G | O6 | a:631 | 4.79 | 1.0 | 36.0 |  | 9.67 |

  
• • **3.20 ± 0.67: the average coodination distance of the 1^st^ six ligands is K^+^ compatible**  
**4: Number of potential Water contacts**   
**2 : Number of second shell contacts involving water or ions**   
**50%: Percentage of satisfied second shell Water contacts**   
• Contact(s): /N7/  
• Contact code: N_b_  
• Contact(s) in the exclusion zone: /O6/O_w_/O_w_/  
→ Missing octahedral coordination for MG ←  
Go to ToC

### MG:a:3197

|  | Res. type | Atom | Residue | Distance | Occupancy | B-factor | Closest atom(s) to a 1^st^ shell water | EM\_dens. |
| --- | --- | --- | --- | --- | --- | --- | --- | --- |
| **Ion** | | | |  | 1.0 | 47.2 |  | 11.53 |
 1 | Water 1 | | a:4106 | 1.95 | 1.0 | 27.0 | 2.59 [A:OP1]a:672 | 12.06 | 2 | Water 2 | | a:7809 | 2.04 | 1.0 | 36.0 | 2.86 [A:OP2]a:806 | 8.31 | 3 | Water 3 | | a:10145 | 2.07 | 1.0 | 35.9 |  | 7.60 | 4 | Water 4 | | a:9958 | 2.11 | 1.0 | 40.2 |  | 6.86 | 5 | Water 5 | | k:270 | 2.23 | 1.0 | 40.1 |  | 9.58 | 6 | Water 6 | | a:5828 | 3.51 | 1.0 | 40.6 | 2.73 [A:OP2]a:806 | 6.84 | 7 | Water 7 | | a:6756 | 3.62 | 1.0 | 41.3 |  | 4.24 | 8 | Water 8 | | a:5777 | 3.85 | 1.0 | 43.4 | 2.73 [A:OP2]a:806 | 6.46 | 9 | Water 9 | | a:9483 | 3.93 | 1.0 | 33.8 |  | 9.53 | 10 | Water 10 | | a:4039 | 3.95 | 1.0 | 26.9 |  | 11.75 | 11 | A | OP1 | a:672 | 3.99 | 1.0 | 36.7 |  | 11.54 | 12 | Water 11 | | a:6104 | 4.04 | 1.0 | 26.5 |  | 11.99 | 13 | A | OP2 | a:806 | 4.11 | 1.0 | 31.4 |  | 11.58 | 14 | Water 12 | | a:5276 | 4.12 | 1.0 | 29.7 |  | 9.21 | 15 | Water 13 | | a:3940 | 4.13 | 1.0 | 25.1 |  | 15.31 | 16 | Water 14 | | a:4279 | 4.18 | 1.0 | 32.2 |  | 11.72 | 17 | Water 15 | | a:9894 | 4.34 | 1.0 | 40.4 |  | 6.29 | 18 | SER | O | k:42 | 4.46 | 1.0 | 26.8 |  | 16.68 | 19 | SER | OG | k:42 | 4.61 | 1.0 | 31.9 |  | 10.00 |

  
**1^st^ shell coordination angles and distances (*trans* in "blue"; *cys* in "green"; deviations in "orange"):**   
• 1-2(W/W): *83º*; 1-3(W/W): 90º; 1-4(W/W): 92º; 1-5(W/W): *83º*; 2-3(W/W): 173º; 2-4(W/W): *96º*; 2-5(W/W): *101º*; 3-4(W/W): *83º*; 3-5(W/W): *79º*; 4-5(W/W): *161º*;   
• 1-2(W/W): 2.7Å; 1-3(W/W): 2.8Å; 1-4(W/W): 2.9Å; 1-5(W/W): 2.8Å; 2-3(W/W): 4.1Å; 2-4(W/W): 3.1Å; 2-5(W/W): *3.3Å*; 3-4(W/W): 2.8Å; 3-5(W/W): 2.7Å; 4-5(W/W): *4.3Å*;   
  
**2.32 ± 0.54: the average coodination distance of the 1^st^ six ligands is Na^+^** compatible  
**12: Number of potential Water contacts**   
**3 : Number of second shell contacts involving water or ions**   
**25%: Percentage of satisfied second shell Water contacts**   
• Contact(s): /O_w_/O_w_/O_w_/O_w_/O_w_/  
• Contact code: 5O_w_  
→ Missing octahedral coordination for MG ←  
→ Slightly distorded coord. octahedron (7.7º) ←  
Go to ToC

### MG:a:3198

|  | Res. type | Atom | Residue | Distance | Occupancy | B-factor | Closest atom(s) to a 1^st^ shell water | EM\_dens. |
| --- | --- | --- | --- | --- | --- | --- | --- | --- |
| **Ion** | | | |  | 1.0 | 40.9 |  | 14.13 |
 1 | Water 1 | | a:4558 | 2.08 | 1.0 | 26.8 | 2.64 [A:OP1]a:831 // 2.92 [G:O4']a:833 | 12.84 | 2 | Water 2 | | a:7807 | 2.21 | 1.0 | 26.7 |  | 13.01 | 3 | Water 3 | | a:9681 | 2.21 | 1.0 | 35.9 |  | 6.78 | 4 | Water 4 | | a:3531 | 2.25 | 1.0 | 37.6 | 2.46 [U:OP2]a:2072 | 6.94 | 5 | Water 5 | | a:9921 | 2.27 | 1.0 | 36.4 |  | 9.71 | 6 | G | O2' | a:832 | 2.28 | 1.0 | 34.0 |  | 11.30 | 7 | A | OP1 | a:831 | 4.16 | 1.0 | 26.9 |  | 12.41 | 8 | Water 6 | | a:3541 | 4.19 | 1.0 | 36.3 |  | 8.29 | 9 | G | O4' | a:833 | 4.23 | 1.0 | 25.6 |  | 17.22 | 10 | Water 7 | | a:8616 | 4.34 | 1.0 | 42.5 |  | 6.12 | 11 | G | O2' | a:807 | 4.37 | 1.0 | 28.7 |  | 13.68 | 12 | G | O3' | a:832 | 4.40 | 1.0 | 27.6 |  | 14.24 | 13 | U | OP2 | a:2072 | 4.48 | 1.0 | 34.6 |  | 6.29 | 14 | Water 8 | | a:9794 | 4.52 | 1.0 | 38.1 |  | 5.83 | 15 | Water 9 | | a:6007 | 4.53 | 1.0 | 35.5 |  | 8.92 |

  
**1^st^ shell coordination angles and distances (*trans* in "blue"; *cys* in "green"; deviations in "orange"):**   
• 1-2(W/W): 90º; 1-3(W/W): 173º; 1-4(W/W): *79º*; 1-5(W/W): 90º; 1-6(W/O2'): 94º; 2-3(W/W): 86º; 2-4(W/W): 90º; 2-5(W/W): 175º; 2-6(W/O2'): *85º*; 3-4(W/W): *96º*; 3-5(W/W): 94º; 3-6(W/O2'): 91º; 4-5(W/W): *84º*; 4-6(W/O2'): 171º; 5-6(W/O2'): *100º*;   
• 1-2(W/W): 3.0Å; 1-3(W/W): *4.3Å*; 1-4(W/W): 2.7Å; 1-5(W/W): 3.1Å; 1-6(W/O2'): 3.2Å; 2-3(W/W): 3.0Å; 2-4(W/W): 3.2Å; 2-5(W/W): *4.5Å*; 2-6(W/O2'): 3.0Å; 3-4(W/W): *3.3Å*; 3-5(W/W): *3.3Å*; 3-6(W/O2'): *3.2Å*; 4-5(W/W): 3.0Å; 4-6(W/O2'): *4.5Å*; 5-6(W/O2'): *3.5Å*;   
  
**2.22 ± 0.07: the average coodination distance of the 1^st^ six ligands is Mg^2+^ compatible**  
**10: Number of potential Water contacts**   
**3 : Number of second shell contacts involving water or ions**   
**30%: Percentage of satisfied second shell Water contacts**   
• Contact(s): /O2'/O_w_/O_w_/O_w_/O_w_/O_w_/  
• Contact code: O_r_5O_w_  
Ooctahedral coordination (5.0º)   
Go to ToC

### MG:a:3199

|  | Res. type | Atom | Residue | Distance | Occupancy | B-factor | Closest atom(s) to a 1^st^ shell water | EM\_dens. |
| --- | --- | --- | --- | --- | --- | --- | --- | --- |
| **Ion** | | | |  | 1.0 | 28.6 |  | 23.09 |
 1 | Water 1 | | a:6499 | 1.94 | 1.0 | 26.2 | 2.77 [C:OP2]a:818 | 13.15 | 2 | Water 2 | | a:9078 | 2.07 | 1.0 | 33.4 |  | 10.62 | 3 | Water 3 | | a:9088 | 2.08 | 1.0 | 24.2 |  | 13.93 | 4 | Water 4 | | a:8095 | 2.11 | 1.0 | 21.2 | 2.89 [G:O6]a:820 | 15.96 | 5 | G | OP2 | a:1189 | 2.14 | 1.0 | 22.2 |  | 13.79 | 6 | Water 5 | | a:4242 | 2.20 | 1.0 | 26.9 | 2.60 [C:OP2]a:819 | 13.49 | 7 | G | OP1 | a:1189 | 3.83 | 1.0 | 25.0 |  | 10.22 | 8 | C | OP2 | a:819 | 3.96 | 1.0 | 30.6 |  | 11.38 | 9 | C | OP2 | a:818 | 4.01 | 1.0 | 26.3 |  | 11.42 | 10 | Water 6 | | a:7747 | 4.26 | 1.0 | 29.9 | 2.86 [C:OP2]a:818 | 10.47 | 11 | Water 7 | | a:9522 | 4.29 | 1.0 | 32.8 |  | 9.21 | 12 | G | O5' | a:1189 | 4.40 | 1.0 | 33.0 |  | 8.42 | 13 | Water 8 | | a:8863 | 4.41 | 1.0 | 38.7 | 2.97 [G:O2']a:1187 | 7.15 | 14 | G | O6 | a:820 | 4.47 | 1.0 | 21.6 |  | 19.69 | 15 | G | O3' | a:1188 | 4.48 | 1.0 | 26.2 |  | 11.82 | 16 | G | O2' | a:1187 | 4.51 | 1.0 | 31.4 |  | 9.19 | 17 | G | N7 | a:1189 | 4.60 | 1.0 | 14.1 |  | 18.87 |

  
**1^st^ shell coordination angles and distances (*trans* in "blue"; *cys* in "green"; deviations in "orange"):**   
• 1-2(W/W): 87º; 1-3(W/W): 91º; 1-4(W/W): *169º*; 1-5(W/OP2): *95º*; 1-6(W/W): *85º*; 2-3(W/W): 177º; 2-4(W/W): *101º*; 2-5(W/OP2): 91º; 2-6(W/W): 94º; 3-4(W/W): *80º*; 3-5(W/OP2): 91º; 3-6(W/W): *84º*; 4-5(W/OP2): 93º; 4-6(W/W): 87º; 5-6(OP2/W): 175º;   
• 1-2(W/W): 2.8Å; 1-3(W/W): 2.9Å; 1-4(W/W): 4.0Å; 1-5(W/OP2): 3.0Å; 1-6(W/W): 2.8Å; 2-3(W/W): 4.1Å; 2-4(W/W): *3.2Å*; 2-5(W/OP2): 3.0Å; 2-6(W/W): 3.1Å; 3-4(W/W): 2.7Å; 3-5(W/OP2): 3.0Å; 3-6(W/W): 2.9Å; 4-5(W/OP2): 3.1Å; 4-6(W/W): 3.0Å; 5-6(OP2/W): *4.3Å*;   
  
**2.09 ± 0.08: the average coodination distance of the 1^st^ six ligands is Mg^2+^ compatible**  
**10: Number of potential Water contacts**   
**3 : Number of second shell contacts involving water or ions**   
**30%: Percentage of satisfied second shell Water contacts**   
• Contact(s): /O_w_/O_w_/O_w_/O_w_/O_w_/OP2/  
• Contact code: O_ph_5O_w_  
Ooctahedral coordination (4.9º)   
Go to ToC

### MG:a:3200

|  | Res. type | Atom | Residue | Distance | Occupancy | B-factor | Closest atom(s) to a 1^st^ shell water | EM\_dens. |
| --- | --- | --- | --- | --- | --- | --- | --- | --- |
| **Ion** | | | |  | 1.0 | 33.6 |  | 13.89 |
 1 | Water 1 | | a:5494 | 1.83 | 1.0 | 35.1 |  | 5.35 | 2 | Water 2 | | a:9037 | 1.90 | 1.0 | 34.7 |  | 6.52 | 3 | Water 3 | | a:4585 | 1.97 | 1.0 | 32.5 |  | 6.29 | 4 | Water 4 | | a:6450 | 1.99 | 1.0 | 31.8 | 2.77 [G:O6]a:978 | 8.88 | 5 | Water 5 | | a:8849 | 2.01 | 1.0 | 29.4 |  | 11.91 | 6 | A | OP2 | a:977 | 2.06 | 1.0 | 28.3 |  | 9.95 | 7 | A | O5' | a:977 | 3.86 | 1.0 | 32.2 |  | 11.58 | 8 | G | O6 | a:978 | 3.96 | 1.0 | 25.4 |  | 17.64 | 9 | Water 6 | | a:8276 | 3.99 | 1.0 | 35.5 |  | 8.29 | 10 | G | O2' | a:976 | 4.08 | 1.0 | 29.2 |  | 13.57 | 11 | Water 7 | | a:5508 | 4.12 | 1.0 | 33.0 |  | 8.42 | 12 | G | N7 | a:978 | 4.13 | 1.0 | 21.6 |  | 15.61 | 13 | Water 8 | | a:8262 | 4.15 | 1.0 | 35.7 | 2.91 [A:OP1]a:977 | 7.51 | 14 | Water 9 | | a:9671 | 4.17 | 1.0 | 37.6 |  | 4.68 | 15 | Water 10 | | a:3603 | 4.22 | 1.0 | 37.8 | 2.49 [A:OP1]a:986 | 6.92 | 16 | G | O3' | a:976 | 4.32 | 1.0 | 28.7 |  | 13.00 | 17 | A | OP1 | a:977 | 4.47 | 1.0 | 33.9 |  | 10.37 | 18 | A | OP1 | a:986 | 4.62 | 1.0 | 24.6 |  | 12.74 |

  
**1^st^ shell coordination angles and distances (*trans* in "blue"; *cys* in "green"; deviations in "orange"):**   
• 1-2(W/W): 172º; 1-3(W/W): 94º; 1-4(W/W): 87º; 1-5(W/W): 85º; 1-6(W/OP2): 88º; 2-3(W/W): 92º; 2-4(W/W): 87º; 2-5(W/W): 89º; 2-6(W/OP2): *98º*; 3-4(W/W): 94º; 3-5(W/W): 176º; 3-6(W/OP2): *82º*; 4-5(W/W): 90º; 4-6(W/OP2): 173º; 5-6(W/OP2): 94º;   
• 1-2(W/W): *3.7Å*; 1-3(W/W): 2.8Å; 1-4(W/W): 2.6Å; 1-5(W/W): 2.6Å; 1-6(W/OP2): 2.7Å; 2-3(W/W): 2.8Å; 2-4(W/W): 2.7Å; 2-5(W/W): 2.7Å; 2-6(W/OP2): 3.0Å; 3-4(W/W): 2.9Å; 3-5(W/W): 4.0Å; 3-6(W/OP2): 2.6Å; 4-5(W/W): 2.8Å; 4-6(W/OP2): 4.0Å; 5-6(W/OP2): 3.0Å;   
  
**1.96 ± 0.07: the average coodination distance of the 1^st^ six ligands is Mg^2+^ compatible**  
**10: Number of potential Water contacts**   
**1 : Number of second shell contacts involving water or ions**   
**10%: Percentage of satisfied second shell Water contacts**   
• Contact(s): /O_w_/O_w_/O_w_/O_w_/OP2/  
• Contact code: O_ph_4O_w_  
• Contact(s) in the exclusion zone: /O_w_/  
→ Missing octahedral coordination for MG ←  
Ooctahedral coordination (4.2º)   
Go to ToC

### MG:a:3201

|  | Res. type | Atom | Residue | Distance | Occupancy | B-factor | Closest atom(s) to a 1^st^ shell water | EM\_dens. |
| --- | --- | --- | --- | --- | --- | --- | --- | --- |
| **Ion** | | | |  | 1.0 | 24.5 |  | 25.16 |
 1 | Water 1 | | a:7342 | 2.02 | 1.0 | 25.1 |  | 14.84 | 2 | C | OP1 | a:2076 | 2.04 | 1.0 | 23.0 |  | 14.03 | 3 | Water 2 | | a:6125 | 2.09 | 1.0 | 24.3 | 2.75 [A:OP2]a:795 | 16.15 | 4 | C | OP1 | a:680 | 2.15 | 1.0 | 25.6 |  | 14.20 | 5 | G | O6 | a:778 | 2.16 | 1.0 | 26.6 |  | 14.70 | 6 | Water 3 | | a:7981 | 2.24 | 1.0 | 27.4 | 2.96 [C:OP2]a:680 | 11.45 | 7 | A | O3' | a:2075 | 3.47 | 1.0 | 27.4 |  | 13.12 | 8 | C | OP2 | a:680 | 3.97 | 1.0 | 24.4 |  | 14.97 | 9 | C | OP2 | a:2076 | 4.26 | 1.0 | 27.1 |  | 13.00 | 10 | C | O5' | a:680 | 4.27 | 1.0 | 23.5 |  | 16.77 | 11 | Water 4 | | a:3855 | 4.29 | 1.0 | 39.8 | 2.55 [C:OP2]a:681 | 5.32 | 12 | Water 5 | | a:9695 | 4.31 | 1.0 | 38.9 |  | 6.69 | 13 | A | O2' | a:795 | 4.31 | 1.0 | 28.1 |  | 14.77 | 14 | C | O5' | a:2076 | 4.32 | 1.0 | 26.5 |  | 14.20 | 15 | Water 6 | | a:7932 | 4.48 | 1.0 | 34.5 | 2.88 [C:OP2]a:2076 | 8.29 | 16 | A | OP2 | a:795 | 4.54 | 1.0 | 30.5 |  | 12.83 | 17 | A | O3' | a:679 | 4.56 | 1.0 | 24.6 |  | 13.65 | 18 | C | OP2 | a:681 | 4.71 | 1.0 | 34.0 |  | 10.63 | 19 | Water 7 | | a:7427 | 4.76 | 1.0 | 42.3 |  | 5.65 |

  
**1^st^ shell coordination angles and distances (*trans* in "blue"; *cys* in "green"; deviations in "orange"):**   
• 1-2(W/OP1): 88º; 1-3(W/W): 173º; 1-4(W/OP1): 92º; 1-5(W/O6): 89º; 1-6(W/W): 91º; 2-3(OP1/W): *98º*; 2-4(OP1/OP1): *102º*; 2-5(OP1/O6): 93º; 2-6(OP1/W): 176º; 3-4(W/OP1): 89º; 3-5(W/O6): 88º; 3-6(W/W): *83º*; 4-5(OP1/O6): *165º*; 4-6(OP1/W): *82º*; 5-6(O6/W): *83º*;   
• 1-2(W/OP1): 2.8Å; 1-3(W/W): 4.1Å; 1-4(W/OP1): 3.0Å; 1-5(W/O6): 2.9Å; 1-6(W/W): 3.0Å; 2-3(OP1/W): 3.1Å; 2-4(OP1/OP1): *3.2Å*; 2-5(OP1/O6): 3.1Å; 2-6(OP1/W): *4.3Å*; 3-4(W/OP1): 3.0Å; 3-5(W/O6): 3.0Å; 3-6(W/W): 2.9Å; 4-5(OP1/O6): *4.3Å*; 4-6(OP1/W): 2.9Å; 5-6(O6/W): 2.9Å;   
  
**2.12 ± 0.07: the average coodination distance of the 1^st^ six ligands is Mg^2+^ compatible**  
**6: Number of potential Water contacts**   
**2 : Number of second shell contacts involving water or ions**   
**33%: Percentage of satisfied second shell Water contacts**   
• Contact(s): /O6/O_w_/O_w_/O_w_/OP1/OP1/  
• Contact code: *cis-*2O_ph_O_b_3O_w_  
→ Slightly distorded coord. octahedron (5.2º) ←  
Go to ToC

### MG:a:3202

|  | Res. type | Atom | Residue | Distance | Occupancy | B-factor | Closest atom(s) to a 1^st^ shell water | EM\_dens. |
| --- | --- | --- | --- | --- | --- | --- | --- | --- |
| **Ion** | | | |  | 1.0 | 29.6 |  | 19.97 |
 1 | U | OP1 | a:2078 | 1.88 | 1.0 | 24.7 |  | 14.53 | 2 | Water 1 | | a:6309 | 1.99 | 1.0 | 31.8 | 2.76 [U:OP2]a:2244 | 8.92 | 3 | Water 2 | | a:4974 | 2.04 | 1.0 | 31.7 | 2.67 [U:OP2]a:2079 | 10.86 | 4 | Water 3 | | a:3616 | 2.26 | 1.0 | 30.9 | 2.49 [G:OP2]a:2243 | 13.17 | 5 | Water 4 | | c:436 | 2.30 | 1.0 | 25.8 | 2.64 [PRO:O]c:227 | 13.60 | 6 | U | OP2 | a:2078 | 3.79 | 1.0 | 22.6 |  | 15.81 | 7 | PRO | O | c:227 | 3.83 | 1.0 | 27.3 |  | 15.79 | 8 | Water 5 | | a:7126 | 3.96 | 1.0 | 35.7 | 2.81 [U:OP2]a:2078 | 5.99 | 9 | G | OP2 | a:2243 | 3.96 | 1.0 | 32.3 |  | 9.86 | 10 | U | O5' | a:2078 | 4.06 | 1.0 | 23.2 |  | 16.61 | 11 | U | OP2 | a:2244 | 4.18 | 1.0 | 37.8 |  | 11.16 | 12 | Water 6 | | a:9830 | 4.25 | 1.0 | 36.5 |  | 6.16 | 13 | C | O3' | a:2077 | 4.28 | 1.0 | 24.5 |  | 15.91 | 14 | U | OP2 | a:2079 | 4.33 | 1.0 | 25.4 |  | 14.76 | 15 | G | O2' | a:2242 | 4.43 | 1.0 | 28.7 |  | 13.47 | 16 | Water 7 | | a:4846 | 4.57 | 1.0 | 36.9 |  | 8.15 | 17 | Water 8 | | c:552 | 4.61 | 1.0 | 34.9 |  | 7.85 | 18 | Water 9 | | a:4797 | 4.61 | 1.0 | 44.3 |  | 6.97 | 19 | G | O3' | a:2242 | 4.66 | 1.0 | 33.5 |  | 11.07 | 20 | Water 10 | | a:4553 | 4.77 | 1.0 | 25.4 | 2.64 [U:OP2]a:2078 // 2.79 [G:O2']a:2242 | 14.06 |

  
**1^st^ shell coordination angles and distances (*trans* in "blue"; *cys* in "green"; deviations in "orange"):**   
• 1-2(OP1/W): *100º*; 1-3(OP1/W): *100º*; 1-4(OP1/W): 173º; 2-3(W/W): 87º; 2-4(W/W): *85º*; 3-4(W/W): *85º*;   
• 1-2(OP1/W): 3.0Å; 1-3(OP1/W): 3.0Å; 1-4(OP1/W): 4.1Å; 2-3(W/W): 2.8Å; 2-4(W/W): 2.9Å; 3-4(W/W): 2.9Å;   
  
**2.38 ± 0.65: the average coodination distance of the 1^st^ six ligands is Na^+^** compatible  
**8: Number of potential Water contacts**   
**4 : Number of second shell contacts involving water or ions**   
**50%: Percentage of satisfied second shell Water contacts**   
• Contact(s): /O_w_/O_w_/O_w_/  
• Contact code: 3O_w_  
• Contact(s) in the exclusion zone: /O_w_/OP1/  
→ Missing octahedral coordination for MG ←  
→ Slightly distorded coord. octahedron (6.8º) ←  
Go to ToC

### MG:a:3203

|  | Res. type | Atom | Residue | Distance | Occupancy | B-factor | Closest atom(s) to a 1^st^ shell water | EM\_dens. |
| --- | --- | --- | --- | --- | --- | --- | --- | --- |
| **Ion** | | | |  | 1.0 | 29.2 |  | 18.40 |
 1 | U | O4 | a:690 | 2.74 | 1.0 | 27.8 |  | 10.95 | 2 | Water 1 | | a:6066 | 2.79 | 1.0 | 40.0 | 2.75 [C:OP2]a:689 | 6.82 | 3 | A | O3' | a:687 | 2.82 | 1.0 | 28.6 |  | 15.12 | 4 | U | OP2 | a:688 | 2.92 | 1.0 | 29.3 |  | 12.17 | 5 | Water 2 | | a:5946 | 3.06 | 1.0 | 31.2 |  | 10.01 | 6 | A | O2' | a:687 | 3.06 | 1.0 | 27.8 |  | 13.88 | 7 | Water 3 | | a:9133 | 3.37 | 1.0 | 34.0 |  | 9.81 | 8 | Water 4 | | a:6541 | 4.38 | 1.0 | 40.8 |  | 5.16 | 9 | C | OP2 | a:689 | 4.39 | 1.0 | 29.4 |  | 10.02 | 10 | U | OP1 | a:688 | 4.42 | 1.0 | 27.8 |  | 13.29 | 11 | U | O5' | a:688 | 4.65 | 1.0 | 29.0 |  | 13.44 | 12 | Water 5 | | a:8980 | 4.73 | 1.0 | 45.9 |  | 5.50 |

  
• • **2.90 ± 0.13: the average coodination distance of the 1^st^ six ligands is K^+^ compatible**  
**4: Number of potential Water contacts**   
**1 : Number of second shell contacts involving water or ions**   
**25%: Percentage of satisfied second shell Water contacts**   
• Contact code: → No contacts ←   
• Contact(s) in the exclusion zone: /O2'/O3'/O4/O_w_/O_w_/O_w_/OP2/  
→ Consider K in place of MG (4 valid contact(s) for K vs. 0 for MG ←)  
Go to ToC

### MG:a:3204

|  | Res. type | Atom | Residue | Distance | Occupancy | B-factor | Closest atom(s) to a 1^st^ shell water | EM\_dens. |
| --- | --- | --- | --- | --- | --- | --- | --- | --- |
| **Ion** | | | |  | 1.0 | 43.4 |  | 17.39 |
 1 | Water 1 | | a:7885 | 1.94 | 1.0 | 39.2 |  | 6.09 | 2 | A | OP1 | a:741 | 2.09 | 1.0 | 34.2 |  | 9.75 | 3 | Water 2 | | a:7842 | 2.12 | 1.0 | 35.0 | 2.87 [U:OP2]a:1780 | 5.45 | 4 | Water 3 | | a:5611 | 2.15 | 1.0 | 30.0 | 2.72 [U:OP1]a:1780 | 10.50 | 5 | G | O3' | a:740 | 3.64 | 1.0 | 32.2 |  | 9.67 | 6 | U | OP2 | a:1781 | 3.87 | 1.0 | 27.0 |  | 14.42 | 7 | A | OP2 | a:741 | 4.03 | 1.0 | 31.6 |  | 12.02 | 8 | U | OP2 | a:1780 | 4.10 | 1.0 | 30.6 |  | 12.07 | 9 | Water 4 | | a:5287 | 4.12 | 1.0 | 30.1 |  | 13.66 | 10 | Water 5 | | a:5509 | 4.26 | 1.0 | 42.1 |  | 5.24 | 11 | U | OP1 | a:1780 | 4.37 | 1.0 | 31.6 |  | 11.10 | 12 | Water 6 | | a:5394 | 4.39 | 1.0 | 44.4 |  | 5.13 | 13 | A | N1 | a:1786 | 4.43 | 1.0 | 20.7 |  | 16.34 | 14 | Water 7 | | a:10100 | 4.45 | 1.0 | 50.1 |  | 3.68 | 15 | A | O5' | a:741 | 4.53 | 1.0 | 29.3 |  | 9.06 |

  
**1^st^ shell coordination angles and distances (*trans* in "blue"; *cys* in "green"; deviations in "orange"):**   
• 1-2(W/OP1): 91º; 1-3(W/W): 94º; 1-4(W/W): 91º; 2-3(OP1/W): *95º*; 2-4(OP1/W): 174º; 3-4(W/W): 90º;   
• 1-2(W/OP1): 2.9Å; 1-3(W/W): 3.0Å; 1-4(W/W): 2.9Å; 2-3(OP1/W): 3.1Å; 2-4(OP1/W): *4.2Å*; 3-4(W/W): 3.0Å;   
  
**2.64 ± 0.80: the average coodination distance of the 1^st^ six ligands is K^+^ compatible**  
**6: Number of potential Water contacts**   
**2 : Number of second shell contacts involving water or ions**   
**33%: Percentage of satisfied second shell Water contacts**   
• Contact(s): /O_w_/O_w_/O_w_/OP1/  
• Contact code: O_ph_3O_w_  
→ Missing octahedral coordination for MG ←  
Ooctahedral coordination (2.8º)   
Go to ToC

### MG:a:3205

|  | Res. type | Atom | Residue | Distance | Occupancy | B-factor | Closest atom(s) to a 1^st^ shell water | EM\_dens. |
| --- | --- | --- | --- | --- | --- | --- | --- | --- |
| **Ion** | | | |  | 1.0 | 33.1 |  | 19.97 |
 1 | Water 1 | | a:9743 | 2.04 | 1.0 | 29.0 |  | 12.10 | 2 | Water 2 | | a:5414 | 2.12 | 1.0 | 23.1 | 2.70 [G:O6]a:1794 // 2.99 [U:O4]a:1829 | 14.40 | 3 | Water 3 | | a:9209 | 2.12 | 1.0 | 28.7 |  | 11.96 | 4 | Water 4 | | c:581 | 2.15 | 1.0 | 24.8 |  | 13.08 | 5 | Water 5 | | a:4582 | 2.20 | 1.0 | 22.8 | 2.64 [G:O6]a:1828 | 16.34 | 6 | Water 6 | | a:5745 | 2.24 | 1.0 | 28.6 | 2.73 [A:OP2]a:1793 | 9.50 | 7 | Water 7 | | a:4913 | 3.74 | 1.0 | 28.1 | 2.85 [A:OP2]a:1793 | 12.73 | 8 | Water 8 | | a:10169 | 3.92 | 1.0 | 43.1 |  | 6.87 | 9 | Water 9 | | c:595 | 4.10 | 1.0 | 40.8 |  | 4.60 | 10 | Water 10 | | a:4366 | 4.10 | 1.0 | 24.0 | 2.62 [U:O4]a:1829 | 16.86 | 11 | Water 11 | | a:6325 | 4.12 | 1.0 | 37.0 |  | 9.51 | 12 | U | O4 | a:1829 | 4.26 | 1.0 | 24.0 |  | 15.87 | 13 | G | O6 | a:1828 | 4.28 | 1.0 | 22.2 |  | 16.69 | 14 | A | OP2 | a:1793 | 4.41 | 1.0 | 25.5 |  | 13.60 | 15 | C | OP1 | a:1792 | 4.44 | 1.0 | 23.4 |  | 15.66 | 16 | Water 12 | | a:8735 | 4.64 | 1.0 | 22.4 |  | 16.37 | 17 | C | OP2 | a:1792 | 4.72 | 1.0 | 24.4 |  | 18.15 | 18 | G | O6 | a:1794 | 4.72 | 1.0 | 25.2 |  | 15.57 |

  
**1^st^ shell coordination angles and distances (*trans* in "blue"; *cys* in "green"; deviations in "orange"):**   
• 1-2(W/W): 90º; 1-3(W/W): 177º; 1-4(W/W): 91º; 1-5(W/W): 91º; 1-6(W/W): *99º*; 2-3(W/W): 87º; 2-4(W/W): 174º; 2-5(W/W): 88º; 2-6(W/W): *97º*; 3-4(W/W): 92º; 3-5(W/W): 89º; 3-6(W/W): *81º*; 4-5(W/W): 86º; 4-6(W/W): 89º; 5-6(W/W): *169º*;   
• 1-2(W/W): 2.9Å; 1-3(W/W): 4.2Å; 1-4(W/W): 3.0Å; 1-5(W/W): 3.0Å; 1-6(W/W): *3.3Å*; 2-3(W/W): 2.9Å; 2-4(W/W): *4.3Å*; 2-5(W/W): 3.0Å; 2-6(W/W): *3.3Å*; 3-4(W/W): 3.1Å; 3-5(W/W): 3.0Å; 3-6(W/W): 2.8Å; 4-5(W/W): 3.0Å; 4-6(W/W): 3.1Å; 5-6(W/W): *4.4Å*;   
  
**2.14 ± 0.06: the average coodination distance of the 1^st^ six ligands is Mg^2+^ compatible**  
**12: Number of potential Water contacts**   
**4 : Number of second shell contacts involving water or ions**   
**33%: Percentage of satisfied second shell Water contacts**   
• Contact(s): /O_w_/O_w_/O_w_/O_w_/O_w_/O_w_/  
• Contact code: 6O_w_  
Ooctahedral coordination (4.0º)   
Go to ToC

### MG:a:3206

|  | Res. type | Atom | Residue | Distance | Occupancy | B-factor | Closest atom(s) to a 1^st^ shell water | EM\_dens. |
| --- | --- | --- | --- | --- | --- | --- | --- | --- |
| **Ion** | | | |  | 1.0 | 51.3 |  | 9.73 |
 1 | G | OP1 | a:968 | 2.04 | 1.0 | 31.2 |  | 11.39 | 2 | Water 1 | | a:8042 | 2.06 | 1.0 | 35.9 | 2.88 [U:OP2]a:969 | 6.86 | 3 | Water 2 | | a:9677 | 2.12 | 1.0 | 41.5 |  | 4.88 | 4 | Water 3 | | a:8396 | 2.13 | 1.0 | 40.3 | 2.92 [G:O2']a:824 | 8.21 | 5 | Water 4 | | a:9729 | 2.29 | 1.0 | 36.1 |  | 6.07 | 6 | Water 5 | | a:9889 | 2.43 | 1.0 | 44.0 |  | 5.40 | 7 | G | O5' | a:968 | 4.07 | 1.0 | 27.6 |  | 12.81 | 8 | Water 6 | | a:5748 | 4.17 | 1.0 | 40.8 |  | 6.11 | 9 | G | OP2 | a:968 | 4.22 | 1.0 | 34.4 |  | 9.18 | 10 | G | O2' | a:824 | 4.32 | 1.0 | 25.6 |  | 12.72 | 11 | Water 7 | | a:9115 | 4.38 | 1.0 | 39.7 | 1.92 [MG:MG]a:3356 | 5.63 | 12 | U | OP2 | a:969 | 4.44 | 1.0 | 38.2 |  | 9.50 | 13 | C | O3' | a:967 | 4.47 | 1.0 | 25.3 |  | 13.84 | 14 | Water 8 | | a:8170 | 4.53 | 1.0 | 36.7 | 2.90 [G:OP2]a:968 | 9.26 | 15 | Water 9 | | a:6746 | 4.53 | 1.0 | 37.9 |  | 6.21 | 16 | **MG** | MG | a:3356 | 4.56 | 1.0 | 45.1 |  | 6.77 | 17 | Water 10 | | a:8588 | 4.72 | 1.0 | 42.2 | 2.27 [MG:MG]a:3356 | 6.20 |

  
**1^st^ shell coordination angles and distances (*trans* in "blue"; *cys* in "green"; deviations in "orange"):**   
• 1-2(OP1/W): 92º; 1-3(OP1/W): *101º*; 1-4(OP1/W): 89º; 1-5(OP1/W): *97º*; 2-3(W/W): *164º*; 2-4(W/W): *100º*; 2-5(W/W): 87º; 3-4(W/W): 89º; 3-5(W/W): *83º*; 4-5(W/W): 171º;   
• 1-2(OP1/W): 3.0Å; 1-3(OP1/W): 3.2Å; 1-4(OP1/W): 2.9Å; 1-5(OP1/W): *3.2Å*; 2-3(W/W): 4.1Å; 2-4(W/W): *3.2Å*; 2-5(W/W): 3.0Å; 3-4(W/W): 3.0Å; 3-5(W/W): 2.9Å; 4-5(W/W): *4.4Å*;   
  
**2.18 ± 0.14: the average coodination distance of the 1^st^ six ligands is Mg^2+^ compatible**  
**10: Number of potential Water contacts**   
**2 : Number of second shell contacts involving water or ions**   
**20%: Percentage of satisfied second shell Water contacts**   
• Contact(s): /O_w_/O_w_/O_w_/O_w_/OP1/  
• Contact code: O_ph_4O_w_  
• Contact(s) in the exclusion zone: /O_w_/  
→ Missing octahedral coordination for MG ←  
→ Slightly distorded coord. octahedron (6.6º) ←  
Go to ToC

### MG:a:3207

|  | Res. type | Atom | Residue | Distance | Occupancy | B-factor | Closest atom(s) to a 1^st^ shell water | EM\_dens. |
| --- | --- | --- | --- | --- | --- | --- | --- | --- |
| **Ion** | | | |  | 1.0 | 31.3 |  | 19.18 |
 1 | Water 1 | | a:8640 | 2.00 | 1.0 | 27.7 | 2.95 [C:OP2]a:1000 | 14.06 | 2 | C | OP2 | a:1155 | 2.07 | 1.0 | 29.0 |  | 11.79 | 3 | Water 2 | | a:8722 | 2.09 | 1.0 | 27.7 | 2.96 [C:OP1]a:1000 | 11.29 | 4 | U | OP2 | a:1001 | 2.15 | 1.0 | 32.9 |  | 11.45 | 5 | Water 3 | | a:4134 | 2.24 | 1.0 | 26.6 | 2.59 [G:O6]a:1156 | 12.26 | 6 | Water 4 | | a:7259 | 2.30 | 1.0 | 30.0 |  | 11.49 | 7 | U | OP1 | a:1001 | 4.04 | 1.0 | 32.3 |  | 8.36 | 8 | C | O3' | a:1000 | 4.15 | 1.0 | 27.5 |  | 10.45 | 9 | Water 5 | | a:6079 | 4.23 | 1.0 | 38.4 |  | 9.37 | 10 | C | OP1 | a:1155 | 4.24 | 1.0 | 27.3 |  | 11.99 | 11 | C | O5' | a:1155 | 4.29 | 1.0 | 30.1 |  | 11.60 | 12 | C | OP1 | a:1000 | 4.35 | 1.0 | 34.3 |  | 9.69 | 13 | G | O6 | a:1156 | 4.48 | 1.0 | 22.5 |  | 14.48 | 14 | C | O3' | a:1154 | 4.49 | 1.0 | 28.1 |  | 12.56 | 15 | C | O5' | a:1000 | 4.49 | 1.0 | 35.1 |  | 8.24 | 16 | C | OP2 | a:1000 | 4.52 | 1.0 | 36.0 |  | 9.17 | 17 | Water 6 | | a:6330 | 4.59 | 1.0 | 39.0 | 2.76 [U:OP1]a:1001 | 6.77 | 18 | U | O5' | a:1001 | 4.65 | 1.0 | 30.4 |  | 9.71 |

  
**1^st^ shell coordination angles and distances (*trans* in "blue"; *cys* in "green"; deviations in "orange"):**   
• 1-2(W/OP2): *99º*; 1-3(W/W): *97º*; 1-4(W/OP2): *97º*; 1-5(W/W): 92º; 1-6(W/W): 179º; 2-3(OP2/W): 94º; 2-4(OP2/OP2): *163º*; 2-5(OP2/W): 88º; 2-6(OP2/W): *80º*; 3-4(W/OP2): 90º; 3-5(W/W): *170º*; 3-6(W/W): *82º*; 4-5(OP2/W): 85º; 4-6(OP2/W): *84º*; 5-6(W/W): 89º;   
• 1-2(W/OP2): 3.1Å; 1-3(W/W): 3.1Å; 1-4(W/OP2): 3.1Å; 1-5(W/W): 3.1Å; 1-6(W/W): *4.3Å*; 2-3(OP2/W): 3.0Å; 2-4(OP2/OP2): 4.2Å; 2-5(OP2/W): 3.0Å; 2-6(OP2/W): 2.8Å; 3-4(W/OP2): 3.0Å; 3-5(W/W): *4.3Å*; 3-6(W/W): 2.9Å; 4-5(OP2/W): 3.0Å; 4-6(OP2/W): 3.0Å; 5-6(W/W): 3.2Å;   
  
**2.14 ± 0.10: the average coodination distance of the 1^st^ six ligands is Mg^2+^ compatible**  
**8: Number of potential Water contacts**   
**3 : Number of second shell contacts involving water or ions**   
**38%: Percentage of satisfied second shell Water contacts**   
• Contact(s): /O_w_/O_w_/O_w_/O_w_/OP2/OP2/  
• Contact code: *trans-*2O_ph_4O_w_  
→ Slightly distorded coord. octahedron (5.9º) ←  
Go to ToC

### MG:a:3208

|  | Res. type | Atom | Residue | Distance | Occupancy | B-factor | Closest atom(s) to a 1^st^ shell water | EM\_dens. |
| --- | --- | --- | --- | --- | --- | --- | --- | --- |
| **Ion** | | | |  | 1.0 | 56.1 |  | 3.90 |
 1 | Water 1 | | a:7851 | 1.90 | 1.0 | 35.0 | 2.87 [G:N7]a:1126 | 8.11 | 2 | Water 2 | | a:8138 | 2.21 | 1.0 | 37.6 | 2.89 [G:O6]a:1126 | 4.71 | 3 | G | O6 | a:1127 | 2.27 | 1.0 | 37.1 |  | 8.67 | 4 | Water 3 | | a:8142 | 2.61 | 1.0 | 41.0 |  | 3.36 | 5 | Water 4 | | a:5343 | 3.67 | 1.0 | 43.5 |  | 5.92 | 6 | Water 5 | | a:8797 | 3.92 | 1.0 | 50.9 |  | 3.41 | 7 | Water 6 | | a:3750 | 3.93 | 1.0 | 40.0 |  | 4.23 | 8 | A | OP2 | a:1030 | 4.24 | 1.0 | 37.3 |  | 8.44 | 9 | Water 7 | | a:4427 | 4.31 | 1.0 | 41.2 |  | 6.85 | 10 | Water 8 | | a:8157 | 4.40 | 1.0 | 36.1 |  | 7.81 | 11 | G | O6 | a:1126 | 4.42 | 1.0 | 32.2 |  | 9.71 | 12 | Water 9 | | a:5706 | 4.49 | 1.0 | 39.7 |  | 7.24 | 13 | G | N7 | a:1126 | 4.51 | 1.0 | 28.3 |  | 10.86 |

  
**1^st^ shell coordination angles and distances (*trans* in "blue"; *cys* in "green"; deviations in "orange"):**   
• 1-2(W/W): *77º*; 1-3(W/O6): 89º; 2-3(W/O6): 88º;   
• 1-2(W/W): 2.6Å; 1-3(W/O6): 2.9Å; 2-3(W/O6): 3.1Å;   
  
**2.76 ± 0.76: the average coodination distance of the 1^st^ six ligands is K^+^ compatible**  
**10: Number of potential Water contacts**   
**2 : Number of second shell contacts involving water or ions**   
**20%: Percentage of satisfied second shell Water contacts**   
• Contact(s): /O6/O_w_/  
• Contact code: O_b_O_w_  
• Contact(s) in the exclusion zone: /O_w_/O_w_/  
→ Missing octahedral coordination for MG ←  
→2Fo-Fc density between 1.0 and 4.0 !   
Go to ToC

### MG:a:3209

|  | Res. type | Atom | Residue | Distance | Occupancy | B-factor | Closest atom(s) to a 1^st^ shell water | EM\_dens. |
| --- | --- | --- | --- | --- | --- | --- | --- | --- |
| **Ion** | | | |  | 1.0 | 40.1 |  | 13.99 |
 1 | Water 1 | | a:9039 | 2.01 | 1.0 | 32.3 |  | 9.39 | 2 | U | OP1 | a:1134 | 2.19 | 1.0 | 32.5 |  | 11.44 | 3 | Water 2 | | a:9481 | 2.23 | 1.0 | 32.4 |  | 8.99 | 4 | Water 3 | | a:10073 | 2.24 | 1.0 | 37.3 |  | 10.08 | 5 | U | OP2 | a:1134 | 4.15 | 1.0 | 26.1 |  | 11.35 | 6 | Water 4 | | a:4762 | 4.20 | 1.0 | 35.3 | 2.65 [A:OP1]a:1131 | 9.96 | 7 | Water 5 | | a:9939 | 4.23 | 1.0 | 39.2 |  | 7.71 | 8 | Water 6 | | a:3495 | 4.31 | 1.0 | 30.5 |  | 11.81 | 9 | A | OP1 | a:1131 | 4.38 | 1.0 | 29.8 |  | 8.99 | 10 | Water 7 | | i:269 | 4.42 | 1.0 | 41.4 |  | 4.67 | 11 | A | OP1 | a:2520 | 4.44 | 1.0 | 29.9 |  | 12.30 | 12 | Water 8 | | a:4125 | 4.45 | 1.0 | 41.4 | 2.59 [U:OP2]a:1134 | 7.75 | 13 | U | O5' | a:1134 | 4.46 | 1.0 | 34.2 |  | 10.61 | 14 | Water 9 | | a:10256 | 4.50 | 1.0 | 44.3 |  | 3.30 | 15 | G | O3' | a:1133 | 4.63 | 1.0 | 29.5 |  | 11.59 |

  
**1^st^ shell coordination angles and distances (*trans* in "blue"; *cys* in "green"; deviations in "orange"):**   
• 1-2(W/OP1): 91º; 1-3(W/W): 87º; 1-4(W/W): 87º; 2-3(OP1/W): 89º; 2-4(OP1/W): *165º*; 3-4(W/W): *76º*;   
• 1-2(W/OP1): 3.0Å; 1-3(W/W): 2.9Å; 1-4(W/W): 2.9Å; 2-3(OP1/W): 3.1Å; 2-4(OP1/W): *4.4Å*; 3-4(W/W): 2.7Å;   
  
**2.84 ± 0.95: the average coodination distance of the 1^st^ six ligands is K^+^ compatible**  
**8: Number of potential Water contacts**   
**1 : Number of second shell contacts involving water or ions**   
**12%: Percentage of satisfied second shell Water contacts**   
• Contact(s): /O_w_/O_w_/O_w_/OP1/  
• Contact code: O_ph_3O_w_  
→ Missing octahedral coordination for MG ←  
→ Slightly distorded coord. octahedron (6.3º) ←  
Go to ToC

### MG:a:3210

|  | Res. type | Atom | Residue | Distance | Occupancy | B-factor | Closest atom(s) to a 1^st^ shell water | EM\_dens. |
| --- | --- | --- | --- | --- | --- | --- | --- | --- |
| **Ion** | | | |  | 1.0 | 43.2 |  | 11.27 |
 1 | Water 1 | | a:4352 | 1.99 | 1.0 | 32.1 | 2.62 [U:O4]a:1205 | 9.21 | 2 | Water 2 | | a:8318 | 2.04 | 1.0 | 31.4 | 2.91 [G:N7]a:1204 // 2.94 [MG:MG]a:3032 | 10.36 | 3 | Water 3 | | a:8198 | 2.05 | 1.0 | 38.6 | 2.90 [A:N1]a:1206 | 7.89 | 4 | Water 4 | | a:10087 | 2.08 | 1.0 | 35.3 |  | 5.60 | 5 | Water 5 | | a:10060 | 2.25 | 1.0 | 41.2 |  | 5.37 | 6 | Water 6 | | a:5859 | 2.29 | 1.0 | 33.3 | 2.73 [G:OP2]a:1204 | 9.37 | 7 | U | O4 | a:1205 | 3.75 | 1.0 | 33.2 |  | 8.25 | 8 | Water 7 | | a:9817 | 3.97 | 1.0 | 47.9 |  | 3.97 | 9 | Water 8 | | a:5820 | 3.98 | 1.0 | 44.3 |  | 4.94 | 10 | **MG** | MG | a:3032 | 4.02 | 1.0 | 25.8 |  | 19.71 | 11 | A | N1 | a:1206 | 4.14 | 1.0 | 29.9 |  | 8.96 | 12 | U | OP2 | a:1203 | 4.28 | 1.0 | 34.8 |  | 9.48 | 13 | Water 9 | | a:7746 | 4.35 | 1.0 | 39.8 | 2.86 [G:OP2]a:1204 | 4.87 | 14 | G | N7 | a:1204 | 4.37 | 1.0 | 26.4 |  | 12.76 | 15 | G | OP2 | a:1204 | 4.54 | 1.0 | 34.9 |  | 8.99 |

  
**1^st^ shell coordination angles and distances (*trans* in "blue"; *cys* in "green"; deviations in "orange"):**   
• 1-2(W/W): 89º; 1-3(W/W): 93º; 1-4(W/W): 91º; 1-5(W/W): 173º; 1-6(W/W): *84º*; 2-3(W/W): 92º; 2-4(W/W): 175º; 2-5(W/W): *84º*; 2-6(W/W): 90º; 3-4(W/W): 93º; 3-5(W/W): 89º; 3-6(W/W): 176º; 4-5(W/W): *96º*; 4-6(W/W): *85º*; 5-6(W/W): 94º;   
• 1-2(W/W): 2.8Å; 1-3(W/W): 2.9Å; 1-4(W/W): 2.9Å; 1-5(W/W): *4.2Å*; 1-6(W/W): 2.9Å; 2-3(W/W): 3.0Å; 2-4(W/W): 4.1Å; 2-5(W/W): 2.9Å; 2-6(W/W): 3.1Å; 3-4(W/W): 3.0Å; 3-5(W/W): 3.0Å; 3-6(W/W): *4.3Å*; 4-5(W/W): *3.2Å*; 4-6(W/W): 2.9Å; 5-6(W/W): *3.3Å*;   
  
**2.12 ± 0.11: the average coodination distance of the 1^st^ six ligands is Mg^2+^ compatible**  
**12: Number of potential Water contacts**   
**5 : Number of second shell contacts involving water or ions**   
**42%: Percentage of satisfied second shell Water contacts**   
• Contact(s): /O_w_/O_w_/O_w_/O_w_/O_w_/O_w_/  
• Contact code: 6O_w_  
Ooctahedral coordination (3.7º)   
Go to ToC

### MG:a:3211

|  | Res. type | Atom | Residue | Distance | Occupancy | B-factor | Closest atom(s) to a 1^st^ shell water | EM\_dens. |
| --- | --- | --- | --- | --- | --- | --- | --- | --- |
| **Ion** | | | |  | 1.0 | 35.9 |  | 16.39 |
 1 | Water 1 | | a:5123 | 1.91 | 1.0 | 32.7 | 2.68 [A:OP1]a:2018 | 11.60 | 2 | Water 2 | | a:6755 | 2.09 | 1.0 | 29.5 |  | 10.63 | 3 | Water 3 | | a:7466 | 2.15 | 1.0 | 34.3 | 2.90 [A:N7]a:1264 | 9.35 | 4 | U | O4 | a:1265 | 2.22 | 1.0 | 23.8 |  | 14.67 | 5 | Water 4 | | a:8799 | 2.28 | 1.0 | 27.6 | 2.97 [U:O4]a:2020 | 9.31 | 6 | A | OP1 | a:2018 | 3.90 | 1.0 | 26.7 |  | 13.31 | 7 | Water 5 | | a:9864 | 4.06 | 1.0 | 43.8 |  | 4.28 | 8 | A | OP2 | a:2019 | 4.07 | 1.0 | 32.6 |  | 10.84 | 9 | A | N1 | a:1266 | 4.23 | 1.0 | 16.6 |  | 16.87 | 10 | U | O4 | a:2020 | 4.27 | 1.0 | 26.6 |  | 16.24 | 11 | A | OP2 | a:2018 | 4.44 | 1.0 | 28.5 |  | 12.84 | 12 | A | N7 | a:1264 | 4.45 | 1.0 | 21.1 |  | 15.74 | 13 | Water 6 | | a:9013 | 4.54 | 1.0 | 45.0 | 3.00 [A:OP1]a:2018 | 4.16 |

  
**1^st^ shell coordination angles and distances (*trans* in "blue"; *cys* in "green"; deviations in "orange"):**   
• 1-2(W/W): *83º*; 1-3(W/W): *105º*; 1-4(W/O4): *163º*; 1-5(W/W): *99º*; 2-3(W/W): 91º; 2-4(W/O4): *81º*; 2-5(W/W): 173º; 3-4(W/O4): *81º*; 3-5(W/W): *95º*; 4-5(O4/W): *96º*;   
• 1-2(W/W): 2.7Å; 1-3(W/W): *3.2Å*; 1-4(W/O4): 4.1Å; 1-5(W/W): 3.2Å; 2-3(W/W): 3.0Å; 2-4(W/O4): 2.8Å; 2-5(W/W): *4.4Å*; 3-4(W/O4): 2.8Å; 3-5(W/W): *3.3Å*; 4-5(O4/W): *3.3Å*;   
  
**2.42 ± 0.67: the average coodination distance of the 1^st^ six ligands is Na^+^** compatible  
**8: Number of potential Water contacts**   
**3 : Number of second shell contacts involving water or ions**   
**38%: Percentage of satisfied second shell Water contacts**   
• Contact(s): /O4/O_w_/O_w_/O_w_/O_w_/  
• Contact code: O_b_4O_w_  
→ Missing octahedral coordination for MG ←  
→ Slightly distorded coord. octahedron (8.7º) ←  
Go to ToC

### MG:a:3212

|  | Res. type | Atom | Residue | Distance | Occupancy | B-factor | Closest atom(s) to a 1^st^ shell water | EM\_dens. |
| --- | --- | --- | --- | --- | --- | --- | --- | --- |
| **Ion** | | | |  | 1.0 | 37.2 |  | 15.10 |
 1 | Water 1 | | a:7510 | 1.93 | 1.0 | 30.0 | 2.84 [G:N7]a:2052 | 12.49 | 2 | Water 2 | | a:7799 | 2.05 | 1.0 | 32.1 | 2.86 [G:N7]a:2053 | 12.72 | 3 | Water 3 | | a:4656 | 2.09 | 1.0 | 32.9 | 2.65 [G:O6]a:2052 // 2.73 [G:O6]a:2053 | 11.17 | 4 | Water 4 | | a:10248 | 2.10 | 1.0 | 32.0 |  | 10.31 | 5 | Water 5 | | a:9957 | 2.17 | 1.0 | 36.5 |  | 10.40 | 6 | Water 6 | | a:10069 | 2.21 | 1.0 | 33.1 |  | 11.17 | 7 | Water 7 | | a:4842 | 3.60 | 1.0 | 39.9 |  | 5.84 | 8 | Water 8 | | a:4801 | 3.94 | 1.0 | 30.5 |  | 9.23 | 9 | Water 9 | | a:9739 | 4.12 | 1.0 | 39.9 |  | 5.65 | 10 | G | O6 | a:2053 | 4.12 | 1.0 | 20.8 |  | 16.61 | 11 | Water 10 | | a:9253 | 4.14 | 1.0 | 33.2 |  | 8.72 | 12 | G | N7 | a:2052 | 4.17 | 1.0 | 19.0 |  | 14.56 | 13 | G | N7 | a:2053 | 4.20 | 1.0 | 19.5 |  | 13.40 | 14 | G | O6 | a:2052 | 4.36 | 1.0 | 22.3 |  | 17.90 | 15 | Water 11 | | a:9806 | 4.37 | 1.0 | 39.3 |  | 7.16 | 16 | Water 12 | | a:4274 | 4.40 | 1.0 | 38.0 |  | 10.29 | 17 | Water 13 | | a:9766 | 4.41 | 1.0 | 40.7 |  | 6.97 | 18 | Water 14 | | a:4104 | 4.52 | 1.0 | 32.0 |  | 11.95 |

  
**1^st^ shell coordination angles and distances (*trans* in "blue"; *cys* in "green"; deviations in "orange"):**   
• 1-2(W/W): 94º; 1-3(W/W): 89º; 1-4(W/W): 90º; 1-5(W/W): 91º; 1-6(W/W): 178º; 2-3(W/W): *98º*; 2-4(W/W): 173º; 2-5(W/W): 89º; 2-6(W/W): 87º; 3-4(W/W): 88º; 3-5(W/W): 173º; 3-6(W/W): 92º; 4-5(W/W): *85º*; 4-6(W/W): 88º; 5-6(W/W): 88º;   
• 1-2(W/W): 2.9Å; 1-3(W/W): 2.8Å; 1-4(W/W): 2.8Å; 1-5(W/W): 2.9Å; 1-6(W/W): 4.1Å; 2-3(W/W): 3.1Å; 2-4(W/W): 4.1Å; 2-5(W/W): 3.0Å; 2-6(W/W): 2.9Å; 3-4(W/W): 2.9Å; 3-5(W/W): *4.3Å*; 3-6(W/W): 3.1Å; 4-5(W/W): 2.9Å; 4-6(W/W): 3.0Å; 5-6(W/W): 3.0Å;   
  
**2.09 ± 0.09: the average coodination distance of the 1^st^ six ligands is Mg^2+^ compatible**  
**12: Number of potential Water contacts**   
**4 : Number of second shell contacts involving water or ions**   
**33%: Percentage of satisfied second shell Water contacts**   
• Contact(s): /O_w_/O_w_/O_w_/O_w_/O_w_/O_w_/  
• Contact code: 6O_w_  
Ooctahedral coordination (3.1º)   
Go to ToC

### MG:a:3213

|  | Res. type | Atom | Residue | Distance | Occupancy | B-factor | Closest atom(s) to a 1^st^ shell water | EM\_dens. |
| --- | --- | --- | --- | --- | --- | --- | --- | --- |
| **Ion** | | | |  | 1.0 | 37.6 |  | 15.21 |
 1 | Water 1 | | a:7697 | 2.03 | 1.0 | 29.8 | 2.86 [G:O4']a:1290 | 11.95 | 2 | Water 2 | | a:8764 | 2.07 | 1.0 | 30.9 |  | 8.70 | 3 | Water 3 | | a:9899 | 2.10 | 1.0 | 31.6 |  | 8.34 | 4 | Water 4 | | a:9982 | 2.20 | 1.0 | 39.8 |  | 5.61 | 5 | Water 5 | | a:9965 | 2.28 | 1.0 | 38.1 |  | 8.21 | 6 | Water 6 | | a:3850 | 2.33 | 1.0 | 37.8 | 2.55 [C:OP2]a:1648 | 7.03 | 7 | Water 7 | | a:9922 | 4.07 | 1.0 | 36.7 |  | 8.03 | 8 | Water 8 | | a:6899 | 4.12 | 1.0 | 37.5 |  | 8.71 | 9 | G | O4' | a:1290 | 4.30 | 1.0 | 31.7 |  | 11.50 | 10 | G | OP1 | a:1651 | 4.31 | 1.0 | 38.3 |  | 4.77 | 11 | A | OP2 | a:1278 | 4.32 | 1.0 | 30.7 |  | 11.36 | 12 | Water 9 | | a:7694 | 4.47 | 1.0 | 33.1 |  | 8.95 | 13 | A | N3 | a:1289 | 4.48 | 1.0 | 23.2 |  | 13.91 | 14 | C | OP2 | a:1648 | 4.48 | 1.0 | 37.8 |  | 8.49 | 15 | Water 10 | | a:6907 | 4.53 | 1.0 | 46.2 |  | 3.82 | 16 | Water 11 | | a:5275 | 4.67 | 1.0 | 42.6 |  | 4.23 |

  
**1^st^ shell coordination angles and distances (*trans* in "blue"; *cys* in "green"; deviations in "orange"):**   
• 1-2(W/W): 89º; 1-3(W/W): 86º; 1-4(W/W): *98º*; 1-5(W/W): 88º; 2-3(W/W): 93º; 2-4(W/W): 94º; 2-5(W/W): 174º; 3-4(W/W): 173º; 3-5(W/W): 92º; 4-5(W/W): *81º*;   
• 1-2(W/W): 2.9Å; 1-3(W/W): 2.8Å; 1-4(W/W): 3.2Å; 1-5(W/W): 3.0Å; 2-3(W/W): 3.0Å; 2-4(W/W): 3.1Å; 2-5(W/W): *4.3Å*; 3-4(W/W): *4.3Å*; 3-5(W/W): 3.2Å; 4-5(W/W): 2.9Å;   
  
**2.17 ± 0.11: the average coodination distance of the 1^st^ six ligands is Mg^2+^ compatible**  
**12: Number of potential Water contacts**   
**2 : Number of second shell contacts involving water or ions**   
**17%: Percentage of satisfied second shell Water contacts**   
• Contact(s): /O_w_/O_w_/O_w_/O_w_/O_w_/  
• Contact code: 5O_w_  
• Contact(s) in the exclusion zone: /O_w_/  
→ Missing octahedral coordination for MG ←  
Ooctahedral coordination (4.7º)   
Go to ToC

### MG:a:3214

|  | Res. type | Atom | Residue | Distance | Occupancy | B-factor | Closest atom(s) to a 1^st^ shell water | EM\_dens. |
| --- | --- | --- | --- | --- | --- | --- | --- | --- |
| **Ion** | | | |  | 1.0 | 25.6 |  | 26.25 |
 1 | Water 1 | | a:7957 | 1.92 | 1.0 | 21.1 | 2.88 [C:OP1]a:946 | 17.34 | 2 | Water 2 | | k:264 | 1.97 | 1.0 | 20.4 |  | 15.61 | 3 | Water 3 | | a:7377 | 2.03 | 1.0 | 22.7 | 2.83 [C:OP2]a:946 | 16.27 | 4 | Water 4 | | a:4402 | 2.13 | 1.0 | 23.6 | 2.62 [U:OP1]a:570 // 2.83 [A:N7]a:947 | 17.75 | 5 | Water 5 | | a:8872 | 2.30 | 1.0 | 21.6 | 2.97 [A:N1]a:2452 | 18.12 | 6 | Water 6 | | a:6468 | 2.38 | 1.0 | 25.3 | 2.77 [G:OP1]a:833 | 15.62 | 7 | Water 7 | | a:3979 | 3.95 | 1.0 | 27.8 |  | 14.53 | 8 | A | N7 | a:947 | 3.97 | 1.0 | 19.1 |  | 16.51 | 9 | Water 8 | | a:5975 | 3.99 | 1.0 | 26.4 |  | 11.80 | 10 | G | OP1 | a:833 | 4.03 | 1.0 | 27.8 |  | 13.26 | 11 | Water 9 | | a:4670 | 4.04 | 1.0 | 27.4 | 2.87 [C:OP2]a:946 | 13.70 | 12 | C | OP2 | a:946 | 4.19 | 1.0 | 25.0 |  | 16.09 | 13 | C | OP1 | a:946 | 4.28 | 1.0 | 25.2 |  | 13.98 | 14 | Water 10 | | a:5144 | 4.46 | 1.0 | 26.9 |  | 14.02 | 15 | U | OP1 | a:570 | 4.58 | 1.0 | 24.9 |  | 13.08 | 16 | A | N1 | a:2452 | 4.64 | 1.0 | 15.7 |  | 17.57 | 17 | HIS | O | k:35 | 4.74 | 1.0 | 30.9 |  | 14.67 |

  
**1^st^ shell coordination angles and distances (*trans* in "blue"; *cys* in "green"; deviations in "orange"):**   
• 1-2(W/W): *97º*; 1-3(W/W): *100º*; 1-4(W/W): 93º; 2-3(W/W): 92º; 2-4(W/W): 91º; 3-4(W/W): *167º*;   
• 1-2(W/W): 2.9Å; 1-3(W/W): 3.0Å; 1-4(W/W): 2.9Å; 2-3(W/W): 2.9Å; 2-4(W/W): 2.9Å; 3-4(W/W): 4.1Å;   
  
**2.12 ± 0.17: the average coodination distance of the 1^st^ six ligands is Mg^2+^ compatible**  
**12: Number of potential Water contacts**   
**6 : Number of second shell contacts involving water or ions**   
**50%: Percentage of satisfied second shell Water contacts**   
• Contact(s): /O_w_/O_w_/O_w_/O_w_/  
• Contact code: 4O_w_  
• Contact(s) in the exclusion zone: /O_w_/O_w_/  
→ Missing octahedral coordination for MG ←  
→ Slightly distorded coord. octahedron (6.0º) ←  
Go to ToC

### MG:a:3215

|  | Res. type | Atom | Residue | Distance | Occupancy | B-factor | Closest atom(s) to a 1^st^ shell water | EM\_dens. |
| --- | --- | --- | --- | --- | --- | --- | --- | --- |
| **Ion** | | | |  | 1.0 | 26.4 |  | 27.58 |
 1 | Water 1 | | a:6268 | 1.91 | 1.0 | 23.0 | 2.76 [G:O6]a:579 | 17.25 | 2 | Water 2 | | a:9352 | 2.08 | 1.0 | 23.2 |  | 17.34 | 3 | Water 3 | | a:9742 | 2.11 | 1.0 | 23.6 |  | 16.88 | 4 | Water 4 | | a:3733 | 2.26 | 1.0 | 19.9 | 2.52 [U:O4]a:578 | 20.01 | 5 | Water 5 | | a:4853 | 2.31 | 1.0 | 26.4 | 2.66 [A:OP2]a:576 // 2.89 [U:OP2]a:578 | 17.88 | 6 | A | OP2 | a:576 | 2.34 | 1.0 | 25.3 |  | 11.76 | 7 | Water 6 | | a:8227 | 3.87 | 1.0 | 31.0 | 2.90 [A:OP1]a:576 | 12.28 | 8 | Water 7 | | a:7597 | 4.05 | 1.0 | 24.8 |  | 15.80 | 9 | Water 8 | | a:6924 | 4.15 | 1.0 | 27.5 |  | 14.71 | 10 | U | O4 | a:578 | 4.15 | 1.0 | 24.6 |  | 17.84 | 11 | A | OP1 | a:576 | 4.16 | 1.0 | 28.0 |  | 11.74 | 12 | Water 9 | | a:5250 | 4.19 | 1.0 | 32.4 | 2.69 [U:OP2]a:578 | 13.54 | 13 | G | O6 | a:579 | 4.20 | 1.0 | 19.8 |  | 15.84 | 14 | Water 10 | | a:5867 | 4.31 | 1.0 | 25.3 | 2.82 [U:OP1]a:575 | 14.49 | 15 | U | OP1 | a:575 | 4.37 | 1.0 | 28.1 |  | 15.79 | 16 | A | O5' | a:576 | 4.51 | 1.0 | 34.2 |  | 3.42 | 17 | U | OP2 | a:578 | 4.72 | 1.0 | 23.3 |  | 15.22 | 18 | U | O3' | a:575 | 4.76 | 1.0 | 29.9 |  | 14.16 |

  
**1^st^ shell coordination angles and distances (*trans* in "blue"; *cys* in "green"; deviations in "orange"):**   
• 1-2(W/W): 94º; 1-3(W/W): *98º*; 1-4(W/W): 89º; 2-3(W/W): 92º; 2-4(W/W): 176º; 3-4(W/W): 90º;   
• 1-2(W/W): 2.9Å; 1-3(W/W): 3.0Å; 1-4(W/W): 2.9Å; 2-3(W/W): 3.0Å; 2-4(W/W): *4.3Å*; 3-4(W/W): 3.1Å;   
  
**2.17 ± 0.15: the average coodination distance of the 1^st^ six ligands is Mg^2+^ compatible**  
**10: Number of potential Water contacts**   
**4 : Number of second shell contacts involving water or ions**   
**40%: Percentage of satisfied second shell Water contacts**   
• Contact(s): /O_w_/O_w_/O_w_/O_w_/  
• Contact code: 4O_w_  
• Contact(s) in the exclusion zone: /O_w_/OP2/  
→ Missing octahedral coordination for MG ←  
Ooctahedral coordination (3.4º)   
Go to ToC

### MG:a:3216

|  | Res. type | Atom | Residue | Distance | Occupancy | B-factor | Closest atom(s) to a 1^st^ shell water | EM\_dens. |
| --- | --- | --- | --- | --- | --- | --- | --- | --- |
| **Ion** | | | |  | 1.0 | 33.3 |  | 22.55 |
 1 | Water 1 | | a:10016 | 2.04 | 1.0 | 32.8 |  | 9.77 | 2 | Water 2 | | a:6228 | 2.12 | 1.0 | 27.0 | 2.76 [G:O6]a:1261 | 12.81 | 3 | Water 3 | | a:10152 | 2.19 | 1.0 | 32.7 |  | 12.88 | 4 | Water 4 | | a:10285 | 2.22 | 1.0 | 33.4 |  | 9.56 | 5 | Water 5 | | a:4090 | 2.22 | 1.0 | 26.2 | 2.59 [U:O4]a:582 | 13.56 | 6 | Water 6 | | a:4805 | 2.22 | 1.0 | 26.3 | 2.66 [U:O4]a:1260 | 14.46 | 7 | Water 7 | | a:6036 | 3.98 | 1.0 | 28.1 |  | 12.52 | 8 | Water 8 | | a:9516 | 4.25 | 1.0 | 23.6 |  | 15.86 | 9 | U | O4 | a:582 | 4.25 | 1.0 | 22.8 |  | 17.09 | 10 | Water 9 | | a:4538 | 4.34 | 1.0 | 24.6 |  | 15.74 | 11 | Water 10 | | a:4799 | 4.36 | 1.0 | 42.5 |  | 6.44 | 12 | U | O4 | a:1260 | 4.37 | 1.0 | 21.6 |  | 17.33 | 13 | Water 11 | | a:4164 | 4.40 | 1.0 | 34.3 |  | 10.88 | 14 | G | O6 | a:1261 | 4.41 | 1.0 | 22.5 |  | 17.95 | 15 | Water 12 | | a:8096 | 4.44 | 1.0 | 40.5 |  | 7.23 | 16 | Water 13 | | a:9544 | 4.44 | 1.0 | 36.3 |  | 6.41 | 17 | Water 14 | | a:6081 | 4.45 | 1.0 | 28.2 |  | 12.77 | 18 | Water 15 | | a:8524 | 4.48 | 1.0 | 24.5 |  | 11.96 | 19 | Water 16 | | a:10209 | 4.66 | 1.0 | 33.0 |  | 9.21 |

  
**1^st^ shell coordination angles and distances (*trans* in "blue"; *cys* in "green"; deviations in "orange"):**   
• 1-2(W/W): 174º; 1-3(W/W): 88º; 1-4(W/W): 94º; 1-5(W/W): 94º; 1-6(W/W): 90º; 2-3(W/W): 87º; 2-4(W/W): 88º; 2-5(W/W): 92º; 2-6(W/W): 87º; 3-4(W/W): 85º; 3-5(W/W): 172º; 3-6(W/W): 90º; 4-5(W/W): 87º; 4-6(W/W): 173º; 5-6(W/W): *98º*;   
• 1-2(W/W): 4.2Å; 1-3(W/W): 2.9Å; 1-4(W/W): 3.1Å; 1-5(W/W): 3.1Å; 1-6(W/W): 3.0Å; 2-3(W/W): 3.0Å; 2-4(W/W): 3.0Å; 2-5(W/W): 3.1Å; 2-6(W/W): 3.0Å; 3-4(W/W): 3.0Å; 3-5(W/W): *4.4Å*; 3-6(W/W): 3.1Å; 4-5(W/W): 3.1Å; 4-6(W/W): *4.4Å*; 5-6(W/W): *3.3Å*;   
  
**2.17 ± 0.07: the average coodination distance of the 1^st^ six ligands is Mg^2+^ compatible**  
**12: Number of potential Water contacts**   
**3 : Number of second shell contacts involving water or ions**   
**25%: Percentage of satisfied second shell Water contacts**   
• Contact(s): /O_w_/O_w_/O_w_/O_w_/O_w_/O_w_/  
• Contact code: 6O_w_  
Ooctahedral coordination (3.9º)   
Go to ToC

### MG:a:3217

|  | Res. type | Atom | Residue | Distance | Occupancy | B-factor | Closest atom(s) to a 1^st^ shell water | EM\_dens. |
| --- | --- | --- | --- | --- | --- | --- | --- | --- |
| **Ion** | | | |  | 1.0 | 26.8 |  | 25.15 |
 1 | Water 1 | | a:9980 | 1.91 | 1.0 | 28.3 |  | 12.71 | 2 | Water 2 | | a:5994 | 2.09 | 1.0 | 22.2 | 2.74 [A:OP2]a:584 | 14.82 | 3 | Water 3 | | a:4056 | 2.11 | 1.0 | 27.0 | 2.58 [A:OP2]a:1255 | 17.76 | 4 | Water 4 | | a:6811 | 2.22 | 1.0 | 22.9 | 2.79 [C:O2']a:1253 | 15.96 | 5 | Water 5 | | a:6745 | 2.29 | 1.0 | 25.0 | 2.79 [A:N7]a:584 | 15.82 | 6 | Water 6 | | a:8556 | 2.39 | 1.0 | 27.7 | 2.94 [G:N7]a:585 | 16.54 | 7 | A | OP2 | a:584 | 4.25 | 1.0 | 27.8 |  | 14.37 | 8 | C | O2' | a:1253 | 4.25 | 1.0 | 22.5 |  | 19.00 | 9 | Water 7 | | p:246 | 4.31 | 1.0 | 29.2 |  | 12.80 | 10 | G | N7 | a:585 | 4.32 | 1.0 | 16.8 |  | 20.40 | 11 | A | OP2 | a:1255 | 4.47 | 1.0 | 23.5 |  | 12.73 | 12 | Water 8 | | a:3745 | 4.49 | 1.0 | 22.3 |  | 16.37 | 13 | Water 9 | | a:5007 | 4.54 | 1.0 | 22.3 | 2.67 [A:OP2]a:584 | 17.47 | 14 | A | N7 | a:584 | 4.60 | 1.0 | 19.3 |  | 18.24 |

  
**1^st^ shell coordination angles and distances (*trans* in "blue"; *cys* in "green"; deviations in "orange"):**   
• 1-2(W/W): 90º; 1-3(W/W): 92º; 1-4(W/W): *99º*; 1-5(W/W): *95º*; 2-3(W/W): 177º; 2-4(W/W): 86º; 2-5(W/W): 94º; 3-4(W/W): *96º*; 3-5(W/W): *84º*; 4-5(W/W): *166º*;   
• 1-2(W/W): 2.8Å; 1-3(W/W): 2.9Å; 1-4(W/W): 3.1Å; 1-5(W/W): 3.1Å; 2-3(W/W): 4.2Å; 2-4(W/W): 2.9Å; 2-5(W/W): *3.2Å*; 3-4(W/W): *3.2Å*; 3-5(W/W): 2.9Å; 4-5(W/W): *4.5Å*;   
  
**2.17 ± 0.16: the average coodination distance of the 1^st^ six ligands is Mg^2+^ compatible**  
**12: Number of potential Water contacts**   
**5 : Number of second shell contacts involving water or ions**   
**42%: Percentage of satisfied second shell Water contacts**   
• Contact(s): /O_w_/O_w_/O_w_/O_w_/O_w_/  
• Contact code: 5O_w_  
• Contact(s) in the exclusion zone: /O_w_/  
→ Missing octahedral coordination for MG ←  
→ Slightly distorded coord. octahedron (5.4º) ←  
Go to ToC

### MG:a:3218

|  | Res. type | Atom | Residue | Distance | Occupancy | B-factor | Closest atom(s) to a 1^st^ shell water | EM\_dens. |
| --- | --- | --- | --- | --- | --- | --- | --- | --- |
| **Ion** | | | |  | 1.0 | 38.8 |  | 12.18 |
 1 | Water 1 | | a:10154 | 1.97 | 1.0 | 43.3 |  | 7.14 | 2 | Water 2 | | a:10090 | 2.01 | 1.0 | 39.0 |  | 6.41 | 3 | Water 3 | | a:4097 | 2.13 | 1.0 | 35.0 | 2.59 [G:O6]a:620 | 8.76 | 4 | Water 4 | | a:7648 | 2.13 | 1.0 | 35.4 | 2.85 [G:N7]a:619 | 7.83 | 5 | Water 5 | | a:5211 | 2.16 | 1.0 | 36.6 | 2.69 [G:N7]a:620 | 10.05 | 6 | G | O6 | a:621 | 3.95 | 1.0 | 36.1 |  | 8.42 | 7 | Water 6 | | a:6003 | 4.07 | 1.0 | 38.0 |  | 6.16 | 8 | Water 7 | | a:3528 | 4.25 | 1.0 | 46.0 | 2.85 [A:OP1]a:618 | 5.56 | 9 | G | O6 | a:620 | 4.30 | 1.0 | 36.4 |  | 9.23 | 10 | Water 8 | | a:7755 | 4.38 | 1.0 | 41.1 |  | 4.36 | 11 | G | N7 | a:619 | 4.42 | 1.0 | 34.1 |  | 8.32 | 12 | G | N7 | a:620 | 4.42 | 1.0 | 33.3 |  | 9.98 | 13 | Water 9 | | a:3563 | 4.43 | 1.0 | 34.3 |  | 8.93 | 14 | A | OP2 | a:618 | 4.59 | 1.0 | 45.8 |  | 6.65 | 15 | G | O6 | a:619 | 4.65 | 1.0 | 37.9 |  | 9.52 | 16 | A | OP1 | a:618 | 4.67 | 1.0 | 46.0 |  | 4.20 | 17 | A | OP2 | a:610 | 4.75 | 1.0 | 35.0 |  | 10.04 |

  
**1^st^ shell coordination angles and distances (*trans* in "blue"; *cys* in "green"; deviations in "orange"):**   
• 1-2(W/W): 89º; 1-3(W/W): 178º; 1-4(W/W): *97º*; 1-5(W/W): *97º*; 2-3(W/W): 93º; 2-4(W/W): 88º; 2-5(W/W): 173º; 3-4(W/W): *82º*; 3-5(W/W): *80º*; 4-5(W/W): 88º;   
• 1-2(W/W): 2.8Å; 1-3(W/W): 4.1Å; 1-4(W/W): 3.1Å; 1-5(W/W): 3.1Å; 2-3(W/W): 3.0Å; 2-4(W/W): 2.9Å; 2-5(W/W): 4.2Å; 3-4(W/W): 2.8Å; 3-5(W/W): 2.8Å; 4-5(W/W): 3.0Å;   
  
**2.39 ± 0.70: the average coodination distance of the 1^st^ six ligands is Na^+^** compatible  
**10: Number of potential Water contacts**   
**3 : Number of second shell contacts involving water or ions**   
**30%: Percentage of satisfied second shell Water contacts**   
• Contact(s): /O_w_/O_w_/O_w_/O_w_/O_w_/  
• Contact code: 5O_w_  
→ Missing octahedral coordination for MG ←  
Ooctahedral coordination (4.9º)   
Go to ToC

### MG:a:3219

|  | Res. type | Atom | Residue | Distance | Occupancy | B-factor | Closest atom(s) to a 1^st^ shell water | EM\_dens. |
| --- | --- | --- | --- | --- | --- | --- | --- | --- |
| **Ion** | | | |  | 1.0 | 39.0 |  | 12.40 |
 1 | Water 1 | | a:8205 | 1.96 | 1.0 | 39.2 |  | 6.54 | 2 | A | OP2 | a:623 | 2.03 | 1.0 | 38.0 |  | 7.02 | 3 | Water 2 | | k:215 | 2.09 | 1.0 | 35.7 |  | 5.54 | 4 | G | OP2 | a:624 | 2.10 | 1.0 | 32.0 |  | 10.23 | 5 | ASN | OD1 | k:99 | 2.14 | 1.0 | 40.0 |  | 7.27 | 6 | Water 3 | | k:238 | 2.18 | 1.0 | 38.6 |  | 6.06 | 7 | A | O5' | a:623 | 4.12 | 1.0 | 41.1 |  | 8.31 | 8 | Water 4 | | a:3902 | 4.22 | 1.0 | 42.7 | 2.56 [G:OP1]a:624 | 6.10 | 9 | G | OP1 | a:624 | 4.23 | 1.0 | 38.8 |  | 6.19 | 10 | Water 5 | | k:206 | 4.24 | 1.0 | 37.7 |  | 6.78 | 11 | A | OP1 | a:623 | 4.24 | 1.0 | 33.5 |  | 8.51 | 12 | Water 6 | | a:8775 | 4.31 | 1.0 | 42.2 |  | 6.53 | 13 | A | O3' | a:623 | 4.32 | 1.0 | 36.7 |  | 8.33 | 14 | Water 7 | | a:3705 | 4.37 | 1.0 | 38.8 |  | 4.43 | 15 | Water 8 | | a:7666 | 4.37 | 1.0 | 43.3 | 2.85 [G:OP1]a:624 | 5.01 | 16 | G | O3' | a:622 | 4.43 | 1.0 | 35.8 |  | 7.24 | 17 | G | O5' | a:624 | 4.51 | 1.0 | 40.6 |  | 7.46 |

  
**1^st^ shell coordination angles and distances (*trans* in "blue"; *cys* in "green"; deviations in "orange"):**   
• 1-2(W/OP2): *102º*; 1-3(W/W): 173º; 1-4(W/OP2): 91º; 1-5(W/OD1): *100º*; 1-6(W/W): 91º; 2-3(OP2/W): *84º*; 2-4(OP2/OP2): 94º; 2-5(OP2/OD1): 89º; 2-6(OP2/W): *166º*; 3-4(W/OP2): 89º; 3-5(W/OD1): *80º*; 3-6(W/W): *82º*; 4-5(OP2/OD1): *168º*; 4-6(OP2/W): 89º; 5-6(OD1/W): *85º*;   
• 1-2(W/OP2): 3.1Å; 1-3(W/W): 4.0Å; 1-4(W/OP2): 2.9Å; 1-5(W/OD1): 3.1Å; 1-6(W/W): 3.0Å; 2-3(OP2/W): 2.8Å; 2-4(OP2/OP2): 3.0Å; 2-5(OP2/OD1): 2.9Å; 2-6(OP2/W): 4.2Å; 3-4(W/OP2): 2.9Å; 3-5(W/OD1): 2.7Å; 3-6(W/W): 2.8Å; 4-5(OP2/OD1): *4.2Å*; 4-6(OP2/W): 3.0Å; 5-6(OD1/W): 2.9Å;   
  
**2.08 ± 0.07: the average coodination distance of the 1^st^ six ligands is Mg^2+^ compatible**  
**6: Number of potential Water contacts**   
**0 : Number of second shell contacts involving water or ions**   
**0%: Percentage of satisfied second shell Water contacts**   
• Contact(s): /O_w_/O_w_/O_w_/OD1_Asn_/OP2/OP2/  
• Contact code: *cis-*2O_ph_O_cno_3O_w_  
→ Slightly distorded coord. octahedron (6.2º) ←  
Go to ToC

### MG:a:3220

|  | Res. type | Atom | Residue | Distance | Occupancy | B-factor | Closest atom(s) to a 1^st^ shell water | EM\_dens. |
| --- | --- | --- | --- | --- | --- | --- | --- | --- |
| **Ion** | | | |  | 1.0 | 22.8 |  | 28.90 |
 1 | A | N7 | a:763 | 2.18 | 1.0 | 18.6 |  | 14.94 | 2 | Water 1 | | a:4849 | 2.19 | 1.0 | 24.2 | 2.66 [A:OP2]a:763 | 15.40 | 3 | Water 2 | | a:3585 | 2.25 | 1.0 | 23.9 | 2.48 [G:O6]a:735 | 14.12 | 4 | Water 3 | | a:8629 | 2.28 | 1.0 | 21.9 | 2.95 [G:OP2]a:735 | 15.42 | 5 | Water 4 | | a:5663 | 2.30 | 1.0 | 26.6 | 2.72 [C:OP2]a:734 // 2.91 [G:OP2]a:735 | 13.55 | 6 | G | N7 | a:735 | 2.43 | 1.0 | 17.6 |  | 17.42 | 7 | G | O6 | a:735 | 3.71 | 1.0 | 25.8 |  | 13.60 | 8 | Water 5 | | a:8445 | 4.01 | 1.0 | 36.7 |  | 7.93 | 9 | A | OP2 | a:763 | 4.03 | 1.0 | 23.5 |  | 14.39 | 10 | G | OP2 | a:735 | 4.05 | 1.0 | 30.6 |  | 9.90 | 11 | G | OP2 | a:762 | 4.45 | 1.0 | 25.2 |  | 13.35 | 12 | C | OP2 | a:734 | 4.46 | 1.0 | 31.4 |  | 10.47 | 13 | Water 6 | | a:8805 | 4.66 | 1.0 | 30.7 | 2.97 [C:OP2]a:734 | 12.83 | 14 | Water 7 | | a:4177 | 4.76 | 1.0 | 28.6 | 2.60 [A:OP2]a:763 | 10.30 |

  
**1^st^ shell coordination angles and distances (*trans* in "blue"; *cys* in "green"; deviations in "orange"):**   
• 1-2(N7/W): 91º; 1-3(N7/W): 90º; 1-4(N7/W): 91º; 1-5(N7/W): 175º; 2-3(W/W): 90º; 2-4(W/W): 95º; 2-5(W/W): 94º; 3-4(W/W): 175º; 3-5(W/W): 89º; 4-5(W/W): 90º;   
• 1-2(N7/W): 3.1Å; 1-3(N7/W): 3.1Å; 1-4(N7/W): 3.2Å; 1-5(N7/W): *4.5Å*; 2-3(W/W): 3.1Å; 2-4(W/W): *3.3Å*; 2-5(W/W): *3.3Å*; 3-4(W/W): *4.5Å*; 3-5(W/W): 3.2Å; 4-5(W/W): *3.2Å*;   
  
**2.27 ± 0.08: the average coodination distance of the 1^st^ six ligands is Mg^2+^ compatible**  
**8: Number of potential Water contacts**   
**5 : Number of second shell contacts involving water or ions**   
**62%: Percentage of satisfied second shell Water contacts**   
• Contact(s): /N7/O_w_/O_w_/O_w_/O_w_/  
• Contact code: N_b_4O_w_  
• Contact(s) in the exclusion zone: /N7/  
→ Missing octahedral coordination for MG ←  
Ooctahedral coordination (2.2º)   
Go to ToC

### MG:a:3221

|  | Res. type | Atom | Residue | Distance | Occupancy | B-factor | Closest atom(s) to a 1^st^ shell water | EM\_dens. |
| --- | --- | --- | --- | --- | --- | --- | --- | --- |
| **Ion** | | | |  | 1.0 | 33.8 |  | 17.77 |
 1 | Water 1 | | a:10282 | 1.93 | 1.0 | 33.9 |  | 11.12 | 2 | Water 2 | | a:7060 | 2.04 | 1.0 | 28.6 | 2.81 [A:N7]a:758 | 12.80 | 3 | Water 3 | | a:5633 | 2.20 | 1.0 | 29.2 | 2.72 [A:N1]a:741 | 10.11 | 4 | Water 4 | | a:4194 | 2.21 | 1.0 | 34.5 | 2.60 [G:O6]a:759 | 14.17 | 5 | Water 5 | | a:7326 | 2.21 | 1.0 | 27.8 | 2.83 [G:N7]a:759 | 12.62 | 6 | Water 6 | | a:10173 | 2.26 | 1.0 | 30.9 |  | 10.75 | 7 | Water 7 | | a:7767 | 3.82 | 1.0 | 29.9 |  | 10.63 | 8 | Water 8 | | a:9816 | 4.21 | 1.0 | 47.6 |  | 4.39 | 9 | Water 9 | | a:7517 | 4.32 | 1.0 | 44.2 |  | 3.80 | 10 | G | O6 | a:759 | 4.33 | 1.0 | 27.2 |  | 15.49 | 11 | G | N7 | a:759 | 4.38 | 1.0 | 17.6 |  | 14.64 | 12 | A | N7 | a:758 | 4.42 | 1.0 | 21.7 |  | 13.94 | 13 | A | N1 | a:741 | 4.45 | 1.0 | 23.0 |  | 13.71 | 14 | Water 10 | | a:8225 | 4.45 | 1.0 | 36.1 |  | 9.85 | 15 | Water 11 | | a:4144 | 4.48 | 1.0 | 38.6 |  | 7.08 |

  
**1^st^ shell coordination angles and distances (*trans* in "blue"; *cys* in "green"; deviations in "orange"):**   
• 1-2(W/W): 91º; 1-3(W/W): 90º; 1-4(W/W): 171º; 1-5(W/W): 92º; 1-6(W/W): 87º; 2-3(W/W): 175º; 2-4(W/W): *96º*; 2-5(W/W): 94º; 2-6(W/W): 91º; 3-4(W/W): *82º*; 3-5(W/W): 90º; 3-6(W/W): *84º*; 4-5(W/W): 92º; 4-6(W/W): 88º; 5-6(W/W): 174º;   
• 1-2(W/W): 2.8Å; 1-3(W/W): 2.9Å; 1-4(W/W): 4.1Å; 1-5(W/W): 3.0Å; 1-6(W/W): 2.9Å; 2-3(W/W): *4.2Å*; 2-4(W/W): 3.2Å; 2-5(W/W): 3.1Å; 2-6(W/W): 3.1Å; 3-4(W/W): 2.9Å; 3-5(W/W): 3.1Å; 3-6(W/W): 3.0Å; 4-5(W/W): 3.2Å; 4-6(W/W): 3.1Å; 5-6(W/W): *4.5Å*;   
  
**2.14 ± 0.12: the average coodination distance of the 1^st^ six ligands is Mg^2+^ compatible**  
**12: Number of potential Water contacts**   
**4 : Number of second shell contacts involving water or ions**   
**33%: Percentage of satisfied second shell Water contacts**   
• Contact(s): /O_w_/O_w_/O_w_/O_w_/O_w_/O_w_/  
• Contact code: 6O_w_  
Ooctahedral coordination (3.6º)   
Go to ToC

### MG:a:3222

|  | Res. type | Atom | Residue | Distance | Occupancy | B-factor | Closest atom(s) to a 1^st^ shell water | EM\_dens. |
| --- | --- | --- | --- | --- | --- | --- | --- | --- |
| **Ion** | | | |  | 1.0 | 35.8 |  | 18.25 |
 1 | Water 1 | | a:6212 | 2.02 | 1.0 | 25.8 | 2.76 [U:OP2]a:746 // 2.89 [SER:OG]d:137 | 14.08 | 2 | Water 2 | | a:4453 | 2.05 | 1.0 | 30.5 | 2.63 [C:OP1]a:2615 | 11.48 | 3 | Water 3 | | a:7548 | 2.14 | 1.0 | 30.6 | 2.85 [1MG:N7]a:747 | 12.08 | 4 | Water 4 | | a:6070 | 2.15 | 1.0 | 27.0 | 2.75 [C:OP1]a:2614 | 10.22 | 5 | PSU | O2 | a:748 | 2.18 | 1.0 | 37.0 |  | 10.67 | 6 | Water 5 | | a:5599 | 2.33 | 1.0 | 25.4 | 2.72 [1MG:O6]a:747 | 12.26 | 7 | Water 6 | | a:9961 | 4.08 | 1.0 | 39.0 |  | 7.67 | 8 | Water 7 | | a:4482 | 4.12 | 1.0 | 31.4 | 2.63 [U:OP2]a:746 | 11.43 | 9 | Water 8 | | a:4303 | 4.14 | 1.0 | 25.0 |  | 11.90 | 10 | C | OP1 | a:2615 | 4.20 | 1.0 | 35.8 |  | 9.30 | 11 | U | OP2 | a:746 | 4.21 | 1.0 | 28.8 |  | 10.79 | 12 | Water 9 | | a:10242 | 4.25 | 1.0 | 35.2 |  | 9.09 | 13 | 1MG | O6 | a:747 | 4.27 | 1.0 | 30.6 |  | 14.68 | 14 | 1MG | N7 | a:747 | 4.33 | 1.0 | 21.9 |  | 12.89 | 15 | C | OP1 | a:2614 | 4.34 | 1.0 | 44.1 |  | 6.56 | 16 | SER | OG | d:137 | 4.43 | 1.0 | 31.4 |  | 12.17 | 17 | Water 10 | | a:6554 | 4.72 | 1.0 | 40.2 |  | 5.64 | 18 | Water 11 | | a:9459 | 4.74 | 1.0 | 42.6 |  | 4.26 |

  
**1^st^ shell coordination angles and distances (*trans* in "blue"; *cys* in "green"; deviations in "orange"):**   
• 1-2(W/W): *97º*; 1-3(W/W): 93º; 1-4(W/W): 93º; 1-5(W/O2): 173º; 2-3(W/W): 94º; 2-4(W/W): 85º; 2-5(W/O2): 89º; 3-4(W/W): 174º; 3-5(W/O2): 86º; 4-5(W/O2): 89º;   
• 1-2(W/W): 3.1Å; 1-3(W/W): 3.0Å; 1-4(W/W): 3.0Å; 1-5(W/O2): 4.2Å; 2-3(W/W): 3.1Å; 2-4(W/W): 2.8Å; 2-5(W/O2): 3.0Å; 3-4(W/W): *4.3Å*; 3-5(W/O2): 2.9Å; 4-5(W/O2): 3.0Å;   
  
**2.14 ± 0.10: the average coodination distance of the 1^st^ six ligands is Mg^2+^ compatible**  
**10: Number of potential Water contacts**   
**6 : Number of second shell contacts involving water or ions**   
**60%: Percentage of satisfied second shell Water contacts**   
• Contact(s): /O2/O_w_/O_w_/O_w_/O_w_/  
• Contact code: O_b_4O_w_  
• Contact(s) in the exclusion zone: /O_w_/  
→ Missing octahedral coordination for MG ←  
Ooctahedral coordination (4.0º)   
Go to ToC

### MG:a:3223

|  | Res. type | Atom | Residue | Distance | Occupancy | B-factor | Closest atom(s) to a 1^st^ shell water | EM\_dens. |
| --- | --- | --- | --- | --- | --- | --- | --- | --- |
| **Ion** | | | |  | 1.0 | 23.4 |  | 25.39 |
 1 | Water 1 | | a:7527 | 1.98 | 1.0 | 24.2 | 2.86 [A:N7]a:753 // 3.00 [A:OP2]a:1616 | 14.78 | 2 | A | OP1 | a:753 | 2.06 | 1.0 | 22.8 |  | 17.08 | 3 | A | OP1 | a:1616 | 2.09 | 1.0 | 27.6 |  | 14.03 | 4 | Water 2 | | a:6551 | 2.14 | 1.0 | 24.2 | 2.78 [A:N1]a:791 // 2.83 [G:O2']a:1615 | 15.41 | 5 | Water 3 | | a:5254 | 2.14 | 1.0 | 21.9 | 2.69 [A:N7]a:1616 | 18.67 | 6 | Water 4 | | a:4541 | 2.42 | 1.0 | 22.6 | 2.64 [C:N3]a:1619 | 15.84 | 7 | A | OP2 | a:1616 | 3.81 | 1.0 | 28.1 |  | 13.63 | 8 | A | O3' | a:752 | 4.05 | 1.0 | 26.8 |  | 13.55 | 9 | Water 5 | | r:219 | 4.06 | 1.0 | 29.6 | 2.81 [A:OP2]a:753 | 12.16 | 10 | A | N7 | a:1616 | 4.07 | 1.0 | 27.6 |  | 13.24 | 11 | G | O3' | a:1615 | 4.17 | 1.0 | 27.7 |  | 15.02 | 12 | Water 6 | | a:8587 | 4.19 | 1.0 | 34.6 | 2.94 [A:O3']a:752 | 9.42 | 13 | A | N7 | a:753 | 4.28 | 1.0 | 22.5 |  | 14.97 | 14 | A | O5' | a:753 | 4.28 | 1.0 | 28.0 |  | 12.06 | 15 | G | O2' | a:1615 | 4.28 | 1.0 | 28.6 |  | 12.73 | 16 | A | N1 | a:791 | 4.40 | 1.0 | 22.8 |  | 12.38 | 17 | C | N3 | a:1619 | 4.44 | 1.0 | 19.5 |  | 16.79 | 18 | A | OP2 | a:753 | 4.46 | 1.0 | 29.2 |  | 13.35 | 19 | A | O5' | a:1616 | 4.49 | 1.0 | 24.9 |  | 15.48 | 20 | Water 7 | | a:8940 | 4.71 | 1.0 | 39.5 |  | 7.24 |

  
**1^st^ shell coordination angles and distances (*trans* in "blue"; *cys* in "green"; deviations in "orange"):**   
• 1-2(W/OP1): *100º*; 1-3(W/OP1): 89º; 1-4(W/W): 91º; 1-5(W/W): 93º; 2-3(OP1/OP1): 171º; 2-4(OP1/W): 94º; 2-5(OP1/W): 86º; 3-4(OP1/W): 88º; 3-5(OP1/W): 91º; 4-5(W/W): 176º;   
• 1-2(W/OP1): 3.1Å; 1-3(W/OP1): 2.8Å; 1-4(W/W): 2.9Å; 1-5(W/W): 3.0Å; 2-3(OP1/OP1): 4.1Å; 2-4(OP1/W): 3.1Å; 2-5(OP1/W): 2.9Å; 3-4(OP1/W): 3.0Å; 3-5(OP1/W): 3.0Å; 4-5(W/W): *4.3Å*;   
  
**2.14 ± 0.14: the average coodination distance of the 1^st^ six ligands is Mg^2+^ compatible**  
**8: Number of potential Water contacts**   
**6 : Number of second shell contacts involving water or ions**   
**75%: Percentage of satisfied second shell Water contacts**   
• Contact(s): /O_w_/O_w_/O_w_/OP1/OP1/  
• Contact code: *trans-*2O_ph_3O_w_  
• Contact(s) in the exclusion zone: /O_w_/  
→ Missing octahedral coordination for MG ←  
Ooctahedral coordination (3.9º)   
Go to ToC

### MG:a:3224

|  | Res. type | Atom | Residue | Distance | Occupancy | B-factor | Closest atom(s) to a 1^st^ shell water | EM\_dens. |
| --- | --- | --- | --- | --- | --- | --- | --- | --- |
| **Ion** | | | |  | 1.0 | 24.7 |  | 27.42 |
 1 | A | OP1 | a:2593 | 1.94 | 1.0 | 21.9 |  | 17.32 | 2 | G | OP2 | a:786 | 2.01 | 1.0 | 20.6 |  | 19.45 | 3 | A | OP2 | a:785 | 2.03 | 1.0 | 20.0 |  | 17.79 | 4 | Water 1 | | a:7901 | 2.18 | 1.0 | 20.9 | 2.89 [ASN:OD1]c:226 | 16.90 | 5 | Water 2 | | c:481 | 2.30 | 1.0 | 22.3 | 2.79 [ALA:O]c:224 | 16.04 | 6 | Water 3 | | a:8778 | 2.33 | 1.0 | 21.6 |  | 19.16 | 7 | A | O5' | a:785 | 3.83 | 1.0 | 23.1 |  | 16.93 | 8 | G | O3' | a:2592 | 3.93 | 1.0 | 22.0 |  | 18.34 | 9 | G | OP1 | a:786 | 3.96 | 1.0 | 22.9 |  | 19.24 | 10 | Water 4 | | a:4703 | 4.03 | 1.0 | 19.2 | 2.65 [A:OP2]a:2593 | 18.20 | 11 | A | O3' | a:784 | 4.06 | 1.0 | 18.9 |  | 18.98 | 12 | A | O3' | a:785 | 4.13 | 1.0 | 18.8 |  | 16.90 | 13 | Water 5 | | a:8187 | 4.22 | 1.0 | 23.4 |  | 14.56 | 14 | A | OP2 | a:2593 | 4.29 | 1.0 | 24.5 |  | 18.24 | 15 | A | O5' | a:2593 | 4.31 | 1.0 | 18.8 |  | 18.19 | 16 | ASN | OD1 | c:226 | 4.34 | 1.0 | 23.8 |  | 19.05 | 17 | ALA | O | c:224 | 4.42 | 1.0 | 27.5 |  | 18.61 | 18 | Water 6 | | a:8298 | 4.43 | 1.0 | 25.7 |  | 14.05 | 19 | G | O5' | a:786 | 4.45 | 1.0 | 22.4 |  | 15.88 | 20 | A | OP1 | a:785 | 4.48 | 1.0 | 25.1 |  | 15.58 | 21 | Water 7 | | c:475 | 4.61 | 1.0 | 27.0 |  | 14.84 |

  
**1^st^ shell coordination angles and distances (*trans* in "blue"; *cys* in "green"; deviations in "orange"):**   
• 1-2(OP1/OP2): *98º*; 1-3(OP1/OP2): *99º*; 1-4(OP1/W): 93º; 1-5(OP1/W): *168º*; 2-3(OP2/OP2): *107º*; 2-4(OP2/W): 87º; 2-5(OP2/W): 90º; 3-4(OP2/W): *160º*; 3-5(OP2/W): 87º; 4-5(W/W): *79º*;   
• 1-2(OP1/OP2): 3.0Å; 1-3(OP1/OP2): 3.0Å; 1-4(OP1/W): 3.0Å; 1-5(OP1/W): *4.2Å*; 2-3(OP2/OP2): *3.2Å*; 2-4(OP2/W): 2.9Å; 2-5(OP2/W): 3.1Å; 3-4(OP2/W): 4.1Å; 3-5(OP2/W): 3.0Å; 4-5(W/W): 2.8Å;   
  
**2.13 ± 0.15: the average coodination distance of the 1^st^ six ligands is Mg^2+^ compatible**  
**6: Number of potential Water contacts**   
**2 : Number of second shell contacts involving water or ions**   
**33%: Percentage of satisfied second shell Water contacts**   
• Contact(s): /O_w_/O_w_/OP1/OP2/OP2/  
• Contact code: *fac-*3O_ph_2O_w_  
• Contact(s) in the exclusion zone: /O_w_/  
→ Missing octahedral coordination for MG ←  
→ Slightly distorded coord. octahedron (8.7º) ←  
Go to ToC

### MG:a:3225

|  | Res. type | Atom | Residue | Distance | Occupancy | B-factor | Closest atom(s) to a 1^st^ shell water | EM\_dens. |
| --- | --- | --- | --- | --- | --- | --- | --- | --- |
| **Ion** | | | |  | 1.0 | 26.7 |  | 24.33 |
 1 | G | OP1 | a:786 | 2.00 | 1.0 | 22.9 |  | 19.24 | 2 | G | OP1 | a:2592 | 2.01 | 1.0 | 22.1 |  | 15.37 | 3 | Water 1 | | a:6700 | 2.08 | 1.0 | 21.1 |  | 16.19 | 4 | Water 2 | | a:4703 | 2.09 | 1.0 | 19.2 | 2.65 [A:OP2]a:2593 // 2.90 [K:K]c:303 | 18.20 | 5 | Water 3 | | c:439 | 2.21 | 1.0 | 24.6 | 2.65 [ASN:OD1]c:226 | 17.61 | 6 | Water 4 | | a:3538 | 2.37 | 1.0 | 25.5 | 2.46 [C:OP2]a:2444 | 14.16 | 7 | Water 5 | | c:478 | 3.76 | 1.0 | 26.2 | 2.94 [G:OP2]a:2592 | 13.14 | 8 | A | OP2 | a:2593 | 3.93 | 1.0 | 24.5 |  | 18.24 | 9 | G | O5' | a:786 | 4.10 | 1.0 | 22.4 |  | 15.88 | 10 | A | O3' | a:2591 | 4.18 | 1.0 | 28.5 |  | 14.17 | 11 | Water 6 | | a:7901 | 4.21 | 1.0 | 20.9 | 2.87 [G:OP2]a:786 // 2.89 [ASN:OD1]c:226 | 16.90 | 12 | G | OP2 | a:786 | 4.24 | 1.0 | 20.6 |  | 19.45 | 13 | G | O5' | a:2592 | 4.25 | 1.0 | 23.3 |  | 16.72 | 14 | Water 7 | | c:426 | 4.35 | 1.0 | 30.9 | 2.95 [K:K]c:303 | 11.97 | 15 | A | O3' | a:785 | 4.40 | 1.0 | 18.8 |  | 16.90 | 16 | G | OP2 | a:2592 | 4.40 | 1.0 | 22.7 |  | 13.56 | 17 | Water 8 | | a:3439 | 4.46 | 1.0 | 28.6 |  | 12.14 | 18 | ASN | OD1 | c:226 | 4.53 | 1.0 | 23.8 |  | 19.05 | 19 | Water 9 | | a:5237 | 4.58 | 1.0 | 28.7 |  | 11.78 | 20 | C | OP2 | a:2444 | 4.62 | 1.0 | 32.2 |  | 9.86 | 21 | A | OP1 | a:2593 | 4.71 | 1.0 | 21.9 |  | 17.32 | 22 | Water 10 | | a:7018 | 4.76 | 1.0 | 31.3 |  | 11.15 | 23 | **K** | K | c:303 | 4.79 | 1.0 | 30.0 |  | 27.12 |

  
**1^st^ shell coordination angles and distances (*trans* in "blue"; *cys* in "green"; deviations in "orange"):**   
• 1-2(OP1/OP1): *100º*; 1-3(OP1/W): 93º; 1-4(OP1/W): 88º; 1-5(OP1/W): *83º*; 2-3(OP1/W): 86º; 2-4(OP1/W): *102º*; 2-5(OP1/W): *169º*; 3-4(W/W): 172º; 3-5(W/W): *83º*; 4-5(W/W): 89º;   
• 1-2(OP1/OP1): 3.1Å; 1-3(OP1/W): 3.0Å; 1-4(OP1/W): 2.8Å; 1-5(OP1/W): 2.8Å; 2-3(OP1/W): 2.8Å; 2-4(OP1/W): 3.2Å; 2-5(OP1/W): 4.2Å; 3-4(W/W): 4.2Å; 3-5(W/W): 2.8Å; 4-5(W/W): 3.0Å;   
  
**2.13 ± 0.13: the average coodination distance of the 1^st^ six ligands is Mg^2+^ compatible**  
**8: Number of potential Water contacts**   
**4 : Number of second shell contacts involving water or ions**   
**50%: Percentage of satisfied second shell Water contacts**   
• Contact(s): /O_w_/O_w_/O_w_/OP1/OP1/  
• Contact code: *cis-*2O_ph_3O_w_  
• Contact(s) in the exclusion zone: /O_w_/  
→ Missing octahedral coordination for MG ←  
→ Slightly distorded coord. octahedron (6.5º) ←  
Go to ToC

### MG:a:3226

|  | Res. type | Atom | Residue | Distance | Occupancy | B-factor | Closest atom(s) to a 1^st^ shell water | EM\_dens. |
| --- | --- | --- | --- | --- | --- | --- | --- | --- |
| **Ion** | | | |  | 1.0 | 29.3 |  | 21.27 |
 1 | A | OP1 | a:802 | 1.93 | 1.0 | 26.1 |  | 12.42 | 2 | Water 1 | | a:6917 | 2.00 | 1.0 | 27.7 |  | 12.44 | 3 | Water 2 | | a:9456 | 2.10 | 1.0 | 26.2 |  | 14.53 | 4 | Water 3 | | a:6208 | 2.12 | 1.0 | 23.1 | 2.76 [G:O2']a:801 | 16.29 | 5 | Water 4 | | a:3508 | 2.29 | 1.0 | 24.5 | 2.45 [C:O2]a:192 // 2.82 [A:N7]a:802 | 15.98 | 6 | Water 5 | | a:5418 | 2.42 | 1.0 | 31.5 | 2.70 [C:N3]a:192 | 13.04 | 7 | A | OP2 | a:802 | 3.91 | 1.0 | 28.5 |  | 13.14 | 8 | G | O2' | a:801 | 4.07 | 1.0 | 26.2 |  | 14.73 | 9 | G | O3' | a:801 | 4.11 | 1.0 | 24.3 |  | 14.20 | 10 | Water 6 | | a:4923 | 4.29 | 1.0 | 43.4 | 2.67 [A:OP2]a:802 | 7.27 | 11 | A | N7 | a:802 | 4.32 | 1.0 | 20.0 |  | 18.53 | 12 | A | O5' | a:802 | 4.38 | 1.0 | 33.2 |  | 12.09 | 13 | C | O2 | a:192 | 4.41 | 1.0 | 26.5 |  | 15.38 | 14 | C | N3 | a:192 | 4.44 | 1.0 | 17.2 |  | 16.96 | 15 | Water 7 | | a:3942 | 4.45 | 1.0 | 29.4 |  | 12.58 | 16 | Water 8 | | a:8438 | 4.50 | 1.0 | 35.0 |  | 9.68 | 17 | A | N1 | a:191 | 4.55 | 1.0 | 15.4 |  | 17.93 | 18 | Water 9 | | a:9214 | 4.56 | 1.0 | 30.6 |  | 10.21 | 19 | Water 10 | | a:6083 | 4.63 | 1.0 | 43.7 |  | 4.46 |

  
**1^st^ shell coordination angles and distances (*trans* in "blue"; *cys* in "green"; deviations in "orange"):**   
• 1-2(OP1/W): 91º; 1-3(OP1/W): *100º*; 1-4(OP1/W): *95º*; 1-5(OP1/W): *100º*; 2-3(W/W): 86º; 2-4(W/W): 173º; 2-5(W/W): *99º*; 3-4(W/W): 90º; 3-5(W/W): *159º*; 4-5(W/W): *83º*;   
• 1-2(OP1/W): 2.8Å; 1-3(OP1/W): 3.1Å; 1-4(OP1/W): 3.0Å; 1-5(OP1/W): *3.2Å*; 2-3(W/W): 2.8Å; 2-4(W/W): 4.1Å; 2-5(W/W): *3.3Å*; 3-4(W/W): 3.0Å; 3-5(W/W): *4.3Å*; 4-5(W/W): 2.9Å;   
  
**2.14 ± 0.17: the average coodination distance of the 1^st^ six ligands is Mg^2+^ compatible**  
**10: Number of potential Water contacts**   
**4 : Number of second shell contacts involving water or ions**   
**40%: Percentage of satisfied second shell Water contacts**   
• Contact(s): /O_w_/O_w_/O_w_/O_w_/OP1/  
• Contact code: O_ph_4O_w_  
• Contact(s) in the exclusion zone: /O_w_/  
→ Missing octahedral coordination for MG ←  
→ Slightly distorded coord. octahedron (7.3º) ←  
Go to ToC

### MG:a:3227

|  | Res. type | Atom | Residue | Distance | Occupancy | B-factor | Closest atom(s) to a 1^st^ shell water | EM\_dens. |
| --- | --- | --- | --- | --- | --- | --- | --- | --- |
| **Ion** | | | |  | 1.0 | 30.3 |  | 22.17 |
 1 | Water 1 | | a:4438 | 2.09 | 1.0 | 24.9 | 2.63 [A:OP2]a:677 | 14.80 | 2 | Water 2 | | a:8745 | 2.10 | 1.0 | 29.1 |  | 14.47 | 3 | Water 3 | | a:8409 | 2.17 | 1.0 | 27.9 | 2.99 [A:OP2]a:804 | 13.65 | 4 | A | OP1 | a:804 | 2.19 | 1.0 | 28.6 |  | 13.30 | 5 | Water 4 | | a:3660 | 2.25 | 1.0 | 25.0 | 2.50 [U:O4]a:805 | 16.36 | 6 | Water 5 | | a:8975 | 2.34 | 1.0 | 24.6 |  | 13.59 | 7 | A | OP2 | a:804 | 3.92 | 1.0 | 33.4 |  | 13.33 | 8 | A | OP2 | a:677 | 3.94 | 1.0 | 28.3 |  | 13.44 | 9 | G | O3' | a:803 | 4.13 | 1.0 | 27.9 |  | 14.72 | 10 | Water 6 | | a:4039 | 4.14 | 1.0 | 26.9 |  | 11.75 | 11 | G | OP2 | a:676 | 4.16 | 1.0 | 23.9 |  | 18.16 | 12 | Water 7 | | a:6034 | 4.20 | 1.0 | 27.4 | 2.74 [A:OP2]a:677 | 12.16 | 13 | U | O4 | a:805 | 4.20 | 1.0 | 23.2 |  | 16.56 | 14 | Water 8 | | a:3940 | 4.24 | 1.0 | 25.1 |  | 15.31 | 15 | Water 9 | | a:7639 | 4.24 | 1.0 | 28.4 | 2.85 [G:OP2]a:676 | 14.79 | 16 | A | O5' | a:804 | 4.56 | 1.0 | 31.5 |  | 10.52 | 17 | Water 10 | | a:9894 | 4.57 | 1.0 | 40.4 |  | 6.29 | 18 | Water 11 | | a:6908 | 4.76 | 1.0 | 36.5 |  | 8.77 |

  
**1^st^ shell coordination angles and distances (*trans* in "blue"; *cys* in "green"; deviations in "orange"):**   
• 1-2(W/W): 93º; 1-3(W/W): 88º; 1-4(W/OP1): 173º; 1-5(W/W): 91º; 2-3(W/W): *96º*; 2-4(W/OP1): 87º; 2-5(W/W): 174º; 3-4(W/OP1): *84º*; 3-5(W/W): 88º; 4-5(OP1/W): 89º;   
• 1-2(W/W): 3.0Å; 1-3(W/W): 3.0Å; 1-4(W/OP1): *4.3Å*; 1-5(W/W): 3.1Å; 2-3(W/W): 3.2Å; 2-4(W/OP1): 3.0Å; 2-5(W/W): *4.3Å*; 3-4(W/OP1): 2.9Å; 3-5(W/W): 3.1Å; 4-5(OP1/W): 3.1Å;   
  
**2.19 ± 0.09: the average coodination distance of the 1^st^ six ligands is Mg^2+^ compatible**  
**10: Number of potential Water contacts**   
**3 : Number of second shell contacts involving water or ions**   
**30%: Percentage of satisfied second shell Water contacts**   
• Contact(s): /O_w_/O_w_/O_w_/O_w_/OP1/  
• Contact code: O_ph_4O_w_  
• Contact(s) in the exclusion zone: /O_w_/  
→ Missing octahedral coordination for MG ←  
Ooctahedral coordination (3.5º)   
Go to ToC

### MG:a:3228

|  | Res. type | Atom | Residue | Distance | Occupancy | B-factor | Closest atom(s) to a 1^st^ shell water | EM\_dens. |
| --- | --- | --- | --- | --- | --- | --- | --- | --- |
| **Ion** | | | |  | 1.0 | 21.8 |  | 28.19 |
 1 | U | OP1 | a:812 | 2.01 | 1.0 | 20.5 |  | 16.77 | 2 | Water 1 | | a:4514 | 2.04 | 1.0 | 23.3 | 2.63 [U:OP1]a:813 | 18.18 | 3 | Water 2 | | a:9406 | 2.17 | 1.0 | 21.3 |  | 17.15 | 4 | Water 3 | | a:7313 | 2.27 | 1.0 | 20.9 |  | 19.28 | 5 | Water 4 | | a:8615 | 2.29 | 1.0 | 28.9 | 2.95 [A:N6]a:1255 | 14.87 | 6 | A | N7 | a:1255 | 2.47 | 1.0 | 15.2 |  | 18.18 | 7 | Water 5 | | a:5984 | 3.73 | 1.0 | 28.3 | 2.86 [U:OP2]a:812 | 14.05 | 8 | U | OP2 | a:812 | 3.89 | 1.0 | 25.3 |  | 17.54 | 9 | U | O5' | a:812 | 4.01 | 1.0 | 22.3 |  | 18.74 | 10 | U | OP1 | a:813 | 4.27 | 1.0 | 22.6 |  | 15.58 | 11 | Water 6 | | a:10167 | 4.28 | 1.0 | 30.6 |  | 9.75 | 12 | Water 7 | | a:5243 | 4.29 | 1.0 | 22.5 | 2.95 [G:O2']a:1254 | 15.97 | 13 | Water 8 | | a:8256 | 4.46 | 1.0 | 34.8 |  | 10.77 | 14 | G | O3' | a:811 | 4.50 | 1.0 | 20.0 |  | 18.33 | 15 | Water 9 | | a:10128 | 4.54 | 1.0 | 34.4 |  | 8.52 | 16 | G | O2' | a:1254 | 4.77 | 1.0 | 24.6 |  | 13.66 | 17 | C | O2' | a:567 | 4.78 | 1.0 | 22.0 |  | 19.46 |

  
**1^st^ shell coordination angles and distances (*trans* in "blue"; *cys* in "green"; deviations in "orange"):**   
• 1-2(OP1/W): *96º*; 1-3(OP1/W): 95º; 1-4(OP1/W): *95º*; 1-5(OP1/W): 177º; 2-3(W/W): *107º*; 2-4(W/W): *83º*; 2-5(W/W): 85º; 3-4(W/W): *164º*; 3-5(W/W): 87º; 4-5(W/W): *83º*;   
• 1-2(OP1/W): 3.0Å; 1-3(OP1/W): 3.1Å; 1-4(OP1/W): 3.2Å; 1-5(OP1/W): *4.3Å*; 2-3(W/W): *3.4Å*; 2-4(W/W): 2.9Å; 2-5(W/W): 2.9Å; 3-4(W/W): *4.4Å*; 3-5(W/W): 3.1Å; 4-5(W/W): 3.0Å;   
  
**2.21 ± 0.16: the average coodination distance of the 1^st^ six ligands is Mg^2+^ compatible**  
**8: Number of potential Water contacts**   
**2 : Number of second shell contacts involving water or ions**   
**25%: Percentage of satisfied second shell Water contacts**   
• Contact(s): /O_w_/O_w_/O_w_/O_w_/OP1/  
• Contact code: O_ph_4O_w_  
• Contact(s) in the exclusion zone: /N7/  
→ Missing octahedral coordination for MG ←  
→ Slightly distorded coord. octahedron (7.4º) ←  
Go to ToC

### MG:a:3229

|  | Res. type | Atom | Residue | Distance | Occupancy | B-factor | Closest atom(s) to a 1^st^ shell water | EM\_dens. |
| --- | --- | --- | --- | --- | --- | --- | --- | --- |
| **Ion** | | | |  | 1.0 | 37.4 |  | 19.61 |
 1 | Water 1 | | a:6192 | 2.06 | 1.0 | 27.9 | 2.75 [G:OP2]a:824 | 12.47 | 2 | Water 2 | | a:10260 | 2.09 | 1.0 | 31.2 |  | 9.95 | 3 | Water 3 | | a:5241 | 2.15 | 1.0 | 26.3 | 2.69 [A:OP2]a:949 | 12.25 | 4 | Water 4 | | a:10010 | 2.15 | 1.0 | 33.6 |  | 9.30 | 5 | Water 5 | | a:3735 | 2.18 | 1.0 | 29.2 | 2.52 [G:OP1]a:824 | 13.26 | 6 | Water 6 | | a:4475 | 2.18 | 1.0 | 28.2 | 2.63 [C:OP2]a:967 | 13.85 | 7 | G | OP1 | a:824 | 4.02 | 1.0 | 32.1 |  | 13.52 | 8 | G | OP2 | a:824 | 4.08 | 1.0 | 25.2 |  | 12.88 | 9 | Water 7 | | a:4295 | 4.17 | 1.0 | 29.1 |  | 12.34 | 10 | Water 8 | | a:8170 | 4.23 | 1.0 | 36.7 |  | 9.26 | 11 | Water 9 | | a:6415 | 4.29 | 1.0 | 36.9 |  | 7.53 | 12 | Water 10 | | a:8214 | 4.31 | 1.0 | 35.0 |  | 9.97 | 13 | Water 11 | | a:10244 | 4.37 | 1.0 | 44.0 |  | 4.96 | 14 | A | OP2 | a:949 | 4.38 | 1.0 | 22.7 |  | 16.90 | 15 | C | OP2 | a:967 | 4.42 | 1.0 | 27.3 |  | 13.88 | 16 | Water 12 | | a:5439 | 4.43 | 1.0 | 30.6 |  | 10.95 | 17 | Water 13 | | a:7100 | 4.49 | 1.0 | 40.8 |  | 8.19 | 18 | Water 14 | | a:5072 | 4.51 | 1.0 | 40.4 | 2.68 [C:OP2]a:967 | 6.25 | 19 | Water 15 | | a:9915 | 4.60 | 1.0 | 44.0 |  | 5.90 | 20 | Water 16 | | a:5090 | 4.67 | 1.0 | 32.0 | 2.68 [G:OP1]a:824 | 9.19 |

  
**1^st^ shell coordination angles and distances (*trans* in "blue"; *cys* in "green"; deviations in "orange"):**   
• 1-2(W/W): 173º; 1-3(W/W): 88º; 1-4(W/W): *104º*; 1-5(W/W): 93º; 1-6(W/W): 88º; 2-3(W/W): *84º*; 2-4(W/W): *84º*; 2-5(W/W): 86º; 2-6(W/W): 94º; 3-4(W/W): *167º*; 3-5(W/W): 90º; 3-6(W/W): *98º*; 4-5(W/W): 85º; 4-6(W/W): 87º; 5-6(W/W): 172º;   
• 1-2(W/W): 4.1Å; 1-3(W/W): 2.9Å; 1-4(W/W): *3.3Å*; 1-5(W/W): 3.1Å; 1-6(W/W): 2.9Å; 2-3(W/W): 2.8Å; 2-4(W/W): 2.8Å; 2-5(W/W): 2.9Å; 2-6(W/W): 3.1Å; 3-4(W/W): *4.3Å*; 3-5(W/W): 3.1Å; 3-6(W/W): *3.3Å*; 4-5(W/W): 2.9Å; 4-6(W/W): 3.0Å; 5-6(W/W): *4.4Å*;   
  
**2.14 ± 0.05: the average coodination distance of the 1^st^ six ligands is Mg^2+^ compatible**  
**12: Number of potential Water contacts**   
**4 : Number of second shell contacts involving water or ions**   
**33%: Percentage of satisfied second shell Water contacts**   
• Contact(s): /O_w_/O_w_/O_w_/O_w_/O_w_/O_w_/  
• Contact code: 6O_w_  
→ Slightly distorded coord. octahedron (5.6º) ←  
Go to ToC

### MG:a:3230

|  | Res. type | Atom | Residue | Distance | Occupancy | B-factor | Closest atom(s) to a 1^st^ shell water | EM\_dens. |
| --- | --- | --- | --- | --- | --- | --- | --- | --- |
| **Ion** | | | |  | 1.0 | 28.0 |  | 19.09 |
 1 | U | OP1 | a:829 | 1.99 | 1.0 | 26.1 |  | 13.52 | 2 | G | OP2 | a:2433 | 2.06 | 1.0 | 24.6 |  | 11.90 | 3 | Water 1 | | a:4253 | 2.15 | 1.0 | 27.2 | 2.61 [U:O3']a:828 | 13.08 | 4 | Water 2 | | a:7520 | 2.36 | 1.0 | 32.3 | 2.90 [A:OP1]a:2434 | 10.85 | 5 | Water 3 | | a:6311 | 2.39 | 1.0 | 30.0 | 2.76 [A:OP1]a:2434 | 11.36 | 6 | A | OP2 | a:2434 | 2.56 | 1.0 | 35.8 |  | 6.80 | 7 | A | OP1 | a:2434 | 3.12 | 1.0 | 45.3 |  | 5.41 | 8 | U | O3' | a:828 | 3.42 | 1.0 | 27.4 |  | 12.92 | 9 | U | OP2 | a:829 | 4.05 | 1.0 | 25.2 |  | 14.42 | 10 | **K** | K | a:3119 | 4.11 | 1.0 | 30.0 |  | 25.79 | 11 | G | O3' | a:2433 | 4.14 | 1.0 | 35.6 |  | 9.62 | 12 | G | O5' | a:2433 | 4.15 | 1.0 | 23.0 |  | 13.75 | 13 | Water 4 | | a:4954 | 4.22 | 1.0 | 39.3 |  | 10.03 | 14 | Water 5 | | a:10097 | 4.27 | 1.0 | 40.2 |  | 5.86 | 15 | G | O3' | a:2432 | 4.32 | 1.0 | 24.7 |  | 14.24 | 16 | Water 6 | | a:9068 | 4.36 | 1.0 | 33.3 |  | 9.58 | 17 | U | O5' | a:829 | 4.41 | 1.0 | 27.4 |  | 11.45 | 18 | G | OP1 | a:2433 | 4.45 | 1.0 | 25.2 |  | 12.03 | 19 | A | O5' | a:2434 | 4.64 | 1.0 | 36.5 |  | 1.65 | 20 | Water 7 | | k:228 | 4.68 | 1.0 | 41.8 |  | 4.11 |

  
**1^st^ shell coordination angles and distances (*trans* in "blue"; *cys* in "green"; deviations in "orange"):**   
• 1-2(OP1/OP2): *97º*; 1-3(OP1/W): 87º; 2-3(OP2/W): 90º;   
• 1-2(OP1/OP2): 3.0Å; 1-3(OP1/W): 2.9Å; 2-3(OP2/W): 3.0Å;   
  
**2.25 ± 0.20: the average coodination distance of the 1^st^ six ligands is Mg^2+^ compatible**  
**6: Number of potential Water contacts**   
**3 : Number of second shell contacts involving water or ions**   
**50%: Percentage of satisfied second shell Water contacts**   
• Contact(s): /O_w_/OP1/OP2/  
• Contact code: *cis-*2O_ph_O_w_  
• Contact(s) in the exclusion zone: /O_w_/O_w_/OP1/OP2/  
→ Missing octahedral coordination for MG ←  
Go to ToC

### MG:a:3231

|  | Res. type | Atom | Residue | Distance | Occupancy | B-factor | Closest atom(s) to a 1^st^ shell water | EM\_dens. |
| --- | --- | --- | --- | --- | --- | --- | --- | --- |
| **Ion** | | | |  | 1.0 | 30.4 |  | 23.88 |
 1 | C | OP2 | a:948 | 1.98 | 1.0 | 29.6 |  | 11.96 | 2 | Water 1 | | a:6926 | 2.05 | 1.0 | 25.8 | 2.80 [G:OP2]a:824 | 15.31 | 3 | Water 2 | | a:8897 | 2.07 | 1.0 | 25.4 |  | 14.49 | 4 | Water 3 | | a:5406 | 2.15 | 1.0 | 25.2 | 2.70 [C:O3']a:946 | 14.02 | 5 | A | OP2 | a:947 | 2.21 | 1.0 | 24.9 |  | 15.58 | 6 | Water 4 | | a:8960 | 2.33 | 1.0 | 23.8 |  | 13.33 | 7 | C | O3' | a:946 | 3.51 | 1.0 | 24.5 |  | 15.37 | 8 | Water 5 | | a:6999 | 3.95 | 1.0 | 37.0 | 2.81 [C:OP1]a:948 | 7.53 | 9 | C | OP1 | a:948 | 4.00 | 1.0 | 25.2 |  | 14.87 | 10 | C | O5' | a:948 | 4.11 | 1.0 | 27.1 |  | 14.14 | 11 | Water 6 | | a:6423 | 4.12 | 1.0 | 31.0 |  | 10.21 | 12 | G | OP2 | a:824 | 4.23 | 1.0 | 25.2 |  | 12.88 | 13 | Water 7 | | a:6371 | 4.23 | 1.0 | 32.6 |  | 9.04 | 14 | A | O5' | a:947 | 4.35 | 1.0 | 28.1 |  | 12.13 | 15 | C | O2' | a:946 | 4.36 | 1.0 | 30.5 |  | 11.53 | 16 | Water 8 | | a:10281 | 4.44 | 1.0 | 40.9 |  | 3.33 | 17 | Water 9 | | a:8020 | 4.45 | 1.0 | 30.8 | 2.90 [C:O2']a:946 | 11.84 | 18 | A | O3' | a:947 | 4.46 | 1.0 | 26.5 |  | 12.60 | 19 | Water 10 | | a:8489 | 4.46 | 1.0 | 35.8 | 2.93 [C:OP1]a:948 | 8.07 | 20 | A | O3' | a:823 | 4.50 | 1.0 | 26.5 |  | 13.86 | 21 | A | OP1 | a:947 | 4.50 | 1.0 | 24.6 |  | 12.82 | 22 | A | O2' | a:823 | 4.52 | 1.0 | 26.1 |  | 15.57 | 23 | Water 11 | | a:8436 | 4.54 | 1.0 | 22.2 | 2.92 [A:OP1]a:947 | 14.61 |

  
**1^st^ shell coordination angles and distances (*trans* in "blue"; *cys* in "green"; deviations in "orange"):**   
• 1-2(OP2/W): 90º; 1-3(OP2/W): 175º; 1-4(OP2/W): 94º; 1-5(OP2/OP2): 87º; 2-3(W/W): 95º; 2-4(W/W): 91º; 2-5(W/OP2): 174º; 3-4(W/W): 85º; 3-5(W/OP2): 88º; 4-5(W/OP2): 94º;   
• 1-2(OP2/W): 2.9Å; 1-3(OP2/W): 4.0Å; 1-4(OP2/W): 3.0Å; 1-5(OP2/OP2): 2.9Å; 2-3(W/W): 3.0Å; 2-4(W/W): 3.0Å; 2-5(W/OP2): *4.3Å*; 3-4(W/W): 2.9Å; 3-5(W/OP2): 3.0Å; 4-5(W/OP2): *3.2Å*;   
  
**2.13 ± 0.12: the average coodination distance of the 1^st^ six ligands is Mg^2+^ compatible**  
**8: Number of potential Water contacts**   
**2 : Number of second shell contacts involving water or ions**   
**25%: Percentage of satisfied second shell Water contacts**   
• Contact(s): /O_w_/O_w_/O_w_/OP2/OP2/  
• Contact code: *cis-*2O_ph_3O_w_  
• Contact(s) in the exclusion zone: /O_w_/  
→ Missing octahedral coordination for MG ←  
Ooctahedral coordination (3.6º)   
Go to ToC

### MG:a:3232

|  | Res. type | Atom | Residue | Distance | Occupancy | B-factor | Closest atom(s) to a 1^st^ shell water | EM\_dens. |
| --- | --- | --- | --- | --- | --- | --- | --- | --- |
| **Ion** | | | |  | 1.0 | 27.2 |  | 24.10 |
 1 | Water 1 | | a:5369 | 2.02 | 1.0 | 21.8 | 2.70 [U:O4]a:834 | 15.72 | 2 | Water 2 | | a:7253 | 2.03 | 1.0 | 20.3 | 2.82 [G:OP2]a:833 | 16.83 | 3 | A | OP1 | a:947 | 2.06 | 1.0 | 24.6 |  | 12.82 | 4 | Water 3 | | a:7379 | 2.18 | 1.0 | 21.8 | 2.83 [G:O6]a:833 | 17.43 | 5 | Water 4 | | a:9134 | 2.19 | 1.0 | 24.7 |  | 15.04 | 6 | Water 5 | | a:8436 | 2.22 | 1.0 | 22.2 |  | 14.61 | 7 | A | OP2 | a:947 | 4.00 | 1.0 | 24.9 |  | 15.58 | 8 | Water 6 | | a:5251 | 4.07 | 1.0 | 41.6 |  | 5.35 | 9 | Water 7 | | a:9307 | 4.08 | 1.0 | 29.8 |  | 12.50 | 10 | U | O4 | a:834 | 4.10 | 1.0 | 24.7 |  | 16.61 | 11 | G | N7 | a:833 | 4.25 | 1.0 | 16.4 |  | 20.13 | 12 | C | O3' | a:946 | 4.28 | 1.0 | 24.5 |  | 15.37 | 13 | G | OP2 | a:833 | 4.37 | 1.0 | 21.1 |  | 14.12 | 14 | A | O5' | a:947 | 4.46 | 1.0 | 28.1 |  | 12.13 | 15 | Water 8 | | a:8995 | 4.51 | 1.0 | 42.9 |  | 5.78 | 16 | Water 9 | | a:6371 | 4.53 | 1.0 | 32.6 | 2.95 [A:OP2]a:947 | 9.04 | 17 | Water 10 | | a:7222 | 4.61 | 1.0 | 30.8 |  | 12.99 | 18 | C | O2' | a:946 | 4.76 | 1.0 | 30.5 |  | 11.53 | 19 | G | O6 | a:833 | 4.76 | 1.0 | 24.7 |  | 16.56 |

  
**1^st^ shell coordination angles and distances (*trans* in "blue"; *cys* in "green"; deviations in "orange"):**   
• 1-2(W/W): *97º*; 1-3(W/OP1): 93º; 1-4(W/W): 89º; 1-5(W/W): 173º; 1-6(W/W): 91º; 2-3(W/OP1): *96º*; 2-4(W/W): 91º; 2-5(W/W): 89º; 2-6(W/W): 172º; 3-4(OP1/W): 173º; 3-5(OP1/W): 90º; 3-6(OP1/W): 86º; 4-5(W/W): 88º; 4-6(W/W): 87º; 5-6(W/W): *83º*;   
• 1-2(W/W): 3.0Å; 1-3(W/OP1): 3.0Å; 1-4(W/W): 2.9Å; 1-5(W/W): *4.2Å*; 1-6(W/W): 3.0Å; 2-3(W/OP1): 3.1Å; 2-4(W/W): 3.0Å; 2-5(W/W): 3.0Å; 2-6(W/W): *4.2Å*; 3-4(OP1/W): *4.2Å*; 3-5(OP1/W): 3.0Å; 3-6(OP1/W): 2.9Å; 4-5(W/W): 3.0Å; 4-6(W/W): 3.0Å; 5-6(W/W): 2.9Å;   
  
**2.12 ± 0.08: the average coodination distance of the 1^st^ six ligands is Mg^2+^ compatible**  
**10: Number of potential Water contacts**   
**3 : Number of second shell contacts involving water or ions**   
**30%: Percentage of satisfied second shell Water contacts**   
• Contact(s): /O_w_/O_w_/O_w_/O_w_/O_w_/OP1/  
• Contact code: O_ph_5O_w_  
Ooctahedral coordination (4.0º)   
Go to ToC

### MG:a:3233

|  | Res. type | Atom | Residue | Distance | Occupancy | B-factor | Closest atom(s) to a 1^st^ shell water | EM\_dens. |
| --- | --- | --- | --- | --- | --- | --- | --- | --- |
| **Ion** | | | |  | 1.0 | 26.0 |  | 21.96 |
 1 | C | OP1 | a:950 | 2.00 | 1.0 | 26.3 |  | 12.88 | 2 | U | OP2 | a:965 | 2.02 | 1.0 | 22.0 |  | 16.00 | 3 | Water 1 | | a:7633 | 2.03 | 1.0 | 27.1 |  | 10.54 | 4 | Water 2 | | a:6618 | 2.10 | 1.0 | 24.4 |  | 12.21 | 5 | G | OP1 | a:964 | 2.12 | 1.0 | 29.2 |  | 13.83 | 6 | Water 3 | | a:6635 | 2.19 | 1.0 | 28.2 | 2.78 [C:OP2]a:950 | 14.75 | 7 | C | OP2 | a:950 | 3.50 | 1.0 | 23.2 |  | 14.45 | 8 | G | OP2 | a:964 | 3.83 | 1.0 | 26.0 |  | 12.63 | 9 | Water 4 | | a:7918 | 3.86 | 1.0 | 34.1 |  | 9.53 | 10 | U | OP1 | a:965 | 4.00 | 1.0 | 19.9 |  | 14.43 | 11 | G | O3' | a:964 | 4.05 | 1.0 | 23.8 |  | 17.93 | 12 | Water 5 | | a:7811 | 4.09 | 1.0 | 34.9 |  | 9.30 | 13 | A | O3' | a:949 | 4.12 | 1.0 | 24.0 |  | 13.35 | 14 | C | O5' | a:950 | 4.31 | 1.0 | 26.1 |  | 14.96 | 15 | G | O5' | a:964 | 4.37 | 1.0 | 26.9 |  | 14.90 | 16 | C | O3' | a:963 | 4.38 | 1.0 | 31.8 |  | 10.07 | 17 | U | O5' | a:965 | 4.47 | 1.0 | 25.0 |  | 14.53 | 18 | Water 6 | | a:9802 | 4.52 | 1.0 | 30.0 |  | 13.00 | 19 | Water 7 | | a:4158 | 4.66 | 1.0 | 42.5 | 2.60 [G:OP2]a:964 | 5.87 | 20 | Water 8 | | a:9987 | 4.68 | 1.0 | 30.5 |  | 11.86 | 21 | Water 9 | | a:5489 | 4.73 | 1.0 | 35.6 | 2.71 [C:OP2]a:950 | 6.43 |

  
**1^st^ shell coordination angles and distances (*trans* in "blue"; *cys* in "green"; deviations in "orange"):**   
• 1-2(OP1/OP2): 173º; 1-3(OP1/W): *97º*; 1-4(OP1/W): 86º; 1-5(OP1/OP1): 89º; 1-6(OP1/W): 93º; 2-3(OP2/W): 90º; 2-4(OP2/W): 88º; 2-5(OP2/OP1): 88º; 2-6(OP2/W): 90º; 3-4(W/W): 177º; 3-5(W/OP1): 90º; 3-6(W/W): 87º; 4-5(W/OP1): 89º; 4-6(W/W): 95º; 5-6(OP1/W): 176º;   
• 1-2(OP1/OP2): 4.0Å; 1-3(OP1/W): 3.0Å; 1-4(OP1/W): 2.8Å; 1-5(OP1/OP1): 2.9Å; 1-6(OP1/W): 3.0Å; 2-3(OP2/W): 2.9Å; 2-4(OP2/W): 2.8Å; 2-5(OP2/OP1): 2.9Å; 2-6(OP2/W): 3.0Å; 3-4(W/W): 4.1Å; 3-5(W/OP1): 2.9Å; 3-6(W/W): 2.9Å; 4-5(W/OP1): 3.0Å; 4-6(W/W): 3.1Å; 5-6(OP1/W): *4.3Å*;   
  
**2.07 ± 0.07: the average coodination distance of the 1^st^ six ligands is Mg^2+^ compatible**  
**6: Number of potential Water contacts**   
**1 : Number of second shell contacts involving water or ions**   
**17%: Percentage of satisfied second shell Water contacts**   
• Contact(s): /O_w_/O_w_/O_w_/OP1/OP1/OP2/  
• Contact code: *mer-*3O_ph_3O_w_  
Ooctahedral coordination (3.0º)   
Go to ToC

### MG:a:3234

|  | Res. type | Atom | Residue | Distance | Occupancy | B-factor | Closest atom(s) to a 1^st^ shell water | EM\_dens. |
| --- | --- | --- | --- | --- | --- | --- | --- | --- |
| **Ion** | | | |  | 1.0 | 28.4 |  | 21.26 |
 1 | A | OP2 | a:1158 | 1.99 | 1.0 | 27.7 |  | 12.20 | 2 | Water 1 | | a:8822 | 2.05 | 1.0 | 25.3 |  | 13.75 | 3 | Water 2 | | a:8134 | 2.09 | 1.0 | 28.3 |  | 13.07 | 4 | Water 3 | | a:5336 | 2.12 | 1.0 | 27.6 | 2.70 [G:O6]a:1159 | 13.50 | 5 | Water 4 | | a:7958 | 2.13 | 1.0 | 27.4 | 2.88 [A:OP2]a:1157 | 14.17 | 6 | Water 5 | | a:8889 | 2.20 | 1.0 | 29.2 | 2.98 [G:O6]a:999 | 11.79 | 7 | Water 6 | | a:4357 | 4.15 | 1.0 | 29.0 | 2.62 [A:OP1]a:1158 | 11.80 | 8 | Water 7 | | a:4201 | 4.16 | 1.0 | 30.5 |  | 12.46 | 9 | A | O5' | a:1158 | 4.20 | 1.0 | 35.5 |  | 9.70 | 10 | Water 8 | | a:3800 | 4.25 | 1.0 | 33.2 | 2.54 [A:OP2]a:1157 | 11.56 | 11 | A | OP1 | a:1158 | 4.27 | 1.0 | 28.1 |  | 11.82 | 12 | Water 9 | | a:9938 | 4.29 | 1.0 | 38.1 |  | 8.04 | 13 | Water 10 | | a:10177 | 4.30 | 1.0 | 46.6 |  | 5.46 | 14 | G | O6 | a:1159 | 4.30 | 1.0 | 30.8 |  | 13.05 | 15 | Water 11 | | p:257 | 4.35 | 1.0 | 33.8 |  | 6.92 | 16 | A | O3' | a:1157 | 4.37 | 1.0 | 26.8 |  | 11.98 | 17 | Water 12 | | p:242 | 4.48 | 1.0 | 31.1 |  | 10.79 | 18 | G | O6 | a:999 | 4.55 | 1.0 | 31.6 |  | 12.76 | 19 | A | O5' | a:1157 | 4.64 | 1.0 | 32.3 |  | 12.27 | 20 | A | OP2 | a:1157 | 4.68 | 1.0 | 29.8 |  | 11.65 |

  
**1^st^ shell coordination angles and distances (*trans* in "blue"; *cys* in "green"; deviations in "orange"):**   
• 1-2(OP2/W): 94º; 1-3(OP2/W): 90º; 1-4(OP2/W): 87º; 1-5(OP2/W): 92º; 1-6(OP2/W): 174º; 2-3(W/W): *97º*; 2-4(W/W): 174º; 2-5(W/W): 88º; 2-6(W/W): 91º; 3-4(W/W): 88º; 3-5(W/W): 174º; 3-6(W/W): 88º; 4-5(W/W): 86º; 4-6(W/W): 88º; 5-6(W/W): 89º;   
• 1-2(OP2/W): 3.0Å; 1-3(OP2/W): 2.9Å; 1-4(OP2/W): 2.8Å; 1-5(OP2/W): 3.0Å; 1-6(OP2/W): 4.2Å; 2-3(W/W): 3.1Å; 2-4(W/W): 4.2Å; 2-5(W/W): 2.9Å; 2-6(W/W): 3.0Å; 3-4(W/W): 2.9Å; 3-5(W/W): *4.2Å*; 3-6(W/W): 3.0Å; 4-5(W/W): 2.9Å; 4-6(W/W): 3.0Å; 5-6(W/W): 3.0Å;   
  
**2.10 ± 0.06: the average coodination distance of the 1^st^ six ligands is Mg^2+^ compatible**  
**10: Number of potential Water contacts**   
**3 : Number of second shell contacts involving water or ions**   
**30%: Percentage of satisfied second shell Water contacts**   
• Contact(s): /O_w_/O_w_/O_w_/O_w_/O_w_/OP2/  
• Contact code: O_ph_5O_w_  
Ooctahedral coordination (3.2º)   
Go to ToC

### MG:a:3235

|  | Res. type | Atom | Residue | Distance | Occupancy | B-factor | Closest atom(s) to a 1^st^ shell water | EM\_dens. |
| --- | --- | --- | --- | --- | --- | --- | --- | --- |
| **Ion** | | | |  | 1.0 | 26.9 |  | 26.03 |
 1 | Water 1 | | a:9967 | 1.92 | 1.0 | 26.0 |  | 14.35 | 2 | Water 2 | | a:8293 | 2.00 | 1.0 | 20.8 | 2.87 [MG:MG]a:3045 | 15.59 | 3 | Water 3 | | a:10092 | 2.16 | 1.0 | 24.4 |  | 14.11 | 4 | Water 4 | | a:6791 | 2.17 | 1.0 | 23.8 | 2.79 [A:N7]a:2013 | 13.29 | 5 | Water 5 | | a:3462 | 2.19 | 1.0 | 23.5 | 2.41 [U:OP2]a:1269 // 2.83 [U:O4]a:2015 | 14.34 | 6 | Water 6 | | a:3488 | 2.39 | 1.0 | 23.9 | 2.43 [G:O6]a:2014 | 14.20 | 7 | **MG** | MG | a:3045 | 3.80 | 1.0 | 37.3 |  | 13.36 | 8 | Water 7 | | a:7699 | 4.07 | 1.0 | 32.2 |  | 11.61 | 9 | Water 8 | | a:4704 | 4.12 | 1.0 | 25.9 |  | 11.12 | 10 | U | O4 | a:2015 | 4.12 | 1.0 | 24.2 |  | 14.38 | 11 | Water 9 | | a:7702 | 4.21 | 1.0 | 37.6 | 1.87 [MG:MG]a:3045 | 8.63 | 12 | A | N7 | a:2013 | 4.21 | 1.0 | 18.6 |  | 17.51 | 13 | U | OP2 | a:1269 | 4.23 | 1.0 | 27.5 |  | 12.88 | 14 | Water 10 | | a:5162 | 4.34 | 1.0 | 27.4 |  | 14.91 | 15 | G | O6 | a:2014 | 4.36 | 1.0 | 23.0 |  | 17.52 | 16 | Water 11 | | a:6675 | 4.41 | 1.0 | 38.1 | 2.78 [U:OP1]a:1269 | 11.13 | 17 | G | N7 | a:2014 | 4.43 | 1.0 | 16.9 |  | 18.14 | 18 | Water 12 | | a:9630 | 4.45 | 1.0 | 29.6 |  | 8.84 | 19 | Water 13 | | a:9498 | 4.55 | 1.0 | 28.9 |  | 9.76 | 20 | U | OP1 | a:1269 | 4.70 | 1.0 | 28.6 |  | 9.85 |

  
**1^st^ shell coordination angles and distances (*trans* in "blue"; *cys* in "green"; deviations in "orange"):**   
• 1-2(W/W): *97º*; 1-3(W/W): *82º*; 1-4(W/W): 87º; 1-5(W/W): 86º; 2-3(W/W): 179º; 2-4(W/W): 91º; 2-5(W/W): 91º; 3-4(W/W): 89º; 3-5(W/W): 88º; 4-5(W/W): 173º;   
• 1-2(W/W): 2.9Å; 1-3(W/W): 2.7Å; 1-4(W/W): 2.8Å; 1-5(W/W): 2.8Å; 2-3(W/W): 4.2Å; 2-4(W/W): 3.0Å; 2-5(W/W): 3.0Å; 3-4(W/W): 3.0Å; 3-5(W/W): 3.0Å; 4-5(W/W): *4.4Å*;   
  
**2.14 ± 0.15: the average coodination distance of the 1^st^ six ligands is Mg^2+^ compatible**  
**12: Number of potential Water contacts**   
**5 : Number of second shell contacts involving water or ions**   
**42%: Percentage of satisfied second shell Water contacts**   
• Contact(s): /O_w_/O_w_/O_w_/O_w_/O_w_/  
• Contact code: 5O_w_  
• Contact(s) in the exclusion zone: /O_w_/  
→ Missing octahedral coordination for MG ←  
Ooctahedral coordination (3.5º)   
Go to ToC

### MG:a:3236

|  | Res. type | Atom | Residue | Distance | Occupancy | B-factor | Closest atom(s) to a 1^st^ shell water | EM\_dens. |
| --- | --- | --- | --- | --- | --- | --- | --- | --- |
| **Ion** | | | |  | 1.0 | 25.8 |  | 24.05 |
 1 | Water 1 | | a:6463 | 1.89 | 1.0 | 27.0 | 2.86 [G:OP1]a:2052 | 13.56 | 2 | C | OP1 | a:2010 | 1.94 | 1.0 | 23.6 |  | 12.71 | 3 | Water 2 | | a:7977 | 2.07 | 1.0 | 28.1 |  | 11.48 | 4 | Water 3 | | a:9050 | 2.10 | 1.0 | 28.4 |  | 13.12 | 5 | Water 4 | | a:8627 | 2.23 | 1.0 | 25.8 | 2.95 [U:O3']a:1269 | 13.79 | 6 | A | OP1 | a:1270 | 2.25 | 1.0 | 24.5 |  | 13.93 | 7 | U | O3' | a:1269 | 3.59 | 1.0 | 30.4 |  | 10.85 | 8 | A | O3' | a:2009 | 3.76 | 1.0 | 25.3 |  | 13.93 | 9 | C | OP2 | a:2010 | 4.06 | 1.0 | 27.1 |  | 14.14 | 10 | G | OP1 | a:2052 | 4.11 | 1.0 | 35.3 |  | 7.14 | 11 | Water 5 | | a:9948 | 4.16 | 1.0 | 38.5 |  | 7.37 | 12 | Water 6 | | a:5856 | 4.35 | 1.0 | 40.8 | 2.73 [G:OP2]a:2053 | 5.14 | 13 | Water 7 | | a:7058 | 4.36 | 1.0 | 36.7 |  | 9.26 | 14 | A | O5' | a:1270 | 4.40 | 1.0 | 30.3 |  | 14.78 | 15 | Water 8 | | a:4507 | 4.43 | 1.0 | 23.1 | 2.63 [C:OP2]a:2010 | 15.10 | 16 | C | O5' | a:2010 | 4.43 | 1.0 | 24.9 |  | 14.04 | 17 | Water 9 | | a:9870 | 4.51 | 1.0 | 39.4 |  | 7.19 | 18 | A | OP2 | a:1270 | 4.53 | 1.0 | 27.5 |  | 14.00 | 19 | Water 10 | | a:6847 | 4.67 | 1.0 | 34.7 |  | 10.65 | 20 | G | OP2 | a:2053 | 4.76 | 1.0 | 23.8 |  | 13.91 | 21 | Water 11 | | a:9672 | 4.77 | 1.0 | 40.6 |  | 4.97 |

  
**1^st^ shell coordination angles and distances (*trans* in "blue"; *cys* in "green"; deviations in "orange"):**   
• 1-2(W/OP1): 93º; 1-3(W/W): 170º; 1-4(W/W): 89º; 1-5(W/W): 88º; 1-6(W/OP1): 91º; 2-3(OP1/W): *96º*; 2-4(OP1/W): *96º*; 2-5(OP1/W): 178º; 2-6(OP1/OP1): 95º; 3-4(W/W): 95º; 3-5(W/W): *83º*; 3-6(W/OP1): *83º*; 4-5(W/W): 86º; 4-6(W/OP1): *169º*; 5-6(W/OP1): *83º*;   
• 1-2(W/OP1): 2.8Å; 1-3(W/W): 4.0Å; 1-4(W/W): 2.8Å; 1-5(W/W): 2.9Å; 1-6(W/OP1): 3.0Å; 2-3(OP1/W): 3.0Å; 2-4(OP1/W): 3.0Å; 2-5(OP1/W): 4.2Å; 2-6(OP1/OP1): 3.1Å; 3-4(W/W): 3.1Å; 3-5(W/W): 2.9Å; 3-6(W/OP1): 2.9Å; 4-5(W/W): 3.0Å; 4-6(W/OP1): *4.3Å*; 5-6(W/OP1): 3.0Å;   
  
**2.08 ± 0.13: the average coodination distance of the 1^st^ six ligands is Mg^2+^ compatible**  
**8: Number of potential Water contacts**   
**2 : Number of second shell contacts involving water or ions**   
**25%: Percentage of satisfied second shell Water contacts**   
• Contact(s): /O_w_/O_w_/O_w_/OP1/OP1/  
• Contact code: *cis-*2O_ph_3O_w_  
• Contact(s) in the exclusion zone: /O_w_/  
→ Missing octahedral coordination for MG ←  
→ Slightly distorded coord. octahedron (5.0º) ←  
Go to ToC

### MG:a:3237

|  | Res. type | Atom | Residue | Distance | Occupancy | B-factor | Closest atom(s) to a 1^st^ shell water | EM\_dens. |
| --- | --- | --- | --- | --- | --- | --- | --- | --- |
| **Ion** | | | |  | 1.0 | 26.6 |  | 25.58 |
 1 | Water 1 | | a:8254 | 1.94 | 1.0 | 26.8 |  | 15.62 | 2 | A | OP1 | a:2009 | 2.05 | 1.0 | 27.7 |  | 13.24 | 3 | Water 2 | | a:4507 | 2.08 | 1.0 | 23.1 | 2.63 [C:OP2]a:2010 | 15.10 | 4 | Water 3 | | a:6147 | 2.19 | 1.0 | 22.1 | 2.75 [A:OP2]a:1270 // 2.84 [A:N7]a:1271 | 16.24 | 5 | A | OP2 | a:1271 | 2.25 | 1.0 | 25.5 |  | 15.83 | 6 | Water 4 | | a:8290 | 2.26 | 1.0 | 25.0 |  | 15.00 | 7 | Water 5 | | a:9630 | 3.94 | 1.0 | 29.6 |  | 8.84 | 8 | A | O5' | a:2009 | 4.14 | 1.0 | 24.9 |  | 15.10 | 9 | A | OP2 | a:2009 | 4.20 | 1.0 | 22.3 |  | 14.68 | 10 | C | OP2 | a:2010 | 4.23 | 1.0 | 27.1 |  | 14.14 | 11 | A | OP2 | a:1270 | 4.27 | 1.0 | 27.5 |  | 14.00 | 12 | Water 6 | | a:3722 | 4.29 | 1.0 | 32.4 |  | 10.90 | 13 | A | N7 | a:1271 | 4.29 | 1.0 | 19.1 |  | 19.01 | 14 | A | O5' | a:1271 | 4.31 | 1.0 | 28.5 |  | 12.46 | 15 | Water 7 | | a:5517 | 4.34 | 1.0 | 27.9 | 2.71 [A:OP2]a:2009 | 13.30 | 16 | G | O3' | a:2008 | 4.52 | 1.0 | 28.6 |  | 13.91 | 17 | A | OP1 | a:1271 | 4.53 | 1.0 | 31.3 |  | 9.96 | 18 | A | O3' | a:1270 | 4.62 | 1.0 | 28.7 |  | 13.28 | 19 | A | OP1 | a:1270 | 4.62 | 1.0 | 24.5 |  | 13.93 | 20 | A | O5' | a:1270 | 4.73 | 1.0 | 30.3 |  | 14.78 |

  
**1^st^ shell coordination angles and distances (*trans* in "blue"; *cys* in "green"; deviations in "orange"):**   
• 1-2(W/OP1): 93º; 1-3(W/W): *95º*; 1-4(W/W): 85º; 1-5(W/OP2): 178º; 1-6(W/W): *97º*; 2-3(OP1/W): 90º; 2-4(OP1/W): 178º; 2-5(OP1/OP2): 88º; 2-6(OP1/W): *85º*; 3-4(W/W): 92º; 3-5(W/OP2): *83º*; 3-6(W/W): *167º*; 4-5(W/OP2): 93º; 4-6(W/W): 93º; 5-6(OP2/W): 85º;   
• 1-2(W/OP1): 2.9Å; 1-3(W/W): 3.0Å; 1-4(W/W): 2.8Å; 1-5(W/OP2): 4.2Å; 1-6(W/W): 3.1Å; 2-3(OP1/W): 2.9Å; 2-4(OP1/W): *4.2Å*; 2-5(OP1/OP2): 3.0Å; 2-6(OP1/W): 2.9Å; 3-4(W/W): 3.1Å; 3-5(W/OP2): 2.9Å; 3-6(W/W): *4.3Å*; 4-5(W/OP2): *3.2Å*; 4-6(W/W): *3.2Å*; 5-6(OP2/W): 3.1Å;   
  
**2.13 ± 0.11: the average coodination distance of the 1^st^ six ligands is Mg^2+^ compatible**  
**8: Number of potential Water contacts**   
**3 : Number of second shell contacts involving water or ions**   
**38%: Percentage of satisfied second shell Water contacts**   
• Contact(s): /O_w_/O_w_/O_w_/O_w_/OP1/OP2/  
• Contact code: *cis-*2O_ph_4O_w_  
Ooctahedral coordination (4.3º)   
Go to ToC

### MG:a:3238

|  | Res. type | Atom | Residue | Distance | Occupancy | B-factor | Closest atom(s) to a 1^st^ shell water | EM\_dens. |
| --- | --- | --- | --- | --- | --- | --- | --- | --- |
| **Ion** | | | |  | 1.0 | 27.5 |  | 23.74 |
 1 | Water 1 | | a:7003 | 1.93 | 1.0 | 27.3 |  | 11.02 | 2 | A | OP1 | a:1274 | 2.03 | 1.0 | 28.1 |  | 14.29 | 3 | G | OP2 | a:1273 | 2.05 | 1.0 | 25.9 |  | 13.01 | 4 | U | OP1 | a:1649 | 2.07 | 1.0 | 30.0 |  | 14.39 | 5 | U | OP1 | a:1650 | 2.09 | 1.0 | 29.7 |  | 12.79 | 6 | Water 2 | | a:6918 | 2.21 | 1.0 | 24.9 | 2.98 [U:OP2]a:1650 | 14.07 | 7 | U | OP2 | a:1650 | 3.87 | 1.0 | 30.5 |  | 8.05 | 8 | Water 3 | | a:4118 | 3.88 | 1.0 | 37.4 |  | 5.90 | 9 | A | OP2 | a:1274 | 3.95 | 1.0 | 28.0 |  | 11.36 | 10 | C | O3' | a:1272 | 4.11 | 1.0 | 23.1 |  | 14.20 | 11 | G | O5' | a:1273 | 4.11 | 1.0 | 30.8 |  | 11.07 | 12 | U | OP2 | a:1649 | 4.12 | 1.0 | 27.7 |  | 11.83 | 13 | Water 4 | | a:8467 | 4.24 | 1.0 | 27.8 | 2.93 [U:OP2]a:1649 | 11.80 | 14 | G | O3' | a:1273 | 4.31 | 1.0 | 29.1 |  | 13.31 | 15 | U | O5' | a:1649 | 4.34 | 1.0 | 24.8 |  | 12.65 | 16 | U | O5' | a:1650 | 4.34 | 1.0 | 27.3 |  | 14.02 | 17 | U | O3' | a:1649 | 4.37 | 1.0 | 29.4 |  | 12.93 | 18 | Water 5 | | a:5892 | 4.37 | 1.0 | 40.1 |  | 4.89 | 19 | A | O5' | a:1274 | 4.40 | 1.0 | 28.0 |  | 11.92 | 20 | G | OP1 | a:1273 | 4.51 | 1.0 | 29.9 |  | 10.81 | 21 | C | O3' | a:1648 | 4.51 | 1.0 | 31.7 |  | 11.79 | 22 | Water 6 | | a:3468 | 4.60 | 1.0 | 32.9 |  | 8.47 |

  
**1^st^ shell coordination angles and distances (*trans* in "blue"; *cys* in "green"; deviations in "orange"):**   
• 1-2(W/OP1): 90º; 1-3(W/OP2): *97º*; 1-4(W/OP1): 90º; 1-5(W/OP1): *96º*; 1-6(W/W): 177º; 2-3(OP1/OP2): 87º; 2-4(OP1/OP1): 95º; 2-5(OP1/OP1): 171º; 2-6(OP1/W): 87º; 3-4(OP2/OP1): 173º; 3-5(OP2/OP1): 85º; 3-6(OP2/W): *82º*; 4-5(OP1/OP1): 92º; 4-6(OP1/W): 91º; 5-6(OP1/W): 86º;   
• 1-2(W/OP1): 2.8Å; 1-3(W/OP2): 3.0Å; 1-4(W/OP1): 2.8Å; 1-5(W/OP1): 3.0Å; 1-6(W/W): 4.1Å; 2-3(OP1/OP2): 2.8Å; 2-4(OP1/OP1): 3.0Å; 2-5(OP1/OP1): 4.1Å; 2-6(OP1/W): 2.9Å; 3-4(OP2/OP1): 4.1Å; 3-5(OP2/OP1): 2.8Å; 3-6(OP2/W): 2.8Å; 4-5(OP1/OP1): 3.0Å; 4-6(OP1/W): 3.0Å; 5-6(OP1/W): 2.9Å;   
  
**2.06 ± 0.08: the average coodination distance of the 1^st^ six ligands is Mg^2+^ compatible**  
**4: Number of potential Water contacts**   
**1 : Number of second shell contacts involving water or ions**   
**25%: Percentage of satisfied second shell Water contacts**   
• Contact(s): /O_w_/O_w_/OP1/OP1/OP1/OP2/  
• Contact code: 4O_ph_2O_w_  
Ooctahedral coordination (4.2º)   
Go to ToC

### MG:a:3239

|  | Res. type | Atom | Residue | Distance | Occupancy | B-factor | Closest atom(s) to a 1^st^ shell water | EM\_dens. |
| --- | --- | --- | --- | --- | --- | --- | --- | --- |
| **Ion** | | | |  | 1.0 | 32.2 |  | 16.71 |
 1 | U | OP1 | a:1315 | 1.95 | 1.0 | 32.5 |  | 11.74 | 2 | Water 1 | | a:7943 | 2.00 | 1.0 | 30.7 |  | 8.91 | 3 | Water 2 | | a:7291 | 2.04 | 1.0 | 30.2 | 2.83 [G:OP1]a:1336 | 10.52 | 4 | Water 3 | | a:8820 | 2.05 | 1.0 | 34.1 |  | 10.43 | 5 | Water 4 | | a:6725 | 2.06 | 1.0 | 27.6 |  | 11.28 | 6 | Water 5 | | a:3562 | 2.34 | 1.0 | 26.9 | 2.48 [G:OP2]a:1336 | 13.74 | 7 | U | OP2 | a:1315 | 4.01 | 1.0 | 27.1 |  | 11.40 | 8 | Water 6 | | a:9125 | 4.13 | 1.0 | 25.8 |  | 14.27 | 9 | Water 7 | | a:4537 | 4.25 | 1.0 | 33.1 | 2.86 [G:OP2]a:1336 | 8.05 | 10 | U | O3' | a:1314 | 4.27 | 1.0 | 27.0 |  | 12.23 | 11 | G | OP2 | a:1336 | 4.36 | 1.0 | 34.1 |  | 11.56 | 12 | U | O5' | a:1315 | 4.37 | 1.0 | 29.7 |  | 11.71 | 13 | G | OP1 | a:1335 | 4.43 | 1.0 | 27.0 |  | 12.43 | 14 | G | OP1 | a:1336 | 4.48 | 1.0 | 30.9 |  | 9.22 | 15 | Water 8 | | s:217 | 4.71 | 1.0 | 44.0 |  | 3.72 |

  
**1^st^ shell coordination angles and distances (*trans* in "blue"; *cys* in "green"; deviations in "orange"):**   
• 1-2(OP1/W): 94º; 1-3(OP1/W): 172º; 1-4(OP1/W): *96º*; 1-5(OP1/W): 88º; 2-3(W/W): 85º; 2-4(W/W): 92º; 2-5(W/W): 178º; 3-4(W/W): 92º; 3-5(W/W): 93º; 4-5(W/W): 90º;   
• 1-2(OP1/W): 2.9Å; 1-3(OP1/W): 4.0Å; 1-4(OP1/W): 3.0Å; 1-5(OP1/W): 2.8Å; 2-3(W/W): 2.7Å; 2-4(W/W): 2.9Å; 2-5(W/W): 4.1Å; 3-4(W/W): 2.9Å; 3-5(W/W): 3.0Å; 4-5(W/W): 2.9Å;   
  
**2.07 ± 0.13: the average coodination distance of the 1^st^ six ligands is Mg^2+^ compatible**  
**10: Number of potential Water contacts**   
**2 : Number of second shell contacts involving water or ions**   
**20%: Percentage of satisfied second shell Water contacts**   
• Contact(s): /O_w_/O_w_/O_w_/O_w_/OP1/  
• Contact code: O_ph_4O_w_  
• Contact(s) in the exclusion zone: /O_w_/  
→ Missing octahedral coordination for MG ←  
Ooctahedral coordination (3.4º)   
Go to ToC

### MG:a:3240

|  | Res. type | Atom | Residue | Distance | Occupancy | B-factor | Closest atom(s) to a 1^st^ shell water | EM\_dens. |
| --- | --- | --- | --- | --- | --- | --- | --- | --- |
| **Ion** | | | |  | 1.0 | 28.9 |  | 18.70 |
 1 | Water 1 | | a:9824 | 1.97 | 1.0 | 30.2 | 2.82 [MG:MG]a:3365 | 11.80 | 2 | Water 2 | | a:9125 | 2.09 | 1.0 | 25.8 |  | 14.27 | 3 | Water 3 | | a:7086 | 2.10 | 1.0 | 24.6 | 2.85 [C:OP2]a:1316 // 2.86 [G:O5']a:1334 | 13.01 | 4 | Water 4 | | a:5908 | 2.12 | 1.0 | 26.9 | 2.74 [G:N7]a:1335 | 13.26 | 5 | G | OP2 | a:1335 | 2.14 | 1.0 | 29.9 |  | 10.28 | 6 | Water 5 | | a:6887 | 2.28 | 1.0 | 27.1 | 2.80 [G:OP2]a:1336 | 12.44 | 7 | **MG** | MG | a:3365 | 4.11 | 1.0 | 39.0 |  | 10.85 | 8 | G | O5' | a:1334 | 4.13 | 1.0 | 27.6 |  | 9.89 | 9 | Water 6 | | a:3562 | 4.17 | 1.0 | 26.9 | 2.48 [G:OP2]a:1336 | 13.74 | 10 | G | O5' | a:1335 | 4.24 | 1.0 | 30.3 |  | 11.06 | 11 | G | OP2 | a:1336 | 4.24 | 1.0 | 34.1 |  | 11.56 | 12 | G | OP2 | a:1334 | 4.26 | 1.0 | 26.0 |  | 11.33 | 13 | G | OP1 | a:1335 | 4.28 | 1.0 | 27.0 |  | 12.43 | 14 | Water 7 | | a:9524 | 4.30 | 1.0 | 41.9 |  | 5.36 | 15 | G | N7 | a:1335 | 4.32 | 1.0 | 21.9 |  | 12.72 | 16 | G | O3' | a:1334 | 4.57 | 1.0 | 33.3 |  | 12.32 | 17 | Water 8 | | a:5539 | 4.63 | 1.0 | 46.2 | 2.71 [G:OP2]a:1334 | 5.09 | 18 | C | OP2 | a:1316 | 4.76 | 1.0 | 29.5 |  | 11.43 |

  
**1^st^ shell coordination angles and distances (*trans* in "blue"; *cys* in "green"; deviations in "orange"):**   
• 1-2(W/W): 94º; 1-3(W/W): 93º; 1-4(W/W): 90º; 1-5(W/OP2): 174º; 1-6(W/W): *95º*; 2-3(W/W): *100º*; 2-4(W/W): 172º; 2-5(W/OP2): 91º; 2-6(W/W): 88º; 3-4(W/W): 86º; 3-5(W/OP2): *83º*; 3-6(W/W): *168º*; 4-5(W/OP2): *85º*; 4-6(W/W): 85º; 5-6(OP2/W): 88º;   
• 1-2(W/W): 3.0Å; 1-3(W/W): 3.0Å; 1-4(W/W): 2.9Å; 1-5(W/OP2): 4.1Å; 1-6(W/W): 3.1Å; 2-3(W/W): *3.2Å*; 2-4(W/W): 4.2Å; 2-5(W/OP2): 3.0Å; 2-6(W/W): 3.0Å; 3-4(W/W): 2.9Å; 3-5(W/OP2): 2.8Å; 3-6(W/W): *4.4Å*; 4-5(W/OP2): 2.9Å; 4-6(W/W): 3.0Å; 5-6(OP2/W): 3.1Å;   
  
**2.12 ± 0.09: the average coodination distance of the 1^st^ six ligands is Mg^2+^ compatible**  
**10: Number of potential Water contacts**   
**5 : Number of second shell contacts involving water or ions**   
**50%: Percentage of satisfied second shell Water contacts**   
• Contact(s): /O_w_/O_w_/O_w_/O_w_/O_w_/OP2/  
• Contact code: O_ph_5O_w_  
→ Slightly distorded coord. octahedron (5.0º) ←  
Go to ToC

### MG:a:3241

|  | Res. type | Atom | Residue | Distance | Occupancy | B-factor | Closest atom(s) to a 1^st^ shell water | EM\_dens. |
| --- | --- | --- | --- | --- | --- | --- | --- | --- |
| **Ion** | | | |  | 1.0 | 33.5 |  | 15.07 |
 1 | Water 1 | | a:7447 | 2.00 | 1.0 | 30.6 | 2.84 [G:N7]a:1379 | 11.26 | 2 | Water 2 | | a:10184 | 2.03 | 1.0 | 31.0 |  | 10.74 | 3 | Water 3 | | a:7420 | 2.08 | 1.0 | 28.8 | 2.84 [C:OP2]a:1353 // 2.84 [U:OP2]a:1354 | 10.89 | 4 | Water 4 | | a:5071 | 2.10 | 1.0 | 26.0 | 2.68 [G:O6]a:1379 | 11.86 | 5 | Water 5 | | a:8677 | 2.25 | 1.0 | 30.2 | 2.95 [C:OP2]a:1378 | 12.94 | 6 | Water 6 | | a:3512 | 2.50 | 1.0 | 32.8 | 2.45 [C:OP2]a:1378 | 10.11 | 7 | U | OP2 | a:1354 | 3.82 | 1.0 | 34.1 |  | 11.13 | 8 | C | OP2 | a:1378 | 4.06 | 1.0 | 32.9 |  | 10.82 | 9 | Water 7 | | a:4933 | 4.08 | 1.0 | 31.7 | 2.67 [C:OP2]a:1353 | 11.48 | 10 | G | N7 | a:1379 | 4.22 | 1.0 | 19.7 |  | 17.82 | 11 | C | OP2 | a:1353 | 4.31 | 1.0 | 35.5 |  | 9.58 | 12 | Water 8 | | a:5810 | 4.34 | 1.0 | 34.0 |  | 8.72 | 13 | Water 9 | | a:10076 | 4.39 | 1.0 | 43.5 |  | 5.63 | 14 | G | O6 | a:1379 | 4.57 | 1.0 | 23.3 |  | 15.48 | 15 | Water 10 | | a:6414 | 4.60 | 1.0 | 42.0 |  | 6.65 | 16 | G | OP2 | a:1379 | 4.67 | 1.0 | 30.0 |  | 12.41 |

  
**1^st^ shell coordination angles and distances (*trans* in "blue"; *cys* in "green"; deviations in "orange"):**   
• 1-2(W/W): *82º*; 1-3(W/W): 177º; 1-4(W/W): 94º; 1-5(W/W): 89º; 2-3(W/W): *99º*; 2-4(W/W): *105º*; 2-5(W/W): 87º; 3-4(W/W): 89º; 3-5(W/W): 88º; 4-5(W/W): *168º*;   
• 1-2(W/W): 2.7Å; 1-3(W/W): 4.1Å; 1-4(W/W): 3.0Å; 1-5(W/W): 3.0Å; 2-3(W/W): 3.1Å; 2-4(W/W): *3.3Å*; 2-5(W/W): 3.0Å; 3-4(W/W): 2.9Å; 3-5(W/W): 3.0Å; 4-5(W/W): *4.3Å*;   
  
**2.16 ± 0.17: the average coodination distance of the 1^st^ six ligands is Mg^2+^ compatible**  
**12: Number of potential Water contacts**   
**6 : Number of second shell contacts involving water or ions**   
**50%: Percentage of satisfied second shell Water contacts**   
• Contact(s): /O_w_/O_w_/O_w_/O_w_/O_w_/  
• Contact code: 5O_w_  
• Contact(s) in the exclusion zone: /O_w_/  
→ Missing octahedral coordination for MG ←  
→ Slightly distorded coord. octahedron (5.8º) ←  
Go to ToC

### MG:a:3242

|  | Res. type | Atom | Residue | Distance | Occupancy | B-factor | Closest atom(s) to a 1^st^ shell water | EM\_dens. |
| --- | --- | --- | --- | --- | --- | --- | --- | --- |
| **Ion** | | | |  | 1.0 | 20.9 |  | 27.91 |
 1 | Water 1 | | a:4314 | 1.94 | 1.0 | 42.7 | 2.61 [G:O6]a:1373 | 4.50 | 2 | Water 2 | | a:9040 | 2.03 | 1.0 | 41.3 | 3.00 [C:OP2]a:1359 | 5.49 | 3 | G | N7 | a:1360 | 2.14 | 1.0 | 19.2 |  | 13.24 | 4 | G | N7 | a:1373 | 2.18 | 1.0 | 18.8 |  | 14.75 | 5 | Water 3 | | a:4463 | 2.28 | 1.0 | 32.7 | 2.63 [G:OP2]a:1373 | 6.00 | 6 | Water 4 | | a:8826 | 2.48 | 1.0 | 43.5 | 2.97 [G:OP2]a:1373 | 4.09 | 7 | G | O6 | a:1373 | 3.41 | 1.0 | 28.5 |  | 12.51 | 8 | G | O6 | a:1360 | 3.64 | 1.0 | 26.8 |  | 13.30 | 9 | G | OP2 | a:1360 | 4.16 | 1.0 | 33.4 |  | 9.65 | 10 | G | OP2 | a:1373 | 4.31 | 1.0 | 29.0 |  | 11.45 | 11 | C | OP2 | a:1359 | 4.62 | 1.0 | 34.2 |  | 8.03 | 12 | Water 5 | | a:6667 | 4.74 | 1.0 | 45.3 | 2.78 [G:OP2]a:1373 | 3.49 |

  
**1^st^ shell coordination angles and distances (*trans* in "blue"; *cys* in "green"; deviations in "orange"):**   
• 1-2(W/W): *76º*; 1-3(W/N7): *107º*; 1-4(W/N7): 92º; 1-5(W/W): *157º*; 2-3(W/N7): *104º*; 2-4(W/N7): *162º*; 2-5(W/W): 94º; 3-4(N7/N7): 92º; 3-5(N7/W): 95º; 4-5(N7/W): 93º;   
• 1-2(W/W): 2.4Å; 1-3(W/N7): *3.3Å*; 1-4(W/N7): 3.0Å; 1-5(W/W): 4.1Å; 2-3(W/N7): *3.3Å*; 2-4(W/N7): 4.2Å; 2-5(W/W): 3.2Å; 3-4(N7/N7): 3.1Å; 3-5(N7/W): *3.3Å*; 4-5(N7/W): *3.2Å*;   
  
**2.17 ± 0.18: the average coodination distance of the 1^st^ six ligands is Mg^2+^ compatible**  
**8: Number of potential Water contacts**   
**4 : Number of second shell contacts involving water or ions**   
**50%: Percentage of satisfied second shell Water contacts**   
• Contact(s): /N7/N7/O_w_/O_w_/O_w_/  
• Contact code: *cis-*2N_b_3O_w_  
• Contact(s) in the exclusion zone: /O_w_/  
→ Missing octahedral coordination for MG ←  
→→→ Distorded coord. octahedron (10.2º) ←←←  
Go to ToC

### MG:a:3243

|  | Res. type | Atom | Residue | Distance | Occupancy | B-factor | Closest atom(s) to a 1^st^ shell water | EM\_dens. |
| --- | --- | --- | --- | --- | --- | --- | --- | --- |
| **Ion** | | | |  | 1.0 | 33.9 |  | 15.23 |
 1 | U | OP1 | a:1396 | 1.97 | 1.0 | 33.2 |  | 10.67 | 2 | C | OP1 | a:1606 | 2.05 | 1.0 | 33.3 |  | 11.02 | 3 | Water 1 | | a:8624 | 2.06 | 1.0 | 37.5 |  | 9.36 | 4 | Water 2 | | a:5201 | 2.07 | 1.0 | 28.6 | 2.69 [A:OP2]a:1397 | 10.64 | 5 | Water 3 | | a:8053 | 2.13 | 1.0 | 30.7 |  | 9.38 | 6 | Water 4 | | a:8469 | 2.41 | 1.0 | 31.5 |  | 10.67 | 7 | A | O3' | a:1395 | 3.67 | 1.0 | 35.7 |  | 8.43 | 8 | Water 5 | | a:8169 | 3.93 | 1.0 | 40.1 | 2.90 [U:OP2]a:1396 | 4.70 | 9 | A | OP2 | a:1397 | 4.13 | 1.0 | 36.5 |  | 9.58 | 10 | A | O3' | a:1605 | 4.19 | 1.0 | 28.8 |  | 12.87 | 11 | U | OP2 | a:1396 | 4.20 | 1.0 | 36.8 |  | 8.40 | 12 | C | O5' | a:1606 | 4.26 | 1.0 | 34.6 |  | 11.91 | 13 | Water 6 | | a:4282 | 4.28 | 1.0 | 32.8 |  | 10.56 | 14 | U | O5' | a:1396 | 4.37 | 1.0 | 32.4 |  | 11.59 | 15 | Water 7 | | a:8135 | 4.44 | 1.0 | 37.3 |  | 8.69 | 16 | C | OP2 | a:1606 | 4.45 | 1.0 | 30.3 |  | 10.81 | 17 | Water 8 | | a:4280 | 4.48 | 1.0 | 37.2 |  | 7.95 | 18 | Water 9 | | a:8762 | 4.51 | 1.0 | 41.8 |  | 5.11 | 19 | Water 10 | | a:5006 | 4.73 | 1.0 | 39.0 | 2.67 [A:OP2]a:1397 | 6.80 |

  
**1^st^ shell coordination angles and distances (*trans* in "blue"; *cys* in "green"; deviations in "orange"):**   
• 1-2(OP1/OP1): *97º*; 1-3(OP1/W): 94º; 1-4(OP1/W): 87º; 1-5(OP1/W): *96º*; 2-3(OP1/W): *168º*; 2-4(OP1/W): 87º; 2-5(OP1/W): 87º; 3-4(W/W): 91º; 3-5(W/W): 94º; 4-5(W/W): 174º;   
• 1-2(OP1/OP1): 3.0Å; 1-3(OP1/W): 2.9Å; 1-4(OP1/W): 2.8Å; 1-5(OP1/W): 3.1Å; 2-3(OP1/W): 4.1Å; 2-4(OP1/W): 2.8Å; 2-5(OP1/W): 2.9Å; 3-4(W/W): 3.0Å; 3-5(W/W): 3.1Å; 4-5(W/W): 4.2Å;   
  
**2.12 ± 0.14: the average coodination distance of the 1^st^ six ligands is Mg^2+^ compatible**  
**8: Number of potential Water contacts**   
**1 : Number of second shell contacts involving water or ions**   
**12%: Percentage of satisfied second shell Water contacts**   
• Contact(s): /O_w_/O_w_/O_w_/OP1/OP1/  
• Contact code: *cis-*2O_ph_3O_w_  
• Contact(s) in the exclusion zone: /O_w_/  
→ Missing octahedral coordination for MG ←  
Ooctahedral coordination (4.9º)   
Go to ToC

### MG:a:3244

|  | Res. type | Atom | Residue | Distance | Occupancy | B-factor | Closest atom(s) to a 1^st^ shell water | EM\_dens. |
| --- | --- | --- | --- | --- | --- | --- | --- | --- |
| **Ion** | | | |  | 1.0 | 30.5 |  | 15.30 |
 1 | A | OP1 | a:1605 | 1.98 | 1.0 | 31.3 |  | 9.79 | 2 | C | OP2 | a:1606 | 1.99 | 1.0 | 30.3 |  | 10.81 | 3 | A | OP1 | a:1397 | 2.04 | 1.0 | 33.2 |  | 8.65 | 4 | Water 1 | | a:8681 | 2.16 | 1.0 | 35.6 |  | 9.32 | 5 | Water 2 | | a:6698 | 2.20 | 1.0 | 32.8 |  | 9.63 | 6 | Water 3 | | a:8135 | 2.25 | 1.0 | 37.3 |  | 8.69 | 7 | A | O5' | a:1605 | 3.86 | 1.0 | 29.2 |  | 13.51 | 8 | C | OP1 | a:1606 | 3.94 | 1.0 | 33.3 |  | 11.02 | 9 | U | O3' | a:1396 | 3.94 | 1.0 | 26.8 |  | 11.86 | 10 | A | OP2 | a:1605 | 4.06 | 1.0 | 33.2 |  | 8.89 | 11 | Water 4 | | a:9660 | 4.07 | 1.0 | 39.2 | 2.73 [K:K]a:3141 | 6.30 | 12 | **K** | K | a:3141 | 4.17 | 1.0 | 30.0 |  | 25.68 | 13 | Water 5 | | a:3744 | 4.23 | 1.0 | 41.8 | 2.52 [A:OP2]a:1605 | 5.23 | 14 | C | O5' | a:1606 | 4.31 | 1.0 | 34.6 |  | 11.91 | 15 | A | OP2 | a:1397 | 4.32 | 1.0 | 36.5 |  | 9.58 | 16 | A | O3' | a:1605 | 4.34 | 1.0 | 28.8 |  | 12.87 | 17 | Water 6 | | a:6102 | 4.37 | 1.0 | 44.2 |  | 6.42 | 18 | A | O5' | a:1397 | 4.44 | 1.0 | 37.0 |  | 9.28 | 19 | U | O3' | a:1604 | 4.44 | 1.0 | 30.7 |  | 11.41 | 20 | C | OP1 | a:1308 | 4.45 | 1.0 | 48.7 |  | 3.24 | 21 | Water 7 | | a:5201 | 4.55 | 1.0 | 28.6 | 2.69 [A:OP2]a:1397 // 2.84 [C:OP1]a:1606 | 10.64 | 22 | Water 8 | | a:4280 | 4.56 | 1.0 | 37.2 |  | 7.95 |

  
**1^st^ shell coordination angles and distances (*trans* in "blue"; *cys* in "green"; deviations in "orange"):**   
• 1-2(OP1/OP2): 95º; 1-3(OP1/OP1): *97º*; 1-4(OP1/W): 91º; 1-5(OP1/W): 88º; 1-6(OP1/W): 175º; 2-3(OP2/OP1): *102º*; 2-4(OP2/W): 93º; 2-5(OP2/W): 175º; 2-6(OP2/W): 90º; 3-4(OP1/W): *162º*; 3-5(OP1/W): *82º*; 3-6(OP1/W): *85º*; 4-5(W/W): *82º*; 4-6(W/W): 86º; 5-6(W/W): 87º;   
• 1-2(OP1/OP2): 2.9Å; 1-3(OP1/OP1): 3.0Å; 1-4(OP1/W): 3.0Å; 1-5(OP1/W): 2.9Å; 1-6(OP1/W): *4.2Å*; 2-3(OP2/OP1): 3.1Å; 2-4(OP2/W): 3.0Å; 2-5(OP2/W): 4.2Å; 2-6(OP2/W): 3.0Å; 3-4(OP1/W): 4.2Å; 3-5(OP1/W): 2.8Å; 3-6(OP1/W): 2.9Å; 4-5(W/W): 2.9Å; 4-6(W/W): 3.0Å; 5-6(W/W): 3.1Å;   
  
**2.10 ± 0.11: the average coodination distance of the 1^st^ six ligands is Mg^2+^ compatible**  
**6: Number of potential Water contacts**   
**0 : Number of second shell contacts involving water or ions**   
**0%: Percentage of satisfied second shell Water contacts**   
• Contact(s): /O_w_/O_w_/O_w_/OP1/OP1/OP2/  
• Contact code: *fac-*3O_ph_3O_w_  
→ Slightly distorded coord. octahedron (5.7º) ←  
Go to ToC

### MG:a:3245

|  | Res. type | Atom | Residue | Distance | Occupancy | B-factor | Closest atom(s) to a 1^st^ shell water | EM\_dens. |
| --- | --- | --- | --- | --- | --- | --- | --- | --- |
| **Ion** | | | |  | 1.0 | 34.6 |  | 17.00 |
 1 | Water 1 | | a:7741 | 1.91 | 1.0 | 33.9 | 2.86 [A:N3]a:1618 | 8.83 | 2 | U | OP2 | a:1649 | 1.97 | 1.0 | 27.7 |  | 11.83 | 3 | Water 2 | | a:5656 | 2.12 | 1.0 | 29.3 | 2.72 [U:OP1]a:1327 | 12.03 | 4 | Water 3 | | a:4595 | 2.16 | 1.0 | 32.3 | 2.64 [A:O2']a:1618 | 9.36 | 5 | Water 4 | | a:8446 | 2.22 | 1.0 | 35.8 | 2.97 [C:O3']a:1648 | 9.05 | 6 | Water 5 | | a:8467 | 2.38 | 1.0 | 27.8 |  | 11.80 | 7 | C | O3' | a:1648 | 3.75 | 1.0 | 31.7 |  | 11.79 | 8 | Water 6 | | a:9200 | 3.82 | 1.0 | 40.9 |  | 5.31 | 9 | A | O2' | a:1618 | 3.99 | 1.0 | 29.1 |  | 10.39 | 10 | Water 7 | | a:4026 | 4.18 | 1.0 | 42.8 | 2.58 [A:O2']a:1618 | 5.42 | 11 | U | OP1 | a:1649 | 4.22 | 1.0 | 30.0 |  | 14.39 | 12 | Water 8 | | a:4127 | 4.24 | 1.0 | 42.4 |  | 4.61 | 13 | U | O5' | a:1649 | 4.29 | 1.0 | 24.8 |  | 12.65 | 14 | Water 9 | | a:6918 | 4.32 | 1.0 | 24.9 |  | 14.07 | 15 | U | OP1 | a:1327 | 4.34 | 1.0 | 34.7 |  | 11.21 | 16 | A | N3 | a:1618 | 4.62 | 1.0 | 28.3 |  | 11.32 | 17 | Water 10 | | a:4543 | 4.66 | 1.0 | 33.1 |  | 11.47 |

  
**1^st^ shell coordination angles and distances (*trans* in "blue"; *cys* in "green"; deviations in "orange"):**   
• 1-2(W/OP2): *104º*; 1-3(W/W): 91º; 1-4(W/W): *85º*; 1-5(W/W): *101º*; 2-3(OP2/W): 91º; 2-4(OP2/W): 170º; 2-5(OP2/W): 88º; 3-4(W/W): 92º; 3-5(W/W): *168º*; 4-5(W/W): 87º;   
• 1-2(W/OP2): 3.1Å; 1-3(W/W): 2.9Å; 1-4(W/W): 2.7Å; 1-5(W/W): 3.2Å; 2-3(OP2/W): 2.9Å; 2-4(OP2/W): 4.1Å; 2-5(OP2/W): 2.9Å; 3-4(W/W): 3.1Å; 3-5(W/W): *4.3Å*; 4-5(W/W): 3.0Å;   
  
**2.13 ± 0.16: the average coodination distance of the 1^st^ six ligands is Mg^2+^ compatible**  
**10: Number of potential Water contacts**   
**4 : Number of second shell contacts involving water or ions**   
**40%: Percentage of satisfied second shell Water contacts**   
• Contact(s): /O_w_/O_w_/O_w_/O_w_/OP2/  
• Contact code: O_ph_4O_w_  
• Contact(s) in the exclusion zone: /O_w_/  
→ Missing octahedral coordination for MG ←  
→ Slightly distorded coord. octahedron (6.0º) ←  
Go to ToC

### K:a:3246

|  | Res. type | Atom | Residue | Distance | Occupancy | B-factor | Closest atom(s) to a 1^st^ shell water | EM\_dens. |
| --- | --- | --- | --- | --- | --- | --- | --- | --- |
| **Ion** | | | |  | 1.0 | 30.0 |  | 12.67 |
 1 | **MG** | MG | a:3155 | 2.42 | 1.0 | 52.6 |  | 6.99 | 2 | G | O6 | a:2649 | 2.60 | 1.0 | 37.2 |  | 6.33 | 3 | G | O6 | a:2648 | 2.71 | 1.0 | 33.5 |  | 12.33 | 4 | Water 1 | | a:5598 | 2.81 | 1.0 | 34.8 | 2.72 [G:N7]a:2648 | 8.75 | 5 | Water 2 | | a:5167 | 3.01 | 1.0 | 36.4 | 2.69 [G:N7]a:2647 | 6.45 | 6 | Water 3 | | a:9145 | 3.31 | 1.0 | 40.8 |  | 5.39 | 7 | Water 4 | | a:9663 | 3.45 | 1.0 | 43.2 |  | 3.93 | 8 | Water 5 | | a:7827 | 3.49 | 1.0 | 48.3 |  | 4.67 | 9 | Water 6 | | a:7715 | 3.50 | 1.0 | 42.1 |  | 4.69 | 10 | Water 7 | | a:7201 | 3.64 | 1.0 | 38.8 |  | 5.50 | 11 | G | N7 | a:2648 | 3.98 | 1.0 | 22.5 |  | 12.49 | 12 | Water 8 | | a:4443 | 4.44 | 1.0 | 43.1 |  | 5.24 | 13 | G | OP2 | a:2647 | 4.54 | 1.0 | 31.9 |  | 12.47 | 14 | G | N7 | a:2647 | 4.75 | 1.0 | 26.7 |  | 12.55 |

  
**1^st^ shell coordination angles and distances (*trans* in "blue"; *cys* in "green"; deviations in "orange"):**   
• 1-2(MG/O6): 115; 1-3(MG/O6): 125; 1-4(MG/W): 44; 1-5(MG/W): 56; 2-3(O6/O6): 68; 2-4(O6/W): 76; 2-5(O6/W): 133; 3-4(O6/W): 92; 3-5(O6/W): 82; 4-5(W/W): 69;   
• 1-2(MG/O6): 4.2; 1-3(MG/O6): 4.6; 1-4(MG/W): 2.0; 1-5(MG/W): 2.6; 2-3(O6/O6): 3.0; 2-4(O6/W): 3.3; 2-5(O6/W): 5.2; 3-4(O6/W): 4.0; 3-5(O6/W): 3.8; 4-5(W/W): 3.3;   
  
**3.97 ± 1.23: the average coodination distance of the 1^st^ six ligands could correspond to an hexahydrated Mg^2+^.**  
**10: Number of potential Water contacts**   
**2 : Number of second shell contacts involving water or ions**   
**20%: Percentage of satisfied second shell Water contacts**   
• Contact(s): /O6/O_w_/O_w_/  
• Contact code: O_b_2O_w_  
• Contact(s) in the exclusion zone: /O6/  
→ Coordination < 6 for K ←  
Go to ToC

### K:a:3247

|  | Res. type | Atom | Residue | Distance | Occupancy | B-factor | Closest atom(s) to a 1^st^ shell water | EM\_dens. |
| --- | --- | --- | --- | --- | --- | --- | --- | --- |
| **Ion** | | | |  | 1.0 | 30.0 |  | 14.30 |
 1 | Water 1 | | a:8389 | 2.83 | 1.0 | 32.0 |  | 12.25 | 2 | G | O5' | a:2254 | 2.87 | 1.0 | 30.7 |  | 11.22 | 3 | Water 2 | | a:9871 | 2.91 | 1.0 | 44.0 |  | 3.46 | 4 | G | N3 | a:2254 | 2.91 | 1.0 | 22.0 |  | 12.13 | 5 | G | O4' | a:2254 | 3.13 | 1.0 | 32.3 |  | 12.30 | 6 | G | OP2 | a:2254 | 3.15 | 1.0 | 30.8 |  | 8.63 | 7 | Water 3 | | a:7044 | 3.50 | 1.0 | 40.8 |  | 5.25 | 8 | Water 4 | | a:6599 | 3.56 | 1.0 | 45.6 |  | 5.01 | 9 | Water 5 | | a:9196 | 3.68 | 1.0 | 37.7 |  | 6.54 | 10 | Water 6 | | a:10084 | 4.37 | 1.0 | 40.7 |  | 6.69 | 11 | Water 7 | | a:6416 | 4.47 | 1.0 | 31.9 |  | 12.05 | 12 | Water 8 | | a:10156 | 4.52 | 1.0 | 44.5 |  | 4.15 | 13 | U | O3' | a:2253 | 4.63 | 1.0 | 30.5 |  | 10.74 | 14 | G | OP1 | a:2254 | 4.79 | 1.0 | 32.8 |  | 10.73 |

  
**1^st^ shell coordination angles and distances (*trans* in "blue"; *cys* in "green"; deviations in "orange"):**   
• 1-2(W/O5'): 138; 1-3(W/W): 102; 1-4(W/N3): 79; 1-5(W/O4'): 89; 1-6(W/OP2): 159; 2-3(O5'/W): 100; 2-4(O5'/N3): 66; 2-5(O5'/O4'): 53; 2-6(O5'/OP2): 47; 3-4(W/N3): 153; 3-5(W/O4'): 96; 3-6(W/OP2): 96; 4-5(N3/O4'): 57; 4-6(N3/OP2): 91; 5-6(O4'/OP2): 100;   
• 1-2(W/O5'): 5.3; 1-3(W/W): 4.4; 1-4(W/N3): 3.6; 1-5(W/O4'): 4.2; 1-6(W/OP2): 5.9; 2-3(O5'/W): 4.4; 2-4(O5'/N3): 3.1; 2-5(O5'/O4'): 2.7; 2-6(O5'/OP2): 2.4; 3-4(W/N3): 5.7; 3-5(W/O4'): 4.5; 3-6(W/OP2): 4.5; 4-5(N3/O4'): 2.9; 4-6(N3/OP2): 4.3; 5-6(O4'/OP2): 4.8;   
  
**4.14 ± 1.23: the average coodination distance of the 1^st^ six ligands could correspond to an hexahydrated Mg^2+^.**  
**8: Number of potential Water contacts**   
**0 : Number of second shell contacts involving water or ions**   
**0%: Percentage of satisfied second shell Water contacts**   
• Contact(s): /N3_G_/O4'/O5'/O_w_/O_w_/OP2/  
• Contact code: O_ph_2O_r_N_b_2O_w_  
Go to ToC

### K:a:3248

|  | Res. type | Atom | Residue | Distance | Occupancy | B-factor | Closest atom(s) to a 1^st^ shell water | EM\_dens. |
| --- | --- | --- | --- | --- | --- | --- | --- | --- |
| **Ion** | | | |  | 1.0 | 30.0 |  | 12.24 |
 1 | Water 1 | | a:8856 | 2.54 | 1.0 | 44.0 | 2.97 [A:OP2]a:226 | 4.49 | 2 | A | OP2 | a:2411 | 2.79 | 1.0 | 33.3 |  | 9.51 | 3 | G | OP2 | a:411 | 2.87 | 1.0 | 35.3 |  | 8.70 | 4 | Water 2 | | a:6353 | 3.03 | 1.0 | 35.5 | 3.00 [A:OP2]a:226 | 7.78 | 5 | G | O3' | a:410 | 3.20 | 1.0 | 32.5 |  | 11.69 | 6 | Water 3 | | a:9474 | 3.27 | 1.0 | 50.8 |  | 3.63 | 7 | A | O3' | a:2410 | 3.48 | 1.0 | 38.2 |  | 8.14 | 8 | Water 4 | | a:5117 | 3.62 | 1.0 | 48.5 |  | 4.60 | 9 | Water 5 | | a:7323 | 3.65 | 1.0 | 47.3 |  | 5.60 | 10 | G | O2' | a:410 | 3.67 | 1.0 | 33.5 |  | 11.11 | 11 | A | OP2 | a:226 | 4.17 | 1.0 | 39.2 |  | 6.13 | 12 | A | O2' | a:2410 | 4.45 | 1.0 | 39.0 |  | 8.64 | 13 | G | OP1 | a:411 | 4.64 | 1.0 | 34.6 |  | 9.36 | 14 | A | OP1 | a:226 | 4.69 | 1.0 | 35.4 |  | 6.71 | 15 | A | O5' | a:2411 | 4.69 | 1.0 | 31.9 |  | 10.31 | 16 | G | O5' | a:411 | 4.79 | 1.0 | 32.8 |  | 9.59 |

  
**1^st^ shell coordination angles and distances (*trans* in "blue"; *cys* in "green"; deviations in "orange"):**   
• 1-2(W/OP2): 128; 1-3(W/OP2): 91; 1-4(W/W): 89; 2-3(OP2/OP2): 75; 2-4(OP2/W): 81; 3-4(OP2/W): 149;   
• 1-2(W/OP2): 4.8; 1-3(W/OP2): 3.9; 1-4(W/W): 3.9; 2-3(OP2/OP2): 3.5; 2-4(OP2/W): 3.8; 3-4(OP2/W): 5.7;   
  
**4.14 ± 1.26: the average coodination distance of the 1^st^ six ligands could correspond to an hexahydrated Mg^2+^.**  
**8: Number of potential Water contacts**   
**2 : Number of second shell contacts involving water or ions**   
**25%: Percentage of satisfied second shell Water contacts**   
• Contact(s): /O_w_/OP2/OP2/  
• Contact code: 2O_ph_O_w_  
• Contact(s) in the exclusion zone: /O_w_/  
→ Coordination < 6 for K ←  
Go to ToC

### K:a:3249

|  | Res. type | Atom | Residue | Distance | Occupancy | B-factor | Closest atom(s) to a 1^st^ shell water | EM\_dens. |
| --- | --- | --- | --- | --- | --- | --- | --- | --- |
| **Ion** | | | |  | 1.0 | 30.0 |  | 25.91 |
 1 | Water 1 | | a:7454 | 2.75 | 1.0 | 45.0 |  | 4.92 | 2 | A | OP2 | a:195 | 2.90 | 1.0 | 27.8 |  | 12.05 | 3 | A | O3' | a:196 | 2.91 | 1.0 | 25.2 |  | 13.34 | 4 | A | O2' | a:196 | 2.97 | 1.0 | 27.0 |  | 13.82 | 5 | A | OP1 | a:197 | 3.18 | 1.0 | 22.5 |  | 17.42 | 6 | Water 2 | | a:7619 | 3.54 | 1.0 | 22.6 | 2.16 [MG:MG]a:3088 | 15.37 | 7 | Water 3 | | a:6019 | 3.78 | 1.0 | 20.2 | 2.11 [MG:MG]a:3088 | 17.41 | 8 | A | O5' | a:197 | 4.10 | 1.0 | 20.0 |  | 16.61 | 9 | Water 4 | | a:9193 | 4.16 | 1.0 | 41.6 |  | 5.29 | 10 | **MG** | MG | a:3088 | 4.29 | 1.0 | 24.4 |  | 27.54 | 11 | Water 5 | | a:9713 | 4.41 | 1.0 | 42.5 |  | 5.26 | 12 | G | OP2 | a:194 | 4.56 | 1.0 | 30.5 |  | 11.62 | 13 | Water 6 | | a:4837 | 4.61 | 1.0 | 36.8 | 2.66 [A:OP1]a:195 | 8.54 | 14 | A | OP1 | a:195 | 4.65 | 1.0 | 30.8 |  | 11.16 | 15 | G | OP1 | a:194 | 4.71 | 1.0 | 30.2 |  | 10.87 |

  
**1^st^ shell coordination angles and distances (*trans* in "blue"; *cys* in "green"; deviations in "orange"):**   
• 1-2(W/OP2): 170; 1-3(W/O3'): 115; 1-4(W/O2'): 73; 1-5(W/OP1): 116; 2-3(OP2/O3'): 74; 2-4(OP2/O2'): 111; 2-5(OP2/OP1): 73; 3-4(O3'/O2'): 55; 3-5(O3'/OP1): 47; 4-5(O2'/OP1): 96;   
• 1-2(W/OP2): 5.6; 1-3(W/O3'): 4.8; 1-4(W/O2'): 3.4; 1-5(W/OP1): 5.0; 2-3(OP2/O3'): 3.5; 2-4(OP2/O2'): 4.8; 2-5(OP2/OP1): 3.6; 3-4(O3'/O2'): 2.7; 3-5(O3'/OP1): 2.4; 4-5(O2'/OP1): 4.6;   
  
**4.36 ± 1.36: the average coodination distance of the 1^st^ six ligands could correspond to an hexahydrated Mg^2+^.**  
**6: Number of potential Water contacts**   
**2 : Number of second shell contacts involving water or ions**   
**33%: Percentage of satisfied second shell Water contacts**   
• Contact(s): /O2'/O3'/O_w_/OP1/OP2/  
• Contact code: 2O_ph_2O_r_O_w_  
→ Coordination < 6 for K ←  
Go to ToC

### K:a:3250

|  | Res. type | Atom | Residue | Distance | Occupancy | B-factor | Closest atom(s) to a 1^st^ shell water | EM\_dens. |
| --- | --- | --- | --- | --- | --- | --- | --- | --- |
| **Ion** | | | |  | 1.0 | 30.0 |  | 13.51 |
 1 | A | N7 | a:515 | 2.90 | 1.0 | 18.5 |  | 17.32 | 2 | Water 1 | | a:8984 | 2.92 | 1.0 | 37.3 | 2.44 [MG:MG]a:3158 | 6.59 | 3 | U | N3 | a:25 | 3.00 | 1.0 | 18.6 |  | 16.25 | 4 | **MG** | MG | a:3158 | 3.31 | 1.0 | 44.2 |  | 7.82 | 5 | A | N7 | a:514 | 3.42 | 1.0 | 16.0 |  | 16.21 | 6 | A | N6 | a:515 | 3.61 | 1.0 | 17.7 |  | 14.89 | 7 | U | O2 | a:25 | 3.64 | 1.0 | 30.2 |  | 13.27 | 8 | C | N4 | a:516 | 3.73 | 1.0 | 18.2 |  | 12.72 | 9 | U | O4 | a:25 | 3.76 | 1.0 | 26.3 |  | 13.10 | 10 | G | O6 | a:24 | 3.88 | 1.0 | 26.3 |  | 13.58 | 11 | C | N3 | a:516 | 3.92 | 1.0 | 19.2 |  | 14.75 | 12 | G | N3 | a:26 | 3.96 | 1.0 | 19.6 |  | 16.52 |

  
**1^st^ shell coordination angles and distances (*trans* in "blue"; *cys* in "green"; deviations in "orange"):**   
• 1-2(N7/W): 103; 1-3(N7/N3): 125; 2-3(W/N3): 124;   
• 1-2(N7/W): 4.6; 1-3(N7/N3): 5.2; 2-3(W/N3): 5.2;   
  
**4.42 ± 1.33: the average coodination distance of the 1^st^ six ligands could correspond to an hexahydrated Mg^2+^.**  
**2: Number of potential Water contacts**   
**1 : Number of second shell contacts involving water or ions**   
**50%: Percentage of satisfied second shell Water contacts**   
• Contact(s): /N3_U_/N7/O_w_/  
• Contact code: N_b_O_w_Anion  
→ Coordination < 6 for K ←  
Go to ToC

### K:a:3251

|  | Res. type | Atom | Residue | Distance | Occupancy | B-factor | Closest atom(s) to a 1^st^ shell water | EM\_dens. |
| --- | --- | --- | --- | --- | --- | --- | --- | --- |
| **Ion** | | | |  | 1.0 | 30.0 |  | 31.56 |
 1 | G | O6 | a:2689 | 2.73 | 1.0 | 24.2 |  | 15.82 | 2 | U | O4 | a:2688 | 2.79 | 1.0 | 28.3 |  | 14.64 | 3 | C | O2 | a:2685 | 2.85 | 1.0 | 24.1 |  | 14.35 | 4 | Water 1 | | a:4304 | 3.00 | 1.0 | 30.4 |  | 13.18 | 5 | Water 2 | | a:7256 | 3.01 | 1.0 | 31.6 | 2.82 [G:N7]a:2689 | 12.03 | 6 | Water 3 | | a:6313 | 3.27 | 1.0 | 31.5 |  | 9.25 | 7 | U | O4 | a:2728 | 3.28 | 1.0 | 22.4 |  | 14.73 | 8 | C | O2' | a:2685 | 3.57 | 1.0 | 30.3 |  | 9.01 | 9 | G | N7 | a:2689 | 4.07 | 1.0 | 18.4 |  | 14.06 | 10 | C | N3 | a:2685 | 4.35 | 1.0 | 15.8 |  | 16.35 | 11 | C | N1 | a:2685 | 4.45 | 1.0 | 19.3 |  | 13.63 | 12 | Water 4 | | a:7357 | 4.77 | 1.0 | 28.7 |  | 13.12 |

  
**1^st^ shell coordination angles and distances (*trans* in "blue"; *cys* in "green"; deviations in "orange"):**   
• 1-2(O6/O4): 77; 1-3(O6/O2): 110; 1-4(O6/W): 170; 1-5(O6/W): 86; 2-3(O4/O2): 83; 2-4(O4/W): 105; 2-5(O4/W): 72; 3-4(O2/W): 80; 3-5(O2/W): 147; 4-5(W/W): 86;   
• 1-2(O6/O4): 3.4; 1-3(O6/O2): 4.6; 1-4(O6/W): 5.7; 1-5(O6/W): 3.9; 2-3(O4/O2): 3.8; 2-4(O4/W): 4.6; 2-5(O4/W): 3.4; 3-4(O2/W): 3.8; 3-5(O2/W): 5.6; 4-5(W/W): 4.1;   
  
**4.08 ± 1.22: the average coodination distance of the 1^st^ six ligands could correspond to an hexahydrated Mg^2+^.**  
**6: Number of potential Water contacts**   
**1 : Number of second shell contacts involving water or ions**   
**17%: Percentage of satisfied second shell Water contacts**   
• Contact(s): /O2/O4/O6/O_w_/O_w_/  
• Contact code: 3O_b_2O_w_  
→ Coordination < 6 for K ←  
Go to ToC

### K:a:3252

|  | Res. type | Atom | Residue | Distance | Occupancy | B-factor | Closest atom(s) to a 1^st^ shell water | EM\_dens. |
| --- | --- | --- | --- | --- | --- | --- | --- | --- |
| **Ion** | | | |  | 1.0 | 30.0 |  | 32.46 |
 1 | Water 1 | | a:6313 | 2.70 | 1.0 | 31.5 |  | 9.25 | 2 | Water 2 | | a:6328 | 2.71 | 1.0 | 35.2 | 2.76 [U:OP2]a:2724 // 2.79 [A:OP2]a:2725 | 9.94 | 3 | G | O6 | a:2690 | 2.81 | 1.0 | 22.1 |  | 16.58 | 4 | Water 3 | | a:5327 | 2.85 | 1.0 | 28.0 | 2.70 [G:N7]a:2726 | 12.66 | 5 | U | O4 | a:2691 | 2.88 | 1.0 | 21.4 |  | 17.66 | 6 | G | O6 | a:2726 | 3.03 | 1.0 | 23.3 |  | 15.99 | 7 | Water 4 | | a:5087 | 3.19 | 1.0 | 32.7 | 2.68 [G:N7]a:2690 | 10.44 | 8 | G | N2 | a:2692 | 3.74 | 1.0 | 17.6 |  | 15.47 | 9 | A | OP2 | a:2725 | 4.19 | 1.0 | 33.3 |  | 10.46 | 10 | Water 5 | | a:7399 | 4.26 | 1.0 | 30.8 |  | 10.33 | 11 | G | N7 | a:2726 | 4.46 | 1.0 | 16.8 |  | 16.52 | 12 | G | N7 | a:2690 | 4.55 | 1.0 | 18.3 |  | 15.31 | 13 | U | OP2 | a:2724 | 4.60 | 1.0 | 35.2 |  | 11.13 | 14 | Water 6 | | a:7202 | 4.80 | 1.0 | 32.6 |  | 7.60 |

  
**1^st^ shell coordination angles and distances (*trans* in "blue"; *cys* in "green"; deviations in "orange"):**   
• 1-2(W/W): 95; 1-3(W/O6): 63; 1-4(W/W): 92; 1-5(W/O4): 125; 1-6(W/O6): 120; 1-7(W/W): 60; 2-3(W/O6): 150; 2-4(W/W): 83; 2-5(W/O4): 99; 2-6(W/O6): 141; 2-7(W/W): 83; 3-4(O6/W): 116; 3-5(O6/O4): 80; 3-6(O6/O6): 69; 3-7(O6/W): 68; 4-5(W/O4): 143; 4-6(W/O6): 79; 4-7(W/W): 147; 5-6(O4/O6): 77; 5-7(O4/W): 69; 6-7(O6/W): 128;   
• 1-2(W/W): 4.0; 1-3(W/O6): 2.9; 1-4(W/W): 4.0; 1-5(W/O4): 5.0; 1-6(W/O6): 5.0; 1-7(W/W): 3.0; 2-3(W/O6): 5.3; 2-4(W/W): 3.7; 2-5(W/O4): 4.3; 2-6(W/O6): 5.4; 2-7(W/W): 3.9; 3-4(O6/W): 4.8; 3-5(O6/O4): 3.6; 3-6(O6/O6): 3.3; 3-7(O6/W): 3.4; 4-5(W/O4): 5.4; 4-6(W/O6): 3.7; 4-7(W/W): 5.8; 5-6(O4/O6): 3.7; 5-7(O4/W): 3.4; 6-7(O6/W): 5.6;   
  
**3.99 ± 1.21: the average coodination distance of the 1^st^ six ligands could correspond to an hexahydrated Mg^2+^.**  
**8: Number of potential Water contacts**   
**4 : Number of second shell contacts involving water or ions**   
**50%: Percentage of satisfied second shell Water contacts**   
• Contact(s): /O4/O6/O6/O_w_/O_w_/O_w_/O_w_/  
• Contact code: 3O_b_4O_w_  
Go to ToC

### K:a:3253

|  | Res. type | Atom | Residue | Distance | Occupancy | B-factor | Closest atom(s) to a 1^st^ shell water | EM\_dens. |
| --- | --- | --- | --- | --- | --- | --- | --- | --- |
| **Ion** | | | |  | 1.0 | 30.0 |  | 37.84 |
 1 | A | OP2 | a:1256 | 2.56 | 1.0 | 22.1 |  | 15.61 | 2 | A | OP1 | a:1255 | 2.77 | 1.0 | 30.8 |  | 14.89 | 3 | G | N7 | a:1258 | 2.88 | 1.0 | 15.6 |  | 17.90 | 4 | Water 1 | | a:6572 | 3.02 | 1.0 | 26.6 |  | 12.24 | 5 | Water 2 | | a:7495 | 3.06 | 1.0 | 28.5 |  | 13.89 | 6 | Water 3 | | a:9516 | 3.15 | 1.0 | 23.6 |  | 15.86 | 7 | Water 4 | | a:3745 | 3.32 | 1.0 | 22.3 | 2.69 [G:O6]a:1258 | 16.37 | 8 | ARG | NH2 | p:33 | 4.17 | 1.0 | 21.9 |  | 16.27 | 9 | A | O3' | a:1255 | 4.47 | 1.0 | 24.2 |  | 15.01 | 10 | G | O6 | a:1258 | 4.57 | 1.0 | 21.6 |  | 21.26 | 11 | A | OP1 | a:1256 | 4.57 | 1.0 | 26.1 |  | 12.62 | 12 | A | O5' | a:1255 | 4.65 | 1.0 | 29.4 |  | 12.60 |

  
**1^st^ shell coordination angles and distances (*trans* in "blue"; *cys* in "green"; deviations in "orange"):**   
• 1-2(OP2/OP1): 104; 1-3(OP2/N7): 80; 1-4(OP2/W): 72; 1-5(OP2/W): 62; 1-6(OP2/W): 127; 2-3(OP1/N7): 85; 2-4(OP1/W): 85; 2-5(OP1/W): 158; 2-6(OP1/W): 126; 3-4(N7/W): 147; 3-5(N7/W): 108; 3-6(N7/W): 86; 4-5(W/W): 75; 4-6(W/W): 125; 5-6(W/W): 74;   
• 1-2(OP2/OP1): 4.2; 1-3(OP2/N7): 3.5; 1-4(OP2/W): 3.3; 1-5(OP2/W): 2.9; 1-6(OP2/W): 5.1; 2-3(OP1/N7): 3.8; 2-4(OP1/W): 3.9; 2-5(OP1/W): 5.7; 2-6(OP1/W): 5.3; 3-4(N7/W): 5.7; 3-5(N7/W): 4.8; 3-6(N7/W): 4.1; 4-5(W/W): 3.7; 4-6(W/W): 5.5; 5-6(W/W): 3.7;   
  
**4.15 ± 1.31: the average coodination distance of the 1^st^ six ligands could correspond to an hexahydrated Mg^2+^.**  
**8: Number of potential Water contacts**   
**1 : Number of second shell contacts involving water or ions**   
**12%: Percentage of satisfied second shell Water contacts**   
• Contact(s): /N7/O_w_/O_w_/O_w_/OP1/  
• Contact code: O_ph_N_b_3O_w_  
• Contact(s) in the exclusion zone: /OP2/  
→ Coordination < 6 for K ←  
Go to ToC

### K:a:3254

|  | Res. type | Atom | Residue | Distance | Occupancy | B-factor | Closest atom(s) to a 1^st^ shell water | EM\_dens. |
| --- | --- | --- | --- | --- | --- | --- | --- | --- |
| **Ion** | | | |  | 1.0 | 30.0 |  | 30.71 |
 1 | A | N1 | a:945 | 2.83 | 1.0 | 21.2 |  | 16.08 | 2 | G | O6 | a:836 | 2.97 | 1.0 | 29.4 |  | 15.99 | 3 | Water 1 | | a:8641 | 3.04 | 1.0 | 31.1 |  | 11.24 | 4 | G | N7 | a:836 | 3.08 | 1.0 | 20.7 |  | 16.42 | 5 | A | N7 | a:835 | 3.17 | 1.0 | 19.3 |  | 15.63 | 6 | C | N4 | a:837 | 3.18 | 1.0 | 21.4 |  | 13.39 | 7 | C | N3 | a:946 | 3.28 | 1.0 | 18.5 |  | 15.78 | 8 | C | N4 | a:946 | 3.41 | 1.0 | 18.1 |  | 14.87 | 9 | Water 2 | | a:9044 | 4.01 | 1.0 | 37.7 |  | 5.20 | 10 | C | O2 | a:946 | 4.48 | 1.0 | 30.7 |  | 14.31 | 11 | G | O6 | a:824 | 4.56 | 1.0 | 28.1 |  | 14.44 |

  
**1^st^ shell coordination angles and distances (*trans* in "blue"; *cys* in "green"; deviations in "orange"):**   
• 1-2(N1/O6): 122; 1-3(N1/W): 91; 1-4(N1/N7): 147; 1-5(N1/N7): 131; 1-6(N1/N4): 73; 2-3(O6/W): 145; 2-4(O6/N7): 60; 2-5(O6/N7): 93; 2-6(O6/N4): 64; 3-4(W/N7): 85; 3-5(W/N7): 72; 3-6(W/N4): 121; 4-5(N7/N7): 79; 4-6(N7/N4): 81; 5-6(N7/N4): 155;   
• 1-2(N1/O6): 5.1; 1-3(N1/W): 4.2; 1-4(N1/N7): 5.7; 1-5(N1/N7): 5.5; 1-6(N1/N4): 3.6; 2-3(O6/W): 5.7; 2-4(O6/N7): 3.0; 2-5(O6/N7): 4.4; 2-6(O6/N4): 3.3; 3-4(W/N7): 4.1; 3-5(W/N7): 3.7; 3-6(W/N4): 5.4; 4-5(N7/N7): 4.0; 4-6(N7/N4): 4.0; 5-6(N7/N4): 6.2;   
  
**4.16 ± 1.22: the average coodination distance of the 1^st^ six ligands could correspond to an hexahydrated Mg^2+^.**  
**2: Number of potential Water contacts**   
**0 : Number of second shell contacts involving water or ions**   
**0%: Percentage of satisfied second shell Water contacts**   
• Contact(s): /N1_A_/N4/N7/N7/O6/O_w_/  
• Contact code: O_b_3N_b_O_w_Anion  
Go to ToC

### K:a:3255

|  | Res. type | Atom | Residue | Distance | Occupancy | B-factor | Closest atom(s) to a 1^st^ shell water | EM\_dens. |
| --- | --- | --- | --- | --- | --- | --- | --- | --- |
| **Ion** | | | |  | 1.0 | 30.0 |  | 8.05 |
 1 | Water 1 | | a:10102 | 2.86 | 1.0 | 49.2 |  | 4.70 | 2 | G | OP2 | a:2462 | 2.87 | 1.0 | 40.0 |  | 8.15 | 3 | G | O6 | a:2494 | 2.90 | 1.0 | 24.8 |  | 11.88 | 4 | Water 2 | | a:6968 | 3.02 | 1.0 | 39.3 | 2.80 [G:N7]a:2494 | 7.50 | 5 | Water 3 | | a:7468 | 3.74 | 1.0 | 36.1 | 2.84 [G:N7]a:2462 | 7.87 | 6 | Water 4 | | a:5098 | 4.01 | 1.0 | 45.3 |  | 3.35 | 7 | G | N7 | a:2462 | 4.04 | 1.0 | 23.5 |  | 12.87 | 8 | G | N7 | a:2494 | 4.07 | 1.0 | 27.2 |  | 10.35 | 9 | Water 5 | | a:8064 | 4.11 | 1.0 | 38.6 |  | 5.40 | 10 | **K** | K | a:3339 | 4.12 | 1.0 | 30.0 |  | 7.10 | 11 | G | O5' | a:2462 | 4.19 | 1.0 | 27.0 |  | 11.74 | 12 | G | OP1 | a:2462 | 4.65 | 1.0 | 30.3 |  | 9.48 |

  
**1^st^ shell coordination angles and distances (*trans* in "blue"; *cys* in "green"; deviations in "orange"):**   
• 1-2(W/OP2): 97; 1-3(W/O6): 113; 1-4(W/W): 57; 2-3(OP2/O6): 114; 2-4(OP2/W): 154; 3-4(O6/W): 80;   
• 1-2(W/OP2): 4.3; 1-3(W/O6): 4.8; 1-4(W/W): 2.8; 2-3(OP2/O6): 4.8; 2-4(OP2/W): 5.7; 3-4(O6/W): 3.8;   
  
**4.58 ± 1.46: the average coodination distance of the 1^st^ six ligands could correspond to an hexahydrated Mg^2+^.**  
**8: Number of potential Water contacts**   
**2 : Number of second shell contacts involving water or ions**   
**25%: Percentage of satisfied second shell Water contacts**   
• Contact(s): /O6/O_w_/O_w_/OP2/  
• Contact code: O_ph_O_b_2O_w_  
→ Coordination < 6 for K ←  
Go to ToC

### K:a:3256

|  | Res. type | Atom | Residue | Distance | Occupancy | B-factor | Closest atom(s) to a 1^st^ shell water | EM\_dens. |
| --- | --- | --- | --- | --- | --- | --- | --- | --- |
| **Ion** | | | |  | 1.0 | 30.0 |  | 16.15 |
 1 | Water 1 | | a:8750 | 2.53 | 1.0 | 46.0 |  | 5.08 | 2 | Water 2 | | a:6155 | 2.88 | 1.0 | 35.8 | 2.75 [U:OP2]a:2526 | 9.33 | 3 | G | O6 | a:2527 | 2.92 | 1.0 | 29.2 |  | 12.32 | 4 | G | N7 | a:2527 | 3.01 | 1.0 | 25.0 |  | 11.52 | 5 | Water 3 | | a:7423 | 3.04 | 1.0 | 38.2 | 2.84 [U:OP2]a:2526 | 7.22 | 6 | Water 4 | | a:4980 | 3.17 | 1.0 | 36.6 |  | 7.03 | 7 | Water 5 | | a:10002 | 3.21 | 1.0 | 43.2 |  | 3.64 | 8 | Water 6 | | a:7931 | 3.31 | 1.0 | 30.3 | 2.88 [A:N1]a:2545 | 10.25 | 9 | Water 7 | | a:10236 | 3.36 | 1.0 | 50.4 |  | 3.11 | 10 | U | O4 | a:2526 | 4.35 | 1.0 | 31.1 |  | 13.06 | 11 | Water 8 | | a:10176 | 4.66 | 1.0 | 42.1 |  | 4.06 | 12 | U | OP2 | a:2526 | 4.68 | 1.0 | 35.9 |  | 10.95 | 13 | A | N1 | a:2545 | 4.73 | 1.0 | 20.2 |  | 13.03 | 14 | Water 9 | | a:5967 | 4.74 | 1.0 | 34.4 |  | 7.38 |

  
**1^st^ shell coordination angles and distances (*trans* in "blue"; *cys* in "green"; deviations in "orange"):**   
• 1-2(W/W): 122; 1-3(W/O6): 66; 1-4(W/N7): 114; 1-5(W/W): 120; 1-6(W/W): 61; 2-3(W/O6): 149; 2-4(W/N7): 124; 2-5(W/W): 68; 2-6(W/W): 136; 3-4(O6/N7): 62; 3-5(O6/W): 138; 3-6(O6/W): 75; 4-5(N7/W): 80; 4-6(N7/W): 68; 5-6(W/W): 74;   
• 1-2(W/W): 4.7; 1-3(W/O6): 3.0; 1-4(W/N7): 4.7; 1-5(W/W): 4.8; 1-6(W/W): 3.0; 2-3(W/O6): 5.6; 2-4(W/N7): 5.2; 2-5(W/W): 3.3; 2-6(W/W): 5.6; 3-4(O6/N7): 3.1; 3-5(O6/W): 5.6; 3-6(O6/W): 3.7; 4-5(N7/W): 3.9; 4-6(N7/W): 3.5; 5-6(W/W): 3.8;   
  
**4.01 ± 1.19: the average coodination distance of the 1^st^ six ligands could correspond to an hexahydrated Mg^2+^.**  
**12: Number of potential Water contacts**   
**3 : Number of second shell contacts involving water or ions**   
**25%: Percentage of satisfied second shell Water contacts**   
• Contact(s): /N7/O6/O_w_/O_w_/O_w_/  
• Contact code: O_b_N_b_3O_w_  
• Contact(s) in the exclusion zone: /O_w_/  
→ Coordination < 6 for K ←  
Go to ToC

### K:a:3257

|  | Res. type | Atom | Residue | Distance | Occupancy | B-factor | Closest atom(s) to a 1^st^ shell water | EM\_dens. |
| --- | --- | --- | --- | --- | --- | --- | --- | --- |
| **Ion** | | | |  | 1.0 | 30.0 |  | 21.39 |
 1 | U | O4 | a:2851 | 2.61 | 1.0 | 23.9 |  | 15.48 | 2 | G | O6 | a:2873 | 2.74 | 1.0 | 28.9 |  | 14.97 | 3 | Water 1 | | a:5730 | 2.91 | 1.0 | 39.6 |  | 5.18 | 4 | Water 2 | | a:4617 | 2.92 | 1.0 | 34.3 | 2.64 [G:N7]a:2873 | 9.67 | 5 | G | O6 | a:2850 | 2.94 | 1.0 | 25.6 |  | 14.84 | 6 | G | N2 | a:2852 | 3.44 | 1.0 | 20.4 |  | 11.61 | 7 | U | N3 | a:2851 | 3.68 | 1.0 | 16.4 |  | 16.41 | 8 | C | N4 | a:2874 | 3.87 | 1.0 | 18.0 |  | 12.31 | 9 | **MG** | MG | a:3029 | 4.01 | 1.0 | 40.0 |  | 9.04 | 10 | G | N7 | a:2873 | 4.38 | 1.0 | 26.6 |  | 12.88 | 11 | Water 3 | | a:8129 | 4.55 | 1.0 | 42.4 | 2.89 [C:N4]a:2874 | 4.62 | 12 | A | OP2 | a:2872 | 4.56 | 1.0 | 39.4 |  | 5.67 | 13 | G | N3 | a:2852 | 4.69 | 1.0 | 23.7 |  | 11.35 |

  
**1^st^ shell coordination angles and distances (*trans* in "blue"; *cys* in "green"; deviations in "orange"):**   
• 1-2(O4/O6): 86; 1-3(O4/W): 91; 1-4(O4/W): 156; 1-5(O4/O6): 76; 2-3(O6/W): 129; 2-4(O6/W): 79; 2-5(O6/O6): 74; 3-4(W/W): 112; 3-5(W/O6): 56; 4-5(W/O6): 116;   
• 1-2(O4/O6): 3.6; 1-3(O4/W): 4.0; 1-4(O4/W): 5.4; 1-5(O4/O6): 3.4; 2-3(O6/W): 5.1; 2-4(O6/W): 3.6; 2-5(O6/O6): 3.4; 3-4(W/W): 4.8; 3-5(W/O6): 2.7; 4-5(W/O6): 5.0;   
  
**4.19 ± 1.31: the average coodination distance of the 1^st^ six ligands could correspond to an hexahydrated Mg^2+^.**  
**4: Number of potential Water contacts**   
**1 : Number of second shell contacts involving water or ions**   
**25%: Percentage of satisfied second shell Water contacts**   
• Contact(s): /O4/O6/O6/O_w_/O_w_/  
• Contact code: 3O_b_2O_w_  
→ Coordination < 6 for K ←  
Go to ToC

### K:a:3258

|  | Res. type | Atom | Residue | Distance | Occupancy | B-factor | Closest atom(s) to a 1^st^ shell water | EM\_dens. |
| --- | --- | --- | --- | --- | --- | --- | --- | --- |
| **Ion** | | | |  | 1.0 | 30.0 |  | 16.80 |
 1 | Water 1 | | a:6028 | 2.73 | 1.0 | 36.5 | 2.74 [G:N7]a:2530 | 7.72 | 2 | G | O6 | a:2529 | 2.90 | 1.0 | 29.2 |  | 11.32 | 3 | G | O6 | a:2530 | 2.91 | 1.0 | 31.5 |  | 10.14 | 4 | U | O4 | a:2541 | 2.92 | 1.0 | 33.0 |  | 9.00 | 5 | G | O6 | a:2540 | 3.29 | 1.0 | 37.3 |  | 8.29 | 6 | C | N4 | a:2531 | 3.74 | 1.0 | 23.6 |  | 9.99 | 7 | U | N3 | a:2541 | 4.02 | 1.0 | 27.5 |  | 10.64 | 8 | C | N4 | a:2542 | 4.10 | 1.0 | 25.6 |  | 10.27 | 9 | G | N7 | a:2530 | 4.18 | 1.0 | 23.2 |  | 11.38 | 10 | Water 2 | | a:9041 | 4.78 | 1.0 | 41.8 | 3.00 [C:N4]a:2531 | 5.32 |

  
**1^st^ shell coordination angles and distances (*trans* in "blue"; *cys* in "green"; deviations in "orange"):**   
• 1-2(W/O6): 77; 1-3(W/O6): 87; 1-4(W/O4): 163; 2-3(O6/O6): 82; 2-4(O6/O4): 96; 3-4(O6/O4): 76;   
• 1-2(W/O6): 3.5; 1-3(W/O6): 3.9; 1-4(W/O4): 5.6; 2-3(O6/O6): 3.8; 2-4(O6/O4): 4.3; 3-4(O6/O4): 3.6;   
  
**4.44 ± 1.41: the average coodination distance of the 1^st^ six ligands could correspond to an hexahydrated Mg^2+^.**  
**2: Number of potential Water contacts**   
**1 : Number of second shell contacts involving water or ions**   
**50%: Percentage of satisfied second shell Water contacts**   
• Contact(s): /O4/O6/O6/O_w_/  
• Contact code: 3O_b_O_w_  
→ Coordination < 6 for K ←  
Go to ToC

### K:a:3259

|  | Res. type | Atom | Residue | Distance | Occupancy | B-factor | Closest atom(s) to a 1^st^ shell water | EM\_dens. |
| --- | --- | --- | --- | --- | --- | --- | --- | --- |
| **Ion** | | | |  | 1.0 | 30.0 |  | 11.88 |
 1 | G | O6 | a:1168 | 2.73 | 1.0 | 35.4 |  | 10.15 | 2 | U | O4 | a:1185 | 2.99 | 1.0 | 43.5 |  | 6.45 | 3 | G | O6 | a:1184 | 3.04 | 1.0 | 48.8 |  | 5.94 | 4 | A | N6 | a:1167 | 3.38 | 1.0 | 23.0 |  | 11.61 | 5 | C | N4 | a:1169 | 3.82 | 1.0 | 32.2 |  | 7.95 | 6 | U | N3 | a:1185 | 3.87 | 1.0 | 30.0 |  | 9.03 | 7 | G | N7 | a:1168 | 4.38 | 1.0 | 29.9 |  | 9.85 | 8 | Water 1 | | a:5082 | 4.54 | 1.0 | 36.6 |  | 5.04 | 9 | Water 2 | | a:9388 | 4.57 | 1.0 | 41.7 |  | 3.24 | 10 | A | N1 | a:1167 | 4.70 | 1.0 | 24.0 |  | 11.60 | 11 | U | O4 | a:1186 | 4.74 | 1.0 | 34.2 |  | 9.96 | 12 | U | O4 | a:1183 | 4.78 | 1.0 | 69.9 |  | 2.37 |

  
**1^st^ shell coordination angles and distances (*trans* in "blue"; *cys* in "green"; deviations in "orange"):**   
• 1-2(O6/O4): 82; 1-3(O6/O6): 69; 2-3(O4/O6): 73;   
• 1-2(O6/O4): 3.7; 1-3(O6/O6): 3.3; 2-3(O4/O6): 3.6;   
  
**4.79 ± 1.56: the average coodination distance of the 1^st^ six ligands could correspond to an hexahydrated Mg^2+^.**  
**2: Number of potential Water contacts**   
**0 : Number of second shell contacts involving water or ions**   
**0%: Percentage of satisfied second shell Water contacts**   
• Contact(s): /O4/O6/O6/  
• Contact code: 3O_b_  
→ Coordination < 6 for K ←  
Go to ToC

### K:a:3260

|  | Res. type | Atom | Residue | Distance | Occupancy | B-factor | Closest atom(s) to a 1^st^ shell water | EM\_dens. |
| --- | --- | --- | --- | --- | --- | --- | --- | --- |
| **Ion** | | | |  | 1.0 | 30.0 |  | 10.91 |
 1 | Water 1 | | a:4806 | 2.75 | 1.0 | 44.9 | 2.66 [G:N7]a:2225 | 4.73 | 2 | G | O6 | a:2225 | 2.86 | 1.0 | 38.7 |  | 6.79 | 3 | U | O4 | a:2206 | 2.89 | 1.0 | 36.3 |  | 7.92 | 4 | U | O4 | a:2224 | 3.15 | 1.0 | 44.0 |  | 6.17 | 5 | G | O6 | a:2205 | 3.66 | 1.0 | 34.7 |  | 9.34 | 6 | C | N4 | a:2226 | 3.69 | 1.0 | 37.3 |  | 7.96 | 7 | U | N3 | a:2206 | 4.01 | 1.0 | 29.6 |  | 11.15 | 8 | Water 2 | | a:6223 | 4.01 | 1.0 | 40.0 |  | 5.58 | 9 | G | N7 | a:2225 | 4.14 | 1.0 | 36.7 |  | 6.91 | 10 | Water 3 | | a:9296 | 4.68 | 1.0 | 42.8 |  | 3.50 |

  
**1^st^ shell coordination angles and distances (*trans* in "blue"; *cys* in "green"; deviations in "orange"):**   
• 1-2(W/O6): 87; 1-3(W/O4): 163; 1-4(W/O4): 71; 2-3(O6/O4): 76; 2-4(O6/O4): 79; 3-4(O4/O4): 108;   
• 1-2(W/O6): 3.9; 1-3(W/O4): 5.6; 1-4(W/O4): 3.4; 2-3(O6/O4): 3.6; 2-4(O6/O4): 3.8; 3-4(O4/O4): 4.9;   
  
**4.50 ± 1.42: the average coodination distance of the 1^st^ six ligands could correspond to an hexahydrated Mg^2+^.**  
**4: Number of potential Water contacts**   
**1 : Number of second shell contacts involving water or ions**   
**25%: Percentage of satisfied second shell Water contacts**   
• Contact(s): /O4/O4/O6/O_w_/  
• Contact code: 3O_b_O_w_  
→ Coordination < 6 for K ←  
Go to ToC

### K:a:3261

|  | Res. type | Atom | Residue | Distance | Occupancy | B-factor | Closest atom(s) to a 1^st^ shell water | EM\_dens. |
| --- | --- | --- | --- | --- | --- | --- | --- | --- |
| **Ion** | | | |  | 1.0 | 30.0 |  | 15.97 |
 1 | Water 1 | | a:4634 | 2.89 | 1.0 | 45.7 |  | 4.49 | 2 | G | O6 | a:2842 | 2.95 | 1.0 | 29.1 |  | 12.36 | 3 | G | N7 | a:2842 | 2.96 | 1.0 | 20.6 |  | 13.02 | 4 | Water 2 | | a:4392 | 2.97 | 1.0 | 32.6 | 2.62 [A:N7]a:2841 | 10.24 | 5 | G | O6 | a:2843 | 3.12 | 1.0 | 34.7 |  | 9.30 | 6 | Water 3 | | a:5710 | 3.14 | 1.0 | 39.8 | 2.72 [G:N7]a:2843 | 5.53 | 7 | U | O4 | a:2882 | 3.27 | 1.0 | 39.4 |  | 8.11 | 8 | G | N7 | a:2843 | 4.02 | 1.0 | 23.2 |  | 10.02 | 9 | Water 4 | | a:8932 | 4.17 | 1.0 | 41.2 | 2.98 [G:O6]a:2881 | 4.08 | 10 | A | N7 | a:2841 | 4.23 | 1.0 | 18.8 |  | 13.63 | 11 | A | N3 | a:2839 | 4.41 | 1.0 | 30.9 |  | 10.05 | 12 | Water 5 | | a:5789 | 4.44 | 1.0 | 37.8 |  | 5.96 | 13 | G | O6 | a:2881 | 4.70 | 1.0 | 36.8 |  | 9.67 | 14 | Water 6 | | a:6308 | 4.77 | 1.0 | 39.9 |  | 5.18 |

  
**1^st^ shell coordination angles and distances (*trans* in "blue"; *cys* in "green"; deviations in "orange"):**   
• 1-2(W/O6): 126; 1-3(W/N7): 65; 1-4(W/W): 82; 1-5(W/O6): 136; 1-6(W/W): 64; 2-3(O6/N7): 63; 2-4(O6/W): 98; 2-5(O6/O6): 68; 2-6(O6/W): 107; 3-4(N7/W): 102; 3-5(N7/O6): 103; 3-6(N7/W): 71; 4-5(W/O6): 142; 4-6(W/W): 145; 5-6(O6/W): 72;   
• 1-2(W/O6): 5.2; 1-3(W/N7): 3.1; 1-4(W/W): 3.8; 1-5(W/O6): 5.6; 1-6(W/W): 3.2; 2-3(O6/N7): 3.1; 2-4(O6/W): 4.5; 2-5(O6/O6): 3.4; 2-6(O6/W): 4.9; 3-4(N7/W): 4.6; 3-5(N7/O6): 4.7; 3-6(N7/W): 3.5; 4-5(W/O6): 5.8; 4-6(W/W): 5.8; 5-6(O6/W): 3.7;   
  
**4.22 ± 1.28: the average coodination distance of the 1^st^ six ligands could correspond to an hexahydrated Mg^2+^.**  
**6: Number of potential Water contacts**   
**2 : Number of second shell contacts involving water or ions**   
**33%: Percentage of satisfied second shell Water contacts**   
• Contact(s): /N7/O6/O6/O_w_/O_w_/O_w_/  
• Contact code: 2O_b_N_b_3O_w_  
Go to ToC

### K:a:3262

|  | Res. type | Atom | Residue | Distance | Occupancy | B-factor | Closest atom(s) to a 1^st^ shell water | EM\_dens. |
| --- | --- | --- | --- | --- | --- | --- | --- | --- |
| **Ion** | | | |  | 1.0 | 30.0 |  | 15.73 |
 1 | G | O6 | a:1987 | 2.74 | 1.0 | 27.6 |  | 13.97 | 2 | U | O4 | a:1986 | 2.78 | 1.0 | 28.6 |  | 13.52 | 3 | G | O6 | a:1772 | 2.98 | 1.0 | 29.1 |  | 13.25 | 4 | Water 1 | | a:7471 | 3.25 | 1.0 | 38.3 |  | 5.88 | 5 | U | O4 | a:1771 | 3.30 | 1.0 | 30.5 |  | 10.60 | 6 | Water 2 | | a:5957 | 3.31 | 1.0 | 40.7 | 2.74 [G:N7]a:1772 | 5.86 | 7 | Water 3 | | a:7616 | 3.33 | 1.0 | 34.6 | 2.85 [G:N7]a:1987 | 8.55 | 8 | Water 4 | | a:6206 | 3.38 | 1.0 | 43.8 | 2.76 [G:O6]a:1984 | 4.85 | 9 | Water 5 | | a:3496 | 3.92 | 1.0 | 40.1 |  | 6.54 | 10 | G | N7 | a:1987 | 4.18 | 1.0 | 19.8 |  | 13.85 | 11 | G | O6 | a:1984 | 4.30 | 1.0 | 25.0 |  | 14.83 | 12 | G | N7 | a:1772 | 4.36 | 1.0 | 21.4 |  | 13.31 |

  
**1^st^ shell coordination angles and distances (*trans* in "blue"; *cys* in "green"; deviations in "orange"):**   
• 1-2(O6/O4): 127; 1-3(O6/O6): 74; 2-3(O4/O6): 73;   
• 1-2(O6/O4): 5.0; 1-3(O6/O6): 3.5; 2-3(O4/O6): 3.4;   
  
**4.18 ± 1.24: the average coodination distance of the 1^st^ six ligands could correspond to an hexahydrated Mg^2+^.**  
**8: Number of potential Water contacts**   
**3 : Number of second shell contacts involving water or ions**   
**38%: Percentage of satisfied second shell Water contacts**   
• Contact(s): /O4/O6/O6/  
• Contact code: 3O_b_  
→ Coordination < 6 for K ←  
Go to ToC

### K:a:3263

|  | Res. type | Atom | Residue | Distance | Occupancy | B-factor | Closest atom(s) to a 1^st^ shell water | EM\_dens. |
| --- | --- | --- | --- | --- | --- | --- | --- | --- |
| **Ion** | | | |  | 1.0 | 30.0 |  | 21.75 |
 1 | U | O4 | a:1843 | 2.67 | 1.0 | 36.0 |  | 10.29 | 2 | Water 1 | | a:9288 | 2.71 | 1.0 | 43.2 |  | 5.97 | 3 | C | O2 | a:1840 | 2.83 | 1.0 | 25.5 |  | 13.27 | 4 | U | O4 | a:1902 | 2.90 | 1.0 | 34.1 |  | 10.65 | 5 | Water 2 | | a:3677 | 3.03 | 1.0 | 48.2 | 2.51 [G:O6]a:1901 | 3.56 | 6 | G | O6 | a:1844 | 3.09 | 1.0 | 33.2 |  | 9.35 | 7 | Water 3 | | a:6502 | 3.14 | 1.0 | 42.3 | 2.77 [G:N7]a:1844 | 5.99 | 8 | Water 4 | | a:9574 | 3.34 | 1.0 | 47.0 |  | 3.53 | 9 | Water 5 | | a:7987 | 4.13 | 1.0 | 34.4 |  | 6.79 | 10 | G | O6 | a:1901 | 4.37 | 1.0 | 34.0 |  | 8.67 | 11 | G | OP2 | a:1842 | 4.43 | 1.0 | 43.7 |  | 6.09 | 12 | C | N1 | a:1840 | 4.54 | 1.0 | 25.6 |  | 12.57 | 13 | **K** | K | a:3264 | 4.60 | 1.0 | 30.0 |  | 22.79 | 14 | C | N3 | a:1840 | 4.63 | 1.0 | 25.2 |  | 12.51 | 15 | G | N7 | a:1844 | 4.67 | 1.0 | 31.4 |  | 9.93 | 16 | G | N7 | a:1842 | 4.73 | 1.0 | 31.7 |  | 8.29 |

  
**1^st^ shell coordination angles and distances (*trans* in "blue"; *cys* in "green"; deviations in "orange"):**   
• 1-2(O4/W): 90; 1-3(O4/O2): 83; 1-4(O4/O4): 137; 1-5(O4/W): 125; 1-6(O4/O6): 74; 1-7(O4/W): 74; 2-3(W/O2): 79; 2-4(W/O4): 113; 2-5(W/W): 119; 2-6(W/O6): 163; 2-7(W/W): 115; 3-4(O2/O4): 68; 3-5(O2/W): 144; 3-6(O2/O6): 94; 3-7(O2/W): 152; 4-5(O4/W): 76; 4-6(O4/O6): 77; 4-7(O4/W): 121; 5-6(W/O6): 76; 5-7(W/W): 52; 6-7(O6/W): 66;   
• 1-2(O4/W): 3.8; 1-3(O4/O2): 3.6; 1-4(O4/O4): 5.2; 1-5(O4/W): 5.1; 1-6(O4/O6): 3.5; 1-7(O4/W): 3.5; 2-3(W/O2): 3.5; 2-4(W/O4): 4.7; 2-5(W/W): 4.9; 2-6(W/O6): 5.7; 2-7(W/W): 4.9; 3-4(O2/O4): 3.2; 3-5(O2/W): 5.6; 3-6(O2/O6): 4.3; 3-7(O2/W): 5.8; 4-5(O4/W): 3.7; 4-6(O4/O6): 3.7; 4-7(O4/W): 5.3; 5-6(W/O6): 3.8; 5-7(W/W): 2.7; 6-7(O6/W): 3.4;   
  
**3.95 ± 1.17: the average coodination distance of the 1^st^ six ligands could correspond to an hexahydrated Mg^2+^.**  
**8: Number of potential Water contacts**   
**2 : Number of second shell contacts involving water or ions**   
**25%: Percentage of satisfied second shell Water contacts**   
• Contact(s): /O2/O4/O4/O6/O_w_/O_w_/O_w_/  
• Contact code: 4O_b_3O_w_  
Go to ToC

### K:a:3264

|  | Res. type | Atom | Residue | Distance | Occupancy | B-factor | Closest atom(s) to a 1^st^ shell water | EM\_dens. |
| --- | --- | --- | --- | --- | --- | --- | --- | --- |
| **Ion** | | | |  | 1.0 | 30.0 |  | 22.79 |
 1 | Water 1 | | a:6846 | 2.64 | 1.0 | 51.0 |  | 4.12 | 2 | G | O6 | a:1842 | 2.86 | 1.0 | 36.7 |  | 9.51 | 3 | C | O4' | a:1840 | 2.92 | 1.0 | 35.7 |  | 10.95 | 4 | C | O2 | a:1840 | 2.95 | 1.0 | 25.5 |  | 13.27 | 5 | A | N1 | a:1903 | 3.06 | 1.0 | 23.4 |  | 12.22 | 6 | G | N7 | a:1841 | 3.07 | 1.0 | 39.5 |  | 7.45 | 7 | G | N7 | a:1842 | 3.10 | 1.0 | 31.7 |  | 8.29 | 8 | G | O6 | a:1841 | 3.12 | 1.0 | 41.8 |  | 8.22 | 9 | Water 2 | | a:9288 | 3.35 | 1.0 | 43.2 | 2.71 [K:K]a:3263 | 5.97 | 10 | C | N1 | a:1840 | 4.07 | 1.0 | 25.6 |  | 12.57 | 11 | U | O4 | a:1843 | 4.41 | 1.0 | 36.0 |  | 10.29 | 12 | C | N3 | a:1840 | 4.46 | 1.0 | 25.2 |  | 12.51 | 13 | **K** | K | a:3263 | 4.60 | 1.0 | 30.0 |  | 21.75 | 14 | A | N3 | a:1903 | 4.73 | 1.0 | 24.4 |  | 13.35 |

  
**1^st^ shell coordination angles and distances (*trans* in "blue"; *cys* in "green"; deviations in "orange"):**   
• 1-2(W/O6): 126; 1-3(W/O4'): 76; 1-4(W/O2): 128; 1-5(W/N1): 64; 1-6(W/N7): 58; 1-7(W/N7): 123; 1-8(W/O6): 58; 2-3(O6/O4'): 150; 2-4(O6/O2): 89; 2-5(O6/N1): 94; 2-6(O6/N7): 129; 2-7(O6/N7): 62; 2-8(O6/O6): 81; 3-4(O4'/O2): 61; 3-5(O4'/N1): 77; 3-6(O4'/N7): 79; 3-7(O4'/N7): 128; 3-8(O4'/O6): 129; 4-5(O2/N1): 78; 4-6(O2/N7): 132; 4-7(O2/N7): 107; 4-8(O2/O6): 169; 5-6(N1/N7): 120; 5-7(N1/N7): 154; 5-8(N1/O6): 99; 6-7(N7/N7): 76; 6-8(N7/O6): 59; 7-8(N7/O6): 71;   
• 1-2(W/O6): 4.9; 1-3(W/O4'): 3.4; 1-4(W/O2): 5.0; 1-5(W/N1): 3.0; 1-6(W/N7): 2.8; 1-7(W/N7): 5.1; 1-8(W/O6): 2.8; 2-3(O6/O4'): 5.6; 2-4(O6/O2): 4.1; 2-5(O6/N1): 4.3; 2-6(O6/N7): 5.4; 2-7(O6/N7): 3.1; 2-8(O6/O6): 3.9; 3-4(O4'/O2): 3.0; 3-5(O4'/N1): 3.7; 3-6(O4'/N7): 3.8; 3-7(O4'/N7): 5.4; 3-8(O4'/O6): 5.4; 4-5(O2/N1): 3.8; 4-6(O2/N7): 5.5; 4-7(O2/N7): 4.9; 4-8(O2/O6): 6.0; 5-6(N1/N7): 5.3; 5-7(N1/N7): 6.0; 5-8(N1/O6): 4.7; 6-7(N7/N7): 3.8; 6-8(N7/O6): 3.0; 7-8(N7/O6): 3.6;   
  
**3.95 ± 1.15: the average coodination distance of the 1^st^ six ligands could correspond to an hexahydrated Mg^2+^.**  
**2: Number of potential Water contacts**   
**0 : Number of second shell contacts involving water or ions**   
**0%: Percentage of satisfied second shell Water contacts**   
• Contact(s): /N1_A_/N7/N7/O2/O4'/O6/O6/O_w_/  
• Contact code: O_r_3O_b_3N_b_O_w_  
Go to ToC

### K:a:3265

|  | Res. type | Atom | Residue | Distance | Occupancy | B-factor | Closest atom(s) to a 1^st^ shell water | EM\_dens. |
| --- | --- | --- | --- | --- | --- | --- | --- | --- |
| **Ion** | | | |  | 1.0 | 30.0 |  | 31.55 |
 1 | C | OP1 | a:2505 | 2.69 | 1.0 | 29.4 |  | 15.09 | 2 | Water 1 | | a:5109 | 2.86 | 1.0 | 24.9 | 2.11 [MG:MG]a:3322 | 17.18 | 3 | 2MA | OP1 | a:2507 | 2.93 | 1.0 | 26.5 |  | 15.71 | 4 | U | OP1 | a:578 | 3.03 | 1.0 | 24.7 |  | 11.99 | 5 | Water 2 | | a:9708 | 3.06 | 1.0 | 39.3 |  | 11.03 | 6 | A | O3' | a:577 | 3.10 | 1.0 | 27.1 |  | 14.85 | 7 | 2MA | O5' | a:2507 | 3.33 | 1.0 | 26.0 |  | 13.16 | 8 | PSU | OP1 | a:2508 | 3.34 | 1.0 | 32.8 |  | 8.78 | 9 | U | O3' | a:2504 | 3.42 | 1.0 | 24.0 |  | 16.64 | 10 | **MG** | MG | a:3322 | 3.64 | 1.0 | 25.5 |  | 25.28 | 11 | PSU | OP2 | a:2508 | 3.87 | 1.0 | 34.5 |  | 9.81 | 12 | Water 3 | | a:4500 | 4.10 | 1.0 | 27.7 |  | 14.91 | 13 | A | O2' | a:577 | 4.25 | 1.0 | 26.0 |  | 14.56 | 14 | U | O5' | a:578 | 4.52 | 1.0 | 22.6 |  | 16.34 | 15 | C | O5' | a:2505 | 4.61 | 1.0 | 23.9 |  | 14.19 | 16 | U | O2' | a:2504 | 4.61 | 1.0 | 33.0 |  | 12.00 | 17 | Water 4 | | a:8860 | 4.72 | 1.0 | 28.3 |  | 13.24 |

  
**1^st^ shell coordination angles and distances (*trans* in "blue"; *cys* in "green"; deviations in "orange"):**   
• 1-2(OP1/W): 179; 1-3(OP1/OP1): 117; 1-4(OP1/OP1): 123; 1-5(OP1/W): 113; 1-6(OP1/O3'): 96; 2-3(W/OP1): 64; 2-4(W/OP1): 58; 2-5(W/W): 67; 2-6(W/O3'): 84; 3-4(OP1/OP1): 56; 3-5(OP1/W): 130; 3-6(OP1/O3'): 102; 4-5(OP1/W): 102; 4-6(OP1/O3'): 47; 5-6(W/O3'): 79;   
• 1-2(OP1/W): 5.5; 1-3(OP1/OP1): 4.8; 1-4(OP1/OP1): 5.0; 1-5(OP1/W): 4.8; 1-6(OP1/O3'): 4.3; 2-3(W/OP1): 3.1; 2-4(W/OP1): 2.8; 2-5(W/W): 3.3; 2-6(W/O3'): 4.0; 3-4(OP1/OP1): 2.8; 3-5(OP1/W): 5.4; 3-6(OP1/O3'): 4.7; 4-5(OP1/W): 4.7; 4-6(OP1/O3'): 2.5; 5-6(W/O3'): 3.9;   
  
**4.06 ± 1.20: the average coodination distance of the 1^st^ six ligands could correspond to an hexahydrated Mg^2+^.**  
**4: Number of potential Water contacts**   
**1 : Number of second shell contacts involving water or ions**   
**25%: Percentage of satisfied second shell Water contacts**   
• Contact(s): /O3'/O_w_/O_w_/OP1/OP1/OP1/  
• Contact code: 3O_ph_O_r_2O_w_  
Go to ToC

### K:a:3266

|  | Res. type | Atom | Residue | Distance | Occupancy | B-factor | Closest atom(s) to a 1^st^ shell water | EM\_dens. |
| --- | --- | --- | --- | --- | --- | --- | --- | --- |
| **Ion** | | | |  | 1.0 | 30.0 |  | 9.94 |
 1 | G | O6 | a:2509 | 2.66 | 1.0 | 42.4 |  | 4.89 | 2 | G | O6 | a:2585 | 2.84 | 1.0 | 33.3 |  | 9.94 | 3 | Water 1 | | a:9512 | 3.06 | 1.0 | 32.9 |  | 9.07 | 4 | C | O2 | a:2614 | 3.21 | 1.0 | 40.6 |  | 7.03 | 5 | G | O2' | a:2580 | 3.25 | 1.0 | 33.8 |  | 11.62 | 6 | Water 2 | | a:7425 | 3.49 | 1.0 | 27.4 | 2.84 [PSU:OP2]a:2584 | 11.86 | 7 | C | O2' | a:2614 | 3.62 | 1.0 | 32.9 |  | 9.28 | 8 | G | N1 | a:2509 | 3.73 | 1.0 | 41.8 |  | 5.25 | 9 | **MG** | MG | a:3304 | 4.25 | 1.0 | 39.7 |  | 15.25 | 10 | C | OP2 | a:2615 | 4.34 | 1.0 | 32.2 |  | 9.34 | 11 | C | N1 | a:2614 | 4.41 | 1.0 | 31.1 |  | 10.57 | 12 | PSU | OP2 | a:2584 | 4.65 | 1.0 | 26.5 |  | 17.47 | 13 | C | O3' | a:2614 | 4.71 | 1.0 | 37.9 |  | 8.87 | 14 | C | N3 | a:2614 | 4.74 | 1.0 | 27.1 |  | 10.00 | 15 | Water 3 | | a:9272 | 4.77 | 1.0 | 38.0 |  | 8.92 |

  
**1^st^ shell coordination angles and distances (*trans* in "blue"; *cys* in "green"; deviations in "orange"):**   
• 1-2(O6/O6): 112; 1-3(O6/W): 75; 2-3(O6/W): 151;   
• 1-2(O6/O6): 4.6; 1-3(O6/W): 3.5; 2-3(O6/W): 5.7;   
  
**4.31 ± 1.31: the average coodination distance of the 1^st^ six ligands could correspond to an hexahydrated Mg^2+^.**  
**4: Number of potential Water contacts**   
**1 : Number of second shell contacts involving water or ions**   
**25%: Percentage of satisfied second shell Water contacts**   
• Contact(s): /O6/O6/O_w_/  
• Contact code: 2O_b_O_w_  
→ Coordination < 6 for K ←  
Go to ToC

### K:a:3267

|  | Res. type | Atom | Residue | Distance | Occupancy | B-factor | Closest atom(s) to a 1^st^ shell water | EM\_dens. |
| --- | --- | --- | --- | --- | --- | --- | --- | --- |
| **Ion** | | | |  | 1.0 | 30.0 |  | 29.60 |
 1 | C | O2 | a:567 | 2.66 | 1.0 | 24.8 |  | 18.06 | 2 | G | OP1 | a:811 | 2.73 | 1.0 | 24.4 |  | 16.07 | 3 | Water 1 | | a:4001 | 2.78 | 1.0 | 26.7 | 2.69 [C:O2']a:567 // 2.82 [A:O2']a:1255 | 13.31 | 4 | U | O2 | a:578 | 2.92 | 1.0 | 25.0 |  | 15.55 | 5 | U | O4' | a:568 | 3.57 | 1.0 | 23.5 |  | 18.35 | 6 | **MG** | MG | a:3354 | 3.74 | 1.0 | 44.6 |  | 7.19 | 7 | G | N3 | a:579 | 3.85 | 1.0 | 15.0 |  | 18.91 | 8 | G | N9 | a:579 | 4.25 | 1.0 | 16.3 |  | 17.92 | 9 | G | O4' | a:579 | 4.28 | 1.0 | 23.7 |  | 18.59 | 10 | U | N1 | a:568 | 4.51 | 1.0 | 14.7 |  | 18.05 | 11 | C | N3 | a:567 | 4.54 | 1.0 | 16.4 |  | 20.41 | 12 | G | O3' | a:810 | 4.55 | 1.0 | 21.0 |  | 17.97 | 13 | U | O2' | a:568 | 4.55 | 1.0 | 23.9 |  | 17.32 | 14 | C | O2' | a:567 | 4.60 | 1.0 | 22.0 |  | 19.46 | 15 | U | O2 | a:568 | 4.63 | 1.0 | 26.6 |  | 16.42 | 16 | A | O2' | a:1255 | 4.68 | 1.0 | 27.5 |  | 16.09 |

  
**1^st^ shell coordination angles and distances (*trans* in "blue"; *cys* in "green"; deviations in "orange"):**   
• 1-2(O2/OP1): 125; 1-3(O2/W): 85; 1-4(O2/O2): 83; 2-3(OP1/W): 56; 2-4(OP1/O2): 130; 3-4(W/O2): 168;   
• 1-2(O2/OP1): 4.8; 1-3(O2/W): 3.7; 1-4(O2/O2): 3.7; 2-3(OP1/W): 2.6; 2-4(OP1/O2): 5.1; 3-4(W/O2): 5.7;   
  
**4.41 ± 1.44: the average coodination distance of the 1^st^ six ligands could correspond to an hexahydrated Mg^2+^.**  
**2: Number of potential Water contacts**   
**2 : Number of second shell contacts involving water or ions**   
**100%: Percentage of satisfied second shell Water contacts**   
• Contact(s): /O2/O2/O_w_/OP1/  
• Contact code: O_ph_2O_b_O_w_  
→ Coordination < 6 for K ←  
Go to ToC

### K:a:3268

|  | Res. type | Atom | Residue | Distance | Occupancy | B-factor | Closest atom(s) to a 1^st^ shell water | EM\_dens. |
| --- | --- | --- | --- | --- | --- | --- | --- | --- |
| **Ion** | | | |  | 1.0 | 30.0 |  | 3.22 |
 1 | G | O6 | a:1720 | 2.62 | 1.0 | 53.1 |  | 5.18 | 2 | G | O6 | a:1721 | 3.27 | 1.0 | 52.6 |  | 5.00 | 3 | U | O4 | a:1744 | 3.41 | 1.0 | 51.5 |  | 2.77 | 4 | C | N4 | a:1743 | 3.84 | 1.0 | 48.3 |  | 4.41 | 5 | G | N7 | a:1720 | 3.99 | 1.0 | 48.9 |  | 4.77 | 6 | G | N2 | a:1745 | 4.00 | 1.0 | 52.3 |  | 3.74 | 7 | G | N1 | a:1745 | 4.06 | 1.0 | 47.8 |  | 4.94 | 8 | U | N3 | a:1744 | 4.09 | 1.0 | 44.1 |  | 6.13 | 9 | U | OP2 | a:1718 | 4.28 | 1.0 | 54.7 |  | 3.14 | 10 | A | OP2 | a:1719 | 4.66 | 1.0 | 58.4 |  | 1.93 | 11 | G | N7 | a:1721 | 4.67 | 1.0 | 49.9 |  | 5.17 |

  
• • **4.88 ± 1.52: the average coodination distance of the 1^st^ six ligands could correspond to an hexahydrated Mg^2+^.**  
**0: Number of potential Water contacts**   
• Contact(s): /O6/  
• Contact code: O_b_  
→ Coordination < 6 for K ←  
→2Fo-Fc density between 1.0 and 4.0 !   
Go to ToC

### K:a:3269

|  | Res. type | Atom | Residue | Distance | Occupancy | B-factor | Closest atom(s) to a 1^st^ shell water | EM\_dens. |
| --- | --- | --- | --- | --- | --- | --- | --- | --- |
| **Ion** | | | |  | 1.0 | 30.0 |  | 17.24 |
 1 | A | N7 | a:2231 | 2.90 | 1.0 | 26.0 |  | 11.91 | 2 | G | O6 | a:2232 | 3.02 | 1.0 | 31.6 |  | 11.31 | 3 | C | OP2 | a:2230 | 3.02 | 1.0 | 34.9 |  | 6.69 | 4 | U | O4 | a:2096 | 3.05 | 1.0 | 37.7 |  | 8.25 | 5 | A | OP2 | a:2231 | 3.15 | 1.0 | 38.9 |  | 7.02 | 6 | C | O5' | a:2230 | 3.44 | 1.0 | 39.1 |  | 6.56 | 7 | Water 1 | | a:5580 | 3.56 | 1.0 | 45.3 |  | 3.37 | 8 | Water 2 | | a:8910 | 3.62 | 1.0 | 44.9 |  | 3.73 | 9 | Water 3 | | a:5772 | 3.65 | 1.0 | 38.9 | 2.93 [G:N7]a:2232 | 4.90 | 10 | G | N7 | a:2232 | 4.15 | 1.0 | 27.4 |  | 11.02 | 11 | A | O3' | a:2229 | 4.48 | 1.0 | 45.7 |  | 6.42 | 12 | A | O5' | a:2231 | 4.65 | 1.0 | 35.5 |  | 8.87 |

  
**1^st^ shell coordination angles and distances (*trans* in "blue"; *cys* in "green"; deviations in "orange"):**   
• 1-2(N7/O6): 85; 1-3(N7/OP2): 131; 1-4(N7/O4): 75; 1-5(N7/OP2): 96; 2-3(O6/OP2): 125; 2-4(O6/O4): 74; 2-5(O6/OP2): 115; 3-4(OP2/O4): 77; 3-5(OP2/OP2): 103; 4-5(O4/OP2): 167;   
• 1-2(N7/O6): 4.0; 1-3(N7/OP2): 5.4; 1-4(N7/O4): 3.6; 1-5(N7/OP2): 4.5; 2-3(O6/OP2): 5.4; 2-4(O6/O4): 3.7; 2-5(O6/OP2): 5.2; 3-4(OP2/O4): 3.8; 3-5(OP2/OP2): 4.8; 4-5(O4/OP2): 6.2;   
  
**4.29 ± 1.28: the average coodination distance of the 1^st^ six ligands could correspond to an hexahydrated Mg^2+^.**  
**4: Number of potential Water contacts**   
**0 : Number of second shell contacts involving water or ions**   
**0%: Percentage of satisfied second shell Water contacts**   
• Contact(s): /N7/O4/O6/OP2/OP2/  
• Contact code: 2O_ph_2O_b_N_b_  
→ Coordination < 6 for K ←  
Go to ToC

### K:a:3270

|  | Res. type | Atom | Residue | Distance | Occupancy | B-factor | Closest atom(s) to a 1^st^ shell water | EM\_dens. |
| --- | --- | --- | --- | --- | --- | --- | --- | --- |
| **Ion** | | | |  | 1.0 | 30.0 |  | 13.26 |
 1 | G | OP1 | a:1684 | 2.63 | 1.0 | 35.9 |  | 8.23 | 2 | Water 1 | | a:9598 | 2.76 | 1.0 | 49.9 |  | 4.34 | 3 | G | O6 | a:1701 | 3.08 | 1.0 | 41.1 |  | 7.90 | 4 | G | O5' | a:1765 | 3.18 | 1.0 | 35.6 |  | 8.91 | 5 | G | O4' | a:1765 | 3.18 | 1.0 | 35.7 |  | 10.05 | 6 | A | O2' | a:1764 | 3.30 | 1.0 | 29.8 |  | 11.22 | 7 | G | OP1 | a:1765 | 3.57 | 1.0 | 36.4 |  | 7.72 | 8 | A | O3' | a:1764 | 3.61 | 1.0 | 30.9 |  | 10.00 | 9 | G | O3' | a:1683 | 3.62 | 1.0 | 37.3 |  | 8.65 | 10 | **MG** | MG | a:3052 | 4.17 | 1.0 | 51.6 |  | 5.95 | 11 | Water 2 | | a:9884 | 4.51 | 1.0 | 44.6 |  | 3.71 | 12 | G | O5' | a:1684 | 4.52 | 1.0 | 30.9 |  | 10.21 |

  
**1^st^ shell coordination angles and distances (*trans* in "blue"; *cys* in "green"; deviations in "orange"):**   
• 1-2(OP1/W): 92; 1-3(OP1/O6): 75; 1-4(OP1/O5'): 128; 1-5(OP1/O4'): 104; 2-3(W/O6): 130; 2-4(W/O5'): 107; 2-5(W/O4'): 160; 3-4(O6/O5'): 119; 3-5(O6/O4'): 66; 4-5(O5'/O4'): 54;   
• 1-2(OP1/W): 3.9; 1-3(OP1/O6): 3.5; 1-4(OP1/O5'): 5.2; 1-5(OP1/O4'): 4.6; 2-3(W/O6): 5.3; 2-4(W/O5'): 4.8; 2-5(W/O4'): 5.9; 3-4(O6/O5'): 5.4; 3-5(O6/O4'): 3.4; 4-5(O5'/O4'): 2.9;   
  
**4.22 ± 1.27: the average coodination distance of the 1^st^ six ligands could correspond to an hexahydrated Mg^2+^.**  
**2: Number of potential Water contacts**   
**0 : Number of second shell contacts involving water or ions**   
**0%: Percentage of satisfied second shell Water contacts**   
• Contact(s): /O4'/O5'/O6/O_w_/OP1/  
• Contact code: O_ph_2O_r_O_b_O_w_  
→ Coordination < 6 for K ←  
Go to ToC

### K:a:3271

|  | Res. type | Atom | Residue | Distance | Occupancy | B-factor | Closest atom(s) to a 1^st^ shell water | EM\_dens. |
| --- | --- | --- | --- | --- | --- | --- | --- | --- |
| **Ion** | | | |  | 1.0 | 30.0 |  | 9.36 |
 1 | U | O4 | a:59 | 2.72 | 1.0 | 35.8 |  | 9.69 | 2 | G | O6 | a:68 | 2.85 | 1.0 | 35.5 |  | 10.25 | 3 | U | O4 | a:67 | 2.90 | 1.0 | 38.2 |  | 8.68 | 4 | G | O6 | a:58 | 3.54 | 1.0 | 34.9 |  | 10.81 | 5 | U | N3 | a:59 | 3.70 | 1.0 | 25.6 |  | 10.24 | 6 | A | N6 | a:74 | 4.32 | 1.0 | 30.6 |  | 8.99 | 7 | G | N7 | a:68 | 4.47 | 1.0 | 29.5 |  | 9.77 |

  
**1^st^ shell coordination angles and distances (*trans* in "blue"; *cys* in "green"; deviations in "orange"):**   
• 1-2(O4/O6): 85; 1-3(O4/O4): 122; 2-3(O6/O4): 72;   
• 1-2(O4/O6): 3.8; 1-3(O4/O4): 4.9; 2-3(O6/O4): 3.4;   
  
**4.08 ± 0.95: the average coodination distance of the 1^st^ six ligands could correspond to an hexahydrated Mg^2+^.**  
**0: Number of potential Water contacts**   
• Contact(s): /O4/O4/O6/  
• Contact code: 3O_b_  
→ Coordination < 6 for K ←  
Go to ToC

### K:a:3272

|  | Res. type | Atom | Residue | Distance | Occupancy | B-factor | Closest atom(s) to a 1^st^ shell water | EM\_dens. |
| --- | --- | --- | --- | --- | --- | --- | --- | --- |
| **Ion** | | | |  | 1.0 | 30.0 |  | 30.18 |
 1 | A | O2' | a:795 | 2.71 | 1.0 | 28.1 |  | 14.77 | 2 | Water 1 | | a:7317 | 2.74 | 1.0 | 27.1 | 2.83 [G:N3]a:777 | 17.41 | 3 | Water 2 | | a:9893 | 2.81 | 1.0 | 45.2 |  | 4.41 | 4 | A | OP2 | a:796 | 2.91 | 1.0 | 25.4 |  | 12.23 | 5 | G | O3' | a:777 | 3.09 | 1.0 | 34.1 |  | 12.62 | 6 | G | O2' | a:777 | 3.12 | 1.0 | 28.4 |  | 14.29 | 7 | G | N7 | a:778 | 3.14 | 1.0 | 26.1 |  | 11.48 | 8 | G | OP2 | a:778 | 3.33 | 1.0 | 32.7 |  | 8.24 | 9 | A | O3' | a:795 | 3.73 | 1.0 | 28.8 |  | 13.29 | 10 | Water 3 | | a:4778 | 4.04 | 1.0 | 28.8 |  | 11.40 | 11 | Water 4 | | a:7695 | 4.05 | 1.0 | 41.1 |  | 5.49 | 12 | G | N3 | a:777 | 4.47 | 1.0 | 15.2 |  | 18.32 | 13 | Water 5 | | a:3855 | 4.63 | 1.0 | 39.8 |  | 5.32 | 14 | G | O5' | a:778 | 4.68 | 1.0 | 38.4 |  | 10.24 |

  
**1^st^ shell coordination angles and distances (*trans* in "blue"; *cys* in "green"; deviations in "orange"):**   
• 1-2(O2'/W): 89; 1-3(O2'/W): 87; 1-4(O2'/OP2): 75; 1-5(O2'/O3'): 156; 1-6(O2'/O2'): 151; 1-7(O2'/N7): 63; 2-3(W/W): 171; 2-4(W/OP2): 106; 2-5(W/O3'): 85; 2-6(W/O2'): 86; 2-7(W/N7): 75; 3-4(W/OP2): 80; 3-5(W/O3'): 96; 3-6(W/O2'): 101; 3-7(W/N7): 96; 4-5(OP2/O3'): 130; 4-6(OP2/O2'): 79; 4-7(OP2/N7): 137; 5-6(O3'/O2'): 52; 5-7(O3'/N7): 93; 6-7(O2'/N7): 142;   
• 1-2(O2'/W): 3.8; 1-3(O2'/W): 3.8; 1-4(O2'/OP2): 3.4; 1-5(O2'/O3'): 5.7; 1-6(O2'/O2'): 5.6; 1-7(O2'/N7): 3.1; 2-3(W/W): 5.5; 2-4(W/OP2): 4.5; 2-5(W/O3'): 3.9; 2-6(W/O2'): 4.0; 2-7(W/N7): 3.6; 3-4(W/OP2): 3.7; 3-5(W/O3'): 4.4; 3-6(W/O2'): 4.6; 3-7(W/N7): 4.4; 4-5(OP2/O3'): 5.4; 4-6(OP2/O2'): 3.8; 4-7(OP2/N7): 5.6; 5-6(O3'/O2'): 2.7; 5-7(O3'/N7): 4.5; 6-7(O2'/N7): 5.9;   
  
**3.97 ± 1.17: the average coodination distance of the 1^st^ six ligands could correspond to an hexahydrated Mg^2+^.**  
**4: Number of potential Water contacts**   
**1 : Number of second shell contacts involving water or ions**   
**25%: Percentage of satisfied second shell Water contacts**   
• Contact(s): /N7/O2'/O2'/O3'/O_w_/O_w_/OP2/  
• Contact code: O_ph_3O_r_N_b_2O_w_  
Go to ToC

### K:a:3273

|  | Res. type | Atom | Residue | Distance | Occupancy | B-factor | Closest atom(s) to a 1^st^ shell water | EM\_dens. |
| --- | --- | --- | --- | --- | --- | --- | --- | --- |
| **Ion** | | | |  | 1.0 | 30.0 |  | 17.21 |
 1 | A | O2' | a:790 | 2.84 | 1.0 | 27.2 |  | 14.60 | 2 | Water 1 | | a:5061 | 2.98 | 1.0 | 40.6 | 2.68 [A:OP2]a:791 | 6.63 | 3 | G | O6 | a:465 | 2.99 | 1.0 | 24.9 |  | 15.27 | 4 | U | O4 | a:792 | 2.99 | 1.0 | 34.4 |  | 9.71 | 5 | U | O4 | a:464 | 3.09 | 1.0 | 26.1 |  | 13.85 | 6 | Water 2 | | a:8790 | 3.31 | 1.0 | 33.0 | 2.96 [G:N1]a:465 | 8.06 | 7 | Water 3 | | a:8411 | 3.36 | 1.0 | 34.5 |  | 8.47 | 8 | G | N1 | a:465 | 3.87 | 1.0 | 15.9 |  | 17.39 | 9 | Water 4 | | a:6769 | 4.01 | 1.0 | 36.2 |  | 7.81 | 10 | Water 5 | | a:7319 | 4.34 | 1.0 | 37.3 |  | 8.09 | 11 | Water 6 | | a:6676 | 4.52 | 1.0 | 36.0 |  | 10.50 | 12 | A | OP2 | a:791 | 4.65 | 1.0 | 32.2 |  | 8.68 |

  
**1^st^ shell coordination angles and distances (*trans* in "blue"; *cys* in "green"; deviations in "orange"):**   
• 1-2(O2'/W): 80; 1-3(O2'/O6): 118; 1-4(O2'/O4): 87; 1-5(O2'/O4): 76; 2-3(W/O6): 111; 2-4(W/O4): 92; 2-5(W/O4): 151; 3-4(O6/O4): 147; 3-5(O6/O4): 69; 4-5(O4/O4): 101;   
• 1-2(O2'/W): 3.7; 1-3(O2'/O6): 5.0; 1-4(O2'/O4): 4.0; 1-5(O2'/O4): 3.6; 2-3(W/O6): 4.9; 2-4(W/O4): 4.3; 2-5(W/O4): 5.9; 3-4(O6/O4): 5.7; 3-5(O6/O4): 3.5; 4-5(O4/O4): 4.7;   
  
**4.24 ± 1.27: the average coodination distance of the 1^st^ six ligands could correspond to an hexahydrated Mg^2+^.**  
**6: Number of potential Water contacts**   
**2 : Number of second shell contacts involving water or ions**   
**33%: Percentage of satisfied second shell Water contacts**   
• Contact(s): /O2'/O4/O4/O6/O_w_/  
• Contact code: O_r_3O_b_O_w_  
→ Coordination < 6 for K ←  
Go to ToC

### K:a:3274

|  | Res. type | Atom | Residue | Distance | Occupancy | B-factor | Closest atom(s) to a 1^st^ shell water | EM\_dens. |
| --- | --- | --- | --- | --- | --- | --- | --- | --- |
| **Ion** | | | |  | 1.0 | 30.0 |  | 13.43 |
 1 | Water 1 | | a:6609 | 2.80 | 1.0 | 40.0 | 2.78 [G:OP2]a:1238 | 7.08 | 2 | G | OP1 | a:1238 | 2.87 | 1.0 | 35.3 |  | 9.76 | 3 | U | OP1 | a:511 | 2.94 | 1.0 | 28.0 |  | 12.34 | 4 | A | O5' | a:505 | 2.98 | 1.0 | 37.0 |  | 5.23 | 5 | Water 2 | | a:8791 | 2.99 | 1.0 | 40.6 |  | 4.79 | 6 | Water 3 | | a:9881 | 3.08 | 1.0 | 42.0 |  | 4.84 | 7 | C | O3' | a:510 | 3.23 | 1.0 | 23.2 |  | 13.26 | 8 | A | O4' | a:505 | 3.30 | 1.0 | 37.0 |  | 8.83 | 9 | A | OP2 | a:505 | 3.77 | 1.0 | 46.3 |  | 5.54 | 10 | G | OP2 | a:1238 | 4.12 | 1.0 | 29.5 |  | 10.44 | 11 | C | O2' | a:510 | 4.44 | 1.0 | 26.4 |  | 13.42 | 12 | Water 4 | | a:3788 | 4.67 | 1.0 | 34.1 |  | 7.60 | 13 | U | O5' | a:511 | 4.68 | 1.0 | 32.8 |  | 12.15 |

  
**1^st^ shell coordination angles and distances (*trans* in "blue"; *cys* in "green"; deviations in "orange"):**   
• 1-2(W/OP1): 77; 1-3(W/OP1): 94; 1-4(W/O5'): 81; 1-5(W/W): 69; 1-6(W/W): 127; 2-3(OP1/OP1): 87; 2-4(OP1/O5'): 157; 2-5(OP1/W): 103; 2-6(OP1/W): 79; 3-4(OP1/O5'): 88; 3-5(OP1/W): 157; 3-6(OP1/W): 131; 4-5(O5'/W): 75; 4-6(O5'/W): 120; 5-6(W/W): 72;   
• 1-2(W/OP1): 3.5; 1-3(W/OP1): 4.2; 1-4(W/O5'): 3.8; 1-5(W/W): 3.3; 1-6(W/W): 5.3; 2-3(OP1/OP1): 4.0; 2-4(OP1/O5'): 5.7; 2-5(OP1/W): 4.6; 2-6(OP1/W): 3.8; 3-4(OP1/O5'): 4.1; 3-5(OP1/W): 5.8; 3-6(OP1/W): 5.5; 4-5(O5'/W): 3.6; 4-6(O5'/W): 5.3; 5-6(W/W): 3.6;   
  
**4.03 ± 1.18: the average coodination distance of the 1^st^ six ligands could correspond to an hexahydrated Mg^2+^.**  
**6: Number of potential Water contacts**   
**1 : Number of second shell contacts involving water or ions**   
**17%: Percentage of satisfied second shell Water contacts**   
• Contact(s): /O5'/O_w_/O_w_/O_w_/OP1/OP1/  
• Contact code: 2O_ph_O_r_3O_w_  
Go to ToC

### K:a:3275

|  | Res. type | Atom | Residue | Distance | Occupancy | B-factor | Closest atom(s) to a 1^st^ shell water | EM\_dens. |
| --- | --- | --- | --- | --- | --- | --- | --- | --- |
| **Ion** | | | |  | 1.0 | 30.0 |  | 31.63 |
 1 | A | OP2 | a:470 | 2.76 | 1.0 | 31.1 |  | 12.59 | 2 | Water 1 | | a:9883 | 2.78 | 1.0 | 36.0 |  | 10.06 | 3 | Water 2 | | a:6121 | 2.84 | 1.0 | 23.6 | 2.75 [G:OP1]a:469 | 14.56 | 4 | Water 3 | | a:7138 | 2.88 | 1.0 | 30.7 | 3.00 [G:O6]a:458 | 12.71 | 5 | G | O4' | a:452 | 2.92 | 1.0 | 33.3 |  | 13.48 | 6 | THR | O | e:53 | 2.95 | 1.0 | 32.3 |  | 12.63 | 7 | A | OP1 | a:470 | 3.19 | 1.0 | 29.1 |  | 11.26 | 8 | Water 4 | | e:411 | 3.73 | 1.0 | 31.7 |  | 9.97 | 9 | Water 5 | | a:8392 | 3.99 | 1.0 | 30.8 |  | 13.32 | 10 | G | O3' | a:469 | 4.27 | 1.0 | 31.6 |  | 11.96 | 11 | Water 6 | | a:3642 | 4.40 | 1.0 | 30.6 |  | 14.28 | 12 | G | O6 | a:458 | 4.61 | 1.0 | 31.9 |  | 12.74 | 13 | G | O5' | a:469 | 4.66 | 1.0 | 22.5 |  | 15.35 | 14 | G | OP1 | a:469 | 4.72 | 1.0 | 28.7 |  | 12.17 | 15 | A | O5' | a:470 | 4.76 | 1.0 | 22.3 |  | 14.35 |

  
**1^st^ shell coordination angles and distances (*trans* in "blue"; *cys* in "green"; deviations in "orange"):**   
• 1-2(OP2/W): 118; 1-3(OP2/W): 104; 1-4(OP2/W): 79; 1-5(OP2/O4'): 82; 1-6(OP2/W): 158; 1-7(OP2/OP1): 50; 2-3(W/W): 79; 2-4(W/W): 161; 2-5(W/O4'): 85; 2-6(W/W): 84; 2-7(W/OP1): 70; 3-4(W/W): 89; 3-5(W/O4'): 164; 3-6(W/W): 81; 3-7(W/OP1): 87; 4-5(W/O4'): 107; 4-6(W/W): 80; 4-7(W/OP1): 124; 5-6(O4'/W): 99; 5-7(O4'/OP1): 87; 6-7(W/OP1): 153;   
• 1-2(OP2/W): 4.8; 1-3(OP2/W): 4.4; 1-4(OP2/W): 3.6; 1-5(OP2/O4'): 3.7; 1-6(OP2/W): 5.6; 1-7(OP2/OP1): 2.5; 2-3(W/W): 3.6; 2-4(W/W): 5.6; 2-5(W/O4'): 3.9; 2-6(W/W): 3.8; 2-7(W/OP1): 3.4; 3-4(W/W): 4.0; 3-5(W/O4'): 5.7; 3-6(W/W): 3.8; 3-7(W/OP1): 4.1; 4-5(W/O4'): 4.7; 4-6(W/W): 3.7; 4-7(W/OP1): 5.4; 5-6(O4'/W): 4.5; 5-7(O4'/OP1): 4.2; 6-7(W/OP1): 6.0;   
  
**4.01 ± 1.21: the average coodination distance of the 1^st^ six ligands could correspond to an hexahydrated Mg^2+^.**  
**10: Number of potential Water contacts**   
**2 : Number of second shell contacts involving water or ions**   
**20%: Percentage of satisfied second shell Water contacts**   
• Contact(s): /O4'/O_back_/O_w_/O_w_/O_w_/OP1/OP2/  
• Contact code: 2O_ph_O_r_O_back_3O_w_  
Go to ToC

### K:a:3276

|  | Res. type | Atom | Residue | Distance | Occupancy | B-factor | Closest atom(s) to a 1^st^ shell water | EM\_dens. |
| --- | --- | --- | --- | --- | --- | --- | --- | --- |
| **Ion** | | | |  | 1.0 | 30.0 |  | 11.91 |
 1 | G | O6 | a:1273 | 2.64 | 1.0 | 32.5 |  | 10.87 | 2 | U | O5' | a:1327 | 2.82 | 1.0 | 31.0 |  | 12.99 | 3 | U | O4' | a:1327 | 3.08 | 1.0 | 33.7 |  | 9.88 | 4 | Water 1 | | a:3946 | 3.14 | 1.0 | 40.1 |  | 3.94 | 5 | Water 2 | | a:9897 | 3.20 | 1.0 | 43.7 |  | 3.07 | 6 | Water 3 | | a:4953 | 3.32 | 1.0 | 39.4 | 2.67 [U:OP2]a:1327 | 9.22 | 7 | Water 4 | | a:4026 | 3.46 | 1.0 | 42.8 | 2.58 [A:O2']a:1618 | 5.42 | 8 | Water 5 | | a:9494 | 3.51 | 1.0 | 39.0 |  | 5.21 | 9 | Water 6 | | a:4543 | 3.54 | 1.0 | 33.1 | 2.64 [G:N7]a:1273 | 11.47 | 10 | U | OP2 | a:1327 | 3.62 | 1.0 | 30.2 |  | 9.52 | 11 | Water 7 | | a:5173 | 3.80 | 1.0 | 35.2 |  | 9.36 | 12 | U | N1 | a:1327 | 4.10 | 1.0 | 23.6 |  | 13.08 | 13 | G | N7 | a:1273 | 4.23 | 1.0 | 26.9 |  | 12.66 | 14 | U | OP1 | a:1327 | 4.63 | 1.0 | 34.7 |  | 11.21 | 15 | A | O2' | a:1618 | 4.78 | 1.0 | 29.1 |  | 10.39 |

  
**1^st^ shell coordination angles and distances (*trans* in "blue"; *cys* in "green"; deviations in "orange"):**   
• 1-2(O6/O5'): 150; 1-3(O6/O4'): 135; 1-4(O6/W): 130; 2-3(O5'/O4'): 59; 2-4(O5'/W): 77; 3-4(O4'/W): 71;   
• 1-2(O6/O5'): 5.3; 1-3(O6/O4'): 5.3; 1-4(O6/W): 5.2; 2-3(O5'/O4'): 2.9; 2-4(O5'/W): 3.7; 3-4(O4'/W): 3.6;   
  
**4.20 ± 1.25: the average coodination distance of the 1^st^ six ligands could correspond to an hexahydrated Mg^2+^.**  
**10: Number of potential Water contacts**   
**2 : Number of second shell contacts involving water or ions**   
**20%: Percentage of satisfied second shell Water contacts**   
• Contact(s): /O4'/O5'/O6/O_w_/  
• Contact code: 2O_r_O_b_O_w_  
→ Coordination < 6 for K ←  
Go to ToC

### K:a:3277

|  | Res. type | Atom | Residue | Distance | Occupancy | B-factor | Closest atom(s) to a 1^st^ shell water | EM\_dens. |
| --- | --- | --- | --- | --- | --- | --- | --- | --- |
| **Ion** | | | |  | 1.0 | 30.0 |  | 31.63 |
 1 | C | OP2 | a:192 | 2.74 | 1.0 | 22.8 |  | 17.54 | 2 | Water 1 | | a:7863 | 2.82 | 1.0 | 35.5 | 2.87 [A:N1]a:203 | 8.47 | 3 | Water 2 | | a:5122 | 2.87 | 1.0 | 23.9 | 2.68 [A:OP2]a:199 | 17.30 | 4 | U | O4 | a:193 | 2.89 | 1.0 | 25.1 |  | 15.02 | 5 | Water 3 | | a:4961 | 2.99 | 1.0 | 22.5 | 2.29 [MG:MG]a:3336 | 16.55 | 6 | U | O4 | a:202 | 3.32 | 1.0 | 26.7 |  | 13.90 | 7 | Water 4 | | a:7731 | 3.42 | 1.0 | 29.6 |  | 10.18 | 8 | Water 5 | | a:9973 | 3.70 | 1.0 | 39.8 |  | 5.43 | 9 | Water 6 | | a:9748 | 3.70 | 1.0 | 25.9 | 2.39 [MG:MG]a:3336 | 15.18 | 10 | C | OP1 | a:192 | 4.19 | 1.0 | 26.0 |  | 15.75 | 11 | Water 7 | | a:9988 | 4.21 | 1.0 | 45.6 |  | 4.02 | 12 | A | N1 | a:203 | 4.25 | 1.0 | 24.6 |  | 13.41 | 13 | **MG** | MG | a:3336 | 4.55 | 1.0 | 26.0 |  | 25.15 | 14 | Water 8 | | a:5911 | 4.57 | 1.0 | 39.3 | 2.74 [U:O4]a:202 | 8.59 | 15 | A | OP2 | a:199 | 4.59 | 1.0 | 26.6 |  | 14.25 | 16 | G | O6 | a:194 | 4.69 | 1.0 | 24.2 |  | 16.46 | 17 | A | OP1 | a:199 | 4.78 | 1.0 | 27.0 |  | 13.58 |

  
**1^st^ shell coordination angles and distances (*trans* in "blue"; *cys* in "green"; deviations in "orange"):**   
• 1-2(OP2/W): 80; 1-3(OP2/W): 108; 1-4(OP2/O4): 85; 1-5(OP2/W): 79; 2-3(W/W): 172; 2-4(W/O4): 110; 2-5(W/W): 114; 3-4(W/O4): 68; 3-5(W/W): 70; 4-5(O4/W): 128;   
• 1-2(OP2/W): 3.6; 1-3(OP2/W): 4.5; 1-4(OP2/O4): 3.8; 1-5(OP2/W): 3.6; 2-3(W/W): 5.7; 2-4(W/O4): 4.7; 2-5(W/W): 4.9; 3-4(W/O4): 3.2; 3-5(W/W): 3.4; 4-5(O4/W): 5.3;   
  
**4.12 ± 1.25: the average coodination distance of the 1^st^ six ligands could correspond to an hexahydrated Mg^2+^.**  
**10: Number of potential Water contacts**   
**3 : Number of second shell contacts involving water or ions**   
**30%: Percentage of satisfied second shell Water contacts**   
• Contact(s): /O4/O_w_/O_w_/O_w_/OP2/  
• Contact code: O_ph_O_b_3O_w_  
→ Coordination < 6 for K ←  
Go to ToC

### K:a:3278

|  | Res. type | Atom | Residue | Distance | Occupancy | B-factor | Closest atom(s) to a 1^st^ shell water | EM\_dens. |
| --- | --- | --- | --- | --- | --- | --- | --- | --- |
| **Ion** | | | |  | 1.0 | 30.0 |  | 22.48 |
 1 | Water 1 | | a:9879 | 2.70 | 1.0 | 44.5 |  | 4.02 | 2 | U | O4 | a:590 | 2.76 | 1.0 | 26.3 |  | 13.87 | 3 | A | O2' | a:672 | 2.91 | 1.0 | 24.8 |  | 14.82 | 4 | A | O3' | a:672 | 3.00 | 1.0 | 23.8 |  | 15.78 | 5 | Water 2 | | a:10080 | 3.02 | 1.0 | 40.0 |  | 6.73 | 6 | Water 3 | | a:4674 | 3.04 | 1.0 | 35.0 |  | 8.86 | 7 | Water 4 | | a:6260 | 3.16 | 1.0 | 44.1 | 2.76 [U:O4]a:591 | 6.01 | 8 | C | OP2 | a:673 | 3.21 | 1.0 | 26.7 |  | 15.72 | 9 | Water 5 | | a:5757 | 3.92 | 1.0 | 39.4 | 2.73 [C:OP2]a:673 | 8.88 | 10 | Water 6 | | a:8950 | 4.35 | 1.0 | 42.4 | 2.99 [C:OP2]a:673 | 6.28 | 11 | Water 7 | | a:7136 | 4.49 | 1.0 | 48.0 |  | 4.92 | 12 | U | O4 | a:591 | 4.50 | 1.0 | 28.4 |  | 13.90 | 13 | Water 8 | | a:5042 | 4.62 | 1.0 | 39.5 |  | 4.97 | 14 | Water 9 | | a:6111 | 4.68 | 1.0 | 40.5 |  | 5.09 |

  
**1^st^ shell coordination angles and distances (*trans* in "blue"; *cys* in "green"; deviations in "orange"):**   
• 1-2(W/O4): 154; 1-3(W/O2'): 79; 1-4(W/O3'): 80; 1-5(W/W): 72; 1-6(W/W): 74; 1-7(W/W): 136; 2-3(O4/O2'): 83; 2-4(O4/O3'): 74; 2-5(O4/W): 126; 2-6(O4/W): 116; 2-7(O4/W): 70; 3-4(O2'/O3'): 56; 3-5(O2'/W): 150; 3-6(O2'/W): 68; 3-7(O2'/W): 128; 4-5(O3'/W): 123; 4-6(O3'/W): 121; 4-7(O3'/W): 143; 5-6(W/W): 98; 5-7(W/W): 73; 6-7(W/W): 85;   
• 1-2(W/O4): 5.3; 1-3(W/O2'): 3.6; 1-4(W/O3'): 3.7; 1-5(W/W): 3.4; 1-6(W/W): 3.4; 1-7(W/W): 5.4; 2-3(O4/O2'): 3.8; 2-4(O4/O3'): 3.5; 2-5(O4/W): 5.1; 2-6(O4/W): 4.9; 2-7(O4/W): 3.4; 3-4(O2'/O3'): 2.8; 3-5(O2'/W): 5.7; 3-6(O2'/W): 3.3; 3-7(O2'/W): 5.5; 4-5(O3'/W): 5.3; 4-6(O3'/W): 5.3; 4-7(O3'/W): 5.8; 5-6(W/W): 4.6; 5-7(W/W): 3.7; 6-7(W/W): 4.2;   
  
**3.97 ± 1.16: the average coodination distance of the 1^st^ six ligands could correspond to an hexahydrated Mg^2+^.**  
**8: Number of potential Water contacts**   
**1 : Number of second shell contacts involving water or ions**   
**12%: Percentage of satisfied second shell Water contacts**   
• Contact(s): /O2'/O3'/O4/O_w_/O_w_/O_w_/O_w_/  
• Contact code: 2O_r_O_b_4O_w_  
Go to ToC

### K:a:3279

|  | Res. type | Atom | Residue | Distance | Occupancy | B-factor | Closest atom(s) to a 1^st^ shell water | EM\_dens. |
| --- | --- | --- | --- | --- | --- | --- | --- | --- |
| **Ion** | | | |  | 1.0 | 30.0 |  | 14.45 |
 1 | Water 1 | | a:5398 | 2.51 | 1.0 | 45.7 | 2.70 [U:O2]a:2445 | 3.43 | 2 | Water 2 | | a:7257 | 2.76 | 1.0 | 35.3 |  | 9.94 | 3 | A | OP1 | a:795 | 2.89 | 1.0 | 34.8 |  | 10.28 | 4 | A | O2' | a:794 | 2.95 | 1.0 | 26.6 |  | 13.38 | 5 | Water 3 | | a:7140 | 3.00 | 1.0 | 42.8 |  | 6.83 | 6 | Water 4 | | a:9155 | 3.36 | 1.0 | 40.8 |  | 5.27 | 7 | A | O2' | a:2075 | 3.47 | 1.0 | 24.6 |  | 16.76 | 8 | Water 5 | | a:5403 | 3.86 | 1.0 | 40.4 |  | 5.62 | 9 | A | O3' | a:794 | 4.05 | 1.0 | 30.1 |  | 13.40 | 10 | A | OP2 | a:795 | 4.10 | 1.0 | 30.5 |  | 12.83 | 11 | U | O2 | a:2445 | 4.21 | 1.0 | 31.5 |  | 14.46 | 12 | Water 6 | | a:5510 | 4.26 | 1.0 | 41.3 |  | 6.92 | 13 | Water 7 | | a:4957 | 4.35 | 1.0 | 41.3 |  | 2.85 | 14 | C | N3 | a:2444 | 4.68 | 1.0 | 18.1 |  | 16.35 |

  
**1^st^ shell coordination angles and distances (*trans* in "blue"; *cys* in "green"; deviations in "orange"):**   
• 1-2(W/W): 89; 1-3(W/OP1): 145; 1-4(W/O2'): 136; 1-5(W/W): 90; 2-3(W/OP1): 118; 2-4(W/O2'): 59; 2-5(W/W): 99; 3-4(OP1/O2'): 79; 3-5(OP1/W): 104; 4-5(O2'/W): 68;   
• 1-2(W/W): 3.7; 1-3(W/OP1): 5.2; 1-4(W/O2'): 5.1; 1-5(W/W): 3.9; 2-3(W/OP1): 4.9; 2-4(W/O2'): 2.8; 2-5(W/W): 4.4; 3-4(OP1/O2'): 3.7; 3-5(OP1/W): 4.7; 4-5(O2'/W): 3.3;   
  
**4.13 ± 1.28: the average coodination distance of the 1^st^ six ligands could correspond to an hexahydrated Mg^2+^.**  
**10: Number of potential Water contacts**   
**1 : Number of second shell contacts involving water or ions**   
**10%: Percentage of satisfied second shell Water contacts**   
• Contact(s): /O2'/O_w_/O_w_/OP1/  
• Contact code: O_ph_O_r_2O_w_  
• Contact(s) in the exclusion zone: /O_w_/  
→ Coordination < 6 for K ←  
Go to ToC

### K:a:3280

|  | Res. type | Atom | Residue | Distance | Occupancy | B-factor | Closest atom(s) to a 1^st^ shell water | EM\_dens. |
| --- | --- | --- | --- | --- | --- | --- | --- | --- |
| **Ion** | | | |  | 1.0 | 30.0 |  | 5.71 |
 1 | G | O6 | a:1859 | 2.61 | 1.0 | 58.8 |  | 2.13 | 2 | Water 1 | | a:6466 | 2.93 | 1.0 | 47.1 |  | 3.20 | 3 | U | O4 | a:1858 | 3.00 | 1.0 | 48.6 |  | 4.15 | 4 | G | N1 | a:1859 | 3.76 | 1.0 | 50.4 |  | 4.91 | 5 | A | OP1 | a:1856 | 4.42 | 1.0 | 39.8 |  | 6.07 | 6 | U | OP2 | a:1857 | 4.46 | 1.0 | 45.0 |  | 4.99 |

  
**1^st^ shell coordination angles and distances (*trans* in "blue"; *cys* in "green"; deviations in "orange"):**   
• 1-2(O6/W): 130; 1-3(O6/O4): 65; 2-3(W/O4): 65;   
• 1-2(O6/W): 5.0; 1-3(O6/O4): 3.0; 2-3(W/O4): 3.2;   
  
**3.53 ± 0.73: the average coodination distance of the 1^st^ six ligands is K^+^ compatible**  
**2: Number of potential Water contacts**   
**0 : Number of second shell contacts involving water or ions**   
**0%: Percentage of satisfied second shell Water contacts**   
• Contact(s): /O4/O6/O_w_/  
• Contact code: 2O_b_O_w_  
→ Coordination < 6 for K ←  
Go to ToC

### K:a:3281

|  | Res. type | Atom | Residue | Distance | Occupancy | B-factor | Closest atom(s) to a 1^st^ shell water | EM\_dens. |
| --- | --- | --- | --- | --- | --- | --- | --- | --- |
| **Ion** | | | |  | 1.0 | 30.0 |  | 5.68 |
 1 | U | O4 | a:2558 | 2.64 | 1.0 | 42.3 |  | 4.76 | 2 | Water 1 | | a:8682 | 2.66 | 1.0 | 43.2 | 2.95 [C:N3]a:2560 | 5.08 | 3 | Water 2 | | a:6153 | 3.00 | 1.0 | 38.3 | 2.93 [U:N3]a:1959 | 4.91 | 4 | U | O4 | a:2559 | 3.26 | 1.0 | 45.5 |  | 3.72 | 5 | C | N3 | a:2560 | 3.33 | 1.0 | 33.1 |  | 11.13 | 6 | C | O2 | a:2560 | 3.33 | 1.0 | 39.7 |  | 6.32 | 7 | U | O4 | a:1959 | 3.43 | 1.0 | 38.8 |  | 8.61 | 8 | U | N3 | a:1959 | 4.22 | 1.0 | 33.5 |  | 10.12 |

  
**1^st^ shell coordination angles and distances (*trans* in "blue"; *cys* in "green"; deviations in "orange"):**   
• 1-2(O4/W): 154; 1-3(O4/W): 58; 2-3(W/W): 115;   
• 1-2(O4/W): 5.2; 1-3(O4/W): 2.8; 2-3(W/W): 4.8;   
  
**4.31 ± 1.36: the average coodination distance of the 1^st^ six ligands could correspond to an hexahydrated Mg^2+^.**  
**4: Number of potential Water contacts**   
**2 : Number of second shell contacts involving water or ions**   
**50%: Percentage of satisfied second shell Water contacts**   
• Contact(s): /O4/O_w_/O_w_/  
• Contact code: O_b_2O_w_  
→ Coordination < 6 for K ←  
Go to ToC

### K:a:3282

|  | Res. type | Atom | Residue | Distance | Occupancy | B-factor | Closest atom(s) to a 1^st^ shell water | EM\_dens. |
| --- | --- | --- | --- | --- | --- | --- | --- | --- |
| **Ion** | | | |  | 1.0 | 30.0 |  | 18.73 |
 1 | Water 1 | | a:10028 | 2.66 | 1.0 | 50.8 |  | 4.87 | 2 | G | O6 | a:1217 | 2.76 | 1.0 | 29.1 |  | 13.12 | 3 | Water 2 | | a:5037 | 2.79 | 1.0 | 32.2 | 2.68 [A:OP2]a:1216 // 2.73 [G:N7]a:1217 | 9.51 | 4 | Water 3 | | a:5740 | 2.80 | 1.0 | 44.8 | 2.73 [A:OP2]a:1215 // 2.93 [A:OP2]a:1216 | 5.69 | 5 | G | O6 | a:1218 | 3.15 | 1.0 | 25.4 |  | 11.80 | 6 | U | O4 | a:1236 | 3.15 | 1.0 | 32.7 |  | 11.55 | 7 | Water 4 | | a:8973 | 3.54 | 1.0 | 46.2 | 2.99 [C:N4]a:1235 | 3.68 | 8 | Water 5 | | a:9392 | 3.58 | 1.0 | 40.9 |  | 4.68 | 9 | G | N7 | a:1217 | 4.18 | 1.0 | 18.1 |  | 14.05 | 10 | A | OP2 | a:1216 | 4.55 | 1.0 | 33.5 |  | 9.50 | 11 | A | OP2 | a:1215 | 4.62 | 1.0 | 34.1 |  | 8.96 | 12 | G | N7 | a:1218 | 4.69 | 1.0 | 22.9 |  | 13.28 | 13 | Water 6 | | a:6841 | 4.71 | 1.0 | 35.9 |  | 6.91 | 14 | G | N3 | a:1237 | 4.73 | 1.0 | 26.7 |  | 12.07 |

  
**1^st^ shell coordination angles and distances (*trans* in "blue"; *cys* in "green"; deviations in "orange"):**   
• 1-2(W/O6): 142; 1-3(W/W): 81; 1-4(W/W): 78; 1-5(W/O6): 80; 1-6(W/O4): 128; 2-3(O6/W): 86; 2-4(O6/W): 130; 2-5(O6/O6): 73; 2-6(O6/O4): 75; 3-4(W/W): 71; 3-5(W/O6): 113; 3-6(W/O4): 151; 4-5(W/O6): 157; 4-6(W/O4): 104; 5-6(O6/O4): 83;   
• 1-2(W/O6): 5.1; 1-3(W/W): 3.5; 1-4(W/W): 3.4; 1-5(W/O6): 3.7; 1-6(W/O4): 5.2; 2-3(O6/W): 3.8; 2-4(O6/W): 5.1; 2-5(O6/O6): 3.5; 2-6(O6/O4): 3.6; 3-4(W/W): 3.2; 3-5(W/O6): 5.0; 3-6(W/O4): 5.7; 4-5(W/O6): 5.8; 4-6(W/O4): 4.7; 5-6(O6/O4): 4.2;   
  
**4.07 ± 1.24: the average coodination distance of the 1^st^ six ligands could correspond to an hexahydrated Mg^2+^.**  
**10: Number of potential Water contacts**   
**5 : Number of second shell contacts involving water or ions**   
**50%: Percentage of satisfied second shell Water contacts**   
• Contact(s): /O4/O6/O6/O_w_/O_w_/O_w_/  
• Contact code: 3O_b_3O_w_  
Go to ToC

### K:a:3283

|  | Res. type | Atom | Residue | Distance | Occupancy | B-factor | Closest atom(s) to a 1^st^ shell water | EM\_dens. |
| --- | --- | --- | --- | --- | --- | --- | --- | --- |
| **Ion** | | | |  | 1.0 | 30.0 |  | 10.74 |
 1 | G | O6 | a:702 | 2.69 | 1.0 | 32.3 |  | 11.32 | 2 | Water 1 | | a:4971 | 2.96 | 1.0 | 44.0 | 2.67 [G:N7]a:702 | 4.31 | 3 | Water 2 | | a:3859 | 3.01 | 1.0 | 45.0 | 2.55 [G:N7]a:703 | 4.84 | 4 | G | O6 | a:703 | 3.13 | 1.0 | 28.9 |  | 9.90 | 5 | Water 3 | | a:6479 | 3.24 | 1.0 | 41.5 |  | 4.31 | 6 | Water 4 | | a:3575 | 3.55 | 1.0 | 40.8 | 2.48 [U:OP2]a:764 | 5.61 | 7 | Water 5 | | a:8356 | 3.83 | 1.0 | 38.9 | 2.92 [U:OP2]a:764 | 4.06 | 8 | G | N7 | a:702 | 3.95 | 1.0 | 26.8 |  | 12.35 | 9 | G | N7 | a:703 | 4.09 | 1.0 | 25.4 |  | 11.62 | 10 | Water 6 | | a:6085 | 4.46 | 1.0 | 40.6 |  | 6.36 | 11 | U | OP2 | a:764 | 4.75 | 1.0 | 30.0 |  | 10.60 |

  
**1^st^ shell coordination angles and distances (*trans* in "blue"; *cys* in "green"; deviations in "orange"):**   
• 1-2(O6/W): 92; 1-3(O6/W): 95; 1-4(O6/O6): 75; 2-3(W/W): 59; 2-4(W/O6): 128; 3-4(W/O6): 72;   
• 1-2(O6/W): 4.1; 1-3(O6/W): 4.2; 1-4(O6/O6): 3.6; 2-3(W/W): 3.0; 2-4(W/O6): 5.5; 3-4(W/O6): 3.6;   
  
**4.39 ± 1.36: the average coodination distance of the 1^st^ six ligands could correspond to an hexahydrated Mg^2+^.**  
**10: Number of potential Water contacts**   
**4 : Number of second shell contacts involving water or ions**   
**40%: Percentage of satisfied second shell Water contacts**   
• Contact(s): /O6/O6/O_w_/O_w_/  
• Contact code: 2O_b_2O_w_  
→ Coordination < 6 for K ←  
Go to ToC

### K:a:3284

|  | Res. type | Atom | Residue | Distance | Occupancy | B-factor | Closest atom(s) to a 1^st^ shell water | EM\_dens. |
| --- | --- | --- | --- | --- | --- | --- | --- | --- |
| **Ion** | | | |  | 1.0 | 30.0 |  | 31.09 |
 1 | Water 1 | | r:215 | 2.62 | 1.0 | 30.1 | 2.72 [GLN:OE1]r:15 | 12.64 | 2 | A | N3 | a:1267 | 2.88 | 1.0 | 15.0 |  | 19.59 | 3 | Water 2 | | r:257 | 2.90 | 1.0 | 33.3 |  | 10.48 | 4 | Water 3 | | a:6832 | 2.91 | 1.0 | 31.7 | 2.79 [A:OP2]a:1266 | 12.67 | 5 | A | O2' | a:1267 | 2.92 | 1.0 | 24.1 |  | 17.82 | 6 | Water 4 | | a:7194 | 2.94 | 1.0 | 25.0 | 2.82 [U:OP2]a:1265 // 2.92 [A:N7]a:1266 | 16.82 | 7 | Water 5 | | z:238 | 3.53 | 1.0 | 36.9 | 2.97 [ARG:NH2]z:17 | 6.72 | 8 | Water 6 | | a:5877 | 3.64 | 1.0 | 36.3 | 2.73 [U:OP2]a:1265 | 6.93 | 9 | Water 7 | | r:239 | 4.33 | 1.0 | 36.6 | 2.92 [GLN:OE1]r:15 | 8.51 | 10 | U | OP2 | a:1265 | 4.33 | 1.0 | 23.2 |  | 16.65 | 11 | A | OP2 | a:1266 | 4.48 | 1.0 | 23.1 |  | 15.55 | 12 | A | N1 | a:1267 | 4.58 | 1.0 | 18.1 |  | 18.78 | 13 | A | O3' | a:1267 | 4.67 | 1.0 | 25.3 |  | 13.67 | 14 | GLN | OE1 | r:15 | 4.71 | 1.0 | 35.6 |  | 11.85 | 15 | A | N7 | a:1266 | 4.78 | 1.0 | 15.2 |  | 18.56 |

  
**1^st^ shell coordination angles and distances (*trans* in "blue"; *cys* in "green"; deviations in "orange"):**   
• 1-2(W/N3): 101; 1-3(W/W): 95; 1-4(W/W): 94; 1-5(W/O2'): 80; 1-6(W/W): 175; 2-3(N3/W): 90; 2-4(N3/W): 136; 2-5(N3/O2'): 78; 2-6(N3/W): 77; 3-4(W/W): 129; 3-5(W/O2'): 165; 3-6(W/W): 90; 4-5(W/O2'): 65; 4-6(W/W): 84; 5-6(O2'/W): 95;   
• 1-2(W/N3): 4.3; 1-3(W/W): 4.1; 1-4(W/W): 4.1; 1-5(W/O2'): 3.6; 1-6(W/W): 5.6; 2-3(N3/W): 4.1; 2-4(N3/W): 5.4; 2-5(N3/O2'): 3.6; 2-6(N3/W): 3.6; 3-4(W/W): 5.3; 3-5(W/O2'): 5.8; 3-6(W/W): 4.1; 4-5(W/O2'): 3.1; 4-6(W/W): 3.9; 5-6(O2'/W): 4.3;   
  
**4.06 ± 1.23: the average coodination distance of the 1^st^ six ligands could correspond to an hexahydrated Mg^2+^.**  
**12: Number of potential Water contacts**   
**6 : Number of second shell contacts involving water or ions**   
**50%: Percentage of satisfied second shell Water contacts**   
• Contact(s): /N3_A_/O2'/O_w_/O_w_/O_w_/O_w_/  
• Contact code: O_r_N_b_4O_w_  
Go to ToC

### MG:a:3285

|  | Res. type | Atom | Residue | Distance | Occupancy | B-factor | Closest atom(s) to a 1^st^ shell water | EM\_dens. |
| --- | --- | --- | --- | --- | --- | --- | --- | --- |
| **Ion** | | | |  | 1.0 | 25.3 |  | 24.13 |
 1 | Water 1 | | a:7095 | 1.99 | 1.0 | 21.4 | 2.81 [G:N7]a:1996 // 2.91 [G:OP1]a:1996 | 17.93 | 2 | A | OP2 | a:1666 | 2.06 | 1.0 | 27.7 |  | 12.63 | 3 | Water 2 | | a:4969 | 2.10 | 1.0 | 21.5 | 2.67 [G:O6]a:1669 | 17.82 | 4 | Water 3 | | a:5626 | 2.25 | 1.0 | 19.5 | 2.72 [A:N7]a:1666 // 2.77 [G:O6]a:1996 | 18.52 | 5 | Water 4 | | a:3657 | 2.31 | 1.0 | 21.2 | 2.50 [G:O6]a:1668 // 2.68 [G:OP2]a:1996 | 15.86 | 6 | Water 5 | | a:6730 | 2.37 | 1.0 | 21.6 | 2.80 [A:N7]a:1667 | 18.55 | 7 | G | N7 | a:1996 | 3.97 | 1.0 | 17.6 |  | 15.12 | 8 | G | O6 | a:1669 | 4.05 | 1.0 | 26.9 |  | 16.18 | 9 | Water 6 | | a:9020 | 4.09 | 1.0 | 26.1 | 3.00 [A:OP1]a:1666 | 13.27 | 10 | G | OP2 | a:1996 | 4.23 | 1.0 | 24.6 |  | 16.09 | 11 | A | OP1 | a:1666 | 4.23 | 1.0 | 29.7 |  | 14.04 | 12 | G | OP1 | a:1996 | 4.24 | 1.0 | 26.8 |  | 13.84 | 13 | Water 7 | | a:10058 | 4.27 | 1.0 | 35.5 |  | 9.73 | 14 | G | O3' | a:1665 | 4.37 | 1.0 | 30.4 |  | 12.27 | 15 | A | N7 | a:1667 | 4.38 | 1.0 | 20.4 |  | 17.10 | 16 | A | O5' | a:1666 | 4.41 | 1.0 | 26.4 |  | 13.62 | 17 | G | O6 | a:1668 | 4.48 | 1.0 | 27.0 |  | 15.05 | 18 | A | N7 | a:1666 | 4.57 | 1.0 | 22.1 |  | 13.63 | 19 | G | O6 | a:1996 | 4.65 | 1.0 | 21.4 |  | 17.98 |

  
**1^st^ shell coordination angles and distances (*trans* in "blue"; *cys* in "green"; deviations in "orange"):**   
• 1-2(W/OP2): *101º*; 1-3(W/W): *95º*; 1-4(W/W): 93º; 2-3(OP2/W): 90º; 2-4(OP2/W): 90º; 3-4(W/W): 172º;   
• 1-2(W/OP2): 3.1Å; 1-3(W/W): 3.0Å; 1-4(W/W): 3.1Å; 2-3(OP2/W): 3.0Å; 2-4(OP2/W): 3.0Å; 3-4(W/W): *4.3Å*;   
  
**2.18 ± 0.14: the average coodination distance of the 1^st^ six ligands is Mg^2+^ compatible**  
**10: Number of potential Water contacts**   
**8 : Number of second shell contacts involving water or ions**   
**80%: Percentage of satisfied second shell Water contacts**   
• Contact(s): /O_w_/O_w_/O_w_/OP2/  
• Contact code: O_ph_3O_w_  
• Contact(s) in the exclusion zone: /O_w_/O_w_/  
→ Missing octahedral coordination for MG ←  
Ooctahedral coordination (4.6º)   
Go to ToC

### MG:a:3286

|  | Res. type | Atom | Residue | Distance | Occupancy | B-factor | Closest atom(s) to a 1^st^ shell water | EM\_dens. |
| --- | --- | --- | --- | --- | --- | --- | --- | --- |
| **Ion** | | | |  | 1.0 | 39.7 |  | 13.95 |
 1 | Water 1 | | a:9872 | 2.02 | 1.0 | 43.2 |  | 4.33 | 2 | Water 2 | | a:4422 | 2.03 | 1.0 | 36.0 | 2.62 [C:OP1]a:1990 | 7.49 | 3 | Water 3 | | a:10085 | 2.04 | 1.0 | 37.9 |  | 6.15 | 4 | Water 4 | | a:8668 | 2.14 | 1.0 | 29.0 | 2.98 [A:OP1]a:1671 | 11.20 | 5 | Water 5 | | a:6275 | 2.16 | 1.0 | 34.6 | 2.76 [C:OP1]a:2555 | 7.71 | 6 | Water 6 | | a:6621 | 2.25 | 1.0 | 30.8 | 2.78 [G:O6]a:1958 // 2.92 [A:OP2]a:1991 | 9.33 | 7 | Water 7 | | a:4397 | 4.09 | 1.0 | 41.6 |  | 5.92 | 8 | C | OP1 | a:1990 | 4.10 | 1.0 | 31.8 |  | 11.53 | 9 | C | OP1 | a:2555 | 4.13 | 1.0 | 29.8 |  | 13.41 | 10 | A | OP1 | a:1671 | 4.31 | 1.0 | 22.9 |  | 16.40 | 11 | G | O6 | a:1958 | 4.31 | 1.0 | 36.0 |  | 10.90 | 12 | A | OP2 | a:1991 | 4.32 | 1.0 | 35.4 |  | 7.41 | 13 | Water 8 | | a:5898 | 4.44 | 1.0 | 32.0 | 2.73 [A:OP1]a:1671 | 10.15 | 14 | Water 9 | | a:9601 | 4.44 | 1.0 | 35.4 |  | 5.65 | 15 | Water 10 | | a:6861 | 4.53 | 1.0 | 22.8 |  | 19.26 | 16 | Water 11 | | a:4845 | 4.53 | 1.0 | 23.3 | 2.66 [C:OP1]a:2555 | 15.70 | 17 | G | O3' | a:2554 | 4.74 | 1.0 | 28.0 |  | 14.10 |

  
**1^st^ shell coordination angles and distances (*trans* in "blue"; *cys* in "green"; deviations in "orange"):**   
• 1-2(W/W): 93º; 1-3(W/W): 86º; 1-4(W/W): 89º; 1-5(W/W): 173º; 1-6(W/W): 88º; 2-3(W/W): 171º; 2-4(W/W): *98º*; 2-5(W/W): *85º*; 2-6(W/W): 89º; 3-4(W/W): 91º; 3-5(W/W): 95º; 3-6(W/W): *81º*; 4-5(W/W): *97º*; 4-6(W/W): 172º; 5-6(W/W): 86º;   
• 1-2(W/W): 2.9Å; 1-3(W/W): 2.8Å; 1-4(W/W): 2.9Å; 1-5(W/W): 4.2Å; 1-6(W/W): 3.0Å; 2-3(W/W): 4.1Å; 2-4(W/W): 3.1Å; 2-5(W/W): 2.8Å; 2-6(W/W): 3.0Å; 3-4(W/W): 3.0Å; 3-5(W/W): 3.1Å; 3-6(W/W): 2.8Å; 4-5(W/W): *3.2Å*; 4-6(W/W): *4.4Å*; 5-6(W/W): 3.0Å;   
  
**2.11 ± 0.08: the average coodination distance of the 1^st^ six ligands is Mg^2+^ compatible**  
**12: Number of potential Water contacts**   
**5 : Number of second shell contacts involving water or ions**   
**42%: Percentage of satisfied second shell Water contacts**   
• Contact(s): /O_w_/O_w_/O_w_/O_w_/O_w_/O_w_/  
• Contact code: 6O_w_  
Ooctahedral coordination (4.8º)   
Go to ToC

### MG:a:3287

|  | Res. type | Atom | Residue | Distance | Occupancy | B-factor | Closest atom(s) to a 1^st^ shell water | EM\_dens. |
| --- | --- | --- | --- | --- | --- | --- | --- | --- |
| **Ion** | | | |  | 1.0 | 28.4 |  | 21.66 |
 1 | Water 1 | | a:7177 | 1.93 | 1.0 | 29.6 | 2.82 [G:OP2]a:2553 | 13.52 | 2 | U | OP2 | a:1673 | 1.98 | 1.0 | 28.5 |  | 11.09 | 3 | C | OP1 | a:1672 | 2.02 | 1.0 | 23.5 |  | 14.67 | 4 | Water 2 | | a:9503 | 2.31 | 1.0 | 30.1 |  | 12.63 | 5 | Water 3 | | a:6298 | 2.35 | 1.0 | 30.1 | 2.76 [G:OP2]a:2554 | 14.40 | 6 | Water 4 | | a:9491 | 2.39 | 1.0 | 24.7 |  | 15.73 | 7 | Water 5 | | a:10211 | 3.99 | 1.0 | 44.4 |  | 5.55 | 8 | U | OP1 | a:1673 | 4.04 | 1.0 | 31.2 |  | 11.37 | 9 | C | O5' | a:1672 | 4.08 | 1.0 | 25.7 |  | 16.55 | 10 | Water 6 | | a:7406 | 4.19 | 1.0 | 22.2 | 2.84 [G:OP2]a:2554 | 16.98 | 11 | U | O5' | a:1673 | 4.25 | 1.0 | 28.9 |  | 11.51 | 12 | C | OP2 | a:1672 | 4.29 | 1.0 | 20.9 |  | 14.04 | 13 | Water 7 | | a:10041 | 4.33 | 1.0 | 36.8 |  | 7.47 | 14 | Water 8 | | a:5862 | 4.34 | 1.0 | 43.0 | 2.73 [U:OP1]a:1673 | 5.32 | 15 | Water 9 | | a:10054 | 4.37 | 1.0 | 44.9 |  | 5.39 | 16 | G | OP2 | a:2554 | 4.38 | 1.0 | 25.6 |  | 14.54 | 17 | C | O3' | a:1672 | 4.39 | 1.0 | 29.2 |  | 13.49 | 18 | G | OP2 | a:2553 | 4.39 | 1.0 | 31.0 |  | 11.33 | 19 | A | O3' | a:1671 | 4.40 | 1.0 | 24.2 |  | 15.89 | 20 | Water 10 | | a:9079 | 4.65 | 1.0 | 44.5 |  | 5.07 | 21 | Water 11 | | a:7412 | 4.71 | 1.0 | 27.1 |  | 11.99 | 22 | Water 12 | | a:4103 | 4.80 | 1.0 | 30.8 |  | 12.84 |

  
**1^st^ shell coordination angles and distances (*trans* in "blue"; *cys* in "green"; deviations in "orange"):**   
• 1-2(W/OP2): 93º; 1-3(W/OP1): 92º; 2-3(OP2/OP1): 92º;   
• 1-2(W/OP2): 2.8Å; 1-3(W/OP1): 2.8Å; 2-3(OP2/OP1): 2.9Å;   
  
**2.16 ± 0.19: the average coodination distance of the 1^st^ six ligands is Mg^2+^ compatible**  
**8: Number of potential Water contacts**   
**2 : Number of second shell contacts involving water or ions**   
**25%: Percentage of satisfied second shell Water contacts**   
• Contact(s): /O_w_/OP1/OP2/  
• Contact code: *cis-*2O_ph_O_w_  
• Contact(s) in the exclusion zone: /O_w_/O_w_/O_w_/  
→ Missing octahedral coordination for MG ←  
Go to ToC

### MG:a:3288

|  | Res. type | Atom | Residue | Distance | Occupancy | B-factor | Closest atom(s) to a 1^st^ shell water | EM\_dens. |
| --- | --- | --- | --- | --- | --- | --- | --- | --- |
| **Ion** | | | |  | 1.0 | 38.2 |  | 16.23 |
 1 | Water 1 | | a:5944 | 1.95 | 1.0 | 31.7 | 2.74 [A:N7]a:1680 | 11.24 | 2 | Water 2 | | a:7612 | 2.02 | 1.0 | 31.6 | 2.85 [A:N7]a:1681 | 10.00 | 3 | A | OP2 | a:1680 | 2.09 | 1.0 | 33.7 |  | 9.84 | 4 | Water 3 | | a:7844 | 2.19 | 1.0 | 32.2 |  | 8.42 | 5 | Water 4 | | a:9378 | 2.23 | 1.0 | 37.3 |  | 10.09 | 6 | Water 5 | | a:8997 | 2.38 | 1.0 | 34.1 |  | 11.31 | 7 | A | O5' | a:1680 | 3.88 | 1.0 | 32.2 |  | 10.31 | 8 | Water 6 | | a:6648 | 3.98 | 1.0 | 34.3 |  | 8.92 | 9 | Water 7 | | a:4687 | 4.07 | 1.0 | 31.3 |  | 12.40 | 10 | A | N7 | a:1680 | 4.12 | 1.0 | 24.5 |  | 15.53 | 11 | A | N7 | a:1681 | 4.20 | 1.0 | 28.5 |  | 11.67 | 12 | Water 8 | | a:10099 | 4.30 | 1.0 | 44.2 |  | 5.39 | 13 | A | OP1 | a:1680 | 4.32 | 1.0 | 35.9 |  | 10.82 | 14 | A | O3' | a:1679 | 4.53 | 1.0 | 33.0 |  | 10.51 | 15 | Water 9 | | a:3882 | 4.61 | 1.0 | 35.2 |  | 5.65 |

  
**1^st^ shell coordination angles and distances (*trans* in "blue"; *cys* in "green"; deviations in "orange"):**   
• 1-2(W/W): *97º*; 1-3(W/OP2): *96º*; 1-4(W/W): 179º; 1-5(W/W): 93º; 2-3(W/OP2): 91º; 2-4(W/W): *82º*; 2-5(W/W): 90º; 3-4(OP2/W): *84º*; 3-5(OP2/W): 171º; 4-5(W/W): 87º;   
• 1-2(W/W): 3.0Å; 1-3(W/OP2): 3.0Å; 1-4(W/W): 4.1Å; 1-5(W/W): 3.0Å; 2-3(W/OP2): 2.9Å; 2-4(W/W): 2.8Å; 2-5(W/W): 3.0Å; 3-4(OP2/W): 2.9Å; 3-5(OP2/W): *4.3Å*; 4-5(W/W): 3.0Å;   
  
**2.14 ± 0.14: the average coodination distance of the 1^st^ six ligands is Mg^2+^ compatible**  
**10: Number of potential Water contacts**   
**2 : Number of second shell contacts involving water or ions**   
**20%: Percentage of satisfied second shell Water contacts**   
• Contact(s): /O_w_/O_w_/O_w_/O_w_/OP2/  
• Contact code: O_ph_4O_w_  
• Contact(s) in the exclusion zone: /O_w_/  
→ Missing octahedral coordination for MG ←  
Ooctahedral coordination (4.4º)   
Go to ToC

### MG:a:3289

|  | Res. type | Atom | Residue | Distance | Occupancy | B-factor | Closest atom(s) to a 1^st^ shell water | EM\_dens. |
| --- | --- | --- | --- | --- | --- | --- | --- | --- |
| **Ion** | | | |  | 1.0 | 27.4 |  | 24.83 |
 1 | Water 1 | | a:6584 | 1.97 | 1.0 | 22.2 | 2.90 [A:N3]a:1774 // 2.97 [A:OP1]a:1788 | 19.43 | 2 | Water 2 | | a:6456 | 2.06 | 1.0 | 26.4 | 2.77 [A:OP2]a:1788 | 17.61 | 3 | C | OP1 | a:1776 | 2.16 | 1.0 | 25.7 |  | 12.91 | 4 | Water 3 | | a:8399 | 2.17 | 1.0 | 24.1 | 2.92 [G:O6]a:1778 | 16.61 | 5 | Water 4 | | a:4123 | 2.18 | 1.0 | 19.7 | 2.59 [U:OP2]a:1777 | 18.01 | 6 | Water 5 | | a:4343 | 2.19 | 1.0 | 21.9 | 2.62 [A:O2']a:1774 | 16.02 | 7 | Water 6 | | a:9920 | 3.93 | 1.0 | 27.7 |  | 12.70 | 8 | C | OP2 | a:1776 | 4.06 | 1.0 | 25.0 |  | 14.23 | 9 | C | O5' | a:1776 | 4.09 | 1.0 | 22.6 |  | 16.40 | 10 | A | O2' | a:1774 | 4.11 | 1.0 | 30.0 |  | 12.94 | 11 | Water 7 | | a:5691 | 4.18 | 1.0 | 23.9 |  | 15.98 | 12 | A | N3 | a:1774 | 4.22 | 1.0 | 19.0 |  | 18.11 | 13 | Water 8 | | a:6470 | 4.28 | 1.0 | 22.6 | 2.97 [A:OP2]a:1788 | 16.02 | 14 | U | OP2 | a:1777 | 4.28 | 1.0 | 26.6 |  | 12.29 | 15 | Water 9 | | a:6952 | 4.28 | 1.0 | 24.0 | 2.95 [C:OP2]a:1776 | 14.28 | 16 | A | OP2 | a:1788 | 4.28 | 1.0 | 23.4 |  | 14.63 | 17 | G | O6 | a:1778 | 4.31 | 1.0 | 20.4 |  | 18.05 | 18 | A | OP1 | a:1788 | 4.40 | 1.0 | 23.8 |  | 14.51 | 19 | Water 10 | | a:5664 | 4.49 | 1.0 | 26.6 |  | 14.67 | 20 | A | O3' | a:1775 | 4.64 | 1.0 | 26.0 |  | 16.24 | 21 | Water 11 | | a:6315 | 4.64 | 1.0 | 23.9 | 2.95 [U:OP2]a:1777 | 14.46 |

  
**1^st^ shell coordination angles and distances (*trans* in "blue"; *cys* in "green"; deviations in "orange"):**   
• 1-2(W/W): 90º; 1-3(W/OP1): *84º*; 1-4(W/W): 178º; 1-5(W/W): *96º*; 1-6(W/W): 88º; 2-3(W/OP1): 171º; 2-4(W/W): 91º; 2-5(W/W): *100º*; 2-6(W/W): 93º; 3-4(OP1/W): 94º; 3-5(OP1/W): 88º; 3-6(OP1/W): *79º*; 4-5(W/W): *85º*; 4-6(W/W): 92º; 5-6(W/W): *167º*;   
• 1-2(W/W): 2.9Å; 1-3(W/OP1): 2.8Å; 1-4(W/W): 4.1Å; 1-5(W/W): 3.1Å; 1-6(W/W): 2.9Å; 2-3(W/OP1): *4.2Å*; 2-4(W/W): 3.0Å; 2-5(W/W): *3.2Å*; 2-6(W/W): 3.1Å; 3-4(OP1/W): 3.2Å; 3-5(OP1/W): 3.0Å; 3-6(OP1/W): 2.8Å; 4-5(W/W): 2.9Å; 4-6(W/W): 3.1Å; 5-6(W/W): *4.3Å*;   
  
**2.12 ± 0.08: the average coodination distance of the 1^st^ six ligands is Mg^2+^ compatible**  
**10: Number of potential Water contacts**   
**6 : Number of second shell contacts involving water or ions**   
**60%: Percentage of satisfied second shell Water contacts**   
• Contact(s): /O_w_/O_w_/O_w_/O_w_/O_w_/OP1/  
• Contact code: O_ph_5O_w_  
→ Slightly distorded coord. octahedron (5.0º) ←  
Go to ToC

### MG:a:3290

|  | Res. type | Atom | Residue | Distance | Occupancy | B-factor | Closest atom(s) to a 1^st^ shell water | EM\_dens. |
| --- | --- | --- | --- | --- | --- | --- | --- | --- |
| **Ion** | | | |  | 1.0 | 28.1 |  | 23.76 |
 1 | G | OP2 | a:1778 | 2.05 | 1.0 | 24.2 |  | 15.69 | 2 | Water 1 | | a:6552 | 2.11 | 1.0 | 24.1 | 2.78 [A:OP1]a:1985 | 12.63 | 3 | Water 2 | | a:9091 | 2.16 | 1.0 | 25.5 |  | 12.78 | 4 | Water 3 | | a:6315 | 2.24 | 1.0 | 23.9 | 2.76 [G:N7]a:1778 | 14.46 | 5 | Water 4 | | a:8667 | 2.41 | 1.0 | 25.4 |  | 15.62 | 6 | G | OP1 | a:1778 | 3.90 | 1.0 | 32.5 |  | 10.39 | 7 | Water 5 | | a:5691 | 3.98 | 1.0 | 23.9 |  | 15.98 | 8 | Water 6 | | a:9768 | 4.03 | 1.0 | 35.5 |  | 7.35 | 9 | U | O3' | a:1777 | 4.14 | 1.0 | 26.0 |  | 14.31 | 10 | A | OP1 | a:1985 | 4.30 | 1.0 | 36.5 |  | 8.94 | 11 | Water 7 | | a:9920 | 4.32 | 1.0 | 27.7 |  | 12.70 | 12 | U | OP1 | a:1777 | 4.32 | 1.0 | 27.1 |  | 14.20 | 13 | U | OP2 | a:1777 | 4.34 | 1.0 | 26.6 |  | 12.29 | 14 | G | N7 | a:1778 | 4.39 | 1.0 | 19.0 |  | 15.43 | 15 | Water 8 | | a:4143 | 4.53 | 1.0 | 31.9 |  | 11.79 | 16 | U | O5' | a:1777 | 4.53 | 1.0 | 25.1 |  | 15.55 | 17 | G | O5' | a:1778 | 4.53 | 1.0 | 31.3 |  | 11.59 | 18 | Water 9 | | a:9537 | 4.57 | 1.0 | 35.5 |  | 6.68 | 19 | Water 10 | | a:7087 | 4.76 | 1.0 | 34.3 | 2.81 [G:OP1]a:1778 | 8.60 | 20 | A | OP2 | a:1985 | 4.79 | 1.0 | 35.2 |  | 7.50 |

  
**1^st^ shell coordination angles and distances (*trans* in "blue"; *cys* in "green"; deviations in "orange"):**   
• 1-2(OP2/W): *168º*; 1-3(OP2/W): 91º; 1-4(OP2/W): 94º; 2-3(W/W): 85º; 2-4(W/W): 87º; 3-4(W/W): *167º*;   
• 1-2(OP2/W): 4.1Å; 1-3(OP2/W): 3.0Å; 1-4(OP2/W): 3.1Å; 2-3(W/W): 2.9Å; 2-4(W/W): 3.0Å; 3-4(W/W): *4.4Å*;   
  
**2.48 ± 0.65: the average coodination distance of the 1^st^ six ligands is Na^+^** compatible  
**8: Number of potential Water contacts**   
**2 : Number of second shell contacts involving water or ions**   
**25%: Percentage of satisfied second shell Water contacts**   
• Contact(s): /O_w_/O_w_/O_w_/OP2/  
• Contact code: O_ph_3O_w_  
• Contact(s) in the exclusion zone: /O_w_/  
→ Missing octahedral coordination for MG ←  
→ Slightly distorded coord. octahedron (6.2º) ←  
Go to ToC

### MG:a:3291

|  | Res. type | Atom | Residue | Distance | Occupancy | B-factor | Closest atom(s) to a 1^st^ shell water | EM\_dens. |
| --- | --- | --- | --- | --- | --- | --- | --- | --- |
| **Ion** | | | |  | 1.0 | 39.5 |  | 12.24 |
 1 | Water 1 | | a:3919 | 2.02 | 1.0 | 36.1 | 2.56 [A:OP1]a:1941 | 9.54 | 2 | Water 2 | | a:6184 | 2.07 | 1.0 | 32.4 | 2.75 [G:N7]a:1963 | 10.21 | 3 | Water 3 | | a:4599 | 2.26 | 1.0 | 34.5 | 2.64 [G:OP2]a:1963 | 9.54 | 4 | Water 4 | | a:6992 | 2.44 | 1.0 | 41.8 | 2.80 [G:OP2]a:1963 | 6.48 | 5 | Water 5 | | a:7668 | 4.10 | 1.0 | 31.2 | 2.85 [C:OP2]a:1962 | 10.52 | 6 | Water 6 | | a:7211 | 4.16 | 1.0 | 49.7 | 2.82 [A:OP1]a:1940 | 4.70 | 7 | G | OP2 | a:1963 | 4.18 | 1.0 | 32.7 |  | 9.94 | 8 | A | OP1 | a:1941 | 4.24 | 1.0 | 36.9 |  | 7.87 | 9 | Water 7 | | a:7967 | 4.30 | 1.0 | 42.3 |  | 5.48 | 10 | A | OP1 | a:1940 | 4.31 | 1.0 | 38.6 |  | 8.52 | 11 | G | N7 | a:1963 | 4.34 | 1.0 | 23.1 |  | 13.55 | 12 | Water 8 | | a:6864 | 4.48 | 1.0 | 43.8 |  | 4.60 | 13 | Water 9 | | a:9701 | 4.51 | 1.0 | 38.3 |  | 8.46 | 14 | C | OP2 | a:1962 | 4.57 | 1.0 | 24.8 |  | 12.65 | 15 | Water 10 | | a:3693 | 4.60 | 1.0 | 45.0 |  | 4.45 |

  
**1^st^ shell coordination angles and distances (*trans* in "blue"; *cys* in "green"; deviations in "orange"):**   
• 1-2(W/W): 91º; 1-3(W/W): *102º*; 2-3(W/W): 95º;   
• 1-2(W/W): 2.9Å; 1-3(W/W): *3.3Å*; 2-3(W/W): 3.2Å;   
  
**2.84 ± 0.92: the average coodination distance of the 1^st^ six ligands is K^+^ compatible**  
**12: Number of potential Water contacts**   
**6 : Number of second shell contacts involving water or ions**   
**50%: Percentage of satisfied second shell Water contacts**   
• Contact(s): /O_w_/O_w_/O_w_/  
• Contact code: 3O_w_  
• Contact(s) in the exclusion zone: /O_w_/  
→ Missing octahedral coordination for MG ←  
Go to ToC

### MG:a:3292

|  | Res. type | Atom | Residue | Distance | Occupancy | B-factor | Closest atom(s) to a 1^st^ shell water | EM\_dens. |
| --- | --- | --- | --- | --- | --- | --- | --- | --- |
| **Ion** | | | |  | 1.0 | 38.8 |  | 15.14 |
 1 | Water 1 | | a:3499 | 1.88 | 1.0 | 42.1 |  | 6.47 | 2 | Water 2 | | a:5357 | 1.90 | 1.0 | 35.9 | 2.76 [2MG:N7]a:1837 | 9.27 | 3 | A | OP2 | a:1974 | 2.07 | 1.0 | 33.9 |  | 7.99 | 4 | Water 3 | | a:8458 | 2.23 | 1.0 | 42.0 |  | 6.15 | 5 | Water 4 | | a:9328 | 2.29 | 1.0 | 40.8 |  | 6.33 | 6 | A | OP2 | a:1904 | 3.86 | 1.0 | 37.1 |  | 7.49 | 7 | Water 5 | | a:5683 | 3.97 | 1.0 | 38.7 |  | 8.28 | 8 | A | OP1 | a:1974 | 3.99 | 1.0 | 34.3 |  | 7.00 | 9 | 2MG | N7 | a:1837 | 4.24 | 1.0 | 29.9 |  | 11.61 | 10 | 2MG | O6 | a:1837 | 4.28 | 1.0 | 33.0 |  | 11.45 | 11 | U | O2' | a:1836 | 4.34 | 1.0 | 34.7 |  | 9.88 | 12 | 2MG | OP2 | a:1837 | 4.34 | 1.0 | 31.9 |  | 9.29 | 13 | Water 6 | | a:3580 | 4.36 | 1.0 | 43.5 | 2.48 [A:OP2]a:1904 | 4.14 | 14 | A | O5' | a:1974 | 4.38 | 1.0 | 35.1 |  | 8.00 | 15 | Water 7 | | a:8274 | 4.39 | 1.0 | 43.1 |  | 6.14 | 16 | A | O3' | a:1973 | 4.47 | 1.0 | 33.9 |  | 10.02 | 17 | Water 8 | | a:8555 | 4.57 | 1.0 | 32.9 | 2.94 [A:O5']a:1974 | 8.46 | 18 | Water 9 | | a:6504 | 4.67 | 1.0 | 42.6 | 2.77 [2MG:OP2]a:1837 | 5.74 |

  
**1^st^ shell coordination angles and distances (*trans* in "blue"; *cys* in "green"; deviations in "orange"):**   
• 1-2(W/W): 85º; 1-3(W/OP2): *76º*; 1-4(W/W): 91º; 1-5(W/W): 94º; 2-3(W/OP2): 86º; 2-4(W/W): 171º; 2-5(W/W): *75º*; 3-4(OP2/W): 85º; 3-5(OP2/W): *159º*; 4-5(W/W): *114º*;   
• 1-2(W/W): 2.6Å; 1-3(W/OP2): 2.4Å; 1-4(W/W): 2.9Å; 1-5(W/W): 3.1Å; 2-3(W/OP2): 2.7Å; 2-4(W/W): 4.1Å; 2-5(W/W): 2.6Å; 3-4(OP2/W): 2.9Å; 3-5(OP2/W): *4.3Å*; 4-5(W/W): *3.8Å*;   
  
**2.37 ± 0.68: the average coodination distance of the 1^st^ six ligands is Na^+^** compatible  
**8: Number of potential Water contacts**   
**1 : Number of second shell contacts involving water or ions**   
**12%: Percentage of satisfied second shell Water contacts**   
• Contact(s): /O_w_/O_w_/OP2/  
• Contact code: O_ph_2O_w_  
• Contact(s) in the exclusion zone: /O_w_/O_w_/  
→ Missing octahedral coordination for MG ←  
→→→ Distorded coord. octahedron (10.3º) ←←←  
Go to ToC

### MG:a:3293

|  | Res. type | Atom | Residue | Distance | Occupancy | B-factor | Closest atom(s) to a 1^st^ shell water | EM\_dens. |
| --- | --- | --- | --- | --- | --- | --- | --- | --- |
| **Ion** | | | |  | 1.0 | 29.3 |  | 21.79 |
 1 | C | OP2 | a:2029 | 1.94 | 1.0 | 24.4 |  | 14.06 | 2 | Water 1 | | a:5404 | 1.98 | 1.0 | 23.1 | 2.70 [G:OP1]a:2028 | 16.73 | 3 | Water 2 | | a:6342 | 2.09 | 1.0 | 30.4 | 2.76 [U:OP2]a:2038 | 11.87 | 4 | Water 3 | | a:8502 | 2.17 | 1.0 | 21.9 |  | 14.41 | 5 | Water 4 | | a:8083 | 2.25 | 1.0 | 24.0 | 2.89 [G:OP2]a:2028 | 16.27 | 6 | Water 5 | | a:8981 | 2.33 | 1.0 | 26.6 |  | 13.97 | 7 | C | OP1 | a:2029 | 3.76 | 1.0 | 31.7 |  | 11.82 | 8 | Water 6 | | d:432 | 3.82 | 1.0 | 32.8 | 2.86 [C:OP1]a:2029 | 8.86 | 9 | Water 7 | | a:4963 | 3.99 | 1.0 | 32.4 |  | 10.98 | 10 | Water 8 | | a:8579 | 4.10 | 1.0 | 34.1 |  | 9.48 | 11 | C | O5' | a:2029 | 4.14 | 1.0 | 22.1 |  | 14.96 | 12 | G | OP1 | a:2028 | 4.20 | 1.0 | 28.9 |  | 11.55 | 13 | U | OP2 | a:2038 | 4.26 | 1.0 | 23.3 |  | 15.50 | 14 | G | O3' | a:2028 | 4.35 | 1.0 | 20.8 |  | 15.54 | 15 | Water 9 | | d:451 | 4.40 | 1.0 | 35.3 |  | 7.25 | 16 | G | OP2 | a:2028 | 4.55 | 1.0 | 26.2 |  | 14.79 | 17 | Water 10 | | a:9489 | 4.61 | 1.0 | 38.7 |  | 6.89 | 18 | Water 11 | | d:468 | 4.68 | 1.0 | 37.9 |  | 7.86 | 19 | G | O5' | a:2028 | 4.70 | 1.0 | 26.0 |  | 13.70 |

  
**1^st^ shell coordination angles and distances (*trans* in "blue"; *cys* in "green"; deviations in "orange"):**   
• 1-2(OP2/W): *96º*; 1-3(OP2/W): 176º; 1-4(OP2/W): 91º; 1-5(OP2/W): 91º; 2-3(W/W): 87º; 2-4(W/W): 90º; 2-5(W/W): 91º; 3-4(W/W): 92º; 3-5(W/W): 86º; 4-5(W/W): 178º;   
• 1-2(OP2/W): 2.9Å; 1-3(OP2/W): 4.0Å; 1-4(OP2/W): 2.9Å; 1-5(OP2/W): 3.0Å; 2-3(W/W): 2.8Å; 2-4(W/W): 2.9Å; 2-5(W/W): 3.0Å; 3-4(W/W): 3.1Å; 3-5(W/W): 3.0Å; 4-5(W/W): *4.4Å*;   
  
**2.12 ± 0.14: the average coodination distance of the 1^st^ six ligands is Mg^2+^ compatible**  
**10: Number of potential Water contacts**   
**3 : Number of second shell contacts involving water or ions**   
**30%: Percentage of satisfied second shell Water contacts**   
• Contact(s): /O_w_/O_w_/O_w_/O_w_/OP2/  
• Contact code: O_ph_4O_w_  
• Contact(s) in the exclusion zone: /O_w_/  
→ Missing octahedral coordination for MG ←  
Ooctahedral coordination (2.3º)   
Go to ToC

### MG:a:3294

|  | Res. type | Atom | Residue | Distance | Occupancy | B-factor | Closest atom(s) to a 1^st^ shell water | EM\_dens. |
| --- | --- | --- | --- | --- | --- | --- | --- | --- |
| **Ion** | | | |  | 1.0 | 28.9 |  | 22.66 |
 1 | G | OP2 | a:2061 | 2.02 | 1.0 | 24.7 |  | 15.03 | 2 | G | OP2 | a:2060 | 2.03 | 1.0 | 26.2 |  | 15.08 | 3 | Water 1 | | a:8044 | 2.06 | 1.0 | 27.8 | 2.89 [G:N7]a:2061 | 14.68 | 4 | Water 2 | | a:5066 | 2.08 | 1.0 | 29.3 | 2.68 [A:N7]a:2062 | 11.16 | 5 | Water 3 | | a:9604 | 2.14 | 1.0 | 29.8 |  | 11.87 | 6 | Water 4 | | a:6577 | 2.34 | 1.0 | 26.5 | 2.78 [PSU:OP2]a:2508 | 13.82 | 7 | G | OP1 | a:2060 | 3.76 | 1.0 | 28.8 |  | 13.15 | 8 | G | O5' | a:2060 | 3.90 | 1.0 | 24.6 |  | 14.20 | 9 | G | O5' | a:2061 | 4.13 | 1.0 | 26.0 |  | 17.07 | 10 | Water 5 | | a:9602 | 4.18 | 1.0 | 41.1 |  | 6.78 | 11 | Water 6 | | a:4281 | 4.25 | 1.0 | 33.5 |  | 7.84 | 12 | G | OP1 | a:2061 | 4.39 | 1.0 | 26.5 |  | 12.14 | 13 | G | N7 | a:2061 | 4.39 | 1.0 | 20.5 |  | 15.27 | 14 | G | O3' | a:2060 | 4.41 | 1.0 | 23.5 |  | 14.10 | 15 | C | O3' | a:2059 | 4.47 | 1.0 | 29.1 |  | 14.03 | 16 | A | N7 | a:2062 | 4.57 | 1.0 | 27.4 |  | 12.36 | 17 | Water 7 | | a:4361 | 4.70 | 1.0 | 33.5 | 2.62 [G:OP1]a:2060 // 2.90 [G:OP1]a:2061 | 11.02 | 18 | PSU | OP2 | a:2508 | 4.70 | 1.0 | 34.5 |  | 9.81 |

  
**1^st^ shell coordination angles and distances (*trans* in "blue"; *cys* in "green"; deviations in "orange"):**   
• 1-2(OP2/OP2): 88º; 1-3(OP2/W): 93º; 1-4(OP2/W): 89º; 1-5(OP2/W): 172º; 2-3(OP2/W): 90º; 2-4(OP2/W): 177º; 2-5(OP2/W): *99º*; 3-4(W/W): 91º; 3-5(W/W): 91º; 4-5(W/W): *85º*;   
• 1-2(OP2/OP2): 2.8Å; 1-3(OP2/W): 3.0Å; 1-4(OP2/W): 2.9Å; 1-5(OP2/W): 4.2Å; 2-3(OP2/W): 2.9Å; 2-4(OP2/W): 4.1Å; 2-5(OP2/W): 3.2Å; 3-4(W/W): 3.0Å; 3-5(W/W): 3.0Å; 4-5(W/W): 2.8Å;   
  
**2.11 ± 0.11: the average coodination distance of the 1^st^ six ligands is Mg^2+^ compatible**  
**8: Number of potential Water contacts**   
**3 : Number of second shell contacts involving water or ions**   
**38%: Percentage of satisfied second shell Water contacts**   
• Contact(s): /O_w_/O_w_/O_w_/OP2/OP2/  
• Contact code: *cis-*2O_ph_3O_w_  
• Contact(s) in the exclusion zone: /O_w_/  
→ Missing octahedral coordination for MG ←  
Ooctahedral coordination (3.4º)   
Go to ToC

### MG:a:3295

|  | Res. type | Atom | Residue | Distance | Occupancy | B-factor | Closest atom(s) to a 1^st^ shell water | EM\_dens. |
| --- | --- | --- | --- | --- | --- | --- | --- | --- |
| **Ion** | | | |  | 1.0 | 26.3 |  | 25.05 |
 1 | G | OP2 | a:2506 | 1.91 | 1.0 | 25.3 |  | 14.68 | 2 | G | OP2 | a:2065 | 1.99 | 1.0 | 22.6 |  | 16.00 | 3 | Water 1 | | a:6024 | 2.16 | 1.0 | 22.3 | 2.74 [A:O2']a:2064 | 15.68 | 4 | Water 2 | | a:8418 | 2.25 | 1.0 | 22.8 |  | 16.82 | 5 | Water 3 | | a:8933 | 2.34 | 1.0 | 23.6 | 2.98 [2MG:N7]a:2449 | 18.94 | 6 | C | O3' | a:2505 | 3.94 | 1.0 | 26.6 |  | 12.95 | 7 | A | O3' | a:2064 | 4.02 | 1.0 | 25.0 |  | 15.99 | 8 | Water 4 | | a:3983 | 4.12 | 1.0 | 22.5 | 2.57 [G:OP1]a:2065 | 15.96 | 9 | A | O2' | a:2064 | 4.18 | 1.0 | 24.2 |  | 16.73 | 10 | C | O2' | a:2505 | 4.25 | 1.0 | 28.7 |  | 15.75 | 11 | G | OP1 | a:2506 | 4.25 | 1.0 | 30.3 |  | 13.98 | 12 | G | O5' | a:2506 | 4.26 | 1.0 | 23.7 |  | 13.54 | 13 | G | OP1 | a:2065 | 4.34 | 1.0 | 23.8 |  | 13.52 | 14 | Water 5 | | a:4835 | 4.38 | 1.0 | 23.6 |  | 16.76 | 15 | G | O5' | a:2065 | 4.39 | 1.0 | 29.3 |  | 13.03 | 16 | G | N7 | a:2450 | 4.59 | 1.0 | 18.8 |  | 17.70 | 17 | Water 6 | | a:5294 | 4.69 | 1.0 | 23.7 |  | 16.38 | 18 | 2MG | N7 | a:2449 | 4.70 | 1.0 | 13.7 |  | 18.25 |

  
**1^st^ shell coordination angles and distances (*trans* in "blue"; *cys* in "green"; deviations in "orange"):**   
• 1-2(OP2/OP2): *99º*; 1-3(OP2/W): *97º*; 1-4(OP2/W): 89º; 2-3(OP2/W): 88º; 2-4(OP2/W): 171º; 3-4(W/W): 86º;   
• 1-2(OP2/OP2): 3.0Å; 1-3(OP2/W): 3.1Å; 1-4(OP2/W): 2.9Å; 2-3(OP2/W): 2.9Å; 2-4(OP2/W): *4.2Å*; 3-4(W/W): 3.0Å;   
  
**2.43 ± 0.69: the average coodination distance of the 1^st^ six ligands is Na^+^** compatible  
**6: Number of potential Water contacts**   
**2 : Number of second shell contacts involving water or ions**   
**33%: Percentage of satisfied second shell Water contacts**   
• Contact(s): /O_w_/O_w_/OP2/OP2/  
• Contact code: *cis-*2O_ph_2O_w_  
• Contact(s) in the exclusion zone: /O_w_/  
→ Missing octahedral coordination for MG ←  
→ Slightly distorded coord. octahedron (5.2º) ←  
Go to ToC

### MG:a:3296

|  | Res. type | Atom | Residue | Distance | Occupancy | B-factor | Closest atom(s) to a 1^st^ shell water | EM\_dens. |
| --- | --- | --- | --- | --- | --- | --- | --- | --- |
| **Ion** | | | |  | 1.0 | 27.3 |  | 24.43 |
 1 | Water 1 | | a:4702 | 2.05 | 1.0 | 24.6 | 2.65 [A:OP1]a:2066 // 2.92 [C:OP2]a:2447 | 15.58 | 2 | Water 2 | | a:9074 | 2.07 | 1.0 | 27.2 |  | 13.71 | 3 | Water 3 | | a:9486 | 2.11 | 1.0 | 25.2 |  | 14.42 | 4 | A | OP2 | a:2066 | 2.15 | 1.0 | 31.4 |  | 11.54 | 5 | Water 4 | | a:6688 | 2.22 | 1.0 | 25.8 | 2.79 [G:OP2]a:2448 | 14.86 | 6 | Water 5 | | a:3983 | 2.37 | 1.0 | 22.5 | 2.57 [G:OP1]a:2065 | 15.96 | 7 | A | OP1 | a:2066 | 3.59 | 1.0 | 43.2 |  | 7.47 | 8 | A | O5' | a:2066 | 3.98 | 1.0 | 38.2 |  | 7.95 | 9 | Water 6 | | a:4835 | 4.04 | 1.0 | 23.6 |  | 16.76 | 10 | Water 7 | | a:8707 | 4.04 | 1.0 | 28.6 |  | 11.54 | 11 | Water 8 | | a:8933 | 4.16 | 1.0 | 23.6 |  | 18.94 | 12 | Water 9 | | a:3795 | 4.18 | 1.0 | 37.5 | 2.53 [C:OP2]a:2447 | 9.85 | 13 | G | OP2 | a:2448 | 4.35 | 1.0 | 22.0 |  | 16.07 | 14 | C | OP2 | a:2447 | 4.38 | 1.0 | 24.9 |  | 14.24 | 15 | G | N7 | a:2448 | 4.40 | 1.0 | 16.9 |  | 17.59 | 16 | G | OP1 | a:2065 | 4.58 | 1.0 | 23.8 |  | 13.52 | 17 | G | O3' | a:2065 | 4.59 | 1.0 | 27.4 |  | 12.70 | 18 | Water 10 | | a:7195 | 4.71 | 1.0 | 30.9 |  | 11.10 |

  
**1^st^ shell coordination angles and distances (*trans* in "blue"; *cys* in "green"; deviations in "orange"):**   
• 1-2(W/W): 178º; 1-3(W/W): *96º*; 1-4(W/OP2): 91º; 1-5(W/W): *96º*; 2-3(W/W): 86º; 2-4(W/OP2): 91º; 2-5(W/W): *82º*; 3-4(W/OP2): 94º; 3-5(W/W): 93º; 4-5(OP2/W): *170º*;   
• 1-2(W/W): 4.1Å; 1-3(W/W): 3.1Å; 1-4(W/OP2): 3.0Å; 1-5(W/W): 3.2Å; 2-3(W/W): 2.9Å; 2-4(W/OP2): 3.0Å; 2-5(W/W): 2.8Å; 3-4(W/OP2): 3.1Å; 3-5(W/W): 3.1Å; 4-5(OP2/W): *4.4Å*;   
  
**2.16 ± 0.11: the average coodination distance of the 1^st^ six ligands is Mg^2+^ compatible**  
**10: Number of potential Water contacts**   
**4 : Number of second shell contacts involving water or ions**   
**40%: Percentage of satisfied second shell Water contacts**   
• Contact(s): /O_w_/O_w_/O_w_/O_w_/OP2/  
• Contact code: O_ph_4O_w_  
• Contact(s) in the exclusion zone: /O_w_/  
→ Missing octahedral coordination for MG ←  
Ooctahedral coordination (4.5º)   
Go to ToC

### MG:a:3297

|  | Res. type | Atom | Residue | Distance | Occupancy | B-factor | Closest atom(s) to a 1^st^ shell water | EM\_dens. |
| --- | --- | --- | --- | --- | --- | --- | --- | --- |
| **Ion** | | | |  | 1.0 | 26.1 |  | 24.95 |
 1 | Water 1 | | a:4473 | 2.11 | 1.0 | 19.5 | 2.63 [U:O4]a:2078 // 2.71 [U:O4]a:2247 | 19.32 | 2 | Water 2 | | a:5258 | 2.17 | 1.0 | 20.7 | 2.69 [G:O6]a:2246 | 17.82 | 3 | Water 3 | | a:7418 | 2.19 | 1.0 | 20.3 | 2.84 [G:N7]a:2246 | 17.66 | 4 | Water 4 | | a:5974 | 2.20 | 1.0 | 20.6 | 2.74 [U:O4]a:2079 | 15.59 | 5 | Water 5 | | a:10114 | 2.20 | 1.0 | 24.5 |  | 16.75 | 6 | Water 6 | | a:8347 | 2.29 | 1.0 | 21.5 | 2.91 [A:N7]a:2245 | 15.88 | 7 | U | O4 | a:2247 | 3.76 | 1.0 | 26.2 |  | 17.58 | 8 | Water 7 | | a:7169 | 3.98 | 1.0 | 34.4 |  | 10.80 | 9 | Water 8 | | a:9617 | 3.98 | 1.0 | 29.4 |  | 9.60 | 10 | U | O4 | a:2078 | 4.15 | 1.0 | 20.4 |  | 17.97 | 11 | U | O4 | a:2079 | 4.22 | 1.0 | 24.3 |  | 16.27 | 12 | G | N7 | a:2242 | 4.22 | 1.0 | 20.2 |  | 15.86 | 13 | G | O6 | a:2246 | 4.29 | 1.0 | 24.7 |  | 15.37 | 14 | A | N7 | a:2245 | 4.41 | 1.0 | 20.1 |  | 14.55 | 15 | Water 9 | | a:6061 | 4.55 | 1.0 | 26.9 |  | 10.70 | 16 | G | N7 | a:2246 | 4.56 | 1.0 | 20.2 |  | 15.68 | 17 | Water 10 | | a:4553 | 4.66 | 1.0 | 25.4 |  | 14.06 | 18 | Water 11 | | a:4024 | 4.67 | 1.0 | 38.0 |  | 7.47 | 19 | Water 12 | | a:4235 | 4.73 | 1.0 | 33.9 |  | 8.04 |

  
**1^st^ shell coordination angles and distances (*trans* in "blue"; *cys* in "green"; deviations in "orange"):**   
• 1-2(W/W): *101º*; 1-3(W/W): *99º*; 1-4(W/W): 88º; 1-5(W/W): 93º; 1-6(W/W): *168º*; 2-3(W/W): *96º*; 2-4(W/W): *84º*; 2-5(W/W): *162º*; 2-6(W/W): 86º; 3-4(W/W): 173º; 3-5(W/W): 91º; 3-6(W/W): 89º; 4-5(W/W): 87º; 4-6(W/W): *84º*; 5-6(W/W): *78º*;   
• 1-2(W/W): *3.3Å*; 1-3(W/W): *3.3Å*; 1-4(W/W): 3.0Å; 1-5(W/W): 3.1Å; 1-6(W/W): *4.4Å*; 2-3(W/W): *3.2Å*; 2-4(W/W): 2.9Å; 2-5(W/W): *4.3Å*; 2-6(W/W): 3.0Å; 3-4(W/W): *4.4Å*; 3-5(W/W): 3.1Å; 3-6(W/W): 3.1Å; 4-5(W/W): 3.0Å; 4-6(W/W): 3.0Å; 5-6(W/W): 2.8Å;   
  
**2.19 ± 0.06: the average coodination distance of the 1^st^ six ligands is Mg^2+^ compatible**  
**12: Number of potential Water contacts**   
**6 : Number of second shell contacts involving water or ions**   
**50%: Percentage of satisfied second shell Water contacts**   
• Contact(s): /O_w_/O_w_/O_w_/O_w_/O_w_/O_w_/  
• Contact code: 6O_w_  
→ Slightly distorded coord. octahedron (6.8º) ←  
Go to ToC

### MG:a:3298

|  | Res. type | Atom | Residue | Distance | Occupancy | B-factor | Closest atom(s) to a 1^st^ shell water | EM\_dens. |
| --- | --- | --- | --- | --- | --- | --- | --- | --- |
| **Ion** | | | |  | 1.0 | 31.8 |  | 21.50 |
 1 | C | OP2 | a:2252 | 1.95 | 1.0 | 27.5 |  | 13.93 | 2 | Water 1 | | a:7678 | 2.04 | 1.0 | 30.6 |  | 15.50 | 3 | U | O4 | a:2253 | 2.05 | 1.0 | 29.3 |  | 14.91 | 4 | Water 2 | | a:7146 | 2.07 | 1.0 | 28.6 |  | 14.12 | 5 | Water 3 | | a:4519 | 2.38 | 1.0 | 29.3 | 2.63 [C:OP1]a:2070 // 2.93 [G:OP1]a:2257 | 14.51 | 6 | C | OP1 | a:2070 | 4.00 | 1.0 | 31.4 |  | 9.86 | 7 | Water 4 | | a:7538 | 4.13 | 1.0 | 32.7 | 2.84 [G:OP1]a:2257 | 12.66 | 8 | C | OP1 | a:2252 | 4.16 | 1.0 | 27.6 |  | 12.25 | 9 | Water 5 | | a:8977 | 4.17 | 1.0 | 40.2 | 2.99 [C:OP1]a:2252 | 5.14 | 10 | A | O3' | a:2251 | 4.22 | 1.0 | 31.8 |  | 13.32 | 11 | C | O5' | a:2252 | 4.31 | 1.0 | 33.2 |  | 13.13 | 12 | A | O5' | a:2251 | 4.35 | 1.0 | 30.6 |  | 13.28 | 13 | Water 6 | | a:7873 | 4.55 | 1.0 | 37.2 |  | 7.76 | 14 | G | OP1 | a:2257 | 4.56 | 1.0 | 28.3 |  | 10.73 | 15 | Water 7 | | a:6535 | 4.63 | 1.0 | 40.8 |  | 8.30 |

  
**1^st^ shell coordination angles and distances (*trans* in "blue"; *cys* in "green"; deviations in "orange"):**   
• 1-2(OP2/W): *96º*; 1-3(OP2/O4): *99º*; 1-4(OP2/W): 89º; 2-3(W/O4): 88º; 2-4(W/W): 89º; 3-4(O4/W): 172º;   
• 1-2(OP2/W): 3.0Å; 1-3(OP2/O4): 3.0Å; 1-4(OP2/W): 2.8Å; 2-3(W/O4): 2.9Å; 2-4(W/W): 2.9Å; 3-4(O4/W): 4.1Å;   
  
**2.42 ± 0.72: the average coodination distance of the 1^st^ six ligands is Na^+^** compatible  
**6: Number of potential Water contacts**   
**2 : Number of second shell contacts involving water or ions**   
**33%: Percentage of satisfied second shell Water contacts**   
• Contact(s): /O4/O_w_/O_w_/OP2/  
• Contact code: O_ph_O_b_2O_w_  
• Contact(s) in the exclusion zone: /O_w_/  
→ Missing octahedral coordination for MG ←  
Ooctahedral coordination (4.3º)   
Go to ToC

### MG:a:3299

|  | Res. type | Atom | Residue | Distance | Occupancy | B-factor | Closest atom(s) to a 1^st^ shell water | EM\_dens. |
| --- | --- | --- | --- | --- | --- | --- | --- | --- |
| **Ion** | | | |  | 1.0 | 30.3 |  | 19.58 |
 1 | A | OP1 | a:2272 | 1.88 | 1.0 | 37.2 |  | 9.02 | 2 | Water 1 | | a:5058 | 2.15 | 1.0 | 26.0 | 2.68 [A:OP2]a:2271 | 15.54 | 3 | Water 2 | | a:9417 | 2.19 | 1.0 | 29.3 |  | 13.43 | 4 | Water 3 | | a:6625 | 2.20 | 1.0 | 23.5 | 2.78 [G:O6]a:2275 | 14.43 | 5 | G | OP1 | a:2273 | 2.22 | 1.0 | 30.6 |  | 10.04 | 6 | Water 4 | | a:6695 | 2.28 | 1.0 | 30.3 | 2.79 [A:N3]a:2271 | 13.33 | 7 | A | OP2 | a:2272 | 3.98 | 1.0 | 25.2 |  | 12.59 | 8 | Water 5 | | a:8258 | 4.06 | 1.0 | 31.9 | 2.94 [G:O5']a:2273 | 11.64 | 9 | A | O3' | a:2272 | 4.14 | 1.0 | 28.8 |  | 11.88 | 10 | Water 6 | | a:6607 | 4.16 | 1.0 | 28.2 | 2.78 [G:O6]a:2275 | 13.23 | 11 | A | O3' | a:2271 | 4.19 | 1.0 | 30.7 |  | 10.95 | 12 | Water 7 | | a:10143 | 4.20 | 1.0 | 34.6 |  | 7.40 | 13 | A | O5' | a:2272 | 4.29 | 1.0 | 30.0 |  | 11.79 | 14 | G | OP2 | a:2273 | 4.30 | 1.0 | 30.9 |  | 11.39 | 15 | G | O6 | a:2275 | 4.33 | 1.0 | 28.8 |  | 12.81 | 16 | A | N3 | a:2271 | 4.37 | 1.0 | 19.5 |  | 12.58 | 17 | Water 8 | | a:5960 | 4.39 | 1.0 | 26.0 | 2.74 [A:OP2]a:2272 // 2.87 [G:OP2]a:2273 | 11.02 | 18 | Water 9 | | a:4666 | 4.39 | 1.0 | 28.9 |  | 11.94 | 19 | A | OP2 | a:2271 | 4.48 | 1.0 | 26.1 |  | 11.19 | 20 | Water 10 | | a:5212 | 4.50 | 1.0 | 38.9 |  | 6.97 | 21 | Water 11 | | a:6775 | 4.51 | 1.0 | 33.5 |  | 9.60 | 22 | G | O5' | a:2273 | 4.65 | 1.0 | 33.4 |  | 10.54 |

  
**1^st^ shell coordination angles and distances (*trans* in "blue"; *cys* in "green"; deviations in "orange"):**   
• 1-2(OP1/W): 87º; 1-3(OP1/W): *101º*; 1-4(OP1/W): *170º*; 1-5(OP1/OP1): 94º; 1-6(OP1/W): 89º; 2-3(W/W): 172º; 2-4(W/W): *83º*; 2-5(W/OP1): 89º; 2-6(W/W): 92º; 3-4(W/W): 89º; 3-5(W/OP1): 88º; 3-6(W/W): 90º; 4-5(W/OP1): 90º; 4-6(W/W): 87º; 5-6(OP1/W): 177º;   
• 1-2(OP1/W): 2.8Å; 1-3(OP1/W): 3.1Å; 1-4(OP1/W): 4.1Å; 1-5(OP1/OP1): 3.0Å; 1-6(OP1/W): 2.9Å; 2-3(W/W): *4.3Å*; 2-4(W/W): 2.9Å; 2-5(W/OP1): 3.1Å; 2-6(W/W): 3.2Å; 3-4(W/W): 3.1Å; 3-5(W/OP1): 3.1Å; 3-6(W/W): 3.2Å; 4-5(W/OP1): 3.1Å; 4-6(W/W): 3.1Å; 5-6(OP1/W): *4.5Å*;   
  
**2.15 ± 0.13: the average coodination distance of the 1^st^ six ligands is Mg^2+^ compatible**  
**8: Number of potential Water contacts**   
**3 : Number of second shell contacts involving water or ions**   
**38%: Percentage of satisfied second shell Water contacts**   
• Contact(s): /O_w_/O_w_/O_w_/O_w_/OP1/  
• Contact code: O_ph_4O_w_  
• Contact(s) in the exclusion zone: /OP1/  
→ Missing octahedral coordination for MG ←  
Ooctahedral coordination (3.7º)   
Go to ToC

### MG:a:3300

|  | Res. type | Atom | Residue | Distance | Occupancy | B-factor | Closest atom(s) to a 1^st^ shell water | EM\_dens. |
| --- | --- | --- | --- | --- | --- | --- | --- | --- |
| **Ion** | | | |  | 1.0 | 29.6 |  | 18.16 |
 1 | A | OP2 | a:2274 | 1.96 | 1.0 | 29.7 |  | 10.37 | 2 | Water 1 | | a:9403 | 2.03 | 1.0 | 29.2 | 2.81 [K:K]a:3345 | 10.84 | 3 | Water 2 | | a:6319 | 2.11 | 1.0 | 27.3 | 2.76 [A:OP2]a:2271 | 13.44 | 4 | Water 3 | | a:6607 | 2.11 | 1.0 | 28.2 | 2.78 [G:O6]a:2275 | 13.23 | 5 | Water 4 | | a:5823 | 2.20 | 1.0 | 29.6 | 2.73 [G:OP2]a:2275 | 12.78 | 6 | Water 5 | | a:4377 | 2.45 | 1.0 | 29.1 | 2.62 [U:O4]a:2276 // 2.91 [K:K]a:3345 | 13.57 | 7 | A | OP1 | a:2274 | 3.87 | 1.0 | 27.7 |  | 10.81 | 8 | A | O5' | a:2274 | 3.89 | 1.0 | 34.2 |  | 12.63 | 9 | **K** | K | a:3345 | 3.95 | 1.0 | 30.0 |  | 16.17 | 10 | A | OP2 | a:2271 | 4.03 | 1.0 | 26.1 |  | 11.19 | 11 | Water 6 | | a:5058 | 4.05 | 1.0 | 26.0 | 2.68 [A:OP2]a:2271 | 15.54 | 12 | Water 7 | | a:8258 | 4.09 | 1.0 | 31.9 |  | 11.64 | 13 | Water 8 | | a:7677 | 4.14 | 1.0 | 33.7 |  | 11.63 | 14 | Water 9 | | a:7552 | 4.32 | 1.0 | 36.9 | 2.85 [G:OP2]a:2275 | 6.95 | 15 | Water 10 | | a:8188 | 4.32 | 1.0 | 38.9 | 2.90 [A:OP1]a:2274 | 6.09 | 16 | G | N7 | a:2275 | 4.32 | 1.0 | 20.1 |  | 13.48 | 17 | U | O4 | a:2276 | 4.34 | 1.0 | 29.2 |  | 12.12 | 18 | Water 11 | | a:6622 | 4.36 | 1.0 | 32.6 | 2.75 [K:K]a:3345 | 9.36 | 19 | G | O3' | a:2273 | 4.47 | 1.0 | 26.1 |  | 13.35 | 20 | Water 12 | | a:10206 | 4.54 | 1.0 | 42.1 |  | 3.28 | 21 | G | OP2 | a:2275 | 4.61 | 1.0 | 28.7 |  | 9.77 | 22 | G | O6 | a:2275 | 4.67 | 1.0 | 28.8 |  | 12.81 |

  
**1^st^ shell coordination angles and distances (*trans* in "blue"; *cys* in "green"; deviations in "orange"):**   
• 1-2(OP2/W): *101º*; 1-3(OP2/W): 89º; 1-4(OP2/W): *95º*; 1-5(OP2/W): *100º*; 2-3(W/W): 89º; 2-4(W/W): *163º*; 2-5(W/W): *84º*; 3-4(W/W): *96º*; 3-5(W/W): *170º*; 4-5(W/W): 88º;   
• 1-2(OP2/W): 3.1Å; 1-3(OP2/W): 2.9Å; 1-4(OP2/W): 3.0Å; 1-5(OP2/W): 3.2Å; 2-3(W/W): 2.9Å; 2-4(W/W): 4.1Å; 2-5(W/W): 2.8Å; 3-4(W/W): 3.1Å; 3-5(W/W): *4.3Å*; 4-5(W/W): 3.0Å;   
  
**2.14 ± 0.15: the average coodination distance of the 1^st^ six ligands is Mg^2+^ compatible**  
**10: Number of potential Water contacts**   
**6 : Number of second shell contacts involving water or ions**   
**60%: Percentage of satisfied second shell Water contacts**   
• Contact(s): /O_w_/O_w_/O_w_/O_w_/OP2/  
• Contact code: O_ph_4O_w_  
• Contact(s) in the exclusion zone: /O_w_/  
→ Missing octahedral coordination for MG ←  
→ Slightly distorded coord. octahedron (6.9º) ←  
Go to ToC

### MG:a:3301

|  | Res. type | Atom | Residue | Distance | Occupancy | B-factor | Closest atom(s) to a 1^st^ shell water | EM\_dens. |
| --- | --- | --- | --- | --- | --- | --- | --- | --- |
| **Ion** | | | |  | 1.0 | 40.4 |  | 15.27 |
 1 | Water 1 | | a:5365 | 1.95 | 1.0 | 32.9 | 2.70 [G:O6]a:2458 | 9.00 | 2 | Water 2 | | a:8998 | 1.97 | 1.0 | 31.3 | 2.99 [C:OP2]a:2456 | 8.57 | 3 | Water 3 | | a:9949 | 2.02 | 1.0 | 34.2 |  | 9.29 | 4 | Water 4 | | a:7803 | 2.13 | 1.0 | 29.4 | 2.86 [G:N7]a:2458 | 11.43 | 5 | Water 5 | | a:6277 | 2.16 | 1.0 | 30.9 | 2.76 [A:N1]a:2501 | 10.92 | 6 | Water 6 | | a:6148 | 2.33 | 1.0 | 31.9 | 2.75 [A:N7]a:2457 | 10.30 | 7 | Water 7 | | a:5515 | 3.89 | 1.0 | 35.6 | 2.79 [C:OP2]a:2456 | 9.00 | 8 | Water 8 | | a:5277 | 3.97 | 1.0 | 37.6 | 2.75 [A:OP2]a:2457 | 10.33 | 9 | A | N7 | a:2457 | 4.21 | 1.0 | 25.0 |  | 14.10 | 10 | Water 9 | | a:4432 | 4.21 | 1.0 | 37.4 | 2.95 [C:OP2]a:2456 | 6.10 | 11 | Water 10 | | a:8442 | 4.29 | 1.0 | 40.7 | 2.92 [C:N4]a:2500 | 6.18 | 12 | C | OP2 | a:2456 | 4.40 | 1.0 | 38.0 |  | 9.21 | 13 | G | O6 | a:2458 | 4.46 | 1.0 | 25.0 |  | 16.18 | 14 | A | N1 | a:2501 | 4.47 | 1.0 | 24.0 |  | 12.13 | 15 | Water 11 | | a:5603 | 4.51 | 1.0 | 40.2 | 2.72 [G:O6]a:2459 | 6.79 | 16 | G | O6 | a:2459 | 4.51 | 1.0 | 24.0 |  | 15.27 | 17 | Water 12 | | a:10279 | 4.54 | 1.0 | 39.9 |  | 5.70 | 18 | Water 13 | | a:9415 | 4.61 | 1.0 | 38.5 |  | 6.40 | 19 | G | N7 | a:2458 | 4.61 | 1.0 | 19.4 |  | 14.35 | 20 | A | OP2 | a:2457 | 4.73 | 1.0 | 32.3 |  | 10.61 |

  
**1^st^ shell coordination angles and distances (*trans* in "blue"; *cys* in "green"; deviations in "orange"):**   
• 1-2(W/W): 173º; 1-3(W/W): 91º; 1-4(W/W): *82º*; 1-5(W/W): 87º; 2-3(W/W): 94º; 2-4(W/W): 94º; 2-5(W/W): *97º*; 3-4(W/W): 93º; 3-5(W/W): 94º; 4-5(W/W): *167º*;   
• 1-2(W/W): 3.9Å; 1-3(W/W): 2.8Å; 1-4(W/W): 2.7Å; 1-5(W/W): 2.8Å; 2-3(W/W): 2.9Å; 2-4(W/W): 3.0Å; 2-5(W/W): 3.1Å; 3-4(W/W): 3.0Å; 3-5(W/W): 3.1Å; 4-5(W/W): *4.3Å*;   
  
**2.10 ± 0.13: the average coodination distance of the 1^st^ six ligands is Mg^2+^ compatible**  
**12: Number of potential Water contacts**   
**5 : Number of second shell contacts involving water or ions**   
**42%: Percentage of satisfied second shell Water contacts**   
• Contact(s): /O_w_/O_w_/O_w_/O_w_/O_w_/  
• Contact code: 5O_w_  
• Contact(s) in the exclusion zone: /O_w_/  
→ Missing octahedral coordination for MG ←  
→ Slightly distorded coord. octahedron (5.5º) ←  
Go to ToC

### MG:a:3302

|  | Res. type | Atom | Residue | Distance | Occupancy | B-factor | Closest atom(s) to a 1^st^ shell water | EM\_dens. |
| --- | --- | --- | --- | --- | --- | --- | --- | --- |
| **Ion** | | | |  | 1.0 | 28.7 |  | 23.55 |
 1 | Water 1 | | a:9526 | 1.96 | 1.0 | 29.4 |  | 11.88 | 2 | C | O2 | a:2503 | 1.97 | 1.0 | 22.6 |  | 13.60 | 3 | Water 2 | | a:5729 | 1.98 | 1.0 | 23.6 | 2.73 [G:OP1]a:2036 // 2.89 [G:N3]a:2458 | 15.63 | 4 | Water 3 | | a:6789 | 2.31 | 1.0 | 25.5 | 2.83 [C:N3]a:2503 | 14.49 | 5 | Water 4 | | a:4176 | 2.31 | 1.0 | 24.1 | 2.60 [G:N7]a:2036 | 13.15 | 6 | Water 5 | | a:5673 | 2.33 | 1.0 | 25.5 | 2.72 [U:O2]a:2504 | 16.33 | 7 | C | N3 | a:2503 | 3.69 | 1.0 | 18.1 |  | 18.05 | 8 | G | N7 | a:2036 | 4.13 | 1.0 | 23.2 |  | 15.39 | 9 | C | N3 | a:2059 | 4.16 | 1.0 | 23.5 |  | 15.60 | 10 | C | N1 | a:2503 | 4.18 | 1.0 | 19.2 |  | 17.06 | 11 | Water 6 | | a:4624 | 4.20 | 1.0 | 29.2 |  | 10.19 | 12 | G | OP1 | a:2036 | 4.21 | 1.0 | 29.0 |  | 10.09 | 13 | Water 7 | | a:5666 | 4.27 | 1.0 | 30.1 |  | 9.11 | 14 | G | N3 | a:2458 | 4.41 | 1.0 | 17.2 |  | 17.88 | 15 | U | O2 | a:2504 | 4.62 | 1.0 | 31.7 |  | 12.16 | 16 | C | O2' | a:2503 | 4.67 | 1.0 | 29.1 |  | 14.03 | 17 | Water 8 | | a:7402 | 4.79 | 1.0 | 35.3 |  | 10.31 |

  
**1^st^ shell coordination angles and distances (*trans* in "blue"; *cys* in "green"; deviations in "orange"):**   
• 1-2(W/O2): *105º*; 1-3(W/W): *97º*; 2-3(O2/W): 94º;   
• 1-2(W/O2): 3.1Å; 1-3(W/W): 2.9Å; 2-3(O2/W): 2.9Å;   
  
**2.14 ± 0.17: the average coodination distance of the 1^st^ six ligands is Mg^2+^ compatible**  
**10: Number of potential Water contacts**   
**5 : Number of second shell contacts involving water or ions**   
**50%: Percentage of satisfied second shell Water contacts**   
• Contact(s): /O2/O_w_/O_w_/  
• Contact code: O_b_2O_w_  
• Contact(s) in the exclusion zone: /O_w_/O_w_/O_w_/  
→ Missing octahedral coordination for MG ←  
Go to ToC

### MG:a:3303

|  | Res. type | Atom | Residue | Distance | Occupancy | B-factor | Closest atom(s) to a 1^st^ shell water | EM\_dens. |
| --- | --- | --- | --- | --- | --- | --- | --- | --- |
| **Ion** | | | |  | 1.0 | 31.5 |  | 17.41 |
 1 | Water 1 | | a:7050 | 1.92 | 1.0 | 32.1 | 2.81 [A:OP2]a:2545 | 9.74 | 2 | Water 2 | | a:8432 | 2.02 | 1.0 | 34.5 | 2.92 [C:OP2]a:2544 | 7.52 | 3 | A | OP2 | a:2522 | 2.14 | 1.0 | 30.7 |  | 11.45 | 4 | Water 3 | | a:7131 | 2.21 | 1.0 | 29.1 | 2.81 [A:O5']a:2522 | 9.51 | 5 | Water 4 | | a:4616 | 2.22 | 1.0 | 25.9 | 2.64 [U:OP2]a:2523 | 13.47 | 6 | Water 5 | | a:4110 | 2.22 | 1.0 | 28.5 | 2.59 [A:N7]a:2545 | 10.81 | 7 | A | O5' | a:2522 | 3.62 | 1.0 | 30.4 |  | 10.20 | 8 | A | OP2 | a:2545 | 3.92 | 1.0 | 27.8 |  | 10.60 | 9 | Water 6 | | a:8528 | 3.96 | 1.0 | 41.9 |  | 4.63 | 10 | U | OP2 | a:2523 | 4.05 | 1.0 | 25.6 |  | 12.42 | 11 | C | O3' | a:2521 | 4.07 | 1.0 | 31.8 |  | 11.43 | 12 | A | N7 | a:2545 | 4.20 | 1.0 | 20.1 |  | 15.94 | 13 | Water 7 | | a:6526 | 4.25 | 1.0 | 36.8 | 2.78 [U:OP2]a:2523 | 7.57 | 14 | Water 8 | | a:3406 | 4.30 | 1.0 | 37.6 | 2.27 [C:OP2]a:2544 | 6.03 | 15 | C | OP2 | a:2544 | 4.33 | 1.0 | 31.9 |  | 9.01 | 16 | C | O2' | a:2521 | 4.38 | 1.0 | 28.2 |  | 14.77 | 17 | Water 9 | | a:5943 | 4.50 | 1.0 | 28.7 |  | 12.44 | 18 | A | OP1 | a:2522 | 4.57 | 1.0 | 34.8 |  | 9.14 | 19 | Water 10 | | a:9285 | 4.60 | 1.0 | 40.4 |  | 3.88 | 20 | Water 11 | | a:7973 | 4.66 | 1.0 | 40.9 | 2.88 [A:OP2]a:2545 | 3.10 |

  
**1^st^ shell coordination angles and distances (*trans* in "blue"; *cys* in "green"; deviations in "orange"):**   
• 1-2(W/W): 94º; 1-3(W/OP2): 94º; 1-4(W/W): 174º; 1-5(W/W): *101º*; 1-6(W/W): 94º; 2-3(W/OP2): 93º; 2-4(W/W): *82º*; 2-5(W/W): *164º*; 2-6(W/W): 95º; 3-4(OP2/W): 90º; 3-5(OP2/W): *81º*; 3-6(OP2/W): *169º*; 4-5(W/W): *83º*; 4-6(W/W): *83º*; 5-6(W/W): 89º;   
• 1-2(W/W): 2.9Å; 1-3(W/OP2): 3.0Å; 1-4(W/W): 4.1Å; 1-5(W/W): *3.2Å*; 1-6(W/W): 3.0Å; 2-3(W/OP2): 3.0Å; 2-4(W/W): 2.8Å; 2-5(W/W): 4.2Å; 2-6(W/W): 3.1Å; 3-4(OP2/W): 3.1Å; 3-5(OP2/W): 2.8Å; 3-6(OP2/W): *4.3Å*; 4-5(W/W): 2.9Å; 4-6(W/W): 2.9Å; 5-6(W/W): 3.1Å;   
  
**2.12 ± 0.11: the average coodination distance of the 1^st^ six ligands is Mg^2+^ compatible**  
**10: Number of potential Water contacts**   
**5 : Number of second shell contacts involving water or ions**   
**50%: Percentage of satisfied second shell Water contacts**   
• Contact(s): /O_w_/O_w_/O_w_/O_w_/O_w_/OP2/  
• Contact code: O_ph_5O_w_  
→ Slightly distorded coord. octahedron (6.4º) ←  
Go to ToC

### MG:a:3304

|  | Res. type | Atom | Residue | Distance | Occupancy | B-factor | Closest atom(s) to a 1^st^ shell water | EM\_dens. |
| --- | --- | --- | --- | --- | --- | --- | --- | --- |
| **Ion** | | | |  | 1.0 | 39.7 |  | 15.25 |
 1 | Water 1 | | a:6484 | 2.00 | 1.0 | 38.8 | 2.87 [PSU:OP2]a:2584 | 7.06 | 2 | C | OP2 | a:2615 | 2.06 | 1.0 | 32.2 |  | 9.34 | 3 | Water 2 | | a:6943 | 2.15 | 1.0 | 32.9 | 2.85 [C:OP2]a:2616 | 8.40 | 4 | C | OP1 | a:2583 | 2.18 | 1.0 | 27.0 |  | 12.53 | 5 | Water 3 | | a:9271 | 2.36 | 1.0 | 34.6 |  | 9.35 | 6 | C | OP1 | a:2615 | 3.89 | 1.0 | 35.8 |  | 9.30 | 7 | C | OP2 | a:2583 | 3.91 | 1.0 | 29.6 |  | 12.65 | 8 | C | O5' | a:2583 | 4.08 | 1.0 | 29.9 |  | 12.99 | 9 | Water 4 | | a:9512 | 4.15 | 1.0 | 32.9 |  | 9.07 | 10 | **MG** | MG | a:3054 | 4.18 | 1.0 | 30.4 |  | 16.41 | 11 | PSU | OP2 | a:2584 | 4.24 | 1.0 | 26.5 |  | 17.47 | 12 | G | O2' | a:2580 | 4.25 | 1.0 | 33.8 |  | 11.62 | 13 | **K** | K | a:3266 | 4.25 | 1.0 | 30.0 |  | 9.94 | 14 | C | O5' | a:2615 | 4.30 | 1.0 | 38.8 |  | 10.12 | 15 | C | OP2 | a:2616 | 4.33 | 1.0 | 35.9 |  | 7.26 | 16 | C | O3' | a:2614 | 4.35 | 1.0 | 37.9 |  | 8.87 | 17 | Water 5 | | a:7425 | 4.41 | 1.0 | 27.4 | 2.84 [PSU:OP2]a:2584 | 11.86 | 18 | C | O2' | a:2614 | 4.42 | 1.0 | 32.9 |  | 9.28 | 19 | Water 6 | | d:497 | 4.43 | 1.0 | 35.7 |  | 7.95 | 20 | Water 7 | | a:9272 | 4.52 | 1.0 | 38.0 |  | 8.92 | 21 | G | O3' | a:2582 | 4.64 | 1.0 | 31.8 |  | 13.56 | 22 | Water 8 | | a:9540 | 4.72 | 1.0 | 38.2 | 2.80 [MG:MG]a:3054 | 6.06 |

  
**1^st^ shell coordination angles and distances (*trans* in "blue"; *cys* in "green"; deviations in "orange"):**   
• 1-2(W/OP2): 86º; 1-3(W/W): *76º*; 1-4(W/OP1): *95º*; 2-3(OP2/W): 91º; 2-4(OP2/OP1): 171º; 3-4(W/OP1): *81º*;   
• 1-2(W/OP2): 2.8Å; 1-3(W/W): 2.6Å; 1-4(W/OP1): 3.1Å; 2-3(OP2/W): 3.0Å; 2-4(OP2/OP1): *4.2Å*; 3-4(W/OP1): 2.8Å;   
  
**2.44 ± 0.66: the average coodination distance of the 1^st^ six ligands is Na^+^** compatible  
**6: Number of potential Water contacts**   
**2 : Number of second shell contacts involving water or ions**   
**33%: Percentage of satisfied second shell Water contacts**   
• Contact(s): /O_w_/O_w_/OP1/OP2/  
• Contact code: *cis-*2O_ph_2O_w_  
• Contact(s) in the exclusion zone: /O_w_/  
→ Missing octahedral coordination for MG ←  
→ Slightly distorded coord. octahedron (7.1º) ←  
Go to ToC

### MG:a:3305

|  | Res. type | Atom | Residue | Distance | Occupancy | B-factor | Closest atom(s) to a 1^st^ shell water | EM\_dens. |
| --- | --- | --- | --- | --- | --- | --- | --- | --- |
| **Ion** | | | |  | 1.0 | 26.2 |  | 23.23 |
 1 | Water 1 | | a:3786 | 1.90 | 1.0 | 30.0 | 2.53 [A:OP1]a:2591 // 2.99 [K:K]a:3117 | 10.91 | 2 | G | OP2 | a:2592 | 1.96 | 1.0 | 22.7 |  | 13.56 | 3 | Water 2 | | a:8754 | 2.04 | 1.0 | 25.8 | 2.97 [G:N7]a:2592 | 14.14 | 4 | Water 3 | | a:4992 | 2.16 | 1.0 | 27.6 | 2.67 [A:OP2]a:2593 | 13.22 | 5 | Water 4 | | c:478 | 2.20 | 1.0 | 26.2 | 2.78 [GLU:OE2]c:236 | 13.14 | 6 | Water 5 | | c:407 | 2.38 | 1.0 | 31.6 | 2.45 [GLU:OE1]c:236 | 10.43 | 7 | G | OP1 | a:2592 | 3.78 | 1.0 | 22.1 |  | 15.37 | 8 | A | OP1 | a:2591 | 3.93 | 1.0 | 33.3 |  | 10.70 | 9 | **K** | K | a:3117 | 3.96 | 1.0 | 30.0 |  | 13.58 | 10 | G | O5' | a:2592 | 4.00 | 1.0 | 23.3 |  | 16.72 | 11 | Water 6 | | c:589 | 4.03 | 1.0 | 46.3 |  | 4.93 | 12 | GLU | OE1 | c:236 | 4.19 | 1.0 | 46.1 |  | 4.35 | 13 | GLU | OE2 | c:236 | 4.24 | 1.0 | 43.8 |  | 6.80 | 14 | A | OP2 | a:2593 | 4.36 | 1.0 | 24.5 |  | 18.24 | 15 | Water 7 | | a:9064 | 4.40 | 1.0 | 40.1 |  | 5.10 | 16 | A | O3' | a:2591 | 4.41 | 1.0 | 28.5 |  | 14.17 | 17 | G | N7 | a:2592 | 4.45 | 1.0 | 17.8 |  | 17.16 | 18 | A | N7 | a:2593 | 4.46 | 1.0 | 16.6 |  | 17.83 | 19 | A | O5' | a:2591 | 4.58 | 1.0 | 30.9 |  | 12.36 |

  
**1^st^ shell coordination angles and distances (*trans* in "blue"; *cys* in "green"; deviations in "orange"):**   
• 1-2(W/OP2): *98º*; 1-3(W/W): 90º; 1-4(W/W): *166º*; 1-5(W/W): *102º*; 2-3(OP2/W): *96º*; 2-4(OP2/W): 95º; 2-5(OP2/W): 90º; 3-4(W/W): *85º*; 3-5(W/W): *166º*; 4-5(W/W): *81º*;   
• 1-2(W/OP2): 2.9Å; 1-3(W/W): 2.8Å; 1-4(W/W): 4.0Å; 1-5(W/W): *3.2Å*; 2-3(OP2/W): 3.0Å; 2-4(OP2/W): 3.0Å; 2-5(OP2/W): 2.9Å; 3-4(W/W): 2.8Å; 3-5(W/W): *4.2Å*; 4-5(W/W): 2.8Å;   
  
**2.11 ± 0.16: the average coodination distance of the 1^st^ six ligands is Mg^2+^ compatible**  
**10: Number of potential Water contacts**   
**6 : Number of second shell contacts involving water or ions**   
**60%: Percentage of satisfied second shell Water contacts**   
• Contact(s): /O_w_/O_w_/O_w_/O_w_/OP2/  
• Contact code: O_ph_4O_w_  
• Contact(s) in the exclusion zone: /O_w_/  
→ Missing octahedral coordination for MG ←  
→ Slightly distorded coord. octahedron (7.3º) ←  
Go to ToC

### MG:a:3306

|  | Res. type | Atom | Residue | Distance | Occupancy | B-factor | Closest atom(s) to a 1^st^ shell water | EM\_dens. |
| --- | --- | --- | --- | --- | --- | --- | --- | --- |
| **Ion** | | | |  | 1.0 | 31.2 |  | 17.19 |
 1 | A | OP2 | a:2715 | 1.89 | 1.0 | 24.9 |  | 13.69 | 2 | G | OP2 | a:2718 | 2.00 | 1.0 | 28.0 |  | 11.51 | 3 | Water 1 | | a:5604 | 2.04 | 1.0 | 29.3 | 2.72 [C:OP2]a:2714 | 8.58 | 4 | Water 2 | | a:8228 | 2.15 | 1.0 | 29.0 | 2.90 [C:OP1]a:2714 | 11.45 | 5 | Water 3 | | a:8333 | 2.23 | 1.0 | 38.9 |  | 8.41 | 6 | A | OP1 | a:2715 | 3.65 | 1.0 | 32.4 |  | 10.33 | 7 | G | OP1 | a:2718 | 3.82 | 1.0 | 28.0 |  | 11.36 | 8 | C | O3' | a:2714 | 4.19 | 1.0 | 28.0 |  | 12.65 | 9 | A | O5' | a:2715 | 4.21 | 1.0 | 27.0 |  | 14.81 | 10 | U | O3' | a:2717 | 4.28 | 1.0 | 34.4 |  | 10.66 | 11 | G | O5' | a:2718 | 4.36 | 1.0 | 26.6 |  | 13.61 | 12 | Water 4 | | m:247 | 4.37 | 1.0 | 43.7 |  | 5.26 | 13 | U | O5' | a:2717 | 4.38 | 1.0 | 36.2 |  | 9.69 | 14 | C | O5' | a:2714 | 4.45 | 1.0 | 29.2 |  | 13.89 | 15 | C | OP2 | a:2714 | 4.48 | 1.0 | 26.1 |  | 15.26 | 16 | Water 5 | | m:252 | 4.60 | 1.0 | 44.8 |  | 4.22 | 17 | C | OP1 | a:2714 | 4.64 | 1.0 | 28.3 |  | 11.37 | 18 | Water 6 | | a:5239 | 4.71 | 1.0 | 25.7 | 2.69 [G:OP1]a:2718 | 13.90 | 19 | U | OP2 | a:2717 | 4.72 | 1.0 | 38.8 |  | 8.65 |

  
**1^st^ shell coordination angles and distances (*trans* in "blue"; *cys* in "green"; deviations in "orange"):**   
• 1-2(OP2/OP2): *100º*; 1-3(OP2/W): 92º; 1-4(OP2/W): 92º; 1-5(OP2/W): 173º; 2-3(OP2/W): 93º; 2-4(OP2/W): *167º*; 2-5(OP2/W): 87º; 3-4(W/W): *82º*; 3-5(W/W): 90º; 4-5(W/W): *81º*;   
• 1-2(OP2/OP2): 3.0Å; 1-3(OP2/W): 2.8Å; 1-4(OP2/W): 2.9Å; 1-5(OP2/W): 4.1Å; 2-3(OP2/W): 2.9Å; 2-4(OP2/W): 4.1Å; 2-5(OP2/W): 2.9Å; 3-4(W/W): 2.7Å; 3-5(W/W): 3.0Å; 4-5(W/W): 2.8Å;   
  
**2.32 ± 0.60: the average coodination distance of the 1^st^ six ligands is Na^+^** compatible  
**6: Number of potential Water contacts**   
**2 : Number of second shell contacts involving water or ions**   
**33%: Percentage of satisfied second shell Water contacts**   
• Contact(s): /O_w_/O_w_/O_w_/OP2/  
• Contact code: O_ph_3O_w_  
• Contact(s) in the exclusion zone: /OP2/  
→ Missing octahedral coordination for MG ←  
→ Slightly distorded coord. octahedron (5.8º) ←  
Go to ToC

### MG:a:3307

|  | Res. type | Atom | Residue | Distance | Occupancy | B-factor | Closest atom(s) to a 1^st^ shell water | EM\_dens. |
| --- | --- | --- | --- | --- | --- | --- | --- | --- |
| **Ion** | | | |  | 1.0 | 42.6 |  | 7.75 |
 1 | Water 1 | | a:7179 | 1.85 | 1.0 | 40.7 | 2.85 [G:O6]a:338 | 5.36 | 2 | Water 2 | | a:8416 | 1.93 | 1.0 | 44.9 | 2.92 [U:O4]a:339 | 4.75 | 3 | A | OP2 | a:299 | 2.17 | 1.0 | 42.9 |  | 4.92 | 4 | Water 3 | | a:4089 | 2.27 | 1.0 | 39.4 | 2.59 [G:N7]a:338 | 5.90 | 5 | A | OP1 | a:299 | 4.16 | 1.0 | 46.5 |  | 3.88 | 6 | Water 4 | | a:8122 | 4.18 | 1.0 | 42.7 |  | 4.05 | 7 | G | N7 | a:338 | 4.28 | 1.0 | 32.6 |  | 8.28 | 8 | G | O3' | a:298 | 4.30 | 1.0 | 45.3 |  | 6.18 | 9 | U | O4 | a:339 | 4.35 | 1.0 | 38.6 |  | 6.20 | 10 | Water 5 | | a:4028 | 4.40 | 1.0 | 44.3 |  | 3.74 | 11 | G | O6 | a:338 | 4.44 | 1.0 | 41.0 |  | 5.94 | 12 | A | N7 | a:299 | 4.46 | 1.0 | 41.5 |  | 7.37 | 13 | A | O5' | a:299 | 4.63 | 1.0 | 43.4 |  | 5.22 |

  
**1^st^ shell coordination angles and distances (*trans* in "blue"; *cys* in "green"; deviations in "orange"):**   
• 1-2(W/W): 86º; 1-3(W/OP2): 89º; 1-4(W/W): 86º; 2-3(W/OP2): *97º*; 2-4(W/W): *83º*; 3-4(OP2/W): 174º;   
• 1-2(W/W): 2.6Å; 1-3(W/OP2): 2.8Å; 1-4(W/W): 2.8Å; 2-3(W/OP2): 3.1Å; 2-4(W/W): 2.8Å; 3-4(OP2/W): *4.4Å*;   
  
**2.76 ± 1.01: the average coodination distance of the 1^st^ six ligands is K^+^ compatible**  
**8: Number of potential Water contacts**   
**3 : Number of second shell contacts involving water or ions**   
**38%: Percentage of satisfied second shell Water contacts**   
• Contact(s): /O_w_/O_w_/OP2/  
• Contact code: O_ph_2O_w_  
• Contact(s) in the exclusion zone: /O_w_/  
→ Missing octahedral coordination for MG ←  
Ooctahedral coordination (4.9º)   
Go to ToC

### MG:a:3308

|  | Res. type | Atom | Residue | Distance | Occupancy | B-factor | Closest atom(s) to a 1^st^ shell water | EM\_dens. |
| --- | --- | --- | --- | --- | --- | --- | --- | --- |
| **Ion** | | | |  | 1.0 | 48.5 |  | 9.39 |
 1 | Water 1 | | a:3639 | 1.95 | 1.0 | 45.0 |  | 4.41 | 2 | A | OP2 | a:422 | 1.97 | 1.0 | 40.1 |  | 5.55 | 3 | Water 2 | | a:4344 | 2.00 | 1.0 | 38.6 | 2.62 [G:N7]a:370 | 6.29 | 4 | G | OP2 | a:370 | 2.06 | 1.0 | 46.8 |  | 4.53 | 5 | Water 3 | | a:4200 | 2.14 | 1.0 | 38.3 | 2.60 [A:OP2]a:423 | 7.34 | 6 | A | O5' | a:422 | 3.82 | 1.0 | 37.3 |  | 6.42 | 7 | G | N7 | a:370 | 3.91 | 1.0 | 41.2 |  | 6.56 | 8 | U | O2' | a:369 | 4.16 | 1.0 | 63.3 |  | 2.49 | 9 | A | OP1 | a:422 | 4.23 | 1.0 | 39.6 |  | 6.56 | 10 | U | O3' | a:369 | 4.25 | 1.0 | 61.0 |  | 4.11 | 11 | Water 4 | | a:7634 | 4.30 | 1.0 | 41.2 | 2.94 [A:N7]a:422 | 5.32 | 12 | A | N7 | a:423 | 4.35 | 1.0 | 31.6 |  | 10.08 | 13 | G | O5' | a:370 | 4.36 | 1.0 | 50.0 |  | 3.74 | 14 | C | O3' | a:421 | 4.37 | 1.0 | 47.8 |  | 5.10 | 15 | A | OP2 | a:423 | 4.37 | 1.0 | 35.6 |  | 8.14 | 16 | C | O2' | a:421 | 4.39 | 1.0 | 39.6 |  | 6.09 | 17 | G | OP1 | a:370 | 4.43 | 1.0 | 45.3 |  | 5.13 | 18 | A | N7 | a:422 | 4.69 | 1.0 | 28.2 |  | 10.41 | 19 | Water 5 | | a:4700 | 4.74 | 1.0 | 46.4 | 2.65 [A:OP2]a:423 | 4.13 |

  
**1^st^ shell coordination angles and distances (*trans* in "blue"; *cys* in "green"; deviations in "orange"):**   
• 1-2(W/OP2): *98º*; 1-3(W/W): 173º; 1-4(W/OP2): *77º*; 1-5(W/W): *100º*; 2-3(OP2/W): 88º; 2-4(OP2/OP2): 172º; 2-5(OP2/W): 91º; 3-4(W/OP2): *97º*; 3-5(W/W): *84º*; 4-5(OP2/W): *95º*;   
• 1-2(W/OP2): 3.0Å; 1-3(W/W): 3.9Å; 1-4(W/OP2): 2.5Å; 1-5(W/W): 3.1Å; 2-3(OP2/W): 2.8Å; 2-4(OP2/OP2): 4.0Å; 2-5(OP2/W): 2.9Å; 3-4(W/OP2): 3.0Å; 3-5(W/W): 2.8Å; 4-5(OP2/W): 3.1Å;   
  
**2.32 ± 0.67: the average coodination distance of the 1^st^ six ligands is Na^+^** compatible  
**6: Number of potential Water contacts**   
**2 : Number of second shell contacts involving water or ions**   
**33%: Percentage of satisfied second shell Water contacts**   
• Contact(s): /O_w_/O_w_/O_w_/OP2/OP2/  
• Contact code: *trans-*2O_ph_3O_w_  
→ Missing octahedral coordination for MG ←  
→ Slightly distorded coord. octahedron (6.6º) ←  
Go to ToC

### MG:a:3309

|  | Res. type | Atom | Residue | Distance | Occupancy | B-factor | Closest atom(s) to a 1^st^ shell water | EM\_dens. |
| --- | --- | --- | --- | --- | --- | --- | --- | --- |
| **Ion** | | | |  | 1.0 | 32.2 |  | 15.50 |
 1 | Water 1 | | a:6590 | 1.95 | 1.0 | 30.0 | 2.78 [A:N7]a:2411 | 12.29 | 2 | G | OP1 | a:411 | 1.98 | 1.0 | 34.6 |  | 9.36 | 3 | A | OP1 | a:2411 | 2.08 | 1.0 | 28.5 |  | 10.12 | 4 | Water 2 | | a:4586 | 2.10 | 1.0 | 29.8 | 2.64 [A:OP1]a:412 | 11.01 | 5 | Water 3 | | a:8866 | 2.27 | 1.0 | 27.6 | 2.97 [U:O4]a:2412 | 10.50 | 6 | Water 4 | | a:8551 | 2.32 | 1.0 | 30.6 | 2.94 [A:O2']a:2410 | 9.54 | 7 | A | OP2 | a:2411 | 3.88 | 1.0 | 33.3 |  | 9.51 | 8 | G | OP2 | a:411 | 4.05 | 1.0 | 35.3 |  | 8.70 | 9 | A | O3' | a:2410 | 4.06 | 1.0 | 38.2 |  | 8.14 | 10 | A | OP1 | a:412 | 4.12 | 1.0 | 34.2 |  | 7.64 | 11 | Water 5 | | a:7768 | 4.12 | 1.0 | 31.5 |  | 11.63 | 12 | A | O2' | a:2410 | 4.17 | 1.0 | 39.0 |  | 8.64 | 13 | G | O3' | a:410 | 4.18 | 1.0 | 32.5 |  | 11.69 | 14 | A | N7 | a:2411 | 4.30 | 1.0 | 29.8 |  | 10.01 | 15 | Water 6 | | a:4112 | 4.40 | 1.0 | 31.1 |  | 9.00 | 16 | Water 7 | | a:9846 | 4.41 | 1.0 | 37.5 |  | 8.58 | 17 | G | O5' | a:411 | 4.45 | 1.0 | 32.8 |  | 9.59 | 18 | A | O5' | a:2411 | 4.53 | 1.0 | 31.9 |  | 10.31 | 19 | Water 8 | | a:8706 | 4.61 | 1.0 | 36.6 |  | 7.82 | 20 | U | O4 | a:2412 | 4.64 | 1.0 | 34.6 |  | 11.24 | 21 | Water 9 | | a:9118 | 4.65 | 1.0 | 36.0 |  | 8.23 | 22 | Water 10 | | a:7553 | 4.77 | 1.0 | 35.5 |  | 8.54 |

  
**1^st^ shell coordination angles and distances (*trans* in "blue"; *cys* in "green"; deviations in "orange"):**   
• 1-2(W/OP1): 94º; 1-3(W/OP1): 93º; 1-4(W/W): 91º; 1-5(W/W): 87º; 2-3(OP1/OP1): *99º*; 2-4(OP1/W): 89º; 2-5(OP1/W): 173º; 3-4(OP1/W): 171º; 3-5(OP1/W): 87º; 4-5(W/W): *84º*;   
• 1-2(W/OP1): 2.9Å; 1-3(W/OP1): 2.9Å; 1-4(W/W): 2.9Å; 1-5(W/W): 2.9Å; 2-3(OP1/OP1): 3.1Å; 2-4(OP1/W): 2.9Å; 2-5(OP1/W): *4.2Å*; 3-4(OP1/W): 4.2Å; 3-5(OP1/W): 3.0Å; 4-5(W/W): 2.9Å;   
  
**2.12 ± 0.14: the average coodination distance of the 1^st^ six ligands is Mg^2+^ compatible**  
**8: Number of potential Water contacts**   
**4 : Number of second shell contacts involving water or ions**   
**50%: Percentage of satisfied second shell Water contacts**   
• Contact(s): /O_w_/O_w_/O_w_/OP1/OP1/  
• Contact code: *cis-*2O_ph_3O_w_  
• Contact(s) in the exclusion zone: /O_w_/  
→ Missing octahedral coordination for MG ←  
Ooctahedral coordination (4.7º)   
Go to ToC

### MG:a:3310

|  | Res. type | Atom | Residue | Distance | Occupancy | B-factor | Closest atom(s) to a 1^st^ shell water | EM\_dens. |
| --- | --- | --- | --- | --- | --- | --- | --- | --- |
| **Ion** | | | |  | 1.0 | 37.0 |  | 19.64 |
 1 | A | OP1 | a:574 | 1.95 | 1.0 | 26.7 |  | 12.39 | 2 | Water 1 | | a:8970 | 2.01 | 1.0 | 32.2 |  | 12.40 | 3 | Water 2 | | a:6874 | 2.12 | 1.0 | 24.4 | 2.80 [A:N7]a:574 | 13.70 | 4 | Water 3 | | a:8545 | 2.32 | 1.0 | 36.4 | 2.94 [A:OP2]a:574 | 8.96 | 5 | Water 4 | | a:3514 | 2.33 | 1.0 | 36.9 | 2.45 [A:OP1]a:985 | 9.64 | 6 | A | OP2 | a:574 | 3.73 | 1.0 | 26.4 |  | 13.43 | 7 | Water 5 | | a:8075 | 3.89 | 1.0 | 43.2 |  | 5.64 | 8 | Water 6 | | a:9694 | 3.96 | 1.0 | 43.5 |  | 5.80 | 9 | Water 7 | | a:5132 | 3.98 | 1.0 | 29.4 |  | 10.88 | 10 | A | O5' | a:574 | 4.07 | 1.0 | 27.7 |  | 15.18 | 11 | Water 8 | | a:8930 | 4.19 | 1.0 | 31.9 | 2.98 [A:OP1]a:985 | 10.58 | 12 | A | N7 | a:574 | 4.34 | 1.0 | 20.2 |  | 17.10 | 13 | Water 9 | | a:5200 | 4.35 | 1.0 | 34.0 |  | 8.50 | 14 | U | O3' | a:573 | 4.35 | 1.0 | 27.0 |  | 14.75 | 15 | A | OP1 | a:985 | 4.37 | 1.0 | 40.4 |  | 7.18 |

  
**1^st^ shell coordination angles and distances (*trans* in "blue"; *cys* in "green"; deviations in "orange"):**   
• 1-2(OP1/W): *98º*; 1-3(OP1/W): 91º; 2-3(W/W): *101º*;   
• 1-2(OP1/W): 3.0Å; 1-3(OP1/W): 2.9Å; 2-3(W/W): 3.2Å;   
  
**2.41 ± 0.61: the average coodination distance of the 1^st^ six ligands is Na^+^** compatible  
**8: Number of potential Water contacts**   
**3 : Number of second shell contacts involving water or ions**   
**38%: Percentage of satisfied second shell Water contacts**   
• Contact(s): /O_w_/O_w_/OP1/  
• Contact code: O_ph_2O_w_  
• Contact(s) in the exclusion zone: /O_w_/O_w_/  
→ Missing octahedral coordination for MG ←  
Go to ToC

### MG:a:3311

|  | Res. type | Atom | Residue | Distance | Occupancy | B-factor | Closest atom(s) to a 1^st^ shell water | EM\_dens. |
| --- | --- | --- | --- | --- | --- | --- | --- | --- |
| **Ion** | | | |  | 1.0 | 25.8 |  | 23.39 |
 1 | Water 1 | | a:4912 | 2.02 | 1.0 | 22.4 | 2.67 [G:N7]a:2039 | 16.71 | 2 | Water 2 | | a:8434 | 2.04 | 1.0 | 25.2 | 2.92 [G:O6]a:2039 | 15.65 | 3 | Water 3 | | a:8202 | 2.11 | 1.0 | 23.4 | 2.90 [A:OP1]a:576 | 15.46 | 4 | Water 4 | | a:8182 | 2.16 | 1.0 | 23.6 | 2.90 [A:N7]a:576 // 2.99 [A:O2']a:2058 | 16.88 | 5 | Water 5 | | a:5065 | 2.17 | 1.0 | 24.8 | 2.68 [C:OP2]a:2059 | 14.58 | 6 | Water 6 | | a:5169 | 2.22 | 1.0 | 28.0 | 2.69 [U:O4]a:2038 | 13.97 | 7 | Water 7 | | a:5268 | 3.94 | 1.0 | 26.0 |  | 15.27 | 8 | Water 8 | | a:10089 | 3.98 | 1.0 | 34.8 |  | 10.91 | 9 | U | O4 | a:2038 | 4.10 | 1.0 | 27.2 |  | 16.79 | 10 | Water 9 | | a:9104 | 4.12 | 1.0 | 26.2 |  | 11.88 | 11 | Water 10 | | a:8227 | 4.15 | 1.0 | 31.0 | 2.90 [A:OP1]a:576 | 12.28 | 12 | Water 11 | | a:6542 | 4.29 | 1.0 | 28.7 |  | 14.62 | 13 | G | N7 | a:2039 | 4.37 | 1.0 | 19.6 |  | 15.88 | 14 | A | N7 | a:576 | 4.37 | 1.0 | 27.4 |  | 12.74 | 15 | C | OP2 | a:2059 | 4.40 | 1.0 | 22.4 |  | 14.43 | 16 | A | OP1 | a:576 | 4.41 | 1.0 | 28.0 |  | 11.74 | 17 | G | O6 | a:2039 | 4.44 | 1.0 | 23.5 |  | 17.91 | 18 | A | O2' | a:2058 | 4.49 | 1.0 | 27.2 |  | 15.23 | 19 | Water 12 | | z:232 | 4.62 | 1.0 | 34.0 |  | 10.94 | 20 | A | O3' | a:2058 | 4.64 | 1.0 | 24.6 |  | 16.59 | 21 | Water 13 | | z:209 | 4.78 | 1.0 | 26.7 |  | 11.99 |

  
**1^st^ shell coordination angles and distances (*trans* in "blue"; *cys* in "green"; deviations in "orange"):**   
• 1-2(W/W): 88º; 1-3(W/W): *98º*; 1-4(W/W): 93º; 1-5(W/W): 175º; 1-6(W/W): *97º*; 2-3(W/W): *100º*; 2-4(W/W): *163º*; 2-5(W/W): *97º*; 2-6(W/W): *82º*; 3-4(W/W): *97º*; 3-5(W/W): *84º*; 3-6(W/W): *164º*; 4-5(W/W): *82º*; 4-6(W/W): *81º*; 5-6(W/W): *80º*;   
• 1-2(W/W): 2.8Å; 1-3(W/W): 3.1Å; 1-4(W/W): 3.0Å; 1-5(W/W): 4.2Å; 1-6(W/W): 3.2Å; 2-3(W/W): 3.2Å; 2-4(W/W): 4.2Å; 2-5(W/W): 3.1Å; 2-6(W/W): 2.8Å; 3-4(W/W): *3.2Å*; 3-5(W/W): 2.9Å; 3-6(W/W): *4.3Å*; 4-5(W/W): 2.8Å; 4-6(W/W): 2.8Å; 5-6(W/W): 2.8Å;   
  
**2.12 ± 0.07: the average coodination distance of the 1^st^ six ligands is Mg^2+^ compatible**  
**12: Number of potential Water contacts**   
**7 : Number of second shell contacts involving water or ions**   
**58%: Percentage of satisfied second shell Water contacts**   
• Contact(s): /O_w_/O_w_/O_w_/O_w_/O_w_/O_w_/  
• Contact code: 6O_w_  
→ Slightly distorded coord. octahedron (8.1º) ←  
Go to ToC

### MG:a:3312

|  | Res. type | Atom | Residue | Distance | Occupancy | B-factor | Closest atom(s) to a 1^st^ shell water | EM\_dens. |
| --- | --- | --- | --- | --- | --- | --- | --- | --- |
| **Ion** | | | |  | 1.0 | 25.6 |  | 22.93 |
 1 | G | O6 | a:1189 | 1.97 | 1.0 | 20.6 |  | 18.43 | 2 | G | OP2 | a:820 | 2.04 | 1.0 | 22.7 |  | 15.67 | 3 | Water 1 | | a:8358 | 2.10 | 1.0 | 24.3 |  | 13.40 | 4 | Water 2 | | a:6394 | 2.20 | 1.0 | 20.5 | 2.77 [G:N7]a:820 | 16.04 | 5 | Water 3 | | a:5636 | 2.23 | 1.0 | 22.2 | 2.72 [G:OP1]a:1188 // 2.81 [G:N7]a:1189 | 14.61 | 6 | Water 4 | | a:3876 | 2.34 | 1.0 | 22.4 | 2.55 [C:OP1]a:819 | 15.48 | 7 | G | N7 | a:1189 | 3.87 | 1.0 | 14.1 |  | 18.87 | 8 | G | O5' | a:820 | 4.07 | 1.0 | 29.5 |  | 12.51 | 9 | G | N7 | a:820 | 4.17 | 1.0 | 15.8 |  | 17.51 | 10 | Water 5 | | a:3949 | 4.19 | 1.0 | 35.0 |  | 9.46 | 11 | Water 6 | | a:9761 | 4.20 | 1.0 | 43.5 |  | 4.98 | 12 | Water 7 | | a:6813 | 4.23 | 1.0 | 40.0 | 2.79 [G:OP1]a:820 | 5.36 | 13 | Water 8 | | a:5298 | 4.29 | 1.0 | 40.0 |  | 5.34 | 14 | Water 9 | | a:4787 | 4.32 | 1.0 | 36.0 | 2.66 [G:OP1]a:1188 | 6.81 | 15 | C | O5' | a:819 | 4.35 | 1.0 | 32.0 |  | 10.37 | 16 | G | OP1 | a:820 | 4.36 | 1.0 | 29.9 |  | 12.47 | 17 | C | O3' | a:819 | 4.43 | 1.0 | 26.9 |  | 14.71 | 18 | C | OP1 | a:819 | 4.45 | 1.0 | 27.2 |  | 13.16 | 19 | G | OP1 | a:1188 | 4.52 | 1.0 | 29.4 |  | 13.60 | 20 | C | OP2 | a:819 | 4.66 | 1.0 | 30.6 |  | 11.38 | 21 | Water 10 | | a:8095 | 4.68 | 1.0 | 21.2 |  | 15.96 | 22 | G | N7 | a:1188 | 4.77 | 1.0 | 16.6 |  | 17.17 |

  
**1^st^ shell coordination angles and distances (*trans* in "blue"; *cys* in "green"; deviations in "orange"):**   
• 1-2(O6/OP2): *97º*; 1-3(O6/W): *98º*; 1-4(O6/W): 87º; 1-5(O6/W): 92º; 2-3(OP2/W): 90º; 2-4(OP2/W): 89º; 2-5(OP2/W): 171º; 3-4(W/W): 175º; 3-5(W/W): 93º; 4-5(W/W): 88º;   
• 1-2(O6/OP2): 3.0Å; 1-3(O6/W): 3.1Å; 1-4(O6/W): 2.9Å; 1-5(O6/W): 3.0Å; 2-3(OP2/W): 2.9Å; 2-4(OP2/W): 3.0Å; 2-5(OP2/W): *4.3Å*; 3-4(W/W): *4.3Å*; 3-5(W/W): 3.1Å; 4-5(W/W): 3.1Å;   
  
**2.15 ± 0.12: the average coodination distance of the 1^st^ six ligands is Mg^2+^ compatible**  
**8: Number of potential Water contacts**   
**4 : Number of second shell contacts involving water or ions**   
**50%: Percentage of satisfied second shell Water contacts**   
• Contact(s): /O6/O_w_/O_w_/O_w_/OP2/  
• Contact code: O_ph_O_b_3O_w_  
• Contact(s) in the exclusion zone: /O_w_/  
→ Missing octahedral coordination for MG ←  
Ooctahedral coordination (4.0º)   
Go to ToC

### MG:a:3313

|  | Res. type | Atom | Residue | Distance | Occupancy | B-factor | Closest atom(s) to a 1^st^ shell water | EM\_dens. |
| --- | --- | --- | --- | --- | --- | --- | --- | --- |
| **Ion** | | | |  | 1.0 | 42.6 |  | 14.14 |
 1 | Water 1 | | a:9655 | 2.11 | 1.0 | 36.5 |  | 6.43 | 2 | A | OP1 | a:983 | 2.12 | 1.0 | 25.9 |  | 11.72 | 3 | Water 2 | | a:6142 | 2.22 | 1.0 | 28.6 | 2.75 [G:OP2]a:980 | 11.19 | 4 | Water 3 | | a:6511 | 2.31 | 1.0 | 40.7 |  | 5.61 | 5 | Water 4 | | a:10047 | 2.34 | 1.0 | 38.3 |  | 7.81 | 6 | A | O5' | a:983 | 3.83 | 1.0 | 29.4 |  | 13.46 | 7 | Water 5 | | a:5951 | 3.89 | 1.0 | 32.2 | 2.74 [A:OP2]a:983 | 10.62 | 8 | Water 6 | | a:6133 | 3.89 | 1.0 | 32.0 |  | 9.70 | 9 | Water 7 | | a:8886 | 4.05 | 1.0 | 46.4 | 2.98 [C:OP2]a:984 | 2.88 | 10 | A | OP2 | a:983 | 4.12 | 1.0 | 25.1 |  | 13.37 | 11 | C | OP2 | a:984 | 4.12 | 1.0 | 25.5 |  | 11.34 | 12 | Water 8 | | a:6384 | 4.29 | 1.0 | 30.7 | 2.80 [G:OP2]a:980 | 10.37 | 13 | Water 9 | | a:3459 | 4.38 | 1.0 | 39.3 |  | 3.90 | 14 | G | OP2 | a:980 | 4.44 | 1.0 | 33.2 |  | 10.84 | 15 | Water 10 | | a:5496 | 4.44 | 1.0 | 42.4 |  | 4.89 | 16 | Water 11 | | a:9107 | 4.45 | 1.0 | 37.9 |  | 5.34 | 17 | A | O3' | a:982 | 4.57 | 1.0 | 28.5 |  | 12.91 | 18 | Water 12 | | a:8325 | 4.75 | 1.0 | 37.7 | 2.91 [A:O3']a:982 | 6.13 |

  
**1^st^ shell coordination angles and distances (*trans* in "blue"; *cys* in "green"; deviations in "orange"):**   
• 1-2(W/OP1): *97º*; 1-3(W/W): *160º*; 2-3(OP1/W): 90º;   
• 1-2(W/OP1): 3.2Å; 1-3(W/W): *4.3Å*; 2-3(OP1/W): 3.1Å;   
  
**2.49 ± 0.61: the average coodination distance of the 1^st^ six ligands is Na^+^** compatible  
**8: Number of potential Water contacts**   
**1 : Number of second shell contacts involving water or ions**   
**12%: Percentage of satisfied second shell Water contacts**   
• Contact(s): /O_w_/O_w_/OP1/  
• Contact code: O_ph_2O_w_  
• Contact(s) in the exclusion zone: /O_w_/O_w_/  
→ Missing octahedral coordination for MG ←  
Go to ToC

### MG:a:3314

|  | Res. type | Atom | Residue | Distance | Occupancy | B-factor | Closest atom(s) to a 1^st^ shell water | EM\_dens. |
| --- | --- | --- | --- | --- | --- | --- | --- | --- |
| **Ion** | | | |  | 1.0 | 25.8 |  | 21.68 |
 1 | A | OP2 | a:992 | 1.93 | 1.0 | 31.0 |  | 10.88 | 2 | C | OP2 | a:993 | 2.08 | 1.0 | 26.4 |  | 14.54 | 3 | Water 1 | | a:7834 | 2.10 | 1.0 | 23.3 | 2.87 [G:OP2]a:1187 | 15.59 | 4 | G | OP2 | a:1188 | 2.15 | 1.0 | 25.6 |  | 15.74 | 5 | Water 2 | | a:8239 | 2.20 | 1.0 | 28.3 |  | 13.80 | 6 | Water 3 | | a:7877 | 2.29 | 1.0 | 27.2 |  | 13.66 | 7 | A | O5' | a:992 | 3.85 | 1.0 | 28.6 |  | 11.49 | 8 | Water 4 | | a:9535 | 3.87 | 1.0 | 43.5 |  | 4.58 | 9 | C | OP1 | a:993 | 3.88 | 1.0 | 28.5 |  | 13.42 | 10 | G | O3' | a:991 | 4.00 | 1.0 | 26.8 |  | 10.88 | 11 | G | O5' | a:1188 | 4.15 | 1.0 | 28.6 |  | 13.07 | 12 | G | O2' | a:991 | 4.19 | 1.0 | 31.4 |  | 11.69 | 13 | C | O5' | a:993 | 4.29 | 1.0 | 27.1 |  | 13.27 | 14 | G | OP1 | a:1188 | 4.30 | 1.0 | 29.4 |  | 13.60 | 15 | Water 5 | | a:10050 | 4.30 | 1.0 | 40.6 |  | 5.84 | 16 | A | OP1 | a:992 | 4.43 | 1.0 | 32.6 |  | 10.40 | 17 | G | OP2 | a:1187 | 4.43 | 1.0 | 32.0 |  | 9.20 | 18 | A | O3' | a:992 | 4.47 | 1.0 | 29.6 |  | 13.60 | 19 | G | OP1 | a:1187 | 4.55 | 1.0 | 33.7 |  | 9.52 | 20 | G | O3' | a:1187 | 4.61 | 1.0 | 29.0 |  | 11.55 | 21 | G | O5' | a:1187 | 4.65 | 1.0 | 31.0 |  | 11.76 |

  
**1^st^ shell coordination angles and distances (*trans* in "blue"; *cys* in "green"; deviations in "orange"):**   
• 1-2(OP2/OP2): *96º*; 1-3(OP2/W): *98º*; 1-4(OP2/OP2): *170º*; 1-5(OP2/W): 89º; 1-6(OP2/W): 85º; 2-3(OP2/W): 94º; 2-4(OP2/OP2): 86º; 2-5(OP2/W): 171º; 2-6(OP2/W): 89º; 3-4(W/OP2): 92º; 3-5(W/W): 93º; 3-6(W/W): 176º; 4-5(OP2/W): 88º; 4-6(OP2/W): *85º*; 5-6(W/W): *84º*;   
• 1-2(OP2/OP2): 3.0Å; 1-3(OP2/W): 3.0Å; 1-4(OP2/OP2): 4.1Å; 1-5(OP2/W): 2.9Å; 1-6(OP2/W): 2.9Å; 2-3(OP2/W): 3.0Å; 2-4(OP2/OP2): 2.9Å; 2-5(OP2/W): *4.3Å*; 2-6(OP2/W): 3.1Å; 3-4(W/OP2): 3.1Å; 3-5(W/W): 3.1Å; 3-6(W/W): *4.4Å*; 4-5(OP2/W): 3.0Å; 4-6(OP2/W): 3.0Å; 5-6(W/W): 3.0Å;   
  
**2.12 ± 0.11: the average coodination distance of the 1^st^ six ligands is Mg^2+^ compatible**  
**6: Number of potential Water contacts**   
**1 : Number of second shell contacts involving water or ions**   
**17%: Percentage of satisfied second shell Water contacts**   
• Contact(s): /O_w_/O_w_/O_w_/OP2/OP2/OP2/  
• Contact code: *mer-*3O_ph_3O_w_  
Ooctahedral coordination (4.8º)   
Go to ToC

### MG:a:3315

|  | Res. type | Atom | Residue | Distance | Occupancy | B-factor | Closest atom(s) to a 1^st^ shell water | EM\_dens. |
| --- | --- | --- | --- | --- | --- | --- | --- | --- |
| **Ion** | | | |  | 1.0 | 33.4 |  | 17.26 |
 1 | A | OP2 | a:1329 | 1.93 | 1.0 | 31.3 |  | 11.13 | 2 | Water 1 | | a:8451 | 1.97 | 1.0 | 28.8 |  | 11.70 | 3 | Water 2 | | a:4268 | 2.18 | 1.0 | 32.2 | 2.61 [A:OP2]a:1330 | 10.76 | 4 | Water 3 | | a:6772 | 2.20 | 1.0 | 28.7 | 2.79 [U:OP1]a:1328 | 11.67 | 5 | Water 4 | | a:4652 | 2.33 | 1.0 | 24.3 | 2.64 [G:N7]a:1326 | 13.59 | 6 | Water 5 | | a:6442 | 2.33 | 1.0 | 30.4 |  | 10.24 | 7 | A | OP1 | a:1329 | 3.78 | 1.0 | 29.4 |  | 11.96 | 8 | A | O5' | a:1329 | 3.92 | 1.0 | 25.1 |  | 11.88 | 9 | Water 6 | | a:4448 | 3.98 | 1.0 | 34.2 | 2.63 [A:OP1]a:1329 | 9.72 | 10 | Water 7 | | a:5842 | 4.18 | 1.0 | 33.1 |  | 11.04 | 11 | Water 8 | | a:5183 | 4.21 | 1.0 | 36.8 |  | 6.16 | 12 | U | OP1 | a:1328 | 4.32 | 1.0 | 25.0 |  | 12.34 | 13 | Water 9 | | a:7302 | 4.32 | 1.0 | 35.7 |  | 8.06 | 14 | G | N7 | a:1326 | 4.42 | 1.0 | 20.8 |  | 15.10 | 15 | A | OP2 | a:1330 | 4.43 | 1.0 | 36.2 |  | 6.27 | 16 | A | N7 | a:1330 | 4.43 | 1.0 | 25.8 |  | 12.65 | 17 | U | O3' | a:1328 | 4.43 | 1.0 | 25.8 |  | 14.00 | 18 | Water 10 | | a:7617 | 4.53 | 1.0 | 29.4 |  | 11.17 | 19 | Water 11 | | a:8703 | 4.57 | 1.0 | 29.1 |  | 11.18 | 20 | U | OP2 | a:1328 | 4.69 | 1.0 | 30.2 |  | 9.43 | 21 | Water 12 | | a:7263 | 4.76 | 1.0 | 37.6 | 2.83 [A:OP1]a:1329 // 2.97 [A:OP2]a:1330 | 6.20 |

  
**1^st^ shell coordination angles and distances (*trans* in "blue"; *cys* in "green"; deviations in "orange"):**   
• 1-2(OP2/W): *97º*; 1-3(OP2/W): 94º; 1-4(OP2/W): *99º*; 2-3(W/W): 94º; 2-4(W/W): 93º; 3-4(W/W): *165º*;   
• 1-2(OP2/W): 2.9Å; 1-3(OP2/W): 3.0Å; 1-4(OP2/W): 3.1Å; 2-3(W/W): 3.0Å; 2-4(W/W): 3.0Å; 3-4(W/W): *4.3Å*;   
  
**2.16 ± 0.16: the average coodination distance of the 1^st^ six ligands is Mg^2+^ compatible**  
**10: Number of potential Water contacts**   
**3 : Number of second shell contacts involving water or ions**   
**30%: Percentage of satisfied second shell Water contacts**   
• Contact(s): /O_w_/O_w_/O_w_/OP2/  
• Contact code: O_ph_3O_w_  
• Contact(s) in the exclusion zone: /O_w_/O_w_/  
→ Missing octahedral coordination for MG ←  
→ Slightly distorded coord. octahedron (6.9º) ←  
Go to ToC

### MG:a:3316

|  | Res. type | Atom | Residue | Distance | Occupancy | B-factor | Closest atom(s) to a 1^st^ shell water | EM\_dens. |
| --- | --- | --- | --- | --- | --- | --- | --- | --- |
| **Ion** | | | |  | 1.0 | 38.2 |  | 11.86 |
 1 | Water 1 | | a:4376 | 1.94 | 1.0 | 39.3 |  | 5.29 | 2 | G | OP2 | a:1362 | 2.08 | 1.0 | 37.8 |  | 8.83 | 3 | A | OP1 | a:1361 | 2.17 | 1.0 | 34.6 |  | 8.45 | 4 | Water 2 | | a:4861 | 2.27 | 1.0 | 38.3 | 2.66 [G:N7]a:1363 | 6.36 | 5 | Water 3 | | a:7652 | 2.35 | 1.0 | 42.1 | 2.85 [G:O6]a:1363 | 5.70 | 6 | A | OP2 | a:1361 | 3.93 | 1.0 | 36.3 |  | 9.15 | 7 | A | O5' | a:1361 | 3.94 | 1.0 | 34.3 |  | 9.51 | 8 | Water 4 | | a:6785 | 4.06 | 1.0 | 39.3 |  | 4.00 | 9 | G | OP1 | a:1362 | 4.11 | 1.0 | 42.5 |  | 5.97 | 10 | A | O3' | a:1361 | 4.33 | 1.0 | 39.8 |  | 8.43 | 11 | Water 5 | | a:6382 | 4.34 | 1.0 | 36.9 |  | 5.96 | 12 | G | N7 | a:1363 | 4.37 | 1.0 | 27.6 |  | 9.95 | 13 | G | O5' | a:1362 | 4.53 | 1.0 | 37.3 |  | 5.73 | 14 | G | O3' | a:1360 | 4.61 | 1.0 | 33.5 |  | 7.69 | 15 | G | O6 | a:1363 | 4.63 | 1.0 | 29.1 |  | 11.25 |

  
**1^st^ shell coordination angles and distances (*trans* in "blue"; *cys* in "green"; deviations in "orange"):**   
• 1-2(W/OP2): *81º*; 1-3(W/OP1): *168º*; 1-4(W/W): *85º*; 2-3(OP2/OP1): *97º*; 2-4(OP2/W): *82º*; 3-4(OP1/W): *83º*;   
• 1-2(W/OP2): 2.6Å; 1-3(W/OP1): 4.1Å; 1-4(W/W): 2.9Å; 2-3(OP2/OP1): 3.2Å; 2-4(OP2/W): 2.9Å; 3-4(OP1/W): 2.9Å;   
  
**2.46 ± 0.67: the average coodination distance of the 1^st^ six ligands is Na^+^** compatible  
**6: Number of potential Water contacts**   
**2 : Number of second shell contacts involving water or ions**   
**33%: Percentage of satisfied second shell Water contacts**   
• Contact(s): /O_w_/O_w_/OP1/OP2/  
• Contact code: *cis-*2O_ph_2O_w_  
• Contact(s) in the exclusion zone: /O_w_/  
→ Missing octahedral coordination for MG ←  
→ Slightly distorded coord. octahedron (8.1º) ←  
Go to ToC

### MG:a:3317

|  | Res. type | Atom | Residue | Distance | Occupancy | B-factor | Closest atom(s) to a 1^st^ shell water | EM\_dens. |
| --- | --- | --- | --- | --- | --- | --- | --- | --- |
| **Ion** | | | |  | 1.0 | 37.7 |  | 11.96 |
 1 | A | OP2 | a:1441 | 1.89 | 1.0 | 39.1 |  | 6.88 | 2 | Water 1 | | a:4915 | 1.94 | 1.0 | 39.9 |  | 6.43 | 3 | Water 2 | | a:6736 | 1.98 | 1.0 | 43.1 |  | 6.38 | 4 | Water 3 | | a:7556 | 2.11 | 1.0 | 41.4 |  | 6.33 | 5 | Water 4 | | a:4542 | 2.12 | 1.0 | 36.7 | 2.64 [A:N1]a:1554 | 7.27 | 6 | U | O4 | a:1442 | 2.28 | 1.0 | 37.2 |  | 9.33 | 7 | A | OP1 | a:1441 | 4.04 | 1.0 | 37.6 |  | 5.32 | 8 | U | O3' | a:1440 | 4.05 | 1.0 | 43.2 |  | 6.08 | 9 | Water 5 | | a:4686 | 4.16 | 1.0 | 42.3 |  | 5.24 | 10 | Water 6 | | a:10292 | 4.27 | 1.0 | 49.2 |  | 2.60 | 11 | A | N1 | a:1554 | 4.30 | 1.0 | 31.0 |  | 9.29 | 12 | A | O5' | a:1441 | 4.31 | 1.0 | 42.0 |  | 6.55 | 13 | U | O5' | a:1440 | 4.44 | 1.0 | 39.2 |  | 6.58 | 14 | Water 7 | | a:9341 | 4.67 | 1.0 | 46.3 |  | 2.92 |

  
**1^st^ shell coordination angles and distances (*trans* in "blue"; *cys* in "green"; deviations in "orange"):**   
• 1-2(OP2/W): 89º; 1-3(OP2/W): 178º; 1-4(OP2/W): 91º; 1-5(OP2/W): *96º*; 1-6(OP2/O4): *97º*; 2-3(W/W): 90º; 2-4(W/W): 94º; 2-5(W/W): 173º; 2-6(W/O4): *84º*; 3-4(W/W): 91º; 3-5(W/W): 86º; 3-6(W/O4): *82º*; 4-5(W/W): 92º; 4-6(W/O4): 172º; 5-6(W/O4): 90º;   
• 1-2(OP2/W): 2.7Å; 1-3(OP2/W): 3.9Å; 1-4(OP2/W): 2.8Å; 1-5(OP2/W): 3.0Å; 1-6(OP2/O4): 3.1Å; 2-3(W/W): 2.8Å; 2-4(W/W): 2.9Å; 2-5(W/W): 4.0Å; 2-6(W/O4): 2.8Å; 3-4(W/W): 2.9Å; 3-5(W/W): 2.8Å; 3-6(W/O4): 2.8Å; 4-5(W/W): 3.0Å; 4-6(W/O4): *4.4Å*; 5-6(W/O4): 3.1Å;   
  
**2.05 ± 0.13: the average coodination distance of the 1^st^ six ligands is Mg^2+^ compatible**  
**8: Number of potential Water contacts**   
**1 : Number of second shell contacts involving water or ions**   
**12%: Percentage of satisfied second shell Water contacts**   
• Contact(s): /O4/O_w_/O_w_/O_w_/O_w_/  
• Contact code: O_b_4O_w_  
• Contact(s) in the exclusion zone: /OP2/  
→ Missing octahedral coordination for MG ←  
Ooctahedral coordination (3.8º)   
Go to ToC

### MG:a:3318

|  | Res. type | Atom | Residue | Distance | Occupancy | B-factor | Closest atom(s) to a 1^st^ shell water | EM\_dens. |
| --- | --- | --- | --- | --- | --- | --- | --- | --- |
| **Ion** | | | |  | 1.0 | 48.2 |  | 6.62 |
 1 | Water 1 | | a:9839 | 1.94 | 1.0 | 44.2 |  | 4.10 | 2 | Water 2 | | a:6615 | 2.01 | 1.0 | 37.7 | 2.78 [C:OP1]a:1619 | 5.71 | 3 | Water 3 | | a:4390 | 2.21 | 1.0 | 43.2 | 2.62 [G:O6]a:1615 | 5.64 | 4 | Water 4 | | a:6057 | 2.32 | 1.0 | 30.0 | 2.75 [G:O6]a:1621 | 8.97 | 5 | Water 5 | | a:4022 | 2.40 | 1.0 | 40.8 | 2.58 [G:OP2]a:1621 | 5.07 | 6 | G | N7 | a:1621 | 4.04 | 1.0 | 25.6 |  | 13.49 | 7 | G | O6 | a:1615 | 4.06 | 1.0 | 27.7 |  | 13.19 | 8 | G | O6 | a:1621 | 4.18 | 1.0 | 24.4 |  | 13.46 | 9 | C | O2' | a:1619 | 4.32 | 1.0 | 29.0 |  | 14.18 | 10 | Water 6 | | a:8853 | 4.35 | 1.0 | 42.4 |  | 5.93 | 11 | Water 7 | | a:9201 | 4.39 | 1.0 | 39.8 |  | 6.56 | 12 | C | OP1 | a:1619 | 4.45 | 1.0 | 34.0 |  | 8.58 | 13 | G | OP2 | a:1621 | 4.62 | 1.0 | 34.9 |  | 7.06 | 14 | A | O2' | a:1610 | 4.75 | 1.0 | 37.8 |  | 7.33 |

  
**1^st^ shell coordination angles and distances (*trans* in "blue"; *cys* in "green"; deviations in "orange"):**   
• 1-2(W/W): 86º; 1-3(W/W): *97º*; 2-3(W/W): *83º*;   
• 1-2(W/W): 2.7Å; 1-3(W/W): 3.1Å; 2-3(W/W): 2.8Å;   
  
**2.49 ± 0.71: the average coodination distance of the 1^st^ six ligands is Na^+^** compatible  
**10: Number of potential Water contacts**   
**4 : Number of second shell contacts involving water or ions**   
**40%: Percentage of satisfied second shell Water contacts**   
• Contact(s): /O_w_/O_w_/O_w_/  
• Contact code: 3O_w_  
• Contact(s) in the exclusion zone: /O_w_/O_w_/  
→ Missing octahedral coordination for MG ←  
Go to ToC

### MG:a:3319

|  | Res. type | Atom | Residue | Distance | Occupancy | B-factor | Closest atom(s) to a 1^st^ shell water | EM\_dens. |
| --- | --- | --- | --- | --- | --- | --- | --- | --- |
| **Ion** | | | |  | 1.0 | 33.2 |  | 19.85 |
 1 | Water 1 | | a:4607 | 1.99 | 1.0 | 31.8 | 2.64 [G:OP1]a:1302 | 11.78 | 2 | Water 2 | | a:7351 | 2.04 | 1.0 | 33.2 | 2.83 [A:OP2]a:1639 | 10.31 | 3 | Water 3 | | a:5980 | 2.16 | 1.0 | 25.7 | 2.74 [A:OP2]a:1637 | 11.01 | 4 | Water 4 | | a:4972 | 2.17 | 1.0 | 33.4 | 2.67 [A:OP1]a:1303 // 2.82 [A:OP1]a:1637 | 10.69 | 5 | U | OP2 | a:1638 | 2.19 | 1.0 | 29.7 |  | 11.02 | 6 | Water 5 | | a:9582 | 2.27 | 1.0 | 31.4 |  | 9.67 | 7 | U | O5' | a:1638 | 3.93 | 1.0 | 26.9 |  | 12.10 | 8 | U | OP1 | a:1638 | 3.95 | 1.0 | 38.3 |  | 9.23 | 9 | A | OP1 | a:1637 | 4.03 | 1.0 | 36.4 |  | 10.71 | 10 | Water 6 | | a:6035 | 4.17 | 1.0 | 31.9 |  | 9.79 | 11 | G | OP1 | a:1302 | 4.18 | 1.0 | 39.2 |  | 8.10 | 12 | G | OP1 | a:1301 | 4.30 | 1.0 | 33.6 |  | 10.42 | 13 | Water 7 | | a:8433 | 4.35 | 1.0 | 34.4 | 2.92 [A:OP2]a:1639 | 6.94 | 14 | A | OP2 | a:1637 | 4.39 | 1.0 | 27.6 |  | 12.89 | 15 | A | OP1 | a:1303 | 4.41 | 1.0 | 33.8 |  | 10.04 | 16 | A | OP2 | a:1639 | 4.51 | 1.0 | 29.0 |  | 10.45 | 17 | A | O3' | a:1637 | 4.67 | 1.0 | 31.5 |  | 11.67 |

  
**1^st^ shell coordination angles and distances (*trans* in "blue"; *cys* in "green"; deviations in "orange"):**   
• 1-2(W/W): 91º; 1-3(W/W): 85º; 1-4(W/W): 94º; 1-5(W/OP2): 174º; 1-6(W/W): 89º; 2-3(W/W): 88º; 2-4(W/W): 174º; 2-5(W/OP2): 94º; 2-6(W/W): 94º; 3-4(W/W): 94º; 3-5(W/OP2): *95º*; 3-6(W/W): 174º; 4-5(W/OP2): *80º*; 4-6(W/W): 85º; 5-6(OP2/W): 90º;   
• 1-2(W/W): 2.9Å; 1-3(W/W): 2.8Å; 1-4(W/W): 3.1Å; 1-5(W/OP2): 4.2Å; 1-6(W/W): 3.0Å; 2-3(W/W): 2.9Å; 2-4(W/W): *4.2Å*; 2-5(W/OP2): 3.1Å; 2-6(W/W): 3.2Å; 3-4(W/W): 3.2Å; 3-5(W/OP2): *3.2Å*; 3-6(W/W): *4.4Å*; 4-5(W/OP2): 2.8Å; 4-6(W/W): 3.0Å; 5-6(OP2/W): 3.1Å;   
  
**2.14 ± 0.09: the average coodination distance of the 1^st^ six ligands is Mg^2+^ compatible**  
**10: Number of potential Water contacts**   
**5 : Number of second shell contacts involving water or ions**   
**50%: Percentage of satisfied second shell Water contacts**   
• Contact(s): /O_w_/O_w_/O_w_/O_w_/O_w_/OP2/  
• Contact code: O_ph_5O_w_  
Ooctahedral coordination (4.2º)   
Go to ToC

### MG:a:3320

|  | Res. type | Atom | Residue | Distance | Occupancy | B-factor | Closest atom(s) to a 1^st^ shell water | EM\_dens. |
| --- | --- | --- | --- | --- | --- | --- | --- | --- |
| **Ion** | | | |  | 1.0 | 29.6 |  | 20.07 |
 1 | Water 1 | | a:7110 | 2.03 | 1.0 | 30.9 | 2.94 [C:OP2]a:2027 | 10.91 | 2 | Water 2 | | a:6481 | 2.10 | 1.0 | 40.8 | 2.77 [A:N3]a:529 | 11.11 | 3 | Water 3 | | a:4999 | 2.13 | 1.0 | 27.6 | 2.67 [C:OP1]a:2027 | 12.95 | 4 | Water 4 | | a:9129 | 2.24 | 1.0 | 30.6 |  | 11.20 | 5 | U | OP1 | a:2026 | 2.26 | 1.0 | 27.2 |  | 13.41 | 6 | Water 5 | | a:9221 | 2.32 | 1.0 | 28.5 |  | 12.60 | 7 | C | O3' | a:2025 | 3.74 | 1.0 | 26.4 |  | 14.55 | 8 | A | N3 | a:529 | 4.00 | 1.0 | 22.8 |  | 14.77 | 9 | C | OP2 | a:2027 | 4.24 | 1.0 | 25.4 |  | 14.30 | 10 | Water 6 | | a:10193 | 4.26 | 1.0 | 40.8 |  | 4.38 | 11 | C | OP1 | a:2027 | 4.44 | 1.0 | 26.9 |  | 14.18 | 12 | U | O5' | a:2026 | 4.45 | 1.0 | 27.2 |  | 14.59 | 13 | Water 7 | | a:6053 | 4.50 | 1.0 | 27.5 |  | 12.70 | 14 | U | OP2 | a:2026 | 4.61 | 1.0 | 32.0 |  | 10.63 | 15 | Water 8 | | a:8161 | 4.67 | 1.0 | 33.6 | 2.89 [U:OP2]a:2026 | 10.23 |

  
**1^st^ shell coordination angles and distances (*trans* in "blue"; *cys* in "green"; deviations in "orange"):**   
• 1-2(W/W): 94º; 1-3(W/W): 90º; 1-4(W/W): 89º; 1-5(W/OP1): *82º*; 2-3(W/W): 93º; 2-4(W/W): 89º; 2-5(W/OP1): 172º; 3-4(W/W): 178º; 3-5(W/OP1): 94º; 4-5(W/OP1): *84º*;   
• 1-2(W/W): 3.0Å; 1-3(W/W): 2.9Å; 1-4(W/W): 3.0Å; 1-5(W/OP1): 2.8Å; 2-3(W/W): 3.1Å; 2-4(W/W): 3.0Å; 2-5(W/OP1): *4.3Å*; 3-4(W/W): *4.4Å*; 3-5(W/OP1): *3.2Å*; 4-5(W/OP1): 3.0Å;   
  
**2.18 ± 0.10: the average coodination distance of the 1^st^ six ligands is Mg^2+^ compatible**  
**10: Number of potential Water contacts**   
**3 : Number of second shell contacts involving water or ions**   
**30%: Percentage of satisfied second shell Water contacts**   
• Contact(s): /O_w_/O_w_/O_w_/O_w_/OP1/  
• Contact code: O_ph_4O_w_  
• Contact(s) in the exclusion zone: /O_w_/  
→ Missing octahedral coordination for MG ←  
Ooctahedral coordination (3.7º)   
Go to ToC

### MG:a:3321

|  | Res. type | Atom | Residue | Distance | Occupancy | B-factor | Closest atom(s) to a 1^st^ shell water | EM\_dens. |
| --- | --- | --- | --- | --- | --- | --- | --- | --- |
| **Ion** | | | |  | 1.0 | 28.4 |  | 17.91 |
 1 | Water 1 | | a:6367 | 1.94 | 1.0 | 26.2 | 2.86 [A:N7]a:2422 | 12.83 | 2 | Water 2 | | 2:122 | 2.01 | 1.0 | 27.6 | 2.77 [LEU:O]2:29 | 12.91 | 3 | U | O4 | a:2423 | 2.07 | 1.0 | 32.5 |  | 11.40 | 4 | Water 3 | | 2:105 | 2.21 | 1.0 | 31.5 | 2.59 [ARG:O]2:30 | 12.24 | 5 | Water 4 | | a:7404 | 2.26 | 1.0 | 31.1 |  | 9.51 | 6 | Water 5 | | 2:139 | 2.36 | 1.0 | 32.4 |  | 10.27 | 7 | Water 6 | | a:4394 | 4.01 | 1.0 | 32.2 |  | 9.79 | 8 | ARG | O | 2:30 | 4.05 | 1.0 | 28.5 |  | 13.45 | 9 | Water 7 | | 2:128 | 4.07 | 1.0 | 33.3 |  | 10.39 | 10 | Water 8 | | a:9720 | 4.24 | 1.0 | 30.3 |  | 8.78 | 11 | Water 9 | | a:6392 | 4.34 | 1.0 | 32.3 |  | 10.40 | 12 | A | N7 | a:2422 | 4.37 | 1.0 | 27.8 |  | 11.32 | 13 | LEU | O | 2:29 | 4.37 | 1.0 | 34.3 |  | 11.47 | 14 | Water 10 | | a:4742 | 4.45 | 1.0 | 36.5 |  | 8.20 | 15 | Water 11 | | a:7306 | 4.46 | 1.0 | 34.0 |  | 9.29 | 16 | Water 12 | | a:7187 | 4.61 | 1.0 | 38.1 |  | 7.72 | 17 | Water 13 | | 2:143 | 4.62 | 1.0 | 41.0 |  | 5.52 |

  
**1^st^ shell coordination angles and distances (*trans* in "blue"; *cys* in "green"; deviations in "orange"):**   
• 1-2(W/W): 94º; 1-3(W/O4): 87º; 1-4(W/W): 176º; 1-5(W/W): *85º*; 2-3(W/O4): *99º*; 2-4(W/W): 90º; 2-5(W/W): 179º; 3-4(O4/W): 93º; 3-5(O4/W): *82º*; 4-5(W/W): 91º;   
• 1-2(W/W): 2.9Å; 1-3(W/O4): 2.8Å; 1-4(W/W): 4.2Å; 1-5(W/W): 2.8Å; 2-3(W/O4): 3.1Å; 2-4(W/W): 3.0Å; 2-5(W/W): *4.3Å*; 3-4(O4/W): 3.1Å; 3-5(O4/W): 2.8Å; 4-5(W/W): *3.2Å*;   
  
**2.14 ± 0.15: the average coodination distance of the 1^st^ six ligands is Mg^2+^ compatible**  
**10: Number of potential Water contacts**   
**3 : Number of second shell contacts involving water or ions**   
**30%: Percentage of satisfied second shell Water contacts**   
• Contact(s): /O4/O_w_/O_w_/O_w_/O_w_/  
• Contact code: O_b_4O_w_  
• Contact(s) in the exclusion zone: /O_w_/  
→ Missing octahedral coordination for MG ←  
Ooctahedral coordination (4.0º)   
Go to ToC

### MG:a:3322

|  | Res. type | Atom | Residue | Distance | Occupancy | B-factor | Closest atom(s) to a 1^st^ shell water | EM\_dens. |
| --- | --- | --- | --- | --- | --- | --- | --- | --- |
| **Ion** | | | |  | 1.0 | 25.5 |  | 25.28 |
 1 | Water 1 | | a:8295 | 2.00 | 1.0 | 24.3 |  | 14.23 | 2 | Water 2 | | a:6644 | 2.01 | 1.0 | 24.1 | 2.78 [2MA:OP2]a:2507 | 15.50 | 3 | U | OP1 | a:578 | 2.08 | 1.0 | 24.7 |  | 11.99 | 4 | 2MA | OP1 | a:2507 | 2.08 | 1.0 | 26.5 |  | 15.71 | 5 | Water 3 | | a:5109 | 2.11 | 1.0 | 24.9 | 2.68 [PSU:OP2]a:2508 // 2.86 [K:K]a:3265 | 17.18 | 6 | Water 4 | | a:4593 | 2.24 | 1.0 | 24.0 | 2.64 [G:OP2]a:579 | 14.87 | 7 | **K** | K | a:3265 | 3.64 | 1.0 | 30.0 |  | 31.55 | 8 | 2MA | OP2 | a:2507 | 3.66 | 1.0 | 25.6 |  | 14.49 | 9 | U | OP2 | a:578 | 4.04 | 1.0 | 23.3 |  | 15.22 | 10 | Water 5 | | a:4361 | 4.12 | 1.0 | 33.5 |  | 11.02 | 11 | Water 6 | | a:6668 | 4.14 | 1.0 | 40.7 |  | 6.21 | 12 | Water 7 | | a:6577 | 4.16 | 1.0 | 26.5 | 2.78 [PSU:OP2]a:2508 | 13.82 | 13 | PSU | OP2 | a:2508 | 4.23 | 1.0 | 34.5 |  | 9.81 | 14 | Water 8 | | a:10057 | 4.29 | 1.0 | 36.0 |  | 9.45 | 15 | A | O3' | a:577 | 4.33 | 1.0 | 27.1 |  | 14.85 | 16 | G | O3' | a:2506 | 4.34 | 1.0 | 25.8 |  | 14.09 | 17 | 2MA | O5' | a:2507 | 4.35 | 1.0 | 26.0 |  | 13.16 | 18 | Water 9 | | a:4095 | 4.41 | 1.0 | 34.0 |  | 11.54 | 19 | U | O5' | a:578 | 4.42 | 1.0 | 22.6 |  | 16.34 | 20 | Water 10 | | a:8860 | 4.56 | 1.0 | 28.3 |  | 13.24 | 21 | Water 11 | | a:7206 | 4.57 | 1.0 | 35.7 | 2.82 [U:OP2]a:578 | 9.41 | 22 | G | OP2 | a:579 | 4.60 | 1.0 | 23.6 |  | 16.04 |

  
**1^st^ shell coordination angles and distances (*trans* in "blue"; *cys* in "green"; deviations in "orange"):**   
• 1-2(W/W): 93º; 1-3(W/OP1): 91º; 1-4(W/OP1): 174º; 1-5(W/W): 89º; 1-6(W/W): 89º; 2-3(W/OP1): 175º; 2-4(W/OP1): 91º; 2-5(W/W): *98º*; 2-6(W/W): 92º; 3-4(OP1/OP1): *85º*; 3-5(OP1/W): 86º; 3-6(OP1/W): *84º*; 4-5(OP1/W): 94º; 4-6(OP1/W): 87º; 5-6(W/W): *170º*;   
• 1-2(W/W): 2.9Å; 1-3(W/OP1): 2.9Å; 1-4(W/OP1): 4.1Å; 1-5(W/W): 2.9Å; 1-6(W/W): 3.0Å; 2-3(W/OP1): 4.1Å; 2-4(W/OP1): 2.9Å; 2-5(W/W): 3.1Å; 2-6(W/W): 3.1Å; 3-4(OP1/OP1): 2.8Å; 3-5(OP1/W): 2.8Å; 3-6(OP1/W): 2.9Å; 4-5(OP1/W): 3.1Å; 4-6(OP1/W): 3.0Å; 5-6(W/W): *4.3Å*;   
  
**2.08 ± 0.08: the average coodination distance of the 1^st^ six ligands is Mg^2+^ compatible**  
**8: Number of potential Water contacts**   
**4 : Number of second shell contacts involving water or ions**   
**50%: Percentage of satisfied second shell Water contacts**   
• Contact(s): /O_w_/O_w_/O_w_/O_w_/OP1/OP1/  
• Contact code: *cis-*2O_ph_4O_w_  
Ooctahedral coordination (4.1º)   
Go to ToC

### MG:a:3323

|  | Res. type | Atom | Residue | Distance | Occupancy | B-factor | Closest atom(s) to a 1^st^ shell water | EM\_dens. |
| --- | --- | --- | --- | --- | --- | --- | --- | --- |
| **Ion** | | | |  | 1.0 | 37.2 |  | 12.37 |
 1 | Water 1 | | a:7401 | 1.89 | 1.0 | 40.9 |  | 6.79 | 2 | A | OP2 | a:2709 | 2.10 | 1.0 | 39.2 |  | 6.65 | 3 | Water 2 | | a:4712 | 2.16 | 1.0 | 31.9 | 2.65 [A:N7]a:2710 // 2.92 [A:OP2]a:2710 | 9.01 | 4 | Water 3 | | a:5342 | 2.22 | 1.0 | 33.9 | 2.70 [C:OP1]a:2708 | 8.22 | 5 | Water 4 | | a:9511 | 2.22 | 1.0 | 35.7 |  | 8.02 | 6 | Water 5 | | a:4728 | 2.35 | 1.0 | 31.1 | 2.65 [A:N7]a:2709 | 9.10 | 7 | A | O5' | a:2709 | 3.78 | 1.0 | 40.5 |  | 6.95 | 8 | Water 6 | | a:7369 | 4.09 | 1.0 | 45.2 | 2.83 [A:OP2]a:2710 | 4.45 | 9 | A | OP1 | a:2709 | 4.15 | 1.0 | 36.0 |  | 7.59 | 10 | A | N7 | a:2710 | 4.20 | 1.0 | 28.3 |  | 11.42 | 11 | Water 7 | | a:9018 | 4.20 | 1.0 | 38.7 |  | 6.91 | 12 | A | N7 | a:2709 | 4.34 | 1.0 | 26.0 |  | 11.01 | 13 | Water 8 | | a:9942 | 4.37 | 1.0 | 41.8 |  | 5.28 | 14 | C | OP2 | a:2708 | 4.52 | 1.0 | 37.7 |  | 7.35 | 15 | C | OP1 | a:2708 | 4.54 | 1.0 | 40.5 |  | 6.41 | 16 | C | O3' | a:2708 | 4.56 | 1.0 | 39.2 |  | 7.56 | 17 | C | O5' | a:2708 | 4.67 | 1.0 | 39.6 |  | 6.73 | 18 | A | OP2 | a:2710 | 4.80 | 1.0 | 36.1 |  | 6.86 |

  
**1^st^ shell coordination angles and distances (*trans* in "blue"; *cys* in "green"; deviations in "orange"):**   
• 1-2(W/OP2): 91º; 1-3(W/W): *97º*; 1-4(W/W): 92º; 1-5(W/W): *96º*; 2-3(OP2/W): *96º*; 2-4(OP2/W): *82º*; 2-5(OP2/W): 171º; 3-4(W/W): 171º; 3-5(W/W): 89º; 4-5(W/W): 92º;   
• 1-2(W/OP2): 2.8Å; 1-3(W/W): 3.0Å; 1-4(W/W): 3.0Å; 1-5(W/W): 3.1Å; 2-3(OP2/W): 3.2Å; 2-4(OP2/W): 2.8Å; 2-5(OP2/W): *4.3Å*; 3-4(W/W): *4.4Å*; 3-5(W/W): 3.1Å; 4-5(W/W): 3.2Å;   
  
**2.16 ± 0.14: the average coodination distance of the 1^st^ six ligands is Mg^2+^ compatible**  
**10: Number of potential Water contacts**   
**4 : Number of second shell contacts involving water or ions**   
**40%: Percentage of satisfied second shell Water contacts**   
• Contact(s): /O_w_/O_w_/O_w_/OP2/  
• Contact code: O_ph_3O_w_  
• Contact(s) in the exclusion zone: /O_w_/O_w_/  
→ Missing octahedral coordination for MG ←  
→ Slightly distorded coord. octahedron (5.1º) ←  
Go to ToC

### MG:a:3324

|  | Res. type | Atom | Residue | Distance | Occupancy | B-factor | Closest atom(s) to a 1^st^ shell water | EM\_dens. |
| --- | --- | --- | --- | --- | --- | --- | --- | --- |
| **Ion** | | | |  | 1.0 | 26.3 |  | 26.90 |
 1 | OMC | OP2 | a:2502 | 1.95 | 1.0 | 28.7 |  | 15.14 | 2 | Water 1 | | a:5885 | 2.02 | 1.0 | 22.6 | 2.73 [H2U:OP2]a:2453 // 2.83 [A:O2']a:2501 | 18.04 | 3 | C | OP2 | a:2503 | 2.10 | 1.0 | 22.8 |  | 18.58 | 4 | Water 2 | | a:6013 | 2.16 | 1.0 | 23.5 | 2.74 [A:O3']a:2501 | 15.49 | 5 | A | OP2 | a:2452 | 2.23 | 1.0 | 21.3 |  | 15.93 | 6 | Water 3 | | a:5854 | 2.34 | 1.0 | 24.0 | 2.73 [G:O2']a:2451 | 17.07 | 7 | A | O3' | a:2501 | 3.37 | 1.0 | 30.7 |  | 14.35 | 8 | G | O3' | a:2451 | 3.82 | 1.0 | 22.5 |  | 17.27 | 9 | OMC | O3' | a:2502 | 4.11 | 1.0 | 21.7 |  | 18.01 | 10 | Water 4 | | a:4644 | 4.14 | 1.0 | 18.7 |  | 18.40 | 11 | OMC | O5' | a:2502 | 4.15 | 1.0 | 22.6 |  | 13.41 | 12 | A | O2' | a:2501 | 4.20 | 1.0 | 28.6 |  | 14.88 | 13 | Water 5 | | a:3794 | 4.23 | 1.0 | 25.8 |  | 14.16 | 14 | C | OP1 | a:2503 | 4.27 | 1.0 | 19.8 |  | 15.81 | 15 | A | OP1 | a:2452 | 4.29 | 1.0 | 19.4 |  | 18.69 | 16 | H2U | OP2 | a:2453 | 4.35 | 1.0 | 35.1 |  | 9.27 | 17 | OMC | OP1 | a:2502 | 4.36 | 1.0 | 26.6 |  | 15.90 | 18 | Water 6 | | a:5628 | 4.41 | 1.0 | 33.5 |  | 8.82 | 19 | C | O5' | a:2503 | 4.51 | 1.0 | 22.7 |  | 16.33 | 20 | G | O2' | a:2451 | 4.56 | 1.0 | 22.8 |  | 16.97 | 21 | A | O5' | a:2452 | 4.65 | 1.0 | 22.1 |  | 16.61 |

  
**1^st^ shell coordination angles and distances (*trans* in "blue"; *cys* in "green"; deviations in "orange"):**   
• 1-2(OP2/W): *98º*; 1-3(OP2/OP2): 90º; 1-4(OP2/W): 89º; 1-5(OP2/OP2): *95º*; 2-3(W/OP2): 171º; 2-4(W/W): 93º; 2-5(W/OP2): *85º*; 3-4(OP2/W): 92º; 3-5(OP2/OP2): 90º; 4-5(W/OP2): 175º;   
• 1-2(OP2/W): 3.0Å; 1-3(OP2/OP2): 2.9Å; 1-4(OP2/W): 2.9Å; 1-5(OP2/OP2): 3.1Å; 2-3(W/OP2): 4.1Å; 2-4(W/W): 3.0Å; 2-5(W/OP2): 2.9Å; 3-4(OP2/W): 3.1Å; 3-5(OP2/OP2): 3.1Å; 4-5(W/OP2): *4.4Å*;   
  
**2.13 ± 0.13: the average coodination distance of the 1^st^ six ligands is Mg^2+^ compatible**  
**6: Number of potential Water contacts**   
**4 : Number of second shell contacts involving water or ions**   
**67%: Percentage of satisfied second shell Water contacts**   
• Contact(s): /O_w_/O_w_/OP2/OP2/OP2/  
• Contact code: *fac-*3O_ph_2O_w_  
• Contact(s) in the exclusion zone: /O3'/O_w_/  
→ Missing octahedral coordination for MG ←  
Ooctahedral coordination (3.8º)   
Go to ToC

### MG:a:3325

|  | Res. type | Atom | Residue | Distance | Occupancy | B-factor | Closest atom(s) to a 1^st^ shell water | EM\_dens. |
| --- | --- | --- | --- | --- | --- | --- | --- | --- |
| **Ion** | | | |  | 1.0 | 23.2 |  | 26.73 |
 1 | A | OP1 | a:2452 | 1.92 | 1.0 | 19.4 |  | 18.69 | 2 | C | OP1 | a:2503 | 2.08 | 1.0 | 19.8 |  | 15.81 | 3 | Water 1 | | a:7008 | 2.14 | 1.0 | 20.7 | 2.95 [6MZ:N1]a:2034 // 2.96 [K:K]a:3111 | 18.60 | 4 | G | O6 | a:572 | 2.20 | 1.0 | 20.6 |  | 17.29 | 5 | Water 2 | | a:6699 | 2.23 | 1.0 | 19.6 |  | 18.66 | 6 | Water 3 | | a:4644 | 2.35 | 1.0 | 18.7 | 2.64 [U:O4]a:571 | 18.40 | 7 | A | OP2 | a:2452 | 3.66 | 1.0 | 21.3 |  | 15.93 | 8 | **K** | K | a:3111 | 3.78 | 1.0 | 30.0 |  | 31.46 | 9 | C | OP2 | a:2503 | 3.87 | 1.0 | 22.8 |  | 18.58 | 10 | Water 4 | | a:6561 | 4.09 | 1.0 | 22.0 | 2.84 [A:O5']a:2452 // 2.92 [K:K]a:3111 | 16.66 | 11 | OMC | O3' | a:2502 | 4.10 | 1.0 | 21.7 |  | 18.01 | 12 | 6MZ | N1 | a:2034 | 4.17 | 1.0 | 17.6 |  | 15.90 | 13 | U | O4 | a:571 | 4.18 | 1.0 | 25.0 |  | 16.51 | 14 | A | O5' | a:2452 | 4.22 | 1.0 | 22.1 |  | 16.61 | 15 | U | OP2 | a:2504 | 4.24 | 1.0 | 21.5 |  | 17.48 | 16 | G | O3' | a:2451 | 4.29 | 1.0 | 22.5 |  | 17.27 | 17 | C | O5' | a:2503 | 4.53 | 1.0 | 22.7 |  | 16.33 | 18 | Water 5 | | a:10007 | 4.57 | 1.0 | 33.7 |  | 10.78 | 19 | U | O2' | a:570 | 4.78 | 1.0 | 24.0 |  | 18.56 |

  
**1^st^ shell coordination angles and distances (*trans* in "blue"; *cys* in "green"; deviations in "orange"):**   
• 1-2(OP1/OP1): *104º*; 1-3(OP1/W): 87º; 1-4(OP1/O6): *164º*; 1-5(OP1/W): *84º*; 2-3(OP1/W): *84º*; 2-4(OP1/O6): 92º; 2-5(OP1/W): 172º; 3-4(W/O6): *98º*; 3-5(W/W): *98º*; 4-5(O6/W): *80º*;   
• 1-2(OP1/OP1): 3.1Å; 1-3(OP1/W): 2.8Å; 1-4(OP1/O6): 4.1Å; 1-5(OP1/W): 2.8Å; 2-3(OP1/W): 2.8Å; 2-4(OP1/O6): 3.1Å; 2-5(OP1/W): *4.3Å*; 3-4(W/O6): *3.3Å*; 3-5(W/W): *3.3Å*; 4-5(O6/W): 2.8Å;   
  
**2.15 ± 0.13: the average coodination distance of the 1^st^ six ligands is Mg^2+^ compatible**  
**6: Number of potential Water contacts**   
**3 : Number of second shell contacts involving water or ions**   
**50%: Percentage of satisfied second shell Water contacts**   
• Contact(s): /O6/O_w_/O_w_/OP1/OP1/  
• Contact code: *trans-*2O_ph_O_b_2O_w_  
• Contact(s) in the exclusion zone: /O_w_/  
→ Missing octahedral coordination for MG ←  
→ Slightly distorded coord. octahedron (8.1º) ←  
Go to ToC

### MG:a:3326

|  | Res. type | Atom | Residue | Distance | Occupancy | B-factor | Closest atom(s) to a 1^st^ shell water | EM\_dens. |
| --- | --- | --- | --- | --- | --- | --- | --- | --- |
| **Ion** | | | |  | 1.0 | 23.7 |  | 25.77 |
 1 | U | OP1 | a:1784 | 1.93 | 1.0 | 27.2 |  | 14.09 | 2 | A | OP2 | a:1785 | 1.99 | 1.0 | 23.2 |  | 13.82 | 3 | A | OP1 | a:1782 | 2.05 | 1.0 | 24.7 |  | 12.59 | 4 | Water 1 | | a:8178 | 2.18 | 1.0 | 25.3 |  | 19.14 | 5 | Water 2 | | a:5750 | 2.23 | 1.0 | 27.2 | 2.73 [A:N7]a:1785 | 16.23 | 6 | Water 3 | | a:3769 | 2.24 | 1.0 | 24.1 | 2.53 [U:OP2]a:1783 | 15.88 | 7 | Water 4 | | a:7438 | 4.04 | 1.0 | 34.4 | 2.84 [A:O5']a:1782 | 9.92 | 8 | U | OP2 | a:1784 | 4.09 | 1.0 | 32.0 |  | 10.94 | 9 | Water 5 | | a:6975 | 4.13 | 1.0 | 19.4 | 2.80 [A:OP1]a:1785 | 16.40 | 10 | A | O5' | a:1782 | 4.14 | 1.0 | 26.0 |  | 14.53 | 11 | Water 6 | | a:10011 | 4.19 | 1.0 | 32.1 |  | 9.89 | 12 | A | O5' | a:1785 | 4.24 | 1.0 | 24.0 |  | 13.74 | 13 | U | O5' | a:1784 | 4.25 | 1.0 | 31.9 |  | 12.39 | 14 | U | O3' | a:1784 | 4.27 | 1.0 | 28.3 |  | 13.28 | 15 | Water 7 | | a:4827 | 4.28 | 1.0 | 32.9 | 2.66 [U:OP2]a:1784 | 9.95 | 16 | A | OP1 | a:1785 | 4.32 | 1.0 | 25.0 |  | 14.39 | 17 | A | OP2 | a:1782 | 4.33 | 1.0 | 26.9 |  | 13.07 | 18 | U | O3' | a:1783 | 4.33 | 1.0 | 28.6 |  | 12.54 | 19 | Water 8 | | a:4899 | 4.40 | 1.0 | 34.4 | 2.81 [A:O5']a:1785 | 14.45 | 20 | U | O3' | a:1781 | 4.43 | 1.0 | 24.1 |  | 15.92 | 21 | U | OP2 | a:1783 | 4.45 | 1.0 | 34.1 |  | 9.91 | 22 | Water 9 | | a:4955 | 4.45 | 1.0 | 23.3 |  | 17.54 | 23 | A | N7 | a:1785 | 4.50 | 1.0 | 20.5 |  | 16.91 | 24 | U | O2' | a:1781 | 4.73 | 1.0 | 25.3 |  | 16.50 |

  
**1^st^ shell coordination angles and distances (*trans* in "blue"; *cys* in "green"; deviations in "orange"):**   
• 1-2(OP1/OP2): 93º; 1-3(OP1/OP1): 90º; 1-4(OP1/W): 176º; 1-5(OP1/W): 94º; 1-6(OP1/W): 89º; 2-3(OP2/OP1): 173º; 2-4(OP2/W): 90º; 2-5(OP2/W): 90º; 2-6(OP2/W): 92º; 3-4(OP1/W): 86º; 3-5(OP1/W): *84º*; 3-6(OP1/W): 93º; 4-5(W/W): 87º; 4-6(W/W): 89º; 5-6(W/W): 176º;   
• 1-2(OP1/OP2): 2.8Å; 1-3(OP1/OP1): 2.8Å; 1-4(OP1/W): 4.1Å; 1-5(OP1/W): 3.0Å; 1-6(OP1/W): 2.9Å; 2-3(OP2/OP1): 4.0Å; 2-4(OP2/W): 3.0Å; 2-5(OP2/W): 3.0Å; 2-6(OP2/W): 3.1Å; 3-4(OP1/W): 2.9Å; 3-5(OP1/W): 2.9Å; 3-6(OP1/W): 3.1Å; 4-5(W/W): 3.0Å; 4-6(W/W): 3.1Å; 5-6(W/W): *4.5Å*;   
  
**2.10 ± 0.12: the average coodination distance of the 1^st^ six ligands is Mg^2+^ compatible**  
**6: Number of potential Water contacts**   
**2 : Number of second shell contacts involving water or ions**   
**33%: Percentage of satisfied second shell Water contacts**   
• Contact(s): /O_w_/O_w_/O_w_/OP1/OP1/OP2/  
• Contact code: *mer-*3O_ph_3O_w_  
Ooctahedral coordination (2.7º)   
Go to ToC

### MG:a:3327

|  | Res. type | Atom | Residue | Distance | Occupancy | B-factor | Closest atom(s) to a 1^st^ shell water | EM\_dens. |
| --- | --- | --- | --- | --- | --- | --- | --- | --- |
| **Ion** | | | |  | 1.0 | 25.9 |  | 23.15 |
 1 | A | OP2 | a:732 | 1.94 | 1.0 | 24.1 |  | 12.80 | 2 | C | OP2 | a:733 | 1.95 | 1.0 | 24.7 |  | 14.44 | 3 | Water 1 | | a:7970 | 2.07 | 1.0 | 28.5 | 2.97 [G:OP1]a:765 | 14.70 | 4 | A | OP1 | a:763 | 2.08 | 1.0 | 22.0 |  | 12.26 | 5 | Water 2 | | a:4421 | 2.19 | 1.0 | 24.1 | 2.62 [G:OP1]a:765 | 14.85 | 6 | Water 3 | | a:6068 | 2.31 | 1.0 | 24.1 | 2.75 [U:OP1]a:764 | 13.76 | 7 | Water 4 | | a:6942 | 3.74 | 1.0 | 31.4 | 2.95 [A:OP1]a:732 | 9.18 | 8 | A | O3' | a:732 | 3.78 | 1.0 | 24.7 |  | 12.21 | 9 | C | OP1 | a:733 | 3.99 | 1.0 | 27.8 |  | 12.09 | 10 | A | O5' | a:763 | 4.00 | 1.0 | 22.2 |  | 14.62 | 11 | G | OP1 | a:765 | 4.02 | 1.0 | 32.2 |  | 10.35 | 12 | G | O3' | a:731 | 4.03 | 1.0 | 25.2 |  | 13.75 | 13 | A | OP2 | a:763 | 4.09 | 1.0 | 23.5 |  | 14.39 | 14 | Water 5 | | a:8805 | 4.11 | 1.0 | 30.7 |  | 12.83 | 15 | Water 6 | | a:5954 | 4.15 | 1.0 | 27.5 |  | 13.49 | 16 | A | O5' | a:732 | 4.22 | 1.0 | 26.9 |  | 10.63 | 17 | A | OP1 | a:732 | 4.31 | 1.0 | 30.0 |  | 14.13 | 18 | C | O5' | a:733 | 4.37 | 1.0 | 27.1 |  | 12.72 | 19 | U | OP1 | a:764 | 4.42 | 1.0 | 30.5 |  | 12.81 | 20 | Water 7 | | a:8268 | 4.52 | 1.0 | 29.1 | 2.91 [U:OP1]a:764 | 11.51 | 21 | G | O3' | a:762 | 4.56 | 1.0 | 27.4 |  | 14.63 |

  
**1^st^ shell coordination angles and distances (*trans* in "blue"; *cys* in "green"; deviations in "orange"):**   
• 1-2(OP2/OP2): 94º; 1-3(OP2/W): 92º; 1-4(OP2/OP1): *101º*; 1-5(OP2/W): *100º*; 2-3(OP2/W): 170º; 2-4(OP2/OP1): *98º*; 2-5(OP2/W): 95º; 3-4(W/OP1): 89º; 3-5(W/W): *76º*; 4-5(OP1/W): *154º*;   
• 1-2(OP2/OP2): 2.8Å; 1-3(OP2/W): 2.9Å; 1-4(OP2/OP1): 3.1Å; 1-5(OP2/W): 3.2Å; 2-3(OP2/W): 4.0Å; 2-4(OP2/OP1): 3.0Å; 2-5(OP2/W): 3.0Å; 3-4(W/OP1): 2.9Å; 3-5(W/W): 2.6Å; 4-5(OP1/W): 4.2Å;   
  
**2.09 ± 0.13: the average coodination distance of the 1^st^ six ligands is Mg^2+^ compatible**  
**6: Number of potential Water contacts**   
**3 : Number of second shell contacts involving water or ions**   
**50%: Percentage of satisfied second shell Water contacts**   
• Contact(s): /O_w_/O_w_/OP1/OP2/OP2/  
• Contact code: *fac-*3O_ph_2O_w_  
• Contact(s) in the exclusion zone: /O_w_/  
→ Missing octahedral coordination for MG ←  
→ Slightly distorded coord. octahedron (8.9º) ←  
Go to ToC

### MG:a:3328

|  | Res. type | Atom | Residue | Distance | Occupancy | B-factor | Closest atom(s) to a 1^st^ shell water | EM\_dens. |
| --- | --- | --- | --- | --- | --- | --- | --- | --- |
| **Ion** | | | |  | 1.0 | 27.0 |  | 23.82 |
 1 | Water 1 | | a:6975 | 1.96 | 1.0 | 19.4 |  | 16.40 | 2 | A | OP1 | a:1785 | 2.03 | 1.0 | 25.0 |  | 14.39 | 3 | C | OP2 | a:742 | 2.08 | 1.0 | 28.4 |  | 13.41 | 4 | Water 2 | | a:7944 | 2.19 | 1.0 | 24.3 |  | 14.60 | 5 | A | OP2 | a:1786 | 2.20 | 1.0 | 24.1 |  | 14.17 | 6 | Water 3 | | a:5246 | 2.44 | 1.0 | 25.8 | 2.69 [U:OP2]a:743 // 2.85 [A:OP1]a:1786 | 13.26 | 7 | Water 4 | | a:5425 | 3.75 | 1.0 | 27.1 | 2.70 [C:OP1]a:742 | 12.85 | 8 | A | OP1 | a:1786 | 3.78 | 1.0 | 26.2 |  | 13.65 | 9 | Water 5 | | a:4955 | 3.94 | 1.0 | 23.3 |  | 17.54 | 10 | A | OP2 | a:1785 | 4.03 | 1.0 | 23.2 |  | 13.82 | 11 | Water 6 | | a:5444 | 4.07 | 1.0 | 36.2 |  | 8.06 | 12 | Water 7 | | a:7536 | 4.12 | 1.0 | 35.6 | 2.84 [U:O3']a:1784 | 5.24 | 13 | C | OP1 | a:742 | 4.21 | 1.0 | 25.7 |  | 12.75 | 14 | A | O3' | a:1785 | 4.24 | 1.0 | 24.6 |  | 13.09 | 15 | A | O5' | a:1785 | 4.29 | 1.0 | 24.0 |  | 13.74 | 16 | U | O3' | a:1784 | 4.38 | 1.0 | 28.3 |  | 13.28 | 17 | C | O5' | a:742 | 4.39 | 1.0 | 25.6 |  | 14.61 | 18 | A | O3' | a:741 | 4.44 | 1.0 | 24.6 |  | 14.94 | 19 | Water 8 | | a:10011 | 4.51 | 1.0 | 32.1 |  | 9.89 | 20 | A | O5' | a:1786 | 4.58 | 1.0 | 30.5 |  | 13.35 | 21 | U | OP2 | a:743 | 4.71 | 1.0 | 32.3 |  | 13.41 | 22 | Water 9 | | a:9045 | 4.74 | 1.0 | 37.7 |  | 7.04 |

  
**1^st^ shell coordination angles and distances (*trans* in "blue"; *cys* in "green"; deviations in "orange"):**   
• 1-2(W/OP1): 89º; 1-3(W/OP2): 94º; 1-4(W/W): 90º; 1-5(W/OP2): *96º*; 2-3(OP1/OP2): 171º; 2-4(OP1/W): 86º; 2-5(OP1/OP2): 90º; 3-4(OP2/W): 86º; 3-5(OP2/OP2): *97º*; 4-5(W/OP2): 172º;   
• 1-2(W/OP1): 2.8Å; 1-3(W/OP2): 3.0Å; 1-4(W/W): 2.9Å; 1-5(W/OP2): 3.1Å; 2-3(OP1/OP2): 4.1Å; 2-4(OP1/W): 2.9Å; 2-5(OP1/OP2): 3.0Å; 3-4(OP2/W): 2.9Å; 3-5(OP2/OP2): *3.2Å*; 4-5(W/OP2): *4.4Å*;   
  
**2.15 ± 0.16: the average coodination distance of the 1^st^ six ligands is Mg^2+^ compatible**  
**6: Number of potential Water contacts**   
**2 : Number of second shell contacts involving water or ions**   
**33%: Percentage of satisfied second shell Water contacts**   
• Contact(s): /O_w_/O_w_/OP1/OP2/OP2/  
• Contact code: *mer-*3O_ph_2O_w_  
• Contact(s) in the exclusion zone: /O_w_/  
→ Missing octahedral coordination for MG ←  
Ooctahedral coordination (4.4º)   
Go to ToC

### MG:a:3329

|  | Res. type | Atom | Residue | Distance | Occupancy | B-factor | Closest atom(s) to a 1^st^ shell water | EM\_dens. |
| --- | --- | --- | --- | --- | --- | --- | --- | --- |
| **Ion** | | | |  | 1.0 | 25.3 |  | 23.10 |
 1 | G | OP1 | a:2433 | 2.02 | 1.0 | 25.2 |  | 12.03 | 2 | U | OP1 | a:828 | 2.03 | 1.0 | 22.6 |  | 14.10 | 3 | G | OP1 | a:2432 | 2.10 | 1.0 | 26.9 |  | 12.92 | 4 | C | OP1 | a:2431 | 2.12 | 1.0 | 29.3 |  | 12.16 | 5 | Water 1 | | a:7954 | 2.17 | 1.0 | 24.1 |  | 13.21 | 6 | Water 2 | | a:3814 | 2.22 | 1.0 | 24.3 | 2.54 [U:OP2]a:829 // 2.93 [K:K]a:3119 | 18.96 | 7 | **K** | K | a:3119 | 3.70 | 1.0 | 30.0 |  | 25.79 | 8 | U | OP2 | a:828 | 3.79 | 1.0 | 24.5 |  | 12.83 | 9 | G | O5' | a:2432 | 3.82 | 1.0 | 26.8 |  | 13.93 | 10 | U | O5' | a:828 | 3.85 | 1.0 | 26.3 |  | 14.41 | 11 | C | O5' | a:2431 | 4.01 | 1.0 | 22.2 |  | 15.84 | 12 | G | O3' | a:2432 | 4.02 | 1.0 | 24.7 |  | 14.24 | 13 | G | OP2 | a:2432 | 4.18 | 1.0 | 25.2 |  | 14.11 | 14 | G | OP2 | a:2433 | 4.28 | 1.0 | 24.6 |  | 11.90 | 15 | C | OP2 | a:2431 | 4.32 | 1.0 | 29.5 |  | 12.57 | 16 | Water 3 | | a:4672 | 4.39 | 1.0 | 35.5 | 2.82 [C:OP2]a:2431 | 9.17 | 17 | G | O5' | a:2433 | 4.44 | 1.0 | 23.0 |  | 13.75 | 18 | A | O3' | a:827 | 4.50 | 1.0 | 22.9 |  | 18.73 | 19 | Water 4 | | a:9485 | 4.50 | 1.0 | 29.4 |  | 9.65 | 20 | U | OP2 | a:829 | 4.52 | 1.0 | 25.2 |  | 14.42 | 21 | A | O3' | a:2430 | 4.55 | 1.0 | 32.9 |  | 12.62 | 22 | C | O3' | a:2431 | 4.59 | 1.0 | 23.1 |  | 15.36 |

  
**1^st^ shell coordination angles and distances (*trans* in "blue"; *cys* in "green"; deviations in "orange"):**   
• 1-2(OP1/OP1): *99º*; 1-3(OP1/OP1): 92º; 1-4(OP1/OP1): 90º; 1-5(OP1/W): 174º; 1-6(OP1/W): 90º; 2-3(OP1/OP1): *102º*; 2-4(OP1/OP1): 170º; 2-5(OP1/W): 87º; 2-6(OP1/W): 92º; 3-4(OP1/OP1): *82º*; 3-5(OP1/W): 88º; 3-6(OP1/W): *165º*; 4-5(OP1/W): *84º*; 4-6(OP1/W): *83º*; 5-6(W/W): 89º;   
• 1-2(OP1/OP1): 3.1Å; 1-3(OP1/OP1): 3.0Å; 1-4(OP1/OP1): 2.9Å; 1-5(OP1/W): 4.2Å; 1-6(OP1/W): 3.0Å; 2-3(OP1/OP1): *3.2Å*; 2-4(OP1/OP1): 4.1Å; 2-5(OP1/W): 2.9Å; 2-6(OP1/W): 3.1Å; 3-4(OP1/OP1): 2.8Å; 3-5(OP1/W): 3.0Å; 3-6(OP1/W): *4.3Å*; 4-5(OP1/W): 2.9Å; 4-6(OP1/W): 2.9Å; 5-6(W/W): 3.1Å;   
  
**2.11 ± 0.07: the average coodination distance of the 1^st^ six ligands is Mg^2+^ compatible**  
**4: Number of potential Water contacts**   
**2 : Number of second shell contacts involving water or ions**   
**50%: Percentage of satisfied second shell Water contacts**   
• Contact(s): /O_w_/O_w_/OP1/OP1/OP1/OP1/  
• Contact code: 4O_ph_2O_w_  
→ Slightly distorded coord. octahedron (5.5º) ←  
Go to ToC

### MG:a:3330

|  | Res. type | Atom | Residue | Distance | Occupancy | B-factor | Closest atom(s) to a 1^st^ shell water | EM\_dens. |
| --- | --- | --- | --- | --- | --- | --- | --- | --- |
| **Ion** | | | |  | 1.0 | 42.7 |  | 7.69 |
 1 | G | OP1 | a:1241 | 2.48 | 1.0 | 41.4 |  | 8.78 | 2 | C | OP1 | a:31 | 2.53 | 1.0 | 29.7 |  | 11.08 | 3 | Water 1 | | a:4540 | 2.65 | 1.0 | 46.1 | 2.64 [C:OP1]a:1202 | 5.16 | 4 | Water 2 | | a:4817 | 3.12 | 1.0 | 44.3 |  | 6.78 | 5 | Water 3 | | a:6290 | 3.13 | 1.0 | 32.4 |  | 7.72 | 6 | U | OP2 | a:1242 | 3.70 | 1.0 | 34.5 |  | 9.30 | 7 | G | O5' | a:1241 | 3.82 | 1.0 | 29.7 |  | 10.74 | 8 | C | OP1 | a:1202 | 4.01 | 1.0 | 36.5 |  | 7.30 | 9 | Water 4 | | a:8698 | 4.05 | 1.0 | 38.9 | 2.95 [G:OP2]a:1241 | 4.12 | 10 | Water 5 | | a:9960 | 4.15 | 1.0 | 37.9 |  | 6.15 | 11 | G | O3' | a:1241 | 4.52 | 1.0 | 37.9 |  | 8.91 | 12 | G | OP2 | a:1241 | 4.57 | 1.0 | 31.4 |  | 11.64 | 13 | G | O3' | a:30 | 4.71 | 1.0 | 29.8 |  | 11.37 | 14 | C | O5' | a:31 | 4.74 | 1.0 | 29.4 |  | 12.78 | 15 | U | OP1 | a:1242 | 4.74 | 1.0 | 42.3 |  | 5.99 |

  
• • **2.93 ± 0.43: the average coodination distance of the 1^st^ six ligands is K^+^ compatible**  
**6: Number of potential Water contacts**   
**1 : Number of second shell contacts involving water or ions**   
**17%: Percentage of satisfied second shell Water contacts**   
• Contact code: → No contacts ←   
• Contact(s) in the exclusion zone: /O_w_/O_w_/O_w_/OP1/OP1/  
→ Firts MG coordination shell missing ←  
Go to ToC

### MG:a:3331

|  | Res. type | Atom | Residue | Distance | Occupancy | B-factor | Closest atom(s) to a 1^st^ shell water | EM\_dens. |
| --- | --- | --- | --- | --- | --- | --- | --- | --- |
| **Ion** | | | |  | 1.0 | 28.6 |  | 21.50 |
 1 | G | OP1 | a:2578 | 1.99 | 1.0 | 26.7 |  | 13.07 | 2 | C | OP1 | a:2577 | 2.00 | 1.0 | 27.5 |  | 13.00 | 3 | Water 1 | | a:7507 | 2.08 | 1.0 | 31.0 |  | 9.91 | 4 | C | OP2 | a:2579 | 2.10 | 1.0 | 24.4 |  | 16.34 | 5 | Water 2 | | a:5263 | 2.19 | 1.0 | 33.4 | 2.69 [C:OP1]a:2511 | 12.59 | 6 | Water 3 | | a:7871 | 2.29 | 1.0 | 32.2 |  | 11.64 | 7 | C | OP2 | a:2577 | 3.84 | 1.0 | 31.5 |  | 10.61 | 8 | G | O3' | a:2578 | 3.98 | 1.0 | 26.0 |  | 16.63 | 9 | C | O5' | a:2577 | 4.01 | 1.0 | 28.1 |  | 13.30 | 10 | C | OP1 | a:2579 | 4.02 | 1.0 | 25.2 |  | 14.03 | 11 | Water 4 | | a:6519 | 4.04 | 1.0 | 30.3 |  | 10.86 | 12 | Water 5 | | a:6383 | 4.11 | 1.0 | 39.7 |  | 5.92 | 13 | G | O5' | a:2578 | 4.15 | 1.0 | 24.8 |  | 13.90 | 14 | C | O3' | a:2577 | 4.22 | 1.0 | 32.9 |  | 12.81 | 15 | Water 6 | | a:9436 | 4.26 | 1.0 | 44.5 |  | 3.03 | 16 | C | OP1 | a:2511 | 4.38 | 1.0 | 45.0 |  | 3.26 | 17 | G | OP2 | a:2578 | 4.42 | 1.0 | 31.5 |  | 11.28 | 18 | A | O3' | a:2576 | 4.52 | 1.0 | 26.5 |  | 13.86 | 19 | C | O5' | a:2579 | 4.55 | 1.0 | 25.1 |  | 14.97 | 20 | A | O2' | a:2576 | 4.69 | 1.0 | 24.4 |  | 14.39 | 21 | Water 7 | | a:7321 | 4.74 | 1.0 | 32.3 | 2.83 [G:OP2]a:2578 | 7.50 |

  
**1^st^ shell coordination angles and distances (*trans* in "blue"; *cys* in "green"; deviations in "orange"):**   
• 1-2(OP1/OP1): 90º; 1-3(OP1/W): 174º; 1-4(OP1/OP2): *99º*; 1-5(OP1/W): 89º; 1-6(OP1/W): *84º*; 2-3(OP1/W): 94º; 2-4(OP1/OP2): *96º*; 2-5(OP1/W): 93º; 2-6(OP1/W): 173º; 3-4(W/OP2): 86º; 3-5(W/W): 85º; 3-6(W/W): 92º; 4-5(OP2/W): *168º*; 4-6(OP2/W): *83º*; 5-6(W/W): 89º;   
• 1-2(OP1/OP1): 2.8Å; 1-3(OP1/W): 4.1Å; 1-4(OP1/OP2): 3.1Å; 1-5(OP1/W): 2.9Å; 1-6(OP1/W): 2.9Å; 2-3(OP1/W): 3.0Å; 2-4(OP1/OP2): 3.1Å; 2-5(OP1/W): 3.0Å; 2-6(OP1/W): *4.3Å*; 3-4(W/OP2): 2.8Å; 3-5(W/W): 2.9Å; 3-6(W/W): 3.2Å; 4-5(OP2/W): *4.3Å*; 4-6(OP2/W): 2.9Å; 5-6(W/W): 3.2Å;   
  
**2.11 ± 0.11: the average coodination distance of the 1^st^ six ligands is Mg^2+^ compatible**  
**6: Number of potential Water contacts**   
**1 : Number of second shell contacts involving water or ions**   
**17%: Percentage of satisfied second shell Water contacts**   
• Contact(s): /O_w_/O_w_/O_w_/OP1/OP1/OP2/  
• Contact code: *fac-*3O_ph_3O_w_  
Ooctahedral coordination (4.9º)   
Go to ToC

### MG:a:3332

|  | Res. type | Atom | Residue | Distance | Occupancy | B-factor | Closest atom(s) to a 1^st^ shell water | EM\_dens. |
| --- | --- | --- | --- | --- | --- | --- | --- | --- |
| **Ion** | | | |  | 1.0 | 29.6 |  | 18.47 |
 1 | A | OP2 | a:1344 | 1.89 | 1.0 | 34.0 |  | 10.97 | 2 | Water 1 | | a:7080 | 1.98 | 1.0 | 28.0 | 2.87 [G:N7]a:1603 | 11.90 | 3 | Water 2 | | a:6334 | 2.07 | 1.0 | 27.1 |  | 11.47 | 4 | U | O4 | a:1604 | 2.14 | 1.0 | 28.5 |  | 11.23 | 5 | Water 3 | | s:201 | 2.19 | 1.0 | 24.8 | 2.58 [THR:O]s:60 | 11.93 | 6 | Water 4 | | a:6023 | 2.20 | 1.0 | 27.4 | 2.87 [G:O6]a:1603 | 12.63 | 7 | THR | O | s:60 | 3.80 | 1.0 | 33.5 |  | 10.04 | 8 | A | OP1 | a:1344 | 3.89 | 1.0 | 31.6 |  | 9.34 | 9 | Water 5 | | a:8854 | 3.97 | 1.0 | 33.7 |  | 10.52 | 10 | A | O5' | a:1344 | 4.00 | 1.0 | 30.3 |  | 9.94 | 11 | Water 6 | | a:5301 | 4.18 | 1.0 | 35.1 |  | 8.55 | 12 | G | N7 | a:1603 | 4.25 | 1.0 | 23.4 |  | 13.07 | 13 | G | O6 | a:1603 | 4.26 | 1.0 | 30.8 |  | 11.31 | 14 | G | O3' | a:1343 | 4.34 | 1.0 | 27.2 |  | 12.75 | 15 | Water 7 | | a:6168 | 4.51 | 1.0 | 31.4 |  | 10.83 | 16 | Water 8 | | a:8115 | 4.77 | 1.0 | 31.8 |  | 11.10 |

  
**1^st^ shell coordination angles and distances (*trans* in "blue"; *cys* in "green"; deviations in "orange"):**   
• 1-2(OP2/W): 174º; 1-3(OP2/W): 88º; 1-4(OP2/O4): *99º*; 1-5(OP2/W): 91º; 1-6(OP2/W): *96º*; 2-3(W/W): 87º; 2-4(W/O4): 86º; 2-5(W/W): *84º*; 2-6(W/W): 89º; 3-4(W/O4): 173º; 3-5(W/W): 87º; 3-6(W/W): *100º*; 4-5(O4/W): 94º; 4-6(O4/W): *78º*; 5-6(W/W): 170º;   
• 1-2(OP2/W): 3.9Å; 1-3(OP2/W): 2.8Å; 1-4(OP2/O4): 3.1Å; 1-5(OP2/W): 2.9Å; 1-6(OP2/W): 3.0Å; 2-3(W/W): 2.8Å; 2-4(W/O4): 2.8Å; 2-5(W/W): 2.8Å; 2-6(W/W): 2.9Å; 3-4(W/O4): *4.2Å*; 3-5(W/W): 2.9Å; 3-6(W/W): *3.3Å*; 4-5(O4/W): 3.2Å; 4-6(O4/W): 2.7Å; 5-6(W/W): *4.4Å*;   
  
**2.08 ± 0.11: the average coodination distance of the 1^st^ six ligands is Mg^2+^ compatible**  
**8: Number of potential Water contacts**   
**3 : Number of second shell contacts involving water or ions**   
**38%: Percentage of satisfied second shell Water contacts**   
• Contact(s): /O4/O_w_/O_w_/O_w_/O_w_/  
• Contact code: O_b_4O_w_  
• Contact(s) in the exclusion zone: /OP2/  
→ Missing octahedral coordination for MG ←  
→ Slightly distorded coord. octahedron (5.5º) ←  
Go to ToC

### MG:a:3333

|  | Res. type | Atom | Residue | Distance | Occupancy | B-factor | Closest atom(s) to a 1^st^ shell water | EM\_dens. |
| --- | --- | --- | --- | --- | --- | --- | --- | --- |
| **Ion** | | | |  | 1.0 | 24.2 |  | 23.66 |
 1 | A | OP2 | a:1671 | 1.97 | 1.0 | 21.8 |  | 14.49 | 2 | C | OP2 | a:1672 | 2.02 | 1.0 | 20.9 |  | 14.04 | 3 | G | OP1 | a:2554 | 2.15 | 1.0 | 25.6 |  | 15.30 | 4 | U | O4 | a:1673 | 2.15 | 1.0 | 25.2 |  | 16.14 | 5 | Water 1 | | a:6861 | 2.26 | 1.0 | 22.8 | 2.80 [C:OP2]a:2555 | 19.26 | 6 | Water 2 | | a:7406 | 2.47 | 1.0 | 22.2 | 2.84 [G:OP2]a:2554 | 16.98 | 7 | Water 3 | | a:8668 | 3.76 | 1.0 | 29.0 | 2.98 [A:OP1]a:1671 | 11.20 | 8 | A | O5' | a:1671 | 3.77 | 1.0 | 24.6 |  | 16.59 | 9 | G | OP2 | a:2554 | 3.78 | 1.0 | 25.6 |  | 14.54 | 10 | Water 4 | | a:7679 | 3.98 | 1.0 | 25.6 | 2.86 [A:O3']a:1670 | 14.63 | 11 | C | OP1 | a:1672 | 4.15 | 1.0 | 23.5 |  | 14.67 | 12 | Water 5 | | a:4867 | 4.26 | 1.0 | 26.5 |  | 12.60 | 13 | A | O3' | a:1670 | 4.27 | 1.0 | 22.1 |  | 17.20 | 14 | A | O3' | a:1671 | 4.29 | 1.0 | 24.2 |  | 15.89 | 15 | A | OP1 | a:1671 | 4.30 | 1.0 | 22.9 |  | 16.40 | 16 | Water 6 | | a:3956 | 4.32 | 1.0 | 38.5 |  | 5.04 | 17 | G | O3' | a:2553 | 4.38 | 1.0 | 25.9 |  | 14.94 | 18 | C | O5' | a:1672 | 4.39 | 1.0 | 25.7 |  | 16.55 | 19 | G | O5' | a:2554 | 4.42 | 1.0 | 24.0 |  | 14.65 | 20 | C | OP2 | a:2555 | 4.47 | 1.0 | 29.6 |  | 13.14 | 21 | Water 7 | | a:9491 | 4.49 | 1.0 | 24.7 |  | 15.73 |

  
**1^st^ shell coordination angles and distances (*trans* in "blue"; *cys* in "green"; deviations in "orange"):**   
• 1-2(OP2/OP2): *97º*; 1-3(OP2/OP1): *95º*; 1-4(OP2/O4): *97º*; 1-5(OP2/W): *96º*; 2-3(OP2/OP1): 92º; 2-4(OP2/O4): *98º*; 2-5(OP2/W): *167º*; 3-4(OP1/O4): *163º*; 3-5(OP1/W): 87º; 4-5(O4/W): *80º*;   
• 1-2(OP2/OP2): 3.0Å; 1-3(OP2/OP1): 3.0Å; 1-4(OP2/O4): 3.1Å; 1-5(OP2/W): 3.1Å; 2-3(OP2/OP1): 3.0Å; 2-4(OP2/O4): 3.2Å; 2-5(OP2/W): *4.3Å*; 3-4(OP1/O4): *4.2Å*; 3-5(OP1/W): 3.0Å; 4-5(O4/W): 2.8Å;   
  
**2.17 ± 0.16: the average coodination distance of the 1^st^ six ligands is Mg^2+^ compatible**  
**4: Number of potential Water contacts**   
**2 : Number of second shell contacts involving water or ions**   
**50%: Percentage of satisfied second shell Water contacts**   
• Contact(s): /O4/O_w_/OP1/OP2/OP2/  
• Contact code: *fac-*3O_ph_O_b_O_w_  
• Contact(s) in the exclusion zone: /O_w_/  
→ Missing octahedral coordination for MG ←  
→ Slightly distorded coord. octahedron (7.9º) ←  
Go to ToC

### MG:a:3334

|  | Res. type | Atom | Residue | Distance | Occupancy | B-factor | Closest atom(s) to a 1^st^ shell water | EM\_dens. |
| --- | --- | --- | --- | --- | --- | --- | --- | --- |
| **Ion** | | | |  | 1.0 | 35.3 |  | 15.57 |
 1 | Water 1 | | a:6911 | 1.92 | 1.0 | 31.2 | 2.80 [A:N7]a:865 | 10.79 | 2 | G | OP2 | a:864 | 1.97 | 1.0 | 30.3 |  | 10.20 | 3 | Water 2 | | a:8215 | 2.15 | 1.0 | 35.3 |  | 7.72 | 4 | Water 3 | | a:9698 | 2.26 | 1.0 | 30.6 |  | 11.19 | 5 | Water 4 | | a:5236 | 2.33 | 1.0 | 26.4 | 2.69 [A:OP1]a:863 // 2.78 [A:OP1]a:912 | 12.88 | 6 | G | O5' | a:864 | 3.65 | 1.0 | 35.0 |  | 10.38 | 7 | Water 5 | | a:9650 | 3.89 | 1.0 | 38.5 |  | 5.82 | 8 | G | N7 | a:864 | 4.03 | 1.0 | 28.5 |  | 12.23 | 9 | G | OP1 | a:864 | 4.04 | 1.0 | 31.3 |  | 9.92 | 10 | A | N7 | a:865 | 4.23 | 1.0 | 29.4 |  | 9.34 | 11 | A | OP2 | a:863 | 4.28 | 1.0 | 36.3 |  | 7.65 | 12 | A | OP1 | a:863 | 4.31 | 1.0 | 29.9 |  | 10.01 | 13 | Water 6 | | a:5430 | 4.39 | 1.0 | 35.7 |  | 9.06 | 14 | A | O3' | a:863 | 4.45 | 1.0 | 30.7 |  | 11.91 | 15 | A | OP1 | a:912 | 4.46 | 1.0 | 30.9 |  | 8.73 | 16 | A | O5' | a:863 | 4.57 | 1.0 | 31.6 |  | 12.23 | 17 | Water 7 | | a:6828 | 4.73 | 1.0 | 42.8 |  | 3.91 |

  
**1^st^ shell coordination angles and distances (*trans* in "blue"; *cys* in "green"; deviations in "orange"):**   
• 1-2(W/OP2): *102º*; 1-3(W/W): *83º*; 1-4(W/W): 91º; 2-3(OP2/W): 89º; 2-4(OP2/W): *162º*; 3-4(W/W): *80º*;   
• 1-2(W/OP2): 3.0Å; 1-3(W/W): 2.7Å; 1-4(W/W): 3.0Å; 2-3(OP2/W): 2.9Å; 2-4(OP2/W): 4.2Å; 3-4(W/W): 2.8Å;   
  
**2.38 ± 0.59: the average coodination distance of the 1^st^ six ligands is Na^+^** compatible  
**8: Number of potential Water contacts**   
**3 : Number of second shell contacts involving water or ions**   
**38%: Percentage of satisfied second shell Water contacts**   
• Contact(s): /O_w_/O_w_/O_w_/OP2/  
• Contact code: O_ph_3O_w_  
• Contact(s) in the exclusion zone: /O_w_/  
→ Missing octahedral coordination for MG ←  
→ Slightly distorded coord. octahedron (8.2º) ←  
Go to ToC

### MG:a:3335

|  | Res. type | Atom | Residue | Distance | Occupancy | B-factor | Closest atom(s) to a 1^st^ shell water | EM\_dens. |
| --- | --- | --- | --- | --- | --- | --- | --- | --- |
| **Ion** | | | |  | 1.0 | 37.9 |  | 14.90 |
 1 | Water 1 | | a:7897 | 1.93 | 1.0 | 35.7 | 2.87 [C:OP2]a:1962 | 7.97 | 2 | Water 2 | | a:8351 | 1.96 | 1.0 | 37.4 |  | 7.83 | 3 | Water 3 | | a:8959 | 2.01 | 1.0 | 32.5 |  | 7.00 | 4 | A | OP2 | a:1942 | 2.03 | 1.0 | 34.0 |  | 7.71 | 5 | Water 4 | | a:8834 | 2.14 | 1.0 | 31.6 | 2.97 [C:OP1]a:1962 | 10.53 | 6 | Water 5 | | a:3560 | 2.32 | 1.0 | 32.1 | 2.48 [C:OP1]a:1961 | 11.59 | 7 | A | O5' | a:1942 | 3.67 | 1.0 | 36.7 |  | 7.61 | 8 | Water 6 | | a:9701 | 3.92 | 1.0 | 38.3 |  | 8.46 | 9 | Water 7 | | a:8878 | 4.06 | 1.0 | 41.6 |  | 5.61 | 10 | Water 8 | | a:8935 | 4.10 | 1.0 | 40.9 |  | 7.32 | 11 | C | OP1 | a:1961 | 4.10 | 1.0 | 34.2 |  | 10.13 | 12 | A | OP1 | a:1942 | 4.30 | 1.0 | 34.8 |  | 9.58 | 13 | C | OP2 | a:1962 | 4.30 | 1.0 | 24.8 |  | 12.65 | 14 | A | O3' | a:1941 | 4.43 | 1.0 | 29.8 |  | 11.00 | 15 | Water 9 | | a:7121 | 4.43 | 1.0 | 33.3 | 2.81 [C:OP1]a:1962 | 9.27 | 16 | Water 10 | | a:6216 | 4.46 | 1.0 | 42.7 |  | 4.20 | 17 | Water 11 | | a:10133 | 4.52 | 1.0 | 47.4 |  | 4.71 | 18 | C | OP1 | a:1962 | 4.58 | 1.0 | 36.6 |  | 8.98 | 19 | G | O4' | a:1772 | 4.65 | 1.0 | 35.0 |  | 11.94 |

  
**1^st^ shell coordination angles and distances (*trans* in "blue"; *cys* in "green"; deviations in "orange"):**   
• 1-2(W/W): 93º; 1-3(W/W): *167º*; 1-4(W/OP2): 94º; 1-5(W/W): *83º*; 2-3(W/W): *95º*; 2-4(W/OP2): 94º; 2-5(W/W): 172º; 3-4(W/OP2): *95º*; 3-5(W/W): 88º; 4-5(OP2/W): 93º;   
• 1-2(W/W): 2.8Å; 1-3(W/W): 3.9Å; 1-4(W/OP2): 2.9Å; 1-5(W/W): 2.7Å; 2-3(W/W): 2.9Å; 2-4(W/OP2): 2.9Å; 2-5(W/W): 4.1Å; 3-4(W/OP2): 3.0Å; 3-5(W/W): 2.9Å; 4-5(OP2/W): 3.0Å;   
  
**2.07 ± 0.13: the average coodination distance of the 1^st^ six ligands is Mg^2+^ compatible**  
**10: Number of potential Water contacts**   
**3 : Number of second shell contacts involving water or ions**   
**30%: Percentage of satisfied second shell Water contacts**   
• Contact(s): /O_w_/O_w_/O_w_/O_w_/OP2/  
• Contact code: O_ph_4O_w_  
• Contact(s) in the exclusion zone: /O_w_/  
→ Missing octahedral coordination for MG ←  
→ Slightly distorded coord. octahedron (5.3º) ←  
Go to ToC

### MG:a:3336

|  | Res. type | Atom | Residue | Distance | Occupancy | B-factor | Closest atom(s) to a 1^st^ shell water | EM\_dens. |
| --- | --- | --- | --- | --- | --- | --- | --- | --- |
| **Ion** | | | |  | 1.0 | 26.0 |  | 25.15 |
 1 | U | OP1 | a:2247 | 1.96 | 1.0 | 23.5 |  | 14.54 | 2 | C | OP1 | a:192 | 2.08 | 1.0 | 26.0 |  | 15.75 | 3 | Water 1 | | a:4401 | 2.12 | 1.0 | 19.9 | 2.62 [U:OP2]a:193 | 16.39 | 4 | Water 2 | | a:8851 | 2.13 | 1.0 | 22.2 |  | 16.31 | 5 | Water 3 | | a:4961 | 2.29 | 1.0 | 22.5 | 2.67 [A:OP1]a:199 // 2.99 [K:K]a:3277 | 16.55 | 6 | Water 4 | | a:9748 | 2.39 | 1.0 | 25.9 |  | 15.18 | 7 | U | O5' | a:2247 | 3.69 | 1.0 | 23.9 |  | 14.58 | 8 | Water 5 | | a:7142 | 3.77 | 1.0 | 39.0 |  | 5.12 | 9 | G | O3' | a:2246 | 3.91 | 1.0 | 25.0 |  | 16.15 | 10 | C | OP2 | a:192 | 3.94 | 1.0 | 22.8 |  | 17.54 | 11 | **MG** | MG | a:3068 | 3.95 | 1.0 | 40.5 |  | 11.77 | 12 | Water 6 | | a:3717 | 3.99 | 1.0 | 28.9 |  | 12.68 | 13 | C | O5' | a:192 | 4.02 | 1.0 | 28.4 |  | 13.72 | 14 | Water 7 | | a:5025 | 4.07 | 1.0 | 29.5 | 2.67 [U:OP2]a:193 | 14.25 | 15 | A | OP1 | a:199 | 4.09 | 1.0 | 27.0 |  | 13.58 | 16 | Water 8 | | a:5122 | 4.23 | 1.0 | 23.9 | 2.87 [K:K]a:3277 | 17.30 | 17 | U | OP2 | a:193 | 4.27 | 1.0 | 27.0 |  | 13.37 | 18 | Water 9 | | a:9973 | 4.39 | 1.0 | 39.8 |  | 5.43 | 19 | U | OP2 | a:2247 | 4.45 | 1.0 | 27.0 |  | 11.36 | 20 | **K** | K | a:3277 | 4.55 | 1.0 | 30.0 |  | 31.63 | 21 | A | O3' | a:191 | 4.55 | 1.0 | 21.8 |  | 15.81 |

  
**1^st^ shell coordination angles and distances (*trans* in "blue"; *cys* in "green"; deviations in "orange"):**   
• 1-2(OP1/OP1): *96º*; 1-3(OP1/W): 94º; 1-4(OP1/W): 93º; 1-5(OP1/W): *102º*; 2-3(OP1/W): *97º*; 2-4(OP1/W): *165º*; 2-5(OP1/W): *83º*; 3-4(W/W): *95º*; 3-5(W/W): *164º*; 4-5(W/W): *83º*;   
• 1-2(OP1/OP1): 3.0Å; 1-3(OP1/W): 3.0Å; 1-4(OP1/W): 3.0Å; 1-5(OP1/W): *3.3Å*; 2-3(OP1/W): 3.1Å; 2-4(OP1/W): 4.2Å; 2-5(OP1/W): 2.9Å; 3-4(W/W): 3.1Å; 3-5(W/W): *4.4Å*; 4-5(W/W): 2.9Å;   
  
**2.16 ± 0.14: the average coodination distance of the 1^st^ six ligands is Mg^2+^ compatible**  
**8: Number of potential Water contacts**   
**3 : Number of second shell contacts involving water or ions**   
**38%: Percentage of satisfied second shell Water contacts**   
• Contact(s): /O_w_/O_w_/O_w_/OP1/OP1/  
• Contact code: *cis-*2O_ph_3O_w_  
• Contact(s) in the exclusion zone: /O_w_/  
→ Missing octahedral coordination for MG ←  
→ Slightly distorded coord. octahedron (8.1º) ←  
Go to ToC

### MG:a:3337

|  | Res. type | Atom | Residue | Distance | Occupancy | B-factor | Closest atom(s) to a 1^st^ shell water | EM\_dens. |
| --- | --- | --- | --- | --- | --- | --- | --- | --- |
| **Ion** | | | |  | 1.0 | 24.8 |  | 24.30 |
 1 | Water 1 | | a:8550 | 2.00 | 1.0 | 21.0 |  | 15.24 | 2 | Water 2 | | a:7076 | 2.09 | 1.0 | 21.6 | 2.81 [G:OP2]a:1976 | 16.07 | 3 | G | OP2 | a:1830 | 2.13 | 1.0 | 28.8 |  | 15.01 | 4 | Water 3 | | a:6433 | 2.19 | 1.0 | 20.4 | 2.77 [U:OP2]a:1829 // 2.80 [G:N7]a:1830 | 15.35 | 5 | Water 4 | | a:9273 | 2.23 | 1.0 | 27.3 |  | 13.60 | 6 | Water 5 | | a:9266 | 2.33 | 1.0 | 24.8 |  | 14.50 | 7 | G | N7 | a:1830 | 3.97 | 1.0 | 18.6 |  | 15.61 | 8 | Water 6 | | a:3681 | 4.05 | 1.0 | 35.6 |  | 7.10 | 9 | G | OP2 | a:1976 | 4.24 | 1.0 | 26.9 |  | 12.40 | 10 | U | O5' | a:1829 | 4.32 | 1.0 | 25.2 |  | 14.12 | 11 | U | O3' | a:1829 | 4.32 | 1.0 | 23.5 |  | 14.64 | 12 | U | OP2 | a:1829 | 4.35 | 1.0 | 25.6 |  | 15.23 | 13 | G | OP1 | a:1830 | 4.37 | 1.0 | 23.5 |  | 14.72 | 14 | G | O5' | a:1830 | 4.40 | 1.0 | 22.4 |  | 13.71 | 15 | U | OP1 | a:1829 | 4.40 | 1.0 | 23.3 |  | 14.91 | 16 | Water 7 | | c:550 | 4.45 | 1.0 | 32.8 | 2.97 [ARG:NH1]c:221 | 9.32 | 17 | Water 8 | | a:9637 | 4.54 | 1.0 | 39.0 |  | 7.97 | 18 | Water 9 | | a:7509 | 4.63 | 1.0 | 24.9 |  | 15.64 |

  
**1^st^ shell coordination angles and distances (*trans* in "blue"; *cys* in "green"; deviations in "orange"):**   
• 1-2(W/W): 171º; 1-3(W/OP2): 91º; 1-4(W/W): 93º; 1-5(W/W): 94º; 2-3(W/OP2): 90º; 2-4(W/W): *96º*; 2-5(W/W): *84º*; 3-4(OP2/W): *97º*; 3-5(OP2/W): 172º; 4-5(W/W): 89º;   
• 1-2(W/W): 4.1Å; 1-3(W/OP2): 2.9Å; 1-4(W/W): 3.0Å; 1-5(W/W): 3.1Å; 2-3(W/OP2): 3.0Å; 2-4(W/W): 3.2Å; 2-5(W/W): 2.9Å; 3-4(OP2/W): *3.2Å*; 3-5(OP2/W): *4.3Å*; 4-5(W/W): 3.1Å;   
  
**2.16 ± 0.11: the average coodination distance of the 1^st^ six ligands is Mg^2+^ compatible**  
**10: Number of potential Water contacts**   
**3 : Number of second shell contacts involving water or ions**   
**30%: Percentage of satisfied second shell Water contacts**   
• Contact(s): /O_w_/O_w_/O_w_/O_w_/OP2/  
• Contact code: O_ph_4O_w_  
• Contact(s) in the exclusion zone: /O_w_/  
→ Missing octahedral coordination for MG ←  
Ooctahedral coordination (4.4º)   
Go to ToC

### K:a:3338

|  | Res. type | Atom | Residue | Distance | Occupancy | B-factor | Closest atom(s) to a 1^st^ shell water | EM\_dens. |
| --- | --- | --- | --- | --- | --- | --- | --- | --- |
| **Ion** | | | |  | 1.0 | 30.0 |  | 13.94 |
 1 | Water 1 | | a:4283 | 2.66 | 1.0 | 34.1 | 2.61 [G:N7]a:697 | 10.21 | 2 | Water 2 | | a:5408 | 2.85 | 1.0 | 35.7 | 2.70 [U:O4]a:768 | 7.57 | 3 | G | O6 | a:697 | 2.98 | 1.0 | 29.2 |  | 12.64 | 4 | U | O4 | a:769 | 3.00 | 1.0 | 31.9 |  | 10.87 | 5 | U | O4 | a:696 | 3.12 | 1.0 | 23.6 |  | 16.54 | 6 | Water 3 | | a:7514 | 3.60 | 1.0 | 33.6 |  | 7.58 | 7 | Water 4 | | a:6117 | 4.02 | 1.0 | 46.2 |  | 4.32 | 8 | U | N3 | a:769 | 4.07 | 1.0 | 20.8 |  | 13.48 | 9 | Water 5 | | a:9905 | 4.13 | 1.0 | 37.2 |  | 7.73 | 10 | G | N7 | a:697 | 4.27 | 1.0 | 24.0 |  | 14.41 | 11 | G | O6 | a:770 | 4.28 | 1.0 | 25.0 |  | 13.95 | 12 | Water 6 | | a:5905 | 4.37 | 1.0 | 35.7 |  | 6.52 | 13 | Water 7 | | a:10056 | 4.41 | 1.0 | 36.4 |  | 6.82 | 14 | U | O4 | a:768 | 4.67 | 1.0 | 28.5 |  | 10.62 | 15 | Water 8 | | a:5896 | 4.68 | 1.0 | 33.1 |  | 6.89 | 16 | Water 9 | | a:6636 | 4.78 | 1.0 | 29.2 |  | 10.06 |

  
**1^st^ shell coordination angles and distances (*trans* in "blue"; *cys* in "green"; deviations in "orange"):**   
• 1-2(W/W): 79; 1-3(W/O6): 74; 1-4(W/O4): 138; 1-5(W/O4): 69; 2-3(W/O6): 78; 2-4(W/O4): 69; 2-5(W/O4): 146; 3-4(O6/O4): 75; 3-5(O6/O4): 103; 4-5(O4/O4): 145;   
• 1-2(W/W): 3.5; 1-3(W/O6): 3.4; 1-4(W/O4): 5.3; 1-5(W/O4): 3.3; 2-3(W/O6): 3.7; 2-4(W/O4): 3.3; 2-5(W/O4): 5.7; 3-4(O6/O4): 3.6; 3-5(O6/O4): 4.8; 4-5(O4/O4): 5.8;   
  
**4.38 ± 1.39: the average coodination distance of the 1^st^ six ligands could correspond to an hexahydrated Mg^2+^.**  
**8: Number of potential Water contacts**   
**2 : Number of second shell contacts involving water or ions**   
**25%: Percentage of satisfied second shell Water contacts**   
• Contact(s): /O4/O4/O6/O_w_/O_w_/  
• Contact code: 3O_b_2O_w_  
→ Coordination < 6 for K ←  
Go to ToC

### K:a:3339

|  | Res. type | Atom | Residue | Distance | Occupancy | B-factor | Closest atom(s) to a 1^st^ shell water | EM\_dens. |
| --- | --- | --- | --- | --- | --- | --- | --- | --- |
| **Ion** | | | |  | 1.0 | 30.0 |  | 7.10 |
 1 | G | O6 | a:2494 | 2.92 | 1.0 | 24.8 |  | 11.88 | 2 | U | O4 | a:2493 | 2.96 | 1.0 | 30.6 |  | 10.44 | 3 | G | O2' | a:2462 | 3.33 | 1.0 | 33.0 |  | 11.74 | 4 | Water 1 | | a:8064 | 3.57 | 1.0 | 38.6 |  | 5.40 | 5 | Water 2 | | a:8130 | 3.71 | 1.0 | 43.5 | 2.89 [G:N7]a:2492 | 5.64 | 6 | **K** | K | a:3255 | 4.12 | 1.0 | 30.0 |  | 8.05 | 7 | G | N1 | a:2494 | 4.28 | 1.0 | 17.0 |  | 15.11 | 8 | A | OP2 | a:2463 | 4.57 | 1.0 | 42.6 |  | 3.09 | 9 | G | N7 | a:2492 | 4.60 | 1.0 | 22.0 |  | 13.52 | 10 | G | O3' | a:2462 | 4.76 | 1.0 | 40.9 |  | 7.61 |

  
**1^st^ shell coordination angles and distances (*trans* in "blue"; *cys* in "green"; deviations in "orange"):**   
• 1-2(O6/O4): 62;   
• 1-2(O6/O4): 3.0;   
  
**4.91 ± 1.56: the average coodination distance of the 1^st^ six ligands could correspond to an hexahydrated Mg^2+^.**  
**4: Number of potential Water contacts**   
**1 : Number of second shell contacts involving water or ions**   
**25%: Percentage of satisfied second shell Water contacts**   
• Contact(s): /O4/O6/  
• Contact code: 2O_b_  
→ Coordination < 6 for K ←  
Go to ToC

### K:a:3340

|  | Res. type | Atom | Residue | Distance | Occupancy | B-factor | Closest atom(s) to a 1^st^ shell water | EM\_dens. |
| --- | --- | --- | --- | --- | --- | --- | --- | --- |
| **Ion** | | | |  | 1.0 | 30.0 |  | 12.70 |
 1 | G | O6 | a:2374 | 2.76 | 1.0 | 34.0 |  | 9.59 | 2 | G | O6 | a:2375 | 2.81 | 1.0 | 37.7 |  | 8.16 | 3 | G | O6 | a:2349 | 2.86 | 1.0 | 32.8 |  | 9.76 | 4 | Water 1 | | a:4638 | 3.16 | 1.0 | 43.9 | 2.64 [G:N7]a:2374 | 4.73 | 5 | Water 2 | | a:6222 | 3.23 | 1.0 | 42.7 | 2.76 [G:N1]a:2349 | 4.84 | 6 | Water 3 | | a:5460 | 3.73 | 1.0 | 38.3 |  | 6.32 | 7 | C | N4 | a:2351 | 3.92 | 1.0 | 27.1 |  | 11.14 | 8 | G | N1 | a:2375 | 4.01 | 1.0 | 26.5 |  | 11.46 | 9 | G | N7 | a:2374 | 4.60 | 1.0 | 31.0 |  | 9.89 |

  
**1^st^ shell coordination angles and distances (*trans* in "blue"; *cys* in "green"; deviations in "orange"):**   
• 1-2(O6/O6): 68; 1-3(O6/O6): 85; 1-4(O6/W): 74; 2-3(O6/O6): 73; 2-4(O6/W): 74; 3-4(O6/W): 146;   
• 1-2(O6/O6): 3.1; 1-3(O6/O6): 3.8; 1-4(O6/W): 3.6; 2-3(O6/O6): 3.4; 2-4(O6/W): 3.6; 3-4(O6/W): 5.8;   
  
**4.41 ± 1.39: the average coodination distance of the 1^st^ six ligands could correspond to an hexahydrated Mg^2+^.**  
**6: Number of potential Water contacts**   
**2 : Number of second shell contacts involving water or ions**   
**33%: Percentage of satisfied second shell Water contacts**   
• Contact(s): /O6/O6/O6/O_w_/  
• Contact code: 3O_b_O_w_  
→ Coordination < 6 for K ←  
Go to ToC

### K:a:3341

|  | Res. type | Atom | Residue | Distance | Occupancy | B-factor | Closest atom(s) to a 1^st^ shell water | EM\_dens. |
| --- | --- | --- | --- | --- | --- | --- | --- | --- |
| **Ion** | | | |  | 1.0 | 30.0 |  | 7.10 |
 1 | G | O6 | a:2751 | 2.70 | 1.0 | 47.0 |  | 7.34 | 2 | Water 1 | | a:7358 | 2.80 | 1.0 | 47.9 |  | 4.34 | 3 | G | N7 | a:2751 | 2.97 | 1.0 | 43.6 |  | 6.55 | 4 | C | OP2 | a:2759 | 3.03 | 1.0 | 51.3 |  | 2.73 | 5 | U | O4 | a:2750 | 4.17 | 1.0 | 45.2 |  | 6.06 | 6 | U | OP2 | a:2758 | 4.39 | 1.0 | 44.4 |  | 4.61 | 7 | C | O5' | a:2759 | 4.46 | 1.0 | 39.7 |  | 6.66 | 8 | U | O5' | a:2758 | 4.61 | 1.0 | 50.0 |  | 4.51 |

  
**1^st^ shell coordination angles and distances (*trans* in "blue"; *cys* in "green"; deviations in "orange"):**   
• 1-2(O6/W): 133; 1-3(O6/N7): 65; 1-4(O6/OP2): 76; 2-3(W/N7): 115; 2-4(W/OP2): 87; 3-4(N7/OP2): 140;   
• 1-2(O6/W): 5.0; 1-3(O6/N7): 3.0; 1-4(O6/OP2): 3.5; 2-3(W/N7): 4.9; 2-4(W/OP2): 4.0; 3-4(N7/OP2): 5.6;   
  
**4.86 ± 1.67: the average coodination distance of the 1^st^ six ligands could correspond to an hexahydrated Mg^2+^.**  
**2: Number of potential Water contacts**   
**0 : Number of second shell contacts involving water or ions**   
**0%: Percentage of satisfied second shell Water contacts**   
• Contact(s): /N7/O6/O_w_/OP2/  
• Contact code: O_ph_O_b_N_b_O_w_  
→ Coordination < 6 for K ←  
Go to ToC

### K:a:3342

|  | Res. type | Atom | Residue | Distance | Occupancy | B-factor | Closest atom(s) to a 1^st^ shell water | EM\_dens. |
| --- | --- | --- | --- | --- | --- | --- | --- | --- |
| **Ion** | | | |  | 1.0 | 30.0 |  | 19.03 |
 1 | A | O2' | a:2780 | 2.78 | 1.0 | 32.0 |  | 9.75 | 2 | A | O2' | a:2785 | 2.78 | 1.0 | 32.2 |  | 11.57 | 3 | G | OP2 | a:2786 | 2.84 | 1.0 | 33.1 |  | 8.22 | 4 | G | N7 | a:2786 | 2.88 | 1.0 | 30.4 |  | 10.19 | 5 | Water 1 | | a:8061 | 2.90 | 1.0 | 42.9 | 2.89 [G:O6]a:2786 | 5.57 | 6 | A | O3' | a:2785 | 3.07 | 1.0 | 36.2 |  | 9.13 | 7 | Water 2 | | a:3927 | 3.63 | 1.0 | 40.2 | 2.91 [C:O2]a:2632 | 3.79 | 8 | A | O3' | a:2780 | 4.07 | 1.0 | 33.7 |  | 9.09 | 9 | G | O6 | a:2786 | 4.23 | 1.0 | 35.2 |  | 10.14 | 10 | G | O5' | a:2786 | 4.43 | 1.0 | 32.7 |  | 9.45 | 11 | G | OP2 | a:2781 | 4.44 | 1.0 | 34.2 |  | 9.71 | 12 | Water 3 | | a:6435 | 4.53 | 1.0 | 39.2 |  | 5.89 | 13 | C | O2 | a:2632 | 4.54 | 1.0 | 32.1 |  | 9.36 | 14 | G | OP1 | a:2786 | 4.76 | 1.0 | 34.3 |  | 9.55 |

  
**1^st^ shell coordination angles and distances (*trans* in "blue"; *cys* in "green"; deviations in "orange"):**   
• 1-2(O2'/O2'): 97; 1-3(O2'/OP2): 163; 1-4(O2'/N7): 73; 1-5(O2'/W): 75; 1-6(O2'/O3'): 122; 2-3(O2'/OP2): 89; 2-4(O2'/N7): 117; 2-5(O2'/W): 155; 2-6(O2'/O3'): 57; 3-4(OP2/N7): 89; 3-5(OP2/W): 105; 3-6(OP2/O3'): 50; 4-5(N7/W): 84; 4-6(N7/O3'): 76; 5-6(W/O3'): 147;   
• 1-2(O2'/O2'): 4.2; 1-3(O2'/OP2): 5.6; 1-4(O2'/N7): 3.4; 1-5(O2'/W): 3.5; 1-6(O2'/O3'): 5.1; 2-3(O2'/OP2): 4.0; 2-4(O2'/N7): 4.8; 2-5(O2'/W): 5.5; 2-6(O2'/O3'): 2.8; 3-4(OP2/N7): 4.0; 3-5(OP2/W): 4.5; 3-6(OP2/O3'): 2.5; 4-5(N7/W): 3.9; 4-6(N7/O3'): 3.7; 5-6(W/O3'): 5.7;   
  
**4.16 ± 1.31: the average coodination distance of the 1^st^ six ligands could correspond to an hexahydrated Mg^2+^.**  
**4: Number of potential Water contacts**   
**2 : Number of second shell contacts involving water or ions**   
**50%: Percentage of satisfied second shell Water contacts**   
• Contact(s): /N7/O2'/O2'/O3'/O_w_/OP2/  
• Contact code: O_ph_3O_r_N_b_O_w_  
Go to ToC

### K:a:3343

|  | Res. type | Atom | Residue | Distance | Occupancy | B-factor | Closest atom(s) to a 1^st^ shell water | EM\_dens. |
| --- | --- | --- | --- | --- | --- | --- | --- | --- |
| **Ion** | | | |  | 1.0 | 30.0 |  | 7.33 |
 1 | Water 1 | | a:5232 | 2.73 | 1.0 | 35.8 | 2.69 [G:N7]a:26 | 6.66 | 2 | G | O6 | a:26 | 2.81 | 1.0 | 21.4 |  | 14.69 | 3 | G | O6 | a:27 | 3.11 | 1.0 | 28.2 |  | 12.54 | 4 | Water 2 | | a:4439 | 3.28 | 1.0 | 43.2 | 2.63 [G:O2']a:512 | 4.23 | 5 | U | OP2 | a:25 | 3.97 | 1.0 | 34.4 |  | 11.12 | 6 | G | N7 | a:26 | 4.24 | 1.0 | 18.6 |  | 18.03 | 7 | G | N1 | a:27 | 4.32 | 1.0 | 17.1 |  | 15.05 | 8 | G | O2' | a:512 | 4.50 | 1.0 | 30.9 |  | 12.61 | 9 | Water 3 | | a:8835 | 4.71 | 1.0 | 24.6 |  | 11.84 | 10 | Water 4 | | a:7165 | 4.73 | 1.0 | 35.4 |  | 6.32 |

  
**1^st^ shell coordination angles and distances (*trans* in "blue"; *cys* in "green"; deviations in "orange"):**   
• 1-2(W/O6): 84; 1-3(W/O6): 69; 2-3(O6/O6): 70;   
• 1-2(W/O6): 3.7; 1-3(W/O6): 3.3; 2-3(O6/O6): 3.4;   
  
**4.83 ± 1.59: the average coodination distance of the 1^st^ six ligands could correspond to an hexahydrated Mg^2+^.**  
**4: Number of potential Water contacts**   
**2 : Number of second shell contacts involving water or ions**   
**50%: Percentage of satisfied second shell Water contacts**   
• Contact(s): /O6/O6/O_w_/  
• Contact code: 2O_b_O_w_  
→ Coordination < 6 for K ←  
Go to ToC

### K:a:3344

|  | Res. type | Atom | Residue | Distance | Occupancy | B-factor | Closest atom(s) to a 1^st^ shell water | EM\_dens. |
| --- | --- | --- | --- | --- | --- | --- | --- | --- |
| **Ion** | | | |  | 1.0 | 30.0 |  | 12.81 |
 1 | A | O2' | a:2827 | 2.74 | 1.0 | 26.8 |  | 14.09 | 2 | C | O2 | a:2010 | 2.79 | 1.0 | 23.8 |  | 14.64 | 3 | Water 1 | | a:7670 | 2.87 | 1.0 | 30.2 | 2.95 [MG:MG]a:3042 | 11.12 | 4 | A | N3 | a:1654 | 2.94 | 1.0 | 18.9 |  | 14.91 | 5 | **MG** | MG | a:3042 | 3.21 | 1.0 | 47.4 |  | 7.50 | 6 | A | O2' | a:1654 | 3.26 | 1.0 | 32.3 |  | 11.46 | 7 | A | O3' | a:2827 | 3.74 | 1.0 | 27.5 |  | 14.59 | 8 | G | N2 | a:1653 | 3.97 | 1.0 | 18.4 |  | 16.69 | 9 | Water 2 | | a:9734 | 4.03 | 1.0 | 37.9 |  | 5.24 | 10 | Water 3 | | a:3674 | 4.27 | 1.0 | 35.4 | 2.51 [C:OP1]a:2828 | 9.80 | 11 | C | N1 | a:2010 | 4.51 | 1.0 | 16.7 |  | 18.09 | 12 | C | N3 | a:2010 | 4.64 | 1.0 | 16.0 |  | 18.30 | 13 | C | O2' | a:2010 | 4.66 | 1.0 | 25.6 |  | 13.98 | 14 | Water 4 | | a:9670 | 4.68 | 1.0 | 31.0 |  | 9.50 | 15 | C | OP1 | a:2828 | 4.75 | 1.0 | 28.7 |  | 13.46 |

  
**1^st^ shell coordination angles and distances (*trans* in "blue"; *cys* in "green"; deviations in "orange"):**   
• 1-2(O2'/O2): 87; 1-3(O2'/W): 75; 1-4(O2'/N3): 156; 2-3(O2/W): 75; 2-4(O2/N3): 75; 3-4(W/N3): 87;   
• 1-2(O2'/O2): 3.8; 1-3(O2'/W): 3.4; 1-4(O2'/N3): 5.6; 2-3(O2/W): 3.4; 2-4(O2/N3): 3.5; 3-4(W/N3): 4.0;   
  
**4.25 ± 1.32: the average coodination distance of the 1^st^ six ligands could correspond to an hexahydrated Mg^2+^.**  
**2: Number of potential Water contacts**   
**1 : Number of second shell contacts involving water or ions**   
**50%: Percentage of satisfied second shell Water contacts**   
• Contact(s): /N3_A_/O2/O2'/O_w_/  
• Contact code: O_r_O_b_N_b_O_w_  
→ Coordination < 6 for K ←  
Go to ToC

### K:a:3345

|  | Res. type | Atom | Residue | Distance | Occupancy | B-factor | Closest atom(s) to a 1^st^ shell water | EM\_dens. |
| --- | --- | --- | --- | --- | --- | --- | --- | --- |
| **Ion** | | | |  | 1.0 | 30.0 |  | 16.17 |
 1 | Water 1 | | a:6622 | 2.75 | 1.0 | 32.6 |  | 9.36 | 2 | Water 2 | | a:9403 | 2.81 | 1.0 | 29.2 | 2.03 [MG:MG]a:3300 | 10.84 | 3 | Water 3 | | a:7244 | 2.82 | 1.0 | 44.1 | 2.82 [U:OP2]a:2269 | 4.96 | 4 | Water 4 | | a:4377 | 2.91 | 1.0 | 29.1 | 2.45 [MG:MG]a:3300 | 13.57 | 5 | A | O2' | a:2270 | 2.98 | 1.0 | 28.2 |  | 12.31 | 6 | Water 5 | | a:5240 | 3.25 | 1.0 | 41.8 | 2.69 [U:OP2]a:2269 | 4.36 | 7 | A | O3' | a:2270 | 3.41 | 1.0 | 28.7 |  | 12.16 | 8 | Water 6 | | a:10206 | 3.58 | 1.0 | 42.1 |  | 3.28 | 9 | **MG** | MG | a:3300 | 3.95 | 1.0 | 29.6 |  | 18.16 | 10 | Water 7 | | a:6319 | 4.04 | 1.0 | 27.3 | 2.11 [MG:MG]a:3300 | 13.44 | 11 | A | OP2 | a:2271 | 4.05 | 1.0 | 26.1 |  | 11.19 | 12 | U | OP2 | a:2269 | 4.19 | 1.0 | 34.1 |  | 9.85 | 13 | A | OP1 | a:2271 | 4.30 | 1.0 | 31.8 |  | 9.26 | 14 | Water 8 | | a:7552 | 4.34 | 1.0 | 36.9 |  | 6.95 | 15 | Water 9 | | a:5823 | 4.62 | 1.0 | 29.6 | 2.20 [MG:MG]a:3300 | 12.78 | 16 | Water 10 | | a:7953 | 4.63 | 1.0 | 44.3 | 2.88 [A:OP1]a:2271 | 5.84 | 17 | Water 11 | | a:10065 | 4.66 | 1.0 | 47.4 |  | 3.82 | 18 | U | O4 | a:2276 | 4.72 | 1.0 | 29.2 |  | 12.12 |

  
**1^st^ shell coordination angles and distances (*trans* in "blue"; *cys* in "green"; deviations in "orange"):**   
• 1-2(W/W): 81; 1-3(W/W): 148; 1-4(W/W): 69; 1-5(W/O2'): 80; 2-3(W/W): 130; 2-4(W/W): 66; 2-5(W/O2'): 154; 3-4(W/W): 124; 3-5(W/O2'): 72; 4-5(W/O2'): 91;   
• 1-2(W/W): 3.6; 1-3(W/W): 5.4; 1-4(W/W): 3.2; 1-5(W/O2'): 3.7; 2-3(W/W): 5.1; 2-4(W/W): 3.1; 2-5(W/O2'): 5.6; 3-4(W/W): 5.0; 3-5(W/O2'): 3.4; 4-5(W/O2'): 4.2;   
  
**4.08 ± 1.23: the average coodination distance of the 1^st^ six ligands could correspond to an hexahydrated Mg^2+^.**  
**12: Number of potential Water contacts**   
**4 : Number of second shell contacts involving water or ions**   
**33%: Percentage of satisfied second shell Water contacts**   
• Contact(s): /O2'/O_w_/O_w_/O_w_/O_w_/  
• Contact code: O_r_4O_w_  
→ Coordination < 6 for K ←  
Go to ToC

### K:a:3346

|  | Res. type | Atom | Residue | Distance | Occupancy | B-factor | Closest atom(s) to a 1^st^ shell water | EM\_dens. |
| --- | --- | --- | --- | --- | --- | --- | --- | --- |
| **Ion** | | | |  | 1.0 | 30.0 |  | 11.43 |
 1 | C | OP2 | a:1640 | 2.68 | 1.0 | 30.2 |  | 12.28 | 2 | Water 1 | | a:3529 | 2.84 | 1.0 | 36.8 | 2.46 [G:OP2]a:1301 | 7.53 | 3 | A | OP2 | a:1639 | 3.07 | 1.0 | 29.0 |  | 10.45 | 4 | A | O5' | a:1639 | 3.11 | 1.0 | 26.6 |  | 13.39 | 5 | Water 2 | | a:6035 | 3.78 | 1.0 | 31.9 | 2.74 [A:N7]a:1639 | 9.79 | 6 | Water 3 | | a:5940 | 3.79 | 1.0 | 38.2 | 2.74 [A:OP1]a:1639 | 5.23 | 7 | A | N7 | a:1639 | 4.01 | 1.0 | 23.4 |  | 15.02 | 8 | G | OP2 | a:1301 | 4.06 | 1.0 | 28.9 |  | 11.74 | 9 | Water 4 | | a:7351 | 4.06 | 1.0 | 33.2 |  | 10.31 | 10 | Water 5 | | a:8433 | 4.06 | 1.0 | 34.4 |  | 6.94 | 11 | Water 6 | | a:4020 | 4.28 | 1.0 | 35.1 |  | 9.89 | 12 | A | O3' | a:1639 | 4.29 | 1.0 | 29.0 |  | 13.14 | 13 | Water 7 | | a:9490 | 4.29 | 1.0 | 37.3 |  | 6.57 | 14 | A | OP1 | a:1639 | 4.45 | 1.0 | 32.8 |  | 8.62 | 15 | C | O5' | a:1640 | 4.55 | 1.0 | 23.6 |  | 12.71 |

  
**1^st^ shell coordination angles and distances (*trans* in "blue"; *cys* in "green"; deviations in "orange"):**   
• 1-2(OP2/W): 99; 1-3(OP2/OP2): 118; 1-4(OP2/O5'): 84; 2-3(W/OP2): 129; 2-4(W/O5'): 175; 3-4(OP2/O5'): 46;   
• 1-2(OP2/W): 4.2; 1-3(OP2/OP2): 4.9; 1-4(OP2/O5'): 3.9; 2-3(W/OP2): 5.3; 2-4(W/O5'): 5.9; 3-4(OP2/O5'): 2.4;   
  
**4.56 ± 1.44: the average coodination distance of the 1^st^ six ligands could correspond to an hexahydrated Mg^2+^.**  
**6: Number of potential Water contacts**   
**3 : Number of second shell contacts involving water or ions**   
**50%: Percentage of satisfied second shell Water contacts**   
• Contact(s): /O5'/O_w_/OP2/OP2/  
• Contact code: 2O_ph_O_r_O_w_  
→ Coordination < 6 for K ←  
Go to ToC

### K:a:3347

|  | Res. type | Atom | Residue | Distance | Occupancy | B-factor | Closest atom(s) to a 1^st^ shell water | EM\_dens. |
| --- | --- | --- | --- | --- | --- | --- | --- | --- |
| **Ion** | | | |  | 1.0 | 30.0 |  | 24.02 |
 1 | G | OP2 | a:973 | 2.73 | 1.0 | 24.8 |  | 14.50 | 2 | G | N7 | a:976 | 2.84 | 1.0 | 21.1 |  | 15.68 | 3 | G | O6 | a:991 | 2.86 | 1.0 | 30.0 |  | 13.43 | 4 | Water 1 | | a:9944 | 2.95 | 1.0 | 40.2 |  | 7.71 | 5 | Water 2 | | a:9578 | 3.01 | 1.0 | 47.7 |  | 2.84 | 6 | G | O6 | a:976 | 3.34 | 1.0 | 29.8 |  | 14.01 | 7 | Water 3 | | a:6575 | 3.58 | 1.0 | 27.1 |  | 12.06 | 8 | Water 4 | | a:10226 | 3.68 | 1.0 | 39.3 |  | 6.47 | 9 | G | OP1 | a:973 | 4.12 | 1.0 | 24.3 |  | 12.14 | 10 | Water 5 | | a:7487 | 4.28 | 1.0 | 26.8 |  | 14.97 | 11 | Water 6 | | a:6185 | 4.38 | 1.0 | 22.3 |  | 16.08 | 12 | Water 7 | | a:6973 | 4.49 | 1.0 | 29.5 |  | 9.53 | 13 | U | O3' | a:972 | 4.51 | 1.0 | 24.0 |  | 13.36 | 14 | U | O5' | a:972 | 4.75 | 1.0 | 33.2 |  | 10.22 |

  
**1^st^ shell coordination angles and distances (*trans* in "blue"; *cys* in "green"; deviations in "orange"):**   
• 1-2(OP2/N7): 82; 1-3(OP2/O6): 83; 1-4(OP2/W): 95; 1-5(OP2/W): 130; 2-3(N7/O6): 92; 2-4(N7/W): 93; 2-5(N7/W): 147; 3-4(O6/W): 175; 3-5(O6/W): 84; 4-5(W/W): 93;   
• 1-2(OP2/N7): 3.7; 1-3(OP2/O6): 3.7; 1-4(OP2/W): 4.2; 1-5(OP2/W): 5.2; 2-3(N7/O6): 4.1; 2-4(N7/W): 4.2; 2-5(N7/W): 5.6; 3-4(O6/W): 5.8; 3-5(O6/W): 3.9; 4-5(W/W): 4.3;   
  
**4.17 ± 1.26: the average coodination distance of the 1^st^ six ligands could correspond to an hexahydrated Mg^2+^.**  
**8: Number of potential Water contacts**   
**0 : Number of second shell contacts involving water or ions**   
**0%: Percentage of satisfied second shell Water contacts**   
• Contact(s): /N7/O6/O_w_/O_w_/OP2/  
• Contact code: O_ph_O_b_N_b_2O_w_  
→ Coordination < 6 for K ←  
Go to ToC

### MG:a:3348

|  | Res. type | Atom | Residue | Distance | Occupancy | B-factor | Closest atom(s) to a 1^st^ shell water | EM\_dens. |
| --- | --- | --- | --- | --- | --- | --- | --- | --- |
| **Ion** | | | |  | 1.0 | 23.2 |  | 28.23 |
 1 | G | OP2 | a:580 | 1.95 | 1.0 | 25.0 |  | 17.14 | 2 | Water 1 | | a:8727 | 2.02 | 1.0 | 21.2 |  | 17.77 | 3 | Water 2 | | a:8000 | 2.11 | 1.0 | 19.8 | 2.88 [G:N7]a:580 | 19.00 | 4 | Water 3 | | a:3868 | 2.11 | 1.0 | 20.1 | 2.55 [G:OP2]a:2022 | 18.51 | 5 | Water 4 | | a:6344 | 2.13 | 1.0 | 24.9 | 2.76 [G:OP2]a:579 | 15.40 | 6 | Water 5 | | a:4571 | 2.28 | 1.0 | 23.5 | 2.64 [U:OP1]a:2021 | 15.29 | 7 | G | O5' | a:580 | 3.73 | 1.0 | 20.4 |  | 16.92 | 8 | G | OP1 | a:580 | 4.09 | 1.0 | 25.2 |  | 15.33 | 9 | G | OP2 | a:2022 | 4.12 | 1.0 | 28.9 |  | 14.22 | 10 | Water 6 | | a:7207 | 4.21 | 1.0 | 40.1 | 2.88 [U:OP1]a:2021 | 5.19 | 11 | U | OP1 | a:2021 | 4.27 | 1.0 | 26.4 |  | 16.71 | 12 | G | N7 | a:580 | 4.33 | 1.0 | 17.2 |  | 19.42 | 13 | Water 7 | | a:3852 | 4.37 | 1.0 | 29.0 |  | 13.09 | 14 | Water 8 | | a:6626 | 4.38 | 1.0 | 38.2 |  | 6.74 | 15 | G | O3' | a:579 | 4.42 | 1.0 | 21.7 |  | 15.81 | 16 | Water 9 | | a:10057 | 4.46 | 1.0 | 36.0 |  | 9.45 | 17 | Water 10 | | a:3694 | 4.49 | 1.0 | 34.4 | 2.51 [G:OP1]a:580 // 2.86 [G:OP2]a:581 | 10.79 | 18 | Water 11 | | a:6202 | 4.49 | 1.0 | 30.8 | 2.84 [G:OP2]a:579 | 12.79 | 19 | U | OP2 | a:2021 | 4.49 | 1.0 | 22.6 |  | 15.04 | 20 | G | OP2 | a:581 | 4.51 | 1.0 | 27.0 |  | 14.88 | 21 | G | OP2 | a:579 | 4.53 | 1.0 | 23.6 |  | 16.04 | 22 | G | O5' | a:579 | 4.71 | 1.0 | 20.9 |  | 16.89 |

  
**1^st^ shell coordination angles and distances (*trans* in "blue"; *cys* in "green"; deviations in "orange"):**   
• 1-2(OP2/W): *96º*; 1-3(OP2/W): 94º; 1-4(OP2/W): 93º; 1-5(OP2/W): 93º; 1-6(OP2/W): 174º; 2-3(W/W): *167º*; 2-4(W/W): *95º*; 2-5(W/W): 89º; 2-6(W/W): 89º; 3-4(W/W): 91º; 3-5(W/W): *83º*; 3-6(W/W): *81º*; 4-5(W/W): 172º; 4-6(W/W): *84º*; 5-6(W/W): 89º;   
• 1-2(OP2/W): 3.0Å; 1-3(OP2/W): 3.0Å; 1-4(OP2/W): 3.0Å; 1-5(OP2/W): 3.0Å; 1-6(OP2/W): *4.2Å*; 2-3(W/W): 4.1Å; 2-4(W/W): 3.1Å; 2-5(W/W): 2.9Å; 2-6(W/W): 3.0Å; 3-4(W/W): 3.0Å; 3-5(W/W): 2.8Å; 3-6(W/W): 2.8Å; 4-5(W/W): *4.2Å*; 4-6(W/W): 3.0Å; 5-6(W/W): 3.1Å;   
  
**2.10 ± 0.10: the average coodination distance of the 1^st^ six ligands is Mg^2+^ compatible**  
**10: Number of potential Water contacts**   
**4 : Number of second shell contacts involving water or ions**   
**40%: Percentage of satisfied second shell Water contacts**   
• Contact(s): /O_w_/O_w_/O_w_/O_w_/O_w_/OP2/  
• Contact code: O_ph_5O_w_  
Ooctahedral coordination (4.9º)   
Go to ToC

### MG:a:3349

|  | Res. type | Atom | Residue | Distance | Occupancy | B-factor | Closest atom(s) to a 1^st^ shell water | EM\_dens. |
| --- | --- | --- | --- | --- | --- | --- | --- | --- |
| **Ion** | | | |  | 1.0 | 28.1 |  | 21.46 |
 1 | U | OP1 | a:965 | 2.03 | 1.0 | 19.9 |  | 14.43 | 2 | Water 1 | | a:9287 | 2.21 | 1.0 | 29.3 |  | 13.41 | 3 | Water 2 | | a:3637 | 2.24 | 1.0 | 25.7 | 2.50 [OMC:OP1]a:2502 | 14.50 | 4 | Water 3 | | a:6017 | 2.30 | 1.0 | 25.4 | 2.74 [A:OP1]a:2501 | 13.40 | 5 | Water 4 | | a:9876 | 2.31 | 1.0 | 28.6 |  | 11.67 | 6 | Water 5 | | a:6618 | 3.90 | 1.0 | 24.4 | 2.85 [U:OP2]a:965 | 12.21 | 7 | OMC | OP1 | a:2502 | 4.01 | 1.0 | 26.6 |  | 15.90 | 8 | Water 6 | | a:9802 | 4.11 | 1.0 | 30.0 |  | 13.00 | 9 | Water 7 | | a:8407 | 4.12 | 1.0 | 41.4 | 2.92 [A:OP1]a:949 | 5.74 | 10 | Water 8 | | a:10017 | 4.15 | 1.0 | 33.0 |  | 11.26 | 11 | U | O5' | a:965 | 4.18 | 1.0 | 25.0 |  | 14.53 | 12 | U | OP2 | a:965 | 4.18 | 1.0 | 22.0 |  | 16.00 | 13 | Water 9 | | a:6782 | 4.26 | 1.0 | 34.0 |  | 8.66 | 14 | Water 10 | | a:4556 | 4.28 | 1.0 | 36.9 | 2.64 [A:OP1]a:949 | 8.61 | 15 | G | O3' | a:964 | 4.41 | 1.0 | 23.8 |  | 17.93 | 16 | A | OP1 | a:949 | 4.47 | 1.0 | 25.7 |  | 13.60 | 17 | Water 11 | | a:8619 | 4.54 | 1.0 | 34.4 |  | 10.99 | 18 | A | OP1 | a:2501 | 4.63 | 1.0 | 23.8 |  | 15.48 | 19 | Water 12 | | a:5284 | 4.67 | 1.0 | 46.2 | 2.70 [OMC:OP1]a:2502 | 6.60 |

  
**1^st^ shell coordination angles and distances (*trans* in "blue"; *cys* in "green"; deviations in "orange"):**   
• 1-2(OP1/W): 92º; 1-3(OP1/W): 89º; 1-4(OP1/W): 90º; 2-3(W/W): 180º; 2-4(W/W): *84º*; 3-4(W/W): *96º*;   
• 1-2(OP1/W): 3.0Å; 1-3(OP1/W): 3.0Å; 1-4(OP1/W): 3.1Å; 2-3(W/W): *4.5Å*; 2-4(W/W): 3.0Å; 3-4(W/W): *3.4Å*;   
  
**2.50 ± 0.63: the average coodination distance of the 1^st^ six ligands is Na^+^** compatible  
**10: Number of potential Water contacts**   
**3 : Number of second shell contacts involving water or ions**   
**30%: Percentage of satisfied second shell Water contacts**   
• Contact(s): /O_w_/O_w_/O_w_/OP1/  
• Contact code: O_ph_3O_w_  
• Contact(s) in the exclusion zone: /O_w_/  
→ Missing octahedral coordination for MG ←  
Ooctahedral coordination (2.7º)   
Go to ToC

### MG:a:3350

|  | Res. type | Atom | Residue | Distance | Occupancy | B-factor | Closest atom(s) to a 1^st^ shell water | EM\_dens. |
| --- | --- | --- | --- | --- | --- | --- | --- | --- |
| **Ion** | | | |  | 1.0 | 51.7 |  | 9.18 |
 1 | Water 1 | | a:5769 | 2.05 | 1.0 | 39.3 | 2.73 [G:O6]a:493 | 8.04 | 2 | Water 2 | | a:8760 | 2.06 | 1.0 | 42.4 | 2.96 [G:N7]a:494 | 7.51 | 3 | Water 3 | | a:10098 | 2.07 | 1.0 | 45.3 |  | 4.69 | 4 | Water 4 | | a:10197 | 2.12 | 1.0 | 42.6 |  | 5.03 | 5 | Water 5 | | a:10230 | 2.15 | 1.0 | 36.8 |  | 5.72 | 6 | G | N7 | a:493 | 3.99 | 1.0 | 26.9 |  | 10.34 | 7 | Water 6 | | a:9412 | 4.04 | 1.0 | 41.7 |  | 4.32 | 8 | Water 7 | | a:5793 | 4.11 | 1.0 | 40.1 |  | 5.79 | 9 | A | OP2 | a:492 | 4.27 | 1.0 | 37.9 |  | 7.57 | 10 | G | O6 | a:493 | 4.32 | 1.0 | 31.1 |  | 9.47 | 11 | G | N7 | a:494 | 4.45 | 1.0 | 25.5 |  | 10.50 | 12 | G | O6 | a:494 | 4.62 | 1.0 | 32.3 |  | 10.86 | 13 | Water 8 | | a:8472 | 4.78 | 1.0 | 41.2 | 2.96 [G:O6]a:494 | 5.41 |

  
**1^st^ shell coordination angles and distances (*trans* in "blue"; *cys* in "green"; deviations in "orange"):**   
• 1-2(W/W): 91º; 1-3(W/W): 89º; 1-4(W/W): 89º; 1-5(W/W): 177º; 2-3(W/W): *169º*; 2-4(W/W): *85º*; 2-5(W/W): 89º; 3-4(W/W): *85º*; 3-5(W/W): 90º; 4-5(W/W): 88º;   
• 1-2(W/W): 2.9Å; 1-3(W/W): 2.9Å; 1-4(W/W): 2.9Å; 1-5(W/W): *4.2Å*; 2-3(W/W): 4.1Å; 2-4(W/W): 2.8Å; 2-5(W/W): 3.0Å; 3-4(W/W): 2.8Å; 3-5(W/W): 3.0Å; 4-5(W/W): 3.0Å;   
  
**2.41 ± 0.71: the average coodination distance of the 1^st^ six ligands is Na^+^** compatible  
**10: Number of potential Water contacts**   
**2 : Number of second shell contacts involving water or ions**   
**20%: Percentage of satisfied second shell Water contacts**   
• Contact(s): /O_w_/O_w_/O_w_/O_w_/O_w_/  
• Contact code: 5O_w_  
→ Missing octahedral coordination for MG ←  
Ooctahedral coordination (3.1º)   
Go to ToC

### MG:a:3351

|  | Res. type | Atom | Residue | Distance | Occupancy | B-factor | Closest atom(s) to a 1^st^ shell water | EM\_dens. |
| --- | --- | --- | --- | --- | --- | --- | --- | --- |
| **Ion** | | | |  | 1.0 | 34.5 |  | 16.29 |
 1 | Water 1 | | a:6037 | 1.95 | 1.0 | 35.2 |  | 9.33 | 2 | A | OP2 | a:1667 | 2.09 | 1.0 | 28.9 |  | 10.40 | 3 | Water 2 | | a:9020 | 2.12 | 1.0 | 26.1 |  | 13.27 | 4 | A | OP1 | a:1666 | 2.16 | 1.0 | 29.7 |  | 14.04 | 5 | Water 3 | | a:8747 | 2.25 | 1.0 | 39.7 |  | 8.39 | 6 | Water 4 | | a:4030 | 4.06 | 1.0 | 40.3 |  | 4.64 | 7 | A | O5' | a:1666 | 4.09 | 1.0 | 26.4 |  | 13.62 | 8 | A | OP1 | a:1667 | 4.10 | 1.0 | 33.2 |  | 10.95 | 9 | A | OP2 | a:1666 | 4.19 | 1.0 | 27.7 |  | 12.63 | 10 | Water 5 | | a:5902 | 4.20 | 1.0 | 35.0 |  | 7.65 | 11 | Water 6 | | a:9311 | 4.30 | 1.0 | 34.8 |  | 6.25 | 12 | A | O5' | a:1667 | 4.31 | 1.0 | 31.9 |  | 11.80 | 13 | Water 7 | | a:7579 | 4.38 | 1.0 | 33.9 |  | 9.79 | 14 | Water 8 | | a:9549 | 4.48 | 1.0 | 40.4 |  | 3.83 | 15 | A | O3' | a:1666 | 4.51 | 1.0 | 26.7 |  | 13.09 | 16 | G | O3' | a:1665 | 4.60 | 1.0 | 30.4 |  | 12.27 | 17 | Water 9 | | a:4969 | 4.62 | 1.0 | 21.5 | 2.95 [A:OP2]a:1666 | 17.82 | 18 | Water 10 | | a:6730 | 4.69 | 1.0 | 21.6 |  | 18.55 | 19 | Water 11 | | a:10058 | 4.76 | 1.0 | 35.5 |  | 9.73 |

  
**1^st^ shell coordination angles and distances (*trans* in "blue"; *cys* in "green"; deviations in "orange"):**   
• 1-2(W/OP2): 85º; 1-3(W/W): 174º; 1-4(W/OP1): 85º; 1-5(W/W): 95º; 2-3(OP2/W): *96º*; 2-4(OP2/OP1): 89º; 2-5(OP2/W): 173º; 3-4(W/OP1): 89º; 3-5(W/W): *83º*; 4-5(OP1/W): *84º*;   
• 1-2(W/OP2): 2.7Å; 1-3(W/W): 4.1Å; 1-4(W/OP1): 2.8Å; 1-5(W/W): 3.1Å; 2-3(OP2/W): 3.1Å; 2-4(OP2/OP1): 3.0Å; 2-5(OP2/W): *4.3Å*; 3-4(W/OP1): 3.0Å; 3-5(W/W): 2.9Å; 4-5(OP1/W): 3.0Å;   
  
**2.44 ± 0.73: the average coodination distance of the 1^st^ six ligands is Na^+^** compatible  
**8: Number of potential Water contacts**   
**0 : Number of second shell contacts involving water or ions**   
**0%: Percentage of satisfied second shell Water contacts**   
• Contact(s): /O_w_/O_w_/O_w_/OP1/OP2/  
• Contact code: *cis-*2O_ph_3O_w_  
→ Missing octahedral coordination for MG ←  
Ooctahedral coordination (4.9º)   
Go to ToC

### MG:a:3352

|  | Res. type | Atom | Residue | Distance | Occupancy | B-factor | Closest atom(s) to a 1^st^ shell water | EM\_dens. |
| --- | --- | --- | --- | --- | --- | --- | --- | --- |
| **Ion** | | | |  | 1.0 | 55.2 |  | 5.06 |
 1 | Water 1 | | a:7695 | 2.02 | 1.0 | 41.1 | 2.86 [G:O2']a:777 | 5.49 | 2 | Water 2 | | a:4757 | 2.03 | 1.0 | 34.3 | 2.65 [G:O6]a:683 | 7.65 | 3 | Water 3 | | a:4924 | 2.28 | 1.0 | 40.8 | 2.67 [G:OP1]a:778 | 5.76 | 4 | Water 4 | | a:6803 | 2.43 | 1.0 | 40.0 | 2.79 [G:N7]a:683 | 7.11 | 5 | Water 5 | | a:9631 | 3.81 | 1.0 | 41.1 |  | 4.91 | 6 | Water 6 | | a:9347 | 3.88 | 1.0 | 33.9 |  | 7.44 | 7 | Water 7 | | a:5469 | 4.02 | 1.0 | 42.8 |  | 4.71 | 8 | G | OP1 | a:778 | 4.16 | 1.0 | 28.1 |  | 12.00 | 9 | G | O6 | a:684 | 4.17 | 1.0 | 26.1 |  | 15.97 | 10 | Water 8 | | a:4501 | 4.26 | 1.0 | 38.8 |  | 7.41 | 11 | G | O6 | a:683 | 4.28 | 1.0 | 22.2 |  | 16.14 | 12 | Water 9 | | a:4778 | 4.35 | 1.0 | 28.8 |  | 11.40 | 13 | G | O2' | a:777 | 4.62 | 1.0 | 28.4 |  | 14.29 | 14 | G | O3' | a:777 | 4.66 | 1.0 | 34.1 |  | 12.62 | 15 | Water 10 | | a:9893 | 4.66 | 1.0 | 45.2 |  | 4.41 | 16 | Water 11 | | a:8030 | 4.66 | 1.0 | 40.4 |  | 7.56 | 17 | G | N7 | a:683 | 4.67 | 1.0 | 19.8 |  | 14.83 |

  
**1^st^ shell coordination angles and distances (*trans* in "blue"; *cys* in "green"; deviations in "orange"):**   
• 1-2(W/W): 90º; 1-3(W/W): *107º*; 2-3(W/W): *161º*;   
• 1-2(W/W): 2.9Å; 1-3(W/W): *3.5Å*; 2-3(W/W): *4.2Å*;   
  
**2.74 ± 0.79: the average coodination distance of the 1^st^ six ligands is K^+^ compatible**  
**12: Number of potential Water contacts**   
**4 : Number of second shell contacts involving water or ions**   
**33%: Percentage of satisfied second shell Water contacts**   
• Contact(s): /O_w_/O_w_/O_w_/  
• Contact code: 3O_w_  
• Contact(s) in the exclusion zone: /O_w_/  
→ Missing octahedral coordination for MG ←  
Go to ToC

### MG:a:3353

|  | Res. type | Atom | Residue | Distance | Occupancy | B-factor | Closest atom(s) to a 1^st^ shell water | EM\_dens. |
| --- | --- | --- | --- | --- | --- | --- | --- | --- |
| **Ion** | | | |  | 1.0 | 43.4 |  | 14.26 |
 1 | Water 1 | | a:7190 | 1.98 | 1.0 | 22.0 | 2.23 [MG:MG]a:3172 | 14.61 | 2 | Water 2 | | a:8014 | 2.15 | 1.0 | 35.9 |  | 7.46 | 3 | G | OP2 | a:2451 | 2.17 | 1.0 | 27.2 |  | 14.53 | 4 | Water 3 | | a:9603 | 2.18 | 1.0 | 30.0 | 1.96 [MG:MG]a:3172 | 13.07 | 5 | Water 4 | | a:5938 | 2.24 | 1.0 | 29.9 | 2.74 [C:OP2]a:2505 | 9.28 | 6 | **MG** | MG | a:3172 | 2.86 | 1.0 | 52.7 |  | 12.18 | 7 | Water 5 | | a:9821 | 3.79 | 1.0 | 37.9 |  | 8.43 | 8 | G | O5' | a:2451 | 3.86 | 1.0 | 24.4 |  | 13.31 | 9 | **K** | K | a:3110 | 3.89 | 1.0 | 30.0 |  | 19.46 | 10 | Water 6 | | a:4500 | 3.91 | 1.0 | 27.7 | 2.73 [G:OP1]a:2506 | 14.91 | 11 | Water 7 | | a:8656 | 3.92 | 1.0 | 34.7 |  | 8.96 | 12 | G | OP1 | a:2506 | 3.93 | 1.0 | 30.3 |  | 13.98 | 13 | G | OP1 | a:2451 | 4.11 | 1.0 | 24.2 |  | 13.46 | 14 | Water 8 | | a:9330 | 4.14 | 1.0 | 32.1 | 2.22 [MG:MG]a:3172 | 9.80 | 15 | G | OP1 | a:2450 | 4.20 | 1.0 | 32.2 |  | 13.70 | 16 | G | OP2 | a:2450 | 4.42 | 1.0 | 24.5 |  | 15.39 | 17 | G | O5' | a:2450 | 4.58 | 1.0 | 24.4 |  | 15.95 | 18 | C | OP2 | a:2505 | 4.60 | 1.0 | 25.7 |  | 13.23 | 19 | Water 9 | | a:6080 | 4.64 | 1.0 | 26.1 | 2.55 [MG:MG]a:3172 | 14.48 | 20 | G | N7 | a:2506 | 4.68 | 1.0 | 20.6 |  | 15.91 | 21 | G | O3' | a:2450 | 4.70 | 1.0 | 25.4 |  | 13.58 |

  
**1^st^ shell coordination angles and distances (*trans* in "blue"; *cys* in "green"; deviations in "orange"):**   
• 1-2(W/W): *98º*; 1-3(W/OP2): 87º; 1-4(W/W): 89º; 1-5(W/W): 171º; 2-3(W/OP2): *84º*; 2-4(W/W): 95º; 2-5(W/W): 91º; 3-4(OP2/W): 175º; 3-5(OP2/W): 94º; 4-5(W/W): 91º;   
• 1-2(W/W): 3.1Å; 1-3(W/OP2): 2.9Å; 1-4(W/W): 2.9Å; 1-5(W/W): *4.2Å*; 2-3(W/OP2): 2.9Å; 2-4(W/W): 3.2Å; 2-5(W/W): 3.1Å; 3-4(OP2/W): *4.4Å*; 3-5(OP2/W): *3.2Å*; 4-5(W/W): 3.2Å;   
  
**2.26 ± 0.28: the average coodination distance of the 1^st^ six ligands is Mg^2+^ compatible**  
**8: Number of potential Water contacts**   
**3 : Number of second shell contacts involving water or ions**   
**38%: Percentage of satisfied second shell Water contacts**   
• Contact(s): /O_w_/O_w_/O_w_/O_w_/OP2/  
• Contact code: O_ph_4O_w_  
→ Missing octahedral coordination for MG ←  
Ooctahedral coordination (4.4º)   
Go to ToC

### MG:a:3354

|  | Res. type | Atom | Residue | Distance | Occupancy | B-factor | Closest atom(s) to a 1^st^ shell water | EM\_dens. |
| --- | --- | --- | --- | --- | --- | --- | --- | --- |
| **Ion** | | | |  | 1.0 | 44.6 |  | 7.19 |
 1 | U | O2' | a:578 | 2.67 | 1.0 | 24.0 |  | 12.23 | 2 | G | O6 | a:2506 | 2.96 | 1.0 | 26.1 |  | 13.64 | 3 | Water 1 | | a:6658 | 3.17 | 1.0 | 34.3 | 2.95 [G:N1]a:2506 | 6.79 | 4 | G | O3' | a:810 | 3.47 | 1.0 | 21.0 |  | 17.97 | 5 | G | O4' | a:579 | 3.73 | 1.0 | 23.7 |  | 18.59 | 6 | **K** | K | a:3267 | 3.74 | 1.0 | 30.0 |  | 29.60 | 7 | U | O2 | a:578 | 3.84 | 1.0 | 25.0 |  | 15.55 | 8 | G | OP1 | a:811 | 4.01 | 1.0 | 24.4 |  | 16.07 | 9 | Water 2 | | a:9330 | 4.06 | 1.0 | 32.1 |  | 9.80 | 10 | Water 3 | | a:4567 | 4.14 | 1.0 | 38.8 |  | 5.99 | 11 | U | O2' | a:568 | 4.80 | 1.0 | 23.9 |  | 17.32 |

  
• • **3.29 ± 0.40: the average coodination distance of the 1^st^ six ligands is K^+^ compatible**  
**2: Number of potential Water contacts**   
**1 : Number of second shell contacts involving water or ions**   
**50%: Percentage of satisfied second shell Water contacts**   
• Contact code: → No contacts ←   
• Contact(s) in the exclusion zone: /O2'/O6/O_w_/  
→ Consider K in place of MG (2 valid contact(s) for K vs. 0 for MG ←)  
Go to ToC

### MG:a:3355

|  | Res. type | Atom | Residue | Distance | Occupancy | B-factor | Closest atom(s) to a 1^st^ shell water | EM\_dens. |
| --- | --- | --- | --- | --- | --- | --- | --- | --- |
| **Ion** | | | |  | 1.0 | 55.8 |  | 7.72 |
 1 | Water 1 | | a:5534 | 1.87 | 1.0 | 38.2 | 2.71 [A:OP2]a:191 | 7.17 | 2 | Water 2 | | a:10194 | 1.88 | 1.0 | 43.6 |  | 5.24 | 3 | Water 3 | | a:6955 | 2.38 | 1.0 | 31.3 |  | 10.91 | 4 | Water 4 | | w:128 | 2.41 | 1.0 | 40.1 |  | 5.56 | 5 | Water 5 | | a:10108 | 3.20 | 1.0 | 46.1 |  | 3.81 | 6 | A | N6 | a:203 | 4.02 | 1.0 | 23.0 |  | 12.21 | 7 | Water 6 | | a:5911 | 4.21 | 1.0 | 39.3 |  | 8.59 | 8 | Water 7 | | a:4564 | 4.22 | 1.0 | 42.0 |  | 6.77 | 9 | A | OP2 | a:191 | 4.22 | 1.0 | 29.2 |  | 13.79 | 10 | Water 8 | | a:7672 | 4.24 | 1.0 | 36.8 |  | 9.15 | 11 | Water 9 | | w:117 | 4.35 | 1.0 | 36.9 |  | 7.68 | 12 | Water 10 | | a:10162 | 4.43 | 1.0 | 40.9 |  | 5.08 | 13 | Water 11 | | a:6719 | 4.44 | 1.0 | 34.2 |  | 9.26 | 14 | Water 12 | | a:6005 | 4.67 | 1.0 | 30.9 |  | 13.05 |

  
**1^st^ shell coordination angles and distances (*trans* in "blue"; *cys* in "green"; deviations in "orange"):**   
• 1-2(W/W): *96º*;   
• 1-2(W/W): 2.8Å;   
  
**2.63 ± 0.76: the average coodination distance of the 1^st^ six ligands is K^+^ compatible**  
**10: Number of potential Water contacts**   
**1 : Number of second shell contacts involving water or ions**   
**10%: Percentage of satisfied second shell Water contacts**   
• Contact code: → No contacts ←   
• Contact(s) in the exclusion zone: /O_w_/O_w_/O_w_/O_w_/O_w_/  
→ Firts MG coordination shell missing ←  
Go to ToC

### MG:a:3356

|  | Res. type | Atom | Residue | Distance | Occupancy | B-factor | Closest atom(s) to a 1^st^ shell water | EM\_dens. |
| --- | --- | --- | --- | --- | --- | --- | --- | --- |
| **Ion** | | | |  | 1.0 | 45.1 |  | 6.77 |
 1 | Water 1 | | a:9115 | 1.92 | 1.0 | 39.7 |  | 5.63 | 2 | C | OP1 | a:825 | 2.02 | 1.0 | 33.5 |  | 9.30 | 3 | Water 2 | | a:8588 | 2.27 | 1.0 | 42.2 |  | 6.20 | 4 | Water 3 | | a:7757 | 2.40 | 1.0 | 37.1 | 2.88 [C:OP1]a:967 | 6.78 | 5 | Water 4 | | a:8396 | 3.51 | 1.0 | 40.3 | 2.13 [MG:MG]a:3206 | 8.21 | 6 | G | O3' | a:824 | 3.67 | 1.0 | 32.0 |  | 12.27 | 7 | Water 5 | | a:9677 | 3.88 | 1.0 | 41.5 | 2.12 [MG:MG]a:3206 | 4.88 | 8 | C | O5' | a:825 | 4.07 | 1.0 | 33.8 |  | 12.47 | 9 | G | OP1 | a:968 | 4.10 | 1.0 | 31.2 |  | 11.39 | 10 | Water 6 | | a:8214 | 4.28 | 1.0 | 35.0 | 2.90 [C:OP1]a:967 | 9.97 | 11 | C | O3' | a:967 | 4.39 | 1.0 | 25.3 |  | 13.84 | 12 | Water 7 | | a:8170 | 4.40 | 1.0 | 36.7 |  | 9.26 | 13 | Water 8 | | a:4893 | 4.42 | 1.0 | 38.3 |  | 6.91 | 14 | C | OP2 | a:825 | 4.51 | 1.0 | 30.9 |  | 10.84 | 15 | A | O4' | a:2277 | 4.54 | 1.0 | 30.8 |  | 13.95 | 16 | **MG** | MG | a:3206 | 4.56 | 1.0 | 51.3 |  | 9.73 | 17 | C | OP1 | a:967 | 4.62 | 1.0 | 29.4 |  | 11.02 |

  
**1^st^ shell coordination angles and distances (*trans* in "blue"; *cys* in "green"; deviations in "orange"):**   
• 1-2(W/OP1): *100º*; 1-3(W/W): *107º*; 2-3(OP1/W): *153º*;   
• 1-2(W/OP1): 3.0Å; 1-3(W/W): *3.4Å*; 2-3(OP1/W): 4.2Å;   
  
**2.63 ± 0.70: the average coodination distance of the 1^st^ six ligands is K^+^ compatible**  
**8: Number of potential Water contacts**   
**2 : Number of second shell contacts involving water or ions**   
**25%: Percentage of satisfied second shell Water contacts**   
• Contact(s): /O_w_/O_w_/OP1/  
• Contact code: O_ph_2O_w_  
• Contact(s) in the exclusion zone: /O_w_/  
→ Missing octahedral coordination for MG ←  
Go to ToC

### MG:a:3357

|  | Res. type | Atom | Residue | Distance | Occupancy | B-factor | Closest atom(s) to a 1^st^ shell water | EM\_dens. |
| --- | --- | --- | --- | --- | --- | --- | --- | --- |
| **Ion** | | | |  | 1.0 | 42.8 |  | 8.99 |
 1 | Water 1 | | a:6134 | 1.90 | 1.0 | 35.2 | 2.85 [G:OP2]a:1458 | 7.28 | 2 | G | OP2 | a:1457 | 1.91 | 1.0 | 39.0 |  | 5.56 | 3 | Water 2 | | a:7077 | 1.96 | 1.0 | 41.4 |  | 5.52 | 4 | Water 3 | | a:5261 | 2.05 | 1.0 | 35.5 | 2.69 [U:O4]a:1459 | 8.07 | 5 | Water 4 | | a:8062 | 2.19 | 1.0 | 41.0 | 2.89 [G:N7]a:1458 // 2.90 [K:K]a:3062 | 7.21 | 6 | Water 5 | | a:3764 | 2.24 | 1.0 | 38.4 | 2.53 [A:OP1]a:1455 // 2.74 [C:OP2]a:1456 | 8.06 | 7 | C | O2' | a:1456 | 4.07 | 1.0 | 41.2 |  | 8.09 | 8 | G | O5' | a:1457 | 4.08 | 1.0 | 36.8 |  | 6.66 | 9 | C | O3' | a:1456 | 4.10 | 1.0 | 44.4 |  | 6.61 | 10 | A | OP1 | a:1455 | 4.23 | 1.0 | 41.4 |  | 6.53 | 11 | G | OP2 | a:1458 | 4.26 | 1.0 | 43.8 |  | 4.58 | 12 | G | OP1 | a:1457 | 4.36 | 1.0 | 41.3 |  | 6.67 | 13 | G | N7 | a:1458 | 4.36 | 1.0 | 33.0 |  | 9.47 | 14 | **K** | K | a:3062 | 4.41 | 1.0 | 30.0 |  | 15.96 | 15 | U | O4 | a:1459 | 4.41 | 1.0 | 42.7 |  | 8.02 | 16 | C | OP2 | a:1456 | 4.44 | 1.0 | 47.9 |  | 4.62 |

  
**1^st^ shell coordination angles and distances (*trans* in "blue"; *cys* in "green"; deviations in "orange"):**   
• 1-2(W/OP2): 92º; 1-3(W/W): 91º; 1-4(W/W): *82º*; 1-5(W/W): 91º; 1-6(W/W): 170º; 2-3(OP2/W): 93º; 2-4(OP2/W): 174º; 2-5(OP2/W): *97º*; 2-6(OP2/W): *96º*; 3-4(W/W): 86º; 3-5(W/W): 170º; 3-6(W/W): 93º; 4-5(W/W): *85º*; 4-6(W/W): 89º; 5-6(W/W): *83º*;   
• 1-2(W/OP2): 2.8Å; 1-3(W/W): 2.7Å; 1-4(W/W): 2.6Å; 1-5(W/W): 2.9Å; 1-6(W/W): 4.1Å; 2-3(OP2/W): 2.8Å; 2-4(OP2/W): 4.0Å; 2-5(OP2/W): 3.1Å; 2-6(OP2/W): 3.1Å; 3-4(W/W): 2.7Å; 3-5(W/W): 4.1Å; 3-6(W/W): 3.1Å; 4-5(W/W): 2.9Å; 4-6(W/W): 3.0Å; 5-6(W/W): 2.9Å;   
  
**2.04 ± 0.13: the average coodination distance of the 1^st^ six ligands is Mg^2+^ compatible**  
**10: Number of potential Water contacts**   
**6 : Number of second shell contacts involving water or ions**   
**60%: Percentage of satisfied second shell Water contacts**   
• Contact(s): /O_w_/O_w_/O_w_/O_w_/O_w_/OP2/  
• Contact code: O_ph_5O_w_  
Ooctahedral coordination (4.9º)   
Go to ToC

### MG:a:3358

|  | Res. type | Atom | Residue | Distance | Occupancy | B-factor | Closest atom(s) to a 1^st^ shell water | EM\_dens. |
| --- | --- | --- | --- | --- | --- | --- | --- | --- |
| **Ion** | | | |  | 1.0 | 30.2 |  | 15.49 |
 1 | G | O6 | a:956 | 2.63 | 1.0 | 23.1 |  | 14.92 | 2 | Water 1 | | a:4719 | 2.79 | 1.0 | 29.6 | 2.65 [G:O6]a:964 | 7.58 | 3 | G | O6 | a:955 | 2.96 | 1.0 | 29.5 |  | 12.73 | 4 | Water 2 | | a:9478 | 3.06 | 1.0 | 48.8 |  | 4.62 | 5 | U | O4 | a:965 | 3.18 | 1.0 | 25.0 |  | 13.65 | 6 | PSU | O2 | a:957 | 3.57 | 1.0 | 25.8 |  | 14.50 | 7 | Water 3 | | a:4892 | 3.86 | 1.0 | 38.8 | 2.67 [G:N7]a:955 | 7.18 | 8 | G | O6 | a:964 | 3.89 | 1.0 | 28.1 |  | 14.02 | 9 | G | N7 | a:956 | 4.16 | 1.0 | 19.9 |  | 14.05 | 10 | Water 4 | | a:6718 | 4.26 | 1.0 | 27.4 | 2.79 [G:O6]a:955 | 11.57 | 11 | Water 5 | | l:339 | 4.30 | 1.0 | 39.4 |  | 7.66 | 12 | G | N7 | a:955 | 4.68 | 1.0 | 21.6 |  | 12.54 |

  
• • **3.03 ± 0.30: the average coodination distance of the 1^st^ six ligands is K^+^ compatible**  
**4: Number of potential Water contacts**   
**1 : Number of second shell contacts involving water or ions**   
**25%: Percentage of satisfied second shell Water contacts**   
• Contact code: → No contacts ←   
• Contact(s) in the exclusion zone: /O4/O6/O6/O_w_/O_w_/  
→ Consider K in place of MG (3 valid contact(s) for K vs. 0 for MG ←)  
Go to ToC

### MG:a:3359

|  | Res. type | Atom | Residue | Distance | Occupancy | B-factor | Closest atom(s) to a 1^st^ shell water | EM\_dens. |
| --- | --- | --- | --- | --- | --- | --- | --- | --- |
| **Ion** | | | |  | 1.0 | 48.5 |  | 8.30 |
 1 | Water 1 | | a:6718 | 2.05 | 1.0 | 27.4 | 2.79 [G:O6]a:954 // 2.79 [G:O6]a:955 | 11.57 | 2 | Water 2 | | a:8464 | 2.06 | 1.0 | 31.9 | 2.93 [G:N7]a:954 | 10.37 | 3 | Water 3 | | a:9774 | 2.16 | 1.0 | 41.4 |  | 7.61 | 4 | Water 4 | | a:4892 | 2.28 | 1.0 | 38.8 | 2.67 [G:N7]a:955 | 7.18 | 5 | Water 5 | | a:10290 | 2.34 | 1.0 | 46.2 |  | 5.06 | 6 | Water 6 | | a:9478 | 3.57 | 1.0 | 48.8 |  | 4.62 | 7 | Water 7 | | a:10204 | 4.14 | 1.0 | 48.8 |  | 4.16 | 8 | Water 8 | | l:350 | 4.15 | 1.0 | 43.9 |  | 4.23 | 9 | G | N7 | a:955 | 4.21 | 1.0 | 21.6 |  | 12.54 | 10 | Water 9 | | a:9950 | 4.22 | 1.0 | 45.4 |  | 4.57 | 11 | Water 10 | | a:8694 | 4.26 | 1.0 | 35.2 |  | 8.03 | 12 | Water 11 | | a:9429 | 4.31 | 1.0 | 37.0 |  | 7.76 | 13 | G | O6 | a:954 | 4.32 | 1.0 | 28.8 |  | 14.45 | 14 | G | O6 | a:955 | 4.49 | 1.0 | 29.5 |  | 12.73 | 15 | G | N7 | a:954 | 4.63 | 1.0 | 21.9 |  | 13.25 | 16 | Water 12 | | a:10283 | 4.70 | 1.0 | 44.6 |  | 4.48 |

  
**1^st^ shell coordination angles and distances (*trans* in "blue"; *cys* in "green"; deviations in "orange"):**   
• 1-2(W/W): *98º*; 1-3(W/W): 92º; 1-4(W/W): *79º*; 2-3(W/W): *102º*; 2-4(W/W): *83º*; 3-4(W/W): 171º;   
• 1-2(W/W): 3.1Å; 1-3(W/W): 3.0Å; 1-4(W/W): 2.8Å; 2-3(W/W): *3.3Å*; 2-4(W/W): 2.9Å; 3-4(W/W): *4.4Å*;   
  
**2.41 ± 0.53: the average coodination distance of the 1^st^ six ligands is Na^+^** compatible  
**12: Number of potential Water contacts**   
**4 : Number of second shell contacts involving water or ions**   
**33%: Percentage of satisfied second shell Water contacts**   
• Contact(s): /O_w_/O_w_/O_w_/O_w_/  
• Contact code: 4O_w_  
• Contact(s) in the exclusion zone: /O_w_/  
→ Missing octahedral coordination for MG ←  
→ Slightly distorded coord. octahedron (8.2º) ←  
Go to ToC

### MG:a:3360

|  | Res. type | Atom | Residue | Distance | Occupancy | B-factor | Closest atom(s) to a 1^st^ shell water | EM\_dens. |
| --- | --- | --- | --- | --- | --- | --- | --- | --- |
| **Ion** | | | |  | 1.0 | 48.0 |  | 12.11 |
 1 | Water 1 | | a:10103 | 1.95 | 1.0 | 42.6 |  | 5.36 | 2 | Water 2 | | a:9740 | 1.99 | 1.0 | 36.2 |  | 7.44 | 3 | Water 3 | | a:4890 | 2.04 | 1.0 | 37.1 | 2.67 [U:OP1]a:2691 | 7.48 | 4 | Water 4 | | a:10008 | 2.10 | 1.0 | 42.9 |  | 5.35 | 5 | Water 5 | | a:4073 | 2.48 | 1.0 | 30.1 | 2.58 [U:OP2]a:2691 | 10.08 | 6 | Water 6 | | a:10113 | 4.09 | 1.0 | 40.0 |  | 7.33 | 7 | Water 7 | | a:9311 | 4.11 | 1.0 | 34.8 |  | 6.25 | 8 | Water 8 | | a:3972 | 4.25 | 1.0 | 38.2 |  | 5.91 | 9 | Water 9 | | a:8417 | 4.27 | 1.0 | 43.7 |  | 5.04 | 10 | G | OP1 | a:2690 | 4.31 | 1.0 | 30.4 |  | 10.91 | 11 | Water 10 | | a:6174 | 4.32 | 1.0 | 42.3 | 2.75 [G:OP1]a:2690 | 5.91 | 12 | U | OP1 | a:2691 | 4.32 | 1.0 | 27.5 |  | 10.58 | 13 | Water 11 | | a:3914 | 4.35 | 1.0 | 40.2 | 2.96 [G:OP2]a:2722 | 5.95 | 14 | U | OP2 | a:2691 | 4.37 | 1.0 | 24.4 |  | 16.71 | 15 | G | OP2 | a:2722 | 4.56 | 1.0 | 34.2 |  | 10.20 | 16 | Water 12 | | a:6272 | 4.59 | 1.0 | 35.8 |  | 8.13 |

  
**1^st^ shell coordination angles and distances (*trans* in "blue"; *cys* in "green"; deviations in "orange"):**   
• 1-2(W/W): *97º*; 1-3(W/W): 93º; 1-4(W/W): *95º*; 2-3(W/W): 90º; 2-4(W/W): *84º*; 3-4(W/W): 170º;   
• 1-2(W/W): 2.9Å; 1-3(W/W): 2.9Å; 1-4(W/W): 3.0Å; 2-3(W/W): 2.8Å; 2-4(W/W): 2.7Å; 3-4(W/W): 4.1Å;   
  
**2.44 ± 0.76: the average coodination distance of the 1^st^ six ligands is Na^+^** compatible  
**12: Number of potential Water contacts**   
**2 : Number of second shell contacts involving water or ions**   
**17%: Percentage of satisfied second shell Water contacts**   
• Contact(s): /O_w_/O_w_/O_w_/O_w_/  
• Contact code: 4O_w_  
• Contact(s) in the exclusion zone: /O_w_/  
→ Missing octahedral coordination for MG ←  
→ Slightly distorded coord. octahedron (5.2º) ←  
Go to ToC

### MG:a:3361

|  | Res. type | Atom | Residue | Distance | Occupancy | B-factor | Closest atom(s) to a 1^st^ shell water | EM\_dens. |
| --- | --- | --- | --- | --- | --- | --- | --- | --- |
| **Ion** | | | |  | 1.0 | 50.4 |  | 8.55 |
 1 | Water 1 | | a:10303 | 2.01 | 1.0 | 47.3 |  | 4.61 | 2 | Water 2 | | a:9994 | 2.08 | 1.0 | 47.8 |  | 4.03 | 3 | Water 3 | | a:7789 | 2.15 | 1.0 | 41.0 | 2.86 [G:N7]a:2847 | 6.99 | 4 | Water 4 | | a:4196 | 2.19 | 1.0 | 36.7 | 2.60 [G:O6]a:2848 | 6.63 | 5 | Water 5 | | a:7225 | 2.34 | 1.0 | 39.8 | 2.82 [G:O6]a:2847 | 7.03 | 6 | Water 6 | | a:8604 | 3.66 | 1.0 | 32.9 |  | 8.34 | 7 | Water 7 | | a:5373 | 3.87 | 1.0 | 36.0 | 2.70 [G:O6]a:2848 | 7.29 | 8 | Water 8 | | a:6744 | 4.03 | 1.0 | 42.8 |  | 4.13 | 9 | Water 9 | | a:10274 | 4.24 | 1.0 | 50.7 |  | 3.42 | 10 | Water 10 | | a:7764 | 4.26 | 1.0 | 40.8 |  | 5.11 | 11 | G | N7 | a:2847 | 4.34 | 1.0 | 28.5 |  | 10.86 | 12 | G | O6 | a:2848 | 4.37 | 1.0 | 39.0 |  | 9.85 | 13 | G | O6 | a:2847 | 4.39 | 1.0 | 31.0 |  | 11.75 | 14 | U | O4 | a:2875 | 4.47 | 1.0 | 31.8 |  | 10.05 | 15 | Water 11 | | a:3691 | 4.49 | 1.0 | 44.8 |  | 3.90 | 16 | Water 12 | | a:7130 | 4.54 | 1.0 | 43.3 |  | 5.62 | 17 | Water 13 | | a:3618 | 4.54 | 1.0 | 43.3 | 2.49 [U:O4]a:2875 | 3.93 |

  
**1^st^ shell coordination angles and distances (*trans* in "blue"; *cys* in "green"; deviations in "orange"):**   
• 1-2(W/W): *96º*; 1-3(W/W): 89º; 1-4(W/W): 173º; 2-3(W/W): 86º; 2-4(W/W): 91º; 3-4(W/W): 90º;   
• 1-2(W/W): 3.0Å; 1-3(W/W): 2.9Å; 1-4(W/W): 4.2Å; 2-3(W/W): 2.9Å; 2-4(W/W): 3.0Å; 3-4(W/W): 3.1Å;   
  
**2.40 ± 0.57: the average coodination distance of the 1^st^ six ligands is Na^+^** compatible  
**12: Number of potential Water contacts**   
**3 : Number of second shell contacts involving water or ions**   
**25%: Percentage of satisfied second shell Water contacts**   
• Contact(s): /O_w_/O_w_/O_w_/O_w_/  
• Contact code: 4O_w_  
• Contact(s) in the exclusion zone: /O_w_/  
→ Missing octahedral coordination for MG ←  
Ooctahedral coordination (3.3º)   
Go to ToC

### MG:a:3362

|  | Res. type | Atom | Residue | Distance | Occupancy | B-factor | Closest atom(s) to a 1^st^ shell water | EM\_dens. |
| --- | --- | --- | --- | --- | --- | --- | --- | --- |
| **Ion** | | | |  | 1.0 | 52.9 |  | 5.74 |
 1 | Water 1 | | a:6326 | 2.01 | 1.0 | 32.7 | 2.76 [G:N7]a:1164 | 8.52 | 2 | Water 2 | | a:6574 | 2.20 | 1.0 | 35.7 | 2.78 [G:O6]a:1165 | 8.69 | 3 | Water 3 | | a:10263 | 2.30 | 1.0 | 46.5 |  | 3.56 | 4 | Water 4 | | a:7290 | 2.59 | 1.0 | 36.2 |  | 7.14 | 5 | Water 5 | | a:8785 | 3.48 | 1.0 | 44.5 |  | 4.93 | 6 | Water 6 | | a:4014 | 4.08 | 1.0 | 36.4 |  | 6.31 | 7 | G | N7 | a:1164 | 4.37 | 1.0 | 26.7 |  | 10.25 | 8 | G | O6 | a:1164 | 4.52 | 1.0 | 28.4 |  | 13.22 | 9 | Water 7 | | a:6329 | 4.54 | 1.0 | 37.9 |  | 5.20 | 10 | G | O6 | a:1165 | 4.59 | 1.0 | 29.1 |  | 13.56 | 11 | Water 8 | | a:5175 | 4.65 | 1.0 | 38.5 |  | 4.65 |

  
**1^st^ shell coordination angles and distances (*trans* in "blue"; *cys* in "green"; deviations in "orange"):**   
• 1-2(W/W): 91º;   
• 1-2(W/W): 3.0Å;   
  
**2.78 ± 0.75: the average coodination distance of the 1^st^ six ligands is K^+^ compatible**  
**12: Number of potential Water contacts**   
**2 : Number of second shell contacts involving water or ions**   
**17%: Percentage of satisfied second shell Water contacts**   
• Contact(s): /O_w_/O_w_/  
• Contact code: 2O_w_  
• Contact(s) in the exclusion zone: /O_w_/O_w_/  
→ Missing octahedral coordination for MG ←  
Go to ToC

### MG:a:3363

|  | Res. type | Atom | Residue | Distance | Occupancy | B-factor | Closest atom(s) to a 1^st^ shell water | EM\_dens. |
| --- | --- | --- | --- | --- | --- | --- | --- | --- |
| **Ion** | | | |  | 1.0 | 52.8 |  | 7.85 |
 1 | Water 1 | | a:8576 | 1.96 | 1.0 | 41.2 |  | 5.98 | 2 | OMU | OP2 | a:2556 | 2.11 | 1.0 | 36.0 |  | 6.41 | 3 | G | OP1 | a:1675 | 2.25 | 1.0 | 36.1 |  | 10.99 | 4 | **K** | K | a:3063 | 2.50 | 1.0 | 30.0 |  | 15.89 | 5 | Water 2 | | a:3956 | 3.38 | 1.0 | 38.5 |  | 5.04 | 6 | Water 3 | | a:3741 | 3.60 | 1.0 | 39.8 | 2.94 [K:K]a:3063 | 6.96 | 7 | Water 4 | | a:4845 | 3.68 | 1.0 | 23.3 | 2.66 [C:OP1]a:2555 | 15.70 | 8 | OMU | OP1 | a:2556 | 3.81 | 1.0 | 36.9 |  | 11.80 | 9 | A | O3' | a:1674 | 3.88 | 1.0 | 32.1 |  | 11.66 | 10 | Water 5 | | a:4867 | 3.92 | 1.0 | 26.5 |  | 12.60 | 11 | Water 6 | | a:6787 | 3.97 | 1.0 | 36.5 | 2.79 [OMU:OP1]a:2556 | 8.22 | 12 | OMU | O5' | a:2556 | 4.05 | 1.0 | 35.0 |  | 12.39 | 13 | Water 7 | | a:3507 | 4.14 | 1.0 | 44.2 |  | 5.89 | 14 | C | OP1 | a:2555 | 4.25 | 1.0 | 29.8 |  | 13.41 | 15 | G | O5' | a:1675 | 4.26 | 1.0 | 28.5 |  | 13.70 | 16 | C | OP2 | a:2555 | 4.37 | 1.0 | 29.6 |  | 13.14 | 17 | C | O5' | a:2555 | 4.48 | 1.0 | 29.8 |  | 12.32 | 18 | C | O3' | a:2555 | 4.58 | 1.0 | 26.8 |  | 12.93 | 19 | G | OP2 | a:1675 | 4.72 | 1.0 | 24.8 |  | 12.44 |

  
**1^st^ shell coordination angles and distances (*trans* in "blue"; *cys* in "green"; deviations in "orange"):**   
• 1-2(W/OP2): 93º; 1-3(W/OP1): *105º*; 2-3(OP2/OP1): *155º*;   
• 1-2(W/OP2): 2.9Å; 1-3(W/OP1): *3.3Å*; 2-3(OP2/OP1): *4.3Å*;   
  
**2.63 ± 0.63: the average coodination distance of the 1^st^ six ligands is K^+^ compatible**  
**6: Number of potential Water contacts**   
**1 : Number of second shell contacts involving water or ions**   
**17%: Percentage of satisfied second shell Water contacts**   
• Contact(s): /O_w_/OP1/OP2/  
• Contact code: *cis-*2O_ph_O_w_  
• Contact(s) in the exclusion zone: /O_w_/  
→ Missing octahedral coordination for MG ←  
Go to ToC

### MG:a:3364

|  | Res. type | Atom | Residue | Distance | Occupancy | B-factor | Closest atom(s) to a 1^st^ shell water | EM\_dens. |
| --- | --- | --- | --- | --- | --- | --- | --- | --- |
| **Ion** | | | |  | 1.0 | 55.2 |  | 6.85 |
 1 | Water 1 | | a:7669 | 1.96 | 1.0 | 36.9 | 2.85 [U:OP2]a:2266 | 8.16 | 2 | Water 2 | | v:117 | 2.09 | 1.0 | 40.3 | 2.80 [SER:O]v:16 | 7.85 | 3 | Water 3 | | v:105 | 2.11 | 1.0 | 43.3 | 2.63 [SER:OG]v:16 | 5.65 | 4 | C | OP2 | a:2267 | 2.50 | 1.0 | 34.9 |  | 10.22 | 5 | SER | OG | v:16 | 3.73 | 1.0 | 33.5 |  | 9.99 | 6 | Water 4 | | v:131 | 3.98 | 1.0 | 44.5 |  | 4.99 | 7 | SER | O | v:16 | 4.32 | 1.0 | 34.3 |  | 10.44 | 8 | Water 5 | | a:9874 | 4.33 | 1.0 | 45.2 |  | 4.22 | 9 | GLU | O | v:17 | 4.44 | 1.0 | 38.8 |  | 9.24 | 10 | Water 6 | | a:10229 | 4.45 | 1.0 | 46.7 |  | 4.56 | 11 | U | OP2 | a:2266 | 4.46 | 1.0 | 36.2 |  | 9.46 | 12 | ASP | OD1 | v:15 | 4.52 | 1.0 | 39.8 |  | 9.42 | 13 | C | O5' | a:2267 | 4.58 | 1.0 | 33.1 |  | 8.86 | 14 | Water 7 | | a:10065 | 4.61 | 1.0 | 47.4 |  | 3.82 | 15 | C | OP1 | a:2267 | 4.67 | 1.0 | 36.2 |  | 8.14 | 16 | Water 8 | | a:7870 | 4.79 | 1.0 | 43.4 |  | 5.15 |

  
**1^st^ shell coordination angles and distances (*trans* in "blue"; *cys* in "green"; deviations in "orange"):**   
• 1-2(W/W): 94º; 1-3(W/W): *96º*; 2-3(W/W): *96º*;   
• 1-2(W/W): 3.0Å; 1-3(W/W): 3.0Å; 2-3(W/W): 3.1Å;   
  
**2.73 ± 0.82: the average coodination distance of the 1^st^ six ligands is K^+^ compatible**  
**8: Number of potential Water contacts**   
**3 : Number of second shell contacts involving water or ions**   
**38%: Percentage of satisfied second shell Water contacts**   
• Contact(s): /O_w_/O_w_/O_w_/  
• Contact code: 3O_w_  
• Contact(s) in the exclusion zone: /OP2/  
→ Missing octahedral coordination for MG ←  
Go to ToC

### MG:a:3365

|  | Res. type | Atom | Residue | Distance | Occupancy | B-factor | Closest atom(s) to a 1^st^ shell water | EM\_dens. |
| --- | --- | --- | --- | --- | --- | --- | --- | --- |
| **Ion** | | | |  | 1.0 | 39.0 |  | 10.85 |
 1 | G | O6 | a:1336 | 2.58 | 1.0 | 28.5 |  | 11.73 | 2 | **MG** | MG | a:3366 | 2.63 | 1.0 | 35.9 |  | 12.93 | 3 | Water 1 | | a:9824 | 2.82 | 1.0 | 30.2 | 1.97 [MG:MG]a:3240 | 11.80 | 4 | G | O6 | a:1335 | 2.94 | 1.0 | 29.5 |  | 11.81 | 5 | Water 2 | | a:5548 | 3.05 | 1.0 | 44.5 | 2.71 [G:OP2]a:1334 | 5.00 | 6 | U | O4 | a:1320 | 3.33 | 1.0 | 32.8 |  | 10.71 | 7 | Water 3 | | a:5908 | 3.45 | 1.0 | 26.9 | 2.12 [MG:MG]a:3240 | 13.26 | 8 | G | O6 | a:1319 | 3.49 | 1.0 | 30.4 |  | 10.38 | 9 | G | N7 | a:1336 | 3.99 | 1.0 | 22.8 |  | 12.43 | 10 | Water 4 | | a:5539 | 4.01 | 1.0 | 46.2 | 2.71 [G:OP2]a:1334 // 2.81 [MG:MG]a:3366 | 5.09 | 11 | **MG** | MG | a:3240 | 4.11 | 1.0 | 28.9 |  | 18.70 | 12 | Water 5 | | a:6887 | 4.20 | 1.0 | 27.1 | 2.28 [MG:MG]a:3240 | 12.44 | 13 | G | OP2 | a:1334 | 4.43 | 1.0 | 26.0 |  | 11.33 | 14 | G | N7 | a:1335 | 4.57 | 1.0 | 21.9 |  | 12.72 |

  
• • **2.89 ± 0.26: the average coodination distance of the 1^st^ six ligands is K^+^ compatible**  
**4: Number of potential Water contacts**   
**2 : Number of second shell contacts involving water or ions**   
**50%: Percentage of satisfied second shell Water contacts**   
• Contact code: → No contacts ←   
• Contact(s) in the exclusion zone: /O4/O6/O6/O_w_/O_w_/  
→ Consider K in place of MG (2 valid contact(s) for K vs. 0 for MG ←)  
Go to ToC

### MG:a:3366

|  | Res. type | Atom | Residue | Distance | Occupancy | B-factor | Closest atom(s) to a 1^st^ shell water | EM\_dens. |
| --- | --- | --- | --- | --- | --- | --- | --- | --- |
| **Ion** | | | |  | 1.0 | 35.9 |  | 12.93 |
 1 | Water 1 | | a:9432 | 2.50 | 1.0 | 40.8 |  | 4.48 | 2 | **MG** | MG | a:3365 | 2.63 | 1.0 | 39.0 |  | 10.85 | 3 | G | O6 | a:1319 | 2.71 | 1.0 | 30.4 |  | 10.38 | 4 | U | O4 | a:1320 | 2.76 | 1.0 | 32.8 |  | 10.71 | 5 | Water 2 | | a:5539 | 2.81 | 1.0 | 46.2 | 2.71 [G:OP2]a:1334 | 5.09 | 6 | Water 3 | | a:5548 | 3.16 | 1.0 | 44.5 | 2.71 [G:OP2]a:1334 | 5.00 | 7 | G | N7 | a:1319 | 3.53 | 1.0 | 30.6 |  | 9.29 | 8 | U | O4 | a:1318 | 3.66 | 1.0 | 33.5 |  | 9.13 | 9 | Water 4 | | a:9824 | 3.79 | 1.0 | 30.2 | 2.82 [MG:MG]a:3365 | 11.80 | 10 | Water 5 | | a:5687 | 3.83 | 1.0 | 26.9 | 2.72 [U:O4]a:1320 | 10.46 | 11 | G | O6 | a:1336 | 4.32 | 1.0 | 28.5 |  | 11.73 | 12 | G | OP2 | a:1334 | 4.63 | 1.0 | 26.0 |  | 11.33 | 13 | Water 6 | | a:4083 | 4.70 | 1.0 | 43.1 |  | 3.59 |

  
• • **2.76 ± 0.20: the average coodination distance of the 1^st^ six ligands is K^+^ compatible**  
**6: Number of potential Water contacts**   
**2 : Number of second shell contacts involving water or ions**   
**33%: Percentage of satisfied second shell Water contacts**   
• Contact code: → No contacts ←   
• Contact(s) in the exclusion zone: /O4/O6/O_w_/O_w_/O_w_/  
→ Consider K in place of MG (3 valid contact(s) for K vs. 0 for MG ←)  
Go to ToC

### MG:b:201

|  | Res. type | Atom | Residue | Distance | Occupancy | B-factor | Closest atom(s) to a 1^st^ shell water | EM\_dens. |
| --- | --- | --- | --- | --- | --- | --- | --- | --- |
| **Ion** | | | |  | 1.0 | 67.3 |  | 2.19 |
 1 | C | OP2 | b:31 | 4.30 | 1.0 | 66.6 |  | 1.91 | 2 | U | O4 | b:32 | 4.31 | 1.0 | 66.6 |  | 2.24 | 3 | A | OP2 | b:46 | 4.37 | 1.0 | 62.2 |  | 2.68 | 4 | C | N4 | b:31 | 4.64 | 1.0 | 58.9 |  | 3.48 | 5 | U | O4 | b:48 | 4.68 | 1.0 | 61.0 |  | 3.26 |

  
• • **4.46 ± 0.17: the average coodination distance of the 1^st^ six ligands could correspond to an hexahydrated Mg^2+^.**  
**0: Number of potential Water contacts**   
• Contact code: → No contacts ←   
→ Firts MG coordination shell missing ←  
→2Fo-Fc density between 1.0 and 4.0 !   
Go to ToC

### MG:b:202

|  | Res. type | Atom | Residue | Distance | Occupancy | B-factor | Closest atom(s) to a 1^st^ shell water | EM\_dens. |
| --- | --- | --- | --- | --- | --- | --- | --- | --- |
| **Ion** | | | |  | 1.0 | 62.5 |  | 3.81 |
 1 | A | OP2 | b:58 | 1.90 | 1.0 | 60.8 |  | 3.80 | 2 | A | OP2 | b:57 | 2.08 | 1.0 | 67.6 |  | 1.99 | 3 | G | O3' | b:56 | 3.82 | 1.0 | 65.3 |  | 2.51 | 4 | A | OP1 | b:58 | 4.00 | 1.0 | 63.6 |  | 2.84 | 5 | G | O6 | b:23 | 4.15 | 1.0 | 63.3 |  | 3.29 | 6 | A | O5' | b:58 | 4.20 | 1.0 | 64.3 |  | 2.90 | 7 | A | O5' | b:57 | 4.23 | 1.0 | 59.6 |  | 4.32 | 8 | A | O3' | b:57 | 4.31 | 1.0 | 64.8 |  | 2.89 | 9 | U | O4 | b:22 | 4.32 | 1.0 | 66.8 |  | 2.72 | 10 | A | OP1 | b:57 | 4.56 | 1.0 | 66.3 |  | 1.99 | 11 | G | O2' | b:56 | 4.68 | 1.0 | 65.8 |  | 2.08 |

  
**1^st^ shell coordination angles and distances (*trans* in "blue"; *cys* in "green"; deviations in "orange"):**   
• 1-2(OP2/OP2): 89º;   
• 1-2(OP2/OP2): 2.8Å;   
  
**3.36 ± 0.98: the average coodination distance of the 1^st^ six ligands is K^+^ compatible**  
**0: Number of potential Water contacts**   
• Contact(s): /OP2/OP2/  
• Contact code: *cis-*2O_ph_  
→ Missing octahedral coordination for MG ←  
→2Fo-Fc density between 1.0 and 4.0 !   
Go to ToC

### MG:b:203

|  | Res. type | Atom | Residue | Distance | Occupancy | B-factor | Closest atom(s) to a 1^st^ shell water | EM\_dens. |
| --- | --- | --- | --- | --- | --- | --- | --- | --- |
| **Ion** | | | |  | 1.0 | 55.1 |  | 2.90 |
 1 | G | O6 | b:18 | 3.19 | 1.0 | 64.6 |  | 3.63 | 2 | G | O6 | b:64 | 3.26 | 1.0 | 65.6 |  | 2.73 | 3 | U | O4 | b:65 | 3.49 | 1.0 | 59.4 |  | 3.67 | 4 | C | N4 | b:19 | 4.02 | 1.0 | 65.4 |  | 3.18 | 5 | G | N7 | b:18 | 4.20 | 1.0 | 61.3 |  | 4.12 | 6 | C | N4 | b:17 | 4.55 | 1.0 | 56.7 |  | 4.32 |

  
• • **3.79 ± 0.51: the average coodination distance of the 1^st^ six ligands could correspond to an hexahydrated Mg^2+^.**  
**0: Number of potential Water contacts**   
• Contact code: → No contacts ←   
• Contact(s) in the exclusion zone: /O6/O6/  
→ Consider K in place of MG (1 valid contact(s) for K vs. 0 for MG ←)  
→2Fo-Fc density between 1.0 and 4.0 !   
Go to ToC

### MG:b:204

|  | Res. type | Atom | Residue | Distance | Occupancy | B-factor | Closest atom(s) to a 1^st^ shell water | EM\_dens. |
| --- | --- | --- | --- | --- | --- | --- | --- | --- |
| **Ion** | | | |  | 1.0 | 46.0 |  | 8.31 |
 1 | Water 1 | | b:312 | 1.83 | 1.0 | 41.3 |  | 4.62 | 2 | Water 2 | | b:336 | 1.97 | 1.0 | 40.8 | 2.76 [A:N7]b:101 // 2.97 [A:OP2]b:101 | 6.43 | 3 | Water 3 | | b:377 | 2.06 | 1.0 | 38.9 |  | 5.95 | 4 | G | OP2 | b:100 | 2.14 | 1.0 | 36.3 |  | 7.75 | 5 | Water 4 | | b:315 | 2.17 | 1.0 | 40.5 | 2.60 [G:O6]b:102 | 6.39 | 6 | G | O5' | b:100 | 3.99 | 1.0 | 34.9 |  | 8.63 | 7 | Water 5 | | b:330 | 4.13 | 1.0 | 37.5 |  | 8.46 | 8 | G | OP1 | b:100 | 4.20 | 1.0 | 40.4 |  | 6.27 | 9 | A | N7 | b:101 | 4.30 | 1.0 | 25.6 |  | 10.33 | 10 | G | O6 | b:102 | 4.32 | 1.0 | 40.1 |  | 6.25 | 11 | Water 6 | | b:304 | 4.45 | 1.0 | 41.2 |  | 3.95 | 12 | A | OP2 | b:101 | 4.46 | 1.0 | 33.9 |  | 9.22 | 13 | A | OP2 | b:73 | 4.47 | 1.0 | 48.3 |  | 4.01 | 14 | Water 7 | | b:361 | 4.48 | 1.0 | 43.1 | 2.93 [A:OP2]b:101 | 4.04 | 15 | A | O3' | b:99 | 4.61 | 1.0 | 38.8 |  | 5.84 |

  
**1^st^ shell coordination angles and distances (*trans* in "blue"; *cys* in "green"; deviations in "orange"):**   
• 1-2(W/W): 171º; 1-3(W/W): *97º*; 1-4(W/OP2): *80º*; 1-5(W/W): 89º; 2-3(W/W): 91º; 2-4(W/OP2): *97º*; 2-5(W/W): 92º; 3-4(W/OP2): *99º*; 3-5(W/W): 91º; 4-5(OP2/W): *166º*;   
• 1-2(W/W): *3.8Å*; 1-3(W/W): 2.9Å; 1-4(W/OP2): 2.6Å; 1-5(W/W): 2.8Å; 2-3(W/W): 2.9Å; 2-4(W/OP2): 3.1Å; 2-5(W/W): 3.0Å; 3-4(W/OP2): *3.2Å*; 3-5(W/W): 3.0Å; 4-5(OP2/W): *4.3Å*;   
  
**2.36 ± 0.74: the average coodination distance of the 1^st^ six ligands is Na^+^** compatible  
**8: Number of potential Water contacts**   
**3 : Number of second shell contacts involving water or ions**   
**38%: Percentage of satisfied second shell Water contacts**   
• Contact(s): /O_w_/O_w_/O_w_/OP2/  
• Contact code: O_ph_3O_w_  
• Contact(s) in the exclusion zone: /O_w_/  
→ Missing octahedral coordination for MG ←  
→ Slightly distorded coord. octahedron (6.1º) ←  
Go to ToC

### MG:b:205

|  | Res. type | Atom | Residue | Distance | Occupancy | B-factor | Closest atom(s) to a 1^st^ shell water | EM\_dens. |
| --- | --- | --- | --- | --- | --- | --- | --- | --- |
| **Ion** | | | |  | 1.0 | 38.7 |  | 12.55 |
 1 | U | O4 | b:95 | 1.92 | 1.0 | 35.3 |  | 10.76 | 2 | Water 1 | | b:355 | 1.94 | 1.0 | 35.0 |  | 8.93 | 3 | Water 2 | | b:339 | 2.07 | 1.0 | 35.8 |  | 8.53 | 4 | Water 3 | | b:356 | 2.10 | 1.0 | 28.6 | 2.89 [A:N7]b:94 | 10.38 | 5 | Water 4 | | b:380 | 2.12 | 1.0 | 37.8 |  | 7.86 | 6 | Water 5 | | b:310 | 2.18 | 1.0 | 34.4 | 2.54 [U:O4]b:80 // 2.67 [G:O6]b:96 | 8.71 | 7 | Water 6 | | b:353 | 3.85 | 1.0 | 40.3 |  | 6.00 | 8 | Water 7 | | b:332 | 3.94 | 1.0 | 30.0 |  | 8.06 | 9 | Water 8 | | b:326 | 4.09 | 1.0 | 43.4 |  | 6.60 | 10 | U | O4 | b:80 | 4.20 | 1.0 | 33.9 |  | 10.67 | 11 | A | N7 | b:94 | 4.23 | 1.0 | 31.3 |  | 9.25 | 12 | Water 9 | | b:345 | 4.31 | 1.0 | 33.2 |  | 6.19 | 13 | G | O6 | b:96 | 4.36 | 1.0 | 30.5 |  | 11.68 |

  
**1^st^ shell coordination angles and distances (*trans* in "blue"; *cys* in "green"; deviations in "orange"):**   
• 1-2(O4/W): *97º*; 1-3(O4/W): 89º; 1-4(O4/W): *95º*; 1-5(O4/W): 173º; 1-6(O4/W): 95º; 2-3(W/W): 173º; 2-4(W/W): 93º; 2-5(W/W): 89º; 2-6(W/W): 94º; 3-4(W/W): *85º*; 3-5(W/W): *84º*; 3-6(W/W): 87º; 4-5(W/W): *81º*; 4-6(W/W): *167º*; 5-6(W/W): 89º;   
• 1-2(O4/W): 2.9Å; 1-3(O4/W): 2.8Å; 1-4(O4/W): 3.0Å; 1-5(O4/W): 4.0Å; 1-6(O4/W): 3.0Å; 2-3(W/W): 4.0Å; 2-4(W/W): 2.9Å; 2-5(W/W): 2.9Å; 2-6(W/W): 3.0Å; 3-4(W/W): 2.8Å; 3-5(W/W): 2.8Å; 3-6(W/W): 2.9Å; 4-5(W/W): 2.7Å; 4-6(W/W): *4.3Å*; 5-6(W/W): 3.0Å;   
  
**2.05 ± 0.09: the average coodination distance of the 1^st^ six ligands is Mg^2+^ compatible**  
**10: Number of potential Water contacts**   
**3 : Number of second shell contacts involving water or ions**   
**30%: Percentage of satisfied second shell Water contacts**   
• Contact(s): /O4/O_w_/O_w_/O_w_/O_w_/O_w_/  
• Contact code: O_b_5O_w_  
→ Slightly distorded coord. octahedron (5.0º) ←  
Go to ToC

### K:b:206

|  | Res. type | Atom | Residue | Distance | Occupancy | B-factor | Closest atom(s) to a 1^st^ shell water | EM\_dens. |
| --- | --- | --- | --- | --- | --- | --- | --- | --- |
| **Ion** | | | |  | 1.0 | 30.0 |  | 6.39 |
 1 | G | O6 | b:16 | 2.55 | 1.0 | 56.6 |  | 4.05 | 2 | G | N7 | b:16 | 2.82 | 1.0 | 53.6 |  | 5.16 | 3 | A | OP1 | b:15 | 2.89 | 1.0 | 55.2 |  | 2.23 | 4 | A | O4' | b:108 | 3.40 | 1.0 | 57.3 |  | 2.89 | 5 | C | N4 | b:17 | 4.15 | 1.0 | 56.7 |  | 4.32 | 6 | A | O5' | b:15 | 4.28 | 1.0 | 54.5 |  | 4.54 | 7 | A | OP2 | b:15 | 4.32 | 1.0 | 51.2 |  | 3.53 | 8 | G | N1 | b:69 | 4.54 | 1.0 | 47.9 |  | 5.68 | 9 | A | N3 | b:108 | 4.66 | 1.0 | 52.7 |  | 5.44 | 10 | A | O3' | b:108 | 4.74 | 1.0 | 59.0 |  | 2.76 | 11 | G | O6 | b:69 | 4.77 | 1.0 | 51.5 |  | 3.87 |

  
**1^st^ shell coordination angles and distances (*trans* in "blue"; *cys* in "green"; deviations in "orange"):**   
• 1-2(O6/N7): 69; 1-3(O6/OP1): 112; 2-3(N7/OP1): 91;   
• 1-2(O6/N7): 3.0; 1-3(O6/OP1): 4.5; 2-3(N7/OP1): 4.1;   
  
**4.83 ± 1.64: the average coodination distance of the 1^st^ six ligands could correspond to an hexahydrated Mg^2+^.**  
**0: Number of potential Water contacts**   
• Contact(s): /N7/OP1/  
• Contact code: O_ph_N_b_  
• Contact(s) in the exclusion zone: /O6/  
→ Coordination < 6 for K ←  
Go to ToC

### K:c:301

|  | Res. type | Atom | Residue | Distance | Occupancy | B-factor | Closest atom(s) to a 1^st^ shell water | EM\_dens. |
| --- | --- | --- | --- | --- | --- | --- | --- | --- |
| **Ion** | | | |  | 1.0 | 30.0 |  | 19.57 |
 1 | ASP | O | c:84 | 2.53 | 1.0 | 32.0 |  | 13.65 | 2 | Water 1 | | c:584 | 2.69 | 1.0 | 41.8 |  | 5.88 | 3 | ARG | O | c:87 | 2.69 | 1.0 | 33.9 |  | 12.55 | 4 | Water 2 | | c:477 | 2.82 | 1.0 | 43.1 | 2.78 [SER:O]c:88 | 5.39 | 5 | Water 3 | | c:464 | 2.95 | 1.0 | 32.0 | 2.73 [ALA:O]c:89 | 11.17 | 6 | TYR | OH | c:83 | 3.66 | 1.0 | 38.2 |  | 10.62 | 7 | PRO | N | c:85 | 4.25 | 1.0 | 21.4 |  | 13.70 | 8 | ASP | N | c:84 | 4.30 | 1.0 | 23.0 |  | 11.62 | 9 | Water 4 | | c:453 | 4.32 | 1.0 | 32.4 | 2.69 [PRO:O]c:85 | 11.43 | 10 | SER | O | c:88 | 4.47 | 1.0 | 33.2 |  | 11.84 | 11 | Water 5 | | c:472 | 4.52 | 1.0 | 37.5 |  | 5.31 | 12 | PRO | O | c:85 | 4.53 | 1.0 | 29.2 |  | 14.68 | 13 | ALA | O | c:89 | 4.58 | 1.0 | 35.1 |  | 10.43 |

  
**1^st^ shell coordination angles and distances (*trans* in "blue"; *cys* in "green"; deviations in "orange"):**   
• 1-2(W/W): 116; 1-3(W/W): 74; 1-4(W/W): 145; 1-5(W/W): 73; 2-3(W/W): 83; 2-4(W/W): 93; 2-5(W/W): 166; 3-4(W/W): 92; 3-5(W/W): 89; 4-5(W/W): 76;   
• 1-2(W/W): 4.4; 1-3(W/W): 3.1; 1-4(W/W): 5.1; 1-5(W/W): 3.3; 2-3(W/W): 3.6; 2-4(W/W): 4.0; 2-5(W/W): 5.6; 3-4(W/W): 4.0; 3-5(W/W): 4.0; 4-5(W/W): 3.5;   
  
**4.32 ± 1.47: the average coodination distance of the 1^st^ six ligands could correspond to an hexahydrated Mg^2+^.**  
**10: Number of potential Water contacts**   
**2 : Number of second shell contacts involving water or ions**   
**20%: Percentage of satisfied second shell Water contacts**   
• Contact(s): /O_back_/O_w_/O_w_/O_w_/  
• Contact code: O_back_3O_w_  
• Contact(s) in the exclusion zone: /O_back_/  
→ Coordination < 6 for K ←  
Go to ToC

### MG:c:302

|  | Res. type | Atom | Residue | Distance | Occupancy | B-factor | Closest atom(s) to a 1^st^ shell water | EM\_dens. |
| --- | --- | --- | --- | --- | --- | --- | --- | --- |
| **Ion** | | | |  | 1.0 | 52.1 |  | 6.08 |
 1 | Water 1 | | c:444 | 1.96 | 1.0 | 40.3 | 2.67 [PRO:O]c:85 | 7.58 | 2 | Water 2 | | a:10125 | 1.99 | 1.0 | 41.8 |  | 5.04 | 3 | Water 3 | | c:579 | 2.03 | 1.0 | 42.2 |  | 5.88 | 4 | Water 4 | | c:465 | 2.08 | 1.0 | 37.0 | 2.74 [GLN:OE1]c:60 | 8.66 | 5 | Water 5 | | c:544 | 3.54 | 1.0 | 42.2 | 2.96 [ASN:O]c:86 | 6.16 | 6 | Water 6 | | a:7524 | 3.84 | 1.0 | 41.2 |  | 4.00 | 7 | GLN | OE1 | c:60 | 4.25 | 1.0 | 33.5 |  | 11.74 | 8 | Water 7 | | a:3889 | 4.26 | 1.0 | 38.3 |  | 9.20 | 9 | PRO | O | c:85 | 4.32 | 1.0 | 29.2 |  | 14.68 | 10 | Water 8 | | c:587 | 4.32 | 1.0 | 39.0 |  | 8.49 | 11 | ASN | O | c:86 | 4.39 | 1.0 | 30.3 |  | 12.44 | 12 | Water 9 | | c:435 | 4.43 | 1.0 | 40.5 |  | 5.97 |

  
**1^st^ shell coordination angles and distances (*trans* in "blue"; *cys* in "green"; deviations in "orange"):**   
• 1-2(W/W): *97º*; 1-3(W/W): 89º; 1-4(W/W): 93º; 2-3(W/W): *83º*; 2-4(W/W): *170º*; 3-4(W/W): *97º*;   
• 1-2(W/W): 3.0Å; 1-3(W/W): 2.8Å; 1-4(W/W): 2.9Å; 2-3(W/W): 2.7Å; 2-4(W/W): 4.1Å; 3-4(W/W): 3.1Å;   
  
**2.57 ± 0.80: the average coodination distance of the 1^st^ six ligands is Na^+^** compatible  
**12: Number of potential Water contacts**   
**3 : Number of second shell contacts involving water or ions**   
**25%: Percentage of satisfied second shell Water contacts**   
• Contact(s): /O_w_/O_w_/O_w_/O_w_/  
• Contact code: 4O_w_  
→ Missing octahedral coordination for MG ←  
→ Slightly distorded coord. octahedron (5.9º) ←  
Go to ToC

### K:c:303

|  | Res. type | Atom | Residue | Distance | Occupancy | B-factor | Closest atom(s) to a 1^st^ shell water | EM\_dens. |
| --- | --- | --- | --- | --- | --- | --- | --- | --- |
| **Ion** | | | |  | 1.0 | 30.0 |  | 27.12 |
 1 | Water 1 | | a:8187 | 2.69 | 1.0 | 23.4 | 2.73 [K:K]c:305 | 14.56 | 2 | Water 2 | | a:5743 | 2.75 | 1.0 | 30.2 | 2.73 [A:OP2]a:2594 | 10.30 | 3 | Water 3 | | a:4703 | 2.90 | 1.0 | 19.2 | 2.09 [MG:MG]a:3225 | 18.20 | 4 | Water 4 | | c:426 | 2.95 | 1.0 | 30.9 |  | 11.97 | 5 | GLU | OE2 | c:236 | 2.99 | 1.0 | 43.8 |  | 6.80 | 6 | GLY | O | c:235 | 3.00 | 1.0 | 29.3 |  | 14.27 | 7 | A | OP2 | a:2593 | 3.12 | 1.0 | 24.5 |  | 18.24 | 8 | Water 5 | | c:483 | 3.31 | 1.0 | 29.6 |  | 10.23 | 9 | Water 6 | | c:478 | 3.42 | 1.0 | 26.2 |  | 13.14 | 10 | Water 7 | | a:4992 | 4.05 | 1.0 | 27.6 |  | 13.22 | 11 | A | OP1 | a:2593 | 4.27 | 1.0 | 21.9 |  | 17.32 | 12 | Water 8 | | a:7901 | 4.27 | 1.0 | 20.9 | 2.98 [A:OP1]a:2593 | 16.90 | 13 | A | OP2 | a:2594 | 4.27 | 1.0 | 30.0 |  | 13.18 | 14 | **K** | K | c:305 | 4.43 | 1.0 | 30.0 |  | 29.54 | 15 | A | O5' | a:2593 | 4.55 | 1.0 | 18.8 |  | 18.19 | 16 | Water 9 | | c:407 | 4.63 | 1.0 | 31.6 | 2.45 [GLU:OE1]c:236 | 10.43 | 17 | Water 10 | | a:3538 | 4.64 | 1.0 | 25.5 | 2.37 [MG:MG]a:3225 | 14.16 | 18 | GLU | OE1 | c:236 | 4.66 | 1.0 | 46.1 |  | 4.35 | 19 | **MG** | MG | a:3225 | 4.79 | 1.0 | 26.7 |  | 24.33 |

  
**1^st^ shell coordination angles and distances (*trans* in "blue"; *cys* in "green"; deviations in "orange"):**   
• 1-2(W/W): 80; 1-3(W/W): 98; 1-4(W/W): 130; 1-5(W/OE2): 155; 1-6(W/W): 73; 1-7(W/OP2): 86; 2-3(W/W): 136; 2-4(W/W): 142; 2-5(W/OE2): 91; 2-6(W/W): 97; 2-7(W/OP2): 84; 3-4(W/W): 71; 3-5(W/OE2): 105; 3-6(W/W): 125; 3-7(W/OP2): 52; 4-5(W/OE2): 52; 4-6(W/W): 75; 4-7(W/OP2): 117; 5-6(OE2/W): 85; 5-7(OE2/OP2): 117; 6-7(W/OP2): 158;   
• 1-2(W/W): 3.5; 1-3(W/W): 4.2; 1-4(W/W): 5.1; 1-5(W/OE2): 5.5; 1-6(W/W): 3.4; 1-7(W/OP2): 4.0; 2-3(W/W): 5.2; 2-4(W/W): 5.4; 2-5(W/OE2): 4.1; 2-6(W/W): 4.3; 2-7(W/OP2): 3.9; 3-4(W/W): 3.4; 3-5(W/OE2): 4.7; 3-6(W/W): 5.2; 3-7(W/OP2): 2.6; 4-5(W/OE2): 2.6; 4-6(W/W): 3.6; 4-7(W/OP2): 5.2; 5-6(OE2/W): 4.1; 5-7(OE2/OP2): 5.2; 6-7(W/OP2): 6.0;   
  
**3.95 ± 1.16: the average coodination distance of the 1^st^ six ligands could correspond to an hexahydrated Mg^2+^.**  
**12: Number of potential Water contacts**   
**3 : Number of second shell contacts involving water or ions**   
**25%: Percentage of satisfied second shell Water contacts**   
• Contact(s): /O_back_/O_w_/O_w_/O_w_/O_w_/OE2_Glu_/OP2/  
• Contact code: O_ph_O_back_O_coo_4O_w_  
Go to ToC

### MG:c:304

|  | Res. type | Atom | Residue | Distance | Occupancy | B-factor | Closest atom(s) to a 1^st^ shell water | EM\_dens. |
| --- | --- | --- | --- | --- | --- | --- | --- | --- |
| **Ion** | | | |  | 1.0 | 22.8 |  | 25.47 |
 1 | G | OP2 | a:1826 | 2.04 | 1.0 | 26.0 |  | 15.33 | 2 | Water 1 | | a:6974 | 2.06 | 1.0 | 20.5 | 2.80 [G:OP1]a:1825 // 2.88 [ARG:O]c:52 | 14.23 | 3 | Water 2 | | c:570 | 2.06 | 1.0 | 25.2 |  | 15.81 | 4 | Water 3 | | c:495 | 2.07 | 1.0 | 24.2 | 2.83 [THR:OG1]c:219 | 14.82 | 5 | Water 4 | | a:9059 | 2.18 | 1.0 | 24.2 |  | 17.02 | 6 | HIS | ND1 | c:53 | 2.26 | 1.0 | 19.0 |  | 15.12 | 7 | G | OP1 | a:1826 | 4.08 | 1.0 | 29.1 |  | 11.72 | 8 | THR | OG1 | c:219 | 4.11 | 1.0 | 30.5 |  | 12.03 | 9 | Water 5 | | a:4471 | 4.15 | 1.0 | 26.3 |  | 15.38 | 10 | G | O3' | a:1825 | 4.21 | 1.0 | 24.7 |  | 15.56 | 11 | Water 6 | | a:10273 | 4.21 | 1.0 | 41.6 |  | 3.47 | 12 | HIS | NE2 | c:53 | 4.28 | 1.0 | 20.4 |  | 17.84 | 13 | Water 7 | | c:419 | 4.29 | 1.0 | 24.7 | 2.55 [THR:OG1]c:219 | 15.55 | 14 | Water 8 | | a:6000 | 4.35 | 1.0 | 24.4 |  | 13.10 | 15 | ARG | O | c:52 | 4.41 | 1.0 | 24.1 |  | 16.41 | 16 | G | O5' | a:1825 | 4.43 | 1.0 | 26.5 |  | 11.77 | 17 | G | OP1 | a:1825 | 4.46 | 1.0 | 32.7 |  | 11.04 | 18 | G | O5' | a:1826 | 4.47 | 1.0 | 23.3 |  | 13.72 | 19 | Water 9 | | a:6922 | 4.49 | 1.0 | 38.8 | 2.80 [G:OP2]a:1825 | 6.45 | 20 | G | OP2 | a:1825 | 4.67 | 1.0 | 28.0 |  | 12.96 |

  
**1^st^ shell coordination angles and distances (*trans* in "blue"; *cys* in "green"; deviations in "orange"):**   
• 1-2(OP2/W): 88º; 1-3(OP2/W): 174º; 1-4(OP2/W): 92º; 1-5(OP2/W): 90º; 1-6(OP2/ND1): 94º; 2-3(W/W): 94º; 2-4(W/W): 178º; 2-5(W/W): 92º; 2-6(W/ND1): 91º; 3-4(W/W): 86º; 3-5(W/W): *84º*; 3-6(W/ND1): 92º; 4-5(W/W): 90º; 4-6(W/ND1): 87º; 5-6(W/ND1): 175º;   
• 1-2(OP2/W): 2.9Å; 1-3(OP2/W): 4.1Å; 1-4(OP2/W): 3.0Å; 1-5(OP2/W): 3.0Å; 1-6(OP2/ND1): 3.2Å; 2-3(W/W): 3.0Å; 2-4(W/W): 4.1Å; 2-5(W/W): 3.0Å; 2-6(W/ND1): 3.1Å; 3-4(W/W): 2.8Å; 3-5(W/W): 2.8Å; 3-6(W/ND1): 3.1Å; 4-5(W/W): 3.0Å; 4-6(W/ND1): 3.0Å; 5-6(W/ND1): *4.4Å*;   
  
**2.11 ± 0.08: the average coodination distance of the 1^st^ six ligands is Mg^2+^ compatible**  
**8: Number of potential Water contacts**   
**3 : Number of second shell contacts involving water or ions**   
**38%: Percentage of satisfied second shell Water contacts**   
• Contact(s): /ND1_His_/O_w_/O_w_/O_w_/O_w_/OP2/  
• Contact code: O_ph_4O_w_N_His_  
Ooctahedral coordination (3.0º)   
Go to ToC

### K:c:305

|  | Res. type | Atom | Residue | Distance | Occupancy | B-factor | Closest atom(s) to a 1^st^ shell water | EM\_dens. |
| --- | --- | --- | --- | --- | --- | --- | --- | --- |
| **Ion** | | | |  | 1.0 | 30.0 |  | 29.54 |
 1 | Water 1 | | a:8187 | 2.73 | 1.0 | 23.4 | 2.69 [K:K]c:303 | 14.56 | 2 | GLY | O | c:237 | 2.77 | 1.0 | 34.1 |  | 13.22 | 3 | GLY | O | c:235 | 2.78 | 1.0 | 29.3 |  | 14.27 | 4 | Water 2 | | c:475 | 2.89 | 1.0 | 27.0 | 2.92 [THR:O]c:223 | 14.84 | 5 | GLY | O | c:234 | 2.94 | 1.0 | 30.8 |  | 15.87 | 6 | Water 3 | | c:483 | 3.02 | 1.0 | 29.6 |  | 10.23 | 7 | Water 4 | | a:4109 | 3.06 | 1.0 | 32.8 | 2.59 [A:OP1]a:2594 | 11.77 | 8 | Water 5 | | c:534 | 3.14 | 1.0 | 31.5 | 2.94 [THR:O]c:223 // 2.99 [K:K]c:306 | 11.27 | 9 | Water 6 | | a:8778 | 4.05 | 1.0 | 21.6 |  | 19.16 | 10 | **K** | K | c:306 | 4.41 | 1.0 | 30.0 |  | 33.16 | 11 | **K** | K | c:303 | 4.43 | 1.0 | 30.0 |  | 27.12 | 12 | THR | O | c:223 | 4.45 | 1.0 | 29.4 |  | 14.49 | 13 | A | OP2 | a:2594 | 4.51 | 1.0 | 30.0 |  | 13.18 | 14 | A | OP1 | a:2594 | 4.70 | 1.0 | 24.5 |  | 14.07 |

  
**1^st^ shell coordination angles and distances (*trans* in "blue"; *cys* in "green"; deviations in "orange"):**   
• 1-2(W/W): 98; 1-3(W/W): 76; 1-4(W/W): 89; 1-5(W/W): 144; 1-6(W/W): 67; 1-7(W/W): 82; 1-8(W/W): 137; 2-3(W/W): 75; 2-4(W/W): 154; 2-5(W/W): 105; 2-6(W/W): 132; 2-7(W/W): 78; 2-8(W/W): 77; 3-4(W/W): 131; 3-5(W/W): 83; 3-6(W/W): 57; 3-7(W/W): 143; 3-8(W/W): 139; 4-5(W/W): 83; 4-6(W/W): 73; 4-7(W/W): 78; 4-8(W/W): 81; 5-6(W/W): 77; 5-7(W/W): 129; 5-8(W/W): 76; 6-7(W/W): 137; 6-8(W/W): 145; 7-8(W/W): 55;   
• 1-2(W/W): 4.1; 1-3(W/W): 3.4; 1-4(W/W): 4.0; 1-5(W/W): 5.4; 1-6(W/W): 3.2; 1-7(W/W): 3.8; 1-8(W/W): 5.5; 2-3(W/W): 3.4; 2-4(W/W): 5.5; 2-5(W/W): 4.5; 2-6(W/W): 5.3; 2-7(W/W): 3.7; 2-8(W/W): 3.7; 3-4(W/W): 5.2; 3-5(W/W): 3.8; 3-6(W/W): 2.8; 3-7(W/W): 5.5; 3-8(W/W): 5.5; 4-5(W/W): 3.8; 4-6(W/W): 3.5; 4-7(W/W): 3.7; 4-8(W/W): 3.9; 5-6(W/W): 3.7; 5-7(W/W): 5.4; 5-8(W/W): 3.8; 6-7(W/W): 5.7; 6-8(W/W): 5.9; 7-8(W/W): 2.9;   
  
**3.89 ± 1.13: the average coodination distance of the 1^st^ six ligands could correspond to an hexahydrated Mg^2+^.**  
**16: Number of potential Water contacts**   
**5 : Number of second shell contacts involving water or ions**   
**31%: Percentage of satisfied second shell Water contacts**   
• Contact(s): /O_back_/O_back_/O_back_/O_w_/O_w_/O_w_/O_w_/O_w_/  
• Contact code: 3O_back_5O_w_  
Go to ToC

### K:c:306

|  | Res. type | Atom | Residue | Distance | Occupancy | B-factor | Closest atom(s) to a 1^st^ shell water | EM\_dens. |
| --- | --- | --- | --- | --- | --- | --- | --- | --- |
| **Ion** | | | |  | 1.0 | 30.0 |  | 33.16 |
 1 | GLY | O | c:234 | 2.58 | 1.0 | 30.8 |  | 15.87 | 2 | HIS | O | c:232 | 2.71 | 1.0 | 27.5 |  | 19.29 | 3 | ASN | O | c:239 | 2.77 | 1.0 | 32.9 |  | 15.44 | 4 | Water 1 | | a:5362 | 2.90 | 1.0 | 30.3 | 2.70 [A:OP1]a:2602 | 12.01 | 5 | Water 2 | | c:534 | 2.99 | 1.0 | 31.5 | 2.94 [THR:O]c:223 | 11.27 | 6 | THR | OG1 | c:223 | 3.11 | 1.0 | 25.3 |  | 12.99 | 7 | THR | O | c:223 | 3.47 | 1.0 | 29.4 |  | 14.49 | 8 | GLY | N | c:234 | 3.75 | 1.0 | 19.2 |  | 16.84 | 9 | GLY | O | c:233 | 4.23 | 1.0 | 24.8 |  | 18.30 | 10 | A | OP1 | a:2602 | 4.38 | 1.0 | 27.3 |  | 13.22 | 11 | **K** | K | c:305 | 4.41 | 1.0 | 30.0 |  | 29.54 | 12 | Water 3 | | c:475 | 4.53 | 1.0 | 27.0 | 2.89 [K:K]c:305 | 14.84 | 13 | Water 4 | | c:466 | 4.67 | 1.0 | 37.0 |  | 8.51 |

  
**1^st^ shell coordination angles and distances (*trans* in "blue"; *cys* in "green"; deviations in "orange"):**   
• 1-2(W/W): 114; 1-3(W/W): 111; 1-4(W/W): 88; 1-5(W/W): 84; 1-6(W/OG1): 143; 2-3(W/W): 125; 2-4(W/W): 74; 2-5(W/W): 142; 2-6(W/OG1): 89; 3-4(W/W): 78; 3-5(W/W): 70; 3-6(W/OG1): 72; 4-5(W/W): 142; 4-6(W/OG1): 128; 5-6(W/OG1): 61;   
• 1-2(W/W): 4.4; 1-3(W/W): 4.4; 1-4(W/W): 3.8; 1-5(W/W): 3.8; 1-6(W/OG1): 5.4; 2-3(W/W): 4.9; 2-4(W/W): 3.4; 2-5(W/W): 5.4; 2-6(W/OG1): 4.1; 3-4(W/W): 3.6; 3-5(W/W): 3.3; 3-6(W/OG1): 3.5; 4-5(W/W): 5.6; 4-6(W/OG1): 5.4; 5-6(W/OG1): 3.1;   
  
**4.05 ± 1.25: the average coodination distance of the 1^st^ six ligands could correspond to an hexahydrated Mg^2+^.**  
**12: Number of potential Water contacts**   
**2 : Number of second shell contacts involving water or ions**   
**17%: Percentage of satisfied second shell Water contacts**   
• Contact(s): /O_back_/O_back_/O_w_/O_w_/OG1_Thr_/  
• Contact code: 2O_back_OH_prot_2O_w_  
• Contact(s) in the exclusion zone: /O_back_/  
→ Coordination < 6 for K ←  
Go to ToC

### MG:d:301

|  | Res. type | Atom | Residue | Distance | Occupancy | B-factor | Closest atom(s) to a 1^st^ shell water | EM\_dens. |
| --- | --- | --- | --- | --- | --- | --- | --- | --- |
| **Ion** | | | |  | 1.0 | 26.6 |  | 23.54 |
 1 | C | OP1 | a:1660 | 1.88 | 1.0 | 29.0 |  | 11.54 | 2 | Water 1 | | a:6115 | 2.04 | 1.0 | 23.6 | 2.75 [G:OP2]a:1661 | 15.32 | 3 | Water 2 | | d:458 | 2.06 | 1.0 | 25.7 | 2.85 [HIS:NE2]d:140 | 12.13 | 4 | Water 3 | | a:3863 | 2.32 | 1.0 | 26.1 | 2.55 [U:OP1]a:746 // 2.88 [LEU:N]d:138 | 15.82 | 5 | Water 4 | | d:491 | 2.32 | 1.0 | 31.4 |  | 12.39 | 6 | HIS | NE2 | d:140 | 2.34 | 1.0 | 23.8 |  | 13.43 | 7 | C | OP2 | a:1660 | 3.80 | 1.0 | 23.5 |  | 14.51 | 8 | C | O5' | a:1660 | 3.90 | 1.0 | 24.1 |  | 14.25 | 9 | Water 5 | | a:7675 | 4.26 | 1.0 | 41.2 | 2.86 [C:OP2]a:1660 | 4.13 | 10 | Water 6 | | d:479 | 4.27 | 1.0 | 35.0 |  | 5.92 | 11 | G | OP2 | a:1661 | 4.32 | 1.0 | 27.6 |  | 13.05 | 12 | ASN | O | d:136 | 4.34 | 1.0 | 29.7 |  | 12.13 | 13 | A | O3' | a:745 | 4.35 | 1.0 | 26.2 |  | 12.90 | 14 | U | OP1 | a:746 | 4.35 | 1.0 | 29.6 |  | 13.08 | 15 | U | O3' | a:1659 | 4.36 | 1.0 | 30.0 |  | 12.96 | 16 | HIS | ND1 | d:140 | 4.44 | 1.0 | 24.9 |  | 12.25 | 17 | Water 7 | | d:444 | 4.55 | 1.0 | 33.0 | 2.75 [ASN:O]d:136 | 9.27 | 18 | Water 8 | | d:428 | 4.62 | 1.0 | 37.9 |  | 4.25 |

  
**1^st^ shell coordination angles and distances (*trans* in "blue"; *cys* in "green"; deviations in "orange"):**   
• 1-2(OP1/W): 94º; 1-3(OP1/W): *101º*; 2-3(W/W): 92º;   
• 1-2(OP1/W): 2.9Å; 1-3(OP1/W): 3.0Å; 2-3(W/W): 3.0Å;   
  
**2.16 ± 0.18: the average coodination distance of the 1^st^ six ligands is Mg^2+^ compatible**  
**8: Number of potential Water contacts**   
**4 : Number of second shell contacts involving water or ions**   
**50%: Percentage of satisfied second shell Water contacts**   
• Contact(s): /O_w_/O_w_/  
• Contact code: 2O_w_  
• Contact(s) in the exclusion zone: /NE2_His_/O_w_/O_w_/OP1/  
→ Missing octahedral coordination for MG ←  
Go to ToC

### K:d:302

|  | Res. type | Atom | Residue | Distance | Occupancy | B-factor | Closest atom(s) to a 1^st^ shell water | EM\_dens. |
| --- | --- | --- | --- | --- | --- | --- | --- | --- |
| **Ion** | | | |  | 1.0 | 30.0 |  | 10.35 |
 1 | ASN | OD1 | d:126 | 2.93 | 1.0 | 38.6 |  | 6.53 | 2 | LYS | O | d:123 | 3.04 | 1.0 | 25.1 |  | 14.28 | 3 | Water 1 | | d:422 | 3.05 | 1.0 | 48.6 |  | 5.43 | 4 | ARG | O | d:124 | 3.23 | 1.0 | 24.6 |  | 16.67 | 5 | C | OP2 | a:2682 | 3.65 | 1.0 | 30.5 |  | 10.39 | 6 | C | OP1 | a:2682 | 3.89 | 1.0 | 38.5 |  | 5.33 | 7 | ASN | ND2 | d:126 | 3.97 | 1.0 | 33.3 |  | 7.40 | 8 | TRP | N | d:125 | 4.34 | 1.0 | 17.7 |  | 16.75 |

  
**1^st^ shell coordination angles and distances (*trans* in "blue"; *cys* in "green"; deviations in "orange"):**   
• 1-2(OD1/W): 88; 1-3(OD1/W): 80; 2-3(W/W): 52;   
• 1-2(OD1/W): 4.2; 1-3(OD1/W): 3.9; 2-3(W/W): 2.7;   
  
**4.68 ± 1.47: the average coodination distance of the 1^st^ six ligands could correspond to an hexahydrated Mg^2+^.**  
**6: Number of potential Water contacts**   
**0 : Number of second shell contacts involving water or ions**   
**0%: Percentage of satisfied second shell Water contacts**   
• Contact(s): /O_back_/O_w_/OD1_Asn_/  
• Contact code: O_back_O_cno_O_w_  
→ Coordination < 6 for K ←  
Go to ToC

### K:e:301

|  | Res. type | Atom | Residue | Distance | Occupancy | B-factor | Closest atom(s) to a 1^st^ shell water | EM\_dens. |
| --- | --- | --- | --- | --- | --- | --- | --- | --- |
| **Ion** | | | |  | 1.0 | 30.0 |  | 10.62 |
 1 | VAL | O | e:167 | 2.52 | 1.0 | 50.8 |  | 3.77 | 2 | LEU | O | e:164 | 2.55 | 1.0 | 46.9 |  | 4.68 | 3 | ALA | O | e:161 | 2.69 | 1.0 | 51.8 |  | 5.31 | 4 | ARG | O | e:162 | 3.28 | 1.0 | 48.8 |  | 5.05 | 5 | LEU | N | e:164 | 4.25 | 1.0 | 40.5 |  | 5.76 | 6 | ARG | N | e:162 | 4.36 | 1.0 | 39.2 |  | 6.91 | 7 | HIS | N | e:165 | 4.46 | 1.0 | 42.5 |  | 4.89 | 8 | VAL | N | e:167 | 4.50 | 1.0 | 40.1 |  | 5.79 | 9 | Water 1 | | a:3772 | 4.57 | 1.0 | 44.2 | 2.85 [ARG:O]e:162 | 3.68 | 10 | ASP | OD1 | e:168 | 4.79 | 1.0 | 57.2 |  | 1.12 |

  
**1^st^ shell coordination angles and distances (*trans* in "blue"; *cys* in "green"; deviations in "orange"):**   
• 1-2(W/W): 77; 1-3(W/W): 97; 2-3(W/W): 78;   
• 1-2(W/W): 3.2; 1-3(W/W): 3.9; 2-3(W/W): 3.3;   
  
**4.77 ± 1.69: the average coodination distance of the 1^st^ six ligands could correspond to an hexahydrated Mg^2+^.**  
**8: Number of potential Water contacts**   
**0 : Number of second shell contacts involving water or ions**   
**0%: Percentage of satisfied second shell Water contacts**   
• Contact(s): /O_back_/  
• Contact code: O_back_  
• Contact(s) in the exclusion zone: /O_back_/O_back_/  
→ Coordination < 6 for K ←  
Go to ToC

### K:f:201

|  | Res. type | Atom | Residue | Distance | Occupancy | B-factor | Closest atom(s) to a 1^st^ shell water | EM\_dens. |
| --- | --- | --- | --- | --- | --- | --- | --- | --- |
| **Ion** | | | |  | 1.0 | 30.0 |  | 2.92 |
 1 | ARG | O | f:125 | 2.27 | 1.0 | 75.2 |  | 1.42 | 2 | THR | O | f:158 | 2.38 | 1.0 | 72.7 |  | 2.28 | 3 | ALA | O | f:160 | 2.44 | 1.0 | 76.5 |  | 0.96 | 4 | ASN | OD1 | f:127 | 3.23 | 1.0 | 72.8 |  | 3.10 | 5 | ALA | N | f:160 | 3.46 | 1.0 | 73.8 |  | 2.79 | 6 | GLY | O | f:126 | 3.96 | 1.0 | 73.2 |  | 2.48 | 7 | ASN | ND2 | f:127 | 4.19 | 1.0 | 70.1 |  | 3.02 | 8 | GLY | N | f:126 | 4.27 | 1.0 | 72.7 |  | 2.72 | 9 | THR | O | f:159 | 4.74 | 1.0 | 70.1 |  | 2.00 |

  
**1^st^ shell coordination angles and distances (*trans* in "blue"; *cys* in "green"; deviations in "orange"):**   
• 1-2(W/W): 146; 1-3(W/W): 92; 2-3(W/W): 107;   
• 1-2(W/W): 4.4; 1-3(W/W): 3.4; 2-3(W/W): 3.9;   
  
**4.37 ± 1.55: the average coodination distance of the 1^st^ six ligands could correspond to an hexahydrated Mg^2+^.**  
**8: Number of potential Water contacts**   
**0 : Number of second shell contacts involving water or ions**   
**0%: Percentage of satisfied second shell Water contacts**   
• Contact code: → No contacts ←   
• Contact(s) in the exclusion zone: /O_back_/O_back_/O_back_/  
→ Consider MG in place of K (1 valid contact(s) for MG vs. 0 for K ←)  
→2Fo-Fc density between 1.0 and 4.0 !   
Go to ToC

### MG:z:101

|  | Res. type | Atom | Residue | Distance | Occupancy | B-factor | Closest atom(s) to a 1^st^ shell water | EM\_dens. |
| --- | --- | --- | --- | --- | --- | --- | --- | --- |
| **Ion** | | | |  | 1.0 | 32.5 |  | 16.90 |
 1 | Water 1 | | z:226 | 1.97 | 1.0 | 30.1 | 2.79 [ARG:O]z:16 | 12.88 | 2 | ASP | OD1 | z:20 | 2.06 | 1.0 | 39.2 |  | 10.70 | 3 | Water 2 | | z:220 | 2.19 | 1.0 | 27.8 | 2.72 [ASP:OD2]z:20 // 2.84 [G:OP1]a:1268 | 13.91 | 4 | Water 3 | | a:4034 | 2.24 | 1.0 | 31.8 | 2.58 [C:OP1]a:2051 | 12.20 | 5 | Water 4 | | z:240 | 2.26 | 1.0 | 36.0 |  | 9.91 | 6 | Water 5 | | z:239 | 2.39 | 1.0 | 30.6 |  | 9.54 | 7 | ASP | OD2 | z:20 | 3.27 | 1.0 | 37.6 |  | 9.32 | 8 | ARG | O | z:16 | 4.10 | 1.0 | 30.8 |  | 13.50 | 9 | Water 6 | | z:249 | 4.21 | 1.0 | 41.6 |  | 4.17 | 10 | Water 7 | | a:8938 | 4.35 | 1.0 | 42.3 |  | 4.72 | 11 | C | OP1 | a:2051 | 4.38 | 1.0 | 29.5 |  | 13.06 | 12 | Water 8 | | a:10266 | 4.50 | 1.0 | 40.0 |  | 7.69 | 13 | Water 9 | | z:214 | 4.51 | 1.0 | 46.3 |  | 4.19 | 14 | Water 10 | | a:5837 | 4.61 | 1.0 | 32.2 | 2.73 [G:OP1]a:1268 | 11.75 | 15 | G | O3' | a:2050 | 4.71 | 1.0 | 25.9 |  | 15.43 | 16 | G | O2' | a:2050 | 4.76 | 1.0 | 28.7 |  | 13.92 | 17 | G | OP1 | a:1268 | 4.78 | 1.0 | 28.7 |  | 13.45 |

  
**1^st^ shell coordination angles and distances (*trans* in "blue"; *cys* in "green"; deviations in "orange"):**   
• 1-2(W/OD1): *99º*; 1-3(W/W): *99º*; 1-4(W/W): 88º; 1-5(W/W): 90º; 2-3(OD1/W): 94º; 2-4(OD1/W): 173º; 2-5(OD1/W): 91º; 3-4(W/W): 88º; 3-5(W/W): *168º*; 4-5(W/W): 85º;   
• 1-2(W/OD1): 3.1Å; 1-3(W/W): 3.2Å; 1-4(W/W): 2.9Å; 1-5(W/W): 3.0Å; 2-3(OD1/W): 3.1Å; 2-4(OD1/W): *4.3Å*; 2-5(OD1/W): 3.1Å; 3-4(W/W): 3.1Å; 3-5(W/W): *4.4Å*; 4-5(W/W): 3.1Å;   
  
**2.18 ± 0.14: the average coodination distance of the 1^st^ six ligands is Mg^2+^ compatible**  
**10: Number of potential Water contacts**   
**4 : Number of second shell contacts involving water or ions**   
**40%: Percentage of satisfied second shell Water contacts**   
• Contact(s): /O_w_/O_w_/O_w_/O_w_/OD1_Asp_/  
• Contact code: O_coo_4O_w_  
• Contact(s) in the exclusion zone: /O_w_/OD2_Asp_/  
→ Missing octahedral coordination for MG ←  
→ Slightly distorded coord. octahedron (5.1º) ←  
\_\_\_\_\_\_\_\_\_\_\_\_\_\_\_\_\_\_\_\_\_\_\_\_\_\_\_\_\_\_\_\_\_\_\_\_\_\_\_\_\_\_\_\_\_\_\_\_\_\_\_\_\_\_\_\_\_\_\_\_\_\_\_\_  

#### 5) Sorted coded contact list

##### Go to ToC

NOTES:

  
- N. → coordination number estimation deduced from the PDB structure;  
- Excl. → existence of contacts in the ion exclusion zone;  
- Bad hyd. → 2^nd^ shell hydration contacts shorter than 2.4 Å.;  
- Lines in red mark stereochemical issues that should be addressed;  
- Lines in green mark ions without stereochemical issues --- Please note that this does not always warrant the ion identification;  
  
(1) N=0 → No 1^st^ shell coordination: difficult to assess ion identity;  
(2) Exclusion contacts → Probable wrong identity assignment --- poor modelization of the ion binding site is also possible;  
(3) Bad 2^nd^ hydration shell contacts → Poor modelization of the hydration shell and/or binding site;  
(4) Check for anion (Cl^-^, ...);  
(5) Mixed contacts with unprotonated O/N and protonated N, suggest poorly modeled binding site;  
(6) Currently the ZN (Zinc), CD (Cadmium) and CO (Cobalt) coordination is not evaluated;  
(7) Check occupancies --- partial occupancies are not reliable at medium to low resolutions, especially for solvent particles;  
(8) Check B-factors;  
  
  

|  |  |  |  |  |  |  |  |
| --- | --- | --- | --- | --- | --- | --- | --- |
| **Ion** | **Count** | **Code** | **N.** | **Excl.** | **Bad hyd.** | **Notes** | **Ion list** |
| MG: | 42 | No 1^st^ shell | >1 | Yes | No | (2) | A:1609 A:1621 A:1625 A:1636 A:1637 A:1640 A:1641 A:1643 A:1682 A:1695 A:1714 A:1719 A:1721 A:1744 a:3005 a:3018 a:3029 a:3031 a:3032 a:3042 a:3044 a:3045 a:3051 a:3052 a:3053 a:3054 a:3055 a:3068 a:3069 a:3071 a:3148 a:3162 a:3169 a:3173 a:3203 a:3330 a:3354 a:3355 a:3358 a:3365 a:3366 b:203 || MG: | 27 | O_ph_5O_w_ | 6 | No | No |  | A:1606 A:1673 a:3036 a:3047 a:3048 a:3078 a:3080 a:3082 a:3086 a:3090 a:3147 a:3154 a:3156 a:3161 a:3165 a:3185 a:3188 a:3193 a:3199 a:3232 a:3234 a:3240 a:3289 a:3303 a:3319 a:3348 a:3357 || MG: | 24 | O_ph_4O_w_ | >5 | Yes | No | (2) | A:1608 A:1669 A:1675 A:1742 a:3007 a:3025 a:3164 a:3183 a:3200 a:3206 a:3226 a:3227 a:3228 a:3239 a:3245 a:3288 a:3293 a:3296 a:3299 a:3300 a:3305 a:3320 a:3335 a:3337 || MG: | 19 | 3O_w_ | >3 | Yes | No | (2) | A:1610 A:1613 A:1639 A:1668 A:1693 A:1703 A:1741 X:101 a:3014 a:3022 a:3075 a:3094 a:3160 a:3163 a:3202 a:3291 a:3318 a:3352 a:3364 || MG: | 18 | 4O_w_ | >4 | Yes | No | (2) | A:1614 A:1631 A:1676 A:1678 a:3013 a:3024 a:3038 a:3065 a:3076 a:3077 a:3092 a:3146 a:3171 a:3214 a:3215 a:3359 a:3360 a:3361 || MG: | 18 | O_w_ | >1 | Yes | No | (2) | A:1617 A:1618 A:1635 A:1697 A:1705 A:1707 A:1709 a:3002 a:3011 a:3012 a:3015 a:3017 a:3043 a:3066 a:3070 a:3073 a:3074 a:3168 || MG: | 18 | 6O_w_ | 6 | No | No |  | A:1670 A:1684 A:1745 a:3016 a:3019 a:3030 a:3067 a:3095 a:3182 a:3205 a:3210 a:3212 a:3216 a:3221 a:3229 a:3286 a:3297 a:3311 || MG: | 18 | O_ph_3O_w_ | >4 | Yes | No | (2) | A:1607 A:1686 A:1716 a:3003 a:3009 a:3079 a:3149 a:3155 a:3166 a:3187 a:3285 a:3290 a:3306 a:3315 a:3323 a:3334 a:3349 b:204 || MG: | 13 | 2O_w_ | >2 | Yes | No | (2) | A:1611 A:1622 A:1630 A:1690 A:1692 a:3006 a:3033 a:3039 a:3041 a:3158 a:3189 a:3362 d:301 || MG: | 11 | 5O_w_ | >5 | Yes | No | (2) | A:1629 A:1632 a:3021 a:3034 a:3035 a:3157 a:3213 a:3217 a:3235 a:3241 a:3301 || MG: | 11 | O_ph_2O_w_ | >3 | Yes | No | (2) | A:1691 a:3004 a:3049 a:3081 a:3091 a:3195 a:3292 a:3307 a:3310 a:3313 a:3356 || MG: | 10 | 2O_ph_3O_w_ | >5 | Yes | No | (2) | A:1672 A:1700 a:3223 a:3225 a:3231 a:3236 a:3243 a:3294 a:3309 a:3336 || MG: | 10 | 2O_ph_4O_w_ | 6 | No | No |  | a:3040 a:3084 a:3087 a:3093 a:3143 a:3150 a:3194 a:3207 a:3237 a:3322 || MG: | 9 | 2O_ph_2O_w_ | >4 | Yes | No | (2) | A:1619 A:1683 a:3046 a:3152 a:3159 a:3167 a:3295 a:3304 a:3316 || MG: | 8 | 3O_ph_3O_w_ | 6 | No | No |  | A:1634 A:1681 a:3089 a:3233 a:3244 a:3314 a:3326 a:3331 || MG: | 6 | No 1^st^ shell | =0 | No | No | (1) | A:1696 A:1711 A:1715 A:1718 a:3026 b:201 || MG: | 5 | 3O_w_ | 3 | No | No |  | A:1616 A:1688 a:3028 a:3037 a:3190 || MG: | 5 | O_ph_4O_w_ | 5 | No | No |  | A:1620 A:1628 A:1680 A:1713 a:3353 || MG: | 5 | 2O_ph_O_w_ | >3 | Yes | No | (2) | A:1702 a:3192 a:3230 a:3287 a:3363 || MG: | 5 | 5O_w_ | 5 | No | No |  | a:3001 a:3085 a:3197 a:3218 a:3350 || MG: | 4 | O_b_4O_w_ | >5 | Yes | No | (2) | a:3222 a:3317 a:3321 a:3332 || MG: | 4 | 3O_ph_2O_w_ | >5 | Yes | No | (2) | a:3224 a:3324 a:3327 a:3328 || MG: | 3 | O_ph_3O_w_ | 4 | No | No |  | A:1677 a:3204 a:3209 || MG: | 3 | 2O_ph_3O_w_ | 5 | No | No |  | A:1679 a:3308 a:3351 || MG: | 3 | O_ph_O_w_ | >2 | Yes | No | (2) | A:1699 A:1706 a:3151 || MG: | 3 | O_b_2O_w_ | >3 | Yes | No | (2) | A:1708 a:3010 a:3302 || MG: | 3 | O_b_5O_w_ | 6 | No | No |  | a:3008 a:3153 b:205 || MG: | 2 | O_b_3O_w_ | >4 | Yes | No | (2) | A:1633 a:3050 || MG: | 2 | O_ph_5O_w_ | >6 | Yes | No | (2) | A:1638 a:3174 || MG: | 2 | O_b_O_w_ | >2 | Yes | No | (2) | A:1710 a:3208 || MG: | 2 | O_w_ | 1 | No | No |  | A:1712 a:3191 || MG: | 2 | 2O_ph_ | >2 | Yes | --- | (2) | A:1717 a:3023 || MG: | 2 | O_ph_O_b_3O_w_ | >5 | Yes | No | (2) | a:3083 a:3312 || MG: | 2 | 4O_ph_2O_w_ | 6 | No | No |  | a:3238 a:3329 || MG: | 2 | 2O_ph_ | 2 | No | --- |  | A:1626 b:202 || MG: | 2 | 4O_w_ | 4 | No | No |  | A:1687 c:302 || MG: | 1 | N_b_2O_w_ | >3 | Yes | No | (2) | A:1612 || MG: | 1 | N_b_3O_w_ | 4 | No | No |  | A:1615 || MG: | 1 | O_ph_O_b_ | >2 | Yes | --- | (2) | A:1623 || MG: | 1 | O_ph_ | >1 | Yes | --- | (2) | A:1624 || MG: | 1 | O_ph_O_b_4O_w_ | 6 | No | No |  | A:1627 || MG: | 1 | 2O_ph_4O_w_ | 6 | No | Yes | (3) | A:1642 || MG: | 1 | 3O_ph_3O_w_ | >6 | Yes | No | (2) | A:1671 || MG: | 1 | 2O_w_ | >2 | Yes | Yes | (2,3) | A:1685 || MG: | 1 | 2O_w_ | 2 | No | No |  | A:1694 || MG: | 1 | 3O_w_ | >3 | Yes | Yes | (2,3) | A:1698 || MG: | 1 | 3O_ph_O_w_ | >4 | Yes | No | (2) | A:1701 || MG: | 1 | O_ph_ | >1 | Yes | --- | (2) | A:1704 || MG: | 1 | O_ph_ | 1 | No | --- |  | A:1720 || MG: | 1 | O_ph_O_b_4O_w_ | 6 | No | Yes | (3) | A:1743 || MG: | 1 | O_ph_5O_w_ | >6 | Yes | Yes | (2,3) | a:3020 || MG: | 1 | O_ph_O_b_2O_w_ | 4 | No | No |  | a:3027 || MG: | 1 | O_ph_2O_w_ | 3 | No | No |  | a:3072 || MG: | 1 | 2O_ph_N_b_3O_w_ | 6 | No | No |  | a:3088 || MG: | 1 | 2O_ph_O_r_O_w_ | >4 | Yes | --- | (2) | a:3144 || MG: | 1 | 2O_ph_O_r_2O_w_ | >5 | Yes | No | (2) | a:3145 || MG: | 1 | O_ph_N_b_3O_w_ | 5 | No | No |  | a:3170 || MG: | 1 | N_b_3O_w_ | >4 | Yes | No | (2) | a:3172 || MG: | 1 | 5O_w_ | >5 | Yes | Yes | (2,3) | a:3184 || MG: | 1 | 2O_ph_O_back_2O_w_N_His_ | 6 | No | No |  | a:3186 || MG: | 1 | N_b_ | >1 | Yes | --- | (2) | a:3196 || MG: | 1 | O_r_5O_w_ | 6 | No | No |  | a:3198 || MG: | 1 | 2O_ph_O_b_3O_w_ | 6 | No | No |  | a:3201 || MG: | 1 | O_b_4O_w_ | 5 | No | No |  | a:3211 || MG: | 1 | 2O_ph_O_cno_3O_w_ | 6 | No | No |  | a:3219 || MG: | 1 | N_b_4O_w_ | >5 | Yes | No | (2) | a:3220 || MG: | 1 | 2N_b_3O_w_ | >5 | Yes | No | (2) | a:3242 || MG: | 1 | O_ph_O_b_2O_w_ | >4 | Yes | No | (2) | a:3298 || MG: | 1 | 3O_ph_O_b_O_w_ | >5 | Yes | No | (2) | a:3333 || MG: | 1 | O_ph_4O_w_N_His_ | 6 | No | No |  | c:304 || K: | 8 | 3O_b_2O_w_ | 5 | No | No |  | A:1731 A:1740 a:3061 a:3099 a:3139 a:3251 a:3257 a:3338 || K: | 7 | 3O_b_O_w_ | 4 | No | --- |  | A:1653 A:1667 A:1734 a:3060 a:3258 a:3260 a:3340 || K: | 5 | 2O_b_O_w_ | 3 | No | No |  | A:1651 a:3057 a:3266 a:3280 a:3343 || K: | 5 | No 1^st^ shell | >1 | Yes | No | (2) | A:1603 A:1647 A:1737 a:3131 f:201 || K: | 4 | O_r_O_b_N_b_O_w_ | 4 | No | --- |  | A:1604 A:1648 A:1725 a:3344 || K: | 3 | 4O_b_ | 4 | No | --- |  | A:1645 A:1656 A:1739 || K: | 3 | 4O_b_O_w_ | 5 | No | No |  | A:1650 a:3124 a:3125 || K: | 3 | 2O_ph_2O_r_O_w_ | 5 | No | No |  | A:1729 a:3117 a:3249 || K: | 3 | 2O_ph_O_r_O_w_ | 4 | No | --- |  | A:1736 a:3109 a:3346 || K: | 3 | 3O_b_2O_w_ | >5 | Yes | No | (2) | a:3102 a:3129 a:3142 || K: | 3 | 3O_b_ | 3 | No | --- |  | a:3259 a:3262 a:3271 || K: | 2 | 2O_b_ | >2 | Yes | --- | (2) | A:1605 a:3175 || K: | 2 | 3O_b_3O_w_ | 6 | No | Yes | (3) | A:1663 a:3180 || K: | 2 | 3O_ph_O_r_O_w_ | 5 | No | No |  | A:1730 a:3063 || K: | 2 | 2O_b_ | 2 | No | --- |  | a:3108 a:3339 || K: | 2 | 3O_b_3O_w_ | 6 | No | No |  | a:3130 a:3282 || K: | 1 | O_ph_2O_r_ | 3 | No | --- |  | A:1601 || K: | 1 | O_ph_2O_w_ | >3 | Yes | No | (2) | A:1602 || K: | 1 | 2O_w_ | >2 | Yes | No | (2) | A:1644 || K: | 1 | O_ph_O_r_O_b_O_w_ | 4 | No | --- |  | A:1649 || K: | 1 | O_ph_N_b_2O_w_ | 4 | No | No |  | A:1654 || K: | 1 | 3O_b_O_w_ | >4 | Yes | --- |  | A:1655 || K: | 1 | O_ph_2O_b_N_b_O_w_ | 5 | No | No |  | A:1657 || K: | 1 | O_ph_O_r_N_b_2O_w_ | 5 | No | No |  | A:1658 || K: | 1 | O_ph_O_b_2O_w_ | >4 | Yes | No | (2) | A:1659 || K: | 1 | No 1^st^ shell | =0 | No | No | (1) | A:1660 || K: | 1 | O_ph_2O_b_ | 3 | No | --- |  | A:1661 || K: | 1 | O_ph_O_r_O_b_O_w_ | >4 | Yes | --- | (2) | A:1662 || K: | 1 | O_b_ | >1 | Yes | --- | (2) | A:1664 || K: | 1 | 3O_ph_O_r_3O_w_ | 7 | No | No |  | A:1724 || K: | 1 | O_r_3O_b_ | >4 | Yes | --- |  | A:1728 || K: | 1 | 2O_b_3O_w_ | 5 | No | No |  | A:1732 || K: | 1 | 4O_r_O_b_N_b_O_w_ | 7 | No | No |  | A:1733 || K: | 1 | 2O_ph_2O_r_2O_w_ | >6 | Yes | No | (2) | A:1735 || K: | 1 | 3O_b_3O_w_ | >6 | Yes | No | (2) | A:1738 || K: | 1 | O_r_3O_b_2O_w_ | 6 | No | No |  | a:3056 || K: | 1 | 2O_ph_3O_r_2O_w_ | 7 | No | No |  | a:3059 || K: | 1 | O_ph_O_b_3O_w_ | >5 | Yes | No | (2) | a:3064 || K: | 1 | O_r_O_b_ | 2 | No | --- |  | a:3097 || K: | 1 | O_ph_O_r_N_b_ | 3 | No | No |  | a:3101 || K: | 1 | O_w_ | >1 | Yes | No | (2) | a:3103 || K: | 1 | O_r_2O_b_N_b_ | 5 | No | No |  | a:3105 || K: | 1 | O_b_5O_w_ | 6 | No | No |  | a:3106 || K: | 1 | 5O_r_O_w_ | 6 | No | No |  | a:3112 || K: | 1 | O_r_O_b_2N_b_O_w_ | 5 | No | No |  | a:3114 || K: | 1 | 3O_b_ | >3 | Yes | --- | (2) | a:3115 || K: | 1 | 3O_ph_O_r_4O_w_ | 8 | No | No |  | a:3119 || K: | 1 | 3O_ph_2O_r_O_w_ | 6 | No | No |  | a:3120 || K: | 1 | O_r_2O_b_3O_w_ | 6 | No | No |  | a:3127 || K: | 1 | O_ph_2O_r_O_b_2N_b_O_w_ | 7 | No | No |  | a:3133 || K: | 1 | 4O_b_2O_w_ | 6 | No | No |  | a:3135 || K: | 1 | 3O_b_O_w_ | 4 | No | --- |  | a:3136 || K: | 1 | 2O_b_N_b_3O_w_ | 6 | No | No |  | a:3137 || K: | 1 | O_ph_2O_b_ | >3 | Yes | --- | (2) | a:3138 || K: | 1 | O_ph_2O_r_3O_w_ | 6 | No | No |  | a:3140 || K: | 1 | 3O_b_N_b_2O_w_ | 6 | No | No |  | a:3177 || K: | 1 | 2O_r_2O_b_N_b_2O_w_ | 7 | No | No |  | a:3179 || K: | 1 | O_b_2O_w_ | >3 | Yes | No | (2) | a:3246 || K: | 1 | 2O_ph_O_w_ | >3 | Yes | No | (2) | a:3248 || K: | 1 | 3O_b_4O_w_ | 7 | No | No |  | a:3252 || K: | 1 | O_ph_N_b_3O_w_ | >5 | Yes | No | (2) | a:3253 || K: | 1 | O_ph_O_b_2O_w_ | 4 | No | No |  | a:3255 || K: | 1 | 2O_b_N_b_3O_w_ | 6 | No | No |  | a:3261 || K: | 1 | 4O_b_3O_w_ | 7 | No | No |  | a:3263 || K: | 1 | O_r_3O_b_3N_b_O_w_ | 8 | No | No |  | a:3264 || K: | 1 | 3O_ph_O_r_2O_w_ | 6 | No | No |  | a:3265 || K: | 1 | O_ph_2O_b_O_w_ | 4 | No | --- |  | a:3267 || K: | 1 | O_b_ | 1 | No | --- |  | a:3268 || K: | 1 | O_ph_3O_r_N_b_2O_w_ | 7 | No | No |  | a:3272 || K: | 1 | O_r_3O_b_O_w_ | 5 | No | No |  | a:3273 || K: | 1 | 2O_ph_O_r_3O_w_ | 6 | No | No |  | a:3274 || K: | 1 | O_ph_O_b_3O_w_ | 5 | No | No |  | a:3277 || K: | 1 | 2O_r_O_b_4O_w_ | 7 | No | No |  | a:3278 || K: | 1 | O_ph_O_r_2O_w_ | >4 | Yes | No | (2) | a:3279 || K: | 1 | O_b_2O_w_ | 3 | No | No |  | a:3281 || K: | 1 | O_ph_O_b_N_b_O_w_ | 4 | No | --- |  | a:3341 || K: | 1 | O_r_4O_w_ | 5 | No | No |  | a:3345 || K: | 1 | O_ph_O_b_N_b_2O_w_ | 5 | No | No |  | a:3347 || K: | 1 | O_ph_N_b_ | >2 | Yes | --- | (2) | b:206 || K: | 1 | O_ph_O_back_O_coo_4O_w_ | 7 | No | No |  | c:303 || K: | 1 | 3O_back_5O_w_ | 8 | No | No |  | c:305 || K: | 1 | O_back_ | >1 | Yes | --- | (2) | e:301 || ZN: | 1 | N_His_3SG_Cys_ | 4 | No | No |  | 3:101 ||  |  |  |  |  |  |  | List of ions with partial occupancies ||  |  |  |  |  |  | Total: | 0 |

  
\_\_\_\_\_\_\_\_\_\_\_\_\_\_\_\_\_\_\_\_\_\_\_\_\_\_\_\_\_\_\_\_\_\_\_\_\_\_\_\_\_\_\_\_\_\_\_\_\_\_\_\_\_\_\_\_\_\_\_\_\_\_\_\_  

#### 6) B-factor and 2FoFc, FoFc density map summary

##### Go to ToC

 **Notes:**  
**• For the B-factor table, all listed atoms were considered;**  
**• For the 2FoFc table, atoms with 1.0 < B-factors > 80.0 were discarded**  
  

| B-factors | Atom | Count | Min | Max | Avg |
| --- | --- | --- | --- | --- | --- |
| 8B0X\_b | P | 4448 | 18.5 | 184.3 | 57.8 (40.7) |
| 8B0X\_b | OP1 | 4446 | 19.3 | 182.7 | 55.1 (37.5) |
| 8B0X\_b | OP2 | 4446 | 18.3 | 182.8 | 54.0 (38.2) |
| 8B0X\_b | O2' | 4448 | 19.2 | 182.3 | 52.9 (38.4) |
| 8B0X\_b | O3' | 4448 | 18.6 | 183.6 | 54.3 (38.6) |
| 8B0X\_b | O4' | 4448 | 18.6 | 183.2 | 54.1 (38.2) |
| 8B0X\_b | O5' | 4448 | 18.2 | 183.4 | 54.3 (38.6) |
| 8B0X\_b | N7 | 2558 | 13.6 | 182.3 | 46.8 (41.0) |
| 8B0X\_b | SG | 27 | 36.2 | 100.8 | 65.3 (27.1) |
| 8B0X\_b | SD | 113 | 24.7 | 120.3 | 70.2 (28.2) |
| 8B0X\_b | HOH | 11461 | 17.4 | 97.2 | 43.3 (14.1) |
| 8B0X\_b | MG | 361 | 20.9 | 85.6 | 44.9 (21.4) |
| 8B0X\_b | K | 168 | 30.0 | 30.0 | 30.0 (0.0) |
| 8B0X\_b | ZN | 1 | 81.2 | 81.2 | 81.2 (0.0) |

  

| 2FoFc | Atom | Count | Min | Max | Avg |
| --- | --- | --- | --- | --- | --- |
| 8B0X\_2FoFc | P | 3638 | 1.9 | 36.1 | 13.5 (7.4) |
| 8B0X\_2FoFc | OP1 | 3805 | 0.4 | 19.5 | 5.5 (3.7) |
| 8B0X\_2FoFc | OP2 | 3814 | 0.1 | 19.4 | 6.3 (3.8) |
| 8B0X\_2FoFc | O2' | 3828 | -0.0 | 21.8 | 7.0 (4.4) |
| 8B0X\_2FoFc | O3' | 3801 | 0.7 | 19.1 | 6.8 (4.0) |
| 8B0X\_2FoFc | O4' | 3815 | 0.8 | 20.6 | 7.5 (4.4) |
| 8B0X\_2FoFc | O5' | 3797 | 0.2 | 19.7 | 6.8 (3.8) |
| 8B0X\_2FoFc | N7 | 2240 | 0.6 | 21.4 | 8.7 (4.4) |
| 8B0X\_2FoFc | SG | 22 | 2.4 | 19.2 | 8.9 (5.5) |
| 8B0X\_2FoFc | SD | 78 | 1.4 | 31.7 | 8.5 (7.6) |
| 8B0X\_2FoFc | HOH | 11433 | -1.2 | 20.0 | 5.6 (3.4) |
| 8B0X\_2FoFc | MG | 355 | 1.1 | 30.1 | 12.3 (7.6) |
| 8B0X\_2FoFc | K | 168 | 1.9 | 39.6 | 15.3 (9.3) |

  
\_\_\_\_\_\_\_\_\_\_\_\_\_\_\_\_\_\_\_\_\_\_\_\_\_\_\_\_\_\_\_\_\_\_\_\_\_\_\_\_\_\_\_\_\_\_\_\_\_\_\_\_\_\_\_\_\_\_\_\_\_\_\_\_  

#### 7) Potential and detected Mg(2+) coodination clamps list

##### Go to ToC

NOTES:

  
- B-factors are in brackets.  
- B-factors > 60 Å^2^ are in red and underlined.  
- A limit of 3.4 Å is set for oxygen to oxygen/nitrogen distances.  
- A limit of 2.4 Å is set for Mg^2+^ to oxygen/nitrogen distances.  
  
**Potential number of OP1...OP1 clamps (with d < 3.4 Å):** **37**   
 1. -- dist(C:OP1:a:31:[29.7], G:OP1:a:1241:[41.4]) = 3.09 **← CHECK**  
2. -- dist(C:OP1:a:192:[26.0], U:OP1:a:2247:[23.5]) = 2.99 -- dist(a:C:192:OP1, a:MG:3336:MG:[26.0]) = 2.08 -- -- dist(a:U:2247:OP1, a:MG:3336:MG:[26.0]) = 1.96 --   
3. -- dist(G:OP1:a:375:[32.0], C:OP1:a:393:[36.2]) = 3.11 -- dist(a:G:375:OP1, a:MG:3143:MG:[35.3]) = 2.09 -- -- dist(a:C:393:OP1, a:MG:3143:MG:[35.3]) = 1.99 --   
4. -- dist(G:OP1:a:411:[34.6], A:OP1:a:2411:[28.5]) = 3.10 -- dist(a:G:411:OP1, a:MG:3309:MG:[32.2]) = 1.98 -- -- dist(a:A:2411:OP1, a:MG:3309:MG:[32.2]) = 2.08 --   
5. -- dist(A:OP1:a:526:[32.3], C:OP1:a:527:[34.5]) = 2.83 -- dist(a:A:526:OP1, a:MG:3084:MG:[33.6]) = 2.03 -- -- dist(a:C:527:OP1, a:MG:3084:MG:[33.6]) = 2.02 --   
6. -- dist(U:OP1:a:578:[24.7], 2MA:OP1:a:2507:[26.5]) = 2.80 -- dist(a:U:578:OP1, a:MG:3322:MG:[25.5]) = 2.08 -- -- dist(a:2MA:2507:OP1, a:MG:3322:MG:[25.5]) = 2.08 --   
7. -- dist(C:OP1:a:680:[25.6], C:OP1:a:2076:[23.0]) = 3.24 -- dist(a:C:680:OP1, a:MG:3201:MG:[24.5]) = 2.15 -- -- dist(a:C:2076:OP1, a:MG:3201:MG:[24.5]) = 2.04 --   
8. -- dist(G:OP1:a:786:[22.9], G:OP1:a:2592:[22.1]) = 3.06 -- dist(a:G:786:OP1, a:MG:3225:MG:[26.7]) = 2.00 -- -- dist(a:G:2592:OP1, a:MG:3225:MG:[26.7]) = 2.01 --   
9. -- dist(U:OP1:a:828:[22.6], G:OP1:a:2432:[26.9]) = 3.21 -- dist(a:U:828:OP1, a:MG:3329:MG:[25.3]) = 2.03 -- -- dist(a:G:2432:OP1, a:MG:3329:MG:[25.3]) = 2.10 --   
10. -- dist(U:OP1:a:828:[22.6], G:OP1:a:2433:[25.2]) = 3.09 -- dist(a:U:828:OP1, a:MG:3329:MG:[25.3]) = 2.03 -- -- dist(a:G:2433:OP1, a:MG:3329:MG:[25.3]) = 2.02 --   
11. -- dist(C:OP1:a:950:[26.3], G:OP1:a:964:[29.2]) = 2.88 -- dist(a:C:950:OP1, a:MG:3233:MG:[26.0]) = 2.00 -- -- dist(a:G:964:OP1, a:MG:3233:MG:[26.0]) = 2.12 --   
12. -- dist(U:OP1:a:1251:[27.0], G:OP1:a:1252:[25.3]) = 3.03 -- dist(a:U:1251:OP1, a:MG:3167:MG:[29.0]) = 2.07 -- -- dist(a:G:1252:OP1, a:MG:3167:MG:[29.0]) = 2.02 --   
13. -- dist(A:OP1:a:1270:[24.5], C:OP1:a:2010:[23.6]) = 3.09 -- dist(a:A:1270:OP1, a:MG:3236:MG:[25.8]) = 2.25 -- -- dist(a:C:2010:OP1, a:MG:3236:MG:[25.8]) = 1.94 --   
14. -- dist(A:OP1:a:1274:[28.1], U:OP1:a:1649:[30.0]) = 3.02 -- dist(a:A:1274:OP1, a:MG:3238:MG:[27.5]) = 2.03 -- -- dist(a:U:1649:OP1, a:MG:3238:MG:[27.5]) = 2.07 --   
15. -- dist(C:OP1:a:1316:[26.7], G:OP1:a:1334:[33.5]) = 3.05 -- dist(a:C:1316:OP1, a:MG:3089:MG:[28.9]) = 2.01 -- -- dist(a:G:1334:OP1, a:MG:3089:MG:[28.9]) = 2.04 --   
16. -- dist(U:OP1:a:1396:[33.2], C:OP1:a:1606:[33.3]) = 3.02 -- dist(a:U:1396:OP1, a:MG:3243:MG:[33.9]) = 1.97 -- -- dist(a:C:1606:OP1, a:MG:3243:MG:[33.9]) = 2.05 --   
17. -- dist(A:OP1:a:1397:[33.2], A:OP1:a:1605:[31.3]) = 3.01 -- dist(a:A:1397:OP1, a:MG:3244:MG:[30.5]) = 2.04 -- -- dist(a:A:1605:OP1, a:MG:3244:MG:[30.5]) = 1.98 --   
18. -- dist(U:OP1:a:1649:[30.0], U:OP1:a:1650:[29.7]) = 3.00 -- dist(a:U:1649:OP1, a:MG:3238:MG:[27.5]) = 2.07 -- -- dist(a:U:1650:OP1, a:MG:3238:MG:[27.5]) = 2.09 --   
19. -- dist(A:OP1:a:1782:[24.7], U:OP1:a:1784:[27.2]) = 2.83 -- dist(a:A:1782:OP1, a:MG:3326:MG:[23.7]) = 2.05 -- -- dist(a:U:1784:OP1, a:MG:3326:MG:[23.7]) = 1.93 --   
20. -- dist(G:OP1:a:1972:[27.0], G:OP1:a:2596:[26.6]) = 2.83 -- dist(a:G:1972:OP1, a:MG:3152:MG:[33.4]) = 2.16 -- -- dist(a:G:2596:OP1, a:MG:3152:MG:[33.4]) = 2.01 --   
21. -- dist(A:OP1:a:1974:[34.3], U:OP1:a:1975:[36.4]) = 3.34 -- dist(a:A:1974:OP1, a:MG:3040:MG:[49.7]) = 2.07 -- -- dist(a:U:1975:OP1, a:MG:3040:MG:[49.7]) = 2.12 --   
22. -- dist(A:OP1:a:2272:[37.2], G:OP1:a:2273:[30.6]) = 3.01 -- dist(a:A:2272:OP1, a:MG:3299:MG:[30.3]) = 1.88 -- -- dist(a:G:2273:OP1, a:MG:3299:MG:[30.3]) = 2.22 --   
23. -- dist(G:OP1:a:2293:[38.1], C:OP1:a:2389:[41.1]) = 3.20 -- dist(a:G:2293:OP1, a:MG:3187:MG:[45.5]) = 2.16 -- -- dist(a:C:2389:OP1, a:MG:3187:MG:[45.5]) = 1.89 --   
24. -- dist(C:OP1:a:2431:[29.3], G:OP1:a:2432:[26.9]) = 2.78 -- dist(a:C:2431:OP1, a:MG:3329:MG:[25.3]) = 2.12 -- -- dist(a:G:2432:OP1, a:MG:3329:MG:[25.3]) = 2.10 --   
25. -- dist(C:OP1:a:2431:[29.3], G:OP1:a:2433:[25.2]) = 2.92 -- dist(a:C:2431:OP1, a:MG:3329:MG:[25.3]) = 2.12 -- -- dist(a:G:2433:OP1, a:MG:3329:MG:[25.3]) = 2.02 --   
26. -- dist(G:OP1:a:2432:[26.9], G:OP1:a:2433:[25.2]) = 2.97 -- dist(a:G:2432:OP1, a:MG:3329:MG:[25.3]) = 2.10 -- -- dist(a:G:2433:OP1, a:MG:3329:MG:[25.3]) = 2.02 --   
27. -- dist(A:OP1:a:2452:[19.4], C:OP1:a:2503:[19.8]) = 3.15 -- dist(a:A:2452:OP1, a:MG:3325:MG:[23.2]) = 1.92 -- -- dist(a:C:2503:OP1, a:MG:3325:MG:[23.2]) = 2.08 --   
28. -- dist(C:OP1:a:2577:[27.5], G:OP1:a:2578:[26.7]) = 2.82 -- dist(a:C:2577:OP1, a:MG:3331:MG:[28.6]) = 2.00 -- -- dist(a:G:2578:OP1, a:MG:3331:MG:[28.6]) = 1.99 --   
29. -- dist(G:OP1:a:2580:[23.4], A:OP1:a:2581:[24.1]) = 3.10 -- dist(a:G:2580:OP1, a:MG:3194:MG:[27.2]) = 2.14 -- -- dist(a:A:2581:OP1, a:MG:3194:MG:[27.2]) = 2.13 --   
30. -- dist(A:OP1:A:59:[54.7], U:OP1:A:387:[56.2]) = 3.16 -- dist(A:A:59:OP1, A:MG:1624:MG:[66.9]) = 1.96 -- **← CHECK**  
31. -- dist(G:OP1:A:450:**[75.3]**, A:OP1:A:452:**[74.8]**) = 2.92 **← CHECK**  
32. -- dist(G:OP1:A:558:[39.5], A:OP1:A:560:[42.4]) = 3.10 -- dist(A:G:558:OP1, A:MG:1627:MG:[54.3]) = 2.16 -- **← CHECK**  
33. -- dist(A:OP1:A:964:**[61.8]**, U:OP1:A:1199:**[56.5]**) = 2.82 -- dist(A:A:964:OP1, A:MG:1700:MG:[65.0]) = 1.93 -- -- dist(A:U:1199:OP1, A:MG:1700:MG:**[65.0]**) = 2.23 --   
34. -- dist(C:OP1:A:1054:**[64.6]**, A:OP1:A:1197:**[56.4]**) = 3.13 -- dist(A:C:1054:OP1, A:MG:1701:MG:[59.2]) = 1.95 -- -- dist(A:A:1197:OP1, A:MG:1701:MG:[59.2]) = 1.92 --   
35. -- dist(G:OP1:A:1068:[53.6], G:OP1:A:1094:[57.2]) = 3.03 -- dist(A:G:1068:OP1, A:MG:1706:MG:[61.1]) = 1.84 -- -- dist(A:G:1094:OP1, A:MG:1706:MG:**[61.1]**) = 1.98 --   
36. -- dist(A:OP1:A:1500:[32.4], G:OP1:A:1505:[36.2]) = 2.89 -- dist(A:A:1500:OP1, A:MG:1634:MG:[37.8]) = 1.99 -- -- dist(A:G:1505:OP1, A:MG:1634:MG:[37.8]) = 2.12 --   
37. -- dist(A:OP1:A:1500:[32.4], A:OP1:A:1508:[31.1]) = 2.88 -- dist(A:A:1500:OP1, A:MG:1634:MG:[37.8]) = 1.99 -- -- dist(A:A:1508:OP1, A:MG:1634:MG:[37.8]) = 2.06 --   
**← Detected number of OP1...OP1 clamps (with d < 3.4 Å):** **33**  
  
**Potential number of OP1...OP2 clamps (with d < 3.4 Å):** **25**   
 1. -- dist(A:OP1:a:197:[22.5], C:OP2:a:198:[21.1]) = 3.13 -- dist(a:A:197:OP1, a:MG:3088:MG:[24.4]) = 2.12 -- -- dist(a:C:198:OP2, a:MG:3088:MG:[24.4]) = 2.14 --   
2. -- dist(C:OP1:a:510:[29.8], U:OP2:a:511:[30.4]) = 2.90 -- dist(a:C:510:OP1, a:MG:3091:MG:[28.5]) = 2.22 -- -- dist(a:U:511:OP2, a:MG:3091:MG:[28.5]) = 1.81 --   
3. -- dist(U:OP1:a:829:[26.1], G:OP2:a:2433:[24.6]) = 3.04 -- dist(a:U:829:OP1, a:MG:3230:MG:[28.0]) = 1.99 -- -- dist(a:G:2433:OP2, a:MG:3230:MG:[28.0]) = 2.06 --   
4. -- dist(G:OP1:a:964:[29.2], U:OP2:a:965:[22.0]) = 2.89 -- dist(a:G:964:OP1, a:MG:3233:MG:[26.0]) = 2.12 -- -- dist(a:U:965:OP2, a:MG:3233:MG:[26.0]) = 2.02 --   
5. -- dist(G:OP1:a:1159:[31.2], C:OP2:a:1160:[32.7]) = 3.18 -- dist(a:G:1159:OP1, a:MG:3192:MG:[41.5]) = 2.04 -- -- dist(a:C:1160:OP2, a:MG:3192:MG:[41.5]) = 2.18 --   
6. -- dist(C:OP1:a:1316:[26.7], C:OP2:a:1317:[41.0]) = 2.86 -- dist(a:C:1316:OP1, a:MG:3089:MG:[28.9]) = 2.01 -- -- dist(a:C:1317:OP2, a:MG:3089:MG:[28.9]) = 1.98 --   
7. -- dist(A:OP1:a:1361:[34.6], G:OP2:a:1362:[37.8]) = 3.19 -- dist(a:A:1361:OP1, a:MG:3316:MG:[38.2]) = 2.17 -- -- dist(a:G:1362:OP2, a:MG:3316:MG:[38.2]) = 2.08 --   
8. -- dist(A:OP1:a:1397:[33.2], C:OP2:a:1606:[30.3]) = 3.13 -- dist(a:A:1397:OP1, a:MG:3244:MG:[30.5]) = 2.04 -- -- dist(a:C:1606:OP2, a:MG:3244:MG:[30.5]) = 1.99 --   
9. -- dist(A:OP1:a:1605:[31.3], C:OP2:a:1606:[30.3]) = 2.91 -- dist(a:A:1605:OP1, a:MG:3244:MG:[30.5]) = 1.98 -- -- dist(a:C:1606:OP2, a:MG:3244:MG:[30.5]) = 1.99 --   
10. -- dist(A:OP1:a:1666:[29.7], A:OP2:a:1667:[28.9]) = 2.98 -- dist(a:A:1666:OP1, a:MG:3351:MG:[34.5]) = 2.16 -- -- dist(a:A:1667:OP2, a:MG:3351:MG:[34.5]) = 2.09 --   
11. -- dist(C:OP1:a:1672:[23.5], U:OP2:a:1673:[28.5]) = 2.87 -- dist(a:C:1672:OP1, a:MG:3287:MG:[28.4]) = 2.02 -- -- dist(a:U:1673:OP2, a:MG:3287:MG:[28.4]) = 1.98 --   
12. -- dist(U:OP1:a:1784:[27.2], A:OP2:a:1785:[23.2]) = 2.85 -- dist(a:U:1784:OP1, a:MG:3326:MG:[23.7]) = 1.93 -- -- dist(a:A:1785:OP2, a:MG:3326:MG:[23.7]) = 1.99 --   
13. -- dist(A:OP1:a:1785:[25.0], A:OP2:a:1786:[24.1]) = 3.00 -- dist(a:A:1785:OP1, a:MG:3328:MG:[27.0]) = 2.03 -- -- dist(a:A:1786:OP2, a:MG:3328:MG:[27.0]) = 2.20 --   
14. -- dist(C:OP1:a:2218:[43.6], G:OP2:a:2219:[41.1]) = 3.37 **← CHECK**  
15. -- dist(C:OP1:a:2577:[27.5], C:OP2:a:2579:[24.4]) = 3.06 -- dist(a:C:2577:OP1, a:MG:3331:MG:[28.6]) = 2.00 -- -- dist(a:C:2579:OP2, a:MG:3331:MG:[28.6]) = 2.10 --   
16. -- dist(G:OP1:a:2578:[26.7], C:OP2:a:2579:[24.4]) = 3.11 -- dist(a:G:2578:OP1, a:MG:3331:MG:[28.6]) = 1.99 -- -- dist(a:C:2579:OP2, a:MG:3331:MG:[28.6]) = 2.10 --   
17. -- dist(C:OP1:a:2650:[30.6], U:OP2:a:2651:[33.5]) = 3.31 -- dist(a:C:2650:OP1, a:MG:3046:MG:[43.8]) = 2.29 -- -- dist(a:U:2651:OP2, a:MG:3046:MG:[43.8]) = 2.23 --   
18. -- dist(A:OP1:a:2770:[34.7], C:OP2:a:2771:[34.6]) = 3.05 -- dist(a:A:2770:OP1, a:MG:3020:MG:[42.7]) = 2.05 -- **← CHECK**  
19. -- dist(A:OP1:A:174:**[71.9]**, C:OP2:A:175:**[76.9]**) = 2.82 -- dist(A:A:174:OP1, A:MG:1626:MG:[66.9]) = 2.12 -- -- dist(A:C:175:OP2, A:MG:1626:MG:**[66.9]**) = 1.93 --   
20. -- dist(U:OP1:A:239:**[57.5]**, G:OP2:A:240:**[63.6]**) = 3.32 **← CHECK**  
21. -- dist(G:OP1:A:944:**[62.4]**, G:OP2:A:945:**[70.2]**) = 3.10 -- dist(A:G:944:OP1, A:MG:1691:MG:[73.6]) = 1.97 -- -- dist(A:G:945:OP2, A:MG:1691:MG:**[73.6]**) = 2.31 --   
22. -- dist(C:OP1:A:979:**[73.9]**, C:OP2:A:980:**[76.7]**) = 3.26 -- dist(A:C:979:OP1, A:MG:1689:MG:[80.7]) = 1.87 -- -- dist(A:C:980:OP2, A:MG:1689:MG:**[80.7]**) = 2.05 --   
23. -- dist(C:OP1:A:1054:**[64.6]**, G:OP2:A:1198:**[57.0]**) = 3.06 -- dist(A:C:1054:OP1, A:MG:1701:MG:[59.2]) = 1.95 -- -- dist(A:G:1198:OP2, A:MG:1701:MG:[59.2]) = 2.07 --   
24. -- dist(A:OP1:A:1197:[56.4], G:OP2:A:1198:[57.0]) = 2.80 -- dist(A:A:1197:OP1, A:MG:1701:MG:[59.2]) = 1.92 -- -- dist(A:G:1198:OP2, A:MG:1701:MG:[59.2]) = 2.07 --   
25. -- dist(C:OP1:A:1303:**[80.6]**, G:OP2:A:1304:**[81.1]**) = 2.66 -- dist(A:C:1303:OP1, A:MG:1717:MG:[76.0]) = 1.96 -- -- dist(A:G:1304:OP2, A:MG:1717:MG:**[76.0]**) = 1.94 --   
**← Detected number of OP1...OP2 clamps (with d < 3.4 Å):** **22**  
  
**Potential number of OP2...OP2 clamps (with d < 3.4 Å):** **41**   
 1. -- dist(A:OP2:a:14:[35.6], G:OP2:a:15:[37.4]) = 2.96 **← CHECK**  
2. -- dist(G:OP2:a:46:[41.5], G:OP2:a:215:[45.8]) = 3.32 **← CHECK**  
3. -- dist(C:OP2:a:163:**[71.1]**, C:OP2:a:164:**[70.7]**) = 3.38 **← CHECK**  
4. -- dist(A:OP2:a:278:**[150.9]**, A:OP2:a:279:**[150.5]**) = 2.86 **← CHECK**  
5. -- dist(A:OP2:a:623:[38.0], G:OP2:a:624:[32.0]) = 3.02 -- dist(a:A:623:OP2, a:MG:3219:MG:[39.0]) = 2.03 -- -- dist(a:G:624:OP2, a:MG:3219:MG:[39.0]) = 2.10 --   
6. -- dist(A:OP2:a:732:[24.1], C:OP2:a:733:[24.7]) = 2.84 -- dist(a:A:732:OP2, a:MG:3327:MG:[25.9]) = 1.94 -- -- dist(a:C:733:OP2, a:MG:3327:MG:[25.9]) = 1.95 --   
7. -- dist(C:OP2:a:742:[28.4], A:OP2:a:1786:[24.1]) = 3.22 -- dist(a:C:742:OP2, a:MG:3328:MG:[27.0]) = 2.08 -- -- dist(a:A:1786:OP2, a:MG:3328:MG:[27.0]) = 2.20 --   
8. -- dist(A:OP2:a:785:[20.0], G:OP2:a:786:[20.6]) = 3.24 -- dist(a:A:785:OP2, a:MG:3224:MG:[24.7]) = 2.03 -- -- dist(a:G:786:OP2, a:MG:3224:MG:[24.7]) = 2.01 --   
9. -- dist(A:OP2:a:947:[24.9], C:OP2:a:948:[29.6]) = 2.89 -- dist(a:A:947:OP2, a:MG:3231:MG:[30.4]) = 2.21 -- -- dist(a:C:948:OP2, a:MG:3231:MG:[30.4]) = 1.98 --   
10. -- dist(A:OP2:a:992:[31.0], C:OP2:a:993:[26.4]) = 2.99 -- dist(a:A:992:OP2, a:MG:3314:MG:[25.8]) = 1.93 -- -- dist(a:C:993:OP2, a:MG:3314:MG:[25.8]) = 2.08 --   
11. -- dist(C:OP2:a:993:[26.4], G:OP2:a:1188:[25.6]) = 2.87 -- dist(a:C:993:OP2, a:MG:3314:MG:[25.8]) = 2.08 -- -- dist(a:G:1188:OP2, a:MG:3314:MG:[25.8]) = 2.15 --   
12. -- dist(A:OP2:a:1011:[28.4], A:OP2:a:1012:[26.8]) = 2.94 -- dist(a:A:1011:OP2, a:MG:3093:MG:[31.2]) = 1.98 -- -- dist(a:A:1012:OP2, a:MG:3093:MG:[31.2]) = 2.06 --   
13. -- dist(C:OP2:a:1015:[35.2], A:OP2:a:1016:[43.8]) = 2.89 **← CHECK**  
14. -- dist(U:OP2:a:1025:[30.2], G:OP2:a:1026:[33.2]) = 2.97 -- dist(a:U:1025:OP2, a:MG:3144:MG:[34.8]) = 2.05 -- -- dist(a:G:1026:OP2, a:MG:3144:MG:[34.8]) = 1.81 --   
15. -- dist(C:OP2:a:1347:[38.4], G:OP2:a:1348:[42.5]) = 3.02 **← CHECK**  
16. -- dist(U:OP2:a:1381:[26.9], G:OP2:a:1382:[30.9]) = 3.00 -- dist(a:U:1381:OP2, a:MG:3145:MG:[31.0]) = 2.02 -- -- dist(a:G:1382:OP2, a:MG:3145:MG:[31.0]) = 2.12 --   
17. -- dist(A:OP2:a:1671:[21.8], C:OP2:a:1672:[20.9]) = 2.99 -- dist(a:A:1671:OP2, a:MG:3333:MG:[24.2]) = 1.97 -- -- dist(a:C:1672:OP2, a:MG:3333:MG:[24.2]) = 2.02 --   
18. -- dist(A:OP2:a:1850:**[61.5]**, G:OP2:a:1851:**[56.0]**) = 3.16 **← CHECK**  
19. -- dist(G:OP2:a:2060:[26.2], G:OP2:a:2061:[24.7]) = 2.81 -- dist(a:G:2060:OP2, a:MG:3294:MG:[28.9]) = 2.03 -- -- dist(a:G:2061:OP2, a:MG:3294:MG:[28.9]) = 2.02 --   
20. -- dist(G:OP2:a:2065:[22.6], G:OP2:a:2506:[25.3]) = 2.96 -- dist(a:G:2065:OP2, a:MG:3295:MG:[26.3]) = 1.99 -- -- dist(a:G:2506:OP2, a:MG:3295:MG:[26.3]) = 1.91 --   
21. -- dist(G:OP2:a:2433:[24.6], A:OP2:a:2434:[35.8]) = 3.26 -- dist(a:G:2433:OP2, a:MG:3230:MG:[28.0]) = 2.06 -- **← CHECK**  
22. -- dist(A:OP2:a:2452:[21.3], OMC:OP2:a:2502:[28.7]) = 3.09 -- dist(a:A:2452:OP2, a:MG:3324:MG:[26.3]) = 2.23 -- -- dist(a:OMC:2502:OP2, a:MG:3324:MG:[26.3]) = 1.95 --   
23. -- dist(A:OP2:a:2452:[21.3], C:OP2:a:2503:[22.8]) = 3.06 -- dist(a:A:2452:OP2, a:MG:3324:MG:[26.3]) = 2.23 -- -- dist(a:C:2503:OP2, a:MG:3324:MG:[26.3]) = 2.10 --   
24. -- dist(OMC:OP2:a:2502:[28.7], C:OP2:a:2503:[22.8]) = 2.86 -- dist(a:OMC:2502:OP2, a:MG:3324:MG:[26.3]) = 1.95 -- -- dist(a:C:2503:OP2, a:MG:3324:MG:[26.3]) = 2.10 --   
25. -- dist(G:OP2:a:2547:[31.9], G:OP2:a:2548:[29.1]) = 3.12 **← CHECK**  
26. -- dist(G:OP2:a:2586:[33.6], G:OP2:a:2587:[37.7]) = 2.80 -- dist(a:G:2586:OP2, a:MG:3023:MG:[48.2]) = 2.11 -- -- dist(a:G:2587:OP2, a:MG:3023:MG:[48.2]) = 2.12 --   
27. -- dist(A:OP2:a:2715:[24.9], G:OP2:a:2718:[28.0]) = 2.98 -- dist(a:A:2715:OP2, a:MG:3306:MG:[31.2]) = 1.89 -- -- dist(a:G:2718:OP2, a:MG:3306:MG:[31.2]) = 2.00 --   
28. -- dist(A:OP2:a:2825:[28.3], G:OP2:a:2826:[23.5]) = 2.78 -- dist(a:A:2825:OP2, a:MG:3186:MG:[26.6]) = 1.97 -- -- dist(a:G:2826:OP2, a:MG:3186:MG:[26.6]) = 1.97 --   
29. -- dist(A:OP2:b:57:**[67.6]**, A:OP2:b:58:**[60.8]**) = 2.78 -- dist(b:A:57:OP2, b:MG:202:MG:[62.5]) = 2.08 -- -- dist(b:A:58:OP2, b:MG:202:MG:**[62.5]**) = 1.90 --   
30. -- dist(A:OP2:A:116:[45.6], G:OP2:A:117:[45.6]) = 2.94 -- dist(A:A:116:OP2, A:MG:1681:MG:[47.7]) = 2.00 -- -- dist(A:G:117:OP2, A:MG:1681:MG:[47.7]) = 2.14 --   
31. -- dist(A:OP2:A:116:[45.6], G:OP2:A:289:[42.5]) = 3.04 -- dist(A:A:116:OP2, A:MG:1681:MG:[47.7]) = 2.00 -- -- dist(A:G:289:OP2, A:MG:1681:MG:[47.7]) = 2.00 --   
32. -- dist(A:OP2:A:412:**[106.6]**, G:OP2:A:413:**[102.9]**) = 3.21 **← CHECK**  
33. -- dist(A:OP2:A:468:**[129.5]**, C:OP2:A:469:**[128.5]**) = 2.79 **← CHECK**  
34. -- dist(G:OP2:A:497:**[89.9]**, A:OP2:A:498:**[81.4]**) = 3.39 **← CHECK**  
35. -- dist(A:OP2:A:509:**[64.9]**, A:OP2:A:510:**[63.8]**) = 2.78 -- dist(A:A:509:OP2, A:MG:1679:MG:[69.9]) = 2.01 -- -- dist(A:A:510:OP2, A:MG:1679:MG:**[69.9]**) = 2.16 --   
36. -- dist(A:OP2:A:572:[40.8], A:OP2:A:573:[34.0]) = 3.09 -- dist(A:A:572:OP2, A:MG:1672:MG:[37.2]) = 2.32 -- -- dist(A:A:573:OP2, A:MG:1672:MG:[37.2]) = 2.01 --   
37. -- dist(A:OP2:A:573:[34.0], A:OP2:A:574:[39.7]) = 2.96 -- dist(A:A:573:OP2, A:MG:1672:MG:[37.2]) = 2.01 -- -- dist(A:A:574:OP2, A:MG:1672:MG:[37.2]) = 2.05 --   
38. -- dist(C:OP2:A:1054:**[60.1]**, A:OP2:A:1197:**[59.9]**) = 2.87 -- dist(A:C:1054:OP2, A:MG:1702:MG:[62.3]) = 1.94 -- -- dist(A:A:1197:OP2, A:MG:1702:MG:**[62.3]**) = 1.91 --   
39. -- dist(A:OP2:A:1499:[35.4], A:OP2:A:1500:[34.8]) = 2.79 -- dist(A:A:1499:OP2, A:MG:1671:MG:[35.6]) = 2.03 -- -- dist(A:A:1500:OP2, A:MG:1671:MG:[35.6]) = 2.07 --   
40. -- dist(A:OP2:A:1499:[35.4], G:OP2:A:1505:[33.4]) = 3.01 -- dist(A:A:1499:OP2, A:MG:1671:MG:[35.6]) = 2.03 -- -- dist(A:G:1505:OP2, A:MG:1671:MG:[35.6]) = 2.17 --   
41. -- dist(A:OP2:A:1500:[34.8], G:OP2:A:1505:[33.4]) = 2.95 -- dist(A:A:1500:OP2, A:MG:1671:MG:[35.6]) = 2.07 -- -- dist(A:G:1505:OP2, A:MG:1671:MG:[35.6]) = 2.17 --   
**← Detected number of OP2...OP2 clamps (with d < 3.4 Å):** **29**  
 **---> Total number of detected of OP...OP clamps (with d < 3.4 Å):** **84 <---**  
  
  
**Water molecules with 2Fo-Fc values > 3.0:**   
  

---

### Total number of ions:

**Total number of Mg^2+^: 361** **Total number of Mg^2+^ with stereochemical issues: 263** **Total number of Mg^2+^ without stereochemical issues: 98** **Total number of K^+^: 168** **Total number of K^+^ with stereochemical issues: 80** **Total number of K^+^ without stereochemical issues: 88** **Total number of Zn^2+^: 1** **Total number of Zn^2+^ with stereochemical issues: 0** **Total number of Zn^2+^ without stereochemical issues: 1** 

### ••• END •••

#### (duration of the job: 0:02:02.807236)* 
